## Supplemental Tables and Figures for "Single cell transcriptomic profiling identifies molecular phenotypes of newborn human lung cells"

Running Title: Newborn human lung cellular diversity

Key Words: single cell RNAseq, matrix fibroblast, lung development, newborn lung

No. of Tables: 0

No. of Figures: 6

No of Supplemental Tables: 11

No. of Supplemental Figures: 9

This work was supported by the Human Developing Lung Molecular Atlas Program (LungMAP) funded by National Heart, Lung, and Blood Institute of National Institutes of Health (U01HL122700 to G.S.P. and U01HL122642 to J.A.W.).

Table S1: Distribution of human cells by sample source and capture

| Cluster | Major Type | Subtype | Donor1 Run1 | Donor1 Run2 | Donor2 Run1 | Donor2 Run2 | Total |
| --- | --- | --- | --- | --- | --- | --- | --- |
| 0 | Mesenchymal | Myofibroblasts | 299 | 267 | 153 | 101 | 820 |
| 1 | Mesenchymal | Matrix Fibroblasts | 157 | 123 | 342 | 192 | 814 |
| 2 | Mesenchymal | Smooth Muscle Cell | 237 | 173 | 114 | 68 | 592 |
| 4 | Mesenchymal | Pericytes | 82 | 147 | 162 | 119 | 510 |
| 5 | Mesenchymal | Stromal Cells | 87 | 72 | 172 | 88 | 419 |
| 6 | Mesenchymal | Pericytes | 121 | 110 | 100 | 67 | 398 |
| 3 | Endothelial | Endothelial Type B | 59 | 107 | 159 | 242 | 567 |
| 8 | Endothelial | Endothelial Type B | 70 | 96 | 87 | 68 | 321 |
| 10 | Endothelial | Endothelial Type C | 40 | 71 | 20 | 15 | 146 |
| 12 | Endothelial | Endothelial Type B | 27 | 28 | 15 | 15 | 85 |
| 7 | Immune | Macrophages | 180 | 144 | 14 | 11 | 349 |
| 9 | Immune | T-Cells | 83 | 58 | 32 | 17 | 190 |
| 13 | Immune | B-Cells | 29 | 23 | 17 | 10 | 79 |
| 11 | Epithelial | AT1 Epithelial | 44 | 29 | 34 | 24 | 131 |
| 14 | Epithelial | AT2 Epithelial | 0 | 38 | 0 | 40 | 78 |
| Total |  |  | 1515 | 1486 | 1421 | 1077 | 5499 |

Table S2: Marker genes for individual clusters of cells from newborn human cells

| Genes | Average Log Fold Change | Unadjusted P-value | Adjusted P-Value | Proportion of cells in Group 1 | Proportion of cells in Group 2 | Cluster No | Sub-Type |
| --- | --- | --- | --- | --- | --- | --- | --- |
| ACTG2 | 2.61 | 0 | 0 | 0.94 | 0.19 | 0 | Myofibroblasts |
| DES | 2.6 | 0 | 0 | 0.87 | 0.10 | 0 | Myofibroblasts |
| ACTA2 | 2.15 | 0 | 0 | 1.00 | 0.70 | 0 | Myofibroblasts |
| TAGLN | 2.05 | 0 | 0 | 0.99 | 0.56 | 0 | Myofibroblasts |
| CNN1 | 1.98 | 0 | 0 | 0.84 | 0.11 | 0 | Myofibroblasts |
| MYLK | 1.93 | 0 | 0 | 0.95 | 0.42 | 0 | Myofibroblasts |
| MYH11 | 1.86 | 0 | 0 | 0.97 | 0.31 | 0 | Myofibroblasts |
| HHIP | 1.81 | 0 | 0 | 0.62 | 0.04 | 0 | Myofibroblasts |
| FILIP1L | 1.45 | 0 | 0 | 0.75 | 0.20 | 0 | Myofibroblasts |
| RAMP1 | 1.45 | 0 | 0 | 0.57 | 0.10 | 0 | Myofibroblasts |
| FGF18 | 1.39 | 0 | 0 | 0.57 | 0.04 | 0 | Myofibroblasts |
| CKB | 1.36 | 0 | 0 | 0.63 | 0.16 | 0 | Myofibroblasts |
| TPM2 | 1.36 | 0 | 0 | 0.99 | 0.61 | 0 | Myofibroblasts |
| MT1X | 1.31 | 0 | 0 | 0.48 | 0.16 | 0 | Myofibroblasts |
| FLNA | 1.22 | 0 | 0 | 0.89 | 0.42 | 0 | Myofibroblasts |
| TNFRSF12A | 1.16 | 0 | 0 | 0.44 | 0.12 | 0 | Myofibroblasts |
| PDLIM7 | 1.16 | 0 | 0 | 0.81 | 0.31 | 0 | Myofibroblasts |
| LPP | 1.14 | 0 | 0 | 0.75 | 0.28 | 0 | Myofibroblasts |
| MYL9 | 1.12 | 0 | 0 | 0.98 | 0.67 | 0 | Myofibroblasts |
| SYNPO2 | 1.07 | 0 | 0 | 0.52 | 0.07 | 0 | Myofibroblasts |
| LTBP1 | 1.06 | 0 | 0 | 0.61 | 0.17 | 0 | Myofibroblasts |
| TPM1 | 1.05 | 0 | 0 | 0.91 | 0.55 | 0 | Myofibroblasts |
| SCX | 1.05 | 0 | 0 | 0.37 | 0.01 | 0 | Myofibroblasts |
| DSTN | 1.04 | 0 | 0 | 0.99 | 0.76 | 0 | Myofibroblasts |
| SELM | 1.02 | 0 | 0 | 0.79 | 0.44 | 0 | Myofibroblasts |
| MGP | 1.02 | 0 | 0 | 0.80 | 0.63 | 0 | Myofibroblasts |
| CARMN | 1.01 | 0 | 0 | 0.76 | 0.28 | 0 | Myofibroblasts |
| CLU | 1 | 0 | 0 | 0.77 | 0.25 | 0 | Myofibroblasts |
| CSRP1 | 0.97 | 0 | 0 | 0.92 | 0.55 | 0 | Myofibroblasts |
| LMCD1 | 0.96 | 0 | 0 | 0.74 | 0.32 | 0 | Myofibroblasts |
| TGFBI | 0.96 | 0 | 0 | 0.59 | 0.32 | 0 | Myofibroblasts |
| NEXN | 0.96 | 0 | 0 | 0.61 | 0.16 | 0 | Myofibroblasts |
| IGFBP5 | 0.95 | 0 | 0 | 0.54 | 0.26 | 0 | Myofibroblasts |
| PDLIM3 | 0.94 | 0 | 0 | 0.68 | 0.23 | 0 | Myofibroblasts |
| LDLRAD4 | 0.93 | 0 | 0 | 0.42 | 0.05 | 0 | Myofibroblasts |
| GREM2 | 0.89 | 0 | 0 | 0.37 | 0.01 | 0 | Myofibroblasts |
| CTGF | 0.89 | 0 | 0 | 0.47 | 0.18 | 0 | Myofibroblasts |
| CALD1 | 0.89 | 0 | 0 | 0.98 | 0.76 | 0 | Myofibroblasts |
| NDUFS5 | 0.87 | 0 | 0 | 0.93 | 0.71 | 0 | Myofibroblasts |
| HSPB1 | 0.87 | 0 | 0 | 0.94 | 0.74 | 0 | Myofibroblasts |
| KCNMB1 | 0.87 | 0 | 0 | 0.46 | 0.06 | 0 | Myofibroblasts |
| LMNA | 0.87 | 0 | 0 | 0.88 | 0.64 | 0 | Myofibroblasts |
| THBS1 | 0.85 | 0 | 0 | 0.37 | 0.13 | 0 | Myofibroblasts |
| PALLD | 0.83 | 0 | 0 | 0.59 | 0.18 | 0 | Myofibroblasts |
| SEMA3C | 0.83 | 0 | 0 | 0.46 | 0.07 | 0 | Myofibroblasts |
| SVIL | 0.82 | 0 | 0 | 0.51 | 0.14 | 0 | Myofibroblasts |
| BAG2 | 0.81 | 0 | 0 | 0.47 | 0.11 | 0 | Myofibroblasts |
| CRYAB | 0.81 | 0 | 0 | 0.69 | 0.29 | 0 | Myofibroblasts |
| HHIP-AS1 | 0.8 | 0 | 0 | 0.40 | 0.02 | 0 | Myofibroblasts |
| SAMD11 | 0.78 | 0 | 0 | 0.37 | 0.03 | 0 | Myofibroblasts |
| FSTL3 | 0.78 | 0 | 0 | 0.39 | 0.08 | 0 | Myofibroblasts |
| SMTN | 0.78 | 0 | 0 | 0.58 | 0.24 | 0 | Myofibroblasts |
| RHOB | 0.77 | 0 | 0 | 0.77 | 0.43 | 0 | Myofibroblasts |
| ALDH1B1 | 0.74 | 0 | 0 | 0.34 | 0.04 | 0 | Myofibroblasts |
| CNN3 | 0.74 | 0 | 0 | 0.78 | 0.56 | 0 | Myofibroblasts |
| SPEG | 0.73 | 0 | 0 | 0.42 | 0.07 | 0 | Myofibroblasts |
| SCARA3 | 0.73 | 0 | 0 | 0.39 | 0.07 | 0 | Myofibroblasts |
| PMEPA1 | 0.72 | 0 | 0 | 0.39 | 0.11 | 0 | Myofibroblasts |
| C9orf3 | 0.71 | 0 | 0 | 0.58 | 0.27 | 0 | Myofibroblasts |
| THBS2 | 0.71 | 0 | 0 | 0.32 | 0.06 | 0 | Myofibroblasts |
| TUBB2A | 0.71 | 0 | 0 | 0.38 | 0.13 | 0 | Myofibroblasts |
| FHL1 | 0.7 | 0 | 0 | 0.86 | 0.53 | 0 | Myofibroblasts |
| LMOD1 | 0.68 | 0 | 0 | 0.42 | 0.10 | 0 | Myofibroblasts |
| TGFB1I1 | 0.68 | 0 | 0 | 0.72 | 0.37 | 0 | Myofibroblasts |
| BCHE | 0.67 | 0 | 0 | 0.39 | 0.13 | 0 | Myofibroblasts |
| SLMAP | 0.66 | 0 | 0 | 0.40 | 0.14 | 0 | Myofibroblasts |
| MRPS6 | 0.66 | 0 | 0 | 0.57 | 0.29 | 0 | Myofibroblasts |
| MYOCD | 0.66 | 0 | 0 | 0.34 | 0.05 | 0 | Myofibroblasts |
| SBSPON | 0.66 | 0 | 0 | 0.31 | 0.06 | 0 | Myofibroblasts |
| NDUFA4 | 0.65 | 0 | 0 | 0.95 | 0.78 | 0 | Myofibroblasts |
| ACTN1 | 0.65 | 0 | 0 | 0.70 | 0.40 | 0 | Myofibroblasts |
| IGF1 | 0.65 | 0 | 0 | 0.31 | 0.05 | 0 | Myofibroblasts |
| TGFBR3 | 0.65 | 0 | 0 | 0.38 | 0.10 | 0 | Myofibroblasts |
| TUBB6 | 0.63 | 0 | 0 | 0.51 | 0.28 | 0 | Myofibroblasts |
| ID4 | 0.63 | 0 | 0 | 0.59 | 0.21 | 0 | Myofibroblasts |
| NUPR1 | 0.62 | 0 | 0 | 0.81 | 0.45 | 0 | Myofibroblasts |
| SH3PXD2A | 0.61 | 0 | 0 | 0.45 | 0.18 | 0 | Myofibroblasts |
| SPARCL1 | 0.61 | 0 | 0 | 0.91 | 0.63 | 0 | Myofibroblasts |
| BOC | 0.61 | 0 | 0 | 0.30 | 0.04 | 0 | Myofibroblasts |
| BTG3 | 0.6 | 0 | 0 | 0.47 | 0.20 | 0 | Myofibroblasts |
| WSB1 | 0.6 | 0 | 0 | 0.83 | 0.63 | 0 | Myofibroblasts |
| MYL6 | 0.6 | 0 | 0 | 1.00 | 0.98 | 0 | Myofibroblasts |
| PDLIM4 | 0.6 | 0 | 0 | 0.32 | 0.07 | 0 | Myofibroblasts |
| PPP1R12B | 0.6 | 0 | 0 | 0.45 | 0.17 | 0 | Myofibroblasts |
| MYADM | 0.59 | 0 | 0 | 0.61 | 0.34 | 0 | Myofibroblasts |
| CRISPLD2 | 0.59 | 0 | 0 | 0.44 | 0.20 | 0 | Myofibroblasts |
| TACC2 | 0.59 | 0 | 0 | 0.31 | 0.05 | 0 | Myofibroblasts |
| CORO1C | 0.59 | 0 | 0 | 0.43 | 0.17 | 0 | Myofibroblasts |
| ADAMTS6 | 0.57 | 0 | 0 | 0.35 | 0.08 | 0 | Myofibroblasts |
| HSPB7 | 0.57 | 0 | 0 | 0.32 | 0.06 | 0 | Myofibroblasts |
| AP000892.6 | 0.56 | 0 | 0 | 0.26 | 0.04 | 0 | Myofibroblasts |
| COL4A5 | 0.55 | 0 | 0 | 0.36 | 0.10 | 0 | Myofibroblasts |
| TUBA1A | 0.55 | 0 | 0 | 0.89 | 0.78 | 0 | Myofibroblasts |
| LGALS3 | 0.54 | 0 | 0 | 0.54 | 0.34 | 0 | Myofibroblasts |
| ANTXR1 | 0.54 | 0 | 0 | 0.46 | 0.20 | 0 | Myofibroblasts |
| FHL2 | 0.54 | 0 | 0 | 0.31 | 0.07 | 0 | Myofibroblasts |
| COL16A1 | 0.54 | 0 | 0 | 0.30 | 0.07 | 0 | Myofibroblasts |
| NFIC | 0.54 | 0 | 0 | 0.59 | 0.33 | 0 | Myofibroblasts |
| GEM | 0.53 | 0 | 0 | 0.37 | 0.12 | 0 | Myofibroblasts |
| H2AFZ | 0.53 | 0 | 0 | 0.80 | 0.66 | 0 | Myofibroblasts |
| ADAMTS8 | 0.52 | 0 | 0 | 0.28 | 0.08 | 0 | Myofibroblasts |
| GADD45A | 0.52 | 0 | 0 | 0.30 | 0.15 | 0 | Myofibroblasts |
| IRS1 | 0.52 | 0 | 0 | 0.31 | 0.08 | 0 | Myofibroblasts |
| NSMCE4A | 0.51 | 0 | 0 | 0.37 | 0.15 | 0 | Myofibroblasts |
| PRNP | 0.5 | 0 | 0 | 0.54 | 0.31 | 0 | Myofibroblasts |
| WDR1 | 0.49 | 0 | 0 | 0.56 | 0.35 | 0 | Myofibroblasts |
| C11orf96 | 0.49 | 0 | 0 | 0.50 | 0.26 | 0 | Myofibroblasts |
| MFAP2 | 0.48 | 0 | 0 | 0.47 | 0.25 | 0 | Myofibroblasts |
| MEG3 | 0.48 | 0 | 0 | 0.60 | 0.42 | 0 | Myofibroblasts |
| RGS2 | 0.48 | 0 | 0 | 0.49 | 0.29 | 0 | Myofibroblasts |
| ID2 | 0.47 | 0 | 0 | 0.79 | 0.59 | 0 | Myofibroblasts |
| CD151 | 0.47 | 0 | 0 | 0.67 | 0.52 | 0 | Myofibroblasts |
| VCL | 0.47 | 0 | 0 | 0.42 | 0.19 | 0 | Myofibroblasts |
| MDFI | 0.45 | 0 | 0 | 0.36 | 0.16 | 0 | Myofibroblasts |
| CYR61 | 0.45 | 0 | 0 | 0.40 | 0.21 | 0 | Myofibroblasts |
| RAB23 | 0.45 | 0 | 0 | 0.28 | 0.08 | 0 | Myofibroblasts |
| CSRP2 | 0.44 | 0 | 0 | 0.43 | 0.21 | 0 | Myofibroblasts |
| CD9 | 0.44 | 0 | 0 | 0.74 | 0.41 | 0 | Myofibroblasts |
| MRGPRF | 0.43 | 0 | 0 | 0.40 | 0.18 | 0 | Myofibroblasts |
| TSPAN2 | 0.43 | 0 | 0 | 0.28 | 0.08 | 0 | Myofibroblasts |
| SEPT6 | 0.42 | 0 | 0 | 0.34 | 0.19 | 0 | Myofibroblasts |
| ILK | 0.42 | 0 | 0 | 0.61 | 0.41 | 0 | Myofibroblasts |
| RP11-756G12.1 | 0.42 | 0 | 0 | 0.28 | 0.08 | 0 | Myofibroblasts |
| FNBP1 | 0.41 | 0 | 0 | 0.39 | 0.19 | 0 | Myofibroblasts |
| CRY1 | 0.41 | 0 | 0 | 0.28 | 0.10 | 0 | Myofibroblasts |
| FAU | 0.41 | 0 | 0 | 1.00 | 0.97 | 0 | Myofibroblasts |
| EEF1B2 | 0.41 | 0 | 0 | 0.88 | 0.71 | 0 | Myofibroblasts |
| RPL9P9 | 0.41 | 0 | 0 | 0.99 | 0.94 | 0 | Myofibroblasts |
| WFDC1 | 0.41 | 0 | 0 | 0.43 | 0.21 | 0 | Myofibroblasts |
| CENPV | 0.4 | 0 | 0 | 0.33 | 0.15 | 0 | Myofibroblasts |
| NENF | 0.4 | 0 | 0 | 0.66 | 0.46 | 0 | Myofibroblasts |
| ECHDC2 | 0.4 | 0 | 0 | 0.33 | 0.14 | 0 | Myofibroblasts |
| CASC15 | 0.4 | 0 | 0 | 0.25 | 0.09 | 0 | Myofibroblasts |
| POLE4 | 0.4 | 0 | 0 | 0.38 | 0.22 | 0 | Myofibroblasts |
| GADD45B | 0.4 | 0 | 0 | 0.53 | 0.40 | 0 | Myofibroblasts |
| MAP1B | 0.4 | 0 | 0 | 0.38 | 0.22 | 0 | Myofibroblasts |
| AC013461.1 | 0.4 | 0 | 0 | 0.57 | 0.34 | 0 | Myofibroblasts |
| RPL5P34 | 0.4 | 0 | 0 | 0.88 | 0.73 | 0 | Myofibroblasts |
| RPL5P4 | 0.39 | 0 | 0 | 0.63 | 0.43 | 0 | Myofibroblasts |
| ST5 | 0.39 | 0 | 0 | 0.41 | 0.21 | 0 | Myofibroblasts |
| SPON1 | 0.39 | 0 | 0 | 0.35 | 0.14 | 0 | Myofibroblasts |
| LTBP3 | 0.39 | 0 | 0 | 0.34 | 0.17 | 0 | Myofibroblasts |
| CCDC107 | 0.39 | 0 | 0 | 0.42 | 0.24 | 0 | Myofibroblasts |
| RPL37A | 0.39 | 0 | 0 | 1.00 | 0.99 | 0 | Myofibroblasts |
| MSRB3 | 0.38 | 0 | 0 | 0.42 | 0.22 | 0 | Myofibroblasts |
| STX8 | 0.38 | 0 | 0 | 0.31 | 0.16 | 0 | Myofibroblasts |
| RPL5 | 0.37 | 0 | 0 | 0.99 | 0.96 | 0 | Myofibroblasts |
| RPS5 | 0.37 | 0 | 0 | 0.99 | 0.95 | 0 | Myofibroblasts |
| SBDS | 0.37 | 0 | 0 | 0.63 | 0.46 | 0 | Myofibroblasts |
| ACTB | 0.37 | 0 | 0 | 1.00 | 0.99 | 0 | Myofibroblasts |
| LDHA | 0.37 | 0 | 0 | 0.71 | 0.56 | 0 | Myofibroblasts |
| NPM1 | 0.36 | 0 | 0 | 0.95 | 0.87 | 0 | Myofibroblasts |
| ARL4A | 0.36 | 0 | 0 | 0.27 | 0.10 | 0 | Myofibroblasts |
| EEF1B2P6 | 0.36 | 0 | 0 | 0.57 | 0.40 | 0 | Myofibroblasts |
| GNG12 | 0.35 | 0 | 0 | 0.27 | 0.14 | 0 | Myofibroblasts |
| PCSK7 | 0.35 | 0 | 0 | 0.46 | 0.26 | 0 | Myofibroblasts |
| CFL2 | 0.35 | 0 | 0 | 0.34 | 0.18 | 0 | Myofibroblasts |
| FAUP1 | 0.34 | 0 | 0 | 0.77 | 0.63 | 0 | Myofibroblasts |
| RPS21 | 0.34 | 0 | 0 | 0.98 | 0.92 | 0 | Myofibroblasts |
| RBMS3 | 0.34 | 0 | 0 | 0.33 | 0.18 | 0 | Myofibroblasts |
| RPL9 | 0.34 | 0 | 0 | 1.00 | 0.98 | 0 | Myofibroblasts |
| RRAD | 0.34 | 0 | 0 | 0.30 | 0.17 | 0 | Myofibroblasts |
| RCN2 | 0.34 | 0 | 0 | 0.55 | 0.39 | 0 | Myofibroblasts |
| TLN1 | 0.34 | 0 | 0 | 0.45 | 0.29 | 0 | Myofibroblasts |
| SOX4 | 0.34 | 0 | 0 | 0.87 | 0.74 | 0 | Myofibroblasts |
| ZSCAN18 | 0.33 | 0 | 0 | 0.43 | 0.26 | 0 | Myofibroblasts |
| RP11-270C12.3 | 0.33 | 0 | 0 | 0.43 | 0.28 | 0 | Myofibroblasts |
| RP11-475C16.1 | 0.33 | 0 | 0 | 0.91 | 0.81 | 0 | Myofibroblasts |
| CRIP2 | 0.32 | 0 | 0 | 0.83 | 0.63 | 0 | Myofibroblasts |
| RP3-486I3.4 | 0.32 | 0 | 0 | 0.79 | 0.64 | 0 | Myofibroblasts |
| NDUFV2 | 0.32 | 0 | 0 | 0.43 | 0.31 | 0 | Myofibroblasts |
| PTK7 | 0.32 | 0 | 0 | 0.35 | 0.19 | 0 | Myofibroblasts |
| RAB34 | 0.32 | 0 | 0 | 0.53 | 0.37 | 0 | Myofibroblasts |
| RPL8 | 0.31 | 0 | 0 | 1.00 | 0.98 | 0 | Myofibroblasts |
| DBN1 | 0.31 | 0 | 0 | 0.35 | 0.20 | 0 | Myofibroblasts |
| EEF1A1P5 | 0.31 | 0 | 0 | 0.78 | 0.64 | 0 | Myofibroblasts |
| ADAMTS1 | 0.31 | 0 | 0 | 0.31 | 0.18 | 0 | Myofibroblasts |
| SYNE1 | 0.31 | 0 | 0 | 0.29 | 0.15 | 0 | Myofibroblasts |
| RBPMS | 0.31 | 0 | 0 | 0.33 | 0.19 | 0 | Myofibroblasts |
| RP11-393N4.2 | 0.3 | 0 | 0 | 0.76 | 0.64 | 0 | Myofibroblasts |
| MXRA7 | 0.3 | 0 | 0 | 0.27 | 0.13 | 0 | Myofibroblasts |
| RPL7 | 0.3 | 0 | 0 | 1.00 | 0.98 | 0 | Myofibroblasts |
| RPL27A | 0.3 | 0 | 0 | 1.00 | 0.99 | 0 | Myofibroblasts |
| FOXN3 | 0.3 | 0 | 0 | 0.34 | 0.21 | 0 | Myofibroblasts |
| RPL18 | 0.3 | 0 | 0 | 1.00 | 0.98 | 0 | Myofibroblasts |
| TCEAL3 | 0.3 | 0 | 0 | 0.31 | 0.18 | 0 | Myofibroblasts |
| RBP1 | 0.3 | 0 | 0 | 0.52 | 0.34 | 0 | Myofibroblasts |
| DAPK3 | 0.3 | 0 | 0 | 0.31 | 0.17 | 0 | Myofibroblasts |
| JUND | 0.29 | 0 | 0 | 0.41 | 0.29 | 0 | Myofibroblasts |
| KANK2 | 0.29 | 0 | 0 | 0.50 | 0.32 | 0 | Myofibroblasts |
| RPL10A | 0.29 | 0 | 0 | 0.99 | 0.96 | 0 | Myofibroblasts |
| RPS24 | 0.29 | 0 | 0 | 1.00 | 0.98 | 0 | Myofibroblasts |
| TERF1 | 0.29 | 0 | 0 | 0.34 | 0.21 | 0 | Myofibroblasts |
| PRDM6 | 0.29 | 0 | 0 | 0.27 | 0.13 | 0 | Myofibroblasts |
| EEF1B2P3 | 0.29 | 0 | 0 | 0.35 | 0.23 | 0 | Myofibroblasts |
| MYH9 | 0.29 | 0 | 0 | 0.57 | 0.43 | 0 | Myofibroblasts |
| RPL21 | 0.28 | 0 | 0 | 1.00 | 0.99 | 0 | Myofibroblasts |
| RPL7P1 | 0.28 | 0 | 0 | 0.80 | 0.66 | 0 | Myofibroblasts |
| DPYSL3 | 0.28 | 0 | 0 | 0.28 | 0.14 | 0 | Myofibroblasts |
| HSP90AB3P | 0.28 | 0 | 0 | 0.61 | 0.49 | 0 | Myofibroblasts |
| RP11-425L10.1 | 0.28 | 0 | 0 | 0.92 | 0.83 | 0 | Myofibroblasts |
| RPL3P4 | 0.28 | 0 | 0 | 0.98 | 0.94 | 0 | Myofibroblasts |
| IMPDH2 | 0.28 | 0 | 0 | 0.54 | 0.38 | 0 | Myofibroblasts |
| RPL10AP6 | 0.28 | 0 | 0 | 0.85 | 0.76 | 0 | Myofibroblasts |
| DRAP1 | 0.28 | 0 | 0 | 0.63 | 0.48 | 0 | Myofibroblasts |
| AC090498.1 | 0.27 | 0 | 0 | 0.98 | 0.93 | 0 | Myofibroblasts |
| RPS10 | 0.27 | 0 | 0 | 0.95 | 0.89 | 0 | Myofibroblasts |
| KLHL23 | 0.27 | 0 | 0 | 0.32 | 0.18 | 0 | Myofibroblasts |
| AEBP1 | 0.27 | 0 | 0 | 0.32 | 0.19 | 0 | Myofibroblasts |
| PLEKHO1 | 0.27 | 0 | 0 | 0.27 | 0.15 | 0 | Myofibroblasts |
| PLS3 | 0.27 | 0 | 0 | 0.54 | 0.41 | 0 | Myofibroblasts |
| SNRPN | 0.27 | 0 | 0 | 0.61 | 0.46 | 0 | Myofibroblasts |
| VDAC2 | 0.27 | 0 | 0 | 0.65 | 0.52 | 0 | Myofibroblasts |
| SNHG14 | 0.27 | 0 | 0 | 0.26 | 0.13 | 0 | Myofibroblasts |
| RP3-417G15.1 | 0.27 | 0 | 0 | 0.87 | 0.75 | 0 | Myofibroblasts |
| GABARAPL2 | 0.27 | 0 | 0 | 0.75 | 0.65 | 0 | Myofibroblasts |
| MTCH1 | 0.27 | 0 | 0 | 0.62 | 0.50 | 0 | Myofibroblasts |
| RHEB | 0.26 | 0 | 0 | 0.56 | 0.47 | 0 | Myofibroblasts |
| SLIRP | 0.26 | 0 | 0 | 0.58 | 0.48 | 0 | Myofibroblasts |
| TNS1 | 0.26 | 0 | 0 | 0.35 | 0.22 | 0 | Myofibroblasts |
| RPL5P1 | 0.26 | 0 | 0 | 0.47 | 0.34 | 0 | Myofibroblasts |
| RPS4XP6 | 0.26 | 0 | 0 | 0.88 | 0.76 | 0 | Myofibroblasts |
| MT2A | 0.26 | 0 | 0 | 0.76 | 0.61 | 0 | Myofibroblasts |
| MLXIP | 0.26 | 0 | 0 | 0.30 | 0.16 | 0 | Myofibroblasts |
| RPL24 | 0.26 | 0 | 0 | 0.98 | 0.96 | 0 | Myofibroblasts |
| ATP1B1 | 0.26 | 0 | 0 | 0.36 | 0.20 | 0 | Myofibroblasts |
| CHCHD2 | 0.26 | 0 | 0 | 0.91 | 0.82 | 0 | Myofibroblasts |
| RPS2P5 | 0.25 | 0 | 0 | 0.86 | 0.72 | 0 | Myofibroblasts |
| AHI1 | 0.25 | 0 | 0 | 0.26 | 0.15 | 0 | Myofibroblasts |
| COX7B | 0.25 | 0 | 0 | 0.72 | 0.61 | 0 | Myofibroblasts |
| TUBB4B | 0.25 | 0 | 0 | 0.52 | 0.39 | 0 | Myofibroblasts |
| RPL41 | 0.25 | 0 | 0 | 1.00 | 1.00 | 0 | Myofibroblasts |
| RPS12 | 0.25 | 0 | 0 | 1.00 | 0.99 | 0 | Myofibroblasts |
| RPL3 | 0.25 | 0 | 0 | 1.00 | 0.99 | 0 | Myofibroblasts |
| RPL24P8 | 0.25 | 0 | 0 | 0.75 | 0.65 | 0 | Myofibroblasts |
| EIF5 | 0.25 | 0 | 0 | 0.59 | 0.49 | 0 | Myofibroblasts |
| EEF1A1 | 0.25 | 0 | 0 | 1.00 | 1.00 | 0 | Myofibroblasts |
| NDUFB9 | 0.25 | 0 | 0 | 0.64 | 0.51 | 0 | Myofibroblasts |
| AXL | 0.25 | 0 | 0 | 0.34 | 0.21 | 0 | Myofibroblasts |
| RPS4X | 0.25 | 0 | 0 | 1.00 | 0.99 | 0 | Myofibroblasts |
| CFD | 2.41 | 0 | 0 | 0.97 | 0.17 | 1 | Matrix Fibroblasts |
| ADH1B | 2.38 | 0 | 0 | 0.94 | 0.15 | 1 | Matrix Fibroblasts |
| LUM | 2.37 | 0 | 0 | 0.97 | 0.28 | 1 | Matrix Fibroblasts |
| GPC3 | 2.11 | 0 | 0 | 0.93 | 0.14 | 1 | Matrix Fibroblasts |
| TCF21 | 2.1 | 0 | 0 | 0.93 | 0.20 | 1 | Matrix Fibroblasts |
| SEPP1 | 1.98 | 0 | 0 | 0.96 | 0.50 | 1 | Matrix Fibroblasts |
| FN1 | 1.91 | 0 | 0 | 0.95 | 0.37 | 1 | Matrix Fibroblasts |
| RARRES2 | 1.75 | 0 | 0 | 0.97 | 0.44 | 1 | Matrix Fibroblasts |
| MFAP4 | 1.72 | 0 | 0 | 0.98 | 0.42 | 1 | Matrix Fibroblasts |
| WNT2 | 1.7 | 0 | 0 | 0.70 | 0.02 | 1 | Matrix Fibroblasts |
| LIMCH1 | 1.66 | 0 | 0 | 0.77 | 0.12 | 1 | Matrix Fibroblasts |
| PLEKHH2 | 1.63 | 0 | 0 | 0.74 | 0.13 | 1 | Matrix Fibroblasts |
| G0S2 | 1.6 | 0 | 0 | 0.63 | 0.11 | 1 | Matrix Fibroblasts |
| COL6A3 | 1.56 | 0 | 0 | 0.91 | 0.30 | 1 | Matrix Fibroblasts |
| FBLN1 | 1.56 | 0 | 0 | 0.94 | 0.20 | 1 | Matrix Fibroblasts |
| RARRES1 | 1.55 | 0 | 0 | 0.52 | 0.12 | 1 | Matrix Fibroblasts |
| APOE | 1.55 | 0 | 0 | 0.83 | 0.23 | 1 | Matrix Fibroblasts |
| INMT | 1.54 | 0 | 0 | 0.69 | 0.14 | 1 | Matrix Fibroblasts |
| C7 | 1.51 | 0 | 0 | 0.77 | 0.07 | 1 | Matrix Fibroblasts |
| VEGFD | 1.45 | 0 | 0 | 0.56 | 0.04 | 1 | Matrix Fibroblasts |
| A2M | 1.41 | 0 | 0 | 0.74 | 0.60 | 1 | Matrix Fibroblasts |
| LMO4 | 1.37 | 0 | 0 | 0.80 | 0.31 | 1 | Matrix Fibroblasts |
| OLFML3 | 1.34 | 0 | 0 | 0.66 | 0.13 | 1 | Matrix Fibroblasts |
| LBH | 1.33 | 0 | 0 | 0.84 | 0.33 | 1 | Matrix Fibroblasts |
| DST | 1.3 | 0 | 0 | 0.86 | 0.36 | 1 | Matrix Fibroblasts |
| CA3 | 1.3 | 0 | 0 | 0.37 | 0.02 | 1 | Matrix Fibroblasts |
| PIEZO2 | 1.26 | 0 | 0 | 0.55 | 0.04 | 1 | Matrix Fibroblasts |
| MACF1 | 1.26 | 0 | 0 | 0.79 | 0.31 | 1 | Matrix Fibroblasts |
| NBL1 | 1.25 | 0 | 0 | 0.85 | 0.29 | 1 | Matrix Fibroblasts |
| DCN | 1.24 | 0 | 0 | 0.97 | 0.47 | 1 | Matrix Fibroblasts |
| MEOX2 | 1.21 | 0 | 0 | 0.56 | 0.07 | 1 | Matrix Fibroblasts |
| ABCA8 | 1.18 | 0 | 0 | 0.53 | 0.06 | 1 | Matrix Fibroblasts |
| CDH11 | 1.18 | 0 | 0 | 0.66 | 0.13 | 1 | Matrix Fibroblasts |
| CAMK2N1 | 1.13 | 0 | 0 | 0.65 | 0.15 | 1 | Matrix Fibroblasts |
| C1R | 1.11 | 0 | 0 | 0.59 | 0.19 | 1 | Matrix Fibroblasts |
| RSPO2 | 1.11 | 0 | 0 | 0.45 | 0.01 | 1 | Matrix Fibroblasts |
| MMP2 | 1.09 | 0 | 0 | 0.82 | 0.27 | 1 | Matrix Fibroblasts |
| PMP22 | 1.08 | 0 | 0 | 0.80 | 0.43 | 1 | Matrix Fibroblasts |
| LITAF | 1.05 | 0 | 0 | 0.71 | 0.43 | 1 | Matrix Fibroblasts |
| GYG1 | 1.05 | 0 | 0 | 0.57 | 0.19 | 1 | Matrix Fibroblasts |
| LRRC17 | 1.04 | 0 | 0 | 0.49 | 0.07 | 1 | Matrix Fibroblasts |
| FGFR4 | 1.03 | 0 | 0 | 0.45 | 0.02 | 1 | Matrix Fibroblasts |
| NR2F1 | 1.03 | 0 | 0 | 0.67 | 0.21 | 1 | Matrix Fibroblasts |
| ABCA6 | 1.03 | 0 | 0 | 0.41 | 0.03 | 1 | Matrix Fibroblasts |
| EMILIN1 | 1.02 | 0 | 0 | 0.63 | 0.18 | 1 | Matrix Fibroblasts |
| METTL7A | 1.01 | 0 | 0 | 0.46 | 0.06 | 1 | Matrix Fibroblasts |
| CEBPD | 1 | 0 | 0 | 0.58 | 0.25 | 1 | Matrix Fibroblasts |
| CCBE1 | 1 | 0 | 0 | 0.48 | 0.06 | 1 | Matrix Fibroblasts |
| SLC40A1 | 0.98 | 0 | 0 | 0.65 | 0.19 | 1 | Matrix Fibroblasts |
| COL6A1 | 0.96 | 0 | 0 | 0.87 | 0.48 | 1 | Matrix Fibroblasts |
| ZYX | 0.96 | 0 | 0 | 0.66 | 0.28 | 1 | Matrix Fibroblasts |
| SMPDL3A | 0.96 | 0 | 0 | 0.42 | 0.05 | 1 | Matrix Fibroblasts |
| COLEC12 | 0.95 | 0 | 0 | 0.48 | 0.08 | 1 | Matrix Fibroblasts |
| TNC | 0.94 | 0 | 0 | 0.44 | 0.13 | 1 | Matrix Fibroblasts |
| FMO2 | 0.92 | 0 | 0 | 0.37 | 0.03 | 1 | Matrix Fibroblasts |
| OGN | 0.91 | 0 | 0 | 0.49 | 0.09 | 1 | Matrix Fibroblasts |
| FBLN5 | 0.91 | 0 | 0 | 0.53 | 0.11 | 1 | Matrix Fibroblasts |
| C1S | 0.91 | 0 | 0 | 0.56 | 0.16 | 1 | Matrix Fibroblasts |
| MDK | 0.9 | 0 | 0 | 0.84 | 0.49 | 1 | Matrix Fibroblasts |
| SLIT2 | 0.89 | 0 | 0 | 0.46 | 0.12 | 1 | Matrix Fibroblasts |
| ANGPT1 | 0.88 | 0 | 0 | 0.46 | 0.10 | 1 | Matrix Fibroblasts |
| PPP1R3C | 0.88 | 0 | 0 | 0.38 | 0.10 | 1 | Matrix Fibroblasts |
| RGS2 | 0.84 | 0 | 0 | 0.62 | 0.26 | 1 | Matrix Fibroblasts |
| ZFP36L2 | 0.84 | 0 | 0 | 0.69 | 0.47 | 1 | Matrix Fibroblasts |
| VCAN | 0.84 | 0 | 0 | 0.76 | 0.28 | 1 | Matrix Fibroblasts |
| MYH10 | 0.84 | 0 | 0 | 0.53 | 0.23 | 1 | Matrix Fibroblasts |
| PPP1R14A | 0.84 | 0 | 0 | 0.73 | 0.38 | 1 | Matrix Fibroblasts |
| GDF10 | 0.83 | 0 | 0 | 0.34 | 0.04 | 1 | Matrix Fibroblasts |
| FGF7 | 0.83 | 0 | 0 | 0.38 | 0.06 | 1 | Matrix Fibroblasts |
| COL1A2 | 0.82 | 0 | 0 | 0.98 | 0.59 | 1 | Matrix Fibroblasts |
| PLPP1 | 0.82 | 0 | 0 | 0.52 | 0.18 | 1 | Matrix Fibroblasts |
| LAMA4 | 0.79 | 0 | 0 | 0.54 | 0.21 | 1 | Matrix Fibroblasts |
| LTBP4 | 0.79 | 0 | 0 | 0.61 | 0.25 | 1 | Matrix Fibroblasts |
| POSTN | 0.79 | 0 | 0 | 0.49 | 0.17 | 1 | Matrix Fibroblasts |
| SNHG6 | 0.77 | 0 | 0 | 0.76 | 0.47 | 1 | Matrix Fibroblasts |
| MEST | 0.77 | 0 | 0 | 0.74 | 0.32 | 1 | Matrix Fibroblasts |
| SCN7A | 0.77 | 0 | 0 | 0.38 | 0.07 | 1 | Matrix Fibroblasts |
| CXCL12 | 0.75 | 0 | 0 | 0.58 | 0.19 | 1 | Matrix Fibroblasts |
| LOXL1 | 0.75 | 0 | 0 | 0.39 | 0.09 | 1 | Matrix Fibroblasts |
| COL13A1 | 0.74 | 0 | 0 | 0.33 | 0.06 | 1 | Matrix Fibroblasts |
| MAF | 0.74 | 0 | 0 | 0.38 | 0.08 | 1 | Matrix Fibroblasts |
| PKN2 | 0.74 | 0 | 0 | 0.42 | 0.15 | 1 | Matrix Fibroblasts |
| DKK3 | 0.73 | 0 | 0 | 0.59 | 0.24 | 1 | Matrix Fibroblasts |
| PLXDC2 | 0.73 | 0 | 0 | 0.43 | 0.11 | 1 | Matrix Fibroblasts |
| SERPING1 | 0.73 | 0 | 0 | 0.69 | 0.35 | 1 | Matrix Fibroblasts |
| FAM213A | 0.73 | 0 | 0 | 0.45 | 0.17 | 1 | Matrix Fibroblasts |
| CHST2 | 0.73 | 0 | 0 | 0.33 | 0.07 | 1 | Matrix Fibroblasts |
| GSTM5 | 0.72 | 0 | 0 | 0.40 | 0.09 | 1 | Matrix Fibroblasts |
| COL5A2 | 0.72 | 0 | 0 | 0.48 | 0.18 | 1 | Matrix Fibroblasts |
| SLC38A5 | 0.72 | 0 | 0 | 0.32 | 0.03 | 1 | Matrix Fibroblasts |
| RGS3 | 0.71 | 0 | 0 | 0.39 | 0.26 | 1 | Matrix Fibroblasts |
| MRC2 | 0.71 | 0 | 0 | 0.33 | 0.08 | 1 | Matrix Fibroblasts |
| CSRP1 | 0.7 | 0 | 0 | 0.86 | 0.56 | 1 | Matrix Fibroblasts |
| MOXD1 | 0.7 | 0 | 0 | 0.31 | 0.03 | 1 | Matrix Fibroblasts |
| ITGA8 | 0.69 | 0 | 0 | 0.39 | 0.09 | 1 | Matrix Fibroblasts |
| COL6A2 | 0.69 | 0 | 0 | 0.88 | 0.58 | 1 | Matrix Fibroblasts |
| SPINT2 | 0.69 | 0 | 0 | 0.41 | 0.10 | 1 | Matrix Fibroblasts |
| LDOC1 | 0.68 | 0 | 0 | 0.34 | 0.09 | 1 | Matrix Fibroblasts |
| KCNQ1OT1 | 0.68 | 0 | 0 | 0.53 | 0.27 | 1 | Matrix Fibroblasts |
| CD302 | 0.68 | 0 | 0 | 0.42 | 0.14 | 1 | Matrix Fibroblasts |
| LRP1 | 0.68 | 0 | 0 | 0.43 | 0.15 | 1 | Matrix Fibroblasts |
| NPNT | 0.68 | 0 | 0 | 0.33 | 0.07 | 1 | Matrix Fibroblasts |
| PCOLCE | 0.68 | 0 | 0 | 0.66 | 0.26 | 1 | Matrix Fibroblasts |
| PDLIM2 | 0.68 | 0 | 0 | 0.46 | 0.18 | 1 | Matrix Fibroblasts |
| SNHG8 | 0.67 | 0 | 0 | 0.77 | 0.53 | 1 | Matrix Fibroblasts |
| AOC3 | 0.67 | 0 | 0 | 0.63 | 0.29 | 1 | Matrix Fibroblasts |
| EPHX1 | 0.67 | 0 | 0 | 0.50 | 0.23 | 1 | Matrix Fibroblasts |
| LSAMP | 0.67 | 0 | 0 | 0.29 | 0.03 | 1 | Matrix Fibroblasts |
| FAT3 | 0.67 | 0 | 0 | 0.27 | 0.03 | 1 | Matrix Fibroblasts |
| TGM2 | 0.66 | 0 | 0 | 0.56 | 0.37 | 1 | Matrix Fibroblasts |
| ALDH1A1 | 0.66 | 0 | 0 | 0.35 | 0.09 | 1 | Matrix Fibroblasts |
| RBP1 | 0.66 | 0 | 0 | 0.60 | 0.32 | 1 | Matrix Fibroblasts |
| SLC1A5 | 0.66 | 0 | 0 | 0.33 | 0.06 | 1 | Matrix Fibroblasts |
| GNAI1 | 0.65 | 0 | 0 | 0.42 | 0.15 | 1 | Matrix Fibroblasts |
| RN7SK | 0.65 | 0 | 0 | 0.43 | 0.27 | 1 | Matrix Fibroblasts |
| FHL1 | 0.65 | 0 | 0 | 0.85 | 0.53 | 1 | Matrix Fibroblasts |
| PTPRD | 0.64 | 0 | 0 | 0.29 | 0.06 | 1 | Matrix Fibroblasts |
| MMP23B | 0.64 | 0 | 0 | 0.39 | 0.10 | 1 | Matrix Fibroblasts |
| TMEM98 | 0.64 | 0 | 0 | 0.58 | 0.29 | 1 | Matrix Fibroblasts |
| CNN2 | 0.63 | 0 | 0 | 0.46 | 0.19 | 1 | Matrix Fibroblasts |
| EIF3E | 0.63 | 0 | 0 | 0.86 | 0.76 | 1 | Matrix Fibroblasts |
| ASPA | 0.63 | 0 | 0 | 0.27 | 0.02 | 1 | Matrix Fibroblasts |
| PRKCB | 0.62 | 0 | 0 | 0.28 | 0.05 | 1 | Matrix Fibroblasts |
| COL3A1 | 0.62 | 0 | 0 | 0.94 | 0.53 | 1 | Matrix Fibroblasts |
| ENPP2 | 0.62 | 0 | 0 | 0.37 | 0.10 | 1 | Matrix Fibroblasts |
| EPB41L4A-AS1 | 0.61 | 0 | 0 | 0.50 | 0.24 | 1 | Matrix Fibroblasts |
| SNCA | 0.61 | 0 | 0 | 0.29 | 0.06 | 1 | Matrix Fibroblasts |
| PEG10 | 0.6 | 0 | 0 | 0.25 | 0.06 | 1 | Matrix Fibroblasts |
| MMP14 | 0.59 | 0 | 0 | 0.44 | 0.20 | 1 | Matrix Fibroblasts |
| QSOX1 | 0.58 | 0 | 0 | 0.42 | 0.19 | 1 | Matrix Fibroblasts |
| GYPC | 0.58 | 0 | 0 | 0.55 | 0.29 | 1 | Matrix Fibroblasts |
| LAMC1 | 0.58 | 0 | 0 | 0.42 | 0.19 | 1 | Matrix Fibroblasts |
| FHOD1 | 0.58 | 0 | 0 | 0.31 | 0.07 | 1 | Matrix Fibroblasts |
| GCSH | 0.57 | 0 | 0 | 0.43 | 0.20 | 1 | Matrix Fibroblasts |
| MDFI | 0.57 | 0 | 0 | 0.40 | 0.16 | 1 | Matrix Fibroblasts |
| BMP4 | 0.57 | 0 | 0 | 0.31 | 0.07 | 1 | Matrix Fibroblasts |
| CDO1 | 0.57 | 0 | 0 | 0.27 | 0.05 | 1 | Matrix Fibroblasts |
| CITED2 | 0.57 | 0 | 0 | 0.44 | 0.21 | 1 | Matrix Fibroblasts |
| SLC44A1 | 0.57 | 0 | 0 | 0.38 | 0.16 | 1 | Matrix Fibroblasts |
| COL1A1 | 0.56 | 0 | 0 | 0.96 | 0.56 | 1 | Matrix Fibroblasts |
| BNIP3L | 0.56 | 0 | 0 | 0.59 | 0.36 | 1 | Matrix Fibroblasts |
| RP11-452N17.1 | 0.56 | 0 | 0 | 0.55 | 0.36 | 1 | Matrix Fibroblasts |
| GJA5 | 0.55 | 0 | 0 | 0.25 | 0.04 | 1 | Matrix Fibroblasts |
| SNAI2 | 0.55 | 0 | 0 | 0.31 | 0.10 | 1 | Matrix Fibroblasts |
| COL5A1 | 0.55 | 0 | 0 | 0.42 | 0.20 | 1 | Matrix Fibroblasts |
| COL14A1 | 0.54 | 0 | 0 | 0.30 | 0.09 | 1 | Matrix Fibroblasts |
| FXYD1 | 0.54 | 0 | 0 | 0.48 | 0.24 | 1 | Matrix Fibroblasts |
| OLFML2A | 0.54 | 0 | 0 | 0.28 | 0.09 | 1 | Matrix Fibroblasts |
| LGALS3BP | 0.53 | 0 | 0 | 0.41 | 0.17 | 1 | Matrix Fibroblasts |
| ALDH9A1 | 0.53 | 0 | 0 | 0.43 | 0.21 | 1 | Matrix Fibroblasts |
| DYNC2LI1 | 0.53 | 0 | 0 | 0.29 | 0.09 | 1 | Matrix Fibroblasts |
| RUNX1T1 | 0.53 | 0 | 0 | 0.37 | 0.15 | 1 | Matrix Fibroblasts |
| FMOD | 0.53 | 0 | 0 | 0.27 | 0.07 | 1 | Matrix Fibroblasts |
| PRDX6 | 0.52 | 0 | 0 | 0.68 | 0.53 | 1 | Matrix Fibroblasts |
| MAOB | 0.52 | 0 | 0 | 0.26 | 0.06 | 1 | Matrix Fibroblasts |
| ENAH | 0.52 | 0 | 0 | 0.49 | 0.27 | 1 | Matrix Fibroblasts |
| PLAGL1 | 0.51 | 0 | 0 | 0.30 | 0.11 | 1 | Matrix Fibroblasts |
| PLPP3 | 0.51 | 0 | 0 | 0.50 | 0.26 | 1 | Matrix Fibroblasts |
| QPRT | 0.51 | 0 | 0 | 0.29 | 0.09 | 1 | Matrix Fibroblasts |
| HIF3A | 0.51 | 0 | 0 | 0.29 | 0.12 | 1 | Matrix Fibroblasts |
| CELF2 | 0.5 | 0 | 0 | 0.41 | 0.21 | 1 | Matrix Fibroblasts |
| MYO10 | 0.5 | 0 | 0 | 0.30 | 0.10 | 1 | Matrix Fibroblasts |
| PTMS | 0.5 | 0 | 0 | 0.91 | 0.75 | 1 | Matrix Fibroblasts |
| COL21A1 | 0.5 | 0 | 0 | 0.32 | 0.12 | 1 | Matrix Fibroblasts |
| SMS | 0.49 | 0 | 0 | 0.40 | 0.20 | 1 | Matrix Fibroblasts |
| PEBP1 | 0.48 | 0 | 0 | 0.81 | 0.65 | 1 | Matrix Fibroblasts |
| MAST4 | 0.48 | 0 | 0 | 0.32 | 0.14 | 1 | Matrix Fibroblasts |
| TMEM261 | 0.48 | 0 | 0 | 0.45 | 0.26 | 1 | Matrix Fibroblasts |
| MXRA8 | 0.48 | 0 | 0 | 0.51 | 0.27 | 1 | Matrix Fibroblasts |
| NMRK1 | 0.48 | 0 | 0 | 0.26 | 0.08 | 1 | Matrix Fibroblasts |
| SOD2 | 0.48 | 0 | 0 | 0.29 | 0.19 | 1 | Matrix Fibroblasts |
| TOMM7 | 0.47 | 0 | 0 | 0.93 | 0.84 | 1 | Matrix Fibroblasts |
| LURAP1L | 0.47 | 0 | 0 | 0.39 | 0.16 | 1 | Matrix Fibroblasts |
| FBLIM1 | 0.46 | 0 | 0 | 0.33 | 0.16 | 1 | Matrix Fibroblasts |
| HACD3 | 0.46 | 0 | 0 | 0.38 | 0.20 | 1 | Matrix Fibroblasts |
| FILIP1 | 0.46 | 0 | 0 | 0.34 | 0.16 | 1 | Matrix Fibroblasts |
| PTGER1 | 0.46 | 0 | 0 | 0.26 | 0.07 | 1 | Matrix Fibroblasts |
| GULP1 | 0.46 | 0 | 0 | 0.28 | 0.11 | 1 | Matrix Fibroblasts |
| CIRBP | 0.46 | 0 | 0 | 0.95 | 0.85 | 1 | Matrix Fibroblasts |
| RND3 | 0.46 | 0 | 0 | 0.34 | 0.17 | 1 | Matrix Fibroblasts |
| FNIP1 | 0.45 | 0 | 0 | 0.35 | 0.20 | 1 | Matrix Fibroblasts |
| MSRB2 | 0.45 | 0 | 0 | 0.38 | 0.20 | 1 | Matrix Fibroblasts |
| PPIB | 0.44 | 0 | 0 | 0.82 | 0.70 | 1 | Matrix Fibroblasts |
| EDNRA | 0.44 | 0 | 0 | 0.39 | 0.21 | 1 | Matrix Fibroblasts |
| SAT1 | 0.44 | 0 | 0 | 0.75 | 0.53 | 1 | Matrix Fibroblasts |
| PLTP | 0.44 | 0 | 0 | 0.30 | 0.12 | 1 | Matrix Fibroblasts |
| NREP | 0.43 | 0 | 0 | 0.51 | 0.36 | 1 | Matrix Fibroblasts |
| FZD1 | 0.43 | 0 | 0 | 0.26 | 0.09 | 1 | Matrix Fibroblasts |
| LPL | 0.43 | 0 | 0 | 0.41 | 0.15 | 1 | Matrix Fibroblasts |
| SOD3 | 0.43 | 0 | 0 | 0.69 | 0.38 | 1 | Matrix Fibroblasts |
| VSTM4 | 0.43 | 0 | 0 | 0.26 | 0.10 | 1 | Matrix Fibroblasts |
| CCDC8 | 0.42 | 0 | 0 | 0.26 | 0.09 | 1 | Matrix Fibroblasts |
| S1PR1 | 0.42 | 0 | 0 | 0.28 | 0.12 | 1 | Matrix Fibroblasts |
| TM2D2 | 0.42 | 0 | 0 | 0.31 | 0.16 | 1 | Matrix Fibroblasts |
| SNRPN | 0.41 | 0 | 0 | 0.60 | 0.46 | 1 | Matrix Fibroblasts |
| RNF146 | 0.41 | 0 | 0 | 0.33 | 0.18 | 1 | Matrix Fibroblasts |
| NEXN | 0.41 | 0 | 0 | 0.39 | 0.20 | 1 | Matrix Fibroblasts |
| CTSC | 0.4 | 0 | 0 | 0.38 | 0.21 | 1 | Matrix Fibroblasts |
| GSTM3 | 0.4 | 0 | 0 | 0.36 | 0.18 | 1 | Matrix Fibroblasts |
| NDN | 0.4 | 0 | 0 | 0.50 | 0.32 | 1 | Matrix Fibroblasts |
| CCL2 | 0.4 | 0 | 0 | 0.36 | 0.14 | 1 | Matrix Fibroblasts |
| MFAP2 | 0.4 | 0 | 0 | 0.42 | 0.26 | 1 | Matrix Fibroblasts |
| RP11-693N9.2 | 0.4 | 0 | 0 | 0.31 | 0.14 | 1 | Matrix Fibroblasts |
| CLEC11A | 0.39 | 0 | 0 | 0.36 | 0.18 | 1 | Matrix Fibroblasts |
| LAPTM4A | 0.39 | 0 | 0 | 0.94 | 0.83 | 1 | Matrix Fibroblasts |
| MYL6B | 0.39 | 0 | 0 | 0.40 | 0.25 | 1 | Matrix Fibroblasts |
| AKAP9 | 0.39 | 0 | 0 | 0.53 | 0.38 | 1 | Matrix Fibroblasts |
| EFEMP2 | 0.39 | 0 | 0 | 0.53 | 0.34 | 1 | Matrix Fibroblasts |
| NR2F1-AS1 | 0.39 | 0 | 0 | 0.26 | 0.09 | 1 | Matrix Fibroblasts |
| RPL11 | 0.39 | 0 | 0 | 1.00 | 0.99 | 1 | Matrix Fibroblasts |
| HSPB6 | 0.38 | 0 | 0 | 0.28 | 0.13 | 1 | Matrix Fibroblasts |
| FSTL1 | 0.38 | 0 | 0 | 0.60 | 0.41 | 1 | Matrix Fibroblasts |
| BTG1 | 0.38 | 0 | 0 | 0.81 | 0.66 | 1 | Matrix Fibroblasts |
| SEC62 | 0.38 | 0 | 0 | 0.82 | 0.69 | 1 | Matrix Fibroblasts |
| NFIA | 0.38 | 0 | 0 | 0.50 | 0.36 | 1 | Matrix Fibroblasts |
| NUPR1 | 0.37 | 0 | 0 | 0.69 | 0.47 | 1 | Matrix Fibroblasts |
| C5orf15 | 0.37 | 0 | 0 | 0.37 | 0.22 | 1 | Matrix Fibroblasts |
| TRIP10 | 0.36 | 0 | 0 | 0.31 | 0.16 | 1 | Matrix Fibroblasts |
| C7orf73 | 0.36 | 0 | 0 | 0.44 | 0.31 | 1 | Matrix Fibroblasts |
| BAG2 | 0.36 | 0 | 0 | 0.31 | 0.14 | 1 | Matrix Fibroblasts |
| 11-Sep | 0.36 | 0 | 0 | 0.45 | 0.29 | 1 | Matrix Fibroblasts |
| MEG3 | 0.35 | 0 | 0 | 0.64 | 0.41 | 1 | Matrix Fibroblasts |
| CREG1 | 0.35 | 0 | 0 | 0.29 | 0.16 | 1 | Matrix Fibroblasts |
| CYBRD1 | 0.35 | 0 | 0 | 0.28 | 0.14 | 1 | Matrix Fibroblasts |
| LRRC75A-AS1 | 0.35 | 0 | 0 | 0.94 | 0.86 | 1 | Matrix Fibroblasts |
| FAM127A | 0.35 | 0 | 0 | 0.49 | 0.35 | 1 | Matrix Fibroblasts |
| MAGED1 | 0.35 | 0 | 0 | 0.30 | 0.18 | 1 | Matrix Fibroblasts |
| NRP1 | 0.35 | 0 | 0 | 0.32 | 0.18 | 1 | Matrix Fibroblasts |
| PLSCR4 | 0.35 | 0 | 0 | 0.30 | 0.16 | 1 | Matrix Fibroblasts |
| TMEM9 | 0.35 | 0 | 0 | 0.31 | 0.17 | 1 | Matrix Fibroblasts |
| RAB13 | 0.34 | 0 | 0 | 0.58 | 0.47 | 1 | Matrix Fibroblasts |
| OSTC | 0.34 | 0 | 0 | 0.63 | 0.51 | 1 | Matrix Fibroblasts |
| SRI | 0.34 | 0 | 0 | 0.51 | 0.36 | 1 | Matrix Fibroblasts |
| DYNC1I2 | 0.34 | 0 | 0 | 0.64 | 0.58 | 1 | Matrix Fibroblasts |
| RPS20 | 0.34 | 0 | 0 | 0.99 | 0.97 | 1 | Matrix Fibroblasts |
| PTK7 | 0.34 | 0 | 0 | 0.32 | 0.19 | 1 | Matrix Fibroblasts |
| ST13 | 0.34 | 0 | 0 | 0.70 | 0.63 | 1 | Matrix Fibroblasts |
| ADAM9 | 0.34 | 0 | 0 | 0.26 | 0.15 | 1 | Matrix Fibroblasts |
| AKAP12 | 0.34 | 0 | 0 | 0.39 | 0.20 | 1 | Matrix Fibroblasts |
| CAPZB | 0.33 | 0 | 0 | 0.73 | 0.62 | 1 | Matrix Fibroblasts |
| HMGN1 | 0.33 | 0 | 0 | 0.72 | 0.60 | 1 | Matrix Fibroblasts |
| FOXO3 | 0.33 | 0 | 0 | 0.32 | 0.20 | 1 | Matrix Fibroblasts |
| ZNF106 | 0.33 | 0 | 0 | 0.26 | 0.13 | 1 | Matrix Fibroblasts |
| ADH5 | 0.33 | 0 | 0 | 0.45 | 0.33 | 1 | Matrix Fibroblasts |
| RRBP1 | 0.33 | 0 | 0 | 0.38 | 0.32 | 1 | Matrix Fibroblasts |
| ECHDC2 | 0.32 | 0 | 0 | 0.27 | 0.15 | 1 | Matrix Fibroblasts |
| GSTA4 | 0.32 | 0 | 0 | 0.26 | 0.15 | 1 | Matrix Fibroblasts |
| IL6ST | 0.32 | 0 | 0 | 0.37 | 0.25 | 1 | Matrix Fibroblasts |
| CCDC80 | 0.32 | 0 | 0 | 0.31 | 0.23 | 1 | Matrix Fibroblasts |
| NUCKS1 | 0.32 | 0 | 0 | 0.79 | 0.71 | 1 | Matrix Fibroblasts |
| FBN1 | 0.32 | 0 | 0 | 0.28 | 0.14 | 1 | Matrix Fibroblasts |
| LIMA1 | 0.32 | 0 | 0 | 0.32 | 0.20 | 1 | Matrix Fibroblasts |
| FAM43A | 0.32 | 0 | 0 | 0.26 | 0.13 | 1 | Matrix Fibroblasts |
| NPM1 | 0.32 | 0 | 0 | 0.93 | 0.87 | 1 | Matrix Fibroblasts |
| ARID5B | 0.31 | 0 | 0 | 0.40 | 0.28 | 1 | Matrix Fibroblasts |
| ACTG1 | 0.31 | 0 | 0 | 0.98 | 0.95 | 1 | Matrix Fibroblasts |
| RPL7AP66 | 0.31 | 0 | 0 | 0.54 | 0.44 | 1 | Matrix Fibroblasts |
| CEBPB | 0.31 | 0 | 0 | 0.41 | 0.29 | 1 | Matrix Fibroblasts |
| CDKN1C | 0.31 | 0 | 0 | 0.32 | 0.19 | 1 | Matrix Fibroblasts |
| ZFAS1 | 0.31 | 0 | 0 | 0.70 | 0.57 | 1 | Matrix Fibroblasts |
| C6orf48 | 0.3 | 0 | 0 | 0.71 | 0.58 | 1 | Matrix Fibroblasts |
| DCXR | 0.3 | 0 | 0 | 0.27 | 0.18 | 1 | Matrix Fibroblasts |
| RP11-114H7.1 | 0.29 | 0 | 0 | 0.66 | 0.57 | 1 | Matrix Fibroblasts |
| AKR7A2 | 0.29 | 0 | 0 | 0.33 | 0.22 | 1 | Matrix Fibroblasts |
| ZEB2 | 0.29 | 0 | 0 | 0.57 | 0.41 | 1 | Matrix Fibroblasts |
| HOXB4 | 0.29 | 0 | 0 | 0.26 | 0.15 | 1 | Matrix Fibroblasts |
| RCN1 | 0.29 | 0 | 0 | 0.51 | 0.37 | 1 | Matrix Fibroblasts |
| MGST3 | 0.29 | 0 | 0 | 0.74 | 0.64 | 1 | Matrix Fibroblasts |
| PNRC1 | 0.29 | 0 | 0 | 0.68 | 0.58 | 1 | Matrix Fibroblasts |
| FBXO17 | 0.29 | 0 | 0 | 0.28 | 0.17 | 1 | Matrix Fibroblasts |
| HSD17B11 | 0.29 | 0 | 0 | 0.34 | 0.22 | 1 | Matrix Fibroblasts |
| HMGN2 | 0.28 | 0 | 0 | 0.76 | 0.67 | 1 | Matrix Fibroblasts |
| ELN | 0.28 | 0 | 0 | 0.79 | 0.50 | 1 | Matrix Fibroblasts |
| COX7A2L | 0.28 | 0 | 0 | 0.54 | 0.47 | 1 | Matrix Fibroblasts |
| UBE2E3 | 0.28 | 0 | 0 | 0.45 | 0.35 | 1 | Matrix Fibroblasts |
| AZI2 | 0.28 | 0 | 0 | 0.26 | 0.16 | 1 | Matrix Fibroblasts |
| PGRMC1 | 0.28 | 0 | 0 | 0.49 | 0.38 | 1 | Matrix Fibroblasts |
| TSC22D3 | 0.28 | 0 | 0 | 0.46 | 0.34 | 1 | Matrix Fibroblasts |
| TECR | 0.28 | 0 | 0 | 0.38 | 0.28 | 1 | Matrix Fibroblasts |
| RPL13AP5 | 0.28 | 0 | 0 | 0.97 | 0.93 | 1 | Matrix Fibroblasts |
| HEBP1 | 0.28 | 0 | 0 | 0.26 | 0.17 | 1 | Matrix Fibroblasts |
| SSR2 | 0.28 | 0 | 0 | 0.78 | 0.71 | 1 | Matrix Fibroblasts |
| MEIS1 | 0.28 | 0 | 0 | 0.28 | 0.18 | 1 | Matrix Fibroblasts |
| TSPYL1 | 0.28 | 0 | 0 | 0.31 | 0.21 | 1 | Matrix Fibroblasts |
| KDELR1 | 0.28 | 0 | 0 | 0.46 | 0.38 | 1 | Matrix Fibroblasts |
| KANK2 | 0.27 | 0 | 0 | 0.45 | 0.33 | 1 | Matrix Fibroblasts |
| HDDC2 | 0.27 | 0 | 0 | 0.32 | 0.21 | 1 | Matrix Fibroblasts |
| SMIM14 | 0.27 | 0 | 0 | 0.31 | 0.21 | 1 | Matrix Fibroblasts |
| CCNG1 | 0.27 | 0 | 0 | 0.29 | 0.19 | 1 | Matrix Fibroblasts |
| MARCKS | 0.27 | 0 | 0 | 0.65 | 0.51 | 1 | Matrix Fibroblasts |
| FKBP7 | 0.27 | 0 | 0 | 0.29 | 0.18 | 1 | Matrix Fibroblasts |
| RPL7AP6 | 0.27 | 0 | 0 | 0.64 | 0.55 | 1 | Matrix Fibroblasts |
| TMEM47 | 0.27 | 0 | 0 | 0.27 | 0.16 | 1 | Matrix Fibroblasts |
| RPL31 | 0.27 | 0 | 0 | 1.00 | 0.98 | 1 | Matrix Fibroblasts |
| MLLT4 | 0.26 | 0 | 0 | 0.25 | 0.16 | 1 | Matrix Fibroblasts |
| GLTSCR2 | 0.26 | 0 | 0 | 0.80 | 0.72 | 1 | Matrix Fibroblasts |
| TMEM59 | 0.26 | 0 | 0 | 0.75 | 0.67 | 1 | Matrix Fibroblasts |
| DPYSL2 | 0.26 | 0 | 0 | 0.37 | 0.29 | 1 | Matrix Fibroblasts |
| CD81 | 0.26 | 0 | 0 | 0.49 | 0.38 | 1 | Matrix Fibroblasts |
| HMGN3 | 0.26 | 0 | 0 | 0.53 | 0.48 | 1 | Matrix Fibroblasts |
| RPL13A | 0.25 | 0 | 0 | 1.00 | 1.00 | 1 | Matrix Fibroblasts |
| HMGB2 | 0.25 | 0 | 0 | 0.36 | 0.24 | 1 | Matrix Fibroblasts |
| NAP1L1 | 0.25 | 0 | 0 | 0.82 | 0.75 | 1 | Matrix Fibroblasts |
| PNISR | 0.25 | 0 | 0 | 0.70 | 0.63 | 1 | Matrix Fibroblasts |
| ADIRF | 2.71 | 0 | 0 | 1.00 | 0.44 | 2 | Smooth Muscle Cells |
| PI15 | 1.96 | 0 | 0 | 0.72 | 0.10 | 2 | Smooth Muscle Cells |
| PTN | 1.77 | 0 | 0 | 0.95 | 0.24 | 2 | Smooth Muscle Cells |
| SOD3 | 1.52 | 0 | 0 | 0.95 | 0.36 | 2 | Smooth Muscle Cells |
| PLN | 1.48 | 0 | 0 | 0.66 | 0.05 | 2 | Smooth Muscle Cells |
| NTRK3 | 1.46 | 0 | 0 | 0.82 | 0.08 | 2 | Smooth Muscle Cells |
| NET1 | 1.43 | 0 | 0 | 0.77 | 0.15 | 2 | Smooth Muscle Cells |
| BCAS3 | 1.37 | 0 | 0 | 0.69 | 0.11 | 2 | Smooth Muscle Cells |
| PTP4A3 | 1.34 | 0 | 0 | 0.76 | 0.19 | 2 | Smooth Muscle Cells |
| IGFBP5 | 1.25 | 0 | 0 | 0.81 | 0.24 | 2 | Smooth Muscle Cells |
| EGFL6 | 1.25 | 0 | 0 | 0.95 | 0.31 | 2 | Smooth Muscle Cells |
| SNCG | 1.19 | 0 | 0 | 0.61 | 0.06 | 2 | Smooth Muscle Cells |
| CRIP1 | 1.15 | 0 | 0 | 0.98 | 0.47 | 2 | Smooth Muscle Cells |
| KCNA5 | 1.14 | 0 | 0 | 0.58 | 0.02 | 2 | Smooth Muscle Cells |
| RERGL | 1.12 | 0 | 0 | 0.43 | 0.01 | 2 | Smooth Muscle Cells |
| CSRP2 | 1.11 | 0 | 0 | 0.67 | 0.19 | 2 | Smooth Muscle Cells |
| MYL9 | 1.11 | 0 | 0 | 0.99 | 0.68 | 2 | Smooth Muscle Cells |
| PTK2 | 1.05 | 0 | 0 | 0.78 | 0.25 | 2 | Smooth Muscle Cells |
| SERPINI1 | 1.04 | 0 | 0 | 0.56 | 0.08 | 2 | Smooth Muscle Cells |
| CRYAB | 1.04 | 0 | 0 | 0.84 | 0.29 | 2 | Smooth Muscle Cells |
| FRZB | 1.04 | 0 | 0 | 0.51 | 0.05 | 2 | Smooth Muscle Cells |
| TINAGL1 | 1.01 | 0 | 0 | 0.90 | 0.33 | 2 | Smooth Muscle Cells |
| MFGE8 | 0.98 | 0 | 0 | 0.78 | 0.28 | 2 | Smooth Muscle Cells |
| WFDC1 | 0.95 | 0 | 0 | 0.68 | 0.19 | 2 | Smooth Muscle Cells |
| MAP3K7CL | 0.94 | 0 | 0 | 0.41 | 0.04 | 2 | Smooth Muscle Cells |
| HES4 | 0.94 | 0 | 0 | 0.74 | 0.21 | 2 | Smooth Muscle Cells |
| ANGPT2 | 0.91 | 0 | 0 | 0.51 | 0.06 | 2 | Smooth Muscle Cells |
| ALDOC | 0.91 | 0 | 0 | 0.47 | 0.05 | 2 | Smooth Muscle Cells |
| TPM2 | 0.91 | 0 | 0 | 0.99 | 0.63 | 2 | Smooth Muscle Cells |
| ALDH1A2 | 0.9 | 0 | 0 | 0.45 | 0.04 | 2 | Smooth Muscle Cells |
| OAZ2 | 0.88 | 0 | 0 | 0.88 | 0.51 | 2 | Smooth Muscle Cells |
| MEF2C | 0.86 | 0 | 0 | 0.82 | 0.38 | 2 | Smooth Muscle Cells |
| PLAC9 | 0.86 | 0 | 0 | 0.85 | 0.36 | 2 | Smooth Muscle Cells |
| TAGLN | 0.84 | 0 | 0 | 0.99 | 0.58 | 2 | Smooth Muscle Cells |
| ADAMTS9 | 0.82 | 0 | 0 | 0.47 | 0.10 | 2 | Smooth Muscle Cells |
| RCAN2 | 0.82 | 0 | 0 | 0.46 | 0.03 | 2 | Smooth Muscle Cells |
| CXCL12 | 0.8 | 0 | 0 | 0.57 | 0.21 | 2 | Smooth Muscle Cells |
| MCAM | 0.8 | 0 | 0 | 0.66 | 0.18 | 2 | Smooth Muscle Cells |
| CRISPLD2 | 0.79 | 0 | 0 | 0.62 | 0.19 | 2 | Smooth Muscle Cells |
| ZFHX3 | 0.79 | 0 | 0 | 0.67 | 0.22 | 2 | Smooth Muscle Cells |
| AOC3 | 0.79 | 0 | 0 | 0.72 | 0.29 | 2 | Smooth Muscle Cells |
| MYH11 | 0.78 | 0 | 0 | 0.91 | 0.35 | 2 | Smooth Muscle Cells |
| PDLIM5 | 0.77 | 0 | 0 | 0.52 | 0.12 | 2 | Smooth Muscle Cells |
| KCNC4 | 0.75 | 0 | 0 | 0.48 | 0.06 | 2 | Smooth Muscle Cells |
| IL17B | 0.75 | 0 | 0 | 0.29 | 0.03 | 2 | Smooth Muscle Cells |
| GSN | 0.73 | 0 | 0 | 0.76 | 0.37 | 2 | Smooth Muscle Cells |
| IGFBP7 | 0.73 | 0 | 0 | 0.99 | 0.64 | 2 | Smooth Muscle Cells |
| GPRC5C | 0.72 | 0 | 0 | 0.46 | 0.07 | 2 | Smooth Muscle Cells |
| PRKCDBP | 0.72 | 0 | 0 | 0.76 | 0.37 | 2 | Smooth Muscle Cells |
| RAN | 0.7 | 0 | 0 | 0.81 | 0.63 | 2 | Smooth Muscle Cells |
| UBA2 | 0.7 | 0 | 0 | 0.64 | 0.24 | 2 | Smooth Muscle Cells |
| NDUFA4 | 0.69 | 0 | 0 | 0.97 | 0.78 | 2 | Smooth Muscle Cells |
| SLIT3 | 0.69 | 0 | 0 | 0.55 | 0.13 | 2 | Smooth Muscle Cells |
| CPE | 0.68 | 0 | 0 | 0.71 | 0.21 | 2 | Smooth Muscle Cells |
| AC013461.1 | 0.68 | 0 | 0 | 0.73 | 0.33 | 2 | Smooth Muscle Cells |
| PGF | 0.67 | 0 | 0 | 0.34 | 0.05 | 2 | Smooth Muscle Cells |
| ARTN | 0.67 | 0 | 0 | 0.33 | 0.01 | 2 | Smooth Muscle Cells |
| ALG9 | 0.67 | 0 | 0 | 0.29 | 0.03 | 2 | Smooth Muscle Cells |
| KLHL23 | 0.66 | 0 | 0 | 0.54 | 0.16 | 2 | Smooth Muscle Cells |
| SPARCL1 | 0.66 | 0 | 0 | 0.95 | 0.63 | 2 | Smooth Muscle Cells |
| PTPN2 | 0.66 | 0 | 0 | 0.54 | 0.17 | 2 | Smooth Muscle Cells |
| SGCA | 0.65 | 0 | 0 | 0.43 | 0.07 | 2 | Smooth Muscle Cells |
| DSTN | 0.65 | 0 | 0 | 0.98 | 0.78 | 2 | Smooth Muscle Cells |
| CASQ2 | 0.65 | 0 | 0 | 0.34 | 0.01 | 2 | Smooth Muscle Cells |
| ISYNA1 | 0.63 | 0 | 0 | 0.53 | 0.18 | 2 | Smooth Muscle Cells |
| RRAD | 0.62 | 0 | 0 | 0.50 | 0.15 | 2 | Smooth Muscle Cells |
| ACTA2 | 0.62 | 0 | 0 | 0.99 | 0.71 | 2 | Smooth Muscle Cells |
| TSC22D1 | 0.61 | 0 | 0 | 0.88 | 0.58 | 2 | Smooth Muscle Cells |
| TGM2 | 0.61 | 0 | 0 | 0.74 | 0.35 | 2 | Smooth Muscle Cells |
| ZFP36L1 | 0.61 | 0 | 0 | 0.94 | 0.72 | 2 | Smooth Muscle Cells |
| THSD4 | 0.6 | 0 | 0 | 0.38 | 0.07 | 2 | Smooth Muscle Cells |
| CBX6 | 0.6 | 0 | 0 | 0.60 | 0.27 | 2 | Smooth Muscle Cells |
| TPM1 | 0.59 | 0 | 0 | 0.91 | 0.56 | 2 | Smooth Muscle Cells |
| GPR20 | 0.59 | 0 | 0 | 0.35 | 0.04 | 2 | Smooth Muscle Cells |
| MGST3 | 0.59 | 0 | 0 | 0.87 | 0.62 | 2 | Smooth Muscle Cells |
| COX7A1 | 0.58 | 0 | 0 | 0.77 | 0.41 | 2 | Smooth Muscle Cells |
| SH3BGRL | 0.57 | 0 | 0 | 0.88 | 0.59 | 2 | Smooth Muscle Cells |
| HSPB2 | 0.57 | 0 | 0 | 0.47 | 0.13 | 2 | Smooth Muscle Cells |
| MYOM2 | 0.56 | 0 | 0 | 0.33 | 0.02 | 2 | Smooth Muscle Cells |
| C1QTNF1 | 0.56 | 0 | 0 | 0.35 | 0.03 | 2 | Smooth Muscle Cells |
| BCAM | 0.56 | 0 | 0 | 0.71 | 0.30 | 2 | Smooth Muscle Cells |
| MSRB3 | 0.55 | 0 | 0 | 0.56 | 0.21 | 2 | Smooth Muscle Cells |
| CHCHD10 | 0.55 | 0 | 0 | 0.50 | 0.16 | 2 | Smooth Muscle Cells |
| PDGFRB | 0.55 | 0 | 0 | 0.86 | 0.34 | 2 | Smooth Muscle Cells |
| NOL3 | 0.55 | 0 | 0 | 0.35 | 0.05 | 2 | Smooth Muscle Cells |
| HCFC1R1 | 0.55 | 0 | 0 | 0.47 | 0.21 | 2 | Smooth Muscle Cells |
| CABP1 | 0.54 | 0 | 0 | 0.27 | 0.01 | 2 | Smooth Muscle Cells |
| SLC25A4 | 0.54 | 0 | 0 | 0.37 | 0.14 | 2 | Smooth Muscle Cells |
| CARMN | 0.54 | 0 | 0 | 0.70 | 0.31 | 2 | Smooth Muscle Cells |
| HRH2 | 0.53 | 0 | 0 | 0.42 | 0.09 | 2 | Smooth Muscle Cells |
| NRGN | 0.52 | 0 | 0 | 0.30 | 0.04 | 2 | Smooth Muscle Cells |
| NDUFA4L2 | 0.52 | 0 | 0 | 0.57 | 0.19 | 2 | Smooth Muscle Cells |
| EDNRA | 0.52 | 0 | 0 | 0.57 | 0.20 | 2 | Smooth Muscle Cells |
| SPATS2L | 0.51 | 0 | 0 | 0.58 | 0.27 | 2 | Smooth Muscle Cells |
| SEPW1 | 0.51 | 0 | 0 | 0.84 | 0.53 | 2 | Smooth Muscle Cells |
| DMPK | 0.51 | 0 | 0 | 0.33 | 0.05 | 2 | Smooth Muscle Cells |
| ENPEP | 0.51 | 0 | 0 | 0.33 | 0.07 | 2 | Smooth Muscle Cells |
| NT5DC2 | 0.5 | 0 | 0 | 0.43 | 0.12 | 2 | Smooth Muscle Cells |
| RASL12 | 0.5 | 0 | 0 | 0.65 | 0.28 | 2 | Smooth Muscle Cells |
| APOE | 0.5 | 0 | 0 | 0.61 | 0.29 | 2 | Smooth Muscle Cells |
| RP11-270C12.3 | 0.49 | 0 | 0 | 0.57 | 0.27 | 2 | Smooth Muscle Cells |
| CDH6 | 0.49 | 0 | 0 | 0.38 | 0.09 | 2 | Smooth Muscle Cells |
| NOTCH3 | 0.49 | 0 | 0 | 0.57 | 0.19 | 2 | Smooth Muscle Cells |
| HACD1 | 0.49 | 0 | 0 | 0.38 | 0.08 | 2 | Smooth Muscle Cells |
| PDE1A | 0.49 | 0 | 0 | 0.27 | 0.01 | 2 | Smooth Muscle Cells |
| 4-Sep | 0.49 | 0 | 0 | 0.40 | 0.13 | 2 | Smooth Muscle Cells |
| NTF3 | 0.49 | 0 | 0 | 0.27 | 0.02 | 2 | Smooth Muscle Cells |
| SLIT2 | 0.49 | 0 | 0 | 0.45 | 0.14 | 2 | Smooth Muscle Cells |
| HEY2 | 0.49 | 0 | 0 | 0.30 | 0.02 | 2 | Smooth Muscle Cells |
| MYL6 | 0.48 | 0 | 0 | 1.00 | 0.98 | 2 | Smooth Muscle Cells |
| HSPB1 | 0.48 | 0 | 0 | 0.97 | 0.74 | 2 | Smooth Muscle Cells |
| C11orf96 | 0.48 | 0 | 0 | 0.56 | 0.27 | 2 | Smooth Muscle Cells |
| YBX3 | 0.48 | 0 | 0 | 0.66 | 0.37 | 2 | Smooth Muscle Cells |
| COX4I2 | 0.48 | 0 | 0 | 0.70 | 0.22 | 2 | Smooth Muscle Cells |
| TBX2 | 0.47 | 0 | 0 | 0.64 | 0.32 | 2 | Smooth Muscle Cells |
| JAG1 | 0.47 | 0 | 0 | 0.34 | 0.08 | 2 | Smooth Muscle Cells |
| LMOD1 | 0.47 | 0 | 0 | 0.41 | 0.11 | 2 | Smooth Muscle Cells |
| ITGA7 | 0.46 | 0 | 0 | 0.29 | 0.03 | 2 | Smooth Muscle Cells |
| EDNRB | 0.46 | 0 | 0 | 0.64 | 0.23 | 2 | Smooth Muscle Cells |
| MTHFD2 | 0.46 | 0 | 0 | 0.36 | 0.09 | 2 | Smooth Muscle Cells |
| NUDT4 | 0.46 | 0 | 0 | 0.54 | 0.23 | 2 | Smooth Muscle Cells |
| COL18A1 | 0.45 | 0 | 0 | 0.45 | 0.15 | 2 | Smooth Muscle Cells |
| FAM107B | 0.45 | 0 | 0 | 0.32 | 0.12 | 2 | Smooth Muscle Cells |
| ADAMTS1 | 0.45 | 0 | 0 | 0.39 | 0.17 | 2 | Smooth Muscle Cells |
| PPP1R12B | 0.45 | 0 | 0 | 0.47 | 0.18 | 2 | Smooth Muscle Cells |
| CAP2 | 0.44 | 0 | 0 | 0.27 | 0.05 | 2 | Smooth Muscle Cells |
| AKAP12 | 0.44 | 0 | 0 | 0.40 | 0.21 | 2 | Smooth Muscle Cells |
| TLE1 | 0.44 | 0 | 0 | 0.39 | 0.13 | 2 | Smooth Muscle Cells |
| DUSP14 | 0.44 | 0 | 0 | 0.32 | 0.08 | 2 | Smooth Muscle Cells |
| PPP1R12A | 0.44 | 0 | 0 | 0.73 | 0.44 | 2 | Smooth Muscle Cells |
| FILIP1 | 0.44 | 0 | 0 | 0.43 | 0.16 | 2 | Smooth Muscle Cells |
| OAT | 0.44 | 0 | 0 | 0.54 | 0.25 | 2 | Smooth Muscle Cells |
| TMEM51 | 0.43 | 0 | 0 | 0.33 | 0.08 | 2 | Smooth Muscle Cells |
| FOXS1 | 0.43 | 0 | 0 | 0.42 | 0.13 | 2 | Smooth Muscle Cells |
| MAP1B | 0.43 | 0 | 0 | 0.50 | 0.21 | 2 | Smooth Muscle Cells |
| FAM13C | 0.43 | 0 | 0 | 0.38 | 0.13 | 2 | Smooth Muscle Cells |
| TBC1D1 | 0.42 | 0 | 0 | 0.34 | 0.09 | 2 | Smooth Muscle Cells |
| SBSPON | 0.42 | 0 | 0 | 0.32 | 0.07 | 2 | Smooth Muscle Cells |
| EEF2 | 0.42 | 0 | 0 | 0.98 | 0.88 | 2 | Smooth Muscle Cells |
| HSPA2 | 0.41 | 0 | 0 | 0.36 | 0.15 | 2 | Smooth Muscle Cells |
| PHLDA1 | 0.41 | 0 | 0 | 0.46 | 0.22 | 2 | Smooth Muscle Cells |
| MRPS6 | 0.41 | 0 | 0 | 0.57 | 0.31 | 2 | Smooth Muscle Cells |
| PDE3A | 0.4 | 0 | 0 | 0.27 | 0.03 | 2 | Smooth Muscle Cells |
| HIF1A | 0.4 | 0 | 0 | 0.48 | 0.26 | 2 | Smooth Muscle Cells |
| PLP2 | 0.4 | 0 | 0 | 0.46 | 0.20 | 2 | Smooth Muscle Cells |
| GAPDH | 0.4 | 0 | 0 | 0.98 | 0.89 | 2 | Smooth Muscle Cells |
| PBX1 | 0.4 | 0 | 0 | 0.52 | 0.25 | 2 | Smooth Muscle Cells |
| SPEG | 0.39 | 0 | 0 | 0.34 | 0.10 | 2 | Smooth Muscle Cells |
| RP4-765C7.2 | 0.39 | 0 | 0 | 0.70 | 0.49 | 2 | Smooth Muscle Cells |
| NR4A1 | 0.39 | 0 | 0 | 0.34 | 0.12 | 2 | Smooth Muscle Cells |
| TBX2-AS1 | 0.39 | 0 | 0 | 0.44 | 0.18 | 2 | Smooth Muscle Cells |
| CDK2AP2 | 0.38 | 0 | 0 | 0.31 | 0.14 | 2 | Smooth Muscle Cells |
| ID4 | 0.38 | 0 | 0 | 0.54 | 0.24 | 2 | Smooth Muscle Cells |
| CALHM2 | 0.38 | 0 | 0 | 0.31 | 0.10 | 2 | Smooth Muscle Cells |
| LGALSL | 0.38 | 0 | 0 | 0.27 | 0.08 | 2 | Smooth Muscle Cells |
| HEYL | 0.37 | 0 | 0 | 0.36 | 0.11 | 2 | Smooth Muscle Cells |
| C1QTNF2 | 0.37 | 0 | 0 | 0.30 | 0.08 | 2 | Smooth Muscle Cells |
| INHBA | 0.37 | 0 | 0 | 0.25 | 0.03 | 2 | Smooth Muscle Cells |
| FAM129A | 0.37 | 0 | 0 | 0.26 | 0.05 | 2 | Smooth Muscle Cells |
| CD9 | 0.37 | 0 | 0 | 0.80 | 0.42 | 2 | Smooth Muscle Cells |
| NUPR1 | 0.37 | 0 | 0 | 0.79 | 0.47 | 2 | Smooth Muscle Cells |
| MT-ND1 | 0.36 | 0 | 0 | 1.00 | 0.97 | 2 | Smooth Muscle Cells |
| C16orf45 | 0.36 | 0 | 0 | 0.37 | 0.12 | 2 | Smooth Muscle Cells |
| YBX1 | 0.36 | 0 | 0 | 0.97 | 0.88 | 2 | Smooth Muscle Cells |
| MT1E | 0.35 | 0 | 0 | 0.47 | 0.22 | 2 | Smooth Muscle Cells |
| EBF1 | 0.35 | 0 | 0 | 0.33 | 0.09 | 2 | Smooth Muscle Cells |
| KANK2 | 0.35 | 0 | 0 | 0.60 | 0.32 | 2 | Smooth Muscle Cells |
| RPS2P5 | 0.35 | 0 | 0 | 0.89 | 0.72 | 2 | Smooth Muscle Cells |
| HSPB6 | 0.35 | 0 | 0 | 0.32 | 0.13 | 2 | Smooth Muscle Cells |
| SDF2L1 | 0.35 | 0 | 0.03 | 0.31 | 0.22 | 2 | Smooth Muscle Cells |
| ALDOA | 0.35 | 0 | 0 | 0.89 | 0.68 | 2 | Smooth Muscle Cells |
| TMEM136 | 0.34 | 0 | 0 | 0.26 | 0.06 | 2 | Smooth Muscle Cells |
| RP11-756G12.1 | 0.34 | 0 | 0 | 0.31 | 0.08 | 2 | Smooth Muscle Cells |
| PHPT1 | 0.34 | 0 | 0 | 0.68 | 0.43 | 2 | Smooth Muscle Cells |
| FOS | 0.34 | 0 | 0 | 0.51 | 0.33 | 2 | Smooth Muscle Cells |
| EEF1A1P5 | 0.34 | 0 | 0 | 0.84 | 0.64 | 2 | Smooth Muscle Cells |
| CYSTM1 | 0.34 | 0 | 0 | 0.58 | 0.31 | 2 | Smooth Muscle Cells |
| ARID5A | 0.34 | 0 | 0 | 0.37 | 0.14 | 2 | Smooth Muscle Cells |
| MT-ND3 | 0.34 | 0 | 0 | 1.00 | 0.98 | 2 | Smooth Muscle Cells |
| PTGIR | 0.34 | 0 | 0 | 0.37 | 0.14 | 2 | Smooth Muscle Cells |
| PLPP3 | 0.34 | 0 | 0 | 0.50 | 0.27 | 2 | Smooth Muscle Cells |
| YWHAQ | 0.34 | 0 | 0 | 0.85 | 0.62 | 2 | Smooth Muscle Cells |
| PLK2 | 0.33 | 0 | 0 | 0.37 | 0.18 | 2 | Smooth Muscle Cells |
| HSPB7 | 0.33 | 0 | 0 | 0.28 | 0.08 | 2 | Smooth Muscle Cells |
| ARL3 | 0.33 | 0 | 0 | 0.52 | 0.28 | 2 | Smooth Muscle Cells |
| CD82 | 0.33 | 0 | 0 | 0.49 | 0.24 | 2 | Smooth Muscle Cells |
| TGFB1I1 | 0.33 | 0 | 0 | 0.67 | 0.40 | 2 | Smooth Muscle Cells |
| ARHGEF7 | 0.33 | 0 | 0 | 0.26 | 0.07 | 2 | Smooth Muscle Cells |
| POLD2 | 0.32 | 0 | 0 | 0.44 | 0.21 | 2 | Smooth Muscle Cells |
| RAD23A | 0.32 | 0 | 0 | 0.66 | 0.43 | 2 | Smooth Muscle Cells |
| COX7A2 | 0.32 | 0 | 0 | 0.93 | 0.78 | 2 | Smooth Muscle Cells |
| ST6GALNAC6 | 0.32 | 0 | 0 | 0.39 | 0.16 | 2 | Smooth Muscle Cells |
| GRK5 | 0.32 | 0 | 0 | 0.39 | 0.17 | 2 | Smooth Muscle Cells |
| PTMAP5 | 0.32 | 0 | 0 | 0.85 | 0.67 | 2 | Smooth Muscle Cells |
| CTNNB1 | 0.32 | 0 | 0 | 0.52 | 0.36 | 2 | Smooth Muscle Cells |
| MPRIP | 0.31 | 0 | 0 | 0.41 | 0.19 | 2 | Smooth Muscle Cells |
| GSTP1 | 0.31 | 0 | 0 | 0.96 | 0.81 | 2 | Smooth Muscle Cells |
| B3GNT2 | 0.31 | 0 | 0 | 0.26 | 0.07 | 2 | Smooth Muscle Cells |
| AC090498.1 | 0.31 | 0 | 0 | 0.99 | 0.93 | 2 | Smooth Muscle Cells |
| MTND1P23 | 0.31 | 0 | 0 | 0.70 | 0.45 | 2 | Smooth Muscle Cells |
| RPL23AP42 | 0.31 | 0 | 0 | 0.85 | 0.65 | 2 | Smooth Muscle Cells |
| DNAJB9 | 0.31 | 0 | 0 | 0.27 | 0.15 | 2 | Smooth Muscle Cells |
| COX7B | 0.31 | 0 | 0 | 0.81 | 0.60 | 2 | Smooth Muscle Cells |
| MAPRE2 | 0.31 | 0 | 0 | 0.32 | 0.12 | 2 | Smooth Muscle Cells |
| MAFB | 0.31 | 0 | 0 | 0.39 | 0.23 | 2 | Smooth Muscle Cells |
| MLEC | 0.31 | 0 | 0 | 0.43 | 0.20 | 2 | Smooth Muscle Cells |
| CTBP2 | 0.31 | 0 | 0 | 0.42 | 0.21 | 2 | Smooth Muscle Cells |
| LDHB | 0.31 | 0 | 0 | 0.84 | 0.59 | 2 | Smooth Muscle Cells |
| CHURC1 | 0.31 | 0 | 0 | 0.56 | 0.35 | 2 | Smooth Muscle Cells |
| RPL5P34 | 0.31 | 0 | 0 | 0.91 | 0.73 | 2 | Smooth Muscle Cells |
| ANGPTL4 | 0.31 | 0 | 0 | 0.30 | 0.12 | 2 | Smooth Muscle Cells |
| NDUFB9 | 0.31 | 0 | 0 | 0.74 | 0.51 | 2 | Smooth Muscle Cells |
| PPP1R14A | 0.31 | 0 | 0 | 0.64 | 0.41 | 2 | Smooth Muscle Cells |
| CAMK2G | 0.3 | 0 | 0 | 0.29 | 0.10 | 2 | Smooth Muscle Cells |
| ATP5D | 0.3 | 0 | 0 | 0.82 | 0.62 | 2 | Smooth Muscle Cells |
| MT-RNR1 | 0.3 | 0 | 0.02 | 0.97 | 0.95 | 2 | Smooth Muscle Cells |
| MT-RNR2 | 0.3 | 0 | 0 | 1.00 | 1.00 | 2 | Smooth Muscle Cells |
| PTMAP2 | 0.3 | 0 | 0 | 0.78 | 0.60 | 2 | Smooth Muscle Cells |
| ARPC5 | 0.3 | 0 | 0 | 0.82 | 0.63 | 2 | Smooth Muscle Cells |
| COX6A1 | 0.3 | 0 | 0 | 0.91 | 0.73 | 2 | Smooth Muscle Cells |
| GPM6B | 0.29 | 0 | 0 | 0.53 | 0.27 | 2 | Smooth Muscle Cells |
| VIM | 0.29 | 0 | 0 | 1.00 | 0.94 | 2 | Smooth Muscle Cells |
| GNG2 | 0.29 | 0 | 0 | 0.30 | 0.14 | 2 | Smooth Muscle Cells |
| RPS2 | 0.29 | 0 | 0 | 1.00 | 0.99 | 2 | Smooth Muscle Cells |
| FKBP4 | 0.29 | 0 | 0 | 0.37 | 0.17 | 2 | Smooth Muscle Cells |
| PTMA | 0.29 | 0 | 0 | 1.00 | 1.00 | 2 | Smooth Muscle Cells |
| PLS3 | 0.29 | 0 | 0 | 0.65 | 0.41 | 2 | Smooth Muscle Cells |
| MOB2 | 0.29 | 0 | 0 | 0.40 | 0.20 | 2 | Smooth Muscle Cells |
| MT-ND4L | 0.29 | 0 | 0 | 0.86 | 0.70 | 2 | Smooth Muscle Cells |
| RPL7P1 | 0.28 | 0 | 0 | 0.88 | 0.65 | 2 | Smooth Muscle Cells |
| PPP2R5E | 0.28 | 0 | 0 | 0.34 | 0.16 | 2 | Smooth Muscle Cells |
| CTB-63M22.1 | 0.28 | 0 | 0 | 0.80 | 0.58 | 2 | Smooth Muscle Cells |
| EMD | 0.28 | 0 | 0 | 0.37 | 0.17 | 2 | Smooth Muscle Cells |
| RPL14 | 0.28 | 0 | 0 | 1.00 | 0.95 | 2 | Smooth Muscle Cells |
| RP11-371A22.1 | 0.28 | 0 | 0 | 0.97 | 0.85 | 2 | Smooth Muscle Cells |
| EIF3D | 0.28 | 0 | 0 | 0.66 | 0.44 | 2 | Smooth Muscle Cells |
| CYC1 | 0.28 | 0 | 0 | 0.62 | 0.40 | 2 | Smooth Muscle Cells |
| FXYD1 | 0.28 | 0 | 0 | 0.48 | 0.25 | 2 | Smooth Muscle Cells |
| HIGD1B | 0.28 | 0 | 0 | 0.70 | 0.24 | 2 | Smooth Muscle Cells |
| RBMS1 | 0.28 | 0 | 0 | 0.57 | 0.39 | 2 | Smooth Muscle Cells |
| ATPAF1 | 0.28 | 0 | 0 | 0.28 | 0.11 | 2 | Smooth Muscle Cells |
| NDUFB7 | 0.28 | 0 | 0 | 0.76 | 0.56 | 2 | Smooth Muscle Cells |
| CTD-2192J16.15 | 0.27 | 0 | 0 | 0.97 | 0.90 | 2 | Smooth Muscle Cells |
| RPL23A | 0.27 | 0 | 0 | 1.00 | 0.98 | 2 | Smooth Muscle Cells |
| MT-CYB | 0.27 | 0 | 0 | 1.00 | 0.99 | 2 | Smooth Muscle Cells |
| MAGEF1 | 0.27 | 0 | 0 | 0.41 | 0.20 | 2 | Smooth Muscle Cells |
| RPL3P4 | 0.27 | 0 | 0 | 1.00 | 0.94 | 2 | Smooth Muscle Cells |
| MT-ND4 | 0.27 | 0 | 0 | 1.00 | 1.00 | 2 | Smooth Muscle Cells |
| ARPC1A | 0.27 | 0 | 0 | 0.47 | 0.26 | 2 | Smooth Muscle Cells |
| PLCE1 | 0.27 | 0 | 0 | 0.26 | 0.08 | 2 | Smooth Muscle Cells |
| SMIM12 | 0.27 | 0 | 0 | 0.33 | 0.17 | 2 | Smooth Muscle Cells |
| PNMA1 | 0.27 | 0 | 0 | 0.33 | 0.15 | 2 | Smooth Muscle Cells |
| RHOB | 0.27 | 0 | 0 | 0.65 | 0.46 | 2 | Smooth Muscle Cells |
| SLMAP | 0.27 | 0 | 0 | 0.33 | 0.16 | 2 | Smooth Muscle Cells |
| FAUP1 | 0.26 | 0 | 0 | 0.81 | 0.63 | 2 | Smooth Muscle Cells |
| EEF1D | 0.26 | 0 | 0 | 0.97 | 0.88 | 2 | Smooth Muscle Cells |
| PLEKHA4 | 0.26 | 0 | 0 | 0.29 | 0.12 | 2 | Smooth Muscle Cells |
| MAP7D3 | 0.26 | 0 | 0 | 0.32 | 0.14 | 2 | Smooth Muscle Cells |
| MT-CO3 | 0.26 | 0 | 0 | 1.00 | 0.99 | 2 | Smooth Muscle Cells |
| FAU | 0.26 | 0 | 0 | 1.00 | 0.97 | 2 | Smooth Muscle Cells |
| NACA | 0.26 | 0 | 0 | 0.99 | 0.93 | 2 | Smooth Muscle Cells |
| NDUFA5 | 0.26 | 0 | 0 | 0.63 | 0.41 | 2 | Smooth Muscle Cells |
| RSL24D1 | 0.26 | 0 | 0 | 0.73 | 0.49 | 2 | Smooth Muscle Cells |
| PRDM6 | 0.26 | 0 | 0 | 0.30 | 0.14 | 2 | Smooth Muscle Cells |
| TPI1 | 0.26 | 0 | 0 | 0.91 | 0.75 | 2 | Smooth Muscle Cells |
| IAH1 | 0.26 | 0 | 0 | 0.43 | 0.24 | 2 | Smooth Muscle Cells |
| RP3-486I3.4 | 0.26 | 0 | 0 | 0.86 | 0.64 | 2 | Smooth Muscle Cells |
| TOB1 | 0.26 | 0 | 0 | 0.29 | 0.12 | 2 | Smooth Muscle Cells |
| MT-ATP6 | 0.26 | 0 | 0 | 1.00 | 0.99 | 2 | Smooth Muscle Cells |
| DACT3 | 0.26 | 0 | 0 | 0.25 | 0.09 | 2 | Smooth Muscle Cells |
| GAS5 | 0.26 | 0 | 0 | 0.89 | 0.70 | 2 | Smooth Muscle Cells |
| COX7C | 0.26 | 0 | 0 | 0.98 | 0.86 | 2 | Smooth Muscle Cells |
| CD151 | 0.25 | 0 | 0 | 0.75 | 0.52 | 2 | Smooth Muscle Cells |
| RPS2P46 | 0.25 | 0 | 0 | 0.68 | 0.47 | 2 | Smooth Muscle Cells |
| CHRAC1 | 0.25 | 0 | 0 | 0.28 | 0.12 | 2 | Smooth Muscle Cells |
| MTATP6P1 | 0.25 | 0 | 0 | 1.00 | 0.98 | 2 | Smooth Muscle Cells |
| RPL41 | 0.25 | 0 | 0 | 1.00 | 1.00 | 2 | Smooth Muscle Cells |
| ANAPC11 | 0.25 | 0 | 0 | 0.74 | 0.55 | 2 | Smooth Muscle Cells |
| RP11-572P18.1 | 0.25 | 0 | 0 | 0.58 | 0.39 | 2 | Smooth Muscle Cells |
| RGCC | 2.26 | 0 | 0 | 0.91 | 0.28 | 3 | Endothelial Cells |
| EDN1 | 2.12 | 0 | 0 | 0.64 | 0.04 | 3 | Endothelial Cells |
| RAMP2 | 1.84 | 0 | 0 | 0.98 | 0.27 | 3 | Endothelial Cells |
| CA4 | 1.84 | 0 | 0 | 0.76 | 0.04 | 3 | Endothelial Cells |
| IFI27 | 1.84 | 0 | 0 | 0.99 | 0.30 | 3 | Endothelial Cells |
| TMEM100 | 1.81 | 0 | 0 | 0.70 | 0.10 | 3 | Endothelial Cells |
| TNFSF10 | 1.79 | 0 | 0 | 0.92 | 0.15 | 3 | Endothelial Cells |
| BST2 | 1.75 | 0 | 0 | 0.96 | 0.23 | 3 | Endothelial Cells |
| CD93 | 1.69 | 0 | 0 | 0.82 | 0.08 | 3 | Endothelial Cells |
| CLEC14A | 1.68 | 0 | 0 | 0.86 | 0.13 | 3 | Endothelial Cells |
| CALCRL | 1.64 | 0 | 0 | 0.83 | 0.13 | 3 | Endothelial Cells |
| EGFL7 | 1.63 | 0 | 0 | 0.95 | 0.19 | 3 | Endothelial Cells |
| GNG11 | 1.62 | 0 | 0 | 0.96 | 0.39 | 3 | Endothelial Cells |
| IL7R | 1.61 | 0 | 0 | 0.56 | 0.04 | 3 | Endothelial Cells |
| RNASE1 | 1.59 | 0 | 0 | 0.93 | 0.20 | 3 | Endothelial Cells |
| CCND1 | 1.57 | 0 | 0 | 0.76 | 0.17 | 3 | Endothelial Cells |
| C8orf4 | 1.56 | 0 | 0 | 0.88 | 0.39 | 3 | Endothelial Cells |
| RDX | 1.56 | 0 | 0 | 0.87 | 0.39 | 3 | Endothelial Cells |
| EPAS1 | 1.54 | 0 | 0 | 0.96 | 0.38 | 3 | Endothelial Cells |
| CLDN5 | 1.54 | 0 | 0 | 0.92 | 0.20 | 3 | Endothelial Cells |
| PCAT19 | 1.47 | 0 | 0 | 0.88 | 0.11 | 3 | Endothelial Cells |
| CDH5 | 1.44 | 0 | 0 | 0.82 | 0.12 | 3 | Endothelial Cells |
| IL33 | 1.44 | 0 | 0 | 0.66 | 0.06 | 3 | Endothelial Cells |
| GIMAP7 | 1.42 | 0 | 0 | 0.73 | 0.11 | 3 | Endothelial Cells |
| HLA-C | 1.39 | 0 | 0 | 0.97 | 0.50 | 3 | Endothelial Cells |
| CD36 | 1.39 | 0 | 0 | 0.66 | 0.10 | 3 | Endothelial Cells |
| HPGD | 1.37 | 0 | 0 | 0.80 | 0.17 | 3 | Endothelial Cells |
| ARHGAP18 | 1.37 | 0 | 0 | 0.65 | 0.07 | 3 | Endothelial Cells |
| ACE | 1.36 | 0 | 0 | 0.68 | 0.08 | 3 | Endothelial Cells |
| TIE1 | 1.33 | 0 | 0 | 0.71 | 0.08 | 3 | Endothelial Cells |
| IGFBP4 | 1.33 | 0 | 0 | 0.98 | 0.45 | 3 | Endothelial Cells |
| PECAM1 | 1.33 | 0 | 0 | 0.92 | 0.19 | 3 | Endothelial Cells |
| HLA-A | 1.32 | 0 | 0 | 1.00 | 0.67 | 3 | Endothelial Cells |
| S100A16 | 1.31 | 0 | 0 | 0.87 | 0.29 | 3 | Endothelial Cells |
| HLA-B | 1.31 | 0 | 0 | 0.99 | 0.49 | 3 | Endothelial Cells |
| IL32 | 1.29 | 0 | 0 | 0.82 | 0.23 | 3 | Endothelial Cells |
| ESAM | 1.26 | 0 | 0 | 0.88 | 0.30 | 3 | Endothelial Cells |
| CD34 | 1.26 | 0 | 0 | 0.65 | 0.10 | 3 | Endothelial Cells |
| RAMP3 | 1.24 | 0 | 0 | 0.67 | 0.11 | 3 | Endothelial Cells |
| THBD | 1.23 | 0 | 0 | 0.62 | 0.10 | 3 | Endothelial Cells |
| ECSCR.1 | 1.23 | 0 | 0 | 0.82 | 0.13 | 3 | Endothelial Cells |
| LXN | 1.22 | 0 | 0 | 0.57 | 0.12 | 3 | Endothelial Cells |
| FABP5 | 1.21 | 0 | 0 | 0.58 | 0.24 | 3 | Endothelial Cells |
| TMEM88 | 1.21 | 0 | 0 | 0.53 | 0.03 | 3 | Endothelial Cells |
| VWF | 1.2 | 0 | 0 | 0.67 | 0.07 | 3 | Endothelial Cells |
| PCDH17 | 1.2 | 0 | 0 | 0.60 | 0.10 | 3 | Endothelial Cells |
| SLC9A3R2 | 1.19 | 0 | 0 | 0.82 | 0.36 | 3 | Endothelial Cells |
| LDB2 | 1.19 | 0 | 0 | 0.73 | 0.13 | 3 | Endothelial Cells |
| NOTCH4 | 1.19 | 0 | 0 | 0.64 | 0.07 | 3 | Endothelial Cells |
| B2M | 1.19 | 0 | 0 | 1.00 | 0.99 | 3 | Endothelial Cells |
| GPIHBP1 | 1.17 | 0 | 0 | 0.45 | 0.01 | 3 | Endothelial Cells |
| IGFBP2 | 1.17 | 0 | 0 | 0.69 | 0.13 | 3 | Endothelial Cells |
| ICAM2 | 1.15 | 0 | 0 | 0.76 | 0.17 | 3 | Endothelial Cells |
| CYB5A | 1.14 | 0 | 0 | 0.80 | 0.30 | 3 | Endothelial Cells |
| SH3BP5 | 1.13 | 0 | 0 | 0.76 | 0.16 | 3 | Endothelial Cells |
| SGK1 | 1.13 | 0 | 0 | 0.53 | 0.12 | 3 | Endothelial Cells |
| FKBP1A | 1.12 | 0 | 0 | 0.97 | 0.69 | 3 | Endothelial Cells |
| ARHGAP29 | 1.12 | 0 | 0 | 0.82 | 0.28 | 3 | Endothelial Cells |
| SOX7 | 1.12 | 0 | 0 | 0.54 | 0.04 | 3 | Endothelial Cells |
| HLA-E | 1.12 | 0 | 0 | 0.98 | 0.56 | 3 | Endothelial Cells |
| TM4SF1 | 1.11 | 0 | 0 | 0.91 | 0.35 | 3 | Endothelial Cells |
| FCN3 | 1.11 | 0 | 0 | 0.43 | 0.03 | 3 | Endothelial Cells |
| CCDC85B | 1.11 | 0 | 0 | 0.86 | 0.36 | 3 | Endothelial Cells |
| ROBO4 | 1.11 | 0 | 0 | 0.61 | 0.06 | 3 | Endothelial Cells |
| ACVRL1 | 1.11 | 0 | 0 | 0.70 | 0.11 | 3 | Endothelial Cells |
| PRCP | 1.1 | 0 | 0 | 0.67 | 0.20 | 3 | Endothelial Cells |
| ITM2A | 1.1 | 0 | 0 | 0.75 | 0.13 | 3 | Endothelial Cells |
| CAV1 | 1.1 | 0 | 0 | 0.99 | 0.81 | 3 | Endothelial Cells |
| GIMAP4 | 1.09 | 0 | 0 | 0.67 | 0.14 | 3 | Endothelial Cells |
| EMCN | 1.08 | 0 | 0 | 0.62 | 0.12 | 3 | Endothelial Cells |
| VAMP5 | 1.08 | 0 | 0 | 0.91 | 0.48 | 3 | Endothelial Cells |
| ID1 | 1.07 | 0 | 0 | 0.75 | 0.30 | 3 | Endothelial Cells |
| SOX18 | 1.07 | 0 | 0 | 0.59 | 0.06 | 3 | Endothelial Cells |
| RALA | 1.07 | 0 | 0 | 0.72 | 0.25 | 3 | Endothelial Cells |
| TSPAN7 | 1.06 | 0 | 0 | 0.58 | 0.08 | 3 | Endothelial Cells |
| SDPR | 1.06 | 0 | 0 | 0.72 | 0.20 | 3 | Endothelial Cells |
| APLNR | 1.05 | 0 | 0 | 0.58 | 0.07 | 3 | Endothelial Cells |
| CLEC2B | 1.05 | 0 | 0 | 0.61 | 0.12 | 3 | Endothelial Cells |
| THSD1 | 1.04 | 0 | 0 | 0.53 | 0.05 | 3 | Endothelial Cells |
| GMFG | 1.03 | 0 | 0 | 0.79 | 0.22 | 3 | Endothelial Cells |
| MARCKSL1 | 1.03 | 0 | 0 | 0.89 | 0.58 | 3 | Endothelial Cells |
| CTNND1 | 1.01 | 0 | 0 | 0.68 | 0.20 | 3 | Endothelial Cells |
| KIT | 1.01 | 0 | 0 | 0.45 | 0.01 | 3 | Endothelial Cells |
| PALMD | 1 | 0 | 0 | 0.48 | 0.04 | 3 | Endothelial Cells |
| CYYR1 | 1 | 0 | 0 | 0.55 | 0.07 | 3 | Endothelial Cells |
| LEPR | 0.99 | 0 | 0 | 0.47 | 0.04 | 3 | Endothelial Cells |
| FENDRR | 0.99 | 0 | 0 | 0.57 | 0.11 | 3 | Endothelial Cells |
| NUDT4 | 0.98 | 0 | 0 | 0.59 | 0.22 | 3 | Endothelial Cells |
| RGS5 | 0.97 | 0 | 0 | 0.71 | 0.19 | 3 | Endothelial Cells |
| H19 | 0.96 | 0 | 0 | 0.46 | 0.13 | 3 | Endothelial Cells |
| CAV2 | 0.95 | 0 | 0 | 0.91 | 0.55 | 3 | Endothelial Cells |
| LYVE1 | 0.94 | 0 | 0 | 0.38 | 0.03 | 3 | Endothelial Cells |
| ID3 | 0.94 | 0 | 0 | 0.86 | 0.50 | 3 | Endothelial Cells |
| DUSP6 | 0.94 | 0 | 0 | 0.55 | 0.15 | 3 | Endothelial Cells |
| EFNA1 | 0.93 | 0 | 0 | 0.55 | 0.10 | 3 | Endothelial Cells |
| PTRF | 0.93 | 0 | 0 | 0.95 | 0.66 | 3 | Endothelial Cells |
| TMSB10 | 0.92 | 0 | 0 | 1.00 | 0.99 | 3 | Endothelial Cells |
| C1orf54 | 0.92 | 0 | 0 | 0.54 | 0.15 | 3 | Endothelial Cells |
| JAM2 | 0.92 | 0 | 0 | 0.52 | 0.08 | 3 | Endothelial Cells |
| WARS | 0.9 | 0 | 0 | 0.55 | 0.16 | 3 | Endothelial Cells |
| HYAL2 | 0.9 | 0 | 0 | 0.61 | 0.19 | 3 | Endothelial Cells |
| LIFR | 0.9 | 0 | 0 | 0.47 | 0.09 | 3 | Endothelial Cells |
| KDR | 0.9 | 0 | 0 | 0.48 | 0.05 | 3 | Endothelial Cells |
| RALB | 0.89 | 0 | 0 | 0.54 | 0.15 | 3 | Endothelial Cells |
| CARHSP1 | 0.89 | 0 | 0 | 0.71 | 0.33 | 3 | Endothelial Cells |
| FAM167B | 0.88 | 0 | 0 | 0.44 | 0.05 | 3 | Endothelial Cells |
| MYZAP | 0.88 | 0 | 0 | 0.51 | 0.06 | 3 | Endothelial Cells |
| STXBP6 | 0.88 | 0 | 0 | 0.54 | 0.11 | 3 | Endothelial Cells |
| POSTN | 0.87 | 0 | 0 | 0.56 | 0.18 | 3 | Endothelial Cells |
| PLK2 | 0.86 | 0 | 0 | 0.56 | 0.16 | 3 | Endothelial Cells |
| ETS2 | 0.86 | 0 | 0 | 0.65 | 0.24 | 3 | Endothelial Cells |
| KIAA1462 | 0.86 | 0 | 0 | 0.53 | 0.09 | 3 | Endothelial Cells |
| PSMB9 | 0.86 | 0 | 0 | 0.63 | 0.17 | 3 | Endothelial Cells |
| BCAM | 0.85 | 0 | 0 | 0.76 | 0.29 | 3 | Endothelial Cells |
| FSCN1 | 0.85 | 0 | 0 | 0.65 | 0.18 | 3 | Endothelial Cells |
| ITGA6 | 0.84 | 0 | 0 | 0.42 | 0.03 | 3 | Endothelial Cells |
| GNAI2 | 0.84 | 0 | 0 | 0.71 | 0.31 | 3 | Endothelial Cells |
| SEC14L1 | 0.84 | 0 | 0 | 0.51 | 0.10 | 3 | Endothelial Cells |
| LY6E | 0.83 | 0 | 0 | 0.78 | 0.42 | 3 | Endothelial Cells |
| FOXF1 | 0.83 | 0 | 0 | 0.65 | 0.22 | 3 | Endothelial Cells |
| FLI1 | 0.83 | 0 | 0 | 0.49 | 0.08 | 3 | Endothelial Cells |
| EFNB2 | 0.82 | 0 | 0 | 0.36 | 0.06 | 3 | Endothelial Cells |
| KCTD12 | 0.82 | 0 | 0 | 0.57 | 0.20 | 3 | Endothelial Cells |
| CTNNAL1 | 0.82 | 0 | 0 | 0.51 | 0.16 | 3 | Endothelial Cells |
| GBP4 | 0.82 | 0 | 0 | 0.46 | 0.06 | 3 | Endothelial Cells |
| S100A13 | 0.81 | 0 | 0 | 0.75 | 0.36 | 3 | Endothelial Cells |
| LEPROTL1 | 0.81 | 0 | 0 | 0.62 | 0.21 | 3 | Endothelial Cells |
| KIAA1217 | 0.8 | 0 | 0 | 0.39 | 0.03 | 3 | Endothelial Cells |
| SPTBN1 | 0.8 | 0 | 0 | 0.76 | 0.39 | 3 | Endothelial Cells |
| TCF4 | 0.8 | 0 | 0 | 0.82 | 0.44 | 3 | Endothelial Cells |
| RAB11A | 0.79 | 0 | 0 | 0.77 | 0.41 | 3 | Endothelial Cells |
| NPDC1 | 0.79 | 0 | 0 | 0.68 | 0.26 | 3 | Endothelial Cells |
| IFITM3 | 0.79 | 0 | 0 | 1.00 | 0.90 | 3 | Endothelial Cells |
| IFITM2 | 0.79 | 0 | 0 | 0.88 | 0.61 | 3 | Endothelial Cells |
| KANK3 | 0.79 | 0 | 0 | 0.50 | 0.11 | 3 | Endothelial Cells |
| EGLN3 | 0.78 | 0 | 0 | 0.40 | 0.05 | 3 | Endothelial Cells |
| ADCY4 | 0.78 | 0 | 0 | 0.43 | 0.06 | 3 | Endothelial Cells |
| APOL3 | 0.78 | 0 | 0 | 0.49 | 0.08 | 3 | Endothelial Cells |
| SOX17 | 0.78 | 0 | 0 | 0.37 | 0.03 | 3 | Endothelial Cells |
| PRPSAP1 | 0.78 | 0 | 0 | 0.52 | 0.15 | 3 | Endothelial Cells |
| MAP3K11 | 0.77 | 0 | 0 | 0.50 | 0.12 | 3 | Endothelial Cells |
| AQP1 | 0.77 | 0 | 0 | 0.59 | 0.17 | 3 | Endothelial Cells |
| ENG | 0.77 | 0 | 0 | 0.65 | 0.26 | 3 | Endothelial Cells |
| CXorf36 | 0.77 | 0 | 0 | 0.40 | 0.05 | 3 | Endothelial Cells |
| APP | 0.77 | 0 | 0 | 0.81 | 0.48 | 3 | Endothelial Cells |
| CFLAR | 0.77 | 0 | 0 | 0.63 | 0.23 | 3 | Endothelial Cells |
| CD200 | 0.77 | 0 | 0 | 0.45 | 0.08 | 3 | Endothelial Cells |
| LMO2 | 0.76 | 0 | 0 | 0.42 | 0.06 | 3 | Endothelial Cells |
| NOSTRIN | 0.75 | 0 | 0 | 0.37 | 0.03 | 3 | Endothelial Cells |
| TEK | 0.75 | 0 | 0 | 0.39 | 0.03 | 3 | Endothelial Cells |
| SERPINB6 | 0.74 | 0 | 0 | 0.69 | 0.39 | 3 | Endothelial Cells |
| PCDH12 | 0.74 | 0 | 0 | 0.42 | 0.06 | 3 | Endothelial Cells |
| PROCR | 0.74 | 0 | 0 | 0.43 | 0.11 | 3 | Endothelial Cells |
| RAPGEF5 | 0.74 | 0 | 0 | 0.47 | 0.09 | 3 | Endothelial Cells |
| ADGRF5 | 0.74 | 0 | 0 | 0.52 | 0.14 | 3 | Endothelial Cells |
| SCD | 0.74 | 0 | 0 | 0.27 | 0.03 | 3 | Endothelial Cells |
| DTL | 0.73 | 0 | 0 | 0.30 | 0.01 | 3 | Endothelial Cells |
| NPR3 | 0.73 | 0 | 0 | 0.38 | 0.06 | 3 | Endothelial Cells |
| ELK3 | 0.73 | 0 | 0 | 0.48 | 0.12 | 3 | Endothelial Cells |
| PSMB8 | 0.73 | 0 | 0 | 0.61 | 0.22 | 3 | Endothelial Cells |
| FDPS | 0.72 | 0 | 0 | 0.64 | 0.31 | 3 | Endothelial Cells |
| PLAT | 0.72 | 0 | 0 | 0.30 | 0.13 | 3 | Endothelial Cells |
| MGST2 | 0.71 | 0 | 0 | 0.59 | 0.23 | 3 | Endothelial Cells |
| ERG | 0.7 | 0 | 0 | 0.44 | 0.08 | 3 | Endothelial Cells |
| ARHGEF15 | 0.7 | 0 | 0 | 0.37 | 0.04 | 3 | Endothelial Cells |
| PODXL | 0.7 | 0 | 0 | 0.40 | 0.06 | 3 | Endothelial Cells |
| MYCT1 | 0.7 | 0 | 0 | 0.37 | 0.03 | 3 | Endothelial Cells |
| PTPRB | 0.7 | 0 | 0 | 0.38 | 0.04 | 3 | Endothelial Cells |
| EMP2 | 0.7 | 0 | 0 | 0.78 | 0.46 | 3 | Endothelial Cells |
| PDGFB | 0.7 | 0 | 0 | 0.35 | 0.03 | 3 | Endothelial Cells |
| BMPR2 | 0.69 | 0 | 0 | 0.53 | 0.19 | 3 | Endothelial Cells |
| ITM2B | 0.69 | 0 | 0 | 0.97 | 0.88 | 3 | Endothelial Cells |
| CARD16 | 0.69 | 0 | 0 | 0.62 | 0.25 | 3 | Endothelial Cells |
| C10orf54 | 0.69 | 0 | 0 | 0.52 | 0.17 | 3 | Endothelial Cells |
| GPR146 | 0.69 | 0 | 0 | 0.41 | 0.06 | 3 | Endothelial Cells |
| IFI6 | 0.69 | 0 | 0 | 0.55 | 0.23 | 3 | Endothelial Cells |
| GIMAP1 | 0.69 | 0 | 0 | 0.43 | 0.07 | 3 | Endothelial Cells |
| MAOA | 0.69 | 0 | 0 | 0.35 | 0.05 | 3 | Endothelial Cells |
| BRI3 | 0.69 | 0 | 0 | 0.59 | 0.26 | 3 | Endothelial Cells |
| TGFBR2 | 0.69 | 0 | 0 | 0.52 | 0.16 | 3 | Endothelial Cells |
| TFPI | 0.69 | 0 | 0 | 0.73 | 0.40 | 3 | Endothelial Cells |
| CALM1 | 0.69 | 0 | 0 | 0.91 | 0.72 | 3 | Endothelial Cells |
| RASGRP3 | 0.69 | 0 | 0 | 0.34 | 0.03 | 3 | Endothelial Cells |
| RASIP1 | 0.68 | 0 | 0 | 0.41 | 0.07 | 3 | Endothelial Cells |
| HOXA5 | 0.68 | 0 | 0 | 0.53 | 0.21 | 3 | Endothelial Cells |
| UACA | 0.68 | 0 | 0 | 0.55 | 0.22 | 3 | Endothelial Cells |
| ETS1 | 0.68 | 0 | 0 | 0.45 | 0.12 | 3 | Endothelial Cells |
| FCGRT | 0.68 | 0 | 0 | 0.83 | 0.54 | 3 | Endothelial Cells |
| TMEM123 | 0.68 | 0 | 0 | 0.65 | 0.35 | 3 | Endothelial Cells |
| TSPAN15 | 0.68 | 0 | 0 | 0.45 | 0.10 | 3 | Endothelial Cells |
| TNFRSF4 | 0.68 | 0 | 0 | 0.35 | 0.05 | 3 | Endothelial Cells |
| WWTR1 | 0.68 | 0 | 0 | 0.49 | 0.18 | 3 | Endothelial Cells |
| HEY1 | 0.67 | 0 | 0 | 0.30 | 0.02 | 3 | Endothelial Cells |
| MEIS2 | 0.67 | 0 | 0 | 0.39 | 0.09 | 3 | Endothelial Cells |
| PRKCH | 0.67 | 0 | 0 | 0.41 | 0.06 | 3 | Endothelial Cells |
| SPEF2 | 0.67 | 0 | 0 | 0.29 | 0.03 | 3 | Endothelial Cells |
| TLE4 | 0.67 | 0 | 0 | 0.48 | 0.18 | 3 | Endothelial Cells |
| TMEM204 | 0.67 | 0 | 0 | 0.48 | 0.16 | 3 | Endothelial Cells |
| PRMT1 | 0.66 | 0 | 0 | 0.77 | 0.49 | 3 | Endothelial Cells |
| CDC25B | 0.66 | 0 | 0 | 0.33 | 0.09 | 3 | Endothelial Cells |
| ADGRL2 | 0.66 | 0 | 0 | 0.47 | 0.12 | 3 | Endothelial Cells |
| VAT1 | 0.66 | 0 | 0 | 0.52 | 0.14 | 3 | Endothelial Cells |
| C10orf10 | 0.66 | 0 | 0 | 0.43 | 0.13 | 3 | Endothelial Cells |
| FOXP1 | 0.66 | 0 | 0 | 0.67 | 0.37 | 3 | Endothelial Cells |
| LAP3 | 0.65 | 0 | 0 | 0.50 | 0.19 | 3 | Endothelial Cells |
| VAMP3 | 0.65 | 0 | 0 | 0.53 | 0.24 | 3 | Endothelial Cells |
| S100A3 | 0.65 | 0 | 0 | 0.32 | 0.09 | 3 | Endothelial Cells |
| HHEX | 0.65 | 0 | 0 | 0.38 | 0.07 | 3 | Endothelial Cells |
| HSPA1A | 0.65 | 0 | 0 | 0.86 | 0.62 | 3 | Endothelial Cells |
| PIM3 | 0.65 | 0 | 0 | 0.46 | 0.14 | 3 | Endothelial Cells |
| STMN1 | 0.65 | 0 | 0 | 0.78 | 0.50 | 3 | Endothelial Cells |
| RAPGEF4 | 0.64 | 0 | 0 | 0.33 | 0.03 | 3 | Endothelial Cells |
| FRY | 0.64 | 0 | 0 | 0.47 | 0.12 | 3 | Endothelial Cells |
| TPST2 | 0.64 | 0 | 0 | 0.46 | 0.11 | 3 | Endothelial Cells |
| SCARF1 | 0.64 | 0 | 0 | 0.37 | 0.05 | 3 | Endothelial Cells |
| MT-RNR1 | 0.64 | 0 | 0 | 0.97 | 0.95 | 3 | Endothelial Cells |
| QKI | 0.64 | 0 | 0 | 0.47 | 0.18 | 3 | Endothelial Cells |
| MRI1 | 0.63 | 0 | 0 | 0.37 | 0.09 | 3 | Endothelial Cells |
| HLA-F | 0.63 | 0 | 0 | 0.44 | 0.10 | 3 | Endothelial Cells |
| ADAM15 | 0.63 | 0 | 0 | 0.32 | 0.09 | 3 | Endothelial Cells |
| MMRN2 | 0.63 | 0 | 0 | 0.39 | 0.06 | 3 | Endothelial Cells |
| ANXA3 | 0.63 | 0 | 0 | 0.43 | 0.09 | 3 | Endothelial Cells |
| HSPA12B | 0.62 | 0 | 0 | 0.39 | 0.06 | 3 | Endothelial Cells |
| MESDC1 | 0.62 | 0 | 0 | 0.42 | 0.13 | 3 | Endothelial Cells |
| TMEM2 | 0.62 | 0 | 0 | 0.30 | 0.06 | 3 | Endothelial Cells |
| TSPAN12 | 0.62 | 0 | 0 | 0.46 | 0.12 | 3 | Endothelial Cells |
| MLF1 | 0.61 | 0 | 0 | 0.40 | 0.12 | 3 | Endothelial Cells |
| DYNLL1 | 0.61 | 0 | 0 | 0.92 | 0.77 | 3 | Endothelial Cells |
| IFI16 | 0.61 | 0 | 0 | 0.63 | 0.32 | 3 | Endothelial Cells |
| DACH1 | 0.61 | 0 | 0 | 0.34 | 0.05 | 3 | Endothelial Cells |
| TBX3 | 0.61 | 0 | 0 | 0.55 | 0.27 | 3 | Endothelial Cells |
| KIAA0355 | 0.61 | 0 | 0 | 0.44 | 0.15 | 3 | Endothelial Cells |
| FAM84A | 0.61 | 0 | 0 | 0.26 | 0.03 | 3 | Endothelial Cells |
| FLT1 | 0.61 | 0 | 0 | 0.35 | 0.05 | 3 | Endothelial Cells |
| GIMAP6 | 0.6 | 0 | 0 | 0.36 | 0.05 | 3 | Endothelial Cells |
| DOCK9 | 0.6 | 0 | 0 | 0.33 | 0.04 | 3 | Endothelial Cells |
| C11orf31 | 0.6 | 0 | 0 | 0.77 | 0.53 | 3 | Endothelial Cells |
| KLF2 | 0.6 | 0 | 0 | 0.42 | 0.15 | 3 | Endothelial Cells |
| PRX | 0.6 | 0 | 0 | 0.39 | 0.06 | 3 | Endothelial Cells |
| RHOC | 0.6 | 0 | 0 | 0.84 | 0.58 | 3 | Endothelial Cells |
| ACTN4 | 0.6 | 0 | 0 | 0.70 | 0.37 | 3 | Endothelial Cells |
| AFAP1L1 | 0.59 | 0 | 0 | 0.34 | 0.05 | 3 | Endothelial Cells |
| MYO6 | 0.59 | 0 | 0 | 0.40 | 0.11 | 3 | Endothelial Cells |
| MYL12A | 0.58 | 0 | 0 | 0.91 | 0.73 | 3 | Endothelial Cells |
| SEMA6A | 0.58 | 0 | 0 | 0.33 | 0.05 | 3 | Endothelial Cells |
| SPHK1 | 0.58 | 0 | 0 | 0.30 | 0.04 | 3 | Endothelial Cells |
| bP-2189O9.2 | 0.58 | 0 | 0 | 0.31 | 0.04 | 3 | Endothelial Cells |
| GATA2 | 0.58 | 0 | 0 | 0.32 | 0.04 | 3 | Endothelial Cells |
| TSPAN4 | 0.57 | 0 | 0 | 0.59 | 0.28 | 3 | Endothelial Cells |
| COL4A1 | 0.57 | 0 | 0 | 0.84 | 0.50 | 3 | Endothelial Cells |
| APOL1 | 0.57 | 0 | 0 | 0.35 | 0.06 | 3 | Endothelial Cells |
| FAM69B | 0.57 | 0 | 0 | 0.30 | 0.03 | 3 | Endothelial Cells |
| NCKAP5 | 0.57 | 0 | 0 | 0.29 | 0.05 | 3 | Endothelial Cells |
| EFNB1 | 0.57 | 0 | 0 | 0.38 | 0.10 | 3 | Endothelial Cells |
| ECE1 | 0.56 | 0 | 0 | 0.41 | 0.13 | 3 | Endothelial Cells |
| PARVB | 0.56 | 0 | 0 | 0.42 | 0.12 | 3 | Endothelial Cells |
| NAA38 | 0.56 | 0 | 0 | 0.61 | 0.36 | 3 | Endothelial Cells |
| TSPAN13 | 0.56 | 0 | 0 | 0.41 | 0.11 | 3 | Endothelial Cells |
| HDAC7 | 0.55 | 0 | 0 | 0.42 | 0.14 | 3 | Endothelial Cells |
| CFAP20 | 0.55 | 0 | 0 | 0.52 | 0.23 | 3 | Endothelial Cells |
| LUZP1 | 0.55 | 0 | 0 | 0.45 | 0.15 | 3 | Endothelial Cells |
| SVIP | 0.55 | 0 | 0 | 0.40 | 0.13 | 3 | Endothelial Cells |
| TMEM255B | 0.55 | 0 | 0 | 0.30 | 0.03 | 3 | Endothelial Cells |
| LYL1 | 0.55 | 0 | 0 | 0.37 | 0.08 | 3 | Endothelial Cells |
| PEA15 | 0.55 | 0 | 0 | 0.53 | 0.24 | 3 | Endothelial Cells |
| IFNGR1 | 0.55 | 0 | 0 | 0.51 | 0.25 | 3 | Endothelial Cells |
| S100A10 | 0.55 | 0 | 0 | 0.94 | 0.62 | 3 | Endothelial Cells |
| BTN3A2 | 0.55 | 0 | 0 | 0.37 | 0.09 | 3 | Endothelial Cells |
| TMEM140 | 0.55 | 0 | 0 | 0.31 | 0.06 | 3 | Endothelial Cells |
| CMTM3 | 0.54 | 0 | 0 | 0.55 | 0.25 | 3 | Endothelial Cells |
| SHE | 0.54 | 0 | 0 | 0.28 | 0.03 | 3 | Endothelial Cells |
| LRRC32 | 0.54 | 0 | 0 | 0.34 | 0.11 | 3 | Endothelial Cells |
| ISG15 | 0.54 | 0 | 0 | 0.53 | 0.24 | 3 | Endothelial Cells |
| NRN1 | 0.54 | 0 | 0 | 0.35 | 0.06 | 3 | Endothelial Cells |
| PLXNC1 | 0.53 | 0 | 0 | 0.28 | 0.03 | 3 | Endothelial Cells |
| CLEC3B | 0.53 | 0 | 0 | 0.26 | 0.03 | 3 | Endothelial Cells |
| HSPA1B | 0.53 | 0 | 0 | 0.67 | 0.35 | 3 | Endothelial Cells |
| FAM101B | 0.53 | 0 | 0 | 0.26 | 0.04 | 3 | Endothelial Cells |
| NUDT14 | 0.53 | 0 | 0 | 0.32 | 0.07 | 3 | Endothelial Cells |
| PON2 | 0.53 | 0 | 0 | 0.48 | 0.22 | 3 | Endothelial Cells |
| SNX3 | 0.53 | 0 | 0 | 0.85 | 0.68 | 3 | Endothelial Cells |
| MLLT4 | 0.52 | 0 | 0 | 0.43 | 0.14 | 3 | Endothelial Cells |
| IGF2BP2 | 0.52 | 0 | 0 | 0.41 | 0.12 | 3 | Endothelial Cells |
| APCDD1 | 0.52 | 0 | 0 | 0.31 | 0.07 | 3 | Endothelial Cells |
| PDE3B | 0.52 | 0 | 0 | 0.28 | 0.05 | 3 | Endothelial Cells |
| ZEB1 | 0.52 | 0 | 0 | 0.40 | 0.12 | 3 | Endothelial Cells |
| STARD3 | 0.52 | 0 | 0 | 0.34 | 0.10 | 3 | Endothelial Cells |
| MAPK3 | 0.52 | 0 | 0 | 0.41 | 0.14 | 3 | Endothelial Cells |
| FXYD6 | 0.52 | 0 | 0 | 0.65 | 0.39 | 3 | Endothelial Cells |
| CASKIN2 | 0.52 | 0 | 0 | 0.34 | 0.07 | 3 | Endothelial Cells |
| POLR2L | 0.51 | 0 | 0 | 0.85 | 0.66 | 3 | Endothelial Cells |
| PPP2R5A | 0.51 | 0 | 0 | 0.33 | 0.08 | 3 | Endothelial Cells |
| KLF4 | 0.51 | 0 | 0 | 0.38 | 0.14 | 3 | Endothelial Cells |
| CASZ1 | 0.51 | 0 | 0 | 0.28 | 0.04 | 3 | Endothelial Cells |
| GUK1 | 0.51 | 0 | 0 | 0.81 | 0.66 | 3 | Endothelial Cells |
| PSME1 | 0.51 | 0 | 0 | 0.78 | 0.58 | 3 | Endothelial Cells |
| ATOX1 | 0.51 | 0 | 0 | 0.61 | 0.38 | 3 | Endothelial Cells |
| ABHD17A | 0.51 | 0 | 0 | 0.39 | 0.13 | 3 | Endothelial Cells |
| STK4 | 0.51 | 0 | 0 | 0.38 | 0.14 | 3 | Endothelial Cells |
| SLC26A2 | 0.51 | 0 | 0 | 0.26 | 0.06 | 3 | Endothelial Cells |
| MAP4K2 | 0.5 | 0 | 0 | 0.29 | 0.05 | 3 | Endothelial Cells |
| SNRK | 0.5 | 0 | 0 | 0.35 | 0.11 | 3 | Endothelial Cells |
| CDC37 | 0.5 | 0 | 0 | 0.70 | 0.47 | 3 | Endothelial Cells |
| SERPINH1 | 0.5 | 0 | 0 | 0.79 | 0.55 | 3 | Endothelial Cells |
| PLEKHA1 | 0.5 | 0 | 0 | 0.41 | 0.15 | 3 | Endothelial Cells |
| PTPRM | 0.5 | 0 | 0 | 0.29 | 0.06 | 3 | Endothelial Cells |
| XAF1 | 0.5 | 0 | 0 | 0.33 | 0.07 | 3 | Endothelial Cells |
| CCDC130 | 0.5 | 0 | 0 | 0.34 | 0.11 | 3 | Endothelial Cells |
| ITGA5 | 0.5 | 0 | 0 | 0.36 | 0.11 | 3 | Endothelial Cells |
| CD320 | 0.5 | 0 | 0 | 0.37 | 0.13 | 3 | Endothelial Cells |
| N4BP3 | 0.5 | 0 | 0 | 0.27 | 0.02 | 3 | Endothelial Cells |
| TSPAN18 | 0.5 | 0 | 0 | 0.30 | 0.06 | 3 | Endothelial Cells |
| GJA4 | 0.5 | 0 | 0 | 0.32 | 0.14 | 3 | Endothelial Cells |
| RBM17 | 0.49 | 0 | 0 | 0.58 | 0.35 | 3 | Endothelial Cells |
| FXYD5 | 0.49 | 0 | 0 | 0.56 | 0.23 | 3 | Endothelial Cells |
| DPYSL3 | 0.49 | 0 | 0 | 0.39 | 0.13 | 3 | Endothelial Cells |
| NEDD4L | 0.49 | 0 | 0 | 0.25 | 0.05 | 3 | Endothelial Cells |
| YES1 | 0.49 | 0 | 0 | 0.32 | 0.09 | 3 | Endothelial Cells |
| HEG1 | 0.49 | 0 | 0 | 0.41 | 0.11 | 3 | Endothelial Cells |
| HPCAL1 | 0.49 | 0 | 0 | 0.38 | 0.13 | 3 | Endothelial Cells |
| HOMER3 | 0.49 | 0 | 0 | 0.30 | 0.07 | 3 | Endothelial Cells |
| GALNT18 | 0.48 | 0 | 0 | 0.34 | 0.10 | 3 | Endothelial Cells |
| COL4A2 | 0.48 | 0 | 0 | 0.82 | 0.52 | 3 | Endothelial Cells |
| NOTCH1 | 0.48 | 0 | 0 | 0.28 | 0.06 | 3 | Endothelial Cells |
| CDC42EP1 | 0.48 | 0 | 0 | 0.46 | 0.23 | 3 | Endothelial Cells |
| OCIAD2 | 0.48 | 0 | 0 | 0.38 | 0.12 | 3 | Endothelial Cells |
| SLC44A2 | 0.48 | 0 | 0 | 0.41 | 0.15 | 3 | Endothelial Cells |
| APH1A | 0.48 | 0 | 0 | 0.53 | 0.30 | 3 | Endothelial Cells |
| LRRFIP1 | 0.48 | 0 | 0 | 0.52 | 0.25 | 3 | Endothelial Cells |
| PSME2 | 0.48 | 0 | 0 | 0.66 | 0.43 | 3 | Endothelial Cells |
| HSPB8 | 0.48 | 0 | 0 | 0.30 | 0.09 | 3 | Endothelial Cells |
| TJP1 | 0.48 | 0 | 0 | 0.54 | 0.26 | 3 | Endothelial Cells |
| UBE2L6 | 0.48 | 0 | 0 | 0.48 | 0.22 | 3 | Endothelial Cells |
| ADGRL3 | 0.47 | 0 | 0 | 0.28 | 0.05 | 3 | Endothelial Cells |
| MEF2A | 0.47 | 0 | 0 | 0.39 | 0.15 | 3 | Endothelial Cells |
| NEDD9 | 0.47 | 0 | 0 | 0.38 | 0.12 | 3 | Endothelial Cells |
| DHRS3 | 0.47 | 0 | 0 | 0.34 | 0.09 | 3 | Endothelial Cells |
| SHROOM4 | 0.47 | 0 | 0 | 0.27 | 0.04 | 3 | Endothelial Cells |
| CD40 | 0.47 | 0 | 0 | 0.32 | 0.08 | 3 | Endothelial Cells |
| JUP | 0.47 | 0 | 0 | 0.41 | 0.14 | 3 | Endothelial Cells |
| PAK4 | 0.47 | 0 | 0 | 0.32 | 0.08 | 3 | Endothelial Cells |
| TSC22D1 | 0.46 | 0 | 0 | 0.79 | 0.59 | 3 | Endothelial Cells |
| CPNE2 | 0.46 | 0 | 0 | 0.28 | 0.07 | 3 | Endothelial Cells |
| NECAP2 | 0.46 | 0 | 0 | 0.45 | 0.20 | 3 | Endothelial Cells |
| RGL2 | 0.46 | 0 | 0 | 0.40 | 0.17 | 3 | Endothelial Cells |
| DGKH | 0.46 | 0 | 0 | 0.30 | 0.07 | 3 | Endothelial Cells |
| SASH1 | 0.46 | 0 | 0 | 0.40 | 0.12 | 3 | Endothelial Cells |
| CTNNB1 | 0.46 | 0 | 0 | 0.55 | 0.36 | 3 | Endothelial Cells |
| TSPAN5 | 0.46 | 0 | 0 | 0.26 | 0.04 | 3 | Endothelial Cells |
| PTTG1IP | 0.46 | 0 | 0 | 0.54 | 0.31 | 3 | Endothelial Cells |
| SCHIP1.1 | 0.46 | 0 | 0 | 0.31 | 0.09 | 3 | Endothelial Cells |
| SH2D3C | 0.46 | 0 | 0 | 0.35 | 0.09 | 3 | Endothelial Cells |
| CLIC1 | 0.46 | 0 | 0 | 0.83 | 0.62 | 3 | Endothelial Cells |
| SYNGR2 | 0.45 | 0 | 0 | 0.49 | 0.25 | 3 | Endothelial Cells |
| DOCK6 | 0.45 | 0 | 0 | 0.35 | 0.13 | 3 | Endothelial Cells |
| PHACTR2 | 0.45 | 0 | 0 | 0.48 | 0.24 | 3 | Endothelial Cells |
| AHR | 0.45 | 0 | 0 | 0.31 | 0.10 | 3 | Endothelial Cells |
| ELK4 | 0.45 | 0 | 0 | 0.33 | 0.11 | 3 | Endothelial Cells |
| HSPG2 | 0.44 | 0 | 0 | 0.44 | 0.17 | 3 | Endothelial Cells |
| LAMTOR2 | 0.44 | 0 | 0 | 0.48 | 0.26 | 3 | Endothelial Cells |
| XIST | 0.44 | 0 | 0 | 0.84 | 0.69 | 3 | Endothelial Cells |
| LPAR6 | 0.44 | 0 | 0 | 0.32 | 0.11 | 3 | Endothelial Cells |
| ARGLU1 | 0.43 | 0 | 0 | 0.63 | 0.45 | 3 | Endothelial Cells |
| SLFN5 | 0.43 | 0 | 0 | 0.29 | 0.07 | 3 | Endothelial Cells |
| SERPINB9 | 0.43 | 0 | 0 | 0.27 | 0.08 | 3 | Endothelial Cells |
| YWHAE | 0.43 | 0 | 0 | 0.76 | 0.62 | 3 | Endothelial Cells |
| NRP1 | 0.43 | 0 | 0 | 0.43 | 0.18 | 3 | Endothelial Cells |
| EPHB4 | 0.43 | 0 | 0 | 0.28 | 0.06 | 3 | Endothelial Cells |
| OCLN | 0.43 | 0 | 0 | 0.27 | 0.05 | 3 | Endothelial Cells |
| TAX1BP1 | 0.43 | 0 | 0 | 0.49 | 0.32 | 3 | Endothelial Cells |
| LSR | 0.43 | 0 | 0 | 0.30 | 0.08 | 3 | Endothelial Cells |
| SPTAN1 | 0.42 | 0 | 0 | 0.44 | 0.20 | 3 | Endothelial Cells |
| GLUL | 0.42 | 0 | 0 | 0.65 | 0.40 | 3 | Endothelial Cells |
| RAC1 | 0.42 | 0 | 0 | 0.90 | 0.80 | 3 | Endothelial Cells |
| SMAGP | 0.42 | 0 | 0 | 0.25 | 0.06 | 3 | Endothelial Cells |
| GPX1 | 0.42 | 0 | 0 | 0.73 | 0.51 | 3 | Endothelial Cells |
| FRYL | 0.42 | 0 | 0 | 0.27 | 0.09 | 3 | Endothelial Cells |
| IRF1 | 0.42 | 0 | 0 | 0.39 | 0.18 | 3 | Endothelial Cells |
| TSPAN14 | 0.42 | 0 | 0 | 0.33 | 0.12 | 3 | Endothelial Cells |
| DUSP1 | 0.42 | 0 | 0 | 0.61 | 0.39 | 3 | Endothelial Cells |
| RALGDS | 0.42 | 0 | 0 | 0.26 | 0.06 | 3 | Endothelial Cells |
| MESDC2 | 0.42 | 0 | 0 | 0.48 | 0.25 | 3 | Endothelial Cells |
| C10orf11 | 0.42 | 0 | 0 | 0.26 | 0.06 | 3 | Endothelial Cells |
| PSMB10 | 0.42 | 0 | 0 | 0.45 | 0.24 | 3 | Endothelial Cells |
| IFIT3 | 0.42 | 0 | 0 | 0.27 | 0.08 | 3 | Endothelial Cells |
| AP1S2 | 0.42 | 0 | 0 | 0.42 | 0.22 | 3 | Endothelial Cells |
| FYN | 0.42 | 0 | 0 | 0.35 | 0.11 | 3 | Endothelial Cells |
| FAM65A | 0.42 | 0 | 0 | 0.28 | 0.07 | 3 | Endothelial Cells |
| PREX1 | 0.42 | 0 | 0 | 0.28 | 0.06 | 3 | Endothelial Cells |
| MFNG | 0.41 | 0 | 0 | 0.25 | 0.05 | 3 | Endothelial Cells |
| ADD3 | 0.41 | 0 | 0 | 0.45 | 0.21 | 3 | Endothelial Cells |
| FAM89A | 0.41 | 0 | 0 | 0.26 | 0.04 | 3 | Endothelial Cells |
| SLC12A2 | 0.41 | 0 | 0 | 0.31 | 0.10 | 3 | Endothelial Cells |
| RP3-449O17.1 | 0.41 | 0 | 0 | 0.26 | 0.06 | 3 | Endothelial Cells |
| GSDMD | 0.41 | 0 | 0 | 0.39 | 0.17 | 3 | Endothelial Cells |
| ELMO1 | 0.41 | 0 | 0 | 0.26 | 0.08 | 3 | Endothelial Cells |
| ARHGEF3 | 0.41 | 0 | 0 | 0.27 | 0.06 | 3 | Endothelial Cells |
| PTPRN2 | 0.41 | 0 | 0 | 0.26 | 0.05 | 3 | Endothelial Cells |
| PXN | 0.41 | 0 | 0 | 0.29 | 0.08 | 3 | Endothelial Cells |
| GBP1 | 0.41 | 0 | 0 | 0.26 | 0.11 | 3 | Endothelial Cells |
| PLXND1 | 0.41 | 0 | 0 | 0.35 | 0.11 | 3 | Endothelial Cells |
| ZNHIT1 | 0.41 | 0 | 0 | 0.61 | 0.43 | 3 | Endothelial Cells |
| SIGIRR | 0.4 | 0 | 0 | 0.37 | 0.15 | 3 | Endothelial Cells |
| DAD1 | 0.4 | 0 | 0 | 0.78 | 0.63 | 3 | Endothelial Cells |
| TMSB4X | 0.4 | 0 | 0 | 1.00 | 1.00 | 3 | Endothelial Cells |
| REEP3 | 0.4 | 0 | 0 | 0.33 | 0.14 | 3 | Endothelial Cells |
| FNBP1L | 0.4 | 0 | 0 | 0.27 | 0.08 | 3 | Endothelial Cells |
| RPS6KA2 | 0.4 | 0 | 0 | 0.30 | 0.09 | 3 | Endothelial Cells |
| STX12 | 0.4 | 0 | 0 | 0.39 | 0.19 | 3 | Endothelial Cells |
| HMBOX1 | 0.4 | 0 | 0 | 0.32 | 0.13 | 3 | Endothelial Cells |
| NECTIN2 | 0.4 | 0 | 0 | 0.42 | 0.22 | 3 | Endothelial Cells |
| CALR | 0.4 | 0 | 0 | 0.76 | 0.56 | 3 | Endothelial Cells |
| CDKN1C | 0.39 | 0 | 0 | 0.37 | 0.19 | 3 | Endothelial Cells |
| TRIM56 | 0.39 | 0 | 0 | 0.39 | 0.20 | 3 | Endothelial Cells |
| DGKZ | 0.39 | 0 | 0 | 0.33 | 0.11 | 3 | Endothelial Cells |
| PPM1F | 0.39 | 0 | 0 | 0.25 | 0.06 | 3 | Endothelial Cells |
| FIS1 | 0.39 | 0 | 0 | 0.67 | 0.49 | 3 | Endothelial Cells |
| LGALS9 | 0.39 | 0 | 0 | 0.29 | 0.08 | 3 | Endothelial Cells |
| AAMDC | 0.39 | 0 | 0 | 0.44 | 0.25 | 3 | Endothelial Cells |
| TXNIP | 0.39 | 0 | 0 | 0.86 | 0.66 | 3 | Endothelial Cells |
| TNFRSF14 | 0.39 | 0 | 0 | 0.34 | 0.10 | 3 | Endothelial Cells |
| RGS12 | 0.39 | 0 | 0 | 0.30 | 0.10 | 3 | Endothelial Cells |
| TNFAIP1 | 0.39 | 0 | 0 | 0.25 | 0.08 | 3 | Endothelial Cells |
| UTRN | 0.39 | 0 | 0 | 0.35 | 0.17 | 3 | Endothelial Cells |
| HERPUD1 | 0.39 | 0 | 0 | 0.59 | 0.40 | 3 | Endothelial Cells |
| DDX17 | 0.39 | 0 | 0 | 0.59 | 0.43 | 3 | Endothelial Cells |
| CHST12 | 0.39 | 0 | 0 | 0.45 | 0.22 | 3 | Endothelial Cells |
| TMEM219 | 0.39 | 0 | 0 | 0.50 | 0.30 | 3 | Endothelial Cells |
| NDUFA8 | 0.38 | 0 | 0 | 0.44 | 0.25 | 3 | Endothelial Cells |
| RP11-467L13.5 | 0.38 | 0 | 0 | 0.34 | 0.18 | 3 | Endothelial Cells |
| KTN1 | 0.38 | 0 | 0 | 0.71 | 0.54 | 3 | Endothelial Cells |
| NTHL1 | 0.38 | 0 | 0 | 0.31 | 0.13 | 3 | Endothelial Cells |
| PHACTR4 | 0.38 | 0 | 0 | 0.31 | 0.13 | 3 | Endothelial Cells |
| PPP3CA | 0.38 | 0 | 0 | 0.37 | 0.17 | 3 | Endothelial Cells |
| TMEM179B | 0.38 | 0 | 0 | 0.46 | 0.26 | 3 | Endothelial Cells |
| CFDP1 | 0.38 | 0 | 0 | 0.57 | 0.42 | 3 | Endothelial Cells |
| SERINC3 | 0.37 | 0 | 0 | 0.42 | 0.23 | 3 | Endothelial Cells |
| ERBIN | 0.37 | 0 | 0 | 0.31 | 0.11 | 3 | Endothelial Cells |
| IFI35 | 0.37 | 0 | 0 | 0.32 | 0.12 | 3 | Endothelial Cells |
| COX17 | 0.37 | 0 | 0 | 0.52 | 0.33 | 3 | Endothelial Cells |
| NFIB | 0.37 | 0 | 0 | 0.69 | 0.48 | 3 | Endothelial Cells |
| RAB5C | 0.37 | 0 | 0 | 0.60 | 0.43 | 3 | Endothelial Cells |
| BAZ2B | 0.37 | 0 | 0 | 0.39 | 0.21 | 3 | Endothelial Cells |
| CSTB | 0.37 | 0 | 0 | 0.68 | 0.47 | 3 | Endothelial Cells |
| EPN2 | 0.37 | 0 | 0 | 0.29 | 0.11 | 3 | Endothelial Cells |
| TSPAN9 | 0.36 | 0 | 0 | 0.26 | 0.09 | 3 | Endothelial Cells |
| SKAP2 | 0.36 | 0 | 0 | 0.41 | 0.24 | 3 | Endothelial Cells |
| HLX | 0.36 | 0 | 0 | 0.30 | 0.11 | 3 | Endothelial Cells |
| NKTR | 0.36 | 0 | 0 | 0.56 | 0.38 | 3 | Endothelial Cells |
| NOP10 | 0.36 | 0 | 0 | 0.66 | 0.46 | 3 | Endothelial Cells |
| NDUFA3 | 0.36 | 0 | 0 | 0.64 | 0.48 | 3 | Endothelial Cells |
| ARHGEF12 | 0.36 | 0 | 0 | 0.46 | 0.23 | 3 | Endothelial Cells |
| DCAF7 | 0.36 | 0 | 0 | 0.39 | 0.21 | 3 | Endothelial Cells |
| SLCO2A1 | 0.36 | 0 | 0 | 0.36 | 0.08 | 3 | Endothelial Cells |
| CYB561 | 0.36 | 0 | 0 | 0.25 | 0.05 | 3 | Endothelial Cells |
| ATP1B3 | 0.36 | 0 | 0 | 0.49 | 0.29 | 3 | Endothelial Cells |
| PAPSS2 | 0.35 | 0 | 0 | 0.27 | 0.09 | 3 | Endothelial Cells |
| ARRB1 | 0.35 | 0 | 0 | 0.26 | 0.07 | 3 | Endothelial Cells |
| SARAF | 0.35 | 0 | 0 | 0.73 | 0.63 | 3 | Endothelial Cells |
| RNF7 | 0.35 | 0 | 0 | 0.58 | 0.40 | 3 | Endothelial Cells |
| SOX4 | 0.35 | 0 | 0 | 0.83 | 0.75 | 3 | Endothelial Cells |
| DNAJA1 | 0.35 | 0 | 0 | 0.64 | 0.47 | 3 | Endothelial Cells |
| TMEM176B | 0.35 | 0 | 0 | 0.27 | 0.17 | 3 | Endothelial Cells |
| ARL2 | 0.35 | 0 | 0 | 0.52 | 0.33 | 3 | Endothelial Cells |
| TPM3 | 0.35 | 0 | 0 | 0.55 | 0.28 | 3 | Endothelial Cells |
| MLLT6 | 0.35 | 0 | 0 | 0.26 | 0.09 | 3 | Endothelial Cells |
| HIPK2 | 0.35 | 0 | 0 | 0.26 | 0.10 | 3 | Endothelial Cells |
| OPTN | 0.35 | 0 | 0 | 0.37 | 0.18 | 3 | Endothelial Cells |
| C9orf16 | 0.35 | 0 | 0 | 0.47 | 0.28 | 3 | Endothelial Cells |
| JAM3 | 0.35 | 0 | 0 | 0.26 | 0.11 | 3 | Endothelial Cells |
| PTMA | 0.35 | 0 | 0 | 1.00 | 1.00 | 3 | Endothelial Cells |
| SRGAP2B | 0.35 | 0 | 0 | 0.30 | 0.12 | 3 | Endothelial Cells |
| FRMD4A | 0.35 | 0 | 0 | 0.28 | 0.12 | 3 | Endothelial Cells |
| S1PR1 | 0.35 | 0 | 0 | 0.32 | 0.12 | 3 | Endothelial Cells |
| IER2 | 0.34 | 0 | 0 | 0.65 | 0.44 | 3 | Endothelial Cells |
| TAPBP | 0.34 | 0 | 0 | 0.41 | 0.24 | 3 | Endothelial Cells |
| TAP1 | 0.34 | 0 | 0 | 0.29 | 0.11 | 3 | Endothelial Cells |
| STAT1 | 0.34 | 0 | 0 | 0.26 | 0.11 | 3 | Endothelial Cells |
| PTPN12 | 0.34 | 0 | 0 | 0.30 | 0.13 | 3 | Endothelial Cells |
| MYL12B | 0.34 | 0 | 0 | 0.85 | 0.77 | 3 | Endothelial Cells |
| TTC28 | 0.34 | 0 | 0 | 0.28 | 0.12 | 3 | Endothelial Cells |
| MAPRE2 | 0.34 | 0 | 0 | 0.28 | 0.12 | 3 | Endothelial Cells |
| CRIP2 | 0.34 | 0 | 0 | 0.84 | 0.64 | 3 | Endothelial Cells |
| SNHG7 | 0.34 | 0 | 0 | 0.57 | 0.42 | 3 | Endothelial Cells |
| NUMB | 0.34 | 0 | 0 | 0.36 | 0.18 | 3 | Endothelial Cells |
| EBPL | 0.34 | 0 | 0 | 0.41 | 0.21 | 3 | Endothelial Cells |
| SDCBP | 0.34 | 0 | 0 | 0.69 | 0.51 | 3 | Endothelial Cells |
| ZNF503 | 0.34 | 0 | 0 | 0.48 | 0.33 | 3 | Endothelial Cells |
| TAX1BP3 | 0.34 | 0 | 0 | 0.53 | 0.35 | 3 | Endothelial Cells |
| RP11-693N9.2 | 0.34 | 0 | 0 | 0.27 | 0.15 | 3 | Endothelial Cells |
| MRPL17 | 0.34 | 0 | 0 | 0.30 | 0.13 | 3 | Endothelial Cells |
| TMBIM6 | 0.33 | 0 | 0 | 0.79 | 0.66 | 3 | Endothelial Cells |
| SDF2L1 | 0.33 | 0 | 0 | 0.38 | 0.21 | 3 | Endothelial Cells |
| CD59 | 0.33 | 0 | 0 | 0.69 | 0.60 | 3 | Endothelial Cells |
| LINC01420 | 0.33 | 0 | 0 | 0.47 | 0.33 | 3 | Endothelial Cells |
| TNIP1 | 0.33 | 0 | 0 | 0.36 | 0.18 | 3 | Endothelial Cells |
| CCDC186 | 0.33 | 0 | 0 | 0.28 | 0.11 | 3 | Endothelial Cells |
| YWHAB | 0.33 | 0 | 0 | 0.72 | 0.57 | 3 | Endothelial Cells |
| MAGEH1 | 0.33 | 0 | 0 | 0.39 | 0.26 | 3 | Endothelial Cells |
| TRIOBP | 0.33 | 0 | 0 | 0.36 | 0.18 | 3 | Endothelial Cells |
| HIF3A | 0.33 | 0 | 0 | 0.30 | 0.12 | 3 | Endothelial Cells |
| IMP3 | 0.32 | 0 | 0 | 0.37 | 0.22 | 3 | Endothelial Cells |
| S100A6 | 0.32 | 0 | 0 | 0.90 | 0.84 | 3 | Endothelial Cells |
| ATP5E | 0.32 | 0 | 0 | 0.94 | 0.87 | 3 | Endothelial Cells |
| TAF11 | 0.32 | 0 | 0 | 0.36 | 0.19 | 3 | Endothelial Cells |
| PSMB3 | 0.32 | 0 | 0 | 0.67 | 0.52 | 3 | Endothelial Cells |
| JUN | 0.32 | 0 | 0 | 0.78 | 0.67 | 3 | Endothelial Cells |
| RRAS | 0.32 | 0 | 0 | 0.30 | 0.14 | 3 | Endothelial Cells |
| RHOA | 0.32 | 0 | 0 | 0.79 | 0.69 | 3 | Endothelial Cells |
| PPP2R2A | 0.32 | 0 | 0 | 0.33 | 0.18 | 3 | Endothelial Cells |
| PSMA7 | 0.32 | 0 | 0 | 0.83 | 0.70 | 3 | Endothelial Cells |
| SREK1IP1 | 0.32 | 0 | 0 | 0.41 | 0.25 | 3 | Endothelial Cells |
| KRTCAP2 | 0.32 | 0 | 0 | 0.68 | 0.53 | 3 | Endothelial Cells |
| ADAR | 0.32 | 0 | 0 | 0.41 | 0.22 | 3 | Endothelial Cells |
| FAM43A | 0.32 | 0 | 0 | 0.29 | 0.13 | 3 | Endothelial Cells |
| PLSCR1 | 0.31 | 0 | 0 | 0.31 | 0.16 | 3 | Endothelial Cells |
| SIPA1L2 | 0.31 | 0 | 0 | 0.27 | 0.10 | 3 | Endothelial Cells |
| RPIA | 0.31 | 0 | 0 | 0.26 | 0.12 | 3 | Endothelial Cells |
| TMEM50A | 0.31 | 0 | 0 | 0.57 | 0.41 | 3 | Endothelial Cells |
| PARP14 | 0.31 | 0 | 0 | 0.27 | 0.09 | 3 | Endothelial Cells |
| PURA | 0.31 | 0 | 0 | 0.35 | 0.18 | 3 | Endothelial Cells |
| GUSBP3 | 0.3 | 0 | 0 | 0.32 | 0.13 | 3 | Endothelial Cells |
| SRGAP2C | 0.3 | 0 | 0 | 0.28 | 0.12 | 3 | Endothelial Cells |
| CAPZA2 | 0.3 | 0 | 0 | 0.66 | 0.50 | 3 | Endothelial Cells |
| ZNF22 | 0.3 | 0 | 0 | 0.49 | 0.31 | 3 | Endothelial Cells |
| GNB1 | 0.3 | 0 | 0 | 0.41 | 0.27 | 3 | Endothelial Cells |
| TMEM109 | 0.3 | 0 | 0 | 0.48 | 0.31 | 3 | Endothelial Cells |
| MAPKAPK5-AS1 | 0.3 | 0 | 0 | 0.27 | 0.11 | 3 | Endothelial Cells |
| MSN | 0.3 | 0 | 0 | 0.62 | 0.45 | 3 | Endothelial Cells |
| SHISA5 | 0.3 | 0 | 0 | 0.40 | 0.25 | 3 | Endothelial Cells |
| CD9 | 0.3 | 0 | 0 | 0.63 | 0.44 | 3 | Endothelial Cells |
| STOM | 0.3 | 0 | 0 | 0.63 | 0.42 | 3 | Endothelial Cells |
| TJP2 | 0.3 | 0 | 0 | 0.26 | 0.09 | 3 | Endothelial Cells |
| MARCKS | 0.3 | 0 | 0 | 0.66 | 0.51 | 3 | Endothelial Cells |
| MT-RNR2 | 0.29 | 0 | 0 | 1.00 | 1.00 | 3 | Endothelial Cells |
| WSB1 | 0.29 | 0 | 0 | 0.80 | 0.65 | 3 | Endothelial Cells |
| PFDN1 | 0.29 | 0 | 0 | 0.47 | 0.30 | 3 | Endothelial Cells |
| PSME2P2 | 0.29 | 0 | 0 | 0.35 | 0.20 | 3 | Endothelial Cells |
| SLC3A2 | 0.29 | 0 | 0 | 0.37 | 0.22 | 3 | Endothelial Cells |
| TPRKB | 0.29 | 0 | 0 | 0.34 | 0.20 | 3 | Endothelial Cells |
| HMGB1 | 0.29 | 0 | 0 | 0.96 | 0.92 | 3 | Endothelial Cells |
| PTBP1 | 0.29 | 0 | 0 | 0.42 | 0.26 | 3 | Endothelial Cells |
| SERPINB1 | 0.29 | 0 | 0 | 0.29 | 0.19 | 3 | Endothelial Cells |
| RYBP | 0.29 | 0 | 0 | 0.26 | 0.11 | 3 | Endothelial Cells |
| PDCD6IP | 0.29 | 0 | 0 | 0.33 | 0.18 | 3 | Endothelial Cells |
| ANP32A | 0.29 | 0 | 0 | 0.44 | 0.29 | 3 | Endothelial Cells |
| CFL1 | 0.29 | 0 | 0 | 0.96 | 0.90 | 3 | Endothelial Cells |
| SOCS3 | 0.29 | 0 | 0 | 0.29 | 0.18 | 3 | Endothelial Cells |
| FOXN3 | 0.29 | 0 | 0 | 0.37 | 0.21 | 3 | Endothelial Cells |
| RTN4 | 0.28 | 0 | 0 | 0.75 | 0.62 | 3 | Endothelial Cells |
| PSMB2 | 0.28 | 0 | 0 | 0.52 | 0.37 | 3 | Endothelial Cells |
| ALDH2 | 0.28 | 0 | 0 | 0.47 | 0.33 | 3 | Endothelial Cells |
| TMEM9B | 0.28 | 0 | 0 | 0.32 | 0.17 | 3 | Endothelial Cells |
| DYNC1LI2 | 0.28 | 0 | 0 | 0.41 | 0.26 | 3 | Endothelial Cells |
| LAPTM4A | 0.28 | 0 | 0 | 0.91 | 0.84 | 3 | Endothelial Cells |
| GABARAPL1 | 0.28 | 0 | 0 | 0.30 | 0.14 | 3 | Endothelial Cells |
| MYLIP | 0.28 | 0 | 0 | 0.46 | 0.28 | 3 | Endothelial Cells |
| SNAP23 | 0.28 | 0 | 0 | 0.33 | 0.18 | 3 | Endothelial Cells |
| CAMTA1 | 0.28 | 0 | 0 | 0.51 | 0.35 | 3 | Endothelial Cells |
| DYNC1H1 | 0.28 | 0 | 0 | 0.35 | 0.20 | 3 | Endothelial Cells |
| PCGF2 | 0.28 | 0 | 0 | 0.27 | 0.12 | 3 | Endothelial Cells |
| SERTAD1 | 0.28 | 0 | 0 | 0.33 | 0.17 | 3 | Endothelial Cells |
| DUSP11 | 0.28 | 0 | 0 | 0.25 | 0.12 | 3 | Endothelial Cells |
| GNAQ | 0.27 | 0 | 0 | 0.32 | 0.15 | 3 | Endothelial Cells |
| POMP | 0.27 | 0 | 0 | 0.73 | 0.61 | 3 | Endothelial Cells |
| EVA1B | 0.27 | 0 | 0 | 0.27 | 0.14 | 3 | Endothelial Cells |
| KLF3 | 0.27 | 0 | 0 | 0.26 | 0.12 | 3 | Endothelial Cells |
| UBE2L3 | 0.27 | 0 | 0 | 0.48 | 0.34 | 3 | Endothelial Cells |
| CD81 | 0.27 | 0 | 0 | 0.56 | 0.38 | 3 | Endothelial Cells |
| SCP2 | 0.27 | 0 | 0 | 0.54 | 0.42 | 3 | Endothelial Cells |
| DDIT4 | 0.27 | 0 | 0 | 0.29 | 0.15 | 3 | Endothelial Cells |
| TXNDC17 | 0.27 | 0 | 0 | 0.39 | 0.26 | 3 | Endothelial Cells |
| MTF2 | 0.27 | 0 | 0 | 0.28 | 0.15 | 3 | Endothelial Cells |
| AP2B1 | 0.27 | 0 | 0 | 0.29 | 0.17 | 3 | Endothelial Cells |
| PLEC | 0.27 | 0 | 0 | 0.30 | 0.15 | 3 | Endothelial Cells |
| SP100 | 0.27 | 0 | 0 | 0.35 | 0.18 | 3 | Endothelial Cells |
| UPF3A | 0.27 | 0 | 0 | 0.38 | 0.24 | 3 | Endothelial Cells |
| HES1 | 0.27 | 0 | 0 | 0.50 | 0.38 | 3 | Endothelial Cells |
| BCAP31 | 0.27 | 0 | 0 | 0.48 | 0.36 | 3 | Endothelial Cells |
| GRAMD1A | 0.27 | 0 | 0 | 0.26 | 0.13 | 3 | Endothelial Cells |
| STX10 | 0.27 | 0 | 0 | 0.33 | 0.17 | 3 | Endothelial Cells |
| MRPL52 | 0.27 | 0 | 0 | 0.48 | 0.35 | 3 | Endothelial Cells |
| ITSN2 | 0.26 | 0 | 0 | 0.29 | 0.15 | 3 | Endothelial Cells |
| PAK2 | 0.26 | 0 | 0 | 0.33 | 0.19 | 3 | Endothelial Cells |
| NDUFB3 | 0.26 | 0 | 0 | 0.48 | 0.35 | 3 | Endothelial Cells |
| BNIP3 | 0.26 | 0 | 0 | 0.31 | 0.15 | 3 | Endothelial Cells |
| CHMP2A | 0.26 | 0 | 0 | 0.58 | 0.46 | 3 | Endothelial Cells |
| CDK11B | 0.26 | 0 | 0 | 0.28 | 0.15 | 3 | Endothelial Cells |
| RNF213 | 0.26 | 0 | 0 | 0.35 | 0.18 | 3 | Endothelial Cells |
| TMEM258 | 0.26 | 0 | 0 | 0.79 | 0.71 | 3 | Endothelial Cells |
| PET100 | 0.26 | 0 | 0 | 0.63 | 0.50 | 3 | Endothelial Cells |
| MOB2 | 0.26 | 0 | 0 | 0.35 | 0.21 | 3 | Endothelial Cells |
| TUSC3 | 0.26 | 0 | 0 | 0.35 | 0.20 | 3 | Endothelial Cells |
| MANF | 0.26 | 0 | 0 | 0.43 | 0.31 | 3 | Endothelial Cells |
| ARPP19 | 0.26 | 0 | 0 | 0.35 | 0.21 | 3 | Endothelial Cells |
| SEPW1 | 0.26 | 0 | 0 | 0.65 | 0.55 | 3 | Endothelial Cells |
| BSG | 0.26 | 0 | 0 | 0.76 | 0.65 | 3 | Endothelial Cells |
| AK2 | 0.26 | 0 | 0 | 0.31 | 0.20 | 3 | Endothelial Cells |
| PDIA6 | 0.26 | 0 | 0 | 0.64 | 0.52 | 3 | Endothelial Cells |
| WDR60 | 0.26 | 0 | 0 | 0.30 | 0.17 | 3 | Endothelial Cells |
| TMEM173 | 0.25 | 0 | 0 | 0.37 | 0.22 | 3 | Endothelial Cells |
| YIPF3 | 0.25 | 0 | 0 | 0.46 | 0.32 | 3 | Endothelial Cells |
| CASP7 | 0.25 | 0 | 0 | 0.27 | 0.12 | 3 | Endothelial Cells |
| BAZ1A | 0.25 | 0 | 0 | 0.32 | 0.18 | 3 | Endothelial Cells |
| TIMM22 | 0.25 | 0 | 0 | 0.26 | 0.15 | 3 | Endothelial Cells |
| ORMDL2 | 0.25 | 0 | 0 | 0.28 | 0.14 | 3 | Endothelial Cells |
| TEAD2 | 0.25 | 0 | 0 | 0.30 | 0.16 | 3 | Endothelial Cells |
| SMARCA2 | 0.25 | 0 | 0 | 0.27 | 0.14 | 3 | Endothelial Cells |
| FAM162B | 2.77 | 0 | 0 | 0.91 | 0.13 | 4 | Pericytes |
| HIGD1B | 2.44 | 0 | 0 | 1.00 | 0.21 | 4 | Pericytes |
| NDUFA4L2 | 2.15 | 0 | 0 | 0.95 | 0.16 | 4 | Pericytes |
| COX4I2 | 2.12 | 0 | 0 | 0.98 | 0.20 | 4 | Pericytes |
| CHN1 | 1.96 | 0 | 0 | 0.96 | 0.20 | 4 | Pericytes |
| PDGFRB | 1.69 | 0 | 0 | 0.98 | 0.33 | 4 | Pericytes |
| CYGB | 1.63 | 0 | 0 | 0.77 | 0.10 | 4 | Pericytes |
| BGN | 1.54 | 0 | 0 | 0.98 | 0.47 | 4 | Pericytes |
| LPL | 1.53 | 0 | 0 | 0.75 | 0.13 | 4 | Pericytes |
| LHFP | 1.52 | 0 | 0 | 0.97 | 0.41 | 4 | Pericytes |
| PAG1 | 1.49 | 0 | 0 | 0.72 | 0.07 | 4 | Pericytes |
| GUCY1A2 | 1.45 | 0 | 0 | 0.76 | 0.08 | 4 | Pericytes |
| MEST | 1.4 | 0 | 0 | 0.92 | 0.33 | 4 | Pericytes |
| EGFL6 | 1.39 | 0 | 0 | 0.98 | 0.32 | 4 | Pericytes |
| SDC2 | 1.39 | 0 | 0 | 0.93 | 0.40 | 4 | Pericytes |
| IMPA2 | 1.38 | 0 | 0 | 0.64 | 0.09 | 4 | Pericytes |
| TPPP3 | 1.36 | 0 | 0 | 0.95 | 0.36 | 4 | Pericytes |
| GPX3 | 1.35 | 0 | 0 | 0.92 | 0.40 | 4 | Pericytes |
| THY1 | 1.26 | 0 | 0 | 0.69 | 0.13 | 4 | Pericytes |
| RFTN1 | 1.2 | 0 | 0 | 0.69 | 0.11 | 4 | Pericytes |
| PTN | 1.2 | 0 | 0 | 0.90 | 0.26 | 4 | Pericytes |
| KCNK3 | 1.18 | 0 | 0 | 0.59 | 0.07 | 4 | Pericytes |
| TACC1 | 1.16 | 0 | 0 | 0.91 | 0.42 | 4 | Pericytes |
| NID1 | 1.15 | 0 | 0 | 0.74 | 0.17 | 4 | Pericytes |
| GAS6 | 1.15 | 0 | 0 | 0.78 | 0.25 | 4 | Pericytes |
| COL4A1 | 1.14 | 0 | 0 | 0.92 | 0.49 | 4 | Pericytes |
| NOTCH3 | 1.11 | 0 | 0 | 0.76 | 0.18 | 4 | Pericytes |
| ITM2C | 1.09 | 0 | 0 | 0.86 | 0.36 | 4 | Pericytes |
| FOXS1 | 1.06 | 0 | 0 | 0.64 | 0.11 | 4 | Pericytes |
| COL4A2 | 1.06 | 0 | 0 | 0.91 | 0.51 | 4 | Pericytes |
| IGFBP7 | 1.05 | 0 | 0 | 0.99 | 0.64 | 4 | Pericytes |
| PCOLCE | 1.05 | 0 | 0 | 0.80 | 0.27 | 4 | Pericytes |
| DUSP4 | 1.03 | 0 | 0 | 0.48 | 0.03 | 4 | Pericytes |
| KCNK17 | 1.03 | 0 | 0 | 0.45 | 0.06 | 4 | Pericytes |
| RERG | 1.03 | 0 | 0 | 0.58 | 0.10 | 4 | Pericytes |
| PERP | 1.02 | 0 | 0 | 0.53 | 0.09 | 4 | Pericytes |
| SPRY1 | 1.02 | 0 | 0 | 0.89 | 0.39 | 4 | Pericytes |
| MYO1B | 1.02 | 0 | 0 | 0.69 | 0.16 | 4 | Pericytes |
| SPARC | 1.01 | 0 | 0 | 1.00 | 0.84 | 4 | Pericytes |
| UXS1 | 1 | 0 | 0 | 0.62 | 0.15 | 4 | Pericytes |
| LAMB1 | 0.98 | 0 | 0 | 0.58 | 0.11 | 4 | Pericytes |
| CADM1 | 0.98 | 0 | 0 | 0.58 | 0.14 | 4 | Pericytes |
| STOM | 0.97 | 0 | 0 | 0.86 | 0.40 | 4 | Pericytes |
| ITGA1 | 0.97 | 0 | 0 | 0.72 | 0.23 | 4 | Pericytes |
| 11-Sep | 0.97 | 0 | 0 | 0.74 | 0.27 | 4 | Pericytes |
| NDRG1 | 0.97 | 0 | 0 | 0.59 | 0.12 | 4 | Pericytes |
| MGLL | 0.96 | 0 | 0 | 0.65 | 0.20 | 4 | Pericytes |
| NCK2 | 0.96 | 0 | 0 | 0.52 | 0.08 | 4 | Pericytes |
| KIAA0040 | 0.94 | 0 | 0 | 0.50 | 0.09 | 4 | Pericytes |
| CCDC102B | 0.94 | 0 | 0 | 0.52 | 0.07 | 4 | Pericytes |
| PCDH18 | 0.94 | 0 | 0 | 0.50 | 0.07 | 4 | Pericytes |
| PDE8B | 0.93 | 0 | 0 | 0.45 | 0.02 | 4 | Pericytes |
| LAMC3 | 0.9 | 0 | 0 | 0.41 | 0.01 | 4 | Pericytes |
| TJP1 | 0.88 | 0 | 0 | 0.67 | 0.25 | 4 | Pericytes |
| NR2F2 | 0.88 | 0 | 0 | 0.76 | 0.31 | 4 | Pericytes |
| SGCE | 0.87 | 0 | 0 | 0.59 | 0.17 | 4 | Pericytes |
| RASL12 | 0.87 | 0 | 0 | 0.74 | 0.28 | 4 | Pericytes |
| ADAMTSL3 | 0.87 | 0 | 0 | 0.45 | 0.04 | 4 | Pericytes |
| CLEC11A | 0.87 | 0 | 0 | 0.58 | 0.17 | 4 | Pericytes |
| GJC1 | 0.86 | 0 | 0 | 0.49 | 0.09 | 4 | Pericytes |
| LURAP1L | 0.85 | 0 | 0 | 0.60 | 0.16 | 4 | Pericytes |
| GUCY1B3 | 0.85 | 0 | 0 | 0.52 | 0.11 | 4 | Pericytes |
| VSNL1 | 0.84 | 0 | 0 | 0.41 | 0.03 | 4 | Pericytes |
| P2RY14 | 0.83 | 0 | 0 | 0.38 | 0.01 | 4 | Pericytes |
| 7-Sep | 0.83 | 0 | 0 | 0.95 | 0.69 | 4 | Pericytes |
| PLCB1 | 0.83 | 0 | 0 | 0.43 | 0.05 | 4 | Pericytes |
| HEYL | 0.83 | 0 | 0 | 0.49 | 0.10 | 4 | Pericytes |
| HRH2 | 0.82 | 0 | 0 | 0.47 | 0.09 | 4 | Pericytes |
| EBF1 | 0.81 | 0 | 0 | 0.45 | 0.08 | 4 | Pericytes |
| TRPC6 | 0.81 | 0 | 0 | 0.42 | 0.04 | 4 | Pericytes |
| ANXA6 | 0.79 | 0 | 0 | 0.73 | 0.39 | 4 | Pericytes |
| NDRG2 | 0.78 | 0 | 0 | 0.70 | 0.30 | 4 | Pericytes |
| 4-Sep | 0.78 | 0 | 0 | 0.50 | 0.12 | 4 | Pericytes |
| TXNIP | 0.78 | 0 | 0 | 0.92 | 0.66 | 4 | Pericytes |
| GPM6B | 0.77 | 0 | 0 | 0.68 | 0.26 | 4 | Pericytes |
| PTEN | 0.77 | 0 | 0 | 0.52 | 0.16 | 4 | Pericytes |
| FARP1 | 0.76 | 0 | 0 | 0.44 | 0.10 | 4 | Pericytes |
| DCN | 0.75 | 0 | 0 | 0.95 | 0.51 | 4 | Pericytes |
| FERMT2 | 0.74 | 0 | 0 | 0.66 | 0.30 | 4 | Pericytes |
| MT2A | 0.71 | 0 | 0 | 0.94 | 0.60 | 4 | Pericytes |
| RGS3 | 0.71 | 0 | 0 | 0.54 | 0.25 | 4 | Pericytes |
| TBX2 | 0.7 | 0 | 0 | 0.71 | 0.32 | 4 | Pericytes |
| ADGRG6 | 0.7 | 0 | 0 | 0.43 | 0.10 | 4 | Pericytes |
| OLFM2 | 0.69 | 0 | 0 | 0.36 | 0.06 | 4 | Pericytes |
| ARHGDIB | 0.69 | 0 | 0 | 0.70 | 0.27 | 4 | Pericytes |
| FHL5 | 0.68 | 0 | 0 | 0.32 | 0.01 | 4 | Pericytes |
| SERPING1 | 0.68 | 0 | 0 | 0.75 | 0.36 | 4 | Pericytes |
| SH3KBP1 | 0.68 | 0 | 0 | 0.51 | 0.16 | 4 | Pericytes |
| GUCY1A3 | 0.67 | 0 | 0 | 0.59 | 0.22 | 4 | Pericytes |
| CCDC68 | 0.67 | 0 | 0 | 0.43 | 0.09 | 4 | Pericytes |
| NR2F1 | 0.67 | 0 | 0 | 0.67 | 0.24 | 4 | Pericytes |
| SLC35B3 | 0.66 | 0 | 0 | 0.38 | 0.07 | 4 | Pericytes |
| MCTP2 | 0.66 | 0 | 0 | 0.39 | 0.06 | 4 | Pericytes |
| CALM2 | 0.66 | 0 | 0 | 0.99 | 0.90 | 4 | Pericytes |
| KLHL23 | 0.65 | 0 | 0 | 0.49 | 0.17 | 4 | Pericytes |
| GRK5 | 0.65 | 0 | 0 | 0.45 | 0.17 | 4 | Pericytes |
| HSPA2 | 0.65 | 0 | 0 | 0.47 | 0.14 | 4 | Pericytes |
| PTGIR | 0.65 | 0 | 0 | 0.45 | 0.13 | 4 | Pericytes |
| KCP | 0.65 | 0 | 0 | 0.27 | 0.01 | 4 | Pericytes |
| MFGE8 | 0.64 | 0 | 0 | 0.72 | 0.30 | 4 | Pericytes |
| MEF2C | 0.64 | 0 | 0 | 0.74 | 0.40 | 4 | Pericytes |
| PPP1R14A | 0.64 | 0 | 0 | 0.77 | 0.40 | 4 | Pericytes |
| IGFBP3 | 0.64 | 0 | 0 | 0.29 | 0.06 | 4 | Pericytes |
| NR3C1 | 0.64 | 0 | 0 | 0.59 | 0.28 | 4 | Pericytes |
| CALD1 | 0.64 | 0 | 0 | 0.98 | 0.77 | 4 | Pericytes |
| PELO | 0.64 | 0 | 0 | 0.43 | 0.14 | 4 | Pericytes |
| ADGRD1 | 0.63 | 0 | 0 | 0.28 | 0.03 | 4 | Pericytes |
| ARHGEF17 | 0.63 | 0 | 0 | 0.36 | 0.08 | 4 | Pericytes |
| COL6A3 | 0.62 | 0 | 0 | 0.84 | 0.34 | 4 | Pericytes |
| CDH6 | 0.62 | 0 | 0 | 0.40 | 0.09 | 4 | Pericytes |
| TBX5 | 0.62 | 0 | 0 | 0.41 | 0.10 | 4 | Pericytes |
| NT5DC2 | 0.61 | 0 | 0 | 0.43 | 0.13 | 4 | Pericytes |
| ITIH5 | 0.61 | 0 | 0 | 0.35 | 0.08 | 4 | Pericytes |
| CD248 | 0.61 | 0 | 0 | 0.34 | 0.06 | 4 | Pericytes |
| PLCE1 | 0.6 | 0 | 0 | 0.34 | 0.08 | 4 | Pericytes |
| HACD4 | 0.6 | 0 | 0 | 0.33 | 0.06 | 4 | Pericytes |
| PPIC | 0.6 | 0 | 0 | 0.65 | 0.34 | 4 | Pericytes |
| EHD2 | 0.6 | 0 | 0 | 0.62 | 0.33 | 4 | Pericytes |
| ECM1 | 0.59 | 0 | 0 | 0.30 | 0.03 | 4 | Pericytes |
| MARCKS | 0.59 | 0 | 0 | 0.81 | 0.50 | 4 | Pericytes |
| H2AFJ | 0.59 | 0 | 0 | 0.67 | 0.37 | 4 | Pericytes |
| GJA4 | 0.59 | 0 | 0 | 0.46 | 0.13 | 4 | Pericytes |
| EPB41L2 | 0.59 | 0 | 0 | 0.54 | 0.24 | 4 | Pericytes |
| ADCY3 | 0.58 | 0 | 0 | 0.33 | 0.06 | 4 | Pericytes |
| RNF152 | 0.58 | 0 | 0 | 0.27 | 0.07 | 4 | Pericytes |
| ZEB2 | 0.58 | 0 | 0 | 0.74 | 0.40 | 4 | Pericytes |
| OAZ2 | 0.58 | 0 | 0 | 0.81 | 0.52 | 4 | Pericytes |
| KCNJ8 | 0.58 | 0 | 0 | 0.42 | 0.13 | 4 | Pericytes |
| DPYSL2 | 0.58 | 0 | 0 | 0.57 | 0.27 | 4 | Pericytes |
| MLLT3 | 0.58 | 0 | 0 | 0.35 | 0.08 | 4 | Pericytes |
| U3.24 | 0.58 | 0 | 0 | 0.28 | 0.03 | 4 | Pericytes |
| EDNRA | 0.58 | 0 | 0 | 0.53 | 0.21 | 4 | Pericytes |
| EZR | 0.58 | 0 | 0 | 0.51 | 0.23 | 4 | Pericytes |
| ADGRA2 | 0.57 | 0 | 0 | 0.39 | 0.11 | 4 | Pericytes |
| ITGB1 | 0.57 | 0 | 0 | 0.86 | 0.61 | 4 | Pericytes |
| CDC42EP2 | 0.57 | 0 | 0 | 0.36 | 0.09 | 4 | Pericytes |
| PTK2 | 0.57 | 0 | 0 | 0.62 | 0.27 | 4 | Pericytes |
| MYO10 | 0.57 | 0 | 0 | 0.38 | 0.10 | 4 | Pericytes |
| F10 | 0.57 | 0 | 0 | 0.38 | 0.10 | 4 | Pericytes |
| TNFRSF21 | 0.57 | 0 | 0 | 0.31 | 0.06 | 4 | Pericytes |
| PRSS23 | 0.57 | 0 | 0 | 0.50 | 0.22 | 4 | Pericytes |
| PLXDC1 | 0.56 | 0 | 0 | 0.27 | 0.02 | 4 | Pericytes |
| TESC | 0.56 | 0 | 0 | 0.29 | 0.04 | 4 | Pericytes |
| COL18A1 | 0.54 | 0 | 0 | 0.42 | 0.16 | 4 | Pericytes |
| CARMN | 0.54 | 0 | 0 | 0.67 | 0.32 | 4 | Pericytes |
| FZD1 | 0.54 | 0 | 0 | 0.34 | 0.10 | 4 | Pericytes |
| CTD-3193K9.4 | 0.54 | 0 | 0 | 0.30 | 0.06 | 4 | Pericytes |
| TFPI | 0.53 | 0 | 0 | 0.74 | 0.41 | 4 | Pericytes |
| WFDC1 | 0.53 | 0 | 0 | 0.50 | 0.22 | 4 | Pericytes |
| FABP5 | 0.53 | 0 | 0 | 0.63 | 0.24 | 4 | Pericytes |
| ARHGAP42 | 0.53 | 0 | 0 | 0.30 | 0.06 | 4 | Pericytes |
| TMEM173 | 0.53 | 0 | 0 | 0.49 | 0.21 | 4 | Pericytes |
| EDIL3 | 0.53 | 0 | 0 | 0.36 | 0.11 | 4 | Pericytes |
| AXL | 0.52 | 0 | 0 | 0.46 | 0.21 | 4 | Pericytes |
| HES4 | 0.52 | 0 | 0 | 0.54 | 0.24 | 4 | Pericytes |
| NR2F2-AS1 | 0.52 | 0 | 0 | 0.36 | 0.10 | 4 | Pericytes |
| TMEM59 | 0.52 | 0 | 0 | 0.87 | 0.67 | 4 | Pericytes |
| DOK6 | 0.51 | 0 | 0 | 0.26 | 0.03 | 4 | Pericytes |
| HOXB-AS1 | 0.51 | 0 | 0 | 0.30 | 0.07 | 4 | Pericytes |
| TNS3 | 0.51 | 0 | 0 | 0.34 | 0.09 | 4 | Pericytes |
| TNS1 | 0.51 | 0 | 0 | 0.46 | 0.22 | 4 | Pericytes |
| EPS8 | 0.51 | 0 | 0 | 0.32 | 0.09 | 4 | Pericytes |
| PON2 | 0.51 | 0 | 0 | 0.50 | 0.22 | 4 | Pericytes |
| PLAC9 | 0.5 | 0 | 0 | 0.72 | 0.38 | 4 | Pericytes |
| AFAP1L2 | 0.5 | 0 | 0 | 0.28 | 0.04 | 4 | Pericytes |
| HIBADH | 0.5 | 0 | 0 | 0.44 | 0.19 | 4 | Pericytes |
| DENND2A | 0.5 | 0 | 0 | 0.31 | 0.08 | 4 | Pericytes |
| LGALS3BP | 0.49 | 0 | 0 | 0.43 | 0.18 | 4 | Pericytes |
| TNS2 | 0.49 | 0 | 0 | 0.38 | 0.15 | 4 | Pericytes |
| LAMA2 | 0.49 | 0 | 0 | 0.27 | 0.04 | 4 | Pericytes |
| GPR183 | 0.49 | 0 | 0 | 0.28 | 0.05 | 4 | Pericytes |
| CLIP1 | 0.48 | 0 | 0 | 0.36 | 0.13 | 4 | Pericytes |
| PPP1R12A | 0.48 | 0 | 0 | 0.71 | 0.45 | 4 | Pericytes |
| MCAM | 0.48 | 0 | 0 | 0.48 | 0.21 | 4 | Pericytes |
| VASN | 0.47 | 0 | 0 | 0.37 | 0.15 | 4 | Pericytes |
| TGFBI | 0.47 | 0 | 0 | 0.75 | 0.32 | 4 | Pericytes |
| GOLIM4 | 0.47 | 0 | 0 | 0.47 | 0.26 | 4 | Pericytes |
| AP2M1 | 0.47 | 0 | 0 | 0.80 | 0.59 | 4 | Pericytes |
| DLC1 | 0.46 | 0 | 0 | 0.46 | 0.21 | 4 | Pericytes |
| ZBTB38 | 0.46 | 0 | 0 | 0.43 | 0.22 | 4 | Pericytes |
| SEMA5A | 0.46 | 0 | 0 | 0.34 | 0.11 | 4 | Pericytes |
| NDN | 0.46 | 0 | 0 | 0.58 | 0.32 | 4 | Pericytes |
| PACSIN3 | 0.45 | 0 | 0 | 0.28 | 0.08 | 4 | Pericytes |
| TMEM204 | 0.45 | 0 | 0 | 0.45 | 0.16 | 4 | Pericytes |
| HOXB5 | 0.45 | 0 | 0 | 0.38 | 0.16 | 4 | Pericytes |
| ARHGAP15 | 0.45 | 0 | 0 | 0.39 | 0.15 | 4 | Pericytes |
| MXRA8 | 0.45 | 0 | 0 | 0.53 | 0.29 | 4 | Pericytes |
| BBX | 0.44 | 0 | 0 | 0.51 | 0.29 | 4 | Pericytes |
| COX7A1 | 0.44 | 0 | 0 | 0.65 | 0.43 | 4 | Pericytes |
| PIK3R1 | 0.44 | 0 | 0 | 0.37 | 0.14 | 4 | Pericytes |
| ARHGEF12 | 0.44 | 0 | 0 | 0.45 | 0.23 | 4 | Pericytes |
| ABTB1 | 0.44 | 0 | 0 | 0.32 | 0.12 | 4 | Pericytes |
| CSPG4 | 0.44 | 0 | 0 | 0.26 | 0.06 | 4 | Pericytes |
| NES | 0.44 | 0 | 0 | 0.32 | 0.11 | 4 | Pericytes |
| EFEMP2 | 0.43 | 0 | 0 | 0.57 | 0.35 | 4 | Pericytes |
| RRAD | 0.43 | 0 | 0 | 0.39 | 0.17 | 4 | Pericytes |
| CD4 | 0.43 | 0 | 0 | 0.30 | 0.09 | 4 | Pericytes |
| FOXF2 | 0.43 | 0 | 0 | 0.27 | 0.07 | 4 | Pericytes |
| FRMD4B | 0.42 | 0 | 0 | 0.35 | 0.14 | 4 | Pericytes |
| AC013461.1 | 0.42 | 0 | 0 | 0.60 | 0.35 | 4 | Pericytes |
| BCAR1 | 0.42 | 0 | 0 | 0.36 | 0.17 | 4 | Pericytes |
| ADAMTS9 | 0.42 | 0 | 0 | 0.26 | 0.13 | 4 | Pericytes |
| COL1A2 | 0.42 | 0 | 0 | 0.98 | 0.62 | 4 | Pericytes |
| COL6A2 | 0.41 | 0 | 0 | 0.87 | 0.60 | 4 | Pericytes |
| TRPV2 | 0.41 | 0 | 0 | 0.25 | 0.07 | 4 | Pericytes |
| CSNK1E | 0.41 | 0 | 0 | 0.39 | 0.22 | 4 | Pericytes |
| TGFB1I1 | 0.4 | 0 | 0 | 0.66 | 0.40 | 4 | Pericytes |
| NDUFAF4 | 0.4 | 0 | 0 | 0.33 | 0.14 | 4 | Pericytes |
| RRAS2 | 0.4 | 0 | 0 | 0.28 | 0.10 | 4 | Pericytes |
| ISYNA1 | 0.4 | 0 | 0 | 0.43 | 0.19 | 4 | Pericytes |
| VAMP5 | 0.4 | 0 | 0 | 0.77 | 0.49 | 4 | Pericytes |
| SMTN | 0.39 | 0 | 0 | 0.48 | 0.27 | 4 | Pericytes |
| TBX2-AS1 | 0.39 | 0 | 0 | 0.40 | 0.19 | 4 | Pericytes |
| ADH5 | 0.39 | 0 | 0 | 0.54 | 0.32 | 4 | Pericytes |
| NR2F1-AS1 | 0.39 | 0 | 0 | 0.28 | 0.10 | 4 | Pericytes |
| MAP1LC3A | 0.39 | 0 | 0 | 0.50 | 0.29 | 4 | Pericytes |
| BNIP3L | 0.39 | 0 | 0 | 0.60 | 0.38 | 4 | Pericytes |
| ITPRIPL2 | 0.39 | 0 | 0 | 0.26 | 0.09 | 4 | Pericytes |
| KLHL42 | 0.39 | 0 | 0 | 0.28 | 0.11 | 4 | Pericytes |
| CTBP2 | 0.39 | 0 | 0 | 0.40 | 0.21 | 4 | Pericytes |
| REM1 | 0.39 | 0 | 0 | 0.26 | 0.09 | 4 | Pericytes |
| PLEKHA5 | 0.39 | 0 | 0 | 0.29 | 0.10 | 4 | Pericytes |
| UBA2 | 0.39 | 0 | 0 | 0.44 | 0.26 | 4 | Pericytes |
| SYDE1 | 0.38 | 0 | 0 | 0.25 | 0.08 | 4 | Pericytes |
| SYPL1 | 0.38 | 0 | 0 | 0.54 | 0.34 | 4 | Pericytes |
| WLS | 0.38 | 0 | 0 | 0.34 | 0.15 | 4 | Pericytes |
| LMCD1 | 0.38 | 0 | 0 | 0.64 | 0.36 | 4 | Pericytes |
| MT1E | 0.38 | 0 | 0 | 0.44 | 0.22 | 4 | Pericytes |
| RPS6KA2 | 0.38 | 0 | 0 | 0.29 | 0.09 | 4 | Pericytes |
| RGS5 | 0.38 | 0 | 0 | 0.40 | 0.23 | 4 | Pericytes |
| EMC2 | 0.38 | 0 | 0 | 0.36 | 0.18 | 4 | Pericytes |
| TAOK3 | 0.38 | 0 | 0 | 0.34 | 0.15 | 4 | Pericytes |
| F2R | 0.37 | 0 | 0 | 0.26 | 0.09 | 4 | Pericytes |
| PHLDA1 | 0.37 | 0 | 0 | 0.41 | 0.23 | 4 | Pericytes |
| HOXB2 | 0.37 | 0 | 0 | 0.34 | 0.16 | 4 | Pericytes |
| NOTCH2 | 0.37 | 0 | 0 | 0.26 | 0.11 | 4 | Pericytes |
| PLEKHA2 | 0.37 | 0 | 0 | 0.26 | 0.09 | 4 | Pericytes |
| LEPROT | 0.37 | 0 | 0 | 0.61 | 0.41 | 4 | Pericytes |
| TXNL1 | 0.36 | 0 | 0 | 0.52 | 0.33 | 4 | Pericytes |
| PSIP1 | 0.36 | 0 | 0 | 0.57 | 0.37 | 4 | Pericytes |
| LIMS1 | 0.36 | 0 | 0 | 0.54 | 0.32 | 4 | Pericytes |
| UBE2E2 | 0.36 | 0 | 0 | 0.34 | 0.16 | 4 | Pericytes |
| MYH9 | 0.36 | 0 | 0 | 0.63 | 0.43 | 4 | Pericytes |
| ASAH1 | 0.36 | 0 | 0 | 0.48 | 0.28 | 4 | Pericytes |
| TMEM47 | 0.36 | 0 | 0 | 0.32 | 0.16 | 4 | Pericytes |
| PRRT2 | 0.35 | 0 | 0 | 0.26 | 0.09 | 4 | Pericytes |
| DOCK6 | 0.35 | 0 | 0 | 0.34 | 0.13 | 4 | Pericytes |
| 2-Sep | 0.35 | 0 | 0 | 0.71 | 0.53 | 4 | Pericytes |
| ITGA8 | 0.35 | 0 | 0 | 0.29 | 0.11 | 4 | Pericytes |
| TCEAL7 | 0.35 | 0 | 0 | 0.30 | 0.11 | 4 | Pericytes |
| PAWR | 0.35 | 0 | 0 | 0.31 | 0.16 | 4 | Pericytes |
| SNAI2 | 0.35 | 0 | 0 | 0.27 | 0.12 | 4 | Pericytes |
| BCHE | 0.35 | 0 | 0 | 0.35 | 0.15 | 4 | Pericytes |
| RPLP0P2 | 0.34 | 0 | 0 | 0.39 | 0.25 | 4 | Pericytes |
| IFI27L2 | 0.34 | 0 | 0 | 0.53 | 0.32 | 4 | Pericytes |
| PKIG | 0.34 | 0 | 0 | 0.57 | 0.37 | 4 | Pericytes |
| SERPINI1 | 0.34 | 0 | 0 | 0.28 | 0.12 | 4 | Pericytes |
| CYB5B | 0.34 | 0 | 0 | 0.39 | 0.24 | 4 | Pericytes |
| ARHGAP24 | 0.33 | 0 | 0 | 0.25 | 0.09 | 4 | Pericytes |
| HOTAIRM1 | 0.33 | 0 | 0 | 0.32 | 0.16 | 4 | Pericytes |
| TCF4 | 0.33 | 0 | 0 | 0.74 | 0.46 | 4 | Pericytes |
| CTDSPL | 0.33 | 0 | 0 | 0.27 | 0.10 | 4 | Pericytes |
| ISCA1 | 0.33 | 0 | 0 | 0.36 | 0.20 | 4 | Pericytes |
| BEX3 | 0.33 | 0 | 0 | 0.87 | 0.66 | 4 | Pericytes |
| FOXF1 | 0.33 | 0 | 0 | 0.48 | 0.24 | 4 | Pericytes |
| PTP4A3 | 0.33 | 0 | 0 | 0.44 | 0.23 | 4 | Pericytes |
| ZFHX3 | 0.33 | 0 | 0 | 0.43 | 0.26 | 4 | Pericytes |
| ECE1 | 0.32 | 0 | 0 | 0.29 | 0.14 | 4 | Pericytes |
| COL6A1 | 0.32 | 0 | 0 | 0.78 | 0.51 | 4 | Pericytes |
| TCEAL8 | 0.32 | 0 | 0 | 0.54 | 0.37 | 4 | Pericytes |
| RHOC | 0.32 | 0 | 0 | 0.80 | 0.59 | 4 | Pericytes |
| PMP22 | 0.32 | 0 | 0 | 0.70 | 0.46 | 4 | Pericytes |
| LAPTM4A | 0.32 | 0 | 0 | 0.96 | 0.84 | 4 | Pericytes |
| LGALS1 | 0.31 | 0 | 0 | 0.98 | 0.89 | 4 | Pericytes |
| HOXB4 | 0.31 | 0 | 0 | 0.31 | 0.15 | 4 | Pericytes |
| DDR2 | 0.31 | 0 | 0 | 0.25 | 0.11 | 4 | Pericytes |
| KLHDC8B | 0.31 | 0 | 0 | 0.26 | 0.13 | 4 | Pericytes |
| ATAD1 | 0.31 | 0 | 0 | 0.28 | 0.14 | 4 | Pericytes |
| ARHGAP29 | 0.31 | 0 | 0 | 0.62 | 0.30 | 4 | Pericytes |
| KLHDC3 | 0.31 | 0 | 0 | 0.43 | 0.26 | 4 | Pericytes |
| DARS | 0.31 | 0 | 0 | 0.41 | 0.27 | 4 | Pericytes |
| HOXB6 | 0.31 | 0 | 0 | 0.27 | 0.11 | 4 | Pericytes |
| AHNAK | 0.3 | 0 | 0 | 0.54 | 0.38 | 4 | Pericytes |
| RPS27L | 0.3 | 0 | 0 | 0.72 | 0.56 | 4 | Pericytes |
| HSD17B12 | 0.3 | 0 | 0 | 0.45 | 0.29 | 4 | Pericytes |
| EIF4A2 | 0.3 | 0 | 0 | 0.90 | 0.74 | 4 | Pericytes |
| CD47 | 0.3 | 0 | 0 | 0.35 | 0.20 | 4 | Pericytes |
| TCAF1 | 0.3 | 0 | 0 | 0.32 | 0.19 | 4 | Pericytes |
| C12orf57 | 0.3 | 0 | 0 | 0.79 | 0.61 | 4 | Pericytes |
| SNW1 | 0.29 | 0 | 0 | 0.42 | 0.27 | 4 | Pericytes |
| EVA1B | 0.29 | 0 | 0 | 0.29 | 0.14 | 4 | Pericytes |
| RPS7 | 0.29 | 0 | 0 | 1.00 | 0.97 | 4 | Pericytes |
| SKP1 | 0.29 | 0 | 0 | 0.93 | 0.83 | 4 | Pericytes |
| ARMCX1 | 0.29 | 0 | 0 | 0.32 | 0.18 | 4 | Pericytes |
| CRTC3 | 0.29 | 0 | 0 | 0.27 | 0.15 | 4 | Pericytes |
| MID1IP1 | 0.28 | 0 | 0 | 0.27 | 0.15 | 4 | Pericytes |
| FAM204A | 0.28 | 0 | 0 | 0.38 | 0.25 | 4 | Pericytes |
| MYLIP | 0.28 | 0 | 0 | 0.46 | 0.29 | 4 | Pericytes |
| VAMP2 | 0.28 | 0 | 0 | 0.67 | 0.52 | 4 | Pericytes |
| WASF2 | 0.28 | 0 | 0 | 0.58 | 0.42 | 4 | Pericytes |
| COMMD7 | 0.28 | 0 | 0 | 0.37 | 0.23 | 4 | Pericytes |
| GYPC | 0.28 | 0 | 0 | 0.48 | 0.31 | 4 | Pericytes |
| RBBP7 | 0.27 | 0 | 0 | 0.44 | 0.29 | 4 | Pericytes |
| MORF4L1 | 0.27 | 0 | 0 | 0.90 | 0.78 | 4 | Pericytes |
| TWSG1 | 0.27 | 0 | 0 | 0.30 | 0.17 | 4 | Pericytes |
| HES1 | 0.27 | 0 | 0 | 0.65 | 0.36 | 4 | Pericytes |
| ARL5A | 0.27 | 0 | 0 | 0.32 | 0.20 | 4 | Pericytes |
| FKBP8 | 0.27 | 0 | 0 | 0.62 | 0.47 | 4 | Pericytes |
| CBR1 | 0.27 | 0 | 0 | 0.28 | 0.16 | 4 | Pericytes |
| ADGRF5 | 0.27 | 0 | 0 | 0.33 | 0.16 | 4 | Pericytes |
| 9-Sep | 0.27 | 0 | 0 | 0.38 | 0.23 | 4 | Pericytes |
| SNRK | 0.26 | 0 | 0 | 0.27 | 0.12 | 4 | Pericytes |
| EID1 | 0.26 | 0 | 0 | 0.90 | 0.78 | 4 | Pericytes |
| ID3 | 0.26 | 0 | 0 | 0.75 | 0.52 | 4 | Pericytes |
| LAMP1 | 0.26 | 0 | 0 | 0.58 | 0.43 | 4 | Pericytes |
| MT-RNR1 | 0.26 | 0 | 0 | 0.97 | 0.95 | 4 | Pericytes |
| NDUFS2 | 0.26 | 0 | 0 | 0.39 | 0.28 | 4 | Pericytes |
| RCN2 | 0.26 | 0 | 0 | 0.57 | 0.40 | 4 | Pericytes |
| MDH1 | 0.26 | 0 | 0 | 0.40 | 0.28 | 4 | Pericytes |
| ADI1 | 0.26 | 0 | 0 | 0.40 | 0.26 | 4 | Pericytes |
| SLC25A37 | 0.26 | 0 | 0 | 0.31 | 0.18 | 4 | Pericytes |
| EMP1 | 0.26 | 0 | 0 | 0.40 | 0.22 | 4 | Pericytes |
| GCC2 | 0.26 | 0 | 0 | 0.33 | 0.20 | 4 | Pericytes |
| 15-Sep | 0.25 | 0 | 0 | 0.66 | 0.51 | 4 | Pericytes |
| TERF2IP | 0.25 | 0 | 0 | 0.50 | 0.38 | 4 | Pericytes |
| CD59 | 0.25 | 0 | 0 | 0.85 | 0.58 | 4 | Pericytes |
| CUTA | 0.25 | 0 | 0 | 0.68 | 0.54 | 4 | Pericytes |
| ATP2B1 | 0.25 | 0 | 0 | 0.37 | 0.25 | 4 | Pericytes |
| FBLIM1 | 0.25 | 0 | 0 | 0.27 | 0.18 | 4 | Pericytes |
| FKBP10 | 0.25 | 0 | 0 | 0.34 | 0.20 | 4 | Pericytes |
| CXCL14 | 2.1 | 0 | 0 | 0.63 | 0.04 | 5 | Stromal Cells |
| FBLN1 | 1.73 | 0 | 0 | 0.89 | 0.26 | 5 | Stromal Cells |
| COL3A1 | 1.73 | 0 | 0 | 0.97 | 0.56 | 5 | Stromal Cells |
| SERPINF1 | 1.52 | 0 | 0 | 0.53 | 0.04 | 5 | Stromal Cells |
| COL1A1 | 1.41 | 0 | 0 | 0.95 | 0.60 | 5 | Stromal Cells |
| CCDC80 | 1.38 | 0 | 0 | 0.69 | 0.20 | 5 | Stromal Cells |
| COL1A2 | 1.21 | 0 | 0 | 0.97 | 0.62 | 5 | Stromal Cells |
| DCN | 1.18 | 0 | 0 | 0.95 | 0.51 | 5 | Stromal Cells |
| NBL1 | 1.15 | 0 | 0 | 0.86 | 0.34 | 5 | Stromal Cells |
| MMP2 | 1.12 | 0 | 0 | 0.85 | 0.31 | 5 | Stromal Cells |
| IGF1 | 1.1 | 0 | 0 | 0.35 | 0.07 | 5 | Stromal Cells |
| OGN | 0.97 | 0 | 0 | 0.53 | 0.12 | 5 | Stromal Cells |
| CFH | 0.94 | 0 | 0 | 0.41 | 0.10 | 5 | Stromal Cells |
| PDGFRA | 0.93 | 0 | 0 | 0.38 | 0.04 | 5 | Stromal Cells |
| MGP | 0.91 | 0 | 0 | 0.79 | 0.65 | 5 | Stromal Cells |
| CTGF | 0.85 | 0 | 0 | 0.39 | 0.21 | 5 | Stromal Cells |
| PMP22 | 0.84 | 0 | 0 | 0.85 | 0.46 | 5 | Stromal Cells |
| A2M | 0.84 | 0 | 0 | 0.89 | 0.60 | 5 | Stromal Cells |
| PRSS23 | 0.82 | 0 | 0 | 0.56 | 0.22 | 5 | Stromal Cells |
| RND3 | 0.82 | 0 | 0 | 0.53 | 0.17 | 5 | Stromal Cells |
| FGF7 | 0.81 | 0 | 0 | 0.40 | 0.08 | 5 | Stromal Cells |
| CLU | 0.81 | 0 | 0 | 0.53 | 0.31 | 5 | Stromal Cells |
| FHL1 | 0.8 | 0 | 0 | 0.87 | 0.55 | 5 | Stromal Cells |
| C1S | 0.8 | 0 | 0 | 0.59 | 0.19 | 5 | Stromal Cells |
| ALDH1A1 | 0.79 | 0 | 0 | 0.35 | 0.11 | 5 | Stromal Cells |
| C2orf40 | 0.77 | 0 | 0 | 0.31 | 0.15 | 5 | Stromal Cells |
| FBN1 | 0.76 | 0 | 0 | 0.50 | 0.13 | 5 | Stromal Cells |
| ADH1B | 0.75 | 0 | 0 | 0.56 | 0.24 | 5 | Stromal Cells |
| C7 | 0.74 | 0 | 0 | 0.38 | 0.16 | 5 | Stromal Cells |
| ASPN | 0.74 | 0 | 0 | 0.32 | 0.07 | 5 | Stromal Cells |
| BMP4 | 0.74 | 0 | 0 | 0.34 | 0.08 | 5 | Stromal Cells |
| PCOLCE | 0.73 | 0 | 0 | 0.60 | 0.29 | 5 | Stromal Cells |
| LUM | 0.73 | 0 | 0 | 0.67 | 0.36 | 5 | Stromal Cells |
| CDH11 | 0.72 | 0 | 0 | 0.57 | 0.18 | 5 | Stromal Cells |
| FSTL1 | 0.71 | 0 | 0 | 0.80 | 0.41 | 5 | Stromal Cells |
| C1R | 0.71 | 0 | 0 | 0.54 | 0.23 | 5 | Stromal Cells |
| HTRA3 | 0.71 | 0 | 0 | 0.30 | 0.04 | 5 | Stromal Cells |
| DPT | 0.7 | 0 | 0 | 0.26 | 0.04 | 5 | Stromal Cells |
| CPE | 0.7 | 0 | 0 | 0.54 | 0.24 | 5 | Stromal Cells |
| LTBP4 | 0.69 | 0 | 0 | 0.66 | 0.27 | 5 | Stromal Cells |
| TNC | 0.69 | 0 | 0 | 0.44 | 0.15 | 5 | Stromal Cells |
| ELN | 0.67 | 0 | 0 | 0.86 | 0.52 | 5 | Stromal Cells |
| MFAP4 | 0.66 | 0 | 0 | 0.89 | 0.47 | 5 | Stromal Cells |
| TSPAN8 | 0.64 | 0 | 0 | 0.27 | 0.02 | 5 | Stromal Cells |
| LMO4 | 0.64 | 0 | 0 | 0.66 | 0.36 | 5 | Stromal Cells |
| COL6A1 | 0.62 | 0 | 0 | 0.88 | 0.51 | 5 | Stromal Cells |
| PTGDS | 0.62 | 0 | 0 | 0.31 | 0.16 | 5 | Stromal Cells |
| ROBO2 | 0.62 | 0 | 0 | 0.34 | 0.05 | 5 | Stromal Cells |
| COL6A2 | 0.61 | 0 | 0 | 0.89 | 0.61 | 5 | Stromal Cells |
| CYR61 | 0.61 | 0 | 0 | 0.50 | 0.21 | 5 | Stromal Cells |
| CCL2 | 0.61 | 0 | 0 | 0.32 | 0.16 | 5 | Stromal Cells |
| TMEM98 | 0.61 | 0 | 0 | 0.66 | 0.30 | 5 | Stromal Cells |
| MMP23B | 0.6 | 0 | 0 | 0.46 | 0.12 | 5 | Stromal Cells |
| DKK3 | 0.6 | 0 | 0 | 0.62 | 0.26 | 5 | Stromal Cells |
| SCN7A | 0.6 | 0 | 0 | 0.38 | 0.09 | 5 | Stromal Cells |
| POSTN | 0.59 | 0 | 0 | 0.44 | 0.20 | 5 | Stromal Cells |
| RARRES2 | 0.59 | 0 | 0 | 0.89 | 0.49 | 5 | Stromal Cells |
| CTSK | 0.59 | 0 | 0 | 0.31 | 0.04 | 5 | Stromal Cells |
| RCN3 | 0.57 | 0 | 0 | 0.32 | 0.06 | 5 | Stromal Cells |
| CFD | 0.57 | 0 | 0 | 0.48 | 0.27 | 5 | Stromal Cells |
| SPON1 | 0.56 | 0 | 0 | 0.45 | 0.15 | 5 | Stromal Cells |
| GSTM5 | 0.55 | 0 | 0 | 0.37 | 0.12 | 5 | Stromal Cells |
| MEG3 | 0.55 | 0 | 0 | 0.72 | 0.42 | 5 | Stromal Cells |
| GFRA2 | 0.54 | 0 | 0 | 0.26 | 0.04 | 5 | Stromal Cells |
| NR2F1 | 0.53 | 0 | 0 | 0.60 | 0.25 | 5 | Stromal Cells |
| RARRES1 | 0.51 | 0 | 0 | 0.42 | 0.16 | 5 | Stromal Cells |
| C1QTNF7 | 0.51 | 0 | 0 | 0.27 | 0.04 | 5 | Stromal Cells |
| MATN2 | 0.5 | 0 | 0 | 0.28 | 0.06 | 5 | Stromal Cells |
| TIMP3 | 0.5 | 0 | 0 | 0.70 | 0.54 | 5 | Stromal Cells |
| CD82 | 0.5 | 0 | 0 | 0.50 | 0.25 | 5 | Stromal Cells |
| LTBP2 | 0.5 | 0 | 0 | 0.38 | 0.12 | 5 | Stromal Cells |
| PLPP3 | 0.49 | 0 | 0 | 0.54 | 0.28 | 5 | Stromal Cells |
| OLFML3 | 0.49 | 0 | 0 | 0.46 | 0.19 | 5 | Stromal Cells |
| LMCD1 | 0.49 | 0 | 0 | 0.62 | 0.36 | 5 | Stromal Cells |
| FBLN5 | 0.48 | 0 | 0 | 0.38 | 0.16 | 5 | Stromal Cells |
| IGFBP6 | 0.48 | 0 | 0 | 0.25 | 0.05 | 5 | Stromal Cells |
| CYBRD1 | 0.48 | 0 | 0 | 0.39 | 0.14 | 5 | Stromal Cells |
| COL5A1 | 0.47 | 0 | 0 | 0.51 | 0.21 | 5 | Stromal Cells |
| ANXA5 | 0.47 | 0 | 0 | 0.78 | 0.67 | 5 | Stromal Cells |
| MIR100HG | 0.47 | 0 | 0 | 0.26 | 0.05 | 5 | Stromal Cells |
| CXCL12 | 0.46 | 0 | 0 | 0.48 | 0.22 | 5 | Stromal Cells |
| GLT8D2 | 0.46 | 0 | 0 | 0.29 | 0.06 | 5 | Stromal Cells |
| MYADM | 0.46 | 0 | 0 | 0.51 | 0.37 | 5 | Stromal Cells |
| TGFBI | 0.46 | 0 | 0 | 0.51 | 0.35 | 5 | Stromal Cells |
| PPIC | 0.46 | 0 | 0 | 0.61 | 0.35 | 5 | Stromal Cells |
| CAMK2N1 | 0.46 | 0 | 0 | 0.48 | 0.20 | 5 | Stromal Cells |
| LOXL1 | 0.46 | 0 | 0 | 0.35 | 0.11 | 5 | Stromal Cells |
| CD302 | 0.45 | 0 | 0 | 0.39 | 0.16 | 5 | Stromal Cells |
| RPL10AP6 | 0.45 | 0 | 0 | 0.91 | 0.76 | 5 | Stromal Cells |
| GPX3 | 0.45 | 0 | 0 | 0.67 | 0.43 | 5 | Stromal Cells |
| LGALS3 | 0.45 | 0 | 0 | 0.61 | 0.35 | 5 | Stromal Cells |
| KDELR2 | 0.44 | 0 | 0 | 0.64 | 0.44 | 5 | Stromal Cells |
| RCN1 | 0.44 | 0 | 0 | 0.60 | 0.37 | 5 | Stromal Cells |
| MYH10 | 0.44 | 0 | 0 | 0.55 | 0.25 | 5 | Stromal Cells |
| NFIB | 0.44 | 0 | 0 | 0.77 | 0.48 | 5 | Stromal Cells |
| ABCA8 | 0.44 | 0 | 0 | 0.33 | 0.11 | 5 | Stromal Cells |
| CCDC68 | 0.43 | 0 | 0 | 0.27 | 0.11 | 5 | Stromal Cells |
| COLEC12 | 0.43 | 0 | 0 | 0.38 | 0.12 | 5 | Stromal Cells |
| ZFP36L2 | 0.43 | 0 | 0 | 0.71 | 0.48 | 5 | Stromal Cells |
| LRRC75A-AS1 | 0.43 | 0 | 0 | 0.97 | 0.86 | 5 | Stromal Cells |
| TSHZ2 | 0.43 | 0 | 0 | 0.31 | 0.11 | 5 | Stromal Cells |
| TSPAN13 | 0.43 | 0 | 0 | 0.33 | 0.12 | 5 | Stromal Cells |
| RBMS3 | 0.43 | 0 | 0 | 0.41 | 0.18 | 5 | Stromal Cells |
| LBH | 0.42 | 0 | 0 | 0.59 | 0.39 | 5 | Stromal Cells |
| ST3GAL4 | 0.42 | 0 | 0 | 0.33 | 0.12 | 5 | Stromal Cells |
| COL13A1 | 0.41 | 0 | 0 | 0.28 | 0.08 | 5 | Stromal Cells |
| NOVA1 | 0.41 | 0 | 0 | 0.35 | 0.10 | 5 | Stromal Cells |
| CST3 | 0.41 | 0 | 0 | 0.84 | 0.57 | 5 | Stromal Cells |
| RPL10A | 0.41 | 0 | 0 | 1.00 | 0.96 | 5 | Stromal Cells |
| KCNQ1OT1 | 0.41 | 0 | 0 | 0.55 | 0.29 | 5 | Stromal Cells |
| TMED3 | 0.4 | 0 | 0 | 0.35 | 0.13 | 5 | Stromal Cells |
| EMP1 | 0.4 | 0 | 0 | 0.33 | 0.23 | 5 | Stromal Cells |
| FKBP10 | 0.4 | 0 | 0 | 0.44 | 0.19 | 5 | Stromal Cells |
| RP11-864N7.2 | 0.4 | 0 | 0 | 0.98 | 0.89 | 5 | Stromal Cells |
| SRPX | 0.4 | 0 | 0 | 0.25 | 0.15 | 5 | Stromal Cells |
| COL21A1 | 0.4 | 0 | 0 | 0.32 | 0.13 | 5 | Stromal Cells |
| NENF | 0.39 | 0 | 0 | 0.74 | 0.46 | 5 | Stromal Cells |
| MDFI | 0.39 | 0 | 0 | 0.43 | 0.17 | 5 | Stromal Cells |
| IL11RA | 0.39 | 0 | 0 | 0.26 | 0.10 | 5 | Stromal Cells |
| NPM1 | 0.39 | 0 | 0 | 0.97 | 0.87 | 5 | Stromal Cells |
| RAB34 | 0.38 | 0 | 0 | 0.65 | 0.37 | 5 | Stromal Cells |
| TTC3 | 0.38 | 0 | 0 | 0.61 | 0.37 | 5 | Stromal Cells |
| RPS12 | 0.38 | 0 | 0 | 1.00 | 0.99 | 5 | Stromal Cells |
| RPLP0 | 0.38 | 0 | 0 | 1.00 | 0.92 | 5 | Stromal Cells |
| RHOBTB3 | 0.38 | 0 | 0 | 0.36 | 0.17 | 5 | Stromal Cells |
| FGFR1 | 0.37 | 0 | 0 | 0.46 | 0.22 | 5 | Stromal Cells |
| LRRC17 | 0.37 | 0 | 0 | 0.28 | 0.12 | 5 | Stromal Cells |
| PRDX4 | 0.37 | 0 | 0 | 0.49 | 0.27 | 5 | Stromal Cells |
| SNHG8 | 0.37 | 0 | 0 | 0.81 | 0.55 | 5 | Stromal Cells |
| PALLD | 0.37 | 0 | 0 | 0.42 | 0.23 | 5 | Stromal Cells |
| GSTM3 | 0.37 | 0 | 0 | 0.40 | 0.19 | 5 | Stromal Cells |
| RPL7P9 | 0.37 | 0 | 0 | 0.91 | 0.74 | 5 | Stromal Cells |
| RRBP1 | 0.36 | 0 | 0 | 0.52 | 0.31 | 5 | Stromal Cells |
| CCBE1 | 0.36 | 0 | 0 | 0.31 | 0.11 | 5 | Stromal Cells |
| SLC40A1 | 0.36 | 0 | 0 | 0.45 | 0.25 | 5 | Stromal Cells |
| EEF1A1 | 0.36 | 0 | 0 | 1.00 | 1.00 | 5 | Stromal Cells |
| CENPV | 0.36 | 0 | 0 | 0.37 | 0.16 | 5 | Stromal Cells |
| EEF2 | 0.36 | 0 | 0 | 0.97 | 0.88 | 5 | Stromal Cells |
| SEPP1 | 0.36 | 0 | 0 | 0.79 | 0.55 | 5 | Stromal Cells |
| LTBP1 | 0.36 | 0 | 0 | 0.40 | 0.22 | 5 | Stromal Cells |
| EFEMP2 | 0.36 | 0 | 0 | 0.62 | 0.35 | 5 | Stromal Cells |
| CNN2 | 0.36 | 0 | 0 | 0.43 | 0.22 | 5 | Stromal Cells |
| HNRNPA1 | 0.36 | 0 | 0 | 0.99 | 0.94 | 5 | Stromal Cells |
| PDLIM3 | 0.36 | 0 | 0 | 0.53 | 0.28 | 5 | Stromal Cells |
| COL5A2 | 0.36 | 0 | 0 | 0.45 | 0.21 | 5 | Stromal Cells |
| RPL13AP5 | 0.36 | 0 | 0 | 0.99 | 0.93 | 5 | Stromal Cells |
| TGFB3 | 0.35 | 0 | 0 | 0.25 | 0.08 | 5 | Stromal Cells |
| FAM3C | 0.35 | 0 | 0 | 0.44 | 0.24 | 5 | Stromal Cells |
| LITAF | 0.35 | 0 | 0 | 0.75 | 0.44 | 5 | Stromal Cells |
| SERPING1 | 0.35 | 0 | 0 | 0.61 | 0.38 | 5 | Stromal Cells |
| SELM | 0.35 | 0 | 0 | 0.75 | 0.47 | 5 | Stromal Cells |
| ZNF503 | 0.35 | 0 | 0 | 0.52 | 0.33 | 5 | Stromal Cells |
| TCF21 | 0.34 | 0 | 0 | 0.55 | 0.29 | 5 | Stromal Cells |
| VIM | 0.34 | 0 | 0 | 1.00 | 0.94 | 5 | Stromal Cells |
| FAM46A | 0.34 | 0 | 0 | 0.30 | 0.12 | 5 | Stromal Cells |
| BEX3 | 0.34 | 0 | 0 | 0.89 | 0.66 | 5 | Stromal Cells |
| MDK | 0.34 | 0 | 0 | 0.73 | 0.52 | 5 | Stromal Cells |
| RPL13A | 0.34 | 0 | 0 | 1.00 | 1.00 | 5 | Stromal Cells |
| RPL4 | 0.34 | 0 | 0 | 0.99 | 0.92 | 5 | Stromal Cells |
| ARF4 | 0.34 | 0 | 0 | 0.68 | 0.50 | 5 | Stromal Cells |
| MEOX2 | 0.33 | 0 | 0 | 0.33 | 0.13 | 5 | Stromal Cells |
| CTB-63M22.1 | 0.33 | 0 | 0 | 0.76 | 0.59 | 5 | Stromal Cells |
| ALDH2 | 0.33 | 0 | 0 | 0.53 | 0.33 | 5 | Stromal Cells |
| CTSF | 0.33 | 0 | 0 | 0.34 | 0.15 | 5 | Stromal Cells |
| FXYD1 | 0.33 | 0 | 0 | 0.49 | 0.25 | 5 | Stromal Cells |
| LRP1 | 0.33 | 0 | 0 | 0.37 | 0.17 | 5 | Stromal Cells |
| TNFRSF12A | 0.33 | 0 | 0 | 0.27 | 0.16 | 5 | Stromal Cells |
| NREP | 0.32 | 0 | 0 | 0.54 | 0.36 | 5 | Stromal Cells |
| ANTXR1 | 0.32 | 0 | 0 | 0.44 | 0.22 | 5 | Stromal Cells |
| C6orf48 | 0.32 | 0 | 0 | 0.81 | 0.59 | 5 | Stromal Cells |
| EEF1A1P5 | 0.32 | 0 | 0 | 0.76 | 0.65 | 5 | Stromal Cells |
| GPX8 | 0.32 | 0 | 0 | 0.29 | 0.10 | 5 | Stromal Cells |
| RPL17 | 0.32 | 0 | 0 | 0.94 | 0.87 | 5 | Stromal Cells |
| HOXA3 | 0.32 | 0 | 0 | 0.26 | 0.11 | 5 | Stromal Cells |
| GAS5 | 0.32 | 0 | 0 | 0.88 | 0.70 | 5 | Stromal Cells |
| NR2F1-AS1 | 0.32 | 0 | 0 | 0.29 | 0.10 | 5 | Stromal Cells |
| EID1 | 0.32 | 0 | 0 | 0.92 | 0.78 | 5 | Stromal Cells |
| RPL15 | 0.32 | 0 | 0 | 1.00 | 0.99 | 5 | Stromal Cells |
| GOLIM4 | 0.31 | 0 | 0 | 0.48 | 0.27 | 5 | Stromal Cells |
| MEST | 0.31 | 0 | 0 | 0.75 | 0.36 | 5 | Stromal Cells |
| RPLP0P6 | 0.31 | 0 | 0 | 0.56 | 0.36 | 5 | Stromal Cells |
| MORF4L2 | 0.31 | 0 | 0 | 0.67 | 0.47 | 5 | Stromal Cells |
| PFN2 | 0.31 | 0 | 0 | 0.42 | 0.23 | 5 | Stromal Cells |
| TWSG1 | 0.31 | 0 | 0 | 0.35 | 0.17 | 5 | Stromal Cells |
| RPS8 | 0.3 | 0 | 0 | 1.00 | 0.99 | 5 | Stromal Cells |
| TUBB2A | 0.3 | 0 | 0 | 0.27 | 0.16 | 5 | Stromal Cells |
| RP11-36C20.1 | 0.3 | 0 | 0 | 0.63 | 0.45 | 5 | Stromal Cells |
| RCN2 | 0.3 | 0 | 0 | 0.58 | 0.40 | 5 | Stromal Cells |
| EMILIN1 | 0.3 | 0 | 0 | 0.46 | 0.23 | 5 | Stromal Cells |
| GPX7 | 0.3 | 0 | 0 | 0.30 | 0.13 | 5 | Stromal Cells |
| HSPB6 | 0.3 | 0 | 0 | 0.32 | 0.14 | 5 | Stromal Cells |
| RPS4X | 0.3 | 0 | 0 | 1.00 | 0.99 | 5 | Stromal Cells |
| EPHX1 | 0.3 | 0 | 0 | 0.39 | 0.26 | 5 | Stromal Cells |
| EIF3L | 0.29 | 0 | 0 | 0.79 | 0.59 | 5 | Stromal Cells |
| ENPP2 | 0.29 | 0 | 0 | 0.27 | 0.13 | 5 | Stromal Cells |
| PDLIM2 | 0.29 | 0 | 0 | 0.41 | 0.21 | 5 | Stromal Cells |
| AC016739.2 | 0.29 | 0 | 0 | 0.97 | 0.91 | 5 | Stromal Cells |
| RPSA | 0.29 | 0 | 0 | 0.97 | 0.89 | 5 | Stromal Cells |
| PLAC9 | 0.29 | 0 | 0 | 0.66 | 0.39 | 5 | Stromal Cells |
| COL27A1 | 0.29 | 0 | 0 | 0.25 | 0.10 | 5 | Stromal Cells |
| NFIA | 0.29 | 0 | 0 | 0.60 | 0.36 | 5 | Stromal Cells |
| AEBP1 | 0.29 | 0 | 0 | 0.38 | 0.19 | 5 | Stromal Cells |
| RPL5P34 | 0.29 | 0 | 0 | 0.90 | 0.74 | 5 | Stromal Cells |
| GPC3 | 0.28 | 0 | 0 | 0.54 | 0.23 | 5 | Stromal Cells |
| FKBP7 | 0.28 | 0 | 0 | 0.36 | 0.18 | 5 | Stromal Cells |
| KRT10 | 0.28 | 0 | 0 | 0.54 | 0.39 | 5 | Stromal Cells |
| RP11-12M9.3 | 0.28 | 0 | 0 | 0.84 | 0.72 | 5 | Stromal Cells |
| RPL3P4 | 0.28 | 0 | 0 | 0.99 | 0.94 | 5 | Stromal Cells |
| IMPDH2 | 0.28 | 0 | 0 | 0.59 | 0.39 | 5 | Stromal Cells |
| RPL6 | 0.28 | 0 | 0 | 1.00 | 0.97 | 5 | Stromal Cells |
| C12orf57 | 0.28 | 0 | 0 | 0.80 | 0.61 | 5 | Stromal Cells |
| RP11-592N21.1 | 0.28 | 0 | 0 | 0.66 | 0.47 | 5 | Stromal Cells |
| RPL6P27 | 0.28 | 0 | 0 | 0.75 | 0.65 | 5 | Stromal Cells |
| BZW2 | 0.28 | 0 | 0 | 0.26 | 0.12 | 5 | Stromal Cells |
| FZD1 | 0.28 | 0 | 0 | 0.27 | 0.11 | 5 | Stromal Cells |
| FAM127A | 0.28 | 0 | 0 | 0.57 | 0.36 | 5 | Stromal Cells |
| RPL5 | 0.28 | 0 | 0 | 0.99 | 0.96 | 5 | Stromal Cells |
| TMED2 | 0.28 | 0 | 0 | 0.69 | 0.51 | 5 | Stromal Cells |
| CNN3 | 0.27 | 0 | 0 | 0.71 | 0.59 | 5 | Stromal Cells |
| SSR2 | 0.27 | 0 | 0 | 0.85 | 0.71 | 5 | Stromal Cells |
| CKAP4 | 0.27 | 0 | 0 | 0.33 | 0.18 | 5 | Stromal Cells |
| RP4-614C10.2 | 0.27 | 0 | 0 | 0.34 | 0.15 | 5 | Stromal Cells |
| EPB41L4A-AS1 | 0.27 | 0 | 0 | 0.48 | 0.27 | 5 | Stromal Cells |
| EIF4A2 | 0.27 | 0 | 0 | 0.87 | 0.75 | 5 | Stromal Cells |
| RPL9 | 0.27 | 0 | 0 | 1.00 | 0.98 | 5 | Stromal Cells |
| PLAGL1 | 0.27 | 0 | 0 | 0.28 | 0.12 | 5 | Stromal Cells |
| CD63 | 0.27 | 0 | 0 | 0.97 | 0.90 | 5 | Stromal Cells |
| HNRNPA1P48 | 0.27 | 0 | 0 | 0.75 | 0.59 | 5 | Stromal Cells |
| KDELR1 | 0.27 | 0 | 0 | 0.56 | 0.37 | 5 | Stromal Cells |
| ZFAS1 | 0.26 | 0 | 0 | 0.75 | 0.58 | 5 | Stromal Cells |
| FN1 | 0.26 | 0 | 0 | 0.73 | 0.43 | 5 | Stromal Cells |
| ARL1 | 0.26 | 0 | 0 | 0.47 | 0.28 | 5 | Stromal Cells |
| RPL7AP66 | 0.26 | 0 | 0 | 0.64 | 0.44 | 5 | Stromal Cells |
| SPRY1 | 0.26 | 0 | 0 | 0.64 | 0.42 | 5 | Stromal Cells |
| ERCC1 | 0.26 | 0 | 0 | 0.38 | 0.23 | 5 | Stromal Cells |
| HOXB5 | 0.26 | 0 | 0 | 0.36 | 0.16 | 5 | Stromal Cells |
| FAM43A | 0.26 | 0 | 0 | 0.25 | 0.14 | 5 | Stromal Cells |
| BASP1 | 0.26 | 0 | 0 | 0.28 | 0.12 | 5 | Stromal Cells |
| RP11-215A21.2 | 0.26 | 0 | 0 | 0.66 | 0.50 | 5 | Stromal Cells |
| FAM114A1 | 0.26 | 0 | 0 | 0.40 | 0.21 | 5 | Stromal Cells |
| RPS4XP6 | 0.26 | 0 | 0 | 0.86 | 0.77 | 5 | Stromal Cells |
| OAT | 0.26 | 0 | 0 | 0.47 | 0.26 | 5 | Stromal Cells |
| TCEAL7 | 0.25 | 0 | 0 | 0.28 | 0.12 | 5 | Stromal Cells |
| RPL7 | 0.25 | 0 | 0 | 1.00 | 0.98 | 5 | Stromal Cells |
| PRSS35 | 1.46 | 0 | 0 | 0.60 | 0.04 | 6 | Pericytes |
| THY1 | 1.41 | 0 | 0 | 0.75 | 0.14 | 6 | Pericytes |
| AGT | 1.4 | 0 | 0 | 0.40 | 0.03 | 6 | Pericytes |
| ID4 | 1.39 | 0 | 0 | 0.74 | 0.23 | 6 | Pericytes |
| COL1A1 | 1.27 | 0 | 0 | 0.99 | 0.59 | 6 | Pericytes |
| C2orf40 | 1.23 | 0 | 0 | 0.62 | 0.13 | 6 | Pericytes |
| KRT18 | 1.23 | 0 | 0 | 0.55 | 0.09 | 6 | Pericytes |
| ELN | 1.23 | 0 | 0 | 0.80 | 0.53 | 6 | Pericytes |
| PLAC9 | 1.17 | 0 | 0 | 0.88 | 0.38 | 6 | Pericytes |
| SEMA5A | 1.16 | 0 | 0 | 0.55 | 0.10 | 6 | Pericytes |
| EGFL6 | 1.07 | 0 | 0 | 0.85 | 0.34 | 6 | Pericytes |
| IGFBP7 | 1.01 | 0 | 0 | 0.99 | 0.65 | 6 | Pericytes |
| OLFML2B | 1.01 | 0 | 0 | 0.44 | 0.06 | 6 | Pericytes |
| SDC2 | 0.99 | 0 | 0 | 0.85 | 0.42 | 6 | Pericytes |
| NTRK3 | 0.99 | 0 | 0 | 0.58 | 0.13 | 6 | Pericytes |
| BGN | 0.98 | 0 | 0 | 0.95 | 0.48 | 6 | Pericytes |
| TINAGL1 | 0.98 | 0 | 0 | 0.83 | 0.36 | 6 | Pericytes |
| SPRY1 | 0.97 | 0 | 0 | 0.82 | 0.41 | 6 | Pericytes |
| FST | 0.95 | 0 | 0 | 0.28 | 0.01 | 6 | Pericytes |
| ARHGAP15 | 0.95 | 0 | 0 | 0.53 | 0.14 | 6 | Pericytes |
| NOTCH3 | 0.93 | 0 | 0 | 0.68 | 0.20 | 6 | Pericytes |
| FOXS1 | 0.92 | 0 | 0 | 0.53 | 0.13 | 6 | Pericytes |
| KCNJ8 | 0.91 | 0 | 0 | 0.50 | 0.13 | 6 | Pericytes |
| GEM | 0.91 | 0 | 0 | 0.43 | 0.13 | 6 | Pericytes |
| ASPN | 0.87 | 0 | 0 | 0.36 | 0.07 | 6 | Pericytes |
| GPM6B | 0.85 | 0 | 0 | 0.67 | 0.27 | 6 | Pericytes |
| ZEB2 | 0.85 | 0 | 0 | 0.78 | 0.40 | 6 | Pericytes |
| GPX3 | 0.84 | 0 | 0 | 0.75 | 0.42 | 6 | Pericytes |
| AC018647.3 | 0.82 | 0 | 0 | 0.27 | 0.02 | 6 | Pericytes |
| GUCY1A3 | 0.8 | 0 | 0 | 0.58 | 0.22 | 6 | Pericytes |
| TIMP1 | 0.8 | 0 | 0 | 0.82 | 0.77 | 6 | Pericytes |
| ISYNA1 | 0.79 | 0 | 0 | 0.48 | 0.19 | 6 | Pericytes |
| SLIT3 | 0.79 | 0 | 0 | 0.48 | 0.16 | 6 | Pericytes |
| COL1A2 | 0.77 | 0 | 0 | 0.97 | 0.63 | 6 | Pericytes |
| 7-Sep | 0.77 | 0 | 0 | 0.91 | 0.70 | 6 | Pericytes |
| NTN4 | 0.76 | 0 | 0 | 0.30 | 0.04 | 6 | Pericytes |
| HES4 | 0.76 | 0 | 0 | 0.62 | 0.24 | 6 | Pericytes |
| SPON2 | 0.75 | 0 | 0 | 0.38 | 0.10 | 6 | Pericytes |
| PRELP | 0.74 | 0 | 0 | 0.36 | 0.07 | 6 | Pericytes |
| COL3A1 | 0.74 | 0 | 0 | 0.89 | 0.57 | 6 | Pericytes |
| EPS8 | 0.74 | 0 | 0 | 0.37 | 0.09 | 6 | Pericytes |
| SPARC | 0.73 | 0 | 0 | 0.99 | 0.84 | 6 | Pericytes |
| TPPP3 | 0.73 | 0 | 0 | 0.79 | 0.39 | 6 | Pericytes |
| CRISPLD2 | 0.7 | 0 | 0 | 0.47 | 0.22 | 6 | Pericytes |
| HIGD1B | 0.7 | 0 | 0 | 0.70 | 0.25 | 6 | Pericytes |
| RGS5 | 0.69 | 0 | 0 | 0.42 | 0.23 | 6 | Pericytes |
| PELO | 0.69 | 0 | 0 | 0.42 | 0.15 | 6 | Pericytes |
| PDGFRB | 0.69 | 0 | 0 | 0.80 | 0.36 | 6 | Pericytes |
| IMPA2 | 0.68 | 0 | 0 | 0.37 | 0.12 | 6 | Pericytes |
| ITGA1 | 0.67 | 0 | 0 | 0.57 | 0.25 | 6 | Pericytes |
| SYTL2 | 0.67 | 0 | 0 | 0.37 | 0.09 | 6 | Pericytes |
| LHFP | 0.67 | 0 | 0 | 0.79 | 0.44 | 6 | Pericytes |
| GUCY1A2 | 0.67 | 0 | 0 | 0.44 | 0.12 | 6 | Pericytes |
| HSPB2 | 0.67 | 0 | 0 | 0.43 | 0.15 | 6 | Pericytes |
| IGFBP5 | 0.67 | 0 | 0 | 0.52 | 0.28 | 6 | Pericytes |
| INPP4B | 0.66 | 0 | 0 | 0.31 | 0.06 | 6 | Pericytes |
| COL14A1 | 0.65 | 0 | 0 | 0.36 | 0.10 | 6 | Pericytes |
| NR2F2 | 0.65 | 0 | 0 | 0.63 | 0.33 | 6 | Pericytes |
| TGFBI | 0.65 | 0 | 0 | 0.69 | 0.33 | 6 | Pericytes |
| MEST | 0.63 | 0 | 0 | 0.71 | 0.36 | 6 | Pericytes |
| PERP | 0.63 | 0 | 0 | 0.37 | 0.11 | 6 | Pericytes |
| APOE | 0.63 | 0 | 0 | 0.55 | 0.30 | 6 | Pericytes |
| ARID5B | 0.63 | 0 | 0 | 0.53 | 0.28 | 6 | Pericytes |
| TIMP3 | 0.62 | 0 | 0 | 0.75 | 0.54 | 6 | Pericytes |
| MRGPRF | 0.61 | 0 | 0 | 0.44 | 0.19 | 6 | Pericytes |
| CPE | 0.6 | 0 | 0 | 0.54 | 0.24 | 6 | Pericytes |
| RERG | 0.6 | 0 | 0 | 0.37 | 0.13 | 6 | Pericytes |
| HEYL | 0.6 | 0 | 0 | 0.35 | 0.12 | 6 | Pericytes |
| REM1 | 0.58 | 0 | 0 | 0.32 | 0.08 | 6 | Pericytes |
| TCEAL7 | 0.57 | 0 | 0 | 0.33 | 0.12 | 6 | Pericytes |
| GRK5 | 0.57 | 0 | 0 | 0.37 | 0.18 | 6 | Pericytes |
| PTK2 | 0.56 | 0 | 0 | 0.56 | 0.29 | 6 | Pericytes |
| ABCC9 | 0.55 | 0 | 0 | 0.28 | 0.05 | 6 | Pericytes |
| HOPX | 0.55 | 0 | 0 | 0.27 | 0.11 | 6 | Pericytes |
| ID3 | 0.55 | 0 | 0 | 0.74 | 0.53 | 6 | Pericytes |
| CD59 | 0.54 | 0 | 0 | 0.83 | 0.59 | 6 | Pericytes |
| GUCY1B3 | 0.53 | 0 | 0 | 0.37 | 0.13 | 6 | Pericytes |
| ARID5A | 0.53 | 0 | 0 | 0.35 | 0.16 | 6 | Pericytes |
| CALD1 | 0.53 | 0 | 0 | 0.97 | 0.78 | 6 | Pericytes |
| LAMB1 | 0.52 | 0 | 0 | 0.37 | 0.14 | 6 | Pericytes |
| FILIP1L | 0.52 | 0 | 0 | 0.59 | 0.26 | 6 | Pericytes |
| TM4SF1 | 0.52 | 0 | 0 | 0.69 | 0.38 | 6 | Pericytes |
| NPR3 | 0.51 | 0 | 0 | 0.25 | 0.08 | 6 | Pericytes |
| MFGE8 | 0.51 | 0 | 0 | 0.56 | 0.32 | 6 | Pericytes |
| GAS6 | 0.5 | 0 | 0 | 0.53 | 0.29 | 6 | Pericytes |
| NDRG2 | 0.5 | 0 | 0 | 0.53 | 0.33 | 6 | Pericytes |
| ENPEP | 0.5 | 0 | 0 | 0.28 | 0.08 | 6 | Pericytes |
| CITED2 | 0.49 | 0 | 0 | 0.36 | 0.23 | 6 | Pericytes |
| FMOD | 0.49 | 0 | 0 | 0.29 | 0.09 | 6 | Pericytes |
| VASN | 0.49 | 0 | 0 | 0.34 | 0.16 | 6 | Pericytes |
| RP4-614C10.2 | 0.49 | 0 | 0 | 0.34 | 0.15 | 6 | Pericytes |
| CRYAB | 0.48 | 0 | 0 | 0.61 | 0.33 | 6 | Pericytes |
| MCAM | 0.47 | 0 | 0 | 0.43 | 0.22 | 6 | Pericytes |
| RP11-693N9.2 | 0.47 | 0 | 0 | 0.34 | 0.15 | 6 | Pericytes |
| CALHM2 | 0.47 | 0 | 0 | 0.26 | 0.11 | 6 | Pericytes |
| MAP1B | 0.47 | 0 | 0 | 0.42 | 0.23 | 6 | Pericytes |
| TMEM176A | 0.47 | 0 | 0 | 0.27 | 0.12 | 6 | Pericytes |
| C6orf48 | 0.47 | 0 | 0 | 0.78 | 0.59 | 6 | Pericytes |
| CCDC102B | 0.46 | 0 | 0 | 0.28 | 0.10 | 6 | Pericytes |
| ADAMTS9 | 0.46 | 0 | 0 | 0.26 | 0.13 | 6 | Pericytes |
| CNN3 | 0.46 | 0 | 0 | 0.79 | 0.58 | 6 | Pericytes |
| C9orf3 | 0.46 | 0 | 0 | 0.50 | 0.30 | 6 | Pericytes |
| OAZ2 | 0.46 | 0 | 0 | 0.73 | 0.53 | 6 | Pericytes |
| CCNI | 0.45 | 0 | 0 | 0.90 | 0.78 | 6 | Pericytes |
| RRAD | 0.45 | 0 | 0 | 0.32 | 0.18 | 6 | Pericytes |
| COL5A1 | 0.44 | 0 | 0 | 0.40 | 0.22 | 6 | Pericytes |
| HSPA2 | 0.44 | 0 | 0 | 0.33 | 0.16 | 6 | Pericytes |
| H1F0 | 0.44 | 0 | 0 | 0.44 | 0.28 | 6 | Pericytes |
| MXRA8 | 0.44 | 0 | 0 | 0.50 | 0.29 | 6 | Pericytes |
| LRRC32 | 0.43 | 0 | 0 | 0.28 | 0.12 | 6 | Pericytes |
| PBX1 | 0.43 | 0 | 0 | 0.40 | 0.27 | 6 | Pericytes |
| SGCE | 0.43 | 0 | 0 | 0.38 | 0.20 | 6 | Pericytes |
| AOC3 | 0.42 | 0 | 0 | 0.51 | 0.33 | 6 | Pericytes |
| COL21A1 | 0.42 | 0 | 0 | 0.28 | 0.13 | 6 | Pericytes |
| FSTL1 | 0.42 | 0 | 0 | 0.59 | 0.43 | 6 | Pericytes |
| 4-Sep | 0.42 | 0 | 0 | 0.30 | 0.14 | 6 | Pericytes |
| SRPX | 0.41 | 0 | 0 | 0.31 | 0.15 | 6 | Pericytes |
| COX4I2 | 0.41 | 0 | 0 | 0.56 | 0.25 | 6 | Pericytes |
| CDH6 | 0.41 | 0 | 0 | 0.26 | 0.11 | 6 | Pericytes |
| LINC00152 | 0.41 | 0 | 0 | 0.26 | 0.10 | 6 | Pericytes |
| UBE2E2 | 0.4 | 0 | 0 | 0.32 | 0.17 | 6 | Pericytes |
| ADGRF5 | 0.4 | 0 | 0 | 0.32 | 0.17 | 6 | Pericytes |
| EBF1 | 0.39 | 0 | 0 | 0.26 | 0.10 | 6 | Pericytes |
| SERPINI1 | 0.39 | 0 | 0 | 0.26 | 0.12 | 6 | Pericytes |
| NID1 | 0.39 | 0 | 0 | 0.39 | 0.21 | 6 | Pericytes |
| AXL | 0.39 | 0 | 0 | 0.37 | 0.22 | 6 | Pericytes |
| FXYD1 | 0.39 | 0 | 0 | 0.44 | 0.26 | 6 | Pericytes |
| CHN1 | 0.39 | 0 | 0 | 0.50 | 0.25 | 6 | Pericytes |
| PLEKHA4 | 0.38 | 0 | 0 | 0.27 | 0.13 | 6 | Pericytes |
| CYR61 | 0.38 | 0 | 0.01 | 0.32 | 0.23 | 6 | Pericytes |
| TXNIP | 0.38 | 0 | 0 | 0.78 | 0.68 | 6 | Pericytes |
| BASP1 | 0.38 | 0 | 0 | 0.25 | 0.13 | 6 | Pericytes |
| ITM2C | 0.38 | 0 | 0 | 0.57 | 0.39 | 6 | Pericytes |
| LTBP2 | 0.37 | 0 | 0 | 0.26 | 0.13 | 6 | Pericytes |
| TUBA1A | 0.37 | 0 | 0 | 0.90 | 0.79 | 6 | Pericytes |
| EEF2 | 0.36 | 0 | 0 | 0.97 | 0.88 | 6 | Pericytes |
| H2AFJ | 0.36 | 0 | 0 | 0.50 | 0.39 | 6 | Pericytes |
| AKAP12 | 0.36 | 0 | 0 | 0.34 | 0.22 | 6 | Pericytes |
| CALU | 0.36 | 0 | 0 | 0.51 | 0.38 | 6 | Pericytes |
| MPRIP | 0.35 | 0 | 0 | 0.34 | 0.20 | 6 | Pericytes |
| CALM2 | 0.35 | 0 | 0 | 0.98 | 0.90 | 6 | Pericytes |
| FILIP1 | 0.35 | 0 | 0 | 0.32 | 0.18 | 6 | Pericytes |
| MAP7D3 | 0.35 | 0 | 0 | 0.28 | 0.15 | 6 | Pericytes |
| ADI1 | 0.35 | 0 | 0 | 0.36 | 0.26 | 6 | Pericytes |
| PHLDA1 | 0.34 | 0 | 0 | 0.35 | 0.23 | 6 | Pericytes |
| CAMLG | 0.34 | 0 | 0 | 0.46 | 0.34 | 6 | Pericytes |
| MT1E | 0.34 | 0 | 0 | 0.38 | 0.23 | 6 | Pericytes |
| TMEM47 | 0.33 | 0 | 0 | 0.30 | 0.17 | 6 | Pericytes |
| PTMA | 0.32 | 0 | 0 | 1.00 | 1.00 | 6 | Pericytes |
| TPM4 | 0.32 | 0 | 0 | 0.74 | 0.59 | 6 | Pericytes |
| RPL4P4 | 0.32 | 0 | 0 | 0.46 | 0.35 | 6 | Pericytes |
| MEF2C | 0.32 | 0 | 0 | 0.57 | 0.42 | 6 | Pericytes |
| TMEM176B | 0.32 | 0 | 0 | 0.30 | 0.17 | 6 | Pericytes |
| NT5DC2 | 0.32 | 0 | 0 | 0.28 | 0.15 | 6 | Pericytes |
| ITGB1 | 0.32 | 0 | 0 | 0.72 | 0.63 | 6 | Pericytes |
| EIF3L | 0.31 | 0 | 0 | 0.73 | 0.60 | 6 | Pericytes |
| PIK3R1 | 0.31 | 0 | 0 | 0.26 | 0.16 | 6 | Pericytes |
| H3F3B | 0.31 | 0 | 0 | 1.00 | 0.97 | 6 | Pericytes |
| PXDN | 0.3 | 0 | 0 | 0.31 | 0.20 | 6 | Pericytes |
| RFTN1 | 0.3 | 0 | 0 | 0.28 | 0.15 | 6 | Pericytes |
| MID1IP1 | 0.3 | 0 | 0 | 0.25 | 0.15 | 6 | Pericytes |
| PTMAP5 | 0.3 | 0 | 0 | 0.76 | 0.69 | 6 | Pericytes |
| RPL3P4 | 0.3 | 0 | 0 | 0.99 | 0.94 | 6 | Pericytes |
| LURAP1L | 0.3 | 0 | 0 | 0.31 | 0.19 | 6 | Pericytes |
| SOD3 | 0.3 | 0 | 0 | 0.71 | 0.41 | 6 | Pericytes |
| PTMAP2 | 0.3 | 0 | 0 | 0.68 | 0.61 | 6 | Pericytes |
| PCOLCE | 0.3 | 0 | 0 | 0.52 | 0.30 | 6 | Pericytes |
| KANK2 | 0.3 | 0 | 0 | 0.49 | 0.34 | 6 | Pericytes |
| PMP22 | 0.29 | 0 | 0 | 0.70 | 0.47 | 6 | Pericytes |
| CBX6 | 0.29 | 0 | 0 | 0.41 | 0.29 | 6 | Pericytes |
| CCDC80 | 0.29 | 0 | 0 | 0.42 | 0.22 | 6 | Pericytes |
| ZNF22 | 0.29 | 0 | 0.02 | 0.41 | 0.33 | 6 | Pericytes |
| RPL6 | 0.29 | 0 | 0 | 0.99 | 0.97 | 6 | Pericytes |
| FOXP1 | 0.29 | 0 | 0 | 0.56 | 0.39 | 6 | Pericytes |
| TCEAL9 | 0.29 | 0 | 0 | 0.67 | 0.53 | 6 | Pericytes |
| MYO1B | 0.29 | 0 | 0 | 0.32 | 0.20 | 6 | Pericytes |
| MRPL32 | 0.29 | 0 | 0.01 | 0.32 | 0.23 | 6 | Pericytes |
| SOX4 | 0.29 | 0 | 0 | 0.86 | 0.75 | 6 | Pericytes |
| RP11-371A22.1 | 0.29 | 0 | 0 | 0.92 | 0.85 | 6 | Pericytes |
| ACTN1 | 0.28 | 0 | 0 | 0.56 | 0.44 | 6 | Pericytes |
| LASP1 | 0.28 | 0 | 0.03 | 0.28 | 0.19 | 6 | Pericytes |
| EEF1A1P5 | 0.28 | 0 | 0 | 0.75 | 0.65 | 6 | Pericytes |
| SORBS3 | 0.28 | 0 | 0 | 0.32 | 0.19 | 6 | Pericytes |
| KLF6 | 0.28 | 0 | 0 | 0.55 | 0.42 | 6 | Pericytes |
| GAS5 | 0.28 | 0 | 0 | 0.82 | 0.71 | 6 | Pericytes |
| BBX | 0.27 | 0 | 0 | 0.39 | 0.30 | 6 | Pericytes |
| PPP1R12A | 0.27 | 0 | 0 | 0.59 | 0.46 | 6 | Pericytes |
| EEF1D | 0.27 | 0 | 0 | 0.94 | 0.89 | 6 | Pericytes |
| RPL14P1 | 0.27 | 0 | 0 | 0.66 | 0.57 | 6 | Pericytes |
| GPC3 | 0.27 | 0 | 0 | 0.50 | 0.24 | 6 | Pericytes |
| RPLP0 | 0.27 | 0 | 0 | 0.97 | 0.93 | 6 | Pericytes |
| RPL6P27 | 0.27 | 0 | 0 | 0.72 | 0.65 | 6 | Pericytes |
| RPL14 | 0.27 | 0 | 0 | 0.99 | 0.95 | 6 | Pericytes |
| EIF2S3 | 0.27 | 0 | 0 | 0.42 | 0.31 | 6 | Pericytes |
| RPL3 | 0.27 | 0 | 0 | 1.00 | 0.99 | 6 | Pericytes |
| TPT1 | 0.26 | 0 | 0 | 0.99 | 0.98 | 6 | Pericytes |
| AP3S1 | 0.26 | 0 | 0.01 | 0.41 | 0.31 | 6 | Pericytes |
| BEX3 | 0.26 | 0 | 0 | 0.77 | 0.67 | 6 | Pericytes |
| LGALS1 | 0.26 | 0 | 0 | 0.97 | 0.89 | 6 | Pericytes |
| H19 | 0.26 | 0 | 0 | 0.31 | 0.15 | 6 | Pericytes |
| SPTSSA | 0.26 | 0 | 0.05 | 0.44 | 0.35 | 6 | Pericytes |
| TPM1 | 0.26 | 0 | 0 | 0.74 | 0.59 | 6 | Pericytes |
| EIF4A2 | 0.25 | 0 | 0 | 0.82 | 0.75 | 6 | Pericytes |
| EEF1A1 | 0.25 | 0 | 0 | 1.00 | 1.00 | 6 | Pericytes |
| 11-Sep | 0.25 | 0 | 0 | 0.42 | 0.30 | 6 | Pericytes |
| FEZ1 | 0.25 | 0 | 0.01 | 0.26 | 0.17 | 6 | Pericytes |
| RPL21 | 0.25 | 0 | 0 | 1.00 | 0.99 | 6 | Pericytes |
| PEBP1 | 0.25 | 0 | 0 | 0.79 | 0.67 | 6 | Pericytes |
| EZR | 0.25 | 0 | 0.01 | 0.34 | 0.25 | 6 | Pericytes |
| S100A8 | 4.21 | 0 | 0 | 0.74 | 0.15 | 7 | Macrophages |
| S100A9 | 3.96 | 0 | 0 | 0.80 | 0.17 | 7 | Macrophages |
| LYZ | 3.84 | 0 | 0 | 0.93 | 0.11 | 7 | Macrophages |
| HLA-DRA | 3.53 | 0 | 0 | 0.97 | 0.22 | 7 | Macrophages |
| HLA-DRB1 | 3.19 | 0 | 0 | 0.90 | 0.07 | 7 | Macrophages |
| TYROBP | 3.12 | 0 | 0 | 0.97 | 0.08 | 7 | Macrophages |
| HLA-DPA1 | 3.07 | 0 | 0 | 0.86 | 0.12 | 7 | Macrophages |
| HLA-DPB1 | 3 | 0 | 0 | 0.83 | 0.14 | 7 | Macrophages |
| AIF1 | 2.95 | 0 | 0 | 0.97 | 0.06 | 7 | Macrophages |
| S100A12 | 2.94 | 0 | 0 | 0.44 | 0.02 | 7 | Macrophages |
| FCER1G | 2.88 | 0 | 0 | 0.97 | 0.09 | 7 | Macrophages |
| CD74 | 2.77 | 0 | 0 | 0.97 | 0.42 | 7 | Macrophages |
| CD14 | 2.75 | 0 | 0 | 0.79 | 0.06 | 7 | Macrophages |
| SRGN | 2.59 | 0 | 0 | 0.97 | 0.18 | 7 | Macrophages |
| TYMP | 2.48 | 0 | 0 | 0.94 | 0.11 | 7 | Macrophages |
| IFI30 | 2.47 | 0 | 0 | 0.90 | 0.02 | 7 | Macrophages |
| FTL | 2.34 | 0 | 0 | 1.00 | 0.96 | 7 | Macrophages |
| LST1 | 2.28 | 0 | 0 | 0.92 | 0.03 | 7 | Macrophages |
| COTL1 | 2.28 | 0 | 0 | 0.92 | 0.16 | 7 | Macrophages |
| CTSS | 2.25 | 0 | 0 | 0.86 | 0.06 | 7 | Macrophages |
| FCN1 | 2.18 | 0 | 0 | 0.66 | 0.02 | 7 | Macrophages |
| CST3 | 2.13 | 0 | 0 | 0.99 | 0.57 | 7 | Macrophages |
| SPI1 | 2.1 | 0 | 0 | 0.92 | 0.02 | 7 | Macrophages |
| FTLP3 | 2.1 | 0 | 0 | 0.95 | 0.48 | 7 | Macrophages |
| MNDA | 2.08 | 0 | 0 | 0.77 | 0.01 | 7 | Macrophages |
| SH3BGRL3 | 2.05 | 0 | 0 | 0.97 | 0.56 | 7 | Macrophages |
| HLA-DQB1 | 2.03 | 0 | 0 | 0.60 | 0.01 | 7 | Macrophages |
| S100A4 | 2.03 | 0 | 0 | 0.92 | 0.36 | 7 | Macrophages |
| RETN | 1.93 | 0 | 0 | 0.42 | 0.01 | 7 | Macrophages |
| MS4A6A | 1.92 | 0 | 0 | 0.71 | 0.02 | 7 | Macrophages |
| CYBA | 1.92 | 0 | 0 | 0.98 | 0.49 | 7 | Macrophages |
| LAPTM5 | 1.89 | 0 | 0 | 0.89 | 0.11 | 7 | Macrophages |
| SERPINA1 | 1.83 | 0 | 0 | 0.71 | 0.02 | 7 | Macrophages |
| CD52 | 1.78 | 0 | 0 | 0.71 | 0.07 | 7 | Macrophages |
| RP11-1143G9.4 | 1.71 | 0 | 0 | 0.44 | 0.03 | 7 | Macrophages |
| UCP2 | 1.7 | 0 | 0 | 0.81 | 0.08 | 7 | Macrophages |
| CORO1A | 1.68 | 0 | 0 | 0.87 | 0.07 | 7 | Macrophages |
| HCST | 1.65 | 0 | 0 | 0.87 | 0.04 | 7 | Macrophages |
| C1orf162 | 1.65 | 0 | 0 | 0.78 | 0.03 | 7 | Macrophages |
| CPVL | 1.62 | 0 | 0 | 0.70 | 0.01 | 7 | Macrophages |
| LSP1 | 1.62 | 0 | 0 | 0.78 | 0.06 | 7 | Macrophages |
| SAT1 | 1.61 | 0 | 0 | 0.98 | 0.54 | 7 | Macrophages |
| CTSB | 1.61 | 0 | 0 | 0.81 | 0.28 | 7 | Macrophages |
| S100A11 | 1.6 | 0 | 0 | 0.97 | 0.58 | 7 | Macrophages |
| HLA-DMA | 1.59 | 0 | 0 | 0.68 | 0.06 | 7 | Macrophages |
| LCP1 | 1.54 | 0 | 0 | 0.77 | 0.04 | 7 | Macrophages |
| ANXA1 | 1.54 | 0 | 0 | 0.89 | 0.37 | 7 | Macrophages |
| PSAP | 1.52 | 0 | 0 | 0.93 | 0.62 | 7 | Macrophages |
| CD68 | 1.51 | 0 | 0 | 0.72 | 0.02 | 7 | Macrophages |
| FCGR3A | 1.49 | 0 | 0 | 0.45 | 0.03 | 7 | Macrophages |
| MS4A7 | 1.48 | 0 | 0 | 0.55 | 0.01 | 7 | Macrophages |
| CD53 | 1.45 | 0 | 0 | 0.80 | 0.05 | 7 | Macrophages |
| STXBP2 | 1.45 | 0 | 0 | 0.69 | 0.04 | 7 | Macrophages |
| CSTA | 1.43 | 0 | 0 | 0.63 | 0.01 | 7 | Macrophages |
| HCK | 1.43 | 0 | 0 | 0.73 | 0.01 | 7 | Macrophages |
| VAMP8 | 1.42 | 0 | 0 | 0.84 | 0.14 | 7 | Macrophages |
| HMOX1 | 1.41 | 0 | 0 | 0.47 | 0.01 | 7 | Macrophages |
| LINC01272 | 1.4 | 0 | 0 | 0.44 | 0.01 | 7 | Macrophages |
| FTH1P10 | 1.38 | 0 | 0 | 0.95 | 0.67 | 7 | Macrophages |
| FTH1 | 1.38 | 0 | 0 | 1.00 | 0.98 | 7 | Macrophages |
| CFP | 1.37 | 0 | 0 | 0.64 | 0.03 | 7 | Macrophages |
| PTPRC | 1.37 | 0 | 0 | 0.76 | 0.04 | 7 | Macrophages |
| FGL2 | 1.34 | 0 | 0 | 0.71 | 0.02 | 7 | Macrophages |
| FCGR2A | 1.34 | 0 | 0 | 0.56 | 0.01 | 7 | Macrophages |
| RNASET2 | 1.34 | 0 | 0 | 0.79 | 0.14 | 7 | Macrophages |
| CLEC10A | 1.33 | 0 | 0 | 0.39 | 0.01 | 7 | Macrophages |
| ITGB2 | 1.32 | 0 | 0 | 0.73 | 0.04 | 7 | Macrophages |
| GPX1 | 1.32 | 0 | 0 | 0.93 | 0.50 | 7 | Macrophages |
| VCAN | 1.32 | 0 | 0 | 0.63 | 0.33 | 7 | Macrophages |
| TIMP1 | 1.31 | 0 | 0 | 0.96 | 0.76 | 7 | Macrophages |
| LILRA5 | 1.3 | 0 | 0 | 0.60 | 0.00 | 7 | Macrophages |
| CSF1R | 1.27 | 0 | 0 | 0.57 | 0.01 | 7 | Macrophages |
| JAML | 1.25 | 0 | 0 | 0.57 | 0.01 | 7 | Macrophages |
| CYBB | 1.25 | 0 | 0 | 0.60 | 0.01 | 7 | Macrophages |
| RHOG | 1.25 | 0 | 0 | 0.79 | 0.17 | 7 | Macrophages |
| HLA-DQA1 | 1.24 | 0 | 0 | 0.47 | 0.01 | 7 | Macrophages |
| NEAT1 | 1.24 | 0 | 0 | 0.94 | 0.62 | 7 | Macrophages |
| CD44 | 1.22 | 0 | 0 | 0.81 | 0.23 | 7 | Macrophages |
| PYCARD | 1.22 | 0 | 0 | 0.71 | 0.11 | 7 | Macrophages |
| ARPC1B | 1.21 | 0 | 0 | 0.87 | 0.27 | 7 | Macrophages |
| CRIP1 | 1.2 | 0 | 0 | 0.82 | 0.51 | 7 | Macrophages |
| LGALS2 | 1.2 | 0 | 0 | 0.49 | 0.01 | 7 | Macrophages |
| PLAUR | 1.2 | 0 | 0 | 0.58 | 0.05 | 7 | Macrophages |
| MPEG1 | 1.18 | 0 | 0 | 0.61 | 0.01 | 7 | Macrophages |
| PTPN6 | 1.17 | 0 | 0 | 0.69 | 0.04 | 7 | Macrophages |
| SAMHD1 | 1.17 | 0 | 0 | 0.72 | 0.10 | 7 | Macrophages |
| DOK2 | 1.17 | 0 | 0 | 0.61 | 0.02 | 7 | Macrophages |
| PLAC8 | 1.16 | 0 | 0 | 0.48 | 0.05 | 7 | Macrophages |
| IGSF6 | 1.16 | 0 | 0 | 0.56 | 0.00 | 7 | Macrophages |
| CTSD | 1.16 | 0 | 0 | 0.64 | 0.30 | 7 | Macrophages |
| FOS | 1.16 | 0 | 0 | 0.82 | 0.32 | 7 | Macrophages |
| ARHGDIB | 1.15 | 0 | 0 | 0.86 | 0.27 | 7 | Macrophages |
| FAM49B | 1.14 | 0 | 0 | 0.76 | 0.10 | 7 | Macrophages |
| FCGR1A | 1.13 | 0 | 0 | 0.51 | 0.01 | 7 | Macrophages |
| CAPG | 1.13 | 0 | 0 | 0.53 | 0.05 | 7 | Macrophages |
| BCL2A1 | 1.12 | 0 | 0 | 0.50 | 0.01 | 7 | Macrophages |
| LILRB2 | 1.12 | 0 | 0 | 0.58 | 0.01 | 7 | Macrophages |
| PKM | 1.12 | 0 | 0 | 0.87 | 0.44 | 7 | Macrophages |
| NPC2 | 1.12 | 0 | 0 | 0.91 | 0.47 | 7 | Macrophages |
| CD300A | 1.11 | 0 | 0 | 0.58 | 0.02 | 7 | Macrophages |
| NCF2 | 1.09 | 0 | 0 | 0.61 | 0.01 | 7 | Macrophages |
| HCLS1 | 1.09 | 0 | 0 | 0.63 | 0.03 | 7 | Macrophages |
| GPR183 | 1.08 | 0 | 0 | 0.48 | 0.05 | 7 | Macrophages |
| CTSL | 1.08 | 0 | 0 | 0.45 | 0.28 | 7 | Macrophages |
| ARPC3 | 1.08 | 0 | 0 | 0.92 | 0.59 | 7 | Macrophages |
| HLA-DQA2 | 1.07 | 0 | 0 | 0.44 | 0.01 | 7 | Macrophages |
| HLA-DMB | 1.07 | 0 | 0 | 0.51 | 0.04 | 7 | Macrophages |
| TPM3 | 1.07 | 0 | 0 | 0.82 | 0.28 | 7 | Macrophages |
| LRRC25 | 1.06 | 0 | 0 | 0.55 | 0.00 | 7 | Macrophages |
| CEBPB | 1.06 | 0 | 0 | 0.77 | 0.28 | 7 | Macrophages |
| TSPO | 1.06 | 0 | 0 | 0.81 | 0.41 | 7 | Macrophages |
| CTSH | 1.06 | 0 | 0 | 0.68 | 0.10 | 7 | Macrophages |
| CD48 | 1.05 | 0 | 0 | 0.56 | 0.03 | 7 | Macrophages |
| EMP3 | 1.05 | 0 | 0 | 0.93 | 0.57 | 7 | Macrophages |
| KYNU | 1.05 | 0 | 0 | 0.54 | 0.01 | 7 | Macrophages |
| FPR1 | 1.03 | 0 | 0 | 0.49 | 0.01 | 7 | Macrophages |
| SLC16A3 | 1.02 | 0 | 0 | 0.63 | 0.07 | 7 | Macrophages |
| THEMIS2 | 1.02 | 0 | 0 | 0.60 | 0.01 | 7 | Macrophages |
| CLIC1 | 1.02 | 0 | 0 | 0.95 | 0.62 | 7 | Macrophages |
| FGR | 1.01 | 0 | 0 | 0.57 | 0.03 | 7 | Macrophages |
| ENO1 | 1.01 | 0 | 0 | 0.92 | 0.54 | 7 | Macrophages |
| PILRA | 1 | 0 | 0 | 0.51 | 0.01 | 7 | Macrophages |
| NAMPT | 1 | 0 | 0 | 0.56 | 0.11 | 7 | Macrophages |
| APOBEC3A | 0.99 | 0 | 0 | 0.26 | 0.01 | 7 | Macrophages |
| ACTB | 0.98 | 0 | 0 | 1.00 | 0.99 | 7 | Macrophages |
| C10orf54 | 0.97 | 0 | 0 | 0.70 | 0.17 | 7 | Macrophages |
| S100A10 | 0.97 | 0 | 0 | 0.88 | 0.64 | 7 | Macrophages |
| CLEC7A | 0.96 | 0 | 0 | 0.51 | 0.00 | 7 | Macrophages |
| GMFG | 0.96 | 0 | 0 | 0.84 | 0.24 | 7 | Macrophages |
| IL1B | 0.96 | 0 | 0 | 0.31 | 0.01 | 7 | Macrophages |
| EFHD2 | 0.96 | 0 | 0 | 0.63 | 0.08 | 7 | Macrophages |
| LY86 | 0.96 | 0 | 0 | 0.54 | 0.01 | 7 | Macrophages |
| RAC2 | 0.95 | 0 | 0 | 0.66 | 0.05 | 7 | Macrophages |
| OAZ1 | 0.95 | 0 | 0 | 0.97 | 0.84 | 7 | Macrophages |
| PTPRE | 0.95 | 0 | 0 | 0.56 | 0.04 | 7 | Macrophages |
| CSTB | 0.95 | 0 | 0 | 0.86 | 0.47 | 7 | Macrophages |
| GCA | 0.95 | 0 | 0 | 0.56 | 0.08 | 7 | Macrophages |
| MS4A4A | 0.94 | 0 | 0 | 0.36 | 0.01 | 7 | Macrophages |
| DUSP1 | 0.94 | 0 | 0 | 0.84 | 0.39 | 7 | Macrophages |
| ARRB2 | 0.94 | 0 | 0 | 0.58 | 0.06 | 7 | Macrophages |
| HSPA1B | 0.93 | 0 | 0 | 0.85 | 0.35 | 7 | Macrophages |
| BID | 0.93 | 0 | 0 | 0.59 | 0.09 | 7 | Macrophages |
| LYN | 0.92 | 0 | 0 | 0.58 | 0.06 | 7 | Macrophages |
| GLIPR2 | 0.91 | 0 | 0 | 0.68 | 0.19 | 7 | Macrophages |
| FERMT3 | 0.91 | 0 | 0 | 0.57 | 0.03 | 7 | Macrophages |
| GAPDH | 0.91 | 0 | 0 | 0.99 | 0.89 | 7 | Macrophages |
| CAP1 | 0.9 | 0 | 0 | 0.84 | 0.44 | 7 | Macrophages |
| CD86 | 0.89 | 0 | 0 | 0.48 | 0.01 | 7 | Macrophages |
| PFN1 | 0.89 | 0 | 0 | 0.98 | 0.85 | 7 | Macrophages |
| TNFSF13B | 0.89 | 0 | 0 | 0.47 | 0.03 | 7 | Macrophages |
| ZFP36 | 0.88 | 0 | 0 | 0.78 | 0.28 | 7 | Macrophages |
| GLIPR1 | 0.88 | 0 | 0 | 0.55 | 0.06 | 7 | Macrophages |
| RGS10 | 0.87 | 0 | 0 | 0.51 | 0.04 | 7 | Macrophages |
| CTSC | 0.87 | 0 | 0 | 0.59 | 0.21 | 7 | Macrophages |
| C4orf48 | 0.87 | 0 | 0 | 0.67 | 0.11 | 7 | Macrophages |
| MAFB | 0.86 | 0 | 0 | 0.66 | 0.22 | 7 | Macrophages |
| FAM26F | 0.86 | 0 | 0 | 0.36 | 0.02 | 7 | Macrophages |
| CTSZ | 0.85 | 0 | 0 | 0.61 | 0.15 | 7 | Macrophages |
| FYB | 0.85 | 0 | 0 | 0.49 | 0.02 | 7 | Macrophages |
| GPSM3 | 0.85 | 0 | 0 | 0.62 | 0.11 | 7 | Macrophages |
| CD1C | 0.85 | 0 | 0 | 0.25 | 0.01 | 7 | Macrophages |
| FTH1P8 | 0.84 | 0 | 0 | 0.71 | 0.32 | 7 | Macrophages |
| TBXAS1 | 0.84 | 0 | 0 | 0.48 | 0.01 | 7 | Macrophages |
| ACTR3 | 0.84 | 0 | 0 | 0.77 | 0.32 | 7 | Macrophages |
| LY96 | 0.84 | 0 | 0 | 0.49 | 0.03 | 7 | Macrophages |
| ABI3 | 0.83 | 0 | 0 | 0.48 | 0.07 | 7 | Macrophages |
| TNFRSF1B | 0.83 | 0 | 0 | 0.47 | 0.04 | 7 | Macrophages |
| BLVRB | 0.83 | 0 | 0 | 0.59 | 0.19 | 7 | Macrophages |
| PLEK | 0.83 | 0 | 0 | 0.43 | 0.02 | 7 | Macrophages |
| FPR3 | 0.82 | 0 | 0 | 0.29 | 0.01 | 7 | Macrophages |
| PGK1 | 0.82 | 0 | 0 | 0.80 | 0.41 | 7 | Macrophages |
| HSPA1A | 0.82 | 0 | 0 | 0.92 | 0.62 | 7 | Macrophages |
| PLIN2 | 0.82 | 0 | 0 | 0.32 | 0.09 | 7 | Macrophages |
| GRN | 0.82 | 0 | 0 | 0.80 | 0.38 | 7 | Macrophages |
| FXYD5 | 0.81 | 0 | 0 | 0.77 | 0.23 | 7 | Macrophages |
| PRELID1 | 0.81 | 0 | 0 | 0.81 | 0.46 | 7 | Macrophages |
| CD55 | 0.81 | 0 | 0 | 0.53 | 0.10 | 7 | Macrophages |
| ARPC2 | 0.81 | 0 | 0 | 0.92 | 0.62 | 7 | Macrophages |
| LIPA | 0.81 | 0 | 0 | 0.49 | 0.12 | 7 | Macrophages |
| TREM1 | 0.8 | 0 | 0 | 0.38 | 0.01 | 7 | Macrophages |
| LINC00152 | 0.8 | 0 | 0 | 0.50 | 0.09 | 7 | Macrophages |
| ABRACL | 0.8 | 0 | 0 | 0.57 | 0.10 | 7 | Macrophages |
| CASP1 | 0.8 | 0 | 0 | 0.56 | 0.10 | 7 | Macrophages |
| RNASE6 | 0.79 | 0 | 0 | 0.44 | 0.01 | 7 | Macrophages |
| SERPINB1 | 0.79 | 0 | 0 | 0.60 | 0.17 | 7 | Macrophages |
| MYO1F | 0.79 | 0 | 0 | 0.48 | 0.03 | 7 | Macrophages |
| SOD2 | 0.79 | 0 | 0 | 0.53 | 0.18 | 7 | Macrophages |
| CD300E | 0.79 | 0 | 0 | 0.36 | 0.00 | 7 | Macrophages |
| LILRB3 | 0.79 | 0 | 0 | 0.48 | 0.01 | 7 | Macrophages |
| AP2S1 | 0.78 | 0 | 0 | 0.83 | 0.43 | 7 | Macrophages |
| LRRFIP1 | 0.78 | 0 | 0 | 0.77 | 0.25 | 7 | Macrophages |
| S100A6 | 0.78 | 0 | 0 | 0.96 | 0.83 | 7 | Macrophages |
| VASP | 0.78 | 0 | 0 | 0.68 | 0.24 | 7 | Macrophages |
| CYTIP | 0.77 | 0 | 0 | 0.40 | 0.02 | 7 | Macrophages |
| CXCL16 | 0.77 | 0 | 0 | 0.41 | 0.06 | 7 | Macrophages |
| CKLF | 0.76 | 0 | 0 | 0.60 | 0.16 | 7 | Macrophages |
| CCR1 | 0.76 | 0 | 0 | 0.40 | 0.01 | 7 | Macrophages |
| STAB1 | 0.76 | 0 | 0 | 0.31 | 0.02 | 7 | Macrophages |
| SLC11A1 | 0.76 | 0 | 0 | 0.37 | 0.02 | 7 | Macrophages |
| ATP6V0B | 0.76 | 0 | 0 | 0.77 | 0.36 | 7 | Macrophages |
| WAS | 0.76 | 0 | 0 | 0.46 | 0.01 | 7 | Macrophages |
| GSTO1 | 0.75 | 0 | 0 | 0.77 | 0.38 | 7 | Macrophages |
| MCL1 | 0.75 | 0 | 0 | 0.68 | 0.27 | 7 | Macrophages |
| CXCR4 | 0.75 | 0 | 0 | 0.51 | 0.08 | 7 | Macrophages |
| CPPED1 | 0.73 | 0 | 0 | 0.45 | 0.04 | 7 | Macrophages |
| MOB1A | 0.73 | 0 | 0 | 0.64 | 0.17 | 7 | Macrophages |
| CYTH4 | 0.73 | 0 | 0 | 0.41 | 0.01 | 7 | Macrophages |
| PRR13 | 0.73 | 0 | 0 | 0.70 | 0.23 | 7 | Macrophages |
| AP1S2 | 0.73 | 0 | 0 | 0.63 | 0.22 | 7 | Macrophages |
| SAMSN1 | 0.73 | 0 | 0 | 0.41 | 0.01 | 7 | Macrophages |
| ACTR2 | 0.73 | 0 | 0 | 0.73 | 0.31 | 7 | Macrophages |
| RGS19 | 0.72 | 0 | 0 | 0.49 | 0.05 | 7 | Macrophages |
| MT-CO1 | 0.72 | 0 | 0 | 0.99 | 1.00 | 7 | Macrophages |
| EVI2B | 0.72 | 0 | 0 | 0.43 | 0.01 | 7 | Macrophages |
| ATP5E | 0.72 | 0 | 0 | 0.99 | 0.87 | 7 | Macrophages |
| MRC1 | 0.71 | 0 | 0 | 0.27 | 0.01 | 7 | Macrophages |
| PLSCR1 | 0.71 | 0 | 0 | 0.61 | 0.15 | 7 | Macrophages |
| PARVG | 0.71 | 0 | 0 | 0.44 | 0.01 | 7 | Macrophages |
| ASAH1 | 0.71 | 0 | 0 | 0.66 | 0.27 | 7 | Macrophages |
| LILRB4 | 0.7 | 0 | 0 | 0.40 | 0.00 | 7 | Macrophages |
| MBOAT7 | 0.7 | 0 | 0 | 0.48 | 0.08 | 7 | Macrophages |
| SERPINB9 | 0.7 | 0 | 0 | 0.44 | 0.08 | 7 | Macrophages |
| TMEM176B | 0.7 | 0 | 0 | 0.46 | 0.16 | 7 | Macrophages |
| STK17B | 0.7 | 0 | 0 | 0.45 | 0.03 | 7 | Macrophages |
| NAPSB | 0.7 | 0 | 0 | 0.35 | 0.01 | 7 | Macrophages |
| DUSP6 | 0.69 | 0 | 0 | 0.59 | 0.17 | 7 | Macrophages |
| C5AR1 | 0.69 | 0 | 0 | 0.36 | 0.01 | 7 | Macrophages |
| LAIR1 | 0.69 | 0 | 0 | 0.37 | 0.02 | 7 | Macrophages |
| ARPC4 | 0.69 | 0 | 0 | 0.72 | 0.32 | 7 | Macrophages |
| VMP1 | 0.69 | 0 | 0 | 0.68 | 0.25 | 7 | Macrophages |
| LGALS1 | 0.68 | 0 | 0 | 0.98 | 0.89 | 7 | Macrophages |
| CDC42 | 0.68 | 0 | 0 | 0.88 | 0.64 | 7 | Macrophages |
| FABP5 | 0.68 | 0 | 0 | 0.50 | 0.26 | 7 | Macrophages |
| RP11-386I14.4 | 0.68 | 0 | 0 | 0.34 | 0.04 | 7 | Macrophages |
| CD37 | 0.67 | 0 | 0 | 0.57 | 0.05 | 7 | Macrophages |
| MFSD1 | 0.67 | 0 | 0 | 0.52 | 0.09 | 7 | Macrophages |
| ANXA2 | 0.67 | 0 | 0 | 0.90 | 0.64 | 7 | Macrophages |
| RNF149 | 0.67 | 0 | 0 | 0.48 | 0.09 | 7 | Macrophages |
| PPT1 | 0.67 | 0 | 0 | 0.59 | 0.22 | 7 | Macrophages |
| PLA2G7 | 0.67 | 0 | 0 | 0.31 | 0.00 | 7 | Macrophages |
| RGS2 | 0.66 | 0 | 0 | 0.65 | 0.29 | 7 | Macrophages |
| TPP1 | 0.66 | 0 | 0 | 0.47 | 0.09 | 7 | Macrophages |
| CFL1 | 0.66 | 0 | 0 | 0.98 | 0.90 | 7 | Macrophages |
| CAPZA1 | 0.66 | 0 | 0 | 0.70 | 0.26 | 7 | Macrophages |
| ATP6V0D1 | 0.66 | 0 | 0 | 0.59 | 0.21 | 7 | Macrophages |
| DBI | 0.66 | 0 | 0 | 0.69 | 0.39 | 7 | Macrophages |
| SLC2A3 | 0.65 | 0 | 0 | 0.43 | 0.16 | 7 | Macrophages |
| TNFAIP2 | 0.65 | 0 | 0 | 0.31 | 0.03 | 7 | Macrophages |
| AOAH | 0.65 | 0 | 0 | 0.40 | 0.01 | 7 | Macrophages |
| SELL | 0.65 | 0 | 0 | 0.28 | 0.03 | 7 | Macrophages |
| SLC7A7 | 0.65 | 0 | 0 | 0.39 | 0.03 | 7 | Macrophages |
| IER5 | 0.65 | 0 | 0 | 0.42 | 0.10 | 7 | Macrophages |
| LIMD2 | 0.65 | 0 | 0 | 0.54 | 0.13 | 7 | Macrophages |
| C20orf24 | 0.65 | 0 | 0 | 0.62 | 0.24 | 7 | Macrophages |
| LILRB1 | 0.64 | 0 | 0 | 0.35 | 0.01 | 7 | Macrophages |
| ALDOA | 0.64 | 0 | 0 | 0.91 | 0.69 | 7 | Macrophages |
| FMNL1 | 0.64 | 0 | 0 | 0.40 | 0.02 | 7 | Macrophages |
| TMEM167A | 0.64 | 0 | 0 | 0.57 | 0.22 | 7 | Macrophages |
| ATP6V1F | 0.64 | 0 | 0 | 0.77 | 0.42 | 7 | Macrophages |
| NOP10 | 0.64 | 0 | 0 | 0.80 | 0.46 | 7 | Macrophages |
| PSME2 | 0.64 | 0 | 0 | 0.76 | 0.43 | 7 | Macrophages |
| ADGRE5 | 0.64 | 0 | 0 | 0.46 | 0.12 | 7 | Macrophages |
| NAGK | 0.63 | 0 | 0 | 0.54 | 0.18 | 7 | Macrophages |
| CD36 | 0.63 | 0 | 0 | 0.51 | 0.13 | 7 | Macrophages |
| SNX10 | 0.63 | 0 | 0 | 0.33 | 0.01 | 7 | Macrophages |
| TKT | 0.63 | 0 | 0 | 0.60 | 0.24 | 7 | Macrophages |
| NCF4 | 0.62 | 0 | 0 | 0.36 | 0.01 | 7 | Macrophages |
| RNF130 | 0.62 | 0 | 0 | 0.55 | 0.15 | 7 | Macrophages |
| EVI2A | 0.62 | 0 | 0 | 0.40 | 0.01 | 7 | Macrophages |
| ARHGAP30 | 0.62 | 0 | 0 | 0.44 | 0.02 | 7 | Macrophages |
| GLRX | 0.61 | 0 | 0 | 0.48 | 0.10 | 7 | Macrophages |
| FCGR1B | 0.61 | 0 | 0 | 0.29 | 0.00 | 7 | Macrophages |
| ADAM8 | 0.61 | 0 | 0 | 0.34 | 0.01 | 7 | Macrophages |
| PLP2 | 0.61 | 0 | 0 | 0.50 | 0.21 | 7 | Macrophages |
| LAT2 | 0.61 | 0 | 0 | 0.41 | 0.02 | 7 | Macrophages |
| PLBD1 | 0.6 | 0 | 0 | 0.38 | 0.04 | 7 | Macrophages |
| ICAM3 | 0.6 | 0 | 0 | 0.39 | 0.07 | 7 | Macrophages |
| CCL3L3 | 0.6 | 0 | 0 | 0.27 | 0.01 | 7 | Macrophages |
| FGD2 | 0.6 | 0 | 0 | 0.31 | 0.01 | 7 | Macrophages |
| CLEC4A | 0.59 | 0 | 0 | 0.30 | 0.01 | 7 | Macrophages |
| ARF6 | 0.59 | 0 | 0 | 0.58 | 0.22 | 7 | Macrophages |
| ASGR1 | 0.59 | 0 | 0 | 0.30 | 0.02 | 7 | Macrophages |
| LILRA1 | 0.59 | 0 | 0 | 0.29 | 0.00 | 7 | Macrophages |
| LILRA2 | 0.59 | 0 | 0 | 0.33 | 0.00 | 7 | Macrophages |
| GABARAP | 0.59 | 0 | 0 | 0.78 | 0.38 | 7 | Macrophages |
| LGMN | 0.58 | 0 | 0 | 0.30 | 0.19 | 7 | Macrophages |
| IER3 | 0.58 | 0 | 0 | 0.35 | 0.13 | 7 | Macrophages |
| LPAR6 | 0.58 | 0 | 0 | 0.42 | 0.11 | 7 | Macrophages |
| NINJ1 | 0.58 | 0 | 0 | 0.43 | 0.16 | 7 | Macrophages |
| BAZ1A | 0.58 | 0 | 0 | 0.57 | 0.17 | 7 | Macrophages |
| FBP1 | 0.57 | 0 | 0 | 0.31 | 0.03 | 7 | Macrophages |
| PCBP1 | 0.57 | 0 | 0 | 0.84 | 0.48 | 7 | Macrophages |
| RPL28 | 0.57 | 0 | 0 | 1.00 | 0.98 | 7 | Macrophages |
| ARPC5 | 0.57 | 0 | 0 | 0.87 | 0.63 | 7 | Macrophages |
| PSMB10 | 0.57 | 0 | 0 | 0.62 | 0.23 | 7 | Macrophages |
| RGS18 | 0.57 | 0 | 0 | 0.34 | 0.00 | 7 | Macrophages |
| RBPJ | 0.57 | 0 | 0 | 0.69 | 0.35 | 7 | Macrophages |
| C3AR1 | 0.56 | 0 | 0 | 0.26 | 0.00 | 7 | Macrophages |
| BLVRA | 0.56 | 0 | 0 | 0.49 | 0.15 | 7 | Macrophages |
| ATF5 | 0.56 | 0 | 0 | 0.32 | 0.05 | 7 | Macrophages |
| DNAJB1 | 0.56 | 0 | 0 | 0.52 | 0.20 | 7 | Macrophages |
| LCP2 | 0.55 | 0 | 0 | 0.32 | 0.02 | 7 | Macrophages |
| NUP214 | 0.55 | 0 | 0 | 0.33 | 0.05 | 7 | Macrophages |
| PPP1CA | 0.55 | 0 | 0 | 0.72 | 0.35 | 7 | Macrophages |
| FAM110A | 0.55 | 0 | 0 | 0.28 | 0.03 | 7 | Macrophages |
| VPS29 | 0.55 | 0 | 0 | 0.73 | 0.37 | 7 | Macrophages |
| UBE2D1 | 0.55 | 0 | 0 | 0.49 | 0.16 | 7 | Macrophages |
| MYO1G | 0.55 | 0 | 0 | 0.33 | 0.02 | 7 | Macrophages |
| UPP1 | 0.55 | 0 | 0 | 0.32 | 0.04 | 7 | Macrophages |
| IL10RA | 0.55 | 0 | 0 | 0.34 | 0.01 | 7 | Macrophages |
| GRB2 | 0.55 | 0 | 0 | 0.57 | 0.17 | 7 | Macrophages |
| RBM47 | 0.55 | 0 | 0 | 0.34 | 0.01 | 7 | Macrophages |
| TCIRG1 | 0.54 | 0 | 0 | 0.42 | 0.06 | 7 | Macrophages |
| SNX2 | 0.54 | 0 | 0 | 0.60 | 0.24 | 7 | Macrophages |
| TALDO1 | 0.54 | 0 | 0 | 0.70 | 0.36 | 7 | Macrophages |
| NAAA | 0.54 | 0 | 0 | 0.36 | 0.06 | 7 | Macrophages |
| IRF8 | 0.54 | 0 | 0 | 0.34 | 0.03 | 7 | Macrophages |
| MXD1 | 0.54 | 0 | 0 | 0.31 | 0.02 | 7 | Macrophages |
| YWHAZ | 0.54 | 0 | 0 | 0.74 | 0.42 | 7 | Macrophages |
| NCKAP1L | 0.54 | 0 | 0 | 0.34 | 0.01 | 7 | Macrophages |
| SDF2L1 | 0.54 | 0 | 0 | 0.49 | 0.21 | 7 | Macrophages |
| GADD45B | 0.54 | 0 | 0 | 0.70 | 0.41 | 7 | Macrophages |
| JUNB | 0.53 | 0 | 0 | 0.84 | 0.50 | 7 | Macrophages |
| ALOX5 | 0.53 | 0 | 0 | 0.34 | 0.01 | 7 | Macrophages |
| GNA15 | 0.53 | 0 | 0 | 0.28 | 0.00 | 7 | Macrophages |
| PTAFR | 0.53 | 0 | 0 | 0.30 | 0.01 | 7 | Macrophages |
| MIR4435-2HG | 0.53 | 0 | 0 | 0.32 | 0.06 | 7 | Macrophages |
| ATG3 | 0.53 | 0 | 0 | 0.51 | 0.17 | 7 | Macrophages |
| ATP6V1B2 | 0.52 | 0 | 0 | 0.43 | 0.11 | 7 | Macrophages |
| OSTF1 | 0.52 | 0 | 0 | 0.62 | 0.21 | 7 | Macrophages |
| ADM | 0.52 | 0 | 0 | 0.28 | 0.07 | 7 | Macrophages |
| TPMT | 0.52 | 0 | 0 | 0.37 | 0.06 | 7 | Macrophages |
| C19orf38 | 0.52 | 0 | 0 | 0.29 | 0.01 | 7 | Macrophages |
| RAP1A | 0.52 | 0 | 0 | 0.65 | 0.28 | 7 | Macrophages |
| AC016734.2 | 0.52 | 0 | 0 | 0.46 | 0.12 | 7 | Macrophages |
| KCNK6 | 0.52 | 0 | 0 | 0.36 | 0.05 | 7 | Macrophages |
| PSME2P2 | 0.52 | 0 | 0 | 0.51 | 0.20 | 7 | Macrophages |
| HLA-B | 0.51 | 0 | 0 | 0.95 | 0.52 | 7 | Macrophages |
| MTHFD2 | 0.51 | 0 | 0 | 0.42 | 0.10 | 7 | Macrophages |
| TWF2 | 0.51 | 0 | 0 | 0.50 | 0.14 | 7 | Macrophages |
| PID1 | 0.51 | 0 | 0 | 0.32 | 0.04 | 7 | Macrophages |
| SERF2 | 0.51 | 0 | 0 | 0.99 | 0.94 | 7 | Macrophages |
| RHOA | 0.51 | 0 | 0 | 0.89 | 0.68 | 7 | Macrophages |
| ENTPD1 | 0.51 | 0 | 0 | 0.41 | 0.12 | 7 | Macrophages |
| DAPP1 | 0.5 | 0 | 0 | 0.30 | 0.01 | 7 | Macrophages |
| CNPY3 | 0.5 | 0 | 0 | 0.57 | 0.18 | 7 | Macrophages |
| PAK1 | 0.5 | 0 | 0 | 0.35 | 0.04 | 7 | Macrophages |
| CCL3 | 0.5 | 0 | 0 | 0.26 | 0.02 | 7 | Macrophages |
| LGALS3 | 0.5 | 0 | 0 | 0.65 | 0.35 | 7 | Macrophages |
| LDHA | 0.5 | 0 | 0 | 0.80 | 0.57 | 7 | Macrophages |
| IKZF1 | 0.5 | 0 | 0 | 0.30 | 0.02 | 7 | Macrophages |
| HAVCR2 | 0.5 | 0 | 0 | 0.29 | 0.01 | 7 | Macrophages |
| RAB10 | 0.5 | 0 | 0 | 0.56 | 0.25 | 7 | Macrophages |
| RP4-765C7.2 | 0.5 | 0 | 0 | 0.86 | 0.49 | 7 | Macrophages |
| HLA-DRB6 | 0.49 | 0 | 0 | 0.28 | 0.01 | 7 | Macrophages |
| LGALS9 | 0.49 | 0 | 0 | 0.42 | 0.08 | 7 | Macrophages |
| RASSF4 | 0.49 | 0 | 0 | 0.29 | 0.02 | 7 | Macrophages |
| SDCBP | 0.49 | 0 | 0 | 0.76 | 0.51 | 7 | Macrophages |
| MSRB1 | 0.49 | 0 | 0 | 0.33 | 0.08 | 7 | Macrophages |
| TMSB4X | 0.49 | 0 | 0 | 1.00 | 1.00 | 7 | Macrophages |
| GK | 0.49 | 0 | 0 | 0.29 | 0.02 | 7 | Macrophages |
| ITGAX | 0.49 | 0 | 0 | 0.30 | 0.01 | 7 | Macrophages |
| ADGRE2 | 0.48 | 0 | 0 | 0.28 | 0.00 | 7 | Macrophages |
| TNFAIP8L2 | 0.48 | 0 | 0 | 0.30 | 0.01 | 7 | Macrophages |
| SCO2 | 0.48 | 0 | 0 | 0.36 | 0.07 | 7 | Macrophages |
| ATP2B1 | 0.48 | 0 | 0 | 0.52 | 0.24 | 7 | Macrophages |
| SELPLG | 0.48 | 0 | 0 | 0.30 | 0.02 | 7 | Macrophages |
| BIN2 | 0.48 | 0 | 0 | 0.35 | 0.02 | 7 | Macrophages |
| HSPA8 | 0.48 | 0 | 0 | 0.75 | 0.59 | 7 | Macrophages |
| PGD | 0.48 | 0 | 0 | 0.38 | 0.08 | 7 | Macrophages |
| RUNX3 | 0.48 | 0 | 0 | 0.28 | 0.02 | 7 | Macrophages |
| RASSF5 | 0.48 | 0 | 0 | 0.32 | 0.03 | 7 | Macrophages |
| EIF4A1 | 0.48 | 0 | 0 | 0.62 | 0.27 | 7 | Macrophages |
| MTCO1P12 | 0.48 | 0 | 0 | 0.66 | 0.29 | 7 | Macrophages |
| STX11 | 0.48 | 0 | 0 | 0.28 | 0.02 | 7 | Macrophages |
| YWHAH | 0.48 | 0 | 0 | 0.55 | 0.35 | 7 | Macrophages |
| IQGAP1 | 0.48 | 0 | 0 | 0.54 | 0.21 | 7 | Macrophages |
| CD33 | 0.48 | 0 | 0 | 0.25 | 0.00 | 7 | Macrophages |
| HSP90AA1 | 0.48 | 0 | 0 | 0.94 | 0.86 | 7 | Macrophages |
| IGFLR1 | 0.47 | 0 | 0 | 0.30 | 0.01 | 7 | Macrophages |
| H2AFY | 0.47 | 0 | 0 | 0.71 | 0.41 | 7 | Macrophages |
| 1-Mar | 0.47 | 0 | 0 | 0.29 | 0.01 | 7 | Macrophages |
| AHR | 0.47 | 0 | 0 | 0.42 | 0.11 | 7 | Macrophages |
| NABP1 | 0.47 | 0 | 0 | 0.39 | 0.09 | 7 | Macrophages |
| GDI2 | 0.47 | 0 | 0 | 0.69 | 0.44 | 7 | Macrophages |
| CIB1 | 0.47 | 0 | 0 | 0.66 | 0.32 | 7 | Macrophages |
| GNG5 | 0.47 | 0 | 0 | 0.80 | 0.51 | 7 | Macrophages |
| SERP1 | 0.47 | 0 | 0 | 0.76 | 0.44 | 7 | Macrophages |
| DBNL | 0.47 | 0 | 0 | 0.60 | 0.27 | 7 | Macrophages |
| BLOC1S1 | 0.46 | 0 | 0 | 0.63 | 0.33 | 7 | Macrophages |
| RAP2B | 0.46 | 0 | 0 | 0.38 | 0.10 | 7 | Macrophages |
| SLA | 0.46 | 0 | 0 | 0.29 | 0.02 | 7 | Macrophages |
| MT-CO2 | 0.46 | 0 | 0 | 0.99 | 0.99 | 7 | Macrophages |
| CECR1 | 0.46 | 0 | 0 | 0.33 | 0.05 | 7 | Macrophages |
| SLC43A2 | 0.46 | 0 | 0 | 0.35 | 0.07 | 7 | Macrophages |
| RP11-186B7.4 | 0.46 | 0 | 0 | 0.62 | 0.28 | 7 | Macrophages |
| TNFAIP8 | 0.46 | 0 | 0 | 0.38 | 0.08 | 7 | Macrophages |
| SLC31A2 | 0.46 | 0 | 0 | 0.31 | 0.02 | 7 | Macrophages |
| HN1 | 0.46 | 0 | 0 | 0.53 | 0.17 | 7 | Macrophages |
| BEST1 | 0.46 | 0 | 0 | 0.36 | 0.09 | 7 | Macrophages |
| CREG1 | 0.45 | 0 | 0 | 0.42 | 0.17 | 7 | Macrophages |
| FAM49A | 0.45 | 0 | 0 | 0.32 | 0.05 | 7 | Macrophages |
| BAX | 0.45 | 0 | 0 | 0.56 | 0.22 | 7 | Macrophages |
| SCIMP | 0.45 | 0 | 0 | 0.27 | 0.01 | 7 | Macrophages |
| CMTM6 | 0.45 | 0 | 0 | 0.61 | 0.31 | 7 | Macrophages |
| KLF6 | 0.45 | 0 | 0 | 0.68 | 0.41 | 7 | Macrophages |
| C9orf72 | 0.45 | 0 | 0 | 0.27 | 0.03 | 7 | Macrophages |
| SQRDL | 0.45 | 0 | 0 | 0.36 | 0.06 | 7 | Macrophages |
| FAM65B | 0.45 | 0 | 0 | 0.27 | 0.03 | 7 | Macrophages |
| MCOLN1 | 0.44 | 0 | 0 | 0.35 | 0.08 | 7 | Macrophages |
| C14orf2 | 0.44 | 0 | 0 | 0.87 | 0.67 | 7 | Macrophages |
| SYNGR2 | 0.44 | 0 | 0 | 0.60 | 0.25 | 7 | Macrophages |
| SNHG5 | 0.44 | 0 | 0 | 0.80 | 0.56 | 7 | Macrophages |
| RBM3 | 0.44 | 0 | 0 | 0.80 | 0.54 | 7 | Macrophages |
| ADAP2 | 0.44 | 0 | 0 | 0.31 | 0.05 | 7 | Macrophages |
| OGFRL1 | 0.44 | 0 | 0 | 0.27 | 0.03 | 7 | Macrophages |
| SUB1 | 0.44 | 0 | 0 | 0.87 | 0.69 | 7 | Macrophages |
| LAMTOR2 | 0.43 | 0 | 0 | 0.57 | 0.26 | 7 | Macrophages |
| CTSA | 0.43 | 0 | 0 | 0.47 | 0.15 | 7 | Macrophages |
| MGST1 | 0.43 | 0 | 0 | 0.26 | 0.02 | 7 | Macrophages |
| WARS | 0.43 | 0 | 0 | 0.48 | 0.19 | 7 | Macrophages |
| RAB31 | 0.43 | 0 | 0 | 0.42 | 0.13 | 7 | Macrophages |
| CARD19 | 0.43 | 0 | 0 | 0.43 | 0.14 | 7 | Macrophages |
| UBA52 | 0.42 | 0 | 0 | 0.99 | 0.95 | 7 | Macrophages |
| DAZAP2 | 0.42 | 0 | 0 | 0.73 | 0.40 | 7 | Macrophages |
| SFT2D1 | 0.42 | 0 | 0 | 0.58 | 0.25 | 7 | Macrophages |
| CCDC109B | 0.42 | 0 | 0 | 0.30 | 0.04 | 7 | Macrophages |
| CSK | 0.42 | 0 | 0 | 0.36 | 0.07 | 7 | Macrophages |
| B2M | 0.42 | 0 | 0 | 1.00 | 0.99 | 7 | Macrophages |
| ODF3B | 0.42 | 0 | 0 | 0.29 | 0.03 | 7 | Macrophages |
| DAB2 | 0.42 | 0 | 0 | 0.26 | 0.11 | 7 | Macrophages |
| GRINA | 0.42 | 0 | 0 | 0.47 | 0.20 | 7 | Macrophages |
| UNC93B1 | 0.42 | 0 | 0 | 0.31 | 0.04 | 7 | Macrophages |
| SIGLEC9 | 0.42 | 0 | 0 | 0.27 | 0.00 | 7 | Macrophages |
| FTLP2 | 0.42 | 0 | 0 | 0.27 | 0.02 | 7 | Macrophages |
| TRABD | 0.42 | 0 | 0 | 0.39 | 0.12 | 7 | Macrophages |
| LAMTOR4 | 0.42 | 0 | 0 | 0.77 | 0.47 | 7 | Macrophages |
| CXorf21 | 0.42 | 0 | 0 | 0.26 | 0.00 | 7 | Macrophages |
| COPE | 0.42 | 0 | 0 | 0.78 | 0.49 | 7 | Macrophages |
| MVP | 0.41 | 0 | 0 | 0.41 | 0.13 | 7 | Macrophages |
| TPI1 | 0.41 | 0 | 0 | 0.89 | 0.76 | 7 | Macrophages |
| CCND3 | 0.41 | 0 | 0 | 0.40 | 0.14 | 7 | Macrophages |
| BTK | 0.41 | 0 | 0 | 0.26 | 0.01 | 7 | Macrophages |
| PFDN5 | 0.41 | 0 | 0 | 0.97 | 0.90 | 7 | Macrophages |
| IFNGR2 | 0.41 | 0 | 0 | 0.42 | 0.14 | 7 | Macrophages |
| NFKBIZ | 0.41 | 0 | 0 | 0.28 | 0.07 | 7 | Macrophages |
| DSE | 0.41 | 0 | 0 | 0.31 | 0.06 | 7 | Macrophages |
| IFNAR2 | 0.41 | 0 | 0 | 0.38 | 0.08 | 7 | Macrophages |
| PPP1R18 | 0.4 | 0 | 0 | 0.44 | 0.14 | 7 | Macrophages |
| RP11-680H20.1 | 0.4 | 0 | 0 | 0.40 | 0.09 | 7 | Macrophages |
| CORO1B | 0.4 | 0 | 0 | 0.47 | 0.19 | 7 | Macrophages |
| ACER3 | 0.4 | 0 | 0 | 0.32 | 0.05 | 7 | Macrophages |
| SYK | 0.4 | 0 | 0 | 0.28 | 0.01 | 7 | Macrophages |
| SHKBP1 | 0.4 | 0 | 0 | 0.39 | 0.10 | 7 | Macrophages |
| CARD16 | 0.4 | 0 | 0 | 0.62 | 0.27 | 7 | Macrophages |
| GPR65 | 0.4 | 0 | 0 | 0.26 | 0.01 | 7 | Macrophages |
| HNMT | 0.4 | 0 | 0 | 0.35 | 0.09 | 7 | Macrophages |
| SPCS3 | 0.39 | 0 | 0 | 0.49 | 0.16 | 7 | Macrophages |
| GLUL | 0.39 | 0 | 0 | 0.68 | 0.41 | 7 | Macrophages |
| FCGRT | 0.39 | 0 | 0 | 0.81 | 0.55 | 7 | Macrophages |
| EZR | 0.39 | 0 | 0 | 0.50 | 0.24 | 7 | Macrophages |
| PPP4C | 0.39 | 0 | 0 | 0.63 | 0.34 | 7 | Macrophages |
| PTP4A2 | 0.39 | 0 | 0 | 0.58 | 0.27 | 7 | Macrophages |
| VPS35 | 0.38 | 0 | 0 | 0.52 | 0.22 | 7 | Macrophages |
| LYST | 0.38 | 0 | 0 | 0.28 | 0.06 | 7 | Macrophages |
| MOB3A | 0.38 | 0 | 0 | 0.30 | 0.07 | 7 | Macrophages |
| SASH3 | 0.38 | 0 | 0 | 0.25 | 0.01 | 7 | Macrophages |
| ERP44 | 0.38 | 0 | 0 | 0.52 | 0.24 | 7 | Macrophages |
| GNAI2 | 0.37 | 0 | 0 | 0.70 | 0.33 | 7 | Macrophages |
| TMSB4XP4 | 0.37 | 0 | 0 | 0.63 | 0.34 | 7 | Macrophages |
| PLIN3 | 0.37 | 0 | 0 | 0.46 | 0.18 | 7 | Macrophages |
| SPINT2 | 0.37 | 0 | 0 | 0.41 | 0.13 | 7 | Macrophages |
| TMEM176A | 0.36 | 0 | 0 | 0.29 | 0.12 | 7 | Macrophages |
| SIRPA | 0.36 | 0 | 0 | 0.30 | 0.04 | 7 | Macrophages |
| FUOM | 0.36 | 0 | 0 | 0.32 | 0.07 | 7 | Macrophages |
| G6PD | 0.36 | 0 | 0 | 0.30 | 0.06 | 7 | Macrophages |
| FAM105A | 0.36 | 0 | 0 | 0.29 | 0.04 | 7 | Macrophages |
| MKNK1 | 0.36 | 0 | 0 | 0.33 | 0.09 | 7 | Macrophages |
| LPCAT2 | 0.36 | 0 | 0 | 0.26 | 0.02 | 7 | Macrophages |
| PGAM1 | 0.35 | 0 | 0 | 0.69 | 0.51 | 7 | Macrophages |
| RPL22L1 | 0.35 | 0 | 0 | 0.46 | 0.21 | 7 | Macrophages |
| MPP1 | 0.35 | 0 | 0 | 0.30 | 0.06 | 7 | Macrophages |
| MYD88 | 0.35 | 0 | 0 | 0.29 | 0.06 | 7 | Macrophages |
| RB1 | 0.35 | 0 | 0 | 0.35 | 0.12 | 7 | Macrophages |
| RNF181 | 0.35 | 0 | 0 | 0.66 | 0.38 | 7 | Macrophages |
| STK10 | 0.35 | 0 | 0 | 0.29 | 0.05 | 7 | Macrophages |
| RIN3 | 0.35 | 0 | 0 | 0.26 | 0.05 | 7 | Macrophages |
| CHCHD7 | 0.35 | 0 | 0 | 0.36 | 0.13 | 7 | Macrophages |
| RILPL2 | 0.35 | 0 | 0 | 0.40 | 0.14 | 7 | Macrophages |
| ATP5EP2 | 0.34 | 0 | 0 | 0.32 | 0.10 | 7 | Macrophages |
| PAK2 | 0.34 | 0 | 0 | 0.48 | 0.18 | 7 | Macrophages |
| SELT | 0.34 | 0 | 0 | 0.63 | 0.35 | 7 | Macrophages |
| RAP1B | 0.34 | 0 | 0 | 0.56 | 0.32 | 7 | Macrophages |
| IFNGR1 | 0.34 | 0 | 0 | 0.54 | 0.26 | 7 | Macrophages |
| CD58 | 0.34 | 0 | 0 | 0.33 | 0.07 | 7 | Macrophages |
| GIMAP4 | 0.34 | 0 | 0 | 0.47 | 0.17 | 7 | Macrophages |
| HEXB | 0.34 | 0 | 0 | 0.42 | 0.18 | 7 | Macrophages |
| TGFB1 | 0.34 | 0 | 0 | 0.42 | 0.14 | 7 | Macrophages |
| BACH1 | 0.34 | 0 | 0 | 0.26 | 0.06 | 7 | Macrophages |
| IRF7 | 0.34 | 0 | 0 | 0.28 | 0.06 | 7 | Macrophages |
| ATP6V0E1 | 0.34 | 0 | 0 | 0.80 | 0.59 | 7 | Macrophages |
| RNH1 | 0.33 | 0 | 0 | 0.71 | 0.49 | 7 | Macrophages |
| CMTM7 | 0.33 | 0 | 0 | 0.33 | 0.07 | 7 | Macrophages |
| NFKBIA | 0.33 | 0 | 0 | 0.56 | 0.39 | 7 | Macrophages |
| RPL39 | 0.33 | 0 | 0 | 1.00 | 0.99 | 7 | Macrophages |
| GBP2 | 0.33 | 0 | 0 | 0.28 | 0.10 | 7 | Macrophages |
| DRAM2 | 0.33 | 0 | 0 | 0.39 | 0.13 | 7 | Macrophages |
| NANS | 0.33 | 0 | 0 | 0.42 | 0.16 | 7 | Macrophages |
| IER2 | 0.33 | 0 | 0 | 0.71 | 0.45 | 7 | Macrophages |
| MYL12A | 0.33 | 0 | 0 | 0.91 | 0.74 | 7 | Macrophages |
| AC016739.2 | 0.33 | 0 | 0 | 0.97 | 0.91 | 7 | Macrophages |
| BLOC1S2 | 0.33 | 0 | 0 | 0.34 | 0.11 | 7 | Macrophages |
| CREM | 0.32 | 0 | 0 | 0.33 | 0.19 | 7 | Macrophages |
| NAGA | 0.32 | 0 | 0 | 0.30 | 0.07 | 7 | Macrophages |
| ISG15 | 0.32 | 0 | 0 | 0.42 | 0.25 | 7 | Macrophages |
| ARHGDIA | 0.32 | 0 | 0 | 0.66 | 0.42 | 7 | Macrophages |
| CCDC88A | 0.32 | 0 | 0 | 0.38 | 0.13 | 7 | Macrophages |
| CNDP2 | 0.32 | 0 | 0 | 0.35 | 0.12 | 7 | Macrophages |
| COMMD8 | 0.32 | 0 | 0 | 0.36 | 0.11 | 7 | Macrophages |
| RAB32 | 0.32 | 0 | 0 | 0.35 | 0.12 | 7 | Macrophages |
| CDKN1A | 0.32 | 0 | 0 | 0.28 | 0.07 | 7 | Macrophages |
| HIGD2A | 0.32 | 0 | 0 | 0.62 | 0.35 | 7 | Macrophages |
| SMCO4 | 0.32 | 0 | 0 | 0.32 | 0.11 | 7 | Macrophages |
| MYO9B | 0.32 | 0 | 0 | 0.28 | 0.07 | 7 | Macrophages |
| MAP2K3 | 0.32 | 0 | 0 | 0.33 | 0.09 | 7 | Macrophages |
| POLD4 | 0.32 | 0 | 0 | 0.33 | 0.11 | 7 | Macrophages |
| IL2RG | 0.32 | 0 | 0 | 0.30 | 0.04 | 7 | Macrophages |
| SSR3 | 0.32 | 0 | 0 | 0.49 | 0.24 | 7 | Macrophages |
| LACTB | 0.32 | 0 | 0 | 0.28 | 0.05 | 7 | Macrophages |
| CAMK1 | 0.31 | 0 | 0 | 0.36 | 0.14 | 7 | Macrophages |
| UBE2D3 | 0.31 | 0 | 0 | 0.79 | 0.59 | 7 | Macrophages |
| PSMB3 | 0.31 | 0 | 0 | 0.80 | 0.52 | 7 | Macrophages |
| M6PR | 0.31 | 0 | 0 | 0.47 | 0.21 | 7 | Macrophages |
| TMSB10 | 0.31 | 0 | 0 | 1.00 | 0.99 | 7 | Macrophages |
| ADPGK | 0.31 | 0 | 0 | 0.36 | 0.12 | 7 | Macrophages |
| PICALM | 0.31 | 0 | 0 | 0.39 | 0.16 | 7 | Macrophages |
| VMA21 | 0.31 | 0 | 0 | 0.32 | 0.08 | 7 | Macrophages |
| LTA4H | 0.31 | 0 | 0 | 0.40 | 0.20 | 7 | Macrophages |
| RXRA | 0.31 | 0 | 0 | 0.26 | 0.07 | 7 | Macrophages |
| HSPH1 | 0.31 | 0 | 0 | 0.31 | 0.10 | 7 | Macrophages |
| IRF1 | 0.31 | 0 | 0 | 0.40 | 0.18 | 7 | Macrophages |
| EIF1 | 0.31 | 0 | 0 | 0.98 | 0.96 | 7 | Macrophages |
| BHLHE40 | 0.31 | 0 | 0 | 0.28 | 0.08 | 7 | Macrophages |
| FTH1P11 | 0.31 | 0 | 0 | 0.27 | 0.08 | 7 | Macrophages |
| PNPLA6 | 0.3 | 0 | 0 | 0.26 | 0.05 | 7 | Macrophages |
| ARHGAP4 | 0.3 | 0 | 0 | 0.25 | 0.04 | 7 | Macrophages |
| SH3BP2 | 0.3 | 0 | 0 | 0.29 | 0.08 | 7 | Macrophages |
| GPX4 | 0.3 | 0 | 0 | 0.75 | 0.56 | 7 | Macrophages |
| YIF1B | 0.3 | 0 | 0 | 0.33 | 0.09 | 7 | Macrophages |
| AMD1 | 0.3 | 0 | 0 | 0.38 | 0.17 | 7 | Macrophages |
| CASP4 | 0.3 | 0 | 0 | 0.38 | 0.15 | 7 | Macrophages |
| COX5A | 0.3 | 0 | 0 | 0.69 | 0.39 | 7 | Macrophages |
| FAM45A | 0.3 | 0 | 0 | 0.32 | 0.11 | 7 | Macrophages |
| SGK1 | 0.3 | 0 | 0 | 0.39 | 0.15 | 7 | Macrophages |
| CDK2AP2 | 0.29 | 0 | 0 | 0.37 | 0.14 | 7 | Macrophages |
| CD47 | 0.29 | 0 | 0 | 0.46 | 0.20 | 7 | Macrophages |
| PLEKHO2 | 0.29 | 0 | 0 | 0.26 | 0.06 | 7 | Macrophages |
| ARL6IP5 | 0.29 | 0 | 0 | 0.65 | 0.41 | 7 | Macrophages |
| TPT1 | 0.29 | 0 | 0 | 1.00 | 0.98 | 7 | Macrophages |
| RNF13 | 0.29 | 0 | 0 | 0.45 | 0.20 | 7 | Macrophages |
| HSPA5 | 0.29 | 0 | 0 | 0.66 | 0.49 | 7 | Macrophages |
| CHCHD10 | 0.29 | 0 | 0 | 0.41 | 0.19 | 7 | Macrophages |
| HMGA1 | 0.29 | 0 | 0 | 0.29 | 0.09 | 7 | Macrophages |
| ATP6AP1 | 0.28 | 0 | 0 | 0.50 | 0.24 | 7 | Macrophages |
| FAM46A | 0.28 | 0 | 0 | 0.29 | 0.13 | 7 | Macrophages |
| CALR | 0.28 | 0 | 0.01 | 0.70 | 0.58 | 7 | Macrophages |
| ACOT9 | 0.28 | 0 | 0 | 0.33 | 0.11 | 7 | Macrophages |
| CD4 | 0.28 | 0 | 0 | 0.30 | 0.10 | 7 | Macrophages |
| ADAM10 | 0.28 | 0 | 0 | 0.28 | 0.11 | 7 | Macrophages |
| CCNL1 | 0.27 | 0 | 0 | 0.54 | 0.31 | 7 | Macrophages |
| CTD-2666L21.3 | 0.27 | 0 | 0 | 0.52 | 0.24 | 7 | Macrophages |
| TRA2B | 0.27 | 0 | 0 | 0.67 | 0.50 | 7 | Macrophages |
| MTMR14 | 0.27 | 0 | 0 | 0.29 | 0.07 | 7 | Macrophages |
| RPLP1 | 0.27 | 0 | 0 | 1.00 | 1.00 | 7 | Macrophages |
| AGTRAP | 0.27 | 0 | 0 | 0.28 | 0.12 | 7 | Macrophages |
| SRA1 | 0.27 | 0 | 0 | 0.38 | 0.16 | 7 | Macrophages |
| LSM10 | 0.27 | 0 | 0 | 0.37 | 0.16 | 7 | Macrophages |
| MGAT1 | 0.27 | 0 | 0 | 0.45 | 0.24 | 7 | Macrophages |
| ATP1B3 | 0.27 | 0 | 0 | 0.52 | 0.30 | 7 | Macrophages |
| JUND | 0.27 | 0 | 0 | 0.53 | 0.29 | 7 | Macrophages |
| YWHAB | 0.27 | 0 | 0 | 0.76 | 0.58 | 7 | Macrophages |
| ENY2 | 0.27 | 0 | 0 | 0.70 | 0.44 | 7 | Macrophages |
| CTC-479C5.12 | 0.26 | 0 | 0 | 0.31 | 0.10 | 7 | Macrophages |
| RPS19 | 0.26 | 0 | 0 | 1.00 | 0.99 | 7 | Macrophages |
| RP3-417G15.1 | 0.26 | 0 | 0 | 0.90 | 0.76 | 7 | Macrophages |
| RPS14P8 | 0.26 | 0 | 0 | 0.46 | 0.19 | 7 | Macrophages |
| DRAP1 | 0.26 | 0 | 0 | 0.70 | 0.49 | 7 | Macrophages |
| MT-CO3 | 0.26 | 0 | 0 | 0.99 | 0.99 | 7 | Macrophages |
| RPL36A | 0.26 | 0 | 0 | 0.97 | 0.91 | 7 | Macrophages |
| ZFAND5 | 0.26 | 0 | 0 | 0.50 | 0.26 | 7 | Macrophages |
| SLC38A2 | 0.26 | 0 | 0 | 0.44 | 0.24 | 7 | Macrophages |
| ANP32A | 0.26 | 0 | 0 | 0.53 | 0.29 | 7 | Macrophages |
| MYDGF | 0.26 | 0 | 0 | 0.63 | 0.42 | 7 | Macrophages |
| C17orf62 | 0.26 | 0 | 0 | 0.33 | 0.14 | 7 | Macrophages |
| NUP62 | 0.26 | 0 | 0 | 0.31 | 0.11 | 7 | Macrophages |
| TGFBI | 0.26 | 0 | 0 | 0.59 | 0.34 | 7 | Macrophages |
| CDC42SE2 | 0.26 | 0 | 0 | 0.27 | 0.08 | 7 | Macrophages |
| COX6B1 | 0.25 | 0 | 0 | 0.85 | 0.71 | 7 | Macrophages |
| LSM6 | 0.25 | 0 | 0 | 0.47 | 0.24 | 7 | Macrophages |
| CISD2 | 0.25 | 0 | 0 | 0.32 | 0.14 | 7 | Macrophages |
| LAP3 | 0.25 | 0 | 0 | 0.43 | 0.21 | 7 | Macrophages |
| RP11-466H18.1 | 0.25 | 0 | 0 | 0.91 | 0.76 | 7 | Macrophages |
| PLEKHB2 | 0.25 | 0 | 0 | 0.30 | 0.12 | 7 | Macrophages |
| WIPF1 | 0.25 | 0 | 0 | 0.33 | 0.11 | 7 | Macrophages |
| MAPKAPK3 | 0.25 | 0 | 0 | 0.31 | 0.09 | 7 | Macrophages |
| SCPEP1 | 0.25 | 0 | 0 | 0.28 | 0.12 | 7 | Macrophages |
| PSMB9 | 0.25 | 0 | 0 | 0.47 | 0.20 | 7 | Macrophages |
| UBC | 0.25 | 0 | 0 | 0.95 | 0.83 | 7 | Macrophages |
| HPGD | 2.76 | 0 | 0 | 0.98 | 0.18 | 8 | Endothelial Cells |
| S100A3 | 2.44 | 0 | 0 | 0.87 | 0.07 | 8 | Endothelial Cells |
| HLA-E | 2.25 | 0 | 0 | 1.00 | 0.58 | 8 | Endothelial Cells |
| ITM2A | 2.19 | 0 | 0 | 0.93 | 0.15 | 8 | Endothelial Cells |
| EMCN | 2.19 | 0 | 0 | 0.78 | 0.13 | 8 | Endothelial Cells |
| CLDN5 | 2.13 | 0 | 0 | 0.96 | 0.23 | 8 | Endothelial Cells |
| SOSTDC1 | 2.03 | 0 | 0 | 0.61 | 0.11 | 8 | Endothelial Cells |
| IFI27 | 2.02 | 0 | 0 | 1.00 | 0.33 | 8 | Endothelial Cells |
| IL32 | 2.01 | 0 | 0 | 0.95 | 0.25 | 8 | Endothelial Cells |
| TSPAN12 | 1.97 | 0 | 0 | 0.80 | 0.12 | 8 | Endothelial Cells |
| RAMP2 | 1.91 | 0 | 0 | 0.99 | 0.30 | 8 | Endothelial Cells |
| EDNRB | 1.8 | 0 | 0 | 0.88 | 0.23 | 8 | Endothelial Cells |
| HLA-C | 1.71 | 0 | 0 | 0.99 | 0.52 | 8 | Endothelial Cells |
| ICAM2 | 1.69 | 0 | 0 | 0.87 | 0.19 | 8 | Endothelial Cells |
| ITM2B | 1.68 | 0 | 0 | 1.00 | 0.88 | 8 | Endothelial Cells |
| HLA-A | 1.67 | 0 | 0 | 1.00 | 0.68 | 8 | Endothelial Cells |
| HLA-B | 1.67 | 0 | 0 | 0.99 | 0.51 | 8 | Endothelial Cells |
| EGFL7 | 1.64 | 0 | 0 | 0.93 | 0.22 | 8 | Endothelial Cells |
| APP | 1.6 | 0 | 0 | 0.93 | 0.49 | 8 | Endothelial Cells |
| ACE | 1.6 | 0 | 0 | 0.78 | 0.10 | 8 | Endothelial Cells |
| S100A4 | 1.59 | 0 | 0 | 0.94 | 0.36 | 8 | Endothelial Cells |
| PECAM1 | 1.59 | 0 | 0 | 0.93 | 0.22 | 8 | Endothelial Cells |
| TNFSF10 | 1.58 | 0 | 0 | 0.82 | 0.19 | 8 | Endothelial Cells |
| B2M | 1.57 | 0 | 0 | 1.00 | 0.99 | 8 | Endothelial Cells |
| S100A6 | 1.56 | 0 | 0 | 0.99 | 0.83 | 8 | Endothelial Cells |
| SH3BP5 | 1.53 | 0 | 0 | 0.81 | 0.18 | 8 | Endothelial Cells |
| GNG11 | 1.52 | 0 | 0 | 0.97 | 0.41 | 8 | Endothelial Cells |
| BST2 | 1.52 | 0 | 0 | 0.94 | 0.26 | 8 | Endothelial Cells |
| SDPR | 1.52 | 0 | 0 | 0.79 | 0.22 | 8 | Endothelial Cells |
| APLN | 1.48 | 0 | 0 | 0.40 | 0.02 | 8 | Endothelial Cells |
| CARD16 | 1.47 | 0 | 0 | 0.80 | 0.26 | 8 | Endothelial Cells |
| CDH5 | 1.47 | 0 | 0 | 0.77 | 0.15 | 8 | Endothelial Cells |
| ESAM | 1.41 | 0 | 0 | 0.89 | 0.32 | 8 | Endothelial Cells |
| SGK1 | 1.4 | 0 | 0 | 0.54 | 0.14 | 8 | Endothelial Cells |
| TM4SF1 | 1.36 | 0 | 0 | 0.87 | 0.37 | 8 | Endothelial Cells |
| ECSCR.1 | 1.36 | 0 | 0 | 0.77 | 0.16 | 8 | Endothelial Cells |
| FRY | 1.35 | 0 | 0 | 0.61 | 0.12 | 8 | Endothelial Cells |
| HSPC324 | 1.33 | 0 | 0 | 0.45 | 0.04 | 8 | Endothelial Cells |
| SLC9A3R2 | 1.31 | 0 | 0 | 0.82 | 0.38 | 8 | Endothelial Cells |
| C10orf10 | 1.3 | 0 | 0 | 0.54 | 0.14 | 8 | Endothelial Cells |
| CAV2 | 1.29 | 0 | 0 | 0.93 | 0.56 | 8 | Endothelial Cells |
| CYB5A | 1.29 | 0 | 0 | 0.82 | 0.33 | 8 | Endothelial Cells |
| SERPINB6 | 1.29 | 0 | 0 | 0.82 | 0.40 | 8 | Endothelial Cells |
| ACVRL1 | 1.28 | 0 | 0 | 0.69 | 0.13 | 8 | Endothelial Cells |
| AQP1 | 1.28 | 0 | 0 | 0.73 | 0.18 | 8 | Endothelial Cells |
| FENDRR | 1.28 | 0 | 0 | 0.62 | 0.13 | 8 | Endothelial Cells |
| PRX | 1.27 | 0 | 0 | 0.54 | 0.06 | 8 | Endothelial Cells |
| MYZAP | 1.26 | 0 | 0 | 0.54 | 0.08 | 8 | Endothelial Cells |
| EMP2 | 1.26 | 0 | 0 | 0.90 | 0.47 | 8 | Endothelial Cells |
| RAMP3 | 1.24 | 0 | 0 | 0.69 | 0.13 | 8 | Endothelial Cells |
| MYO6 | 1.24 | 0 | 0 | 0.54 | 0.12 | 8 | Endothelial Cells |
| CARHSP1 | 1.23 | 0 | 0 | 0.85 | 0.34 | 8 | Endothelial Cells |
| EPAS1 | 1.21 | 0 | 0 | 0.88 | 0.41 | 8 | Endothelial Cells |
| BCAM | 1.21 | 0 | 0 | 0.79 | 0.31 | 8 | Endothelial Cells |
| HES1 | 1.18 | 0 | 0 | 0.60 | 0.38 | 8 | Endothelial Cells |
| TSPAN15 | 1.16 | 0 | 0 | 0.54 | 0.11 | 8 | Endothelial Cells |
| ROBO4 | 1.15 | 0 | 0 | 0.56 | 0.09 | 8 | Endothelial Cells |
| LY6E | 1.15 | 0 | 0 | 0.78 | 0.44 | 8 | Endothelial Cells |
| IFI6 | 1.11 | 0 | 0 | 0.61 | 0.24 | 8 | Endothelial Cells |
| RAB11A | 1.11 | 0 | 0 | 0.80 | 0.43 | 8 | Endothelial Cells |
| TNFRSF4 | 1.09 | 0 | 0 | 0.41 | 0.06 | 8 | Endothelial Cells |
| GPR146 | 1.09 | 0 | 0 | 0.44 | 0.07 | 8 | Endothelial Cells |
| PCDH12 | 1.08 | 0 | 0 | 0.46 | 0.07 | 8 | Endothelial Cells |
| PCDH17 | 1.08 | 0 | 0 | 0.58 | 0.13 | 8 | Endothelial Cells |
| FOXF1 | 1.07 | 0 | 0 | 0.65 | 0.24 | 8 | Endothelial Cells |
| ISG15 | 1.07 | 0 | 0 | 0.54 | 0.25 | 8 | Endothelial Cells |
| VAT1 | 1.06 | 0 | 0 | 0.54 | 0.15 | 8 | Endothelial Cells |
| PCAT19 | 1.05 | 0 | 0 | 0.68 | 0.16 | 8 | Endothelial Cells |
| ARHGAP29 | 1.05 | 0 | 0 | 0.73 | 0.31 | 8 | Endothelial Cells |
| ADGRL2 | 1.04 | 0 | 0 | 0.55 | 0.13 | 8 | Endothelial Cells |
| S100A13 | 1.04 | 0 | 0 | 0.73 | 0.38 | 8 | Endothelial Cells |
| SLCO2A1 | 1.04 | 0 | 0 | 0.47 | 0.09 | 8 | Endothelial Cells |
| UBE2L6 | 1.03 | 0 | 0 | 0.58 | 0.23 | 8 | Endothelial Cells |
| RCSD1 | 1.03 | 0 | 0 | 0.53 | 0.17 | 8 | Endothelial Cells |
| CCDC85B | 1.03 | 0 | 0 | 0.75 | 0.39 | 8 | Endothelial Cells |
| SPTBN1 | 1.03 | 0 | 0 | 0.81 | 0.40 | 8 | Endothelial Cells |
| VIPR1 | 1.02 | 0 | 0 | 0.40 | 0.03 | 8 | Endothelial Cells |
| KIAA1462 | 1.01 | 0 | 0 | 0.50 | 0.11 | 8 | Endothelial Cells |
| VAMP5 | 1.01 | 0 | 0 | 0.87 | 0.50 | 8 | Endothelial Cells |
| TMEM123 | 1.01 | 0 | 0 | 0.71 | 0.36 | 8 | Endothelial Cells |
| C8orf4 | 1.01 | 0 | 0 | 0.81 | 0.41 | 8 | Endothelial Cells |
| RAPGEF5 | 1 | 0 | 0 | 0.45 | 0.11 | 8 | Endothelial Cells |
| HSPA12B | 1 | 0 | 0 | 0.45 | 0.07 | 8 | Endothelial Cells |
| DUSP6 | 0.99 | 0 | 0 | 0.53 | 0.17 | 8 | Endothelial Cells |
| STXBP6 | 0.99 | 0 | 0 | 0.49 | 0.13 | 8 | Endothelial Cells |
| PSME2 | 0.98 | 0 | 0 | 0.74 | 0.44 | 8 | Endothelial Cells |
| IGFBP4 | 0.98 | 0 | 0 | 0.94 | 0.48 | 8 | Endothelial Cells |
| APOL3 | 0.98 | 0 | 0 | 0.45 | 0.10 | 8 | Endothelial Cells |
| TMEM204 | 0.97 | 0 | 0 | 0.52 | 0.17 | 8 | Endothelial Cells |
| NPDC1 | 0.96 | 0 | 0 | 0.67 | 0.29 | 8 | Endothelial Cells |
| TSPAN4 | 0.96 | 0 | 0 | 0.64 | 0.29 | 8 | Endothelial Cells |
| FKBP1A | 0.95 | 0 | 0 | 0.94 | 0.71 | 8 | Endothelial Cells |
| TBX3 | 0.95 | 0 | 0 | 0.61 | 0.28 | 8 | Endothelial Cells |
| IRF1 | 0.95 | 0 | 0 | 0.43 | 0.18 | 8 | Endothelial Cells |
| PSMB9 | 0.95 | 0 | 0 | 0.57 | 0.20 | 8 | Endothelial Cells |
| PDLIM1 | 0.94 | 0 | 0 | 0.65 | 0.38 | 8 | Endothelial Cells |
| TIE1 | 0.94 | 0 | 0 | 0.55 | 0.12 | 8 | Endothelial Cells |
| KDR | 0.93 | 0 | 0 | 0.40 | 0.07 | 8 | Endothelial Cells |
| LEPROTL1 | 0.93 | 0 | 0 | 0.60 | 0.23 | 8 | Endothelial Cells |
| CDKAL1 | 0.92 | 0 | 0 | 0.36 | 0.06 | 8 | Endothelial Cells |
| PIM3 | 0.92 | 0 | 0 | 0.47 | 0.16 | 8 | Endothelial Cells |
| GBP4 | 0.92 | 0 | 0 | 0.33 | 0.09 | 8 | Endothelial Cells |
| RALA | 0.92 | 0 | 0 | 0.60 | 0.27 | 8 | Endothelial Cells |
| FXYD5 | 0.92 | 0 | 0 | 0.59 | 0.25 | 8 | Endothelial Cells |
| ANXA3 | 0.89 | 0 | 0 | 0.48 | 0.10 | 8 | Endothelial Cells |
| OPTN | 0.87 | 0 | 0 | 0.46 | 0.18 | 8 | Endothelial Cells |
| GUK1 | 0.87 | 0 | 0 | 0.88 | 0.66 | 8 | Endothelial Cells |
| GMFG | 0.86 | 0 | 0 | 0.68 | 0.26 | 8 | Endothelial Cells |
| APOL1 | 0.86 | 0 | 0 | 0.34 | 0.08 | 8 | Endothelial Cells |
| GNAI2 | 0.86 | 0 | 0 | 0.65 | 0.33 | 8 | Endothelial Cells |
| TMEM140 | 0.85 | 0 | 0 | 0.34 | 0.07 | 8 | Endothelial Cells |
| SYNGR2 | 0.85 | 0 | 0 | 0.55 | 0.25 | 8 | Endothelial Cells |
| CALCRL | 0.85 | 0 | 0 | 0.51 | 0.18 | 8 | Endothelial Cells |
| ERG | 0.85 | 0 | 0 | 0.40 | 0.10 | 8 | Endothelial Cells |
| RGS9 | 0.84 | 0 | 0 | 0.36 | 0.09 | 8 | Endothelial Cells |
| GIMAP4 | 0.83 | 0 | 0 | 0.56 | 0.17 | 8 | Endothelial Cells |
| ID3 | 0.83 | 0 | 0 | 0.82 | 0.52 | 8 | Endothelial Cells |
| HLA-F | 0.82 | 0 | 0 | 0.39 | 0.12 | 8 | Endothelial Cells |
| PSMB8 | 0.81 | 0 | 0 | 0.57 | 0.25 | 8 | Endothelial Cells |
| KANK3 | 0.81 | 0 | 0 | 0.45 | 0.13 | 8 | Endothelial Cells |
| GSDMD | 0.81 | 0 | 0 | 0.44 | 0.18 | 8 | Endothelial Cells |
| CFAP20 | 0.8 | 0 | 0 | 0.46 | 0.25 | 8 | Endothelial Cells |
| CYB561 | 0.8 | 0 | 0 | 0.32 | 0.05 | 8 | Endothelial Cells |
| RBM17 | 0.8 | 0 | 0 | 0.54 | 0.36 | 8 | Endothelial Cells |
| FCGRT | 0.8 | 0 | 0 | 0.78 | 0.55 | 8 | Endothelial Cells |
| SARAF | 0.8 | 0 | 0 | 0.77 | 0.63 | 8 | Endothelial Cells |
| AFAP1L1 | 0.8 | 0 | 0 | 0.34 | 0.06 | 8 | Endothelial Cells |
| KLF4 | 0.79 | 0 | 0 | 0.40 | 0.15 | 8 | Endothelial Cells |
| PARP14 | 0.79 | 0 | 0 | 0.33 | 0.10 | 8 | Endothelial Cells |
| bP-2189O9.2 | 0.79 | 0 | 0 | 0.30 | 0.06 | 8 | Endothelial Cells |
| PSMB10 | 0.79 | 0 | 0 | 0.49 | 0.24 | 8 | Endothelial Cells |
| DHRS7 | 0.79 | 0 | 0 | 0.49 | 0.23 | 8 | Endothelial Cells |
| BCL6B | 0.79 | 0 | 0 | 0.28 | 0.04 | 8 | Endothelial Cells |
| S100A16 | 0.79 | 0 | 0 | 0.68 | 0.33 | 8 | Endothelial Cells |
| TAP1 | 0.78 | 0 | 0 | 0.33 | 0.12 | 8 | Endothelial Cells |
| PSME1 | 0.78 | 0 | 0 | 0.77 | 0.59 | 8 | Endothelial Cells |
| CAV1 | 0.78 | 0 | 0 | 0.98 | 0.82 | 8 | Endothelial Cells |
| PSME2P2 | 0.78 | 0 | 0 | 0.42 | 0.21 | 8 | Endothelial Cells |
| CDH13 | 0.77 | 0 | 0 | 0.28 | 0.07 | 8 | Endothelial Cells |
| SCHIP1.1 | 0.77 | 0 | 0 | 0.35 | 0.10 | 8 | Endothelial Cells |
| NRP1 | 0.77 | 0 | 0 | 0.43 | 0.19 | 8 | Endothelial Cells |
| SASH1 | 0.77 | 0 | 0 | 0.40 | 0.14 | 8 | Endothelial Cells |
| CD34 | 0.76 | 0 | 0 | 0.46 | 0.14 | 8 | Endothelial Cells |
| MSN | 0.76 | 0 | 0 | 0.71 | 0.46 | 8 | Endothelial Cells |
| ITGA3 | 0.76 | 0 | 0 | 0.35 | 0.10 | 8 | Endothelial Cells |
| LYL1 | 0.76 | 0 | 0 | 0.36 | 0.10 | 8 | Endothelial Cells |
| IFITM3 | 0.76 | 0 | 0 | 0.99 | 0.90 | 8 | Endothelial Cells |
| BTN3A2 | 0.75 | 0 | 0 | 0.36 | 0.10 | 8 | Endothelial Cells |
| XAF1 | 0.75 | 0 | 0 | 0.30 | 0.09 | 8 | Endothelial Cells |
| CMTM3 | 0.75 | 0 | 0 | 0.50 | 0.27 | 8 | Endothelial Cells |
| PTRF | 0.75 | 0 | 0 | 0.94 | 0.68 | 8 | Endothelial Cells |
| GALNT18 | 0.75 | 0 | 0 | 0.37 | 0.11 | 8 | Endothelial Cells |
| NRN1 | 0.75 | 0 | 0 | 0.31 | 0.07 | 8 | Endothelial Cells |
| IFI35 | 0.74 | 0 | 0 | 0.34 | 0.13 | 8 | Endothelial Cells |
| TAP2 | 0.74 | 0 | 0 | 0.30 | 0.07 | 8 | Endothelial Cells |
| LAMP1 | 0.74 | 0 | 0 | 0.65 | 0.43 | 8 | Endothelial Cells |
| LDB2 | 0.74 | 0 | 0 | 0.48 | 0.17 | 8 | Endothelial Cells |
| BSG | 0.74 | 0 | 0 | 0.78 | 0.65 | 8 | Endothelial Cells |
| TNFRSF14 | 0.74 | 0 | 0 | 0.34 | 0.12 | 8 | Endothelial Cells |
| ITGA1 | 0.74 | 0 | 0 | 0.54 | 0.25 | 8 | Endothelial Cells |
| ARHGAP18 | 0.73 | 0 | 0 | 0.42 | 0.11 | 8 | Endothelial Cells |
| FSCN1 | 0.73 | 0 | 0 | 0.48 | 0.21 | 8 | Endothelial Cells |
| GATA2 | 0.73 | 0 | 0 | 0.30 | 0.06 | 8 | Endothelial Cells |
| RTN4 | 0.73 | 0 | 0 | 0.76 | 0.63 | 8 | Endothelial Cells |
| JUP | 0.72 | 0 | 0 | 0.38 | 0.16 | 8 | Endothelial Cells |
| SOCS3 | 0.72 | 0 | 0 | 0.33 | 0.18 | 8 | Endothelial Cells |
| CLEC14A | 0.72 | 0 | 0 | 0.60 | 0.18 | 8 | Endothelial Cells |
| PPA1 | 0.72 | 0 | 0 | 0.56 | 0.36 | 8 | Endothelial Cells |
| DHRS3 | 0.71 | 0 | 0 | 0.32 | 0.10 | 8 | Endothelial Cells |
| PROCR | 0.71 | 0 | 0 | 0.37 | 0.13 | 8 | Endothelial Cells |
| IFNGR1 | 0.71 | 0 | 0 | 0.48 | 0.26 | 8 | Endothelial Cells |
| DLL4 | 0.71 | 0 | 0 | 0.27 | 0.05 | 8 | Endothelial Cells |
| ANXA1 | 0.7 | 0 | 0 | 0.70 | 0.39 | 8 | Endothelial Cells |
| CTNND1 | 0.7 | 0 | 0 | 0.50 | 0.24 | 8 | Endothelial Cells |
| ZNF331 | 0.7 | 0 | 0 | 0.29 | 0.10 | 8 | Endothelial Cells |
| QKI | 0.7 | 0 | 0 | 0.43 | 0.19 | 8 | Endothelial Cells |
| EFNA1 | 0.69 | 0 | 0 | 0.41 | 0.13 | 8 | Endothelial Cells |
| FLT1 | 0.69 | 0 | 0 | 0.30 | 0.07 | 8 | Endothelial Cells |
| MAPKAPK5-AS1 | 0.69 | 0 | 0 | 0.33 | 0.11 | 8 | Endothelial Cells |
| PREX1 | 0.69 | 0 | 0 | 0.27 | 0.07 | 8 | Endothelial Cells |
| SH2D3C | 0.69 | 0 | 0 | 0.33 | 0.11 | 8 | Endothelial Cells |
| OCLN | 0.69 | 0 | 0 | 0.26 | 0.06 | 8 | Endothelial Cells |
| TMSB10 | 0.69 | 0 | 0 | 1.00 | 0.99 | 8 | Endothelial Cells |
| KITLG | 0.69 | 0 | 0 | 0.27 | 0.07 | 8 | Endothelial Cells |
| PLA2G4C | 0.69 | 0 | 0 | 0.28 | 0.03 | 8 | Endothelial Cells |
| AFMID | 0.68 | 0 | 0 | 0.26 | 0.07 | 8 | Endothelial Cells |
| THSD1 | 0.68 | 0 | 0 | 0.34 | 0.08 | 8 | Endothelial Cells |
| WARS | 0.68 | 0 | 0 | 0.39 | 0.19 | 8 | Endothelial Cells |
| PTPRN2 | 0.68 | 0 | 0 | 0.30 | 0.06 | 8 | Endothelial Cells |
| LAP3 | 0.68 | 0 | 0 | 0.41 | 0.21 | 8 | Endothelial Cells |
| TFPI | 0.67 | 0 | 0 | 0.64 | 0.42 | 8 | Endothelial Cells |
| SLC12A2 | 0.67 | 0 | 0 | 0.30 | 0.11 | 8 | Endothelial Cells |
| PIK3IP1 | 0.67 | 0 | 0 | 0.28 | 0.08 | 8 | Endothelial Cells |
| PEA15 | 0.67 | 0 | 0 | 0.46 | 0.26 | 8 | Endothelial Cells |
| CAPZA2 | 0.66 | 0 | 0 | 0.68 | 0.51 | 8 | Endothelial Cells |
| LTC4S | 0.66 | 0 | 0 | 0.27 | 0.07 | 8 | Endothelial Cells |
| TMEM179B | 0.66 | 0 | 0 | 0.43 | 0.27 | 8 | Endothelial Cells |
| PTTG1IP | 0.66 | 0 | 0 | 0.53 | 0.32 | 8 | Endothelial Cells |
| HLX | 0.65 | 0 | 0 | 0.30 | 0.12 | 8 | Endothelial Cells |
| PHLDA1 | 0.65 | 0 | 0 | 0.41 | 0.23 | 8 | Endothelial Cells |
| TAPBP | 0.65 | 0 | 0 | 0.46 | 0.24 | 8 | Endothelial Cells |
| ID1 | 0.64 | 0 | 0 | 0.59 | 0.34 | 8 | Endothelial Cells |
| SEC14L1 | 0.63 | 0 | 0 | 0.33 | 0.13 | 8 | Endothelial Cells |
| ARL2 | 0.63 | 0 | 0 | 0.52 | 0.34 | 8 | Endothelial Cells |
| RASIP1 | 0.63 | 0 | 0 | 0.30 | 0.09 | 8 | Endothelial Cells |
| LMO2 | 0.63 | 0 | 0 | 0.30 | 0.09 | 8 | Endothelial Cells |
| MAP3K11 | 0.63 | 0 | 0 | 0.36 | 0.14 | 8 | Endothelial Cells |
| CXorf36 | 0.62 | 0 | 0 | 0.29 | 0.07 | 8 | Endothelial Cells |
| SOX18 | 0.62 | 0 | 0 | 0.31 | 0.10 | 8 | Endothelial Cells |
| LSR | 0.62 | 0 | 0 | 0.27 | 0.09 | 8 | Endothelial Cells |
| JAM2 | 0.62 | 0 | 0 | 0.35 | 0.11 | 8 | Endothelial Cells |
| CASP1 | 0.61 | 0 | 0 | 0.31 | 0.12 | 8 | Endothelial Cells |
| MESDC1 | 0.61 | 0 | 0 | 0.35 | 0.15 | 8 | Endothelial Cells |
| CAPN2 | 0.61 | 0 | 0 | 0.46 | 0.31 | 8 | Endothelial Cells |
| NECAP2 | 0.61 | 0 | 0 | 0.40 | 0.22 | 8 | Endothelial Cells |
| SLC3A2 | 0.61 | 0 | 0 | 0.38 | 0.23 | 8 | Endothelial Cells |
| CASP4 | 0.61 | 0 | 0 | 0.33 | 0.15 | 8 | Endothelial Cells |
| MYLIP | 0.61 | 0 | 0 | 0.46 | 0.29 | 8 | Endothelial Cells |
| ARHGEF15 | 0.61 | 0 | 0 | 0.26 | 0.06 | 8 | Endothelial Cells |
| MFNG | 0.61 | 0 | 0 | 0.27 | 0.06 | 8 | Endothelial Cells |
| PRPSAP1 | 0.61 | 0 | 0 | 0.36 | 0.18 | 8 | Endothelial Cells |
| MLLT4 | 0.6 | 0 | 0 | 0.36 | 0.16 | 8 | Endothelial Cells |
| RPN1 | 0.6 | 0 | 0 | 0.45 | 0.25 | 8 | Endothelial Cells |
| LAPTM4A | 0.6 | 0 | 0 | 0.93 | 0.84 | 8 | Endothelial Cells |
| HHEX | 0.59 | 0 | 0 | 0.27 | 0.09 | 8 | Endothelial Cells |
| VAMP3 | 0.59 | 0 | 0 | 0.43 | 0.26 | 8 | Endothelial Cells |
| MTIF3 | 0.59 | 0 | 0 | 0.37 | 0.22 | 8 | Endothelial Cells |
| SIRT2 | 0.59 | 0 | 0 | 0.36 | 0.19 | 8 | Endothelial Cells |
| SP100 | 0.59 | 0 | 0 | 0.35 | 0.19 | 8 | Endothelial Cells |
| APPL1 | 0.59 | 0 | 0 | 0.34 | 0.17 | 8 | Endothelial Cells |
| NFKBIA | 0.58 | 0 | 0 | 0.54 | 0.39 | 8 | Endothelial Cells |
| JUN | 0.58 | 0 | 0 | 0.86 | 0.67 | 8 | Endothelial Cells |
| GIMAP6 | 0.58 | 0 | 0 | 0.27 | 0.07 | 8 | Endothelial Cells |
| RAC1 | 0.58 | 0 | 0 | 0.90 | 0.80 | 8 | Endothelial Cells |
| RPS6KA2 | 0.58 | 0 | 0 | 0.27 | 0.10 | 8 | Endothelial Cells |
| MGST2 | 0.58 | 0 | 0 | 0.43 | 0.25 | 8 | Endothelial Cells |
| SPARC | 0.57 | 0 | 0 | 0.98 | 0.84 | 8 | Endothelial Cells |
| HOPX | 0.57 | 0 | 0 | 0.32 | 0.11 | 8 | Endothelial Cells |
| TMBIM1 | 0.57 | 0 | 0 | 0.38 | 0.24 | 8 | Endothelial Cells |
| ZEB1 | 0.57 | 0 | 0 | 0.31 | 0.14 | 8 | Endothelial Cells |
| CD46 | 0.57 | 0 | 0 | 0.42 | 0.27 | 8 | Endothelial Cells |
| REEP3 | 0.56 | 0 | 0 | 0.32 | 0.15 | 8 | Endothelial Cells |
| NUMB | 0.56 | 0 | 0 | 0.35 | 0.19 | 8 | Endothelial Cells |
| DAAM1 | 0.56 | 0 | 0 | 0.36 | 0.23 | 8 | Endothelial Cells |
| LMO7 | 0.56 | 0 | 0 | 0.27 | 0.07 | 8 | Endothelial Cells |
| LUZP1 | 0.56 | 0 | 0 | 0.33 | 0.17 | 8 | Endothelial Cells |
| SCARF1 | 0.55 | 0 | 0 | 0.26 | 0.07 | 8 | Endothelial Cells |
| RAC1P2 | 0.55 | 0 | 0 | 0.58 | 0.43 | 8 | Endothelial Cells |
| GRN | 0.55 | 0 | 0 | 0.58 | 0.40 | 8 | Endothelial Cells |
| CA4 | 0.55 | 0 | 0 | 0.32 | 0.11 | 8 | Endothelial Cells |
| CD200 | 0.55 | 0 | 0 | 0.28 | 0.11 | 8 | Endothelial Cells |
| PODXL | 0.55 | 0 | 0 | 0.29 | 0.08 | 8 | Endothelial Cells |
| TMEM50A | 0.55 | 0 | 0 | 0.55 | 0.42 | 8 | Endothelial Cells |
| GBP1 | 0.54 | 0 | 0 | 0.25 | 0.12 | 8 | Endothelial Cells |
| TCF4 | 0.54 | 0 | 0 | 0.62 | 0.47 | 8 | Endothelial Cells |
| GPX1 | 0.54 | 0 | 0 | 0.69 | 0.52 | 8 | Endothelial Cells |
| ETS1 | 0.54 | 0 | 0 | 0.31 | 0.14 | 8 | Endothelial Cells |
| TMEM100 | 0.54 | 0 | 0 | 0.32 | 0.15 | 8 | Endothelial Cells |
| CLIC1 | 0.54 | 0 | 0 | 0.82 | 0.63 | 8 | Endothelial Cells |
| AES | 0.53 | 0 | 0 | 0.57 | 0.43 | 8 | Endothelial Cells |
| CTC-479C5.12 | 0.53 | 0 | 0 | 0.26 | 0.10 | 8 | Endothelial Cells |
| MYL12A | 0.53 | 0 | 0 | 0.85 | 0.74 | 8 | Endothelial Cells |
| SERPINB1 | 0.53 | 0 | 0 | 0.34 | 0.19 | 8 | Endothelial Cells |
| TGFBR2 | 0.53 | 0 | 0 | 0.38 | 0.19 | 8 | Endothelial Cells |
| GUSBP3 | 0.53 | 0 | 0 | 0.29 | 0.15 | 8 | Endothelial Cells |
| RGL2 | 0.53 | 0 | 0 | 0.33 | 0.18 | 8 | Endothelial Cells |
| TLE4 | 0.53 | 0 | 0 | 0.32 | 0.20 | 8 | Endothelial Cells |
| CLEC2B | 0.53 | 0 | 0 | 0.34 | 0.16 | 8 | Endothelial Cells |
| RRAS | 0.53 | 0 | 0 | 0.32 | 0.15 | 8 | Endothelial Cells |
| POLR2F | 0.52 | 0 | 0 | 0.46 | 0.36 | 8 | Endothelial Cells |
| SNRK | 0.52 | 0 | 0 | 0.26 | 0.12 | 8 | Endothelial Cells |
| TSPO | 0.52 | 0 | 0 | 0.61 | 0.43 | 8 | Endothelial Cells |
| TJP1 | 0.52 | 0 | 0 | 0.44 | 0.28 | 8 | Endothelial Cells |
| TMEM219 | 0.52 | 0 | 0 | 0.44 | 0.31 | 8 | Endothelial Cells |
| SYNE2 | 0.52 | 0 | 0 | 0.32 | 0.18 | 8 | Endothelial Cells |
| DYNLL1 | 0.52 | 0 | 0 | 0.86 | 0.78 | 8 | Endothelial Cells |
| PRKCH | 0.52 | 0 | 0 | 0.25 | 0.09 | 8 | Endothelial Cells |
| KIAA0355 | 0.52 | 0 | 0 | 0.34 | 0.17 | 8 | Endothelial Cells |
| C5orf56 | 0.52 | 0 | 0 | 0.26 | 0.10 | 8 | Endothelial Cells |
| MID1IP1 | 0.51 | 0 | 0 | 0.27 | 0.15 | 8 | Endothelial Cells |
| S1PR1 | 0.51 | 0 | 0 | 0.31 | 0.13 | 8 | Endothelial Cells |
| DNAJA1 | 0.51 | 0 | 0 | 0.57 | 0.48 | 8 | Endothelial Cells |
| NEAT1 | 0.51 | 0 | 0 | 0.83 | 0.62 | 8 | Endothelial Cells |
| RNF13 | 0.51 | 0 | 0 | 0.35 | 0.21 | 8 | Endothelial Cells |
| FLI1 | 0.5 | 0 | 0 | 0.27 | 0.11 | 8 | Endothelial Cells |
| TRIOBP | 0.5 | 0 | 0 | 0.34 | 0.19 | 8 | Endothelial Cells |
| MYL12B | 0.5 | 0 | 0 | 0.85 | 0.77 | 8 | Endothelial Cells |
| ETS2 | 0.49 | 0 | 0 | 0.44 | 0.27 | 8 | Endothelial Cells |
| HDAC7 | 0.49 | 0 | 0 | 0.30 | 0.16 | 8 | Endothelial Cells |
| UACA | 0.49 | 0 | 0 | 0.40 | 0.24 | 8 | Endothelial Cells |
| MARCKS | 0.49 | 0 | 0 | 0.65 | 0.52 | 8 | Endothelial Cells |
| NES | 0.49 | 0 | 0 | 0.26 | 0.12 | 8 | Endothelial Cells |
| NOTCH4 | 0.49 | 0 | 0 | 0.28 | 0.12 | 8 | Endothelial Cells |
| PDCL | 0.49 | 0 | 0 | 0.26 | 0.14 | 8 | Endothelial Cells |
| PSMA3 | 0.49 | 0 | 0 | 0.42 | 0.33 | 8 | Endothelial Cells |
| CSTB | 0.48 | 0 | 0 | 0.64 | 0.49 | 8 | Endothelial Cells |
| CRBN | 0.48 | 0 | 0 | 0.32 | 0.20 | 8 | Endothelial Cells |
| FXYD6 | 0.48 | 0 | 0 | 0.53 | 0.41 | 8 | Endothelial Cells |
| BMPR2 | 0.48 | 0 | 0 | 0.36 | 0.21 | 8 | Endothelial Cells |
| TBX2 | 0.47 | 0 | 0 | 0.48 | 0.35 | 8 | Endothelial Cells |
| RAB5A | 0.47 | 0 | 0 | 0.33 | 0.20 | 8 | Endothelial Cells |
| MEF2A | 0.47 | 0 | 0 | 0.30 | 0.16 | 8 | Endothelial Cells |
| EVA1B | 0.47 | 0 | 0 | 0.29 | 0.15 | 8 | Endothelial Cells |
| GABARAPL1 | 0.47 | 0 | 0 | 0.27 | 0.15 | 8 | Endothelial Cells |
| SIGIRR | 0.46 | 0 | 0 | 0.28 | 0.16 | 8 | Endothelial Cells |
| TPM3 | 0.46 | 0 | 0 | 0.46 | 0.30 | 8 | Endothelial Cells |
| SPPL2A | 0.46 | 0 | 0 | 0.26 | 0.18 | 8 | Endothelial Cells |
| CKLF | 0.46 | 0 | 0 | 0.31 | 0.18 | 8 | Endothelial Cells |
| EPB41L2 | 0.46 | 0 | 0 | 0.37 | 0.26 | 8 | Endothelial Cells |
| YWHAE | 0.46 | 0 | 0 | 0.74 | 0.63 | 8 | Endothelial Cells |
| SERTAD1 | 0.45 | 0 | 0 | 0.29 | 0.18 | 8 | Endothelial Cells |
| TMEM173 | 0.45 | 0 | 0 | 0.36 | 0.22 | 8 | Endothelial Cells |
| TNFRSF1A | 0.45 | 0 | 0 | 0.46 | 0.38 | 8 | Endothelial Cells |
| TMEM9B | 0.45 | 0 | 0 | 0.30 | 0.17 | 8 | Endothelial Cells |
| MAPK3 | 0.45 | 0 | 0 | 0.29 | 0.16 | 8 | Endothelial Cells |
| MOB2 | 0.45 | 0 | 0 | 0.33 | 0.21 | 8 | Endothelial Cells |
| AAMDC | 0.45 | 0 | 0 | 0.37 | 0.26 | 8 | Endothelial Cells |
| GIMAP1 | 0.45 | 0 | 0 | 0.25 | 0.10 | 8 | Endothelial Cells |
| GNL1 | 0.45 | 0 | 0 | 0.27 | 0.19 | 8 | Endothelial Cells |
| SOX4 | 0.44 | 0 | 0 | 0.78 | 0.76 | 8 | Endothelial Cells |
| IFI27L2 | 0.44 | 0 | 0 | 0.44 | 0.34 | 8 | Endothelial Cells |
| NDFIP1 | 0.44 | 0 | 0 | 0.57 | 0.50 | 8 | Endothelial Cells |
| ARHGEF12 | 0.44 | 0 | 0 | 0.35 | 0.25 | 8 | Endothelial Cells |
| TUSC3 | 0.44 | 0 | 0.01 | 0.29 | 0.21 | 8 | Endothelial Cells |
| CD320 | 0.44 | 0 | 0 | 0.28 | 0.15 | 8 | Endothelial Cells |
| PHACTR4 | 0.43 | 0 | 0 | 0.26 | 0.14 | 8 | Endothelial Cells |
| TXNIP | 0.43 | 0 | 0 | 0.84 | 0.67 | 8 | Endothelial Cells |
| HSPA1A | 0.43 | 0 | 0 | 0.73 | 0.64 | 8 | Endothelial Cells |
| PLEC | 0.43 | 0 | 0 | 0.25 | 0.16 | 8 | Endothelial Cells |
| NINJ1 | 0.43 | 0 | 0 | 0.28 | 0.17 | 8 | Endothelial Cells |
| TAGLN2 | 0.43 | 0 | 0 | 0.81 | 0.75 | 8 | Endothelial Cells |
| UBC | 0.43 | 0 | 0 | 0.85 | 0.84 | 8 | Endothelial Cells |
| CHMP5 | 0.43 | 0 | 0 | 0.49 | 0.42 | 8 | Endothelial Cells |
| MT-ND2 | 0.43 | 0 | 0 | 0.99 | 0.99 | 8 | Endothelial Cells |
| ACTN4 | 0.43 | 0 | 0 | 0.49 | 0.40 | 8 | Endothelial Cells |
| SHISA5 | 0.43 | 0 | 0 | 0.36 | 0.26 | 8 | Endothelial Cells |
| ITGA5 | 0.43 | 0 | 0 | 0.26 | 0.12 | 8 | Endothelial Cells |
| MAGED2 | 0.43 | 0 | 0 | 0.73 | 0.62 | 8 | Endothelial Cells |
| VIMP | 0.43 | 0 | 0 | 0.47 | 0.38 | 8 | Endothelial Cells |
| TMEM50B | 0.42 | 0 | 0 | 0.33 | 0.23 | 8 | Endothelial Cells |
| RHOA | 0.42 | 0 | 0 | 0.81 | 0.69 | 8 | Endothelial Cells |
| CDC37 | 0.42 | 0 | 0 | 0.54 | 0.49 | 8 | Endothelial Cells |
| BRI3 | 0.42 | 0 | 0 | 0.40 | 0.29 | 8 | Endothelial Cells |
| CFLAR | 0.42 | 0 | 0 | 0.39 | 0.26 | 8 | Endothelial Cells |
| FIS1 | 0.42 | 0 | 0 | 0.56 | 0.51 | 8 | Endothelial Cells |
| RAB1A | 0.41 | 0 | 0 | 0.54 | 0.48 | 8 | Endothelial Cells |
| YIPF3 | 0.41 | 0 | 0 | 0.43 | 0.33 | 8 | Endothelial Cells |
| ADGRF5 | 0.41 | 0 | 0 | 0.30 | 0.17 | 8 | Endothelial Cells |
| NECTIN2 | 0.4 | 0 | 0.01 | 0.33 | 0.24 | 8 | Endothelial Cells |
| DYNC1LI2 | 0.4 | 0 | 0 | 0.38 | 0.27 | 8 | Endothelial Cells |
| RP11-467L13.5 | 0.4 | 0 | 0 | 0.28 | 0.19 | 8 | Endothelial Cells |
| LSM2 | 0.4 | 0 | 0.01 | 0.39 | 0.32 | 8 | Endothelial Cells |
| PLEKHA1 | 0.4 | 0 | 0.01 | 0.26 | 0.17 | 8 | Endothelial Cells |
| IER2 | 0.4 | 0 | 0 | 0.56 | 0.46 | 8 | Endothelial Cells |
| CHMP1B | 0.4 | 0 | 0 | 0.31 | 0.22 | 8 | Endothelial Cells |
| SULT1A1 | 0.4 | 0 | 0 | 0.27 | 0.15 | 8 | Endothelial Cells |
| SLC44A2 | 0.4 | 0 | 0 | 0.28 | 0.17 | 8 | Endothelial Cells |
| CDKN1C | 0.39 | 0 | 0.01 | 0.30 | 0.20 | 8 | Endothelial Cells |
| POLR2L | 0.39 | 0 | 0 | 0.75 | 0.67 | 8 | Endothelial Cells |
| PXDN | 0.39 | 0 | 0 | 0.30 | 0.20 | 8 | Endothelial Cells |
| JUNB | 0.39 | 0 | 0 | 0.60 | 0.51 | 8 | Endothelial Cells |
| NAA38 | 0.39 | 0 | 0 | 0.49 | 0.38 | 8 | Endothelial Cells |
| SPTAN1 | 0.39 | 0 | 0 | 0.33 | 0.22 | 8 | Endothelial Cells |
| YWHAB | 0.39 | 0 | 0 | 0.61 | 0.59 | 8 | Endothelial Cells |
| CALM1 | 0.38 | 0 | 0 | 0.79 | 0.74 | 8 | Endothelial Cells |
| HPCAL1 | 0.38 | 0 | 0 | 0.28 | 0.15 | 8 | Endothelial Cells |
| DCAF7 | 0.38 | 0 | 0.01 | 0.31 | 0.22 | 8 | Endothelial Cells |
| VIM | 0.38 | 0 | 0 | 0.98 | 0.94 | 8 | Endothelial Cells |
| AK1 | 0.38 | 0 | 0 | 0.27 | 0.17 | 8 | Endothelial Cells |
| MARCKSL1 | 0.38 | 0 | 0 | 0.74 | 0.60 | 8 | Endothelial Cells |
| ERICH1 | 0.38 | 0 | 0.03 | 0.27 | 0.18 | 8 | Endothelial Cells |
| RHOC | 0.38 | 0 | 0 | 0.69 | 0.60 | 8 | Endothelial Cells |
| STK4 | 0.38 | 0 | 0 | 0.28 | 0.16 | 8 | Endothelial Cells |
| DYNLRB1 | 0.38 | 0 | 0 | 0.54 | 0.50 | 8 | Endothelial Cells |
| GLUD1 | 0.38 | 0 | 0 | 0.35 | 0.26 | 8 | Endothelial Cells |
| KLC1 | 0.37 | 0 | 0 | 0.31 | 0.20 | 8 | Endothelial Cells |
| STOM | 0.37 | 0 | 0 | 0.60 | 0.43 | 8 | Endothelial Cells |
| TMEM109 | 0.37 | 0 | 0.01 | 0.40 | 0.32 | 8 | Endothelial Cells |
| KRTCAP2 | 0.37 | 0 | 0 | 0.59 | 0.54 | 8 | Endothelial Cells |
| GPSM3 | 0.37 | 0 | 0 | 0.27 | 0.14 | 8 | Endothelial Cells |
| TSC22D1 | 0.37 | 0 | 0 | 0.66 | 0.61 | 8 | Endothelial Cells |
| HNRNPF | 0.36 | 0 | 0.01 | 0.51 | 0.47 | 8 | Endothelial Cells |
| POMP | 0.36 | 0 | 0 | 0.64 | 0.62 | 8 | Endothelial Cells |
| TMBIM6 | 0.36 | 0 | 0 | 0.73 | 0.67 | 8 | Endothelial Cells |
| SERINC3 | 0.36 | 0 | 0 | 0.36 | 0.24 | 8 | Endothelial Cells |
| KLF2 | 0.36 | 0 | 0 | 0.28 | 0.17 | 8 | Endothelial Cells |
| C11orf31 | 0.36 | 0 | 0 | 0.60 | 0.55 | 8 | Endothelial Cells |
| CIB1 | 0.35 | 0 | 0.01 | 0.41 | 0.33 | 8 | Endothelial Cells |
| ZNHIT1 | 0.35 | 0 | 0.03 | 0.49 | 0.45 | 8 | Endothelial Cells |
| CYSTM1 | 0.35 | 0 | 0.01 | 0.42 | 0.33 | 8 | Endothelial Cells |
| NDUFB4 | 0.35 | 0 | 0 | 0.64 | 0.61 | 8 | Endothelial Cells |
| RABAC1 | 0.35 | 0 | 0 | 0.65 | 0.59 | 8 | Endothelial Cells |
| NDUFAF3 | 0.35 | 0 | 0 | 0.51 | 0.45 | 8 | Endothelial Cells |
| TPST2 | 0.35 | 0 | 0 | 0.26 | 0.14 | 8 | Endothelial Cells |
| PSMB1 | 0.35 | 0 | 0 | 0.68 | 0.65 | 8 | Endothelial Cells |
| DDAH2 | 0.34 | 0 | 0 | 0.59 | 0.54 | 8 | Endothelial Cells |
| NTPCR | 0.34 | 0 | 0.01 | 0.26 | 0.17 | 8 | Endothelial Cells |
| ANXA7 | 0.34 | 0 | 0.05 | 0.38 | 0.31 | 8 | Endothelial Cells |
| KTN1 | 0.34 | 0 | 0 | 0.62 | 0.56 | 8 | Endothelial Cells |
| DAD1 | 0.33 | 0 | 0 | 0.68 | 0.64 | 8 | Endothelial Cells |
| HYAL2 | 0.33 | 0 | 0 | 0.40 | 0.22 | 8 | Endothelial Cells |
| ATXN3 | 0.33 | 0 | 0.04 | 0.27 | 0.18 | 8 | Endothelial Cells |
| CCND1 | 0.33 | 0 | 0 | 0.40 | 0.22 | 8 | Endothelial Cells |
| RPS27L | 0.33 | 0 | 0 | 0.61 | 0.58 | 8 | Endothelial Cells |
| RGCC | 0.33 | 0 | 0 | 0.60 | 0.33 | 8 | Endothelial Cells |
| S100A10 | 0.32 | 0 | 0 | 0.90 | 0.64 | 8 | Endothelial Cells |
| PSMA7 | 0.32 | 0 | 0 | 0.75 | 0.72 | 8 | Endothelial Cells |
| ENG | 0.31 | 0 | 0 | 0.43 | 0.30 | 8 | Endothelial Cells |
| ARPC3 | 0.31 | 0 | 0 | 0.70 | 0.61 | 8 | Endothelial Cells |
| S100A11 | 0.31 | 0 | 0 | 0.73 | 0.60 | 8 | Endothelial Cells |
| ATP6V0E1 | 0.31 | 0 | 0 | 0.65 | 0.60 | 8 | Endothelial Cells |
| ADIRF | 0.3 | 0 | 0 | 0.87 | 0.48 | 8 | Endothelial Cells |
| TIMP3 | 0.29 | 0 | 0 | 0.65 | 0.55 | 8 | Endothelial Cells |
| RNASE1 | 0.28 | 0 | 0 | 0.63 | 0.26 | 8 | Endothelial Cells |
| PDIA3 | 0.28 | 0 | 0.04 | 0.61 | 0.58 | 8 | Endothelial Cells |
| GNLY | 3.41 | 0 | 0 | 0.47 | 0.02 | 9 | T Cells |
| NKG7 | 3.36 | 0 | 0 | 0.69 | 0.05 | 9 | T Cells |
| KLRB1 | 2.93 | 0 | 0 | 0.69 | 0.01 | 9 | T Cells |
| GZMB | 2.76 | 0 | 0 | 0.53 | 0.01 | 9 | T Cells |
| CCL4 | 2.74 | 0 | 0 | 0.39 | 0.02 | 9 | T Cells |
| PRF1 | 2.61 | 0 | 0 | 0.57 | 0.01 | 9 | T Cells |
| CCL5 | 2.42 | 0 | 0 | 0.42 | 0.02 | 9 | T Cells |
| CCL3 | 2.42 | 0 | 0 | 0.26 | 0.03 | 9 | T Cells |
| CTSW | 2.39 | 0 | 0 | 0.71 | 0.01 | 9 | T Cells |
| CD7 | 2.35 | 0 | 0 | 0.82 | 0.01 | 9 | T Cells |
| GZMA | 2.33 | 0 | 0 | 0.62 | 0.01 | 9 | T Cells |
| CST7 | 2.3 | 0 | 0 | 0.58 | 0.02 | 9 | T Cells |
| XCL2 | 2.3 | 0 | 0 | 0.40 | 0.00 | 9 | T Cells |
| IFITM1 | 2.18 | 0 | 0 | 0.87 | 0.15 | 9 | T Cells |
| TRBC1 | 2.13 | 0 | 0 | 0.65 | 0.01 | 9 | T Cells |
| CORO1A | 1.96 | 0 | 0 | 0.83 | 0.10 | 9 | T Cells |
| CD247 | 1.95 | 0 | 0 | 0.66 | 0.00 | 9 | T Cells |
| KLRD1 | 1.9 | 0 | 0 | 0.48 | 0.01 | 9 | T Cells |
| TRDC | 1.88 | 0 | 0 | 0.46 | 0.00 | 9 | T Cells |
| PLAC8 | 1.87 | 0 | 0 | 0.70 | 0.06 | 9 | T Cells |
| HCST | 1.83 | 0 | 0 | 0.71 | 0.07 | 9 | T Cells |
| CD3D | 1.81 | 0 | 0 | 0.47 | 0.00 | 9 | T Cells |
| RAC2 | 1.74 | 0 | 0 | 0.73 | 0.07 | 9 | T Cells |
| LTB | 1.69 | 0 | 0 | 0.45 | 0.04 | 9 | T Cells |
| GZMM | 1.68 | 0 | 0 | 0.57 | 0.01 | 9 | T Cells |
| ITGB2 | 1.62 | 0 | 0 | 0.64 | 0.07 | 9 | T Cells |
| TRAC | 1.6 | 0 | 0 | 0.35 | 0.01 | 9 | T Cells |
| LCK | 1.6 | 0 | 0 | 0.59 | 0.01 | 9 | T Cells |
| XCL1 | 1.58 | 0 | 0 | 0.25 | 0.00 | 9 | T Cells |
| LIMD2 | 1.58 | 0 | 0 | 0.72 | 0.13 | 9 | T Cells |
| CMC1 | 1.57 | 0 | 0 | 0.43 | 0.13 | 9 | T Cells |
| PTPRC | 1.56 | 0 | 0 | 0.70 | 0.07 | 9 | T Cells |
| FCGR3A | 1.55 | 0 | 0 | 0.42 | 0.04 | 9 | T Cells |
| IL2RG | 1.54 | 0 | 0 | 0.62 | 0.03 | 9 | T Cells |
| TRBC2 | 1.53 | 0 | 0 | 0.50 | 0.01 | 9 | T Cells |
| PCED1B-AS1 | 1.46 | 0 | 0 | 0.57 | 0.02 | 9 | T Cells |
| EVL | 1.45 | 0 | 0 | 0.72 | 0.18 | 9 | T Cells |
| FGFBP2 | 1.44 | 0 | 0 | 0.30 | 0.01 | 9 | T Cells |
| SPON2 | 1.41 | 0 | 0 | 0.40 | 0.11 | 9 | T Cells |
| LSP1 | 1.39 | 0 | 0 | 0.66 | 0.09 | 9 | T Cells |
| ARHGDIB | 1.36 | 0 | 0 | 0.82 | 0.29 | 9 | T Cells |
| KLRF1 | 1.35 | 0 | 0 | 0.35 | 0.00 | 9 | T Cells |
| BIN2 | 1.31 | 0 | 0 | 0.45 | 0.03 | 9 | T Cells |
| IER2 | 1.29 | 0 | 0 | 0.77 | 0.45 | 9 | T Cells |
| RARRES3 | 1.29 | 0 | 0 | 0.52 | 0.11 | 9 | T Cells |
| CD160 | 1.28 | 0 | 0 | 0.32 | 0.00 | 9 | T Cells |
| KLRC1 | 1.28 | 0 | 0 | 0.27 | 0.00 | 9 | T Cells |
| CD53 | 1.28 | 0 | 0 | 0.54 | 0.08 | 9 | T Cells |
| GZMH | 1.27 | 0 | 0 | 0.28 | 0.00 | 9 | T Cells |
| MATK | 1.25 | 0 | 0 | 0.42 | 0.01 | 9 | T Cells |
| CD3E | 1.22 | 0 | 0 | 0.38 | 0.00 | 9 | T Cells |
| FYB | 1.22 | 0 | 0 | 0.45 | 0.04 | 9 | T Cells |
| ACAP1 | 1.21 | 0 | 0 | 0.46 | 0.04 | 9 | T Cells |
| MYO1F | 1.21 | 0 | 0 | 0.40 | 0.05 | 9 | T Cells |
| HOPX | 1.19 | 0 | 0 | 0.44 | 0.11 | 9 | T Cells |
| CD48 | 1.18 | 0 | 0 | 0.51 | 0.05 | 9 | T Cells |
| CD52 | 1.17 | 0 | 0 | 0.48 | 0.09 | 9 | T Cells |
| IL2RB | 1.17 | 0 | 0 | 0.35 | 0.00 | 9 | T Cells |
| HLA-B | 1.17 | 0 | 0 | 0.99 | 0.53 | 9 | T Cells |
| GIMAP7 | 1.16 | 0 | 0 | 0.67 | 0.16 | 9 | T Cells |
| LCP1 | 1.16 | 0 | 0 | 0.52 | 0.07 | 9 | T Cells |
| S1PR5 | 1.16 | 0 | 0 | 0.28 | 0.00 | 9 | T Cells |
| TRAF3IP3 | 1.15 | 0 | 0 | 0.42 | 0.02 | 9 | T Cells |
| PYHIN1 | 1.14 | 0 | 0 | 0.36 | 0.00 | 9 | T Cells |
| TBC1D10C | 1.14 | 0 | 0 | 0.45 | 0.02 | 9 | T Cells |
| DUSP2 | 1.13 | 0 | 0 | 0.31 | 0.01 | 9 | T Cells |
| CLIC3 | 1.11 | 0 | 0 | 0.32 | 0.05 | 9 | T Cells |
| DENND2D | 1.11 | 0 | 0 | 0.35 | 0.01 | 9 | T Cells |
| ARL4C | 1.11 | 0 | 0 | 0.36 | 0.06 | 9 | T Cells |
| CD37 | 1.09 | 0 | 0 | 0.53 | 0.07 | 9 | T Cells |
| UCP2 | 1.08 | 0 | 0 | 0.53 | 0.11 | 9 | T Cells |
| STK17A | 1.06 | 0 | 0 | 0.39 | 0.06 | 9 | T Cells |
| ZAP70 | 1.06 | 0 | 0 | 0.38 | 0.00 | 9 | T Cells |
| HLA-A | 1.06 | 0 | 0 | 1.00 | 0.69 | 9 | T Cells |
| CYBA | 1.05 | 0 | 0 | 0.86 | 0.51 | 9 | T Cells |
| B2M | 1.04 | 0 | 0 | 1.00 | 0.99 | 9 | T Cells |
| SELL | 1.04 | 0 | 0 | 0.27 | 0.03 | 9 | T Cells |
| DNAJB1 | 1.04 | 0 | 0 | 0.51 | 0.21 | 9 | T Cells |
| TXK | 1.03 | 0 | 0 | 0.33 | 0.00 | 9 | T Cells |
| ID2 | 1.03 | 0 | 0 | 0.83 | 0.61 | 9 | T Cells |
| RP11-347P5.1 | 1.02 | 0 | 0 | 0.32 | 0.03 | 9 | T Cells |
| BTG1 | 1.02 | 0 | 0 | 0.86 | 0.68 | 9 | T Cells |
| CD3G | 1.02 | 0 | 0 | 0.31 | 0.00 | 9 | T Cells |
| PLEK | 1 | 0 | 0 | 0.29 | 0.04 | 9 | T Cells |
| FCER1G | 1 | 0 | 0 | 0.58 | 0.13 | 9 | T Cells |
| JUN | 1 | 0 | 0 | 0.85 | 0.68 | 9 | T Cells |
| IL16 | 0.99 | 0 | 0 | 0.36 | 0.03 | 9 | T Cells |
| STK4 | 0.99 | 0 | 0 | 0.51 | 0.15 | 9 | T Cells |
| IRF1 | 0.98 | 0 | 0 | 0.54 | 0.19 | 9 | T Cells |
| HAVCR2 | 0.98 | 0 | 0 | 0.28 | 0.02 | 9 | T Cells |
| AC092580.4 | 0.98 | 0 | 0 | 0.26 | 0.00 | 9 | T Cells |
| LAPTM5 | 0.97 | 0 | 0 | 0.58 | 0.14 | 9 | T Cells |
| JUNB | 0.97 | 0 | 0 | 0.73 | 0.51 | 9 | T Cells |
| DOK2 | 0.97 | 0 | 0 | 0.35 | 0.05 | 9 | T Cells |
| HSPA1B | 0.96 | 0 | 0 | 0.64 | 0.37 | 9 | T Cells |
| CD69 | 0.94 | 0 | 0 | 0.26 | 0.01 | 9 | T Cells |
| GPSM3 | 0.94 | 0 | 0 | 0.47 | 0.13 | 9 | T Cells |
| PSMB9 | 0.94 | 0 | 0 | 0.56 | 0.21 | 9 | T Cells |
| RHOH | 0.93 | 0 | 0 | 0.32 | 0.01 | 9 | T Cells |
| TBX21 | 0.93 | 0 | 0 | 0.26 | 0.00 | 9 | T Cells |
| PFN1 | 0.93 | 0 | 0 | 0.95 | 0.85 | 9 | T Cells |
| LINC00861 | 0.93 | 0 | 0 | 0.28 | 0.00 | 9 | T Cells |
| ALOX5AP | 0.92 | 0 | 0 | 0.27 | 0.02 | 9 | T Cells |
| CLEC2D | 0.92 | 0 | 0 | 0.41 | 0.08 | 9 | T Cells |
| SAMD3 | 0.92 | 0 | 0 | 0.32 | 0.00 | 9 | T Cells |
| CD27 | 0.92 | 0 | 0 | 0.25 | 0.00 | 9 | T Cells |
| CYTIP | 0.92 | 0 | 0 | 0.34 | 0.04 | 9 | T Cells |
| APOBEC3G | 0.92 | 0 | 0 | 0.28 | 0.05 | 9 | T Cells |
| TYROBP | 0.91 | 0 | 0 | 0.55 | 0.12 | 9 | T Cells |
| ICAM3 | 0.91 | 0 | 0 | 0.42 | 0.08 | 9 | T Cells |
| FAM49B | 0.91 | 0 | 0 | 0.41 | 0.13 | 9 | T Cells |
| 1-Sep | 0.91 | 0 | 0 | 0.34 | 0.04 | 9 | T Cells |
| HLA-C | 0.91 | 0 | 0 | 0.92 | 0.54 | 9 | T Cells |
| SYTL1 | 0.91 | 0 | 0 | 0.28 | 0.01 | 9 | T Cells |
| RHOF | 0.91 | 0 | 0 | 0.31 | 0.05 | 9 | T Cells |
| SKAP1 | 0.9 | 0 | 0 | 0.29 | 0.01 | 9 | T Cells |
| CD300A | 0.9 | 0 | 0 | 0.27 | 0.05 | 9 | T Cells |
| CD2 | 0.9 | 0 | 0 | 0.28 | 0.00 | 9 | T Cells |
| RNF213 | 0.9 | 0 | 0 | 0.44 | 0.19 | 9 | T Cells |
| ARHGAP45 | 0.89 | 0 | 0 | 0.33 | 0.03 | 9 | T Cells |
| FYN | 0.88 | 0 | 0 | 0.40 | 0.12 | 9 | T Cells |
| ARHGEF1 | 0.88 | 0 | 0 | 0.44 | 0.13 | 9 | T Cells |
| FMNL1 | 0.88 | 0 | 0 | 0.31 | 0.03 | 9 | T Cells |
| ARPC1B | 0.87 | 0 | 0 | 0.59 | 0.30 | 9 | T Cells |
| IL7R | 0.86 | 0 | 0 | 0.32 | 0.09 | 9 | T Cells |
| FGR | 0.86 | 0 | 0 | 0.26 | 0.06 | 9 | T Cells |
| ARHGAP30 | 0.86 | 0 | 0 | 0.32 | 0.04 | 9 | T Cells |
| FAM65B | 0.86 | 0 | 0 | 0.30 | 0.04 | 9 | T Cells |
| RUNX3 | 0.85 | 0 | 0 | 0.30 | 0.03 | 9 | T Cells |
| SRGN | 0.85 | 0 | 0 | 0.72 | 0.21 | 9 | T Cells |
| TCF7 | 0.85 | 0 | 0 | 0.29 | 0.06 | 9 | T Cells |
| NCR3 | 0.84 | 0 | 0 | 0.27 | 0.00 | 9 | T Cells |
| RP11-796E2.4 | 0.84 | 0 | 0 | 0.30 | 0.09 | 9 | T Cells |
| HCLS1 | 0.84 | 0 | 0 | 0.37 | 0.06 | 9 | T Cells |
| GIMAP4 | 0.84 | 0 | 0 | 0.51 | 0.18 | 9 | T Cells |
| ATM | 0.83 | 0 | 0 | 0.35 | 0.09 | 9 | T Cells |
| DEF6 | 0.82 | 0 | 0 | 0.30 | 0.02 | 9 | T Cells |
| PTPN7 | 0.81 | 0 | 0 | 0.28 | 0.02 | 9 | T Cells |
| PRKCH | 0.8 | 0 | 0 | 0.36 | 0.09 | 9 | T Cells |
| WIPF1 | 0.8 | 0 | 0 | 0.36 | 0.12 | 9 | T Cells |
| CCND3 | 0.8 | 0 | 0 | 0.39 | 0.15 | 9 | T Cells |
| AKNA | 0.79 | 0 | 0 | 0.28 | 0.06 | 9 | T Cells |
| RASSF5 | 0.78 | 0 | 0 | 0.29 | 0.03 | 9 | T Cells |
| GADD45B | 0.78 | 0 | 0 | 0.63 | 0.42 | 9 | T Cells |
| STK17B | 0.78 | 0 | 0 | 0.28 | 0.05 | 9 | T Cells |
| SLC2A3 | 0.78 | 0 | 0 | 0.42 | 0.17 | 9 | T Cells |
| PIM1 | 0.77 | 0 | 0 | 0.34 | 0.08 | 9 | T Cells |
| EFHD2 | 0.77 | 0 | 0 | 0.32 | 0.11 | 9 | T Cells |
| CDKN2D | 0.74 | 0 | 0 | 0.26 | 0.05 | 9 | T Cells |
| AAK1 | 0.74 | 0 | 0 | 0.44 | 0.23 | 9 | T Cells |
| ADGRE5 | 0.73 | 0 | 0 | 0.34 | 0.13 | 9 | T Cells |
| PTPN4 | 0.73 | 0 | 0 | 0.29 | 0.07 | 9 | T Cells |
| PPP2R5C | 0.73 | 0 | 0 | 0.34 | 0.13 | 9 | T Cells |
| CH17-189H20.1 | 0.72 | 0 | 0 | 0.28 | 0.07 | 9 | T Cells |
| CDC42SE1 | 0.72 | 0 | 0 | 0.31 | 0.14 | 9 | T Cells |
| ARPC2 | 0.72 | 0 | 0 | 0.82 | 0.63 | 9 | T Cells |
| HEXIM1 | 0.71 | 0 | 0 | 0.42 | 0.26 | 9 | T Cells |
| MBP | 0.71 | 0 | 0 | 0.27 | 0.04 | 9 | T Cells |
| PSMB8 | 0.71 | 0 | 0 | 0.50 | 0.26 | 9 | T Cells |
| PPP1R18 | 0.71 | 0 | 0 | 0.36 | 0.15 | 9 | T Cells |
| APMAP | 0.71 | 0 | 0 | 0.37 | 0.23 | 9 | T Cells |
| SUN2 | 0.71 | 0 | 0 | 0.35 | 0.16 | 9 | T Cells |
| FCMR | 0.7 | 0 | 0 | 0.26 | 0.03 | 9 | T Cells |
| SLC38A2 | 0.7 | 0 | 0 | 0.42 | 0.25 | 9 | T Cells |
| TSTD1 | 0.7 | 0 | 0 | 0.31 | 0.07 | 9 | T Cells |
| JAK1 | 0.69 | 0 | 0 | 0.44 | 0.31 | 9 | T Cells |
| GBP2 | 0.69 | 0 | 0 | 0.32 | 0.11 | 9 | T Cells |
| MYL12A | 0.69 | 0 | 0 | 0.86 | 0.75 | 9 | T Cells |
| IL27RA | 0.68 | 0 | 0 | 0.26 | 0.05 | 9 | T Cells |
| GMFG | 0.68 | 0 | 0 | 0.58 | 0.27 | 9 | T Cells |
| SNHG9 | 0.68 | 0 | 0 | 0.36 | 0.16 | 9 | T Cells |
| ISG20 | 0.67 | 0 | 0 | 0.27 | 0.05 | 9 | T Cells |
| FOSB | 0.67 | 0 | 0 | 0.32 | 0.14 | 9 | T Cells |
| ZFP36L2 | 0.66 | 0 | 0 | 0.65 | 0.49 | 9 | T Cells |
| SLC9A3R1 | 0.65 | 0 | 0 | 0.27 | 0.06 | 9 | T Cells |
| RPS29 | 0.65 | 0 | 0 | 1.00 | 0.99 | 9 | T Cells |
| CDC42SE2 | 0.65 | 0 | 0 | 0.26 | 0.09 | 9 | T Cells |
| PTPN6 | 0.65 | 0 | 0 | 0.30 | 0.07 | 9 | T Cells |
| REL | 0.65 | 0 | 0 | 0.26 | 0.08 | 9 | T Cells |
| 9-Sep | 0.65 | 0 | 0 | 0.43 | 0.24 | 9 | T Cells |
| PRPF38B | 0.65 | 0 | 0 | 0.53 | 0.35 | 9 | T Cells |
| HLA-F | 0.64 | 0 | 0 | 0.36 | 0.12 | 9 | T Cells |
| CHST12 | 0.64 | 0 | 0 | 0.37 | 0.24 | 9 | T Cells |
| PSMB10 | 0.64 | 0 | 0 | 0.47 | 0.25 | 9 | T Cells |
| BTN3A2 | 0.64 | 0 | 0 | 0.31 | 0.11 | 9 | T Cells |
| GNG2 | 0.63 | 0 | 0 | 0.31 | 0.15 | 9 | T Cells |
| TGFB1 | 0.63 | 0 | 0 | 0.36 | 0.15 | 9 | T Cells |
| LEF1 | 0.63 | 0 | 0 | 0.25 | 0.07 | 9 | T Cells |
| GIMAP1 | 0.62 | 0 | 0 | 0.30 | 0.10 | 9 | T Cells |
| RP4-765C7.2 | 0.62 | 0 | 0 | 0.66 | 0.51 | 9 | T Cells |
| FXYD5 | 0.62 | 0 | 0 | 0.51 | 0.26 | 9 | T Cells |
| OSTF1 | 0.61 | 0 | 0 | 0.40 | 0.23 | 9 | T Cells |
| ABRACL | 0.6 | 0 | 0 | 0.35 | 0.13 | 9 | T Cells |
| C1orf56 | 0.59 | 0 | 0.01 | 0.30 | 0.18 | 9 | T Cells |
| FOS | 0.59 | 0 | 0 | 0.48 | 0.35 | 9 | T Cells |
| C9orf142 | 0.59 | 0 | 0 | 0.36 | 0.22 | 9 | T Cells |
| RCSD1 | 0.59 | 0 | 0 | 0.39 | 0.18 | 9 | T Cells |
| SRSF7 | 0.59 | 0 | 0 | 0.57 | 0.50 | 9 | T Cells |
| RPL28 | 0.58 | 0 | 0 | 0.98 | 0.98 | 9 | T Cells |
| RPS27 | 0.58 | 0 | 0 | 1.00 | 1.00 | 9 | T Cells |
| FKBP11 | 0.58 | 0 | 0 | 0.25 | 0.09 | 9 | T Cells |
| ARPC5L | 0.58 | 0 | 0 | 0.32 | 0.19 | 9 | T Cells |
| RAP1B | 0.58 | 0 | 0 | 0.45 | 0.33 | 9 | T Cells |
| FERMT3 | 0.58 | 0 | 0 | 0.26 | 0.06 | 9 | T Cells |
| ELF1 | 0.57 | 0 | 0 | 0.33 | 0.16 | 9 | T Cells |
| CLEC2B | 0.56 | 0 | 0 | 0.34 | 0.17 | 9 | T Cells |
| SIGIRR | 0.56 | 0 | 0 | 0.28 | 0.17 | 9 | T Cells |
| ABI3 | 0.56 | 0 | 0 | 0.26 | 0.09 | 9 | T Cells |
| UBE2F | 0.56 | 0 | 0 | 0.27 | 0.13 | 9 | T Cells |
| ABHD17A | 0.56 | 0 | 0 | 0.31 | 0.16 | 9 | T Cells |
| PPP1CA | 0.55 | 0 | 0 | 0.50 | 0.37 | 9 | T Cells |
| ETS1 | 0.55 | 0 | 0 | 0.32 | 0.14 | 9 | T Cells |
| C5orf56 | 0.55 | 0 | 0 | 0.25 | 0.10 | 9 | T Cells |
| PABPC1 | 0.54 | 0 | 0 | 0.80 | 0.74 | 9 | T Cells |
| TAP1 | 0.54 | 0 | 0 | 0.30 | 0.12 | 9 | T Cells |
| GTF3A | 0.54 | 0 | 0 | 0.42 | 0.25 | 9 | T Cells |
| DGKZ | 0.53 | 0 | 0 | 0.30 | 0.13 | 9 | T Cells |
| PCSK7 | 0.53 | 0 | 0 | 0.41 | 0.28 | 9 | T Cells |
| ARPC3 | 0.53 | 0 | 0 | 0.74 | 0.61 | 9 | T Cells |
| 6-Sep | 0.53 | 0 | 0 | 0.38 | 0.21 | 9 | T Cells |
| TLE4 | 0.52 | 0 | 0 | 0.37 | 0.21 | 9 | T Cells |
| VAMP8 | 0.52 | 0 | 0 | 0.43 | 0.18 | 9 | T Cells |
| HMOX2 | 0.52 | 0 | 0 | 0.32 | 0.18 | 9 | T Cells |
| HSP90AA1 | 0.52 | 0 | 0 | 0.91 | 0.86 | 9 | T Cells |
| RP11-761N21.2 | 0.51 | 0 | 0 | 0.71 | 0.62 | 9 | T Cells |
| PRMT2 | 0.51 | 0 | 0.02 | 0.43 | 0.32 | 9 | T Cells |
| EIF4A1 | 0.51 | 0 | 0.03 | 0.41 | 0.29 | 9 | T Cells |
| KLF2 | 0.5 | 0 | 0 | 0.34 | 0.18 | 9 | T Cells |
| TPST2 | 0.5 | 0 | 0 | 0.28 | 0.14 | 9 | T Cells |
| CAP1 | 0.5 | 0 | 0 | 0.60 | 0.46 | 9 | T Cells |
| IL32 | 0.49 | 0 | 0 | 0.48 | 0.28 | 9 | T Cells |
| XBP1 | 0.49 | 0 | 0.03 | 0.42 | 0.30 | 9 | T Cells |
| RHOG | 0.49 | 0 | 0 | 0.38 | 0.21 | 9 | T Cells |
| BRD2 | 0.48 | 0 | 0.05 | 0.41 | 0.31 | 9 | T Cells |
| DDX39A | 0.48 | 0 | 0.04 | 0.27 | 0.16 | 9 | T Cells |
| INTS6 | 0.47 | 0 | 0.01 | 0.32 | 0.19 | 9 | T Cells |
| HSPA8 | 0.47 | 0 | 0 | 0.70 | 0.60 | 9 | T Cells |
| RPSA | 0.46 | 0 | 0 | 0.86 | 0.90 | 9 | T Cells |
| AES | 0.46 | 0 | 0 | 0.56 | 0.43 | 9 | T Cells |
| RPL13P12 | 0.46 | 0 | 0 | 0.95 | 0.96 | 9 | T Cells |
| C9orf16 | 0.46 | 0 | 0.01 | 0.42 | 0.30 | 9 | T Cells |
| SNHG5 | 0.46 | 0 | 0 | 0.66 | 0.57 | 9 | T Cells |
| CD44 | 0.46 | 0 | 0 | 0.43 | 0.26 | 9 | T Cells |
| APRT | 0.45 | 0 | 0 | 0.54 | 0.42 | 9 | T Cells |
| TPM3 | 0.45 | 0 | 0 | 0.48 | 0.30 | 9 | T Cells |
| CLIC1 | 0.45 | 0 | 0 | 0.68 | 0.64 | 9 | T Cells |
| PSME1 | 0.44 | 0 | 0.01 | 0.65 | 0.60 | 9 | T Cells |
| RPS21 | 0.44 | 0 | 0 | 0.92 | 0.93 | 9 | T Cells |
| DDX5 | 0.43 | 0 | 0 | 0.90 | 0.79 | 9 | T Cells |
| HLA-E | 0.43 | 0 | 0 | 0.90 | 0.59 | 9 | T Cells |
| GYPC | 0.42 | 0 | 0.03 | 0.44 | 0.33 | 9 | T Cells |
| EEF1B2 | 0.42 | 0 | 0 | 0.73 | 0.74 | 9 | T Cells |
| RPLP2 | 0.42 | 0 | 0 | 1.00 | 0.99 | 9 | T Cells |
| PSME2 | 0.41 | 0 | 0 | 0.56 | 0.45 | 9 | T Cells |
| OCIAD2 | 0.41 | 0 | 0 | 0.29 | 0.14 | 9 | T Cells |
| DDIT4 | 0.4 | 0 | 0.03 | 0.28 | 0.16 | 9 | T Cells |
| TWF2 | 0.4 | 0 | 0.01 | 0.28 | 0.16 | 9 | T Cells |
| SH3BGRL3 | 0.39 | 0 | 0 | 0.73 | 0.58 | 9 | T Cells |
| TMSB4X | 0.38 | 0 | 0 | 1.00 | 1.00 | 9 | T Cells |
| RPS19 | 0.38 | 0 | 0 | 1.00 | 0.99 | 9 | T Cells |
| RPS15A | 0.37 | 0 | 0 | 0.99 | 0.99 | 9 | T Cells |
| RP11-889L3.1 | 0.37 | 0 | 0 | 0.92 | 0.86 | 9 | T Cells |
| PYCARD | 0.37 | 0 | 0.02 | 0.26 | 0.14 | 9 | T Cells |
| EMP3 | 0.36 | 0 | 0 | 0.70 | 0.59 | 9 | T Cells |
| RBM39 | 0.35 | 0 | 0.04 | 0.67 | 0.64 | 9 | T Cells |
| RPL37 | 0.34 | 0 | 0 | 0.95 | 0.98 | 9 | T Cells |
| RPL19 | 0.33 | 0 | 0 | 1.00 | 0.99 | 9 | T Cells |
| RPL18A | 0.33 | 0 | 0 | 0.98 | 0.99 | 9 | T Cells |
| RPL13 | 0.32 | 0 | 0 | 1.00 | 1.00 | 9 | T Cells |
| RPS18 | 0.32 | 0 | 0.03 | 1.00 | 1.00 | 9 | T Cells |
| CFL1 | 0.32 | 0 | 0 | 0.90 | 0.91 | 9 | T Cells |
| RPS2 | 0.31 | 0 | 0 | 1.00 | 0.99 | 9 | T Cells |
| CTD-2192J16.15 | 0.3 | 0 | 0 | 0.91 | 0.91 | 9 | T Cells |
| UBA52 | 0.3 | 0 | 0 | 0.95 | 0.95 | 9 | T Cells |
| RPL12 | 0.3 | 0 | 0.01 | 0.95 | 0.97 | 9 | T Cells |
| RPS10 | 0.29 | 0 | 0 | 0.86 | 0.90 | 9 | T Cells |
| RPS3 | 0.28 | 0 | 0 | 0.98 | 0.98 | 9 | T Cells |
| RPL23A | 0.27 | 0 | 0 | 0.98 | 0.98 | 9 | T Cells |
| EIF1 | 0.27 | 0 | 0.01 | 0.96 | 0.96 | 9 | T Cells |
| RPL10 | 0.27 | 0 | 0 | 1.00 | 1.00 | 9 | T Cells |
| RPL30 | 0.26 | 0 | 0 | 0.98 | 0.98 | 9 | T Cells |
| MALAT1 | 0.26 | 0 | 0 | 1.00 | 1.00 | 9 | T Cells |
| RPL27 | 0.26 | 0 | 0 | 0.93 | 0.96 | 9 | T Cells |
| ACKR1 | 3.96 | 0 | 0 | 0.83 | 0.08 | 10 | Endothelial Cells |
| PTGDS | 2.17 | 0 | 0 | 0.83 | 0.15 | 10 | Endothelial Cells |
| RNASE1 | 2.08 | 0 | 0 | 0.99 | 0.26 | 10 | Endothelial Cells |
| VWF | 2.08 | 0 | 0 | 0.92 | 0.11 | 10 | Endothelial Cells |
| HYAL2 | 2.04 | 0 | 0 | 0.86 | 0.22 | 10 | Endothelial Cells |
| SLCO2A1 | 1.99 | 0 | 0 | 0.82 | 0.09 | 10 | Endothelial Cells |
| APLNR | 1.94 | 0 | 0 | 0.80 | 0.11 | 10 | Endothelial Cells |
| THBD | 1.91 | 0 | 0 | 0.86 | 0.13 | 10 | Endothelial Cells |
| ACKR3 | 1.79 | 0 | 0 | 0.73 | 0.02 | 10 | Endothelial Cells |
| DUSP23 | 1.75 | 0 | 0 | 0.73 | 0.11 | 10 | Endothelial Cells |
| VCAM1 | 1.75 | 0 | 0 | 0.58 | 0.03 | 10 | Endothelial Cells |
| NNMT | 1.74 | 0 | 0 | 0.63 | 0.03 | 10 | Endothelial Cells |
| CD59 | 1.7 | 0 | 0 | 0.88 | 0.60 | 10 | Endothelial Cells |
| RGS5 | 1.63 | 0 | 0 | 0.91 | 0.23 | 10 | Endothelial Cells |
| AQP1 | 1.63 | 0 | 0 | 0.85 | 0.20 | 10 | Endothelial Cells |
| MMRN2 | 1.6 | 0 | 0 | 0.79 | 0.07 | 10 | Endothelial Cells |
| CPE | 1.58 | 0 | 0 | 0.76 | 0.25 | 10 | Endothelial Cells |
| PECAM1 | 1.58 | 0 | 0 | 0.98 | 0.25 | 10 | Endothelial Cells |
| RAMP3 | 1.56 | 0 | 0 | 0.82 | 0.15 | 10 | Endothelial Cells |
| IGFBP4 | 1.55 | 0 | 0 | 0.99 | 0.49 | 10 | Endothelial Cells |
| PCAT19 | 1.55 | 0 | 0 | 0.92 | 0.17 | 10 | Endothelial Cells |
| ICAM1 | 1.48 | 0 | 0 | 0.60 | 0.10 | 10 | Endothelial Cells |
| HSPG2 | 1.48 | 0 | 0 | 0.83 | 0.18 | 10 | Endothelial Cells |
| CLU | 1.47 | 0 | 0 | 0.77 | 0.31 | 10 | Endothelial Cells |
| S100P | 1.42 | 0 | 0 | 0.45 | 0.01 | 10 | Endothelial Cells |
| LYPD2 | 1.41 | 0 | 0 | 0.47 | 0.02 | 10 | Endothelial Cells |
| HEG1 | 1.4 | 0 | 0 | 0.75 | 0.13 | 10 | Endothelial Cells |
| SELP | 1.39 | 0 | 0 | 0.65 | 0.01 | 10 | Endothelial Cells |
| CCL14 | 1.38 | 0 | 0 | 0.43 | 0.03 | 10 | Endothelial Cells |
| IFITM2 | 1.38 | 0 | 0 | 0.96 | 0.63 | 10 | Endothelial Cells |
| CALCRL | 1.36 | 0 | 0 | 0.95 | 0.18 | 10 | Endothelial Cells |
| ECSCR.1 | 1.35 | 0 | 0 | 0.84 | 0.18 | 10 | Endothelial Cells |
| MALL | 1.35 | 0 | 0 | 0.64 | 0.04 | 10 | Endothelial Cells |
| IFITM3 | 1.34 | 0 | 0 | 1.00 | 0.91 | 10 | Endothelial Cells |
| CLEC14A | 1.29 | 0 | 0 | 0.93 | 0.18 | 10 | Endothelial Cells |
| FBLN2 | 1.26 | 0 | 0 | 0.49 | 0.05 | 10 | Endothelial Cells |
| PTPRB | 1.25 | 0 | 0 | 0.69 | 0.06 | 10 | Endothelial Cells |
| ADAM15 | 1.24 | 0 | 0 | 0.66 | 0.10 | 10 | Endothelial Cells |
| FAM167B | 1.24 | 0 | 0 | 0.66 | 0.07 | 10 | Endothelial Cells |
| SRPX | 1.24 | 0 | 0 | 0.67 | 0.14 | 10 | Endothelial Cells |
| PCDH17 | 1.24 | 0 | 0 | 0.69 | 0.14 | 10 | Endothelial Cells |
| TIMP1 | 1.23 | 0 | 0 | 0.97 | 0.77 | 10 | Endothelial Cells |
| IFI27 | 1.2 | 0 | 0 | 1.00 | 0.35 | 10 | Endothelial Cells |
| ID1 | 1.2 | 0 | 0 | 0.75 | 0.34 | 10 | Endothelial Cells |
| FKBP1A | 1.2 | 0 | 0 | 0.97 | 0.72 | 10 | Endothelial Cells |
| IFITM1 | 1.16 | 0 | 0 | 0.64 | 0.17 | 10 | Endothelial Cells |
| CLDN11 | 1.16 | 0 | 0 | 0.49 | 0.03 | 10 | Endothelial Cells |
| CD93 | 1.15 | 0 | 0 | 0.71 | 0.14 | 10 | Endothelial Cells |
| PRCP | 1.15 | 0 | 0 | 0.76 | 0.23 | 10 | Endothelial Cells |
| EMP3 | 1.14 | 0 | 0 | 0.83 | 0.59 | 10 | Endothelial Cells |
| TPST2 | 1.14 | 0 | 0 | 0.65 | 0.14 | 10 | Endothelial Cells |
| ENTPD1 | 1.13 | 0 | 0 | 0.62 | 0.12 | 10 | Endothelial Cells |
| FOXP1 | 1.11 | 0 | 0 | 0.88 | 0.39 | 10 | Endothelial Cells |
| LTC4S | 1.11 | 0 | 0 | 0.57 | 0.07 | 10 | Endothelial Cells |
| TGM2 | 1.1 | 0 | 0 | 0.87 | 0.38 | 10 | Endothelial Cells |
| KLF2 | 1.09 | 0 | 0 | 0.62 | 0.17 | 10 | Endothelial Cells |
| RAMP2 | 1.09 | 0 | 0 | 0.99 | 0.33 | 10 | Endothelial Cells |
| VEGFC | 1.09 | 0 | 0 | 0.47 | 0.03 | 10 | Endothelial Cells |
| CYP1B1 | 1.08 | 0 | 0 | 0.33 | 0.01 | 10 | Endothelial Cells |
| NPDC1 | 1.07 | 0 | 0 | 0.86 | 0.29 | 10 | Endothelial Cells |
| CD320 | 1.04 | 0 | 0 | 0.61 | 0.14 | 10 | Endothelial Cells |
| TIMP3 | 1.04 | 0 | 0 | 0.75 | 0.55 | 10 | Endothelial Cells |
| NRN1 | 1.02 | 0 | 0 | 0.55 | 0.07 | 10 | Endothelial Cells |
| TEK | 0.98 | 0 | 0 | 0.56 | 0.06 | 10 | Endothelial Cells |
| ANXA2 | 0.97 | 0 | 0 | 0.93 | 0.65 | 10 | Endothelial Cells |
| IGF2 | 0.96 | 0 | 0 | 0.55 | 0.05 | 10 | Endothelial Cells |
| IL32 | 0.96 | 0 | 0 | 0.86 | 0.27 | 10 | Endothelial Cells |
| IL33 | 0.96 | 0 | 0 | 0.68 | 0.11 | 10 | Endothelial Cells |
| WARS | 0.95 | 0 | 0 | 0.62 | 0.19 | 10 | Endothelial Cells |
| TMSB10 | 0.94 | 0 | 0 | 1.00 | 0.99 | 10 | Endothelial Cells |
| SH3BP5 | 0.94 | 0 | 0 | 0.75 | 0.20 | 10 | Endothelial Cells |
| EGFL7 | 0.94 | 0 | 0 | 0.89 | 0.25 | 10 | Endothelial Cells |
| ENG | 0.93 | 0 | 0 | 0.78 | 0.29 | 10 | Endothelial Cells |
| LHX6 | 0.92 | 0 | 0 | 0.46 | 0.01 | 10 | Endothelial Cells |
| FXYD5 | 0.91 | 0 | 0 | 0.80 | 0.25 | 10 | Endothelial Cells |
| CTNNAL1 | 0.91 | 0 | 0 | 0.64 | 0.19 | 10 | Endothelial Cells |
| TMEM176B | 0.89 | 0 | 0 | 0.60 | 0.17 | 10 | Endothelial Cells |
| ADCY4 | 0.89 | 0 | 0 | 0.58 | 0.09 | 10 | Endothelial Cells |
| CARHSP1 | 0.89 | 0 | 0 | 0.77 | 0.36 | 10 | Endothelial Cells |
| PROCR | 0.88 | 0 | 0 | 0.57 | 0.13 | 10 | Endothelial Cells |
| UGCG | 0.88 | 0 | 0 | 0.47 | 0.12 | 10 | Endothelial Cells |
| MEIS2 | 0.87 | 0 | 0 | 0.55 | 0.11 | 10 | Endothelial Cells |
| ASRGL1 | 0.86 | 0 | 0 | 0.45 | 0.04 | 10 | Endothelial Cells |
| CRIP2 | 0.86 | 0 | 0 | 0.95 | 0.65 | 10 | Endothelial Cells |
| LIMS2 | 0.86 | 0 | 0 | 0.53 | 0.08 | 10 | Endothelial Cells |
| IL1R1 | 0.86 | 0 | 0 | 0.50 | 0.11 | 10 | Endothelial Cells |
| ARL4A | 0.84 | 0 | 0 | 0.47 | 0.11 | 10 | Endothelial Cells |
| GUK1 | 0.84 | 0 | 0 | 0.93 | 0.66 | 10 | Endothelial Cells |
| PLVAP | 0.82 | 0 | 0 | 0.35 | 0.01 | 10 | Endothelial Cells |
| EDN1 | 0.82 | 0 | 0 | 0.38 | 0.10 | 10 | Endothelial Cells |
| FLT1 | 0.81 | 0 | 0 | 0.46 | 0.07 | 10 | Endothelial Cells |
| ARPC1B | 0.81 | 0 | 0 | 0.73 | 0.30 | 10 | Endothelial Cells |
| TSPAN7 | 0.8 | 0 | 0 | 0.58 | 0.12 | 10 | Endothelial Cells |
| EHD4 | 0.79 | 0 | 0 | 0.46 | 0.06 | 10 | Endothelial Cells |
| GIMAP7 | 0.79 | 0 | 0 | 0.64 | 0.16 | 10 | Endothelial Cells |
| TM4SF1 | 0.79 | 0 | 0 | 0.87 | 0.39 | 10 | Endothelial Cells |
| NDUFA8 | 0.78 | 0 | 0 | 0.63 | 0.26 | 10 | Endothelial Cells |
| HLA-B | 0.78 | 0 | 0 | 0.98 | 0.53 | 10 | Endothelial Cells |
| S100A10 | 0.78 | 0 | 0 | 0.99 | 0.65 | 10 | Endothelial Cells |
| NUAK1 | 0.78 | 0 | 0 | 0.38 | 0.04 | 10 | Endothelial Cells |
| KCTD12 | 0.77 | 0 | 0 | 0.71 | 0.23 | 10 | Endothelial Cells |
| TPM3 | 0.77 | 0 | 0 | 0.69 | 0.30 | 10 | Endothelial Cells |
| IGFBP2 | 0.77 | 0 | 0 | 0.64 | 0.18 | 10 | Endothelial Cells |
| FAM110D | 0.77 | 0 | 0 | 0.45 | 0.04 | 10 | Endothelial Cells |
| SDCBP | 0.76 | 0 | 0 | 0.84 | 0.52 | 10 | Endothelial Cells |
| FLNB | 0.76 | 0 | 0 | 0.43 | 0.05 | 10 | Endothelial Cells |
| NDRG1 | 0.76 | 0 | 0 | 0.53 | 0.16 | 10 | Endothelial Cells |
| MARCKSL1 | 0.76 | 0 | 0 | 0.90 | 0.60 | 10 | Endothelial Cells |
| TAGLN2 | 0.75 | 0 | 0 | 0.94 | 0.75 | 10 | Endothelial Cells |
| ANXA3 | 0.74 | 0 | 0 | 0.53 | 0.11 | 10 | Endothelial Cells |
| PLAT | 0.74 | 0 | 0 | 0.49 | 0.14 | 10 | Endothelial Cells |
| TRIOBP | 0.74 | 0 | 0 | 0.58 | 0.19 | 10 | Endothelial Cells |
| PLXND1 | 0.74 | 0 | 0 | 0.54 | 0.12 | 10 | Endothelial Cells |
| SEMA3F | 0.73 | 0 | 0 | 0.33 | 0.03 | 10 | Endothelial Cells |
| CDC25B | 0.73 | 0 | 0 | 0.44 | 0.11 | 10 | Endothelial Cells |
| ITGA5 | 0.73 | 0 | 0 | 0.49 | 0.12 | 10 | Endothelial Cells |
| LEPR | 0.72 | 0 | 0 | 0.41 | 0.07 | 10 | Endothelial Cells |
| CLDN5 | 0.72 | 0 | 0 | 0.83 | 0.26 | 10 | Endothelial Cells |
| LY6E | 0.72 | 0 | 0 | 0.82 | 0.45 | 10 | Endothelial Cells |
| ARL4D | 0.72 | 0 | 0 | 0.41 | 0.10 | 10 | Endothelial Cells |
| EPHB4 | 0.72 | 0 | 0 | 0.43 | 0.07 | 10 | Endothelial Cells |
| LYVE1 | 0.72 | 0 | 0 | 0.36 | 0.06 | 10 | Endothelial Cells |
| BST2 | 0.72 | 0 | 0 | 0.82 | 0.29 | 10 | Endothelial Cells |
| ICAM2 | 0.71 | 0 | 0 | 0.71 | 0.22 | 10 | Endothelial Cells |
| CD81 | 0.71 | 0 | 0 | 0.73 | 0.39 | 10 | Endothelial Cells |
| EVA1C | 0.71 | 0 | 0 | 0.43 | 0.02 | 10 | Endothelial Cells |
| H19 | 0.7 | 0 | 0 | 0.45 | 0.15 | 10 | Endothelial Cells |
| NFKBIA | 0.7 | 0 | 0 | 0.62 | 0.39 | 10 | Endothelial Cells |
| CDC37 | 0.7 | 0 | 0 | 0.80 | 0.49 | 10 | Endothelial Cells |
| NOSTRIN | 0.69 | 0 | 0 | 0.35 | 0.06 | 10 | Endothelial Cells |
| SOX18 | 0.69 | 0 | 0 | 0.43 | 0.10 | 10 | Endothelial Cells |
| NOS3 | 0.69 | 0 | 0 | 0.26 | 0.01 | 10 | Endothelial Cells |
| CTSH | 0.69 | 0 | 0 | 0.54 | 0.13 | 10 | Endothelial Cells |
| RAB11A | 0.68 | 0 | 0 | 0.75 | 0.44 | 10 | Endothelial Cells |
| CYYR1 | 0.68 | 0 | 0 | 0.52 | 0.11 | 10 | Endothelial Cells |
| CHODL | 0.68 | 0 | 0 | 0.27 | 0.01 | 10 | Endothelial Cells |
| SEPN1 | 0.67 | 0 | 0 | 0.41 | 0.09 | 10 | Endothelial Cells |
| SRGN | 0.67 | 0 | 0 | 0.82 | 0.21 | 10 | Endothelial Cells |
| TLL1 | 0.67 | 0 | 0 | 0.30 | 0.00 | 10 | Endothelial Cells |
| ECE1 | 0.66 | 0 | 0 | 0.47 | 0.15 | 10 | Endothelial Cells |
| CALR | 0.66 | 0 | 0 | 0.86 | 0.58 | 10 | Endothelial Cells |
| PALMD | 0.65 | 0 | 0 | 0.45 | 0.07 | 10 | Endothelial Cells |
| ELK3 | 0.65 | 0 | 0 | 0.51 | 0.14 | 10 | Endothelial Cells |
| CD9 | 0.65 | 0 | 0 | 0.71 | 0.45 | 10 | Endothelial Cells |
| NPR1 | 0.64 | 0 | 0 | 0.36 | 0.04 | 10 | Endothelial Cells |
| JAM2 | 0.64 | 0 | 0 | 0.48 | 0.12 | 10 | Endothelial Cells |
| EPAS1 | 0.64 | 0 | 0 | 0.82 | 0.43 | 10 | Endothelial Cells |
| RHOA | 0.64 | 0 | 0 | 0.89 | 0.69 | 10 | Endothelial Cells |
| TCN2 | 0.64 | 0 | 0 | 0.39 | 0.06 | 10 | Endothelial Cells |
| TSPO | 0.63 | 0 | 0 | 0.76 | 0.43 | 10 | Endothelial Cells |
| CCDC69 | 0.63 | 0 | 0 | 0.38 | 0.05 | 10 | Endothelial Cells |
| SLC44A2 | 0.63 | 0 | 0 | 0.52 | 0.17 | 10 | Endothelial Cells |
| RP11-467L13.5 | 0.63 | 0 | 0 | 0.55 | 0.18 | 10 | Endothelial Cells |
| S1PR1 | 0.63 | 0 | 0 | 0.47 | 0.13 | 10 | Endothelial Cells |
| PTTG1IP | 0.62 | 0 | 0 | 0.64 | 0.33 | 10 | Endothelial Cells |
| TPM4 | 0.62 | 0 | 0 | 0.82 | 0.60 | 10 | Endothelial Cells |
| SYNGR2 | 0.62 | 0 | 0 | 0.63 | 0.26 | 10 | Endothelial Cells |
| LDB2 | 0.62 | 0 | 0 | 0.59 | 0.18 | 10 | Endothelial Cells |
| CMIP | 0.62 | 0 | 0 | 0.45 | 0.09 | 10 | Endothelial Cells |
| TNFSF10 | 0.62 | 0 | 0 | 0.69 | 0.22 | 10 | Endothelial Cells |
| S100A16 | 0.62 | 0 | 0 | 0.75 | 0.34 | 10 | Endothelial Cells |
| RHOC | 0.62 | 0 | 0 | 0.83 | 0.60 | 10 | Endothelial Cells |
| MYL12A | 0.61 | 0 | 0 | 0.93 | 0.75 | 10 | Endothelial Cells |
| HDAC9 | 0.61 | 0 | 0 | 0.31 | 0.02 | 10 | Endothelial Cells |
| HPCAL1 | 0.61 | 0 | 0 | 0.45 | 0.15 | 10 | Endothelial Cells |
| EVL | 0.6 | 0 | 0 | 0.52 | 0.19 | 10 | Endothelial Cells |
| HLA-A | 0.6 | 0 | 0 | 0.98 | 0.69 | 10 | Endothelial Cells |
| ADGRL4 | 0.6 | 0 | 0 | 0.34 | 0.03 | 10 | Endothelial Cells |
| S100A11 | 0.6 | 0 | 0 | 0.89 | 0.60 | 10 | Endothelial Cells |
| ADAMTSL1 | 0.59 | 0 | 0 | 0.27 | 0.01 | 10 | Endothelial Cells |
| ESAM | 0.59 | 0 | 0 | 0.74 | 0.35 | 10 | Endothelial Cells |
| BRI3 | 0.59 | 0 | 0 | 0.60 | 0.29 | 10 | Endothelial Cells |
| NR2F2 | 0.59 | 0 | 0 | 0.63 | 0.34 | 10 | Endothelial Cells |
| BCL3 | 0.59 | 0 | 0 | 0.34 | 0.07 | 10 | Endothelial Cells |
| HSP90B1 | 0.59 | 0 | 0 | 0.84 | 0.59 | 10 | Endothelial Cells |
| MRAS | 0.59 | 0 | 0 | 0.27 | 0.04 | 10 | Endothelial Cells |
| LAP3 | 0.59 | 0 | 0 | 0.50 | 0.22 | 10 | Endothelial Cells |
| EMCN | 0.59 | 0 | 0 | 0.67 | 0.15 | 10 | Endothelial Cells |
| PLPP3 | 0.59 | 0 | 0 | 0.53 | 0.29 | 10 | Endothelial Cells |
| CD109 | 0.59 | 0 | 0 | 0.32 | 0.05 | 10 | Endothelial Cells |
| TUBA1B | 0.59 | 0 | 0 | 0.93 | 0.78 | 10 | Endothelial Cells |
| FKBP1C | 0.58 | 0 | 0 | 0.45 | 0.10 | 10 | Endothelial Cells |
| FSCN1 | 0.58 | 0 | 0 | 0.56 | 0.22 | 10 | Endothelial Cells |
| TGFBR2 | 0.58 | 0 | 0 | 0.49 | 0.19 | 10 | Endothelial Cells |
| SAMD5 | 0.58 | 0 | 0 | 0.30 | 0.04 | 10 | Endothelial Cells |
| LIFR | 0.58 | 0 | 0 | 0.41 | 0.12 | 10 | Endothelial Cells |
| MAML1 | 0.58 | 0 | 0 | 0.40 | 0.09 | 10 | Endothelial Cells |
| MESDC1 | 0.58 | 0 | 0 | 0.46 | 0.15 | 10 | Endothelial Cells |
| CD36 | 0.58 | 0 | 0 | 0.51 | 0.15 | 10 | Endothelial Cells |
| MYL12B | 0.58 | 0 | 0 | 0.91 | 0.77 | 10 | Endothelial Cells |
| FLI1 | 0.58 | 0 | 0 | 0.46 | 0.11 | 10 | Endothelial Cells |
| PRSS23 | 0.57 | 0 | 0 | 0.56 | 0.24 | 10 | Endothelial Cells |
| CHSY1 | 0.57 | 0 | 0 | 0.34 | 0.07 | 10 | Endothelial Cells |
| PLA2G16 | 0.57 | 0 | 0 | 0.40 | 0.10 | 10 | Endothelial Cells |
| STOM | 0.57 | 0 | 0 | 0.76 | 0.44 | 10 | Endothelial Cells |
| C1orf54 | 0.57 | 0 | 0 | 0.49 | 0.18 | 10 | Endothelial Cells |
| FAM129B | 0.57 | 0 | 0 | 0.37 | 0.10 | 10 | Endothelial Cells |
| ADGRG6 | 0.57 | 0 | 0 | 0.41 | 0.12 | 10 | Endothelial Cells |
| RBP5 | 0.56 | 0 | 0 | 0.28 | 0.01 | 10 | Endothelial Cells |
| SORBS2 | 0.56 | 0 | 0 | 0.30 | 0.07 | 10 | Endothelial Cells |
| ABI3BP | 0.56 | 0 | 0 | 0.29 | 0.08 | 10 | Endothelial Cells |
| HSPB1 | 0.56 | 0 | 0 | 0.95 | 0.76 | 10 | Endothelial Cells |
| GNAQ | 0.56 | 0 | 0 | 0.45 | 0.16 | 10 | Endothelial Cells |
| BMPR2 | 0.56 | 0 | 0 | 0.56 | 0.21 | 10 | Endothelial Cells |
| LARP6 | 0.56 | 0 | 0 | 0.36 | 0.11 | 10 | Endothelial Cells |
| TUBB | 0.56 | 0 | 0 | 0.92 | 0.79 | 10 | Endothelial Cells |
| FAM155A | 0.55 | 0 | 0 | 0.33 | 0.05 | 10 | Endothelial Cells |
| FDPS | 0.55 | 0 | 0 | 0.69 | 0.34 | 10 | Endothelial Cells |
| EHBP1L1 | 0.55 | 0 | 0 | 0.33 | 0.06 | 10 | Endothelial Cells |
| PTRF | 0.55 | 0 | 0 | 0.95 | 0.69 | 10 | Endothelial Cells |
| SNCG | 0.55 | 0 | 0 | 0.47 | 0.11 | 10 | Endothelial Cells |
| GNAI2 | 0.54 | 0 | 0 | 0.65 | 0.34 | 10 | Endothelial Cells |
| KLF4 | 0.54 | 0 | 0 | 0.37 | 0.16 | 10 | Endothelial Cells |
| TNFRSF1A | 0.54 | 0 | 0 | 0.62 | 0.38 | 10 | Endothelial Cells |
| APOL1 | 0.54 | 0 | 0 | 0.42 | 0.08 | 10 | Endothelial Cells |
| SPTAN1 | 0.54 | 0 | 0 | 0.48 | 0.22 | 10 | Endothelial Cells |
| C10orf11 | 0.54 | 0 | 0 | 0.26 | 0.07 | 10 | Endothelial Cells |
| PODXL | 0.54 | 0 | 0 | 0.36 | 0.09 | 10 | Endothelial Cells |
| RAI14 | 0.54 | 0 | 0 | 0.36 | 0.10 | 10 | Endothelial Cells |
| F2R | 0.54 | 0 | 0 | 0.34 | 0.10 | 10 | Endothelial Cells |
| PEA15 | 0.53 | 0 | 0 | 0.55 | 0.26 | 10 | Endothelial Cells |
| CCND3 | 0.53 | 0 | 0 | 0.45 | 0.15 | 10 | Endothelial Cells |
| MMRN1 | 0.53 | 0 | 0 | 0.27 | 0.01 | 10 | Endothelial Cells |
| PPP3CA | 0.53 | 0 | 0 | 0.45 | 0.19 | 10 | Endothelial Cells |
| MLLT4 | 0.53 | 0 | 0 | 0.48 | 0.16 | 10 | Endothelial Cells |
| CXorf36 | 0.53 | 0 | 0 | 0.40 | 0.07 | 10 | Endothelial Cells |
| ERG | 0.53 | 0 | 0 | 0.41 | 0.11 | 10 | Endothelial Cells |
| COX17 | 0.53 | 0 | 0 | 0.61 | 0.35 | 10 | Endothelial Cells |
| CAPNS1 | 0.53 | 0 | 0 | 0.56 | 0.29 | 10 | Endothelial Cells |
| KANK3 | 0.53 | 0 | 0 | 0.45 | 0.14 | 10 | Endothelial Cells |
| ETS2 | 0.52 | 0 | 0 | 0.58 | 0.27 | 10 | Endothelial Cells |
| SCARF1 | 0.52 | 0 | 0 | 0.33 | 0.08 | 10 | Endothelial Cells |
| HSPA5 | 0.52 | 0 | 0 | 0.72 | 0.49 | 10 | Endothelial Cells |
| EIF2S2 | 0.52 | 0 | 0 | 0.70 | 0.47 | 10 | Endothelial Cells |
| ITGB4 | 0.52 | 0 | 0 | 0.27 | 0.01 | 10 | Endothelial Cells |
| ACVRL1 | 0.51 | 0 | 0 | 0.55 | 0.16 | 10 | Endothelial Cells |
| HS3ST1 | 0.51 | 0 | 0 | 0.25 | 0.05 | 10 | Endothelial Cells |
| NOP10 | 0.51 | 0 | 0 | 0.77 | 0.48 | 10 | Endothelial Cells |
| MYCT1 | 0.51 | 0 | 0 | 0.31 | 0.06 | 10 | Endothelial Cells |
| FABP5 | 0.51 | 0 | 0 | 0.59 | 0.26 | 10 | Endothelial Cells |
| ARHGAP23 | 0.51 | 0 | 0 | 0.27 | 0.03 | 10 | Endothelial Cells |
| FAM171A1 | 0.5 | 0 | 0 | 0.30 | 0.06 | 10 | Endothelial Cells |
| PLK2 | 0.5 | 0 | 0 | 0.47 | 0.20 | 10 | Endothelial Cells |
| CASKIN2 | 0.5 | 0 | 0 | 0.38 | 0.09 | 10 | Endothelial Cells |
| LRRC32 | 0.5 | 0 | 0 | 0.44 | 0.12 | 10 | Endothelial Cells |
| NECTIN2 | 0.5 | 0 | 0 | 0.50 | 0.24 | 10 | Endothelial Cells |
| COL9A3 | 0.5 | 0 | 0 | 0.30 | 0.02 | 10 | Endothelial Cells |
| IFI44L | 0.5 | 0 | 0 | 0.29 | 0.04 | 10 | Endothelial Cells |
| ABI3 | 0.5 | 0 | 0 | 0.35 | 0.09 | 10 | Endothelial Cells |
| KPNA2 | 0.5 | 0 | 0 | 0.30 | 0.08 | 10 | Endothelial Cells |
| TXN | 0.5 | 0 | 0 | 0.76 | 0.55 | 10 | Endothelial Cells |
| RAPGEF5 | 0.5 | 0 | 0 | 0.39 | 0.13 | 10 | Endothelial Cells |
| GNG5 | 0.5 | 0 | 0 | 0.76 | 0.52 | 10 | Endothelial Cells |
| B2M | 0.5 | 0 | 0 | 1.00 | 0.99 | 10 | Endothelial Cells |
| POSTN | 0.5 | 0 | 0 | 0.50 | 0.21 | 10 | Endothelial Cells |
| RASIP1 | 0.5 | 0 | 0 | 0.38 | 0.09 | 10 | Endothelial Cells |
| HSPA12B | 0.49 | 0 | 0 | 0.38 | 0.09 | 10 | Endothelial Cells |
| PIM3 | 0.49 | 0 | 0 | 0.38 | 0.17 | 10 | Endothelial Cells |
| INO80C | 0.49 | 0 | 0 | 0.34 | 0.10 | 10 | Endothelial Cells |
| TSPAN9 | 0.49 | 0 | 0 | 0.36 | 0.10 | 10 | Endothelial Cells |
| PLSCR1 | 0.49 | 0 | 0 | 0.45 | 0.17 | 10 | Endothelial Cells |
| WNK1 | 0.49 | 0 | 0 | 0.36 | 0.14 | 10 | Endothelial Cells |
| IFI16 | 0.49 | 0 | 0 | 0.56 | 0.34 | 10 | Endothelial Cells |
| SULF2 | 0.49 | 0 | 0 | 0.31 | 0.04 | 10 | Endothelial Cells |
| CDH5 | 0.49 | 0 | 0 | 0.62 | 0.18 | 10 | Endothelial Cells |
| MYH9 | 0.49 | 0 | 0 | 0.73 | 0.44 | 10 | Endothelial Cells |
| HHEX | 0.49 | 0 | 0 | 0.35 | 0.09 | 10 | Endothelial Cells |
| BCAM | 0.48 | 0 | 0 | 0.71 | 0.33 | 10 | Endothelial Cells |
| SHE | 0.48 | 0 | 0 | 0.33 | 0.05 | 10 | Endothelial Cells |
| MSN | 0.48 | 0 | 0 | 0.69 | 0.46 | 10 | Endothelial Cells |
| RAC1 | 0.48 | 0 | 0 | 0.92 | 0.81 | 10 | Endothelial Cells |
| GIMAP4 | 0.48 | 0 | 0 | 0.55 | 0.18 | 10 | Endothelial Cells |
| TMEM88 | 0.48 | 0 | 0 | 0.35 | 0.08 | 10 | Endothelial Cells |
| CD47 | 0.48 | 0 | 0 | 0.51 | 0.21 | 10 | Endothelial Cells |
| ITGA3 | 0.48 | 0 | 0 | 0.39 | 0.10 | 10 | Endothelial Cells |
| TNXB | 0.48 | 0 | 0 | 0.32 | 0.08 | 10 | Endothelial Cells |
| LGALS9 | 0.47 | 0 | 0 | 0.37 | 0.10 | 10 | Endothelial Cells |
| DDIT4 | 0.47 | 0 | 0 | 0.38 | 0.16 | 10 | Endothelial Cells |
| CLSTN3 | 0.47 | 0 | 0 | 0.27 | 0.03 | 10 | Endothelial Cells |
| SERPINH1 | 0.47 | 0 | 0 | 0.82 | 0.57 | 10 | Endothelial Cells |
| PARVB | 0.47 | 0 | 0 | 0.41 | 0.15 | 10 | Endothelial Cells |
| C1orf115 | 0.47 | 0 | 0 | 0.31 | 0.05 | 10 | Endothelial Cells |
| ITGA6 | 0.47 | 0 | 0 | 0.32 | 0.07 | 10 | Endothelial Cells |
| DNAJA1 | 0.47 | 0 | 0 | 0.67 | 0.48 | 10 | Endothelial Cells |
| DYNLL1 | 0.47 | 0 | 0 | 0.89 | 0.79 | 10 | Endothelial Cells |
| SOX7 | 0.46 | 0 | 0 | 0.36 | 0.08 | 10 | Endothelial Cells |
| VAMP5 | 0.46 | 0 | 0 | 0.82 | 0.51 | 10 | Endothelial Cells |
| PDLIM4 | 0.46 | 0 | 0 | 0.34 | 0.10 | 10 | Endothelial Cells |
| NEAT1 | 0.46 | 0 | 0 | 0.84 | 0.63 | 10 | Endothelial Cells |
| SERTAD4-AS1 | 0.46 | 0 | 0 | 0.27 | 0.05 | 10 | Endothelial Cells |
| RALGAPA2 | 0.46 | 0 | 0 | 0.28 | 0.05 | 10 | Endothelial Cells |
| SDPR | 0.46 | 0 | 0 | 0.64 | 0.24 | 10 | Endothelial Cells |
| HDAC1 | 0.46 | 0 | 0 | 0.37 | 0.17 | 10 | Endothelial Cells |
| S100A6 | 0.46 | 0 | 0 | 0.99 | 0.84 | 10 | Endothelial Cells |
| SH3BGRL3 | 0.46 | 0 | 0 | 0.88 | 0.58 | 10 | Endothelial Cells |
| AHR | 0.46 | 0 | 0 | 0.36 | 0.12 | 10 | Endothelial Cells |
| SLC3A2 | 0.46 | 0 | 0 | 0.49 | 0.23 | 10 | Endothelial Cells |
| TIE1 | 0.45 | 0 | 0 | 0.47 | 0.14 | 10 | Endothelial Cells |
| PDLIM1 | 0.45 | 0 | 0 | 0.56 | 0.39 | 10 | Endothelial Cells |
| GIMAP6 | 0.45 | 0 | 0 | 0.34 | 0.08 | 10 | Endothelial Cells |
| ADAMTS6 | 0.45 | 0 | 0 | 0.31 | 0.12 | 10 | Endothelial Cells |
| INPP1 | 0.45 | 0 | 0 | 0.35 | 0.10 | 10 | Endothelial Cells |
| RASSF9 | 0.45 | 0 | 0 | 0.25 | 0.02 | 10 | Endothelial Cells |
| PCDH12 | 0.45 | 0 | 0 | 0.34 | 0.09 | 10 | Endothelial Cells |
| SHISA5 | 0.45 | 0 | 0 | 0.52 | 0.26 | 10 | Endothelial Cells |
| NAA10 | 0.44 | 0 | 0 | 0.51 | 0.30 | 10 | Endothelial Cells |
| GRN | 0.44 | 0 | 0 | 0.69 | 0.40 | 10 | Endothelial Cells |
| ACTN4 | 0.44 | 0 | 0 | 0.64 | 0.40 | 10 | Endothelial Cells |
| PLEC | 0.43 | 0 | 0 | 0.40 | 0.16 | 10 | Endothelial Cells |
| FOXO1 | 0.43 | 0 | 0 | 0.27 | 0.06 | 10 | Endothelial Cells |
| PON2 | 0.43 | 0 | 0 | 0.53 | 0.24 | 10 | Endothelial Cells |
| B4GALT5 | 0.43 | 0 | 0 | 0.27 | 0.05 | 10 | Endothelial Cells |
| TMEM176A | 0.43 | 0 | 0 | 0.36 | 0.12 | 10 | Endothelial Cells |
| MAP3K11 | 0.42 | 0 | 0 | 0.41 | 0.15 | 10 | Endothelial Cells |
| RP4-725G10.3 | 0.42 | 0 | 0 | 0.27 | 0.05 | 10 | Endothelial Cells |
| STMN1 | 0.42 | 0 | 0 | 0.75 | 0.52 | 10 | Endothelial Cells |
| TMBIM1 | 0.42 | 0 | 0 | 0.46 | 0.24 | 10 | Endothelial Cells |
| FIS1 | 0.42 | 0 | 0 | 0.74 | 0.50 | 10 | Endothelial Cells |
| SEMA6A | 0.42 | 0 | 0 | 0.34 | 0.07 | 10 | Endothelial Cells |
| SPTBN1 | 0.42 | 0 | 0 | 0.69 | 0.42 | 10 | Endothelial Cells |
| ACTN1 | 0.42 | 0 | 0 | 0.71 | 0.44 | 10 | Endothelial Cells |
| RRAS | 0.42 | 0 | 0 | 0.40 | 0.15 | 10 | Endothelial Cells |
| SLC9A3R2 | 0.42 | 0 | 0 | 0.72 | 0.40 | 10 | Endothelial Cells |
| YES1 | 0.42 | 0 | 0 | 0.36 | 0.11 | 10 | Endothelial Cells |
| DLL1 | 0.42 | 0 | 0 | 0.25 | 0.04 | 10 | Endothelial Cells |
| IFI6 | 0.41 | 0 | 0 | 0.52 | 0.26 | 10 | Endothelial Cells |
| MAGEH1 | 0.41 | 0 | 0 | 0.51 | 0.27 | 10 | Endothelial Cells |
| GAPDH | 0.41 | 0 | 0 | 0.96 | 0.89 | 10 | Endothelial Cells |
| POLR2L | 0.41 | 0 | 0 | 0.80 | 0.67 | 10 | Endothelial Cells |
| HSPA1A | 0.41 | 0 | 0 | 0.79 | 0.64 | 10 | Endothelial Cells |
| SMAGP | 0.41 | 0 | 0 | 0.32 | 0.07 | 10 | Endothelial Cells |
| VAT1 | 0.41 | 0 | 0 | 0.41 | 0.17 | 10 | Endothelial Cells |
| LSR | 0.41 | 0 | 0 | 0.34 | 0.10 | 10 | Endothelial Cells |
| HSPB8 | 0.41 | 0 | 0 | 0.32 | 0.11 | 10 | Endothelial Cells |
| TRIB2 | 0.4 | 0 | 0 | 0.30 | 0.10 | 10 | Endothelial Cells |
| PRKCH | 0.4 | 0 | 0 | 0.34 | 0.09 | 10 | Endothelial Cells |
| PRDX1 | 0.4 | 0 | 0 | 0.77 | 0.66 | 10 | Endothelial Cells |
| AGRN | 0.4 | 0 | 0 | 0.28 | 0.07 | 10 | Endothelial Cells |
| PALM | 0.4 | 0 | 0 | 0.35 | 0.11 | 10 | Endothelial Cells |
| DPYSL3 | 0.4 | 0 | 0 | 0.36 | 0.15 | 10 | Endothelial Cells |
| MAPK3 | 0.4 | 0 | 0 | 0.40 | 0.16 | 10 | Endothelial Cells |
| LMO2 | 0.4 | 0 | 0 | 0.33 | 0.09 | 10 | Endothelial Cells |
| CANX | 0.4 | 0 | 0 | 0.63 | 0.40 | 10 | Endothelial Cells |
| LAMB2 | 0.39 | 0 | 0 | 0.39 | 0.20 | 10 | Endothelial Cells |
| PDIA3 | 0.39 | 0 | 0 | 0.74 | 0.58 | 10 | Endothelial Cells |
| NUCB2 | 0.39 | 0 | 0 | 0.45 | 0.25 | 10 | Endothelial Cells |
| DLC1 | 0.39 | 0 | 0 | 0.43 | 0.23 | 10 | Endothelial Cells |
| CTNND1 | 0.39 | 0 | 0 | 0.51 | 0.25 | 10 | Endothelial Cells |
| NRP1 | 0.39 | 0 | 0 | 0.47 | 0.20 | 10 | Endothelial Cells |
| IGF2BP2 | 0.39 | 0 | 0 | 0.37 | 0.15 | 10 | Endothelial Cells |
| RCAN1 | 0.38 | 0 | 0 | 0.28 | 0.12 | 10 | Endothelial Cells |
| IL6ST | 0.38 | 0 | 0 | 0.46 | 0.26 | 10 | Endothelial Cells |
| PSMB9 | 0.38 | 0 | 0 | 0.53 | 0.21 | 10 | Endothelial Cells |
| EGLN2 | 0.38 | 0 | 0 | 0.41 | 0.19 | 10 | Endothelial Cells |
| MESDC2 | 0.38 | 0 | 0 | 0.49 | 0.27 | 10 | Endothelial Cells |
| TMEM2 | 0.38 | 0 | 0 | 0.25 | 0.08 | 10 | Endothelial Cells |
| CFLAR | 0.38 | 0 | 0 | 0.45 | 0.26 | 10 | Endothelial Cells |
| YWHAE | 0.38 | 0 | 0 | 0.82 | 0.63 | 10 | Endothelial Cells |
| APP | 0.38 | 0 | 0 | 0.77 | 0.51 | 10 | Endothelial Cells |
| SPTLC2 | 0.38 | 0 | 0 | 0.28 | 0.09 | 10 | Endothelial Cells |
| GADD45GIP1 | 0.37 | 0 | 0 | 0.73 | 0.54 | 10 | Endothelial Cells |
| PPP2R5A | 0.37 | 0 | 0 | 0.29 | 0.10 | 10 | Endothelial Cells |
| RCN1 | 0.37 | 0 | 0.01 | 0.58 | 0.38 | 10 | Endothelial Cells |
| HLA-C | 0.37 | 0 | 0 | 0.90 | 0.54 | 10 | Endothelial Cells |
| SHC1 | 0.37 | 0 | 0 | 0.29 | 0.10 | 10 | Endothelial Cells |
| CARD16 | 0.37 | 0 | 0 | 0.58 | 0.28 | 10 | Endothelial Cells |
| SEC61G | 0.37 | 0 | 0 | 0.86 | 0.68 | 10 | Endothelial Cells |
| ATOX1 | 0.37 | 0 | 0 | 0.60 | 0.40 | 10 | Endothelial Cells |
| HN1 | 0.37 | 0 | 0 | 0.38 | 0.19 | 10 | Endothelial Cells |
| ATP5E | 0.36 | 0 | 0 | 0.99 | 0.87 | 10 | Endothelial Cells |
| ANXA1 | 0.36 | 0 | 0 | 0.73 | 0.40 | 10 | Endothelial Cells |
| MDK | 0.36 | 0 | 0 | 0.75 | 0.53 | 10 | Endothelial Cells |
| EGLN3 | 0.36 | 0 | 0 | 0.30 | 0.08 | 10 | Endothelial Cells |
| TMEM179B | 0.36 | 0 | 0 | 0.51 | 0.27 | 10 | Endothelial Cells |
| CLEC3B | 0.36 | 0 | 0 | 0.27 | 0.05 | 10 | Endothelial Cells |
| SNRPG | 0.36 | 0 | 0 | 0.70 | 0.53 | 10 | Endothelial Cells |
| KRAS | 0.36 | 0 | 0 | 0.33 | 0.14 | 10 | Endothelial Cells |
| CCDC85B | 0.35 | 0 | 0 | 0.67 | 0.41 | 10 | Endothelial Cells |
| GRAMD1A | 0.35 | 0 | 0 | 0.34 | 0.14 | 10 | Endothelial Cells |
| FAM65A | 0.35 | 0 | 0 | 0.30 | 0.09 | 10 | Endothelial Cells |
| PSMD14 | 0.35 | 0 | 0 | 0.34 | 0.17 | 10 | Endothelial Cells |
| DOCK6 | 0.35 | 0 | 0 | 0.32 | 0.15 | 10 | Endothelial Cells |
| FNDC3B | 0.35 | 0 | 0 | 0.36 | 0.14 | 10 | Endothelial Cells |
| NPR3 | 0.35 | 0 | 0 | 0.29 | 0.09 | 10 | Endothelial Cells |
| NUCB1 | 0.35 | 0 | 0.01 | 0.60 | 0.42 | 10 | Endothelial Cells |
| TTYH3 | 0.35 | 0 | 0 | 0.27 | 0.07 | 10 | Endothelial Cells |
| GIMAP1 | 0.35 | 0 | 0 | 0.34 | 0.10 | 10 | Endothelial Cells |
| PDIA4 | 0.35 | 0 | 0.01 | 0.47 | 0.28 | 10 | Endothelial Cells |
| NXN | 0.35 | 0 | 0 | 0.28 | 0.07 | 10 | Endothelial Cells |
| PIK3C2A | 0.35 | 0 | 0 | 0.26 | 0.09 | 10 | Endothelial Cells |
| RPS28 | 0.35 | 0 | 0 | 1.00 | 0.99 | 10 | Endothelial Cells |
| SERINC3 | 0.35 | 0 | 0 | 0.43 | 0.24 | 10 | Endothelial Cells |
| CTNNA1 | 0.35 | 0 | 0 | 0.55 | 0.34 | 10 | Endothelial Cells |
| GNG11 | 0.35 | 0 | 0 | 0.82 | 0.44 | 10 | Endothelial Cells |
| CLIC2 | 0.35 | 0 | 0 | 0.25 | 0.05 | 10 | Endothelial Cells |
| GOSR2 | 0.34 | 0 | 0 | 0.26 | 0.08 | 10 | Endothelial Cells |
| TUBB4B | 0.34 | 0 | 0 | 0.60 | 0.40 | 10 | Endothelial Cells |
| NPC2 | 0.34 | 0 | 0 | 0.69 | 0.50 | 10 | Endothelial Cells |
| KRTCAP2 | 0.34 | 0 | 0 | 0.77 | 0.54 | 10 | Endothelial Cells |
| RRBP1 | 0.34 | 0 | 0 | 0.54 | 0.32 | 10 | Endothelial Cells |
| HLA-E | 0.34 | 0 | 0 | 0.96 | 0.59 | 10 | Endothelial Cells |
| TMSB4X | 0.34 | 0 | 0 | 0.99 | 1.00 | 10 | Endothelial Cells |
| VWA1 | 0.34 | 0 | 0 | 0.29 | 0.09 | 10 | Endothelial Cells |
| NME1 | 0.34 | 0 | 0 | 0.47 | 0.25 | 10 | Endothelial Cells |
| TFPI | 0.34 | 0 | 0 | 0.66 | 0.43 | 10 | Endothelial Cells |
| FLYWCH2 | 0.34 | 0 | 0 | 0.36 | 0.16 | 10 | Endothelial Cells |
| PPIA | 0.34 | 0 | 0 | 0.95 | 0.84 | 10 | Endothelial Cells |
| SH2D3C | 0.34 | 0 | 0 | 0.29 | 0.12 | 10 | Endothelial Cells |
| PTPRM | 0.33 | 0 | 0 | 0.30 | 0.08 | 10 | Endothelial Cells |
| NMI | 0.33 | 0 | 0 | 0.34 | 0.12 | 10 | Endothelial Cells |
| G6PC3 | 0.33 | 0 | 0 | 0.29 | 0.11 | 10 | Endothelial Cells |
| MGST2 | 0.33 | 0 | 0 | 0.47 | 0.26 | 10 | Endothelial Cells |
| THSD1 | 0.33 | 0 | 0 | 0.28 | 0.09 | 10 | Endothelial Cells |
| BCAP31 | 0.33 | 0 | 0 | 0.59 | 0.37 | 10 | Endothelial Cells |
| TSPAN18 | 0.33 | 0 | 0 | 0.25 | 0.08 | 10 | Endothelial Cells |
| NKTR | 0.33 | 0 | 0 | 0.62 | 0.39 | 10 | Endothelial Cells |
| ZMIZ1 | 0.33 | 0 | 0 | 0.26 | 0.09 | 10 | Endothelial Cells |
| ID3 | 0.33 | 0 | 0.02 | 0.75 | 0.54 | 10 | Endothelial Cells |
| LDHA | 0.33 | 0 | 0 | 0.73 | 0.58 | 10 | Endothelial Cells |
| TNKS1BP1 | 0.33 | 0 | 0 | 0.27 | 0.08 | 10 | Endothelial Cells |
| C16orf45 | 0.33 | 0 | 0 | 0.32 | 0.14 | 10 | Endothelial Cells |
| ARRB1 | 0.33 | 0 | 0 | 0.27 | 0.08 | 10 | Endothelial Cells |
| ADAR | 0.32 | 0 | 0 | 0.44 | 0.23 | 10 | Endothelial Cells |
| AP2S1 | 0.32 | 0 | 0.01 | 0.63 | 0.45 | 10 | Endothelial Cells |
| PSMA4 | 0.32 | 0 | 0.01 | 0.77 | 0.58 | 10 | Endothelial Cells |
| ITM2A | 0.32 | 0 | 0 | 0.59 | 0.18 | 10 | Endothelial Cells |
| LYST | 0.32 | 0 | 0 | 0.27 | 0.07 | 10 | Endothelial Cells |
| KIAA0355 | 0.32 | 0 | 0 | 0.38 | 0.17 | 10 | Endothelial Cells |
| PTPRG | 0.32 | 0 | 0 | 0.30 | 0.10 | 10 | Endothelial Cells |
| JAG1 | 0.32 | 0 | 0 | 0.26 | 0.11 | 10 | Endothelial Cells |
| APOL3 | 0.32 | 0 | 0 | 0.34 | 0.12 | 10 | Endothelial Cells |
| PDLIM5 | 0.32 | 0 | 0 | 0.32 | 0.16 | 10 | Endothelial Cells |
| ZNF521 | 0.31 | 0 | 0 | 0.27 | 0.07 | 10 | Endothelial Cells |
| NFE2L1 | 0.31 | 0 | 0 | 0.32 | 0.13 | 10 | Endothelial Cells |
| HDAC7 | 0.31 | 0 | 0 | 0.34 | 0.16 | 10 | Endothelial Cells |
| MRPL57 | 0.31 | 0 | 0.01 | 0.56 | 0.35 | 10 | Endothelial Cells |
| FAM89B | 0.31 | 0 | 0.01 | 0.31 | 0.15 | 10 | Endothelial Cells |
| SERF2 | 0.31 | 0 | 0 | 0.96 | 0.95 | 10 | Endothelial Cells |
| TAP1 | 0.31 | 0 | 0 | 0.32 | 0.12 | 10 | Endothelial Cells |
| SEC11C | 0.31 | 0 | 0 | 0.34 | 0.12 | 10 | Endothelial Cells |
| PSME2 | 0.31 | 0 | 0 | 0.67 | 0.45 | 10 | Endothelial Cells |
| CFL1 | 0.31 | 0 | 0 | 0.97 | 0.91 | 10 | Endothelial Cells |
| ANGPTL4 | 0.31 | 0 | 0 | 0.31 | 0.14 | 10 | Endothelial Cells |
| ELOVL1 | 0.3 | 0 | 0 | 0.33 | 0.14 | 10 | Endothelial Cells |
| TNIP1 | 0.3 | 0 | 0.02 | 0.36 | 0.19 | 10 | Endothelial Cells |
| TNFAIP1 | 0.3 | 0 | 0 | 0.29 | 0.09 | 10 | Endothelial Cells |
| LINC00116 | 0.3 | 0 | 0 | 0.35 | 0.17 | 10 | Endothelial Cells |
| TSPAN14 | 0.3 | 0 | 0 | 0.33 | 0.14 | 10 | Endothelial Cells |
| DCTPP1 | 0.3 | 0 | 0 | 0.32 | 0.15 | 10 | Endothelial Cells |
| NPNT | 0.3 | 0 | 0 | 0.30 | 0.10 | 10 | Endothelial Cells |
| WWTR1 | 0.3 | 0 | 0 | 0.38 | 0.20 | 10 | Endothelial Cells |
| MCAM | 0.29 | 0 | 0 | 0.43 | 0.23 | 10 | Endothelial Cells |
| EIF1 | 0.29 | 0 | 0 | 0.97 | 0.96 | 10 | Endothelial Cells |
| PSMB8 | 0.29 | 0 | 0 | 0.49 | 0.26 | 10 | Endothelial Cells |
| C10orf54 | 0.29 | 0 | 0 | 0.42 | 0.20 | 10 | Endothelial Cells |
| RPN2 | 0.29 | 0 | 0.01 | 0.57 | 0.38 | 10 | Endothelial Cells |
| PEAK1 | 0.29 | 0 | 0 | 0.26 | 0.10 | 10 | Endothelial Cells |
| YBX3 | 0.29 | 0 | 0.03 | 0.60 | 0.40 | 10 | Endothelial Cells |
| PXN | 0.29 | 0 | 0 | 0.30 | 0.10 | 10 | Endothelial Cells |
| BOLA3 | 0.29 | 0 | 0 | 0.30 | 0.14 | 10 | Endothelial Cells |
| SERPINE2 | 0.28 | 0 | 0 | 0.45 | 0.17 | 10 | Endothelial Cells |
| IGFBP7 | 0.28 | 0 | 0 | 0.90 | 0.67 | 10 | Endothelial Cells |
| PSME2P2 | 0.28 | 0 | 0 | 0.40 | 0.21 | 10 | Endothelial Cells |
| SDF2L1 | 0.28 | 0 | 0.01 | 0.40 | 0.23 | 10 | Endothelial Cells |
| VAMP3 | 0.28 | 0 | 0.01 | 0.45 | 0.26 | 10 | Endothelial Cells |
| SNX3 | 0.28 | 0 | 0 | 0.80 | 0.69 | 10 | Endothelial Cells |
| IER5 | 0.28 | 0 | 0 | 0.27 | 0.11 | 10 | Endothelial Cells |
| CDV3 | 0.28 | 0 | 0.02 | 0.31 | 0.16 | 10 | Endothelial Cells |
| PRAF2 | 0.28 | 0 | 0 | 0.36 | 0.17 | 10 | Endothelial Cells |
| XIST | 0.28 | 0 | 0.01 | 0.84 | 0.70 | 10 | Endothelial Cells |
| AC012354.8 | 0.28 | 0 | 0.04 | 0.36 | 0.20 | 10 | Endothelial Cells |
| DNAJB11 | 0.28 | 0 | 0.02 | 0.40 | 0.23 | 10 | Endothelial Cells |
| PSMA7 | 0.28 | 0 | 0 | 0.86 | 0.71 | 10 | Endothelial Cells |
| RNF181 | 0.27 | 0 | 0 | 0.62 | 0.39 | 10 | Endothelial Cells |
| RPL36 | 0.27 | 0 | 0 | 1.00 | 0.98 | 10 | Endothelial Cells |
| LUZP1 | 0.27 | 0 | 0 | 0.36 | 0.18 | 10 | Endothelial Cells |
| FYN | 0.27 | 0 | 0 | 0.30 | 0.13 | 10 | Endothelial Cells |
| COX16 | 0.27 | 0 | 0 | 0.35 | 0.16 | 10 | Endothelial Cells |
| PRKD2 | 0.27 | 0 | 0 | 0.26 | 0.08 | 10 | Endothelial Cells |
| MVP | 0.27 | 0 | 0 | 0.32 | 0.14 | 10 | Endothelial Cells |
| ITGB1 | 0.27 | 0 | 0.04 | 0.79 | 0.63 | 10 | Endothelial Cells |
| PPDPF | 0.26 | 0 | 0.03 | 0.88 | 0.80 | 10 | Endothelial Cells |
| HSPA8 | 0.26 | 0 | 0.03 | 0.77 | 0.59 | 10 | Endothelial Cells |
| ORAI1 | 0.26 | 0 | 0 | 0.25 | 0.06 | 10 | Endothelial Cells |
| ZEB1 | 0.26 | 0 | 0 | 0.33 | 0.15 | 10 | Endothelial Cells |
| RBX1 | 0.26 | 0 | 0 | 0.77 | 0.59 | 10 | Endothelial Cells |
| RP11-220D10.1 | 0.26 | 0 | 0 | 0.27 | 0.11 | 10 | Endothelial Cells |
| TMEM9B | 0.26 | 0 | 0 | 0.36 | 0.18 | 10 | Endothelial Cells |
| SWAP70 | 0.26 | 0 | 0 | 0.27 | 0.10 | 10 | Endothelial Cells |
| HLA-F | 0.25 | 0 | 0 | 0.36 | 0.13 | 10 | Endothelial Cells |
| ITSN2 | 0.25 | 0 | 0.02 | 0.32 | 0.16 | 10 | Endothelial Cells |
| OST4 | 0.25 | 0 | 0 | 0.90 | 0.78 | 10 | Endothelial Cells |
| ABHD17A | 0.25 | 0 | 0 | 0.33 | 0.16 | 10 | Endothelial Cells |
| PAK4 | 0.25 | 0 | 0 | 0.25 | 0.10 | 10 | Endothelial Cells |
| CCL21 | 3.84 | 0 | 0 | 0.31 | 0.07 | 11 | AT1 Epithelial Cells |
| TFF3 | 2.03 | 0 | 0 | 0.34 | 0.01 | 11 | AT1 Epithelial Cells |
| SFTPB | 2.01 | 0 | 0 | 0.27 | 0.08 | 11 | AT1 Epithelial Cells |
| AGR3 | 1.78 | 0 | 0 | 0.31 | 0.02 | 11 | AT1 Epithelial Cells |
| AGER | 1.77 | 0 | 0 | 0.26 | 0.05 | 11 | AT1 Epithelial Cells |
| ELF3 | 1.58 | 0 | 0 | 0.40 | 0.02 | 11 | AT1 Epithelial Cells |
| AGR2 | 1.41 | 0 | 0 | 0.31 | 0.02 | 11 | AT1 Epithelial Cells |
| KRT19 | 1.38 | 0 | 0 | 0.34 | 0.03 | 11 | AT1 Epithelial Cells |
| KRT8 | 1.38 | 0 | 0 | 0.41 | 0.07 | 11 | AT1 Epithelial Cells |
| FXYD3 | 1.36 | 0 | 0 | 0.38 | 0.02 | 11 | AT1 Epithelial Cells |
| GPRC5A | 1.28 | 0 | 0 | 0.28 | 0.03 | 11 | AT1 Epithelial Cells |
| KRT18 | 1.17 | 0 | 0 | 0.47 | 0.12 | 11 | AT1 Epithelial Cells |
| FOLR1 | 1.05 | 0 | 0 | 0.31 | 0.04 | 11 | AT1 Epithelial Cells |
| TUBB4B | 1.03 | 0 | 0 | 0.63 | 0.40 | 11 | AT1 Epithelial Cells |
| ANXA2 | 1.03 | 0 | 0 | 0.78 | 0.65 | 11 | AT1 Epithelial Cells |
| LMO7 | 1.02 | 0 | 0 | 0.30 | 0.08 | 11 | AT1 Epithelial Cells |
| S100A10 | 0.98 | 0 | 0 | 0.79 | 0.65 | 11 | AT1 Epithelial Cells |
| CLDN4 | 0.98 | 0 | 0 | 0.34 | 0.02 | 11 | AT1 Epithelial Cells |
| DMKN | 0.97 | 0 | 0 | 0.34 | 0.03 | 11 | AT1 Epithelial Cells |
| NKX2-1 | 0.97 | 0 | 0 | 0.31 | 0.02 | 11 | AT1 Epithelial Cells |
| C19orf33 | 0.95 | 0 | 0 | 0.28 | 0.03 | 11 | AT1 Epithelial Cells |
| CD24 | 0.91 | 0 | 0 | 0.26 | 0.02 | 11 | AT1 Epithelial Cells |
| EFEMP1 | 0.9 | 0 | 0 | 0.42 | 0.07 | 11 | AT1 Epithelial Cells |
| TACSTD2 | 0.89 | 0 | 0 | 0.28 | 0.03 | 11 | AT1 Epithelial Cells |
| AQP4 | 0.88 | 0 | 0 | 0.27 | 0.02 | 11 | AT1 Epithelial Cells |
| MT-CO2 | 0.87 | 0 | 0 | 0.95 | 0.99 | 11 | AT1 Epithelial Cells |
| SDC4 | 0.86 | 0 | 0 | 0.36 | 0.04 | 11 | AT1 Epithelial Cells |
| CPM | 0.86 | 0 | 0 | 0.28 | 0.04 | 11 | AT1 Epithelial Cells |
| IFT57 | 0.85 | 0 | 0 | 0.46 | 0.21 | 11 | AT1 Epithelial Cells |
| CD9 | 0.85 | 0 | 0 | 0.69 | 0.45 | 11 | AT1 Epithelial Cells |
| CTSH | 0.84 | 0 | 0 | 0.49 | 0.13 | 11 | AT1 Epithelial Cells |
| EPCAM | 0.84 | 0 | 0 | 0.30 | 0.02 | 11 | AT1 Epithelial Cells |
| MORN2 | 0.83 | 0 | 0 | 0.30 | 0.07 | 11 | AT1 Epithelial Cells |
| GJA1 | 0.82 | 0 | 0 | 0.34 | 0.04 | 11 | AT1 Epithelial Cells |
| MT-ND4 | 0.82 | 0 | 0 | 0.96 | 1.00 | 11 | AT1 Epithelial Cells |
| MT-ND2 | 0.82 | 0 | 0.03 | 0.97 | 0.99 | 11 | AT1 Epithelial Cells |
| TSTD1 | 0.81 | 0 | 0 | 0.32 | 0.07 | 11 | AT1 Epithelial Cells |
| HOPX | 0.8 | 0 | 0 | 0.27 | 0.12 | 11 | AT1 Epithelial Cells |
| PRDX5 | 0.8 | 0 | 0 | 0.70 | 0.56 | 11 | AT1 Epithelial Cells |
| MRPS31 | 0.78 | 0 | 0 | 0.30 | 0.12 | 11 | AT1 Epithelial Cells |
| EFHC1 | 0.76 | 0 | 0 | 0.34 | 0.15 | 11 | AT1 Epithelial Cells |
| HIST1H1C | 0.75 | 0 | 0 | 0.30 | 0.09 | 11 | AT1 Epithelial Cells |
| ARL4A | 0.75 | 0 | 0 | 0.35 | 0.12 | 11 | AT1 Epithelial Cells |
| CD55 | 0.73 | 0 | 0 | 0.37 | 0.12 | 11 | AT1 Epithelial Cells |
| TSPAN13 | 0.72 | 0 | 0 | 0.32 | 0.13 | 11 | AT1 Epithelial Cells |
| RAB11FIP1 | 0.71 | 0 | 0 | 0.37 | 0.04 | 11 | AT1 Epithelial Cells |
| NEAT1 | 0.71 | 0 | 0.02 | 0.79 | 0.63 | 11 | AT1 Epithelial Cells |
| VAMP8 | 0.7 | 0 | 0 | 0.46 | 0.18 | 11 | AT1 Epithelial Cells |
| MT-ND5 | 0.66 | 0 | 0 | 0.94 | 0.89 | 11 | AT1 Epithelial Cells |
| PDPN | 0.66 | 0 | 0 | 0.34 | 0.05 | 11 | AT1 Epithelial Cells |
| EFNA1 | 0.65 | 0 | 0 | 0.46 | 0.14 | 11 | AT1 Epithelial Cells |
| CRNDE | 0.64 | 0 | 0 | 0.28 | 0.01 | 11 | AT1 Epithelial Cells |
| SEC11C | 0.62 | 0 | 0 | 0.30 | 0.13 | 11 | AT1 Epithelial Cells |
| PTPRF | 0.62 | 0 | 0 | 0.31 | 0.06 | 11 | AT1 Epithelial Cells |
| CKB | 0.61 | 0 | 0 | 0.47 | 0.22 | 11 | AT1 Epithelial Cells |
| ANXA4 | 0.59 | 0 | 0 | 0.38 | 0.16 | 11 | AT1 Epithelial Cells |
| CD47 | 0.58 | 0 | 0 | 0.47 | 0.21 | 11 | AT1 Epithelial Cells |
| ATP11A | 0.58 | 0 | 0 | 0.29 | 0.06 | 11 | AT1 Epithelial Cells |
| SPINT2 | 0.58 | 0 | 0 | 0.33 | 0.15 | 11 | AT1 Epithelial Cells |
| CLDN7 | 0.57 | 0 | 0 | 0.25 | 0.03 | 11 | AT1 Epithelial Cells |
| ODF2L | 0.57 | 0 | 0 | 0.38 | 0.18 | 11 | AT1 Epithelial Cells |
| DYNLT1 | 0.53 | 0 | 0.03 | 0.60 | 0.43 | 11 | AT1 Epithelial Cells |
| NRP2 | 0.53 | 0 | 0 | 0.31 | 0.08 | 11 | AT1 Epithelial Cells |
| TUBA1C | 0.51 | 0 | 0 | 0.34 | 0.14 | 11 | AT1 Epithelial Cells |
| ELK3 | 0.51 | 0 | 0 | 0.35 | 0.15 | 11 | AT1 Epithelial Cells |
| LGALS3 | 0.51 | 0 | 0 | 0.58 | 0.37 | 11 | AT1 Epithelial Cells |
| PRKCZ | 0.51 | 0 | 0 | 0.30 | 0.02 | 11 | AT1 Epithelial Cells |
| WDR34 | 0.51 | 0 | 0 | 0.32 | 0.10 | 11 | AT1 Epithelial Cells |
| ATP1A1 | 0.51 | 0 | 0 | 0.45 | 0.21 | 11 | AT1 Epithelial Cells |
| MRPL14 | 0.5 | 0 | 0 | 0.50 | 0.23 | 11 | AT1 Epithelial Cells |
| HSPB11 | 0.49 | 0 | 0 | 0.34 | 0.17 | 11 | AT1 Epithelial Cells |
| ATP5E | 0.49 | 0 | 0.01 | 0.81 | 0.87 | 11 | AT1 Epithelial Cells |
| TXN | 0.48 | 0 | 0.02 | 0.67 | 0.55 | 11 | AT1 Epithelial Cells |
| SMC4 | 0.48 | 0 | 0 | 0.29 | 0.11 | 11 | AT1 Epithelial Cells |
| C4orf48 | 0.44 | 0 | 0 | 0.37 | 0.14 | 11 | AT1 Epithelial Cells |
| CSTB | 0.44 | 0 | 0 | 0.71 | 0.49 | 11 | AT1 Epithelial Cells |
| DNAJC10 | 0.44 | 0 | 0 | 0.32 | 0.11 | 11 | AT1 Epithelial Cells |
| PPFIBP1 | 0.44 | 0 | 0.04 | 0.30 | 0.15 | 11 | AT1 Epithelial Cells |
| FAM174A | 0.43 | 0 | 0 | 0.31 | 0.14 | 11 | AT1 Epithelial Cells |
| ITGA3 | 0.42 | 0 | 0 | 0.29 | 0.11 | 11 | AT1 Epithelial Cells |
| SRI | 0.42 | 0 | 0.01 | 0.57 | 0.38 | 11 | AT1 Epithelial Cells |
| HMGA1 | 0.42 | 0 | 0 | 0.29 | 0.10 | 11 | AT1 Epithelial Cells |
| MAGI1 | 0.41 | 0 | 0 | 0.31 | 0.07 | 11 | AT1 Epithelial Cells |
| PERP | 0.41 | 0 | 0 | 0.31 | 0.13 | 11 | AT1 Epithelial Cells |
| F11R | 0.4 | 0 | 0 | 0.31 | 0.07 | 11 | AT1 Epithelial Cells |
| ATXN2 | 0.4 | 0 | 0 | 0.37 | 0.16 | 11 | AT1 Epithelial Cells |
| PPDPF | 0.4 | 0 | 0.01 | 0.84 | 0.80 | 11 | AT1 Epithelial Cells |
| RPA3 | 0.4 | 0 | 0 | 0.31 | 0.14 | 11 | AT1 Epithelial Cells |
| GLS | 0.4 | 0 | 0 | 0.25 | 0.10 | 11 | AT1 Epithelial Cells |
| UCP2 | 0.39 | 0 | 0 | 0.33 | 0.12 | 11 | AT1 Epithelial Cells |
| GRAMD1A | 0.39 | 0 | 0 | 0.37 | 0.14 | 11 | AT1 Epithelial Cells |
| SMARCA5 | 0.38 | 0 | 0.04 | 0.37 | 0.20 | 11 | AT1 Epithelial Cells |
| PCBD1 | 0.38 | 0 | 0 | 0.45 | 0.21 | 11 | AT1 Epithelial Cells |
| HPCAL1 | 0.38 | 0 | 0 | 0.36 | 0.16 | 11 | AT1 Epithelial Cells |
| C1orf198 | 0.37 | 0 | 0.01 | 0.30 | 0.14 | 11 | AT1 Epithelial Cells |
| MBIP | 0.36 | 0 | 0 | 0.26 | 0.11 | 11 | AT1 Epithelial Cells |
| ADD3 | 0.35 | 0 | 0 | 0.45 | 0.23 | 11 | AT1 Epithelial Cells |
| CLU | 0.35 | 0 | 0.01 | 0.56 | 0.32 | 11 | AT1 Epithelial Cells |
| WNK1 | 0.34 | 0 | 0.03 | 0.29 | 0.14 | 11 | AT1 Epithelial Cells |
| ANXA2P2 | 0.34 | 0 | 0 | 0.28 | 0.08 | 11 | AT1 Epithelial Cells |
| HSPH1 | 0.34 | 0 | 0 | 0.26 | 0.11 | 11 | AT1 Epithelial Cells |
| PTP4A1 | 0.33 | 0 | 0 | 0.30 | 0.12 | 11 | AT1 Epithelial Cells |
| CTNNA1 | 0.33 | 0 | 0.04 | 0.53 | 0.34 | 11 | AT1 Epithelial Cells |
| FLNB | 0.33 | 0 | 0 | 0.26 | 0.06 | 11 | AT1 Epithelial Cells |
| CNDP2 | 0.31 | 0 | 0 | 0.29 | 0.13 | 11 | AT1 Epithelial Cells |
| SHC1 | 0.31 | 0 | 0 | 0.26 | 0.10 | 11 | AT1 Epithelial Cells |
| FILIP1 | 0.31 | 0 | 0 | 0.40 | 0.18 | 11 | AT1 Epithelial Cells |
| ABHD2 | 0.3 | 0 | 0 | 0.26 | 0.07 | 11 | AT1 Epithelial Cells |
| UTRN | 0.3 | 0 | 0 | 0.41 | 0.18 | 11 | AT1 Epithelial Cells |
| TJP2 | 0.29 | 0 | 0.01 | 0.25 | 0.11 | 11 | AT1 Epithelial Cells |
| COQ4 | 0.27 | 0 | 0 | 0.34 | 0.15 | 11 | AT1 Epithelial Cells |
| MLLT4 | 0.27 | 0 | 0 | 0.35 | 0.17 | 11 | AT1 Epithelial Cells |
| ENOSF1 | 0.27 | 0 | 0 | 0.28 | 0.07 | 11 | AT1 Epithelial Cells |
| FAM129B | 0.26 | 0 | 0 | 0.28 | 0.11 | 11 | AT1 Epithelial Cells |
| CTSD | 0.26 | 0 | 0.01 | 0.53 | 0.31 | 11 | AT1 Epithelial Cells |
| DPP7 | 0.25 | 0 | 0.05 | 0.38 | 0.20 | 11 | AT1 Epithelial Cells |
| SERPINE2 | 3.29 | 0 | 0 | 0.84 | 0.17 | 12 | Endothelial Cells |
| GLUL | 2.64 | 0 | 0 | 0.99 | 0.42 | 12 | Endothelial Cells |
| ID1 | 2.17 | 0 | 0 | 0.95 | 0.34 | 12 | Endothelial Cells |
| GJA4 | 2.06 | 0 | 0 | 0.91 | 0.14 | 12 | Endothelial Cells |
| SLC9A3R2 | 2.03 | 0 | 0 | 0.99 | 0.40 | 12 | Endothelial Cells |
| AQP1 | 2.03 | 0 | 0 | 0.99 | 0.20 | 12 | Endothelial Cells |
| KCTD12 | 1.99 | 0 | 0 | 0.91 | 0.23 | 12 | Endothelial Cells |
| ATP13A3 | 1.95 | 0 | 0 | 0.75 | 0.05 | 12 | Endothelial Cells |
| CLEC14A | 1.83 | 0 | 0 | 0.92 | 0.19 | 12 | Endothelial Cells |
| SOX17 | 1.83 | 0 | 0 | 0.72 | 0.05 | 12 | Endothelial Cells |
| SRGN | 1.82 | 0 | 0 | 0.97 | 0.22 | 12 | Endothelial Cells |
| H19 | 1.82 | 0 | 0 | 0.72 | 0.15 | 12 | Endothelial Cells |
| SULT1E1 | 1.77 | 0 | 0 | 0.47 | 0.05 | 12 | Endothelial Cells |
| IGF2 | 1.76 | 0 | 0 | 0.69 | 0.06 | 12 | Endothelial Cells |
| GJA5 | 1.72 | 0 | 0 | 0.65 | 0.06 | 12 | Endothelial Cells |
| PODXL | 1.67 | 0 | 0 | 0.73 | 0.08 | 12 | Endothelial Cells |
| CXCL12 | 1.64 | 0 | 0 | 0.82 | 0.24 | 12 | Endothelial Cells |
| CD9 | 1.63 | 0 | 0 | 0.92 | 0.45 | 12 | Endothelial Cells |
| HEG1 | 1.62 | 0 | 0 | 0.79 | 0.13 | 12 | Endothelial Cells |
| MMRN2 | 1.61 | 0 | 0 | 0.78 | 0.08 | 12 | Endothelial Cells |
| C10orf10 | 1.54 | 0 | 0 | 0.78 | 0.15 | 12 | Endothelial Cells |
| IFI27 | 1.53 | 0 | 0 | 1.00 | 0.36 | 12 | Endothelial Cells |
| CLDN5 | 1.5 | 0 | 0 | 0.95 | 0.26 | 12 | Endothelial Cells |
| STMN1 | 1.5 | 0 | 0 | 0.95 | 0.52 | 12 | Endothelial Cells |
| KLF2 | 1.49 | 0 | 0 | 0.73 | 0.17 | 12 | Endothelial Cells |
| TSPAN7 | 1.48 | 0 | 0 | 0.75 | 0.12 | 12 | Endothelial Cells |
| ENG | 1.48 | 0 | 0 | 0.80 | 0.30 | 12 | Endothelial Cells |
| CWC27 | 1.48 | 0 | 0 | 0.64 | 0.12 | 12 | Endothelial Cells |
| PECAM1 | 1.46 | 0 | 0 | 0.97 | 0.25 | 12 | Endothelial Cells |
| FBLN2 | 1.45 | 0 | 0 | 0.57 | 0.05 | 12 | Endothelial Cells |
| IFITM2 | 1.45 | 0 | 0 | 0.98 | 0.63 | 12 | Endothelial Cells |
| PI16 | 1.45 | 0 | 0 | 0.40 | 0.01 | 12 | Endothelial Cells |
| TMEM100 | 1.44 | 0 | 0 | 0.77 | 0.15 | 12 | Endothelial Cells |
| TM4SF1 | 1.44 | 0 | 0 | 0.97 | 0.39 | 12 | Endothelial Cells |
| NPDC1 | 1.44 | 0 | 0 | 0.85 | 0.30 | 12 | Endothelial Cells |
| HYAL2 | 1.43 | 0 | 0 | 0.87 | 0.22 | 12 | Endothelial Cells |
| SULF1 | 1.42 | 0 | 0 | 0.49 | 0.02 | 12 | Endothelial Cells |
| SRP14 | 1.42 | 0 | 0 | 1.00 | 0.92 | 12 | Endothelial Cells |
| JAM2 | 1.39 | 0 | 0 | 0.77 | 0.12 | 12 | Endothelial Cells |
| PTPRB | 1.39 | 0 | 0 | 0.66 | 0.07 | 12 | Endothelial Cells |
| EFNB2 | 1.39 | 0 | 0 | 0.69 | 0.08 | 12 | Endothelial Cells |
| CALCRL | 1.37 | 0 | 0 | 0.89 | 0.19 | 12 | Endothelial Cells |
| RNASE1 | 1.36 | 0 | 0 | 0.95 | 0.27 | 12 | Endothelial Cells |
| SLC6A6 | 1.33 | 0 | 0 | 0.49 | 0.04 | 12 | Endothelial Cells |
| RAMP2 | 1.32 | 0 | 0 | 0.99 | 0.33 | 12 | Endothelial Cells |
| ADAMTS6 | 1.31 | 0 | 0 | 0.65 | 0.12 | 12 | Endothelial Cells |
| VWF | 1.31 | 0 | 0 | 0.79 | 0.12 | 12 | Endothelial Cells |
| EDN1 | 1.31 | 0 | 0 | 0.75 | 0.10 | 12 | Endothelial Cells |
| ADAM15 | 1.3 | 0 | 0 | 0.67 | 0.11 | 12 | Endothelial Cells |
| ADGRL4 | 1.29 | 0 | 0 | 0.51 | 0.03 | 12 | Endothelial Cells |
| CD34 | 1.27 | 0 | 0 | 0.75 | 0.15 | 12 | Endothelial Cells |
| ICAM2 | 1.26 | 0 | 0 | 0.81 | 0.22 | 12 | Endothelial Cells |
| FOXP1 | 1.26 | 0 | 0 | 0.87 | 0.40 | 12 | Endothelial Cells |
| SEMA3G | 1.23 | 0 | 0 | 0.53 | 0.01 | 12 | Endothelial Cells |
| DKK2 | 1.23 | 0 | 0 | 0.46 | 0.00 | 12 | Endothelial Cells |
| ATP1B3 | 1.23 | 0 | 0 | 0.75 | 0.31 | 12 | Endothelial Cells |
| SOX18 | 1.23 | 0 | 0 | 0.62 | 0.10 | 12 | Endothelial Cells |
| BST2 | 1.22 | 0 | 0 | 0.97 | 0.29 | 12 | Endothelial Cells |
| EXOC6 | 1.22 | 0 | 0 | 0.58 | 0.07 | 12 | Endothelial Cells |
| OCIAD2 | 1.21 | 0 | 0 | 0.60 | 0.14 | 12 | Endothelial Cells |
| CRIP2 | 1.2 | 0 | 0 | 0.98 | 0.65 | 12 | Endothelial Cells |
| LTC4S | 1.18 | 0 | 0 | 0.53 | 0.08 | 12 | Endothelial Cells |
| TGFBR2 | 1.17 | 0 | 0 | 0.71 | 0.19 | 12 | Endothelial Cells |
| PDLIM1 | 1.17 | 0 | 0 | 0.85 | 0.38 | 12 | Endothelial Cells |
| SMS | 1.17 | 0 | 0 | 0.59 | 0.22 | 12 | Endothelial Cells |
| IFITM3 | 1.17 | 0 | 0 | 1.00 | 0.91 | 12 | Endothelial Cells |
| MECOM | 1.16 | 0 | 0 | 0.53 | 0.05 | 12 | Endothelial Cells |
| SSUH2 | 1.15 | 0 | 0 | 0.38 | 0.00 | 12 | Endothelial Cells |
| CLIC3 | 1.14 | 0 | 0 | 0.47 | 0.05 | 12 | Endothelial Cells |
| CFAP36 | 1.14 | 0 | 0 | 0.61 | 0.19 | 12 | Endothelial Cells |
| STOM | 1.13 | 0 | 0 | 0.92 | 0.44 | 12 | Endothelial Cells |
| JAG1 | 1.13 | 0 | 0 | 0.53 | 0.11 | 12 | Endothelial Cells |
| FBLN5 | 1.11 | 0 | 0 | 0.60 | 0.17 | 12 | Endothelial Cells |
| CTNNAL1 | 1.1 | 0 | 0 | 0.58 | 0.19 | 12 | Endothelial Cells |
| IFI6 | 1.09 | 0 | 0 | 0.65 | 0.26 | 12 | Endothelial Cells |
| MYCT1 | 1.09 | 0 | 0 | 0.55 | 0.06 | 12 | Endothelial Cells |
| CD93 | 1.09 | 0 | 0 | 0.68 | 0.15 | 12 | Endothelial Cells |
| CCDC88A | 1.08 | 0 | 0 | 0.45 | 0.14 | 12 | Endothelial Cells |
| ECSCR.1 | 1.08 | 0 | 0 | 0.80 | 0.19 | 12 | Endothelial Cells |
| LYPD2 | 1.08 | 0 | 0 | 0.38 | 0.02 | 12 | Endothelial Cells |
| ADCY4 | 1.07 | 0 | 0 | 0.59 | 0.09 | 12 | Endothelial Cells |
| ASRGL1 | 1.07 | 0 | 0 | 0.53 | 0.05 | 12 | Endothelial Cells |
| PCSK5 | 1.07 | 0 | 0 | 0.46 | 0.05 | 12 | Endothelial Cells |
| EPAS1 | 1.06 | 0 | 0 | 0.92 | 0.43 | 12 | Endothelial Cells |
| SEC11C | 1.06 | 0 | 0 | 0.58 | 0.12 | 12 | Endothelial Cells |
| IFITM1 | 1.05 | 0 | 0 | 0.53 | 0.17 | 12 | Endothelial Cells |
| C4orf48 | 1.04 | 0 | 0 | 0.58 | 0.14 | 12 | Endothelial Cells |
| AC004947.2 | 1.04 | 0 | 0 | 0.37 | 0.00 | 12 | Endothelial Cells |
| PCAT19 | 1.04 | 0 | 0 | 0.74 | 0.18 | 12 | Endothelial Cells |
| CDH5 | 1.04 | 0 | 0 | 0.82 | 0.18 | 12 | Endothelial Cells |
| FKBP1A | 1.03 | 0 | 0 | 0.93 | 0.72 | 12 | Endothelial Cells |
| CD81 | 1.03 | 0 | 0 | 0.72 | 0.39 | 12 | Endothelial Cells |
| DUSP1 | 1.03 | 0 | 0 | 0.74 | 0.41 | 12 | Endothelial Cells |
| LEPROTL1 | 1.03 | 0 | 0 | 0.74 | 0.24 | 12 | Endothelial Cells |
| CTSH | 1.02 | 0 | 0 | 0.68 | 0.13 | 12 | Endothelial Cells |
| IGFBP4 | 1.02 | 0 | 0 | 0.94 | 0.50 | 12 | Endothelial Cells |
| TSPO | 1.01 | 0 | 0 | 0.82 | 0.43 | 12 | Endothelial Cells |
| JAG2 | 1.01 | 0 | 0 | 0.44 | 0.03 | 12 | Endothelial Cells |
| EMP1 | 1 | 0 | 0 | 0.64 | 0.23 | 12 | Endothelial Cells |
| APOA1 | 1 | 0 | 0 | 0.39 | 0.02 | 12 | Endothelial Cells |
| MALL | 0.99 | 0 | 0 | 0.45 | 0.05 | 12 | Endothelial Cells |
| KLF4 | 0.99 | 0 | 0 | 0.54 | 0.16 | 12 | Endothelial Cells |
| SHE | 0.98 | 0 | 0 | 0.54 | 0.05 | 12 | Endothelial Cells |
| ACVRL1 | 0.98 | 0 | 0 | 0.64 | 0.16 | 12 | Endothelial Cells |
| HEY1 | 0.97 | 0 | 0 | 0.41 | 0.05 | 12 | Endothelial Cells |
| ECE1 | 0.96 | 0 | 0 | 0.60 | 0.15 | 12 | Endothelial Cells |
| CAV1 | 0.94 | 0 | 0 | 1.00 | 0.83 | 12 | Endothelial Cells |
| FCN3 | 0.93 | 0 | 0 | 0.41 | 0.06 | 12 | Endothelial Cells |
| PTPRM | 0.93 | 0 | 0 | 0.46 | 0.08 | 12 | Endothelial Cells |
| IL33 | 0.92 | 0 | 0 | 0.53 | 0.12 | 12 | Endothelial Cells |
| PIK3C2B | 0.92 | 0 | 0 | 0.45 | 0.04 | 12 | Endothelial Cells |
| NUDT14 | 0.92 | 0 | 0 | 0.54 | 0.09 | 12 | Endothelial Cells |
| AMD1 | 0.91 | 0 | 0 | 0.39 | 0.18 | 12 | Endothelial Cells |
| PALMD | 0.9 | 0 | 0 | 0.51 | 0.08 | 12 | Endothelial Cells |
| SNX3 | 0.89 | 0 | 0 | 0.91 | 0.69 | 12 | Endothelial Cells |
| COX17 | 0.89 | 0 | 0 | 0.72 | 0.35 | 12 | Endothelial Cells |
| ID3 | 0.89 | 0 | 0 | 0.88 | 0.54 | 12 | Endothelial Cells |
| PCMTD1 | 0.88 | 0 | 0 | 0.57 | 0.22 | 12 | Endothelial Cells |
| TAGLN2 | 0.88 | 0 | 0 | 0.94 | 0.75 | 12 | Endothelial Cells |
| BCAM | 0.88 | 0 | 0 | 0.84 | 0.33 | 12 | Endothelial Cells |
| SSTR1 | 0.87 | 0 | 0 | 0.40 | 0.01 | 12 | Endothelial Cells |
| LTBP4 | 0.87 | 0 | 0 | 0.68 | 0.29 | 12 | Endothelial Cells |
| MAML1 | 0.87 | 0 | 0 | 0.47 | 0.09 | 12 | Endothelial Cells |
| ITPR1 | 0.86 | 0 | 0 | 0.39 | 0.06 | 12 | Endothelial Cells |
| MIR503HG | 0.86 | 0 | 0 | 0.34 | 0.02 | 12 | Endothelial Cells |
| UACA | 0.85 | 0 | 0 | 0.55 | 0.24 | 12 | Endothelial Cells |
| AIF1L | 0.85 | 0 | 0 | 0.42 | 0.02 | 12 | Endothelial Cells |
| PCDH17 | 0.85 | 0 | 0 | 0.60 | 0.15 | 12 | Endothelial Cells |
| RHOC | 0.85 | 0 | 0 | 0.87 | 0.61 | 12 | Endothelial Cells |
| MRPL33 | 0.85 | 0 | 0 | 0.69 | 0.44 | 12 | Endothelial Cells |
| PARVB | 0.84 | 0 | 0 | 0.52 | 0.15 | 12 | Endothelial Cells |
| SPTAN1 | 0.84 | 0 | 0 | 0.57 | 0.22 | 12 | Endothelial Cells |
| TMSB10 | 0.84 | 0 | 0 | 1.00 | 0.99 | 12 | Endothelial Cells |
| FXYD5 | 0.84 | 0 | 0 | 0.74 | 0.26 | 12 | Endothelial Cells |
| IGFBP2 | 0.84 | 0 | 0 | 0.62 | 0.18 | 12 | Endothelial Cells |
| GATA2 | 0.84 | 0 | 0 | 0.48 | 0.07 | 12 | Endothelial Cells |
| PLLP | 0.83 | 0 | 0 | 0.38 | 0.03 | 12 | Endothelial Cells |
| TEK | 0.83 | 0 | 0 | 0.42 | 0.06 | 12 | Endothelial Cells |
| CD109 | 0.83 | 0 | 0 | 0.41 | 0.05 | 12 | Endothelial Cells |
| SWAP70 | 0.82 | 0 | 0 | 0.49 | 0.10 | 12 | Endothelial Cells |
| PTPRG | 0.82 | 0 | 0 | 0.45 | 0.10 | 12 | Endothelial Cells |
| CD320 | 0.82 | 0 | 0 | 0.54 | 0.15 | 12 | Endothelial Cells |
| PEMT | 0.81 | 0 | 0 | 0.41 | 0.11 | 12 | Endothelial Cells |
| CYYR1 | 0.81 | 0 | 0 | 0.51 | 0.12 | 12 | Endothelial Cells |
| APLNR | 0.8 | 0 | 0 | 0.53 | 0.12 | 12 | Endothelial Cells |
| FAM107A | 0.8 | 0 | 0 | 0.40 | 0.02 | 12 | Endothelial Cells |
| CD59 | 0.8 | 0 | 0 | 0.87 | 0.60 | 12 | Endothelial Cells |
| NDRG1 | 0.8 | 0 | 0 | 0.54 | 0.16 | 12 | Endothelial Cells |
| S100A16 | 0.8 | 0 | 0 | 0.79 | 0.34 | 12 | Endothelial Cells |
| HSPA12B | 0.79 | 0 | 0 | 0.48 | 0.09 | 12 | Endothelial Cells |
| CALR | 0.79 | 0 | 0 | 0.87 | 0.58 | 12 | Endothelial Cells |
| NRN1 | 0.79 | 0 | 0 | 0.51 | 0.08 | 12 | Endothelial Cells |
| SYNPO | 0.79 | 0 | 0 | 0.45 | 0.12 | 12 | Endothelial Cells |
| LAP3 | 0.79 | 0 | 0 | 0.58 | 0.22 | 12 | Endothelial Cells |
| APOL4 | 0.79 | 0 | 0 | 0.34 | 0.02 | 12 | Endothelial Cells |
| LIMS2 | 0.78 | 0 | 0 | 0.44 | 0.08 | 12 | Endothelial Cells |
| PRKCDBP | 0.78 | 0 | 0 | 0.79 | 0.40 | 12 | Endothelial Cells |
| EGFL7 | 0.78 | 0 | 0 | 0.81 | 0.26 | 12 | Endothelial Cells |
| PPA1 | 0.77 | 0 | 0 | 0.68 | 0.37 | 12 | Endothelial Cells |
| FKBP9 | 0.77 | 0 | 0 | 0.49 | 0.14 | 12 | Endothelial Cells |
| CAPZA2 | 0.77 | 0 | 0 | 0.74 | 0.51 | 12 | Endothelial Cells |
| NOTCH4 | 0.77 | 0 | 0 | 0.51 | 0.12 | 12 | Endothelial Cells |
| HSPG2 | 0.76 | 0 | 0 | 0.53 | 0.19 | 12 | Endothelial Cells |
| RDX | 0.75 | 0 | 0 | 0.79 | 0.43 | 12 | Endothelial Cells |
| SMAD6 | 0.75 | 0 | 0 | 0.37 | 0.04 | 12 | Endothelial Cells |
| ESAM | 0.75 | 0 | 0 | 0.82 | 0.35 | 12 | Endothelial Cells |
| C1orf54 | 0.75 | 0 | 0 | 0.52 | 0.18 | 12 | Endothelial Cells |
| NAA10 | 0.74 | 0 | 0 | 0.68 | 0.30 | 12 | Endothelial Cells |
| FABP5 | 0.73 | 0 | 0 | 0.53 | 0.27 | 12 | Endothelial Cells |
| SLC14A1 | 0.73 | 0 | 0 | 0.28 | 0.02 | 12 | Endothelial Cells |
| SARAF | 0.73 | 0 | 0 | 0.86 | 0.64 | 12 | Endothelial Cells |
| GIMAP1 | 0.73 | 0 | 0 | 0.44 | 0.10 | 12 | Endothelial Cells |
| HSPC324 | 0.72 | 0 | 0 | 0.39 | 0.06 | 12 | Endothelial Cells |
| SLFN5 | 0.72 | 0 | 0 | 0.39 | 0.08 | 12 | Endothelial Cells |
| AC079776.2 | 0.72 | 0 | 0 | 0.29 | 0.00 | 12 | Endothelial Cells |
| IL32 | 0.72 | 0 | 0 | 0.80 | 0.28 | 12 | Endothelial Cells |
| ANXA2 | 0.72 | 0 | 0 | 0.92 | 0.65 | 12 | Endothelial Cells |
| EMP3 | 0.72 | 0 | 0 | 0.86 | 0.59 | 12 | Endothelial Cells |
| ITGB4 | 0.72 | 0 | 0 | 0.33 | 0.02 | 12 | Endothelial Cells |
| SENP7 | 0.71 | 0 | 0 | 0.26 | 0.06 | 12 | Endothelial Cells |
| TUBB | 0.71 | 0 | 0 | 0.91 | 0.79 | 12 | Endothelial Cells |
| LGALS9 | 0.71 | 0 | 0 | 0.42 | 0.10 | 12 | Endothelial Cells |
| KIAA1462 | 0.71 | 0 | 0 | 0.49 | 0.13 | 12 | Endothelial Cells |
| TUBA4A | 0.71 | 0 | 0 | 0.42 | 0.08 | 12 | Endothelial Cells |
| RP11-467L13.5 | 0.71 | 0 | 0 | 0.55 | 0.19 | 12 | Endothelial Cells |
| BMPR2 | 0.7 | 0 | 0 | 0.58 | 0.22 | 12 | Endothelial Cells |
| TMBIM1 | 0.7 | 0 | 0 | 0.58 | 0.24 | 12 | Endothelial Cells |
| FZD6 | 0.7 | 0 | 0 | 0.35 | 0.02 | 12 | Endothelial Cells |
| TIE1 | 0.7 | 0 | 0 | 0.58 | 0.14 | 12 | Endothelial Cells |
| MAGEH1 | 0.69 | 0 | 0 | 0.55 | 0.27 | 12 | Endothelial Cells |
| C1orf115 | 0.69 | 0 | 0 | 0.34 | 0.05 | 12 | Endothelial Cells |
| ITGA5 | 0.69 | 0 | 0 | 0.45 | 0.13 | 12 | Endothelial Cells |
| ANXA3 | 0.68 | 0 | 0 | 0.46 | 0.12 | 12 | Endothelial Cells |
| ETS1 | 0.68 | 0 | 0 | 0.44 | 0.15 | 12 | Endothelial Cells |
| NR4A1 | 0.68 | 0 | 0 | 0.34 | 0.14 | 12 | Endothelial Cells |
| TNFSF10 | 0.68 | 0 | 0 | 0.69 | 0.22 | 12 | Endothelial Cells |
| LIMA1 | 0.68 | 0 | 0 | 0.55 | 0.21 | 12 | Endothelial Cells |
| ELMO1 | 0.68 | 0 | 0 | 0.41 | 0.09 | 12 | Endothelial Cells |
| ISG15 | 0.68 | 0 | 0 | 0.53 | 0.26 | 12 | Endothelial Cells |
| PTRF | 0.68 | 0 | 0 | 0.92 | 0.69 | 12 | Endothelial Cells |
| LRRFIP1 | 0.67 | 0 | 0 | 0.53 | 0.27 | 12 | Endothelial Cells |
| COBLL1 | 0.67 | 0 | 0 | 0.42 | 0.07 | 12 | Endothelial Cells |
| S100A11 | 0.67 | 0 | 0 | 0.93 | 0.60 | 12 | Endothelial Cells |
| NEDD9 | 0.67 | 0 | 0 | 0.41 | 0.15 | 12 | Endothelial Cells |
| ATP1A1 | 0.66 | 0 | 0 | 0.54 | 0.22 | 12 | Endothelial Cells |
| FLT1 | 0.66 | 0 | 0 | 0.40 | 0.08 | 12 | Endothelial Cells |
| EVL | 0.66 | 0 | 0 | 0.47 | 0.19 | 12 | Endothelial Cells |
| ELK3 | 0.66 | 0 | 0 | 0.49 | 0.15 | 12 | Endothelial Cells |
| SSBP4 | 0.66 | 0 | 0 | 0.52 | 0.18 | 12 | Endothelial Cells |
| PIK3R3 | 0.66 | 0 | 0 | 0.33 | 0.03 | 12 | Endothelial Cells |
| KAZALD1 | 0.66 | 0 | 0 | 0.31 | 0.02 | 12 | Endothelial Cells |
| SULT1B1 | 0.66 | 0 | 0 | 0.28 | 0.01 | 12 | Endothelial Cells |
| NCKAP1 | 0.65 | 0 | 0 | 0.48 | 0.17 | 12 | Endothelial Cells |
| HOXA5 | 0.65 | 0 | 0 | 0.58 | 0.23 | 12 | Endothelial Cells |
| ADGRF5 | 0.65 | 0 | 0 | 0.53 | 0.17 | 12 | Endothelial Cells |
| LRP6 | 0.65 | 0 | 0 | 0.40 | 0.11 | 12 | Endothelial Cells |
| ITSN2 | 0.65 | 0 | 0 | 0.47 | 0.16 | 12 | Endothelial Cells |
| GIMAP4 | 0.65 | 0 | 0 | 0.58 | 0.19 | 12 | Endothelial Cells |
| GNAI2 | 0.65 | 0 | 0 | 0.65 | 0.34 | 12 | Endothelial Cells |
| MRPL17 | 0.64 | 0 | 0 | 0.44 | 0.14 | 12 | Endothelial Cells |
| SKAP2 | 0.64 | 0 | 0 | 0.49 | 0.26 | 12 | Endothelial Cells |
| FRYL | 0.64 | 0 | 0 | 0.38 | 0.11 | 12 | Endothelial Cells |
| F2R | 0.64 | 0 | 0 | 0.42 | 0.10 | 12 | Endothelial Cells |
| GADD45G | 0.64 | 0 | 0 | 0.31 | 0.11 | 12 | Endothelial Cells |
| LRRC32 | 0.64 | 0 | 0 | 0.46 | 0.13 | 12 | Endothelial Cells |
| VEGFC | 0.64 | 0 | 0 | 0.33 | 0.04 | 12 | Endothelial Cells |
| POMP | 0.63 | 0 | 0 | 0.80 | 0.62 | 12 | Endothelial Cells |
| PAPSS1 | 0.63 | 0 | 0 | 0.42 | 0.13 | 12 | Endothelial Cells |
| SYNGR2 | 0.63 | 0 | 0 | 0.57 | 0.27 | 12 | Endothelial Cells |
| F11R | 0.63 | 0 | 0 | 0.35 | 0.07 | 12 | Endothelial Cells |
| CD58 | 0.63 | 0 | 0 | 0.35 | 0.09 | 12 | Endothelial Cells |
| HSPA5 | 0.63 | 0 | 0 | 0.77 | 0.49 | 12 | Endothelial Cells |
| PLPP1 | 0.63 | 0 | 0 | 0.47 | 0.23 | 12 | Endothelial Cells |
| RAPGEF5 | 0.63 | 0 | 0 | 0.47 | 0.13 | 12 | Endothelial Cells |
| ACTN4 | 0.63 | 0 | 0 | 0.65 | 0.40 | 12 | Endothelial Cells |
| ARGLU1 | 0.63 | 0 | 0 | 0.67 | 0.46 | 12 | Endothelial Cells |
| MAP3K11 | 0.62 | 0 | 0 | 0.46 | 0.15 | 12 | Endothelial Cells |
| CPNE8 | 0.62 | 0 | 0 | 0.28 | 0.04 | 12 | Endothelial Cells |
| RAPGEF1 | 0.62 | 0 | 0 | 0.32 | 0.06 | 12 | Endothelial Cells |
| NAA38 | 0.62 | 0 | 0 | 0.68 | 0.38 | 12 | Endothelial Cells |
| ENTPD1 | 0.62 | 0 | 0 | 0.47 | 0.13 | 12 | Endothelial Cells |
| TSPAN2 | 0.62 | 0 | 0 | 0.35 | 0.10 | 12 | Endothelial Cells |
| RASIP1 | 0.62 | 0 | 0 | 0.40 | 0.10 | 12 | Endothelial Cells |
| LDB2 | 0.62 | 0 | 0 | 0.54 | 0.18 | 12 | Endothelial Cells |
| PTTG1IP | 0.61 | 0 | 0 | 0.65 | 0.33 | 12 | Endothelial Cells |
| PLSCR1 | 0.61 | 0 | 0 | 0.46 | 0.17 | 12 | Endothelial Cells |
| SOX7 | 0.61 | 0 | 0 | 0.39 | 0.09 | 12 | Endothelial Cells |
| SDCBP | 0.61 | 0 | 0 | 0.75 | 0.52 | 12 | Endothelial Cells |
| WARS | 0.61 | 0 | 0 | 0.52 | 0.20 | 12 | Endothelial Cells |
| MGST2 | 0.61 | 0 | 0 | 0.60 | 0.26 | 12 | Endothelial Cells |
| SLC44A2 | 0.6 | 0 | 0 | 0.46 | 0.17 | 12 | Endothelial Cells |
| GIMAP7 | 0.6 | 0 | 0 | 0.59 | 0.17 | 12 | Endothelial Cells |
| CALM1 | 0.6 | 0 | 0 | 0.85 | 0.74 | 12 | Endothelial Cells |
| PREX2 | 0.6 | 0 | 0 | 0.32 | 0.05 | 12 | Endothelial Cells |
| THSD7A | 0.6 | 0 | 0 | 0.27 | 0.03 | 12 | Endothelial Cells |
| C8orf33 | 0.6 | 0 | 0 | 0.41 | 0.21 | 12 | Endothelial Cells |
| ASS1 | 0.6 | 0 | 0 | 0.32 | 0.05 | 12 | Endothelial Cells |
| PDLIM5 | 0.59 | 0 | 0 | 0.49 | 0.16 | 12 | Endothelial Cells |
| FLI1 | 0.59 | 0 | 0 | 0.40 | 0.12 | 12 | Endothelial Cells |
| SSFA2 | 0.59 | 0 | 0 | 0.38 | 0.09 | 12 | Endothelial Cells |
| ARL15 | 0.59 | 0 | 0 | 0.33 | 0.03 | 12 | Endothelial Cells |
| BHLHE40 | 0.59 | 0 | 0 | 0.39 | 0.08 | 12 | Endothelial Cells |
| SIPA1L2 | 0.59 | 0 | 0 | 0.35 | 0.11 | 12 | Endothelial Cells |
| GPRC5B | 0.59 | 0 | 0 | 0.33 | 0.03 | 12 | Endothelial Cells |
| PRCP | 0.59 | 0 | 0 | 0.58 | 0.24 | 12 | Endothelial Cells |
| PROCR | 0.59 | 0 | 0 | 0.44 | 0.14 | 12 | Endothelial Cells |
| ABLIM1 | 0.59 | 0 | 0 | 0.35 | 0.07 | 12 | Endothelial Cells |
| COL9A3 | 0.58 | 0 | 0 | 0.29 | 0.02 | 12 | Endothelial Cells |
| MEIS2 | 0.58 | 0 | 0 | 0.40 | 0.12 | 12 | Endothelial Cells |
| RPS6KA2 | 0.58 | 0 | 0 | 0.40 | 0.11 | 12 | Endothelial Cells |
| RAC1 | 0.58 | 0 | 0 | 0.88 | 0.81 | 12 | Endothelial Cells |
| SH3BP5 | 0.58 | 0 | 0 | 0.55 | 0.21 | 12 | Endothelial Cells |
| APOL3 | 0.58 | 0 | 0 | 0.44 | 0.12 | 12 | Endothelial Cells |
| PRICKLE1 | 0.58 | 0 | 0 | 0.29 | 0.05 | 12 | Endothelial Cells |
| UGCG | 0.58 | 0 | 0 | 0.34 | 0.13 | 12 | Endothelial Cells |
| C10orf11 | 0.57 | 0 | 0 | 0.35 | 0.07 | 12 | Endothelial Cells |
| ARPC1B | 0.57 | 0 | 0 | 0.58 | 0.31 | 12 | Endothelial Cells |
| DOCK9 | 0.57 | 0 | 0 | 0.26 | 0.07 | 12 | Endothelial Cells |
| GNAQ | 0.57 | 0 | 0 | 0.41 | 0.16 | 12 | Endothelial Cells |
| SLC12A2 | 0.57 | 0 | 0 | 0.31 | 0.12 | 12 | Endothelial Cells |
| CTTNBP2NL | 0.57 | 0 | 0 | 0.35 | 0.08 | 12 | Endothelial Cells |
| SREK1IP1 | 0.57 | 0 | 0 | 0.51 | 0.26 | 12 | Endothelial Cells |
| ARHGEF3 | 0.56 | 0 | 0 | 0.39 | 0.08 | 12 | Endothelial Cells |
| PKN3 | 0.56 | 0 | 0 | 0.27 | 0.02 | 12 | Endothelial Cells |
| LSR | 0.56 | 0 | 0 | 0.34 | 0.10 | 12 | Endothelial Cells |
| RALB | 0.56 | 0 | 0 | 0.48 | 0.19 | 12 | Endothelial Cells |
| SPTBN1 | 0.56 | 0 | 0 | 0.78 | 0.42 | 12 | Endothelial Cells |
| ADAR | 0.56 | 0 | 0 | 0.52 | 0.23 | 12 | Endothelial Cells |
| CLIC4 | 0.56 | 0 | 0 | 0.59 | 0.33 | 12 | Endothelial Cells |
| PREX1 | 0.55 | 0 | 0 | 0.32 | 0.08 | 12 | Endothelial Cells |
| PLCG2 | 0.55 | 0 | 0 | 0.26 | 0.03 | 12 | Endothelial Cells |
| MYL12B | 0.55 | 0 | 0 | 0.86 | 0.77 | 12 | Endothelial Cells |
| C1QTNF6 | 0.55 | 0 | 0 | 0.28 | 0.05 | 12 | Endothelial Cells |
| KNOP1 | 0.55 | 0 | 0 | 0.40 | 0.17 | 12 | Endothelial Cells |
| KIAA1671 | 0.55 | 0 | 0 | 0.27 | 0.03 | 12 | Endothelial Cells |
| JUNB | 0.55 | 0 | 0.01 | 0.71 | 0.52 | 12 | Endothelial Cells |
| THSD1 | 0.55 | 0 | 0 | 0.39 | 0.09 | 12 | Endothelial Cells |
| C19orf33 | 0.55 | 0 | 0 | 0.28 | 0.03 | 12 | Endothelial Cells |
| AC132217.4 | 0.54 | 0 | 0 | 0.26 | 0.01 | 12 | Endothelial Cells |
| OSBPL8 | 0.54 | 0 | 0.01 | 0.39 | 0.18 | 12 | Endothelial Cells |
| TTC28 | 0.54 | 0 | 0 | 0.35 | 0.13 | 12 | Endothelial Cells |
| FURIN | 0.54 | 0 | 0 | 0.29 | 0.07 | 12 | Endothelial Cells |
| CTNNA1 | 0.54 | 0 | 0 | 0.57 | 0.34 | 12 | Endothelial Cells |
| TACC1 | 0.53 | 0 | 0 | 0.71 | 0.46 | 12 | Endothelial Cells |
| IER2 | 0.53 | 0 | 0.05 | 0.62 | 0.46 | 12 | Endothelial Cells |
| COL18A1 | 0.53 | 0 | 0 | 0.40 | 0.18 | 12 | Endothelial Cells |
| MYO1C | 0.53 | 0 | 0 | 0.48 | 0.23 | 12 | Endothelial Cells |
| CFLAR | 0.53 | 0 | 0 | 0.54 | 0.26 | 12 | Endothelial Cells |
| CRIM1 | 0.53 | 0 | 0.01 | 0.31 | 0.13 | 12 | Endothelial Cells |
| TUBB4B | 0.53 | 0 | 0 | 0.62 | 0.40 | 12 | Endothelial Cells |
| WWTR1 | 0.53 | 0 | 0.01 | 0.41 | 0.21 | 12 | Endothelial Cells |
| DHH | 0.53 | 0 | 0 | 0.28 | 0.01 | 12 | Endothelial Cells |
| FGL2 | 0.53 | 0 | 0 | 0.26 | 0.06 | 12 | Endothelial Cells |
| TCN2 | 0.53 | 0 | 0 | 0.27 | 0.06 | 12 | Endothelial Cells |
| SORBS2 | 0.52 | 0 | 0 | 0.27 | 0.07 | 12 | Endothelial Cells |
| TNFAIP1 | 0.52 | 0 | 0 | 0.29 | 0.10 | 12 | Endothelial Cells |
| TMEM30A | 0.52 | 0 | 0 | 0.46 | 0.21 | 12 | Endothelial Cells |
| TES | 0.52 | 0 | 0 | 0.28 | 0.09 | 12 | Endothelial Cells |
| KLF3 | 0.52 | 0 | 0 | 0.38 | 0.13 | 12 | Endothelial Cells |
| ERC1 | 0.52 | 0 | 0 | 0.31 | 0.09 | 12 | Endothelial Cells |
| TBC1D1 | 0.52 | 0 | 0 | 0.31 | 0.11 | 12 | Endothelial Cells |
| ARHGEF15 | 0.51 | 0 | 0 | 0.38 | 0.07 | 12 | Endothelial Cells |
| FDPS | 0.51 | 0 | 0 | 0.60 | 0.34 | 12 | Endothelial Cells |
| KANK3 | 0.51 | 0 | 0 | 0.38 | 0.14 | 12 | Endothelial Cells |
| CYB5R3 | 0.51 | 0 | 0.03 | 0.64 | 0.45 | 12 | Endothelial Cells |
| ORAI1 | 0.5 | 0 | 0 | 0.32 | 0.07 | 12 | Endothelial Cells |
| LEPR | 0.5 | 0 | 0 | 0.31 | 0.08 | 12 | Endothelial Cells |
| SERINC3 | 0.5 | 0 | 0 | 0.47 | 0.24 | 12 | Endothelial Cells |
| LYL1 | 0.5 | 0 | 0 | 0.33 | 0.11 | 12 | Endothelial Cells |
| FEZ2 | 0.5 | 0 | 0 | 0.47 | 0.24 | 12 | Endothelial Cells |
| FLNB | 0.5 | 0 | 0 | 0.31 | 0.06 | 12 | Endothelial Cells |
| FUT8 | 0.5 | 0 | 0 | 0.28 | 0.05 | 12 | Endothelial Cells |
| NDUFC2 | 0.5 | 0 | 0 | 0.77 | 0.64 | 12 | Endothelial Cells |
| ELF1 | 0.5 | 0 | 0 | 0.44 | 0.16 | 12 | Endothelial Cells |
| CASKIN2 | 0.49 | 0 | 0 | 0.35 | 0.09 | 12 | Endothelial Cells |
| RHOA | 0.49 | 0 | 0 | 0.81 | 0.70 | 12 | Endothelial Cells |
| EPHB4 | 0.49 | 0 | 0 | 0.32 | 0.08 | 12 | Endothelial Cells |
| DLL4 | 0.49 | 0 | 0 | 0.28 | 0.06 | 12 | Endothelial Cells |
| MLLT4 | 0.49 | 0 | 0 | 0.41 | 0.17 | 12 | Endothelial Cells |
| G3BP1 | 0.49 | 0 | 0 | 0.48 | 0.23 | 12 | Endothelial Cells |
| ITGB1 | 0.49 | 0 | 0 | 0.84 | 0.63 | 12 | Endothelial Cells |
| BIRC6 | 0.49 | 0 | 0 | 0.35 | 0.14 | 12 | Endothelial Cells |
| TFPI | 0.49 | 0 | 0.03 | 0.67 | 0.43 | 12 | Endothelial Cells |
| SOX13 | 0.49 | 0 | 0 | 0.28 | 0.05 | 12 | Endothelial Cells |
| EFNA1 | 0.48 | 0 | 0 | 0.40 | 0.14 | 12 | Endothelial Cells |
| NOTCH1 | 0.48 | 0 | 0 | 0.35 | 0.08 | 12 | Endothelial Cells |
| SLC2A1 | 0.48 | 0 | 0 | 0.26 | 0.04 | 12 | Endothelial Cells |
| MAST4 | 0.48 | 0 | 0 | 0.39 | 0.16 | 12 | Endothelial Cells |
| NPR1 | 0.48 | 0 | 0 | 0.28 | 0.04 | 12 | Endothelial Cells |
| AP1S2 | 0.47 | 0 | 0.01 | 0.46 | 0.24 | 12 | Endothelial Cells |
| SCRN2 | 0.47 | 0 | 0 | 0.31 | 0.12 | 12 | Endothelial Cells |
| TPM3 | 0.47 | 0 | 0 | 0.57 | 0.31 | 12 | Endothelial Cells |
| GUK1 | 0.47 | 0 | 0.03 | 0.77 | 0.67 | 12 | Endothelial Cells |
| HSP90B1 | 0.47 | 0 | 0 | 0.79 | 0.60 | 12 | Endothelial Cells |
| AHR | 0.47 | 0 | 0 | 0.33 | 0.12 | 12 | Endothelial Cells |
| RPN2 | 0.47 | 0 | 0 | 0.64 | 0.38 | 12 | Endothelial Cells |
| RP11-693N9.2 | 0.47 | 0 | 0 | 0.39 | 0.16 | 12 | Endothelial Cells |
| ELN | 0.47 | 0 | 0 | 0.86 | 0.54 | 12 | Endothelial Cells |
| CDC37 | 0.47 | 0 | 0.01 | 0.68 | 0.49 | 12 | Endothelial Cells |
| CMTM8 | 0.46 | 0 | 0 | 0.27 | 0.06 | 12 | Endothelial Cells |
| MMP15 | 0.46 | 0 | 0 | 0.29 | 0.05 | 12 | Endothelial Cells |
| AGRN | 0.46 | 0 | 0 | 0.27 | 0.07 | 12 | Endothelial Cells |
| TNKS1BP1 | 0.46 | 0 | 0 | 0.27 | 0.08 | 12 | Endothelial Cells |
| RICTOR | 0.46 | 0 | 0 | 0.39 | 0.15 | 12 | Endothelial Cells |
| S1PR1 | 0.46 | 0 | 0 | 0.39 | 0.14 | 12 | Endothelial Cells |
| UTRN | 0.46 | 0 | 0 | 0.42 | 0.18 | 12 | Endothelial Cells |
| AZIN1 | 0.46 | 0 | 0 | 0.35 | 0.13 | 12 | Endothelial Cells |
| MAPK3 | 0.45 | 0 | 0 | 0.39 | 0.17 | 12 | Endothelial Cells |
| S100A6 | 0.45 | 0 | 0 | 0.98 | 0.84 | 12 | Endothelial Cells |
| FAM69B | 0.45 | 0 | 0 | 0.26 | 0.05 | 12 | Endothelial Cells |
| FES | 0.45 | 0 | 0 | 0.28 | 0.07 | 12 | Endothelial Cells |
| HHEX | 0.45 | 0 | 0 | 0.32 | 0.10 | 12 | Endothelial Cells |
| KRAS | 0.45 | 0 | 0 | 0.37 | 0.15 | 12 | Endothelial Cells |
| CCDC85B | 0.45 | 0 | 0.02 | 0.62 | 0.41 | 12 | Endothelial Cells |
| TUSC3 | 0.45 | 0 | 0.01 | 0.41 | 0.21 | 12 | Endothelial Cells |
| WNK1 | 0.45 | 0 | 0.03 | 0.32 | 0.14 | 12 | Endothelial Cells |
| FKBP1C | 0.45 | 0 | 0 | 0.28 | 0.10 | 12 | Endothelial Cells |
| MMP2 | 0.44 | 0 | 0 | 0.65 | 0.34 | 12 | Endothelial Cells |
| DYNLL1 | 0.44 | 0 | 0 | 0.85 | 0.79 | 12 | Endothelial Cells |
| SNAP23 | 0.44 | 0 | 0.01 | 0.40 | 0.19 | 12 | Endothelial Cells |
| LUZP1 | 0.44 | 0 | 0 | 0.40 | 0.18 | 12 | Endothelial Cells |
| GFOD1 | 0.44 | 0 | 0 | 0.26 | 0.06 | 12 | Endothelial Cells |
| RAB11A | 0.44 | 0 | 0 | 0.69 | 0.44 | 12 | Endothelial Cells |
| THBD | 0.44 | 0 | 0 | 0.45 | 0.15 | 12 | Endothelial Cells |
| ABHD17A | 0.44 | 0 | 0 | 0.40 | 0.16 | 12 | Endothelial Cells |
| MGAT1 | 0.44 | 0 | 0.02 | 0.46 | 0.25 | 12 | Endothelial Cells |
| PPP2R2A | 0.44 | 0 | 0 | 0.40 | 0.19 | 12 | Endothelial Cells |
| TMEM88 | 0.44 | 0 | 0 | 0.33 | 0.08 | 12 | Endothelial Cells |
| CCDC88C | 0.44 | 0 | 0 | 0.35 | 0.10 | 12 | Endothelial Cells |
| ARHGAP27 | 0.43 | 0 | 0 | 0.27 | 0.05 | 12 | Endothelial Cells |
| PRKD2 | 0.43 | 0 | 0 | 0.28 | 0.08 | 12 | Endothelial Cells |
| VAMP5 | 0.43 | 0 | 0 | 0.78 | 0.52 | 12 | Endothelial Cells |
| CXCR4 | 0.43 | 0 | 0 | 0.35 | 0.10 | 12 | Endothelial Cells |
| ITGA6 | 0.43 | 0 | 0 | 0.28 | 0.07 | 12 | Endothelial Cells |
| TUBA1B | 0.43 | 0 | 0 | 0.89 | 0.78 | 12 | Endothelial Cells |
| KRTCAP2 | 0.43 | 0 | 0.04 | 0.72 | 0.54 | 12 | Endothelial Cells |
| MLLT1 | 0.43 | 0 | 0 | 0.28 | 0.08 | 12 | Endothelial Cells |
| ZFAND2B | 0.43 | 0 | 0 | 0.32 | 0.12 | 12 | Endothelial Cells |
| PLS3 | 0.43 | 0 | 0 | 0.67 | 0.43 | 12 | Endothelial Cells |
| RAB3GAP1 | 0.43 | 0 | 0 | 0.26 | 0.10 | 12 | Endothelial Cells |
| TSPAN14 | 0.42 | 0 | 0 | 0.37 | 0.14 | 12 | Endothelial Cells |
| QKI | 0.42 | 0 | 0 | 0.45 | 0.20 | 12 | Endothelial Cells |
| C10orf54 | 0.42 | 0 | 0 | 0.45 | 0.20 | 12 | Endothelial Cells |
| SERTAD4-AS1 | 0.42 | 0 | 0 | 0.27 | 0.05 | 12 | Endothelial Cells |
| PEA15 | 0.42 | 0 | 0.02 | 0.49 | 0.27 | 12 | Endothelial Cells |
| TRIB2 | 0.42 | 0 | 0 | 0.29 | 0.10 | 12 | Endothelial Cells |
| PHACTR2 | 0.42 | 0 | 0 | 0.51 | 0.26 | 12 | Endothelial Cells |
| DTD1 | 0.41 | 0 | 0 | 0.29 | 0.11 | 12 | Endothelial Cells |
| ARHGAP17 | 0.41 | 0 | 0 | 0.27 | 0.07 | 12 | Endothelial Cells |
| FRMD4B | 0.41 | 0 | 0 | 0.41 | 0.15 | 12 | Endothelial Cells |
| MRPS28 | 0.41 | 0 | 0.01 | 0.32 | 0.13 | 12 | Endothelial Cells |
| IGF2BP2 | 0.4 | 0 | 0 | 0.38 | 0.15 | 12 | Endothelial Cells |
| APOL1 | 0.4 | 0 | 0 | 0.26 | 0.09 | 12 | Endothelial Cells |
| TCEAL9 | 0.4 | 0 | 0.03 | 0.75 | 0.54 | 12 | Endothelial Cells |
| IL11RA | 0.4 | 0 | 0 | 0.28 | 0.11 | 12 | Endothelial Cells |
| CEP68 | 0.4 | 0 | 0 | 0.29 | 0.11 | 12 | Endothelial Cells |
| FAM129B | 0.4 | 0 | 0 | 0.32 | 0.11 | 12 | Endothelial Cells |
| USHBP1 | 0.4 | 0 | 0 | 0.26 | 0.04 | 12 | Endothelial Cells |
| FYN | 0.39 | 0 | 0.01 | 0.32 | 0.13 | 12 | Endothelial Cells |
| CITED4 | 0.39 | 0 | 0 | 0.26 | 0.05 | 12 | Endothelial Cells |
| KDR | 0.39 | 0 | 0 | 0.28 | 0.09 | 12 | Endothelial Cells |
| SH3BGRL3 | 0.39 | 0 | 0 | 0.87 | 0.58 | 12 | Endothelial Cells |
| CCNY | 0.39 | 0 | 0 | 0.26 | 0.08 | 12 | Endothelial Cells |
| PAK4 | 0.39 | 0 | 0 | 0.28 | 0.10 | 12 | Endothelial Cells |
| TUBG1 | 0.39 | 0 | 0 | 0.26 | 0.09 | 12 | Endothelial Cells |
| FGD5 | 0.39 | 0 | 0 | 0.26 | 0.06 | 12 | Endothelial Cells |
| KIF5B | 0.39 | 0 | 0.01 | 0.71 | 0.47 | 12 | Endothelial Cells |
| ELOVL5 | 0.39 | 0 | 0.04 | 0.31 | 0.13 | 12 | Endothelial Cells |
| ARRB1 | 0.38 | 0 | 0 | 0.27 | 0.08 | 12 | Endothelial Cells |
| GAS2L1 | 0.38 | 0 | 0 | 0.26 | 0.07 | 12 | Endothelial Cells |
| CNKSR3 | 0.38 | 0 | 0 | 0.26 | 0.07 | 12 | Endothelial Cells |
| SH2D3C | 0.38 | 0 | 0 | 0.35 | 0.12 | 12 | Endothelial Cells |
| HLA-B | 0.38 | 0 | 0 | 0.95 | 0.54 | 12 | Endothelial Cells |
| RAI2 | 0.37 | 0 | 0 | 0.27 | 0.06 | 12 | Endothelial Cells |
| HPCAL1 | 0.37 | 0 | 0 | 0.38 | 0.16 | 12 | Endothelial Cells |
| MIER2 | 0.37 | 0 | 0 | 0.27 | 0.08 | 12 | Endothelial Cells |
| TMEM181 | 0.37 | 0 | 0 | 0.26 | 0.05 | 12 | Endothelial Cells |
| RAI14 | 0.36 | 0 | 0 | 0.28 | 0.10 | 12 | Endothelial Cells |
| VAMP3 | 0.36 | 0 | 0.01 | 0.51 | 0.26 | 12 | Endothelial Cells |
| MFNG | 0.36 | 0 | 0 | 0.27 | 0.07 | 12 | Endothelial Cells |
| GABARAPL2 | 0.35 | 0 | 0.03 | 0.81 | 0.66 | 12 | Endothelial Cells |
| IFNAR2 | 0.35 | 0 | 0 | 0.27 | 0.10 | 12 | Endothelial Cells |
| FRY | 0.35 | 0 | 0 | 0.37 | 0.15 | 12 | Endothelial Cells |
| DOCK6 | 0.35 | 0 | 0 | 0.39 | 0.15 | 12 | Endothelial Cells |
| SERF2 | 0.35 | 0 | 0.01 | 0.98 | 0.95 | 12 | Endothelial Cells |
| TSPAN13 | 0.35 | 0 | 0 | 0.34 | 0.13 | 12 | Endothelial Cells |
| A4GALT | 0.35 | 0 | 0 | 0.26 | 0.08 | 12 | Endothelial Cells |
| ELK4 | 0.34 | 0 | 0 | 0.33 | 0.13 | 12 | Endothelial Cells |
| PRKCH | 0.34 | 0 | 0 | 0.31 | 0.09 | 12 | Endothelial Cells |
| ERG | 0.34 | 0 | 0 | 0.37 | 0.12 | 12 | Endothelial Cells |
| RGS5 | 0.34 | 0 | 0.02 | 0.48 | 0.24 | 12 | Endothelial Cells |
| SLCO2A1 | 0.34 | 0 | 0 | 0.32 | 0.11 | 12 | Endothelial Cells |
| PPIA | 0.34 | 0 | 0.02 | 0.93 | 0.85 | 12 | Endothelial Cells |
| GNG11 | 0.34 | 0 | 0 | 0.85 | 0.44 | 12 | Endothelial Cells |
| SVIP | 0.33 | 0 | 0.04 | 0.35 | 0.16 | 12 | Endothelial Cells |
| INPP5K | 0.33 | 0 | 0 | 0.31 | 0.10 | 12 | Endothelial Cells |
| SHKBP1 | 0.33 | 0 | 0.02 | 0.28 | 0.11 | 12 | Endothelial Cells |
| A2M | 0.32 | 0 | 0 | 0.93 | 0.61 | 12 | Endothelial Cells |
| TPST2 | 0.32 | 0 | 0 | 0.38 | 0.15 | 12 | Endothelial Cells |
| HLA-E | 0.32 | 0 | 0 | 0.88 | 0.60 | 12 | Endothelial Cells |
| EPS8 | 0.31 | 0 | 0.02 | 0.27 | 0.10 | 12 | Endothelial Cells |
| MANSC1 | 0.31 | 0 | 0 | 0.26 | 0.05 | 12 | Endothelial Cells |
| SPTLC2 | 0.3 | 0 | 0 | 0.26 | 0.09 | 12 | Endothelial Cells |
| CYFIP1 | 0.3 | 0 | 0.03 | 0.35 | 0.16 | 12 | Endothelial Cells |
| RAMP3 | 0.3 | 0 | 0 | 0.40 | 0.16 | 12 | Endothelial Cells |
| PICALM | 0.3 | 0 | 0.04 | 0.38 | 0.17 | 12 | Endothelial Cells |
| CDC42EP3 | 0.29 | 0 | 0 | 0.26 | 0.09 | 12 | Endothelial Cells |
| CFL1 | 0.29 | 0 | 0.03 | 0.97 | 0.91 | 12 | Endothelial Cells |
| AGPAT1 | 0.28 | 0 | 0.05 | 0.28 | 0.11 | 12 | Endothelial Cells |
| NMI | 0.27 | 0 | 0.01 | 0.31 | 0.12 | 12 | Endothelial Cells |
| C8orf4 | 0.27 | 0 | 0.02 | 0.68 | 0.43 | 12 | Endothelial Cells |
| PTMA | 0.27 | 0 | 0 | 1.00 | 1.00 | 12 | Endothelial Cells |
| IGHM | 4.27 | 0 | 0 | 0.99 | 0.02 | 13 | B Cells |
| IGKC | 4.12 | 0 | 0 | 0.61 | 0.02 | 13 | B Cells |
| IGLC2 | 3.57 | 0 | 0 | 0.77 | 0.00 | 13 | B Cells |
| IGLC3 | 3.24 | 0 | 0 | 0.63 | 0.00 | 13 | B Cells |
| CD79B | 2.92 | 0 | 0 | 0.94 | 0.02 | 13 | B Cells |
| CD37 | 2.77 | 0 | 0 | 0.98 | 0.07 | 13 | B Cells |
| LTB | 2.7 | 0 | 0 | 0.92 | 0.04 | 13 | B Cells |
| MS4A1 | 2.68 | 0 | 0 | 0.90 | 0.00 | 13 | B Cells |
| TCL1A | 2.6 | 0 | 0 | 0.75 | 0.00 | 13 | B Cells |
| CD79A | 2.46 | 0 | 0 | 0.94 | 0.01 | 13 | B Cells |
| CD52 | 2.42 | 0 | 0 | 0.99 | 0.09 | 13 | B Cells |
| CD74 | 2.31 | 0 | 0 | 1.00 | 0.44 | 13 | B Cells |
| IGHD | 2.14 | 0 | 0 | 0.75 | 0.00 | 13 | B Cells |
| VPREB3 | 1.88 | 0 | 0 | 0.63 | 0.00 | 13 | B Cells |
| CXCR4 | 1.84 | 0 | 0 | 0.72 | 0.10 | 13 | B Cells |
| IGLL5 | 1.78 | 0 | 0 | 0.38 | 0.00 | 13 | B Cells |
| HLA-DRA | 1.64 | 0 | 0 | 0.98 | 0.26 | 13 | B Cells |
| SPIB | 1.56 | 0 | 0 | 0.54 | 0.01 | 13 | B Cells |
| FAM129C | 1.56 | 0 | 0 | 0.61 | 0.01 | 13 | B Cells |
| CORO1A | 1.55 | 0 | 0 | 0.84 | 0.11 | 13 | B Cells |
| MZB1 | 1.53 | 0 | 0 | 0.57 | 0.00 | 13 | B Cells |
| HLA-DPB1 | 1.5 | 0 | 0 | 0.94 | 0.18 | 13 | B Cells |
| LIMD2 | 1.5 | 0 | 0 | 0.79 | 0.14 | 13 | B Cells |
| HLA-DRB1 | 1.48 | 0 | 0 | 0.91 | 0.11 | 13 | B Cells |
| PLAC8 | 1.46 | 0 | 0 | 0.67 | 0.07 | 13 | B Cells |
| JCHAIN | 1.44 | 0 | 0 | 0.48 | 0.02 | 13 | B Cells |
| HLA-DMB | 1.43 | 0 | 0 | 0.70 | 0.06 | 13 | B Cells |
| LAPTM5 | 1.43 | 0 | 0 | 0.76 | 0.15 | 13 | B Cells |
| FCRLA | 1.42 | 0 | 0 | 0.56 | 0.00 | 13 | B Cells |
| HLA-DPA1 | 1.36 | 0 | 0 | 0.89 | 0.15 | 13 | B Cells |
| UCP2 | 1.34 | 0 | 0 | 0.72 | 0.11 | 13 | B Cells |
| NCF1C | 1.33 | 0 | 0 | 0.54 | 0.01 | 13 | B Cells |
| CD24 | 1.31 | 0 | 0 | 0.39 | 0.02 | 13 | B Cells |
| PLD4 | 1.28 | 0 | 0 | 0.51 | 0.01 | 13 | B Cells |
| BIRC3 | 1.27 | 0 | 0 | 0.49 | 0.03 | 13 | B Cells |
| HLA-DMA | 1.24 | 0 | 0 | 0.66 | 0.09 | 13 | B Cells |
| RAC2 | 1.24 | 0 | 0 | 0.68 | 0.08 | 13 | B Cells |
| LINC00926 | 1.22 | 0 | 0 | 0.49 | 0.01 | 13 | B Cells |
| CYB561A3 | 1.22 | 0 | 0 | 0.56 | 0.09 | 13 | B Cells |
| IRF8 | 1.22 | 0 | 0 | 0.53 | 0.04 | 13 | B Cells |
| BANK1 | 1.21 | 0 | 0 | 0.53 | 0.00 | 13 | B Cells |
| RCSD1 | 1.2 | 0 | 0 | 0.67 | 0.18 | 13 | B Cells |
| PTPN6 | 1.2 | 0 | 0 | 0.57 | 0.07 | 13 | B Cells |
| ISG20 | 1.18 | 0 | 0 | 0.57 | 0.05 | 13 | B Cells |
| FCMR | 1.18 | 0 | 0 | 0.52 | 0.03 | 13 | B Cells |
| CLEC2D | 1.16 | 0 | 0 | 0.62 | 0.08 | 13 | B Cells |
| CD19 | 1.15 | 0 | 0 | 0.49 | 0.00 | 13 | B Cells |
| SMIM14 | 1.07 | 0 | 0 | 0.58 | 0.22 | 13 | B Cells |
| PCDH9 | 1.07 | 0 | 0 | 0.28 | 0.04 | 13 | B Cells |
| FCRL1 | 1.06 | 0 | 0 | 0.42 | 0.00 | 13 | B Cells |
| RP5-887A10.1 | 1.05 | 0 | 0 | 0.33 | 0.00 | 13 | B Cells |
| HLA-DQA1 | 1.05 | 0 | 0 | 0.47 | 0.03 | 13 | B Cells |
| CSK | 1 | 0 | 0 | 0.49 | 0.08 | 13 | B Cells |
| CD22 | 1 | 0 | 0 | 0.41 | 0.01 | 13 | B Cells |
| CD53 | 0.99 | 0 | 0 | 0.60 | 0.09 | 13 | B Cells |
| RHOH | 0.99 | 0 | 0 | 0.44 | 0.02 | 13 | B Cells |
| KIAA0226L | 0.98 | 0 | 0 | 0.42 | 0.01 | 13 | B Cells |
| ARHGDIB | 0.97 | 0 | 0 | 0.77 | 0.30 | 13 | B Cells |
| LSP1 | 0.96 | 0 | 0 | 0.60 | 0.10 | 13 | B Cells |
| BTG1 | 0.96 | 0 | 0 | 0.99 | 0.68 | 13 | B Cells |
| FCER2 | 0.95 | 0 | 0 | 0.33 | 0.01 | 13 | B Cells |
| GPSM3 | 0.94 | 0 | 0 | 0.60 | 0.14 | 13 | B Cells |
| JUNB | 0.93 | 0 | 0 | 0.80 | 0.52 | 13 | B Cells |
| TNFRSF13C | 0.93 | 0 | 0 | 0.41 | 0.00 | 13 | B Cells |
| LY86 | 0.93 | 0 | 0 | 0.47 | 0.04 | 13 | B Cells |
| CD72 | 0.93 | 0 | 0 | 0.38 | 0.01 | 13 | B Cells |
| RALGPS2 | 0.91 | 0 | 0 | 0.34 | 0.02 | 13 | B Cells |
| HLA-DQA2 | 0.9 | 0 | 0 | 0.43 | 0.03 | 13 | B Cells |
| PTPRC | 0.9 | 0 | 0 | 0.49 | 0.08 | 13 | B Cells |
| SP140 | 0.9 | 0 | 0 | 0.39 | 0.01 | 13 | B Cells |
| IL2RG | 0.9 | 0 | 0 | 0.43 | 0.05 | 13 | B Cells |
| DNAJB1 | 0.9 | 0 | 0 | 0.49 | 0.21 | 13 | B Cells |
| CYBA | 0.9 | 0 | 0 | 0.90 | 0.52 | 13 | B Cells |
| TBC1D10C | 0.88 | 0 | 0 | 0.37 | 0.03 | 13 | B Cells |
| DRAM2 | 0.84 | 0 | 0 | 0.47 | 0.14 | 13 | B Cells |
| POU2F2 | 0.83 | 0 | 0 | 0.35 | 0.02 | 13 | B Cells |
| LYN | 0.83 | 0 | 0 | 0.44 | 0.09 | 13 | B Cells |
| SMCHD1 | 0.83 | 0 | 0 | 0.44 | 0.12 | 13 | B Cells |
| LAT2 | 0.83 | 0 | 0 | 0.30 | 0.04 | 13 | B Cells |
| TSTD1 | 0.83 | 0 | 0 | 0.44 | 0.07 | 13 | B Cells |
| DUSP1 | 0.83 | 0 | 0 | 0.63 | 0.41 | 13 | B Cells |
| BTK | 0.83 | 0 | 0 | 0.34 | 0.02 | 13 | B Cells |
| HSH2D | 0.83 | 0 | 0 | 0.34 | 0.01 | 13 | B Cells |
| PRKCB | 0.82 | 0 | 0 | 0.39 | 0.08 | 13 | B Cells |
| NAPSB | 0.82 | 0 | 0 | 0.34 | 0.03 | 13 | B Cells |
| RPS27 | 0.82 | 0 | 0 | 1.00 | 1.00 | 13 | B Cells |
| SWAP70 | 0.82 | 0 | 0 | 0.42 | 0.10 | 13 | B Cells |
| ALOX5 | 0.82 | 0 | 0 | 0.30 | 0.03 | 13 | B Cells |
| RNASET2 | 0.81 | 0 | 0 | 0.56 | 0.18 | 13 | B Cells |
| FCRL5 | 0.81 | 0 | 0 | 0.27 | 0.00 | 13 | B Cells |
| IER2 | 0.81 | 0 | 0 | 0.70 | 0.46 | 13 | B Cells |
| RPL18AP3 | 0.8 | 0 | 0 | 0.96 | 0.94 | 13 | B Cells |
| ERP29 | 0.8 | 0 | 0 | 0.67 | 0.37 | 13 | B Cells |
| RPL28 | 0.8 | 0 | 0 | 0.99 | 0.98 | 13 | B Cells |
| LRMP | 0.79 | 0 | 0 | 0.25 | 0.01 | 13 | B Cells |
| SP110 | 0.79 | 0 | 0 | 0.33 | 0.09 | 13 | B Cells |
| NCF1B | 0.79 | 0 | 0 | 0.27 | 0.00 | 13 | B Cells |
| SIT1 | 0.78 | 0 | 0 | 0.32 | 0.00 | 13 | B Cells |
| CAPG | 0.77 | 0 | 0 | 0.38 | 0.08 | 13 | B Cells |
| FAM65B | 0.77 | 0 | 0 | 0.34 | 0.04 | 13 | B Cells |
| FAM26F | 0.77 | 0 | 0 | 0.37 | 0.03 | 13 | B Cells |
| CD48 | 0.77 | 0 | 0 | 0.41 | 0.06 | 13 | B Cells |
| RPS29 | 0.77 | 0 | 0 | 1.00 | 0.99 | 13 | B Cells |
| CD69 | 0.77 | 0 | 0 | 0.28 | 0.02 | 13 | B Cells |
| C16orf74 | 0.76 | 0 | 0 | 0.28 | 0.03 | 13 | B Cells |
| SELL | 0.76 | 0 | 0 | 0.29 | 0.04 | 13 | B Cells |
| EEF1B2 | 0.76 | 0 | 0 | 0.84 | 0.74 | 13 | B Cells |
| RPL18A | 0.76 | 0 | 0 | 0.99 | 0.99 | 13 | B Cells |
| RPSA | 0.76 | 0 | 0 | 0.91 | 0.90 | 13 | B Cells |
| RPS2P5 | 0.75 | 0 | 0 | 0.84 | 0.74 | 13 | B Cells |
| HSPA1B | 0.75 | 0 | 0 | 0.66 | 0.38 | 13 | B Cells |
| GADD45B | 0.74 | 0 | 0 | 0.73 | 0.42 | 13 | B Cells |
| PHACTR1 | 0.74 | 0 | 0 | 0.25 | 0.05 | 13 | B Cells |
| MYLIP | 0.74 | 0 | 0.03 | 0.52 | 0.30 | 13 | B Cells |
| 1-Mar | 0.73 | 0 | 0 | 0.29 | 0.02 | 13 | B Cells |
| FOSB | 0.72 | 0 | 0 | 0.43 | 0.14 | 13 | B Cells |
| RP11-761N21.2 | 0.72 | 0 | 0 | 0.82 | 0.63 | 13 | B Cells |
| BCL11A | 0.72 | 0 | 0 | 0.25 | 0.01 | 13 | B Cells |
| ZCCHC7 | 0.72 | 0 | 0 | 0.30 | 0.09 | 13 | B Cells |
| IKZF1 | 0.71 | 0 | 0 | 0.29 | 0.03 | 13 | B Cells |
| LCP1 | 0.71 | 0 | 0 | 0.42 | 0.08 | 13 | B Cells |
| STX7 | 0.71 | 0 | 0 | 0.37 | 0.15 | 13 | B Cells |
| RP4-765C7.2 | 0.71 | 0 | 0.02 | 0.63 | 0.51 | 13 | B Cells |
| STK4 | 0.71 | 0 | 0 | 0.48 | 0.16 | 13 | B Cells |
| RPS19 | 0.71 | 0 | 0 | 0.99 | 0.99 | 13 | B Cells |
| RASGRP2 | 0.7 | 0 | 0 | 0.33 | 0.06 | 13 | B Cells |
| EEF1B2P6 | 0.7 | 0 | 0 | 0.72 | 0.42 | 13 | B Cells |
| RPS21 | 0.69 | 0 | 0 | 0.99 | 0.93 | 13 | B Cells |
| ACAP1 | 0.69 | 0 | 0 | 0.32 | 0.05 | 13 | B Cells |
| EVI2B | 0.69 | 0 | 0 | 0.35 | 0.03 | 13 | B Cells |
| ARHGAP4 | 0.69 | 0 | 0 | 0.32 | 0.05 | 13 | B Cells |
| AFF3 | 0.69 | 0 | 0 | 0.30 | 0.05 | 13 | B Cells |
| QRSL1 | 0.69 | 0 | 0 | 0.29 | 0.05 | 13 | B Cells |
| BLNK | 0.68 | 0 | 0 | 0.29 | 0.00 | 13 | B Cells |
| ARPC1B | 0.68 | 0 | 0 | 0.73 | 0.31 | 13 | B Cells |
| RPL22L1 | 0.68 | 0 | 0 | 0.54 | 0.22 | 13 | B Cells |
| HLA-DQB1 | 0.68 | 0 | 0 | 0.43 | 0.05 | 13 | B Cells |
| ARPC3 | 0.67 | 0 | 0 | 0.81 | 0.61 | 13 | B Cells |
| CXXC5 | 0.67 | 0 | 0 | 0.32 | 0.09 | 13 | B Cells |
| RP11-386I14.4 | 0.67 | 0 | 0 | 0.27 | 0.06 | 13 | B Cells |
| HLA-DOB | 0.66 | 0 | 0 | 0.28 | 0.00 | 13 | B Cells |
| DAPP1 | 0.66 | 0 | 0 | 0.28 | 0.02 | 13 | B Cells |
| NCF1 | 0.66 | 0 | 0 | 0.30 | 0.01 | 13 | B Cells |
| PARP1 | 0.66 | 0 | 0 | 0.43 | 0.20 | 13 | B Cells |
| PABPC1 | 0.66 | 0 | 0 | 0.94 | 0.74 | 13 | B Cells |
| HIST1H1C | 0.65 | 0 | 0 | 0.27 | 0.09 | 13 | B Cells |
| 1-Sep | 0.65 | 0 | 0 | 0.29 | 0.05 | 13 | B Cells |
| RPSAP58 | 0.65 | 0 | 0 | 0.63 | 0.47 | 13 | B Cells |
| IKZF3 | 0.65 | 0 | 0 | 0.25 | 0.01 | 13 | B Cells |
| TAPBP | 0.65 | 0 | 0 | 0.58 | 0.25 | 13 | B Cells |
| ARHGAP30 | 0.65 | 0 | 0 | 0.32 | 0.04 | 13 | B Cells |
| RGS19 | 0.64 | 0 | 0 | 0.27 | 0.08 | 13 | B Cells |
| ALOX5AP | 0.64 | 0 | 0 | 0.29 | 0.03 | 13 | B Cells |
| ICAM3 | 0.64 | 0 | 0 | 0.38 | 0.08 | 13 | B Cells |
| ST6GAL1 | 0.64 | 0 | 0 | 0.25 | 0.04 | 13 | B Cells |
| AC097523.1 | 0.64 | 0 | 0 | 0.68 | 0.43 | 13 | B Cells |
| HCLS1 | 0.64 | 0 | 0 | 0.38 | 0.07 | 13 | B Cells |
| RPL39 | 0.63 | 0 | 0 | 1.00 | 0.99 | 13 | B Cells |
| HCK | 0.63 | 0 | 0 | 0.25 | 0.06 | 13 | B Cells |
| PSMB9 | 0.63 | 0 | 0 | 0.54 | 0.21 | 13 | B Cells |
| CD83 | 0.63 | 0 | 0 | 0.28 | 0.06 | 13 | B Cells |
| CYFIP2 | 0.63 | 0 | 0 | 0.27 | 0.03 | 13 | B Cells |
| RPS2 | 0.62 | 0 | 0 | 1.00 | 0.99 | 13 | B Cells |
| SP100 | 0.62 | 0 | 0 | 0.47 | 0.19 | 13 | B Cells |
| HVCN1 | 0.62 | 0 | 0 | 0.29 | 0.05 | 13 | B Cells |
| ELF1 | 0.62 | 0 | 0 | 0.37 | 0.16 | 13 | B Cells |
| TNFAIP8 | 0.61 | 0 | 0 | 0.33 | 0.09 | 13 | B Cells |
| RP11-796E2.4 | 0.61 | 0 | 0 | 0.29 | 0.10 | 13 | B Cells |
| ORAI2 | 0.61 | 0 | 0 | 0.27 | 0.06 | 13 | B Cells |
| RPL13P12 | 0.61 | 0 | 0 | 0.96 | 0.96 | 13 | B Cells |
| RPLP2 | 0.61 | 0 | 0 | 1.00 | 0.99 | 13 | B Cells |
| RP11-889L3.1 | 0.6 | 0 | 0 | 0.92 | 0.86 | 13 | B Cells |
| PIM2 | 0.6 | 0 | 0 | 0.28 | 0.02 | 13 | B Cells |
| EVL | 0.6 | 0 | 0 | 0.49 | 0.19 | 13 | B Cells |
| HIP1R | 0.59 | 0 | 0 | 0.25 | 0.02 | 13 | B Cells |
| HLA-F | 0.59 | 0 | 0 | 0.39 | 0.13 | 13 | B Cells |
| CTSS | 0.59 | 0 | 0 | 0.51 | 0.10 | 13 | B Cells |
| DCK | 0.59 | 0 | 0 | 0.28 | 0.09 | 13 | B Cells |
| SNX2 | 0.59 | 0 | 0.03 | 0.46 | 0.26 | 13 | B Cells |
| 6-Sep | 0.59 | 0 | 0 | 0.46 | 0.21 | 13 | B Cells |
| SLC38A1 | 0.59 | 0 | 0 | 0.28 | 0.03 | 13 | B Cells |
| RPL37 | 0.58 | 0 | 0 | 1.00 | 0.98 | 13 | B Cells |
| RPL12 | 0.58 | 0 | 0 | 0.99 | 0.97 | 13 | B Cells |
| CYTIP | 0.58 | 0 | 0 | 0.29 | 0.04 | 13 | B Cells |
| RPS11 | 0.58 | 0 | 0 | 0.99 | 0.97 | 13 | B Cells |
| STRBP | 0.57 | 0 | 0 | 0.27 | 0.04 | 13 | B Cells |
| SYTL1 | 0.57 | 0 | 0 | 0.27 | 0.02 | 13 | B Cells |
| RP11-234A1.1 | 0.57 | 0 | 0 | 0.96 | 0.94 | 13 | B Cells |
| RPL13AP5 | 0.56 | 0 | 0 | 0.95 | 0.94 | 13 | B Cells |
| FOS | 0.56 | 0 | 0.01 | 0.60 | 0.35 | 13 | B Cells |
| BTG2 | 0.56 | 0 | 0.02 | 0.42 | 0.21 | 13 | B Cells |
| P2RX5 | 0.56 | 0 | 0 | 0.25 | 0.00 | 13 | B Cells |
| RP11-864N7.2 | 0.55 | 0 | 0 | 0.94 | 0.90 | 13 | B Cells |
| CD55 | 0.55 | 0 | 0 | 0.39 | 0.13 | 13 | B Cells |
| RPL12P4 | 0.55 | 0 | 0 | 0.82 | 0.79 | 13 | B Cells |
| RPS12 | 0.54 | 0 | 0 | 0.99 | 0.99 | 13 | B Cells |
| STK17B | 0.54 | 0 | 0 | 0.25 | 0.06 | 13 | B Cells |
| ODC1 | 0.54 | 0 | 0 | 0.35 | 0.09 | 13 | B Cells |
| ARHGEF1 | 0.54 | 0 | 0 | 0.37 | 0.14 | 13 | B Cells |
| HLA-B | 0.53 | 0 | 0 | 0.96 | 0.54 | 13 | B Cells |
| RPS18 | 0.53 | 0 | 0 | 1.00 | 1.00 | 13 | B Cells |
| PLEKHA2 | 0.53 | 0 | 0 | 0.30 | 0.10 | 13 | B Cells |
| ADPGK | 0.53 | 0 | 0.01 | 0.30 | 0.13 | 13 | B Cells |
| SNX29 | 0.52 | 0 | 0 | 0.25 | 0.06 | 13 | B Cells |
| RPS23 | 0.52 | 0 | 0 | 1.00 | 0.99 | 13 | B Cells |
| RPS2P46 | 0.52 | 0 | 0.01 | 0.65 | 0.49 | 13 | B Cells |
| RPL13A | 0.52 | 0 | 0 | 0.99 | 1.00 | 13 | B Cells |
| RPL19 | 0.52 | 0 | 0 | 1.00 | 0.99 | 13 | B Cells |
| AC004453.8 | 0.52 | 0 | 0 | 0.92 | 0.86 | 13 | B Cells |
| RPL13 | 0.52 | 0 | 0 | 1.00 | 1.00 | 13 | B Cells |
| RPS10 | 0.52 | 0 | 0 | 0.95 | 0.90 | 13 | B Cells |
| RPL23A | 0.52 | 0 | 0 | 0.99 | 0.98 | 13 | B Cells |
| SIPA1 | 0.51 | 0 | 0 | 0.30 | 0.08 | 13 | B Cells |
| CD40 | 0.51 | 0 | 0 | 0.28 | 0.10 | 13 | B Cells |
| RPS5 | 0.51 | 0 | 0 | 0.95 | 0.96 | 13 | B Cells |
| WAS | 0.51 | 0 | 0 | 0.25 | 0.04 | 13 | B Cells |
| HMGA1 | 0.5 | 0 | 0 | 0.33 | 0.10 | 13 | B Cells |
| LYL1 | 0.5 | 0 | 0.01 | 0.28 | 0.11 | 13 | B Cells |
| RPL23AP42 | 0.5 | 0 | 0.01 | 0.79 | 0.67 | 13 | B Cells |
| RP11-425L10.1 | 0.49 | 0 | 0 | 0.89 | 0.84 | 13 | B Cells |
| RP11-475C16.1 | 0.49 | 0 | 0 | 0.90 | 0.83 | 13 | B Cells |
| CTD-2192J16.15 | 0.49 | 0 | 0 | 0.92 | 0.91 | 13 | B Cells |
| COTL1 | 0.49 | 0 | 0 | 0.52 | 0.20 | 13 | B Cells |
| IRF7 | 0.48 | 0 | 0 | 0.25 | 0.08 | 13 | B Cells |
| KLF2 | 0.48 | 0 | 0 | 0.39 | 0.18 | 13 | B Cells |
| PSMB10 | 0.48 | 0 | 0 | 0.52 | 0.25 | 13 | B Cells |
| SPINT2 | 0.47 | 0 | 0 | 0.39 | 0.15 | 13 | B Cells |
| RPS23P8 | 0.47 | 0 | 0.01 | 0.82 | 0.83 | 13 | B Cells |
| GGA2 | 0.47 | 0 | 0 | 0.25 | 0.09 | 13 | B Cells |
| HLA-C | 0.47 | 0 | 0 | 0.90 | 0.54 | 13 | B Cells |
| AB019441.29 | 0.46 | 0 | 0.01 | 0.66 | 0.47 | 13 | B Cells |
| DDX5 | 0.46 | 0 | 0 | 0.92 | 0.79 | 13 | B Cells |
| RP11-832N8.1 | 0.46 | 0 | 0.01 | 0.39 | 0.19 | 13 | B Cells |
| RPLP0 | 0.46 | 0 | 0 | 0.96 | 0.93 | 13 | B Cells |
| SMC6 | 0.46 | 0 | 0 | 0.29 | 0.08 | 13 | B Cells |
| LGALS9 | 0.45 | 0 | 0 | 0.34 | 0.10 | 13 | B Cells |
| RPS15A | 0.45 | 0 | 0 | 0.99 | 0.99 | 13 | B Cells |
| RPL32 | 0.44 | 0 | 0 | 1.00 | 0.99 | 13 | B Cells |
| RPL27A | 0.43 | 0 | 0 | 1.00 | 0.99 | 13 | B Cells |
| GLTSCR2 | 0.43 | 0 | 0.03 | 0.82 | 0.73 | 13 | B Cells |
| RPL34 | 0.43 | 0 | 0 | 1.00 | 1.00 | 13 | B Cells |
| RPL26 | 0.43 | 0 | 0 | 0.98 | 0.99 | 13 | B Cells |
| RPL10 | 0.42 | 0 | 0 | 1.00 | 1.00 | 13 | B Cells |
| RPL29 | 0.42 | 0 | 0 | 0.98 | 0.97 | 13 | B Cells |
| IRF1 | 0.42 | 0 | 0.04 | 0.41 | 0.20 | 13 | B Cells |
| RPS13 | 0.41 | 0 | 0 | 1.00 | 0.97 | 13 | B Cells |
| PDE7A | 0.41 | 0 | 0 | 0.25 | 0.05 | 13 | B Cells |
| RACK1 | 0.41 | 0 | 0 | 0.98 | 0.96 | 13 | B Cells |
| RPL30 | 0.41 | 0 | 0 | 0.99 | 0.98 | 13 | B Cells |
| RPS24 | 0.4 | 0 | 0 | 1.00 | 0.98 | 13 | B Cells |
| RPS28 | 0.4 | 0 | 0 | 0.99 | 0.99 | 13 | B Cells |
| ELOVL5 | 0.4 | 0 | 0.02 | 0.32 | 0.13 | 13 | B Cells |
| DHPS | 0.4 | 0 | 0.03 | 0.42 | 0.21 | 13 | B Cells |
| RPL26P19 | 0.4 | 0 | 0.01 | 0.87 | 0.86 | 13 | B Cells |
| RPS3 | 0.4 | 0 | 0 | 0.99 | 0.98 | 13 | B Cells |
| RP11-371A22.1 | 0.4 | 0 | 0.02 | 0.87 | 0.86 | 13 | B Cells |
| RPS14 | 0.39 | 0 | 0 | 1.00 | 1.00 | 13 | B Cells |
| B2M | 0.38 | 0 | 0 | 1.00 | 0.99 | 13 | B Cells |
| RPS3A | 0.37 | 0 | 0 | 1.00 | 0.99 | 13 | B Cells |
| RPS17 | 0.37 | 0 | 0 | 0.94 | 0.98 | 13 | B Cells |
| ZNF581 | 0.37 | 0 | 0.03 | 0.25 | 0.10 | 13 | B Cells |
| C17orf62 | 0.37 | 0 | 0.04 | 0.33 | 0.15 | 13 | B Cells |
| RPL18 | 0.36 | 0 | 0 | 0.98 | 0.98 | 13 | B Cells |
| ITGB2 | 0.36 | 0 | 0 | 0.29 | 0.08 | 13 | B Cells |
| HLA-A | 0.35 | 0 | 0 | 0.96 | 0.70 | 13 | B Cells |
| RPL27 | 0.35 | 0 | 0 | 0.96 | 0.96 | 13 | B Cells |
| RPL11 | 0.35 | 0 | 0 | 1.00 | 0.99 | 13 | B Cells |
| RPS8 | 0.33 | 0 | 0 | 1.00 | 0.99 | 13 | B Cells |
| UBA52 | 0.32 | 0 | 0 | 0.98 | 0.95 | 13 | B Cells |
| RPL35 | 0.32 | 0 | 0 | 0.98 | 0.98 | 13 | B Cells |
| RPLP1 | 0.32 | 0 | 0 | 1.00 | 1.00 | 13 | B Cells |
| RPS25 | 0.32 | 0 | 0 | 0.96 | 0.98 | 13 | B Cells |
| RPL36 | 0.31 | 0 | 0 | 1.00 | 0.99 | 13 | B Cells |
| RPS6 | 0.31 | 0 | 0 | 0.95 | 0.99 | 13 | B Cells |
| RPL31 | 0.3 | 0 | 0 | 0.98 | 0.98 | 13 | B Cells |
| SPI1 | 0.29 | 0 | 0 | 0.28 | 0.07 | 13 | B Cells |
| RPL8 | 0.28 | 0 | 0 | 0.98 | 0.99 | 13 | B Cells |
| RPS9 | 0.28 | 0 | 0 | 0.99 | 0.98 | 13 | B Cells |
| RPS4X | 0.27 | 0 | 0 | 0.99 | 0.99 | 13 | B Cells |
| SFTPC | 4.13 | 0 | 0 | 0.40 | 0.09 | 14 | AT2 Epithelial Cells |
| SFTPB | 3.51 | 0 | 0 | 0.62 | 0.08 | 14 | AT2 Epithelial Cells |
| AGER | 3.24 | 0 | 0 | 0.82 | 0.04 | 14 | AT2 Epithelial Cells |
| SFTPA1 | 2.57 | 0 | 0 | 0.32 | 0.02 | 14 | AT2 Epithelial Cells |
| KRT19 | 2.43 | 0 | 0 | 0.87 | 0.02 | 14 | AT2 Epithelial Cells |
| GPRC5A | 2.27 | 0 | 0 | 0.86 | 0.02 | 14 | AT2 Epithelial Cells |
| KRT8 | 2.14 | 0 | 0 | 0.83 | 0.07 | 14 | AT2 Epithelial Cells |
| NAPSA | 2.11 | 0 | 0 | 0.49 | 0.02 | 14 | AT2 Epithelial Cells |
| ELF3 | 2.11 | 0 | 0 | 0.82 | 0.02 | 14 | AT2 Epithelial Cells |
| FOLR1 | 2.09 | 0 | 0 | 0.86 | 0.04 | 14 | AT2 Epithelial Cells |
| SLC34A2 | 2.03 | 0 | 0 | 0.58 | 0.01 | 14 | AT2 Epithelial Cells |
| KRT18 | 2.03 | 0 | 0 | 0.83 | 0.12 | 14 | AT2 Epithelial Cells |
| FXYD3 | 2.02 | 0 | 0 | 0.89 | 0.02 | 14 | AT2 Epithelial Cells |
| C19orf33 | 2.02 | 0 | 0 | 0.82 | 0.02 | 14 | AT2 Epithelial Cells |
| LMO7 | 2 | 0 | 0 | 0.81 | 0.07 | 14 | AT2 Epithelial Cells |
| CLIC3 | 1.97 | 0 | 0 | 0.74 | 0.05 | 14 | AT2 Epithelial Cells |
| TACSTD2 | 1.89 | 0 | 0 | 0.74 | 0.02 | 14 | AT2 Epithelial Cells |
| AQP4 | 1.89 | 0 | 0 | 0.78 | 0.01 | 14 | AT2 Epithelial Cells |
| AGR3 | 1.87 | 0 | 0 | 0.46 | 0.02 | 14 | AT2 Epithelial Cells |
| KRT7 | 1.86 | 0 | 0 | 0.53 | 0.01 | 14 | AT2 Epithelial Cells |
| CLDN18 | 1.85 | 0 | 0 | 0.81 | 0.01 | 14 | AT2 Epithelial Cells |
| CLDN4 | 1.82 | 0 | 0 | 0.76 | 0.02 | 14 | AT2 Epithelial Cells |
| HOPX | 1.81 | 0 | 0 | 0.83 | 0.11 | 14 | AT2 Epithelial Cells |
| MSLN | 1.8 | 0 | 0 | 0.71 | 0.01 | 14 | AT2 Epithelial Cells |
| TSPAN13 | 1.74 | 0 | 0 | 0.83 | 0.13 | 14 | AT2 Epithelial Cells |
| MUC1 | 1.69 | 0 | 0 | 0.64 | 0.01 | 14 | AT2 Epithelial Cells |
| NKX2-1 | 1.69 | 0 | 0 | 0.85 | 0.01 | 14 | AT2 Epithelial Cells |
| EPCAM | 1.68 | 0 | 0 | 0.78 | 0.02 | 14 | AT2 Epithelial Cells |
| NPC2 | 1.63 | 0 | 0 | 0.72 | 0.50 | 14 | AT2 Epithelial Cells |
| SFTPD | 1.51 | 0 | 0 | 0.42 | 0.01 | 14 | AT2 Epithelial Cells |
| AGR2 | 1.51 | 0 | 0 | 0.54 | 0.02 | 14 | AT2 Epithelial Cells |
| EMP2 | 1.5 | 0 | 0 | 0.89 | 0.49 | 14 | AT2 Epithelial Cells |
| RTKN2 | 1.5 | 0 | 0 | 0.45 | 0.01 | 14 | AT2 Epithelial Cells |
| CPM | 1.49 | 0 | 0 | 0.73 | 0.04 | 14 | AT2 Epithelial Cells |
| CD55 | 1.48 | 0 | 0 | 0.72 | 0.12 | 14 | AT2 Epithelial Cells |
| S100A10 | 1.47 | 0 | 0 | 0.95 | 0.65 | 14 | AT2 Epithelial Cells |
| SLPI | 1.46 | 0 | 0 | 0.37 | 0.04 | 14 | AT2 Epithelial Cells |
| SFTA2 | 1.46 | 0 | 0 | 0.64 | 0.01 | 14 | AT2 Epithelial Cells |
| HIST1H1C | 1.41 | 0 | 0 | 0.45 | 0.09 | 14 | AT2 Epithelial Cells |
| CLDN7 | 1.4 | 0 | 0 | 0.56 | 0.03 | 14 | AT2 Epithelial Cells |
| ANXA2 | 1.38 | 0 | 0 | 0.99 | 0.65 | 14 | AT2 Epithelial Cells |
| SDC4 | 1.38 | 0 | 0 | 0.59 | 0.04 | 14 | AT2 Epithelial Cells |
| ANXA3 | 1.36 | 0 | 0 | 0.68 | 0.12 | 14 | AT2 Epithelial Cells |
| MGST1 | 1.35 | 0 | 0 | 0.64 | 0.03 | 14 | AT2 Epithelial Cells |
| TNNC1 | 1.35 | 0 | 0 | 0.39 | 0.01 | 14 | AT2 Epithelial Cells |
| CYP4B1 | 1.32 | 0 | 0 | 0.44 | 0.01 | 14 | AT2 Epithelial Cells |
| SDC1 | 1.31 | 0 | 0 | 0.63 | 0.02 | 14 | AT2 Epithelial Cells |
| ATP1B1 | 1.3 | 0 | 0 | 0.78 | 0.22 | 14 | AT2 Epithelial Cells |
| HSD17B6 | 1.29 | 0 | 0 | 0.58 | 0.01 | 14 | AT2 Epithelial Cells |
| S100A14 | 1.29 | 0 | 0 | 0.60 | 0.01 | 14 | AT2 Epithelial Cells |
| SPINT2 | 1.28 | 0 | 0 | 0.80 | 0.14 | 14 | AT2 Epithelial Cells |
| ID1 | 1.22 | 0 | 0 | 0.91 | 0.34 | 14 | AT2 Epithelial Cells |
| F3 | 1.22 | 0 | 0 | 0.49 | 0.03 | 14 | AT2 Epithelial Cells |
| EFEMP1 | 1.21 | 0 | 0 | 0.67 | 0.07 | 14 | AT2 Epithelial Cells |
| MRPL14 | 1.2 | 0 | 0 | 0.74 | 0.23 | 14 | AT2 Epithelial Cells |
| ATP11A | 1.2 | 0 | 0 | 0.68 | 0.05 | 14 | AT2 Epithelial Cells |
| C3 | 1.17 | 0 | 0 | 0.40 | 0.04 | 14 | AT2 Epithelial Cells |
| SAT1 | 1.16 | 0 | 0 | 0.89 | 0.56 | 14 | AT2 Epithelial Cells |
| MALL | 1.16 | 0 | 0 | 0.47 | 0.05 | 14 | AT2 Epithelial Cells |
| EFNA1 | 1.16 | 0 | 0 | 0.78 | 0.14 | 14 | AT2 Epithelial Cells |
| CD47 | 1.14 | 0 | 0 | 0.68 | 0.21 | 14 | AT2 Epithelial Cells |
| SPINT1 | 1.14 | 0 | 0 | 0.60 | 0.02 | 14 | AT2 Epithelial Cells |
| RAB11FIP1 | 1.14 | 0 | 0 | 0.54 | 0.04 | 14 | AT2 Epithelial Cells |
| ATP1A1 | 1.14 | 0 | 0 | 0.77 | 0.21 | 14 | AT2 Epithelial Cells |
| CYB5A | 1.13 | 0 | 0 | 0.87 | 0.35 | 14 | AT2 Epithelial Cells |
| TSTD1 | 1.13 | 0 | 0 | 0.68 | 0.07 | 14 | AT2 Epithelial Cells |
| AQP5 | 1.12 | 0 | 0 | 0.42 | 0.01 | 14 | AT2 Epithelial Cells |
| VAMP8 | 1.12 | 0 | 0 | 0.80 | 0.18 | 14 | AT2 Epithelial Cells |
| ICAM1 | 1.1 | 0 | 0 | 0.64 | 0.10 | 14 | AT2 Epithelial Cells |
| SELENBP1 | 1.1 | 0 | 0 | 0.67 | 0.17 | 14 | AT2 Epithelial Cells |
| IFT57 | 1.09 | 0 | 0 | 0.59 | 0.21 | 14 | AT2 Epithelial Cells |
| CTSH | 1.09 | 0 | 0 | 0.53 | 0.14 | 14 | AT2 Epithelial Cells |
| NDNF | 1.09 | 0 | 0 | 0.62 | 0.10 | 14 | AT2 Epithelial Cells |
| PRSS8 | 1.07 | 0 | 0 | 0.51 | 0.00 | 14 | AT2 Epithelial Cells |
| PPDPF | 1.05 | 0 | 0 | 0.96 | 0.80 | 14 | AT2 Epithelial Cells |
| ANKRD29 | 1.05 | 0 | 0 | 0.44 | 0.01 | 14 | AT2 Epithelial Cells |
| DUSP1 | 1.04 | 0 | 0 | 0.67 | 0.41 | 14 | AT2 Epithelial Cells |
| MMP7 | 1.04 | 0 | 0 | 0.30 | 0.01 | 14 | AT2 Epithelial Cells |
| CXADR | 1.02 | 0 | 0 | 0.53 | 0.03 | 14 | AT2 Epithelial Cells |
| CADM1 | 1.02 | 0 | 0 | 0.63 | 0.17 | 14 | AT2 Epithelial Cells |
| VSIG2 | 1.01 | 0 | 0 | 0.54 | 0.02 | 14 | AT2 Epithelial Cells |
| DMKN | 0.98 | 0 | 0 | 0.60 | 0.03 | 14 | AT2 Epithelial Cells |
| GJA1 | 0.97 | 0 | 0 | 0.50 | 0.04 | 14 | AT2 Epithelial Cells |
| VEGFA | 0.97 | 0 | 0 | 0.50 | 0.05 | 14 | AT2 Epithelial Cells |
| SEMA3B | 0.96 | 0 | 0 | 0.41 | 0.03 | 14 | AT2 Epithelial Cells |
| GSTP1 | 0.96 | 0 | 0 | 0.97 | 0.83 | 14 | AT2 Epithelial Cells |
| SLC39A8 | 0.95 | 0 | 0 | 0.46 | 0.02 | 14 | AT2 Epithelial Cells |
| COL4A4 | 0.95 | 0 | 0 | 0.35 | 0.01 | 14 | AT2 Epithelial Cells |
| SMARCA5 | 0.95 | 0 | 0 | 0.64 | 0.20 | 14 | AT2 Epithelial Cells |
| PERP | 0.94 | 0 | 0 | 0.56 | 0.12 | 14 | AT2 Epithelial Cells |
| PEG10 | 0.94 | 0 | 0 | 0.44 | 0.08 | 14 | AT2 Epithelial Cells |
| LAMP3 | 0.94 | 0 | 0 | 0.39 | 0.01 | 14 | AT2 Epithelial Cells |
| ICAM4 | 0.93 | 0 | 0 | 0.35 | 0.01 | 14 | AT2 Epithelial Cells |
| PGC | 0.93 | 0 | 0 | 0.27 | 0.00 | 14 | AT2 Epithelial Cells |
| AQP3 | 0.92 | 0 | 0 | 0.37 | 0.05 | 14 | AT2 Epithelial Cells |
| RAB25 | 0.92 | 0 | 0 | 0.55 | 0.01 | 14 | AT2 Epithelial Cells |
| KLF5 | 0.91 | 0 | 0 | 0.49 | 0.01 | 14 | AT2 Epithelial Cells |
| RNU12.1 | 0.91 | 0 | 0 | 0.40 | 0.11 | 14 | AT2 Epithelial Cells |
| ERBB3 | 0.91 | 0 | 0 | 0.55 | 0.02 | 14 | AT2 Epithelial Cells |
| PTP4A1 | 0.9 | 0 | 0 | 0.53 | 0.12 | 14 | AT2 Epithelial Cells |
| MT-ND3 | 0.89 | 0 | 0 | 1.00 | 0.98 | 14 | AT2 Epithelial Cells |
| ITGA3 | 0.88 | 0 | 0 | 0.54 | 0.11 | 14 | AT2 Epithelial Cells |
| LAMA3 | 0.88 | 0 | 0 | 0.36 | 0.01 | 14 | AT2 Epithelial Cells |
| ASAH1 | 0.87 | 0 | 0 | 0.72 | 0.29 | 14 | AT2 Epithelial Cells |
| C1orf198 | 0.87 | 0 | 0 | 0.49 | 0.13 | 14 | AT2 Epithelial Cells |
| ABCA3 | 0.87 | 0 | 0 | 0.35 | 0.02 | 14 | AT2 Epithelial Cells |
| UPK3B | 0.87 | 0 | 0 | 0.26 | 0.00 | 14 | AT2 Epithelial Cells |
| APLP2 | 0.87 | 0 | 0 | 0.89 | 0.57 | 14 | AT2 Epithelial Cells |
| CYSTM1 | 0.86 | 0 | 0 | 0.71 | 0.33 | 14 | AT2 Epithelial Cells |
| H19 | 0.86 | 0 | 0 | 0.47 | 0.16 | 14 | AT2 Epithelial Cells |
| CRNDE | 0.86 | 0 | 0 | 0.50 | 0.01 | 14 | AT2 Epithelial Cells |
| MBIP | 0.85 | 0 | 0 | 0.55 | 0.10 | 14 | AT2 Epithelial Cells |
| TMEM125 | 0.85 | 0 | 0 | 0.49 | 0.02 | 14 | AT2 Epithelial Cells |
| PTPRF | 0.85 | 0 | 0 | 0.54 | 0.06 | 14 | AT2 Epithelial Cells |
| HES1 | 0.83 | 0 | 0 | 0.83 | 0.38 | 14 | AT2 Epithelial Cells |
| LAMA5 | 0.83 | 0 | 0 | 0.44 | 0.11 | 14 | AT2 Epithelial Cells |
| WIF1 | 0.83 | 0 | 0 | 0.37 | 0.05 | 14 | AT2 Epithelial Cells |
| C1orf116 | 0.83 | 0 | 0 | 0.49 | 0.01 | 14 | AT2 Epithelial Cells |
| RASSF7 | 0.82 | 0 | 0 | 0.44 | 0.02 | 14 | AT2 Epithelial Cells |
| DLC1 | 0.82 | 0 | 0 | 0.69 | 0.23 | 14 | AT2 Epithelial Cells |
| IRX2 | 0.81 | 0 | 0 | 0.40 | 0.01 | 14 | AT2 Epithelial Cells |
| LIPH | 0.81 | 0 | 0 | 0.37 | 0.00 | 14 | AT2 Epithelial Cells |
| CD9 | 0.8 | 0 | 0 | 0.89 | 0.45 | 14 | AT2 Epithelial Cells |
| PLD3 | 0.8 | 0 | 0 | 0.78 | 0.35 | 14 | AT2 Epithelial Cells |
| ZFAS1 | 0.8 | 0 | 0 | 0.91 | 0.59 | 14 | AT2 Epithelial Cells |
| SUSD2 | 0.79 | 0 | 0 | 0.40 | 0.01 | 14 | AT2 Epithelial Cells |
| CXCL17 | 0.79 | 0 | 0 | 0.30 | 0.00 | 14 | AT2 Epithelial Cells |
| PEBP4 | 0.79 | 0 | 0 | 0.36 | 0.01 | 14 | AT2 Epithelial Cells |
| ALCAM | 0.79 | 0 | 0 | 0.47 | 0.08 | 14 | AT2 Epithelial Cells |
| PHACTR1 | 0.79 | 0 | 0 | 0.41 | 0.04 | 14 | AT2 Epithelial Cells |
| CLDN3 | 0.79 | 0 | 0 | 0.45 | 0.01 | 14 | AT2 Epithelial Cells |
| BCAM | 0.78 | 0 | 0 | 0.89 | 0.33 | 14 | AT2 Epithelial Cells |
| BANCR | 0.78 | 0 | 0 | 0.27 | 0.00 | 14 | AT2 Epithelial Cells |
| EVA1A | 0.78 | 0 | 0 | 0.45 | 0.01 | 14 | AT2 Epithelial Cells |
| LGALS3 | 0.78 | 0 | 0 | 0.67 | 0.37 | 14 | AT2 Epithelial Cells |
| ADIRF | 0.78 | 0 | 0 | 0.95 | 0.50 | 14 | AT2 Epithelial Cells |
| EPS8L2 | 0.78 | 0 | 0 | 0.40 | 0.01 | 14 | AT2 Epithelial Cells |
| HIST1H2BG | 0.77 | 0 | 0 | 0.50 | 0.14 | 14 | AT2 Epithelial Cells |
| SFTA3 | 0.77 | 0 | 0 | 0.41 | 0.01 | 14 | AT2 Epithelial Cells |
| PRDX5 | 0.76 | 0 | 0 | 0.86 | 0.56 | 14 | AT2 Epithelial Cells |
| CDH1 | 0.76 | 0 | 0 | 0.37 | 0.01 | 14 | AT2 Epithelial Cells |
| GKN2 | 0.76 | 0 | 0 | 0.32 | 0.00 | 14 | AT2 Epithelial Cells |
| MPZL2 | 0.76 | 0 | 0 | 0.40 | 0.01 | 14 | AT2 Epithelial Cells |
| LAMB3 | 0.75 | 0 | 0 | 0.36 | 0.01 | 14 | AT2 Epithelial Cells |
| FAM129B | 0.75 | 0 | 0 | 0.53 | 0.10 | 14 | AT2 Epithelial Cells |
| ANXA4 | 0.75 | 0 | 0 | 0.53 | 0.16 | 14 | AT2 Epithelial Cells |
| FILIP1 | 0.75 | 0 | 0 | 0.47 | 0.18 | 14 | AT2 Epithelial Cells |
| RNASE1 | 0.75 | 0 | 0 | 0.80 | 0.27 | 14 | AT2 Epithelial Cells |
| C5orf38 | 0.73 | 0 | 0 | 0.35 | 0.01 | 14 | AT2 Epithelial Cells |
| TMPRSS2 | 0.72 | 0 | 0 | 0.33 | 0.00 | 14 | AT2 Epithelial Cells |
| PDPN | 0.72 | 0 | 0 | 0.39 | 0.05 | 14 | AT2 Epithelial Cells |
| WFDC2 | 0.71 | 0 | 0 | 0.37 | 0.01 | 14 | AT2 Epithelial Cells |
| CLIC5 | 0.7 | 0 | 0 | 0.37 | 0.06 | 14 | AT2 Epithelial Cells |
| MAL2 | 0.69 | 0 | 0 | 0.40 | 0.01 | 14 | AT2 Epithelial Cells |
| MT-CO3 | 0.69 | 0 | 0 | 1.00 | 0.99 | 14 | AT2 Epithelial Cells |
| FLRT3 | 0.69 | 0 | 0 | 0.39 | 0.01 | 14 | AT2 Epithelial Cells |
| ANOS1 | 0.69 | 0 | 0 | 0.26 | 0.02 | 14 | AT2 Epithelial Cells |
| GLS | 0.69 | 0 | 0 | 0.42 | 0.10 | 14 | AT2 Epithelial Cells |
| MT-CO2 | 0.69 | 0 | 0 | 1.00 | 0.99 | 14 | AT2 Epithelial Cells |
| CKB | 0.69 | 0 | 0 | 0.69 | 0.22 | 14 | AT2 Epithelial Cells |
| SECISBP2L | 0.69 | 0 | 0 | 0.47 | 0.11 | 14 | AT2 Epithelial Cells |
| MAGI3 | 0.68 | 0 | 0 | 0.36 | 0.03 | 14 | AT2 Epithelial Cells |
| MT-ND1 | 0.68 | 0 | 0 | 0.99 | 0.97 | 14 | AT2 Epithelial Cells |
| LAMC2 | 0.68 | 0 | 0 | 0.28 | 0.01 | 14 | AT2 Epithelial Cells |
| UTRN | 0.68 | 0 | 0 | 0.47 | 0.18 | 14 | AT2 Epithelial Cells |
| GOLGA8B | 0.67 | 0 | 0 | 0.33 | 0.04 | 14 | AT2 Epithelial Cells |
| MT-ND4 | 0.67 | 0 | 0 | 1.00 | 1.00 | 14 | AT2 Epithelial Cells |
| MT-CO1 | 0.67 | 0 | 0 | 0.99 | 1.00 | 14 | AT2 Epithelial Cells |
| TKT | 0.66 | 0 | 0 | 0.64 | 0.26 | 14 | AT2 Epithelial Cells |
| STARD10 | 0.66 | 0 | 0 | 0.47 | 0.07 | 14 | AT2 Epithelial Cells |
| HSPG2 | 0.66 | 0 | 0 | 0.47 | 0.19 | 14 | AT2 Epithelial Cells |
| EPB41L5 | 0.65 | 0 | 0 | 0.30 | 0.03 | 14 | AT2 Epithelial Cells |
| TMEM63B | 0.65 | 0 | 0 | 0.36 | 0.03 | 14 | AT2 Epithelial Cells |
| YPEL3 | 0.65 | 0 | 0 | 0.53 | 0.18 | 14 | AT2 Epithelial Cells |
| CD24 | 0.65 | 0 | 0 | 0.35 | 0.02 | 14 | AT2 Epithelial Cells |
| TMC5 | 0.65 | 0 | 0 | 0.40 | 0.01 | 14 | AT2 Epithelial Cells |
| OCLN | 0.64 | 0 | 0 | 0.45 | 0.07 | 14 | AT2 Epithelial Cells |
| AGPAT2 | 0.64 | 0 | 0 | 0.50 | 0.13 | 14 | AT2 Epithelial Cells |
| MT-CYB | 0.63 | 0 | 0 | 0.99 | 0.99 | 14 | AT2 Epithelial Cells |
| MTATP6P1 | 0.63 | 0 | 0 | 1.00 | 0.98 | 14 | AT2 Epithelial Cells |
| ST3GAL5 | 0.63 | 0 | 0 | 0.30 | 0.02 | 14 | AT2 Epithelial Cells |
| C16orf89 | 0.62 | 0 | 0 | 0.27 | 0.05 | 14 | AT2 Epithelial Cells |
| HPCAL1 | 0.62 | 0 | 0 | 0.50 | 0.16 | 14 | AT2 Epithelial Cells |
| PPL | 0.62 | 0 | 0 | 0.33 | 0.01 | 14 | AT2 Epithelial Cells |
| FOXA2 | 0.62 | 0 | 0 | 0.26 | 0.00 | 14 | AT2 Epithelial Cells |
| IRX3 | 0.62 | 0 | 0 | 0.31 | 0.01 | 14 | AT2 Epithelial Cells |
| AGRN | 0.62 | 0 | 0 | 0.35 | 0.07 | 14 | AT2 Epithelial Cells |
| LPCAT1 | 0.62 | 0 | 0 | 0.41 | 0.06 | 14 | AT2 Epithelial Cells |
| BAIAP2 | 0.61 | 0 | 0 | 0.41 | 0.08 | 14 | AT2 Epithelial Cells |
| S100A6 | 0.61 | 0 | 0 | 1.00 | 0.84 | 14 | AT2 Epithelial Cells |
| RIN2 | 0.61 | 0 | 0 | 0.35 | 0.12 | 14 | AT2 Epithelial Cells |
| MT-ND2 | 0.61 | 0 | 0 | 1.00 | 0.99 | 14 | AT2 Epithelial Cells |
| KCNJ15 | 0.6 | 0 | 0 | 0.32 | 0.01 | 14 | AT2 Epithelial Cells |
| PATJ | 0.6 | 0 | 0 | 0.30 | 0.02 | 14 | AT2 Epithelial Cells |
| SHROOM3 | 0.6 | 0 | 0 | 0.33 | 0.02 | 14 | AT2 Epithelial Cells |
| UNC13D | 0.6 | 0 | 0 | 0.27 | 0.03 | 14 | AT2 Epithelial Cells |
| AKR1C1 | 0.6 | 0 | 0 | 0.27 | 0.01 | 14 | AT2 Epithelial Cells |
| NCOA7 | 0.59 | 0 | 0 | 0.35 | 0.03 | 14 | AT2 Epithelial Cells |
| SESN1 | 0.59 | 0 | 0 | 0.32 | 0.08 | 14 | AT2 Epithelial Cells |
| ERRFI1 | 0.59 | 0 | 0 | 0.32 | 0.04 | 14 | AT2 Epithelial Cells |
| F11R | 0.59 | 0 | 0 | 0.40 | 0.07 | 14 | AT2 Epithelial Cells |
| NHSL1 | 0.58 | 0 | 0 | 0.31 | 0.04 | 14 | AT2 Epithelial Cells |
| NEDD4L | 0.58 | 0 | 0 | 0.45 | 0.06 | 14 | AT2 Epithelial Cells |
| WWC2 | 0.58 | 0 | 0 | 0.30 | 0.06 | 14 | AT2 Epithelial Cells |
| PRDX1 | 0.58 | 0 | 0 | 0.87 | 0.66 | 14 | AT2 Epithelial Cells |
| ORMDL1 | 0.58 | 0 | 0.01 | 0.45 | 0.23 | 14 | AT2 Epithelial Cells |
| PDGFA | 0.57 | 0 | 0 | 0.33 | 0.06 | 14 | AT2 Epithelial Cells |
| PCYOX1 | 0.57 | 0 | 0 | 0.39 | 0.10 | 14 | AT2 Epithelial Cells |
| SLC2A3 | 0.57 | 0 | 0 | 0.42 | 0.18 | 14 | AT2 Epithelial Cells |
| PPP1R14C | 0.57 | 0 | 0 | 0.31 | 0.01 | 14 | AT2 Epithelial Cells |
| NECAB3 | 0.56 | 0 | 0 | 0.30 | 0.03 | 14 | AT2 Epithelial Cells |
| MT-ATP6 | 0.55 | 0 | 0 | 1.00 | 0.99 | 14 | AT2 Epithelial Cells |
| RP11-357D18.1 | 0.55 | 0 | 0 | 0.26 | 0.00 | 14 | AT2 Epithelial Cells |
| GOLGA8A | 0.54 | 0 | 0 | 0.32 | 0.07 | 14 | AT2 Epithelial Cells |
| MMP28 | 0.54 | 0 | 0 | 0.26 | 0.02 | 14 | AT2 Epithelial Cells |
| NSG1 | 0.54 | 0 | 0 | 0.31 | 0.01 | 14 | AT2 Epithelial Cells |
| PLLP | 0.54 | 0 | 0 | 0.27 | 0.03 | 14 | AT2 Epithelial Cells |
| KLHL24 | 0.54 | 0 | 0 | 0.36 | 0.10 | 14 | AT2 Epithelial Cells |
| FAM20A | 0.54 | 0 | 0 | 0.27 | 0.02 | 14 | AT2 Epithelial Cells |
| CCDC12 | 0.54 | 0 | 0 | 0.56 | 0.28 | 14 | AT2 Epithelial Cells |
| POLR2L | 0.53 | 0 | 0 | 0.89 | 0.67 | 14 | AT2 Epithelial Cells |
| LGALS3BP | 0.53 | 0 | 0 | 0.50 | 0.20 | 14 | AT2 Epithelial Cells |
| COX7C | 0.53 | 0 | 0 | 0.97 | 0.87 | 14 | AT2 Epithelial Cells |
| ABLIM1 | 0.53 | 0 | 0 | 0.32 | 0.07 | 14 | AT2 Epithelial Cells |
| MRPS36 | 0.52 | 0 | 0 | 0.51 | 0.25 | 14 | AT2 Epithelial Cells |
| RHOF | 0.52 | 0 | 0 | 0.36 | 0.06 | 14 | AT2 Epithelial Cells |
| ITGB6 | 0.52 | 0 | 0 | 0.27 | 0.01 | 14 | AT2 Epithelial Cells |
| TUBA1C | 0.52 | 0 | 0 | 0.40 | 0.14 | 14 | AT2 Epithelial Cells |
| P3H2 | 0.52 | 0 | 0 | 0.33 | 0.03 | 14 | AT2 Epithelial Cells |
| ATP8A1 | 0.51 | 0 | 0 | 0.28 | 0.02 | 14 | AT2 Epithelial Cells |
| LRRFIP1 | 0.51 | 0 | 0 | 0.59 | 0.27 | 14 | AT2 Epithelial Cells |
| MISP | 0.51 | 0 | 0 | 0.28 | 0.00 | 14 | AT2 Epithelial Cells |
| HMGA1 | 0.51 | 0 | 0 | 0.45 | 0.10 | 14 | AT2 Epithelial Cells |
| ST6GALNAC5 | 0.51 | 0 | 0 | 0.26 | 0.01 | 14 | AT2 Epithelial Cells |
| SOX11 | 0.5 | 0 | 0 | 0.35 | 0.10 | 14 | AT2 Epithelial Cells |
| CMTM8 | 0.5 | 0 | 0 | 0.40 | 0.06 | 14 | AT2 Epithelial Cells |
| RBM47 | 0.5 | 0 | 0 | 0.33 | 0.03 | 14 | AT2 Epithelial Cells |
| SPRYD7 | 0.49 | 0 | 0 | 0.27 | 0.07 | 14 | AT2 Epithelial Cells |
| MT-ND5 | 0.49 | 0 | 0.01 | 0.96 | 0.89 | 14 | AT2 Epithelial Cells |
| INTS6 | 0.49 | 0 | 0 | 0.47 | 0.20 | 14 | AT2 Epithelial Cells |
| PCBD1 | 0.49 | 0 | 0 | 0.51 | 0.21 | 14 | AT2 Epithelial Cells |
| ERBB2 | 0.49 | 0 | 0 | 0.36 | 0.09 | 14 | AT2 Epithelial Cells |
| DDAH1 | 0.48 | 0 | 0 | 0.36 | 0.03 | 14 | AT2 Epithelial Cells |
| DHCR24 | 0.48 | 0 | 0 | 0.31 | 0.01 | 14 | AT2 Epithelial Cells |
| NFKBIZ | 0.48 | 0 | 0 | 0.28 | 0.08 | 14 | AT2 Epithelial Cells |
| CEBPD | 0.48 | 0 | 0 | 0.58 | 0.30 | 14 | AT2 Epithelial Cells |
| CRB3 | 0.48 | 0 | 0 | 0.26 | 0.00 | 14 | AT2 Epithelial Cells |
| PDXK | 0.48 | 0 | 0 | 0.33 | 0.08 | 14 | AT2 Epithelial Cells |
| ANXA1 | 0.47 | 0 | 0 | 0.74 | 0.40 | 14 | AT2 Epithelial Cells |
| TUBB4B | 0.47 | 0 | 0 | 0.71 | 0.40 | 14 | AT2 Epithelial Cells |
| CAT | 0.47 | 0 | 0 | 0.53 | 0.25 | 14 | AT2 Epithelial Cells |
| DSP | 0.47 | 0 | 0 | 0.33 | 0.02 | 14 | AT2 Epithelial Cells |
| CTSD | 0.47 | 0 | 0 | 0.68 | 0.31 | 14 | AT2 Epithelial Cells |
| TMEM87A | 0.46 | 0 | 0 | 0.47 | 0.22 | 14 | AT2 Epithelial Cells |
| GNG5 | 0.46 | 0 | 0 | 0.77 | 0.53 | 14 | AT2 Epithelial Cells |
| PON2 | 0.46 | 0 | 0 | 0.58 | 0.24 | 14 | AT2 Epithelial Cells |
| GADD45B | 0.46 | 0 | 0 | 0.73 | 0.42 | 14 | AT2 Epithelial Cells |
| CXCL16 | 0.46 | 0 | 0 | 0.40 | 0.08 | 14 | AT2 Epithelial Cells |
| HSBP1L1 | 0.46 | 0 | 0 | 0.36 | 0.04 | 14 | AT2 Epithelial Cells |
| ARL4C | 0.45 | 0 | 0 | 0.36 | 0.07 | 14 | AT2 Epithelial Cells |
| BTG3 | 0.45 | 0 | 0 | 0.53 | 0.24 | 14 | AT2 Epithelial Cells |
| BLVRB | 0.45 | 0 | 0 | 0.51 | 0.21 | 14 | AT2 Epithelial Cells |
| TMEM30B | 0.45 | 0 | 0 | 0.28 | 0.01 | 14 | AT2 Epithelial Cells |
| TJP2 | 0.45 | 0 | 0 | 0.32 | 0.11 | 14 | AT2 Epithelial Cells |
| ISOC1 | 0.45 | 0 | 0 | 0.33 | 0.09 | 14 | AT2 Epithelial Cells |
| MYO9A | 0.45 | 0 | 0 | 0.32 | 0.10 | 14 | AT2 Epithelial Cells |
| ADK | 0.44 | 0 | 0 | 0.39 | 0.08 | 14 | AT2 Epithelial Cells |
| SLC7A8 | 0.44 | 0 | 0 | 0.26 | 0.03 | 14 | AT2 Epithelial Cells |
| HEXIM1 | 0.44 | 0 | 0.03 | 0.50 | 0.26 | 14 | AT2 Epithelial Cells |
| CTNNBIP1 | 0.44 | 0 | 0 | 0.41 | 0.11 | 14 | AT2 Epithelial Cells |
| PLXNB2 | 0.43 | 0 | 0 | 0.36 | 0.12 | 14 | AT2 Epithelial Cells |
| DDR1 | 0.43 | 0 | 0 | 0.32 | 0.04 | 14 | AT2 Epithelial Cells |
| GRAMD1A | 0.43 | 0 | 0 | 0.45 | 0.14 | 14 | AT2 Epithelial Cells |
| SH3YL1 | 0.43 | 0 | 0 | 0.32 | 0.09 | 14 | AT2 Epithelial Cells |
| CCND3 | 0.43 | 0 | 0 | 0.44 | 0.15 | 14 | AT2 Epithelial Cells |
| SNHG16 | 0.43 | 0 | 0 | 0.50 | 0.23 | 14 | AT2 Epithelial Cells |
| BEX2 | 0.43 | 0 | 0 | 0.26 | 0.03 | 14 | AT2 Epithelial Cells |
| SSR2 | 0.42 | 0 | 0.01 | 0.86 | 0.72 | 14 | AT2 Epithelial Cells |
| LPCAT3 | 0.42 | 0 | 0 | 0.35 | 0.10 | 14 | AT2 Epithelial Cells |
| JUP | 0.42 | 0 | 0 | 0.41 | 0.17 | 14 | AT2 Epithelial Cells |
| LRBA | 0.42 | 0 | 0 | 0.31 | 0.07 | 14 | AT2 Epithelial Cells |
| RPL39P3 | 0.41 | 0 | 0 | 0.62 | 0.34 | 14 | AT2 Epithelial Cells |
| MSMO1 | 0.41 | 0 | 0 | 0.30 | 0.08 | 14 | AT2 Epithelial Cells |
| STT3B | 0.41 | 0 | 0 | 0.46 | 0.18 | 14 | AT2 Epithelial Cells |
| TSPO | 0.41 | 0 | 0 | 0.76 | 0.43 | 14 | AT2 Epithelial Cells |
| SCD5 | 0.41 | 0 | 0 | 0.30 | 0.08 | 14 | AT2 Epithelial Cells |
| ADIPOR1 | 0.41 | 0 | 0 | 0.42 | 0.17 | 14 | AT2 Epithelial Cells |
| RBPMS | 0.41 | 0 | 0 | 0.47 | 0.20 | 14 | AT2 Epithelial Cells |
| TXNDC17 | 0.41 | 0 | 0 | 0.54 | 0.27 | 14 | AT2 Epithelial Cells |
| B4GALT1 | 0.41 | 0 | 0 | 0.32 | 0.11 | 14 | AT2 Epithelial Cells |
| RP11-532F12.5 | 0.41 | 0 | 0 | 0.26 | 0.01 | 14 | AT2 Epithelial Cells |
| HMGN3 | 0.41 | 0 | 0.02 | 0.71 | 0.49 | 14 | AT2 Epithelial Cells |
| MAPK13 | 0.4 | 0 | 0 | 0.27 | 0.02 | 14 | AT2 Epithelial Cells |
| DYNLT1 | 0.4 | 0 | 0.01 | 0.67 | 0.43 | 14 | AT2 Epithelial Cells |
| CNDP2 | 0.4 | 0 | 0 | 0.39 | 0.13 | 14 | AT2 Epithelial Cells |
| VWA1 | 0.4 | 0 | 0 | 0.32 | 0.10 | 14 | AT2 Epithelial Cells |
| CELF2 | 0.39 | 0 | 0 | 0.50 | 0.24 | 14 | AT2 Epithelial Cells |
| ICA1 | 0.39 | 0 | 0 | 0.27 | 0.04 | 14 | AT2 Epithelial Cells |
| HIST1H2AC | 0.39 | 0 | 0 | 0.27 | 0.09 | 14 | AT2 Epithelial Cells |
| KCNQ1OT1 | 0.39 | 0 | 0.03 | 0.59 | 0.31 | 14 | AT2 Epithelial Cells |
| ALDH3A2 | 0.38 | 0 | 0 | 0.26 | 0.08 | 14 | AT2 Epithelial Cells |
| RPS20 | 0.38 | 0 | 0 | 1.00 | 0.97 | 14 | AT2 Epithelial Cells |
| SNHG3 | 0.38 | 0 | 0 | 0.35 | 0.13 | 14 | AT2 Epithelial Cells |
| ADGRF5 | 0.38 | 0 | 0 | 0.46 | 0.17 | 14 | AT2 Epithelial Cells |
| TOR1AIP2 | 0.37 | 0 | 0 | 0.41 | 0.15 | 14 | AT2 Epithelial Cells |
| OCIAD2 | 0.37 | 0 | 0 | 0.42 | 0.14 | 14 | AT2 Epithelial Cells |
| HOOK2 | 0.36 | 0 | 0 | 0.39 | 0.12 | 14 | AT2 Epithelial Cells |
| C11orf1 | 0.36 | 0 | 0 | 0.26 | 0.06 | 14 | AT2 Epithelial Cells |
| PTRHD1 | 0.36 | 0 | 0 | 0.49 | 0.18 | 14 | AT2 Epithelial Cells |
| RPL39 | 0.36 | 0 | 0 | 0.99 | 0.99 | 14 | AT2 Epithelial Cells |
| MYO6 | 0.36 | 0 | 0 | 0.44 | 0.14 | 14 | AT2 Epithelial Cells |
| ANXA2P2 | 0.35 | 0 | 0 | 0.30 | 0.09 | 14 | AT2 Epithelial Cells |
| FNBP1L | 0.35 | 0 | 0.02 | 0.26 | 0.10 | 14 | AT2 Epithelial Cells |
| WBP2 | 0.35 | 0 | 0.01 | 0.41 | 0.18 | 14 | AT2 Epithelial Cells |
| TMC4 | 0.35 | 0 | 0 | 0.28 | 0.01 | 14 | AT2 Epithelial Cells |
| PLEKHJ1 | 0.34 | 0 | 0.05 | 0.40 | 0.19 | 14 | AT2 Epithelial Cells |
| RPL37 | 0.34 | 0 | 0 | 1.00 | 0.98 | 14 | AT2 Epithelial Cells |
| AK1 | 0.34 | 0 | 0 | 0.42 | 0.17 | 14 | AT2 Epithelial Cells |
| RAB5B | 0.34 | 0 | 0 | 0.39 | 0.14 | 14 | AT2 Epithelial Cells |
| PRKCI | 0.34 | 0 | 0 | 0.32 | 0.06 | 14 | AT2 Epithelial Cells |
| FARP1 | 0.34 | 0 | 0 | 0.36 | 0.13 | 14 | AT2 Epithelial Cells |
| ZDHHC12 | 0.34 | 0 | 0 | 0.26 | 0.06 | 14 | AT2 Epithelial Cells |
| TNFRSF12A | 0.34 | 0 | 0 | 0.42 | 0.17 | 14 | AT2 Epithelial Cells |
| LIMCH1 | 0.34 | 0 | 0 | 0.50 | 0.21 | 14 | AT2 Epithelial Cells |
| HMGN1 | 0.33 | 0 | 0.03 | 0.86 | 0.61 | 14 | AT2 Epithelial Cells |
| TES | 0.33 | 0 | 0 | 0.27 | 0.09 | 14 | AT2 Epithelial Cells |
| CPAMD8 | 0.33 | 0 | 0 | 0.27 | 0.04 | 14 | AT2 Epithelial Cells |
| PBX2 | 0.33 | 0 | 0.04 | 0.27 | 0.10 | 14 | AT2 Epithelial Cells |
| C2orf68 | 0.32 | 0 | 0.01 | 0.32 | 0.12 | 14 | AT2 Epithelial Cells |
| RPL31P49 | 0.32 | 0 | 0.04 | 0.26 | 0.10 | 14 | AT2 Epithelial Cells |
| RP1-34B20.21 | 0.32 | 0 | 0 | 0.30 | 0.07 | 14 | AT2 Epithelial Cells |
| SMC4 | 0.32 | 0 | 0 | 0.31 | 0.11 | 14 | AT2 Epithelial Cells |
| C14orf1 | 0.32 | 0 | 0.01 | 0.39 | 0.17 | 14 | AT2 Epithelial Cells |
| RAB27A | 0.31 | 0 | 0 | 0.30 | 0.10 | 14 | AT2 Epithelial Cells |
| AHR | 0.31 | 0 | 0 | 0.36 | 0.12 | 14 | AT2 Epithelial Cells |
| ODC1 | 0.31 | 0 | 0.03 | 0.26 | 0.10 | 14 | AT2 Epithelial Cells |
| RPL37A | 0.31 | 0 | 0 | 1.00 | 0.99 | 14 | AT2 Epithelial Cells |
| RPL28 | 0.3 | 0 | 0 | 1.00 | 0.98 | 14 | AT2 Epithelial Cells |
| SYNGR2 | 0.3 | 0 | 0 | 0.56 | 0.27 | 14 | AT2 Epithelial Cells |
| PIAS1 | 0.29 | 0 | 0.01 | 0.32 | 0.12 | 14 | AT2 Epithelial Cells |
| TPD52 | 0.29 | 0 | 0 | 0.26 | 0.04 | 14 | AT2 Epithelial Cells |
| PARP14 | 0.29 | 0 | 0.01 | 0.30 | 0.11 | 14 | AT2 Epithelial Cells |
| RPS19 | 0.29 | 0 | 0 | 1.00 | 0.99 | 14 | AT2 Epithelial Cells |
| PACSIN3 | 0.29 | 0 | 0.01 | 0.28 | 0.10 | 14 | AT2 Epithelial Cells |
| NDRG1 | 0.28 | 0 | 0.03 | 0.39 | 0.17 | 14 | AT2 Epithelial Cells |
| RPL24 | 0.28 | 0 | 0 | 1.00 | 0.97 | 14 | AT2 Epithelial Cells |
| NUDT14 | 0.28 | 0 | 0 | 0.30 | 0.09 | 14 | AT2 Epithelial Cells |
| WDR34 | 0.26 | 0 | 0.01 | 0.28 | 0.10 | 14 | AT2 Epithelial Cells |
| UBL3 | 0.26 | 0 | 0 | 0.33 | 0.13 | 14 | AT2 Epithelial Cells |
| PTPN1 | 0.26 | 0 | 0 | 0.26 | 0.07 | 14 | AT2 Epithelial Cells |
| USF2 | 0.25 | 0 | 0 | 0.33 | 0.12 | 14 | AT2 Epithelial Cells |

Table S3: Distribution of mouse cells across age

| **Cluster** | **Cell Type** | **Cellular Subtype** | **PND1** | **PND10** | **PND3** | **PND7** | **Total** |
| --- | --- | --- | --- | --- | --- | --- | --- |
| 0 | Endothelial | Endothelial | 603 | 1652 | 1056 | 724 | 4035 |
| 7 | Endothelial | Endothelial | 27 | 650 | 473 | 126 | 1276 |
| 8 | Endothelial | Endothelial | 1234 | 1 | 8 | 1 | 1244 |
| 14 | Endothelial | Endothelial | 56 | 534 | 213 | 299 | 1102 |
| 16 | Endothelial | Endothelial | 150 | 233 | 224 | 139 | 746 |
| 19 | Endothelial | Endothelial | 101 | 108 | 111 | 168 | 488 |
| 3 | Epithelial | AT2 Cells | 178 | 1046 | 990 | 274 | 2488 |
| 9 | Epithelial | AT2 Cells | 1080 | 36 | 106 | 18 | 1240 |
| 10 | Epithelial | AT1 Cells | 435 | 117 | 663 | 17 | 1232 |
| 17 | Epithelial | Ciliated Cells | 92 | 235 | 351 | 8 | 686 |
| 27 | Epithelial | Mixed Epithelial | 39 | 109 | 70 | 7 | 225 |
| 29 | Epithelial | AT2 Cells | 9 | 93 | 14 | 66 | 182 |
| 31 | Epithelial | AT2 Cells | 17 | 27 | 13 | 23 | 80 |
| 4 | Immune | Monocytes | 101 | 331 | 465 | 897 | 1794 |
| 5 | Immune | B-Cells | 62 | 904 | 262 | 292 | 1520 |
| 6 | Immune | T-Cells | 38 | 743 | 275 | 367 | 1520 |
| 12 | Immune | Macrophages | 292 | 59 | 606 | 177 | 1134 |
| 21 | Immune | Macrophages | 72 | 125 | 117 | 107 | 421 |
| 22 | Immune | Neutrophils | 121 | 17 | 194 | 57 | 389 |
| 25 | Immune | T-Cells | 1 | 108 | 102 | 52 | 263 |
| 28 | Immune | Myeloid Cells | 18 | 37 | 75 | 73 | 203 |
| 1 | Mesenchymal | Fibroblasts | 21 | 1075 | 765 | 984 | 2845 |
| 2 | Mesenchymal | Stromal Cells | 113 | 1289 | 285 | 840 | 2527 |
| 11 | Mesenchymal | Matrix Fibroblasts | 1216 | 0 | 1 | 0 | 1217 |
| 13 | Mesenchymal | Matrix Fibroblasts | 238 | 347 | 341 | 181 | 1107 |
| 15 | Mesenchymal | Matrix Fibroblasts | 905 | 0 | 36 | 2 | 943 |
| 18 | Mesenchymal | Matrix Fibroblasts | 256 | 170 | 94 | 159 | 679 |
| 20 | Mesenchymal | Matrix Fibroblasts | 463 | 0 | 1 | 0 | 464 |
| 23 | Mesenchymal | Matrix Fibroblasts | 15 | 73 | 113 | 83 | 284 |
| 24 | Mesenchymal | Matrix Fibroblasts | 1 | 116 | 13 | 144 | 274 |
| 26 | Mesenchymal | Fibroblasts | 6 | 168 | 41 | 24 | 239 |
| 30 | Mesenchymal | Myo-fibroblasts | 43 | 29 | 12 | 15 | 99 |
| Total |  |  | 8003 | 10432 | 8090 | 6324 | 32946 |

Table S4: Distribution of human and mouse cells by sample source and capture

| **Major Type** | **Subclass** | **Cluster** | **Lung1 Run1** | **Lung1 Run2** | **Lung2 Run1** | **Lung2 Run2** | **Human** | **PND**  **1** | **PND**  **3** | **PND**  **7** | **PND**  **10** | **Mouse** | **Total** |
| --- | --- | --- | --- | --- | --- | --- | --- | --- | --- | --- | --- | --- | --- |
| Endothelial | Endothelial | 0 | 47 | 29 | 20 | 49 | 145 | 1446 | 1094 | 771 | 1685 | 4996 | 5141 |
| Endothelial | Endothelial | 5 | 75 | 59 | 70 | 91 | 295 | 349 | 496 | 178 | 716 | 1739 | 2034 |
| Endothelial | Endothelial | 9 | 31 | 41 | 24 | 24 | 120 | 182 | 270 | 151 | 313 | 916 | 1036 |
| Endothelial | Endothelial | 15 | 0 | 1 | 29 | 51 | 81 | 0 | 0 | 0 | 0 | 0 | 81 |
| Mesenchymal | Matrix Fibroblasts | 1 | 61 | 41 | 41 | 11 | 154 | 1295 | 842 | 965 | 1074 | 4176 | 4330 |
| Mesenchymal | Matrix Fibroblasts | 14 | 121 | 48 | 130 | 98 | 397 | 2 | 5 | 7 | 2 | 16 | 413 |
| Mesenchymal | Myo-fibroblasts | 3 | 192 | 90 | 36 | 81 | 399 | 249 | 308 | 846 | 1276 | 2679 | 3078 |
| Mesenchymal | Myo-fibroblasts | 10 | 86 | 74 | 4 | 8 | 172 | 169 | 195 | 152 | 69 | 585 | 757 |
| Mesenchymal | Myo-fibroblasts | 11 | 39 | 40 | 80 | 29 | 188 | 102 | 153 | 28 | 286 | 569 | 757 |
| Mesenchymal | Pericytes | 12 | 45 | 43 | 40 | 38 | 166 | 196 | 59 | 124 | 100 | 479 | 645 |
| Epithelial | AT1 Cells | 6 | 1 | 8 | 0 | 1 | 10 | 790 | 707 | 22 | 140 | 1659 | 1669 |
| Epithelial | AT2 Cells | 2 | 4 | 7 | 2 | 1 | 14 | 833 | 1102 | 331 | 1197 | 3463 | 3477 |
| Immune Cells | Macrophages | 4 | 60 | 36 | 6 | 4 | 106 | 413 | 1128 | 1012 | 343 | 2896 | 3002 |
| Immune Cells | B Cells | 7 | 10 | 3 | 6 | 4 | 23 | 49 | 242 | 253 | 821 | 1365 | 1388 |
| Immune Cells | CD4 T Cells | 8 | 0 | 0 | 16 | 3 | 19 | 31 | 292 | 335 | 653 | 1311 | 1330 |
| Immune Cells | Myeloid Cells | 13 | 1 | 0 | 0 | 0 | 1 | 56 | 140 | 215 | 174 | 585 | 586 |
| Immune Cells | NK Cells | 16 | 28 | 9 | 0 | 0 | 37 | 0 | 0 | 1 | 0 | 1 | 38 |

Table S5: Principal components (PCs) and associated statistics involved with age estimation of human cells

| Cell Type | Principal Component (PC) Associated with Age | % variance | Absolute Correlation with Age (r) |
| --- | --- | --- | --- |
| Endothelial Cells | 3 | 12 | 0.46 |
| Myo-fibroblasts | 8 | 1 | 0.41 |
| Matrix Fibroblasts | 2 | 22 | 0.62 |
| Macrophages | 2 | 27 | 0.41 |
| B-Cells | 8 | 2 | 0.31 |
| CD4 T-Cells | 7 | 2 | 0.12 |
| Epithelial Cells | 3 | 14 | 0.42 |

Table S6: Sample Characteristics

| Donor | GAB (Weeks) | Age at donation | Gender | scSeq | Flow Cytometry | FISH |
| --- | --- | --- | --- | --- | --- | --- |
| D038 | 38 | 1 Day | F | Y | Y |  |
| D051 | 31 | 1 Day | F | Y | Y |  |
| D019 | 39 | 1 Day | M |  | Y |  |
| D109 | 41 | 1 Day | F |  |  | Y |
| D005 | 39 | 1 Day | M |  | Y | Y |
| D102 | 37 | 1 Day | M |  |  | Y |

Table S7: Marker genes for individual clusters of cells from mouse cells

| Genes | Average Log Fold Change | Unadjusted P-value | Adjusted P-Value | Proportion of cells in Group 1 | Proportion of cells in Group 2 | Cluster No | Sub-Type |
| --- | --- | --- | --- | --- | --- | --- | --- |
| Hpgd | 1.84 | 0.00 | 0.00 | 0.75 | 0.19 | 0 | Endothelial |
| Scn7a | 1.75 | 0.00 | 0.00 | 0.88 | 0.22 | 0 | Endothelial |
| Calcrl | 1.74 | 0.00 | 0.00 | 0.80 | 0.18 | 0 | Endothelial |
| Cd93 | 1.71 | 0.00 | 0.00 | 0.76 | 0.15 | 0 | Endothelial |
| Gpihbp1 | 1.71 | 0.00 | 0.00 | 0.63 | 0.10 | 0 | Endothelial |
| Tmem100 | 1.70 | 0.00 | 0.00 | 0.87 | 0.22 | 0 | Endothelial |
| Cxcl12 | 1.57 | 0.00 | 0.00 | 0.35 | 0.09 | 0 | Endothelial |
| Cdh5 | 1.56 | 0.00 | 0.00 | 0.77 | 0.16 | 0 | Endothelial |
| Pcdh17 | 1.55 | 0.00 | 0.00 | 0.66 | 0.15 | 0 | Endothelial |
| Ptprb | 1.54 | 0.00 | 0.00 | 0.71 | 0.15 | 0 | Endothelial |
| Clec14a | 1.54 | 0.00 | 0.00 | 0.62 | 0.12 | 0 | Endothelial |
| BC028528 | 1.54 | 0.00 | 0.00 | 0.77 | 0.23 | 0 | Endothelial |
| Egfl7 | 1.47 | 0.00 | 0.00 | 0.85 | 0.20 | 0 | Endothelial |
| Sox17 | 1.45 | 0.00 | 0.00 | 0.50 | 0.09 | 0 | Endothelial |
| Gpr116 | 1.43 | 0.00 | 0.00 | 0.87 | 0.31 | 0 | Endothelial |
| Cyyr1 | 1.41 | 0.00 | 0.00 | 0.50 | 0.10 | 0 | Endothelial |
| Epas1 | 1.38 | 0.00 | 0.00 | 0.71 | 0.19 | 0 | Endothelial |
| Pecam1 | 1.37 | 0.00 | 0.00 | 0.71 | 0.18 | 0 | Endothelial |
| Rasgrp3 | 1.35 | 0.00 | 0.00 | 0.47 | 0.08 | 0 | Endothelial |
| Sema3c | 1.32 | 0.00 | 0.00 | 0.50 | 0.16 | 0 | Endothelial |
| Ldb2 | 1.31 | 0.00 | 0.00 | 0.51 | 0.12 | 0 | Endothelial |
| S100a16 | 1.31 | 0.00 | 0.00 | 0.62 | 0.19 | 0 | Endothelial |
| Tm4sf1 | 1.31 | 0.00 | 0.00 | 0.59 | 0.16 | 0 | Endothelial |
| Tmem2 | 1.31 | 0.00 | 0.00 | 0.45 | 0.11 | 0 | Endothelial |
| Ctla2a | 1.31 | 0.00 | 0.00 | 0.62 | 0.15 | 0 | Endothelial |
| Tek | 1.29 | 0.00 | 0.00 | 0.44 | 0.09 | 0 | Endothelial |
| Ramp2 | 1.28 | 0.00 | 0.00 | 0.81 | 0.21 | 0 | Endothelial |
| Plvap | 1.26 | 0.00 | 0.00 | 0.43 | 0.09 | 0 | Endothelial |
| Cav1 | 1.26 | 0.00 | 0.00 | 0.73 | 0.26 | 0 | Endothelial |
| Cldn5 | 1.26 | 0.00 | 0.00 | 0.68 | 0.16 | 0 | Endothelial |
| Tspan7 | 1.24 | 0.00 | 0.00 | 0.61 | 0.16 | 0 | Endothelial |
| Edn1 | 1.23 | 0.00 | 0.00 | 0.33 | 0.07 | 0 | Endothelial |
| Stmn2 | 1.20 | 0.00 | 0.00 | 0.46 | 0.11 | 0 | Endothelial |
| Clec1a | 1.20 | 0.00 | 0.00 | 0.53 | 0.12 | 0 | Endothelial |
| Cd36 | 1.19 | 0.00 | 0.00 | 0.72 | 0.28 | 0 | Endothelial |
| Kit | 1.18 | 0.00 | 0.00 | 0.33 | 0.05 | 0 | Endothelial |
| Efnb2 | 1.18 | 0.00 | 0.00 | 0.41 | 0.11 | 0 | Endothelial |
| Sdpr | 1.17 | 0.00 | 0.00 | 0.89 | 0.47 | 0 | Endothelial |
| Tspan18 | 1.17 | 0.00 | 0.00 | 0.43 | 0.10 | 0 | Endothelial |
| Fendrr | 1.17 | 0.00 | 0.00 | 0.52 | 0.16 | 0 | Endothelial |
| Lyve1 | 1.14 | 0.00 | 0.00 | 0.62 | 0.17 | 0 | Endothelial |
| Rasip1 | 1.13 | 0.00 | 0.00 | 0.39 | 0.09 | 0 | Endothelial |
| Ecscr | 1.13 | 0.00 | 0.00 | 0.52 | 0.12 | 0 | Endothelial |
| Ace | 1.12 | 0.00 | 0.00 | 0.41 | 0.10 | 0 | Endothelial |
| Rnf144a | 1.12 | 0.00 | 0.00 | 0.42 | 0.12 | 0 | Endothelial |
| Afap1l1 | 1.12 | 0.00 | 0.00 | 0.46 | 0.14 | 0 | Endothelial |
| Ehd4 | 1.12 | 0.00 | 0.00 | 0.63 | 0.23 | 0 | Endothelial |
| Tspan13 | 1.11 | 0.00 | 0.00 | 0.65 | 0.20 | 0 | Endothelial |
| Gja4 | 1.10 | 0.00 | 0.00 | 0.28 | 0.04 | 0 | Endothelial |
| Palmd | 1.09 | 0.00 | 0.00 | 0.38 | 0.10 | 0 | Endothelial |
| Nostrin | 1.08 | 0.00 | 0.00 | 0.43 | 0.11 | 0 | Endothelial |
| Esam | 1.08 | 0.00 | 0.00 | 0.49 | 0.14 | 0 | Endothelial |
| Sgk1 | 1.08 | 0.00 | 0.00 | 0.45 | 0.16 | 0 | Endothelial |
| Myzap | 1.07 | 0.00 | 0.00 | 0.43 | 0.11 | 0 | Endothelial |
| Ushbp1 | 1.05 | 0.00 | 0.00 | 0.32 | 0.06 | 0 | Endothelial |
| Mest | 1.05 | 0.00 | 0.00 | 0.49 | 0.16 | 0 | Endothelial |
| Sema3g | 1.04 | 0.00 | 0.00 | 0.28 | 0.05 | 0 | Endothelial |
| Itga6 | 1.04 | 0.00 | 0.00 | 0.37 | 0.11 | 0 | Endothelial |
| Myct1 | 1.04 | 0.00 | 0.00 | 0.41 | 0.10 | 0 | Endothelial |
| Thbd | 1.04 | 0.00 | 0.00 | 0.56 | 0.21 | 0 | Endothelial |
| Tubb2a | 1.04 | 0.00 | 0.00 | 0.46 | 0.18 | 0 | Endothelial |
| Acer2 | 1.02 | 0.00 | 0.00 | 0.31 | 0.07 | 0 | Endothelial |
| Ripply3 | 1.02 | 0.00 | 0.00 | 0.38 | 0.09 | 0 | Endothelial |
| Fmo1 | 1.01 | 0.00 | 0.00 | 0.34 | 0.09 | 0 | Endothelial |
| Ppap2a | 1.01 | 0.00 | 0.00 | 0.46 | 0.16 | 0 | Endothelial |
| Flt1 | 1.01 | 0.00 | 0.00 | 0.42 | 0.12 | 0 | Endothelial |
| Slc9a3r2 | 1.01 | 0.00 | 0.00 | 0.49 | 0.17 | 0 | Endothelial |
| Acvrl1 | 1.00 | 0.00 | 0.00 | 0.47 | 0.14 | 0 | Endothelial |
| Tnfsf10 | 1.00 | 0.00 | 0.00 | 0.30 | 0.06 | 0 | Endothelial |
| Erg | 1.00 | 0.00 | 0.00 | 0.37 | 0.10 | 0 | Endothelial |
| Ccnd1 | 0.98 | 0.00 | 0.00 | 0.42 | 0.16 | 0 | Endothelial |
| Ets1 | 0.97 | 0.00 | 0.00 | 0.52 | 0.21 | 0 | Endothelial |
| Tie1 | 0.97 | 0.00 | 0.00 | 0.35 | 0.08 | 0 | Endothelial |
| Aqp1 | 0.97 | 0.00 | 0.00 | 0.48 | 0.16 | 0 | Endothelial |
| Scn3b | 0.96 | 0.00 | 0.00 | 0.26 | 0.05 | 0 | Endothelial |
| Podxl | 0.96 | 0.00 | 0.00 | 0.31 | 0.08 | 0 | Endothelial |
| Aplnr | 0.94 | 0.00 | 0.00 | 0.25 | 0.05 | 0 | Endothelial |
| Emcn | 0.93 | 0.00 | 0.00 | 0.33 | 0.08 | 0 | Endothelial |
| Tjp1 | 0.93 | 0.00 | 0.00 | 0.37 | 0.13 | 0 | Endothelial |
| Ece1 | 0.92 | 0.00 | 0.00 | 0.53 | 0.22 | 0 | Endothelial |
| Icam2 | 0.92 | 0.00 | 0.00 | 0.44 | 0.12 | 0 | Endothelial |
| Pde4b | 0.92 | 0.00 | 0.00 | 0.34 | 0.12 | 0 | Endothelial |
| Slc43a3 | 0.92 | 0.00 | 0.00 | 0.48 | 0.18 | 0 | Endothelial |
| Fkbp1a | 0.91 | 0.00 | 0.00 | 0.68 | 0.37 | 0 | Endothelial |
| Tubb2b | 0.89 | 0.00 | 0.00 | 0.32 | 0.11 | 0 | Endothelial |
| Gata2 | 0.89 | 0.00 | 0.00 | 0.37 | 0.11 | 0 | Endothelial |
| Cd34 | 0.89 | 0.00 | 0.00 | 0.45 | 0.17 | 0 | Endothelial |
| Id3 | 0.89 | 0.00 | 0.00 | 0.75 | 0.42 | 0 | Endothelial |
| Klhl5 | 0.88 | 0.00 | 0.00 | 0.34 | 0.12 | 0 | Endothelial |
| Tbx3 | 0.88 | 0.00 | 0.00 | 0.47 | 0.17 | 0 | Endothelial |
| Ly6e | 0.87 | 0.00 | 0.00 | 0.75 | 0.38 | 0 | Endothelial |
| Slco2a1 | 0.87 | 0.00 | 0.00 | 0.30 | 0.09 | 0 | Endothelial |
| Ddah2 | 0.87 | 0.00 | 0.00 | 0.50 | 0.23 | 0 | Endothelial |
| Ptprm | 0.86 | 0.00 | 0.00 | 0.26 | 0.08 | 0 | Endothelial |
| Foxp1 | 0.86 | 0.00 | 0.00 | 0.69 | 0.42 | 0 | Endothelial |
| Col4a2 | 0.86 | 0.00 | 0.00 | 0.69 | 0.40 | 0 | Endothelial |
| Casz1 | 0.86 | 0.00 | 0.00 | 0.32 | 0.11 | 0 | Endothelial |
| Col4a1 | 0.86 | 0.00 | 0.00 | 0.75 | 0.46 | 0 | Endothelial |
| Srgn | 0.85 | 0.00 | 0.00 | 0.65 | 0.28 | 0 | Endothelial |
| Piezo2 | 0.85 | 0.00 | 0.00 | 0.29 | 0.09 | 0 | Endothelial |
| 4931406P16Rik | 0.85 | 0.00 | 0.00 | 0.38 | 0.16 | 0 | Endothelial |
| Cav2 | 0.84 | 0.00 | 0.00 | 0.41 | 0.16 | 0 | Endothelial |
| Gng11 | 0.84 | 0.00 | 0.00 | 0.57 | 0.25 | 0 | Endothelial |
| Ly6a | 0.84 | 0.00 | 0.00 | 0.46 | 0.16 | 0 | Endothelial |
| 9430020K01Rik | 0.84 | 0.00 | 0.00 | 0.31 | 0.11 | 0 | Endothelial |
| Cxx1a | 0.84 | 0.00 | 0.00 | 0.45 | 0.19 | 0 | Endothelial |
| Ppp1r16b | 0.82 | 0.00 | 0.00 | 0.26 | 0.07 | 0 | Endothelial |
| Ahr | 0.81 | 0.00 | 0.00 | 0.26 | 0.07 | 0 | Endothelial |
| Kdr | 0.80 | 0.00 | 0.00 | 0.45 | 0.13 | 0 | Endothelial |
| Gimap4 | 0.80 | 0.00 | 0.00 | 0.42 | 0.14 | 0 | Endothelial |
| Gimap1 | 0.79 | 0.00 | 0.00 | 0.34 | 0.11 | 0 | Endothelial |
| Anp32a | 0.78 | 0.00 | 0.00 | 0.78 | 0.58 | 0 | Endothelial |
| Rhoj | 0.77 | 0.00 | 0.00 | 0.39 | 0.16 | 0 | Endothelial |
| Sptbn1 | 0.77 | 0.00 | 0.00 | 0.76 | 0.52 | 0 | Endothelial |
| Mcam | 0.75 | 0.00 | 0.00 | 0.32 | 0.11 | 0 | Endothelial |
| Crip2 | 0.72 | 0.00 | 0.00 | 0.65 | 0.38 | 0 | Endothelial |
| Itm2b | 0.64 | 0.00 | 0.00 | 0.87 | 0.74 | 0 | Endothelial |
| Clic5 | 0.57 | 0.00 | 0.00 | 0.48 | 0.20 | 0 | Endothelial |
| Kitl | 0.56 | 0.00 | 0.00 | 0.45 | 0.20 | 0 | Endothelial |
| Vim | 0.55 | 0.00 | 0.00 | 0.87 | 0.63 | 0 | Endothelial |
| Emp1 | 0.78 | 0.00 | 0.00 | 0.43 | 0.19 | 0 | Endothelial |
| Tmem88 | 0.80 | 0.00 | 0.00 | 0.28 | 0.09 | 0 | Endothelial |
| Elk3 | 0.82 | 0.00 | 0.00 | 0.32 | 0.11 | 0 | Endothelial |
| Cxx1b | 0.80 | 0.00 | 0.00 | 0.43 | 0.20 | 0 | Endothelial |
| 2810025M15Rik | 0.79 | 0.00 | 0.00 | 0.30 | 0.10 | 0 | Endothelial |
| S1pr1 | 0.76 | 0.00 | 0.00 | 0.39 | 0.16 | 0 | Endothelial |
| Prcp | 0.79 | 0.00 | 0.00 | 0.25 | 0.08 | 0 | Endothelial |
| Cxx1c | 0.80 | 0.00 | 0.00 | 0.43 | 0.19 | 0 | Endothelial |
| Sparcl1 | 0.78 | 0.00 | 0.00 | 0.61 | 0.36 | 0 | Endothelial |
| Tcf4 | 0.74 | 0.00 | 0.00 | 0.62 | 0.38 | 0 | Endothelial |
| Nrp1 | 0.70 | 0.00 | 0.00 | 0.61 | 0.37 | 0 | Endothelial |
| Heg1 | 0.83 | 0.00 | 0.00 | 0.35 | 0.14 | 0 | Endothelial |
| Lxn | 0.81 | 0.00 | 0.00 | 0.29 | 0.10 | 0 | Endothelial |
| Prx | 0.66 | 0.00 | 0.00 | 0.27 | 0.08 | 0 | Endothelial |
| Cd200 | 0.71 | 0.00 | 0.00 | 0.49 | 0.25 | 0 | Endothelial |
| Ptrf | 0.74 | 0.00 | 0.00 | 0.55 | 0.33 | 0 | Endothelial |
| Myl12b | 0.61 | 0.00 | 0.00 | 0.73 | 0.59 | 0 | Endothelial |
| Ccdc85b | 0.77 | 0.00 | 0.00 | 0.28 | 0.10 | 0 | Endothelial |
| Guk1 | 0.85 | 0.00 | 0.00 | 0.32 | 0.13 | 0 | Endothelial |
| Prex2 | 0.81 | 0.00 | 0.00 | 0.37 | 0.16 | 0 | Endothelial |
| Pitpnc1 | 0.77 | 0.00 | 0.00 | 0.28 | 0.10 | 0 | Endothelial |
| Rdx | 0.71 | 0.00 | 0.00 | 0.63 | 0.43 | 0 | Endothelial |
| B2m | 0.44 | 0.00 | 0.00 | 0.83 | 0.65 | 0 | Endothelial |
| Hoxa5 | 0.79 | 0.00 | 0.00 | 0.29 | 0.11 | 0 | Endothelial |
| Gimap5 | 0.70 | 0.00 | 0.00 | 0.27 | 0.10 | 0 | Endothelial |
| Eng | 0.70 | 0.00 | 0.00 | 0.27 | 0.10 | 0 | Endothelial |
| Cd97 | 0.73 | 0.00 | 0.00 | 0.36 | 0.16 | 0 | Endothelial |
| Arhgap29 | 0.76 | 0.00 | 0.00 | 0.40 | 0.19 | 0 | Endothelial |
| Ybx1 | 0.51 | 0.00 | 0.00 | 0.77 | 0.62 | 0 | Endothelial |
| Slc6a6 | 0.91 | 0.00 | 0.00 | 0.43 | 0.23 | 0 | Endothelial |
| Sema6a | 0.70 | 0.00 | 0.00 | 0.27 | 0.10 | 0 | Endothelial |
| Id1 | 0.72 | 0.00 | 0.00 | 0.42 | 0.22 | 0 | Endothelial |
| Grap | 0.68 | 0.00 | 0.00 | 0.29 | 0.11 | 0 | Endothelial |
| Rapgef5 | 0.72 | 0.00 | 0.00 | 0.25 | 0.09 | 0 | Endothelial |
| Kank3 | 0.73 | 0.00 | 0.00 | 0.30 | 0.12 | 0 | Endothelial |
| Klf4 | 0.66 | 0.00 | 0.00 | 0.42 | 0.21 | 0 | Endothelial |
| Jup | 0.73 | 0.00 | 0.00 | 0.32 | 0.14 | 0 | Endothelial |
| Bst2 | 0.70 | 0.00 | 0.00 | 0.29 | 0.12 | 0 | Endothelial |
| Gnai2 | 0.66 | 0.00 | 0.00 | 0.56 | 0.36 | 0 | Endothelial |
| Cnn3 | 0.64 | 0.00 | 0.00 | 0.49 | 0.28 | 0 | Endothelial |
| Hip1 | 0.73 | 0.00 | 0.00 | 0.32 | 0.14 | 0 | Endothelial |
| Itgb1 | 0.49 | 0.00 | 0.00 | 0.77 | 0.61 | 0 | Endothelial |
| Rgs12 | 0.69 | 0.00 | 0.00 | 0.27 | 0.10 | 0 | Endothelial |
| Gimap6 | 0.62 | 0.00 | 0.00 | 0.33 | 0.14 | 0 | Endothelial |
| Ptprg | 0.74 | 0.00 | 0.00 | 0.27 | 0.11 | 0 | Endothelial |
| Sox4 | 0.67 | 0.00 | 0.00 | 0.55 | 0.34 | 0 | Endothelial |
| Ly6c1 | 0.62 | 0.00 | 0.00 | 0.36 | 0.17 | 0 | Endothelial |
| Notch1 | 0.74 | 0.00 | 0.00 | 0.28 | 0.12 | 0 | Endothelial |
| Slk | 0.67 | 0.00 | 0.00 | 0.59 | 0.41 | 0 | Endothelial |
| H2-D1 | 0.49 | 0.00 | 0.00 | 0.68 | 0.47 | 0 | Endothelial |
| Pltp | 0.71 | 0.00 | 0.00 | 0.34 | 0.16 | 0 | Endothelial |
| Snrk | 0.72 | 0.00 | 0.00 | 0.31 | 0.14 | 0 | Endothelial |
| Dlc1 | 0.71 | 0.00 | 0.00 | 0.33 | 0.15 | 0 | Endothelial |
| Nes | 0.53 | 0.00 | 0.00 | 0.40 | 0.19 | 0 | Endothelial |
| Cyth3 | 0.68 | 0.00 | 0.00 | 0.41 | 0.22 | 0 | Endothelial |
| Foxf1 | 0.66 | 0.00 | 0.00 | 0.26 | 0.10 | 0 | Endothelial |
| Snx3 | 0.67 | 0.00 | 0.00 | 0.49 | 0.31 | 0 | Endothelial |
| Rgcc | 0.53 | 0.00 | 0.00 | 0.34 | 0.16 | 0 | Endothelial |
| Jun | 0.42 | 0.00 | 0.00 | 0.56 | 0.34 | 0 | Endothelial |
| Dusp3 | 0.67 | 0.00 | 0.00 | 0.35 | 0.17 | 0 | Endothelial |
| Arhgef12 | 0.74 | 0.00 | 0.00 | 0.39 | 0.22 | 0 | Endothelial |
| Tmsb10 | 0.31 | 0.00 | 0.00 | 0.94 | 0.85 | 0 | Endothelial |
| Ablim1 | 0.72 | 0.00 | 0.00 | 0.41 | 0.23 | 0 | Endothelial |
| Lgals9 | 0.64 | 0.00 | 0.00 | 0.28 | 0.13 | 0 | Endothelial |
| Cd151 | 0.66 | 0.00 | 0.00 | 0.31 | 0.14 | 0 | Endothelial |
| Smarca2 | 0.71 | 0.00 | 0.00 | 0.49 | 0.33 | 0 | Endothelial |
| Pea15a | 0.65 | 0.00 | 0.00 | 0.28 | 0.13 | 0 | Endothelial |
| Marcks | 0.40 | 0.00 | 0.00 | 0.68 | 0.48 | 0 | Endothelial |
| Ctnna1 | 0.64 | 0.00 | 0.00 | 0.51 | 0.34 | 0 | Endothelial |
| Stmn1 | 0.55 | 0.00 | 0.00 | 0.39 | 0.22 | 0 | Endothelial |
| Cd9 | 0.43 | 0.00 | 0.00 | 0.52 | 0.33 | 0 | Endothelial |
| Uaca | 0.67 | 0.00 | 0.00 | 0.27 | 0.12 | 0 | Endothelial |
| Ifitm3 | 0.50 | 0.00 | 0.00 | 0.55 | 0.38 | 0 | Endothelial |
| Klf7 | 0.61 | 0.00 | 0.00 | 0.32 | 0.17 | 0 | Endothelial |
| Marcksl1 | 0.63 | 0.00 | 0.00 | 0.33 | 0.18 | 0 | Endothelial |
| Fnbp1l | 0.63 | 0.00 | 0.00 | 0.39 | 0.24 | 0 | Endothelial |
| Fermt2 | 0.47 | 0.00 | 0.00 | 0.54 | 0.38 | 0 | Endothelial |
| Rhoc | 0.59 | 0.00 | 0.00 | 0.34 | 0.19 | 0 | Endothelial |
| Lphn2 | 0.57 | 0.00 | 0.00 | 0.26 | 0.12 | 0 | Endothelial |
| Mgll | 0.62 | 0.00 | 0.00 | 0.29 | 0.14 | 0 | Endothelial |
| Dnajc8 | 0.53 | 0.00 | 0.00 | 0.54 | 0.41 | 0 | Endothelial |
| Hes1 | 0.65 | 0.00 | 0.00 | 0.43 | 0.28 | 0 | Endothelial |
| Anxa3 | 0.50 | 0.00 | 0.00 | 0.36 | 0.21 | 0 | Endothelial |
| Plk2 | 0.64 | 0.00 | 0.00 | 0.28 | 0.15 | 0 | Endothelial |
| Aplp2 | 0.54 | 0.00 | 0.00 | 0.51 | 0.37 | 0 | Endothelial |
| Gcc2 | 0.65 | 0.00 | 0.00 | 0.38 | 0.23 | 0 | Endothelial |
| Rab11a | 0.57 | 0.00 | 0.00 | 0.38 | 0.25 | 0 | Endothelial |
| Luzp1 | 0.61 | 0.00 | 0.00 | 0.32 | 0.18 | 0 | Endothelial |
| Wasf2 | 0.59 | 0.00 | 0.00 | 0.42 | 0.29 | 0 | Endothelial |
| Mllt4 | 0.61 | 0.00 | 0.00 | 0.33 | 0.20 | 0 | Endothelial |
| Dusp6 | 0.57 | 0.00 | 0.00 | 0.30 | 0.17 | 0 | Endothelial |
| Tpm4 | 0.43 | 0.00 | 0.00 | 0.60 | 0.48 | 0 | Endothelial |
| Tubb5 | 0.42 | 0.00 | 0.00 | 0.56 | 0.43 | 0 | Endothelial |
| Klf2 | 0.53 | 0.00 | 0.00 | 0.32 | 0.18 | 0 | Endothelial |
| Actn4 | 0.57 | 0.00 | 0.00 | 0.40 | 0.28 | 0 | Endothelial |
| Hsp90ab1 | 0.25 | 0.00 | 0.00 | 0.91 | 0.89 | 0 | Endothelial |
| Xiap | 0.60 | 0.00 | 0.00 | 0.31 | 0.19 | 0 | Endothelial |
| Cyb5r3 | 0.53 | 0.00 | 0.00 | 0.49 | 0.38 | 0 | Endothelial |
| Rap1a | 0.51 | 0.00 | 0.00 | 0.42 | 0.29 | 0 | Endothelial |
| Efnb1 | 0.55 | 0.00 | 0.00 | 0.25 | 0.14 | 0 | Endothelial |
| Ppp1r2 | 0.46 | 0.00 | 0.00 | 0.38 | 0.25 | 0 | Endothelial |
| Oaz2 | 0.58 | 0.00 | 0.00 | 0.39 | 0.27 | 0 | Endothelial |
| Ctnnd1 | 0.54 | 0.00 | 0.00 | 0.27 | 0.15 | 0 | Endothelial |
| Atp1b3 | 0.55 | 0.00 | 0.00 | 0.36 | 0.24 | 0 | Endothelial |
| Sptan1 | 0.56 | 0.00 | 0.00 | 0.27 | 0.16 | 0 | Endothelial |
| Clic1 | 0.46 | 0.00 | 0.00 | 0.48 | 0.37 | 0 | Endothelial |
| Zfp503 | 0.55 | 0.00 | 0.00 | 0.25 | 0.14 | 0 | Endothelial |
| Peg3 | 0.39 | 0.00 | 0.00 | 0.43 | 0.29 | 0 | Endothelial |
| Bnip2 | 0.53 | 0.00 | 0.00 | 0.41 | 0.29 | 0 | Endothelial |
| Msn | 0.40 | 0.00 | 0.00 | 0.53 | 0.42 | 0 | Endothelial |
| Map7d1 | 0.55 | 0.00 | 0.00 | 0.28 | 0.17 | 0 | Endothelial |
| Tuba1a | 0.32 | 0.00 | 0.00 | 0.62 | 0.49 | 0 | Endothelial |
| Ubb | 0.25 | 0.00 | 0.00 | 0.83 | 0.80 | 0 | Endothelial |
| Gm26735 | 0.54 | 0.00 | 0.00 | 0.30 | 0.18 | 0 | Endothelial |
| Sypl | 0.52 | 0.00 | 0.00 | 0.36 | 0.25 | 0 | Endothelial |
| F11r | 0.49 | 0.00 | 0.00 | 0.26 | 0.15 | 0 | Endothelial |
| Leprot | 0.48 | 0.00 | 0.00 | 0.43 | 0.32 | 0 | Endothelial |
| Plcg1 | 0.54 | 0.00 | 0.00 | 0.26 | 0.15 | 0 | Endothelial |
| Myl12a | 0.30 | 0.00 | 0.00 | 0.70 | 0.65 | 0 | Endothelial |
| Dynll1 | 0.40 | 0.00 | 0.00 | 0.54 | 0.44 | 0 | Endothelial |
| Tmem204 | 0.44 | 0.00 | 0.00 | 0.25 | 0.15 | 0 | Endothelial |
| Mfap2 | 0.31 | 0.00 | 0.00 | 0.48 | 0.33 | 0 | Endothelial |
| Add3 | 0.48 | 0.00 | 0.00 | 0.38 | 0.27 | 0 | Endothelial |
| Serpinh1 | 0.32 | 0.00 | 0.00 | 0.51 | 0.38 | 0 | Endothelial |
| Hnrnpa2b1 | 0.29 | 0.00 | 0.00 | 0.75 | 0.72 | 0 | Endothelial |
| Ywhab | 0.35 | 0.00 | 0.00 | 0.62 | 0.56 | 0 | Endothelial |
| Itpripl2 | 0.50 | 0.00 | 0.00 | 0.26 | 0.16 | 0 | Endothelial |
| Acap2 | 0.49 | 0.00 | 0.00 | 0.28 | 0.19 | 0 | Endothelial |
| Cfl1 | 0.34 | 0.00 | 0.00 | 0.63 | 0.58 | 0 | Endothelial |
| Sh3glb1 | 0.41 | 0.00 | 0.00 | 0.48 | 0.39 | 0 | Endothelial |
| H2-K1 | 0.37 | 0.00 | 0.00 | 0.42 | 0.31 | 0 | Endothelial |
| Ctnnb1 | 0.44 | 0.00 | 0.00 | 0.42 | 0.34 | 0 | Endothelial |
| Tax1bp1 | 0.39 | 0.00 | 0.00 | 0.67 | 0.63 | 0 | Endothelial |
| Hspa8 | 0.25 | 0.00 | 0.00 | 0.70 | 0.67 | 0 | Endothelial |
| Arf1 | 0.39 | 0.00 | 0.00 | 0.49 | 0.42 | 0 | Endothelial |
| Dnaja1 | 0.33 | 0.00 | 0.00 | 0.56 | 0.51 | 0 | Endothelial |
| Csnk1a1 | 0.32 | 0.00 | 0.00 | 0.58 | 0.53 | 0 | Endothelial |
| Eif4g2 | 0.40 | 0.00 | 0.00 | 0.47 | 0.40 | 0 | Endothelial |
| Rhoa | 0.35 | 0.00 | 0.00 | 0.51 | 0.46 | 0 | Endothelial |
| Fubp1 | 0.45 | 0.00 | 0.00 | 0.32 | 0.24 | 0 | Endothelial |
| Rnf7 | 0.42 | 0.00 | 0.00 | 0.31 | 0.23 | 0 | Endothelial |
| Akr1a1 | 0.32 | 0.00 | 0.00 | 0.53 | 0.49 | 0 | Endothelial |
| Hbb-bs | 0.41 | 0.00 | 0.00 | 0.81 | 0.81 | 0 | Endothelial |
| Adamts1 | 0.33 | 0.00 | 0.00 | 0.29 | 0.20 | 0 | Endothelial |
| Hook3 | 0.43 | 0.00 | 0.00 | 0.29 | 0.22 | 0 | Endothelial |
| Hbb-bt | 0.43 | 0.00 | 0.00 | 0.72 | 0.71 | 0 | Endothelial |
| Snrpb | 0.37 | 0.00 | 0.00 | 0.41 | 0.35 | 0 | Endothelial |
| Phldb2 | 0.38 | 0.00 | 0.00 | 0.36 | 0.28 | 0 | Endothelial |
| Glul | 0.35 | 0.00 | 0.00 | 0.29 | 0.22 | 0 | Endothelial |
| Ywhae | 0.28 | 0.00 | 0.00 | 0.57 | 0.54 | 0 | Endothelial |
| Ralb | 0.37 | 0.00 | 0.00 | 0.26 | 0.19 | 0 | Endothelial |
| Dync1i2 | 0.33 | 0.00 | 0.00 | 0.51 | 0.48 | 0 | Endothelial |
| Hmg20b | 0.39 | 0.00 | 0.00 | 0.39 | 0.33 | 0 | Endothelial |
| Tmco1 | 0.39 | 0.00 | 0.00 | 0.35 | 0.29 | 0 | Endothelial |
| 11-Sep | 0.36 | 0.00 | 0.00 | 0.29 | 0.22 | 0 | Endothelial |
| Cdc42 | 0.27 | 0.00 | 0.00 | 0.57 | 0.54 | 0 | Endothelial |
| Clic4 | 0.35 | 0.00 | 0.00 | 0.35 | 0.29 | 0 | Endothelial |
| Zmiz1 | 0.40 | 0.00 | 0.00 | 0.28 | 0.21 | 0 | Endothelial |
| Top1 | 0.34 | 0.00 | 0.00 | 0.49 | 0.45 | 0 | Endothelial |
| 1700020I14Rik | 0.39 | 0.00 | 0.00 | 0.28 | 0.22 | 0 | Endothelial |
| Arpc3 | 0.37 | 0.00 | 0.00 | 0.32 | 0.26 | 0 | Endothelial |
| Zbtb20 | 0.36 | 0.00 | 0.00 | 0.33 | 0.26 | 0 | Endothelial |
| H1f0 | 0.38 | 0.00 | 0.00 | 0.32 | 0.26 | 0 | Endothelial |
| Ndufa8 | 0.38 | 0.00 | 0.00 | 0.30 | 0.25 | 0 | Endothelial |
| Ifngr1 | 0.32 | 0.00 | 0.00 | 0.30 | 0.23 | 0 | Endothelial |
| Rab6a | 0.39 | 0.00 | 0.00 | 0.26 | 0.21 | 0 | Endothelial |
| Nap1l4 | 0.38 | 0.00 | 0.00 | 0.31 | 0.26 | 0 | Endothelial |
| Cyb5 | 0.36 | 0.00 | 0.00 | 0.31 | 0.26 | 0 | Endothelial |
| Cd2ap | 0.35 | 0.00 | 0.00 | 0.39 | 0.34 | 0 | Endothelial |
| Mef2a | 0.30 | 0.00 | 0.00 | 0.41 | 0.36 | 0 | Endothelial |
| Hba-a2 | 0.41 | 0.00 | 0.00 | 0.70 | 0.70 | 0 | Endothelial |
| Gnb1 | 0.36 | 0.00 | 0.00 | 0.27 | 0.21 | 0 | Endothelial |
| Rab1 | 0.34 | 0.00 | 0.00 | 0.35 | 0.30 | 0 | Endothelial |
| Rac1 | 0.35 | 0.00 | 0.00 | 0.34 | 0.30 | 0 | Endothelial |
| Cbx1 | 0.30 | 0.00 | 0.00 | 0.45 | 0.41 | 0 | Endothelial |
| Hba-a1 | 0.41 | 0.00 | 0.00 | 0.70 | 0.71 | 0 | Endothelial |
| Qk | 0.32 | 0.00 | 0.00 | 0.34 | 0.29 | 0 | Endothelial |
| Gnaq | 0.35 | 0.00 | 0.00 | 0.26 | 0.21 | 0 | Endothelial |
| Prdx2 | 0.30 | 0.00 | 0.00 | 0.37 | 0.33 | 0 | Endothelial |
| G3bp2 | 0.33 | 0.00 | 0.00 | 0.35 | 0.31 | 0 | Endothelial |
| S100a13 | 0.32 | 0.00 | 0.00 | 0.30 | 0.25 | 0 | Endothelial |
| Arglu1 | 0.29 | 0.00 | 0.00 | 0.45 | 0.42 | 0 | Endothelial |
| Ccdc50 | 0.35 | 0.00 | 0.00 | 0.30 | 0.26 | 0 | Endothelial |
| Ivns1abp | 0.42 | 0.00 | 0.00 | 0.27 | 0.22 | 0 | Endothelial |
| Hnrnpf | 0.29 | 0.00 | 0.00 | 0.38 | 0.34 | 0 | Endothelial |
| Itsn2 | 0.31 | 0.00 | 0.00 | 0.29 | 0.24 | 0 | Endothelial |
| Ktn1 | 0.30 | 0.00 | 0.00 | 0.41 | 0.38 | 0 | Endothelial |
| Mxd4 | 0.27 | 0.00 | 0.00 | 0.41 | 0.37 | 0 | Endothelial |
| Ttc3 | 0.29 | 0.00 | 0.00 | 0.32 | 0.29 | 0 | Endothelial |
| 2-Sep | 0.26 | 0.00 | 0.00 | 0.38 | 0.35 | 0 | Endothelial |
| Ywhaq | 0.26 | 0.00 | 0.00 | 0.39 | 0.36 | 0 | Endothelial |
| Prkar1a | 0.29 | 0.00 | 0.00 | 0.35 | 0.32 | 0 | Endothelial |
| Apc | 0.30 | 0.00 | 0.00 | 0.27 | 0.23 | 0 | Endothelial |
| Ppp1ca | 0.27 | 0.00 | 0.00 | 0.35 | 0.33 | 0 | Endothelial |
| Serinc1 | 0.26 | 0.00 | 0.00 | 0.32 | 0.29 | 0 | Endothelial |
| B230219D22Rik | 0.29 | 0.00 | 0.00 | 0.30 | 0.27 | 0 | Endothelial |
| Csnk2a1 | 0.26 | 0.00 | 0.00 | 0.33 | 0.31 | 0 | Endothelial |
| Zcrb1 | 0.29 | 0.00 | 0.00 | 0.31 | 0.29 | 0 | Endothelial |
| Zc3h7a | 0.27 | 0.00 | 0.00 | 0.31 | 0.29 | 0 | Endothelial |
| Vps36 | 0.27 | 0.00 | 0.00 | 0.31 | 0.29 | 0 | Endothelial |
| Chmp5 | 0.27 | 0.00 | 0.00 | 0.26 | 0.24 | 0 | Endothelial |
| Rab10 | 0.26 | 0.00 | 0.02 | 0.26 | 0.24 | 0 | Endothelial |
| Dpt | 2.69 | 0.00 | 0.00 | 0.93 | 0.11 | 1 | Fibroblast |
| Mfap4 | 2.62 | 0.00 | 0.00 | 0.98 | 0.18 | 1 | Fibroblast |
| Adh1 | 2.41 | 0.00 | 0.00 | 0.93 | 0.10 | 1 | Fibroblast |
| Cdh11 | 2.06 | 0.00 | 0.00 | 0.89 | 0.13 | 1 | Fibroblast |
| Npnt | 1.99 | 0.00 | 0.00 | 0.89 | 0.18 | 1 | Fibroblast |
| Macf1 | 1.94 | 0.00 | 0.00 | 0.98 | 0.48 | 1 | Fibroblast |
| Fhl1 | 1.92 | 0.00 | 0.00 | 0.94 | 0.23 | 1 | Fibroblast |
| Limch1 | 1.88 | 0.00 | 0.00 | 0.97 | 0.25 | 1 | Fibroblast |
| Gyg | 1.85 | 0.00 | 0.00 | 0.84 | 0.13 | 1 | Fibroblast |
| Rbp1 | 1.83 | 0.00 | 0.00 | 0.84 | 0.14 | 1 | Fibroblast |
| Fn1 | 1.82 | 0.00 | 0.00 | 0.87 | 0.18 | 1 | Fibroblast |
| Enpep | 1.78 | 0.00 | 0.00 | 0.66 | 0.05 | 1 | Fibroblast |
| Cp | 1.73 | 0.00 | 0.00 | 0.67 | 0.08 | 1 | Fibroblast |
| Lbh | 1.66 | 0.00 | 0.00 | 0.79 | 0.14 | 1 | Fibroblast |
| Gpc3 | 1.64 | 0.00 | 0.00 | 0.75 | 0.11 | 1 | Fibroblast |
| Cpm | 1.64 | 0.00 | 0.00 | 0.72 | 0.13 | 1 | Fibroblast |
| Plxdc2 | 1.63 | 0.00 | 0.00 | 0.66 | 0.08 | 1 | Fibroblast |
| Meox2 | 1.60 | 0.00 | 0.00 | 0.59 | 0.03 | 1 | Fibroblast |
| Mdk | 1.60 | 0.00 | 0.00 | 0.77 | 0.13 | 1 | Fibroblast |
| Tcf21 | 1.59 | 0.00 | 0.00 | 0.86 | 0.11 | 1 | Fibroblast |
| Colec12 | 1.59 | 0.00 | 0.00 | 0.60 | 0.06 | 1 | Fibroblast |
| Pcolce2 | 1.58 | 0.00 | 0.00 | 0.59 | 0.05 | 1 | Fibroblast |
| Slit2 | 1.56 | 0.00 | 0.00 | 0.53 | 0.03 | 1 | Fibroblast |
| Maf | 1.54 | 0.00 | 0.00 | 0.82 | 0.14 | 1 | Fibroblast |
| Ptprd | 1.53 | 0.00 | 0.00 | 0.61 | 0.07 | 1 | Fibroblast |
| Plac9b | 1.52 | 0.00 | 0.00 | 0.90 | 0.25 | 1 | Fibroblast |
| Plac9a | 1.50 | 0.00 | 0.00 | 0.90 | 0.25 | 1 | Fibroblast |
| Nexn | 1.50 | 0.00 | 0.00 | 0.79 | 0.17 | 1 | Fibroblast |
| Ogn | 1.49 | 0.00 | 0.00 | 0.65 | 0.09 | 1 | Fibroblast |
| Sh3bgrl | 1.46 | 0.00 | 0.00 | 0.90 | 0.40 | 1 | Fibroblast |
| Col1a2 | 1.44 | 0.00 | 0.00 | 1.00 | 0.34 | 1 | Fibroblast |
| Figf | 1.42 | 0.00 | 0.00 | 0.49 | 0.03 | 1 | Fibroblast |
| Serpine2 | 1.42 | 0.00 | 0.00 | 0.85 | 0.20 | 1 | Fibroblast |
| Mmp2 | 1.42 | 0.00 | 0.00 | 0.63 | 0.09 | 1 | Fibroblast |
| Akap12 | 1.40 | 0.00 | 0.00 | 0.76 | 0.19 | 1 | Fibroblast |
| Vcan | 1.36 | 0.00 | 0.00 | 0.54 | 0.07 | 1 | Fibroblast |
| Itga8 | 1.35 | 0.00 | 0.00 | 0.52 | 0.05 | 1 | Fibroblast |
| Slc27a6 | 1.34 | 0.00 | 0.00 | 0.41 | 0.01 | 1 | Fibroblast |
| Cped1 | 1.31 | 0.00 | 0.00 | 0.51 | 0.06 | 1 | Fibroblast |
| Palld | 1.29 | 0.00 | 0.00 | 0.66 | 0.13 | 1 | Fibroblast |
| Myh10 | 1.28 | 0.00 | 0.00 | 0.72 | 0.18 | 1 | Fibroblast |
| Col13a1 | 1.28 | 0.00 | 0.00 | 0.52 | 0.06 | 1 | Fibroblast |
| G0s2 | 1.26 | 0.00 | 0.00 | 0.60 | 0.09 | 1 | Fibroblast |
| Nrep | 1.23 | 0.00 | 0.00 | 0.91 | 0.39 | 1 | Fibroblast |
| Slc38a5 | 1.22 | 0.00 | 0.00 | 0.43 | 0.03 | 1 | Fibroblast |
| Wnt2 | 1.20 | 0.00 | 0.00 | 0.38 | 0.02 | 1 | Fibroblast |
| Mylk | 1.19 | 0.00 | 0.00 | 0.78 | 0.23 | 1 | Fibroblast |
| Rap2a | 1.19 | 0.00 | 0.00 | 0.55 | 0.12 | 1 | Fibroblast |
| Vcam1 | 1.18 | 0.00 | 0.00 | 0.40 | 0.03 | 1 | Fibroblast |
| Spon1 | 1.17 | 0.00 | 0.00 | 0.42 | 0.04 | 1 | Fibroblast |
| Mxra8 | 1.15 | 0.00 | 0.00 | 0.58 | 0.13 | 1 | Fibroblast |
| Serping1 | 1.13 | 0.00 | 0.00 | 0.53 | 0.09 | 1 | Fibroblast |
| Phex | 1.12 | 0.00 | 0.00 | 0.33 | 0.01 | 1 | Fibroblast |
| Iigp1 | 1.11 | 0.00 | 0.00 | 0.49 | 0.10 | 1 | Fibroblast |
| Snai2 | 1.11 | 0.00 | 0.00 | 0.48 | 0.09 | 1 | Fibroblast |
| Specc1l | 1.11 | 0.00 | 0.00 | 0.46 | 0.09 | 1 | Fibroblast |
| Mgp | 1.10 | 0.00 | 0.00 | 0.89 | 0.26 | 1 | Fibroblast |
| Atp1a2 | 1.10 | 0.00 | 0.00 | 0.46 | 0.07 | 1 | Fibroblast |
| Olfml3 | 1.09 | 0.00 | 0.00 | 0.43 | 0.05 | 1 | Fibroblast |
| Fblim1 | 1.05 | 0.00 | 0.00 | 0.56 | 0.13 | 1 | Fibroblast |
| Tns3 | 1.05 | 0.00 | 0.00 | 0.43 | 0.08 | 1 | Fibroblast |
| Col1a1 | 1.05 | 0.00 | 0.00 | 0.81 | 0.21 | 1 | Fibroblast |
| Gpm6b | 1.04 | 0.00 | 0.00 | 0.43 | 0.08 | 1 | Fibroblast |
| Ablim3 | 1.04 | 0.00 | 0.00 | 0.34 | 0.02 | 1 | Fibroblast |
| Igfbp4 | 1.03 | 0.00 | 0.00 | 0.66 | 0.22 | 1 | Fibroblast |
| Bgn | 1.03 | 0.00 | 0.00 | 0.70 | 0.20 | 1 | Fibroblast |
| Fbln5 | 1.03 | 0.00 | 0.00 | 0.59 | 0.14 | 1 | Fibroblast |
| Col3a1 | 1.02 | 0.00 | 0.00 | 0.96 | 0.27 | 1 | Fibroblast |
| Phlda1 | 1.02 | 0.00 | 0.00 | 0.59 | 0.17 | 1 | Fibroblast |
| Adarb1 | 1.01 | 0.00 | 0.00 | 0.37 | 0.05 | 1 | Fibroblast |
| Col6a3 | 1.01 | 0.00 | 0.00 | 0.42 | 0.07 | 1 | Fibroblast |
| 4-Sep | 1.00 | 0.00 | 0.00 | 0.67 | 0.19 | 1 | Fibroblast |
| Hivep3 | 1.00 | 0.00 | 0.00 | 0.37 | 0.05 | 1 | Fibroblast |
| Nebl | 0.99 | 0.00 | 0.00 | 0.41 | 0.07 | 1 | Fibroblast |
| Ces1d | 0.97 | 0.00 | 0.00 | 0.47 | 0.14 | 1 | Fibroblast |
| Clec3b | 0.97 | 0.00 | 0.00 | 0.37 | 0.04 | 1 | Fibroblast |
| Chst2 | 0.97 | 0.00 | 0.00 | 0.33 | 0.04 | 1 | Fibroblast |
| Fbn1 | 0.96 | 0.00 | 0.00 | 0.53 | 0.14 | 1 | Fibroblast |
| Tril | 0.96 | 0.00 | 0.00 | 0.35 | 0.05 | 1 | Fibroblast |
| Ednra | 0.95 | 0.00 | 0.00 | 0.37 | 0.06 | 1 | Fibroblast |
| Cacna1d | 0.95 | 0.00 | 0.00 | 0.33 | 0.03 | 1 | Fibroblast |
| 0610007N19Rik | 0.95 | 0.00 | 0.00 | 0.48 | 0.12 | 1 | Fibroblast |
| Cpxm1 | 0.94 | 0.00 | 0.00 | 0.28 | 0.02 | 1 | Fibroblast |
| Csrp1 | 0.94 | 0.00 | 0.00 | 0.49 | 0.14 | 1 | Fibroblast |
| Angpt1 | 0.92 | 0.00 | 0.00 | 0.34 | 0.05 | 1 | Fibroblast |
| Zyx | 0.91 | 0.00 | 0.00 | 0.53 | 0.17 | 1 | Fibroblast |
| Ccdc80 | 0.91 | 0.00 | 0.00 | 0.41 | 0.09 | 1 | Fibroblast |
| Frem1 | 0.91 | 0.00 | 0.00 | 0.31 | 0.04 | 1 | Fibroblast |
| Acadsb | 0.90 | 0.00 | 0.00 | 0.38 | 0.08 | 1 | Fibroblast |
| Pdlim2 | 0.90 | 0.00 | 0.00 | 0.48 | 0.15 | 1 | Fibroblast |
| Slc36a2 | 0.89 | 0.00 | 0.00 | 0.28 | 0.02 | 1 | Fibroblast |
| Gsn | 0.89 | 0.00 | 0.00 | 0.69 | 0.21 | 1 | Fibroblast |
| Robo2 | 0.88 | 0.00 | 0.00 | 0.36 | 0.07 | 1 | Fibroblast |
| Sned1 | 0.88 | 0.00 | 0.00 | 0.28 | 0.04 | 1 | Fibroblast |
| Mesdc2 | 0.88 | 0.00 | 0.00 | 0.49 | 0.16 | 1 | Fibroblast |
| Loxl1 | 0.87 | 0.00 | 0.00 | 0.41 | 0.08 | 1 | Fibroblast |
| Nr2f2 | 0.87 | 0.00 | 0.00 | 0.55 | 0.18 | 1 | Fibroblast |
| Nfib | 0.86 | 0.00 | 0.00 | 0.83 | 0.44 | 1 | Fibroblast |
| Adamts17 | 0.85 | 0.00 | 0.00 | 0.26 | 0.02 | 1 | Fibroblast |
| Cnn2 | 0.85 | 0.00 | 0.00 | 0.73 | 0.36 | 1 | Fibroblast |
| Agpat4 | 0.85 | 0.00 | 0.00 | 0.41 | 0.11 | 1 | Fibroblast |
| Nrcam | 0.85 | 0.00 | 0.00 | 0.29 | 0.04 | 1 | Fibroblast |
| Ppp1r3c | 0.84 | 0.00 | 0.00 | 0.26 | 0.02 | 1 | Fibroblast |
| Col6a1 | 0.84 | 0.00 | 0.00 | 0.53 | 0.16 | 1 | Fibroblast |
| Vldlr | 0.84 | 0.00 | 0.00 | 0.38 | 0.09 | 1 | Fibroblast |
| Pcolce | 0.84 | 0.00 | 0.00 | 0.35 | 0.06 | 1 | Fibroblast |
| Col6a2 | 0.83 | 0.00 | 0.00 | 0.45 | 0.12 | 1 | Fibroblast |
| Spats2l | 0.83 | 0.00 | 0.00 | 0.32 | 0.07 | 1 | Fibroblast |
| 6030408B16Rik | 0.83 | 0.00 | 0.00 | 0.29 | 0.03 | 1 | Fibroblast |
| Stbd1 | 0.83 | 0.00 | 0.00 | 0.30 | 0.05 | 1 | Fibroblast |
| Tns1 | 0.83 | 0.00 | 0.00 | 0.57 | 0.20 | 1 | Fibroblast |
| Zeb2 | 0.83 | 0.00 | 0.00 | 0.68 | 0.28 | 1 | Fibroblast |
| Tmem119 | 0.82 | 0.00 | 0.00 | 0.29 | 0.03 | 1 | Fibroblast |
| Hadh | 0.82 | 0.00 | 0.00 | 0.43 | 0.15 | 1 | Fibroblast |
| Isoc1 | 0.81 | 0.00 | 0.00 | 0.30 | 0.06 | 1 | Fibroblast |
| Pcdhga9 | 0.81 | 0.00 | 0.00 | 0.45 | 0.15 | 1 | Fibroblast |
| C230081A13Rik | 0.81 | 0.00 | 0.00 | 0.34 | 0.08 | 1 | Fibroblast |
| Tmsb10 | 0.79 | 0.00 | 0.00 | 0.99 | 0.84 | 1 | Fibroblast |
| Hhip | 0.79 | 0.00 | 0.00 | 0.30 | 0.05 | 1 | Fibroblast |
| Gfra2 | 0.78 | 0.00 | 0.00 | 0.28 | 0.04 | 1 | Fibroblast |
| Dkk3 | 0.77 | 0.00 | 0.00 | 0.40 | 0.09 | 1 | Fibroblast |
| Fibin | 0.77 | 0.00 | 0.00 | 0.42 | 0.10 | 1 | Fibroblast |
| Bmp5 | 0.77 | 0.00 | 0.00 | 0.26 | 0.04 | 1 | Fibroblast |
| Lgals1 | 0.77 | 0.00 | 0.00 | 0.90 | 0.37 | 1 | Fibroblast |
| Ism1 | 0.76 | 0.00 | 0.00 | 0.32 | 0.07 | 1 | Fibroblast |
| Ppp1r14a | 0.76 | 0.00 | 0.00 | 0.34 | 0.06 | 1 | Fibroblast |
| Etv1 | 0.74 | 0.00 | 0.00 | 0.29 | 0.05 | 1 | Fibroblast |
| Lamb1 | 0.74 | 0.00 | 0.00 | 0.42 | 0.14 | 1 | Fibroblast |
| Rbms3 | 0.74 | 0.00 | 0.00 | 0.43 | 0.15 | 1 | Fibroblast |
| Prex2 | 0.72 | 0.00 | 0.00 | 0.46 | 0.16 | 1 | Fibroblast |
| Maged1 | 0.72 | 0.00 | 0.00 | 0.44 | 0.16 | 1 | Fibroblast |
| Rras2 | 0.71 | 0.00 | 0.00 | 0.26 | 0.05 | 1 | Fibroblast |
| Fkbp9 | 0.71 | 0.00 | 0.00 | 0.38 | 0.12 | 1 | Fibroblast |
| Xist | 0.70 | 0.00 | 0.00 | 0.79 | 0.48 | 1 | Fibroblast |
| Sh3bp5 | 0.70 | 0.00 | 0.00 | 0.47 | 0.17 | 1 | Fibroblast |
| Rarres2 | 0.70 | 0.00 | 0.00 | 0.41 | 0.11 | 1 | Fibroblast |
| Fxyd1 | 0.70 | 0.00 | 0.00 | 0.32 | 0.07 | 1 | Fibroblast |
| Col5a2 | 0.70 | 0.00 | 0.00 | 0.46 | 0.15 | 1 | Fibroblast |
| Fam171a1 | 0.69 | 0.00 | 0.00 | 0.29 | 0.07 | 1 | Fibroblast |
| Rnd3 | 0.68 | 0.00 | 0.00 | 0.26 | 0.06 | 1 | Fibroblast |
| Sparc | 0.68 | 0.00 | 0.00 | 0.99 | 0.71 | 1 | Fibroblast |
| Cdo1 | 0.66 | 0.00 | 0.00 | 0.27 | 0.05 | 1 | Fibroblast |
| Egflam | 0.66 | 0.00 | 0.00 | 0.28 | 0.06 | 1 | Fibroblast |
| Hsd11b1 | 0.64 | 0.00 | 0.00 | 0.32 | 0.07 | 1 | Fibroblast |
| Cryab | 0.60 | 0.00 | 0.00 | 0.39 | 0.12 | 1 | Fibroblast |
| Tpm1 | 0.59 | 0.00 | 0.00 | 0.87 | 0.49 | 1 | Fibroblast |
| Igf1 | 0.57 | 0.00 | 0.00 | 0.38 | 0.12 | 1 | Fibroblast |
| Cdkn1c | 0.52 | 0.00 | 0.00 | 0.45 | 0.16 | 1 | Fibroblast |
| Igfbp7 | 0.51 | 0.00 | 0.00 | 0.79 | 0.36 | 1 | Fibroblast |
| Sparcl1 | 0.48 | 0.00 | 0.00 | 0.77 | 0.36 | 1 | Fibroblast |
| Fstl1 | 0.46 | 0.00 | 0.00 | 0.84 | 0.41 | 1 | Fibroblast |
| Tnc | 0.44 | 0.00 | 0.00 | 0.40 | 0.12 | 1 | Fibroblast |
| Fmo2 | 0.37 | 0.00 | 0.00 | 0.49 | 0.14 | 1 | Fibroblast |
| Ugdh | 0.67 | 0.00 | 0.00 | 0.28 | 0.07 | 1 | Fibroblast |
| Mfap2 | 0.59 | 0.00 | 0.00 | 0.68 | 0.32 | 1 | Fibroblast |
| Nedd4 | 0.63 | 0.00 | 0.00 | 0.81 | 0.51 | 1 | Fibroblast |
| Lims1 | 0.71 | 0.00 | 0.00 | 0.56 | 0.26 | 1 | Fibroblast |
| Limd1 | 0.75 | 0.00 | 0.00 | 0.42 | 0.16 | 1 | Fibroblast |
| Antxr1 | 0.67 | 0.00 | 0.00 | 0.33 | 0.10 | 1 | Fibroblast |
| Nrp2 | 0.65 | 0.00 | 0.00 | 0.25 | 0.06 | 1 | Fibroblast |
| Tbx2 | 0.60 | 0.00 | 0.00 | 0.33 | 0.10 | 1 | Fibroblast |
| Rps4x | 0.43 | 0.00 | 0.00 | 0.96 | 0.86 | 1 | Fibroblast |
| Kdelc2 | 0.68 | 0.00 | 0.00 | 0.31 | 0.09 | 1 | Fibroblast |
| Timp2 | 0.66 | 0.00 | 0.00 | 0.47 | 0.19 | 1 | Fibroblast |
| Pbx1 | 0.72 | 0.00 | 0.00 | 0.44 | 0.18 | 1 | Fibroblast |
| Selenbp1 | 0.38 | 0.00 | 0.00 | 0.51 | 0.20 | 1 | Fibroblast |
| Nid1 | 0.63 | 0.00 | 0.00 | 0.49 | 0.21 | 1 | Fibroblast |
| Gng11 | 0.54 | 0.00 | 0.00 | 0.59 | 0.26 | 1 | Fibroblast |
| Cd81 | 0.59 | 0.00 | 0.00 | 0.69 | 0.38 | 1 | Fibroblast |
| H6pd | 0.58 | 0.00 | 0.00 | 0.27 | 0.08 | 1 | Fibroblast |
| Serpinh1 | 0.55 | 0.00 | 0.00 | 0.69 | 0.36 | 1 | Fibroblast |
| Lama4 | 0.63 | 0.00 | 0.00 | 0.34 | 0.12 | 1 | Fibroblast |
| Mbnl2 | 0.68 | 0.00 | 0.00 | 0.61 | 0.32 | 1 | Fibroblast |
| Hmgn1 | 0.61 | 0.00 | 0.00 | 0.71 | 0.43 | 1 | Fibroblast |
| Pltp | 0.61 | 0.00 | 0.00 | 0.42 | 0.16 | 1 | Fibroblast |
| Adamts2 | 0.53 | 0.00 | 0.00 | 0.29 | 0.09 | 1 | Fibroblast |
| Fkbp7 | 0.60 | 0.00 | 0.00 | 0.33 | 0.12 | 1 | Fibroblast |
| Lmo4 | 0.66 | 0.00 | 0.00 | 0.50 | 0.24 | 1 | Fibroblast |
| Eif4ebp1 | 0.58 | 0.00 | 0.00 | 0.33 | 0.12 | 1 | Fibroblast |
| Eid1 | 0.65 | 0.00 | 0.00 | 0.54 | 0.27 | 1 | Fibroblast |
| Adamts1 | 0.54 | 0.00 | 0.00 | 0.46 | 0.19 | 1 | Fibroblast |
| Tspan3 | 0.60 | 0.00 | 0.00 | 0.36 | 0.14 | 1 | Fibroblast |
| Ywhaq | 0.63 | 0.00 | 0.00 | 0.61 | 0.35 | 1 | Fibroblast |
| Fgfr1 | 0.60 | 0.00 | 0.00 | 0.28 | 0.09 | 1 | Fibroblast |
| Nisch | 0.63 | 0.00 | 0.00 | 0.54 | 0.28 | 1 | Fibroblast |
| Cdc16 | 0.63 | 0.00 | 0.00 | 0.36 | 0.14 | 1 | Fibroblast |
| 1700025G04Rik | 0.60 | 0.00 | 0.00 | 0.27 | 0.09 | 1 | Fibroblast |
| Tagln2 | 0.56 | 0.00 | 0.00 | 0.69 | 0.43 | 1 | Fibroblast |
| Tpm2 | 0.34 | 0.00 | 0.00 | 0.34 | 0.12 | 1 | Fibroblast |
| Emilin1 | 0.52 | 0.00 | 0.00 | 0.26 | 0.08 | 1 | Fibroblast |
| Prdx5 | 0.46 | 0.00 | 0.00 | 0.57 | 0.30 | 1 | Fibroblast |
| Tmem254a | 0.60 | 0.00 | 0.00 | 0.40 | 0.18 | 1 | Fibroblast |
| Rcn3 | 0.51 | 0.00 | 0.00 | 0.32 | 0.11 | 1 | Fibroblast |
| Bmp4 | 0.56 | 0.00 | 0.00 | 0.27 | 0.09 | 1 | Fibroblast |
| Cox6c | 0.48 | 0.00 | 0.00 | 0.82 | 0.64 | 1 | Fibroblast |
| Myl9 | 0.33 | 0.00 | 0.00 | 0.31 | 0.11 | 1 | Fibroblast |
| Pmp22 | 0.34 | 0.00 | 0.00 | 0.58 | 0.29 | 1 | Fibroblast |
| Tmem176a | 0.59 | 0.00 | 0.00 | 0.41 | 0.19 | 1 | Fibroblast |
| Lox | 0.41 | 0.00 | 0.00 | 0.33 | 0.12 | 1 | Fibroblast |
| Mrps6 | 0.58 | 0.00 | 0.00 | 0.27 | 0.09 | 1 | Fibroblast |
| Lima1 | 0.57 | 0.00 | 0.00 | 0.32 | 0.13 | 1 | Fibroblast |
| Tmem254c | 0.59 | 0.00 | 0.00 | 0.38 | 0.17 | 1 | Fibroblast |
| Ccnd2 | 0.53 | 0.00 | 0.00 | 0.52 | 0.26 | 1 | Fibroblast |
| Tmem254b | 0.60 | 0.00 | 0.00 | 0.40 | 0.18 | 1 | Fibroblast |
| Tgfb1i1 | 0.52 | 0.00 | 0.00 | 0.31 | 0.12 | 1 | Fibroblast |
| Gtf2h5 | 0.60 | 0.00 | 0.00 | 0.50 | 0.27 | 1 | Fibroblast |
| Gpr126 | 0.45 | 0.00 | 0.00 | 0.27 | 0.09 | 1 | Fibroblast |
| Flna | 0.50 | 0.00 | 0.00 | 0.45 | 0.21 | 1 | Fibroblast |
| Zmynd11 | 0.56 | 0.00 | 0.00 | 0.55 | 0.31 | 1 | Fibroblast |
| Ltbp4 | 0.47 | 0.00 | 0.00 | 0.27 | 0.09 | 1 | Fibroblast |
| S100a1 | 0.54 | 0.00 | 0.00 | 0.33 | 0.13 | 1 | Fibroblast |
| Cbx5 | 0.57 | 0.00 | 0.00 | 0.47 | 0.24 | 1 | Fibroblast |
| N6amt1 | 0.53 | 0.00 | 0.00 | 0.29 | 0.11 | 1 | Fibroblast |
| Lrp1 | 0.49 | 0.00 | 0.00 | 0.28 | 0.10 | 1 | Fibroblast |
| Slc25a4 | 0.45 | 0.00 | 0.00 | 0.72 | 0.48 | 1 | Fibroblast |
| Eva1b | 0.50 | 0.00 | 0.00 | 0.39 | 0.18 | 1 | Fibroblast |
| Nenf | 0.57 | 0.00 | 0.00 | 0.37 | 0.17 | 1 | Fibroblast |
| mt-Nd1 | 0.32 | 0.00 | 0.00 | 0.97 | 0.91 | 1 | Fibroblast |
| Cd63 | 0.45 | 0.00 | 0.00 | 0.52 | 0.28 | 1 | Fibroblast |
| Tacc1 | 0.56 | 0.00 | 0.00 | 0.42 | 0.21 | 1 | Fibroblast |
| Sox4 | 0.46 | 0.00 | 0.00 | 0.60 | 0.35 | 1 | Fibroblast |
| Ralb | 0.55 | 0.00 | 0.00 | 0.38 | 0.18 | 1 | Fibroblast |
| Hspa5 | 0.47 | 0.00 | 0.00 | 0.75 | 0.54 | 1 | Fibroblast |
| Slc25a5 | 0.47 | 0.00 | 0.00 | 0.76 | 0.58 | 1 | Fibroblast |
| Vcl | 0.49 | 0.00 | 0.00 | 0.28 | 0.12 | 1 | Fibroblast |
| Wnt5a | 0.28 | 0.00 | 0.00 | 0.26 | 0.09 | 1 | Fibroblast |
| Celf2 | 0.45 | 0.00 | 0.00 | 0.48 | 0.26 | 1 | Fibroblast |
| Gm4204 | 0.35 | 0.00 | 0.00 | 0.71 | 0.48 | 1 | Fibroblast |
| Blmh | 0.51 | 0.00 | 0.00 | 0.34 | 0.16 | 1 | Fibroblast |
| Twsg1 | 0.48 | 0.00 | 0.00 | 0.26 | 0.11 | 1 | Fibroblast |
| Scp2 | 0.51 | 0.00 | 0.00 | 0.52 | 0.31 | 1 | Fibroblast |
| Atpif1 | 0.41 | 0.00 | 0.00 | 0.72 | 0.51 | 1 | Fibroblast |
| Pls3 | 0.49 | 0.00 | 0.00 | 0.37 | 0.19 | 1 | Fibroblast |
| Myo6 | 0.49 | 0.00 | 0.00 | 0.29 | 0.13 | 1 | Fibroblast |
| Tuba1a | 0.35 | 0.00 | 0.00 | 0.73 | 0.49 | 1 | Fibroblast |
| Selenbp2 | 0.26 | 0.00 | 0.00 | 0.34 | 0.15 | 1 | Fibroblast |
| Socs2 | 0.37 | 0.00 | 0.00 | 0.55 | 0.32 | 1 | Fibroblast |
| Ptms | 0.47 | 0.00 | 0.00 | 0.59 | 0.38 | 1 | Fibroblast |
| Ech1 | 0.44 | 0.00 | 0.00 | 0.31 | 0.14 | 1 | Fibroblast |
| Cnpy2 | 0.48 | 0.00 | 0.00 | 0.36 | 0.18 | 1 | Fibroblast |
| Rps4y2 | 0.56 | 0.00 | 0.00 | 0.33 | 0.17 | 1 | Fibroblast |
| Gas5 | 0.38 | 0.00 | 0.00 | 0.72 | 0.52 | 1 | Fibroblast |
| Calr | 0.45 | 0.00 | 0.00 | 0.62 | 0.42 | 1 | Fibroblast |
| Ndufa4 | 0.40 | 0.00 | 0.00 | 0.72 | 0.53 | 1 | Fibroblast |
| Anxa6 | 0.45 | 0.00 | 0.00 | 0.37 | 0.19 | 1 | Fibroblast |
| A630007B06Rik | 0.50 | 0.00 | 0.00 | 0.36 | 0.18 | 1 | Fibroblast |
| Wsb1 | 0.45 | 0.00 | 0.00 | 0.38 | 0.20 | 1 | Fibroblast |
| Sec61b | 0.43 | 0.00 | 0.00 | 0.64 | 0.45 | 1 | Fibroblast |
| Cd302 | 0.36 | 0.00 | 0.00 | 0.30 | 0.13 | 1 | Fibroblast |
| Gm26735 | 0.42 | 0.00 | 0.00 | 0.36 | 0.18 | 1 | Fibroblast |
| Gpx8 | 0.41 | 0.00 | 0.00 | 0.29 | 0.13 | 1 | Fibroblast |
| Clic4 | 0.41 | 0.00 | 0.00 | 0.48 | 0.28 | 1 | Fibroblast |
| Ppic | 0.40 | 0.00 | 0.00 | 0.58 | 0.37 | 1 | Fibroblast |
| Smim4 | 0.45 | 0.00 | 0.00 | 0.26 | 0.11 | 1 | Fibroblast |
| Tgfb2 | 0.35 | 0.00 | 0.00 | 0.29 | 0.13 | 1 | Fibroblast |
| Maged2 | 0.40 | 0.00 | 0.00 | 0.33 | 0.16 | 1 | Fibroblast |
| Ckap4 | 0.46 | 0.00 | 0.00 | 0.33 | 0.17 | 1 | Fibroblast |
| Sub1 | 0.37 | 0.00 | 0.00 | 0.68 | 0.50 | 1 | Fibroblast |
| Npr3 | 0.36 | 0.00 | 0.00 | 0.27 | 0.12 | 1 | Fibroblast |
| Hmcn1 | 0.34 | 0.00 | 0.00 | 0.28 | 0.13 | 1 | Fibroblast |
| Tuba1b | 0.32 | 0.00 | 0.00 | 0.66 | 0.44 | 1 | Fibroblast |
| Nap1l1 | 0.28 | 0.00 | 0.00 | 0.67 | 0.45 | 1 | Fibroblast |
| Lamc1 | 0.45 | 0.00 | 0.00 | 0.30 | 0.15 | 1 | Fibroblast |
| Gas6 | 0.47 | 0.00 | 0.00 | 0.29 | 0.14 | 1 | Fibroblast |
| Dnaja1 | 0.40 | 0.00 | 0.00 | 0.67 | 0.50 | 1 | Fibroblast |
| Rad50 | 0.49 | 0.00 | 0.00 | 0.34 | 0.18 | 1 | Fibroblast |
| Ywhab | 0.35 | 0.00 | 0.00 | 0.73 | 0.55 | 1 | Fibroblast |
| Trove2 | 0.46 | 0.00 | 0.00 | 0.27 | 0.13 | 1 | Fibroblast |
| Tpm4 | 0.33 | 0.00 | 0.00 | 0.68 | 0.47 | 1 | Fibroblast |
| Rpl39 | 0.27 | 0.00 | 0.00 | 0.91 | 0.79 | 1 | Fibroblast |
| Hsbp1 | 0.43 | 0.00 | 0.00 | 0.48 | 0.30 | 1 | Fibroblast |
| Mettl7a1 | 0.29 | 0.00 | 0.00 | 0.39 | 0.20 | 1 | Fibroblast |
| Zbtb20 | 0.43 | 0.00 | 0.00 | 0.44 | 0.25 | 1 | Fibroblast |
| 7-Sep | 0.32 | 0.00 | 0.00 | 0.75 | 0.57 | 1 | Fibroblast |
| Pebp1 | 0.42 | 0.00 | 0.00 | 0.41 | 0.24 | 1 | Fibroblast |
| Atxn7l3b | 0.39 | 0.00 | 0.00 | 0.57 | 0.39 | 1 | Fibroblast |
| Prdx2 | 0.39 | 0.00 | 0.00 | 0.50 | 0.32 | 1 | Fibroblast |
| Hist1h2bc | 0.43 | 0.00 | 0.00 | 0.26 | 0.13 | 1 | Fibroblast |
| Rcn1 | 0.41 | 0.00 | 0.00 | 0.27 | 0.13 | 1 | Fibroblast |
| Ndufc1 | 0.41 | 0.00 | 0.00 | 0.47 | 0.29 | 1 | Fibroblast |
| Mmp14 | 0.37 | 0.00 | 0.00 | 0.42 | 0.24 | 1 | Fibroblast |
| Romo1 | 0.39 | 0.00 | 0.00 | 0.46 | 0.29 | 1 | Fibroblast |
| Thra | 0.41 | 0.00 | 0.00 | 0.31 | 0.16 | 1 | Fibroblast |
| Tcf12 | 0.44 | 0.00 | 0.00 | 0.31 | 0.16 | 1 | Fibroblast |
| Tsix | 0.42 | 0.00 | 0.00 | 0.32 | 0.17 | 1 | Fibroblast |
| Nfia | 0.38 | 0.00 | 0.00 | 0.39 | 0.23 | 1 | Fibroblast |
| Idh2 | 0.40 | 0.00 | 0.00 | 0.33 | 0.18 | 1 | Fibroblast |
| Gm10320 | 0.39 | 0.00 | 0.00 | 0.54 | 0.38 | 1 | Fibroblast |
| Cox8a | 0.34 | 0.00 | 0.00 | 0.70 | 0.55 | 1 | Fibroblast |
| Gnas | 0.26 | 0.00 | 0.00 | 0.81 | 0.66 | 1 | Fibroblast |
| Cox6a1 | 0.34 | 0.00 | 0.00 | 0.69 | 0.54 | 1 | Fibroblast |
| Eny2 | 0.41 | 0.00 | 0.00 | 0.37 | 0.22 | 1 | Fibroblast |
| Smim15 | 0.39 | 0.00 | 0.00 | 0.33 | 0.19 | 1 | Fibroblast |
| Cox7c | 0.31 | 0.00 | 0.00 | 0.76 | 0.62 | 1 | Fibroblast |
| Plin2 | 0.30 | 0.00 | 0.00 | 0.25 | 0.12 | 1 | Fibroblast |
| Ssr1 | 0.38 | 0.00 | 0.00 | 0.39 | 0.23 | 1 | Fibroblast |
| Sqstm1 | 0.35 | 0.00 | 0.00 | 0.47 | 0.31 | 1 | Fibroblast |
| Parva | 0.38 | 0.00 | 0.00 | 0.28 | 0.15 | 1 | Fibroblast |
| Gstp2 | 0.35 | 0.00 | 0.00 | 0.30 | 0.16 | 1 | Fibroblast |
| Gstp1 | 0.35 | 0.00 | 0.00 | 0.33 | 0.18 | 1 | Fibroblast |
| mt-Nd5 | 0.26 | 0.00 | 0.00 | 0.87 | 0.75 | 1 | Fibroblast |
| Nr3c1 | 0.38 | 0.00 | 0.00 | 0.39 | 0.24 | 1 | Fibroblast |
| Uqcrb | 0.33 | 0.00 | 0.00 | 0.58 | 0.43 | 1 | Fibroblast |
| Tmem204 | 0.27 | 0.00 | 0.00 | 0.28 | 0.15 | 1 | Fibroblast |
| Cfdp1 | 0.34 | 0.00 | 0.00 | 0.53 | 0.37 | 1 | Fibroblast |
| Ndufb9 | 0.35 | 0.00 | 0.00 | 0.51 | 0.36 | 1 | Fibroblast |
| Emc2 | 0.36 | 0.00 | 0.00 | 0.26 | 0.14 | 1 | Fibroblast |
| Comt | 0.38 | 0.00 | 0.00 | 0.26 | 0.14 | 1 | Fibroblast |
| Mtss1 | 0.36 | 0.00 | 0.00 | 0.26 | 0.14 | 1 | Fibroblast |
| Ostc | 0.35 | 0.00 | 0.00 | 0.36 | 0.22 | 1 | Fibroblast |
| Abracl | 0.33 | 0.00 | 0.00 | 0.36 | 0.22 | 1 | Fibroblast |
| Nktr | 0.33 | 0.00 | 0.00 | 0.51 | 0.35 | 1 | Fibroblast |
| Mprip | 0.37 | 0.00 | 0.00 | 0.33 | 0.20 | 1 | Fibroblast |
| Fus | 0.31 | 0.00 | 0.00 | 0.62 | 0.47 | 1 | Fibroblast |
| Ifrd1 | 0.38 | 0.00 | 0.00 | 0.27 | 0.15 | 1 | Fibroblast |
| Eif4a2 | 0.34 | 0.00 | 0.00 | 0.46 | 0.31 | 1 | Fibroblast |
| Tmem176b | 0.31 | 0.00 | 0.00 | 0.49 | 0.33 | 1 | Fibroblast |
| Marcksl1 | 0.29 | 0.00 | 0.00 | 0.32 | 0.19 | 1 | Fibroblast |
| Zranb2 | 0.35 | 0.00 | 0.00 | 0.38 | 0.24 | 1 | Fibroblast |
| Gm15013 | 0.35 | 0.00 | 0.00 | 0.50 | 0.36 | 1 | Fibroblast |
| Cox7b | 0.33 | 0.00 | 0.00 | 0.59 | 0.46 | 1 | Fibroblast |
| Kctd10 | 0.34 | 0.00 | 0.00 | 0.26 | 0.14 | 1 | Fibroblast |
| Son | 0.26 | 0.00 | 0.00 | 0.83 | 0.72 | 1 | Fibroblast |
| Gm9847 | 0.32 | 0.00 | 0.00 | 0.27 | 0.15 | 1 | Fibroblast |
| 15-Sep | 0.31 | 0.00 | 0.00 | 0.48 | 0.33 | 1 | Fibroblast |
| Atp5g1 | 0.34 | 0.00 | 0.00 | 0.34 | 0.21 | 1 | Fibroblast |
| Hmgn5 | 0.39 | 0.00 | 0.00 | 0.39 | 0.25 | 1 | Fibroblast |
| Atp5e | 0.28 | 0.00 | 0.00 | 0.64 | 0.49 | 1 | Fibroblast |
| Chd3 | 0.32 | 0.00 | 0.00 | 0.30 | 0.18 | 1 | Fibroblast |
| Gm21092 | 0.33 | 0.00 | 0.00 | 0.31 | 0.19 | 1 | Fibroblast |
| Mpc2 | 0.32 | 0.00 | 0.00 | 0.36 | 0.23 | 1 | Fibroblast |
| Eef1g | 0.26 | 0.00 | 0.00 | 0.63 | 0.47 | 1 | Fibroblast |
| Calu | 0.29 | 0.00 | 0.00 | 0.36 | 0.23 | 1 | Fibroblast |
| Eml4 | 0.32 | 0.00 | 0.00 | 0.29 | 0.17 | 1 | Fibroblast |
| Arf4 | 0.31 | 0.00 | 0.00 | 0.40 | 0.26 | 1 | Fibroblast |
| Ndufb3 | 0.32 | 0.00 | 0.00 | 0.44 | 0.30 | 1 | Fibroblast |
| Ndufb6 | 0.32 | 0.00 | 0.00 | 0.35 | 0.22 | 1 | Fibroblast |
| Uqcr10 | 0.30 | 0.00 | 0.00 | 0.49 | 0.35 | 1 | Fibroblast |
| Nfix | 0.30 | 0.00 | 0.00 | 0.28 | 0.16 | 1 | Fibroblast |
| Mrfap1 | 0.30 | 0.00 | 0.00 | 0.46 | 0.32 | 1 | Fibroblast |
| Pdcd4 | 0.27 | 0.00 | 0.00 | 0.35 | 0.22 | 1 | Fibroblast |
| Tbl1x | 0.31 | 0.00 | 0.00 | 0.37 | 0.24 | 1 | Fibroblast |
| Mtus1 | 0.29 | 0.00 | 0.00 | 0.28 | 0.16 | 1 | Fibroblast |
| Tceal8 | 0.31 | 0.00 | 0.00 | 0.30 | 0.19 | 1 | Fibroblast |
| Cox6b1 | 0.26 | 0.00 | 0.00 | 0.70 | 0.56 | 1 | Fibroblast |
| Csnk1a1 | 0.26 | 0.00 | 0.00 | 0.66 | 0.53 | 1 | Fibroblast |
| Hnrnph1 | 0.27 | 0.00 | 0.00 | 0.52 | 0.37 | 1 | Fibroblast |
| Myo1b | 0.29 | 0.00 | 0.00 | 0.27 | 0.16 | 1 | Fibroblast |
| Ppib | 0.27 | 0.00 | 0.00 | 0.56 | 0.42 | 1 | Fibroblast |
| Psmc5 | 0.32 | 0.00 | 0.00 | 0.34 | 0.22 | 1 | Fibroblast |
| Snrpd2 | 0.29 | 0.00 | 0.00 | 0.37 | 0.24 | 1 | Fibroblast |
| Ndufv3 | 0.34 | 0.00 | 0.00 | 0.35 | 0.23 | 1 | Fibroblast |
| Pdia4 | 0.30 | 0.00 | 0.00 | 0.32 | 0.20 | 1 | Fibroblast |
| Ndufa5 | 0.29 | 0.00 | 0.00 | 0.37 | 0.25 | 1 | Fibroblast |
| 9530068E07Rik | 0.30 | 0.00 | 0.00 | 0.27 | 0.17 | 1 | Fibroblast |
| Cul1 | 0.32 | 0.00 | 0.00 | 0.28 | 0.18 | 1 | Fibroblast |
| Ndufb11 | 0.28 | 0.00 | 0.00 | 0.45 | 0.33 | 1 | Fibroblast |
| Ier3ip1 | 0.27 | 0.00 | 0.00 | 0.42 | 0.29 | 1 | Fibroblast |
| Pura | 0.30 | 0.00 | 0.00 | 0.31 | 0.20 | 1 | Fibroblast |
| 1110004F10Rik | 0.27 | 0.00 | 0.00 | 0.40 | 0.28 | 1 | Fibroblast |
| Kif1b | 0.31 | 0.00 | 0.00 | 0.34 | 0.23 | 1 | Fibroblast |
| Pdap1 | 0.25 | 0.00 | 0.00 | 0.57 | 0.44 | 1 | Fibroblast |
| Acaa2 | 0.30 | 0.00 | 0.00 | 0.26 | 0.16 | 1 | Fibroblast |
| Gm6483 | 0.28 | 0.00 | 0.00 | 0.27 | 0.16 | 1 | Fibroblast |
| Pfdn5 | 0.25 | 0.00 | 0.00 | 0.61 | 0.49 | 1 | Fibroblast |
| Atp5k | 0.29 | 0.00 | 0.00 | 0.43 | 0.31 | 1 | Fibroblast |
| Smim11 | 0.29 | 0.00 | 0.00 | 0.28 | 0.17 | 1 | Fibroblast |
| Slirp | 0.27 | 0.00 | 0.00 | 0.32 | 0.21 | 1 | Fibroblast |
| Ndufb2 | 0.28 | 0.00 | 0.00 | 0.31 | 0.20 | 1 | Fibroblast |
| Srpk2 | 0.28 | 0.00 | 0.00 | 0.25 | 0.15 | 1 | Fibroblast |
| Atp5j2 | 0.25 | 0.00 | 0.00 | 0.55 | 0.43 | 1 | Fibroblast |
| 2410015M20Rik | 0.29 | 0.00 | 0.00 | 0.33 | 0.22 | 1 | Fibroblast |
| Hdlbp | 0.30 | 0.00 | 0.00 | 0.30 | 0.20 | 1 | Fibroblast |
| Smarcc2 | 0.29 | 0.00 | 0.00 | 0.26 | 0.16 | 1 | Fibroblast |
| Htatsf1 | 0.29 | 0.00 | 0.00 | 0.36 | 0.26 | 1 | Fibroblast |
| Sar1a | 0.29 | 0.00 | 0.00 | 0.30 | 0.20 | 1 | Fibroblast |
| Lamp2 | 0.26 | 0.00 | 0.00 | 0.37 | 0.26 | 1 | Fibroblast |
| Swi5 | 0.27 | 0.00 | 0.00 | 0.37 | 0.26 | 1 | Fibroblast |
| Pdia3 | 0.25 | 0.00 | 0.00 | 0.55 | 0.43 | 1 | Fibroblast |
| Cox20 | 0.27 | 0.00 | 0.00 | 0.27 | 0.17 | 1 | Fibroblast |
| Ube2a | 0.31 | 0.00 | 0.00 | 0.27 | 0.18 | 1 | Fibroblast |
| Sec61g | 0.27 | 0.00 | 0.00 | 0.45 | 0.34 | 1 | Fibroblast |
| Ing4 | 0.27 | 0.00 | 0.00 | 0.32 | 0.21 | 1 | Fibroblast |
| Mrpl52 | 0.25 | 0.00 | 0.00 | 0.43 | 0.32 | 1 | Fibroblast |
| Mocs2 | 0.25 | 0.00 | 0.00 | 0.29 | 0.19 | 1 | Fibroblast |
| Copb2 | 0.28 | 0.00 | 0.00 | 0.27 | 0.17 | 1 | Fibroblast |
| Pdia6 | 0.26 | 0.00 | 0.00 | 0.40 | 0.29 | 1 | Fibroblast |
| Bccip | 0.26 | 0.00 | 0.00 | 0.29 | 0.20 | 1 | Fibroblast |
| Anapc13 | 0.27 | 0.00 | 0.00 | 0.30 | 0.21 | 1 | Fibroblast |
| Ufc1 | 0.25 | 0.00 | 0.00 | 0.26 | 0.18 | 1 | Fibroblast |
| Aspn | 2.90 | 0.00 | 0.00 | 0.48 | 0.07 | 2 | Stromal Cell |
| Tgfbi | 2.66 | 0.00 | 0.00 | 0.97 | 0.13 | 2 | Stromal Cell |
| Tnc | 2.25 | 0.00 | 0.00 | 0.84 | 0.08 | 2 | Stromal Cell |
| Htra1 | 2.13 | 0.00 | 0.00 | 0.62 | 0.04 | 2 | Stromal Cell |
| P2ry14 | 2.07 | 0.00 | 0.00 | 0.71 | 0.03 | 2 | Stromal Cell |
| Tagln | 1.97 | 0.00 | 0.00 | 0.77 | 0.10 | 2 | Stromal Cell |
| Loxl2 | 1.96 | 0.00 | 0.00 | 0.72 | 0.09 | 2 | Stromal Cell |
| Igf1 | 1.94 | 0.00 | 0.00 | 0.72 | 0.09 | 2 | Stromal Cell |
| Myh11 | 1.93 | 0.00 | 0.00 | 0.74 | 0.06 | 2 | Stromal Cell |
| Wnt5a | 1.92 | 0.00 | 0.00 | 0.67 | 0.06 | 2 | Stromal Cell |
| Prss35 | 1.92 | 0.00 | 0.00 | 0.59 | 0.03 | 2 | Stromal Cell |
| Acta2 | 1.89 | 0.00 | 0.00 | 0.85 | 0.14 | 2 | Stromal Cell |
| Agt | 1.89 | 0.00 | 0.00 | 0.71 | 0.05 | 2 | Stromal Cell |
| Ltbp2 | 1.84 | 0.00 | 0.00 | 0.65 | 0.05 | 2 | Stromal Cell |
| Filip1l | 1.78 | 0.00 | 0.00 | 0.87 | 0.29 | 2 | Stromal Cell |
| Fstl1 | 1.73 | 0.00 | 0.00 | 0.98 | 0.40 | 2 | Stromal Cell |
| Serpine2 | 1.72 | 0.00 | 0.00 | 0.81 | 0.21 | 2 | Stromal Cell |
| Cald1 | 1.71 | 0.00 | 0.00 | 0.95 | 0.37 | 2 | Stromal Cell |
| Net1 | 1.69 | 0.00 | 0.00 | 0.64 | 0.14 | 2 | Stromal Cell |
| Col3a1 | 1.67 | 0.00 | 0.00 | 0.98 | 0.28 | 2 | Stromal Cell |
| Pdlim3 | 1.67 | 0.00 | 0.00 | 0.55 | 0.03 | 2 | Stromal Cell |
| Actg2 | 1.66 | 0.00 | 0.00 | 0.52 | 0.04 | 2 | Stromal Cell |
| Egfem1 | 1.63 | 0.00 | 0.00 | 0.53 | 0.02 | 2 | Stromal Cell |
| Spon2 | 1.61 | 0.00 | 0.00 | 0.53 | 0.03 | 2 | Stromal Cell |
| RP23-103I12.13 | 1.61 | 0.00 | 0.00 | 0.57 | 0.06 | 2 | Stromal Cell |
| Lgals1 | 1.58 | 0.00 | 0.00 | 0.96 | 0.37 | 2 | Stromal Cell |
| Mustn1 | 1.53 | 0.00 | 0.00 | 0.49 | 0.04 | 2 | Stromal Cell |
| Nrep | 1.49 | 0.00 | 0.00 | 0.93 | 0.39 | 2 | Stromal Cell |
| Nes | 1.48 | 0.00 | 0.00 | 0.70 | 0.18 | 2 | Stromal Cell |
| Pde5a | 1.45 | 0.00 | 0.00 | 0.58 | 0.08 | 2 | Stromal Cell |
| Sdc2 | 1.43 | 0.00 | 0.00 | 0.59 | 0.09 | 2 | Stromal Cell |
| Olfml2b | 1.42 | 0.00 | 0.00 | 0.45 | 0.03 | 2 | Stromal Cell |
| Ckb | 1.40 | 0.00 | 0.00 | 0.59 | 0.11 | 2 | Stromal Cell |
| Mdk | 1.37 | 0.00 | 0.00 | 0.70 | 0.15 | 2 | Stromal Cell |
| Car2 | 1.34 | 0.00 | 0.00 | 0.72 | 0.27 | 2 | Stromal Cell |
| Des | 1.33 | 0.00 | 0.00 | 0.41 | 0.03 | 2 | Stromal Cell |
| Gpr64 | 1.32 | 0.00 | 0.00 | 0.36 | 0.03 | 2 | Stromal Cell |
| Thbs1 | 1.31 | 0.00 | 0.00 | 0.50 | 0.10 | 2 | Stromal Cell |
| Tpm2 | 1.31 | 0.00 | 0.00 | 0.60 | 0.10 | 2 | Stromal Cell |
| Cd248 | 1.29 | 0.00 | 0.00 | 0.38 | 0.02 | 2 | Stromal Cell |
| Eln | 1.27 | 0.00 | 0.00 | 0.50 | 0.13 | 2 | Stromal Cell |
| Myl9 | 1.26 | 0.00 | 0.00 | 0.56 | 0.09 | 2 | Stromal Cell |
| Tpm1 | 1.25 | 0.00 | 0.00 | 0.92 | 0.49 | 2 | Stromal Cell |
| Tgfb2 | 1.25 | 0.00 | 0.00 | 0.51 | 0.11 | 2 | Stromal Cell |
| Col1a2 | 1.23 | 0.00 | 0.00 | 0.99 | 0.35 | 2 | Stromal Cell |
| Myh10 | 1.21 | 0.00 | 0.00 | 0.68 | 0.19 | 2 | Stromal Cell |
| Mylk | 1.21 | 0.00 | 0.00 | 0.75 | 0.24 | 2 | Stromal Cell |
| 6330403K07Rik | 1.20 | 0.00 | 0.00 | 0.38 | 0.02 | 2 | Stromal Cell |
| Dkk3 | 1.19 | 0.00 | 0.00 | 0.51 | 0.09 | 2 | Stromal Cell |
| Bgn | 1.19 | 0.00 | 0.00 | 0.73 | 0.20 | 2 | Stromal Cell |
| Col5a1 | 1.18 | 0.00 | 0.00 | 0.45 | 0.06 | 2 | Stromal Cell |
| Fndc1 | 1.17 | 0.00 | 0.00 | 0.37 | 0.03 | 2 | Stromal Cell |
| Hlf | 1.16 | 0.00 | 0.00 | 0.34 | 0.03 | 2 | Stromal Cell |
| Lsp1 | 1.16 | 0.00 | 0.00 | 0.56 | 0.16 | 2 | Stromal Cell |
| Mfap2 | 1.15 | 0.00 | 0.00 | 0.82 | 0.31 | 2 | Stromal Cell |
| Robo2 | 1.15 | 0.00 | 0.00 | 0.41 | 0.07 | 2 | Stromal Cell |
| Igfbp5 | 1.15 | 0.00 | 0.00 | 0.40 | 0.08 | 2 | Stromal Cell |
| Lpar1 | 1.15 | 0.00 | 0.00 | 0.35 | 0.05 | 2 | Stromal Cell |
| Pdgfra | 1.14 | 0.00 | 0.00 | 0.34 | 0.04 | 2 | Stromal Cell |
| C1qtnf7 | 1.12 | 0.00 | 0.00 | 0.41 | 0.07 | 2 | Stromal Cell |
| Pamr1 | 1.11 | 0.00 | 0.00 | 0.31 | 0.03 | 2 | Stromal Cell |
| Map1b | 1.10 | 0.00 | 0.00 | 0.40 | 0.08 | 2 | Stromal Cell |
| Rcn3 | 1.08 | 0.00 | 0.00 | 0.50 | 0.10 | 2 | Stromal Cell |
| Fam129a | 1.08 | 0.00 | 0.00 | 0.38 | 0.06 | 2 | Stromal Cell |
| Hmcn1 | 1.06 | 0.00 | 0.00 | 0.41 | 0.12 | 2 | Stromal Cell |
| Pam | 1.04 | 0.00 | 0.00 | 0.47 | 0.15 | 2 | Stromal Cell |
| Fbn1 | 1.03 | 0.00 | 0.00 | 0.55 | 0.14 | 2 | Stromal Cell |
| Adamts2 | 1.03 | 0.00 | 0.00 | 0.38 | 0.08 | 2 | Stromal Cell |
| Tgfb3 | 1.02 | 0.00 | 0.00 | 0.31 | 0.03 | 2 | Stromal Cell |
| Palld | 1.00 | 0.00 | 0.00 | 0.55 | 0.15 | 2 | Stromal Cell |
| Mgp | 1.00 | 0.00 | 0.00 | 0.71 | 0.29 | 2 | Stromal Cell |
| Angptl2 | 0.99 | 0.00 | 0.00 | 0.29 | 0.02 | 2 | Stromal Cell |
| Col1a1 | 0.99 | 0.00 | 0.00 | 0.78 | 0.22 | 2 | Stromal Cell |
| Cygb | 0.98 | 0.00 | 0.00 | 0.32 | 0.04 | 2 | Stromal Cell |
| Socs2 | 0.98 | 0.00 | 0.00 | 0.72 | 0.31 | 2 | Stromal Cell |
| Col27a1 | 0.97 | 0.00 | 0.00 | 0.30 | 0.04 | 2 | Stromal Cell |
| Lhfp | 0.97 | 0.00 | 0.00 | 0.50 | 0.13 | 2 | Stromal Cell |
| Creb3l2 | 0.96 | 0.00 | 0.00 | 0.41 | 0.11 | 2 | Stromal Cell |
| Tshz2 | 0.96 | 0.00 | 0.00 | 0.36 | 0.07 | 2 | Stromal Cell |
| Rarres2 | 0.95 | 0.00 | 0.00 | 0.47 | 0.11 | 2 | Stromal Cell |
| Ddr2 | 0.94 | 0.00 | 0.00 | 0.29 | 0.04 | 2 | Stromal Cell |
| Lhfpl2 | 0.94 | 0.00 | 0.00 | 0.35 | 0.07 | 2 | Stromal Cell |
| Tenm4 | 0.93 | 0.00 | 0.00 | 0.27 | 0.02 | 2 | Stromal Cell |
| Rbms3 | 0.92 | 0.00 | 0.00 | 0.47 | 0.15 | 2 | Stromal Cell |
| Itga9 | 0.92 | 0.00 | 0.00 | 0.36 | 0.07 | 2 | Stromal Cell |
| Serpinh1 | 0.92 | 0.00 | 0.00 | 0.76 | 0.36 | 2 | Stromal Cell |
| Samd4 | 0.91 | 0.00 | 0.00 | 0.32 | 0.07 | 2 | Stromal Cell |
| Grb10 | 0.91 | 0.00 | 0.00 | 0.46 | 0.14 | 2 | Stromal Cell |
| Col5a2 | 0.91 | 0.00 | 0.00 | 0.49 | 0.15 | 2 | Stromal Cell |
| Col6a3 | 0.91 | 0.00 | 0.00 | 0.37 | 0.08 | 2 | Stromal Cell |
| Nfib | 0.91 | 0.00 | 0.00 | 0.83 | 0.44 | 2 | Stromal Cell |
| Golim4 | 0.90 | 0.00 | 0.00 | 0.49 | 0.18 | 2 | Stromal Cell |
| Nnat | 0.90 | 0.00 | 0.00 | 0.29 | 0.06 | 2 | Stromal Cell |
| Pdzrn3 | 0.89 | 0.00 | 0.00 | 0.29 | 0.04 | 2 | Stromal Cell |
| Plat | 0.88 | 0.00 | 0.00 | 0.26 | 0.04 | 2 | Stromal Cell |
| Calu | 0.87 | 0.00 | 0.00 | 0.55 | 0.22 | 2 | Stromal Cell |
| Gpc4 | 0.87 | 0.00 | 0.00 | 0.26 | 0.03 | 2 | Stromal Cell |
| Mxra7 | 0.87 | 0.00 | 0.00 | 0.41 | 0.12 | 2 | Stromal Cell |
| Postn | 0.87 | 0.00 | 0.00 | 0.33 | 0.06 | 2 | Stromal Cell |
| Sox4 | 0.86 | 0.00 | 0.00 | 0.71 | 0.34 | 2 | Stromal Cell |
| Anxa5 | 0.85 | 0.00 | 0.00 | 0.70 | 0.38 | 2 | Stromal Cell |
| Pdgfrl | 0.85 | 0.00 | 0.00 | 0.25 | 0.02 | 2 | Stromal Cell |
| Etv1 | 0.85 | 0.00 | 0.00 | 0.29 | 0.06 | 2 | Stromal Cell |
| Cnn3 | 0.84 | 0.00 | 0.00 | 0.65 | 0.27 | 2 | Stromal Cell |
| Mmp14 | 0.82 | 0.00 | 0.00 | 0.55 | 0.23 | 2 | Stromal Cell |
| Leprel2 | 0.81 | 0.00 | 0.00 | 0.30 | 0.06 | 2 | Stromal Cell |
| Selm | 0.79 | 0.00 | 0.00 | 0.32 | 0.07 | 2 | Stromal Cell |
| Nexn | 0.79 | 0.00 | 0.00 | 0.55 | 0.19 | 2 | Stromal Cell |
| Sparc | 0.78 | 0.00 | 0.00 | 0.99 | 0.71 | 2 | Stromal Cell |
| Gpr126 | 0.78 | 0.00 | 0.00 | 0.34 | 0.09 | 2 | Stromal Cell |
| Col16a1 | 0.76 | 0.00 | 0.00 | 0.25 | 0.04 | 2 | Stromal Cell |
| Gm13889 | 0.75 | 0.00 | 0.00 | 0.27 | 0.05 | 2 | Stromal Cell |
| Pros1 | 0.75 | 0.00 | 0.00 | 0.32 | 0.08 | 2 | Stromal Cell |
| Cd44 | 0.73 | 0.00 | 0.00 | 0.49 | 0.18 | 2 | Stromal Cell |
| Prdm6 | 0.69 | 0.00 | 0.00 | 0.26 | 0.05 | 2 | Stromal Cell |
| Plac9a | 0.67 | 0.00 | 0.00 | 0.71 | 0.27 | 2 | Stromal Cell |
| Plac9b | 0.64 | 0.00 | 0.00 | 0.71 | 0.27 | 2 | Stromal Cell |
| Mex3a | 0.71 | 0.00 | 0.00 | 0.28 | 0.07 | 2 | Stromal Cell |
| Hsp90b1 | 0.59 | 0.00 | 0.00 | 0.92 | 0.75 | 2 | Stromal Cell |
| Ghr | 0.82 | 0.00 | 0.00 | 0.39 | 0.13 | 2 | Stromal Cell |
| Smtn | 0.67 | 0.00 | 0.00 | 0.27 | 0.07 | 2 | Stromal Cell |
| Ism1 | 0.70 | 0.00 | 0.00 | 0.29 | 0.07 | 2 | Stromal Cell |
| Fkbp7 | 0.75 | 0.00 | 0.00 | 0.37 | 0.12 | 2 | Stromal Cell |
| Sema3c | 0.67 | 0.00 | 0.00 | 0.49 | 0.18 | 2 | Stromal Cell |
| Prnp | 0.89 | 0.00 | 0.00 | 0.34 | 0.10 | 2 | Stromal Cell |
| Iigp1 | 0.75 | 0.00 | 0.00 | 0.37 | 0.12 | 2 | Stromal Cell |
| Rgs2 | 0.78 | 0.00 | 0.00 | 0.58 | 0.26 | 2 | Stromal Cell |
| Mmp2 | 0.67 | 0.00 | 0.00 | 0.38 | 0.12 | 2 | Stromal Cell |
| Egflam | 0.67 | 0.00 | 0.00 | 0.27 | 0.07 | 2 | Stromal Cell |
| Itgb1 | 0.61 | 0.00 | 0.00 | 0.86 | 0.61 | 2 | Stromal Cell |
| Ctgf | 0.85 | 0.00 | 0.00 | 0.27 | 0.07 | 2 | Stromal Cell |
| Timp2 | 0.76 | 0.00 | 0.00 | 0.48 | 0.19 | 2 | Stromal Cell |
| Gm9847 | 0.79 | 0.00 | 0.00 | 0.40 | 0.14 | 2 | Stromal Cell |
| Loxl1 | 0.76 | 0.00 | 0.00 | 0.32 | 0.09 | 2 | Stromal Cell |
| Pmepa1 | 0.78 | 0.00 | 0.00 | 0.32 | 0.09 | 2 | Stromal Cell |
| Celf2 | 0.77 | 0.00 | 0.00 | 0.56 | 0.26 | 2 | Stromal Cell |
| Adam19 | 0.69 | 0.00 | 0.00 | 0.29 | 0.08 | 2 | Stromal Cell |
| Lpp | 0.79 | 0.00 | 0.00 | 0.47 | 0.20 | 2 | Stromal Cell |
| Gm26735 | 0.77 | 0.00 | 0.00 | 0.44 | 0.17 | 2 | Stromal Cell |
| Zeb2 | 0.67 | 0.00 | 0.00 | 0.60 | 0.29 | 2 | Stromal Cell |
| Gja1 | 1.13 | 0.00 | 0.00 | 0.27 | 0.08 | 2 | Stromal Cell |
| Pdlim7 | 0.67 | 0.00 | 0.00 | 0.30 | 0.09 | 2 | Stromal Cell |
| Calm2 | 0.59 | 0.00 | 0.00 | 0.85 | 0.63 | 2 | Stromal Cell |
| Tbx5 | 0.56 | 0.00 | 0.00 | 0.26 | 0.07 | 2 | Stromal Cell |
| Tpm4 | 0.64 | 0.00 | 0.00 | 0.74 | 0.47 | 2 | Stromal Cell |
| Lmna | 0.75 | 0.00 | 0.00 | 0.45 | 0.18 | 2 | Stromal Cell |
| Rcn1 | 0.68 | 0.00 | 0.00 | 0.36 | 0.12 | 2 | Stromal Cell |
| Ppic | 0.65 | 0.00 | 0.00 | 0.65 | 0.36 | 2 | Stromal Cell |
| Gpm6b | 0.63 | 0.00 | 0.00 | 0.31 | 0.10 | 2 | Stromal Cell |
| Rora | 0.78 | 0.00 | 0.00 | 0.29 | 0.09 | 2 | Stromal Cell |
| P4hb | 0.66 | 0.00 | 0.00 | 0.65 | 0.38 | 2 | Stromal Cell |
| Tgfb1i1 | 0.61 | 0.00 | 0.00 | 0.34 | 0.12 | 2 | Stromal Cell |
| Tuba1a | 0.56 | 0.00 | 0.00 | 0.76 | 0.49 | 2 | Stromal Cell |
| Flna | 0.67 | 0.00 | 0.00 | 0.48 | 0.21 | 2 | Stromal Cell |
| Myl6 | 0.54 | 0.00 | 0.00 | 0.84 | 0.68 | 2 | Stromal Cell |
| Adamts10 | 0.66 | 0.00 | 0.00 | 0.30 | 0.10 | 2 | Stromal Cell |
| Pxdn | 0.63 | 0.00 | 0.00 | 0.29 | 0.09 | 2 | Stromal Cell |
| Serf1 | 0.63 | 0.00 | 0.00 | 0.29 | 0.10 | 2 | Stromal Cell |
| Lamb1 | 0.64 | 0.00 | 0.00 | 0.37 | 0.14 | 2 | Stromal Cell |
| Ptma | 0.37 | 0.00 | 0.00 | 0.97 | 0.90 | 2 | Stromal Cell |
| Ctsl | 0.57 | 0.00 | 0.00 | 0.65 | 0.37 | 2 | Stromal Cell |
| Col6a1 | 0.61 | 0.00 | 0.00 | 0.42 | 0.17 | 2 | Stromal Cell |
| Ryk | 0.67 | 0.00 | 0.00 | 0.31 | 0.11 | 2 | Stromal Cell |
| Mllt3 | 0.68 | 0.00 | 0.00 | 0.29 | 0.10 | 2 | Stromal Cell |
| Rrbp1 | 0.60 | 0.00 | 0.00 | 0.77 | 0.53 | 2 | Stromal Cell |
| Tspan3 | 0.64 | 0.00 | 0.00 | 0.36 | 0.14 | 2 | Stromal Cell |
| Cd81 | 0.54 | 0.00 | 0.00 | 0.66 | 0.39 | 2 | Stromal Cell |
| Fndc3b | 0.64 | 0.00 | 0.00 | 0.30 | 0.10 | 2 | Stromal Cell |
| Lama4 | 0.63 | 0.00 | 0.00 | 0.32 | 0.12 | 2 | Stromal Cell |
| Maged2 | 0.57 | 0.00 | 0.00 | 0.38 | 0.16 | 2 | Stromal Cell |
| Snai2 | 0.62 | 0.00 | 0.00 | 0.31 | 0.11 | 2 | Stromal Cell |
| Col6a2 | 0.56 | 0.00 | 0.00 | 0.34 | 0.13 | 2 | Stromal Cell |
| Cd63 | 0.54 | 0.00 | 0.00 | 0.54 | 0.28 | 2 | Stromal Cell |
| Ttc3 | 0.66 | 0.00 | 0.00 | 0.51 | 0.27 | 2 | Stromal Cell |
| Ilk | 0.61 | 0.00 | 0.00 | 0.44 | 0.21 | 2 | Stromal Cell |
| Csrp2 | 0.59 | 0.00 | 0.00 | 0.37 | 0.15 | 2 | Stromal Cell |
| Actn1 | 0.66 | 0.00 | 0.00 | 0.43 | 0.21 | 2 | Stromal Cell |
| Amotl1 | 0.68 | 0.00 | 0.00 | 0.34 | 0.15 | 2 | Stromal Cell |
| Cst3 | 0.31 | 0.00 | 0.00 | 0.80 | 0.57 | 2 | Stromal Cell |
| Mxra8 | 0.51 | 0.00 | 0.00 | 0.36 | 0.15 | 2 | Stromal Cell |
| Nfix | 0.63 | 0.00 | 0.00 | 0.36 | 0.16 | 2 | Stromal Cell |
| Cyr61 | 0.53 | 0.00 | 0.00 | 0.30 | 0.11 | 2 | Stromal Cell |
| Tshz1 | 0.62 | 0.00 | 0.00 | 0.29 | 0.11 | 2 | Stromal Cell |
| Nedd4 | 0.50 | 0.00 | 0.00 | 0.75 | 0.52 | 2 | Stromal Cell |
| Gpx8 | 0.57 | 0.00 | 0.00 | 0.32 | 0.13 | 2 | Stromal Cell |
| Atpif1 | 0.50 | 0.00 | 0.00 | 0.73 | 0.51 | 2 | Stromal Cell |
| Phldb2 | 0.61 | 0.00 | 0.00 | 0.51 | 0.28 | 2 | Stromal Cell |
| Fgfr1 | 0.56 | 0.00 | 0.00 | 0.27 | 0.10 | 2 | Stromal Cell |
| Rcn2 | 0.59 | 0.00 | 0.00 | 0.32 | 0.14 | 2 | Stromal Cell |
| Arf4 | 0.57 | 0.00 | 0.00 | 0.47 | 0.26 | 2 | Stromal Cell |
| Antxr1 | 0.53 | 0.00 | 0.00 | 0.28 | 0.11 | 2 | Stromal Cell |
| Vim | 0.38 | 0.00 | 0.00 | 0.89 | 0.64 | 2 | Stromal Cell |
| Ccnd2 | 0.65 | 0.00 | 0.00 | 0.48 | 0.27 | 2 | Stromal Cell |
| Tjp2 | 0.63 | 0.00 | 0.00 | 0.33 | 0.15 | 2 | Stromal Cell |
| Minos1 | 0.53 | 0.00 | 0.00 | 0.63 | 0.43 | 2 | Stromal Cell |
| Slc25a4 | 0.47 | 0.00 | 0.00 | 0.69 | 0.49 | 2 | Stromal Cell |
| Dstn | 0.47 | 0.00 | 0.00 | 0.71 | 0.49 | 2 | Stromal Cell |
| Maged1 | 0.54 | 0.00 | 0.00 | 0.36 | 0.17 | 2 | Stromal Cell |
| Mapre2 | 0.50 | 0.00 | 0.00 | 0.25 | 0.10 | 2 | Stromal Cell |
| Fam115a | 0.55 | 0.00 | 0.00 | 0.30 | 0.13 | 2 | Stromal Cell |
| Hmgn1 | 0.47 | 0.00 | 0.00 | 0.65 | 0.44 | 2 | Stromal Cell |
| Anxa6 | 0.52 | 0.00 | 0.00 | 0.38 | 0.19 | 2 | Stromal Cell |
| Picalm | 0.60 | 0.00 | 0.00 | 0.38 | 0.20 | 2 | Stromal Cell |
| Parva | 0.52 | 0.00 | 0.00 | 0.32 | 0.15 | 2 | Stromal Cell |
| mt-Nd1 | 0.28 | 0.00 | 0.00 | 0.97 | 0.91 | 2 | Stromal Cell |
| Fkbp9 | 0.47 | 0.00 | 0.00 | 0.29 | 0.13 | 2 | Stromal Cell |
| Ckap4 | 0.53 | 0.00 | 0.00 | 0.35 | 0.17 | 2 | Stromal Cell |
| Tbx2 | 0.44 | 0.00 | 0.00 | 0.27 | 0.11 | 2 | Stromal Cell |
| Hspa5 | 0.46 | 0.00 | 0.00 | 0.73 | 0.54 | 2 | Stromal Cell |
| Tsc22d1 | 0.49 | 0.00 | 0.00 | 0.53 | 0.32 | 2 | Stromal Cell |
| Cdk4 | 0.53 | 0.00 | 0.00 | 0.42 | 0.23 | 2 | Stromal Cell |
| Cgnl1 | 0.55 | 0.00 | 0.00 | 0.26 | 0.11 | 2 | Stromal Cell |
| Laptm4a | 0.41 | 0.00 | 0.00 | 0.70 | 0.50 | 2 | Stromal Cell |
| Atp2b1 | 0.45 | 0.00 | 0.00 | 0.73 | 0.57 | 2 | Stromal Cell |
| Mpzl1 | 0.46 | 0.00 | 0.00 | 0.25 | 0.10 | 2 | Stromal Cell |
| Morf4l2 | 0.53 | 0.00 | 0.00 | 0.39 | 0.21 | 2 | Stromal Cell |
| Sparcl1 | 0.28 | 0.00 | 0.00 | 0.64 | 0.37 | 2 | Stromal Cell |
| Hsp90ab1 | 0.29 | 0.00 | 0.00 | 0.94 | 0.88 | 2 | Stromal Cell |
| Ptms | 0.50 | 0.00 | 0.00 | 0.58 | 0.39 | 2 | Stromal Cell |
| Twsg1 | 0.51 | 0.00 | 0.00 | 0.25 | 0.11 | 2 | Stromal Cell |
| Nfic | 0.46 | 0.00 | 0.00 | 0.42 | 0.23 | 2 | Stromal Cell |
| Fam101b | 0.47 | 0.00 | 0.00 | 0.34 | 0.17 | 2 | Stromal Cell |
| Itm2c | 0.50 | 0.00 | 0.00 | 0.39 | 0.21 | 2 | Stromal Cell |
| Prdx2 | 0.47 | 0.00 | 0.00 | 0.51 | 0.32 | 2 | Stromal Cell |
| Nfia | 0.49 | 0.00 | 0.00 | 0.41 | 0.23 | 2 | Stromal Cell |
| Dynlrb1 | 0.48 | 0.00 | 0.00 | 0.51 | 0.33 | 2 | Stromal Cell |
| Marcksl1 | 0.44 | 0.00 | 0.00 | 0.36 | 0.18 | 2 | Stromal Cell |
| Ndufa11 | 0.50 | 0.00 | 0.00 | 0.44 | 0.26 | 2 | Stromal Cell |
| Cdc42ep3 | 0.55 | 0.00 | 0.00 | 0.28 | 0.14 | 2 | Stromal Cell |
| Nupr1 | 0.36 | 0.00 | 0.00 | 0.34 | 0.17 | 2 | Stromal Cell |
| Gng12 | 0.45 | 0.00 | 0.00 | 0.25 | 0.11 | 2 | Stromal Cell |
| Serf2 | 0.41 | 0.00 | 0.00 | 0.71 | 0.55 | 2 | Stromal Cell |
| Fibin | 0.39 | 0.00 | 0.00 | 0.26 | 0.12 | 2 | Stromal Cell |
| Fblim1 | 0.33 | 0.00 | 0.00 | 0.32 | 0.15 | 2 | Stromal Cell |
| Nptn | 0.46 | 0.00 | 0.00 | 0.37 | 0.20 | 2 | Stromal Cell |
| Calr | 0.47 | 0.00 | 0.00 | 0.59 | 0.42 | 2 | Stromal Cell |
| Dst | 0.52 | 0.00 | 0.00 | 0.26 | 0.13 | 2 | Stromal Cell |
| Lamc1 | 0.43 | 0.00 | 0.00 | 0.30 | 0.15 | 2 | Stromal Cell |
| 2-Sep | 0.43 | 0.00 | 0.00 | 0.52 | 0.34 | 2 | Stromal Cell |
| Gnb4 | 0.44 | 0.00 | 0.00 | 0.26 | 0.12 | 2 | Stromal Cell |
| Mgll | 0.38 | 0.00 | 0.00 | 0.30 | 0.15 | 2 | Stromal Cell |
| Zfp260 | 0.45 | 0.00 | 0.00 | 0.25 | 0.12 | 2 | Stromal Cell |
| 11-Sep | 0.44 | 0.00 | 0.00 | 0.38 | 0.21 | 2 | Stromal Cell |
| Pls3 | 0.44 | 0.00 | 0.00 | 0.35 | 0.19 | 2 | Stromal Cell |
| mt-Cytb | 0.26 | 0.00 | 0.00 | 0.91 | 0.82 | 2 | Stromal Cell |
| Prdx4 | 0.43 | 0.00 | 0.00 | 0.30 | 0.15 | 2 | Stromal Cell |
| 0610007N19Rik | 0.38 | 0.00 | 0.00 | 0.28 | 0.14 | 2 | Stromal Cell |
| Npm1 | 0.33 | 0.00 | 0.00 | 0.72 | 0.57 | 2 | Stromal Cell |
| Pdia6 | 0.43 | 0.00 | 0.00 | 0.46 | 0.29 | 2 | Stromal Cell |
| Epb4.1l2 | 0.41 | 0.00 | 0.00 | 0.33 | 0.18 | 2 | Stromal Cell |
| Svil | 0.50 | 0.00 | 0.00 | 0.30 | 0.16 | 2 | Stromal Cell |
| Arid5b | 0.46 | 0.00 | 0.00 | 0.37 | 0.21 | 2 | Stromal Cell |
| Ssr2 | 0.41 | 0.00 | 0.00 | 0.46 | 0.30 | 2 | Stromal Cell |
| F2r | 0.37 | 0.00 | 0.00 | 0.27 | 0.13 | 2 | Stromal Cell |
| Pebp1 | 0.40 | 0.00 | 0.00 | 0.40 | 0.24 | 2 | Stromal Cell |
| Rsu1 | 0.40 | 0.00 | 0.00 | 0.34 | 0.19 | 2 | Stromal Cell |
| 9530068E07Rik | 0.40 | 0.00 | 0.00 | 0.30 | 0.17 | 2 | Stromal Cell |
| Ppib | 0.38 | 0.00 | 0.00 | 0.58 | 0.42 | 2 | Stromal Cell |
| Ostc | 0.39 | 0.00 | 0.00 | 0.37 | 0.22 | 2 | Stromal Cell |
| Pdap1 | 0.35 | 0.00 | 0.00 | 0.59 | 0.44 | 2 | Stromal Cell |
| Ddah2 | 0.32 | 0.00 | 0.00 | 0.42 | 0.25 | 2 | Stromal Cell |
| Tmed9 | 0.40 | 0.00 | 0.00 | 0.40 | 0.25 | 2 | Stromal Cell |
| Ddost | 0.41 | 0.00 | 0.00 | 0.32 | 0.18 | 2 | Stromal Cell |
| Cfdp1 | 0.36 | 0.00 | 0.00 | 0.53 | 0.37 | 2 | Stromal Cell |
| Csnk1a1 | 0.33 | 0.00 | 0.00 | 0.67 | 0.53 | 2 | Stromal Cell |
| Thoc7 | 0.40 | 0.00 | 0.00 | 0.47 | 0.31 | 2 | Stromal Cell |
| Myl12a | 0.29 | 0.00 | 0.00 | 0.76 | 0.65 | 2 | Stromal Cell |
| Yipf5 | 0.36 | 0.00 | 0.00 | 0.25 | 0.13 | 2 | Stromal Cell |
| Pdia3 | 0.34 | 0.00 | 0.00 | 0.58 | 0.43 | 2 | Stromal Cell |
| Lman1 | 0.39 | 0.00 | 0.00 | 0.31 | 0.18 | 2 | Stromal Cell |
| 15-Sep | 0.38 | 0.00 | 0.00 | 0.48 | 0.33 | 2 | Stromal Cell |
| Sfr1 | 0.37 | 0.00 | 0.00 | 0.48 | 0.33 | 2 | Stromal Cell |
| Oxct1 | 0.40 | 0.00 | 0.00 | 0.26 | 0.14 | 2 | Stromal Cell |
| Dnajc10 | 0.37 | 0.00 | 0.00 | 0.26 | 0.14 | 2 | Stromal Cell |
| Tia1 | 0.36 | 0.00 | 0.00 | 0.35 | 0.21 | 2 | Stromal Cell |
| Nr2f2 | 0.27 | 0.00 | 0.00 | 0.35 | 0.20 | 2 | Stromal Cell |
| Cbx1 | 0.34 | 0.00 | 0.00 | 0.56 | 0.41 | 2 | Stromal Cell |
| Eprs | 0.39 | 0.00 | 0.00 | 0.35 | 0.22 | 2 | Stromal Cell |
| Kdelr2 | 0.37 | 0.00 | 0.00 | 0.31 | 0.18 | 2 | Stromal Cell |
| Slmap | 0.43 | 0.00 | 0.00 | 0.27 | 0.15 | 2 | Stromal Cell |
| Myh9 | 0.34 | 0.00 | 0.00 | 0.47 | 0.32 | 2 | Stromal Cell |
| Tmem59 | 0.36 | 0.00 | 0.00 | 0.54 | 0.40 | 2 | Stromal Cell |
| mt-Nd2 | 0.26 | 0.00 | 0.00 | 0.80 | 0.69 | 2 | Stromal Cell |
| Cox20 | 0.33 | 0.00 | 0.00 | 0.30 | 0.17 | 2 | Stromal Cell |
| Eif4g2 | 0.31 | 0.00 | 0.00 | 0.55 | 0.40 | 2 | Stromal Cell |
| Arl1 | 0.37 | 0.00 | 0.00 | 0.32 | 0.20 | 2 | Stromal Cell |
| Rbbp7 | 0.33 | 0.00 | 0.00 | 0.38 | 0.25 | 2 | Stromal Cell |
| Myo1b | 0.33 | 0.00 | 0.00 | 0.28 | 0.16 | 2 | Stromal Cell |
| Bccip | 0.37 | 0.00 | 0.00 | 0.32 | 0.20 | 2 | Stromal Cell |
| Rtn4 | 0.36 | 0.00 | 0.00 | 0.35 | 0.23 | 2 | Stromal Cell |
| Psmd7 | 0.33 | 0.00 | 0.00 | 0.41 | 0.28 | 2 | Stromal Cell |
| Nars | 0.35 | 0.00 | 0.00 | 0.45 | 0.32 | 2 | Stromal Cell |
| Zfp36l1 | 0.28 | 0.00 | 0.00 | 0.58 | 0.45 | 2 | Stromal Cell |
| Mrfap1 | 0.32 | 0.00 | 0.00 | 0.46 | 0.33 | 2 | Stromal Cell |
| Map1lc3a | 0.39 | 0.00 | 0.00 | 0.31 | 0.20 | 2 | Stromal Cell |
| Erh | 0.34 | 0.00 | 0.00 | 0.40 | 0.28 | 2 | Stromal Cell |
| Gm4540 | 0.35 | 0.00 | 0.00 | 0.38 | 0.26 | 2 | Stromal Cell |
| Copb2 | 0.35 | 0.00 | 0.00 | 0.28 | 0.17 | 2 | Stromal Cell |
| Vezf1 | 0.35 | 0.00 | 0.00 | 0.35 | 0.23 | 2 | Stromal Cell |
| Pdia4 | 0.34 | 0.00 | 0.00 | 0.32 | 0.20 | 2 | Stromal Cell |
| Larp7 | 0.33 | 0.00 | 0.00 | 0.27 | 0.17 | 2 | Stromal Cell |
| Cnpy2 | 0.33 | 0.00 | 0.00 | 0.30 | 0.19 | 2 | Stromal Cell |
| Sec13 | 0.32 | 0.00 | 0.00 | 0.28 | 0.17 | 2 | Stromal Cell |
| Gstp1 | 0.28 | 0.00 | 0.00 | 0.30 | 0.19 | 2 | Stromal Cell |
| Glg1 | 0.33 | 0.00 | 0.00 | 0.30 | 0.19 | 2 | Stromal Cell |
| Sar1a | 0.33 | 0.00 | 0.00 | 0.31 | 0.20 | 2 | Stromal Cell |
| Spcs1 | 0.30 | 0.00 | 0.00 | 0.41 | 0.29 | 2 | Stromal Cell |
| Hdlbp | 0.34 | 0.00 | 0.00 | 0.31 | 0.20 | 2 | Stromal Cell |
| Zbtb20 | 0.36 | 0.00 | 0.00 | 0.38 | 0.26 | 2 | Stromal Cell |
| Gm10222 | 0.27 | 0.00 | 0.00 | 0.56 | 0.42 | 2 | Stromal Cell |
| Ccdc41 | 0.30 | 0.00 | 0.00 | 0.27 | 0.16 | 2 | Stromal Cell |
| Purb | 0.33 | 0.00 | 0.00 | 0.48 | 0.37 | 2 | Stromal Cell |
| Rpn2 | 0.33 | 0.00 | 0.00 | 0.31 | 0.20 | 2 | Stromal Cell |
| Romo1 | 0.32 | 0.00 | 0.00 | 0.41 | 0.29 | 2 | Stromal Cell |
| Pura | 0.38 | 0.00 | 0.00 | 0.30 | 0.20 | 2 | Stromal Cell |
| Dynll1 | 0.26 | 0.00 | 0.00 | 0.57 | 0.44 | 2 | Stromal Cell |
| Tmem167 | 0.30 | 0.00 | 0.00 | 0.32 | 0.21 | 2 | Stromal Cell |
| Ssr3 | 0.27 | 0.00 | 0.00 | 0.42 | 0.30 | 2 | Stromal Cell |
| Canx | 0.27 | 0.00 | 0.00 | 0.60 | 0.48 | 2 | Stromal Cell |
| Selk | 0.28 | 0.00 | 0.00 | 0.43 | 0.31 | 2 | Stromal Cell |
| mt-Nd4l | 0.28 | 0.00 | 0.00 | 0.49 | 0.37 | 2 | Stromal Cell |
| Cbx3 | 0.28 | 0.00 | 0.00 | 0.46 | 0.34 | 2 | Stromal Cell |
| Ccdc23 | 0.29 | 0.00 | 0.00 | 0.25 | 0.16 | 2 | Stromal Cell |
| Idh2 | 0.29 | 0.00 | 0.00 | 0.29 | 0.19 | 2 | Stromal Cell |
| Gm5611 | 0.27 | 0.00 | 0.00 | 0.46 | 0.35 | 2 | Stromal Cell |
| Ranbp1 | 0.27 | 0.00 | 0.00 | 0.40 | 0.28 | 2 | Stromal Cell |
| Ldha | 0.26 | 0.00 | 0.00 | 0.40 | 0.29 | 2 | Stromal Cell |
| Snrpd3 | 0.27 | 0.00 | 0.00 | 0.43 | 0.31 | 2 | Stromal Cell |
| Hnrnpr | 0.30 | 0.00 | 0.00 | 0.35 | 0.24 | 2 | Stromal Cell |
| Tln1 | 0.25 | 0.00 | 0.00 | 0.51 | 0.39 | 2 | Stromal Cell |
| Ift20 | 0.29 | 0.00 | 0.00 | 0.35 | 0.25 | 2 | Stromal Cell |
| Serinc1 | 0.28 | 0.00 | 0.00 | 0.39 | 0.29 | 2 | Stromal Cell |
| Gm6563 | 0.27 | 0.00 | 0.00 | 0.44 | 0.33 | 2 | Stromal Cell |
| Fundc2 | 0.28 | 0.00 | 0.00 | 0.25 | 0.16 | 2 | Stromal Cell |
| Gm10068 | 0.26 | 0.00 | 0.00 | 0.41 | 0.30 | 2 | Stromal Cell |
| Ginm1 | 0.30 | 0.00 | 0.00 | 0.33 | 0.23 | 2 | Stromal Cell |
| Swi5 | 0.28 | 0.00 | 0.00 | 0.36 | 0.26 | 2 | Stromal Cell |
| Ddx1 | 0.30 | 0.00 | 0.00 | 0.25 | 0.16 | 2 | Stromal Cell |
| Txnl1 | 0.28 | 0.00 | 0.00 | 0.28 | 0.18 | 2 | Stromal Cell |
| Gsk3b | 0.28 | 0.00 | 0.00 | 0.37 | 0.27 | 2 | Stromal Cell |
| Khdrbs1 | 0.27 | 0.00 | 0.00 | 0.41 | 0.31 | 2 | Stromal Cell |
| Rab1 | 0.25 | 0.00 | 0.00 | 0.41 | 0.30 | 2 | Stromal Cell |
| Chd4 | 0.26 | 0.00 | 0.00 | 0.57 | 0.48 | 2 | Stromal Cell |
| Rbpms | 0.28 | 0.00 | 0.00 | 0.27 | 0.18 | 2 | Stromal Cell |
| Hsbp1 | 0.25 | 0.00 | 0.00 | 0.41 | 0.30 | 2 | Stromal Cell |
| Vcp | 0.25 | 0.00 | 0.00 | 0.45 | 0.35 | 2 | Stromal Cell |
| Nr3c1 | 0.28 | 0.00 | 0.00 | 0.34 | 0.25 | 2 | Stromal Cell |
| Id2 | 0.26 | 0.00 | 0.00 | 0.43 | 0.33 | 2 | Stromal Cell |
| 1810022K09Rik | 0.27 | 0.00 | 0.00 | 0.32 | 0.23 | 2 | Stromal Cell |
| Kif1b | 0.27 | 0.00 | 0.00 | 0.32 | 0.23 | 2 | Stromal Cell |
| Pbx1 | 0.26 | 0.00 | 0.00 | 0.28 | 0.19 | 2 | Stromal Cell |
| Cfl2 | 0.26 | 0.00 | 0.00 | 0.34 | 0.24 | 2 | Stromal Cell |
| Myadm | 0.25 | 0.00 | 0.00 | 0.28 | 0.19 | 2 | Stromal Cell |
| Ilf2 | 0.28 | 0.00 | 0.00 | 0.27 | 0.19 | 2 | Stromal Cell |
| Psma4 | 0.25 | 0.00 | 0.00 | 0.41 | 0.31 | 2 | Stromal Cell |
| Vimp | 0.26 | 0.00 | 0.00 | 0.39 | 0.29 | 2 | Stromal Cell |
| Arcn1 | 0.26 | 0.00 | 0.00 | 0.29 | 0.21 | 2 | Stromal Cell |
| Mpc2 | 0.25 | 0.00 | 0.00 | 0.32 | 0.23 | 2 | Stromal Cell |
| Copb1 | 0.26 | 0.00 | 0.00 | 0.29 | 0.20 | 2 | Stromal Cell |
| Vdac1 | 0.25 | 0.00 | 0.00 | 0.28 | 0.20 | 2 | Stromal Cell |
| Pik3ca | 0.28 | 0.00 | 0.00 | 0.27 | 0.19 | 2 | Stromal Cell |
| Syncrip | 0.26 | 0.00 | 0.00 | 0.33 | 0.24 | 2 | Stromal Cell |
| Fip1l1 | 0.28 | 0.00 | 0.00 | 0.25 | 0.18 | 2 | Stromal Cell |
| 2700089E24Rik | 0.26 | 0.00 | 0.00 | 0.31 | 0.23 | 2 | Stromal Cell |
| Golgb1 | 0.28 | 0.00 | 0.00 | 0.40 | 0.32 | 2 | Stromal Cell |
| Clip1 | 0.29 | 0.00 | 0.00 | 0.27 | 0.20 | 2 | Stromal Cell |
| Sftpc | 3.45 | 0.00 | 0.00 | 1.00 | 0.60 | 3 | AT2 Cell |
| Sftpa1 | 3.25 | 0.00 | 0.00 | 1.00 | 0.15 | 3 | AT2 Cell |
| Sftpb | 2.96 | 0.00 | 0.00 | 1.00 | 0.12 | 3 | AT2 Cell |
| Cxcl15 | 2.90 | 0.00 | 0.00 | 0.96 | 0.09 | 3 | AT2 Cell |
| Chi3l1 | 2.77 | 0.00 | 0.00 | 0.82 | 0.06 | 3 | AT2 Cell |
| Hc | 2.63 | 0.00 | 0.00 | 0.87 | 0.06 | 3 | AT2 Cell |
| Slc34a2 | 2.62 | 0.00 | 0.00 | 0.99 | 0.09 | 3 | AT2 Cell |
| Lyz1 | 2.52 | 0.00 | 0.00 | 0.79 | 0.14 | 3 | AT2 Cell |
| Retnla | 2.50 | 0.00 | 0.00 | 0.37 | 0.09 | 3 | AT2 Cell |
| Sftpd | 2.44 | 0.00 | 0.00 | 0.95 | 0.10 | 3 | AT2 Cell |
| S100g | 2.37 | 0.00 | 0.00 | 0.87 | 0.07 | 3 | AT2 Cell |
| Lamp3 | 2.35 | 0.00 | 0.00 | 0.84 | 0.06 | 3 | AT2 Cell |
| Bex2 | 2.19 | 0.00 | 0.00 | 0.73 | 0.06 | 3 | AT2 Cell |
| Lpcat1 | 2.09 | 0.00 | 0.00 | 0.89 | 0.13 | 3 | AT2 Cell |
| Sfta2 | 2.06 | 0.00 | 0.00 | 0.87 | 0.10 | 3 | AT2 Cell |
| Ppp1r14c | 2.04 | 0.00 | 0.00 | 0.74 | 0.06 | 3 | AT2 Cell |
| Npc2 | 1.99 | 0.00 | 0.00 | 0.99 | 0.51 | 3 | AT2 Cell |
| Lcn2 | 1.98 | 0.00 | 0.00 | 0.50 | 0.04 | 3 | AT2 Cell |
| Wfdc2 | 1.97 | 0.00 | 0.00 | 0.88 | 0.11 | 3 | AT2 Cell |
| Napsa | 1.94 | 0.00 | 0.00 | 0.91 | 0.15 | 3 | AT2 Cell |
| Lyz2 | 1.93 | 0.00 | 0.00 | 0.87 | 0.22 | 3 | AT2 Cell |
| Egfl6 | 1.91 | 0.00 | 0.00 | 0.65 | 0.04 | 3 | AT2 Cell |
| Scd1 | 1.88 | 0.00 | 0.00 | 0.57 | 0.05 | 3 | AT2 Cell |
| Dram1 | 1.86 | 0.00 | 0.00 | 0.73 | 0.07 | 3 | AT2 Cell |
| Abca3 | 1.83 | 0.00 | 0.00 | 0.71 | 0.08 | 3 | AT2 Cell |
| Bex4 | 1.81 | 0.00 | 0.00 | 0.74 | 0.08 | 3 | AT2 Cell |
| Spink5 | 1.79 | 0.00 | 0.00 | 0.49 | 0.04 | 3 | AT2 Cell |
| Muc1 | 1.70 | 0.00 | 0.00 | 0.61 | 0.05 | 3 | AT2 Cell |
| Rbpjl | 1.63 | 0.00 | 0.00 | 0.51 | 0.03 | 3 | AT2 Cell |
| Cbr2 | 1.63 | 0.00 | 0.00 | 0.82 | 0.12 | 3 | AT2 Cell |
| Bex1 | 1.63 | 0.00 | 0.00 | 0.61 | 0.06 | 3 | AT2 Cell |
| Lgi3 | 1.59 | 0.00 | 0.00 | 0.55 | 0.04 | 3 | AT2 Cell |
| Fabp5 | 1.57 | 0.00 | 0.00 | 0.59 | 0.10 | 3 | AT2 Cell |
| Neat1 | 1.49 | 0.00 | 0.00 | 0.79 | 0.23 | 3 | AT2 Cell |
| Ptprf | 1.45 | 0.00 | 0.00 | 0.65 | 0.12 | 3 | AT2 Cell |
| Ctsh | 1.43 | 0.00 | 0.00 | 0.72 | 0.17 | 3 | AT2 Cell |
| Alcam | 1.43 | 0.00 | 0.00 | 0.77 | 0.19 | 3 | AT2 Cell |
| Sdc4 | 1.42 | 0.00 | 0.00 | 0.72 | 0.19 | 3 | AT2 Cell |
| Il33 | 1.40 | 0.00 | 0.00 | 0.36 | 0.01 | 3 | AT2 Cell |
| Car8 | 1.33 | 0.00 | 0.00 | 0.46 | 0.05 | 3 | AT2 Cell |
| Pi4k2b | 1.31 | 0.00 | 0.00 | 0.48 | 0.07 | 3 | AT2 Cell |
| Acsl4 | 1.31 | 0.00 | 0.00 | 0.50 | 0.09 | 3 | AT2 Cell |
| Ppp1r9a | 1.30 | 0.00 | 0.00 | 0.53 | 0.11 | 3 | AT2 Cell |
| Tgoln1 | 1.28 | 0.00 | 0.00 | 0.58 | 0.16 | 3 | AT2 Cell |
| Mid1ip1 | 1.27 | 0.00 | 0.00 | 0.49 | 0.12 | 3 | AT2 Cell |
| Ces1d | 1.26 | 0.00 | 0.00 | 0.64 | 0.13 | 3 | AT2 Cell |
| Fasn | 1.25 | 0.00 | 0.00 | 0.41 | 0.05 | 3 | AT2 Cell |
| Etv5 | 1.25 | 0.00 | 0.00 | 0.41 | 0.04 | 3 | AT2 Cell |
| Cldn18 | 1.25 | 0.00 | 0.00 | 0.84 | 0.12 | 3 | AT2 Cell |
| Atp1b1 | 1.24 | 0.00 | 0.00 | 0.73 | 0.14 | 3 | AT2 Cell |
| Ank3 | 1.23 | 0.00 | 0.00 | 0.42 | 0.05 | 3 | AT2 Cell |
| Pla2g1b | 1.21 | 0.00 | 0.00 | 0.36 | 0.03 | 3 | AT2 Cell |
| Atp6v1c2 | 1.21 | 0.00 | 0.00 | 0.37 | 0.03 | 3 | AT2 Cell |
| Atp8a1 | 1.19 | 0.00 | 0.00 | 0.54 | 0.12 | 3 | AT2 Cell |
| Rnase4 | 1.17 | 0.00 | 0.00 | 0.57 | 0.16 | 3 | AT2 Cell |
| Abcd3 | 1.17 | 0.00 | 0.00 | 0.56 | 0.16 | 3 | AT2 Cell |
| Sdc1 | 1.15 | 0.00 | 0.00 | 0.48 | 0.09 | 3 | AT2 Cell |
| Irx3 | 1.14 | 0.00 | 0.00 | 0.38 | 0.05 | 3 | AT2 Cell |
| Nkx2-1 | 1.14 | 0.00 | 0.00 | 0.45 | 0.07 | 3 | AT2 Cell |
| Snhg11 | 1.11 | 0.00 | 0.00 | 0.29 | 0.02 | 3 | AT2 Cell |
| Wbp5 | 1.10 | 0.00 | 0.00 | 0.87 | 0.45 | 3 | AT2 Cell |
| Zdhhc3 | 1.10 | 0.00 | 0.00 | 0.48 | 0.13 | 3 | AT2 Cell |
| Ngfrap1 | 1.09 | 0.00 | 0.00 | 0.59 | 0.21 | 3 | AT2 Cell |
| Fgfr2 | 1.08 | 0.00 | 0.00 | 0.36 | 0.05 | 3 | AT2 Cell |
| Nucb2 | 1.07 | 0.00 | 0.00 | 0.48 | 0.14 | 3 | AT2 Cell |
| Mgst1 | 1.07 | 0.00 | 0.00 | 0.62 | 0.17 | 3 | AT2 Cell |
| Dcxr | 1.06 | 0.00 | 0.00 | 0.44 | 0.08 | 3 | AT2 Cell |
| Soat1 | 1.05 | 0.00 | 0.00 | 0.41 | 0.08 | 3 | AT2 Cell |
| Myo5c | 1.02 | 0.00 | 0.00 | 0.36 | 0.06 | 3 | AT2 Cell |
| Trf | 1.02 | 0.00 | 0.00 | 0.35 | 0.08 | 3 | AT2 Cell |
| Brd7 | 1.01 | 0.00 | 0.00 | 0.60 | 0.23 | 3 | AT2 Cell |
| Epcam | 1.01 | 0.00 | 0.00 | 0.40 | 0.08 | 3 | AT2 Cell |
| Tinag | 1.00 | 0.00 | 0.00 | 0.28 | 0.02 | 3 | AT2 Cell |
| Cldn3 | 1.00 | 0.00 | 0.00 | 0.48 | 0.09 | 3 | AT2 Cell |
| Cat | 0.99 | 0.00 | 0.00 | 0.40 | 0.10 | 3 | AT2 Cell |
| Atp11a | 0.99 | 0.00 | 0.00 | 0.39 | 0.09 | 3 | AT2 Cell |
| Gclc | 0.96 | 0.00 | 0.00 | 0.31 | 0.07 | 3 | AT2 Cell |
| Ctsc | 0.95 | 0.00 | 0.00 | 0.54 | 0.17 | 3 | AT2 Cell |
| Spint2 | 0.94 | 0.00 | 0.00 | 0.40 | 0.09 | 3 | AT2 Cell |
| Exosc7 | 0.93 | 0.00 | 0.00 | 0.40 | 0.11 | 3 | AT2 Cell |
| Meg3 | 0.92 | 0.00 | 0.00 | 0.64 | 0.19 | 3 | AT2 Cell |
| Mecom | 0.92 | 0.00 | 0.00 | 0.35 | 0.09 | 3 | AT2 Cell |
| Gas6 | 0.91 | 0.00 | 0.00 | 0.42 | 0.13 | 3 | AT2 Cell |
| Ptplad1 | 0.90 | 0.00 | 0.00 | 0.32 | 0.08 | 3 | AT2 Cell |
| Scd2 | 0.89 | 0.00 | 0.00 | 0.52 | 0.19 | 3 | AT2 Cell |
| Smim6 | 0.88 | 0.00 | 0.00 | 0.25 | 0.02 | 3 | AT2 Cell |
| Prr15l | 0.87 | 0.00 | 0.00 | 0.28 | 0.05 | 3 | AT2 Cell |
| Cmtm8 | 0.86 | 0.00 | 0.00 | 0.38 | 0.11 | 3 | AT2 Cell |
| H2afj | 0.86 | 0.00 | 0.00 | 0.62 | 0.30 | 3 | AT2 Cell |
| Tmem243 | 0.86 | 0.00 | 0.00 | 0.29 | 0.06 | 3 | AT2 Cell |
| Prnp | 0.85 | 0.00 | 0.00 | 0.36 | 0.10 | 3 | AT2 Cell |
| Rab27b | 0.85 | 0.00 | 0.00 | 0.29 | 0.06 | 3 | AT2 Cell |
| Snx25 | 0.84 | 0.00 | 0.00 | 0.26 | 0.05 | 3 | AT2 Cell |
| Id2 | 0.84 | 0.00 | 0.00 | 0.67 | 0.31 | 3 | AT2 Cell |
| Chchd10 | 0.84 | 0.00 | 0.00 | 0.44 | 0.11 | 3 | AT2 Cell |
| Mettl7a1 | 0.84 | 0.00 | 0.00 | 0.52 | 0.19 | 3 | AT2 Cell |
| Cdh1 | 0.84 | 0.00 | 0.00 | 0.30 | 0.06 | 3 | AT2 Cell |
| Slc39a8 | 0.80 | 0.00 | 0.00 | 0.30 | 0.06 | 3 | AT2 Cell |
| Cadm1 | 0.79 | 0.00 | 0.00 | 0.41 | 0.12 | 3 | AT2 Cell |
| Cd74 | 0.74 | 0.00 | 0.00 | 0.46 | 0.10 | 3 | AT2 Cell |
| Mt1 | 0.72 | 0.00 | 0.00 | 0.44 | 0.14 | 3 | AT2 Cell |
| H2-Aa | 0.69 | 0.00 | 0.00 | 0.33 | 0.06 | 3 | AT2 Cell |
| Ager | 0.68 | 0.00 | 0.00 | 0.72 | 0.13 | 3 | AT2 Cell |
| Malat1 | 0.41 | 0.00 | 0.00 | 1.00 | 1.00 | 3 | AT2 Cell |
| Serpinb6b | 0.78 | 0.00 | 0.00 | 0.43 | 0.15 | 3 | AT2 Cell |
| Slc12a2 | 0.79 | 0.00 | 0.00 | 0.40 | 0.13 | 3 | AT2 Cell |
| Mbip | 0.80 | 0.00 | 0.00 | 0.33 | 0.10 | 3 | AT2 Cell |
| Acot1 | 0.74 | 0.00 | 0.00 | 0.26 | 0.06 | 3 | AT2 Cell |
| Gde1 | 0.77 | 0.00 | 0.00 | 0.38 | 0.13 | 3 | AT2 Cell |
| H2-Ab1 | 0.39 | 0.00 | 0.00 | 0.29 | 0.08 | 3 | AT2 Cell |
| Cldn7 | 0.65 | 0.00 | 0.00 | 0.26 | 0.07 | 3 | AT2 Cell |
| Taok3 | 0.75 | 0.00 | 0.00 | 0.43 | 0.17 | 3 | AT2 Cell |
| Itga9 | 0.69 | 0.00 | 0.00 | 0.28 | 0.08 | 3 | AT2 Cell |
| Nupr1 | 0.75 | 0.00 | 0.00 | 0.42 | 0.17 | 3 | AT2 Cell |
| Dpp4 | 0.66 | 0.00 | 0.00 | 0.34 | 0.11 | 3 | AT2 Cell |
| Pon3 | 0.69 | 0.00 | 0.00 | 0.29 | 0.09 | 3 | AT2 Cell |
| Rps19 | 0.48 | 0.00 | 0.00 | 0.94 | 0.82 | 3 | AT2 Cell |
| Scp2 | 0.77 | 0.00 | 0.00 | 0.57 | 0.31 | 3 | AT2 Cell |
| Vamp8 | 0.67 | 0.00 | 0.00 | 0.61 | 0.33 | 3 | AT2 Cell |
| Avpi1 | 0.69 | 0.00 | 0.00 | 0.26 | 0.07 | 3 | AT2 Cell |
| Uhrf1bp1l | 0.70 | 0.00 | 0.00 | 0.28 | 0.08 | 3 | AT2 Cell |
| Rai14 | 0.70 | 0.00 | 0.00 | 0.29 | 0.10 | 3 | AT2 Cell |
| Myh14 | 0.60 | 0.00 | 0.00 | 0.28 | 0.08 | 3 | AT2 Cell |
| Emb | 0.63 | 0.00 | 0.00 | 0.30 | 0.10 | 3 | AT2 Cell |
| Oat | 0.72 | 0.00 | 0.00 | 0.32 | 0.11 | 3 | AT2 Cell |
| App | 0.47 | 0.00 | 0.00 | 0.83 | 0.58 | 3 | AT2 Cell |
| Elovl1 | 0.72 | 0.00 | 0.00 | 0.35 | 0.14 | 3 | AT2 Cell |
| Cd36 | 0.38 | 0.00 | 0.00 | 0.63 | 0.31 | 3 | AT2 Cell |
| Bri3 | 0.68 | 0.00 | 0.00 | 0.42 | 0.19 | 3 | AT2 Cell |
| Acsl5 | 0.64 | 0.00 | 0.00 | 0.28 | 0.10 | 3 | AT2 Cell |
| Iqgap1 | 0.54 | 0.00 | 0.00 | 0.69 | 0.43 | 3 | AT2 Cell |
| Iah1 | 0.65 | 0.00 | 0.00 | 0.29 | 0.11 | 3 | AT2 Cell |
| Adk | 0.62 | 0.00 | 0.00 | 0.29 | 0.11 | 3 | AT2 Cell |
| Usp8 | 0.66 | 0.00 | 0.00 | 0.35 | 0.15 | 3 | AT2 Cell |
| Ptgs1 | 0.63 | 0.00 | 0.00 | 0.26 | 0.09 | 3 | AT2 Cell |
| Tmem245 | 0.59 | 0.00 | 0.00 | 0.26 | 0.09 | 3 | AT2 Cell |
| Chka | 0.59 | 0.00 | 0.00 | 0.26 | 0.09 | 3 | AT2 Cell |
| Secisbp2l | 0.61 | 0.00 | 0.00 | 0.36 | 0.17 | 3 | AT2 Cell |
| Cpm | 0.45 | 0.00 | 0.00 | 0.38 | 0.17 | 3 | AT2 Cell |
| Tmbim6 | 0.54 | 0.00 | 0.00 | 0.57 | 0.36 | 3 | AT2 Cell |
| Cd63 | 0.53 | 0.00 | 0.00 | 0.50 | 0.28 | 3 | AT2 Cell |
| Gm26924 | 0.62 | 0.00 | 0.00 | 0.73 | 0.56 | 3 | AT2 Cell |
| Dbi | 0.56 | 0.00 | 0.00 | 0.56 | 0.36 | 3 | AT2 Cell |
| Lpin2 | 0.50 | 0.00 | 0.00 | 0.31 | 0.14 | 3 | AT2 Cell |
| Fkbp4 | 0.56 | 0.00 | 0.00 | 0.30 | 0.14 | 3 | AT2 Cell |
| Mtch1 | 0.58 | 0.00 | 0.00 | 0.29 | 0.14 | 3 | AT2 Cell |
| Arl6ip1 | 0.36 | 0.00 | 0.00 | 0.71 | 0.51 | 3 | AT2 Cell |
| Bsg | 0.46 | 0.00 | 0.00 | 0.54 | 0.35 | 3 | AT2 Cell |
| Ccz1 | 0.86 | 0.00 | 0.00 | 0.27 | 0.12 | 3 | AT2 Cell |
| Ctnnb1 | 0.51 | 0.00 | 0.00 | 0.52 | 0.33 | 3 | AT2 Cell |
| Swt1 | 0.49 | 0.00 | 0.00 | 0.29 | 0.15 | 3 | AT2 Cell |
| Rnaset2b | 0.50 | 0.00 | 0.00 | 0.41 | 0.25 | 3 | AT2 Cell |
| Rnaset2a | 0.49 | 0.00 | 0.00 | 0.41 | 0.25 | 3 | AT2 Cell |
| Atp6v1a | 0.46 | 0.00 | 0.00 | 0.43 | 0.27 | 3 | AT2 Cell |
| Tmem30a | 0.48 | 0.00 | 0.00 | 0.43 | 0.27 | 3 | AT2 Cell |
| Ezr | 0.33 | 0.00 | 0.00 | 0.37 | 0.21 | 3 | AT2 Cell |
| Chd3 | 0.45 | 0.00 | 0.00 | 0.32 | 0.18 | 3 | AT2 Cell |
| Sepp1 | 0.26 | 0.00 | 0.00 | 0.59 | 0.40 | 3 | AT2 Cell |
| Tnfaip1 | 0.45 | 0.00 | 0.00 | 0.29 | 0.16 | 3 | AT2 Cell |
| Etfb | 0.43 | 0.00 | 0.00 | 0.42 | 0.27 | 3 | AT2 Cell |
| Dstn | 0.30 | 0.00 | 0.00 | 0.66 | 0.49 | 3 | AT2 Cell |
| 1500012F01Rik | 0.43 | 0.00 | 0.00 | 0.44 | 0.29 | 3 | AT2 Cell |
| Arf6 | 0.43 | 0.00 | 0.00 | 0.31 | 0.18 | 3 | AT2 Cell |
| Pcnt | 0.48 | 0.00 | 0.00 | 0.25 | 0.14 | 3 | AT2 Cell |
| Phldb2 | 0.36 | 0.00 | 0.00 | 0.42 | 0.28 | 3 | AT2 Cell |
| Tspo | 0.38 | 0.00 | 0.00 | 0.38 | 0.25 | 3 | AT2 Cell |
| Golga4 | 0.38 | 0.00 | 0.00 | 0.41 | 0.29 | 3 | AT2 Cell |
| Rpl21 | 0.26 | 0.00 | 0.00 | 0.84 | 0.74 | 3 | AT2 Cell |
| Snx4 | 0.40 | 0.00 | 0.00 | 0.37 | 0.24 | 3 | AT2 Cell |
| Ift20 | 0.39 | 0.00 | 0.00 | 0.37 | 0.25 | 3 | AT2 Cell |
| Rpl36al | 0.30 | 0.00 | 0.00 | 0.63 | 0.52 | 3 | AT2 Cell |
| Nfe2l2 | 0.35 | 0.00 | 0.00 | 0.36 | 0.24 | 3 | AT2 Cell |
| Apc | 0.34 | 0.00 | 0.00 | 0.34 | 0.23 | 3 | AT2 Cell |
| Vimp | 0.32 | 0.00 | 0.00 | 0.41 | 0.29 | 3 | AT2 Cell |
| Akap9 | 0.25 | 0.00 | 0.00 | 0.48 | 0.35 | 3 | AT2 Cell |
| Atp6v1g1 | 0.32 | 0.00 | 0.00 | 0.42 | 0.31 | 3 | AT2 Cell |
| Wls | 0.30 | 0.00 | 0.00 | 0.32 | 0.22 | 3 | AT2 Cell |
| Ddx17 | 0.33 | 0.00 | 0.00 | 0.34 | 0.24 | 3 | AT2 Cell |
| Use1 | 0.29 | 0.00 | 0.00 | 0.35 | 0.26 | 3 | AT2 Cell |
| Ndufc2 | 0.25 | 0.00 | 0.00 | 0.47 | 0.38 | 3 | AT2 Cell |
| Tceal8 | 0.34 | 0.00 | 0.00 | 0.27 | 0.19 | 3 | AT2 Cell |
| Higd1a | 0.27 | 0.00 | 0.00 | 0.26 | 0.18 | 3 | AT2 Cell |
| Fkbp2 | 0.30 | 0.00 | 0.00 | 0.25 | 0.18 | 3 | AT2 Cell |
| Cyb5 | 0.26 | 0.00 | 0.00 | 0.34 | 0.26 | 3 | AT2 Cell |
| Pik3ca | 0.27 | 0.00 | 0.00 | 0.26 | 0.19 | 3 | AT2 Cell |
| Plac8 | 2.61 | 0.00 | 0.00 | 0.87 | 0.13 | 4 | Monocyte |
| Lyz2 | 2.32 | 0.00 | 0.00 | 0.91 | 0.23 | 4 | Monocyte |
| Tyrobp | 2.26 | 0.00 | 0.00 | 0.87 | 0.08 | 4 | Monocyte |
| Fcer1g | 2.23 | 0.00 | 0.00 | 0.82 | 0.08 | 4 | Monocyte |
| Ccr2 | 2.22 | 0.00 | 0.00 | 0.63 | 0.04 | 4 | Monocyte |
| Alox5ap | 2.01 | 0.00 | 0.00 | 0.76 | 0.06 | 4 | Monocyte |
| Cybb | 2.01 | 0.00 | 0.00 | 0.71 | 0.06 | 4 | Monocyte |
| Lst1 | 2.00 | 0.00 | 0.00 | 0.63 | 0.04 | 4 | Monocyte |
| Ifi27l2a | 2.00 | 0.00 | 0.00 | 0.59 | 0.07 | 4 | Monocyte |
| Ctss | 1.94 | 0.00 | 0.00 | 0.89 | 0.11 | 4 | Monocyte |
| S100a4 | 1.92 | 0.00 | 0.00 | 0.60 | 0.04 | 4 | Monocyte |
| Ms4a6c | 1.89 | 0.00 | 0.00 | 0.57 | 0.04 | 4 | Monocyte |
| Ifitm6 | 1.73 | 0.00 | 0.00 | 0.46 | 0.01 | 4 | Monocyte |
| Fyb | 1.72 | 0.00 | 0.00 | 0.66 | 0.16 | 4 | Monocyte |
| Lgals3 | 1.71 | 0.00 | 0.00 | 0.71 | 0.12 | 4 | Monocyte |
| Cst3 | 1.71 | 0.00 | 0.00 | 0.95 | 0.57 | 4 | Monocyte |
| Plbd1 | 1.69 | 0.00 | 0.00 | 0.47 | 0.02 | 4 | Monocyte |
| Samsn1 | 1.67 | 0.00 | 0.00 | 0.49 | 0.04 | 4 | Monocyte |
| Clec4a3 | 1.66 | 0.00 | 0.00 | 0.44 | 0.02 | 4 | Monocyte |
| Msrb1 | 1.66 | 0.00 | 0.00 | 0.63 | 0.12 | 4 | Monocyte |
| Mpeg1 | 1.64 | 0.00 | 0.00 | 0.56 | 0.05 | 4 | Monocyte |
| Spi1 | 1.64 | 0.00 | 0.00 | 0.52 | 0.04 | 4 | Monocyte |
| Pla2g7 | 1.63 | 0.00 | 0.00 | 0.47 | 0.03 | 4 | Monocyte |
| Mnda | 1.62 | 0.00 | 0.00 | 0.46 | 0.04 | 4 | Monocyte |
| Rnase6 | 1.62 | 0.00 | 0.00 | 0.46 | 0.03 | 4 | Monocyte |
| Laptm5 | 1.60 | 0.00 | 0.00 | 0.72 | 0.11 | 4 | Monocyte |
| Psap | 1.58 | 0.00 | 0.00 | 0.87 | 0.33 | 4 | Monocyte |
| Ptprc | 1.56 | 0.00 | 0.00 | 0.68 | 0.12 | 4 | Monocyte |
| Wfdc17 | 1.55 | 0.00 | 0.00 | 0.38 | 0.02 | 4 | Monocyte |
| Ifi204 | 1.55 | 0.00 | 0.00 | 0.44 | 0.03 | 4 | Monocyte |
| Samhd1 | 1.51 | 0.00 | 0.00 | 0.66 | 0.17 | 4 | Monocyte |
| Cd52 | 1.50 | 0.00 | 0.00 | 0.86 | 0.17 | 4 | Monocyte |
| Cyba | 1.50 | 0.00 | 0.00 | 0.79 | 0.27 | 4 | Monocyte |
| Gm4070 | 1.48 | 0.00 | 0.00 | 0.51 | 0.10 | 4 | Monocyte |
| Gvin1 | 1.48 | 0.00 | 0.00 | 0.51 | 0.10 | 4 | Monocyte |
| Igsf6 | 1.47 | 0.00 | 0.00 | 0.41 | 0.03 | 4 | Monocyte |
| Pld4 | 1.45 | 0.00 | 0.00 | 0.45 | 0.03 | 4 | Monocyte |
| Slfn2 | 1.44 | 0.00 | 0.00 | 0.61 | 0.11 | 4 | Monocyte |
| Gpx1 | 1.43 | 0.00 | 0.00 | 0.86 | 0.41 | 4 | Monocyte |
| Coro1a | 1.43 | 0.00 | 0.00 | 0.52 | 0.08 | 4 | Monocyte |
| Lyz1 | 1.43 | 0.00 | 0.00 | 0.78 | 0.16 | 4 | Monocyte |
| Ccl6 | 1.41 | 0.00 | 0.00 | 0.65 | 0.07 | 4 | Monocyte |
| Gngt2 | 1.41 | 0.00 | 0.00 | 0.37 | 0.09 | 4 | Monocyte |
| Ifi205 | 1.40 | 0.00 | 0.00 | 0.37 | 0.03 | 4 | Monocyte |
| Apbb1ip | 1.40 | 0.00 | 0.00 | 0.49 | 0.07 | 4 | Monocyte |
| Itga4 | 1.37 | 0.00 | 0.00 | 0.46 | 0.08 | 4 | Monocyte |
| Ifitm3 | 1.37 | 0.00 | 0.00 | 0.79 | 0.38 | 4 | Monocyte |
| Sat1 | 1.37 | 0.00 | 0.00 | 0.75 | 0.31 | 4 | Monocyte |
| Cd300a | 1.36 | 0.00 | 0.00 | 0.37 | 0.02 | 4 | Monocyte |
| Bcl2a1d | 1.35 | 0.00 | 0.00 | 0.37 | 0.07 | 4 | Monocyte |
| Mndal | 1.34 | 0.00 | 0.00 | 0.57 | 0.14 | 4 | Monocyte |
| Gm2a | 1.34 | 0.00 | 0.00 | 0.45 | 0.07 | 4 | Monocyte |
| Arhgap30 | 1.32 | 0.00 | 0.00 | 0.50 | 0.08 | 4 | Monocyte |
| Cx3cr1 | 1.32 | 0.00 | 0.00 | 0.30 | 0.02 | 4 | Monocyte |
| Prdx5 | 1.31 | 0.00 | 0.00 | 0.69 | 0.30 | 4 | Monocyte |
| Arhgdib | 1.31 | 0.00 | 0.00 | 0.62 | 0.14 | 4 | Monocyte |
| Cytip | 1.28 | 0.00 | 0.00 | 0.45 | 0.08 | 4 | Monocyte |
| Bcl2a1a | 1.28 | 0.00 | 0.00 | 0.32 | 0.05 | 4 | Monocyte |
| Sirpb1c | 1.27 | 0.00 | 0.00 | 0.33 | 0.02 | 4 | Monocyte |
| Itgal | 1.24 | 0.00 | 0.00 | 0.30 | 0.03 | 4 | Monocyte |
| Tmsb4x | 1.24 | 0.00 | 0.00 | 1.00 | 0.88 | 4 | Monocyte |
| Lsp1 | 1.23 | 0.00 | 0.00 | 0.64 | 0.17 | 4 | Monocyte |
| Gpr141 | 1.19 | 0.00 | 0.00 | 0.28 | 0.01 | 4 | Monocyte |
| Lilrb4 | 1.18 | 0.00 | 0.00 | 0.40 | 0.04 | 4 | Monocyte |
| Taldo1 | 1.17 | 0.00 | 0.00 | 0.63 | 0.24 | 4 | Monocyte |
| Gsr | 1.17 | 0.00 | 0.00 | 0.37 | 0.07 | 4 | Monocyte |
| Arpc1b | 1.17 | 0.00 | 0.00 | 0.74 | 0.35 | 4 | Monocyte |
| Gda | 1.17 | 0.00 | 0.00 | 0.30 | 0.04 | 4 | Monocyte |
| Ptpn6 | 1.16 | 0.00 | 0.00 | 0.41 | 0.07 | 4 | Monocyte |
| Fam111a | 1.15 | 0.00 | 0.00 | 0.48 | 0.14 | 4 | Monocyte |
| Tpd52 | 1.14 | 0.00 | 0.00 | 0.40 | 0.09 | 4 | Monocyte |
| Csf1r | 1.13 | 0.00 | 0.00 | 0.31 | 0.03 | 4 | Monocyte |
| Slc11a1 | 1.13 | 0.00 | 0.00 | 0.27 | 0.02 | 4 | Monocyte |
| Hcls1 | 1.11 | 0.00 | 0.00 | 0.36 | 0.06 | 4 | Monocyte |
| Ly86 | 1.11 | 0.00 | 0.00 | 0.37 | 0.05 | 4 | Monocyte |
| B2m | 1.11 | 0.00 | 0.00 | 0.93 | 0.66 | 4 | Monocyte |
| Csf2ra | 1.11 | 0.00 | 0.00 | 0.34 | 0.05 | 4 | Monocyte |
| Bcl2a1b | 1.11 | 0.00 | 0.00 | 0.29 | 0.06 | 4 | Monocyte |
| Ptpn18 | 1.10 | 0.00 | 0.00 | 0.47 | 0.13 | 4 | Monocyte |
| Aif1 | 1.09 | 0.00 | 0.00 | 0.28 | 0.02 | 4 | Monocyte |
| Ncf2 | 1.08 | 0.00 | 0.00 | 0.31 | 0.03 | 4 | Monocyte |
| 1-Mar | 1.06 | 0.00 | 0.00 | 0.28 | 0.03 | 4 | Monocyte |
| AF251705 | 1.05 | 0.00 | 0.00 | 0.25 | 0.02 | 4 | Monocyte |
| Psmb8 | 1.05 | 0.00 | 0.00 | 0.51 | 0.17 | 4 | Monocyte |
| Sirpb1b | 1.05 | 0.00 | 0.00 | 0.26 | 0.02 | 4 | Monocyte |
| Nr4a1 | 1.04 | 0.00 | 0.00 | 0.28 | 0.06 | 4 | Monocyte |
| Fgr | 1.04 | 0.00 | 0.00 | 0.25 | 0.01 | 4 | Monocyte |
| Cotl1 | 1.04 | 0.00 | 0.00 | 0.51 | 0.16 | 4 | Monocyte |
| Hck | 1.03 | 0.00 | 0.00 | 0.28 | 0.03 | 4 | Monocyte |
| Pirb | 1.03 | 0.00 | 0.00 | 0.29 | 0.03 | 4 | Monocyte |
| Lcp1 | 1.02 | 0.00 | 0.00 | 0.58 | 0.21 | 4 | Monocyte |
| 1810033B17Rik | 1.00 | 0.00 | 0.00 | 0.28 | 0.02 | 4 | Monocyte |
| Fxyd5 | 1.00 | 0.00 | 0.00 | 0.50 | 0.16 | 4 | Monocyte |
| Itgb2 | 0.99 | 0.00 | 0.00 | 0.27 | 0.03 | 4 | Monocyte |
| S100a6 | 0.99 | 0.00 | 0.00 | 0.77 | 0.29 | 4 | Monocyte |
| Pygl | 0.99 | 0.00 | 0.00 | 0.39 | 0.10 | 4 | Monocyte |
| 9930111J21Rik2 | 0.99 | 0.00 | 0.00 | 0.40 | 0.10 | 4 | Monocyte |
| Ptpre | 0.96 | 0.00 | 0.00 | 0.29 | 0.05 | 4 | Monocyte |
| Plek | 0.96 | 0.00 | 0.00 | 0.32 | 0.06 | 4 | Monocyte |
| Pip4k2a | 0.95 | 0.00 | 0.00 | 0.28 | 0.05 | 4 | Monocyte |
| Il6ra | 0.93 | 0.00 | 0.00 | 0.38 | 0.09 | 4 | Monocyte |
| Ncf1 | 0.92 | 0.00 | 0.00 | 0.31 | 0.06 | 4 | Monocyte |
| Cfp | 0.91 | 0.00 | 0.00 | 0.27 | 0.04 | 4 | Monocyte |
| Hp | 0.90 | 0.00 | 0.00 | 0.40 | 0.10 | 4 | Monocyte |
| Arpc2 | 0.89 | 0.00 | 0.00 | 0.86 | 0.60 | 4 | Monocyte |
| Gp49a | 0.88 | 0.00 | 0.00 | 0.29 | 0.04 | 4 | Monocyte |
| Ms4a6b | 0.81 | 0.00 | 0.00 | 0.29 | 0.05 | 4 | Monocyte |
| Gm9843 | 0.77 | 0.00 | 0.00 | 0.99 | 0.92 | 4 | Monocyte |
| Fau | 0.73 | 0.00 | 0.00 | 0.98 | 0.86 | 4 | Monocyte |
| Actb | 0.59 | 0.00 | 0.00 | 1.00 | 0.98 | 4 | Monocyte |
| AI662270 | 0.90 | 0.00 | 0.00 | 0.29 | 0.06 | 4 | Monocyte |
| Prkcd | 1.01 | 0.00 | 0.00 | 0.37 | 0.10 | 4 | Monocyte |
| 4632428N05Rik | 0.87 | 0.00 | 0.00 | 0.27 | 0.05 | 4 | Monocyte |
| Rgs2 | 0.83 | 0.00 | 0.00 | 0.64 | 0.26 | 4 | Monocyte |
| Unc93b1 | 0.85 | 0.00 | 0.00 | 0.29 | 0.06 | 4 | Monocyte |
| Prr13 | 1.07 | 0.00 | 0.00 | 0.40 | 0.12 | 4 | Monocyte |
| H2-D1 | 0.93 | 0.00 | 0.00 | 0.80 | 0.48 | 4 | Monocyte |
| Gm10116 | 0.85 | 0.00 | 0.00 | 0.83 | 0.61 | 4 | Monocyte |
| Emp3 | 1.06 | 0.00 | 0.00 | 0.51 | 0.20 | 4 | Monocyte |
| Arid3a | 0.96 | 0.00 | 0.00 | 0.34 | 0.09 | 4 | Monocyte |
| Ctsb | 1.03 | 0.00 | 0.00 | 0.58 | 0.26 | 4 | Monocyte |
| Cd53 | 0.81 | 0.00 | 0.00 | 0.33 | 0.08 | 4 | Monocyte |
| Smpdl3a | 0.99 | 0.00 | 0.00 | 0.34 | 0.10 | 4 | Monocyte |
| Emb | 0.95 | 0.00 | 0.00 | 0.36 | 0.10 | 4 | Monocyte |
| Sirpa | 0.80 | 0.00 | 0.00 | 0.27 | 0.06 | 4 | Monocyte |
| Ifi203 | 0.95 | 0.00 | 0.00 | 0.41 | 0.13 | 4 | Monocyte |
| Crip1 | 0.93 | 0.00 | 0.00 | 0.77 | 0.46 | 4 | Monocyte |
| Ptpn1 | 1.05 | 0.00 | 0.00 | 0.49 | 0.19 | 4 | Monocyte |
| Arpc5 | 0.83 | 0.00 | 0.00 | 0.80 | 0.55 | 4 | Monocyte |
| Coro2a | 0.85 | 0.00 | 0.00 | 0.26 | 0.06 | 4 | Monocyte |
| Cdk2ap2 | 1.09 | 0.00 | 0.00 | 0.42 | 0.14 | 4 | Monocyte |
| Ly6e | 0.73 | 0.00 | 0.00 | 0.75 | 0.41 | 4 | Monocyte |
| Cd44 | 0.88 | 0.00 | 0.00 | 0.48 | 0.19 | 4 | Monocyte |
| Gmfg | 0.84 | 0.00 | 0.00 | 0.39 | 0.13 | 4 | Monocyte |
| Ifngr1 | 0.95 | 0.00 | 0.00 | 0.52 | 0.22 | 4 | Monocyte |
| Actr3 | 0.87 | 0.00 | 0.00 | 0.61 | 0.32 | 4 | Monocyte |
| Shfm1 | 0.71 | 0.00 | 0.00 | 0.84 | 0.63 | 4 | Monocyte |
| Ftl1 | 0.81 | 0.00 | 0.00 | 0.78 | 0.56 | 4 | Monocyte |
| Itgam | 0.96 | 0.00 | 0.00 | 0.55 | 0.27 | 4 | Monocyte |
| Rac2 | 0.70 | 0.00 | 0.00 | 0.34 | 0.10 | 4 | Monocyte |
| Ctsc | 0.87 | 0.00 | 0.00 | 0.46 | 0.18 | 4 | Monocyte |
| Trim30a | 0.84 | 0.00 | 0.00 | 0.28 | 0.07 | 4 | Monocyte |
| Sh3bgrl3 | 0.81 | 0.00 | 0.00 | 0.63 | 0.33 | 4 | Monocyte |
| Ly6c2 | 1.24 | 0.00 | 0.00 | 0.38 | 0.14 | 4 | Monocyte |
| Pycard | 0.78 | 0.00 | 0.00 | 0.29 | 0.08 | 4 | Monocyte |
| Fam49b | 0.82 | 0.00 | 0.00 | 0.31 | 0.10 | 4 | Monocyte |
| Ifi30 | 0.85 | 0.00 | 0.00 | 0.31 | 0.10 | 4 | Monocyte |
| Psma7 | 0.75 | 0.00 | 0.00 | 0.76 | 0.54 | 4 | Monocyte |
| Fam46a | 0.80 | 0.00 | 0.00 | 0.29 | 0.09 | 4 | Monocyte |
| Lyn | 0.83 | 0.00 | 0.00 | 0.34 | 0.12 | 4 | Monocyte |
| Apoe | 0.65 | 0.00 | 0.00 | 0.54 | 0.27 | 4 | Monocyte |
| Zfp36l2 | 0.79 | 0.00 | 0.00 | 0.31 | 0.10 | 4 | Monocyte |
| Zeb2 | 0.76 | 0.00 | 0.00 | 0.57 | 0.30 | 4 | Monocyte |
| Lrrc58 | 0.44 | 0.00 | 0.00 | 0.98 | 0.91 | 4 | Monocyte |
| Rps27a | 0.52 | 0.00 | 0.00 | 0.88 | 0.76 | 4 | Monocyte |
| Ikbkb | 0.77 | 0.00 | 0.00 | 0.29 | 0.10 | 4 | Monocyte |
| Aprt | 0.83 | 0.00 | 0.00 | 0.38 | 0.16 | 4 | Monocyte |
| Fos | 0.68 | 0.00 | 0.00 | 0.43 | 0.18 | 4 | Monocyte |
| Cd302 | 0.81 | 0.00 | 0.00 | 0.35 | 0.14 | 4 | Monocyte |
| Rps29 | 0.42 | 0.00 | 0.00 | 0.97 | 0.90 | 4 | Monocyte |
| Ap1s2 | 0.75 | 0.00 | 0.00 | 0.31 | 0.11 | 4 | Monocyte |
| Capg | 0.72 | 0.00 | 0.00 | 0.28 | 0.10 | 4 | Monocyte |
| Akap13 | 0.79 | 0.00 | 0.00 | 0.49 | 0.26 | 4 | Monocyte |
| Calm1 | 0.41 | 0.00 | 0.00 | 0.95 | 0.86 | 4 | Monocyte |
| Sec11c | 0.77 | 0.00 | 0.00 | 0.38 | 0.17 | 4 | Monocyte |
| Atox1 | 0.78 | 0.00 | 0.00 | 0.57 | 0.36 | 4 | Monocyte |
| Mcl1 | 0.78 | 0.00 | 0.00 | 0.44 | 0.22 | 4 | Monocyte |
| S100a8 | 0.59 | 0.00 | 0.00 | 0.27 | 0.10 | 4 | Monocyte |
| Fam96a | 0.76 | 0.00 | 0.00 | 0.33 | 0.14 | 4 | Monocyte |
| Napsa | 0.29 | 0.00 | 0.00 | 0.45 | 0.19 | 4 | Monocyte |
| Psme2b | 0.76 | 0.00 | 0.00 | 0.51 | 0.29 | 4 | Monocyte |
| Jhdm1d | 0.79 | 0.00 | 0.00 | 0.34 | 0.15 | 4 | Monocyte |
| Tpm3-rs7 | 0.51 | 0.00 | 0.00 | 0.84 | 0.69 | 4 | Monocyte |
| S100a11 | 0.64 | 0.00 | 0.00 | 0.67 | 0.48 | 4 | Monocyte |
| Capza2 | 0.71 | 0.00 | 0.00 | 0.54 | 0.34 | 4 | Monocyte |
| Psmb10 | 0.67 | 0.00 | 0.00 | 0.29 | 0.12 | 4 | Monocyte |
| Actr2 | 0.71 | 0.00 | 0.00 | 0.47 | 0.27 | 4 | Monocyte |
| Rps9 | 0.33 | 0.00 | 0.00 | 0.96 | 0.91 | 4 | Monocyte |
| Pomp | 0.68 | 0.00 | 0.00 | 0.51 | 0.31 | 4 | Monocyte |
| Tln1 | 0.63 | 0.00 | 0.00 | 0.59 | 0.39 | 4 | Monocyte |
| Coro1b | 0.71 | 0.00 | 0.00 | 0.43 | 0.24 | 4 | Monocyte |
| Mrpl33 | 0.74 | 0.00 | 0.00 | 0.48 | 0.29 | 4 | Monocyte |
| Capzb | 0.63 | 0.00 | 0.00 | 0.59 | 0.41 | 4 | Monocyte |
| Clta | 0.62 | 0.00 | 0.00 | 0.61 | 0.42 | 4 | Monocyte |
| Ahnak | 0.60 | 0.00 | 0.00 | 0.63 | 0.43 | 4 | Monocyte |
| Pabpc1 | 0.35 | 0.00 | 0.00 | 0.90 | 0.80 | 4 | Monocyte |
| Eno1 | 0.65 | 0.00 | 0.00 | 0.33 | 0.15 | 4 | Monocyte |
| Pfn1 | 0.59 | 0.00 | 0.00 | 0.64 | 0.46 | 4 | Monocyte |
| Ccdc88a | 0.71 | 0.00 | 0.00 | 0.43 | 0.24 | 4 | Monocyte |
| Psme2 | 0.70 | 0.00 | 0.00 | 0.42 | 0.23 | 4 | Monocyte |
| Stk38 | 0.70 | 0.00 | 0.00 | 0.37 | 0.19 | 4 | Monocyte |
| Tpm3 | 0.43 | 0.00 | 0.00 | 0.82 | 0.69 | 4 | Monocyte |
| Srgn | 0.48 | 0.00 | 0.00 | 0.56 | 0.32 | 4 | Monocyte |
| Ctsz | 0.64 | 0.00 | 0.00 | 0.31 | 0.14 | 4 | Monocyte |
| Junb | 0.62 | 0.00 | 0.00 | 0.38 | 0.19 | 4 | Monocyte |
| Serp1 | 0.66 | 0.00 | 0.00 | 0.40 | 0.22 | 4 | Monocyte |
| Rap1b | 0.69 | 0.00 | 0.00 | 0.40 | 0.22 | 4 | Monocyte |
| Wdr89 | 0.30 | 0.00 | 0.00 | 0.95 | 0.89 | 4 | Monocyte |
| mt-Rnr2 | 0.31 | 0.00 | 0.00 | 1.00 | 0.99 | 4 | Monocyte |
| Cd97 | 0.60 | 0.00 | 0.00 | 0.35 | 0.17 | 4 | Monocyte |
| Ctsa | 0.64 | 0.00 | 0.00 | 0.27 | 0.12 | 4 | Monocyte |
| H3f3a | 0.46 | 0.00 | 0.00 | 0.72 | 0.59 | 4 | Monocyte |
| Klf4 | 0.54 | 0.00 | 0.00 | 0.42 | 0.23 | 4 | Monocyte |
| Lrp1 | 0.55 | 0.00 | 0.00 | 0.26 | 0.11 | 4 | Monocyte |
| Anxa1 | 0.45 | 0.00 | 0.00 | 0.36 | 0.18 | 4 | Monocyte |
| Cfl1 | 0.47 | 0.00 | 0.00 | 0.71 | 0.57 | 4 | Monocyte |
| Psme1 | 0.60 | 0.00 | 0.00 | 0.48 | 0.31 | 4 | Monocyte |
| Gm5506 | 0.59 | 0.00 | 0.00 | 0.29 | 0.14 | 4 | Monocyte |
| H2-K1 | 0.52 | 0.00 | 0.00 | 0.51 | 0.32 | 4 | Monocyte |
| Gm7676 | 0.57 | 0.00 | 0.00 | 0.35 | 0.18 | 4 | Monocyte |
| Creg1 | 0.67 | 0.00 | 0.00 | 0.32 | 0.17 | 4 | Monocyte |
| Arpc4 | 0.63 | 0.00 | 0.00 | 0.33 | 0.17 | 4 | Monocyte |
| Btg1 | 0.64 | 0.00 | 0.00 | 0.45 | 0.28 | 4 | Monocyte |
| Actg1 | 0.38 | 0.00 | 0.00 | 0.85 | 0.75 | 4 | Monocyte |
| Taok3 | 0.62 | 0.00 | 0.00 | 0.34 | 0.18 | 4 | Monocyte |
| Gnai2 | 0.50 | 0.00 | 0.00 | 0.55 | 0.38 | 4 | Monocyte |
| Dusp6 | 0.64 | 0.00 | 0.00 | 0.33 | 0.18 | 4 | Monocyte |
| Ostf1 | 0.55 | 0.00 | 0.00 | 0.41 | 0.25 | 4 | Monocyte |
| Rap1a | 0.59 | 0.00 | 0.00 | 0.46 | 0.30 | 4 | Monocyte |
| Fth1 | 0.35 | 0.00 | 0.00 | 0.89 | 0.81 | 4 | Monocyte |
| Ctage5 | 0.62 | 0.00 | 0.00 | 0.43 | 0.28 | 4 | Monocyte |
| Rpsa | 0.39 | 0.00 | 0.00 | 0.73 | 0.61 | 4 | Monocyte |
| Pkm | 0.58 | 0.00 | 0.00 | 0.37 | 0.22 | 4 | Monocyte |
| Neat1 | 0.45 | 0.00 | 0.00 | 0.44 | 0.26 | 4 | Monocyte |
| Zfp36 | 0.51 | 0.00 | 0.00 | 0.31 | 0.16 | 4 | Monocyte |
| Rpsa-ps10 | 0.38 | 0.00 | 0.00 | 0.77 | 0.67 | 4 | Monocyte |
| Sppl2a | 0.58 | 0.00 | 0.00 | 0.27 | 0.13 | 4 | Monocyte |
| Tkt | 0.56 | 0.00 | 0.00 | 0.27 | 0.14 | 4 | Monocyte |
| Msn | 0.47 | 0.00 | 0.00 | 0.59 | 0.43 | 4 | Monocyte |
| Arhgap17 | 0.58 | 0.00 | 0.00 | 0.31 | 0.17 | 4 | Monocyte |
| mt-Rnr1 | 0.50 | 0.00 | 0.00 | 0.60 | 0.46 | 4 | Monocyte |
| Rplp2 | 0.28 | 0.00 | 0.00 | 0.89 | 0.81 | 4 | Monocyte |
| Mgst1 | 0.52 | 0.00 | 0.00 | 0.34 | 0.19 | 4 | Monocyte |
| Erp29 | 0.53 | 0.00 | 0.00 | 0.40 | 0.25 | 4 | Monocyte |
| Rps11 | 0.30 | 0.00 | 0.00 | 0.83 | 0.76 | 4 | Monocyte |
| Fam32a | 0.60 | 0.00 | 0.00 | 0.37 | 0.24 | 4 | Monocyte |
| Plin2 | 0.50 | 0.00 | 0.00 | 0.25 | 0.13 | 4 | Monocyte |
| Nmt1 | 0.58 | 0.00 | 0.00 | 0.36 | 0.22 | 4 | Monocyte |
| Iqgap1 | 0.43 | 0.00 | 0.00 | 0.59 | 0.44 | 4 | Monocyte |
| Diap1 | 0.57 | 0.00 | 0.00 | 0.26 | 0.14 | 4 | Monocyte |
| Ccdc12 | 0.57 | 0.00 | 0.00 | 0.31 | 0.18 | 4 | Monocyte |
| Arpc3 | 0.50 | 0.00 | 0.00 | 0.40 | 0.26 | 4 | Monocyte |
| Anxa2 | 0.44 | 0.00 | 0.00 | 0.60 | 0.47 | 4 | Monocyte |
| Sub1 | 0.42 | 0.00 | 0.00 | 0.62 | 0.51 | 4 | Monocyte |
| Mycbp2 | 0.53 | 0.00 | 0.00 | 0.37 | 0.23 | 4 | Monocyte |
| Slk | 0.43 | 0.00 | 0.00 | 0.57 | 0.43 | 4 | Monocyte |
| Serf2 | 0.41 | 0.00 | 0.00 | 0.66 | 0.56 | 4 | Monocyte |
| Tspo | 0.56 | 0.00 | 0.00 | 0.38 | 0.25 | 4 | Monocyte |
| Rps10-ps1 | 0.27 | 0.00 | 0.00 | 0.84 | 0.77 | 4 | Monocyte |
| Llph | 0.50 | 0.00 | 0.00 | 0.46 | 0.33 | 4 | Monocyte |
| Cap1 | 0.52 | 0.00 | 0.00 | 0.29 | 0.16 | 4 | Monocyte |
| Rpl35a | 0.26 | 0.00 | 0.00 | 0.86 | 0.79 | 4 | Monocyte |
| Tor1aip1 | 0.53 | 0.00 | 0.00 | 0.31 | 0.18 | 4 | Monocyte |
| Flna | 0.47 | 0.00 | 0.00 | 0.36 | 0.23 | 4 | Monocyte |
| Gm22751 | 0.54 | 0.00 | 0.00 | 0.27 | 0.16 | 4 | Monocyte |
| Ptbp3 | 0.47 | 0.00 | 0.00 | 0.48 | 0.36 | 4 | Monocyte |
| Arid4a | 0.53 | 0.00 | 0.00 | 0.41 | 0.29 | 4 | Monocyte |
| Clic1 | 0.41 | 0.00 | 0.00 | 0.50 | 0.37 | 4 | Monocyte |
| Klf13 | 0.49 | 0.00 | 0.00 | 0.28 | 0.17 | 4 | Monocyte |
| Mir692-2a | 0.55 | 0.00 | 0.00 | 0.31 | 0.19 | 4 | Monocyte |
| Lrrfip1 | 0.49 | 0.00 | 0.00 | 0.37 | 0.25 | 4 | Monocyte |
| 2810474O19Rik | 0.49 | 0.00 | 0.00 | 0.38 | 0.26 | 4 | Monocyte |
| Rpl22 | 0.25 | 0.00 | 0.00 | 0.85 | 0.79 | 4 | Monocyte |
| Glud1 | 0.49 | 0.00 | 0.00 | 0.28 | 0.17 | 4 | Monocyte |
| Reep5 | 0.48 | 0.00 | 0.00 | 0.35 | 0.24 | 4 | Monocyte |
| Atp5e | 0.38 | 0.00 | 0.00 | 0.60 | 0.50 | 4 | Monocyte |
| H2afy | 0.50 | 0.00 | 0.00 | 0.30 | 0.19 | 4 | Monocyte |
| Rps10 | 0.25 | 0.00 | 0.00 | 0.77 | 0.70 | 4 | Monocyte |
| Hspa8 | 0.33 | 0.00 | 0.00 | 0.73 | 0.67 | 4 | Monocyte |
| Rpl36al | 0.36 | 0.00 | 0.00 | 0.61 | 0.52 | 4 | Monocyte |
| Gm22774 | 0.51 | 0.00 | 0.00 | 0.27 | 0.17 | 4 | Monocyte |
| Tpr | 0.40 | 0.00 | 0.00 | 0.58 | 0.49 | 4 | Monocyte |
| Celf2 | 0.40 | 0.00 | 0.00 | 0.39 | 0.27 | 4 | Monocyte |
| Anp32a | 0.37 | 0.00 | 0.00 | 0.68 | 0.60 | 4 | Monocyte |
| Grb2 | 0.46 | 0.00 | 0.00 | 0.28 | 0.18 | 4 | Monocyte |
| Sec61b | 0.38 | 0.00 | 0.00 | 0.54 | 0.46 | 4 | Monocyte |
| Cdc42se1 | 0.48 | 0.00 | 0.00 | 0.29 | 0.19 | 4 | Monocyte |
| Cox4i1 | 0.26 | 0.00 | 0.00 | 0.78 | 0.73 | 4 | Monocyte |
| Nupr1 | 0.30 | 0.00 | 0.00 | 0.29 | 0.18 | 4 | Monocyte |
| Cdc42 | 0.34 | 0.00 | 0.00 | 0.62 | 0.54 | 4 | Monocyte |
| Ptp4a2 | 0.38 | 0.00 | 0.00 | 0.52 | 0.43 | 4 | Monocyte |
| Cd47 | 0.34 | 0.00 | 0.00 | 0.52 | 0.43 | 4 | Monocyte |
| Tsc22d4 | 0.44 | 0.00 | 0.00 | 0.37 | 0.27 | 4 | Monocyte |
| Klf6 | 0.39 | 0.00 | 0.00 | 0.46 | 0.36 | 4 | Monocyte |
| Ywhaz | 0.38 | 0.00 | 0.00 | 0.57 | 0.51 | 4 | Monocyte |
| Tmed5 | 0.45 | 0.00 | 0.00 | 0.26 | 0.17 | 4 | Monocyte |
| Dbi | 0.38 | 0.00 | 0.00 | 0.46 | 0.37 | 4 | Monocyte |
| Add3 | 0.42 | 0.00 | 0.00 | 0.37 | 0.28 | 4 | Monocyte |
| Gm26924 | 0.44 | 0.00 | 0.00 | 0.63 | 0.57 | 4 | Monocyte |
| Atp6v0b | 0.40 | 0.00 | 0.00 | 0.28 | 0.19 | 4 | Monocyte |
| Cmip | 0.43 | 0.00 | 0.00 | 0.29 | 0.20 | 4 | Monocyte |
| Slc25a5 | 0.29 | 0.00 | 0.00 | 0.66 | 0.59 | 4 | Monocyte |
| Irf2 | 0.43 | 0.00 | 0.00 | 0.33 | 0.24 | 4 | Monocyte |
| Abracl | 0.39 | 0.00 | 0.00 | 0.32 | 0.23 | 4 | Monocyte |
| Pak2 | 0.39 | 0.00 | 0.00 | 0.37 | 0.29 | 4 | Monocyte |
| Gm10320 | 0.34 | 0.00 | 0.00 | 0.46 | 0.39 | 4 | Monocyte |
| Atp5h | 0.32 | 0.00 | 0.00 | 0.53 | 0.47 | 4 | Monocyte |
| Atp6v0c | 0.38 | 0.00 | 0.00 | 0.33 | 0.25 | 4 | Monocyte |
| Atp5l | 0.35 | 0.00 | 0.00 | 0.50 | 0.44 | 4 | Monocyte |
| Gm10221 | 0.34 | 0.00 | 0.00 | 0.51 | 0.46 | 4 | Monocyte |
| Atp6v1e1 | 0.38 | 0.00 | 0.00 | 0.33 | 0.25 | 4 | Monocyte |
| Wdr26 | 0.39 | 0.00 | 0.00 | 0.34 | 0.26 | 4 | Monocyte |
| Brk1 | 0.35 | 0.00 | 0.00 | 0.34 | 0.26 | 4 | Monocyte |
| Cmpk1 | 0.40 | 0.00 | 0.00 | 0.28 | 0.21 | 4 | Monocyte |
| Lmo4 | 0.37 | 0.00 | 0.00 | 0.33 | 0.26 | 4 | Monocyte |
| Gabarap | 0.29 | 0.00 | 0.00 | 0.50 | 0.44 | 4 | Monocyte |
| Prpf40a | 0.37 | 0.00 | 0.00 | 0.40 | 0.32 | 4 | Monocyte |
| Ubl5 | 0.33 | 0.00 | 0.00 | 0.39 | 0.32 | 4 | Monocyte |
| Nsa2 | 0.38 | 0.00 | 0.00 | 0.37 | 0.30 | 4 | Monocyte |
| Zbtb7a | 0.36 | 0.00 | 0.00 | 0.27 | 0.20 | 4 | Monocyte |
| Atp6v0e | 0.37 | 0.00 | 0.00 | 0.27 | 0.20 | 4 | Monocyte |
| Cox6b1 | 0.26 | 0.00 | 0.00 | 0.61 | 0.57 | 4 | Monocyte |
| 2010107E04Rik | 0.31 | 0.00 | 0.00 | 0.47 | 0.40 | 4 | Monocyte |
| Nfe2l2 | 0.41 | 0.00 | 0.00 | 0.32 | 0.25 | 4 | Monocyte |
| S100a10 | 0.30 | 0.00 | 0.00 | 0.45 | 0.38 | 4 | Monocyte |
| BC005537 | 0.37 | 0.00 | 0.00 | 0.40 | 0.33 | 4 | Monocyte |
| Aldoa | 0.36 | 0.00 | 0.00 | 0.26 | 0.19 | 4 | Monocyte |
| Dazap2 | 0.33 | 0.00 | 0.00 | 0.27 | 0.21 | 4 | Monocyte |
| Mbnl1 | 0.33 | 0.00 | 0.00 | 0.33 | 0.26 | 4 | Monocyte |
| Gm10250 | 0.29 | 0.00 | 0.00 | 0.55 | 0.51 | 4 | Monocyte |
| Cox7b | 0.30 | 0.00 | 0.00 | 0.51 | 0.47 | 4 | Monocyte |
| Vamp8 | 0.25 | 0.00 | 0.00 | 0.42 | 0.34 | 4 | Monocyte |
| Tab2 | 0.37 | 0.00 | 0.00 | 0.29 | 0.23 | 4 | Monocyte |
| Ifitm2 | 0.27 | 0.00 | 0.00 | 0.39 | 0.32 | 4 | Monocyte |
| Eif3k | 0.31 | 0.00 | 0.00 | 0.39 | 0.34 | 4 | Monocyte |
| Tmem14c | 0.33 | 0.00 | 0.00 | 0.27 | 0.21 | 4 | Monocyte |
| D8Ertd738e | 0.33 | 0.00 | 0.00 | 0.33 | 0.27 | 4 | Monocyte |
| Gm17087 | 0.33 | 0.00 | 0.00 | 0.30 | 0.24 | 4 | Monocyte |
| Tmem50a | 0.25 | 0.00 | 0.00 | 0.49 | 0.45 | 4 | Monocyte |
| Magt1 | 0.35 | 0.00 | 0.00 | 0.25 | 0.20 | 4 | Monocyte |
| Myeov2 | 0.29 | 0.00 | 0.00 | 0.35 | 0.30 | 4 | Monocyte |
| Tma7 | 0.27 | 0.00 | 0.00 | 0.36 | 0.32 | 4 | Monocyte |
| Uqcrq | 0.28 | 0.00 | 0.00 | 0.38 | 0.34 | 4 | Monocyte |
| Ndufb7 | 0.29 | 0.00 | 0.00 | 0.33 | 0.28 | 4 | Monocyte |
| Degs1 | 0.33 | 0.00 | 0.00 | 0.28 | 0.23 | 4 | Monocyte |
| M6pr | 0.29 | 0.00 | 0.00 | 0.28 | 0.23 | 4 | Monocyte |
| Cox5a | 0.27 | 0.00 | 0.00 | 0.36 | 0.31 | 4 | Monocyte |
| Fis1 | 0.28 | 0.00 | 0.00 | 0.33 | 0.29 | 4 | Monocyte |
| Chmp2a | 0.26 | 0.00 | 0.00 | 0.35 | 0.31 | 4 | Monocyte |
| Abi1 | 0.28 | 0.00 | 0.00 | 0.27 | 0.22 | 4 | Monocyte |
| Atp6v1f | 0.27 | 0.00 | 0.00 | 0.29 | 0.25 | 4 | Monocyte |
| Psmb3 | 0.27 | 0.00 | 0.00 | 0.34 | 0.31 | 4 | Monocyte |
| Ccdc50 | 0.26 | 0.00 | 0.00 | 0.30 | 0.26 | 4 | Monocyte |
| Smdt1 | 0.26 | 0.00 | 0.00 | 0.33 | 0.29 | 4 | Monocyte |
| Atp5l-ps1 | 0.26 | 0.00 | 0.00 | 0.31 | 0.28 | 4 | Monocyte |
| Frg1 | 0.26 | 0.00 | 0.01 | 0.29 | 0.26 | 4 | Monocyte |
| Srp9 | 0.25 | 0.00 | 0.05 | 0.31 | 0.28 | 4 | Monocyte |
| Igkc | 3.20 | 0.00 | 0.00 | 0.86 | 0.02 | 5 | B Cell |
| Ms4a1 | 3.06 | 0.00 | 0.00 | 0.85 | 0.02 | 5 | B Cell |
| Ighm | 2.86 | 0.00 | 0.00 | 0.89 | 0.03 | 5 | B Cell |
| Cd79a | 2.65 | 0.00 | 0.00 | 0.80 | 0.01 | 5 | B Cell |
| Ly6d | 2.65 | 0.00 | 0.00 | 0.79 | 0.01 | 5 | B Cell |
| Iglc2 | 2.53 | 0.00 | 0.00 | 0.59 | 0.01 | 5 | B Cell |
| Cd79b | 2.53 | 0.00 | 0.00 | 0.78 | 0.03 | 5 | B Cell |
| Cd74 | 2.32 | 0.00 | 0.00 | 0.82 | 0.10 | 5 | B Cell |
| Cd37 | 2.24 | 0.00 | 0.00 | 0.72 | 0.05 | 5 | B Cell |
| H2-Ab1 | 1.90 | 0.00 | 0.00 | 0.62 | 0.07 | 5 | B Cell |
| H2-Eb1 | 1.88 | 0.00 | 0.00 | 0.53 | 0.05 | 5 | B Cell |
| H2-Aa | 1.80 | 0.00 | 0.00 | 0.55 | 0.06 | 5 | B Cell |
| Cd52 | 1.74 | 0.00 | 0.00 | 0.86 | 0.17 | 5 | B Cell |
| Ptprcap | 1.72 | 0.00 | 0.00 | 0.58 | 0.05 | 5 | B Cell |
| Mzb1 | 1.68 | 0.00 | 0.00 | 0.43 | 0.01 | 5 | B Cell |
| Bank1 | 1.65 | 0.00 | 0.00 | 0.40 | 0.01 | 5 | B Cell |
| Vpreb3 | 1.65 | 0.00 | 0.00 | 0.28 | 0.01 | 5 | B Cell |
| Cd19 | 1.57 | 0.00 | 0.00 | 0.41 | 0.01 | 5 | B Cell |
| Plac8 | 1.56 | 0.00 | 0.00 | 0.74 | 0.14 | 5 | B Cell |
| Siglecg | 1.56 | 0.00 | 0.00 | 0.40 | 0.01 | 5 | B Cell |
| Fcrla | 1.51 | 0.00 | 0.00 | 0.39 | 0.01 | 5 | B Cell |
| Pax5 | 1.46 | 0.00 | 0.00 | 0.35 | 0.01 | 5 | B Cell |
| Sp140 | 1.37 | 0.00 | 0.00 | 0.39 | 0.06 | 5 | B Cell |
| Tnfrsf13c | 1.35 | 0.00 | 0.00 | 0.29 | 0.01 | 5 | B Cell |
| Cd2 | 1.33 | 0.00 | 0.00 | 0.36 | 0.03 | 5 | B Cell |
| Mndal | 1.31 | 0.00 | 0.00 | 0.57 | 0.14 | 5 | B Cell |
| Fam129c | 1.30 | 0.00 | 0.00 | 0.28 | 0.01 | 5 | B Cell |
| Ebf1 | 1.29 | 0.00 | 0.00 | 0.33 | 0.03 | 5 | B Cell |
| Cmah | 1.26 | 0.00 | 0.00 | 0.30 | 0.03 | 5 | B Cell |
| Ly86 | 1.25 | 0.00 | 0.00 | 0.40 | 0.05 | 5 | B Cell |
| Ifi203 | 1.24 | 0.00 | 0.00 | 0.50 | 0.12 | 5 | B Cell |
| Syk | 1.23 | 0.00 | 0.00 | 0.34 | 0.05 | 5 | B Cell |
| D17H6S56E-5 | 1.23 | 0.00 | 0.00 | 0.40 | 0.09 | 5 | B Cell |
| Arhgdib | 1.23 | 0.00 | 0.00 | 0.59 | 0.14 | 5 | B Cell |
| Rac2 | 1.20 | 0.00 | 0.00 | 0.48 | 0.09 | 5 | B Cell |
| Rnase6 | 1.18 | 0.00 | 0.00 | 0.34 | 0.04 | 5 | B Cell |
| Ralgps2 | 1.17 | 0.00 | 0.00 | 0.34 | 0.06 | 5 | B Cell |
| Ltb | 1.17 | 0.00 | 0.00 | 0.28 | 0.02 | 5 | B Cell |
| H2-DMb2 | 1.14 | 0.00 | 0.00 | 0.25 | 0.01 | 5 | B Cell |
| Ptpn6 | 1.13 | 0.00 | 0.00 | 0.37 | 0.07 | 5 | B Cell |
| Tnfrsf13b | 1.11 | 0.00 | 0.00 | 0.30 | 0.03 | 5 | B Cell |
| Cd53 | 1.08 | 0.00 | 0.00 | 0.39 | 0.08 | 5 | B Cell |
| A430104N18Rik | 1.07 | 0.00 | 0.00 | 0.32 | 0.05 | 5 | B Cell |
| Arhgap30 | 1.05 | 0.00 | 0.00 | 0.39 | 0.09 | 5 | B Cell |
| Pyhin1 | 1.04 | 0.00 | 0.00 | 0.27 | 0.04 | 5 | B Cell |
| Unc93b1 | 1.02 | 0.00 | 0.00 | 0.32 | 0.06 | 5 | B Cell |
| Lsp1 | 1.00 | 0.00 | 0.00 | 0.56 | 0.18 | 5 | B Cell |
| Rps3a1 | 0.71 | 0.00 | 0.00 | 0.97 | 0.85 | 5 | B Cell |
| Rps26 | 0.66 | 0.00 | 0.00 | 0.97 | 0.89 | 5 | B Cell |
| Rps9 | 0.64 | 0.00 | 0.00 | 0.98 | 0.91 | 5 | B Cell |
| Rpl13a | 0.69 | 0.00 | 0.00 | 0.96 | 0.86 | 5 | B Cell |
| Apobec3 | 1.07 | 0.00 | 0.00 | 0.32 | 0.07 | 5 | B Cell |
| Ptprc | 0.90 | 0.00 | 0.00 | 0.48 | 0.13 | 5 | B Cell |
| Gm9843 | 0.61 | 0.00 | 0.00 | 0.98 | 0.92 | 5 | B Cell |
| Cd55 | 1.12 | 0.00 | 0.00 | 0.30 | 0.07 | 5 | B Cell |
| Rpl32 | 0.68 | 0.00 | 0.00 | 0.95 | 0.84 | 5 | B Cell |
| Coro1a | 0.91 | 0.00 | 0.00 | 0.36 | 0.09 | 5 | B Cell |
| Rps3a2 | 0.67 | 0.00 | 0.00 | 0.94 | 0.82 | 5 | B Cell |
| Rps26-ps1 | 0.64 | 0.00 | 0.00 | 0.97 | 0.87 | 5 | B Cell |
| Rpl4 | 0.69 | 0.00 | 0.00 | 0.94 | 0.83 | 5 | B Cell |
| Gm9846 | 0.64 | 0.00 | 0.00 | 0.98 | 0.89 | 5 | B Cell |
| Gm10275 | 0.81 | 0.00 | 0.00 | 0.86 | 0.66 | 5 | B Cell |
| Rps11 | 0.72 | 0.00 | 0.00 | 0.90 | 0.75 | 5 | B Cell |
| Ifi30 | 1.08 | 0.00 | 0.00 | 0.36 | 0.10 | 5 | B Cell |
| Ctss | 0.26 | 0.00 | 0.00 | 0.48 | 0.14 | 5 | B Cell |
| Wdr89 | 0.59 | 0.00 | 0.00 | 0.96 | 0.89 | 5 | B Cell |
| Ly6e | 0.78 | 0.00 | 0.00 | 0.78 | 0.41 | 5 | B Cell |
| Sp110 | 0.90 | 0.00 | 0.00 | 0.28 | 0.06 | 5 | B Cell |
| Rps3 | 0.60 | 0.00 | 0.00 | 0.93 | 0.82 | 5 | B Cell |
| Slfn2 | 0.91 | 0.00 | 0.00 | 0.40 | 0.12 | 5 | B Cell |
| Rps5 | 0.66 | 0.00 | 0.00 | 0.91 | 0.77 | 5 | B Cell |
| mt-Co1 | 0.50 | 0.00 | 0.00 | 0.99 | 0.95 | 5 | B Cell |
| Rpl18a | 0.60 | 0.00 | 0.00 | 0.94 | 0.82 | 5 | B Cell |
| Rpl10a | 0.67 | 0.00 | 0.00 | 0.89 | 0.74 | 5 | B Cell |
| Rps3a3 | 0.64 | 0.00 | 0.00 | 0.91 | 0.77 | 5 | B Cell |
| Ncf1 | 0.85 | 0.00 | 0.00 | 0.26 | 0.06 | 5 | B Cell |
| Rplp1 | 0.52 | 0.00 | 0.00 | 0.97 | 0.89 | 5 | B Cell |
| Vars | 0.91 | 0.00 | 0.00 | 0.27 | 0.07 | 5 | B Cell |
| Rps10-ps1 | 0.64 | 0.00 | 0.00 | 0.90 | 0.77 | 5 | B Cell |
| 9930111J21Rik2 | 0.96 | 0.00 | 0.00 | 0.35 | 0.11 | 5 | B Cell |
| Hcls1 | 0.79 | 0.00 | 0.00 | 0.27 | 0.07 | 5 | B Cell |
| Rps15a | 0.55 | 0.00 | 0.00 | 0.92 | 0.81 | 5 | B Cell |
| Psmb8 | 0.85 | 0.00 | 0.00 | 0.45 | 0.17 | 5 | B Cell |
| Fau | 0.52 | 0.00 | 0.00 | 0.94 | 0.86 | 5 | B Cell |
| Rps16 | 0.52 | 0.00 | 0.00 | 0.94 | 0.83 | 5 | B Cell |
| Rpsa-ps10 | 0.67 | 0.00 | 0.00 | 0.83 | 0.67 | 5 | B Cell |
| Parp1 | 0.94 | 0.00 | 0.00 | 0.31 | 0.09 | 5 | B Cell |
| Gm6472 | 0.57 | 0.00 | 0.00 | 0.92 | 0.79 | 5 | B Cell |
| Rps6 | 0.54 | 0.00 | 0.00 | 0.93 | 0.82 | 5 | B Cell |
| Gm2000 | 0.54 | 0.00 | 0.00 | 0.93 | 0.84 | 5 | B Cell |
| Rpl8 | 0.60 | 0.00 | 0.00 | 0.90 | 0.79 | 5 | B Cell |
| Rps20 | 0.61 | 0.00 | 0.00 | 0.90 | 0.78 | 5 | B Cell |
| Serp1 | 0.97 | 0.00 | 0.00 | 0.48 | 0.22 | 5 | B Cell |
| Tmsb4x | 0.43 | 0.00 | 0.00 | 0.97 | 0.88 | 5 | B Cell |
| Ifi27l2a | 0.77 | 0.00 | 0.00 | 0.30 | 0.08 | 5 | B Cell |
| Rps24 | 0.49 | 0.00 | 0.00 | 0.96 | 0.88 | 5 | B Cell |
| Cd24a | 0.97 | 0.00 | 0.00 | 0.36 | 0.12 | 5 | B Cell |
| Sh3bgrl3 | 0.86 | 0.00 | 0.00 | 0.61 | 0.33 | 5 | B Cell |
| Gm10269 | 0.57 | 0.00 | 0.00 | 0.89 | 0.79 | 5 | B Cell |
| Mef2c | 1.00 | 0.00 | 0.00 | 0.41 | 0.15 | 5 | B Cell |
| Gm10073 | 0.50 | 0.00 | 0.00 | 0.94 | 0.85 | 5 | B Cell |
| Rplp0 | 0.55 | 0.00 | 0.00 | 0.93 | 0.83 | 5 | B Cell |
| Gm7536 | 0.53 | 0.00 | 0.00 | 0.92 | 0.81 | 5 | B Cell |
| Limd2 | 0.93 | 0.00 | 0.00 | 0.39 | 0.14 | 5 | B Cell |
| Gnb2l1 | 0.66 | 0.00 | 0.00 | 0.81 | 0.65 | 5 | B Cell |
| Rpl12 | 0.73 | 0.00 | 0.00 | 0.75 | 0.54 | 5 | B Cell |
| Gm9396 | 0.74 | 0.00 | 0.00 | 0.74 | 0.53 | 5 | B Cell |
| Pabpc1 | 0.55 | 0.00 | 0.00 | 0.92 | 0.80 | 5 | B Cell |
| Rps12-ps3 | 0.57 | 0.00 | 0.00 | 0.89 | 0.78 | 5 | B Cell |
| Gm8730 | 0.55 | 0.00 | 0.00 | 0.92 | 0.82 | 5 | B Cell |
| Sh3bp5 | 0.90 | 0.00 | 0.00 | 0.44 | 0.18 | 5 | B Cell |
| Rplp2 | 0.52 | 0.00 | 0.00 | 0.91 | 0.81 | 5 | B Cell |
| Rpl22 | 0.55 | 0.00 | 0.00 | 0.90 | 0.79 | 5 | B Cell |
| Sp100 | 0.78 | 0.00 | 0.00 | 0.27 | 0.08 | 5 | B Cell |
| Rpl14 | 0.53 | 0.00 | 0.00 | 0.92 | 0.83 | 5 | B Cell |
| Gimap4 | 0.83 | 0.00 | 0.00 | 0.42 | 0.17 | 5 | B Cell |
| Gimap6 | 0.84 | 0.00 | 0.00 | 0.40 | 0.15 | 5 | B Cell |
| Rps10 | 0.59 | 0.00 | 0.00 | 0.84 | 0.70 | 5 | B Cell |
| Actb | 0.37 | 0.00 | 0.00 | 1.00 | 0.98 | 5 | B Cell |
| mt-Co3 | 0.47 | 0.00 | 0.00 | 0.95 | 0.88 | 5 | B Cell |
| Snx5 | 0.88 | 0.00 | 0.00 | 0.34 | 0.12 | 5 | B Cell |
| Rpl6 | 0.46 | 0.00 | 0.00 | 0.95 | 0.87 | 5 | B Cell |
| Sub1 | 0.94 | 0.00 | 0.00 | 0.69 | 0.50 | 5 | B Cell |
| Ptpn18 | 0.77 | 0.00 | 0.00 | 0.36 | 0.13 | 5 | B Cell |
| Rpl5 | 0.56 | 0.00 | 0.00 | 0.87 | 0.74 | 5 | B Cell |
| Rps27 | 0.55 | 0.00 | 0.00 | 0.93 | 0.81 | 5 | B Cell |
| Shisa5 | 0.89 | 0.00 | 0.00 | 0.43 | 0.20 | 5 | B Cell |
| Rps18 | 0.61 | 0.00 | 0.00 | 0.80 | 0.65 | 5 | B Cell |
| Rps19 | 0.46 | 0.00 | 0.00 | 0.92 | 0.82 | 5 | B Cell |
| Rps7 | 0.53 | 0.00 | 0.00 | 0.87 | 0.73 | 5 | B Cell |
| H2-K1 | 0.70 | 0.00 | 0.00 | 0.58 | 0.31 | 5 | B Cell |
| Rps25 | 0.49 | 0.00 | 0.00 | 0.90 | 0.80 | 5 | B Cell |
| H2-D1 | 0.56 | 0.00 | 0.00 | 0.75 | 0.48 | 5 | B Cell |
| Rpl35 | 0.48 | 0.00 | 0.00 | 0.89 | 0.80 | 5 | B Cell |
| Rps4x | 0.41 | 0.00 | 0.00 | 0.95 | 0.86 | 5 | B Cell |
| Snx2 | 0.88 | 0.00 | 0.00 | 0.37 | 0.16 | 5 | B Cell |
| Gimap1 | 0.72 | 0.00 | 0.00 | 0.34 | 0.13 | 5 | B Cell |
| Il2rg | 0.74 | 0.00 | 0.00 | 0.27 | 0.09 | 5 | B Cell |
| Crip1 | 0.86 | 0.00 | 0.00 | 0.70 | 0.47 | 5 | B Cell |
| Rpl13 | 0.48 | 0.00 | 0.00 | 0.88 | 0.79 | 5 | B Cell |
| Fam111a | 0.83 | 0.00 | 0.00 | 0.36 | 0.15 | 5 | B Cell |
| Npm1 | 0.70 | 0.00 | 0.00 | 0.73 | 0.57 | 5 | B Cell |
| Eef1b2 | 0.57 | 0.00 | 0.00 | 0.79 | 0.64 | 5 | B Cell |
| Lrrc58 | 0.38 | 0.00 | 0.00 | 0.98 | 0.91 | 5 | B Cell |
| Gm9493 | 0.50 | 0.00 | 0.00 | 0.87 | 0.74 | 5 | B Cell |
| Rpl39 | 0.46 | 0.00 | 0.00 | 0.90 | 0.80 | 5 | B Cell |
| Itga4 | 0.69 | 0.00 | 0.00 | 0.27 | 0.09 | 5 | B Cell |
| mt-Co2 | 0.48 | 0.00 | 0.00 | 0.87 | 0.77 | 5 | B Cell |
| Rpl27a | 0.46 | 0.00 | 0.00 | 0.88 | 0.78 | 5 | B Cell |
| Nol7 | 0.77 | 0.00 | 0.00 | 0.56 | 0.35 | 5 | B Cell |
| Gm10288 | 0.47 | 0.00 | 0.00 | 0.88 | 0.78 | 5 | B Cell |
| Gmfg | 0.76 | 0.00 | 0.00 | 0.32 | 0.13 | 5 | B Cell |
| Rps13 | 0.56 | 0.00 | 0.00 | 0.77 | 0.64 | 5 | B Cell |
| Eef1g | 0.69 | 0.00 | 0.00 | 0.65 | 0.48 | 5 | B Cell |
| Rpl19 | 0.46 | 0.00 | 0.00 | 0.85 | 0.75 | 5 | B Cell |
| Lcp1 | 0.68 | 0.00 | 0.00 | 0.44 | 0.22 | 5 | B Cell |
| Rpsa | 0.57 | 0.00 | 0.00 | 0.75 | 0.61 | 5 | B Cell |
| Rps18-ps3 | 0.60 | 0.00 | 0.00 | 0.71 | 0.55 | 5 | B Cell |
| mt-Nd5 | 0.51 | 0.00 | 0.00 | 0.86 | 0.75 | 5 | B Cell |
| Napsa | 0.37 | 0.00 | 0.00 | 0.45 | 0.19 | 5 | B Cell |
| Ncl | 0.59 | 0.00 | 0.00 | 0.85 | 0.78 | 5 | B Cell |
| Rpl3 | 0.55 | 0.00 | 0.00 | 0.75 | 0.63 | 5 | B Cell |
| Gm10263 | 0.44 | 0.00 | 0.00 | 0.88 | 0.76 | 5 | B Cell |
| Rps28 | 0.39 | 0.00 | 0.00 | 0.94 | 0.83 | 5 | B Cell |
| Grap | 0.69 | 0.00 | 0.00 | 0.31 | 0.13 | 5 | B Cell |
| Gm10036 | 0.47 | 0.00 | 0.00 | 0.84 | 0.74 | 5 | B Cell |
| Gm5428 | 0.42 | 0.00 | 0.00 | 0.87 | 0.79 | 5 | B Cell |
| Rps15 | 0.41 | 0.00 | 0.00 | 0.87 | 0.78 | 5 | B Cell |
| H3f3a | 0.60 | 0.00 | 0.00 | 0.72 | 0.59 | 5 | B Cell |
| Rpl13-ps3 | 0.45 | 0.00 | 0.00 | 0.85 | 0.75 | 5 | B Cell |
| Arpc5l | 0.81 | 0.00 | 0.00 | 0.34 | 0.16 | 5 | B Cell |
| mt-Nd1 | 0.37 | 0.00 | 0.00 | 0.96 | 0.91 | 5 | B Cell |
| Rabgap1l | 0.75 | 0.00 | 0.00 | 0.26 | 0.10 | 5 | B Cell |
| Rpl27-ps3 | 0.44 | 0.00 | 0.00 | 0.85 | 0.76 | 5 | B Cell |
| Cyba | 0.56 | 0.00 | 0.00 | 0.51 | 0.29 | 5 | B Cell |
| Rps12 | 0.51 | 0.00 | 0.00 | 0.80 | 0.68 | 5 | B Cell |
| Rps23 | 0.38 | 0.00 | 0.00 | 0.88 | 0.79 | 5 | B Cell |
| Capg | 0.71 | 0.00 | 0.00 | 0.25 | 0.10 | 5 | B Cell |
| Gm4070 | 0.55 | 0.00 | 0.00 | 0.28 | 0.11 | 5 | B Cell |
| Rps2 | 0.49 | 0.00 | 0.00 | 0.76 | 0.67 | 5 | B Cell |
| Gm4204 | 0.61 | 0.00 | 0.00 | 0.65 | 0.50 | 5 | B Cell |
| Mycbp2 | 0.80 | 0.00 | 0.00 | 0.41 | 0.23 | 5 | B Cell |
| Tcof1 | 0.69 | 0.00 | 0.00 | 0.26 | 0.10 | 5 | B Cell |
| Rpl23 | 0.35 | 0.00 | 0.00 | 0.94 | 0.86 | 5 | B Cell |
| Bin1 | 0.69 | 0.00 | 0.00 | 0.25 | 0.10 | 5 | B Cell |
| Sec11c | 0.74 | 0.00 | 0.00 | 0.35 | 0.17 | 5 | B Cell |
| Rps27a | 0.40 | 0.00 | 0.00 | 0.85 | 0.76 | 5 | B Cell |
| mt-Atp6 | 0.37 | 0.00 | 0.00 | 0.92 | 0.84 | 5 | B Cell |
| Gm11808 | 0.40 | 0.00 | 0.00 | 0.86 | 0.77 | 5 | B Cell |
| Klf2 | 0.64 | 0.00 | 0.00 | 0.38 | 0.19 | 5 | B Cell |
| Rpl26 | 0.38 | 0.00 | 0.00 | 0.88 | 0.81 | 5 | B Cell |
| Rpl35a | 0.37 | 0.00 | 0.00 | 0.87 | 0.79 | 5 | B Cell |
| Rpl34 | 0.38 | 0.00 | 0.00 | 0.87 | 0.80 | 5 | B Cell |
| Nme1 | 0.73 | 0.00 | 0.00 | 0.35 | 0.18 | 5 | B Cell |
| Rps21 | 0.40 | 0.00 | 0.00 | 0.83 | 0.75 | 5 | B Cell |
| Nap1l1 | 0.56 | 0.00 | 0.00 | 0.62 | 0.47 | 5 | B Cell |
| Ezr | 0.61 | 0.00 | 0.00 | 0.39 | 0.21 | 5 | B Cell |
| Tpm3 | 0.44 | 0.00 | 0.00 | 0.81 | 0.70 | 5 | B Cell |
| Serbp1 | 0.45 | 0.00 | 0.00 | 0.81 | 0.74 | 5 | B Cell |
| Rpl18 | 0.47 | 0.00 | 0.00 | 0.73 | 0.64 | 5 | B Cell |
| Rpl13a-ps1 | 0.52 | 0.00 | 0.00 | 0.66 | 0.55 | 5 | B Cell |
| Cotl1 | 0.62 | 0.00 | 0.00 | 0.34 | 0.17 | 5 | B Cell |
| Hsp90ab1 | 0.36 | 0.00 | 0.00 | 0.92 | 0.89 | 5 | B Cell |
| Gm10260 | 0.54 | 0.00 | 0.00 | 0.62 | 0.48 | 5 | B Cell |
| Calm1 | 0.34 | 0.00 | 0.00 | 0.92 | 0.86 | 5 | B Cell |
| Gvin1 | 0.53 | 0.00 | 0.00 | 0.26 | 0.11 | 5 | B Cell |
| Tpm3-rs7 | 0.43 | 0.00 | 0.00 | 0.81 | 0.69 | 5 | B Cell |
| Gm10335 | 0.37 | 0.00 | 0.00 | 0.82 | 0.73 | 5 | B Cell |
| Rpl11 | 0.42 | 0.00 | 0.00 | 0.77 | 0.68 | 5 | B Cell |
| Rpl41 | 0.26 | 0.00 | 0.00 | 0.98 | 0.94 | 5 | B Cell |
| Arhgap17 | 0.69 | 0.00 | 0.00 | 0.32 | 0.17 | 5 | B Cell |
| Rpl9-ps6 | 0.37 | 0.00 | 0.00 | 0.82 | 0.73 | 5 | B Cell |
| Rps14 | 0.28 | 0.00 | 0.00 | 0.95 | 0.92 | 5 | B Cell |
| Rpl7 | 0.35 | 0.00 | 0.00 | 0.82 | 0.74 | 5 | B Cell |
| Zfp706 | 0.62 | 0.00 | 0.00 | 0.47 | 0.32 | 5 | B Cell |
| Gm7589 | 0.42 | 0.00 | 0.00 | 0.73 | 0.65 | 5 | B Cell |
| Ets1 | 0.57 | 0.00 | 0.00 | 0.41 | 0.24 | 5 | B Cell |
| Rpl27 | 0.38 | 0.00 | 0.00 | 0.77 | 0.69 | 5 | B Cell |
| Itsn2 | 0.66 | 0.00 | 0.00 | 0.39 | 0.24 | 5 | B Cell |
| Crlf3 | 0.62 | 0.00 | 0.00 | 0.27 | 0.13 | 5 | B Cell |
| Gm10132 | 0.36 | 0.00 | 0.00 | 0.80 | 0.72 | 5 | B Cell |
| Rps8 | 0.31 | 0.00 | 0.00 | 0.87 | 0.81 | 5 | B Cell |
| Srgn | 0.26 | 0.00 | 0.00 | 0.53 | 0.32 | 5 | B Cell |
| Pdcd4 | 0.64 | 0.00 | 0.00 | 0.36 | 0.22 | 5 | B Cell |
| Lyn | 0.57 | 0.00 | 0.00 | 0.26 | 0.13 | 5 | B Cell |
| Rpl23a-ps3 | 0.33 | 0.00 | 0.00 | 0.82 | 0.75 | 5 | B Cell |
| Arpc5 | 0.42 | 0.00 | 0.00 | 0.67 | 0.56 | 5 | B Cell |
| Actr3 | 0.56 | 0.00 | 0.00 | 0.47 | 0.33 | 5 | B Cell |
| Zcchc11 | 0.65 | 0.00 | 0.00 | 0.34 | 0.20 | 5 | B Cell |
| Rps29 | 0.26 | 0.00 | 0.00 | 0.96 | 0.90 | 5 | B Cell |
| Gltscr2 | 0.49 | 0.00 | 0.00 | 0.59 | 0.48 | 5 | B Cell |
| Rpl37 | 0.31 | 0.00 | 0.00 | 0.85 | 0.80 | 5 | B Cell |
| Dnajc2 | 0.56 | 0.00 | 0.00 | 0.33 | 0.20 | 5 | B Cell |
| Nop58 | 0.65 | 0.00 | 0.00 | 0.39 | 0.26 | 5 | B Cell |
| Elf1 | 0.60 | 0.00 | 0.00 | 0.34 | 0.20 | 5 | B Cell |
| Pfn1 | 0.51 | 0.00 | 0.00 | 0.57 | 0.46 | 5 | B Cell |
| Rps17 | 0.28 | 0.00 | 0.00 | 0.83 | 0.77 | 5 | B Cell |
| Rpl17 | 0.40 | 0.00 | 0.00 | 0.64 | 0.53 | 5 | B Cell |
| Eef1a1 | 0.27 | 0.00 | 0.00 | 0.90 | 0.87 | 5 | B Cell |
| Erp29 | 0.58 | 0.00 | 0.00 | 0.39 | 0.26 | 5 | B Cell |
| Llph | 0.54 | 0.00 | 0.00 | 0.46 | 0.33 | 5 | B Cell |
| Gm5611 | 0.55 | 0.00 | 0.00 | 0.47 | 0.35 | 5 | B Cell |
| Arpc2 | 0.32 | 0.00 | 0.00 | 0.72 | 0.61 | 5 | B Cell |
| AC123611.1 | 0.49 | 0.00 | 0.00 | 0.50 | 0.39 | 5 | B Cell |
| Btg1 | 0.61 | 0.00 | 0.00 | 0.40 | 0.28 | 5 | B Cell |
| Mcl1 | 0.53 | 0.00 | 0.00 | 0.35 | 0.22 | 5 | B Cell |
| Nop56 | 0.61 | 0.00 | 0.00 | 0.32 | 0.20 | 5 | B Cell |
| Ddx21 | 0.58 | 0.00 | 0.00 | 0.38 | 0.26 | 5 | B Cell |
| Smim14 | 0.63 | 0.00 | 0.00 | 0.30 | 0.19 | 5 | B Cell |
| Hspe1 | 0.54 | 0.00 | 0.00 | 0.40 | 0.28 | 5 | B Cell |
| Rpl23a | 0.32 | 0.00 | 0.00 | 0.74 | 0.66 | 5 | B Cell |
| Ptbp3 | 0.47 | 0.00 | 0.00 | 0.47 | 0.36 | 5 | B Cell |
| Dnajc7 | 0.58 | 0.00 | 0.00 | 0.32 | 0.21 | 5 | B Cell |
| Rpl36al | 0.40 | 0.00 | 0.00 | 0.60 | 0.52 | 5 | B Cell |
| Gpx1 | 0.34 | 0.00 | 0.00 | 0.55 | 0.43 | 5 | B Cell |
| 4930523C07Rik | 0.56 | 0.00 | 0.00 | 0.29 | 0.18 | 5 | B Cell |
| S100a10 | 0.48 | 0.00 | 0.00 | 0.49 | 0.38 | 5 | B Cell |
| Arpc1b | 0.41 | 0.00 | 0.00 | 0.49 | 0.37 | 5 | B Cell |
| Rpl36 | 0.32 | 0.00 | 0.00 | 0.69 | 0.63 | 5 | B Cell |
| mt-Cytb | 0.26 | 0.00 | 0.00 | 0.87 | 0.82 | 5 | B Cell |
| Orc5 | 0.38 | 0.00 | 0.00 | 0.62 | 0.55 | 5 | B Cell |
| Rpl28 | 0.38 | 0.00 | 0.00 | 0.60 | 0.53 | 5 | B Cell |
| Naca | 0.36 | 0.00 | 0.00 | 0.64 | 0.58 | 5 | B Cell |
| Rpl30 | 0.36 | 0.00 | 0.00 | 0.66 | 0.58 | 5 | B Cell |
| Chd1 | 0.55 | 0.00 | 0.00 | 0.27 | 0.17 | 5 | B Cell |
| Ddx5 | 0.32 | 0.00 | 0.00 | 0.75 | 0.71 | 5 | B Cell |
| 1500012F01Rik | 0.49 | 0.00 | 0.00 | 0.40 | 0.30 | 5 | B Cell |
| mt-Nd4 | 0.27 | 0.00 | 0.00 | 0.79 | 0.75 | 5 | B Cell |
| Foxp1 | 0.35 | 0.00 | 0.00 | 0.56 | 0.44 | 5 | B Cell |
| Rpl36a | 0.30 | 0.00 | 0.00 | 0.68 | 0.62 | 5 | B Cell |
| Eml4 | 0.57 | 0.00 | 0.00 | 0.27 | 0.17 | 5 | B Cell |
| Rpl22l1 | 0.35 | 0.00 | 0.00 | 0.62 | 0.56 | 5 | B Cell |
| Mif | 0.53 | 0.00 | 0.00 | 0.28 | 0.18 | 5 | B Cell |
| Prpf38b | 0.39 | 0.00 | 0.00 | 0.58 | 0.49 | 5 | B Cell |
| Samhd1 | 0.37 | 0.00 | 0.00 | 0.30 | 0.19 | 5 | B Cell |
| Ccnd2 | 0.51 | 0.00 | 0.00 | 0.38 | 0.28 | 5 | B Cell |
| Lsm4 | 0.45 | 0.00 | 0.00 | 0.30 | 0.20 | 5 | B Cell |
| AC121131.2 | 0.43 | 0.00 | 0.00 | 0.47 | 0.40 | 5 | B Cell |
| Gm17511 | 0.31 | 0.00 | 0.00 | 0.66 | 0.61 | 5 | B Cell |
| Hnrnpa2b1 | 0.29 | 0.00 | 0.00 | 0.75 | 0.72 | 5 | B Cell |
| Msn | 0.34 | 0.00 | 0.00 | 0.52 | 0.43 | 5 | B Cell |
| Gas5 | 0.34 | 0.00 | 0.00 | 0.60 | 0.53 | 5 | B Cell |
| Pgls | 0.50 | 0.00 | 0.00 | 0.25 | 0.17 | 5 | B Cell |
| Stk38 | 0.46 | 0.00 | 0.00 | 0.29 | 0.20 | 5 | B Cell |
| Grb2 | 0.47 | 0.00 | 0.00 | 0.26 | 0.18 | 5 | B Cell |
| Set | 0.48 | 0.00 | 0.00 | 0.41 | 0.34 | 5 | B Cell |
| Gm10222 | 0.37 | 0.00 | 0.00 | 0.50 | 0.43 | 5 | B Cell |
| Gm7808 | 0.28 | 0.00 | 0.00 | 0.66 | 0.60 | 5 | B Cell |
| Rbm3 | 0.38 | 0.00 | 0.00 | 0.48 | 0.41 | 5 | B Cell |
| Yeats4 | 0.47 | 0.00 | 0.00 | 0.27 | 0.19 | 5 | B Cell |
| Akap13 | 0.46 | 0.00 | 0.00 | 0.35 | 0.27 | 5 | B Cell |
| Cdc42se1 | 0.43 | 0.00 | 0.00 | 0.28 | 0.19 | 5 | B Cell |
| Zfp106 | 0.43 | 0.00 | 0.00 | 0.28 | 0.20 | 5 | B Cell |
| Xrn2 | 0.44 | 0.00 | 0.00 | 0.37 | 0.30 | 5 | B Cell |
| Rpl9 | 0.26 | 0.00 | 0.00 | 0.66 | 0.63 | 5 | B Cell |
| Gm5093 | 0.33 | 0.00 | 0.00 | 0.52 | 0.47 | 5 | B Cell |
| Ppdpf | 0.44 | 0.00 | 0.00 | 0.26 | 0.18 | 5 | B Cell |
| Ptpn1 | 0.40 | 0.00 | 0.00 | 0.28 | 0.20 | 5 | B Cell |
| Cep110 | 0.42 | 0.00 | 0.00 | 0.26 | 0.19 | 5 | B Cell |
| Oaz1 | 0.28 | 0.00 | 0.00 | 0.64 | 0.62 | 5 | B Cell |
| mt-Nd4l | 0.39 | 0.00 | 0.00 | 0.45 | 0.38 | 5 | B Cell |
| Clta | 0.32 | 0.00 | 0.00 | 0.49 | 0.43 | 5 | B Cell |
| Ivns1abp | 0.35 | 0.00 | 0.00 | 0.30 | 0.22 | 5 | B Cell |
| Eif5a | 0.39 | 0.00 | 0.00 | 0.41 | 0.34 | 5 | B Cell |
| Sf3b1 | 0.33 | 0.00 | 0.00 | 0.52 | 0.46 | 5 | B Cell |
| Emp3 | 0.36 | 0.00 | 0.00 | 0.29 | 0.21 | 5 | B Cell |
| Srsf3 | 0.30 | 0.00 | 0.00 | 0.50 | 0.45 | 5 | B Cell |
| Rpl36-ps3 | 0.26 | 0.00 | 0.00 | 0.58 | 0.53 | 5 | B Cell |
| Gdi2 | 0.40 | 0.00 | 0.00 | 0.39 | 0.33 | 5 | B Cell |
| Gm10020 | 0.28 | 0.00 | 0.00 | 0.56 | 0.52 | 5 | B Cell |
| Anp32b | 0.28 | 0.00 | 0.00 | 0.54 | 0.50 | 5 | B Cell |
| Psme2b | 0.36 | 0.00 | 0.00 | 0.36 | 0.30 | 5 | B Cell |
| Psme1 | 0.36 | 0.00 | 0.00 | 0.37 | 0.31 | 5 | B Cell |
| mt-Nd3 | 0.28 | 0.00 | 0.00 | 0.51 | 0.48 | 5 | B Cell |
| Taf1d | 0.40 | 0.00 | 0.00 | 0.29 | 0.23 | 5 | B Cell |
| Hnrnpc | 0.32 | 0.00 | 0.00 | 0.48 | 0.44 | 5 | B Cell |
| Rps2-ps10 | 0.31 | 0.00 | 0.00 | 0.48 | 0.43 | 5 | B Cell |
| Gm5239 | 0.26 | 0.00 | 0.00 | 0.55 | 0.52 | 5 | B Cell |
| Rrp1 | 0.34 | 0.00 | 0.00 | 0.42 | 0.37 | 5 | B Cell |
| Mbnl1 | 0.32 | 0.00 | 0.00 | 0.33 | 0.27 | 5 | B Cell |
| Gm6139 | 0.31 | 0.00 | 0.00 | 0.46 | 0.42 | 5 | B Cell |
| Gtf2f1 | 0.40 | 0.00 | 0.00 | 0.27 | 0.21 | 5 | B Cell |
| Rsl1d1 | 0.39 | 0.00 | 0.00 | 0.33 | 0.27 | 5 | B Cell |
| Eif5 | 0.36 | 0.00 | 0.00 | 0.41 | 0.37 | 5 | B Cell |
| Pa2g4 | 0.42 | 0.00 | 0.00 | 0.31 | 0.26 | 5 | B Cell |
| Fkbp3 | 0.38 | 0.00 | 0.00 | 0.35 | 0.30 | 5 | B Cell |
| Snrpf | 0.36 | 0.00 | 0.00 | 0.33 | 0.28 | 5 | B Cell |
| Polr1d | 0.34 | 0.00 | 0.00 | 0.33 | 0.28 | 5 | B Cell |
| Pdia4 | 0.36 | 0.00 | 0.00 | 0.27 | 0.21 | 5 | B Cell |
| Cnbp | 0.30 | 0.00 | 0.00 | 0.48 | 0.45 | 5 | B Cell |
| Rps2-ps6 | 0.30 | 0.00 | 0.00 | 0.44 | 0.41 | 5 | B Cell |
| Snrpe | 0.32 | 0.00 | 0.00 | 0.40 | 0.36 | 5 | B Cell |
| Hspd1 | 0.37 | 0.00 | 0.00 | 0.31 | 0.25 | 5 | B Cell |
| Ranbp1 | 0.37 | 0.00 | 0.00 | 0.34 | 0.29 | 5 | B Cell |
| AC167036.1 | 0.33 | 0.00 | 0.00 | 0.35 | 0.30 | 5 | B Cell |
| Ran | 0.29 | 0.00 | 0.00 | 0.42 | 0.38 | 5 | B Cell |
| Safb2 | 0.37 | 0.00 | 0.00 | 0.29 | 0.24 | 5 | B Cell |
| Tmod3 | 0.37 | 0.00 | 0.00 | 0.31 | 0.26 | 5 | B Cell |
| Hnrnpm | 0.27 | 0.00 | 0.00 | 0.52 | 0.49 | 5 | B Cell |
| Hnrnpf | 0.30 | 0.00 | 0.00 | 0.38 | 0.34 | 5 | B Cell |
| Psmb1 | 0.28 | 0.00 | 0.00 | 0.48 | 0.47 | 5 | B Cell |
| Rpl10a-ps2 | 0.30 | 0.00 | 0.00 | 0.33 | 0.29 | 5 | B Cell |
| Eif3c | 0.25 | 0.00 | 0.00 | 0.50 | 0.48 | 5 | B Cell |
| Ddx46 | 0.29 | 0.00 | 0.00 | 0.34 | 0.30 | 5 | B Cell |
| Atp5a1 | 0.27 | 0.00 | 0.00 | 0.44 | 0.41 | 5 | B Cell |
| Fgfr1op2 | 0.30 | 0.00 | 0.00 | 0.32 | 0.28 | 5 | B Cell |
| Nop10 | 0.33 | 0.00 | 0.00 | 0.32 | 0.28 | 5 | B Cell |
| Eif3e | 0.27 | 0.00 | 0.00 | 0.43 | 0.40 | 5 | B Cell |
| Kmt2e | 0.28 | 0.00 | 0.00 | 0.45 | 0.43 | 5 | B Cell |
| Irf2 | 0.34 | 0.00 | 0.00 | 0.28 | 0.24 | 5 | B Cell |
| Psma1 | 0.28 | 0.00 | 0.00 | 0.30 | 0.26 | 5 | B Cell |
| Eif3k | 0.30 | 0.00 | 0.00 | 0.37 | 0.34 | 5 | B Cell |
| Cct2 | 0.30 | 0.00 | 0.00 | 0.30 | 0.27 | 5 | B Cell |
| Gm15013 | 0.26 | 0.00 | 0.00 | 0.39 | 0.37 | 5 | B Cell |
| mt-Atp8 | 0.31 | 0.00 | 0.00 | 0.31 | 0.28 | 5 | B Cell |
| Gm10126 | 0.25 | 0.00 | 0.01 | 0.42 | 0.40 | 5 | B Cell |
| Cct5 | 0.32 | 0.00 | 0.01 | 0.27 | 0.23 | 5 | B Cell |
| Add3 | 0.26 | 0.00 | 0.01 | 0.32 | 0.28 | 5 | B Cell |
| Eef1d | 0.29 | 0.00 | 0.01 | 0.29 | 0.26 | 5 | B Cell |
| Gm17087 | 0.28 | 0.00 | 0.03 | 0.28 | 0.24 | 5 | B Cell |
| Ms4a4b | 2.91 | 0.00 | 0.00 | 0.79 | 0.02 | 6 | T Cell |
| Trbc2 | 2.66 | 0.00 | 0.00 | 0.78 | 0.02 | 6 | T Cell |
| Trbc1 | 2.40 | 0.00 | 0.00 | 0.67 | 0.01 | 6 | T Cell |
| Cd3d | 2.08 | 0.00 | 0.00 | 0.59 | 0.01 | 6 | T Cell |
| Cd3g | 2.07 | 0.00 | 0.00 | 0.62 | 0.01 | 6 | T Cell |
| Ms4a6b | 1.92 | 0.00 | 0.00 | 0.56 | 0.04 | 6 | T Cell |
| Il7r | 1.70 | 0.00 | 0.00 | 0.37 | 0.01 | 6 | T Cell |
| Ptprc | 1.68 | 0.00 | 0.00 | 0.72 | 0.12 | 6 | T Cell |
| Tcf7 | 1.67 | 0.00 | 0.00 | 0.46 | 0.05 | 6 | T Cell |
| Cd52 | 1.63 | 0.00 | 0.00 | 0.86 | 0.17 | 6 | T Cell |
| Ptprcap | 1.61 | 0.00 | 0.00 | 0.58 | 0.05 | 6 | T Cell |
| Nkg7 | 1.59 | 0.00 | 0.00 | 0.30 | 0.01 | 6 | T Cell |
| AW112010 | 1.55 | 0.00 | 0.00 | 0.31 | 0.03 | 6 | T Cell |
| Lck | 1.51 | 0.00 | 0.00 | 0.40 | 0.01 | 6 | T Cell |
| Rac2 | 1.51 | 0.00 | 0.00 | 0.61 | 0.09 | 6 | T Cell |
| Shisa5 | 1.41 | 0.00 | 0.00 | 0.64 | 0.19 | 6 | T Cell |
| Ccr7 | 1.41 | 0.00 | 0.00 | 0.30 | 0.02 | 6 | T Cell |
| H2-K1 | 1.39 | 0.00 | 0.00 | 0.81 | 0.30 | 6 | T Cell |
| Thy1 | 1.37 | 0.00 | 0.00 | 0.34 | 0.01 | 6 | T Cell |
| Selplg | 1.36 | 0.00 | 0.00 | 0.42 | 0.04 | 6 | T Cell |
| Lef1 | 1.32 | 0.00 | 0.00 | 0.32 | 0.03 | 6 | T Cell |
| Hcst | 1.32 | 0.00 | 0.00 | 0.36 | 0.03 | 6 | T Cell |
| Arhgdib | 1.32 | 0.00 | 0.00 | 0.65 | 0.14 | 6 | T Cell |
| Cd2 | 1.28 | 0.00 | 0.00 | 0.37 | 0.03 | 6 | T Cell |
| Limd2 | 1.27 | 0.00 | 0.00 | 0.54 | 0.14 | 6 | T Cell |
| Gimap4 | 1.24 | 0.00 | 0.00 | 0.55 | 0.16 | 6 | T Cell |
| Sept1 | 1.23 | 0.00 | 0.00 | 0.36 | 0.03 | 6 | T Cell |
| Cd27 | 1.23 | 0.00 | 0.00 | 0.30 | 0.01 | 6 | T Cell |
| Laptm5 | 1.20 | 0.00 | 0.00 | 0.60 | 0.12 | 6 | T Cell |
| Bcl11b | 1.19 | 0.00 | 0.00 | 0.28 | 0.01 | 6 | T Cell |
| Cd53 | 1.19 | 0.00 | 0.00 | 0.45 | 0.08 | 6 | T Cell |
| Sp100 | 1.19 | 0.00 | 0.00 | 0.40 | 0.07 | 6 | T Cell |
| Epsti1 | 1.17 | 0.00 | 0.00 | 0.30 | 0.03 | 6 | T Cell |
| Skap1 | 1.17 | 0.00 | 0.00 | 0.31 | 0.01 | 6 | T Cell |
| Il2rg | 1.13 | 0.00 | 0.00 | 0.41 | 0.09 | 6 | T Cell |
| Sh3kbp1 | 1.13 | 0.00 | 0.00 | 0.40 | 0.09 | 6 | T Cell |
| Cd3e | 1.12 | 0.00 | 0.00 | 0.28 | 0.01 | 6 | T Cell |
| Sell | 1.11 | 0.00 | 0.00 | 0.28 | 0.03 | 6 | T Cell |
| Hmha1 | 1.10 | 0.00 | 0.00 | 0.37 | 0.07 | 6 | T Cell |
| Cytip | 1.09 | 0.00 | 0.00 | 0.39 | 0.08 | 6 | T Cell |
| Gm10275 | 1.07 | 0.00 | 0.00 | 0.90 | 0.65 | 6 | T Cell |
| Lat | 1.07 | 0.00 | 0.00 | 0.26 | 0.01 | 6 | T Cell |
| Ptpn18 | 1.07 | 0.00 | 0.00 | 0.48 | 0.13 | 6 | T Cell |
| Slfn2 | 1.06 | 0.00 | 0.00 | 0.48 | 0.12 | 6 | T Cell |
| Ms4a4c | 1.05 | 0.00 | 0.00 | 0.28 | 0.02 | 6 | T Cell |
| Satb1 | 1.02 | 0.00 | 0.00 | 0.25 | 0.03 | 6 | T Cell |
| Dgka | 1.01 | 0.00 | 0.00 | 0.28 | 0.04 | 6 | T Cell |
| Itgb7 | 1.00 | 0.00 | 0.00 | 0.28 | 0.03 | 6 | T Cell |
| Ltb | 0.99 | 0.00 | 0.00 | 0.27 | 0.03 | 6 | T Cell |
| Bin2 | 0.96 | 0.00 | 0.00 | 0.28 | 0.04 | 6 | T Cell |
| Rplp0 | 0.95 | 0.00 | 0.00 | 0.97 | 0.83 | 6 | T Cell |
| Gm8730 | 0.94 | 0.00 | 0.00 | 0.97 | 0.81 | 6 | T Cell |
| B2m | 0.93 | 0.00 | 0.00 | 0.97 | 0.66 | 6 | T Cell |
| Rpsa-ps10 | 0.93 | 0.00 | 0.00 | 0.90 | 0.67 | 6 | T Cell |
| Rpl13a | 0.89 | 0.00 | 0.00 | 0.98 | 0.86 | 6 | T Cell |
| Rplp1 | 0.87 | 0.00 | 0.00 | 0.98 | 0.89 | 6 | T Cell |
| Rps15a | 0.86 | 0.00 | 0.00 | 0.97 | 0.81 | 6 | T Cell |
| Gm10073 | 0.85 | 0.00 | 0.00 | 0.97 | 0.85 | 6 | T Cell |
| Rpl32 | 0.82 | 0.00 | 0.00 | 0.97 | 0.84 | 6 | T Cell |
| Rps11 | 0.81 | 0.00 | 0.00 | 0.93 | 0.75 | 6 | T Cell |
| Pabpc1 | 0.81 | 0.00 | 0.00 | 0.95 | 0.80 | 6 | T Cell |
| Wdr89 | 0.81 | 0.00 | 0.00 | 0.98 | 0.88 | 6 | T Cell |
| Rps26 | 0.80 | 0.00 | 0.00 | 0.99 | 0.89 | 6 | T Cell |
| Rps26-ps1 | 0.79 | 0.00 | 0.00 | 0.97 | 0.87 | 6 | T Cell |
| Rps3 | 0.79 | 0.00 | 0.00 | 0.96 | 0.82 | 6 | T Cell |
| Rps16 | 0.78 | 0.00 | 0.00 | 0.97 | 0.83 | 6 | T Cell |
| Rpl4 | 0.76 | 0.00 | 0.00 | 0.97 | 0.83 | 6 | T Cell |
| Rps3a1 | 0.70 | 0.00 | 0.00 | 0.97 | 0.85 | 6 | T Cell |
| Tmsb4x | 0.67 | 0.00 | 0.00 | 0.99 | 0.88 | 6 | T Cell |
| Rps9 | 0.64 | 0.00 | 0.00 | 0.99 | 0.91 | 6 | T Cell |
| Coro1a | 0.94 | 0.00 | 0.00 | 0.39 | 0.09 | 6 | T Cell |
| Rps6 | 0.73 | 0.00 | 0.00 | 0.95 | 0.82 | 6 | T Cell |
| Gm6472 | 0.77 | 0.00 | 0.00 | 0.94 | 0.79 | 6 | T Cell |
| Gm9396 | 1.02 | 0.00 | 0.00 | 0.82 | 0.53 | 6 | T Cell |
| Arhgap30 | 0.95 | 0.00 | 0.00 | 0.39 | 0.09 | 6 | T Cell |
| H2-D1 | 0.91 | 0.00 | 0.00 | 0.86 | 0.48 | 6 | T Cell |
| Gm9846 | 0.69 | 0.00 | 0.00 | 0.98 | 0.89 | 6 | T Cell |
| Rpl18a | 0.75 | 0.00 | 0.00 | 0.96 | 0.82 | 6 | T Cell |
| Rps24 | 0.69 | 0.00 | 0.00 | 0.98 | 0.88 | 6 | T Cell |
| Rplp2 | 0.74 | 0.00 | 0.00 | 0.95 | 0.81 | 6 | T Cell |
| Rps10-ps1 | 0.75 | 0.00 | 0.00 | 0.94 | 0.77 | 6 | T Cell |
| Cotl1 | 1.05 | 0.00 | 0.00 | 0.51 | 0.16 | 6 | T Cell |
| Mndal | 0.93 | 0.00 | 0.00 | 0.50 | 0.15 | 6 | T Cell |
| Gm9843 | 0.60 | 0.00 | 0.00 | 0.99 | 0.92 | 6 | T Cell |
| Rpl5 | 0.77 | 0.00 | 0.00 | 0.92 | 0.74 | 6 | T Cell |
| Rpl12 | 1.00 | 0.00 | 0.00 | 0.82 | 0.54 | 6 | T Cell |
| Ifi203 | 0.99 | 0.00 | 0.00 | 0.45 | 0.13 | 6 | T Cell |
| Tmsb10 | 0.68 | 0.00 | 0.00 | 0.98 | 0.85 | 6 | T Cell |
| Psmb8 | 0.98 | 0.00 | 0.00 | 0.52 | 0.17 | 6 | T Cell |
| Apbb1ip | 0.88 | 0.00 | 0.00 | 0.36 | 0.08 | 6 | T Cell |
| Sp110 | 0.91 | 0.00 | 0.00 | 0.30 | 0.06 | 6 | T Cell |
| A430104N18Rik | 0.86 | 0.00 | 0.00 | 0.29 | 0.06 | 6 | T Cell |
| Rps3a2 | 0.67 | 0.00 | 0.00 | 0.95 | 0.82 | 6 | T Cell |
| Fyb | 0.99 | 0.00 | 0.00 | 0.51 | 0.17 | 6 | T Cell |
| Stk17b | 0.94 | 0.00 | 0.00 | 0.28 | 0.06 | 6 | T Cell |
| Pdcd4 | 1.14 | 0.00 | 0.00 | 0.55 | 0.21 | 6 | T Cell |
| 6-Sep | 0.90 | 0.00 | 0.00 | 0.29 | 0.06 | 6 | T Cell |
| Rpl8 | 0.70 | 0.00 | 0.00 | 0.93 | 0.79 | 6 | T Cell |
| Rpsa | 0.86 | 0.00 | 0.00 | 0.85 | 0.61 | 6 | T Cell |
| Gimap6 | 1.00 | 0.00 | 0.00 | 0.47 | 0.15 | 6 | T Cell |
| Pycard | 0.94 | 0.00 | 0.00 | 0.33 | 0.08 | 6 | T Cell |
| Rps20 | 0.71 | 0.00 | 0.00 | 0.91 | 0.78 | 6 | T Cell |
| Rps5 | 0.70 | 0.00 | 0.00 | 0.92 | 0.77 | 6 | T Cell |
| Eef1a1 | 0.61 | 0.00 | 0.00 | 0.95 | 0.87 | 6 | T Cell |
| Rps7 | 0.72 | 0.00 | 0.00 | 0.90 | 0.73 | 6 | T Cell |
| Rpl39 | 0.64 | 0.00 | 0.00 | 0.94 | 0.80 | 6 | T Cell |
| Gnb2l1 | 0.76 | 0.00 | 0.00 | 0.85 | 0.64 | 6 | T Cell |
| Gm9493 | 0.71 | 0.00 | 0.00 | 0.90 | 0.74 | 6 | T Cell |
| Eef1b2 | 0.78 | 0.00 | 0.00 | 0.85 | 0.64 | 6 | T Cell |
| Gm11808 | 0.66 | 0.00 | 0.00 | 0.92 | 0.77 | 6 | T Cell |
| Gm2000 | 0.60 | 0.00 | 0.00 | 0.94 | 0.84 | 6 | T Cell |
| Fxyd5 | 0.85 | 0.00 | 0.00 | 0.48 | 0.17 | 6 | T Cell |
| Gm7536 | 0.61 | 0.00 | 0.00 | 0.94 | 0.81 | 6 | T Cell |
| Rpl19 | 0.66 | 0.00 | 0.00 | 0.90 | 0.75 | 6 | T Cell |
| Rpl10a | 0.68 | 0.00 | 0.00 | 0.89 | 0.74 | 6 | T Cell |
| Rps3a3 | 0.63 | 0.00 | 0.00 | 0.92 | 0.77 | 6 | T Cell |
| Rps14 | 0.52 | 0.00 | 0.00 | 0.99 | 0.92 | 6 | T Cell |
| Rps4x | 0.55 | 0.00 | 0.00 | 0.96 | 0.86 | 6 | T Cell |
| Gimap9 | 0.89 | 0.00 | 0.00 | 0.28 | 0.07 | 6 | T Cell |
| Rps18 | 0.71 | 0.00 | 0.00 | 0.84 | 0.65 | 6 | T Cell |
| Tmem71 | 0.86 | 0.00 | 0.00 | 0.25 | 0.06 | 6 | T Cell |
| Rpl22 | 0.61 | 0.00 | 0.00 | 0.92 | 0.79 | 6 | T Cell |
| Rps10 | 0.66 | 0.00 | 0.00 | 0.87 | 0.70 | 6 | T Cell |
| Rps21 | 0.64 | 0.00 | 0.00 | 0.91 | 0.75 | 6 | T Cell |
| Rps13 | 0.73 | 0.00 | 0.00 | 0.83 | 0.63 | 6 | T Cell |
| Rpl13 | 0.59 | 0.00 | 0.00 | 0.93 | 0.79 | 6 | T Cell |
| Rpl23 | 0.55 | 0.00 | 0.00 | 0.96 | 0.86 | 6 | T Cell |
| Gm10288 | 0.60 | 0.00 | 0.00 | 0.91 | 0.78 | 6 | T Cell |
| Rpl3 | 0.72 | 0.00 | 0.00 | 0.82 | 0.62 | 6 | T Cell |
| Gm10269 | 0.61 | 0.00 | 0.00 | 0.92 | 0.79 | 6 | T Cell |
| Rpl6 | 0.52 | 0.00 | 0.00 | 0.95 | 0.87 | 6 | T Cell |
| Rps23 | 0.61 | 0.00 | 0.00 | 0.92 | 0.79 | 6 | T Cell |
| Ets1 | 0.92 | 0.00 | 0.00 | 0.55 | 0.24 | 6 | T Cell |
| Gmfg | 0.85 | 0.00 | 0.00 | 0.39 | 0.13 | 6 | T Cell |
| Rpl14 | 0.57 | 0.00 | 0.00 | 0.93 | 0.83 | 6 | T Cell |
| Rpl23a-ps3 | 0.61 | 0.00 | 0.00 | 0.90 | 0.74 | 6 | T Cell |
| Fau | 0.52 | 0.00 | 0.00 | 0.96 | 0.86 | 6 | T Cell |
| Rps27 | 0.59 | 0.00 | 0.00 | 0.94 | 0.81 | 6 | T Cell |
| Actb | 0.51 | 0.00 | 0.00 | 1.00 | 0.98 | 6 | T Cell |
| Rps28 | 0.58 | 0.00 | 0.00 | 0.95 | 0.83 | 6 | T Cell |
| Gm10036 | 0.60 | 0.00 | 0.00 | 0.89 | 0.74 | 6 | T Cell |
| Gm10335 | 0.59 | 0.00 | 0.00 | 0.89 | 0.72 | 6 | T Cell |
| Rpl34 | 0.56 | 0.00 | 0.00 | 0.93 | 0.79 | 6 | T Cell |
| Rps29 | 0.55 | 0.00 | 0.00 | 0.97 | 0.90 | 6 | T Cell |
| Cd37 | 0.57 | 0.00 | 0.00 | 0.28 | 0.07 | 6 | T Cell |
| Gm10263 | 0.64 | 0.00 | 0.00 | 0.90 | 0.76 | 6 | T Cell |
| Rpl35 | 0.54 | 0.00 | 0.00 | 0.91 | 0.80 | 6 | T Cell |
| Gm10132 | 0.57 | 0.00 | 0.00 | 0.88 | 0.72 | 6 | T Cell |
| Rpl27-ps3 | 0.57 | 0.00 | 0.00 | 0.90 | 0.76 | 6 | T Cell |
| Rpl13a-ps1 | 0.76 | 0.00 | 0.00 | 0.75 | 0.54 | 6 | T Cell |
| Rpl13-ps3 | 0.56 | 0.00 | 0.00 | 0.89 | 0.75 | 6 | T Cell |
| Rpl27a | 0.54 | 0.00 | 0.00 | 0.91 | 0.78 | 6 | T Cell |
| Rpl9-ps6 | 0.58 | 0.00 | 0.00 | 0.86 | 0.73 | 6 | T Cell |
| Gm17511 | 0.71 | 0.00 | 0.00 | 0.80 | 0.60 | 6 | T Cell |
| Lcp1 | 0.74 | 0.00 | 0.00 | 0.50 | 0.22 | 6 | T Cell |
| Jak1 | 0.91 | 0.00 | 0.00 | 0.51 | 0.24 | 6 | T Cell |
| Rpl18 | 0.64 | 0.00 | 0.00 | 0.81 | 0.63 | 6 | T Cell |
| Emb | 0.80 | 0.00 | 0.00 | 0.32 | 0.10 | 6 | T Cell |
| Rpl41 | 0.38 | 0.00 | 0.00 | 0.98 | 0.95 | 6 | T Cell |
| Lrrc58 | 0.54 | 0.00 | 0.00 | 0.97 | 0.91 | 6 | T Cell |
| Rapgef6 | 0.96 | 0.00 | 0.00 | 0.35 | 0.13 | 6 | T Cell |
| Rps15 | 0.53 | 0.00 | 0.00 | 0.91 | 0.78 | 6 | T Cell |
| Mbnl1 | 0.87 | 0.00 | 0.00 | 0.51 | 0.26 | 6 | T Cell |
| Rps25 | 0.50 | 0.00 | 0.00 | 0.91 | 0.80 | 6 | T Cell |
| Rps18-ps3 | 0.66 | 0.00 | 0.00 | 0.75 | 0.55 | 6 | T Cell |
| Gm5428 | 0.50 | 0.00 | 0.00 | 0.90 | 0.78 | 6 | T Cell |
| Rpl23a | 0.56 | 0.00 | 0.00 | 0.82 | 0.66 | 6 | T Cell |
| Lsp1 | 0.59 | 0.00 | 0.00 | 0.45 | 0.18 | 6 | T Cell |
| Gimap1 | 0.73 | 0.00 | 0.00 | 0.36 | 0.13 | 6 | T Cell |
| Gm10260 | 0.70 | 0.00 | 0.00 | 0.69 | 0.48 | 6 | T Cell |
| Actr3 | 0.75 | 0.00 | 0.00 | 0.57 | 0.32 | 6 | T Cell |
| Crlf3 | 0.87 | 0.00 | 0.00 | 0.34 | 0.13 | 6 | T Cell |
| Rgs10 | 0.82 | 0.00 | 0.00 | 0.28 | 0.09 | 6 | T Cell |
| Pfn1 | 0.72 | 0.00 | 0.00 | 0.67 | 0.46 | 6 | T Cell |
| Rpl27 | 0.51 | 0.00 | 0.00 | 0.84 | 0.69 | 6 | T Cell |
| Leprotl1 | 0.79 | 0.00 | 0.00 | 0.35 | 0.14 | 6 | T Cell |
| Rpl7 | 0.52 | 0.00 | 0.00 | 0.87 | 0.74 | 6 | T Cell |
| Rpl35a | 0.48 | 0.00 | 0.00 | 0.89 | 0.79 | 6 | T Cell |
| Smc4 | 0.93 | 0.00 | 0.00 | 0.57 | 0.34 | 6 | T Cell |
| Sh3bgrl3 | 0.70 | 0.00 | 0.00 | 0.58 | 0.34 | 6 | T Cell |
| Gm7589 | 0.53 | 0.00 | 0.00 | 0.80 | 0.65 | 6 | T Cell |
| Npm1 | 0.62 | 0.00 | 0.00 | 0.75 | 0.57 | 6 | T Cell |
| Gm10076 | 0.36 | 0.00 | 0.00 | 0.96 | 0.92 | 6 | T Cell |
| Elf1 | 0.80 | 0.00 | 0.00 | 0.41 | 0.20 | 6 | T Cell |
| Itga4 | 0.73 | 0.00 | 0.00 | 0.26 | 0.09 | 6 | T Cell |
| Atp1b3 | 0.74 | 0.00 | 0.00 | 0.47 | 0.25 | 6 | T Cell |
| Rpl26 | 0.43 | 0.00 | 0.00 | 0.90 | 0.81 | 6 | T Cell |
| Rpl37 | 0.43 | 0.00 | 0.00 | 0.90 | 0.80 | 6 | T Cell |
| Rpl15 | 0.56 | 0.00 | 0.00 | 0.73 | 0.58 | 6 | T Cell |
| Srpk1 | 0.76 | 0.00 | 0.00 | 0.30 | 0.12 | 6 | T Cell |
| Rpl11 | 0.51 | 0.00 | 0.00 | 0.79 | 0.67 | 6 | T Cell |
| Rps27a | 0.44 | 0.00 | 0.00 | 0.87 | 0.76 | 6 | T Cell |
| Gm7808 | 0.56 | 0.00 | 0.00 | 0.75 | 0.59 | 6 | T Cell |
| Arpc2 | 0.51 | 0.00 | 0.00 | 0.79 | 0.61 | 6 | T Cell |
| Ifi27l2a | 0.85 | 0.00 | 0.00 | 0.25 | 0.09 | 6 | T Cell |
| Rps12-ps3 | 0.44 | 0.00 | 0.00 | 0.88 | 0.78 | 6 | T Cell |
| Gm10020 | 0.60 | 0.00 | 0.00 | 0.68 | 0.51 | 6 | T Cell |
| Tpt1 | 0.36 | 0.00 | 0.00 | 0.95 | 0.88 | 6 | T Cell |
| Ablim1 | 0.71 | 0.00 | 0.00 | 0.46 | 0.24 | 6 | T Cell |
| Rps2 | 0.50 | 0.00 | 0.00 | 0.79 | 0.67 | 6 | T Cell |
| Gm5239 | 0.60 | 0.00 | 0.00 | 0.68 | 0.51 | 6 | T Cell |
| Mir682 | 0.38 | 0.00 | 0.00 | 0.93 | 0.85 | 6 | T Cell |
| S100a10 | 0.77 | 0.00 | 0.00 | 0.57 | 0.38 | 6 | T Cell |
| Rps17 | 0.42 | 0.00 | 0.00 | 0.85 | 0.77 | 6 | T Cell |
| Rpl30 | 0.55 | 0.00 | 0.00 | 0.72 | 0.58 | 6 | T Cell |
| Rps19 | 0.35 | 0.00 | 0.00 | 0.92 | 0.82 | 6 | T Cell |
| Fkbp3 | 0.74 | 0.00 | 0.00 | 0.49 | 0.29 | 6 | T Cell |
| AC121131.2 | 0.63 | 0.00 | 0.00 | 0.57 | 0.39 | 6 | T Cell |
| AC123611.1 | 0.63 | 0.00 | 0.00 | 0.56 | 0.39 | 6 | T Cell |
| Rpl9 | 0.48 | 0.00 | 0.00 | 0.74 | 0.62 | 6 | T Cell |
| Rpl17 | 0.55 | 0.00 | 0.00 | 0.69 | 0.53 | 6 | T Cell |
| Srgn | 0.49 | 0.00 | 0.00 | 0.56 | 0.32 | 6 | T Cell |
| Gvin1 | 0.60 | 0.00 | 0.00 | 0.27 | 0.11 | 6 | T Cell |
| Rps8 | 0.37 | 0.00 | 0.00 | 0.89 | 0.81 | 6 | T Cell |
| Orc5 | 0.51 | 0.00 | 0.00 | 0.69 | 0.55 | 6 | T Cell |
| Rpl37a | 0.33 | 0.00 | 0.00 | 0.90 | 0.83 | 6 | T Cell |
| Cdk2ap2 | 0.62 | 0.00 | 0.00 | 0.31 | 0.15 | 6 | T Cell |
| Rpl36a | 0.46 | 0.00 | 0.00 | 0.74 | 0.62 | 6 | T Cell |
| Gm4070 | 0.60 | 0.00 | 0.00 | 0.27 | 0.11 | 6 | T Cell |
| Gimap5 | 0.58 | 0.00 | 0.00 | 0.26 | 0.11 | 6 | T Cell |
| Rpl36 | 0.43 | 0.00 | 0.00 | 0.76 | 0.63 | 6 | T Cell |
| Rpl22l1 | 0.48 | 0.00 | 0.00 | 0.69 | 0.56 | 6 | T Cell |
| Samhd1 | 0.58 | 0.00 | 0.00 | 0.36 | 0.19 | 6 | T Cell |
| Gltscr2 | 0.52 | 0.00 | 0.00 | 0.63 | 0.47 | 6 | T Cell |
| Nt5c | 0.59 | 0.00 | 0.00 | 0.30 | 0.15 | 6 | T Cell |
| Ldha | 0.59 | 0.00 | 0.00 | 0.45 | 0.29 | 6 | T Cell |
| Rpl38 | 0.36 | 0.00 | 0.00 | 0.82 | 0.73 | 6 | T Cell |
| Bzw2 | 0.64 | 0.00 | 0.00 | 0.29 | 0.14 | 6 | T Cell |
| Arpc1b | 0.48 | 0.00 | 0.00 | 0.53 | 0.37 | 6 | T Cell |
| Rpl21 | 0.36 | 0.00 | 0.00 | 0.85 | 0.75 | 6 | T Cell |
| Rps12 | 0.38 | 0.00 | 0.00 | 0.77 | 0.68 | 6 | T Cell |
| Rpl28 | 0.45 | 0.00 | 0.00 | 0.65 | 0.53 | 6 | T Cell |
| Stk4 | 0.62 | 0.00 | 0.00 | 0.27 | 0.14 | 6 | T Cell |
| Psme2b | 0.54 | 0.00 | 0.00 | 0.45 | 0.29 | 6 | T Cell |
| Ccnd2 | 0.52 | 0.00 | 0.00 | 0.45 | 0.28 | 6 | T Cell |
| H2-T23 | 0.53 | 0.00 | 0.00 | 0.25 | 0.12 | 6 | T Cell |
| Naca | 0.42 | 0.00 | 0.00 | 0.68 | 0.57 | 6 | T Cell |
| Gm5611 | 0.51 | 0.00 | 0.00 | 0.49 | 0.35 | 6 | T Cell |
| Psme1 | 0.53 | 0.00 | 0.00 | 0.46 | 0.31 | 6 | T Cell |
| Rpl36al | 0.42 | 0.00 | 0.00 | 0.64 | 0.52 | 6 | T Cell |
| Serbp1 | 0.36 | 0.00 | 0.00 | 0.81 | 0.74 | 6 | T Cell |
| Eef1g | 0.45 | 0.00 | 0.00 | 0.60 | 0.48 | 6 | T Cell |
| Gm10126 | 0.55 | 0.00 | 0.00 | 0.52 | 0.39 | 6 | T Cell |
| Eif3e | 0.47 | 0.00 | 0.00 | 0.53 | 0.40 | 6 | T Cell |
| Arpc5 | 0.42 | 0.00 | 0.00 | 0.66 | 0.56 | 6 | T Cell |
| Tmem66 | 0.56 | 0.00 | 0.00 | 0.33 | 0.20 | 6 | T Cell |
| AC167036.1 | 0.56 | 0.00 | 0.00 | 0.42 | 0.30 | 6 | T Cell |
| mt-Nd5 | 0.33 | 0.00 | 0.00 | 0.83 | 0.75 | 6 | T Cell |
| Gm17541 | 0.54 | 0.00 | 0.00 | 0.30 | 0.18 | 6 | T Cell |
| Gm5093 | 0.43 | 0.00 | 0.00 | 0.58 | 0.46 | 6 | T Cell |
| Eno1 | 0.47 | 0.00 | 0.00 | 0.28 | 0.16 | 6 | T Cell |
| Ncl | 0.31 | 0.00 | 0.00 | 0.84 | 0.78 | 6 | T Cell |
| mt-Atp6 | 0.25 | 0.00 | 0.00 | 0.90 | 0.84 | 6 | T Cell |
| Klf2 | 0.46 | 0.00 | 0.00 | 0.33 | 0.19 | 6 | T Cell |
| Nme1 | 0.48 | 0.00 | 0.00 | 0.31 | 0.19 | 6 | T Cell |
| Rpl36-ps3 | 0.35 | 0.00 | 0.00 | 0.64 | 0.53 | 6 | T Cell |
| Cd47 | 0.44 | 0.00 | 0.00 | 0.55 | 0.43 | 6 | T Cell |
| Psme2 | 0.47 | 0.00 | 0.00 | 0.36 | 0.23 | 6 | T Cell |
| Crip1 | 0.29 | 0.00 | 0.00 | 0.62 | 0.47 | 6 | T Cell |
| Stk38 | 0.55 | 0.00 | 0.00 | 0.31 | 0.20 | 6 | T Cell |
| Cfl1 | 0.36 | 0.00 | 0.00 | 0.67 | 0.58 | 6 | T Cell |
| Emg1 | 0.47 | 0.00 | 0.00 | 0.26 | 0.15 | 6 | T Cell |
| Eif3h | 0.45 | 0.00 | 0.00 | 0.46 | 0.35 | 6 | T Cell |
| Snrpf | 0.44 | 0.00 | 0.00 | 0.40 | 0.28 | 6 | T Cell |
| Akap13 | 0.46 | 0.00 | 0.00 | 0.38 | 0.27 | 6 | T Cell |
| Abracl | 0.47 | 0.00 | 0.00 | 0.35 | 0.23 | 6 | T Cell |
| Eef2 | 0.36 | 0.00 | 0.00 | 0.64 | 0.57 | 6 | T Cell |
| Sec11c | 0.45 | 0.00 | 0.00 | 0.29 | 0.17 | 6 | T Cell |
| Xrn2 | 0.52 | 0.00 | 0.00 | 0.40 | 0.30 | 6 | T Cell |
| Mif | 0.46 | 0.00 | 0.00 | 0.29 | 0.18 | 6 | T Cell |
| Emp3 | 0.40 | 0.00 | 0.00 | 0.33 | 0.21 | 6 | T Cell |
| Rpl31 | 0.32 | 0.00 | 0.00 | 0.67 | 0.58 | 6 | T Cell |
| Ddx5 | 0.30 | 0.00 | 0.00 | 0.77 | 0.70 | 6 | T Cell |
| Gm24865 | 0.32 | 0.00 | 0.00 | 0.69 | 0.63 | 6 | T Cell |
| Anp32a | 0.26 | 0.00 | 0.00 | 0.70 | 0.60 | 6 | T Cell |
| Rpl29 | 0.37 | 0.00 | 0.00 | 0.54 | 0.46 | 6 | T Cell |
| Gas5 | 0.31 | 0.00 | 0.00 | 0.62 | 0.53 | 6 | T Cell |
| Gm22758 | 0.25 | 0.00 | 0.00 | 0.76 | 0.69 | 6 | T Cell |
| Sf3b1 | 0.40 | 0.00 | 0.00 | 0.55 | 0.46 | 6 | T Cell |
| Hnrnpf | 0.41 | 0.00 | 0.00 | 0.44 | 0.34 | 6 | T Cell |
| Tpm3-rs7 | 0.25 | 0.00 | 0.00 | 0.76 | 0.69 | 6 | T Cell |
| Oaz1 | 0.31 | 0.00 | 0.00 | 0.67 | 0.62 | 6 | T Cell |
| Trp53 | 0.42 | 0.00 | 0.00 | 0.25 | 0.16 | 6 | T Cell |
| Gm8186 | 0.35 | 0.00 | 0.00 | 0.48 | 0.39 | 6 | T Cell |
| Nop10 | 0.42 | 0.00 | 0.00 | 0.37 | 0.27 | 6 | T Cell |
| Gm3940 | 0.37 | 0.00 | 0.00 | 0.45 | 0.36 | 6 | T Cell |
| Gm5244 | 0.47 | 0.00 | 0.00 | 0.37 | 0.28 | 6 | T Cell |
| Add3 | 0.37 | 0.00 | 0.00 | 0.38 | 0.28 | 6 | T Cell |
| Gm15013 | 0.39 | 0.00 | 0.00 | 0.45 | 0.37 | 6 | T Cell |
| Ddx21 | 0.43 | 0.00 | 0.00 | 0.35 | 0.26 | 6 | T Cell |
| Hspe1 | 0.40 | 0.00 | 0.00 | 0.37 | 0.28 | 6 | T Cell |
| Snrpg | 0.34 | 0.00 | 0.00 | 0.47 | 0.39 | 6 | T Cell |
| Gm17087 | 0.41 | 0.00 | 0.00 | 0.33 | 0.24 | 6 | T Cell |
| Cep110 | 0.48 | 0.00 | 0.00 | 0.27 | 0.19 | 6 | T Cell |
| Eif3m | 0.37 | 0.00 | 0.00 | 0.33 | 0.24 | 6 | T Cell |
| Fgfr1op2 | 0.39 | 0.00 | 0.00 | 0.36 | 0.28 | 6 | T Cell |
| Snrpd3 | 0.37 | 0.00 | 0.00 | 0.40 | 0.32 | 6 | T Cell |
| Cox7a2l | 0.34 | 0.00 | 0.00 | 0.49 | 0.41 | 6 | T Cell |
| Btf3 | 0.30 | 0.00 | 0.00 | 0.58 | 0.51 | 6 | T Cell |
| Rnaset2a | 0.38 | 0.00 | 0.00 | 0.35 | 0.26 | 6 | T Cell |
| Tsc22d4 | 0.37 | 0.00 | 0.00 | 0.36 | 0.27 | 6 | T Cell |
| Cdc42se1 | 0.42 | 0.00 | 0.00 | 0.28 | 0.19 | 6 | T Cell |
| Rps2-ps6 | 0.31 | 0.00 | 0.00 | 0.48 | 0.41 | 6 | T Cell |
| Gm8226 | 0.34 | 0.00 | 0.00 | 0.47 | 0.40 | 6 | T Cell |
| Tpr | 0.32 | 0.00 | 0.00 | 0.57 | 0.49 | 6 | T Cell |
| Nop58 | 0.36 | 0.00 | 0.00 | 0.34 | 0.26 | 6 | T Cell |
| Cdkn1b | 0.43 | 0.00 | 0.00 | 0.29 | 0.21 | 6 | T Cell |
| Rps2-ps10 | 0.31 | 0.00 | 0.00 | 0.49 | 0.43 | 6 | T Cell |
| Psma2 | 0.32 | 0.00 | 0.00 | 0.50 | 0.44 | 6 | T Cell |
| Gm6133 | 0.36 | 0.00 | 0.00 | 0.41 | 0.34 | 6 | T Cell |
| Eif3k | 0.35 | 0.00 | 0.00 | 0.41 | 0.34 | 6 | T Cell |
| Dusp11 | 0.37 | 0.00 | 0.00 | 0.33 | 0.25 | 6 | T Cell |
| Rpl10a-ps2 | 0.31 | 0.00 | 0.00 | 0.36 | 0.28 | 6 | T Cell |
| Pkm | 0.38 | 0.00 | 0.00 | 0.30 | 0.22 | 6 | T Cell |
| Rpl21-ps6 | 0.28 | 0.00 | 0.00 | 0.50 | 0.43 | 6 | T Cell |
| Tspo | 0.34 | 0.00 | 0.00 | 0.33 | 0.26 | 6 | T Cell |
| Btg1 | 0.34 | 0.00 | 0.00 | 0.36 | 0.28 | 6 | T Cell |
| Nop56 | 0.37 | 0.00 | 0.00 | 0.27 | 0.20 | 6 | T Cell |
| Ifngr1 | 0.41 | 0.00 | 0.00 | 0.31 | 0.24 | 6 | T Cell |
| Rnaset2b | 0.37 | 0.00 | 0.00 | 0.33 | 0.26 | 6 | T Cell |
| Ddx46 | 0.34 | 0.00 | 0.00 | 0.37 | 0.30 | 6 | T Cell |
| Rbm3 | 0.30 | 0.00 | 0.00 | 0.47 | 0.41 | 6 | T Cell |
| Mdh1 | 0.33 | 0.00 | 0.00 | 0.26 | 0.19 | 6 | T Cell |
| Ppp1ca | 0.32 | 0.00 | 0.00 | 0.40 | 0.33 | 6 | T Cell |
| Gm6139 | 0.29 | 0.00 | 0.00 | 0.47 | 0.42 | 6 | T Cell |
| Lsm4 | 0.33 | 0.00 | 0.00 | 0.28 | 0.20 | 6 | T Cell |
| Polr1d | 0.34 | 0.00 | 0.00 | 0.35 | 0.28 | 6 | T Cell |
| Lrrfip1 | 0.35 | 0.00 | 0.00 | 0.32 | 0.25 | 6 | T Cell |
| Eif3i | 0.30 | 0.00 | 0.00 | 0.29 | 0.22 | 6 | T Cell |
| Gm5786 | 0.28 | 0.00 | 0.00 | 0.44 | 0.39 | 6 | T Cell |
| Actr2 | 0.28 | 0.00 | 0.00 | 0.34 | 0.27 | 6 | T Cell |
| Eif3f | 0.32 | 0.00 | 0.00 | 0.42 | 0.36 | 6 | T Cell |
| Ncor1 | 0.33 | 0.00 | 0.00 | 0.45 | 0.40 | 6 | T Cell |
| Cnbp | 0.29 | 0.00 | 0.00 | 0.49 | 0.45 | 6 | T Cell |
| Uqcrh | 0.25 | 0.00 | 0.00 | 0.52 | 0.47 | 6 | T Cell |
| Tma7 | 0.30 | 0.00 | 0.00 | 0.38 | 0.32 | 6 | T Cell |
| Pak2 | 0.31 | 0.00 | 0.00 | 0.35 | 0.29 | 6 | T Cell |
| Rpl21-ps4 | 0.29 | 0.00 | 0.00 | 0.40 | 0.35 | 6 | T Cell |
| Gm16477 | 0.26 | 0.00 | 0.00 | 0.43 | 0.39 | 6 | T Cell |
| G3bp2 | 0.25 | 0.00 | 0.00 | 0.37 | 0.31 | 6 | T Cell |
| Smek2 | 0.32 | 0.00 | 0.00 | 0.27 | 0.22 | 6 | T Cell |
| Top2b | 0.31 | 0.00 | 0.00 | 0.32 | 0.27 | 6 | T Cell |
| Spcs2 | 0.26 | 0.00 | 0.00 | 0.36 | 0.31 | 6 | T Cell |
| Ppp2r5c | 0.29 | 0.00 | 0.00 | 0.26 | 0.21 | 6 | T Cell |
| Thrap3 | 0.29 | 0.00 | 0.00 | 0.41 | 0.37 | 6 | T Cell |
| Rsl1d1 | 0.30 | 0.00 | 0.00 | 0.32 | 0.27 | 6 | T Cell |
| 2810474O19Rik | 0.34 | 0.00 | 0.00 | 0.31 | 0.26 | 6 | T Cell |
| Gm6576 | 0.27 | 0.00 | 0.00 | 0.36 | 0.31 | 6 | T Cell |
| Gm17669 | 0.32 | 0.00 | 0.00 | 0.27 | 0.22 | 6 | T Cell |
| Cct2 | 0.28 | 0.00 | 0.00 | 0.31 | 0.27 | 6 | T Cell |
| Cdc37 | 0.25 | 0.00 | 0.00 | 0.32 | 0.28 | 6 | T Cell |
| Itsn2 | 0.25 | 0.00 | 0.01 | 0.29 | 0.25 | 6 | T Cell |
| Rsbn1l | 0.32 | 0.00 | 0.03 | 0.29 | 0.25 | 6 | T Cell |
| Car4 | 2.74 | 0.00 | 0.00 | 0.88 | 0.05 | 7 | Endothelial |
| Kdr | 2.43 | 0.00 | 0.00 | 0.94 | 0.14 | 7 | Endothelial |
| Kitl | 2.42 | 0.00 | 0.00 | 0.95 | 0.20 | 7 | Endothelial |
| Ednrb | 2.25 | 0.00 | 0.00 | 0.86 | 0.13 | 7 | Endothelial |
| Scn7a | 2.13 | 0.00 | 0.00 | 1.00 | 0.27 | 7 | Endothelial |
| Cyp4b1 | 2.13 | 0.00 | 0.00 | 0.76 | 0.09 | 7 | Endothelial |
| Apln | 2.03 | 0.00 | 0.00 | 0.64 | 0.03 | 7 | Endothelial |
| Pmp22 | 1.95 | 0.00 | 0.00 | 0.92 | 0.29 | 7 | Endothelial |
| Igfbp7 | 1.94 | 0.00 | 0.00 | 0.96 | 0.37 | 7 | Endothelial |
| Cldn5 | 1.89 | 0.00 | 0.00 | 0.91 | 0.20 | 7 | Endothelial |
| Tspan13 | 1.89 | 0.00 | 0.00 | 0.92 | 0.23 | 7 | Endothelial |
| Ly6a | 1.78 | 0.00 | 0.00 | 0.84 | 0.17 | 7 | Endothelial |
| Tbx3 | 1.76 | 0.00 | 0.00 | 0.85 | 0.18 | 7 | Endothelial |
| Clec1a | 1.73 | 0.00 | 0.00 | 0.81 | 0.14 | 7 | Endothelial |
| Ecscr | 1.72 | 0.00 | 0.00 | 0.80 | 0.15 | 7 | Endothelial |
| Prx | 1.69 | 0.00 | 0.00 | 0.67 | 0.08 | 7 | Endothelial |
| Mest | 1.64 | 0.00 | 0.00 | 0.78 | 0.18 | 7 | Endothelial |
| Cdh5 | 1.63 | 0.00 | 0.00 | 0.88 | 0.21 | 7 | Endothelial |
| Icam2 | 1.62 | 0.00 | 0.00 | 0.74 | 0.14 | 7 | Endothelial |
| Egfl7 | 1.54 | 0.00 | 0.00 | 0.95 | 0.25 | 7 | Endothelial |
| Nrp1 | 1.53 | 0.00 | 0.00 | 0.92 | 0.37 | 7 | Endothelial |
| Cd34 | 1.52 | 0.00 | 0.00 | 0.74 | 0.18 | 7 | Endothelial |
| Sema6a | 1.51 | 0.00 | 0.00 | 0.61 | 0.10 | 7 | Endothelial |
| Myct1 | 1.49 | 0.00 | 0.00 | 0.68 | 0.11 | 7 | Endothelial |
| Tmem100 | 1.48 | 0.00 | 0.00 | 0.94 | 0.28 | 7 | Endothelial |
| Hpgd | 1.48 | 0.00 | 0.00 | 0.88 | 0.24 | 7 | Endothelial |
| Stmn2 | 1.46 | 0.00 | 0.00 | 0.72 | 0.13 | 7 | Endothelial |
| Ehd4 | 1.43 | 0.00 | 0.00 | 0.84 | 0.26 | 7 | Endothelial |
| Nhlrc2 | 1.42 | 0.00 | 0.00 | 0.50 | 0.07 | 7 | Endothelial |
| Fendrr | 1.42 | 0.00 | 0.00 | 0.75 | 0.18 | 7 | Endothelial |
| Acvrl1 | 1.42 | 0.00 | 0.00 | 0.69 | 0.16 | 7 | Endothelial |
| Cyth3 | 1.42 | 0.00 | 0.00 | 0.74 | 0.23 | 7 | Endothelial |
| Rgs12 | 1.41 | 0.00 | 0.00 | 0.60 | 0.10 | 7 | Endothelial |
| Emp2 | 1.41 | 0.00 | 0.00 | 0.90 | 0.33 | 7 | Endothelial |
| Ramp2 | 1.39 | 0.00 | 0.00 | 0.91 | 0.26 | 7 | Endothelial |
| Lphn2 | 1.38 | 0.00 | 0.00 | 0.60 | 0.12 | 7 | Endothelial |
| Pecam1 | 1.35 | 0.00 | 0.00 | 0.83 | 0.22 | 7 | Endothelial |
| Fmo1 | 1.35 | 0.00 | 0.00 | 0.58 | 0.10 | 7 | Endothelial |
| Flt1 | 1.35 | 0.00 | 0.00 | 0.67 | 0.14 | 7 | Endothelial |
| Piezo2 | 1.32 | 0.00 | 0.00 | 0.55 | 0.10 | 7 | Endothelial |
| Pcdh17 | 1.32 | 0.00 | 0.00 | 0.74 | 0.19 | 7 | Endothelial |
| 4-Sep | 1.32 | 0.00 | 0.00 | 0.74 | 0.21 | 7 | Endothelial |
| Pcdh1 | 1.31 | 0.00 | 0.00 | 0.44 | 0.06 | 7 | Endothelial |
| Tmem47 | 1.31 | 0.00 | 0.00 | 0.48 | 0.09 | 7 | Endothelial |
| S1pr1 | 1.30 | 0.00 | 0.00 | 0.67 | 0.17 | 7 | Endothelial |
| Gpr116 | 1.29 | 0.00 | 0.00 | 0.92 | 0.36 | 7 | Endothelial |
| Fkbp1a | 1.29 | 0.00 | 0.00 | 0.86 | 0.39 | 7 | Endothelial |
| Ly6c1 | 1.29 | 0.00 | 0.00 | 0.66 | 0.17 | 7 | Endothelial |
| Pcdha2 | 1.28 | 0.00 | 0.00 | 0.29 | 0.02 | 7 | Endothelial |
| Ctla2a | 1.28 | 0.00 | 0.00 | 0.73 | 0.19 | 7 | Endothelial |
| Tspan7 | 1.27 | 0.00 | 0.00 | 0.77 | 0.19 | 7 | Endothelial |
| Tmem204 | 1.27 | 0.00 | 0.00 | 0.57 | 0.14 | 7 | Endothelial |
| Dusp3 | 1.26 | 0.00 | 0.00 | 0.64 | 0.18 | 7 | Endothelial |
| Ahr | 1.26 | 0.00 | 0.00 | 0.50 | 0.08 | 7 | Endothelial |
| Thbd | 1.26 | 0.00 | 0.00 | 0.76 | 0.23 | 7 | Endothelial |
| Calcrl | 1.25 | 0.00 | 0.00 | 0.82 | 0.23 | 7 | Endothelial |
| Slc9a3r2 | 1.23 | 0.00 | 0.00 | 0.69 | 0.19 | 7 | Endothelial |
| Ppap2a | 1.23 | 0.00 | 0.00 | 0.64 | 0.18 | 7 | Endothelial |
| Tspan18 | 1.22 | 0.00 | 0.00 | 0.56 | 0.12 | 7 | Endothelial |
| Esam | 1.22 | 0.00 | 0.00 | 0.67 | 0.16 | 7 | Endothelial |
| Afap1l1 | 1.22 | 0.00 | 0.00 | 0.63 | 0.16 | 7 | Endothelial |
| Rtn1 | 1.19 | 0.00 | 0.00 | 0.32 | 0.02 | 7 | Endothelial |
| Timp3 | 1.19 | 0.00 | 0.00 | 0.80 | 0.35 | 7 | Endothelial |
| Emp1 | 1.18 | 0.00 | 0.00 | 0.66 | 0.20 | 7 | Endothelial |
| Ccnd1 | 1.17 | 0.00 | 0.00 | 0.60 | 0.18 | 7 | Endothelial |
| Chst1 | 1.16 | 0.00 | 0.00 | 0.31 | 0.01 | 7 | Endothelial |
| Ripply3 | 1.16 | 0.00 | 0.00 | 0.53 | 0.11 | 7 | Endothelial |
| Sdpr | 1.15 | 0.00 | 0.00 | 0.96 | 0.51 | 7 | Endothelial |
| Ly6e | 1.12 | 0.00 | 0.00 | 0.90 | 0.41 | 7 | Endothelial |
| Klf7 | 1.12 | 0.00 | 0.00 | 0.58 | 0.17 | 7 | Endothelial |
| Nostrin | 1.11 | 0.00 | 0.00 | 0.57 | 0.13 | 7 | Endothelial |
| Podxl | 1.11 | 0.00 | 0.00 | 0.49 | 0.10 | 7 | Endothelial |
| Mgll | 1.10 | 0.00 | 0.00 | 0.52 | 0.15 | 7 | Endothelial |
| Gimap4 | 1.10 | 0.00 | 0.00 | 0.60 | 0.16 | 7 | Endothelial |
| Scn3b | 1.09 | 0.00 | 0.00 | 0.39 | 0.07 | 7 | Endothelial |
| 9430020K01Rik | 1.08 | 0.00 | 0.00 | 0.51 | 0.12 | 7 | Endothelial |
| App | 1.08 | 0.00 | 0.00 | 0.93 | 0.58 | 7 | Endothelial |
| Col4a2 | 1.07 | 0.00 | 0.00 | 0.85 | 0.42 | 7 | Endothelial |
| Trib2 | 1.07 | 0.00 | 0.00 | 0.45 | 0.10 | 7 | Endothelial |
| Prickle1 | 1.07 | 0.00 | 0.00 | 0.38 | 0.06 | 7 | Endothelial |
| Slc16a9 | 1.06 | 0.00 | 0.00 | 0.32 | 0.05 | 7 | Endothelial |
| Klf4 | 1.06 | 0.00 | 0.00 | 0.67 | 0.22 | 7 | Endothelial |
| Lmo2 | 1.06 | 0.00 | 0.00 | 0.38 | 0.06 | 7 | Endothelial |
| Cd36 | 1.05 | 0.00 | 0.00 | 0.78 | 0.32 | 7 | Endothelial |
| Jun | 1.03 | 0.00 | 0.00 | 0.83 | 0.35 | 7 | Endothelial |
| Foxf1 | 1.03 | 0.00 | 0.00 | 0.46 | 0.11 | 7 | Endothelial |
| Impdh1 | 1.01 | 0.00 | 0.00 | 0.38 | 0.07 | 7 | Endothelial |
| Rapgef5 | 1.01 | 0.00 | 0.00 | 0.43 | 0.10 | 7 | Endothelial |
| Myzap | 0.99 | 0.00 | 0.00 | 0.53 | 0.13 | 7 | Endothelial |
| Epas1 | 0.99 | 0.00 | 0.00 | 0.72 | 0.24 | 7 | Endothelial |
| Emcn | 0.99 | 0.00 | 0.00 | 0.46 | 0.10 | 7 | Endothelial |
| Enho | 0.99 | 0.00 | 0.00 | 0.26 | 0.02 | 7 | Endothelial |
| Ccdc68 | 0.98 | 0.00 | 0.00 | 0.29 | 0.05 | 7 | Endothelial |
| Arhgef3 | 0.98 | 0.00 | 0.00 | 0.36 | 0.06 | 7 | Endothelial |
| Tnfsf10 | 0.96 | 0.00 | 0.00 | 0.38 | 0.08 | 7 | Endothelial |
| Stard8 | 0.95 | 0.00 | 0.00 | 0.35 | 0.06 | 7 | Endothelial |
| Dcbld1 | 0.95 | 0.00 | 0.00 | 0.26 | 0.04 | 7 | Endothelial |
| Edil3 | 0.95 | 0.00 | 0.00 | 0.26 | 0.02 | 7 | Endothelial |
| Arap2 | 0.95 | 0.00 | 0.00 | 0.36 | 0.07 | 7 | Endothelial |
| Mapt | 0.92 | 0.00 | 0.00 | 0.28 | 0.04 | 7 | Endothelial |
| Clic5 | 0.92 | 0.00 | 0.00 | 0.70 | 0.22 | 7 | Endothelial |
| Gata2 | 0.92 | 0.00 | 0.00 | 0.51 | 0.13 | 7 | Endothelial |
| Adrb1 | 0.85 | 0.00 | 0.00 | 0.29 | 0.05 | 7 | Endothelial |
| Ybx1 | 0.93 | 0.00 | 0.00 | 0.91 | 0.63 | 7 | Endothelial |
| Arhgef12 | 1.09 | 0.00 | 0.00 | 0.63 | 0.22 | 7 | Endothelial |
| Bmpr2 | 1.00 | 0.00 | 0.00 | 0.39 | 0.09 | 7 | Endothelial |
| Gimap6 | 0.99 | 0.00 | 0.00 | 0.53 | 0.15 | 7 | Endothelial |
| Pvrl3 | 1.03 | 0.00 | 0.00 | 0.40 | 0.09 | 7 | Endothelial |
| Ppp1r16b | 0.94 | 0.00 | 0.00 | 0.39 | 0.08 | 7 | Endothelial |
| Agfg1 | 0.93 | 0.00 | 0.00 | 0.30 | 0.05 | 7 | Endothelial |
| Cav2 | 0.99 | 0.00 | 0.00 | 0.56 | 0.18 | 7 | Endothelial |
| BC028528 | 0.89 | 0.00 | 0.00 | 0.72 | 0.28 | 7 | Endothelial |
| Col4a1 | 0.93 | 0.00 | 0.00 | 0.86 | 0.48 | 7 | Endothelial |
| Ddah1 | 0.89 | 0.00 | 0.00 | 0.30 | 0.05 | 7 | Endothelial |
| Nipal3 | 0.88 | 0.00 | 0.00 | 0.29 | 0.05 | 7 | Endothelial |
| Slco2a1 | 0.96 | 0.00 | 0.00 | 0.42 | 0.10 | 7 | Endothelial |
| Arhgap29 | 1.02 | 0.00 | 0.00 | 0.60 | 0.20 | 7 | Endothelial |
| Cd24a | 0.97 | 0.00 | 0.00 | 0.46 | 0.12 | 7 | Endothelial |
| Mfng | 0.84 | 0.00 | 0.00 | 0.26 | 0.04 | 7 | Endothelial |
| Cyyr1 | 0.85 | 0.00 | 0.00 | 0.49 | 0.14 | 7 | Endothelial |
| Marcks | 0.89 | 0.00 | 0.00 | 0.88 | 0.49 | 7 | Endothelial |
| Sptbn1 | 0.92 | 0.00 | 0.00 | 0.87 | 0.53 | 7 | Endothelial |
| Nav1 | 0.99 | 0.00 | 0.00 | 0.41 | 0.10 | 7 | Endothelial |
| Id3 | 0.84 | 0.00 | 0.00 | 0.84 | 0.44 | 7 | Endothelial |
| Rasip1 | 0.85 | 0.00 | 0.00 | 0.42 | 0.11 | 7 | Endothelial |
| Pitpnc1 | 0.96 | 0.00 | 0.00 | 0.42 | 0.11 | 7 | Endothelial |
| S100a16 | 0.89 | 0.00 | 0.00 | 0.62 | 0.23 | 7 | Endothelial |
| Tubb2a | 0.94 | 0.00 | 0.00 | 0.57 | 0.20 | 7 | Endothelial |
| Tmcc3 | 0.91 | 0.00 | 0.00 | 0.34 | 0.08 | 7 | Endothelial |
| Hspa12b | 0.75 | 0.00 | 0.00 | 0.28 | 0.05 | 7 | Endothelial |
| Sepp1 | 0.91 | 0.00 | 0.00 | 0.78 | 0.40 | 7 | Endothelial |
| 4931406P16Rik | 0.93 | 0.00 | 0.00 | 0.52 | 0.17 | 7 | Endothelial |
| Plekha1 | 0.93 | 0.00 | 0.00 | 0.40 | 0.11 | 7 | Endothelial |
| Ace | 0.83 | 0.00 | 0.00 | 0.45 | 0.13 | 7 | Endothelial |
| Tspan12 | 0.82 | 0.00 | 0.00 | 0.27 | 0.05 | 7 | Endothelial |
| Mllt4 | 0.98 | 0.00 | 0.00 | 0.54 | 0.20 | 7 | Endothelial |
| Jup | 0.94 | 0.00 | 0.00 | 0.47 | 0.15 | 7 | Endothelial |
| Id1 | 0.97 | 0.00 | 0.00 | 0.59 | 0.23 | 7 | Endothelial |
| Ecm1 | 0.89 | 0.00 | 0.00 | 0.27 | 0.05 | 7 | Endothelial |
| Add3 | 0.91 | 0.00 | 0.00 | 0.64 | 0.27 | 7 | Endothelial |
| Ushbp1 | 0.80 | 0.00 | 0.00 | 0.34 | 0.08 | 7 | Endothelial |
| Rasal2 | 0.76 | 0.00 | 0.00 | 0.26 | 0.05 | 7 | Endothelial |
| Nkd1 | 0.91 | 0.00 | 0.00 | 0.38 | 0.10 | 7 | Endothelial |
| Anxa3 | 0.88 | 0.00 | 0.00 | 0.57 | 0.21 | 7 | Endothelial |
| Fermt2 | 0.84 | 0.00 | 0.00 | 0.75 | 0.38 | 7 | Endothelial |
| Nes | 0.84 | 0.00 | 0.00 | 0.57 | 0.20 | 7 | Endothelial |
| Tbx2 | 0.91 | 0.00 | 0.00 | 0.39 | 0.11 | 7 | Endothelial |
| Cnn3 | 0.84 | 0.00 | 0.00 | 0.66 | 0.29 | 7 | Endothelial |
| Bpgm | 0.88 | 0.00 | 0.00 | 0.28 | 0.06 | 7 | Endothelial |
| Rhoj | 0.85 | 0.00 | 0.00 | 0.51 | 0.17 | 7 | Endothelial |
| Acbd5 | 0.92 | 0.00 | 0.00 | 0.43 | 0.13 | 7 | Endothelial |
| Sema3c | 0.86 | 0.00 | 0.00 | 0.53 | 0.19 | 7 | Endothelial |
| Btbd3 | 0.91 | 0.00 | 0.00 | 0.44 | 0.14 | 7 | Endothelial |
| Cd151 | 0.88 | 0.00 | 0.00 | 0.46 | 0.15 | 7 | Endothelial |
| Crip2 | 0.79 | 0.00 | 0.00 | 0.77 | 0.40 | 7 | Endothelial |
| Snrk | 0.87 | 0.00 | 0.00 | 0.46 | 0.15 | 7 | Endothelial |
| Lyn | 0.82 | 0.00 | 0.00 | 0.41 | 0.12 | 7 | Endothelial |
| Pea15a | 0.86 | 0.00 | 0.00 | 0.42 | 0.13 | 7 | Endothelial |
| Arhgef15 | 0.72 | 0.00 | 0.00 | 0.29 | 0.07 | 7 | Endothelial |
| Ccdc85b | 0.80 | 0.00 | 0.00 | 0.38 | 0.11 | 7 | Endothelial |
| Slfn5 | 0.74 | 0.00 | 0.00 | 0.34 | 0.09 | 7 | Endothelial |
| Cd38 | 0.72 | 0.00 | 0.00 | 0.25 | 0.05 | 7 | Endothelial |
| Abcg1 | 0.72 | 0.00 | 0.00 | 0.26 | 0.06 | 7 | Endothelial |
| Mxra7 | 0.82 | 0.00 | 0.00 | 0.42 | 0.13 | 7 | Endothelial |
| Ptprg | 0.83 | 0.00 | 0.00 | 0.39 | 0.12 | 7 | Endothelial |
| Kank3 | 0.80 | 0.00 | 0.00 | 0.42 | 0.13 | 7 | Endothelial |
| Arap3 | 0.74 | 0.00 | 0.00 | 0.29 | 0.07 | 7 | Endothelial |
| Cd200 | 0.64 | 0.00 | 0.00 | 0.62 | 0.26 | 7 | Endothelial |
| Rcsd1 | 0.77 | 0.00 | 0.00 | 0.30 | 0.08 | 7 | Endothelial |
| Tie1 | 0.69 | 0.00 | 0.00 | 0.37 | 0.11 | 7 | Endothelial |
| Ly6c2 | 0.74 | 0.00 | 0.00 | 0.43 | 0.14 | 7 | Endothelial |
| Cxx1b | 0.76 | 0.00 | 0.00 | 0.53 | 0.21 | 7 | Endothelial |
| Arrdc3 | 0.83 | 0.00 | 0.00 | 0.36 | 0.11 | 7 | Endothelial |
| Aqp1 | 0.67 | 0.00 | 0.00 | 0.51 | 0.19 | 7 | Endothelial |
| Ets1 | 0.76 | 0.00 | 0.00 | 0.57 | 0.24 | 7 | Endothelial |
| Adcy4 | 0.73 | 0.00 | 0.00 | 0.30 | 0.08 | 7 | Endothelial |
| Gng11 | 0.74 | 0.00 | 0.00 | 0.62 | 0.28 | 7 | Endothelial |
| Pde4b | 0.80 | 0.00 | 0.00 | 0.41 | 0.13 | 7 | Endothelial |
| Ppap2b | 0.77 | 0.00 | 0.00 | 0.55 | 0.23 | 7 | Endothelial |
| Syn3 | 0.94 | 0.00 | 0.00 | 0.42 | 0.15 | 7 | Endothelial |
| Tjp1 | 0.81 | 0.00 | 0.00 | 0.43 | 0.15 | 7 | Endothelial |
| Cxx1a | 0.73 | 0.00 | 0.00 | 0.53 | 0.21 | 7 | Endothelial |
| Vim | 0.58 | 0.00 | 0.00 | 0.92 | 0.65 | 7 | Endothelial |
| Casz1 | 0.74 | 0.00 | 0.00 | 0.40 | 0.13 | 7 | Endothelial |
| Smarca2 | 0.78 | 0.00 | 0.00 | 0.65 | 0.33 | 7 | Endothelial |
| Sgk1 | 0.73 | 0.00 | 0.00 | 0.48 | 0.18 | 7 | Endothelial |
| Srgn | 0.60 | 0.00 | 0.00 | 0.68 | 0.32 | 7 | Endothelial |
| Ece1 | 0.71 | 0.00 | 0.00 | 0.57 | 0.24 | 7 | Endothelial |
| Clic4 | 0.84 | 0.00 | 0.00 | 0.59 | 0.28 | 7 | Endothelial |
| Rell1 | 0.83 | 0.00 | 0.00 | 0.49 | 0.19 | 7 | Endothelial |
| 2810025M15Rik | 0.73 | 0.00 | 0.00 | 0.37 | 0.12 | 7 | Endothelial |
| Sox17 | 0.63 | 0.00 | 0.00 | 0.40 | 0.13 | 7 | Endothelial |
| Tpst2 | 0.78 | 0.00 | 0.00 | 0.35 | 0.11 | 7 | Endothelial |
| Klf2 | 0.78 | 0.00 | 0.00 | 0.49 | 0.19 | 7 | Endothelial |
| Rasgrp3 | 0.65 | 0.00 | 0.00 | 0.38 | 0.12 | 7 | Endothelial |
| Ceacam1 | 0.71 | 0.00 | 0.00 | 0.31 | 0.09 | 7 | Endothelial |
| Bicd2 | 0.73 | 0.00 | 0.00 | 0.28 | 0.08 | 7 | Endothelial |
| Calm1 | 0.43 | 0.00 | 0.00 | 0.97 | 0.86 | 7 | Endothelial |
| Mef2a | 0.74 | 0.00 | 0.00 | 0.65 | 0.35 | 7 | Endothelial |
| Ptprm | 0.65 | 0.00 | 0.00 | 0.31 | 0.09 | 7 | Endothelial |
| Itm2b | 0.53 | 0.00 | 0.00 | 0.90 | 0.75 | 7 | Endothelial |
| Serpinh1 | 0.67 | 0.00 | 0.00 | 0.70 | 0.38 | 7 | Endothelial |
| Cd97 | 0.73 | 0.00 | 0.00 | 0.45 | 0.17 | 7 | Endothelial |
| Hes1 | 0.78 | 0.00 | 0.00 | 0.58 | 0.28 | 7 | Endothelial |
| B2m | 0.50 | 0.00 | 0.00 | 0.90 | 0.67 | 7 | Endothelial |
| Tmem140 | 0.68 | 0.00 | 0.00 | 0.26 | 0.07 | 7 | Endothelial |
| Snx3 | 0.73 | 0.00 | 0.00 | 0.61 | 0.32 | 7 | Endothelial |
| Zeb1 | 0.72 | 0.00 | 0.00 | 0.33 | 0.11 | 7 | Endothelial |
| Lyve1 | 0.41 | 0.00 | 0.00 | 0.53 | 0.21 | 7 | Endothelial |
| Tubb2b | 0.68 | 0.00 | 0.00 | 0.36 | 0.12 | 7 | Endothelial |
| Nid1 | 0.74 | 0.00 | 0.00 | 0.51 | 0.22 | 7 | Endothelial |
| Prkce | 0.70 | 0.00 | 0.00 | 0.26 | 0.07 | 7 | Endothelial |
| Ddah2 | 0.69 | 0.00 | 0.00 | 0.54 | 0.25 | 7 | Endothelial |
| Skil | 0.75 | 0.00 | 0.00 | 0.37 | 0.14 | 7 | Endothelial |
| Galnt18 | 0.70 | 0.00 | 0.00 | 0.32 | 0.10 | 7 | Endothelial |
| Qk | 0.73 | 0.00 | 0.00 | 0.57 | 0.29 | 7 | Endothelial |
| Acap2 | 0.79 | 0.00 | 0.00 | 0.45 | 0.19 | 7 | Endothelial |
| St3gal6 | 0.67 | 0.00 | 0.00 | 0.28 | 0.09 | 7 | Endothelial |
| Tuba1a | 0.61 | 0.00 | 0.00 | 0.78 | 0.50 | 7 | Endothelial |
| Hilpda | 0.67 | 0.00 | 0.00 | 0.27 | 0.08 | 7 | Endothelial |
| Abi1 | 0.74 | 0.00 | 0.00 | 0.48 | 0.21 | 7 | Endothelial |
| Dlc1 | 0.72 | 0.00 | 0.00 | 0.42 | 0.17 | 7 | Endothelial |
| Clec14a | 0.53 | 0.00 | 0.00 | 0.44 | 0.17 | 7 | Endothelial |
| Dpp4 | 0.71 | 0.00 | 0.00 | 0.34 | 0.12 | 7 | Endothelial |
| Cxx1c | 0.66 | 0.00 | 0.00 | 0.48 | 0.21 | 7 | Endothelial |
| Gimap1 | 0.68 | 0.00 | 0.00 | 0.36 | 0.13 | 7 | Endothelial |
| Rhoc | 0.72 | 0.00 | 0.00 | 0.45 | 0.20 | 7 | Endothelial |
| Cltb | 0.70 | 0.00 | 0.00 | 0.31 | 0.11 | 7 | Endothelial |
| Fas | 0.71 | 0.00 | 0.00 | 0.29 | 0.09 | 7 | Endothelial |
| Itgb1 | 0.52 | 0.00 | 0.00 | 0.85 | 0.62 | 7 | Endothelial |
| Arhgap31 | 0.69 | 0.00 | 0.00 | 0.32 | 0.11 | 7 | Endothelial |
| Klhl5 | 0.66 | 0.00 | 0.00 | 0.36 | 0.14 | 7 | Endothelial |
| Rap1a | 0.70 | 0.00 | 0.00 | 0.56 | 0.29 | 7 | Endothelial |
| Mgst3 | 0.69 | 0.00 | 0.00 | 0.28 | 0.09 | 7 | Endothelial |
| Erg | 0.60 | 0.00 | 0.00 | 0.35 | 0.13 | 7 | Endothelial |
| Rasgef1a | 0.58 | 0.00 | 0.00 | 0.26 | 0.08 | 7 | Endothelial |
| Map7d1 | 0.70 | 0.00 | 0.00 | 0.41 | 0.17 | 7 | Endothelial |
| Hip1 | 0.63 | 0.00 | 0.00 | 0.38 | 0.15 | 7 | Endothelial |
| Myo10 | 0.64 | 0.00 | 0.00 | 0.33 | 0.12 | 7 | Endothelial |
| Gngt2 | 0.55 | 0.00 | 0.00 | 0.29 | 0.10 | 7 | Endothelial |
| Ldb2 | 0.54 | 0.00 | 0.00 | 0.39 | 0.16 | 7 | Endothelial |
| Cmtm3 | 0.62 | 0.00 | 0.00 | 0.33 | 0.13 | 7 | Endothelial |
| Rdx | 0.60 | 0.00 | 0.00 | 0.68 | 0.45 | 7 | Endothelial |
| Elmo1 | 0.70 | 0.00 | 0.00 | 0.30 | 0.11 | 7 | Endothelial |
| Bin1 | 0.65 | 0.00 | 0.00 | 0.29 | 0.10 | 7 | Endothelial |
| Myl12a | 0.50 | 0.00 | 0.00 | 0.81 | 0.65 | 7 | Endothelial |
| Grap | 0.57 | 0.00 | 0.00 | 0.34 | 0.13 | 7 | Endothelial |
| Nfic | 0.66 | 0.00 | 0.00 | 0.47 | 0.24 | 7 | Endothelial |
| Myl12b | 0.53 | 0.00 | 0.00 | 0.76 | 0.60 | 7 | Endothelial |
| Gpihbp1 | 0.49 | 0.00 | 0.00 | 0.39 | 0.16 | 7 | Endothelial |
| Rab11a | 0.63 | 0.00 | 0.00 | 0.49 | 0.25 | 7 | Endothelial |
| Bst2 | 0.62 | 0.00 | 0.00 | 0.34 | 0.13 | 7 | Endothelial |
| Oraov1 | 0.60 | 0.00 | 0.00 | 0.25 | 0.09 | 7 | Endothelial |
| Efnb2 | 0.52 | 0.00 | 0.00 | 0.35 | 0.14 | 7 | Endothelial |
| Pam | 0.59 | 0.00 | 0.00 | 0.38 | 0.17 | 7 | Endothelial |
| Eng | 0.60 | 0.00 | 0.00 | 0.29 | 0.11 | 7 | Endothelial |
| Gnai2 | 0.58 | 0.00 | 0.00 | 0.62 | 0.38 | 7 | Endothelial |
| Pxdn | 0.58 | 0.00 | 0.00 | 0.28 | 0.10 | 7 | Endothelial |
| Vcl | 0.72 | 0.00 | 0.00 | 0.31 | 0.12 | 7 | Endothelial |
| Tmem88 | 0.52 | 0.00 | 0.00 | 0.28 | 0.10 | 7 | Endothelial |
| Cav1 | 0.41 | 0.00 | 0.00 | 0.59 | 0.31 | 7 | Endothelial |
| Cd47 | 0.61 | 0.00 | 0.00 | 0.66 | 0.42 | 7 | Endothelial |
| Notch1 | 0.55 | 0.00 | 0.00 | 0.32 | 0.13 | 7 | Endothelial |
| Reep3 | 0.63 | 0.00 | 0.00 | 0.39 | 0.18 | 7 | Endothelial |
| Gcc2 | 0.64 | 0.00 | 0.00 | 0.47 | 0.24 | 7 | Endothelial |
| Dynll1 | 0.52 | 0.00 | 0.00 | 0.67 | 0.44 | 7 | Endothelial |
| Cbfa2t3 | 0.49 | 0.00 | 0.00 | 0.34 | 0.14 | 7 | Endothelial |
| Zfhx3 | 0.68 | 0.00 | 0.00 | 0.36 | 0.17 | 7 | Endothelial |
| Eva1b | 0.67 | 0.00 | 0.00 | 0.40 | 0.19 | 7 | Endothelial |
| Myo18a | 0.58 | 0.00 | 0.00 | 0.27 | 0.10 | 7 | Endothelial |
| Mcam | 0.48 | 0.00 | 0.00 | 0.31 | 0.13 | 7 | Endothelial |
| Carhsp1 | 0.57 | 0.00 | 0.00 | 0.31 | 0.13 | 7 | Endothelial |
| Mocs2 | 0.64 | 0.00 | 0.00 | 0.39 | 0.19 | 7 | Endothelial |
| Sptan1 | 0.58 | 0.00 | 0.00 | 0.36 | 0.16 | 7 | Endothelial |
| Smad7 | 0.55 | 0.00 | 0.00 | 0.28 | 0.11 | 7 | Endothelial |
| Tcf4 | 0.50 | 0.00 | 0.00 | 0.64 | 0.40 | 7 | Endothelial |
| Phactr2 | 0.56 | 0.00 | 0.00 | 0.30 | 0.13 | 7 | Endothelial |
| Ctnna1 | 0.54 | 0.00 | 0.00 | 0.58 | 0.36 | 7 | Endothelial |
| Atp8a1 | 0.59 | 0.00 | 0.00 | 0.34 | 0.15 | 7 | Endothelial |
| Plcg1 | 0.59 | 0.00 | 0.00 | 0.35 | 0.16 | 7 | Endothelial |
| Ets2 | 0.57 | 0.00 | 0.00 | 0.29 | 0.12 | 7 | Endothelial |
| Dnajc8 | 0.53 | 0.00 | 0.00 | 0.63 | 0.42 | 7 | Endothelial |
| F11r | 0.52 | 0.00 | 0.00 | 0.35 | 0.16 | 7 | Endothelial |
| Cyb5r3 | 0.60 | 0.00 | 0.00 | 0.59 | 0.38 | 7 | Endothelial |
| H2-D1 | 0.38 | 0.00 | 0.00 | 0.73 | 0.49 | 7 | Endothelial |
| Arpc2 | 0.41 | 0.00 | 0.00 | 0.78 | 0.61 | 7 | Endothelial |
| Ctnnd1 | 0.60 | 0.00 | 0.00 | 0.34 | 0.16 | 7 | Endothelial |
| Yaf2 | 0.56 | 0.00 | 0.00 | 0.25 | 0.10 | 7 | Endothelial |
| Tppp3 | 0.40 | 0.00 | 0.00 | 0.29 | 0.12 | 7 | Endothelial |
| Guk1 | 0.56 | 0.00 | 0.00 | 0.33 | 0.15 | 7 | Endothelial |
| Rbms1 | 0.58 | 0.00 | 0.00 | 0.59 | 0.38 | 7 | Endothelial |
| Wasf2 | 0.55 | 0.00 | 0.00 | 0.51 | 0.30 | 7 | Endothelial |
| Aplp2 | 0.52 | 0.00 | 0.00 | 0.60 | 0.38 | 7 | Endothelial |
| Fibin | 0.73 | 0.00 | 0.00 | 0.28 | 0.12 | 7 | Endothelial |
| Srpr | 0.59 | 0.00 | 0.00 | 0.44 | 0.24 | 7 | Endothelial |
| Elk3 | 0.51 | 0.00 | 0.00 | 0.31 | 0.13 | 7 | Endothelial |
| Mtus1 | 0.60 | 0.00 | 0.00 | 0.35 | 0.17 | 7 | Endothelial |
| Tspan8 | 0.50 | 0.00 | 0.00 | 0.28 | 0.12 | 7 | Endothelial |
| Gstm1 | 0.50 | 0.00 | 0.00 | 0.32 | 0.15 | 7 | Endothelial |
| Rap1b | 0.56 | 0.00 | 0.00 | 0.42 | 0.23 | 7 | Endothelial |
| Cd93 | 0.42 | 0.00 | 0.00 | 0.43 | 0.21 | 7 | Endothelial |
| Tmem176a | 0.57 | 0.00 | 0.00 | 0.39 | 0.20 | 7 | Endothelial |
| Luzp1 | 0.52 | 0.00 | 0.00 | 0.38 | 0.19 | 7 | Endothelial |
| Laptm4a | 0.48 | 0.00 | 0.00 | 0.70 | 0.51 | 7 | Endothelial |
| Vps36 | 0.59 | 0.00 | 0.00 | 0.48 | 0.28 | 7 | Endothelial |
| Plec | 0.53 | 0.00 | 0.00 | 0.28 | 0.12 | 7 | Endothelial |
| Ccnd3 | 0.56 | 0.00 | 0.00 | 0.36 | 0.18 | 7 | Endothelial |
| Plscr3 | 0.49 | 0.00 | 0.00 | 0.28 | 0.12 | 7 | Endothelial |
| Arl2bp | 0.53 | 0.00 | 0.00 | 0.29 | 0.13 | 7 | Endothelial |
| Nras | 0.54 | 0.00 | 0.00 | 0.33 | 0.16 | 7 | Endothelial |
| Daam1 | 0.52 | 0.00 | 0.00 | 0.32 | 0.15 | 7 | Endothelial |
| Itpripl2 | 0.52 | 0.00 | 0.00 | 0.34 | 0.17 | 7 | Endothelial |
| Tmod3 | 0.54 | 0.00 | 0.00 | 0.45 | 0.26 | 7 | Endothelial |
| Bnip2 | 0.53 | 0.00 | 0.00 | 0.49 | 0.29 | 7 | Endothelial |
| Plk2 | 0.51 | 0.00 | 0.00 | 0.33 | 0.16 | 7 | Endothelial |
| Serinc1 | 0.58 | 0.00 | 0.00 | 0.47 | 0.29 | 7 | Endothelial |
| Nck1 | 0.50 | 0.00 | 0.00 | 0.28 | 0.13 | 7 | Endothelial |
| Fnbp1l | 0.48 | 0.00 | 0.00 | 0.44 | 0.25 | 7 | Endothelial |
| Cdc42 | 0.46 | 0.00 | 0.00 | 0.69 | 0.54 | 7 | Endothelial |
| Pon2 | 0.51 | 0.00 | 0.00 | 0.30 | 0.14 | 7 | Endothelial |
| Msn | 0.45 | 0.00 | 0.00 | 0.62 | 0.43 | 7 | Endothelial |
| Hook3 | 0.54 | 0.00 | 0.00 | 0.40 | 0.22 | 7 | Endothelial |
| Tmem66 | 0.54 | 0.00 | 0.00 | 0.36 | 0.20 | 7 | Endothelial |
| Tjp2 | 0.49 | 0.00 | 0.00 | 0.32 | 0.16 | 7 | Endothelial |
| Rassf1 | 0.49 | 0.00 | 0.00 | 0.29 | 0.14 | 7 | Endothelial |
| Gng2 | 0.43 | 0.00 | 0.00 | 0.26 | 0.12 | 7 | Endothelial |
| Msi2 | 0.52 | 0.00 | 0.00 | 0.33 | 0.17 | 7 | Endothelial |
| Gnaq | 0.53 | 0.00 | 0.00 | 0.38 | 0.21 | 7 | Endothelial |
| Sash1 | 0.46 | 0.00 | 0.00 | 0.27 | 0.13 | 7 | Endothelial |
| Sypl | 0.49 | 0.00 | 0.00 | 0.43 | 0.25 | 7 | Endothelial |
| Ppp2r5a | 0.50 | 0.00 | 0.00 | 0.29 | 0.14 | 7 | Endothelial |
| Mfap2 | 0.35 | 0.00 | 0.00 | 0.56 | 0.34 | 7 | Endothelial |
| Nin | 0.49 | 0.00 | 0.00 | 0.29 | 0.14 | 7 | Endothelial |
| Csnk1a1 | 0.41 | 0.00 | 0.00 | 0.69 | 0.53 | 7 | Endothelial |
| Wwc2 | 0.48 | 0.00 | 0.00 | 0.27 | 0.13 | 7 | Endothelial |
| Fcho2 | 0.53 | 0.00 | 0.00 | 0.35 | 0.19 | 7 | Endothelial |
| Tmco1 | 0.48 | 0.00 | 0.00 | 0.46 | 0.29 | 7 | Endothelial |
| Lmo7 | 0.27 | 0.00 | 0.00 | 0.27 | 0.12 | 7 | Endothelial |
| Klc1 | 0.47 | 0.00 | 0.00 | 0.29 | 0.14 | 7 | Endothelial |
| Anp32a | 0.33 | 0.00 | 0.00 | 0.73 | 0.60 | 7 | Endothelial |
| Rac1 | 0.48 | 0.00 | 0.00 | 0.46 | 0.30 | 7 | Endothelial |
| Ap1s2 | 0.38 | 0.00 | 0.00 | 0.25 | 0.12 | 7 | Endothelial |
| Zmiz1 | 0.42 | 0.00 | 0.00 | 0.38 | 0.21 | 7 | Endothelial |
| Slc43a3 | 0.38 | 0.00 | 0.00 | 0.38 | 0.21 | 7 | Endothelial |
| Heg1 | 0.37 | 0.00 | 0.00 | 0.31 | 0.16 | 7 | Endothelial |
| Cyfip1 | 0.45 | 0.00 | 0.00 | 0.30 | 0.16 | 7 | Endothelial |
| Ywhaz | 0.40 | 0.00 | 0.00 | 0.65 | 0.51 | 7 | Endothelial |
| Arf1 | 0.42 | 0.00 | 0.00 | 0.58 | 0.42 | 7 | Endothelial |
| Ywhab | 0.33 | 0.00 | 0.00 | 0.70 | 0.56 | 7 | Endothelial |
| Rhoa | 0.41 | 0.00 | 0.00 | 0.61 | 0.46 | 7 | Endothelial |
| Palmd | 0.34 | 0.00 | 0.00 | 0.26 | 0.13 | 7 | Endothelial |
| Ier2 | 0.41 | 0.00 | 0.00 | 0.41 | 0.25 | 7 | Endothelial |
| B230219D22Rik | 0.44 | 0.00 | 0.00 | 0.43 | 0.27 | 7 | Endothelial |
| Tmem176b | 0.43 | 0.00 | 0.00 | 0.50 | 0.34 | 7 | Endothelial |
| Ubb | 0.26 | 0.00 | 0.00 | 0.87 | 0.81 | 7 | Endothelial |
| Zcrb1 | 0.45 | 0.00 | 0.00 | 0.44 | 0.29 | 7 | Endothelial |
| Kif1b | 0.48 | 0.00 | 0.00 | 0.38 | 0.23 | 7 | Endothelial |
| Hdac7 | 0.42 | 0.00 | 0.00 | 0.26 | 0.14 | 7 | Endothelial |
| Eif4g2 | 0.43 | 0.00 | 0.00 | 0.55 | 0.41 | 7 | Endothelial |
| Mapk3 | 0.43 | 0.00 | 0.00 | 0.26 | 0.13 | 7 | Endothelial |
| Efnb1 | 0.41 | 0.00 | 0.00 | 0.28 | 0.15 | 7 | Endothelial |
| Lamp1 | 0.42 | 0.00 | 0.00 | 0.57 | 0.42 | 7 | Endothelial |
| Sdcbp | 0.42 | 0.00 | 0.00 | 0.43 | 0.27 | 7 | Endothelial |
| Itsn2 | 0.43 | 0.00 | 0.00 | 0.39 | 0.24 | 7 | Endothelial |
| Fxyd5 | 0.34 | 0.00 | 0.00 | 0.32 | 0.18 | 7 | Endothelial |
| Ctsl | 0.34 | 0.00 | 0.00 | 0.56 | 0.39 | 7 | Endothelial |
| Mxd4 | 0.38 | 0.00 | 0.00 | 0.53 | 0.37 | 7 | Endothelial |
| Mbnl2 | 0.39 | 0.00 | 0.00 | 0.50 | 0.34 | 7 | Endothelial |
| Rpn1 | 0.44 | 0.00 | 0.00 | 0.34 | 0.21 | 7 | Endothelial |
| Lgals9 | 0.34 | 0.00 | 0.00 | 0.27 | 0.14 | 7 | Endothelial |
| Eid1 | 0.41 | 0.00 | 0.00 | 0.44 | 0.29 | 7 | Endothelial |
| Gna13 | 0.47 | 0.00 | 0.00 | 0.26 | 0.14 | 7 | Endothelial |
| Leprotl1 | 0.41 | 0.00 | 0.00 | 0.27 | 0.15 | 7 | Endothelial |
| Nap1l4 | 0.41 | 0.00 | 0.00 | 0.40 | 0.26 | 7 | Endothelial |
| Zfp706 | 0.44 | 0.00 | 0.00 | 0.46 | 0.32 | 7 | Endothelial |
| Rock2 | 0.42 | 0.00 | 0.00 | 0.47 | 0.32 | 7 | Endothelial |
| Hpcal1 | 0.40 | 0.00 | 0.00 | 0.27 | 0.15 | 7 | Endothelial |
| Tubb5 | 0.28 | 0.00 | 0.00 | 0.61 | 0.44 | 7 | Endothelial |
| Epb4.1l2 | 0.40 | 0.00 | 0.00 | 0.32 | 0.19 | 7 | Endothelial |
| Zbtb20 | 0.40 | 0.00 | 0.00 | 0.41 | 0.26 | 7 | Endothelial |
| Dram2 | 0.40 | 0.00 | 0.00 | 0.29 | 0.16 | 7 | Endothelial |
| Myh9 | 0.41 | 0.00 | 0.00 | 0.47 | 0.33 | 7 | Endothelial |
| Dpysl2 | 0.42 | 0.00 | 0.00 | 0.30 | 0.17 | 7 | Endothelial |
| Tpm4 | 0.31 | 0.00 | 0.00 | 0.62 | 0.49 | 7 | Endothelial |
| Ccser2 | 0.36 | 0.00 | 0.00 | 0.26 | 0.14 | 7 | Endothelial |
| Leprot | 0.38 | 0.00 | 0.00 | 0.47 | 0.32 | 7 | Endothelial |
| Fubp1 | 0.38 | 0.00 | 0.00 | 0.38 | 0.24 | 7 | Endothelial |
| Rab10 | 0.42 | 0.00 | 0.00 | 0.38 | 0.24 | 7 | Endothelial |
| Oaz2 | 0.40 | 0.00 | 0.00 | 0.42 | 0.28 | 7 | Endothelial |
| Elf2 | 0.43 | 0.00 | 0.00 | 0.30 | 0.18 | 7 | Endothelial |
| Myadm | 0.44 | 0.00 | 0.00 | 0.32 | 0.20 | 7 | Endothelial |
| Pak1ip1 | 0.38 | 0.00 | 0.00 | 0.26 | 0.14 | 7 | Endothelial |
| Dusp6 | 0.35 | 0.00 | 0.00 | 0.31 | 0.18 | 7 | Endothelial |
| Pcdhga9 | 0.36 | 0.00 | 0.00 | 0.30 | 0.17 | 7 | Endothelial |
| Shisa5 | 0.30 | 0.00 | 0.00 | 0.34 | 0.20 | 7 | Endothelial |
| Tmem59 | 0.38 | 0.00 | 0.00 | 0.53 | 0.40 | 7 | Endothelial |
| Twf1 | 0.39 | 0.00 | 0.00 | 0.27 | 0.16 | 7 | Endothelial |
| Tnfaip1 | 0.39 | 0.00 | 0.00 | 0.28 | 0.16 | 7 | Endothelial |
| Mtdh | 0.32 | 0.00 | 0.00 | 0.61 | 0.48 | 7 | Endothelial |
| Rab5a | 0.40 | 0.00 | 0.00 | 0.25 | 0.15 | 7 | Endothelial |
| Xiap | 0.39 | 0.00 | 0.00 | 0.32 | 0.20 | 7 | Endothelial |
| 1110008F13Rik | 0.43 | 0.00 | 0.00 | 0.27 | 0.16 | 7 | Endothelial |
| Clic1 | 0.35 | 0.00 | 0.00 | 0.51 | 0.38 | 7 | Endothelial |
| Tmem50a | 0.32 | 0.00 | 0.00 | 0.57 | 0.45 | 7 | Endothelial |
| Tln1 | 0.33 | 0.00 | 0.00 | 0.52 | 0.39 | 7 | Endothelial |
| Selk | 0.37 | 0.00 | 0.00 | 0.44 | 0.31 | 7 | Endothelial |
| Serinc3 | 0.30 | 0.00 | 0.00 | 0.45 | 0.32 | 7 | Endothelial |
| Ppp1ca | 0.37 | 0.00 | 0.00 | 0.45 | 0.33 | 7 | Endothelial |
| Rap2a | 0.31 | 0.00 | 0.00 | 0.27 | 0.15 | 7 | Endothelial |
| 1700020I14Rik | 0.35 | 0.00 | 0.00 | 0.34 | 0.22 | 7 | Endothelial |
| Cdkn1a | 0.36 | 0.00 | 0.00 | 0.27 | 0.16 | 7 | Endothelial |
| Crk | 0.38 | 0.00 | 0.00 | 0.27 | 0.16 | 7 | Endothelial |
| Lhfp | 0.26 | 0.00 | 0.00 | 0.26 | 0.15 | 7 | Endothelial |
| Timp2 | 0.35 | 0.00 | 0.00 | 0.32 | 0.21 | 7 | Endothelial |
| Psmd12 | 0.34 | 0.00 | 0.00 | 0.34 | 0.23 | 7 | Endothelial |
| Gnb2 | 0.37 | 0.00 | 0.00 | 0.49 | 0.37 | 7 | Endothelial |
| Rtn4 | 0.34 | 0.00 | 0.00 | 0.35 | 0.23 | 7 | Endothelial |
| Rab6a | 0.33 | 0.00 | 0.00 | 0.32 | 0.21 | 7 | Endothelial |
| Ablim1 | 0.27 | 0.00 | 0.00 | 0.37 | 0.25 | 7 | Endothelial |
| Pttg1ip | 0.37 | 0.00 | 0.00 | 0.27 | 0.17 | 7 | Endothelial |
| Tsg101 | 0.35 | 0.00 | 0.00 | 0.26 | 0.16 | 7 | Endothelial |
| Tuba1c | 0.27 | 0.00 | 0.00 | 0.45 | 0.32 | 7 | Endothelial |
| Pak2 | 0.34 | 0.00 | 0.00 | 0.41 | 0.29 | 7 | Endothelial |
| Rala | 0.34 | 0.00 | 0.00 | 0.34 | 0.23 | 7 | Endothelial |
| Atxn7l3b | 0.32 | 0.00 | 0.00 | 0.51 | 0.40 | 7 | Endothelial |
| Chmp5 | 0.35 | 0.00 | 0.00 | 0.34 | 0.24 | 7 | Endothelial |
| Tmed10 | 0.38 | 0.00 | 0.00 | 0.46 | 0.36 | 7 | Endothelial |
| Dhrs7 | 0.35 | 0.00 | 0.00 | 0.25 | 0.16 | 7 | Endothelial |
| Cggbp1 | 0.37 | 0.00 | 0.00 | 0.27 | 0.18 | 7 | Endothelial |
| Pomp | 0.30 | 0.00 | 0.00 | 0.43 | 0.31 | 7 | Endothelial |
| Dnaja1 | 0.26 | 0.00 | 0.00 | 0.62 | 0.51 | 7 | Endothelial |
| Map1lc3a | 0.37 | 0.00 | 0.00 | 0.30 | 0.20 | 7 | Endothelial |
| Hmg20b | 0.32 | 0.00 | 0.00 | 0.45 | 0.34 | 7 | Endothelial |
| Prkar1a | 0.31 | 0.00 | 0.00 | 0.43 | 0.32 | 7 | Endothelial |
| Pdcd10 | 0.33 | 0.00 | 0.00 | 0.31 | 0.21 | 7 | Endothelial |
| Hnrnpf | 0.29 | 0.00 | 0.00 | 0.45 | 0.34 | 7 | Endothelial |
| Rab2a | 0.33 | 0.00 | 0.00 | 0.37 | 0.27 | 7 | Endothelial |
| Amotl1 | 0.28 | 0.00 | 0.00 | 0.25 | 0.16 | 7 | Endothelial |
| Pls3 | 0.31 | 0.00 | 0.00 | 0.30 | 0.20 | 7 | Endothelial |
| Cnih1 | 0.33 | 0.00 | 0.00 | 0.28 | 0.18 | 7 | Endothelial |
| Cd2ap | 0.27 | 0.00 | 0.00 | 0.45 | 0.34 | 7 | Endothelial |
| Dync1i2 | 0.27 | 0.00 | 0.00 | 0.57 | 0.48 | 7 | Endothelial |
| Shoc2 | 0.30 | 0.00 | 0.00 | 0.30 | 0.20 | 7 | Endothelial |
| Gsk3b | 0.31 | 0.00 | 0.00 | 0.38 | 0.27 | 7 | Endothelial |
| Tmem30a | 0.32 | 0.00 | 0.00 | 0.38 | 0.28 | 7 | Endothelial |
| Sbds | 0.30 | 0.00 | 0.00 | 0.32 | 0.22 | 7 | Endothelial |
| Phf20l1 | 0.28 | 0.00 | 0.00 | 0.39 | 0.28 | 7 | Endothelial |
| Vps35 | 0.30 | 0.00 | 0.00 | 0.27 | 0.18 | 7 | Endothelial |
| Gnb1 | 0.31 | 0.00 | 0.00 | 0.31 | 0.22 | 7 | Endothelial |
| Tmbim6 | 0.28 | 0.00 | 0.00 | 0.47 | 0.38 | 7 | Endothelial |
| Rab1 | 0.29 | 0.00 | 0.00 | 0.40 | 0.31 | 7 | Endothelial |
| 2700089E24Rik | 0.29 | 0.00 | 0.00 | 0.32 | 0.23 | 7 | Endothelial |
| Wasl | 0.27 | 0.00 | 0.00 | 0.27 | 0.18 | 7 | Endothelial |
| Ginm1 | 0.30 | 0.00 | 0.00 | 0.33 | 0.23 | 7 | Endothelial |
| Sec62 | 0.27 | 0.00 | 0.00 | 0.54 | 0.45 | 7 | Endothelial |
| Mapre1 | 0.28 | 0.00 | 0.00 | 0.39 | 0.29 | 7 | Endothelial |
| Irf2 | 0.31 | 0.00 | 0.00 | 0.33 | 0.24 | 7 | Endothelial |
| Ddx3x | 0.26 | 0.00 | 0.00 | 0.39 | 0.30 | 7 | Endothelial |
| Sh3glb1 | 0.25 | 0.00 | 0.00 | 0.49 | 0.40 | 7 | Endothelial |
| Prpf4b | 0.26 | 0.00 | 0.00 | 0.54 | 0.46 | 7 | Endothelial |
| Eif4g3 | 0.29 | 0.00 | 0.00 | 0.36 | 0.27 | 7 | Endothelial |
| Ccdc47 | 0.35 | 0.00 | 0.00 | 0.29 | 0.21 | 7 | Endothelial |
| Dcun1d5 | 0.31 | 0.00 | 0.00 | 0.25 | 0.17 | 7 | Endothelial |
| G3bp2 | 0.26 | 0.00 | 0.00 | 0.40 | 0.31 | 7 | Endothelial |
| Sri | 0.28 | 0.00 | 0.00 | 0.37 | 0.27 | 7 | Endothelial |
| Snrpb | 0.27 | 0.00 | 0.00 | 0.44 | 0.35 | 7 | Endothelial |
| Zfand5 | 0.29 | 0.00 | 0.00 | 0.34 | 0.26 | 7 | Endothelial |
| Csnk2a1 | 0.28 | 0.00 | 0.00 | 0.39 | 0.31 | 7 | Endothelial |
| Dusp11 | 0.28 | 0.00 | 0.00 | 0.34 | 0.25 | 7 | Endothelial |
| Rab7 | 0.30 | 0.00 | 0.00 | 0.33 | 0.25 | 7 | Endothelial |
| Gabarapl2 | 0.26 | 0.00 | 0.00 | 0.30 | 0.22 | 7 | Endothelial |
| Psmd7 | 0.27 | 0.00 | 0.00 | 0.37 | 0.28 | 7 | Endothelial |
| Pafah1b1 | 0.26 | 0.00 | 0.00 | 0.42 | 0.33 | 7 | Endothelial |
| Ube2a | 0.29 | 0.00 | 0.00 | 0.26 | 0.18 | 7 | Endothelial |
| Rtn3 | 0.28 | 0.00 | 0.00 | 0.34 | 0.26 | 7 | Endothelial |
| Rsu1 | 0.25 | 0.00 | 0.00 | 0.28 | 0.20 | 7 | Endothelial |
| Psmd6 | 0.27 | 0.00 | 0.00 | 0.27 | 0.19 | 7 | Endothelial |
| Ppdpf | 0.27 | 0.00 | 0.00 | 0.26 | 0.18 | 7 | Endothelial |
| Lman1 | 0.28 | 0.00 | 0.00 | 0.26 | 0.18 | 7 | Endothelial |
| Emc7 | 0.26 | 0.00 | 0.00 | 0.29 | 0.21 | 7 | Endothelial |
| Atp6v1a | 0.27 | 0.00 | 0.00 | 0.36 | 0.28 | 7 | Endothelial |
| Atf4 | 0.25 | 0.00 | 0.00 | 0.36 | 0.28 | 7 | Endothelial |
| Adam10 | 0.28 | 0.00 | 0.00 | 0.26 | 0.19 | 7 | Endothelial |
| Ociad1 | 0.26 | 0.00 | 0.00 | 0.39 | 0.31 | 7 | Endothelial |
| Taok1 | 0.25 | 0.00 | 0.00 | 0.26 | 0.19 | 7 | Endothelial |
| Pcbp1 | 0.26 | 0.00 | 0.00 | 0.29 | 0.22 | 7 | Endothelial |
| Lyve1 | 2.37 | 0.00 | 0.00 | 0.92 | 0.20 | 8 | Endothelial |
| Cd200 | 2.11 | 0.00 | 0.00 | 0.94 | 0.25 | 8 | Endothelial |
| Cdkn1a | 1.95 | 0.00 | 0.00 | 0.80 | 0.14 | 8 | Endothelial |
| Ly6c1 | 1.92 | 0.00 | 0.00 | 0.83 | 0.17 | 8 | Endothelial |
| Ramp2 | 1.84 | 0.00 | 0.00 | 0.98 | 0.26 | 8 | Endothelial |
| Cxx1c | 1.68 | 0.00 | 0.00 | 0.84 | 0.20 | 8 | Endothelial |
| Cxx1a | 1.66 | 0.00 | 0.00 | 0.84 | 0.20 | 8 | Endothelial |
| Ly6a | 1.66 | 0.00 | 0.00 | 0.81 | 0.17 | 8 | Endothelial |
| Lpl | 1.65 | 0.00 | 0.00 | 0.68 | 0.13 | 8 | Endothelial |
| Ednrb | 1.63 | 0.00 | 0.00 | 0.73 | 0.13 | 8 | Endothelial |
| Cxx1b | 1.60 | 0.00 | 0.00 | 0.82 | 0.20 | 8 | Endothelial |
| Klf4 | 1.60 | 0.00 | 0.00 | 0.80 | 0.21 | 8 | Endothelial |
| Aqp1 | 1.53 | 0.00 | 0.00 | 0.79 | 0.17 | 8 | Endothelial |
| Gata2 | 1.47 | 0.00 | 0.00 | 0.72 | 0.12 | 8 | Endothelial |
| Cav1 | 1.45 | 0.00 | 0.00 | 0.91 | 0.29 | 8 | Endothelial |
| Marcks | 1.44 | 0.00 | 0.00 | 0.95 | 0.49 | 8 | Endothelial |
| Klf2 | 1.39 | 0.00 | 0.00 | 0.72 | 0.18 | 8 | Endothelial |
| Tspan7 | 1.39 | 0.00 | 0.00 | 0.83 | 0.19 | 8 | Endothelial |
| Epas1 | 1.36 | 0.00 | 0.00 | 0.86 | 0.23 | 8 | Endothelial |
| Btbd3 | 1.35 | 0.00 | 0.00 | 0.64 | 0.13 | 8 | Endothelial |
| Ly6c2 | 1.34 | 0.00 | 0.00 | 0.69 | 0.13 | 8 | Endothelial |
| Slc43a3 | 1.33 | 0.00 | 0.00 | 0.76 | 0.20 | 8 | Endothelial |
| Serinc3 | 1.32 | 0.00 | 0.00 | 0.82 | 0.30 | 8 | Endothelial |
| Plcb4 | 1.27 | 0.00 | 0.00 | 0.59 | 0.13 | 8 | Endothelial |
| Slc9a3r2 | 1.25 | 0.00 | 0.00 | 0.76 | 0.19 | 8 | Endothelial |
| Cldn5 | 1.25 | 0.00 | 0.00 | 0.87 | 0.20 | 8 | Endothelial |
| Myzap | 1.23 | 0.00 | 0.00 | 0.66 | 0.13 | 8 | Endothelial |
| Clec1a | 1.23 | 0.00 | 0.00 | 0.71 | 0.15 | 8 | Endothelial |
| Gpr116 | 1.20 | 0.00 | 0.00 | 0.94 | 0.36 | 8 | Endothelial |
| Cd36 | 1.19 | 0.00 | 0.00 | 0.88 | 0.31 | 8 | Endothelial |
| Scarb1 | 1.18 | 0.00 | 0.00 | 0.49 | 0.08 | 8 | Endothelial |
| Fas | 1.17 | 0.00 | 0.00 | 0.54 | 0.08 | 8 | Endothelial |
| Peg3 | 1.16 | 0.00 | 0.00 | 0.73 | 0.29 | 8 | Endothelial |
| Egfl7 | 1.16 | 0.00 | 0.00 | 0.91 | 0.26 | 8 | Endothelial |
| S1pr1 | 1.13 | 0.00 | 0.00 | 0.67 | 0.17 | 8 | Endothelial |
| Sema7a | 1.13 | 0.00 | 0.00 | 0.42 | 0.03 | 8 | Endothelial |
| Thbd | 1.12 | 0.00 | 0.00 | 0.75 | 0.23 | 8 | Endothelial |
| Ybx1 | 1.11 | 0.00 | 0.00 | 0.95 | 0.63 | 8 | Endothelial |
| Nfkbia | 1.11 | 0.00 | 0.00 | 0.82 | 0.36 | 8 | Endothelial |
| Plvap | 1.11 | 0.00 | 0.00 | 0.58 | 0.11 | 8 | Endothelial |
| BC028528 | 1.10 | 0.00 | 0.00 | 0.85 | 0.27 | 8 | Endothelial |
| Mxd4 | 1.09 | 0.00 | 0.00 | 0.81 | 0.36 | 8 | Endothelial |
| Igfbp3 | 1.09 | 0.00 | 0.00 | 0.43 | 0.06 | 8 | Endothelial |
| Tcn2 | 1.07 | 0.00 | 0.00 | 0.49 | 0.10 | 8 | Endothelial |
| Clic5 | 1.07 | 0.00 | 0.00 | 0.75 | 0.21 | 8 | Endothelial |
| Gimap5 | 1.07 | 0.00 | 0.00 | 0.53 | 0.10 | 8 | Endothelial |
| Icam2 | 1.07 | 0.00 | 0.00 | 0.65 | 0.14 | 8 | Endothelial |
| Cav2 | 1.05 | 0.00 | 0.00 | 0.66 | 0.17 | 8 | Endothelial |
| Ddit4 | 1.05 | 0.00 | 0.00 | 0.42 | 0.06 | 8 | Endothelial |
| Ppp1r2 | 1.05 | 0.00 | 0.00 | 0.71 | 0.25 | 8 | Endothelial |
| Ppap2a | 1.04 | 0.00 | 0.00 | 0.66 | 0.18 | 8 | Endothelial |
| Eltd1 | 1.04 | 0.00 | 0.00 | 0.45 | 0.06 | 8 | Endothelial |
| Sult1a1 | 1.03 | 0.00 | 0.00 | 0.41 | 0.05 | 8 | Endothelial |
| Esam | 1.03 | 0.00 | 0.00 | 0.68 | 0.16 | 8 | Endothelial |
| Tmem100 | 1.02 | 0.00 | 0.00 | 0.85 | 0.28 | 8 | Endothelial |
| Timp3 | 1.02 | 0.00 | 0.00 | 0.84 | 0.34 | 8 | Endothelial |
| Itm2b | 1.02 | 0.00 | 0.00 | 0.97 | 0.75 | 8 | Endothelial |
| Pecam1 | 1.02 | 0.00 | 0.00 | 0.80 | 0.22 | 8 | Endothelial |
| Emcn | 1.02 | 0.00 | 0.00 | 0.50 | 0.10 | 8 | Endothelial |
| Acvrl1 | 1.01 | 0.00 | 0.00 | 0.66 | 0.16 | 8 | Endothelial |
| Clec14a | 1.00 | 0.00 | 0.00 | 0.67 | 0.16 | 8 | Endothelial |
| Ace | 0.98 | 0.00 | 0.00 | 0.57 | 0.12 | 8 | Endothelial |
| Cd93 | 0.98 | 0.00 | 0.00 | 0.76 | 0.20 | 8 | Endothelial |
| Sox17 | 0.97 | 0.00 | 0.00 | 0.58 | 0.12 | 8 | Endothelial |
| Cep85l | 0.96 | 0.00 | 0.00 | 0.39 | 0.06 | 8 | Endothelial |
| Ehd4 | 0.96 | 0.00 | 0.00 | 0.77 | 0.26 | 8 | Endothelial |
| Crip2 | 0.96 | 0.00 | 0.00 | 0.85 | 0.40 | 8 | Endothelial |
| Ece1 | 0.96 | 0.00 | 0.00 | 0.70 | 0.24 | 8 | Endothelial |
| Mylip | 0.94 | 0.00 | 0.00 | 0.38 | 0.06 | 8 | Endothelial |
| Prss23 | 0.92 | 0.00 | 0.00 | 0.38 | 0.06 | 8 | Endothelial |
| Foxf1 | 0.91 | 0.00 | 0.00 | 0.47 | 0.11 | 8 | Endothelial |
| Pcdh17 | 0.91 | 0.00 | 0.00 | 0.68 | 0.19 | 8 | Endothelial |
| Clca1 | 0.90 | 0.00 | 0.00 | 0.31 | 0.03 | 8 | Endothelial |
| Acer2 | 0.90 | 0.00 | 0.00 | 0.46 | 0.09 | 8 | Endothelial |
| Prx | 0.90 | 0.00 | 0.00 | 0.44 | 0.09 | 8 | Endothelial |
| Erg | 0.90 | 0.00 | 0.00 | 0.53 | 0.12 | 8 | Endothelial |
| Slfn5 | 0.89 | 0.00 | 0.00 | 0.45 | 0.09 | 8 | Endothelial |
| Gimap6 | 0.89 | 0.00 | 0.00 | 0.58 | 0.15 | 8 | Endothelial |
| Tie1 | 0.88 | 0.00 | 0.00 | 0.49 | 0.10 | 8 | Endothelial |
| Grrp1 | 0.88 | 0.00 | 0.00 | 0.34 | 0.03 | 8 | Endothelial |
| Tek | 0.87 | 0.00 | 0.00 | 0.51 | 0.12 | 8 | Endothelial |
| Ptprb | 0.86 | 0.00 | 0.00 | 0.72 | 0.20 | 8 | Endothelial |
| Rasip1 | 0.85 | 0.00 | 0.00 | 0.51 | 0.11 | 8 | Endothelial |
| Slco2a1 | 0.85 | 0.00 | 0.00 | 0.47 | 0.10 | 8 | Endothelial |
| Unc45b | 0.84 | 0.00 | 0.00 | 0.39 | 0.05 | 8 | Endothelial |
| Cdh5 | 0.82 | 0.00 | 0.00 | 0.76 | 0.21 | 8 | Endothelial |
| Zbtb16 | 0.81 | 0.00 | 0.00 | 0.34 | 0.05 | 8 | Endothelial |
| Calcrl | 0.78 | 0.00 | 0.00 | 0.75 | 0.24 | 8 | Endothelial |
| Adam15 | 0.78 | 0.00 | 0.00 | 0.37 | 0.07 | 8 | Endothelial |
| Ackr3 | 0.75 | 0.00 | 0.00 | 0.29 | 0.04 | 8 | Endothelial |
| Arl4d | 0.73 | 0.00 | 0.00 | 0.25 | 0.03 | 8 | Endothelial |
| 6430548M08Rik | 0.70 | 0.00 | 0.00 | 0.31 | 0.04 | 8 | Endothelial |
| Robo4 | 0.69 | 0.00 | 0.00 | 0.32 | 0.05 | 8 | Endothelial |
| Tsc22d3 | 0.92 | 0.00 | 0.00 | 0.68 | 0.23 | 8 | Endothelial |
| Tspan13 | 0.80 | 0.00 | 0.00 | 0.72 | 0.23 | 8 | Endothelial |
| Gpihbp1 | 0.93 | 0.00 | 0.00 | 0.55 | 0.15 | 8 | Endothelial |
| Fendrr | 0.81 | 0.00 | 0.00 | 0.63 | 0.19 | 8 | Endothelial |
| Qsox1 | 0.84 | 0.00 | 0.00 | 0.41 | 0.09 | 8 | Endothelial |
| Lgals9 | 0.90 | 0.00 | 0.00 | 0.50 | 0.13 | 8 | Endothelial |
| Flt1 | 0.86 | 0.00 | 0.00 | 0.54 | 0.14 | 8 | Endothelial |
| Mcam | 0.74 | 0.00 | 0.00 | 0.49 | 0.12 | 8 | Endothelial |
| Txnip | 0.95 | 0.00 | 0.00 | 0.68 | 0.24 | 8 | Endothelial |
| Ier3 | 0.89 | 0.00 | 0.00 | 0.48 | 0.12 | 8 | Endothelial |
| Grap | 0.86 | 0.00 | 0.00 | 0.49 | 0.12 | 8 | Endothelial |
| Ier2 | 0.96 | 0.00 | 0.00 | 0.67 | 0.24 | 8 | Endothelial |
| Ctla2a | 0.87 | 0.00 | 0.00 | 0.63 | 0.19 | 8 | Endothelial |
| Ldb2 | 0.81 | 0.00 | 0.00 | 0.55 | 0.15 | 8 | Endothelial |
| S100a16 | 0.79 | 0.00 | 0.00 | 0.67 | 0.22 | 8 | Endothelial |
| Carhsp1 | 0.84 | 0.00 | 0.00 | 0.48 | 0.13 | 8 | Endothelial |
| Tpt1 | 0.66 | 0.00 | 0.00 | 0.97 | 0.87 | 8 | Endothelial |
| Sgms1 | 0.83 | 0.00 | 0.00 | 0.37 | 0.08 | 8 | Endothelial |
| Jun | 0.89 | 0.00 | 0.00 | 0.79 | 0.35 | 8 | Endothelial |
| Cyb5r3 | 0.95 | 0.00 | 0.00 | 0.77 | 0.37 | 8 | Endothelial |
| Dok4 | 0.62 | 0.00 | 0.00 | 0.30 | 0.05 | 8 | Endothelial |
| Klhl5 | 0.78 | 0.00 | 0.00 | 0.48 | 0.13 | 8 | Endothelial |
| Ankrd33b | 0.69 | 0.00 | 0.00 | 0.29 | 0.05 | 8 | Endothelial |
| Nostrin | 0.78 | 0.00 | 0.00 | 0.50 | 0.14 | 8 | Endothelial |
| Emp1 | 0.87 | 0.00 | 0.00 | 0.61 | 0.21 | 8 | Endothelial |
| Smagp | 0.62 | 0.00 | 0.00 | 0.28 | 0.05 | 8 | Endothelial |
| Ripply3 | 0.68 | 0.00 | 0.00 | 0.46 | 0.12 | 8 | Endothelial |
| Dusp1 | 0.81 | 0.00 | 0.00 | 0.52 | 0.16 | 8 | Endothelial |
| Npr3 | 0.85 | 0.00 | 0.00 | 0.45 | 0.12 | 8 | Endothelial |
| Ptrf | 0.90 | 0.00 | 0.00 | 0.74 | 0.34 | 8 | Endothelial |
| Rps6 | 0.72 | 0.00 | 0.00 | 0.95 | 0.82 | 8 | Endothelial |
| Casz1 | 0.83 | 0.00 | 0.00 | 0.45 | 0.13 | 8 | Endothelial |
| Tcf4 | 0.91 | 0.00 | 0.00 | 0.79 | 0.40 | 8 | Endothelial |
| Hey1 | 0.68 | 0.00 | 0.00 | 0.26 | 0.04 | 8 | Endothelial |
| Foxp1 | 0.94 | 0.00 | 0.00 | 0.80 | 0.43 | 8 | Endothelial |
| Efnb2 | 0.77 | 0.00 | 0.00 | 0.48 | 0.14 | 8 | Endothelial |
| Gm26384 | 0.79 | 0.00 | 0.00 | 0.84 | 0.47 | 8 | Endothelial |
| Klf7 | 0.86 | 0.00 | 0.00 | 0.54 | 0.17 | 8 | Endothelial |
| Slk | 0.91 | 0.00 | 0.00 | 0.79 | 0.42 | 8 | Endothelial |
| Grasp | 0.67 | 0.00 | 0.00 | 0.32 | 0.07 | 8 | Endothelial |
| Tmem88 | 0.72 | 0.00 | 0.00 | 0.40 | 0.10 | 8 | Endothelial |
| Stmn2 | 0.87 | 0.00 | 0.00 | 0.49 | 0.14 | 8 | Endothelial |
| Palmd | 0.81 | 0.00 | 0.00 | 0.44 | 0.12 | 8 | Endothelial |
| Ly6e | 0.72 | 0.00 | 0.00 | 0.85 | 0.41 | 8 | Endothelial |
| Cyyr1 | 0.62 | 0.00 | 0.00 | 0.48 | 0.14 | 8 | Endothelial |
| Meis2 | 0.78 | 0.00 | 0.00 | 0.41 | 0.11 | 8 | Endothelial |
| Efcab4a | 0.61 | 0.00 | 0.00 | 0.30 | 0.06 | 8 | Endothelial |
| Fxyd5 | 0.72 | 0.00 | 0.00 | 0.52 | 0.17 | 8 | Endothelial |
| 9430020K01Rik | 0.66 | 0.00 | 0.00 | 0.43 | 0.12 | 8 | Endothelial |
| Trp53i11 | 0.62 | 0.00 | 0.00 | 0.26 | 0.05 | 8 | Endothelial |
| Ppp1r16b | 0.63 | 0.00 | 0.00 | 0.36 | 0.09 | 8 | Endothelial |
| Sepp1 | 0.86 | 0.00 | 0.00 | 0.77 | 0.40 | 8 | Endothelial |
| Kdsr | 0.69 | 0.00 | 0.00 | 0.33 | 0.08 | 8 | Endothelial |
| Plaur | 0.54 | 0.00 | 0.00 | 0.26 | 0.05 | 8 | Endothelial |
| Id1 | 0.84 | 0.00 | 0.00 | 0.59 | 0.23 | 8 | Endothelial |
| Hspb1 | 0.90 | 0.00 | 0.00 | 0.43 | 0.13 | 8 | Endothelial |
| Fam117b | 0.60 | 0.00 | 0.00 | 0.31 | 0.07 | 8 | Endothelial |
| Rhoj | 0.74 | 0.00 | 0.00 | 0.52 | 0.17 | 8 | Endothelial |
| Sash1 | 0.75 | 0.00 | 0.00 | 0.42 | 0.12 | 8 | Endothelial |
| Stt3b | 0.83 | 0.00 | 0.00 | 0.50 | 0.18 | 8 | Endothelial |
| Rdx | 0.86 | 0.00 | 0.00 | 0.79 | 0.44 | 8 | Endothelial |
| Jam2 | 0.58 | 0.00 | 0.00 | 0.27 | 0.06 | 8 | Endothelial |
| Ecscr | 0.61 | 0.00 | 0.00 | 0.51 | 0.16 | 8 | Endothelial |
| Kank3 | 0.66 | 0.00 | 0.00 | 0.45 | 0.13 | 8 | Endothelial |
| Rps27 | 0.56 | 0.00 | 0.00 | 0.96 | 0.81 | 8 | Endothelial |
| Tm4sf1 | 0.73 | 0.00 | 0.00 | 0.57 | 0.20 | 8 | Endothelial |
| Arap3 | 0.54 | 0.00 | 0.00 | 0.30 | 0.07 | 8 | Endothelial |
| Vat1 | 0.63 | 0.00 | 0.00 | 0.33 | 0.08 | 8 | Endothelial |
| Sdpr | 0.61 | 0.00 | 0.00 | 0.90 | 0.51 | 8 | Endothelial |
| Cish | 0.58 | 0.00 | 0.00 | 0.30 | 0.07 | 8 | Endothelial |
| Dock9 | 0.62 | 0.00 | 0.00 | 0.34 | 0.09 | 8 | Endothelial |
| Tspan18 | 0.61 | 0.00 | 0.00 | 0.43 | 0.13 | 8 | Endothelial |
| Fnbp1l | 0.84 | 0.00 | 0.00 | 0.59 | 0.24 | 8 | Endothelial |
| 2810025M15Rik | 0.63 | 0.00 | 0.00 | 0.40 | 0.12 | 8 | Endothelial |
| Hyal2 | 0.60 | 0.00 | 0.00 | 0.30 | 0.07 | 8 | Endothelial |
| Hoxa5 | 0.67 | 0.00 | 0.00 | 0.41 | 0.12 | 8 | Endothelial |
| Efnb1 | 0.68 | 0.00 | 0.00 | 0.44 | 0.14 | 8 | Endothelial |
| Tpst2 | 0.66 | 0.00 | 0.00 | 0.39 | 0.11 | 8 | Endothelial |
| Fli1 | 0.69 | 0.00 | 0.00 | 0.38 | 0.11 | 8 | Endothelial |
| Fbxw2 | 0.75 | 0.00 | 0.00 | 0.47 | 0.16 | 8 | Endothelial |
| Cd59a | 0.75 | 0.00 | 0.00 | 0.32 | 0.08 | 8 | Endothelial |
| Sh2b3 | 0.58 | 0.00 | 0.00 | 0.32 | 0.08 | 8 | Endothelial |
| Sema6a | 0.66 | 0.00 | 0.00 | 0.38 | 0.11 | 8 | Endothelial |
| Tenc1 | 0.68 | 0.00 | 0.00 | 0.42 | 0.14 | 8 | Endothelial |
| Tbx3 | 0.71 | 0.00 | 0.00 | 0.53 | 0.20 | 8 | Endothelial |
| Rasgrp2 | 0.57 | 0.00 | 0.00 | 0.34 | 0.09 | 8 | Endothelial |
| Rhoc | 0.69 | 0.00 | 0.00 | 0.53 | 0.20 | 8 | Endothelial |
| Dennd3 | 0.53 | 0.00 | 0.00 | 0.29 | 0.07 | 8 | Endothelial |
| Galnt18 | 0.63 | 0.00 | 0.00 | 0.36 | 0.10 | 8 | Endothelial |
| Gcc2 | 0.75 | 0.00 | 0.00 | 0.58 | 0.24 | 8 | Endothelial |
| Adcy4 | 0.55 | 0.00 | 0.00 | 0.31 | 0.08 | 8 | Endothelial |
| Sec14l1 | 0.66 | 0.00 | 0.00 | 0.38 | 0.12 | 8 | Endothelial |
| Fgd5 | 0.46 | 0.00 | 0.00 | 0.26 | 0.06 | 8 | Endothelial |
| Rpl31 | 0.62 | 0.00 | 0.00 | 0.86 | 0.57 | 8 | Endothelial |
| Rab11a | 0.71 | 0.00 | 0.00 | 0.59 | 0.25 | 8 | Endothelial |
| Nsg1 | 0.60 | 0.00 | 0.00 | 0.30 | 0.08 | 8 | Endothelial |
| Gm9846 | 0.44 | 0.00 | 0.00 | 0.98 | 0.89 | 8 | Endothelial |
| Snrk | 0.65 | 0.00 | 0.00 | 0.45 | 0.15 | 8 | Endothelial |
| Cyp4b1 | 0.60 | 0.00 | 0.00 | 0.36 | 0.10 | 8 | Endothelial |
| Rps29 | 0.44 | 0.00 | 0.00 | 0.98 | 0.90 | 8 | Endothelial |
| Ucp2 | 0.59 | 0.00 | 0.00 | 0.37 | 0.11 | 8 | Endothelial |
| Pim3 | 0.57 | 0.00 | 0.00 | 0.27 | 0.07 | 8 | Endothelial |
| Ivns1abp | 1.11 | 0.00 | 0.00 | 0.51 | 0.22 | 8 | Endothelial |
| Sgk1 | 0.76 | 0.00 | 0.00 | 0.49 | 0.18 | 8 | Endothelial |
| Eng | 0.57 | 0.00 | 0.00 | 0.36 | 0.11 | 8 | Endothelial |
| Ceacam1 | 0.53 | 0.00 | 0.00 | 0.32 | 0.09 | 8 | Endothelial |
| Podxl | 0.52 | 0.00 | 0.00 | 0.36 | 0.10 | 8 | Endothelial |
| Trib2 | 0.61 | 0.00 | 0.00 | 0.35 | 0.10 | 8 | Endothelial |
| Fkbp5 | 0.58 | 0.00 | 0.00 | 0.31 | 0.08 | 8 | Endothelial |
| Rasgef1a | 0.57 | 0.00 | 0.00 | 0.31 | 0.08 | 8 | Endothelial |
| Ets2 | 0.58 | 0.00 | 0.00 | 0.37 | 0.11 | 8 | Endothelial |
| Cnn3 | 0.57 | 0.00 | 0.00 | 0.65 | 0.29 | 8 | Endothelial |
| S100a13 | 0.74 | 0.00 | 0.00 | 0.56 | 0.25 | 8 | Endothelial |
| Fkbp1a | 0.62 | 0.00 | 0.00 | 0.74 | 0.40 | 8 | Endothelial |
| Afap1l1 | 0.54 | 0.00 | 0.00 | 0.47 | 0.17 | 8 | Endothelial |
| Rgs12 | 0.60 | 0.00 | 0.00 | 0.37 | 0.11 | 8 | Endothelial |
| Arl4a | 0.63 | 0.00 | 0.00 | 0.35 | 0.11 | 8 | Endothelial |
| Clic4 | 0.70 | 0.00 | 0.00 | 0.61 | 0.28 | 8 | Endothelial |
| Clic1 | 0.67 | 0.00 | 0.00 | 0.70 | 0.37 | 8 | Endothelial |
| Tjp1 | 0.57 | 0.00 | 0.00 | 0.44 | 0.15 | 8 | Endothelial |
| Rapgef5 | 0.53 | 0.00 | 0.00 | 0.34 | 0.10 | 8 | Endothelial |
| Plcg1 | 0.63 | 0.00 | 0.00 | 0.43 | 0.15 | 8 | Endothelial |
| Syn3 | 0.70 | 0.00 | 0.00 | 0.42 | 0.15 | 8 | Endothelial |
| Edn1 | 0.73 | 0.00 | 0.00 | 0.31 | 0.09 | 8 | Endothelial |
| Ifitm2 | 0.65 | 0.00 | 0.00 | 0.65 | 0.31 | 8 | Endothelial |
| Cd97 | 0.57 | 0.00 | 0.00 | 0.47 | 0.17 | 8 | Endothelial |
| Gstm1 | 0.61 | 0.00 | 0.00 | 0.41 | 0.14 | 8 | Endothelial |
| Prkce | 0.51 | 0.00 | 0.00 | 0.27 | 0.07 | 8 | Endothelial |
| Cd9 | 0.56 | 0.00 | 0.00 | 0.70 | 0.34 | 8 | Endothelial |
| Srgn | 0.51 | 0.00 | 0.00 | 0.69 | 0.32 | 8 | Endothelial |
| 4931406P16Rik | 0.60 | 0.00 | 0.00 | 0.46 | 0.17 | 8 | Endothelial |
| Socs3 | 0.49 | 0.00 | 0.00 | 0.26 | 0.07 | 8 | Endothelial |
| Tmem204 | 0.58 | 0.00 | 0.00 | 0.42 | 0.15 | 8 | Endothelial |
| Tax1bp1 | 0.68 | 0.00 | 0.00 | 0.85 | 0.63 | 8 | Endothelial |
| Fth1 | 0.52 | 0.00 | 0.00 | 0.94 | 0.81 | 8 | Endothelial |
| St3gal6 | 0.51 | 0.00 | 0.00 | 0.30 | 0.09 | 8 | Endothelial |
| Tmem2 | 0.55 | 0.00 | 0.00 | 0.40 | 0.14 | 8 | Endothelial |
| Prkd2 | 0.51 | 0.00 | 0.00 | 0.30 | 0.09 | 8 | Endothelial |
| Ccdc28b | 0.55 | 0.00 | 0.00 | 0.32 | 0.10 | 8 | Endothelial |
| Car4 | 0.66 | 0.00 | 0.00 | 0.27 | 0.07 | 8 | Endothelial |
| H2-T23 | 0.55 | 0.00 | 0.00 | 0.36 | 0.12 | 8 | Endothelial |
| Cbfa2t3 | 0.54 | 0.00 | 0.00 | 0.40 | 0.14 | 8 | Endothelial |
| Cttnbp2nl | 0.51 | 0.00 | 0.00 | 0.33 | 0.10 | 8 | Endothelial |
| Arhgef15 | 0.42 | 0.00 | 0.00 | 0.26 | 0.07 | 8 | Endothelial |
| Fmo1 | 0.49 | 0.00 | 0.00 | 0.34 | 0.11 | 8 | Endothelial |
| Ehd1 | 0.54 | 0.00 | 0.00 | 0.34 | 0.11 | 8 | Endothelial |
| Ctla2b | 0.47 | 0.00 | 0.00 | 0.25 | 0.07 | 8 | Endothelial |
| Nkd1 | 0.53 | 0.00 | 0.00 | 0.33 | 0.11 | 8 | Endothelial |
| Arhgap29 | 0.61 | 0.00 | 0.00 | 0.50 | 0.21 | 8 | Endothelial |
| Mest | 0.42 | 0.00 | 0.00 | 0.49 | 0.19 | 8 | Endothelial |
| Slc6a6 | 0.65 | 0.00 | 0.00 | 0.54 | 0.25 | 8 | Endothelial |
| Hdac7 | 0.55 | 0.00 | 0.00 | 0.37 | 0.13 | 8 | Endothelial |
| Sptbn1 | 0.52 | 0.00 | 0.00 | 0.82 | 0.54 | 8 | Endothelial |
| Ifitm3 | 0.54 | 0.00 | 0.00 | 0.71 | 0.39 | 8 | Endothelial |
| Mob2 | 0.51 | 0.00 | 0.00 | 0.28 | 0.08 | 8 | Endothelial |
| Lphn3 | 0.48 | 0.00 | 0.00 | 0.26 | 0.07 | 8 | Endothelial |
| Ubald2 | 0.57 | 0.00 | 0.00 | 0.38 | 0.14 | 8 | Endothelial |
| Dnajc8 | 0.64 | 0.00 | 0.00 | 0.71 | 0.42 | 8 | Endothelial |
| 2900026A02Rik | 0.46 | 0.00 | 0.00 | 0.26 | 0.07 | 8 | Endothelial |
| Pttg1ip | 0.56 | 0.00 | 0.00 | 0.41 | 0.16 | 8 | Endothelial |
| Mknk2 | 0.53 | 0.00 | 0.00 | 0.33 | 0.11 | 8 | Endothelial |
| Jup | 0.52 | 0.00 | 0.00 | 0.41 | 0.15 | 8 | Endothelial |
| Ets1 | 0.47 | 0.00 | 0.00 | 0.55 | 0.24 | 8 | Endothelial |
| Cyth3 | 0.54 | 0.00 | 0.00 | 0.53 | 0.24 | 8 | Endothelial |
| Dusp3 | 0.57 | 0.00 | 0.00 | 0.45 | 0.19 | 8 | Endothelial |
| Fam63a | 0.50 | 0.00 | 0.00 | 0.28 | 0.09 | 8 | Endothelial |
| Itga6 | 0.50 | 0.00 | 0.00 | 0.36 | 0.13 | 8 | Endothelial |
| Myct1 | 0.41 | 0.00 | 0.00 | 0.36 | 0.12 | 8 | Endothelial |
| Ppap2b | 0.56 | 0.00 | 0.00 | 0.52 | 0.23 | 8 | Endothelial |
| Efna1 | 0.46 | 0.00 | 0.00 | 0.25 | 0.07 | 8 | Endothelial |
| Gngt2 | 0.35 | 0.00 | 0.00 | 0.31 | 0.10 | 8 | Endothelial |
| Hilpda | 0.48 | 0.00 | 0.00 | 0.26 | 0.08 | 8 | Endothelial |
| Bst2 | 0.48 | 0.00 | 0.00 | 0.36 | 0.13 | 8 | Endothelial |
| Rasgrp3 | 0.38 | 0.00 | 0.00 | 0.36 | 0.12 | 8 | Endothelial |
| Eef1a1 | 0.36 | 0.00 | 0.00 | 0.96 | 0.87 | 8 | Endothelial |
| mt-Rnr2 | 0.38 | 0.00 | 0.00 | 1.00 | 0.99 | 8 | Endothelial |
| Leprot | 0.62 | 0.00 | 0.00 | 0.60 | 0.32 | 8 | Endothelial |
| Hpcal1 | 0.53 | 0.00 | 0.00 | 0.37 | 0.14 | 8 | Endothelial |
| Ptgs1 | 0.54 | 0.00 | 0.00 | 0.30 | 0.10 | 8 | Endothelial |
| Gm11808 | 0.42 | 0.00 | 0.00 | 0.90 | 0.77 | 8 | Endothelial |
| Rap1a | 0.55 | 0.00 | 0.00 | 0.58 | 0.29 | 8 | Endothelial |
| Rin2 | 0.58 | 0.00 | 0.00 | 0.38 | 0.15 | 8 | Endothelial |
| Rcsd1 | 0.44 | 0.00 | 0.00 | 0.26 | 0.08 | 8 | Endothelial |
| Gm7808 | 0.49 | 0.00 | 0.00 | 0.82 | 0.59 | 8 | Endothelial |
| Gnb2 | 0.55 | 0.00 | 0.00 | 0.65 | 0.37 | 8 | Endothelial |
| Junb | 0.44 | 0.00 | 0.00 | 0.45 | 0.19 | 8 | Endothelial |
| Smarca2 | 0.60 | 0.00 | 0.00 | 0.62 | 0.34 | 8 | Endothelial |
| Klf9 | 0.52 | 0.00 | 0.00 | 0.44 | 0.19 | 8 | Endothelial |
| Clec2d | 0.43 | 0.00 | 0.00 | 0.27 | 0.09 | 8 | Endothelial |
| Crip1 | 0.54 | 0.00 | 0.00 | 0.75 | 0.47 | 8 | Endothelial |
| Sparcl1 | 0.48 | 0.00 | 0.00 | 0.69 | 0.38 | 8 | Endothelial |
| Ptprg | 0.49 | 0.00 | 0.00 | 0.33 | 0.12 | 8 | Endothelial |
| Kit | 0.33 | 0.00 | 0.00 | 0.26 | 0.08 | 8 | Endothelial |
| Itpripl2 | 0.61 | 0.00 | 0.00 | 0.39 | 0.16 | 8 | Endothelial |
| Numb | 0.42 | 0.00 | 0.00 | 0.25 | 0.08 | 8 | Endothelial |
| Rps11 | 0.39 | 0.00 | 0.00 | 0.91 | 0.75 | 8 | Endothelial |
| Arglu1 | 0.55 | 0.00 | 0.00 | 0.68 | 0.41 | 8 | Endothelial |
| Cd47 | 0.57 | 0.00 | 0.00 | 0.69 | 0.42 | 8 | Endothelial |
| Arpc3 | 0.53 | 0.00 | 0.00 | 0.52 | 0.25 | 8 | Endothelial |
| Znrf1 | 0.45 | 0.00 | 0.00 | 0.30 | 0.11 | 8 | Endothelial |
| Bcam | 0.43 | 0.00 | 0.00 | 0.26 | 0.09 | 8 | Endothelial |
| Hmgn5 | 0.69 | 0.00 | 0.00 | 0.50 | 0.26 | 8 | Endothelial |
| Cdk14 | 0.48 | 0.00 | 0.00 | 0.30 | 0.11 | 8 | Endothelial |
| Col4a3bp | 0.53 | 0.00 | 0.00 | 0.36 | 0.15 | 8 | Endothelial |
| Pitpnc1 | 0.40 | 0.00 | 0.00 | 0.32 | 0.12 | 8 | Endothelial |
| Lphn2 | 0.39 | 0.00 | 0.00 | 0.33 | 0.13 | 8 | Endothelial |
| Gng11 | 0.38 | 0.00 | 0.00 | 0.57 | 0.28 | 8 | Endothelial |
| Vim | 0.39 | 0.00 | 0.00 | 0.90 | 0.65 | 8 | Endothelial |
| Zmiz1 | 0.50 | 0.00 | 0.00 | 0.45 | 0.21 | 8 | Endothelial |
| Plk2 | 0.49 | 0.00 | 0.00 | 0.37 | 0.16 | 8 | Endothelial |
| Gm10126 | 0.47 | 0.00 | 0.00 | 0.64 | 0.39 | 8 | Endothelial |
| Klf3 | 0.44 | 0.00 | 0.00 | 0.30 | 0.11 | 8 | Endothelial |
| Id3 | 0.41 | 0.00 | 0.00 | 0.74 | 0.45 | 8 | Endothelial |
| Klf13 | 0.44 | 0.00 | 0.00 | 0.38 | 0.16 | 8 | Endothelial |
| Arl2bp | 0.44 | 0.00 | 0.00 | 0.32 | 0.13 | 8 | Endothelial |
| Gm5239 | 0.44 | 0.00 | 0.00 | 0.75 | 0.51 | 8 | Endothelial |
| Atp2b1 | 0.46 | 0.00 | 0.00 | 0.80 | 0.58 | 8 | Endothelial |
| Tinagl1 | 0.29 | 0.00 | 0.00 | 0.28 | 0.10 | 8 | Endothelial |
| Rps23 | 0.33 | 0.00 | 0.00 | 0.92 | 0.79 | 8 | Endothelial |
| Tle4 | 0.50 | 0.00 | 0.00 | 0.31 | 0.12 | 8 | Endothelial |
| Igfbp4 | 0.34 | 0.00 | 0.00 | 0.51 | 0.24 | 8 | Endothelial |
| Atp1b3 | 0.45 | 0.00 | 0.00 | 0.49 | 0.25 | 8 | Endothelial |
| Arhgap31 | 0.36 | 0.00 | 0.00 | 0.29 | 0.12 | 8 | Endothelial |
| Zcchc6 | 0.53 | 0.00 | 0.00 | 0.47 | 0.25 | 8 | Endothelial |
| Rps25 | 0.32 | 0.00 | 0.00 | 0.92 | 0.80 | 8 | Endothelial |
| Elovl5 | 0.41 | 0.00 | 0.00 | 0.34 | 0.15 | 8 | Endothelial |
| Zfp36 | 0.36 | 0.00 | 0.00 | 0.36 | 0.16 | 8 | Endothelial |
| Zfp945 | 0.44 | 0.00 | 0.00 | 0.30 | 0.12 | 8 | Endothelial |
| Rab5c | 0.40 | 0.00 | 0.00 | 0.29 | 0.12 | 8 | Endothelial |
| Gimap1 | 0.35 | 0.00 | 0.00 | 0.32 | 0.13 | 8 | Endothelial |
| Rps14 | 0.27 | 0.00 | 0.00 | 0.97 | 0.92 | 8 | Endothelial |
| Cd34 | 0.33 | 0.00 | 0.00 | 0.42 | 0.19 | 8 | Endothelial |
| Gnaq | 0.43 | 0.00 | 0.00 | 0.42 | 0.21 | 8 | Endothelial |
| Zeb1 | 0.35 | 0.00 | 0.00 | 0.28 | 0.11 | 8 | Endothelial |
| Cirbp | 0.40 | 0.00 | 0.00 | 0.42 | 0.21 | 8 | Endothelial |
| Aldh2 | 0.42 | 0.00 | 0.00 | 0.56 | 0.33 | 8 | Endothelial |
| Acap2 | 0.43 | 0.00 | 0.00 | 0.40 | 0.19 | 8 | Endothelial |
| Gm24276 | 0.40 | 0.00 | 0.00 | 0.43 | 0.22 | 8 | Endothelial |
| Mgll | 0.34 | 0.00 | 0.00 | 0.35 | 0.15 | 8 | Endothelial |
| Dram2 | 0.35 | 0.00 | 0.00 | 0.35 | 0.16 | 8 | Endothelial |
| Tmem128 | 0.38 | 0.00 | 0.00 | 0.28 | 0.12 | 8 | Endothelial |
| Arhgef12 | 0.33 | 0.00 | 0.00 | 0.45 | 0.23 | 8 | Endothelial |
| Arhgef2 | 0.43 | 0.00 | 0.00 | 0.40 | 0.20 | 8 | Endothelial |
| Adamts1 | 0.35 | 0.00 | 0.00 | 0.42 | 0.21 | 8 | Endothelial |
| Rps8 | 0.28 | 0.00 | 0.00 | 0.93 | 0.81 | 8 | Endothelial |
| Aph1a | 0.35 | 0.00 | 0.00 | 0.28 | 0.12 | 8 | Endothelial |
| Gltscr2 | 0.39 | 0.00 | 0.00 | 0.70 | 0.47 | 8 | Endothelial |
| Cd151 | 0.32 | 0.00 | 0.00 | 0.35 | 0.16 | 8 | Endothelial |
| Mef2a | 0.41 | 0.00 | 0.00 | 0.59 | 0.35 | 8 | Endothelial |
| Rnf144a | 0.31 | 0.00 | 0.00 | 0.34 | 0.15 | 8 | Endothelial |
| Rac1 | 0.41 | 0.00 | 0.00 | 0.52 | 0.29 | 8 | Endothelial |
| Crk | 0.41 | 0.00 | 0.00 | 0.34 | 0.16 | 8 | Endothelial |
| Rbpms | 0.38 | 0.00 | 0.00 | 0.38 | 0.18 | 8 | Endothelial |
| Btg2 | 0.31 | 0.00 | 0.00 | 0.44 | 0.23 | 8 | Endothelial |
| Mpzl1 | 0.30 | 0.00 | 0.00 | 0.27 | 0.11 | 8 | Endothelial |
| Adipor2 | 0.41 | 0.00 | 0.00 | 0.26 | 0.11 | 8 | Endothelial |
| Csrp2 | 0.34 | 0.00 | 0.00 | 0.35 | 0.16 | 8 | Endothelial |
| Lxn | 0.33 | 0.00 | 0.00 | 0.28 | 0.12 | 8 | Endothelial |
| Top1 | 0.47 | 0.00 | 0.00 | 0.64 | 0.44 | 8 | Endothelial |
| Rhoa | 0.38 | 0.00 | 0.00 | 0.68 | 0.46 | 8 | Endothelial |
| Ddah2 | 0.35 | 0.00 | 0.00 | 0.48 | 0.26 | 8 | Endothelial |
| Med13l | 0.33 | 0.00 | 0.00 | 0.27 | 0.12 | 8 | Endothelial |
| RPL24 | 0.34 | 0.00 | 0.00 | 0.69 | 0.45 | 8 | Endothelial |
| Hmgb1 | 0.35 | 0.00 | 0.00 | 0.79 | 0.61 | 8 | Endothelial |
| Gm10020 | 0.35 | 0.00 | 0.00 | 0.72 | 0.51 | 8 | Endothelial |
| Wasf2 | 0.35 | 0.00 | 0.00 | 0.52 | 0.30 | 8 | Endothelial |
| Notch1 | 0.33 | 0.00 | 0.00 | 0.29 | 0.13 | 8 | Endothelial |
| Ccdc85b | 0.26 | 0.00 | 0.00 | 0.27 | 0.11 | 8 | Endothelial |
| Luzp1 | 0.39 | 0.00 | 0.00 | 0.38 | 0.19 | 8 | Endothelial |
| Ppp2r5a | 0.36 | 0.00 | 0.00 | 0.31 | 0.14 | 8 | Endothelial |
| Skap2 | 0.28 | 0.00 | 0.00 | 0.28 | 0.12 | 8 | Endothelial |
| Hmg20b | 0.41 | 0.00 | 0.00 | 0.54 | 0.33 | 8 | Endothelial |
| Sbds | 0.43 | 0.00 | 0.00 | 0.41 | 0.22 | 8 | Endothelial |
| Ptpn12 | 0.32 | 0.00 | 0.00 | 0.29 | 0.13 | 8 | Endothelial |
| Eif1 | 0.32 | 0.00 | 0.00 | 0.83 | 0.67 | 8 | Endothelial |
| Laptm4a | 0.35 | 0.00 | 0.00 | 0.73 | 0.51 | 8 | Endothelial |
| Becn1 | 0.36 | 0.00 | 0.00 | 0.30 | 0.14 | 8 | Endothelial |
| Uaca | 0.33 | 0.00 | 0.00 | 0.30 | 0.14 | 8 | Endothelial |
| Pmp22 | 0.38 | 0.00 | 0.00 | 0.53 | 0.31 | 8 | Endothelial |
| Rpl28 | 0.37 | 0.00 | 0.00 | 0.71 | 0.53 | 8 | Endothelial |
| Dpysl2 | 0.37 | 0.00 | 0.00 | 0.35 | 0.17 | 8 | Endothelial |
| Nav1 | 0.29 | 0.00 | 0.00 | 0.26 | 0.11 | 8 | Endothelial |
| Gnai2 | 0.34 | 0.00 | 0.00 | 0.60 | 0.38 | 8 | Endothelial |
| Arpc1b | 0.27 | 0.00 | 0.00 | 0.60 | 0.37 | 8 | Endothelial |
| Heg1 | 0.25 | 0.00 | 0.00 | 0.33 | 0.16 | 8 | Endothelial |
| Tmem50a | 0.39 | 0.00 | 0.00 | 0.65 | 0.44 | 8 | Endothelial |
| Rpl15 | 0.31 | 0.00 | 0.00 | 0.78 | 0.58 | 8 | Endothelial |
| Kras | 0.38 | 0.00 | 0.00 | 0.35 | 0.18 | 8 | Endothelial |
| Tmf1 | 0.33 | 0.00 | 0.00 | 0.35 | 0.18 | 8 | Endothelial |
| Rassf1 | 0.35 | 0.00 | 0.00 | 0.29 | 0.14 | 8 | Endothelial |
| Map7d1 | 0.34 | 0.00 | 0.00 | 0.35 | 0.18 | 8 | Endothelial |
| mt-Atp6 | 0.26 | 0.00 | 0.00 | 0.93 | 0.84 | 8 | Endothelial |
| Qk | 0.39 | 0.00 | 0.00 | 0.49 | 0.29 | 8 | Endothelial |
| Phf20l1 | 0.42 | 0.00 | 0.00 | 0.47 | 0.28 | 8 | Endothelial |
| Clk1 | 0.35 | 0.00 | 0.00 | 0.55 | 0.35 | 8 | Endothelial |
| Gpi1 | 0.29 | 0.00 | 0.00 | 0.29 | 0.14 | 8 | Endothelial |
| Abi1 | 0.39 | 0.00 | 0.00 | 0.40 | 0.22 | 8 | Endothelial |
| Rpl38 | 0.28 | 0.00 | 0.00 | 0.86 | 0.73 | 8 | Endothelial |
| Zmym5 | 0.36 | 0.00 | 0.00 | 0.37 | 0.20 | 8 | Endothelial |
| Sh3glb1 | 0.39 | 0.00 | 0.00 | 0.60 | 0.40 | 8 | Endothelial |
| Tsc22d1 | 0.36 | 0.00 | 0.00 | 0.53 | 0.33 | 8 | Endothelial |
| Smc4 | 0.26 | 0.00 | 0.00 | 0.56 | 0.34 | 8 | Endothelial |
| Rpl9-ps6 | 0.27 | 0.00 | 0.00 | 0.87 | 0.73 | 8 | Endothelial |
| Actn4 | 0.35 | 0.00 | 0.00 | 0.48 | 0.29 | 8 | Endothelial |
| Rpl27 | 0.29 | 0.00 | 0.00 | 0.84 | 0.69 | 8 | Endothelial |
| Gm10263 | 0.26 | 0.00 | 0.00 | 0.89 | 0.76 | 8 | Endothelial |
| Rpl9 | 0.30 | 0.00 | 0.00 | 0.80 | 0.62 | 8 | Endothelial |
| Hes1 | 0.37 | 0.00 | 0.00 | 0.48 | 0.29 | 8 | Endothelial |
| Tacc1 | 0.34 | 0.00 | 0.00 | 0.40 | 0.22 | 8 | Endothelial |
| Zfp326 | 0.41 | 0.00 | 0.00 | 0.41 | 0.23 | 8 | Endothelial |
| Plec | 0.29 | 0.00 | 0.00 | 0.27 | 0.12 | 8 | Endothelial |
| Dazap2 | 0.31 | 0.00 | 0.00 | 0.38 | 0.20 | 8 | Endothelial |
| Tead2 | 0.27 | 0.00 | 0.00 | 0.26 | 0.12 | 8 | Endothelial |
| Cfl1 | 0.28 | 0.00 | 0.00 | 0.78 | 0.57 | 8 | Endothelial |
| Ctnna1 | 0.30 | 0.00 | 0.00 | 0.56 | 0.36 | 8 | Endothelial |
| Fcho2 | 0.31 | 0.00 | 0.00 | 0.36 | 0.19 | 8 | Endothelial |
| Clip1 | 0.40 | 0.00 | 0.00 | 0.37 | 0.20 | 8 | Endothelial |
| Rpl17 | 0.31 | 0.00 | 0.00 | 0.73 | 0.53 | 8 | Endothelial |
| Twf1 | 0.33 | 0.00 | 0.00 | 0.31 | 0.16 | 8 | Endothelial |
| Gm6133 | 0.30 | 0.00 | 0.00 | 0.54 | 0.33 | 8 | Endothelial |
| Baz2b | 0.37 | 0.00 | 0.00 | 0.47 | 0.29 | 8 | Endothelial |
| Gkap1 | 0.32 | 0.00 | 0.00 | 0.26 | 0.12 | 8 | Endothelial |
| D19Ertd737e | 0.33 | 0.00 | 0.00 | 0.30 | 0.16 | 8 | Endothelial |
| Rpl29 | 0.31 | 0.00 | 0.00 | 0.65 | 0.45 | 8 | Endothelial |
| Rell1 | 0.36 | 0.00 | 0.00 | 0.36 | 0.20 | 8 | Endothelial |
| Sypl | 0.31 | 0.00 | 0.00 | 0.43 | 0.25 | 8 | Endothelial |
| Pde4b | 0.27 | 0.00 | 0.00 | 0.28 | 0.14 | 8 | Endothelial |
| Maf1 | 0.30 | 0.00 | 0.00 | 0.32 | 0.17 | 8 | Endothelial |
| Cmtm3 | 0.26 | 0.00 | 0.00 | 0.27 | 0.13 | 8 | Endothelial |
| Ndufc2 | 0.34 | 0.00 | 0.00 | 0.57 | 0.38 | 8 | Endothelial |
| Add3 | 0.28 | 0.00 | 0.00 | 0.46 | 0.28 | 8 | Endothelial |
| Gna13 | 0.29 | 0.00 | 0.00 | 0.27 | 0.14 | 8 | Endothelial |
| Klf6 | 0.31 | 0.00 | 0.00 | 0.54 | 0.35 | 8 | Endothelial |
| Cr1l | 0.31 | 0.00 | 0.00 | 0.26 | 0.13 | 8 | Endothelial |
| Syf2 | 0.29 | 0.00 | 0.00 | 0.48 | 0.29 | 8 | Endothelial |
| Pttg1 | 0.27 | 0.00 | 0.00 | 0.28 | 0.14 | 8 | Endothelial |
| Eif3f | 0.31 | 0.00 | 0.00 | 0.55 | 0.36 | 8 | Endothelial |
| Ctnnd1 | 0.25 | 0.00 | 0.00 | 0.31 | 0.16 | 8 | Endothelial |
| Zfp503 | 0.27 | 0.00 | 0.00 | 0.29 | 0.15 | 8 | Endothelial |
| Pdcd10 | 0.33 | 0.00 | 0.00 | 0.36 | 0.21 | 8 | Endothelial |
| Dnaja1 | 0.32 | 0.00 | 0.00 | 0.68 | 0.51 | 8 | Endothelial |
| Prrc2c | 0.34 | 0.00 | 0.00 | 0.77 | 0.63 | 8 | Endothelial |
| Vamp3 | 0.26 | 0.00 | 0.00 | 0.28 | 0.14 | 8 | Endothelial |
| Serinc1 | 0.31 | 0.00 | 0.00 | 0.46 | 0.29 | 8 | Endothelial |
| Strn3 | 0.31 | 0.00 | 0.00 | 0.37 | 0.21 | 8 | Endothelial |
| Gm20091 | 0.29 | 0.00 | 0.00 | 0.38 | 0.22 | 8 | Endothelial |
| Cdc42bpa | 0.33 | 0.00 | 0.00 | 0.28 | 0.15 | 8 | Endothelial |
| Rwdd1 | 0.27 | 0.00 | 0.00 | 0.44 | 0.27 | 8 | Endothelial |
| Dynlt1f | 0.30 | 0.00 | 0.00 | 0.39 | 0.24 | 8 | Endothelial |
| Cd2ap | 0.33 | 0.00 | 0.00 | 0.52 | 0.34 | 8 | Endothelial |
| Bnip2 | 0.26 | 0.00 | 0.00 | 0.47 | 0.30 | 8 | Endothelial |
| Cnih1 | 0.29 | 0.00 | 0.00 | 0.33 | 0.18 | 8 | Endothelial |
| Rb1cc1 | 0.32 | 0.00 | 0.00 | 0.46 | 0.30 | 8 | Endothelial |
| Irf2 | 0.29 | 0.00 | 0.00 | 0.40 | 0.24 | 8 | Endothelial |
| Trim56 | 0.31 | 0.00 | 0.00 | 0.31 | 0.17 | 8 | Endothelial |
| 1110008F13Rik | 0.26 | 0.00 | 0.00 | 0.29 | 0.16 | 8 | Endothelial |
| Pnpla8 | 0.27 | 0.00 | 0.00 | 0.32 | 0.18 | 8 | Endothelial |
| Atrx | 0.31 | 0.00 | 0.00 | 0.77 | 0.65 | 8 | Endothelial |
| Rtf1 | 0.30 | 0.00 | 0.00 | 0.50 | 0.33 | 8 | Endothelial |
| Ghr | 0.26 | 0.00 | 0.00 | 0.27 | 0.15 | 8 | Endothelial |
| Zc3h7a | 0.28 | 0.00 | 0.00 | 0.44 | 0.28 | 8 | Endothelial |
| Gabarapl2 | 0.26 | 0.00 | 0.00 | 0.36 | 0.22 | 8 | Endothelial |
| Bclaf1 | 0.29 | 0.00 | 0.00 | 0.58 | 0.41 | 8 | Endothelial |
| Dynlt1b | 0.28 | 0.00 | 0.00 | 0.40 | 0.25 | 8 | Endothelial |
| Dynlt1a | 0.30 | 0.00 | 0.00 | 0.37 | 0.22 | 8 | Endothelial |
| Ubn1 | 0.28 | 0.00 | 0.00 | 0.28 | 0.16 | 8 | Endothelial |
| Dynlt1c | 0.29 | 0.00 | 0.00 | 0.42 | 0.26 | 8 | Endothelial |
| Rab2a | 0.26 | 0.00 | 0.00 | 0.42 | 0.27 | 8 | Endothelial |
| Capns1 | 0.26 | 0.00 | 0.00 | 0.39 | 0.24 | 8 | Endothelial |
| Cox7a2l | 0.27 | 0.00 | 0.00 | 0.57 | 0.41 | 8 | Endothelial |
| Smap1 | 0.27 | 0.00 | 0.00 | 0.34 | 0.20 | 8 | Endothelial |
| Ccar1 | 0.32 | 0.00 | 0.00 | 0.47 | 0.32 | 8 | Endothelial |
| Bnip3l | 0.25 | 0.00 | 0.00 | 0.53 | 0.37 | 8 | Endothelial |
| Wbp4 | 0.28 | 0.00 | 0.00 | 0.38 | 0.24 | 8 | Endothelial |
| Arid4a | 0.29 | 0.00 | 0.00 | 0.44 | 0.29 | 8 | Endothelial |
| Ube2d3 | 0.28 | 0.00 | 0.00 | 0.47 | 0.32 | 8 | Endothelial |
| Ik | 0.25 | 0.00 | 0.00 | 0.51 | 0.35 | 8 | Endothelial |
| Zcrb1 | 0.26 | 0.00 | 0.00 | 0.43 | 0.29 | 8 | Endothelial |
| Eci2 | 0.25 | 0.00 | 0.00 | 0.28 | 0.17 | 8 | Endothelial |
| Apc | 0.25 | 0.00 | 0.00 | 0.36 | 0.23 | 8 | Endothelial |
| Anapc16 | 0.26 | 0.00 | 0.00 | 0.29 | 0.18 | 8 | Endothelial |
| Rbbp6 | 0.28 | 0.00 | 0.00 | 0.50 | 0.35 | 8 | Endothelial |
| Ppig | 0.27 | 0.00 | 0.00 | 0.53 | 0.39 | 8 | Endothelial |
| Dlk1 | 2.26 | 0.00 | 0.00 | 0.80 | 0.04 | 9 | AT2 Cell |
| Meg3 | 2.23 | 0.00 | 0.00 | 0.95 | 0.19 | 9 | AT2 Cell |
| Slc34a2 | 2.15 | 0.00 | 0.00 | 0.97 | 0.13 | 9 | AT2 Cell |
| Atp1b1 | 2.08 | 0.00 | 0.00 | 0.95 | 0.16 | 9 | AT2 Cell |
| Sftpd | 2.06 | 0.00 | 0.00 | 0.98 | 0.14 | 9 | AT2 Cell |
| Ctsh | 2.05 | 0.00 | 0.00 | 0.98 | 0.18 | 9 | AT2 Cell |
| Rgcc | 1.96 | 0.00 | 0.00 | 0.86 | 0.15 | 9 | AT2 Cell |
| Napsa | 1.91 | 0.00 | 0.00 | 0.98 | 0.17 | 9 | AT2 Cell |
| Bex4 | 1.87 | 0.00 | 0.00 | 0.89 | 0.10 | 9 | AT2 Cell |
| Sftpb | 1.82 | 0.00 | 0.00 | 0.99 | 0.15 | 9 | AT2 Cell |
| Npc2 | 1.82 | 0.00 | 0.00 | 1.00 | 0.53 | 9 | AT2 Cell |
| S100g | 1.80 | 0.00 | 0.00 | 0.85 | 0.10 | 9 | AT2 Cell |
| Pmvk | 1.77 | 0.00 | 0.00 | 0.67 | 0.07 | 9 | AT2 Cell |
| Sftpc | 1.65 | 0.00 | 0.00 | 1.00 | 0.62 | 9 | AT2 Cell |
| Errfi1 | 1.63 | 0.00 | 0.00 | 0.79 | 0.07 | 9 | AT2 Cell |
| Sftpa1 | 1.62 | 0.00 | 0.00 | 0.99 | 0.19 | 9 | AT2 Cell |
| Lpcat1 | 1.62 | 0.00 | 0.00 | 0.93 | 0.15 | 9 | AT2 Cell |
| Sfta2 | 1.58 | 0.00 | 0.00 | 0.94 | 0.13 | 9 | AT2 Cell |
| Cldn3 | 1.57 | 0.00 | 0.00 | 0.87 | 0.09 | 9 | AT2 Cell |
| Mt1 | 1.53 | 0.00 | 0.00 | 0.80 | 0.13 | 9 | AT2 Cell |
| Lpin2 | 1.50 | 0.00 | 0.00 | 0.80 | 0.13 | 9 | AT2 Cell |
| Spink5 | 1.46 | 0.00 | 0.00 | 0.64 | 0.06 | 9 | AT2 Cell |
| Selenbp1 | 1.45 | 0.00 | 0.00 | 0.86 | 0.20 | 9 | AT2 Cell |
| Bex1 | 1.44 | 0.00 | 0.00 | 0.78 | 0.08 | 9 | AT2 Cell |
| Dcxr | 1.39 | 0.00 | 0.00 | 0.77 | 0.08 | 9 | AT2 Cell |
| Cxcl15 | 1.38 | 0.00 | 0.00 | 0.91 | 0.12 | 9 | AT2 Cell |
| Prdx6 | 1.35 | 0.00 | 0.00 | 0.87 | 0.25 | 9 | AT2 Cell |
| Lmo7 | 1.35 | 0.00 | 0.00 | 0.75 | 0.11 | 9 | AT2 Cell |
| Scd2 | 1.34 | 0.00 | 0.00 | 0.85 | 0.19 | 9 | AT2 Cell |
| Dram1 | 1.32 | 0.00 | 0.00 | 0.82 | 0.09 | 9 | AT2 Cell |
| Selenbp2 | 1.30 | 0.00 | 0.00 | 0.79 | 0.15 | 9 | AT2 Cell |
| Alcam | 1.26 | 0.00 | 0.00 | 0.92 | 0.20 | 9 | AT2 Cell |
| Mt2 | 1.24 | 0.00 | 0.00 | 0.57 | 0.05 | 9 | AT2 Cell |
| Wfdc2 | 1.21 | 0.00 | 0.00 | 0.88 | 0.14 | 9 | AT2 Cell |
| Fabp5 | 1.20 | 0.00 | 0.00 | 0.70 | 0.11 | 9 | AT2 Cell |
| Gprc5a | 1.19 | 0.00 | 0.00 | 0.66 | 0.05 | 9 | AT2 Cell |
| Pi4k2b | 1.18 | 0.00 | 0.00 | 0.70 | 0.08 | 9 | AT2 Cell |
| Tinagl1 | 1.18 | 0.00 | 0.00 | 0.67 | 0.09 | 9 | AT2 Cell |
| Scnn1a | 1.17 | 0.00 | 0.00 | 0.64 | 0.05 | 9 | AT2 Cell |
| Ager | 1.16 | 0.00 | 0.00 | 0.85 | 0.15 | 9 | AT2 Cell |
| Cbr2 | 1.14 | 0.00 | 0.00 | 0.79 | 0.14 | 9 | AT2 Cell |
| Cldn18 | 1.13 | 0.00 | 0.00 | 0.90 | 0.15 | 9 | AT2 Cell |
| Mettl7a1 | 1.13 | 0.00 | 0.00 | 0.82 | 0.20 | 9 | AT2 Cell |
| Serpinb6b | 1.12 | 0.00 | 0.00 | 0.71 | 0.15 | 9 | AT2 Cell |
| Myh14 | 1.12 | 0.00 | 0.00 | 0.64 | 0.08 | 9 | AT2 Cell |
| Chchd10 | 1.11 | 0.00 | 0.00 | 0.78 | 0.11 | 9 | AT2 Cell |
| Lgi3 | 1.10 | 0.00 | 0.00 | 0.66 | 0.06 | 9 | AT2 Cell |
| Tfcp2l1 | 1.08 | 0.00 | 0.00 | 0.53 | 0.02 | 9 | AT2 Cell |
| H2afj | 1.08 | 0.00 | 0.00 | 0.87 | 0.30 | 9 | AT2 Cell |
| Nkx2-1 | 1.07 | 0.00 | 0.00 | 0.70 | 0.08 | 9 | AT2 Cell |
| Swt1 | 1.07 | 0.00 | 0.00 | 0.72 | 0.14 | 9 | AT2 Cell |
| Ptprf | 1.06 | 0.00 | 0.00 | 0.76 | 0.14 | 9 | AT2 Cell |
| Irx1 | 1.05 | 0.00 | 0.00 | 0.57 | 0.03 | 9 | AT2 Cell |
| Tmem243 | 1.04 | 0.00 | 0.00 | 0.61 | 0.06 | 9 | AT2 Cell |
| Exosc7 | 1.04 | 0.00 | 0.00 | 0.67 | 0.11 | 9 | AT2 Cell |
| Slc12a2 | 1.03 | 0.00 | 0.00 | 0.71 | 0.13 | 9 | AT2 Cell |
| Hc | 1.03 | 0.00 | 0.00 | 0.78 | 0.09 | 9 | AT2 Cell |
| Gde1 | 1.02 | 0.00 | 0.00 | 0.72 | 0.13 | 9 | AT2 Cell |
| Myo5c | 1.02 | 0.00 | 0.00 | 0.60 | 0.06 | 9 | AT2 Cell |
| Timp3 | 1.01 | 0.00 | 0.00 | 0.84 | 0.34 | 9 | AT2 Cell |
| Rnase4 | 1.01 | 0.00 | 0.00 | 0.75 | 0.16 | 9 | AT2 Cell |
| Vamp8 | 1.00 | 0.00 | 0.00 | 0.89 | 0.33 | 9 | AT2 Cell |
| Hopx | 1.00 | 0.00 | 0.00 | 0.57 | 0.09 | 9 | AT2 Cell |
| Abcd3 | 1.00 | 0.00 | 0.00 | 0.75 | 0.17 | 9 | AT2 Cell |
| Clic3 | 0.99 | 0.00 | 0.00 | 0.64 | 0.07 | 9 | AT2 Cell |
| Chka | 0.99 | 0.00 | 0.00 | 0.61 | 0.09 | 9 | AT2 Cell |
| Sdc1 | 0.99 | 0.00 | 0.00 | 0.70 | 0.10 | 9 | AT2 Cell |
| Scd1 | 0.99 | 0.00 | 0.00 | 0.59 | 0.07 | 9 | AT2 Cell |
| Slc15a2 | 0.98 | 0.00 | 0.00 | 0.56 | 0.04 | 9 | AT2 Cell |
| Epcam | 0.97 | 0.00 | 0.00 | 0.66 | 0.08 | 9 | AT2 Cell |
| Krt18 | 0.97 | 0.00 | 0.00 | 0.59 | 0.06 | 9 | AT2 Cell |
| Bend7 | 0.97 | 0.00 | 0.00 | 0.57 | 0.05 | 9 | AT2 Cell |
| Herpud1 | 0.97 | 0.00 | 0.00 | 0.60 | 0.12 | 9 | AT2 Cell |
| Taok3 | 0.96 | 0.00 | 0.00 | 0.73 | 0.17 | 9 | AT2 Cell |
| Atp6v0a4 | 0.96 | 0.00 | 0.00 | 0.43 | 0.01 | 9 | AT2 Cell |
| Akap5 | 0.96 | 0.00 | 0.00 | 0.56 | 0.08 | 9 | AT2 Cell |
| Mettl7a2 | 0.94 | 0.00 | 0.00 | 0.61 | 0.09 | 9 | AT2 Cell |
| Eif4h | 0.93 | 0.00 | 0.00 | 0.76 | 0.25 | 9 | AT2 Cell |
| Insig1 | 0.92 | 0.00 | 0.00 | 0.51 | 0.06 | 9 | AT2 Cell |
| Sdc4 | 0.92 | 0.00 | 0.00 | 0.83 | 0.21 | 9 | AT2 Cell |
| Krt8 | 0.91 | 0.00 | 0.00 | 0.57 | 0.06 | 9 | AT2 Cell |
| Zdhhc3 | 0.91 | 0.00 | 0.00 | 0.69 | 0.14 | 9 | AT2 Cell |
| Krt7 | 0.91 | 0.00 | 0.00 | 0.56 | 0.06 | 9 | AT2 Cell |
| 1500012F01Rik | 0.91 | 0.00 | 0.00 | 0.80 | 0.28 | 9 | AT2 Cell |
| Ces1d | 0.90 | 0.00 | 0.00 | 0.77 | 0.15 | 9 | AT2 Cell |
| Mgst1 | 0.90 | 0.00 | 0.00 | 0.79 | 0.18 | 9 | AT2 Cell |
| Npc1 | 0.88 | 0.00 | 0.00 | 0.56 | 0.07 | 9 | AT2 Cell |
| Cystm1 | 0.87 | 0.00 | 0.00 | 0.56 | 0.06 | 9 | AT2 Cell |
| Icam1 | 0.86 | 0.00 | 0.00 | 0.57 | 0.08 | 9 | AT2 Cell |
| Aqp5 | 0.86 | 0.00 | 0.00 | 0.46 | 0.05 | 9 | AT2 Cell |
| Socs2 | 0.86 | 0.00 | 0.00 | 0.83 | 0.32 | 9 | AT2 Cell |
| 8430408G22Rik | 0.85 | 0.00 | 0.00 | 0.42 | 0.03 | 9 | AT2 Cell |
| Muc1 | 0.85 | 0.00 | 0.00 | 0.59 | 0.07 | 9 | AT2 Cell |
| Cebpa | 0.85 | 0.00 | 0.00 | 0.48 | 0.04 | 9 | AT2 Cell |
| Ank3 | 0.85 | 0.00 | 0.00 | 0.55 | 0.06 | 9 | AT2 Cell |
| Ptp4a1 | 0.84 | 0.00 | 0.00 | 0.54 | 0.09 | 9 | AT2 Cell |
| Pip5k1b | 0.83 | 0.00 | 0.00 | 0.44 | 0.02 | 9 | AT2 Cell |
| Ccz1 | 0.83 | 0.00 | 0.00 | 0.61 | 0.11 | 9 | AT2 Cell |
| Ppp1r14c | 0.83 | 0.00 | 0.00 | 0.67 | 0.09 | 9 | AT2 Cell |
| Mid1ip1 | 0.83 | 0.00 | 0.00 | 0.59 | 0.13 | 9 | AT2 Cell |
| Cstb | 0.83 | 0.00 | 0.00 | 0.78 | 0.26 | 9 | AT2 Cell |
| Tspan2 | 0.83 | 0.00 | 0.00 | 0.59 | 0.11 | 9 | AT2 Cell |
| Rps19 | 0.82 | 0.00 | 0.00 | 0.99 | 0.82 | 9 | AT2 Cell |
| 2200002D01Rik | 0.82 | 0.00 | 0.00 | 0.47 | 0.06 | 9 | AT2 Cell |
| Npnt | 0.81 | 0.00 | 0.00 | 0.87 | 0.22 | 9 | AT2 Cell |
| Egfl6 | 0.81 | 0.00 | 0.00 | 0.59 | 0.07 | 9 | AT2 Cell |
| Gadd45g | 0.81 | 0.00 | 0.00 | 0.45 | 0.07 | 9 | AT2 Cell |
| Rai14 | 0.81 | 0.00 | 0.00 | 0.57 | 0.09 | 9 | AT2 Cell |
| Acot1 | 0.80 | 0.00 | 0.00 | 0.52 | 0.06 | 9 | AT2 Cell |
| Secisbp2l | 0.80 | 0.00 | 0.00 | 0.66 | 0.16 | 9 | AT2 Cell |
| Irx3 | 0.79 | 0.00 | 0.00 | 0.52 | 0.06 | 9 | AT2 Cell |
| D230025D16Rik | 0.79 | 0.00 | 0.00 | 0.52 | 0.06 | 9 | AT2 Cell |
| Gclc | 0.79 | 0.00 | 0.00 | 0.47 | 0.07 | 9 | AT2 Cell |
| Abca3 | 0.78 | 0.00 | 0.00 | 0.63 | 0.10 | 9 | AT2 Cell |
| Cldn7 | 0.78 | 0.00 | 0.00 | 0.55 | 0.06 | 9 | AT2 Cell |
| Brd7 | 0.76 | 0.00 | 0.00 | 0.74 | 0.24 | 9 | AT2 Cell |
| Spint2 | 0.76 | 0.00 | 0.00 | 0.60 | 0.10 | 9 | AT2 Cell |
| Lamp3 | 0.76 | 0.00 | 0.00 | 0.67 | 0.10 | 9 | AT2 Cell |
| Ctsc | 0.76 | 0.00 | 0.00 | 0.72 | 0.18 | 9 | AT2 Cell |
| Etv5 | 0.76 | 0.00 | 0.00 | 0.48 | 0.05 | 9 | AT2 Cell |
| Ezr | 0.75 | 0.00 | 0.00 | 0.76 | 0.20 | 9 | AT2 Cell |
| Bex2 | 0.75 | 0.00 | 0.00 | 0.53 | 0.09 | 9 | AT2 Cell |
| Tmem213 | 0.74 | 0.00 | 0.00 | 0.36 | 0.02 | 9 | AT2 Cell |
| Ndnf | 0.74 | 0.00 | 0.00 | 0.41 | 0.04 | 9 | AT2 Cell |
| Wdr45 | 0.74 | 0.00 | 0.00 | 0.49 | 0.07 | 9 | AT2 Cell |
| Emp2 | 0.73 | 0.00 | 0.00 | 0.84 | 0.33 | 9 | AT2 Cell |
| Col4a3 | 0.73 | 0.00 | 0.00 | 0.42 | 0.04 | 9 | AT2 Cell |
| Kcnj15 | 0.73 | 0.00 | 0.00 | 0.41 | 0.03 | 9 | AT2 Cell |
| Bri3 | 0.73 | 0.00 | 0.00 | 0.69 | 0.19 | 9 | AT2 Cell |
| Krt19 | 0.72 | 0.00 | 0.00 | 0.46 | 0.05 | 9 | AT2 Cell |
| Malat1 | 0.72 | 0.00 | 0.00 | 1.00 | 1.00 | 9 | AT2 Cell |
| Wwc1 | 0.72 | 0.00 | 0.00 | 0.45 | 0.05 | 9 | AT2 Cell |
| Pcnt | 0.72 | 0.00 | 0.00 | 0.55 | 0.13 | 9 | AT2 Cell |
| Fasn | 0.72 | 0.00 | 0.00 | 0.48 | 0.06 | 9 | AT2 Cell |
| Uhrf1bp1l | 0.71 | 0.00 | 0.00 | 0.51 | 0.08 | 9 | AT2 Cell |
| Mbip | 0.71 | 0.00 | 0.00 | 0.55 | 0.10 | 9 | AT2 Cell |
| Tmem245 | 0.70 | 0.00 | 0.00 | 0.53 | 0.09 | 9 | AT2 Cell |
| Prdx6b | 0.70 | 0.00 | 0.00 | 0.43 | 0.05 | 9 | AT2 Cell |
| Pde7a | 0.69 | 0.00 | 0.00 | 0.45 | 0.06 | 9 | AT2 Cell |
| Atp8a1 | 0.69 | 0.00 | 0.00 | 0.62 | 0.14 | 9 | AT2 Cell |
| Cdkn2b | 0.68 | 0.00 | 0.00 | 0.41 | 0.04 | 9 | AT2 Cell |
| Gas6 | 0.68 | 0.00 | 0.00 | 0.60 | 0.13 | 9 | AT2 Cell |
| Acsl4 | 0.68 | 0.00 | 0.00 | 0.53 | 0.10 | 9 | AT2 Cell |
| Irx2 | 0.68 | 0.00 | 0.00 | 0.40 | 0.03 | 9 | AT2 Cell |
| AA986860 | 0.68 | 0.00 | 0.00 | 0.37 | 0.02 | 9 | AT2 Cell |
| Gramd2 | 0.68 | 0.00 | 0.00 | 0.40 | 0.04 | 9 | AT2 Cell |
| Rtkn2 | 0.66 | 0.00 | 0.00 | 0.37 | 0.05 | 9 | AT2 Cell |
| Msln | 0.66 | 0.00 | 0.00 | 0.33 | 0.04 | 9 | AT2 Cell |
| Fgfr2 | 0.66 | 0.00 | 0.00 | 0.44 | 0.06 | 9 | AT2 Cell |
| Ppp1r9a | 0.66 | 0.00 | 0.00 | 0.56 | 0.13 | 9 | AT2 Cell |
| Fdx1 | 0.66 | 0.00 | 0.00 | 0.55 | 0.13 | 9 | AT2 Cell |
| Tmem63b | 0.65 | 0.00 | 0.00 | 0.44 | 0.05 | 9 | AT2 Cell |
| Fam189a2 | 0.65 | 0.00 | 0.00 | 0.37 | 0.03 | 9 | AT2 Cell |
| Slc39a8 | 0.65 | 0.00 | 0.00 | 0.44 | 0.06 | 9 | AT2 Cell |
| Prss8 | 0.65 | 0.00 | 0.00 | 0.38 | 0.03 | 9 | AT2 Cell |
| S100a14 | 0.65 | 0.00 | 0.00 | 0.33 | 0.03 | 9 | AT2 Cell |
| Elf5 | 0.65 | 0.00 | 0.00 | 0.34 | 0.02 | 9 | AT2 Cell |
| Gm26809 | 0.65 | 0.00 | 0.00 | 0.33 | 0.04 | 9 | AT2 Cell |
| Gm12840 | 0.64 | 0.00 | 0.00 | 0.38 | 0.06 | 9 | AT2 Cell |
| Rbm47 | 0.64 | 0.00 | 0.00 | 0.40 | 0.04 | 9 | AT2 Cell |
| Kcnc3 | 0.64 | 0.00 | 0.00 | 0.35 | 0.02 | 9 | AT2 Cell |
| Pim3 | 0.64 | 0.00 | 0.00 | 0.44 | 0.06 | 9 | AT2 Cell |
| C77080 | 0.63 | 0.00 | 0.00 | 0.37 | 0.03 | 9 | AT2 Cell |
| C77370 | 0.62 | 0.00 | 0.00 | 0.31 | 0.02 | 9 | AT2 Cell |
| Pla2g1b | 0.62 | 0.00 | 0.00 | 0.39 | 0.04 | 9 | AT2 Cell |
| F3 | 0.62 | 0.00 | 0.00 | 0.35 | 0.04 | 9 | AT2 Cell |
| Mal2 | 0.62 | 0.00 | 0.00 | 0.43 | 0.05 | 9 | AT2 Cell |
| Atp13a3 | 0.61 | 0.00 | 0.00 | 0.48 | 0.08 | 9 | AT2 Cell |
| Rasl11a | 0.61 | 0.00 | 0.00 | 0.35 | 0.05 | 9 | AT2 Cell |
| Sik1 | 0.61 | 0.00 | 0.00 | 0.35 | 0.03 | 9 | AT2 Cell |
| Scnn1g | 0.61 | 0.00 | 0.00 | 0.33 | 0.03 | 9 | AT2 Cell |
| Btg3 | 0.61 | 0.00 | 0.00 | 0.50 | 0.10 | 9 | AT2 Cell |
| Lamc2 | 0.60 | 0.00 | 0.00 | 0.38 | 0.03 | 9 | AT2 Cell |
| Tspan8 | 0.60 | 0.00 | 0.00 | 0.54 | 0.11 | 9 | AT2 Cell |
| Tspan11 | 0.60 | 0.00 | 0.00 | 0.34 | 0.03 | 9 | AT2 Cell |
| Ppl | 0.60 | 0.00 | 0.00 | 0.35 | 0.03 | 9 | AT2 Cell |
| Mal | 0.60 | 0.00 | 0.00 | 0.37 | 0.04 | 9 | AT2 Cell |
| Prr15l | 0.60 | 0.00 | 0.00 | 0.41 | 0.05 | 9 | AT2 Cell |
| Tmem37 | 0.60 | 0.00 | 0.00 | 0.34 | 0.04 | 9 | AT2 Cell |
| Gm7334 | 0.59 | 0.00 | 0.00 | 0.50 | 0.10 | 9 | AT2 Cell |
| Nedd4l | 0.59 | 0.00 | 0.00 | 0.37 | 0.04 | 9 | AT2 Cell |
| Ccdc68 | 0.59 | 0.00 | 0.00 | 0.37 | 0.04 | 9 | AT2 Cell |
| Atp6v1c2 | 0.59 | 0.00 | 0.00 | 0.37 | 0.04 | 9 | AT2 Cell |
| Crb3 | 0.59 | 0.00 | 0.00 | 0.36 | 0.03 | 9 | AT2 Cell |
| Cmtm8 | 0.59 | 0.00 | 0.00 | 0.52 | 0.11 | 9 | AT2 Cell |
| Ccdc141 | 0.59 | 0.00 | 0.00 | 0.46 | 0.09 | 9 | AT2 Cell |
| Prkci | 0.59 | 0.00 | 0.00 | 0.52 | 0.12 | 9 | AT2 Cell |
| Txlng | 0.59 | 0.00 | 0.00 | 0.42 | 0.08 | 9 | AT2 Cell |
| Eps8l2 | 0.57 | 0.00 | 0.00 | 0.33 | 0.03 | 9 | AT2 Cell |
| Llgl2 | 0.57 | 0.00 | 0.00 | 0.35 | 0.03 | 9 | AT2 Cell |
| Kcnk1 | 0.57 | 0.00 | 0.00 | 0.32 | 0.02 | 9 | AT2 Cell |
| Avpi1 | 0.57 | 0.00 | 0.00 | 0.44 | 0.08 | 9 | AT2 Cell |
| Cdh1 | 0.56 | 0.00 | 0.00 | 0.44 | 0.07 | 9 | AT2 Cell |
| Rbpjl | 0.56 | 0.00 | 0.00 | 0.38 | 0.05 | 9 | AT2 Cell |
| Wwp1 | 0.56 | 0.00 | 0.00 | 0.53 | 0.12 | 9 | AT2 Cell |
| Matn4 | 0.55 | 0.00 | 0.00 | 0.33 | 0.03 | 9 | AT2 Cell |
| Pfkfb2 | 0.55 | 0.00 | 0.00 | 0.37 | 0.05 | 9 | AT2 Cell |
| Slc1a5 | 0.55 | 0.00 | 0.00 | 0.48 | 0.10 | 9 | AT2 Cell |
| Col4a4 | 0.55 | 0.00 | 0.00 | 0.36 | 0.03 | 9 | AT2 Cell |
| Srebf1 | 0.55 | 0.00 | 0.00 | 0.36 | 0.05 | 9 | AT2 Cell |
| Erbb3 | 0.54 | 0.00 | 0.00 | 0.32 | 0.02 | 9 | AT2 Cell |
| Acot2 | 0.53 | 0.00 | 0.00 | 0.37 | 0.05 | 9 | AT2 Cell |
| Traf1 | 0.52 | 0.00 | 0.00 | 0.34 | 0.03 | 9 | AT2 Cell |
| Nrbp2 | 0.52 | 0.00 | 0.00 | 0.35 | 0.05 | 9 | AT2 Cell |
| Arg2 | 0.52 | 0.00 | 0.00 | 0.26 | 0.01 | 9 | AT2 Cell |
| Mthfd1 | 0.51 | 0.00 | 0.00 | 0.41 | 0.07 | 9 | AT2 Cell |
| Rian | 0.51 | 0.00 | 0.00 | 0.31 | 0.03 | 9 | AT2 Cell |
| Ano1 | 0.51 | 0.00 | 0.00 | 0.34 | 0.04 | 9 | AT2 Cell |
| Dhrs3 | 0.51 | 0.00 | 0.00 | 0.37 | 0.06 | 9 | AT2 Cell |
| Rab11fip1 | 0.50 | 0.00 | 0.00 | 0.34 | 0.04 | 9 | AT2 Cell |
| Slc6a14 | 0.50 | 0.00 | 0.00 | 0.28 | 0.03 | 9 | AT2 Cell |
| Flcn | 0.49 | 0.00 | 0.00 | 0.34 | 0.04 | 9 | AT2 Cell |
| Mir22hg | 0.48 | 0.00 | 0.00 | 0.35 | 0.06 | 9 | AT2 Cell |
| Spock2 | 0.48 | 0.00 | 0.00 | 0.30 | 0.04 | 9 | AT2 Cell |
| Tspan1 | 0.48 | 0.00 | 0.00 | 0.31 | 0.03 | 9 | AT2 Cell |
| Pnpla2 | 0.48 | 0.00 | 0.00 | 0.38 | 0.06 | 9 | AT2 Cell |
| Hk2 | 0.48 | 0.00 | 0.00 | 0.33 | 0.04 | 9 | AT2 Cell |
| Foxa2 | 0.47 | 0.00 | 0.00 | 0.31 | 0.04 | 9 | AT2 Cell |
| Gja1 | 0.47 | 0.00 | 0.00 | 0.45 | 0.08 | 9 | AT2 Cell |
| Snx25 | 0.47 | 0.00 | 0.00 | 0.35 | 0.05 | 9 | AT2 Cell |
| Ralgapa2 | 0.47 | 0.00 | 0.00 | 0.31 | 0.04 | 9 | AT2 Cell |
| Foxo3 | 0.47 | 0.00 | 0.00 | 0.36 | 0.06 | 9 | AT2 Cell |
| Per1 | 0.46 | 0.00 | 0.00 | 0.31 | 0.04 | 9 | AT2 Cell |
| Pgs1 | 0.46 | 0.00 | 0.00 | 0.38 | 0.06 | 9 | AT2 Cell |
| Fam129b | 0.46 | 0.00 | 0.00 | 0.39 | 0.07 | 9 | AT2 Cell |
| Slc43a2 | 0.46 | 0.00 | 0.00 | 0.34 | 0.05 | 9 | AT2 Cell |
| Klf15 | 0.46 | 0.00 | 0.00 | 0.28 | 0.02 | 9 | AT2 Cell |
| Bnip3 | 0.45 | 0.00 | 0.00 | 0.41 | 0.08 | 9 | AT2 Cell |
| Ldlr | 0.45 | 0.00 | 0.00 | 0.30 | 0.04 | 9 | AT2 Cell |
| Pard6b | 0.45 | 0.00 | 0.00 | 0.26 | 0.02 | 9 | AT2 Cell |
| Arhgap44 | 0.45 | 0.00 | 0.00 | 0.27 | 0.02 | 9 | AT2 Cell |
| Sh3d21 | 0.44 | 0.00 | 0.00 | 0.27 | 0.03 | 9 | AT2 Cell |
| Perp | 0.44 | 0.00 | 0.00 | 0.27 | 0.02 | 9 | AT2 Cell |
| Mapkapk2 | 0.44 | 0.00 | 0.00 | 0.33 | 0.05 | 9 | AT2 Cell |
| Pvrl2 | 0.44 | 0.00 | 0.00 | 0.32 | 0.04 | 9 | AT2 Cell |
| Baiap2l1 | 0.44 | 0.00 | 0.00 | 0.28 | 0.03 | 9 | AT2 Cell |
| Epb4.1l5 | 0.43 | 0.00 | 0.00 | 0.29 | 0.03 | 9 | AT2 Cell |
| Cxadr | 0.43 | 0.00 | 0.00 | 0.33 | 0.05 | 9 | AT2 Cell |
| Spry2 | 0.42 | 0.00 | 0.00 | 0.31 | 0.05 | 9 | AT2 Cell |
| Arhgef28 | 0.42 | 0.00 | 0.00 | 0.26 | 0.03 | 9 | AT2 Cell |
| Cds1 | 0.42 | 0.00 | 0.00 | 0.26 | 0.02 | 9 | AT2 Cell |
| Nceh1 | 0.42 | 0.00 | 0.00 | 0.34 | 0.06 | 9 | AT2 Cell |
| Igf2bp2 | 0.42 | 0.00 | 0.00 | 0.29 | 0.03 | 9 | AT2 Cell |
| Slc26a9 | 0.42 | 0.00 | 0.00 | 0.25 | 0.02 | 9 | AT2 Cell |
| Klf5 | 0.41 | 0.00 | 0.00 | 0.27 | 0.02 | 9 | AT2 Cell |
| Cdc25b | 0.41 | 0.00 | 0.00 | 0.28 | 0.04 | 9 | AT2 Cell |
| Map7 | 0.41 | 0.00 | 0.00 | 0.30 | 0.04 | 9 | AT2 Cell |
| Rasgef1b | 0.41 | 0.00 | 0.00 | 0.27 | 0.02 | 9 | AT2 Cell |
| Vsig2 | 0.41 | 0.00 | 0.00 | 0.26 | 0.03 | 9 | AT2 Cell |
| Pkp2 | 0.40 | 0.00 | 0.00 | 0.27 | 0.03 | 9 | AT2 Cell |
| Igfbp6 | 0.39 | 0.00 | 0.00 | 0.26 | 0.03 | 9 | AT2 Cell |
| Pgm2 | 0.39 | 0.00 | 0.00 | 0.27 | 0.03 | 9 | AT2 Cell |
| Mpv17l2 | 0.39 | 0.00 | 0.00 | 0.29 | 0.04 | 9 | AT2 Cell |
| Flrt3 | 0.38 | 0.00 | 0.00 | 0.27 | 0.03 | 9 | AT2 Cell |
| Lamb3 | 0.38 | 0.00 | 0.00 | 0.26 | 0.03 | 9 | AT2 Cell |
| Tc2n | 0.38 | 0.00 | 0.00 | 0.26 | 0.03 | 9 | AT2 Cell |
| Tmc4 | 0.38 | 0.00 | 0.00 | 0.26 | 0.03 | 9 | AT2 Cell |
| Sec14l3 | 0.29 | 0.00 | 0.00 | 0.50 | 0.10 | 9 | AT2 Cell |
| Cadm1 | 0.55 | 0.00 | 0.00 | 0.54 | 0.13 | 9 | AT2 Cell |
| Il18r1 | 0.44 | 0.00 | 0.00 | 0.35 | 0.06 | 9 | AT2 Cell |
| Siva1 | 0.59 | 0.00 | 0.00 | 0.54 | 0.13 | 9 | AT2 Cell |
| Ypel5 | 0.53 | 0.00 | 0.00 | 0.46 | 0.10 | 9 | AT2 Cell |
| Wbp5 | 0.81 | 0.00 | 0.00 | 0.94 | 0.46 | 9 | AT2 Cell |
| Fdps | 0.42 | 0.00 | 0.00 | 0.31 | 0.05 | 9 | AT2 Cell |
| Gpx4 | 0.81 | 0.00 | 0.00 | 0.86 | 0.36 | 9 | AT2 Cell |
| Col23a1 | 0.45 | 0.00 | 0.00 | 0.28 | 0.04 | 9 | AT2 Cell |
| Spry1 | 0.46 | 0.00 | 0.00 | 0.35 | 0.06 | 9 | AT2 Cell |
| St6galnac2 | 0.39 | 0.00 | 0.00 | 0.30 | 0.05 | 9 | AT2 Cell |
| Vapa | 0.70 | 0.00 | 0.00 | 0.67 | 0.21 | 9 | AT2 Cell |
| Rpl21 | 0.71 | 0.00 | 0.00 | 0.97 | 0.74 | 9 | AT2 Cell |
| Atp6v0a1 | 0.45 | 0.00 | 0.00 | 0.34 | 0.06 | 9 | AT2 Cell |
| Tef | 0.46 | 0.00 | 0.00 | 0.35 | 0.07 | 9 | AT2 Cell |
| Dnase2a | 0.42 | 0.00 | 0.00 | 0.31 | 0.05 | 9 | AT2 Cell |
| Lzts2 | 0.50 | 0.00 | 0.00 | 0.42 | 0.09 | 9 | AT2 Cell |
| Arl6ip1 | 0.82 | 0.00 | 0.00 | 0.93 | 0.51 | 9 | AT2 Cell |
| Plp2 | 0.50 | 0.00 | 0.00 | 0.49 | 0.12 | 9 | AT2 Cell |
| Pdpn | 0.42 | 0.00 | 0.00 | 0.28 | 0.04 | 9 | AT2 Cell |
| P4ha1 | 0.51 | 0.00 | 0.00 | 0.38 | 0.08 | 9 | AT2 Cell |
| Pdgfa | 0.50 | 0.00 | 0.00 | 0.43 | 0.09 | 9 | AT2 Cell |
| Plxna2 | 0.39 | 0.00 | 0.00 | 0.35 | 0.07 | 9 | AT2 Cell |
| Chp1 | 0.51 | 0.00 | 0.00 | 0.51 | 0.13 | 9 | AT2 Cell |
| Mecom | 0.51 | 0.00 | 0.00 | 0.43 | 0.09 | 9 | AT2 Cell |
| Lrrc16a | 0.35 | 0.00 | 0.00 | 0.26 | 0.04 | 9 | AT2 Cell |
| Gm6169 | 0.49 | 0.00 | 0.00 | 0.48 | 0.11 | 9 | AT2 Cell |
| Chi3l1 | 0.28 | 0.00 | 0.00 | 0.45 | 0.10 | 9 | AT2 Cell |
| Prpf18 | 0.53 | 0.00 | 0.00 | 0.48 | 0.12 | 9 | AT2 Cell |
| Anxa1 | 0.65 | 0.00 | 0.00 | 0.60 | 0.17 | 9 | AT2 Cell |
| Ldhb | 0.44 | 0.00 | 0.00 | 0.36 | 0.07 | 9 | AT2 Cell |
| Dpp4 | 0.45 | 0.00 | 0.00 | 0.49 | 0.12 | 9 | AT2 Cell |
| Zfp36 | 0.58 | 0.00 | 0.00 | 0.56 | 0.15 | 9 | AT2 Cell |
| Ppfibp2 | 0.36 | 0.00 | 0.00 | 0.31 | 0.05 | 9 | AT2 Cell |
| Arrdc1 | 0.43 | 0.00 | 0.00 | 0.41 | 0.09 | 9 | AT2 Cell |
| Tmem164 | 0.44 | 0.00 | 0.00 | 0.32 | 0.06 | 9 | AT2 Cell |
| Fos | 0.67 | 0.00 | 0.00 | 0.59 | 0.18 | 9 | AT2 Cell |
| Neat1 | 0.70 | 0.00 | 0.00 | 0.75 | 0.26 | 9 | AT2 Cell |
| Agpat3 | 0.39 | 0.00 | 0.00 | 0.35 | 0.07 | 9 | AT2 Cell |
| Jund | 0.67 | 0.00 | 0.00 | 0.73 | 0.26 | 9 | AT2 Cell |
| Emc6 | 0.50 | 0.00 | 0.00 | 0.50 | 0.13 | 9 | AT2 Cell |
| Klf9 | 0.53 | 0.00 | 0.00 | 0.61 | 0.18 | 9 | AT2 Cell |
| Vegfa | 0.60 | 0.00 | 0.00 | 0.54 | 0.15 | 9 | AT2 Cell |
| Spg21 | 0.46 | 0.00 | 0.00 | 0.35 | 0.07 | 9 | AT2 Cell |
| Srebf2 | 0.36 | 0.00 | 0.00 | 0.27 | 0.05 | 9 | AT2 Cell |
| Gm9847 | 0.52 | 0.00 | 0.00 | 0.53 | 0.15 | 9 | AT2 Cell |
| Anxa3 | 0.57 | 0.00 | 0.00 | 0.65 | 0.21 | 9 | AT2 Cell |
| Slc38a2 | 0.58 | 0.00 | 0.00 | 0.59 | 0.18 | 9 | AT2 Cell |
| Tacc2 | 0.41 | 0.00 | 0.00 | 0.34 | 0.07 | 9 | AT2 Cell |
| Tpt1 | 0.57 | 0.00 | 0.00 | 0.99 | 0.87 | 9 | AT2 Cell |
| Eif3b | 0.46 | 0.00 | 0.00 | 0.44 | 0.11 | 9 | AT2 Cell |
| Lrrk2 | 0.35 | 0.00 | 0.00 | 0.26 | 0.04 | 9 | AT2 Cell |
| Atp11a | 0.42 | 0.00 | 0.00 | 0.43 | 0.10 | 9 | AT2 Cell |
| Per3 | 0.33 | 0.00 | 0.00 | 0.26 | 0.04 | 9 | AT2 Cell |
| Dmtf1 | 0.48 | 0.00 | 0.00 | 0.42 | 0.10 | 9 | AT2 Cell |
| Hdgf | 0.58 | 0.00 | 0.00 | 0.68 | 0.23 | 9 | AT2 Cell |
| Atp6v1g1 | 0.65 | 0.00 | 0.00 | 0.77 | 0.30 | 9 | AT2 Cell |
| Syn3 | 0.58 | 0.00 | 0.00 | 0.51 | 0.15 | 9 | AT2 Cell |
| Iqgap1 | 0.66 | 0.00 | 0.00 | 0.89 | 0.43 | 9 | AT2 Cell |
| Gnas | 0.65 | 0.00 | 0.00 | 0.96 | 0.66 | 9 | AT2 Cell |
| Wdfy1 | 0.32 | 0.00 | 0.00 | 0.25 | 0.04 | 9 | AT2 Cell |
| Carkd | 0.46 | 0.00 | 0.00 | 0.51 | 0.15 | 9 | AT2 Cell |
| Rpl21-ps4 | 0.64 | 0.00 | 0.00 | 0.80 | 0.34 | 9 | AT2 Cell |
| Tspan15 | 0.34 | 0.00 | 0.00 | 0.31 | 0.06 | 9 | AT2 Cell |
| Asah1 | 0.41 | 0.00 | 0.00 | 0.44 | 0.11 | 9 | AT2 Cell |
| Tbcel | 0.34 | 0.00 | 0.00 | 0.30 | 0.06 | 9 | AT2 Cell |
| Gstk1 | 0.41 | 0.00 | 0.00 | 0.32 | 0.07 | 9 | AT2 Cell |
| Senp2 | 0.44 | 0.00 | 0.00 | 0.37 | 0.09 | 9 | AT2 Cell |
| Klhl24 | 0.41 | 0.00 | 0.00 | 0.43 | 0.11 | 9 | AT2 Cell |
| Mpp5 | 0.44 | 0.00 | 0.00 | 0.42 | 0.11 | 9 | AT2 Cell |
| Glrx | 0.39 | 0.00 | 0.00 | 0.32 | 0.07 | 9 | AT2 Cell |
| Ppa2 | 0.38 | 0.00 | 0.00 | 0.39 | 0.09 | 9 | AT2 Cell |
| Cux1 | 0.47 | 0.00 | 0.00 | 0.52 | 0.15 | 9 | AT2 Cell |
| Tnfrsf12a | 0.38 | 0.00 | 0.00 | 0.27 | 0.05 | 9 | AT2 Cell |
| Nfe2l2 | 0.59 | 0.00 | 0.00 | 0.66 | 0.24 | 9 | AT2 Cell |
| Gata6 | 0.32 | 0.00 | 0.00 | 0.35 | 0.08 | 9 | AT2 Cell |
| Nupr1 | 1.32 | 0.00 | 0.00 | 0.52 | 0.17 | 9 | AT2 Cell |
| Rps8 | 0.55 | 0.00 | 0.00 | 0.98 | 0.81 | 9 | AT2 Cell |
| Gm26384 | 0.65 | 0.00 | 0.00 | 0.88 | 0.47 | 9 | AT2 Cell |
| Agrn | 0.48 | 0.00 | 0.00 | 0.45 | 0.12 | 9 | AT2 Cell |
| Rpl21-ps6 | 0.67 | 0.00 | 0.00 | 0.85 | 0.42 | 9 | AT2 Cell |
| Cers2 | 0.44 | 0.00 | 0.00 | 0.48 | 0.13 | 9 | AT2 Cell |
| Atp1a1 | 0.46 | 0.00 | 0.00 | 0.48 | 0.14 | 9 | AT2 Cell |
| Irf2bp2 | 0.46 | 0.00 | 0.00 | 0.50 | 0.15 | 9 | AT2 Cell |
| Ifi30 | 0.30 | 0.00 | 0.00 | 0.41 | 0.10 | 9 | AT2 Cell |
| Cited2 | 0.44 | 0.00 | 0.00 | 0.38 | 0.09 | 9 | AT2 Cell |
| Lyz1 | 0.50 | 0.00 | 0.00 | 0.56 | 0.18 | 9 | AT2 Cell |
| Kank2 | 0.37 | 0.00 | 0.00 | 0.41 | 0.11 | 9 | AT2 Cell |
| Stard3 | 0.30 | 0.00 | 0.00 | 0.27 | 0.05 | 9 | AT2 Cell |
| Tanc2 | 0.35 | 0.00 | 0.00 | 0.32 | 0.07 | 9 | AT2 Cell |
| Rps28 | 0.53 | 0.00 | 0.00 | 0.98 | 0.83 | 9 | AT2 Cell |
| Por | 0.44 | 0.00 | 0.00 | 0.43 | 0.12 | 9 | AT2 Cell |
| Rab27b | 0.32 | 0.00 | 0.00 | 0.30 | 0.06 | 9 | AT2 Cell |
| Tob1 | 0.31 | 0.00 | 0.00 | 0.28 | 0.06 | 9 | AT2 Cell |
| Lama5 | 0.35 | 0.00 | 0.00 | 0.35 | 0.08 | 9 | AT2 Cell |
| Gm10263 | 0.54 | 0.00 | 0.00 | 0.96 | 0.76 | 9 | AT2 Cell |
| F11r | 0.40 | 0.00 | 0.00 | 0.51 | 0.15 | 9 | AT2 Cell |
| Bcam | 0.32 | 0.00 | 0.00 | 0.35 | 0.09 | 9 | AT2 Cell |
| Cpm | 0.32 | 0.00 | 0.00 | 0.55 | 0.17 | 9 | AT2 Cell |
| Camk2n1 | 0.40 | 0.00 | 0.00 | 0.47 | 0.13 | 9 | AT2 Cell |
| Rnf114 | 0.32 | 0.00 | 0.00 | 0.34 | 0.08 | 9 | AT2 Cell |
| Ptplad1 | 0.33 | 0.00 | 0.00 | 0.35 | 0.08 | 9 | AT2 Cell |
| Ppm1b | 0.32 | 0.00 | 0.00 | 0.26 | 0.05 | 9 | AT2 Cell |
| Mir682 | 0.47 | 0.00 | 0.00 | 0.98 | 0.85 | 9 | AT2 Cell |
| Ctage5 | 0.53 | 0.00 | 0.00 | 0.69 | 0.27 | 9 | AT2 Cell |
| Cd9 | 0.46 | 0.00 | 0.00 | 0.81 | 0.33 | 9 | AT2 Cell |
| Gm22758 | 0.55 | 0.00 | 0.00 | 0.95 | 0.68 | 9 | AT2 Cell |
| Atp6v1a | 0.51 | 0.00 | 0.00 | 0.69 | 0.26 | 9 | AT2 Cell |
| Gm22759 | 0.57 | 0.00 | 0.00 | 0.94 | 0.63 | 9 | AT2 Cell |
| Rpl37a | 0.49 | 0.00 | 0.00 | 0.98 | 0.83 | 9 | AT2 Cell |
| Polr2e | 0.40 | 0.00 | 0.00 | 0.48 | 0.15 | 9 | AT2 Cell |
| Bmp1 | 0.34 | 0.00 | 0.00 | 0.31 | 0.07 | 9 | AT2 Cell |
| Fam177a | 0.47 | 0.00 | 0.00 | 0.62 | 0.22 | 9 | AT2 Cell |
| Rps27 | 0.50 | 0.00 | 0.00 | 0.98 | 0.81 | 9 | AT2 Cell |
| Mdfic | 0.27 | 0.00 | 0.00 | 0.28 | 0.06 | 9 | AT2 Cell |
| Nkd1 | 0.35 | 0.00 | 0.00 | 0.39 | 0.10 | 9 | AT2 Cell |
| Calcoco1 | 0.26 | 0.00 | 0.00 | 0.25 | 0.05 | 9 | AT2 Cell |
| 1700047I17Rik2 | 0.39 | 0.00 | 0.00 | 0.54 | 0.18 | 9 | AT2 Cell |
| Gm22761 | 0.42 | 0.00 | 0.00 | 0.51 | 0.16 | 9 | AT2 Cell |
| Anxa4 | 0.32 | 0.00 | 0.00 | 0.35 | 0.09 | 9 | AT2 Cell |
| Gm24865 | 0.55 | 0.00 | 0.00 | 0.93 | 0.62 | 9 | AT2 Cell |
| Gm6133 | 0.54 | 0.00 | 0.00 | 0.76 | 0.33 | 9 | AT2 Cell |
| Cd36 | 0.40 | 0.00 | 0.00 | 0.79 | 0.32 | 9 | AT2 Cell |
| Crebrf | 0.33 | 0.00 | 0.00 | 0.40 | 0.11 | 9 | AT2 Cell |
| Dusp1 | 0.43 | 0.00 | 0.00 | 0.49 | 0.16 | 9 | AT2 Cell |
| Tnfaip1 | 0.38 | 0.00 | 0.00 | 0.49 | 0.15 | 9 | AT2 Cell |
| Gdpd1 | 0.31 | 0.00 | 0.00 | 0.27 | 0.06 | 9 | AT2 Cell |
| Cd55 | 0.28 | 0.00 | 0.00 | 0.29 | 0.07 | 9 | AT2 Cell |
| Soat1 | 0.33 | 0.00 | 0.00 | 0.37 | 0.10 | 9 | AT2 Cell |
| Gm22426 | 0.55 | 0.00 | 0.00 | 0.89 | 0.47 | 9 | AT2 Cell |
| Lcn2 | 0.27 | 0.00 | 0.00 | 0.28 | 0.06 | 9 | AT2 Cell |
| Ppp3ca | 0.52 | 0.00 | 0.00 | 0.66 | 0.26 | 9 | AT2 Cell |
| Arf6 | 0.42 | 0.00 | 0.00 | 0.53 | 0.18 | 9 | AT2 Cell |
| Nav2 | 0.29 | 0.00 | 0.00 | 0.32 | 0.08 | 9 | AT2 Cell |
| Abhd5 | 0.32 | 0.00 | 0.00 | 0.28 | 0.06 | 9 | AT2 Cell |
| Tgoln1 | 0.37 | 0.00 | 0.00 | 0.53 | 0.18 | 9 | AT2 Cell |
| Gm26300 | 0.54 | 0.00 | 0.00 | 0.79 | 0.36 | 9 | AT2 Cell |
| Ypel3 | 0.35 | 0.00 | 0.00 | 0.57 | 0.19 | 9 | AT2 Cell |
| Hmga1-rs1 | 0.27 | 0.00 | 0.00 | 0.29 | 0.07 | 9 | AT2 Cell |
| Diap1 | 0.31 | 0.00 | 0.00 | 0.44 | 0.13 | 9 | AT2 Cell |
| Mtfr1l | 0.31 | 0.00 | 0.00 | 0.37 | 0.10 | 9 | AT2 Cell |
| Polr1c | 0.32 | 0.00 | 0.00 | 0.34 | 0.09 | 9 | AT2 Cell |
| Cat | 0.36 | 0.00 | 0.00 | 0.40 | 0.12 | 9 | AT2 Cell |
| Pdlim2 | 0.37 | 0.00 | 0.00 | 0.50 | 0.17 | 9 | AT2 Cell |
| Hmga1 | 0.27 | 0.00 | 0.00 | 0.30 | 0.07 | 9 | AT2 Cell |
| Pink1 | 0.30 | 0.00 | 0.00 | 0.37 | 0.10 | 9 | AT2 Cell |
| Wls | 0.37 | 0.00 | 0.00 | 0.59 | 0.21 | 9 | AT2 Cell |
| Ddt | 0.27 | 0.00 | 0.00 | 0.30 | 0.07 | 9 | AT2 Cell |
| Gm24951 | 0.27 | 0.00 | 0.00 | 0.31 | 0.08 | 9 | AT2 Cell |
| Atp6v1b2 | 0.32 | 0.00 | 0.00 | 0.40 | 0.12 | 9 | AT2 Cell |
| Sos2 | 0.28 | 0.00 | 0.00 | 0.26 | 0.06 | 9 | AT2 Cell |
| Rpl17 | 0.53 | 0.00 | 0.00 | 0.89 | 0.53 | 9 | AT2 Cell |
| Magi3 | 0.36 | 0.00 | 0.00 | 0.37 | 0.11 | 9 | AT2 Cell |
| Dcaf11 | 0.26 | 0.00 | 0.00 | 0.26 | 0.06 | 9 | AT2 Cell |
| Glul | 0.34 | 0.00 | 0.00 | 0.58 | 0.21 | 9 | AT2 Cell |
| Pptc7 | 0.28 | 0.00 | 0.00 | 0.30 | 0.08 | 9 | AT2 Cell |
| Slc25a51 | 0.27 | 0.00 | 0.00 | 0.32 | 0.08 | 9 | AT2 Cell |
| Pnrc1 | 0.35 | 0.00 | 0.00 | 0.55 | 0.19 | 9 | AT2 Cell |
| Elovl5 | 0.29 | 0.00 | 0.00 | 0.45 | 0.14 | 9 | AT2 Cell |
| Gm25593 | 0.37 | 0.00 | 0.00 | 0.56 | 0.20 | 9 | AT2 Cell |
| Clk4 | 0.29 | 0.00 | 0.00 | 0.36 | 0.10 | 9 | AT2 Cell |
| Sod1 | 0.56 | 0.00 | 0.00 | 0.85 | 0.46 | 9 | AT2 Cell |
| Cdc42ep3 | 0.29 | 0.00 | 0.00 | 0.44 | 0.14 | 9 | AT2 Cell |
| Dazap1 | 0.30 | 0.00 | 0.00 | 0.37 | 0.11 | 9 | AT2 Cell |
| Gm9846 | 0.40 | 0.00 | 0.00 | 0.99 | 0.89 | 9 | AT2 Cell |
| Csgalnact2 | 0.28 | 0.00 | 0.00 | 0.30 | 0.08 | 9 | AT2 Cell |
| Rps12-ps3 | 0.45 | 0.00 | 0.00 | 0.97 | 0.78 | 9 | AT2 Cell |
| Rps12 | 0.49 | 0.00 | 0.00 | 0.94 | 0.68 | 9 | AT2 Cell |
| Gzf1 | 0.31 | 0.00 | 0.00 | 0.30 | 0.08 | 9 | AT2 Cell |
| Idi1 | 0.33 | 0.00 | 0.00 | 0.29 | 0.07 | 9 | AT2 Cell |
| Rpl31 | 0.53 | 0.00 | 0.00 | 0.90 | 0.57 | 9 | AT2 Cell |
| Slc31a1 | 0.30 | 0.00 | 0.00 | 0.36 | 0.11 | 9 | AT2 Cell |
| Dapk1 | 0.29 | 0.00 | 0.00 | 0.36 | 0.10 | 9 | AT2 Cell |
| Ngfrap1 | 0.44 | 0.00 | 0.00 | 0.59 | 0.22 | 9 | AT2 Cell |
| Echs1 | 0.32 | 0.00 | 0.00 | 0.44 | 0.14 | 9 | AT2 Cell |
| Gm24276 | 0.37 | 0.00 | 0.00 | 0.57 | 0.21 | 9 | AT2 Cell |
| Pon3 | 0.28 | 0.00 | 0.00 | 0.33 | 0.09 | 9 | AT2 Cell |
| Amotl2 | 0.28 | 0.00 | 0.00 | 0.32 | 0.09 | 9 | AT2 Cell |
| Adk | 0.26 | 0.00 | 0.00 | 0.38 | 0.11 | 9 | AT2 Cell |
| Hist1h2al | 0.27 | 0.00 | 0.00 | 0.33 | 0.09 | 9 | AT2 Cell |
| Ctgf | 0.35 | 0.00 | 0.00 | 0.30 | 0.08 | 9 | AT2 Cell |
| Gm8973 | 0.26 | 0.00 | 0.00 | 0.30 | 0.08 | 9 | AT2 Cell |
| Sgms1 | 0.27 | 0.00 | 0.00 | 0.31 | 0.08 | 9 | AT2 Cell |
| Itga9 | 0.27 | 0.00 | 0.00 | 0.32 | 0.09 | 9 | AT2 Cell |
| Ptov1 | 0.27 | 0.00 | 0.00 | 0.43 | 0.13 | 9 | AT2 Cell |
| Ier3 | 0.33 | 0.00 | 0.00 | 0.41 | 0.13 | 9 | AT2 Cell |
| Rnf181 | 0.31 | 0.00 | 0.00 | 0.38 | 0.12 | 9 | AT2 Cell |
| Actn1 | 0.37 | 0.00 | 0.00 | 0.57 | 0.22 | 9 | AT2 Cell |
| Zfyve21 | 0.27 | 0.00 | 0.00 | 0.31 | 0.09 | 9 | AT2 Cell |
| Snx7 | 0.27 | 0.00 | 0.00 | 0.27 | 0.07 | 9 | AT2 Cell |
| Nucb2 | 0.33 | 0.00 | 0.00 | 0.45 | 0.15 | 9 | AT2 Cell |
| Dstn | 0.47 | 0.00 | 0.00 | 0.88 | 0.49 | 9 | AT2 Cell |
| mt-Rnr2 | 0.45 | 0.00 | 0.00 | 1.00 | 0.99 | 9 | AT2 Cell |
| Pcyt1a | 0.32 | 0.00 | 0.00 | 0.34 | 0.10 | 9 | AT2 Cell |
| Trp53bp2 | 0.29 | 0.00 | 0.00 | 0.31 | 0.08 | 9 | AT2 Cell |
| Rpl27a | 0.44 | 0.00 | 0.00 | 0.96 | 0.78 | 9 | AT2 Cell |
| Usp8 | 0.33 | 0.00 | 0.00 | 0.46 | 0.16 | 9 | AT2 Cell |
| Cirbp | 0.30 | 0.00 | 0.00 | 0.55 | 0.20 | 9 | AT2 Cell |
| Junb | 0.33 | 0.00 | 0.00 | 0.52 | 0.19 | 9 | AT2 Cell |
| Clic5 | 0.34 | 0.00 | 0.00 | 0.57 | 0.22 | 9 | AT2 Cell |
| Tob2 | 0.29 | 0.00 | 0.00 | 0.33 | 0.10 | 9 | AT2 Cell |
| Osbpl9 | 0.28 | 0.00 | 0.00 | 0.39 | 0.12 | 9 | AT2 Cell |
| Drap1 | 0.32 | 0.00 | 0.00 | 0.60 | 0.24 | 9 | AT2 Cell |
| Col6a1 | 0.36 | 0.00 | 0.00 | 0.51 | 0.18 | 9 | AT2 Cell |
| Abhd17b | 0.26 | 0.00 | 0.00 | 0.30 | 0.08 | 9 | AT2 Cell |
| Qk | 0.39 | 0.00 | 0.00 | 0.66 | 0.29 | 9 | AT2 Cell |
| Mir692-2a | 0.27 | 0.00 | 0.00 | 0.51 | 0.19 | 9 | AT2 Cell |
| Gm22760 | 0.28 | 0.00 | 0.00 | 0.42 | 0.14 | 9 | AT2 Cell |
| Tead1 | 0.33 | 0.00 | 0.00 | 0.39 | 0.13 | 9 | AT2 Cell |
| Phlda1 | 0.37 | 0.00 | 0.00 | 0.52 | 0.19 | 9 | AT2 Cell |
| Atp2a2 | 0.26 | 0.00 | 0.00 | 0.38 | 0.12 | 9 | AT2 Cell |
| Pik3ca | 0.34 | 0.00 | 0.00 | 0.50 | 0.18 | 9 | AT2 Cell |
| mt-Td | 0.28 | 0.00 | 0.00 | 0.45 | 0.16 | 9 | AT2 Cell |
| Ddx39b | 0.36 | 0.00 | 0.00 | 0.51 | 0.20 | 9 | AT2 Cell |
| Chd2 | 0.32 | 0.00 | 0.00 | 0.44 | 0.15 | 9 | AT2 Cell |
| Eif1 | 0.47 | 0.00 | 0.00 | 0.92 | 0.67 | 9 | AT2 Cell |
| Las1l | 0.44 | 0.00 | 0.00 | 0.39 | 0.13 | 9 | AT2 Cell |
| Gm22751 | 0.27 | 0.00 | 0.00 | 0.44 | 0.15 | 9 | AT2 Cell |
| RPL24 | 0.42 | 0.00 | 0.00 | 0.84 | 0.45 | 9 | AT2 Cell |
| Cir1 | 0.30 | 0.00 | 0.00 | 0.52 | 0.20 | 9 | AT2 Cell |
| Gm20721 | 0.38 | 0.00 | 0.00 | 0.40 | 0.14 | 9 | AT2 Cell |
| Rpl36 | 0.44 | 0.00 | 0.00 | 0.91 | 0.62 | 9 | AT2 Cell |
| Rock1 | 0.43 | 0.00 | 0.00 | 0.71 | 0.34 | 9 | AT2 Cell |
| Rpl36-ps3 | 0.45 | 0.00 | 0.00 | 0.86 | 0.52 | 9 | AT2 Cell |
| Rpl41 | 0.30 | 0.00 | 0.00 | 0.99 | 0.94 | 9 | AT2 Cell |
| Cdc42bpa | 0.27 | 0.00 | 0.00 | 0.41 | 0.14 | 9 | AT2 Cell |
| Rb1cc1 | 0.37 | 0.00 | 0.00 | 0.65 | 0.29 | 9 | AT2 Cell |
| Gm7536 | 0.39 | 0.00 | 0.00 | 0.97 | 0.81 | 9 | AT2 Cell |
| Mrpl14 | 0.28 | 0.00 | 0.00 | 0.41 | 0.14 | 9 | AT2 Cell |
| Creg1 | 0.27 | 0.00 | 0.00 | 0.45 | 0.16 | 9 | AT2 Cell |
| Ube2i | 0.36 | 0.00 | 0.00 | 0.64 | 0.28 | 9 | AT2 Cell |
| Txnrd1 | 0.30 | 0.00 | 0.00 | 0.33 | 0.11 | 9 | AT2 Cell |
| Actn4 | 0.33 | 0.00 | 0.00 | 0.64 | 0.28 | 9 | AT2 Cell |
| Ing2 | 0.26 | 0.00 | 0.00 | 0.32 | 0.10 | 9 | AT2 Cell |
| Maf1 | 0.25 | 0.00 | 0.00 | 0.44 | 0.16 | 9 | AT2 Cell |
| Wnk1 | 0.28 | 0.00 | 0.00 | 0.46 | 0.17 | 9 | AT2 Cell |
| Degs1 | 0.28 | 0.00 | 0.00 | 0.56 | 0.22 | 9 | AT2 Cell |
| Rps15 | 0.37 | 0.00 | 0.00 | 0.96 | 0.78 | 9 | AT2 Cell |
| Phldb2 | 0.29 | 0.00 | 0.00 | 0.65 | 0.28 | 9 | AT2 Cell |
| Ftl1 | 0.39 | 0.00 | 0.00 | 0.90 | 0.56 | 9 | AT2 Cell |
| Acadl | 0.29 | 0.00 | 0.00 | 0.54 | 0.22 | 9 | AT2 Cell |
| Rpl34 | 0.37 | 0.00 | 0.00 | 0.97 | 0.79 | 9 | AT2 Cell |
| Apc | 0.26 | 0.00 | 0.00 | 0.56 | 0.22 | 9 | AT2 Cell |
| Sepp1 | 0.31 | 0.00 | 0.00 | 0.79 | 0.40 | 9 | AT2 Cell |
| Rps14 | 0.31 | 0.00 | 0.00 | 0.99 | 0.92 | 9 | AT2 Cell |
| Tmbim6 | 0.34 | 0.00 | 0.00 | 0.76 | 0.36 | 9 | AT2 Cell |
| Gm10076 | 0.30 | 0.00 | 0.00 | 0.99 | 0.92 | 9 | AT2 Cell |
| Rpl18a | 0.34 | 0.00 | 0.00 | 0.98 | 0.82 | 9 | AT2 Cell |
| Rabac1 | 0.31 | 0.00 | 0.00 | 0.62 | 0.27 | 9 | AT2 Cell |
| Rpl10a-ps2 | 0.29 | 0.00 | 0.00 | 0.63 | 0.27 | 9 | AT2 Cell |
| Slco3a1 | 0.26 | 0.00 | 0.00 | 0.26 | 0.08 | 9 | AT2 Cell |
| Rpl37 | 0.34 | 0.00 | 0.00 | 0.96 | 0.79 | 9 | AT2 Cell |
| Atp6v0d1 | 0.25 | 0.00 | 0.00 | 0.44 | 0.17 | 9 | AT2 Cell |
| Tceal8 | 0.27 | 0.00 | 0.00 | 0.47 | 0.19 | 9 | AT2 Cell |
| App | 0.33 | 0.00 | 0.00 | 0.93 | 0.58 | 9 | AT2 Cell |
| Cyb5 | 0.27 | 0.00 | 0.00 | 0.59 | 0.25 | 9 | AT2 Cell |
| Arid4b | 0.31 | 0.00 | 0.00 | 0.69 | 0.32 | 9 | AT2 Cell |
| Rpl26 | 0.33 | 0.00 | 0.00 | 0.97 | 0.80 | 9 | AT2 Cell |
| Phf3 | 0.28 | 0.00 | 0.00 | 0.55 | 0.24 | 9 | AT2 Cell |
| Gm10116 | 0.35 | 0.00 | 0.00 | 0.91 | 0.62 | 9 | AT2 Cell |
| Use1 | 0.26 | 0.00 | 0.00 | 0.59 | 0.26 | 9 | AT2 Cell |
| Gm9493 | 0.34 | 0.00 | 0.00 | 0.95 | 0.73 | 9 | AT2 Cell |
| Rpl22l1 | 0.41 | 0.00 | 0.00 | 0.88 | 0.55 | 9 | AT2 Cell |
| Fau | 0.28 | 0.00 | 0.00 | 0.98 | 0.86 | 9 | AT2 Cell |
| Golga4 | 0.32 | 0.00 | 0.00 | 0.62 | 0.28 | 9 | AT2 Cell |
| Ankrd12 | 0.30 | 0.00 | 0.00 | 0.57 | 0.26 | 9 | AT2 Cell |
| Gm7589 | 0.37 | 0.00 | 0.00 | 0.92 | 0.64 | 9 | AT2 Cell |
| Zfp91 | 0.29 | 0.00 | 0.00 | 0.64 | 0.29 | 9 | AT2 Cell |
| Gm5093 | 0.37 | 0.00 | 0.00 | 0.81 | 0.46 | 9 | AT2 Cell |
| Klf6 | 0.39 | 0.00 | 0.00 | 0.68 | 0.35 | 9 | AT2 Cell |
| Gm10335 | 0.35 | 0.00 | 0.00 | 0.95 | 0.72 | 9 | AT2 Cell |
| Gm10132 | 0.35 | 0.00 | 0.00 | 0.94 | 0.72 | 9 | AT2 Cell |
| Bsg | 0.32 | 0.00 | 0.00 | 0.71 | 0.35 | 9 | AT2 Cell |
| Rps7 | 0.33 | 0.00 | 0.00 | 0.95 | 0.73 | 9 | AT2 Cell |
| Ubc | 0.40 | 0.00 | 0.00 | 0.81 | 0.47 | 9 | AT2 Cell |
| Rps24 | 0.29 | 0.00 | 0.00 | 0.98 | 0.88 | 9 | AT2 Cell |
| Mtch2 | 0.25 | 0.00 | 0.00 | 0.42 | 0.17 | 9 | AT2 Cell |
| Gm11808 | 0.31 | 0.00 | 0.00 | 0.96 | 0.77 | 9 | AT2 Cell |
| Gm3940 | 0.30 | 0.00 | 0.00 | 0.69 | 0.35 | 9 | AT2 Cell |
| Akap9 | 0.30 | 0.00 | 0.00 | 0.70 | 0.35 | 9 | AT2 Cell |
| Gm7808 | 0.35 | 0.00 | 0.00 | 0.89 | 0.59 | 9 | AT2 Cell |
| Srsf11 | 0.38 | 0.00 | 0.00 | 0.81 | 0.48 | 9 | AT2 Cell |
| S100a6 | 0.50 | 0.00 | 0.00 | 0.58 | 0.31 | 9 | AT2 Cell |
| Jun | 0.31 | 0.00 | 0.00 | 0.69 | 0.35 | 9 | AT2 Cell |
| Gm5239 | 0.33 | 0.00 | 0.00 | 0.84 | 0.50 | 9 | AT2 Cell |
| Rpl23a-ps3 | 0.31 | 0.00 | 0.00 | 0.95 | 0.74 | 9 | AT2 Cell |
| Gm6472 | 0.29 | 0.00 | 0.00 | 0.96 | 0.79 | 9 | AT2 Cell |
| Rpl32 | 0.27 | 0.00 | 0.00 | 0.97 | 0.84 | 9 | AT2 Cell |
| Rps26-ps1 | 0.28 | 0.00 | 0.00 | 0.98 | 0.87 | 9 | AT2 Cell |
| Slc25a5 | 0.39 | 0.00 | 0.00 | 0.89 | 0.58 | 9 | AT2 Cell |
| Orc5 | 0.36 | 0.00 | 0.00 | 0.86 | 0.54 | 9 | AT2 Cell |
| Rpl38 | 0.30 | 0.00 | 0.00 | 0.95 | 0.72 | 9 | AT2 Cell |
| Rpl28 | 0.33 | 0.00 | 0.00 | 0.85 | 0.52 | 9 | AT2 Cell |
| Sqstm1 | 0.30 | 0.00 | 0.00 | 0.63 | 0.31 | 9 | AT2 Cell |
| Rpl11 | 0.32 | 0.00 | 0.00 | 0.92 | 0.67 | 9 | AT2 Cell |
| Rpl10a | 0.29 | 0.00 | 0.00 | 0.94 | 0.74 | 9 | AT2 Cell |
| Gm5428 | 0.28 | 0.00 | 0.00 | 0.96 | 0.78 | 9 | AT2 Cell |
| Cox7a2l | 0.28 | 0.00 | 0.00 | 0.76 | 0.40 | 9 | AT2 Cell |
| 2810474O19Rik | 0.27 | 0.00 | 0.00 | 0.54 | 0.26 | 9 | AT2 Cell |
| Rpl29 | 0.29 | 0.00 | 0.00 | 0.80 | 0.45 | 9 | AT2 Cell |
| Rpl23a | 0.32 | 0.00 | 0.00 | 0.92 | 0.66 | 9 | AT2 Cell |
| Gm17511 | 0.29 | 0.00 | 0.00 | 0.91 | 0.60 | 9 | AT2 Cell |
| Emd | 0.26 | 0.00 | 0.00 | 0.75 | 0.39 | 9 | AT2 Cell |
| Egr1 | 0.29 | 0.00 | 0.00 | 0.33 | 0.13 | 9 | AT2 Cell |
| Rpl5 | 0.28 | 0.00 | 0.00 | 0.94 | 0.74 | 9 | AT2 Cell |
| Rps17 | 0.29 | 0.00 | 0.00 | 0.95 | 0.76 | 9 | AT2 Cell |
| Rpl10 | 0.28 | 0.00 | 0.00 | 0.94 | 0.70 | 9 | AT2 Cell |
| Dbi | 0.26 | 0.00 | 0.00 | 0.68 | 0.36 | 9 | AT2 Cell |
| Gm10288 | 0.27 | 0.00 | 0.00 | 0.96 | 0.78 | 9 | AT2 Cell |
| Gm10036 | 0.28 | 0.00 | 0.00 | 0.94 | 0.74 | 9 | AT2 Cell |
| Rpl10-ps3 | 0.27 | 0.00 | 0.00 | 0.95 | 0.72 | 9 | AT2 Cell |
| Rbm25 | 0.30 | 0.00 | 0.00 | 0.92 | 0.68 | 9 | AT2 Cell |
| Eef2 | 0.30 | 0.00 | 0.00 | 0.87 | 0.56 | 9 | AT2 Cell |
| Rpl19 | 0.25 | 0.00 | 0.00 | 0.95 | 0.74 | 9 | AT2 Cell |
| Rpl18 | 0.25 | 0.00 | 0.00 | 0.90 | 0.63 | 9 | AT2 Cell |
| Atrx | 0.26 | 0.00 | 0.00 | 0.88 | 0.65 | 9 | AT2 Cell |
| Txn1 | 0.25 | 0.00 | 0.00 | 0.79 | 0.49 | 9 | AT2 Cell |
| Ahnak | 0.26 | 0.00 | 0.00 | 0.65 | 0.43 | 9 | AT2 Cell |
| Akap5 | 3.47 | 0.00 | 0.00 | 0.97 | 0.06 | 10 | AT1 Cell |
| Ager | 3.37 | 0.00 | 0.00 | 0.99 | 0.14 | 10 | AT1 Cell |
| Cldn18 | 2.97 | 0.00 | 0.00 | 0.98 | 0.15 | 10 | AT1 Cell |
| Rtkn2 | 2.69 | 0.00 | 0.00 | 0.83 | 0.03 | 10 | AT1 Cell |
| Emp2 | 2.46 | 0.00 | 0.00 | 0.98 | 0.32 | 10 | AT1 Cell |
| Lmo7 | 2.34 | 0.00 | 0.00 | 0.86 | 0.10 | 10 | AT1 Cell |
| Aqp5 | 2.32 | 0.00 | 0.00 | 0.73 | 0.04 | 10 | AT1 Cell |
| Vegfa | 2.31 | 0.00 | 0.00 | 0.86 | 0.14 | 10 | AT1 Cell |
| Krt7 | 2.28 | 0.00 | 0.00 | 0.77 | 0.05 | 10 | AT1 Cell |
| Clic5 | 2.28 | 0.00 | 0.00 | 0.95 | 0.21 | 10 | AT1 Cell |
| Col4a3 | 2.23 | 0.00 | 0.00 | 0.67 | 0.03 | 10 | AT1 Cell |
| Spock2 | 2.22 | 0.00 | 0.00 | 0.71 | 0.03 | 10 | AT1 Cell |
| S100a6 | 2.22 | 0.00 | 0.00 | 0.95 | 0.29 | 10 | AT1 Cell |
| Gprc5a | 2.08 | 0.00 | 0.00 | 0.70 | 0.05 | 10 | AT1 Cell |
| Col4a4 | 2.03 | 0.00 | 0.00 | 0.60 | 0.02 | 10 | AT1 Cell |
| Msln | 2.03 | 0.00 | 0.00 | 0.61 | 0.03 | 10 | AT1 Cell |
| Clic3 | 2.01 | 0.00 | 0.00 | 0.72 | 0.07 | 10 | AT1 Cell |
| Hopx | 1.98 | 0.00 | 0.00 | 0.70 | 0.08 | 10 | AT1 Cell |
| Serpinb6b | 1.95 | 0.00 | 0.00 | 0.73 | 0.15 | 10 | AT1 Cell |
| Myh14 | 1.94 | 0.00 | 0.00 | 0.70 | 0.07 | 10 | AT1 Cell |
| 2200002D01Rik | 1.92 | 0.00 | 0.00 | 0.64 | 0.05 | 10 | AT1 Cell |
| Pdgfa | 1.92 | 0.00 | 0.00 | 0.67 | 0.09 | 10 | AT1 Cell |
| Tinagl1 | 1.91 | 0.00 | 0.00 | 0.63 | 0.09 | 10 | AT1 Cell |
| Sec14l3 | 1.86 | 0.00 | 0.00 | 0.88 | 0.08 | 10 | AT1 Cell |
| Sema3e | 1.85 | 0.00 | 0.00 | 0.54 | 0.03 | 10 | AT1 Cell |
| Scnn1g | 1.84 | 0.00 | 0.00 | 0.56 | 0.02 | 10 | AT1 Cell |
| Pdpn | 1.83 | 0.00 | 0.00 | 0.58 | 0.03 | 10 | AT1 Cell |
| S100a14 | 1.81 | 0.00 | 0.00 | 0.51 | 0.02 | 10 | AT1 Cell |
| Krt19 | 1.81 | 0.00 | 0.00 | 0.59 | 0.05 | 10 | AT1 Cell |
| Timp3 | 1.81 | 0.00 | 0.00 | 0.90 | 0.34 | 10 | AT1 Cell |
| Ndnf | 1.78 | 0.00 | 0.00 | 0.48 | 0.03 | 10 | AT1 Cell |
| Cdkn2b | 1.78 | 0.00 | 0.00 | 0.58 | 0.04 | 10 | AT1 Cell |
| Tspan8 | 1.76 | 0.00 | 0.00 | 0.68 | 0.10 | 10 | AT1 Cell |
| Sema3a | 1.73 | 0.00 | 0.00 | 0.47 | 0.02 | 10 | AT1 Cell |
| Slc39a8 | 1.72 | 0.00 | 0.00 | 0.56 | 0.06 | 10 | AT1 Cell |
| Scnn1a | 1.69 | 0.00 | 0.00 | 0.58 | 0.05 | 10 | AT1 Cell |
| Icam1 | 1.65 | 0.00 | 0.00 | 0.62 | 0.08 | 10 | AT1 Cell |
| Ndst1 | 1.65 | 0.00 | 0.00 | 0.56 | 0.10 | 10 | AT1 Cell |
| Prdx6 | 1.63 | 0.00 | 0.00 | 0.79 | 0.26 | 10 | AT1 Cell |
| Ano1 | 1.61 | 0.00 | 0.00 | 0.48 | 0.04 | 10 | AT1 Cell |
| Cryab | 1.59 | 0.00 | 0.00 | 0.67 | 0.12 | 10 | AT1 Cell |
| Gipc2 | 1.58 | 0.00 | 0.00 | 0.47 | 0.03 | 10 | AT1 Cell |
| Ahnak | 1.58 | 0.00 | 0.00 | 0.90 | 0.42 | 10 | AT1 Cell |
| Itgb6 | 1.55 | 0.00 | 0.00 | 0.39 | 0.01 | 10 | AT1 Cell |
| Fam174b | 1.55 | 0.00 | 0.00 | 0.46 | 0.04 | 10 | AT1 Cell |
| Anxa3 | 1.54 | 0.00 | 0.00 | 0.74 | 0.21 | 10 | AT1 Cell |
| Flrt3 | 1.53 | 0.00 | 0.00 | 0.41 | 0.02 | 10 | AT1 Cell |
| Tmem37 | 1.50 | 0.00 | 0.00 | 0.43 | 0.03 | 10 | AT1 Cell |
| Krt8 | 1.49 | 0.00 | 0.00 | 0.50 | 0.07 | 10 | AT1 Cell |
| Anxa1 | 1.49 | 0.00 | 0.00 | 0.67 | 0.17 | 10 | AT1 Cell |
| Mthfd1 | 1.49 | 0.00 | 0.00 | 0.49 | 0.07 | 10 | AT1 Cell |
| Rab11fip1 | 1.45 | 0.00 | 0.00 | 0.46 | 0.04 | 10 | AT1 Cell |
| Agrn | 1.44 | 0.00 | 0.00 | 0.58 | 0.12 | 10 | AT1 Cell |
| Syn3 | 1.44 | 0.00 | 0.00 | 0.57 | 0.14 | 10 | AT1 Cell |
| Alcam | 1.44 | 0.00 | 0.00 | 0.80 | 0.21 | 10 | AT1 Cell |
| Gramd2 | 1.44 | 0.00 | 0.00 | 0.45 | 0.04 | 10 | AT1 Cell |
| Cadm1 | 1.44 | 0.00 | 0.00 | 0.57 | 0.13 | 10 | AT1 Cell |
| Mal2 | 1.44 | 0.00 | 0.00 | 0.45 | 0.05 | 10 | AT1 Cell |
| F3 | 1.42 | 0.00 | 0.00 | 0.41 | 0.03 | 10 | AT1 Cell |
| Cystm1 | 1.41 | 0.00 | 0.00 | 0.49 | 0.07 | 10 | AT1 Cell |
| Dag1 | 1.40 | 0.00 | 0.00 | 0.48 | 0.08 | 10 | AT1 Cell |
| Rgcc | 1.40 | 0.00 | 0.00 | 0.58 | 0.16 | 10 | AT1 Cell |
| Ptprf | 1.38 | 0.00 | 0.00 | 0.63 | 0.14 | 10 | AT1 Cell |
| Krt18 | 1.34 | 0.00 | 0.00 | 0.46 | 0.07 | 10 | AT1 Cell |
| Igfbp7 | 1.33 | 0.00 | 0.00 | 0.89 | 0.38 | 10 | AT1 Cell |
| Tspan15 | 1.31 | 0.00 | 0.00 | 0.46 | 0.06 | 10 | AT1 Cell |
| Cd9 | 1.29 | 0.00 | 0.00 | 0.83 | 0.33 | 10 | AT1 Cell |
| Ctgf | 1.29 | 0.00 | 0.00 | 0.42 | 0.07 | 10 | AT1 Cell |
| Lamc2 | 1.27 | 0.00 | 0.00 | 0.37 | 0.03 | 10 | AT1 Cell |
| Scd2 | 1.27 | 0.00 | 0.00 | 0.61 | 0.20 | 10 | AT1 Cell |
| Ezr | 1.26 | 0.00 | 0.00 | 0.65 | 0.20 | 10 | AT1 Cell |
| Slc6a14 | 1.26 | 0.00 | 0.00 | 0.35 | 0.03 | 10 | AT1 Cell |
| Il18r1 | 1.24 | 0.00 | 0.00 | 0.42 | 0.06 | 10 | AT1 Cell |
| Cxadr | 1.24 | 0.00 | 0.00 | 0.37 | 0.05 | 10 | AT1 Cell |
| Wwp1 | 1.23 | 0.00 | 0.00 | 0.49 | 0.12 | 10 | AT1 Cell |
| Fbln5 | 1.23 | 0.00 | 0.00 | 0.63 | 0.16 | 10 | AT1 Cell |
| Atp1b1 | 1.22 | 0.00 | 0.00 | 0.61 | 0.17 | 10 | AT1 Cell |
| Mmp11 | 1.22 | 0.00 | 0.00 | 0.37 | 0.04 | 10 | AT1 Cell |
| Trp53bp2 | 1.22 | 0.00 | 0.00 | 0.42 | 0.08 | 10 | AT1 Cell |
| Nkx2-1 | 1.20 | 0.00 | 0.00 | 0.46 | 0.09 | 10 | AT1 Cell |
| Tead1 | 1.20 | 0.00 | 0.00 | 0.48 | 0.13 | 10 | AT1 Cell |
| Bcam | 1.18 | 0.00 | 0.00 | 0.41 | 0.08 | 10 | AT1 Cell |
| Epcam | 1.17 | 0.00 | 0.00 | 0.44 | 0.09 | 10 | AT1 Cell |
| Scnn1b | 1.17 | 0.00 | 0.00 | 0.32 | 0.02 | 10 | AT1 Cell |
| Nav2 | 1.15 | 0.00 | 0.00 | 0.40 | 0.08 | 10 | AT1 Cell |
| Pmp22 | 1.14 | 0.00 | 0.00 | 0.76 | 0.30 | 10 | AT1 Cell |
| Dennd3 | 1.14 | 0.00 | 0.00 | 0.39 | 0.06 | 10 | AT1 Cell |
| Serpinb9 | 1.13 | 0.00 | 0.00 | 0.35 | 0.07 | 10 | AT1 Cell |
| Llgl2 | 1.12 | 0.00 | 0.00 | 0.33 | 0.03 | 10 | AT1 Cell |
| Wwc1 | 1.11 | 0.00 | 0.00 | 0.36 | 0.05 | 10 | AT1 Cell |
| Hs2st1 | 1.11 | 0.00 | 0.00 | 0.32 | 0.05 | 10 | AT1 Cell |
| Spint2 | 1.11 | 0.00 | 0.00 | 0.45 | 0.10 | 10 | AT1 Cell |
| Akr1c14 | 1.10 | 0.00 | 0.00 | 0.27 | 0.02 | 10 | AT1 Cell |
| Gja1 | 1.09 | 0.00 | 0.00 | 0.42 | 0.08 | 10 | AT1 Cell |
| Cldn7 | 1.09 | 0.00 | 0.00 | 0.37 | 0.07 | 10 | AT1 Cell |
| Pkp2 | 1.08 | 0.00 | 0.00 | 0.30 | 0.03 | 10 | AT1 Cell |
| Arhgef26 | 1.07 | 0.00 | 0.00 | 0.36 | 0.06 | 10 | AT1 Cell |
| Pard6b | 1.06 | 0.00 | 0.00 | 0.28 | 0.02 | 10 | AT1 Cell |
| Eps8l2 | 1.06 | 0.00 | 0.00 | 0.30 | 0.03 | 10 | AT1 Cell |
| Slco3a1 | 1.06 | 0.00 | 0.00 | 0.36 | 0.07 | 10 | AT1 Cell |
| Tacstd2 | 1.05 | 0.00 | 0.00 | 0.27 | 0.02 | 10 | AT1 Cell |
| Col4a5 | 1.01 | 0.00 | 0.00 | 0.26 | 0.03 | 10 | AT1 Cell |
| Fam189a2 | 1.01 | 0.00 | 0.00 | 0.30 | 0.04 | 10 | AT1 Cell |
| Osbpl6 | 0.99 | 0.00 | 0.00 | 0.30 | 0.05 | 10 | AT1 Cell |
| Sparc | 0.97 | 0.00 | 0.00 | 0.99 | 0.73 | 10 | AT1 Cell |
| Ctsh | 0.95 | 0.00 | 0.00 | 0.64 | 0.19 | 10 | AT1 Cell |
| Npnt | 0.92 | 0.00 | 0.00 | 0.70 | 0.23 | 10 | AT1 Cell |
| Fads3 | 0.91 | 0.00 | 0.00 | 0.27 | 0.04 | 10 | AT1 Cell |
| Limch1 | 0.85 | 0.00 | 0.00 | 0.85 | 0.29 | 10 | AT1 Cell |
| Cldn3 | 1.09 | 0.00 | 0.00 | 0.45 | 0.11 | 10 | AT1 Cell |
| Malat1 | 0.67 | 0.00 | 0.00 | 1.00 | 1.00 | 10 | AT1 Cell |
| Neat1 | 1.13 | 0.00 | 0.00 | 0.68 | 0.26 | 10 | AT1 Cell |
| Wdr45 | 0.98 | 0.00 | 0.00 | 0.35 | 0.07 | 10 | AT1 Cell |
| Anxa2 | 1.06 | 0.00 | 0.00 | 0.83 | 0.46 | 10 | AT1 Cell |
| Ppp3ca | 1.17 | 0.00 | 0.00 | 0.66 | 0.26 | 10 | AT1 Cell |
| Pvrl3 | 1.06 | 0.00 | 0.00 | 0.39 | 0.09 | 10 | AT1 Cell |
| App | 0.95 | 0.00 | 0.00 | 0.91 | 0.59 | 10 | AT1 Cell |
| Slc1a5 | 1.09 | 0.00 | 0.00 | 0.40 | 0.10 | 10 | AT1 Cell |
| Cttnbp2nl | 1.11 | 0.00 | 0.00 | 0.41 | 0.10 | 10 | AT1 Cell |
| Mpp5 | 1.12 | 0.00 | 0.00 | 0.42 | 0.11 | 10 | AT1 Cell |
| Msn | 1.00 | 0.00 | 0.00 | 0.81 | 0.42 | 10 | AT1 Cell |
| Bend7 | 0.98 | 0.00 | 0.00 | 0.31 | 0.06 | 10 | AT1 Cell |
| Myl12a | 0.92 | 0.00 | 0.00 | 0.89 | 0.65 | 10 | AT1 Cell |
| Fam134b | 0.92 | 0.00 | 0.00 | 0.30 | 0.06 | 10 | AT1 Cell |
| Pde7a | 0.97 | 0.00 | 0.00 | 0.31 | 0.06 | 10 | AT1 Cell |
| F11r | 1.02 | 0.00 | 0.00 | 0.49 | 0.16 | 10 | AT1 Cell |
| Fgf1 | 0.95 | 0.00 | 0.00 | 0.29 | 0.06 | 10 | AT1 Cell |
| Nbl1 | 0.85 | 0.00 | 0.00 | 0.30 | 0.06 | 10 | AT1 Cell |
| Bsg | 1.06 | 0.00 | 0.00 | 0.70 | 0.35 | 10 | AT1 Cell |
| Gata6 | 0.95 | 0.00 | 0.00 | 0.34 | 0.08 | 10 | AT1 Cell |
| Magi3 | 1.08 | 0.00 | 0.00 | 0.39 | 0.11 | 10 | AT1 Cell |
| Cdh1 | 0.86 | 0.00 | 0.00 | 0.32 | 0.07 | 10 | AT1 Cell |
| Ier3 | 1.03 | 0.00 | 0.00 | 0.43 | 0.13 | 10 | AT1 Cell |
| S100a11 | 0.94 | 0.00 | 0.00 | 0.80 | 0.48 | 10 | AT1 Cell |
| Fzd2 | 0.91 | 0.00 | 0.00 | 0.27 | 0.05 | 10 | AT1 Cell |
| Tmsb4x | 0.54 | 0.00 | 0.00 | 0.99 | 0.88 | 10 | AT1 Cell |
| Klf6 | 1.01 | 0.00 | 0.00 | 0.69 | 0.35 | 10 | AT1 Cell |
| Crip2 | 0.85 | 0.00 | 0.00 | 0.77 | 0.40 | 10 | AT1 Cell |
| Samhd1 | 0.92 | 0.00 | 0.00 | 0.52 | 0.18 | 10 | AT1 Cell |
| Sdc1 | 0.95 | 0.00 | 0.00 | 0.39 | 0.11 | 10 | AT1 Cell |
| Tmem63b | 0.83 | 0.00 | 0.00 | 0.28 | 0.06 | 10 | AT1 Cell |
| Vamp8 | 0.91 | 0.00 | 0.00 | 0.67 | 0.33 | 10 | AT1 Cell |
| Prkci | 0.92 | 0.00 | 0.00 | 0.40 | 0.12 | 10 | AT1 Cell |
| Prdx6b | 0.79 | 0.00 | 0.00 | 0.25 | 0.05 | 10 | AT1 Cell |
| Hspg2 | 0.79 | 0.00 | 0.00 | 0.26 | 0.06 | 10 | AT1 Cell |
| Lgals3 | 0.68 | 0.00 | 0.00 | 0.44 | 0.14 | 10 | AT1 Cell |
| Mir22hg | 0.81 | 0.00 | 0.00 | 0.27 | 0.06 | 10 | AT1 Cell |
| Pgrmc1 | 0.96 | 0.00 | 0.00 | 0.47 | 0.18 | 10 | AT1 Cell |
| Gde1 | 0.91 | 0.00 | 0.00 | 0.41 | 0.14 | 10 | AT1 Cell |
| Nceh1 | 0.81 | 0.00 | 0.00 | 0.26 | 0.06 | 10 | AT1 Cell |
| Fam177a | 0.92 | 0.00 | 0.00 | 0.53 | 0.23 | 10 | AT1 Cell |
| Wwc2 | 0.89 | 0.00 | 0.00 | 0.38 | 0.12 | 10 | AT1 Cell |
| Dstn | 0.75 | 0.00 | 0.00 | 0.78 | 0.50 | 10 | AT1 Cell |
| Lrpap1 | 1.00 | 0.00 | 0.00 | 0.41 | 0.14 | 10 | AT1 Cell |
| Aqp1 | 0.80 | 0.00 | 0.00 | 0.49 | 0.19 | 10 | AT1 Cell |
| Qk | 0.89 | 0.00 | 0.00 | 0.60 | 0.29 | 10 | AT1 Cell |
| Gm6169 | 0.88 | 0.00 | 0.00 | 0.36 | 0.12 | 10 | AT1 Cell |
| Cyr61 | 0.79 | 0.00 | 0.00 | 0.38 | 0.12 | 10 | AT1 Cell |
| Plp2 | 0.91 | 0.00 | 0.00 | 0.37 | 0.12 | 10 | AT1 Cell |
| Myo1c | 0.77 | 0.00 | 0.00 | 0.31 | 0.09 | 10 | AT1 Cell |
| Nbeal1 | 0.81 | 0.00 | 0.00 | 0.30 | 0.09 | 10 | AT1 Cell |
| Phactr1 | 0.81 | 0.00 | 0.00 | 0.27 | 0.07 | 10 | AT1 Cell |
| 1700047I17Rik2 | 0.84 | 0.00 | 0.00 | 0.46 | 0.18 | 10 | AT1 Cell |
| Ctdspl | 0.79 | 0.00 | 0.00 | 0.33 | 0.10 | 10 | AT1 Cell |
| Cav1 | 0.60 | 0.00 | 0.00 | 0.66 | 0.30 | 10 | AT1 Cell |
| Creb3l2 | 0.86 | 0.00 | 0.00 | 0.37 | 0.13 | 10 | AT1 Cell |
| Cux1 | 0.90 | 0.00 | 0.00 | 0.42 | 0.16 | 10 | AT1 Cell |
| Sdpr | 0.59 | 0.00 | 0.00 | 0.84 | 0.51 | 10 | AT1 Cell |
| Tppp3 | 0.72 | 0.00 | 0.00 | 0.36 | 0.12 | 10 | AT1 Cell |
| Tns1 | 0.79 | 0.00 | 0.00 | 0.51 | 0.22 | 10 | AT1 Cell |
| Tspan2 | 0.83 | 0.00 | 0.00 | 0.35 | 0.12 | 10 | AT1 Cell |
| Efna1 | 0.75 | 0.00 | 0.00 | 0.27 | 0.07 | 10 | AT1 Cell |
| Chka | 0.76 | 0.00 | 0.00 | 0.31 | 0.10 | 10 | AT1 Cell |
| Bex4 | 0.52 | 0.00 | 0.00 | 0.38 | 0.12 | 10 | AT1 Cell |
| Gadd45b | 0.75 | 0.00 | 0.00 | 0.28 | 0.08 | 10 | AT1 Cell |
| Tes | 0.72 | 0.00 | 0.00 | 0.30 | 0.09 | 10 | AT1 Cell |
| Asah1 | 0.81 | 0.00 | 0.00 | 0.34 | 0.12 | 10 | AT1 Cell |
| Chpt1 | 0.80 | 0.00 | 0.00 | 0.34 | 0.11 | 10 | AT1 Cell |
| Cnn2 | 0.72 | 0.00 | 0.00 | 0.66 | 0.38 | 10 | AT1 Cell |
| Ptgs1 | 0.78 | 0.00 | 0.00 | 0.31 | 0.10 | 10 | AT1 Cell |
| Utrn | 0.80 | 0.00 | 0.00 | 0.40 | 0.16 | 10 | AT1 Cell |
| Galnt18 | 0.72 | 0.00 | 0.00 | 0.32 | 0.10 | 10 | AT1 Cell |
| Rabac1 | 0.78 | 0.00 | 0.00 | 0.54 | 0.27 | 10 | AT1 Cell |
| Atp8a1 | 0.82 | 0.00 | 0.00 | 0.38 | 0.15 | 10 | AT1 Cell |
| Wnk1 | 0.84 | 0.00 | 0.00 | 0.41 | 0.17 | 10 | AT1 Cell |
| Chchd10 | 0.63 | 0.00 | 0.00 | 0.36 | 0.13 | 10 | AT1 Cell |
| Meg3 | 0.49 | 0.00 | 0.00 | 0.50 | 0.21 | 10 | AT1 Cell |
| Tfdp2 | 0.77 | 0.00 | 0.00 | 0.28 | 0.09 | 10 | AT1 Cell |
| Prkcdbp | 0.78 | 0.00 | 0.00 | 0.31 | 0.11 | 10 | AT1 Cell |
| Tacc2 | 0.71 | 0.00 | 0.00 | 0.25 | 0.07 | 10 | AT1 Cell |
| Ctnna1 | 0.72 | 0.00 | 0.00 | 0.62 | 0.35 | 10 | AT1 Cell |
| Dpysl2 | 0.82 | 0.00 | 0.00 | 0.40 | 0.17 | 10 | AT1 Cell |
| Wls | 0.83 | 0.00 | 0.00 | 0.46 | 0.22 | 10 | AT1 Cell |
| Tjp2 | 0.76 | 0.00 | 0.00 | 0.39 | 0.16 | 10 | AT1 Cell |
| Gm7334 | 0.76 | 0.00 | 0.00 | 0.31 | 0.11 | 10 | AT1 Cell |
| Arl4c | 0.70 | 0.00 | 0.00 | 0.29 | 0.10 | 10 | AT1 Cell |
| Hes1 | 0.70 | 0.00 | 0.00 | 0.55 | 0.28 | 10 | AT1 Cell |
| Fnbp1l | 0.74 | 0.00 | 0.00 | 0.50 | 0.25 | 10 | AT1 Cell |
| Ddx47 | 0.78 | 0.00 | 0.00 | 0.36 | 0.15 | 10 | AT1 Cell |
| Ehd2 | 0.72 | 0.00 | 0.00 | 0.35 | 0.14 | 10 | AT1 Cell |
| Anxa4 | 0.71 | 0.00 | 0.00 | 0.27 | 0.09 | 10 | AT1 Cell |
| Capn2 | 0.72 | 0.00 | 0.00 | 0.36 | 0.15 | 10 | AT1 Cell |
| Calm2 | 0.58 | 0.00 | 0.00 | 0.82 | 0.64 | 10 | AT1 Cell |
| Sptbn1 | 0.57 | 0.00 | 0.00 | 0.78 | 0.54 | 10 | AT1 Cell |
| Cers2 | 0.76 | 0.00 | 0.00 | 0.34 | 0.14 | 10 | AT1 Cell |
| Actb | 0.30 | 0.00 | 0.00 | 1.00 | 0.98 | 10 | AT1 Cell |
| Tpm3 | 0.52 | 0.00 | 0.00 | 0.86 | 0.69 | 10 | AT1 Cell |
| Amotl2 | 0.63 | 0.00 | 0.00 | 0.26 | 0.09 | 10 | AT1 Cell |
| Gng5 | 0.70 | 0.00 | 0.00 | 0.65 | 0.45 | 10 | AT1 Cell |
| Btg3 | 0.73 | 0.00 | 0.00 | 0.29 | 0.11 | 10 | AT1 Cell |
| Cav2 | 0.60 | 0.00 | 0.00 | 0.42 | 0.18 | 10 | AT1 Cell |
| Rps19 | 0.53 | 0.00 | 0.00 | 0.91 | 0.82 | 10 | AT1 Cell |
| Mrpl14 | 0.73 | 0.00 | 0.00 | 0.35 | 0.15 | 10 | AT1 Cell |
| Gpx4 | 0.69 | 0.00 | 0.00 | 0.58 | 0.37 | 10 | AT1 Cell |
| Hdgf | 0.73 | 0.00 | 0.00 | 0.46 | 0.24 | 10 | AT1 Cell |
| Ece1 | 0.60 | 0.00 | 0.00 | 0.50 | 0.25 | 10 | AT1 Cell |
| Ptrf | 0.57 | 0.00 | 0.00 | 0.60 | 0.35 | 10 | AT1 Cell |
| Epn2 | 0.69 | 0.00 | 0.00 | 0.30 | 0.12 | 10 | AT1 Cell |
| Errfi1 | 0.56 | 0.00 | 0.00 | 0.27 | 0.09 | 10 | AT1 Cell |
| Mbip | 0.67 | 0.00 | 0.00 | 0.29 | 0.11 | 10 | AT1 Cell |
| Gsn | 0.32 | 0.00 | 0.00 | 0.50 | 0.24 | 10 | AT1 Cell |
| Tgfb2 | 0.60 | 0.00 | 0.00 | 0.33 | 0.14 | 10 | AT1 Cell |
| Ssr2 | 0.68 | 0.00 | 0.00 | 0.53 | 0.30 | 10 | AT1 Cell |
| Selenbp1 | 0.80 | 0.00 | 0.00 | 0.43 | 0.22 | 10 | AT1 Cell |
| Apc | 0.71 | 0.00 | 0.00 | 0.44 | 0.23 | 10 | AT1 Cell |
| Lrrc58 | 0.31 | 0.00 | 0.00 | 0.98 | 0.91 | 10 | AT1 Cell |
| Nckap5 | 0.64 | 0.00 | 0.00 | 0.31 | 0.12 | 10 | AT1 Cell |
| Col4a2 | 0.51 | 0.00 | 0.00 | 0.67 | 0.42 | 10 | AT1 Cell |
| Rhoa | 0.62 | 0.00 | 0.00 | 0.65 | 0.46 | 10 | AT1 Cell |
| Tpm3-rs7 | 0.45 | 0.00 | 0.00 | 0.84 | 0.69 | 10 | AT1 Cell |
| Myo1b | 0.65 | 0.00 | 0.00 | 0.36 | 0.16 | 10 | AT1 Cell |
| F2r | 0.61 | 0.00 | 0.00 | 0.32 | 0.14 | 10 | AT1 Cell |
| Fdx1 | 0.71 | 0.00 | 0.00 | 0.31 | 0.14 | 10 | AT1 Cell |
| Arhgap5 | 0.71 | 0.00 | 0.00 | 0.38 | 0.18 | 10 | AT1 Cell |
| Rabgap1l | 0.63 | 0.00 | 0.00 | 0.26 | 0.10 | 10 | AT1 Cell |
| Iqgap1 | 0.55 | 0.00 | 0.00 | 0.66 | 0.44 | 10 | AT1 Cell |
| Myl6 | 0.43 | 0.00 | 0.00 | 0.81 | 0.69 | 10 | AT1 Cell |
| Sod1 | 0.59 | 0.00 | 0.00 | 0.64 | 0.47 | 10 | AT1 Cell |
| Selenbp2 | 0.74 | 0.00 | 0.00 | 0.34 | 0.16 | 10 | AT1 Cell |
| Rnf181 | 0.60 | 0.00 | 0.00 | 0.27 | 0.12 | 10 | AT1 Cell |
| Crlf3 | 0.52 | 0.00 | 0.00 | 0.30 | 0.13 | 10 | AT1 Cell |
| Sptlc2 | 0.61 | 0.00 | 0.00 | 0.31 | 0.14 | 10 | AT1 Cell |
| 2810474O19Rik | 0.62 | 0.00 | 0.00 | 0.45 | 0.26 | 10 | AT1 Cell |
| Rbms1 | 0.56 | 0.00 | 0.00 | 0.57 | 0.38 | 10 | AT1 Cell |
| Anxa5 | 0.57 | 0.00 | 0.00 | 0.58 | 0.40 | 10 | AT1 Cell |
| Tjp1 | 0.50 | 0.00 | 0.00 | 0.33 | 0.16 | 10 | AT1 Cell |
| Stx7 | 0.58 | 0.00 | 0.00 | 0.30 | 0.14 | 10 | AT1 Cell |
| Son | 0.40 | 0.00 | 0.00 | 0.83 | 0.73 | 10 | AT1 Cell |
| Mprip | 0.54 | 0.00 | 0.00 | 0.38 | 0.21 | 10 | AT1 Cell |
| 4930523C07Rik | 0.58 | 0.00 | 0.00 | 0.34 | 0.18 | 10 | AT1 Cell |
| Kank2 | 0.52 | 0.00 | 0.00 | 0.25 | 0.11 | 10 | AT1 Cell |
| Ralbp1 | 0.53 | 0.00 | 0.00 | 0.49 | 0.32 | 10 | AT1 Cell |
| Arhgef2 | 0.56 | 0.00 | 0.00 | 0.36 | 0.20 | 10 | AT1 Cell |
| Pdlim2 | 0.56 | 0.00 | 0.00 | 0.33 | 0.17 | 10 | AT1 Cell |
| Tm9sf3 | 0.55 | 0.00 | 0.00 | 0.50 | 0.34 | 10 | AT1 Cell |
| Cdkn1a | 0.36 | 0.00 | 0.00 | 0.32 | 0.16 | 10 | AT1 Cell |
| Fam101b | 0.55 | 0.00 | 0.00 | 0.33 | 0.18 | 10 | AT1 Cell |
| P4hb | 0.51 | 0.00 | 0.00 | 0.56 | 0.40 | 10 | AT1 Cell |
| Lamp1 | 0.48 | 0.00 | 0.00 | 0.58 | 0.42 | 10 | AT1 Cell |
| Tmbim6 | 0.47 | 0.00 | 0.00 | 0.54 | 0.37 | 10 | AT1 Cell |
| Sypl | 0.49 | 0.00 | 0.00 | 0.41 | 0.25 | 10 | AT1 Cell |
| Swt1 | 0.57 | 0.00 | 0.00 | 0.29 | 0.15 | 10 | AT1 Cell |
| Myl12b | 0.34 | 0.00 | 0.00 | 0.73 | 0.60 | 10 | AT1 Cell |
| Cdkn1b | 0.53 | 0.00 | 0.00 | 0.36 | 0.21 | 10 | AT1 Cell |
| Elovl5 | 0.48 | 0.00 | 0.00 | 0.29 | 0.15 | 10 | AT1 Cell |
| Actn4 | 0.50 | 0.00 | 0.00 | 0.44 | 0.29 | 10 | AT1 Cell |
| Gnas | 0.30 | 0.00 | 0.00 | 0.79 | 0.67 | 10 | AT1 Cell |
| Tmem59 | 0.50 | 0.00 | 0.00 | 0.54 | 0.40 | 10 | AT1 Cell |
| 4933426M11Rik | 0.50 | 0.00 | 0.00 | 0.28 | 0.15 | 10 | AT1 Cell |
| Zfhx3 | 0.49 | 0.00 | 0.00 | 0.31 | 0.17 | 10 | AT1 Cell |
| Atp2b1 | 0.34 | 0.00 | 0.00 | 0.72 | 0.58 | 10 | AT1 Cell |
| Rb1cc1 | 0.50 | 0.00 | 0.00 | 0.45 | 0.30 | 10 | AT1 Cell |
| 1810037I17Rik | 0.46 | 0.00 | 0.00 | 0.47 | 0.32 | 10 | AT1 Cell |
| Eif1a | 0.49 | 0.00 | 0.00 | 0.27 | 0.14 | 10 | AT1 Cell |
| Ddb1 | 0.50 | 0.00 | 0.00 | 0.32 | 0.18 | 10 | AT1 Cell |
| Carkd | 0.50 | 0.00 | 0.00 | 0.28 | 0.15 | 10 | AT1 Cell |
| Sptan1 | 0.47 | 0.00 | 0.00 | 0.30 | 0.16 | 10 | AT1 Cell |
| Klf9 | 0.46 | 0.00 | 0.00 | 0.33 | 0.19 | 10 | AT1 Cell |
| Pdcd6ip | 0.50 | 0.00 | 0.00 | 0.29 | 0.16 | 10 | AT1 Cell |
| Rin2 | 0.47 | 0.00 | 0.00 | 0.28 | 0.15 | 10 | AT1 Cell |
| Actg1 | 0.30 | 0.00 | 0.00 | 0.84 | 0.76 | 10 | AT1 Cell |
| Kras | 0.51 | 0.00 | 0.00 | 0.32 | 0.18 | 10 | AT1 Cell |
| Vapa | 0.47 | 0.00 | 0.00 | 0.36 | 0.22 | 10 | AT1 Cell |
| Parva | 0.46 | 0.00 | 0.00 | 0.28 | 0.15 | 10 | AT1 Cell |
| Dnajc3 | 0.49 | 0.00 | 0.00 | 0.40 | 0.27 | 10 | AT1 Cell |
| Serpinb6a | 0.48 | 0.00 | 0.00 | 0.31 | 0.18 | 10 | AT1 Cell |
| Srrm2 | 0.38 | 0.00 | 0.00 | 0.71 | 0.60 | 10 | AT1 Cell |
| Akap13 | 0.43 | 0.00 | 0.00 | 0.41 | 0.26 | 10 | AT1 Cell |
| Ppic | 0.43 | 0.00 | 0.00 | 0.53 | 0.38 | 10 | AT1 Cell |
| 15-Sep | 0.49 | 0.00 | 0.00 | 0.47 | 0.33 | 10 | AT1 Cell |
| Rnase4 | 0.43 | 0.00 | 0.00 | 0.32 | 0.18 | 10 | AT1 Cell |
| Spop | 0.41 | 0.00 | 0.00 | 0.47 | 0.33 | 10 | AT1 Cell |
| Tbrg1 | 0.45 | 0.00 | 0.00 | 0.43 | 0.30 | 10 | AT1 Cell |
| Pkm | 0.45 | 0.00 | 0.00 | 0.36 | 0.22 | 10 | AT1 Cell |
| Gm3150 | 0.52 | 0.00 | 0.00 | 0.28 | 0.16 | 10 | AT1 Cell |
| 1110008F13Rik | 0.43 | 0.00 | 0.00 | 0.28 | 0.16 | 10 | AT1 Cell |
| Wwtr1 | 0.44 | 0.00 | 0.00 | 0.26 | 0.14 | 10 | AT1 Cell |
| Diap1 | 0.43 | 0.00 | 0.00 | 0.26 | 0.14 | 10 | AT1 Cell |
| Btg2 | 0.33 | 0.00 | 0.00 | 0.37 | 0.23 | 10 | AT1 Cell |
| Mllt4 | 0.41 | 0.00 | 0.00 | 0.34 | 0.21 | 10 | AT1 Cell |
| Syne2 | 0.45 | 0.00 | 0.00 | 0.25 | 0.14 | 10 | AT1 Cell |
| Svil | 0.50 | 0.00 | 0.00 | 0.28 | 0.17 | 10 | AT1 Cell |
| Lrrfip1 | 0.46 | 0.00 | 0.00 | 0.38 | 0.25 | 10 | AT1 Cell |
| Ptms | 0.37 | 0.00 | 0.00 | 0.53 | 0.40 | 10 | AT1 Cell |
| Cd2ap | 0.38 | 0.00 | 0.00 | 0.48 | 0.34 | 10 | AT1 Cell |
| Srsf11 | 0.47 | 0.00 | 0.00 | 0.58 | 0.49 | 10 | AT1 Cell |
| Hmgn1 | 0.34 | 0.00 | 0.00 | 0.57 | 0.45 | 10 | AT1 Cell |
| Aplp2 | 0.44 | 0.00 | 0.00 | 0.50 | 0.39 | 10 | AT1 Cell |
| Selk | 0.38 | 0.00 | 0.00 | 0.44 | 0.31 | 10 | AT1 Cell |
| Atp6v1a | 0.41 | 0.00 | 0.00 | 0.40 | 0.28 | 10 | AT1 Cell |
| Pls3 | 0.41 | 0.00 | 0.00 | 0.31 | 0.20 | 10 | AT1 Cell |
| Lpin2 | 0.36 | 0.00 | 0.00 | 0.25 | 0.15 | 10 | AT1 Cell |
| S100a13 | 0.39 | 0.00 | 0.00 | 0.37 | 0.25 | 10 | AT1 Cell |
| Bag1 | 0.43 | 0.00 | 0.00 | 0.35 | 0.23 | 10 | AT1 Cell |
| Slc38a2 | 0.41 | 0.00 | 0.00 | 0.30 | 0.19 | 10 | AT1 Cell |
| Tmem176b | 0.41 | 0.00 | 0.00 | 0.45 | 0.34 | 10 | AT1 Cell |
| Actn1 | 0.40 | 0.00 | 0.00 | 0.34 | 0.22 | 10 | AT1 Cell |
| Akap9 | 0.37 | 0.00 | 0.00 | 0.48 | 0.36 | 10 | AT1 Cell |
| Tmed2 | 0.39 | 0.00 | 0.00 | 0.36 | 0.24 | 10 | AT1 Cell |
| Kif5b | 0.33 | 0.00 | 0.00 | 0.58 | 0.48 | 10 | AT1 Cell |
| Mmp14 | 0.36 | 0.00 | 0.00 | 0.37 | 0.25 | 10 | AT1 Cell |
| Cab39 | 0.39 | 0.00 | 0.00 | 0.27 | 0.17 | 10 | AT1 Cell |
| Adam10 | 0.40 | 0.00 | 0.00 | 0.29 | 0.18 | 10 | AT1 Cell |
| Mettl7a1 | 0.31 | 0.00 | 0.00 | 0.33 | 0.21 | 10 | AT1 Cell |
| Lamc1 | 0.36 | 0.00 | 0.00 | 0.25 | 0.16 | 10 | AT1 Cell |
| Tbl1x | 0.37 | 0.00 | 0.00 | 0.36 | 0.25 | 10 | AT1 Cell |
| Drap1 | 0.33 | 0.00 | 0.00 | 0.36 | 0.25 | 10 | AT1 Cell |
| Os9 | 0.42 | 0.00 | 0.00 | 0.30 | 0.20 | 10 | AT1 Cell |
| Wasl | 0.40 | 0.00 | 0.00 | 0.28 | 0.18 | 10 | AT1 Cell |
| Rock1 | 0.35 | 0.00 | 0.00 | 0.45 | 0.35 | 10 | AT1 Cell |
| Phldb2 | 0.38 | 0.00 | 0.00 | 0.40 | 0.29 | 10 | AT1 Cell |
| Sdc4 | 0.29 | 0.00 | 0.00 | 0.34 | 0.23 | 10 | AT1 Cell |
| Eif1 | 0.25 | 0.00 | 0.00 | 0.72 | 0.67 | 10 | AT1 Cell |
| Snx4 | 0.39 | 0.00 | 0.00 | 0.35 | 0.25 | 10 | AT1 Cell |
| Tra2a | 0.39 | 0.00 | 0.00 | 0.31 | 0.21 | 10 | AT1 Cell |
| Timp2 | 0.34 | 0.00 | 0.00 | 0.31 | 0.21 | 10 | AT1 Cell |
| Bcap31 | 0.39 | 0.00 | 0.00 | 0.30 | 0.21 | 10 | AT1 Cell |
| Spcs2 | 0.37 | 0.00 | 0.00 | 0.40 | 0.31 | 10 | AT1 Cell |
| Rtn3 | 0.41 | 0.00 | 0.00 | 0.35 | 0.25 | 10 | AT1 Cell |
| Atp6v0e | 0.39 | 0.00 | 0.00 | 0.29 | 0.20 | 10 | AT1 Cell |
| Laptm4a | 0.27 | 0.00 | 0.00 | 0.61 | 0.52 | 10 | AT1 Cell |
| Skp1a | 0.37 | 0.00 | 0.00 | 0.48 | 0.40 | 10 | AT1 Cell |
| Ubl3 | 0.34 | 0.00 | 0.00 | 0.28 | 0.19 | 10 | AT1 Cell |
| Sec62 | 0.28 | 0.00 | 0.00 | 0.54 | 0.45 | 10 | AT1 Cell |
| Myh9 | 0.30 | 0.00 | 0.00 | 0.43 | 0.33 | 10 | AT1 Cell |
| Mbnl1 | 0.34 | 0.00 | 0.00 | 0.35 | 0.27 | 10 | AT1 Cell |
| Mcl1 | 0.27 | 0.00 | 0.00 | 0.32 | 0.23 | 10 | AT1 Cell |
| Ktn1 | 0.30 | 0.00 | 0.00 | 0.47 | 0.38 | 10 | AT1 Cell |
| Zc3h7a | 0.33 | 0.00 | 0.00 | 0.37 | 0.29 | 10 | AT1 Cell |
| Tceb2 | 0.26 | 0.00 | 0.00 | 0.53 | 0.47 | 10 | AT1 Cell |
| Tmed10 | 0.30 | 0.00 | 0.00 | 0.44 | 0.36 | 10 | AT1 Cell |
| Cyb5 | 0.31 | 0.00 | 0.00 | 0.35 | 0.26 | 10 | AT1 Cell |
| Arf1 | 0.26 | 0.00 | 0.00 | 0.50 | 0.42 | 10 | AT1 Cell |
| Ypel3 | 0.28 | 0.00 | 0.00 | 0.28 | 0.20 | 10 | AT1 Cell |
| Rab7 | 0.28 | 0.00 | 0.00 | 0.33 | 0.25 | 10 | AT1 Cell |
| Vps28 | 0.32 | 0.00 | 0.00 | 0.25 | 0.18 | 10 | AT1 Cell |
| Chmp2a | 0.29 | 0.00 | 0.00 | 0.39 | 0.31 | 10 | AT1 Cell |
| Tubb4b | 0.27 | 0.00 | 0.00 | 0.26 | 0.19 | 10 | AT1 Cell |
| Adipor1 | 0.30 | 0.00 | 0.00 | 0.31 | 0.24 | 10 | AT1 Cell |
| Rab6a | 0.33 | 0.00 | 0.00 | 0.28 | 0.21 | 10 | AT1 Cell |
| Itm2c | 0.31 | 0.00 | 0.00 | 0.30 | 0.23 | 10 | AT1 Cell |
| Cox17 | 0.28 | 0.00 | 0.00 | 0.31 | 0.24 | 10 | AT1 Cell |
| Cfl2 | 0.27 | 0.00 | 0.00 | 0.31 | 0.25 | 10 | AT1 Cell |
| Gm22759 | 0.25 | 0.00 | 0.00 | 0.66 | 0.64 | 10 | AT1 Cell |
| Ankrd12 | 0.31 | 0.00 | 0.00 | 0.33 | 0.27 | 10 | AT1 Cell |
| Usp34 | 0.26 | 0.00 | 0.00 | 0.32 | 0.25 | 10 | AT1 Cell |
| Nfe2l1 | 0.26 | 0.00 | 0.00 | 0.29 | 0.23 | 10 | AT1 Cell |
| Gm26300 | 0.25 | 0.00 | 0.00 | 0.42 | 0.38 | 10 | AT1 Cell |
| Ppib | 0.25 | 0.00 | 0.00 | 0.48 | 0.44 | 10 | AT1 Cell |
| Fmo2 | 2.67 | 0.00 | 0.00 | 0.95 | 0.14 | 11 | Matrix Fibroblast |
| Gpx3 | 2.52 | 0.00 | 0.00 | 0.93 | 0.11 | 11 | Matrix Fibroblast |
| Inmt | 2.49 | 0.00 | 0.00 | 0.62 | 0.03 | 11 | Matrix Fibroblast |
| Ifitm1 | 2.17 | 0.00 | 0.00 | 0.93 | 0.11 | 11 | Matrix Fibroblast |
| Lox | 2.00 | 0.00 | 0.00 | 0.80 | 0.11 | 11 | Matrix Fibroblast |
| Fos | 1.98 | 0.00 | 0.00 | 0.79 | 0.17 | 11 | Matrix Fibroblast |
| Gsn | 1.87 | 0.00 | 0.00 | 0.88 | 0.23 | 11 | Matrix Fibroblast |
| Tsc22d3 | 1.86 | 0.00 | 0.00 | 0.86 | 0.22 | 11 | Matrix Fibroblast |
| Tcf21 | 1.83 | 0.00 | 0.00 | 0.91 | 0.15 | 11 | Matrix Fibroblast |
| Peg3 | 1.82 | 0.00 | 0.00 | 0.90 | 0.28 | 11 | Matrix Fibroblast |
| Hsd11b1 | 1.74 | 0.00 | 0.00 | 0.65 | 0.07 | 11 | Matrix Fibroblast |
| Limch1 | 1.70 | 0.00 | 0.00 | 0.97 | 0.28 | 11 | Matrix Fibroblast |
| Cyr61 | 1.68 | 0.00 | 0.00 | 0.57 | 0.11 | 11 | Matrix Fibroblast |
| Selenbp1 | 1.66 | 0.00 | 0.00 | 0.82 | 0.20 | 11 | Matrix Fibroblast |
| Dpep1 | 1.66 | 0.00 | 0.00 | 0.58 | 0.05 | 11 | Matrix Fibroblast |
| Glul | 1.59 | 0.00 | 0.00 | 0.76 | 0.20 | 11 | Matrix Fibroblast |
| G0s2 | 1.58 | 0.00 | 0.00 | 0.62 | 0.12 | 11 | Matrix Fibroblast |
| Selenbp2 | 1.58 | 0.00 | 0.00 | 0.70 | 0.15 | 11 | Matrix Fibroblast |
| Ppp1r2 | 1.52 | 0.00 | 0.00 | 0.78 | 0.24 | 11 | Matrix Fibroblast |
| Il6ra | 1.50 | 0.00 | 0.00 | 0.60 | 0.09 | 11 | Matrix Fibroblast |
| Cdo1 | 1.45 | 0.00 | 0.00 | 0.48 | 0.05 | 11 | Matrix Fibroblast |
| Nap1l1 | 1.44 | 0.00 | 0.00 | 0.88 | 0.46 | 11 | Matrix Fibroblast |
| Ndrg2 | 1.43 | 0.00 | 0.00 | 0.55 | 0.06 | 11 | Matrix Fibroblast |
| Smoc2 | 1.43 | 0.00 | 0.00 | 0.39 | 0.02 | 11 | Matrix Fibroblast |
| Drap1 | 1.41 | 0.00 | 0.00 | 0.71 | 0.23 | 11 | Matrix Fibroblast |
| Gm4204 | 1.38 | 0.00 | 0.00 | 0.89 | 0.49 | 11 | Matrix Fibroblast |
| Adamts1 | 1.33 | 0.00 | 0.00 | 0.64 | 0.20 | 11 | Matrix Fibroblast |
| Klf9 | 1.33 | 0.00 | 0.00 | 0.65 | 0.18 | 11 | Matrix Fibroblast |
| Gadd45b | 1.28 | 0.00 | 0.00 | 0.43 | 0.07 | 11 | Matrix Fibroblast |
| Tns1 | 1.28 | 0.00 | 0.00 | 0.71 | 0.22 | 11 | Matrix Fibroblast |
| Mark3 | 1.27 | 0.00 | 0.00 | 0.56 | 0.15 | 11 | Matrix Fibroblast |
| Ogn | 1.25 | 0.00 | 0.00 | 0.57 | 0.12 | 11 | Matrix Fibroblast |
| Npr3 | 1.23 | 0.00 | 0.00 | 0.50 | 0.12 | 11 | Matrix Fibroblast |
| Bnip3 | 1.22 | 0.00 | 0.00 | 0.43 | 0.07 | 11 | Matrix Fibroblast |
| Nebl | 1.22 | 0.00 | 0.00 | 0.45 | 0.09 | 11 | Matrix Fibroblast |
| Apoe | 1.21 | 0.00 | 0.00 | 0.91 | 0.26 | 11 | Matrix Fibroblast |
| Mettl21e | 1.20 | 0.00 | 0.00 | 0.32 | 0.02 | 11 | Matrix Fibroblast |
| Ifitm2 | 1.18 | 0.00 | 0.00 | 0.75 | 0.31 | 11 | Matrix Fibroblast |
| H6pd | 1.18 | 0.00 | 0.00 | 0.43 | 0.08 | 11 | Matrix Fibroblast |
| Vldlr | 1.17 | 0.00 | 0.00 | 0.46 | 0.10 | 11 | Matrix Fibroblast |
| Gm7676 | 1.16 | 0.00 | 0.00 | 0.62 | 0.17 | 11 | Matrix Fibroblast |
| Serping1 | 1.13 | 0.00 | 0.00 | 0.54 | 0.11 | 11 | Matrix Fibroblast |
| Zbtb16 | 1.13 | 0.00 | 0.00 | 0.37 | 0.05 | 11 | Matrix Fibroblast |
| Fhl1 | 1.09 | 0.00 | 0.00 | 0.84 | 0.28 | 11 | Matrix Fibroblast |
| D4Wsu53e | 1.09 | 0.00 | 0.00 | 0.92 | 0.59 | 11 | Matrix Fibroblast |
| Fkbp5 | 1.09 | 0.00 | 0.00 | 0.41 | 0.08 | 11 | Matrix Fibroblast |
| Maf | 1.09 | 0.00 | 0.00 | 0.67 | 0.18 | 11 | Matrix Fibroblast |
| Macf1 | 1.05 | 0.00 | 0.00 | 0.92 | 0.51 | 11 | Matrix Fibroblast |
| Atp1a2 | 1.04 | 0.00 | 0.00 | 0.43 | 0.09 | 11 | Matrix Fibroblast |
| Eln | 1.04 | 0.00 | 0.00 | 0.60 | 0.14 | 11 | Matrix Fibroblast |
| Mgp | 1.03 | 0.00 | 0.00 | 0.92 | 0.30 | 11 | Matrix Fibroblast |
| Ptgis | 1.02 | 0.00 | 0.00 | 0.34 | 0.04 | 11 | Matrix Fibroblast |
| Pla2g12a | 0.99 | 0.00 | 0.00 | 0.27 | 0.04 | 11 | Matrix Fibroblast |
| Ahcy | 0.99 | 0.00 | 0.00 | 0.28 | 0.04 | 11 | Matrix Fibroblast |
| Gm4737 | 0.92 | 0.00 | 0.00 | 0.25 | 0.03 | 11 | Matrix Fibroblast |
| Ifitm7 | 0.91 | 0.00 | 0.00 | 0.27 | 0.02 | 11 | Matrix Fibroblast |
| Mfap4 | 0.85 | 0.00 | 0.00 | 0.90 | 0.22 | 11 | Matrix Fibroblast |
| Adh1 | 0.69 | 0.00 | 0.00 | 0.62 | 0.16 | 11 | Matrix Fibroblast |
| Gyg | 0.91 | 0.00 | 0.00 | 0.61 | 0.18 | 11 | Matrix Fibroblast |
| Hspb1 | 1.15 | 0.00 | 0.00 | 0.48 | 0.13 | 11 | Matrix Fibroblast |
| Zfp36 | 1.22 | 0.00 | 0.00 | 0.52 | 0.15 | 11 | Matrix Fibroblast |
| Sesn1 | 0.92 | 0.00 | 0.00 | 0.31 | 0.05 | 11 | Matrix Fibroblast |
| Egr1 | 1.40 | 0.00 | 0.00 | 0.47 | 0.13 | 11 | Matrix Fibroblast |
| Rps4x | 0.69 | 0.00 | 0.00 | 0.98 | 0.86 | 11 | Matrix Fibroblast |
| Nr2f2 | 1.05 | 0.00 | 0.00 | 0.59 | 0.20 | 11 | Matrix Fibroblast |
| Ctsl | 1.03 | 0.00 | 0.00 | 0.77 | 0.38 | 11 | Matrix Fibroblast |
| Fbxo32 | 0.92 | 0.00 | 0.00 | 0.26 | 0.04 | 11 | Matrix Fibroblast |
| Malat1 | 0.50 | 0.00 | 0.00 | 1.00 | 1.00 | 11 | Matrix Fibroblast |
| Col13a1 | 0.87 | 0.00 | 0.00 | 0.38 | 0.09 | 11 | Matrix Fibroblast |
| Lbh | 0.85 | 0.00 | 0.00 | 0.56 | 0.18 | 11 | Matrix Fibroblast |
| Nampt | 1.01 | 0.00 | 0.00 | 0.33 | 0.07 | 11 | Matrix Fibroblast |
| Dusp1 | 1.11 | 0.00 | 0.00 | 0.49 | 0.16 | 11 | Matrix Fibroblast |
| Fn1 | 0.75 | 0.00 | 0.00 | 0.63 | 0.22 | 11 | Matrix Fibroblast |
| Plac9b | 0.78 | 0.00 | 0.00 | 0.73 | 0.29 | 11 | Matrix Fibroblast |
| Tpt1 | 0.59 | 0.00 | 0.00 | 0.99 | 0.87 | 11 | Matrix Fibroblast |
| Plac9a | 0.77 | 0.00 | 0.00 | 0.73 | 0.29 | 11 | Matrix Fibroblast |
| Palld | 0.88 | 0.00 | 0.00 | 0.51 | 0.16 | 11 | Matrix Fibroblast |
| Angptl4 | 0.97 | 0.00 | 0.00 | 0.29 | 0.06 | 11 | Matrix Fibroblast |
| Rbp1 | 0.76 | 0.00 | 0.00 | 0.56 | 0.18 | 11 | Matrix Fibroblast |
| Mertk | 0.84 | 0.00 | 0.00 | 0.28 | 0.06 | 11 | Matrix Fibroblast |
| Cacna1d | 0.77 | 0.00 | 0.00 | 0.26 | 0.05 | 11 | Matrix Fibroblast |
| Rps27 | 0.60 | 0.00 | 0.00 | 0.97 | 0.81 | 11 | Matrix Fibroblast |
| Spon1 | 0.73 | 0.00 | 0.00 | 0.29 | 0.06 | 11 | Matrix Fibroblast |
| Ppap2b | 0.98 | 0.00 | 0.00 | 0.57 | 0.23 | 11 | Matrix Fibroblast |
| Fxyd1 | 0.82 | 0.00 | 0.00 | 0.34 | 0.08 | 11 | Matrix Fibroblast |
| Cnst | 0.79 | 0.00 | 0.00 | 0.25 | 0.05 | 11 | Matrix Fibroblast |
| Tgm2 | 0.91 | 0.00 | 0.00 | 0.33 | 0.08 | 11 | Matrix Fibroblast |
| Zfp36l1 | 0.99 | 0.00 | 0.00 | 0.77 | 0.44 | 11 | Matrix Fibroblast |
| Glud1 | 0.99 | 0.00 | 0.00 | 0.47 | 0.16 | 11 | Matrix Fibroblast |
| Tgfbr3 | 0.79 | 0.00 | 0.00 | 0.31 | 0.08 | 11 | Matrix Fibroblast |
| Trp53inp1 | 0.81 | 0.00 | 0.00 | 0.30 | 0.07 | 11 | Matrix Fibroblast |
| Gm9846 | 0.50 | 0.00 | 0.00 | 0.99 | 0.89 | 11 | Matrix Fibroblast |
| Txnip | 0.91 | 0.00 | 0.00 | 0.57 | 0.24 | 11 | Matrix Fibroblast |
| Myo1b | 0.95 | 0.00 | 0.00 | 0.45 | 0.16 | 11 | Matrix Fibroblast |
| Gm26669 | 1.09 | 0.00 | 0.00 | 0.53 | 0.22 | 11 | Matrix Fibroblast |
| Echdc2 | 0.77 | 0.00 | 0.00 | 0.25 | 0.05 | 11 | Matrix Fibroblast |
| Kank3 | 0.79 | 0.00 | 0.00 | 0.42 | 0.13 | 11 | Matrix Fibroblast |
| Aldh2 | 0.85 | 0.00 | 0.00 | 0.65 | 0.32 | 11 | Matrix Fibroblast |
| Rpl23a-ps3 | 0.59 | 0.00 | 0.00 | 0.94 | 0.74 | 11 | Matrix Fibroblast |
| Cp | 0.78 | 0.00 | 0.00 | 0.39 | 0.12 | 11 | Matrix Fibroblast |
| Hp | 0.79 | 0.00 | 0.00 | 0.36 | 0.10 | 11 | Matrix Fibroblast |
| Gm10335 | 0.59 | 0.00 | 0.00 | 0.93 | 0.72 | 11 | Matrix Fibroblast |
| Cebpb | 0.80 | 0.00 | 0.00 | 0.32 | 0.08 | 11 | Matrix Fibroblast |
| Gm10132 | 0.60 | 0.00 | 0.00 | 0.92 | 0.72 | 11 | Matrix Fibroblast |
| Fibin | 0.78 | 0.00 | 0.00 | 0.38 | 0.12 | 11 | Matrix Fibroblast |
| Angpt1 | 0.87 | 0.00 | 0.00 | 0.28 | 0.07 | 11 | Matrix Fibroblast |
| Rpl13 | 0.54 | 0.00 | 0.00 | 0.95 | 0.79 | 11 | Matrix Fibroblast |
| Igfbp3 | 0.72 | 0.00 | 0.00 | 0.27 | 0.06 | 11 | Matrix Fibroblast |
| Rpl13-ps3 | 0.57 | 0.00 | 0.00 | 0.94 | 0.75 | 11 | Matrix Fibroblast |
| Rpl23 | 0.50 | 0.00 | 0.00 | 0.97 | 0.86 | 11 | Matrix Fibroblast |
| Junb | 0.97 | 0.00 | 0.00 | 0.48 | 0.19 | 11 | Matrix Fibroblast |
| Pabpc1 | 0.58 | 0.00 | 0.00 | 0.95 | 0.80 | 11 | Matrix Fibroblast |
| Phlda1 | 0.89 | 0.00 | 0.00 | 0.49 | 0.20 | 11 | Matrix Fibroblast |
| Rpl34 | 0.55 | 0.00 | 0.00 | 0.95 | 0.79 | 11 | Matrix Fibroblast |
| Gas5 | 0.73 | 0.00 | 0.00 | 0.79 | 0.53 | 11 | Matrix Fibroblast |
| Cdh11 | 0.54 | 0.00 | 0.00 | 0.51 | 0.18 | 11 | Matrix Fibroblast |
| Rpl39 | 0.53 | 0.00 | 0.00 | 0.95 | 0.80 | 11 | Matrix Fibroblast |
| Sparcl1 | 0.57 | 0.00 | 0.00 | 0.76 | 0.38 | 11 | Matrix Fibroblast |
| Lama4 | 0.79 | 0.00 | 0.00 | 0.38 | 0.13 | 11 | Matrix Fibroblast |
| Cir1 | 0.92 | 0.00 | 0.00 | 0.47 | 0.20 | 11 | Matrix Fibroblast |
| Mxra8 | 0.72 | 0.00 | 0.00 | 0.43 | 0.16 | 11 | Matrix Fibroblast |
| Nexn | 0.70 | 0.00 | 0.00 | 0.52 | 0.21 | 11 | Matrix Fibroblast |
| Lamb1 | 0.77 | 0.00 | 0.00 | 0.42 | 0.15 | 11 | Matrix Fibroblast |
| Clec3b | 0.65 | 0.00 | 0.00 | 0.26 | 0.06 | 11 | Matrix Fibroblast |
| Rps26-ps1 | 0.46 | 0.00 | 0.00 | 0.98 | 0.87 | 11 | Matrix Fibroblast |
| 4-Sep | 0.63 | 0.00 | 0.00 | 0.54 | 0.22 | 11 | Matrix Fibroblast |
| Rpl23a | 0.57 | 0.00 | 0.00 | 0.88 | 0.66 | 11 | Matrix Fibroblast |
| Rps8 | 0.47 | 0.00 | 0.00 | 0.96 | 0.81 | 11 | Matrix Fibroblast |
| Rps14 | 0.42 | 0.00 | 0.00 | 0.98 | 0.92 | 11 | Matrix Fibroblast |
| Prex2 | 0.74 | 0.00 | 0.00 | 0.45 | 0.18 | 11 | Matrix Fibroblast |
| Col6a1 | 0.69 | 0.00 | 0.00 | 0.46 | 0.18 | 11 | Matrix Fibroblast |
| Ifrd1 | 0.82 | 0.00 | 0.00 | 0.39 | 0.15 | 11 | Matrix Fibroblast |
| Rps17 | 0.51 | 0.00 | 0.00 | 0.92 | 0.77 | 11 | Matrix Fibroblast |
| Snai2 | 0.78 | 0.00 | 0.00 | 0.34 | 0.11 | 11 | Matrix Fibroblast |
| mt-Rnr2 | 0.41 | 0.00 | 0.00 | 1.00 | 0.99 | 11 | Matrix Fibroblast |
| Mettl7a1 | 0.72 | 0.00 | 0.00 | 0.49 | 0.21 | 11 | Matrix Fibroblast |
| Rpl9 | 0.57 | 0.00 | 0.00 | 0.84 | 0.62 | 11 | Matrix Fibroblast |
| Gm26384 | 0.64 | 0.00 | 0.00 | 0.75 | 0.48 | 11 | Matrix Fibroblast |
| Rps23 | 0.47 | 0.00 | 0.00 | 0.95 | 0.79 | 11 | Matrix Fibroblast |
| Cd302 | 0.77 | 0.00 | 0.00 | 0.38 | 0.14 | 11 | Matrix Fibroblast |
| RPL24 | 0.64 | 0.00 | 0.00 | 0.74 | 0.45 | 11 | Matrix Fibroblast |
| Rps28 | 0.44 | 0.00 | 0.00 | 0.97 | 0.83 | 11 | Matrix Fibroblast |
| Camk2n1 | 0.75 | 0.00 | 0.00 | 0.37 | 0.14 | 11 | Matrix Fibroblast |
| Specc1l | 0.74 | 0.00 | 0.00 | 0.34 | 0.12 | 11 | Matrix Fibroblast |
| Tacc1 | 0.77 | 0.00 | 0.00 | 0.48 | 0.22 | 11 | Matrix Fibroblast |
| Btg2 | 1.03 | 0.00 | 0.00 | 0.48 | 0.23 | 11 | Matrix Fibroblast |
| Rpl9-ps6 | 0.50 | 0.00 | 0.00 | 0.91 | 0.73 | 11 | Matrix Fibroblast |
| Rpl37a | 0.43 | 0.00 | 0.00 | 0.95 | 0.83 | 11 | Matrix Fibroblast |
| Rps26 | 0.39 | 0.00 | 0.00 | 0.98 | 0.90 | 11 | Matrix Fibroblast |
| Snhg9 | 0.74 | 0.00 | 0.00 | 0.29 | 0.10 | 11 | Matrix Fibroblast |
| Ifngr1 | 0.73 | 0.00 | 0.00 | 0.50 | 0.23 | 11 | Matrix Fibroblast |
| Tenc1 | 0.84 | 0.00 | 0.00 | 0.36 | 0.14 | 11 | Matrix Fibroblast |
| Mir682 | 0.41 | 0.00 | 0.00 | 0.96 | 0.85 | 11 | Matrix Fibroblast |
| Ivd | 0.75 | 0.00 | 0.00 | 0.27 | 0.08 | 11 | Matrix Fibroblast |
| Rpl37 | 0.45 | 0.00 | 0.00 | 0.94 | 0.80 | 11 | Matrix Fibroblast |
| Gm22759 | 0.50 | 0.00 | 0.00 | 0.85 | 0.64 | 11 | Matrix Fibroblast |
| Fblim1 | 0.65 | 0.00 | 0.00 | 0.39 | 0.16 | 11 | Matrix Fibroblast |
| Gm22758 | 0.47 | 0.00 | 0.00 | 0.90 | 0.68 | 11 | Matrix Fibroblast |
| 2410004N09Rik | 0.71 | 0.00 | 0.00 | 0.30 | 0.10 | 11 | Matrix Fibroblast |
| Cdkn1c | 0.54 | 0.00 | 0.00 | 0.42 | 0.18 | 11 | Matrix Fibroblast |
| Gm10263 | 0.45 | 0.00 | 0.00 | 0.94 | 0.76 | 11 | Matrix Fibroblast |
| Gm24865 | 0.51 | 0.00 | 0.00 | 0.85 | 0.62 | 11 | Matrix Fibroblast |
| Med13l | 0.73 | 0.00 | 0.00 | 0.32 | 0.12 | 11 | Matrix Fibroblast |
| Jun | 0.89 | 0.00 | 0.00 | 0.61 | 0.36 | 11 | Matrix Fibroblast |
| Rps25 | 0.41 | 0.00 | 0.00 | 0.94 | 0.80 | 11 | Matrix Fibroblast |
| Bgn | 0.53 | 0.00 | 0.00 | 0.51 | 0.23 | 11 | Matrix Fibroblast |
| Snhg6 | 0.78 | 0.00 | 0.00 | 0.40 | 0.18 | 11 | Matrix Fibroblast |
| Pkn2 | 0.74 | 0.00 | 0.00 | 0.43 | 0.20 | 11 | Matrix Fibroblast |
| 2410006H16Rik | 0.71 | 0.00 | 0.00 | 0.50 | 0.25 | 11 | Matrix Fibroblast |
| Gm8730 | 0.38 | 0.00 | 0.00 | 0.94 | 0.82 | 11 | Matrix Fibroblast |
| Sms | 0.78 | 0.00 | 0.00 | 0.30 | 0.11 | 11 | Matrix Fibroblast |
| Rps29 | 0.33 | 0.00 | 0.00 | 0.99 | 0.90 | 11 | Matrix Fibroblast |
| Rps15a | 0.41 | 0.00 | 0.00 | 0.94 | 0.81 | 11 | Matrix Fibroblast |
| Tpm1 | 0.45 | 0.00 | 0.00 | 0.78 | 0.51 | 11 | Matrix Fibroblast |
| C230081A13Rik | 0.66 | 0.00 | 0.00 | 0.28 | 0.09 | 11 | Matrix Fibroblast |
| Rpl17 | 0.54 | 0.00 | 0.00 | 0.77 | 0.53 | 11 | Matrix Fibroblast |
| Clk1 | 0.70 | 0.00 | 0.00 | 0.58 | 0.34 | 11 | Matrix Fibroblast |
| Rpl36 | 0.51 | 0.00 | 0.00 | 0.83 | 0.62 | 11 | Matrix Fibroblast |
| Plxdc2 | 0.55 | 0.00 | 0.00 | 0.33 | 0.12 | 11 | Matrix Fibroblast |
| D19Bwg1357e | 0.75 | 0.00 | 0.00 | 0.33 | 0.13 | 11 | Matrix Fibroblast |
| Rplp0 | 0.36 | 0.00 | 0.00 | 0.94 | 0.83 | 11 | Matrix Fibroblast |
| Plcb4 | 0.74 | 0.00 | 0.00 | 0.34 | 0.14 | 11 | Matrix Fibroblast |
| Rpl36-ps3 | 0.55 | 0.00 | 0.00 | 0.76 | 0.53 | 11 | Matrix Fibroblast |
| Wipi1 | 0.70 | 0.00 | 0.00 | 0.32 | 0.12 | 11 | Matrix Fibroblast |
| Pik3r1 | 0.79 | 0.00 | 0.00 | 0.31 | 0.12 | 11 | Matrix Fibroblast |
| Thbs1 | 0.61 | 0.00 | 0.00 | 0.33 | 0.12 | 11 | Matrix Fibroblast |
| Limd1 | 0.70 | 0.00 | 0.00 | 0.39 | 0.17 | 11 | Matrix Fibroblast |
| Rpl10-ps3 | 0.46 | 0.00 | 0.00 | 0.89 | 0.72 | 11 | Matrix Fibroblast |
| Npnt | 0.35 | 0.00 | 0.00 | 0.51 | 0.24 | 11 | Matrix Fibroblast |
| Dapk1 | 0.66 | 0.00 | 0.00 | 0.28 | 0.11 | 11 | Matrix Fibroblast |
| Gabarapl1 | 0.72 | 0.00 | 0.00 | 0.35 | 0.15 | 11 | Matrix Fibroblast |
| Map1lc3b | 0.66 | 0.00 | 0.00 | 0.61 | 0.39 | 11 | Matrix Fibroblast |
| Rpl10 | 0.45 | 0.00 | 0.00 | 0.87 | 0.71 | 11 | Matrix Fibroblast |
| Lrp1 | 0.58 | 0.00 | 0.00 | 0.30 | 0.11 | 11 | Matrix Fibroblast |
| Rps27a | 0.41 | 0.00 | 0.00 | 0.92 | 0.76 | 11 | Matrix Fibroblast |
| Col6a2 | 0.65 | 0.00 | 0.00 | 0.34 | 0.14 | 11 | Matrix Fibroblast |
| Rpl31 | 0.48 | 0.00 | 0.00 | 0.80 | 0.57 | 11 | Matrix Fibroblast |
| Rpl21 | 0.38 | 0.00 | 0.00 | 0.92 | 0.74 | 11 | Matrix Fibroblast |
| Snhg5 | 0.72 | 0.00 | 0.00 | 0.37 | 0.17 | 11 | Matrix Fibroblast |
| 1810058I24Rik | 0.69 | 0.00 | 0.00 | 0.40 | 0.20 | 11 | Matrix Fibroblast |
| Rps16 | 0.35 | 0.00 | 0.00 | 0.94 | 0.84 | 11 | Matrix Fibroblast |
| Ybx3 | 0.73 | 0.00 | 0.00 | 0.39 | 0.19 | 11 | Matrix Fibroblast |
| Mettl7a2 | 0.63 | 0.00 | 0.00 | 0.27 | 0.10 | 11 | Matrix Fibroblast |
| Gm17511 | 0.45 | 0.00 | 0.00 | 0.80 | 0.60 | 11 | Matrix Fibroblast |
| Csrp1 | 0.61 | 0.00 | 0.00 | 0.36 | 0.16 | 11 | Matrix Fibroblast |
| Rps3a3 | 0.38 | 0.00 | 0.00 | 0.90 | 0.77 | 11 | Matrix Fibroblast |
| Rock2 | 0.72 | 0.00 | 0.00 | 0.52 | 0.32 | 11 | Matrix Fibroblast |
| Rpl35a | 0.36 | 0.00 | 0.00 | 0.91 | 0.79 | 11 | Matrix Fibroblast |
| Sh3bgrl | 0.46 | 0.00 | 0.00 | 0.68 | 0.44 | 11 | Matrix Fibroblast |
| Rps15 | 0.38 | 0.00 | 0.00 | 0.91 | 0.78 | 11 | Matrix Fibroblast |
| Gm7589 | 0.44 | 0.00 | 0.00 | 0.81 | 0.65 | 11 | Matrix Fibroblast |
| Rps3a2 | 0.34 | 0.00 | 0.00 | 0.93 | 0.82 | 11 | Matrix Fibroblast |
| Ier2 | 0.76 | 0.00 | 0.00 | 0.45 | 0.25 | 11 | Matrix Fibroblast |
| Rps18-ps3 | 0.46 | 0.00 | 0.00 | 0.75 | 0.55 | 11 | Matrix Fibroblast |
| Ptms | 0.49 | 0.00 | 0.00 | 0.62 | 0.39 | 11 | Matrix Fibroblast |
| Gm6133 | 0.50 | 0.00 | 0.00 | 0.56 | 0.33 | 11 | Matrix Fibroblast |
| Btg1 | 0.55 | 0.00 | 0.00 | 0.49 | 0.28 | 11 | Matrix Fibroblast |
| Adamts2 | 0.44 | 0.00 | 0.00 | 0.26 | 0.10 | 11 | Matrix Fibroblast |
| Gm5428 | 0.36 | 0.00 | 0.00 | 0.91 | 0.78 | 11 | Matrix Fibroblast |
| Rps13 | 0.40 | 0.00 | 0.00 | 0.82 | 0.64 | 11 | Matrix Fibroblast |
| Gm22426 | 0.47 | 0.00 | 0.00 | 0.70 | 0.48 | 11 | Matrix Fibroblast |
| Mef2a | 0.60 | 0.00 | 0.00 | 0.56 | 0.36 | 11 | Matrix Fibroblast |
| Rps12-ps3 | 0.35 | 0.00 | 0.00 | 0.91 | 0.78 | 11 | Matrix Fibroblast |
| Rpl32 | 0.31 | 0.00 | 0.00 | 0.95 | 0.84 | 11 | Matrix Fibroblast |
| Luc7l2 | 0.59 | 0.00 | 0.00 | 0.58 | 0.39 | 11 | Matrix Fibroblast |
| Rps18 | 0.41 | 0.00 | 0.00 | 0.83 | 0.65 | 11 | Matrix Fibroblast |
| Fgfr1 | 0.57 | 0.00 | 0.00 | 0.26 | 0.10 | 11 | Matrix Fibroblast |
| Bnip3l | 0.61 | 0.00 | 0.00 | 0.56 | 0.36 | 11 | Matrix Fibroblast |
| Myh10 | 0.43 | 0.00 | 0.00 | 0.44 | 0.22 | 11 | Matrix Fibroblast |
| Ezh2 | 0.67 | 0.00 | 0.00 | 0.36 | 0.18 | 11 | Matrix Fibroblast |
| Gm26300 | 0.47 | 0.00 | 0.00 | 0.59 | 0.37 | 11 | Matrix Fibroblast |
| Rpl18a | 0.31 | 0.00 | 0.00 | 0.92 | 0.82 | 11 | Matrix Fibroblast |
| Rplp2 | 0.33 | 0.00 | 0.00 | 0.92 | 0.81 | 11 | Matrix Fibroblast |
| Klf6 | 0.55 | 0.00 | 0.00 | 0.56 | 0.35 | 11 | Matrix Fibroblast |
| Wbp5 | 0.59 | 0.00 | 0.00 | 0.66 | 0.47 | 11 | Matrix Fibroblast |
| Rpl30 | 0.45 | 0.00 | 0.00 | 0.74 | 0.58 | 11 | Matrix Fibroblast |
| Gm9493 | 0.34 | 0.00 | 0.00 | 0.89 | 0.74 | 11 | Matrix Fibroblast |
| Snhg1 | 0.63 | 0.00 | 0.00 | 0.31 | 0.15 | 11 | Matrix Fibroblast |
| Eif4ebp1 | 0.58 | 0.00 | 0.00 | 0.29 | 0.13 | 11 | Matrix Fibroblast |
| Rps12 | 0.37 | 0.00 | 0.00 | 0.84 | 0.68 | 11 | Matrix Fibroblast |
| Rpl26 | 0.34 | 0.00 | 0.00 | 0.90 | 0.81 | 11 | Matrix Fibroblast |
| Nfkbia | 0.47 | 0.00 | 0.00 | 0.58 | 0.37 | 11 | Matrix Fibroblast |
| Rps10 | 0.37 | 0.00 | 0.00 | 0.85 | 0.70 | 11 | Matrix Fibroblast |
| Rpl28 | 0.42 | 0.00 | 0.00 | 0.72 | 0.53 | 11 | Matrix Fibroblast |
| Gm3940 | 0.48 | 0.00 | 0.00 | 0.56 | 0.35 | 11 | Matrix Fibroblast |
| Rpl11 | 0.37 | 0.00 | 0.00 | 0.83 | 0.67 | 11 | Matrix Fibroblast |
| Rpl5 | 0.35 | 0.00 | 0.00 | 0.87 | 0.74 | 11 | Matrix Fibroblast |
| Rpl27a | 0.32 | 0.00 | 0.00 | 0.91 | 0.78 | 11 | Matrix Fibroblast |
| Rps3a1 | 0.28 | 0.00 | 0.00 | 0.94 | 0.85 | 11 | Matrix Fibroblast |
| Arid5b | 0.62 | 0.00 | 0.00 | 0.40 | 0.22 | 11 | Matrix Fibroblast |
| Gm7808 | 0.40 | 0.00 | 0.00 | 0.78 | 0.59 | 11 | Matrix Fibroblast |
| Gm10260 | 0.44 | 0.00 | 0.00 | 0.67 | 0.48 | 11 | Matrix Fibroblast |
| Gm15013 | 0.52 | 0.00 | 0.00 | 0.55 | 0.36 | 11 | Matrix Fibroblast |
| Zyx | 0.49 | 0.00 | 0.00 | 0.37 | 0.19 | 11 | Matrix Fibroblast |
| Smc3 | 0.58 | 0.00 | 0.00 | 0.55 | 0.37 | 11 | Matrix Fibroblast |
| Rin2 | 0.61 | 0.00 | 0.00 | 0.31 | 0.15 | 11 | Matrix Fibroblast |
| Gnas | 0.36 | 0.00 | 0.00 | 0.82 | 0.66 | 11 | Matrix Fibroblast |
| Gm11808 | 0.30 | 0.00 | 0.00 | 0.90 | 0.77 | 11 | Matrix Fibroblast |
| Hmgn5 | 0.80 | 0.00 | 0.00 | 0.43 | 0.26 | 11 | Matrix Fibroblast |
| mt-Cytb | 0.31 | 0.00 | 0.00 | 0.91 | 0.82 | 11 | Matrix Fibroblast |
| Rps3 | 0.29 | 0.00 | 0.00 | 0.92 | 0.83 | 11 | Matrix Fibroblast |
| Gm10036 | 0.32 | 0.00 | 0.00 | 0.88 | 0.74 | 11 | Matrix Fibroblast |
| Tgfbr2 | 0.51 | 0.00 | 0.00 | 0.26 | 0.12 | 11 | Matrix Fibroblast |
| Ifitm3 | 0.31 | 0.00 | 0.00 | 0.62 | 0.39 | 11 | Matrix Fibroblast |
| Rps21 | 0.32 | 0.00 | 0.00 | 0.87 | 0.75 | 11 | Matrix Fibroblast |
| Pnrc1 | 0.51 | 0.00 | 0.00 | 0.36 | 0.20 | 11 | Matrix Fibroblast |
| Prdx5 | 0.39 | 0.00 | 0.00 | 0.51 | 0.32 | 11 | Matrix Fibroblast |
| Rpl6 | 0.28 | 0.00 | 0.00 | 0.95 | 0.87 | 11 | Matrix Fibroblast |
| Rps7 | 0.32 | 0.00 | 0.00 | 0.88 | 0.73 | 11 | Matrix Fibroblast |
| Gm5239 | 0.39 | 0.00 | 0.00 | 0.70 | 0.51 | 11 | Matrix Fibroblast |
| Gm6472 | 0.29 | 0.00 | 0.00 | 0.91 | 0.80 | 11 | Matrix Fibroblast |
| Rpl36a | 0.39 | 0.00 | 0.00 | 0.76 | 0.62 | 11 | Matrix Fibroblast |
| Epn2 | 0.52 | 0.00 | 0.00 | 0.26 | 0.12 | 11 | Matrix Fibroblast |
| Tax1bp1 | 0.41 | 0.00 | 0.00 | 0.77 | 0.63 | 11 | Matrix Fibroblast |
| Jhdm1d | 0.55 | 0.00 | 0.00 | 0.31 | 0.15 | 11 | Matrix Fibroblast |
| Gm10288 | 0.29 | 0.00 | 0.00 | 0.90 | 0.78 | 11 | Matrix Fibroblast |
| Eif3e | 0.47 | 0.00 | 0.00 | 0.56 | 0.40 | 11 | Matrix Fibroblast |
| Ube2e3 | 0.53 | 0.00 | 0.00 | 0.28 | 0.14 | 11 | Matrix Fibroblast |
| Tsc22d1 | 0.48 | 0.00 | 0.00 | 0.51 | 0.33 | 11 | Matrix Fibroblast |
| Pbx1 | 0.55 | 0.00 | 0.00 | 0.35 | 0.20 | 11 | Matrix Fibroblast |
| Gm5093 | 0.39 | 0.00 | 0.00 | 0.64 | 0.46 | 11 | Matrix Fibroblast |
| Rb1cc1 | 0.53 | 0.00 | 0.00 | 0.46 | 0.30 | 11 | Matrix Fibroblast |
| Rps10-ps1 | 0.28 | 0.00 | 0.00 | 0.88 | 0.77 | 11 | Matrix Fibroblast |
| Gpc3 | 0.30 | 0.00 | 0.00 | 0.33 | 0.16 | 11 | Matrix Fibroblast |
| Rpl7 | 0.31 | 0.00 | 0.00 | 0.85 | 0.74 | 11 | Matrix Fibroblast |
| Rpl21-ps6 | 0.39 | 0.00 | 0.00 | 0.61 | 0.43 | 11 | Matrix Fibroblast |
| Tbrg1 | 0.52 | 0.00 | 0.00 | 0.46 | 0.30 | 11 | Matrix Fibroblast |
| Cdkn1a | 0.35 | 0.00 | 0.00 | 0.32 | 0.16 | 11 | Matrix Fibroblast |
| Nfia | 0.53 | 0.00 | 0.00 | 0.39 | 0.24 | 11 | Matrix Fibroblast |
| Sqstm1 | 0.48 | 0.00 | 0.00 | 0.47 | 0.31 | 11 | Matrix Fibroblast |
| Eif1 | 0.34 | 0.00 | 0.00 | 0.80 | 0.67 | 11 | Matrix Fibroblast |
| Iscu | 0.51 | 0.00 | 0.00 | 0.29 | 0.15 | 11 | Matrix Fibroblast |
| Rpl27 | 0.31 | 0.00 | 0.00 | 0.83 | 0.69 | 11 | Matrix Fibroblast |
| Ankrd11 | 0.48 | 0.00 | 0.00 | 0.53 | 0.37 | 11 | Matrix Fibroblast |
| Fkbp9 | 0.47 | 0.00 | 0.00 | 0.27 | 0.14 | 11 | Matrix Fibroblast |
| Luc7l3 | 0.44 | 0.00 | 0.00 | 0.70 | 0.56 | 11 | Matrix Fibroblast |
| Thra | 0.48 | 0.00 | 0.00 | 0.31 | 0.17 | 11 | Matrix Fibroblast |
| Jund | 0.51 | 0.00 | 0.00 | 0.42 | 0.27 | 11 | Matrix Fibroblast |
| Srrm1 | 0.49 | 0.00 | 0.00 | 0.61 | 0.48 | 11 | Matrix Fibroblast |
| Gm10126 | 0.38 | 0.00 | 0.00 | 0.56 | 0.39 | 11 | Matrix Fibroblast |
| Tcf25 | 0.44 | 0.00 | 0.00 | 0.64 | 0.51 | 11 | Matrix Fibroblast |
| Gstm1 | 0.45 | 0.00 | 0.00 | 0.28 | 0.15 | 11 | Matrix Fibroblast |
| Rbm39 | 0.32 | 0.00 | 0.00 | 0.81 | 0.71 | 11 | Matrix Fibroblast |
| Celf2 | 0.43 | 0.00 | 0.00 | 0.44 | 0.28 | 11 | Matrix Fibroblast |
| Rpsa | 0.29 | 0.00 | 0.00 | 0.75 | 0.62 | 11 | Matrix Fibroblast |
| Nisch | 0.44 | 0.00 | 0.00 | 0.45 | 0.30 | 11 | Matrix Fibroblast |
| Gltscr2 | 0.36 | 0.00 | 0.00 | 0.63 | 0.48 | 11 | Matrix Fibroblast |
| Echs1 | 0.48 | 0.00 | 0.00 | 0.27 | 0.15 | 11 | Matrix Fibroblast |
| Cnn2 | 0.34 | 0.00 | 0.00 | 0.55 | 0.39 | 11 | Matrix Fibroblast |
| Rpl21-ps4 | 0.39 | 0.00 | 0.00 | 0.50 | 0.35 | 11 | Matrix Fibroblast |
| Srrm2 | 0.37 | 0.00 | 0.00 | 0.73 | 0.60 | 11 | Matrix Fibroblast |
| Pdcd4 | 0.44 | 0.00 | 0.00 | 0.36 | 0.22 | 11 | Matrix Fibroblast |
| Arglu1 | 0.42 | 0.00 | 0.00 | 0.55 | 0.42 | 11 | Matrix Fibroblast |
| Kif1b | 0.44 | 0.00 | 0.00 | 0.37 | 0.23 | 11 | Matrix Fibroblast |
| Gm22761 | 0.43 | 0.00 | 0.00 | 0.29 | 0.17 | 11 | Matrix Fibroblast |
| Nenf | 0.41 | 0.00 | 0.00 | 0.31 | 0.18 | 11 | Matrix Fibroblast |
| Nktr | 0.47 | 0.00 | 0.00 | 0.49 | 0.36 | 11 | Matrix Fibroblast |
| Cox7c | 0.33 | 0.00 | 0.00 | 0.74 | 0.63 | 11 | Matrix Fibroblast |
| Pcbp2 | 0.40 | 0.00 | 0.00 | 0.54 | 0.42 | 11 | Matrix Fibroblast |
| Mbnl2 | 0.37 | 0.00 | 0.00 | 0.48 | 0.34 | 11 | Matrix Fibroblast |
| Sod1 | 0.35 | 0.00 | 0.00 | 0.60 | 0.47 | 11 | Matrix Fibroblast |
| Sdc4 | 0.34 | 0.00 | 0.00 | 0.36 | 0.22 | 11 | Matrix Fibroblast |
| Smc4 | 0.33 | 0.00 | 0.00 | 0.49 | 0.34 | 11 | Matrix Fibroblast |
| Nedd4 | 0.31 | 0.00 | 0.00 | 0.68 | 0.53 | 11 | Matrix Fibroblast |
| Hbp1 | 0.43 | 0.00 | 0.00 | 0.28 | 0.16 | 11 | Matrix Fibroblast |
| Zmym5 | 0.48 | 0.00 | 0.00 | 0.32 | 0.20 | 11 | Matrix Fibroblast |
| Rpl29 | 0.33 | 0.00 | 0.00 | 0.59 | 0.46 | 11 | Matrix Fibroblast |
| Hmgb1 | 0.27 | 0.00 | 0.00 | 0.73 | 0.61 | 11 | Matrix Fibroblast |
| Cirbp | 0.46 | 0.00 | 0.00 | 0.33 | 0.21 | 11 | Matrix Fibroblast |
| Gm10221 | 0.38 | 0.00 | 0.00 | 0.57 | 0.46 | 11 | Matrix Fibroblast |
| Hnrnpl | 0.44 | 0.00 | 0.00 | 0.37 | 0.25 | 11 | Matrix Fibroblast |
| Dynlt1a | 0.41 | 0.00 | 0.00 | 0.34 | 0.22 | 11 | Matrix Fibroblast |
| Btf3 | 0.32 | 0.00 | 0.00 | 0.63 | 0.51 | 11 | Matrix Fibroblast |
| Rbbp7 | 0.41 | 0.00 | 0.00 | 0.38 | 0.26 | 11 | Matrix Fibroblast |
| Ypel3 | 0.35 | 0.00 | 0.00 | 0.32 | 0.20 | 11 | Matrix Fibroblast |
| Atp5l | 0.37 | 0.00 | 0.00 | 0.54 | 0.44 | 11 | Matrix Fibroblast |
| Prdx6 | 0.28 | 0.00 | 0.00 | 0.40 | 0.27 | 11 | Matrix Fibroblast |
| Lmo4 | 0.35 | 0.00 | 0.00 | 0.38 | 0.26 | 11 | Matrix Fibroblast |
| Gm10923 | 0.37 | 0.00 | 0.00 | 0.33 | 0.21 | 11 | Matrix Fibroblast |
| Rbbp6 | 0.44 | 0.00 | 0.00 | 0.46 | 0.35 | 11 | Matrix Fibroblast |
| Scp2 | 0.36 | 0.00 | 0.00 | 0.45 | 0.32 | 11 | Matrix Fibroblast |
| Pnpla8 | 0.42 | 0.00 | 0.00 | 0.29 | 0.18 | 11 | Matrix Fibroblast |
| Sfrs18 | 0.32 | 0.00 | 0.00 | 0.68 | 0.59 | 11 | Matrix Fibroblast |
| Atp5l-ps1 | 0.41 | 0.00 | 0.00 | 0.39 | 0.27 | 11 | Matrix Fibroblast |
| Igfbp4 | 0.29 | 0.00 | 0.00 | 0.38 | 0.25 | 11 | Matrix Fibroblast |
| Gm10020 | 0.26 | 0.00 | 0.00 | 0.63 | 0.51 | 11 | Matrix Fibroblast |
| Zfp326 | 0.42 | 0.00 | 0.00 | 0.34 | 0.24 | 11 | Matrix Fibroblast |
| Socs2 | 0.28 | 0.00 | 0.00 | 0.47 | 0.34 | 11 | Matrix Fibroblast |
| Tmem254b | 0.38 | 0.00 | 0.00 | 0.30 | 0.19 | 11 | Matrix Fibroblast |
| Ythdc1 | 0.36 | 0.00 | 0.00 | 0.54 | 0.43 | 11 | Matrix Fibroblast |
| Tsix | 0.44 | 0.00 | 0.00 | 0.28 | 0.18 | 11 | Matrix Fibroblast |
| Eif3f | 0.32 | 0.00 | 0.00 | 0.48 | 0.36 | 11 | Matrix Fibroblast |
| Dynlt1c | 0.38 | 0.00 | 0.00 | 0.37 | 0.27 | 11 | Matrix Fibroblast |
| Smc5 | 0.40 | 0.00 | 0.00 | 0.26 | 0.16 | 11 | Matrix Fibroblast |
| Ccnl1 | 0.35 | 0.00 | 0.00 | 0.44 | 0.33 | 11 | Matrix Fibroblast |
| Dnajc3 | 0.40 | 0.00 | 0.00 | 0.37 | 0.27 | 11 | Matrix Fibroblast |
| Hip1 | 0.32 | 0.00 | 0.00 | 0.25 | 0.16 | 11 | Matrix Fibroblast |
| Sfr1 | 0.35 | 0.00 | 0.00 | 0.44 | 0.33 | 11 | Matrix Fibroblast |
| Ccnl2 | 0.36 | 0.00 | 0.00 | 0.34 | 0.24 | 11 | Matrix Fibroblast |
| 4931406P16Rik | 0.28 | 0.00 | 0.00 | 0.28 | 0.18 | 11 | Matrix Fibroblast |
| Gm17669 | 0.33 | 0.00 | 0.00 | 0.31 | 0.22 | 11 | Matrix Fibroblast |
| Rap2a | 0.28 | 0.00 | 0.00 | 0.25 | 0.16 | 11 | Matrix Fibroblast |
| Dynlt1b | 0.32 | 0.00 | 0.00 | 0.35 | 0.25 | 11 | Matrix Fibroblast |
| Ilk | 0.32 | 0.00 | 0.00 | 0.33 | 0.23 | 11 | Matrix Fibroblast |
| Gm10709 | 0.35 | 0.00 | 0.00 | 0.37 | 0.27 | 11 | Matrix Fibroblast |
| Ccar1 | 0.39 | 0.00 | 0.00 | 0.42 | 0.33 | 11 | Matrix Fibroblast |
| Lamp2 | 0.32 | 0.00 | 0.00 | 0.36 | 0.26 | 11 | Matrix Fibroblast |
| Kmt2e | 0.32 | 0.00 | 0.00 | 0.52 | 0.42 | 11 | Matrix Fibroblast |
| Tnrc6a | 0.36 | 0.00 | 0.00 | 0.35 | 0.26 | 11 | Matrix Fibroblast |
| 1500012F01Rik | 0.32 | 0.00 | 0.00 | 0.40 | 0.30 | 11 | Matrix Fibroblast |
| Rpl7a-ps5 | 0.29 | 0.00 | 0.00 | 0.46 | 0.37 | 11 | Matrix Fibroblast |
| St13 | 0.31 | 0.00 | 0.00 | 0.44 | 0.34 | 11 | Matrix Fibroblast |
| Eif5b | 0.32 | 0.00 | 0.00 | 0.59 | 0.51 | 11 | Matrix Fibroblast |
| Ankrd12 | 0.45 | 0.00 | 0.00 | 0.35 | 0.27 | 11 | Matrix Fibroblast |
| Tmem254c | 0.34 | 0.00 | 0.00 | 0.27 | 0.18 | 11 | Matrix Fibroblast |
| Tmem254a | 0.34 | 0.00 | 0.00 | 0.28 | 0.19 | 11 | Matrix Fibroblast |
| Atf4 | 0.35 | 0.00 | 0.00 | 0.36 | 0.28 | 11 | Matrix Fibroblast |
| Dynlt1f | 0.32 | 0.00 | 0.00 | 0.33 | 0.24 | 11 | Matrix Fibroblast |
| Gm20091 | 0.33 | 0.00 | 0.00 | 0.31 | 0.22 | 11 | Matrix Fibroblast |
| Pnn | 0.27 | 0.00 | 0.00 | 0.61 | 0.54 | 11 | Matrix Fibroblast |
| Ash1l | 0.37 | 0.00 | 0.00 | 0.38 | 0.29 | 11 | Matrix Fibroblast |
| Use1 | 0.32 | 0.00 | 0.00 | 0.36 | 0.27 | 11 | Matrix Fibroblast |
| Gm24276 | 0.30 | 0.00 | 0.00 | 0.31 | 0.22 | 11 | Matrix Fibroblast |
| Psmc1 | 0.33 | 0.00 | 0.00 | 0.40 | 0.31 | 11 | Matrix Fibroblast |
| Zfp91 | 0.33 | 0.00 | 0.00 | 0.39 | 0.30 | 11 | Matrix Fibroblast |
| Gm5619 | 0.28 | 0.00 | 0.00 | 0.42 | 0.33 | 11 | Matrix Fibroblast |
| Cul1 | 0.36 | 0.00 | 0.00 | 0.26 | 0.18 | 11 | Matrix Fibroblast |
| Ppil4 | 0.32 | 0.00 | 0.00 | 0.27 | 0.19 | 11 | Matrix Fibroblast |
| Hint1 | 0.27 | 0.00 | 0.00 | 0.54 | 0.46 | 11 | Matrix Fibroblast |
| Map1lc3a | 0.26 | 0.00 | 0.00 | 0.28 | 0.20 | 11 | Matrix Fibroblast |
| H2afv | 0.27 | 0.00 | 0.00 | 0.37 | 0.28 | 11 | Matrix Fibroblast |
| Matr3 | 0.33 | 0.00 | 0.00 | 0.48 | 0.41 | 11 | Matrix Fibroblast |
| Vezf1 | 0.30 | 0.00 | 0.00 | 0.32 | 0.24 | 11 | Matrix Fibroblast |
| Idh2 | 0.31 | 0.00 | 0.00 | 0.27 | 0.19 | 11 | Matrix Fibroblast |
| Prpf4b | 0.29 | 0.00 | 0.00 | 0.52 | 0.46 | 11 | Matrix Fibroblast |
| Gm8226 | 0.25 | 0.00 | 0.00 | 0.48 | 0.40 | 11 | Matrix Fibroblast |
| Ube2b | 0.26 | 0.00 | 0.00 | 0.44 | 0.36 | 11 | Matrix Fibroblast |
| Eea1 | 0.32 | 0.00 | 0.00 | 0.28 | 0.21 | 11 | Matrix Fibroblast |
| Nipbl | 0.30 | 0.00 | 0.00 | 0.47 | 0.39 | 11 | Matrix Fibroblast |
| Arid4a | 0.35 | 0.00 | 0.00 | 0.37 | 0.29 | 11 | Matrix Fibroblast |
| Srsf10 | 0.30 | 0.00 | 0.00 | 0.33 | 0.25 | 11 | Matrix Fibroblast |
| Gm25593 | 0.30 | 0.00 | 0.00 | 0.28 | 0.21 | 11 | Matrix Fibroblast |
| Ppig | 0.32 | 0.00 | 0.00 | 0.46 | 0.40 | 11 | Matrix Fibroblast |
| Ndufa6 | 0.26 | 0.00 | 0.00 | 0.49 | 0.42 | 11 | Matrix Fibroblast |
| Ewsr1 | 0.28 | 0.00 | 0.00 | 0.34 | 0.27 | 11 | Matrix Fibroblast |
| Bclaf1 | 0.26 | 0.00 | 0.00 | 0.48 | 0.41 | 11 | Matrix Fibroblast |
| Stt3b | 0.31 | 0.00 | 0.00 | 0.25 | 0.19 | 11 | Matrix Fibroblast |
| Prpf38b | 0.25 | 0.00 | 0.00 | 0.57 | 0.50 | 11 | Matrix Fibroblast |
| Eif4a1 | 0.26 | 0.00 | 0.00 | 0.42 | 0.36 | 11 | Matrix Fibroblast |
| Arhgap5 | 0.29 | 0.00 | 0.00 | 0.25 | 0.19 | 11 | Matrix Fibroblast |
| Zfr | 0.32 | 0.00 | 0.00 | 0.35 | 0.29 | 11 | Matrix Fibroblast |
| Fxr1 | 0.31 | 0.00 | 0.00 | 0.30 | 0.23 | 11 | Matrix Fibroblast |
| Brd3 | 0.29 | 0.00 | 0.00 | 0.32 | 0.25 | 11 | Matrix Fibroblast |
| Cast | 0.30 | 0.00 | 0.00 | 0.26 | 0.20 | 11 | Matrix Fibroblast |
| Ralbp1 | 0.26 | 0.00 | 0.00 | 0.39 | 0.33 | 11 | Matrix Fibroblast |
| Rtf1 | 0.29 | 0.00 | 0.00 | 0.40 | 0.33 | 11 | Matrix Fibroblast |
| Pdcd5 | 0.32 | 0.00 | 0.00 | 0.34 | 0.28 | 11 | Matrix Fibroblast |
| Nsa2 | 0.28 | 0.00 | 0.00 | 0.36 | 0.30 | 11 | Matrix Fibroblast |
| Tbl1x | 0.28 | 0.00 | 0.00 | 0.31 | 0.25 | 11 | Matrix Fibroblast |
| Zcchc11 | 0.26 | 0.00 | 0.00 | 0.27 | 0.21 | 11 | Matrix Fibroblast |
| Ggnbp2 | 0.26 | 0.00 | 0.00 | 0.34 | 0.28 | 11 | Matrix Fibroblast |
| Mier1 | 0.25 | 0.00 | 0.00 | 0.30 | 0.24 | 11 | Matrix Fibroblast |
| Snrpb2 | 0.29 | 0.00 | 0.00 | 0.30 | 0.24 | 11 | Matrix Fibroblast |
| Phf3 | 0.27 | 0.00 | 0.01 | 0.30 | 0.25 | 11 | Matrix Fibroblast |
| Smap1 | 0.27 | 0.00 | 0.01 | 0.25 | 0.21 | 11 | Matrix Fibroblast |
| Clip1 | 0.28 | 0.00 | 0.02 | 0.25 | 0.21 | 11 | Matrix Fibroblast |
| Esf1 | 0.30 | 0.00 | 0.04 | 0.28 | 0.24 | 11 | Matrix Fibroblast |
| Ctss | 2.69 | 0.00 | 0.00 | 0.94 | 0.13 | 12 | Macrophage |
| Ccl6 | 2.68 | 0.00 | 0.00 | 0.73 | 0.08 | 12 | Macrophage |
| Psap | 2.06 | 0.00 | 0.00 | 0.93 | 0.34 | 12 | Macrophage |
| Lgals3 | 2.01 | 0.00 | 0.00 | 0.81 | 0.13 | 12 | Macrophage |
| Bcl2a1d | 1.99 | 0.00 | 0.00 | 0.72 | 0.06 | 12 | Macrophage |
| Cybb | 1.92 | 0.00 | 0.00 | 0.70 | 0.08 | 12 | Macrophage |
| Mpeg1 | 1.90 | 0.00 | 0.00 | 0.72 | 0.05 | 12 | Macrophage |
| Bcl2a1b | 1.87 | 0.00 | 0.00 | 0.65 | 0.05 | 12 | Macrophage |
| Bcl2a1a | 1.81 | 0.00 | 0.00 | 0.63 | 0.05 | 12 | Macrophage |
| Ctsd | 1.78 | 0.00 | 0.00 | 0.78 | 0.23 | 12 | Macrophage |
| Plek | 1.59 | 0.00 | 0.00 | 0.56 | 0.05 | 12 | Macrophage |
| Cebpb | 1.59 | 0.00 | 0.00 | 0.48 | 0.08 | 12 | Macrophage |
| Fth1 | 1.58 | 0.00 | 0.00 | 0.95 | 0.81 | 12 | Macrophage |
| Atp6v0d2 | 1.53 | 0.00 | 0.00 | 0.28 | 0.01 | 12 | Macrophage |
| Fcer1g | 1.52 | 0.00 | 0.00 | 0.70 | 0.10 | 12 | Macrophage |
| Tyrobp | 1.48 | 0.00 | 0.00 | 0.73 | 0.10 | 12 | Macrophage |
| Cd300lf | 1.47 | 0.00 | 0.00 | 0.46 | 0.03 | 12 | Macrophage |
| Hebp1 | 1.45 | 0.00 | 0.00 | 0.37 | 0.03 | 12 | Macrophage |
| Ccr2 | 1.45 | 0.00 | 0.00 | 0.39 | 0.06 | 12 | Macrophage |
| Cd44 | 1.45 | 0.00 | 0.00 | 0.68 | 0.18 | 12 | Macrophage |
| Laptm5 | 1.43 | 0.00 | 0.00 | 0.71 | 0.12 | 12 | Macrophage |
| Cyba | 1.42 | 0.00 | 0.00 | 0.81 | 0.28 | 12 | Macrophage |
| Tnfaip2 | 1.42 | 0.00 | 0.00 | 0.42 | 0.04 | 12 | Macrophage |
| Ear2 | 1.40 | 0.00 | 0.00 | 0.32 | 0.02 | 12 | Macrophage |
| Mrc1 | 1.38 | 0.00 | 0.00 | 0.35 | 0.02 | 12 | Macrophage |
| Sirpa | 1.36 | 0.00 | 0.00 | 0.49 | 0.05 | 12 | Macrophage |
| Plin2 | 1.34 | 0.00 | 0.00 | 0.54 | 0.12 | 12 | Macrophage |
| Lilrb4 | 1.33 | 0.00 | 0.00 | 0.47 | 0.05 | 12 | Macrophage |
| Pygl | 1.33 | 0.00 | 0.00 | 0.51 | 0.10 | 12 | Macrophage |
| Lcp1 | 1.32 | 0.00 | 0.00 | 0.72 | 0.21 | 12 | Macrophage |
| Snx10 | 1.28 | 0.00 | 0.00 | 0.44 | 0.03 | 12 | Macrophage |
| Clec4n | 1.26 | 0.00 | 0.00 | 0.34 | 0.01 | 12 | Macrophage |
| Baz1a | 1.25 | 0.00 | 0.00 | 0.51 | 0.12 | 12 | Macrophage |
| Gda | 1.24 | 0.00 | 0.00 | 0.42 | 0.04 | 12 | Macrophage |
| Alox5ap | 1.24 | 0.00 | 0.00 | 0.58 | 0.08 | 12 | Macrophage |
| 1810033B17Rik | 1.21 | 0.00 | 0.00 | 0.39 | 0.03 | 12 | Macrophage |
| Ctsc | 1.20 | 0.00 | 0.00 | 0.63 | 0.18 | 12 | Macrophage |
| Fcgr3 | 1.20 | 0.00 | 0.00 | 0.44 | 0.03 | 12 | Macrophage |
| Fpr2 | 1.18 | 0.00 | 0.00 | 0.30 | 0.02 | 12 | Macrophage |
| Csf2rb | 1.17 | 0.00 | 0.00 | 0.41 | 0.03 | 12 | Macrophage |
| Ccr1 | 1.16 | 0.00 | 0.00 | 0.39 | 0.02 | 12 | Macrophage |
| Gp49a | 1.14 | 0.00 | 0.00 | 0.39 | 0.04 | 12 | Macrophage |
| Abcg1 | 1.14 | 0.00 | 0.00 | 0.34 | 0.05 | 12 | Macrophage |
| Slfn2 | 1.13 | 0.00 | 0.00 | 0.57 | 0.12 | 12 | Macrophage |
| Trf | 1.10 | 0.00 | 0.00 | 0.43 | 0.09 | 12 | Macrophage |
| 4930506M07Rik | 1.10 | 0.00 | 0.00 | 0.37 | 0.04 | 12 | Macrophage |
| Lyz2 | 1.09 | 0.00 | 0.00 | 0.85 | 0.25 | 12 | Macrophage |
| Msrb1 | 1.09 | 0.00 | 0.00 | 0.55 | 0.13 | 12 | Macrophage |
| Clec4a2 | 1.05 | 0.00 | 0.00 | 0.31 | 0.02 | 12 | Macrophage |
| Fam49b | 1.04 | 0.00 | 0.00 | 0.45 | 0.10 | 12 | Macrophage |
| Spi1 | 1.04 | 0.00 | 0.00 | 0.44 | 0.05 | 12 | Macrophage |
| Ms4a6c | 1.03 | 0.00 | 0.00 | 0.37 | 0.05 | 12 | Macrophage |
| AI662270 | 1.03 | 0.00 | 0.00 | 0.40 | 0.06 | 12 | Macrophage |
| Sirpb1c | 1.02 | 0.00 | 0.00 | 0.34 | 0.03 | 12 | Macrophage |
| Cd84 | 1.01 | 0.00 | 0.00 | 0.36 | 0.04 | 12 | Macrophage |
| Il18 | 1.01 | 0.00 | 0.00 | 0.25 | 0.01 | 12 | Macrophage |
| Cfp | 1.01 | 0.00 | 0.00 | 0.29 | 0.04 | 12 | Macrophage |
| Coro2a | 1.00 | 0.00 | 0.00 | 0.35 | 0.06 | 12 | Macrophage |
| Fcgr2b | 0.99 | 0.00 | 0.00 | 0.30 | 0.03 | 12 | Macrophage |
| Tmsb4x | 0.98 | 0.00 | 0.00 | 0.99 | 0.88 | 12 | Macrophage |
| Ccl9 | 0.98 | 0.00 | 0.00 | 0.29 | 0.02 | 12 | Macrophage |
| Slc15a3 | 0.97 | 0.00 | 0.00 | 0.31 | 0.02 | 12 | Macrophage |
| Apbb1ip | 0.96 | 0.00 | 0.00 | 0.45 | 0.08 | 12 | Macrophage |
| Ncf1 | 0.96 | 0.00 | 0.00 | 0.39 | 0.06 | 12 | Macrophage |
| Itgb2 | 0.95 | 0.00 | 0.00 | 0.33 | 0.04 | 12 | Macrophage |
| Arhgap30 | 0.94 | 0.00 | 0.00 | 0.46 | 0.09 | 12 | Macrophage |
| Sh3bp2 | 0.89 | 0.00 | 0.00 | 0.26 | 0.03 | 12 | Macrophage |
| Csf2ra | 0.88 | 0.00 | 0.00 | 0.34 | 0.05 | 12 | Macrophage |
| Axl | 0.87 | 0.00 | 0.00 | 0.28 | 0.03 | 12 | Macrophage |
| Clec4a3 | 0.87 | 0.00 | 0.00 | 0.31 | 0.03 | 12 | Macrophage |
| Pld4 | 0.83 | 0.00 | 0.00 | 0.32 | 0.05 | 12 | Macrophage |
| Sirpb1b | 0.82 | 0.00 | 0.00 | 0.28 | 0.02 | 12 | Macrophage |
| Cd68 | 0.81 | 0.00 | 0.00 | 0.28 | 0.02 | 12 | Macrophage |
| AF251705 | 0.80 | 0.00 | 0.00 | 0.26 | 0.02 | 12 | Macrophage |
| Ncf2 | 0.77 | 0.00 | 0.00 | 0.28 | 0.04 | 12 | Macrophage |
| Igsf6 | 0.76 | 0.00 | 0.00 | 0.28 | 0.04 | 12 | Macrophage |
| Klhdc4 | 1.12 | 0.00 | 0.00 | 0.26 | 0.04 | 12 | Macrophage |
| Cd302 | 1.07 | 0.00 | 0.00 | 0.52 | 0.13 | 12 | Macrophage |
| Cstb | 1.74 | 0.00 | 0.00 | 0.67 | 0.26 | 12 | Macrophage |
| Hcls1 | 0.81 | 0.00 | 0.00 | 0.37 | 0.07 | 12 | Macrophage |
| Sla | 0.80 | 0.00 | 0.00 | 0.26 | 0.04 | 12 | Macrophage |
| Cd300a | 0.70 | 0.00 | 0.00 | 0.25 | 0.03 | 12 | Macrophage |
| Cotl1 | 1.05 | 0.00 | 0.00 | 0.57 | 0.16 | 12 | Macrophage |
| Rassf4 | 0.71 | 0.00 | 0.00 | 0.26 | 0.03 | 12 | Macrophage |
| Cyth4 | 0.74 | 0.00 | 0.00 | 0.27 | 0.04 | 12 | Macrophage |
| Naaa | 0.81 | 0.00 | 0.00 | 0.26 | 0.04 | 12 | Macrophage |
| Gm10116 | 1.24 | 0.00 | 0.00 | 0.88 | 0.62 | 12 | Macrophage |
| Ptprc | 0.74 | 0.00 | 0.00 | 0.53 | 0.14 | 12 | Macrophage |
| Gm13139 | 0.83 | 0.00 | 0.00 | 0.34 | 0.06 | 12 | Macrophage |
| Aprt | 1.04 | 0.00 | 0.00 | 0.53 | 0.16 | 12 | Macrophage |
| Ifi204 | 0.87 | 0.00 | 0.00 | 0.28 | 0.05 | 12 | Macrophage |
| Abhd12 | 0.77 | 0.00 | 0.00 | 0.26 | 0.04 | 12 | Macrophage |
| Ptpn6 | 0.81 | 0.00 | 0.00 | 0.37 | 0.08 | 12 | Macrophage |
| Ifi30 | 0.91 | 0.00 | 0.00 | 0.42 | 0.10 | 12 | Macrophage |
| Ftl1 | 1.25 | 0.00 | 0.00 | 0.84 | 0.56 | 12 | Macrophage |
| Gns | 1.00 | 0.00 | 0.00 | 0.45 | 0.12 | 12 | Macrophage |
| Efhd2 | 0.89 | 0.00 | 0.00 | 0.37 | 0.08 | 12 | Macrophage |
| Pla2g7 | 1.02 | 0.00 | 0.00 | 0.29 | 0.05 | 12 | Macrophage |
| Gm13248 | 0.77 | 0.00 | 0.00 | 0.29 | 0.05 | 12 | Macrophage |
| Gsr | 0.96 | 0.00 | 0.00 | 0.36 | 0.08 | 12 | Macrophage |
| Tgfbi | 0.35 | 0.00 | 0.00 | 0.60 | 0.18 | 12 | Macrophage |
| Ctsz | 1.07 | 0.00 | 0.00 | 0.47 | 0.14 | 12 | Macrophage |
| Cd52 | 0.80 | 0.00 | 0.00 | 0.60 | 0.19 | 12 | Macrophage |
| Ctsb | 1.31 | 0.00 | 0.00 | 0.64 | 0.27 | 12 | Macrophage |
| Fxyd5 | 0.86 | 0.00 | 0.00 | 0.54 | 0.17 | 12 | Macrophage |
| Mnda | 0.98 | 0.00 | 0.00 | 0.28 | 0.05 | 12 | Macrophage |
| Lyz1 | 0.57 | 0.00 | 0.00 | 0.58 | 0.18 | 12 | Macrophage |
| Dab2 | 0.81 | 0.00 | 0.00 | 0.32 | 0.07 | 12 | Macrophage |
| Cd53 | 0.73 | 0.00 | 0.00 | 0.37 | 0.08 | 12 | Macrophage |
| Atp6v0c | 1.02 | 0.00 | 0.00 | 0.61 | 0.24 | 12 | Macrophage |
| Zfp36l2 | 0.92 | 0.00 | 0.00 | 0.40 | 0.11 | 12 | Macrophage |
| Mt1 | 1.04 | 0.00 | 0.00 | 0.49 | 0.15 | 12 | Macrophage |
| Gpx1 | 0.89 | 0.00 | 0.00 | 0.78 | 0.42 | 12 | Macrophage |
| Pycard | 0.80 | 0.00 | 0.00 | 0.35 | 0.08 | 12 | Macrophage |
| Pld3 | 0.94 | 0.00 | 0.00 | 0.28 | 0.06 | 12 | Macrophage |
| Taldo1 | 0.96 | 0.00 | 0.00 | 0.62 | 0.25 | 12 | Macrophage |
| Sat1 | 0.98 | 0.00 | 0.00 | 0.69 | 0.32 | 12 | Macrophage |
| Lst1 | 0.73 | 0.00 | 0.00 | 0.29 | 0.06 | 12 | Macrophage |
| Unc93b1 | 0.72 | 0.00 | 0.00 | 0.31 | 0.07 | 12 | Macrophage |
| Lrp1 | 0.83 | 0.00 | 0.00 | 0.39 | 0.11 | 12 | Macrophage |
| Hexa | 0.78 | 0.00 | 0.00 | 0.31 | 0.07 | 12 | Macrophage |
| Lpl | 1.53 | 0.00 | 0.00 | 0.42 | 0.14 | 12 | Macrophage |
| Nceh1 | 0.86 | 0.00 | 0.00 | 0.28 | 0.06 | 12 | Macrophage |
| Gm4070 | 0.85 | 0.00 | 0.00 | 0.40 | 0.11 | 12 | Macrophage |
| Gvin1 | 0.86 | 0.00 | 0.00 | 0.39 | 0.11 | 12 | Macrophage |
| Tpd52 | 0.77 | 0.00 | 0.00 | 0.37 | 0.10 | 12 | Macrophage |
| Ctsa | 0.84 | 0.00 | 0.00 | 0.41 | 0.12 | 12 | Macrophage |
| Llph | 0.99 | 0.00 | 0.00 | 0.67 | 0.33 | 12 | Macrophage |
| M6pr | 0.92 | 0.00 | 0.00 | 0.56 | 0.22 | 12 | Macrophage |
| Pabpc1 | 0.61 | 0.00 | 0.00 | 0.94 | 0.80 | 12 | Macrophage |
| Hmga1-rs1 | 0.74 | 0.00 | 0.00 | 0.29 | 0.07 | 12 | Macrophage |
| Creg1 | 0.90 | 0.00 | 0.00 | 0.47 | 0.16 | 12 | Macrophage |
| Hmga1 | 0.71 | 0.00 | 0.00 | 0.29 | 0.07 | 12 | Macrophage |
| Arhgdib | 0.60 | 0.00 | 0.00 | 0.47 | 0.15 | 12 | Macrophage |
| Arpc5 | 0.73 | 0.00 | 0.00 | 0.84 | 0.56 | 12 | Macrophage |
| Ubash3b | 0.65 | 0.00 | 0.00 | 0.27 | 0.06 | 12 | Macrophage |
| Hexb | 0.68 | 0.00 | 0.00 | 0.28 | 0.07 | 12 | Macrophage |
| Tcf7l2 | 0.80 | 0.00 | 0.00 | 0.30 | 0.08 | 12 | Macrophage |
| Arrb2 | 0.61 | 0.00 | 0.00 | 0.26 | 0.06 | 12 | Macrophage |
| Prdx5 | 0.94 | 0.00 | 0.00 | 0.63 | 0.31 | 12 | Macrophage |
| Grn | 0.80 | 0.00 | 0.00 | 0.39 | 0.13 | 12 | Macrophage |
| Fermt3 | 0.57 | 0.00 | 0.00 | 0.25 | 0.06 | 12 | Macrophage |
| Btg1 | 0.88 | 0.00 | 0.00 | 0.59 | 0.28 | 12 | Macrophage |
| Fam96a | 0.80 | 0.00 | 0.00 | 0.41 | 0.14 | 12 | Macrophage |
| Apoe | 1.55 | 0.00 | 0.00 | 0.57 | 0.28 | 12 | Macrophage |
| Coro1a | 0.55 | 0.00 | 0.00 | 0.34 | 0.09 | 12 | Macrophage |
| mt-Rnr1 | 1.34 | 0.00 | 0.00 | 0.72 | 0.46 | 12 | Macrophage |
| Pgd | 0.76 | 0.00 | 0.00 | 0.34 | 0.10 | 12 | Macrophage |
| Atp6v1b2 | 0.85 | 0.00 | 0.00 | 0.37 | 0.12 | 12 | Macrophage |
| Gm2a | 0.64 | 0.00 | 0.00 | 0.31 | 0.08 | 12 | Macrophage |
| Fos | 0.84 | 0.00 | 0.00 | 0.48 | 0.18 | 12 | Macrophage |
| Rps27a | 0.61 | 0.00 | 0.00 | 0.91 | 0.76 | 12 | Macrophage |
| Shfm1 | 0.71 | 0.00 | 0.00 | 0.86 | 0.64 | 12 | Macrophage |
| Actr2 | 0.79 | 0.00 | 0.00 | 0.57 | 0.27 | 12 | Macrophage |
| Runx1 | 0.68 | 0.00 | 0.00 | 0.26 | 0.07 | 12 | Macrophage |
| Actb | 0.48 | 0.00 | 0.00 | 1.00 | 0.98 | 12 | Macrophage |
| Prkcd | 0.68 | 0.00 | 0.00 | 0.34 | 0.10 | 12 | Macrophage |
| Cd9 | 1.09 | 0.00 | 0.00 | 0.62 | 0.34 | 12 | Macrophage |
| Mir692-2a | 0.92 | 0.00 | 0.00 | 0.46 | 0.19 | 12 | Macrophage |
| Adipor2 | 0.84 | 0.00 | 0.00 | 0.34 | 0.11 | 12 | Macrophage |
| Ppt1 | 0.70 | 0.00 | 0.00 | 0.36 | 0.12 | 12 | Macrophage |
| Atp6v1e1 | 0.88 | 0.00 | 0.00 | 0.53 | 0.25 | 12 | Macrophage |
| Samhd1 | 0.67 | 0.00 | 0.00 | 0.47 | 0.19 | 12 | Macrophage |
| Tgfbr1 | 0.71 | 0.00 | 0.00 | 0.26 | 0.07 | 12 | Macrophage |
| Psmb10 | 0.65 | 0.00 | 0.00 | 0.35 | 0.12 | 12 | Macrophage |
| Mcl1 | 0.78 | 0.00 | 0.00 | 0.49 | 0.22 | 12 | Macrophage |
| Iqgap1 | 0.75 | 0.00 | 0.00 | 0.71 | 0.44 | 12 | Macrophage |
| Gm22774 | 0.82 | 0.00 | 0.00 | 0.41 | 0.16 | 12 | Macrophage |
| Wdr89 | 0.44 | 0.00 | 0.00 | 0.97 | 0.89 | 12 | Macrophage |
| Gm26445 | 0.84 | 0.00 | 0.00 | 0.38 | 0.15 | 12 | Macrophage |
| Reep5 | 0.79 | 0.00 | 0.00 | 0.50 | 0.23 | 12 | Macrophage |
| Capza2 | 0.70 | 0.00 | 0.00 | 0.62 | 0.34 | 12 | Macrophage |
| Gusb | 0.58 | 0.00 | 0.00 | 0.27 | 0.08 | 12 | Macrophage |
| Atp6v0b | 0.70 | 0.00 | 0.00 | 0.44 | 0.18 | 12 | Macrophage |
| Rplp1 | 0.44 | 0.00 | 0.00 | 0.96 | 0.90 | 12 | Macrophage |
| Capg | 0.58 | 0.00 | 0.00 | 0.31 | 0.10 | 12 | Macrophage |
| Lamp1 | 0.67 | 0.00 | 0.00 | 0.69 | 0.42 | 12 | Macrophage |
| Cndp2 | 0.65 | 0.00 | 0.00 | 0.27 | 0.08 | 12 | Macrophage |
| Anp32b | 0.77 | 0.00 | 0.00 | 0.74 | 0.49 | 12 | Macrophage |
| Lgmn | 0.76 | 0.00 | 0.00 | 0.34 | 0.12 | 12 | Macrophage |
| Emb | 0.64 | 0.00 | 0.00 | 0.32 | 0.11 | 12 | Macrophage |
| Rplp2 | 0.45 | 0.00 | 0.00 | 0.93 | 0.81 | 12 | Macrophage |
| Plgrkt | 0.61 | 0.00 | 0.00 | 0.31 | 0.11 | 12 | Macrophage |
| Tkt | 0.73 | 0.00 | 0.00 | 0.36 | 0.14 | 12 | Macrophage |
| Arpc1b | 0.69 | 0.00 | 0.00 | 0.63 | 0.37 | 12 | Macrophage |
| Neat1 | 0.61 | 0.00 | 0.00 | 0.56 | 0.26 | 12 | Macrophage |
| Gm10073 | 0.44 | 0.00 | 0.00 | 0.95 | 0.85 | 12 | Macrophage |
| Gm22751 | 0.82 | 0.00 | 0.00 | 0.38 | 0.16 | 12 | Macrophage |
| Soat1 | 0.68 | 0.00 | 0.00 | 0.30 | 0.10 | 12 | Macrophage |
| Gm9843 | 0.40 | 0.00 | 0.00 | 0.97 | 0.92 | 12 | Macrophage |
| Sdcbp | 0.76 | 0.00 | 0.00 | 0.52 | 0.27 | 12 | Macrophage |
| Rab8b | 0.60 | 0.00 | 0.00 | 0.28 | 0.09 | 12 | Macrophage |
| Lipa | 0.66 | 0.00 | 0.00 | 0.32 | 0.12 | 12 | Macrophage |
| Akap13 | 0.66 | 0.00 | 0.00 | 0.52 | 0.26 | 12 | Macrophage |
| Rpsa | 0.53 | 0.00 | 0.00 | 0.81 | 0.61 | 12 | Macrophage |
| H2-D1 | 0.54 | 0.00 | 0.00 | 0.76 | 0.49 | 12 | Macrophage |
| Idh1 | 0.67 | 0.00 | 0.00 | 0.27 | 0.09 | 12 | Macrophage |
| mt-Rnr2 | 0.40 | 0.00 | 0.00 | 1.00 | 0.99 | 12 | Macrophage |
| Dbi | 0.69 | 0.00 | 0.00 | 0.61 | 0.36 | 12 | Macrophage |
| Rpl37a | 0.44 | 0.00 | 0.00 | 0.93 | 0.83 | 12 | Macrophage |
| Gngt2 | 0.53 | 0.00 | 0.00 | 0.29 | 0.10 | 12 | Macrophage |
| Fau | 0.43 | 0.00 | 0.00 | 0.93 | 0.87 | 12 | Macrophage |
| Gabarap | 0.68 | 0.00 | 0.00 | 0.65 | 0.44 | 12 | Macrophage |
| Gm26924 | 1.51 | 0.00 | 0.00 | 0.76 | 0.56 | 12 | Macrophage |
| 9-Sep | 0.58 | 0.00 | 0.00 | 0.26 | 0.09 | 12 | Macrophage |
| Mir682 | 0.43 | 0.00 | 0.00 | 0.94 | 0.85 | 12 | Macrophage |
| Sgk1 | 0.77 | 0.00 | 0.00 | 0.41 | 0.18 | 12 | Macrophage |
| Rpl35a | 0.44 | 0.00 | 0.00 | 0.91 | 0.79 | 12 | Macrophage |
| Rpl38 | 0.49 | 0.00 | 0.00 | 0.86 | 0.73 | 12 | Macrophage |
| Clta | 0.58 | 0.00 | 0.00 | 0.66 | 0.42 | 12 | Macrophage |
| Rpsa-ps10 | 0.46 | 0.00 | 0.00 | 0.84 | 0.67 | 12 | Macrophage |
| Actr3 | 0.60 | 0.00 | 0.00 | 0.57 | 0.32 | 12 | Macrophage |
| Gm22758 | 0.50 | 0.00 | 0.00 | 0.84 | 0.68 | 12 | Macrophage |
| Lmo4 | 0.75 | 0.00 | 0.00 | 0.48 | 0.25 | 12 | Macrophage |
| Arpc2 | 0.52 | 0.00 | 0.00 | 0.79 | 0.61 | 12 | Macrophage |
| Tmed5 | 0.56 | 0.00 | 0.00 | 0.38 | 0.17 | 12 | Macrophage |
| Rps9 | 0.34 | 0.00 | 0.00 | 0.96 | 0.91 | 12 | Macrophage |
| 1810037I17Rik | 0.71 | 0.00 | 0.00 | 0.55 | 0.31 | 12 | Macrophage |
| Smpdl3a | 0.58 | 0.00 | 0.00 | 0.28 | 0.10 | 12 | Macrophage |
| Pitpna | 0.54 | 0.00 | 0.00 | 0.30 | 0.12 | 12 | Macrophage |
| Ptbp3 | 0.60 | 0.00 | 0.00 | 0.59 | 0.36 | 12 | Macrophage |
| Zeb2 | 0.52 | 0.00 | 0.00 | 0.56 | 0.30 | 12 | Macrophage |
| Txn1 | 0.64 | 0.00 | 0.00 | 0.70 | 0.49 | 12 | Macrophage |
| Rnh1 | 0.64 | 0.00 | 0.00 | 0.34 | 0.15 | 12 | Macrophage |
| Fabp5 | 0.88 | 0.00 | 0.00 | 0.31 | 0.13 | 12 | Macrophage |
| Fam111a | 0.50 | 0.00 | 0.00 | 0.35 | 0.15 | 12 | Macrophage |
| Nme1 | 0.64 | 0.00 | 0.00 | 0.39 | 0.18 | 12 | Macrophage |
| Asah1 | 0.54 | 0.00 | 0.00 | 0.30 | 0.12 | 12 | Macrophage |
| Cox5a | 0.62 | 0.00 | 0.00 | 0.54 | 0.31 | 12 | Macrophage |
| Serp1 | 0.58 | 0.00 | 0.00 | 0.44 | 0.22 | 12 | Macrophage |
| Il6ra | 0.46 | 0.00 | 0.00 | 0.27 | 0.10 | 12 | Macrophage |
| Gm10076 | 0.35 | 0.00 | 0.00 | 0.97 | 0.92 | 12 | Macrophage |
| Ccdc88a | 0.65 | 0.00 | 0.00 | 0.46 | 0.24 | 12 | Macrophage |
| BC005537 | 0.58 | 0.00 | 0.00 | 0.55 | 0.33 | 12 | Macrophage |
| Sh3bgrl3 | 0.56 | 0.00 | 0.00 | 0.57 | 0.34 | 12 | Macrophage |
| Cap1 | 0.55 | 0.00 | 0.00 | 0.36 | 0.16 | 12 | Macrophage |
| Cox4i1 | 0.43 | 0.00 | 0.00 | 0.87 | 0.73 | 12 | Macrophage |
| Psme2b | 0.63 | 0.00 | 0.00 | 0.50 | 0.29 | 12 | Macrophage |
| Coro1b | 0.53 | 0.00 | 0.00 | 0.46 | 0.24 | 12 | Macrophage |
| Gm5506 | 0.55 | 0.00 | 0.00 | 0.32 | 0.14 | 12 | Macrophage |
| Myeov2 | 0.59 | 0.00 | 0.00 | 0.51 | 0.29 | 12 | Macrophage |
| Stat1 | 0.58 | 0.00 | 0.00 | 0.26 | 0.10 | 12 | Macrophage |
| Gm17511 | 0.51 | 0.00 | 0.00 | 0.76 | 0.61 | 12 | Macrophage |
| Eno1 | 0.54 | 0.00 | 0.00 | 0.34 | 0.16 | 12 | Macrophage |
| Tpm3-rs7 | 0.42 | 0.00 | 0.00 | 0.86 | 0.69 | 12 | Macrophage |
| Colgalt1 | 0.57 | 0.00 | 0.00 | 0.27 | 0.11 | 12 | Macrophage |
| B2m | 0.64 | 0.00 | 0.00 | 0.81 | 0.67 | 12 | Macrophage |
| Cd47 | 0.58 | 0.00 | 0.00 | 0.64 | 0.43 | 12 | Macrophage |
| Gm22759 | 0.49 | 0.00 | 0.00 | 0.79 | 0.64 | 12 | Macrophage |
| Rps29 | 0.39 | 0.00 | 0.00 | 0.94 | 0.91 | 12 | Macrophage |
| Bola2 | 0.59 | 0.00 | 0.00 | 0.34 | 0.16 | 12 | Macrophage |
| Psme2 | 0.60 | 0.00 | 0.00 | 0.43 | 0.23 | 12 | Macrophage |
| Junb | 0.61 | 0.00 | 0.00 | 0.38 | 0.19 | 12 | Macrophage |
| Snx1 | 0.52 | 0.00 | 0.00 | 0.27 | 0.11 | 12 | Macrophage |
| Fyb | 0.48 | 0.00 | 0.00 | 0.37 | 0.18 | 12 | Macrophage |
| Eif3a | 0.55 | 0.00 | 0.00 | 0.77 | 0.60 | 12 | Macrophage |
| Rpl22 | 0.37 | 0.00 | 0.00 | 0.89 | 0.79 | 12 | Macrophage |
| Psmb8 | 0.49 | 0.00 | 0.00 | 0.38 | 0.18 | 12 | Macrophage |
| Mycbp2 | 0.55 | 0.00 | 0.00 | 0.43 | 0.23 | 12 | Macrophage |
| Atp6ap2 | 0.57 | 0.00 | 0.00 | 0.32 | 0.15 | 12 | Macrophage |
| Nmt1 | 0.53 | 0.00 | 0.00 | 0.42 | 0.22 | 12 | Macrophage |
| Ncl | 0.45 | 0.00 | 0.00 | 0.88 | 0.78 | 12 | Macrophage |
| Rpl41 | 0.30 | 0.00 | 0.00 | 0.97 | 0.95 | 12 | Macrophage |
| Snx5 | 0.47 | 0.00 | 0.00 | 0.29 | 0.13 | 12 | Macrophage |
| Cox17 | 0.63 | 0.00 | 0.00 | 0.43 | 0.24 | 12 | Macrophage |
| Itgam | 0.56 | 0.00 | 0.00 | 0.47 | 0.28 | 12 | Macrophage |
| Gm24865 | 0.44 | 0.00 | 0.00 | 0.77 | 0.62 | 12 | Macrophage |
| Lrrfip1 | 0.51 | 0.00 | 0.00 | 0.45 | 0.25 | 12 | Macrophage |
| Ucp2 | 0.42 | 0.00 | 0.00 | 0.28 | 0.12 | 12 | Macrophage |
| Abracl | 0.57 | 0.00 | 0.00 | 0.42 | 0.23 | 12 | Macrophage |
| 2010107E04Rik | 0.48 | 0.00 | 0.00 | 0.60 | 0.40 | 12 | Macrophage |
| Tmem160 | 0.44 | 0.00 | 0.00 | 0.26 | 0.11 | 12 | Macrophage |
| Prdx1 | 0.62 | 0.00 | 0.00 | 0.67 | 0.50 | 12 | Macrophage |
| Lsp1 | 0.28 | 0.00 | 0.00 | 0.39 | 0.19 | 12 | Macrophage |
| Fn1 | 0.34 | 0.00 | 0.00 | 0.45 | 0.23 | 12 | Macrophage |
| Ptpn18 | 0.40 | 0.00 | 0.00 | 0.31 | 0.14 | 12 | Macrophage |
| Mndal | 0.54 | 0.00 | 0.00 | 0.34 | 0.16 | 12 | Macrophage |
| Arpc4 | 0.45 | 0.00 | 0.00 | 0.35 | 0.17 | 12 | Macrophage |
| Lyn | 0.41 | 0.00 | 0.00 | 0.29 | 0.13 | 12 | Macrophage |
| Etfb | 0.55 | 0.00 | 0.00 | 0.47 | 0.28 | 12 | Macrophage |
| Pfn1 | 0.45 | 0.00 | 0.00 | 0.65 | 0.46 | 12 | Macrophage |
| S100a6 | 0.26 | 0.00 | 0.00 | 0.53 | 0.31 | 12 | Macrophage |
| Skap2 | 0.42 | 0.00 | 0.00 | 0.28 | 0.12 | 12 | Macrophage |
| Gm22426 | 0.46 | 0.00 | 0.00 | 0.65 | 0.48 | 12 | Macrophage |
| Gm10221 | 0.47 | 0.00 | 0.00 | 0.63 | 0.46 | 12 | Macrophage |
| Tmem14c | 0.52 | 0.00 | 0.00 | 0.38 | 0.21 | 12 | Macrophage |
| Hspa8 | 0.41 | 0.00 | 0.00 | 0.81 | 0.67 | 12 | Macrophage |
| Eif2s2 | 0.51 | 0.00 | 0.00 | 0.64 | 0.47 | 12 | Macrophage |
| Ptpn12 | 0.55 | 0.00 | 0.00 | 0.28 | 0.13 | 12 | Macrophage |
| Lamtor2 | 0.51 | 0.00 | 0.00 | 0.31 | 0.15 | 12 | Macrophage |
| Eif5b | 0.45 | 0.00 | 0.00 | 0.69 | 0.51 | 12 | Macrophage |
| Atp6ap1 | 0.52 | 0.00 | 0.00 | 0.27 | 0.12 | 12 | Macrophage |
| Gm5244 | 0.44 | 0.00 | 0.00 | 0.47 | 0.28 | 12 | Macrophage |
| Chmp4b | 0.52 | 0.00 | 0.00 | 0.37 | 0.20 | 12 | Macrophage |
| Atp5l | 0.46 | 0.00 | 0.00 | 0.61 | 0.43 | 12 | Macrophage |
| Tpm3 | 0.34 | 0.00 | 0.00 | 0.84 | 0.70 | 12 | Macrophage |
| Tapbp | 0.50 | 0.00 | 0.00 | 0.30 | 0.14 | 12 | Macrophage |
| S100a10 | 0.39 | 0.00 | 0.00 | 0.58 | 0.38 | 12 | Macrophage |
| Gm26300 | 0.49 | 0.00 | 0.00 | 0.55 | 0.37 | 12 | Macrophage |
| H2-K1 | 0.38 | 0.00 | 0.00 | 0.52 | 0.32 | 12 | Macrophage |
| Sod2 | 0.57 | 0.00 | 0.00 | 0.27 | 0.13 | 12 | Macrophage |
| Rgs2 | 0.32 | 0.00 | 0.00 | 0.47 | 0.28 | 12 | Macrophage |
| 9930111J21Rik2 | 0.32 | 0.00 | 0.00 | 0.26 | 0.12 | 12 | Macrophage |
| Cd164 | 0.59 | 0.00 | 0.00 | 0.27 | 0.14 | 12 | Macrophage |
| Ctage5 | 0.49 | 0.00 | 0.00 | 0.46 | 0.28 | 12 | Macrophage |
| Gmfg | 0.33 | 0.00 | 0.00 | 0.29 | 0.14 | 12 | Macrophage |
| Cmpk1 | 0.48 | 0.00 | 0.00 | 0.37 | 0.21 | 12 | Macrophage |
| Anxa1 | 0.28 | 0.00 | 0.00 | 0.35 | 0.18 | 12 | Macrophage |
| Erp29 | 0.45 | 0.00 | 0.00 | 0.43 | 0.26 | 12 | Macrophage |
| Gbp7 | 0.46 | 0.00 | 0.00 | 0.28 | 0.14 | 12 | Macrophage |
| Ap1s2 | 0.54 | 0.00 | 0.00 | 0.25 | 0.12 | 12 | Macrophage |
| Diap1 | 0.44 | 0.00 | 0.00 | 0.28 | 0.14 | 12 | Macrophage |
| Rps10-ps1 | 0.28 | 0.00 | 0.00 | 0.87 | 0.77 | 12 | Macrophage |
| Zfp36 | 0.48 | 0.00 | 0.00 | 0.31 | 0.16 | 12 | Macrophage |
| Pkm | 0.48 | 0.00 | 0.00 | 0.38 | 0.22 | 12 | Macrophage |
| Ddx21 | 0.47 | 0.00 | 0.00 | 0.42 | 0.26 | 12 | Macrophage |
| Akr1a1 | 0.41 | 0.00 | 0.00 | 0.65 | 0.49 | 12 | Macrophage |
| Rps10 | 0.29 | 0.00 | 0.00 | 0.81 | 0.70 | 12 | Macrophage |
| Psmd13 | 0.47 | 0.00 | 0.00 | 0.31 | 0.17 | 12 | Macrophage |
| Rpl23 | 0.27 | 0.00 | 0.00 | 0.91 | 0.87 | 12 | Macrophage |
| Znhit1 | 0.41 | 0.00 | 0.00 | 0.27 | 0.14 | 12 | Macrophage |
| Atp6v1g1 | 0.48 | 0.00 | 0.00 | 0.48 | 0.32 | 12 | Macrophage |
| Atp6v0e | 0.38 | 0.00 | 0.00 | 0.35 | 0.20 | 12 | Macrophage |
| Utp14a | 0.42 | 0.00 | 0.00 | 0.30 | 0.15 | 12 | Macrophage |
| Rrbp1 | 0.36 | 0.00 | 0.00 | 0.70 | 0.55 | 12 | Macrophage |
| Prr13 | 0.37 | 0.00 | 0.00 | 0.26 | 0.13 | 12 | Macrophage |
| Atp1b3 | 0.43 | 0.00 | 0.00 | 0.41 | 0.25 | 12 | Macrophage |
| Gnai2 | 0.34 | 0.00 | 0.00 | 0.56 | 0.38 | 12 | Macrophage |
| Pomp | 0.40 | 0.00 | 0.00 | 0.47 | 0.31 | 12 | Macrophage |
| Ptpn1 | 0.40 | 0.00 | 0.00 | 0.35 | 0.20 | 12 | Macrophage |
| Tmbim6 | 0.42 | 0.00 | 0.00 | 0.53 | 0.37 | 12 | Macrophage |
| Aldoa | 0.41 | 0.00 | 0.00 | 0.33 | 0.19 | 12 | Macrophage |
| Ahnak | 0.34 | 0.00 | 0.00 | 0.61 | 0.43 | 12 | Macrophage |
| Atp5l-ps1 | 0.42 | 0.00 | 0.00 | 0.42 | 0.27 | 12 | Macrophage |
| Gm25593 | 0.41 | 0.00 | 0.00 | 0.35 | 0.21 | 12 | Macrophage |
| Eif5a | 0.42 | 0.00 | 0.00 | 0.49 | 0.34 | 12 | Macrophage |
| Rexo2 | 0.44 | 0.00 | 0.00 | 0.27 | 0.14 | 12 | Macrophage |
| Rbm25 | 0.36 | 0.00 | 0.00 | 0.80 | 0.68 | 12 | Macrophage |
| Hsp90b1 | 0.40 | 0.00 | 0.00 | 0.84 | 0.76 | 12 | Macrophage |
| Anxa2 | 0.44 | 0.00 | 0.00 | 0.62 | 0.47 | 12 | Macrophage |
| Rpl36al | 0.34 | 0.00 | 0.00 | 0.67 | 0.52 | 12 | Macrophage |
| Ubl3 | 0.38 | 0.00 | 0.00 | 0.33 | 0.19 | 12 | Macrophage |
| Ivns1abp | 0.34 | 0.00 | 0.00 | 0.37 | 0.22 | 12 | Macrophage |
| Emp3 | 0.34 | 0.00 | 0.00 | 0.36 | 0.21 | 12 | Macrophage |
| Arpc3 | 0.35 | 0.00 | 0.00 | 0.41 | 0.26 | 12 | Macrophage |
| Glud1 | 0.35 | 0.00 | 0.00 | 0.31 | 0.17 | 12 | Macrophage |
| Acaa1a | 0.42 | 0.00 | 0.00 | 0.29 | 0.16 | 12 | Macrophage |
| Ifngr1 | 0.43 | 0.00 | 0.00 | 0.38 | 0.24 | 12 | Macrophage |
| Tmem256 | 0.38 | 0.00 | 0.00 | 0.48 | 0.33 | 12 | Macrophage |
| Slc6a6 | 0.29 | 0.00 | 0.00 | 0.41 | 0.25 | 12 | Macrophage |
| Cdk2ap2 | 0.40 | 0.00 | 0.00 | 0.28 | 0.15 | 12 | Macrophage |
| Cst3 | 0.45 | 0.00 | 0.00 | 0.68 | 0.59 | 12 | Macrophage |
| Vamp8 | 0.32 | 0.00 | 0.00 | 0.51 | 0.34 | 12 | Macrophage |
| 2810474O19Rik | 0.38 | 0.00 | 0.00 | 0.41 | 0.26 | 12 | Macrophage |
| Sh3bgrl | 0.28 | 0.00 | 0.00 | 0.62 | 0.44 | 12 | Macrophage |
| Gapdh | 0.41 | 0.00 | 0.00 | 0.26 | 0.14 | 12 | Macrophage |
| Nsa2 | 0.36 | 0.00 | 0.00 | 0.45 | 0.30 | 12 | Macrophage |
| Ran | 0.39 | 0.00 | 0.00 | 0.54 | 0.38 | 12 | Macrophage |
| Tbca | 0.44 | 0.00 | 0.00 | 0.47 | 0.34 | 12 | Macrophage |
| Arpp19 | 0.41 | 0.00 | 0.00 | 0.45 | 0.31 | 12 | Macrophage |
| Actg1 | 0.26 | 0.00 | 0.00 | 0.85 | 0.76 | 12 | Macrophage |
| Esd | 0.46 | 0.00 | 0.00 | 0.33 | 0.20 | 12 | Macrophage |
| Gm10126 | 0.39 | 0.00 | 0.00 | 0.53 | 0.40 | 12 | Macrophage |
| Chmp3 | 0.37 | 0.00 | 0.00 | 0.34 | 0.21 | 12 | Macrophage |
| Ostf1 | 0.33 | 0.00 | 0.00 | 0.40 | 0.25 | 12 | Macrophage |
| Gm20091 | 0.36 | 0.00 | 0.00 | 0.35 | 0.22 | 12 | Macrophage |
| Tspo | 0.42 | 0.00 | 0.00 | 0.38 | 0.25 | 12 | Macrophage |
| Hmgn2 | 0.33 | 0.00 | 0.00 | 0.58 | 0.43 | 12 | Macrophage |
| Rbm3 | 0.37 | 0.00 | 0.00 | 0.54 | 0.41 | 12 | Macrophage |
| Dnajc2 | 0.39 | 0.00 | 0.00 | 0.33 | 0.20 | 12 | Macrophage |
| Irf2bp2 | 0.41 | 0.00 | 0.00 | 0.27 | 0.16 | 12 | Macrophage |
| Cox7b | 0.38 | 0.00 | 0.00 | 0.59 | 0.46 | 12 | Macrophage |
| Sec11c | 0.30 | 0.00 | 0.00 | 0.30 | 0.18 | 12 | Macrophage |
| Hspe1 | 0.39 | 0.00 | 0.00 | 0.42 | 0.28 | 12 | Macrophage |
| Dazap2 | 0.36 | 0.00 | 0.00 | 0.33 | 0.21 | 12 | Macrophage |
| Ifi203 | 0.28 | 0.00 | 0.00 | 0.26 | 0.14 | 12 | Macrophage |
| Timm10b | 0.35 | 0.00 | 0.00 | 0.29 | 0.17 | 12 | Macrophage |
| Gm22761 | 0.37 | 0.00 | 0.00 | 0.29 | 0.17 | 12 | Macrophage |
| Srp9 | 0.38 | 0.00 | 0.00 | 0.42 | 0.28 | 12 | Macrophage |
| Uqcrq | 0.36 | 0.00 | 0.00 | 0.47 | 0.33 | 12 | Macrophage |
| Stx7 | 0.34 | 0.00 | 0.00 | 0.26 | 0.15 | 12 | Macrophage |
| Jhdm1d | 0.35 | 0.00 | 0.00 | 0.27 | 0.16 | 12 | Macrophage |
| Ldha | 0.36 | 0.00 | 0.00 | 0.42 | 0.29 | 12 | Macrophage |
| Flna | 0.34 | 0.00 | 0.00 | 0.36 | 0.23 | 12 | Macrophage |
| Klf6 | 0.30 | 0.00 | 0.00 | 0.51 | 0.36 | 12 | Macrophage |
| Tor1aip1 | 0.29 | 0.00 | 0.00 | 0.31 | 0.19 | 12 | Macrophage |
| Gm10282 | 0.34 | 0.00 | 0.00 | 0.54 | 0.40 | 12 | Macrophage |
| Scand1 | 0.30 | 0.00 | 0.00 | 0.27 | 0.16 | 12 | Macrophage |
| Polr2l | 0.38 | 0.00 | 0.00 | 0.27 | 0.16 | 12 | Macrophage |
| Eif3j1 | 0.40 | 0.00 | 0.00 | 0.42 | 0.29 | 12 | Macrophage |
| Cox6b1 | 0.27 | 0.00 | 0.00 | 0.68 | 0.57 | 12 | Macrophage |
| Gdi2 | 0.36 | 0.00 | 0.00 | 0.46 | 0.33 | 12 | Macrophage |
| Atp6v1d | 0.35 | 0.00 | 0.00 | 0.28 | 0.17 | 12 | Macrophage |
| Snrpe | 0.32 | 0.00 | 0.00 | 0.49 | 0.36 | 12 | Macrophage |
| Vasp | 0.33 | 0.00 | 0.00 | 0.31 | 0.19 | 12 | Macrophage |
| Cltc | 0.37 | 0.00 | 0.00 | 0.30 | 0.18 | 12 | Macrophage |
| Eif3j2 | 0.40 | 0.00 | 0.00 | 0.44 | 0.32 | 12 | Macrophage |
| Tpr | 0.29 | 0.00 | 0.00 | 0.63 | 0.49 | 12 | Macrophage |
| Atg3 | 0.30 | 0.00 | 0.00 | 0.30 | 0.18 | 12 | Macrophage |
| Ddx46 | 0.33 | 0.00 | 0.00 | 0.42 | 0.29 | 12 | Macrophage |
| Cdc5l | 0.38 | 0.00 | 0.00 | 0.34 | 0.22 | 12 | Macrophage |
| 0610009D07Rik | 0.30 | 0.00 | 0.00 | 0.34 | 0.22 | 12 | Macrophage |
| Celf2 | 0.26 | 0.00 | 0.00 | 0.41 | 0.28 | 12 | Macrophage |
| Prpf40a | 0.32 | 0.00 | 0.00 | 0.45 | 0.32 | 12 | Macrophage |
| Psme1 | 0.35 | 0.00 | 0.00 | 0.43 | 0.31 | 12 | Macrophage |
| Sh3glb1 | 0.30 | 0.00 | 0.00 | 0.53 | 0.40 | 12 | Macrophage |
| Calm3 | 0.33 | 0.00 | 0.00 | 0.30 | 0.19 | 12 | Macrophage |
| Anxa5 | 0.37 | 0.00 | 0.00 | 0.52 | 0.40 | 12 | Macrophage |
| Hspa9 | 0.41 | 0.00 | 0.00 | 0.26 | 0.16 | 12 | Macrophage |
| Hnrnpm | 0.30 | 0.00 | 0.00 | 0.62 | 0.49 | 12 | Macrophage |
| Stk38 | 0.28 | 0.00 | 0.00 | 0.31 | 0.20 | 12 | Macrophage |
| Romo1 | 0.33 | 0.00 | 0.00 | 0.42 | 0.30 | 12 | Macrophage |
| Ywhah | 0.32 | 0.00 | 0.00 | 0.38 | 0.26 | 12 | Macrophage |
| Grb2 | 0.26 | 0.00 | 0.00 | 0.29 | 0.18 | 12 | Macrophage |
| Ndufa2 | 0.30 | 0.00 | 0.00 | 0.55 | 0.43 | 12 | Macrophage |
| Hspd1 | 0.38 | 0.00 | 0.00 | 0.37 | 0.25 | 12 | Macrophage |
| Atox1 | 0.30 | 0.00 | 0.00 | 0.48 | 0.36 | 12 | Macrophage |
| Canx | 0.32 | 0.00 | 0.00 | 0.60 | 0.49 | 12 | Macrophage |
| Cox7c | 0.25 | 0.00 | 0.00 | 0.73 | 0.63 | 12 | Macrophage |
| Snrpb2 | 0.29 | 0.00 | 0.00 | 0.36 | 0.24 | 12 | Macrophage |
| Cast | 0.31 | 0.00 | 0.00 | 0.30 | 0.19 | 12 | Macrophage |
| Atp5g3 | 0.33 | 0.00 | 0.00 | 0.48 | 0.36 | 12 | Macrophage |
| Tbcb | 0.28 | 0.00 | 0.00 | 0.35 | 0.23 | 12 | Macrophage |
| Gm20594 | 0.30 | 0.00 | 0.00 | 0.27 | 0.17 | 12 | Macrophage |
| Atp6v1f | 0.30 | 0.00 | 0.00 | 0.35 | 0.24 | 12 | Macrophage |
| F11r | 0.29 | 0.00 | 0.00 | 0.26 | 0.16 | 12 | Macrophage |
| Hspa4 | 0.34 | 0.00 | 0.00 | 0.34 | 0.23 | 12 | Macrophage |
| Atp6v0d1 | 0.31 | 0.00 | 0.00 | 0.27 | 0.18 | 12 | Macrophage |
| Nars | 0.29 | 0.00 | 0.00 | 0.44 | 0.32 | 12 | Macrophage |
| Arf5 | 0.29 | 0.00 | 0.00 | 0.38 | 0.28 | 12 | Macrophage |
| Rtn3 | 0.26 | 0.00 | 0.00 | 0.36 | 0.25 | 12 | Macrophage |
| Ndufa1 | 0.28 | 0.00 | 0.00 | 0.42 | 0.31 | 12 | Macrophage |
| Wdr1 | 0.27 | 0.00 | 0.00 | 0.26 | 0.17 | 12 | Macrophage |
| Utp3 | 0.28 | 0.00 | 0.00 | 0.32 | 0.22 | 12 | Macrophage |
| H2afy | 0.27 | 0.00 | 0.00 | 0.29 | 0.20 | 12 | Macrophage |
| Adipor1 | 0.28 | 0.00 | 0.00 | 0.34 | 0.23 | 12 | Macrophage |
| Jtb | 0.29 | 0.00 | 0.00 | 0.28 | 0.18 | 12 | Macrophage |
| Nol7 | 0.25 | 0.00 | 0.00 | 0.47 | 0.35 | 12 | Macrophage |
| Mtdh | 0.30 | 0.00 | 0.00 | 0.59 | 0.48 | 12 | Macrophage |
| Chmp2a | 0.27 | 0.00 | 0.00 | 0.42 | 0.31 | 12 | Macrophage |
| Psmb5 | 0.25 | 0.00 | 0.00 | 0.33 | 0.22 | 12 | Macrophage |
| Atp6v1a | 0.29 | 0.00 | 0.00 | 0.38 | 0.28 | 12 | Macrophage |
| Mtpn | 0.29 | 0.00 | 0.00 | 0.37 | 0.27 | 12 | Macrophage |
| Rap1b | 0.27 | 0.00 | 0.00 | 0.33 | 0.23 | 12 | Macrophage |
| S100a11 | 0.30 | 0.00 | 0.00 | 0.58 | 0.49 | 12 | Macrophage |
| Snrpd1 | 0.27 | 0.00 | 0.00 | 0.29 | 0.20 | 12 | Macrophage |
| Timm13 | 0.27 | 0.00 | 0.00 | 0.32 | 0.23 | 12 | Macrophage |
| Eea1 | 0.29 | 0.00 | 0.00 | 0.30 | 0.21 | 12 | Macrophage |
| Sdhd | 0.26 | 0.00 | 0.00 | 0.26 | 0.17 | 12 | Macrophage |
| Jund | 0.31 | 0.00 | 0.00 | 0.36 | 0.28 | 12 | Macrophage |
| Smarca5 | 0.27 | 0.00 | 0.00 | 0.40 | 0.30 | 12 | Macrophage |
| Zc3h13 | 0.27 | 0.00 | 0.00 | 0.42 | 0.33 | 12 | Macrophage |
| Btg2 | 0.30 | 0.00 | 0.00 | 0.31 | 0.23 | 12 | Macrophage |
| B020018G12Rik | 0.25 | 0.00 | 0.00 | 0.34 | 0.26 | 12 | Macrophage |
| Mif | 0.25 | 0.00 | 0.00 | 0.27 | 0.19 | 12 | Macrophage |
| Pcna-ps2 | 0.29 | 0.00 | 0.00 | 0.25 | 0.18 | 12 | Macrophage |
| Ghitm | 0.29 | 0.00 | 0.00 | 0.29 | 0.22 | 12 | Macrophage |
| Acta2 | 2.39 | 0.00 | 0.00 | 0.66 | 0.18 | 13 | Matrix Fibroblast |
| Dcn | 2.38 | 0.00 | 0.00 | 0.36 | 0.01 | 13 | Matrix Fibroblast |
| Col1a1 | 2.30 | 0.00 | 0.00 | 0.79 | 0.24 | 13 | Matrix Fibroblast |
| Col3a1 | 2.25 | 0.00 | 0.00 | 0.81 | 0.31 | 13 | Matrix Fibroblast |
| Igfbp5 | 2.25 | 0.00 | 0.00 | 0.56 | 0.09 | 13 | Matrix Fibroblast |
| Col1a2 | 2.17 | 0.00 | 0.00 | 0.98 | 0.38 | 13 | Matrix Fibroblast |
| Myl9 | 1.83 | 0.00 | 0.00 | 0.55 | 0.11 | 13 | Matrix Fibroblast |
| Tpm2 | 1.82 | 0.00 | 0.00 | 0.58 | 0.12 | 13 | Matrix Fibroblast |
| Col14a1 | 1.81 | 0.00 | 0.00 | 0.50 | 0.04 | 13 | Matrix Fibroblast |
| Actc1 | 1.80 | 0.00 | 0.00 | 0.26 | 0.02 | 13 | Matrix Fibroblast |
| Tagln | 1.75 | 0.00 | 0.00 | 0.55 | 0.14 | 13 | Matrix Fibroblast |
| Lmod1 | 1.72 | 0.00 | 0.00 | 0.38 | 0.03 | 13 | Matrix Fibroblast |
| Mfap5 | 1.71 | 0.00 | 0.00 | 0.41 | 0.05 | 13 | Matrix Fibroblast |
| Mgp | 1.67 | 0.00 | 0.00 | 0.88 | 0.30 | 13 | Matrix Fibroblast |
| Postn | 1.67 | 0.00 | 0.00 | 0.51 | 0.07 | 13 | Matrix Fibroblast |
| Pi15 | 1.60 | 0.00 | 0.00 | 0.34 | 0.02 | 13 | Matrix Fibroblast |
| Gpc6 | 1.47 | 0.00 | 0.00 | 0.35 | 0.01 | 13 | Matrix Fibroblast |
| Fstl1 | 1.39 | 0.00 | 0.00 | 0.94 | 0.43 | 13 | Matrix Fibroblast |
| Eln | 1.38 | 0.00 | 0.00 | 0.62 | 0.14 | 13 | Matrix Fibroblast |
| Ccdc80 | 1.36 | 0.00 | 0.00 | 0.51 | 0.10 | 13 | Matrix Fibroblast |
| Ogn | 1.29 | 0.00 | 0.00 | 0.54 | 0.12 | 13 | Matrix Fibroblast |
| Sparc | 1.28 | 0.00 | 0.00 | 0.99 | 0.73 | 13 | Matrix Fibroblast |
| Igfbp7 | 1.26 | 0.00 | 0.00 | 0.90 | 0.38 | 13 | Matrix Fibroblast |
| Lgals1 | 1.24 | 0.00 | 0.00 | 0.93 | 0.40 | 13 | Matrix Fibroblast |
| Lox | 1.21 | 0.00 | 0.00 | 0.54 | 0.12 | 13 | Matrix Fibroblast |
| Adamts2 | 1.18 | 0.00 | 0.00 | 0.50 | 0.09 | 13 | Matrix Fibroblast |
| Rcn3 | 1.17 | 0.00 | 0.00 | 0.54 | 0.12 | 13 | Matrix Fibroblast |
| Osr1 | 1.14 | 0.00 | 0.00 | 0.31 | 0.02 | 13 | Matrix Fibroblast |
| Loxl1 | 1.06 | 0.00 | 0.00 | 0.48 | 0.10 | 13 | Matrix Fibroblast |
| Prrx1 | 0.94 | 0.00 | 0.00 | 0.27 | 0.02 | 13 | Matrix Fibroblast |
| Gpc3 | 1.66 | 0.00 | 0.00 | 0.55 | 0.15 | 13 | Matrix Fibroblast |
| Myh11 | 1.94 | 0.00 | 0.00 | 0.43 | 0.10 | 13 | Matrix Fibroblast |
| Aspn | 0.63 | 0.00 | 0.00 | 0.43 | 0.09 | 13 | Matrix Fibroblast |
| 0610007N19Rik | 1.01 | 0.00 | 0.00 | 0.52 | 0.13 | 13 | Matrix Fibroblast |
| Plac9b | 0.99 | 0.00 | 0.00 | 0.78 | 0.29 | 13 | Matrix Fibroblast |
| Aebp1 | 1.19 | 0.00 | 0.00 | 0.36 | 0.07 | 13 | Matrix Fibroblast |
| Rarres2 | 1.26 | 0.00 | 0.00 | 0.48 | 0.12 | 13 | Matrix Fibroblast |
| Plac9a | 0.98 | 0.00 | 0.00 | 0.77 | 0.29 | 13 | Matrix Fibroblast |
| Col5a1 | 0.98 | 0.00 | 0.00 | 0.39 | 0.08 | 13 | Matrix Fibroblast |
| Cygb | 1.02 | 0.00 | 0.00 | 0.30 | 0.05 | 13 | Matrix Fibroblast |
| Fbn1 | 1.01 | 0.00 | 0.00 | 0.55 | 0.16 | 13 | Matrix Fibroblast |
| Ndrg2 | 0.91 | 0.00 | 0.00 | 0.35 | 0.07 | 13 | Matrix Fibroblast |
| Bgn | 1.04 | 0.00 | 0.00 | 0.65 | 0.23 | 13 | Matrix Fibroblast |
| Actg2 | 1.38 | 0.00 | 0.00 | 0.33 | 0.07 | 13 | Matrix Fibroblast |
| Cald1 | 1.26 | 0.00 | 0.00 | 0.81 | 0.40 | 13 | Matrix Fibroblast |
| Selm | 0.86 | 0.00 | 0.00 | 0.36 | 0.08 | 13 | Matrix Fibroblast |
| Pcolce | 0.98 | 0.00 | 0.00 | 0.36 | 0.08 | 13 | Matrix Fibroblast |
| Itga8 | 1.22 | 0.00 | 0.00 | 0.37 | 0.08 | 13 | Matrix Fibroblast |
| Fam129a | 0.97 | 0.00 | 0.00 | 0.35 | 0.08 | 13 | Matrix Fibroblast |
| Ptgis | 0.86 | 0.00 | 0.00 | 0.27 | 0.05 | 13 | Matrix Fibroblast |
| Igf2 | 1.26 | 0.00 | 0.00 | 0.31 | 0.07 | 13 | Matrix Fibroblast |
| Serping1 | 0.91 | 0.00 | 0.00 | 0.43 | 0.11 | 13 | Matrix Fibroblast |
| Gucy1a3 | 0.70 | 0.00 | 0.00 | 0.38 | 0.09 | 13 | Matrix Fibroblast |
| Fxyd1 | 0.82 | 0.00 | 0.00 | 0.36 | 0.08 | 13 | Matrix Fibroblast |
| Tpm1 | 1.29 | 0.00 | 0.00 | 0.84 | 0.51 | 13 | Matrix Fibroblast |
| Gsn | 1.03 | 0.00 | 0.00 | 0.61 | 0.24 | 13 | Matrix Fibroblast |
| Prkcdbp | 0.89 | 0.00 | 0.00 | 0.39 | 0.11 | 13 | Matrix Fibroblast |
| Ltbp4 | 0.80 | 0.00 | 0.00 | 0.38 | 0.10 | 13 | Matrix Fibroblast |
| Cxcl12 | 0.47 | 0.00 | 0.00 | 0.41 | 0.11 | 13 | Matrix Fibroblast |
| Serpinh1 | 0.82 | 0.00 | 0.00 | 0.75 | 0.38 | 13 | Matrix Fibroblast |
| Map1b | 0.97 | 0.00 | 0.00 | 0.35 | 0.10 | 13 | Matrix Fibroblast |
| Meg3 | 0.85 | 0.00 | 0.00 | 0.56 | 0.21 | 13 | Matrix Fibroblast |
| Fblim1 | 1.02 | 0.00 | 0.00 | 0.44 | 0.16 | 13 | Matrix Fibroblast |
| Col6a2 | 0.83 | 0.00 | 0.00 | 0.42 | 0.14 | 13 | Matrix Fibroblast |
| Col6a1 | 0.86 | 0.00 | 0.00 | 0.49 | 0.18 | 13 | Matrix Fibroblast |
| Col5a2 | 0.87 | 0.00 | 0.00 | 0.47 | 0.17 | 13 | Matrix Fibroblast |
| Cst3 | 0.64 | 0.00 | 0.00 | 0.85 | 0.58 | 13 | Matrix Fibroblast |
| Gata6 | 0.73 | 0.00 | 0.00 | 0.30 | 0.08 | 13 | Matrix Fibroblast |
| Timp2 | 0.79 | 0.00 | 0.00 | 0.50 | 0.20 | 13 | Matrix Fibroblast |
| Il11ra1 | 0.71 | 0.00 | 0.00 | 0.27 | 0.07 | 13 | Matrix Fibroblast |
| Csrp2 | 0.99 | 0.00 | 0.00 | 0.42 | 0.16 | 13 | Matrix Fibroblast |
| Maged2 | 0.69 | 0.00 | 0.00 | 0.44 | 0.17 | 13 | Matrix Fibroblast |
| Cd63 | 0.72 | 0.00 | 0.00 | 0.59 | 0.29 | 13 | Matrix Fibroblast |
| Vkorc1 | 0.62 | 0.00 | 0.00 | 0.31 | 0.10 | 13 | Matrix Fibroblast |
| Prss35 | 0.56 | 0.00 | 0.00 | 0.26 | 0.07 | 13 | Matrix Fibroblast |
| Cd81 | 0.63 | 0.00 | 0.00 | 0.72 | 0.40 | 13 | Matrix Fibroblast |
| Myh10 | 0.66 | 0.00 | 0.00 | 0.52 | 0.22 | 13 | Matrix Fibroblast |
| Mustn1 | 0.94 | 0.00 | 0.00 | 0.26 | 0.07 | 13 | Matrix Fibroblast |
| Akap12 | 1.13 | 0.00 | 0.00 | 0.50 | 0.23 | 13 | Matrix Fibroblast |
| Leprel2 | 0.59 | 0.00 | 0.00 | 0.25 | 0.07 | 13 | Matrix Fibroblast |
| Pls3 | 0.78 | 0.00 | 0.00 | 0.45 | 0.20 | 13 | Matrix Fibroblast |
| Crip1 | 1.04 | 0.00 | 0.00 | 0.72 | 0.47 | 13 | Matrix Fibroblast |
| Lhfp | 0.71 | 0.00 | 0.00 | 0.39 | 0.15 | 13 | Matrix Fibroblast |
| Fn1 | 0.52 | 0.00 | 0.00 | 0.53 | 0.23 | 13 | Matrix Fibroblast |
| Nexn | 0.58 | 0.00 | 0.00 | 0.50 | 0.21 | 13 | Matrix Fibroblast |
| Slc25a4 | 0.70 | 0.00 | 0.00 | 0.74 | 0.50 | 13 | Matrix Fibroblast |
| Fam114a1 | 0.60 | 0.00 | 0.00 | 0.25 | 0.08 | 13 | Matrix Fibroblast |
| Col6a3 | 0.62 | 0.00 | 0.00 | 0.29 | 0.09 | 13 | Matrix Fibroblast |
| Flna | 0.81 | 0.00 | 0.00 | 0.48 | 0.23 | 13 | Matrix Fibroblast |
| Myl6 | 0.74 | 0.00 | 0.00 | 0.86 | 0.69 | 13 | Matrix Fibroblast |
| Dnajb4 | 0.75 | 0.00 | 0.00 | 0.29 | 0.10 | 13 | Matrix Fibroblast |
| Antxr1 | 0.64 | 0.00 | 0.00 | 0.32 | 0.11 | 13 | Matrix Fibroblast |
| Fbln5 | 0.60 | 0.00 | 0.00 | 0.41 | 0.17 | 13 | Matrix Fibroblast |
| Gpx8 | 0.64 | 0.00 | 0.00 | 0.36 | 0.14 | 13 | Matrix Fibroblast |
| Nfia | 0.70 | 0.00 | 0.00 | 0.48 | 0.23 | 13 | Matrix Fibroblast |
| Mmp14 | 0.68 | 0.00 | 0.00 | 0.50 | 0.25 | 13 | Matrix Fibroblast |
| S100a11 | 0.58 | 0.00 | 0.00 | 0.73 | 0.48 | 13 | Matrix Fibroblast |
| Loxl2 | 0.41 | 0.00 | 0.00 | 0.36 | 0.13 | 13 | Matrix Fibroblast |
| Plxdc2 | 0.52 | 0.00 | 0.00 | 0.33 | 0.12 | 13 | Matrix Fibroblast |
| Atp1a2 | 0.54 | 0.00 | 0.00 | 0.29 | 0.10 | 13 | Matrix Fibroblast |
| S100a6 | 0.37 | 0.00 | 0.00 | 0.61 | 0.31 | 13 | Matrix Fibroblast |
| Ilk | 0.62 | 0.00 | 0.00 | 0.47 | 0.22 | 13 | Matrix Fibroblast |
| Rbp1 | 0.47 | 0.00 | 0.00 | 0.44 | 0.19 | 13 | Matrix Fibroblast |
| Cryab | 0.59 | 0.00 | 0.00 | 0.35 | 0.14 | 13 | Matrix Fibroblast |
| Fhl1 | 0.30 | 0.00 | 0.00 | 0.59 | 0.29 | 13 | Matrix Fibroblast |
| Rrbp1 | 0.59 | 0.00 | 0.00 | 0.75 | 0.54 | 13 | Matrix Fibroblast |
| Pam | 0.60 | 0.00 | 0.00 | 0.39 | 0.17 | 13 | Matrix Fibroblast |
| Dstn | 0.81 | 0.00 | 0.00 | 0.71 | 0.50 | 13 | Matrix Fibroblast |
| Vim | 0.49 | 0.00 | 0.00 | 0.87 | 0.66 | 13 | Matrix Fibroblast |
| Mylk | 0.81 | 0.00 | 0.00 | 0.51 | 0.27 | 13 | Matrix Fibroblast |
| Cdh11 | 0.46 | 0.00 | 0.00 | 0.42 | 0.18 | 13 | Matrix Fibroblast |
| Nfib | 0.55 | 0.00 | 0.00 | 0.71 | 0.46 | 13 | Matrix Fibroblast |
| Igfbp4 | 0.71 | 0.00 | 0.00 | 0.47 | 0.25 | 13 | Matrix Fibroblast |
| Ckap4 | 0.57 | 0.00 | 0.00 | 0.38 | 0.17 | 13 | Matrix Fibroblast |
| Fkbp9 | 0.50 | 0.00 | 0.00 | 0.32 | 0.13 | 13 | Matrix Fibroblast |
| Cav1 | 0.43 | 0.00 | 0.00 | 0.59 | 0.31 | 13 | Matrix Fibroblast |
| Lpp | 0.79 | 0.00 | 0.00 | 0.42 | 0.21 | 13 | Matrix Fibroblast |
| Rbms3 | 0.52 | 0.00 | 0.00 | 0.37 | 0.16 | 13 | Matrix Fibroblast |
| Tgfb2 | 0.63 | 0.00 | 0.00 | 0.32 | 0.14 | 13 | Matrix Fibroblast |
| Fkbp7 | 0.52 | 0.00 | 0.00 | 0.31 | 0.13 | 13 | Matrix Fibroblast |
| Ptrf | 0.54 | 0.00 | 0.00 | 0.60 | 0.35 | 13 | Matrix Fibroblast |
| Vcan | 0.55 | 0.00 | 0.00 | 0.27 | 0.11 | 13 | Matrix Fibroblast |
| Gpx7 | 0.46 | 0.00 | 0.00 | 0.25 | 0.09 | 13 | Matrix Fibroblast |
| Tmed3 | 0.54 | 0.00 | 0.00 | 0.29 | 0.12 | 13 | Matrix Fibroblast |
| Mdk | 0.43 | 0.00 | 0.00 | 0.40 | 0.18 | 13 | Matrix Fibroblast |
| Pmepa1 | 0.51 | 0.00 | 0.00 | 0.27 | 0.11 | 13 | Matrix Fibroblast |
| Ctsl | 0.46 | 0.00 | 0.00 | 0.62 | 0.38 | 13 | Matrix Fibroblast |
| Lmna | 0.54 | 0.00 | 0.00 | 0.40 | 0.20 | 13 | Matrix Fibroblast |
| Nupr1 | 0.54 | 0.00 | 0.00 | 0.38 | 0.18 | 13 | Matrix Fibroblast |
| Dpt | 0.74 | 0.00 | 0.00 | 0.36 | 0.17 | 13 | Matrix Fibroblast |
| Sdc2 | 0.39 | 0.00 | 0.00 | 0.29 | 0.12 | 13 | Matrix Fibroblast |
| Fgfr1 | 0.42 | 0.00 | 0.00 | 0.26 | 0.10 | 13 | Matrix Fibroblast |
| Pdlim7 | 0.49 | 0.00 | 0.00 | 0.26 | 0.10 | 13 | Matrix Fibroblast |
| Hcfc1r1 | 0.59 | 0.00 | 0.00 | 0.31 | 0.14 | 13 | Matrix Fibroblast |
| Pbx1 | 0.52 | 0.00 | 0.00 | 0.39 | 0.19 | 13 | Matrix Fibroblast |
| Nid1 | 0.50 | 0.00 | 0.00 | 0.43 | 0.23 | 13 | Matrix Fibroblast |
| Anxa5 | 0.45 | 0.00 | 0.00 | 0.61 | 0.40 | 13 | Matrix Fibroblast |
| Rcn1 | 0.51 | 0.00 | 0.00 | 0.31 | 0.14 | 13 | Matrix Fibroblast |
| Dkk3 | 0.41 | 0.00 | 0.00 | 0.28 | 0.11 | 13 | Matrix Fibroblast |
| Nenf | 0.49 | 0.00 | 0.00 | 0.37 | 0.18 | 13 | Matrix Fibroblast |
| Eid1 | 0.51 | 0.00 | 0.00 | 0.50 | 0.28 | 13 | Matrix Fibroblast |
| Mfap2 | 0.51 | 0.00 | 0.00 | 0.57 | 0.34 | 13 | Matrix Fibroblast |
| Calu | 0.51 | 0.00 | 0.00 | 0.44 | 0.24 | 13 | Matrix Fibroblast |
| Cbx6 | 0.45 | 0.00 | 0.00 | 0.25 | 0.10 | 13 | Matrix Fibroblast |
| Slmap | 0.60 | 0.00 | 0.00 | 0.32 | 0.15 | 13 | Matrix Fibroblast |
| Ppic | 0.45 | 0.00 | 0.00 | 0.60 | 0.38 | 13 | Matrix Fibroblast |
| Cdc42ep3 | 0.63 | 0.00 | 0.00 | 0.30 | 0.14 | 13 | Matrix Fibroblast |
| Nr2f2 | 0.48 | 0.00 | 0.00 | 0.40 | 0.21 | 13 | Matrix Fibroblast |
| Actn1 | 0.66 | 0.00 | 0.00 | 0.40 | 0.22 | 13 | Matrix Fibroblast |
| Nedd4 | 0.39 | 0.00 | 0.00 | 0.74 | 0.53 | 13 | Matrix Fibroblast |
| Daam1 | 0.62 | 0.00 | 0.00 | 0.31 | 0.15 | 13 | Matrix Fibroblast |
| Golim4 | 0.45 | 0.00 | 0.00 | 0.38 | 0.20 | 13 | Matrix Fibroblast |
| Tm4sf1 | 0.31 | 0.00 | 0.00 | 0.41 | 0.21 | 13 | Matrix Fibroblast |
| Peg3 | 0.52 | 0.00 | 0.00 | 0.49 | 0.30 | 13 | Matrix Fibroblast |
| Thra | 0.42 | 0.00 | 0.00 | 0.34 | 0.17 | 13 | Matrix Fibroblast |
| Gnas | 0.36 | 0.00 | 0.00 | 0.82 | 0.67 | 13 | Matrix Fibroblast |
| Maged1 | 0.44 | 0.00 | 0.00 | 0.35 | 0.18 | 13 | Matrix Fibroblast |
| Sparcl1 | 0.42 | 0.00 | 0.00 | 0.59 | 0.39 | 13 | Matrix Fibroblast |
| Twsg1 | 0.41 | 0.00 | 0.00 | 0.25 | 0.12 | 13 | Matrix Fibroblast |
| Serpinb6a | 0.42 | 0.00 | 0.00 | 0.34 | 0.18 | 13 | Matrix Fibroblast |
| Svil | 0.45 | 0.00 | 0.00 | 0.32 | 0.17 | 13 | Matrix Fibroblast |
| P4hb | 0.41 | 0.00 | 0.00 | 0.57 | 0.40 | 13 | Matrix Fibroblast |
| Laptm4a | 0.38 | 0.00 | 0.00 | 0.69 | 0.51 | 13 | Matrix Fibroblast |
| Tspan3 | 0.41 | 0.00 | 0.00 | 0.30 | 0.15 | 13 | Matrix Fibroblast |
| Arf4 | 0.44 | 0.00 | 0.00 | 0.43 | 0.27 | 13 | Matrix Fibroblast |
| Rhoj | 0.37 | 0.00 | 0.00 | 0.34 | 0.18 | 13 | Matrix Fibroblast |
| Filip1l | 0.45 | 0.00 | 0.00 | 0.50 | 0.33 | 13 | Matrix Fibroblast |
| Hdlbp | 0.40 | 0.00 | 0.00 | 0.36 | 0.21 | 13 | Matrix Fibroblast |
| Dnajc10 | 0.41 | 0.00 | 0.00 | 0.28 | 0.15 | 13 | Matrix Fibroblast |
| Rps2 | 0.34 | 0.00 | 0.00 | 0.78 | 0.67 | 13 | Matrix Fibroblast |
| S100a10 | 0.43 | 0.00 | 0.00 | 0.54 | 0.38 | 13 | Matrix Fibroblast |
| Sepw1 | 0.40 | 0.00 | 0.00 | 0.34 | 0.19 | 13 | Matrix Fibroblast |
| Ppap2b | 0.45 | 0.00 | 0.00 | 0.39 | 0.23 | 13 | Matrix Fibroblast |
| Sh3bgrl | 0.29 | 0.00 | 0.00 | 0.63 | 0.44 | 13 | Matrix Fibroblast |
| Rbpms | 0.53 | 0.00 | 0.00 | 0.32 | 0.19 | 13 | Matrix Fibroblast |
| Emp3 | 0.35 | 0.00 | 0.00 | 0.36 | 0.21 | 13 | Matrix Fibroblast |
| Arl1 | 0.40 | 0.00 | 0.00 | 0.35 | 0.20 | 13 | Matrix Fibroblast |
| Nfix | 0.43 | 0.00 | 0.00 | 0.30 | 0.17 | 13 | Matrix Fibroblast |
| Kdelr2 | 0.41 | 0.00 | 0.00 | 0.32 | 0.19 | 13 | Matrix Fibroblast |
| Myadm | 0.38 | 0.00 | 0.00 | 0.34 | 0.20 | 13 | Matrix Fibroblast |
| Rnase4 | 0.39 | 0.00 | 0.00 | 0.32 | 0.18 | 13 | Matrix Fibroblast |
| Celf2 | 0.31 | 0.00 | 0.00 | 0.45 | 0.28 | 13 | Matrix Fibroblast |
| Tuba1a | 0.28 | 0.00 | 0.00 | 0.68 | 0.50 | 13 | Matrix Fibroblast |
| Zfhx3 | 0.44 | 0.00 | 0.00 | 0.30 | 0.17 | 13 | Matrix Fibroblast |
| 7-Sep | 0.29 | 0.00 | 0.00 | 0.73 | 0.58 | 13 | Matrix Fibroblast |
| Serf2 | 0.32 | 0.00 | 0.00 | 0.69 | 0.56 | 13 | Matrix Fibroblast |
| Grb10 | 0.32 | 0.00 | 0.00 | 0.29 | 0.16 | 13 | Matrix Fibroblast |
| Serpine2 | 0.25 | 0.00 | 0.00 | 0.41 | 0.25 | 13 | Matrix Fibroblast |
| Lamc1 | 0.32 | 0.00 | 0.00 | 0.28 | 0.16 | 13 | Matrix Fibroblast |
| Oxct1 | 0.44 | 0.00 | 0.00 | 0.26 | 0.15 | 13 | Matrix Fibroblast |
| Parva | 0.33 | 0.00 | 0.00 | 0.28 | 0.16 | 13 | Matrix Fibroblast |
| Ppp1cb | 0.42 | 0.00 | 0.00 | 0.32 | 0.20 | 13 | Matrix Fibroblast |
| Mxra8 | 0.28 | 0.00 | 0.00 | 0.29 | 0.16 | 13 | Matrix Fibroblast |
| Ppib | 0.33 | 0.00 | 0.00 | 0.57 | 0.43 | 13 | Matrix Fibroblast |
| Ppp1r12a | 0.54 | 0.00 | 0.00 | 0.37 | 0.25 | 13 | Matrix Fibroblast |
| Nfic | 0.31 | 0.00 | 0.00 | 0.38 | 0.24 | 13 | Matrix Fibroblast |
| Ckb | 0.34 | 0.00 | 0.00 | 0.25 | 0.14 | 13 | Matrix Fibroblast |
| Hmgn1 | 0.30 | 0.00 | 0.00 | 0.59 | 0.45 | 13 | Matrix Fibroblast |
| Gabarapl1 | 0.37 | 0.00 | 0.00 | 0.27 | 0.16 | 13 | Matrix Fibroblast |
| Mef2c | 0.32 | 0.00 | 0.00 | 0.28 | 0.16 | 13 | Matrix Fibroblast |
| Sec61g | 0.34 | 0.00 | 0.00 | 0.47 | 0.35 | 13 | Matrix Fibroblast |
| Lamb1 | 0.29 | 0.00 | 0.00 | 0.27 | 0.16 | 13 | Matrix Fibroblast |
| Tceal8 | 0.34 | 0.00 | 0.00 | 0.31 | 0.19 | 13 | Matrix Fibroblast |
| Hspa5 | 0.30 | 0.00 | 0.00 | 0.68 | 0.55 | 13 | Matrix Fibroblast |
| Cbx5 | 0.33 | 0.00 | 0.00 | 0.39 | 0.26 | 13 | Matrix Fibroblast |
| Rsu1 | 0.33 | 0.00 | 0.00 | 0.31 | 0.20 | 13 | Matrix Fibroblast |
| Ndufv3 | 0.35 | 0.00 | 0.00 | 0.35 | 0.24 | 13 | Matrix Fibroblast |
| Gm5786 | 0.32 | 0.00 | 0.00 | 0.51 | 0.39 | 13 | Matrix Fibroblast |
| Tbrg1 | 0.29 | 0.00 | 0.00 | 0.43 | 0.30 | 13 | Matrix Fibroblast |
| Sar1a | 0.29 | 0.00 | 0.00 | 0.31 | 0.20 | 13 | Matrix Fibroblast |
| Gstp1 | 0.27 | 0.00 | 0.00 | 0.31 | 0.19 | 13 | Matrix Fibroblast |
| Sfr1 | 0.29 | 0.00 | 0.00 | 0.46 | 0.33 | 13 | Matrix Fibroblast |
| Ssr4 | 0.31 | 0.00 | 0.00 | 0.36 | 0.24 | 13 | Matrix Fibroblast |
| Ttc3 | 0.27 | 0.00 | 0.00 | 0.42 | 0.29 | 13 | Matrix Fibroblast |
| Calr | 0.29 | 0.00 | 0.00 | 0.56 | 0.43 | 13 | Matrix Fibroblast |
| Cfl2 | 0.37 | 0.00 | 0.00 | 0.36 | 0.25 | 13 | Matrix Fibroblast |
| Idh2 | 0.31 | 0.00 | 0.00 | 0.30 | 0.19 | 13 | Matrix Fibroblast |
| Morf4l2 | 0.30 | 0.00 | 0.00 | 0.33 | 0.22 | 13 | Matrix Fibroblast |
| Rps2-ps10 | 0.31 | 0.00 | 0.00 | 0.54 | 0.43 | 13 | Matrix Fibroblast |
| Rps2-ps6 | 0.33 | 0.00 | 0.00 | 0.51 | 0.41 | 13 | Matrix Fibroblast |
| Hsp90b1 | 0.34 | 0.00 | 0.00 | 0.84 | 0.76 | 13 | Matrix Fibroblast |
| Ddost | 0.28 | 0.00 | 0.00 | 0.29 | 0.19 | 13 | Matrix Fibroblast |
| Erlec1 | 0.29 | 0.00 | 0.00 | 0.25 | 0.16 | 13 | Matrix Fibroblast |
| Gm6139 | 0.32 | 0.00 | 0.00 | 0.52 | 0.42 | 13 | Matrix Fibroblast |
| Mif | 0.29 | 0.00 | 0.00 | 0.29 | 0.19 | 13 | Matrix Fibroblast |
| Prdx4 | 0.27 | 0.00 | 0.00 | 0.26 | 0.16 | 13 | Matrix Fibroblast |
| Copb2 | 0.29 | 0.00 | 0.00 | 0.27 | 0.18 | 13 | Matrix Fibroblast |
| Mrfap1 | 0.28 | 0.00 | 0.00 | 0.44 | 0.33 | 13 | Matrix Fibroblast |
| Ostc | 0.26 | 0.00 | 0.00 | 0.33 | 0.23 | 13 | Matrix Fibroblast |
| Eprs | 0.29 | 0.00 | 0.00 | 0.32 | 0.22 | 13 | Matrix Fibroblast |
| Map1lc3a | 0.25 | 0.00 | 0.00 | 0.30 | 0.20 | 13 | Matrix Fibroblast |
| Htatsf1 | 0.25 | 0.00 | 0.00 | 0.37 | 0.26 | 13 | Matrix Fibroblast |
| 2810428I15Rik | 0.28 | 0.00 | 0.00 | 0.29 | 0.19 | 13 | Matrix Fibroblast |
| Zbtb20 | 0.32 | 0.00 | 0.00 | 0.37 | 0.27 | 13 | Matrix Fibroblast |
| Atp5g1 | 0.30 | 0.00 | 0.00 | 0.32 | 0.22 | 13 | Matrix Fibroblast |
| Cdkn1c | 0.27 | 0.00 | 0.00 | 0.27 | 0.18 | 13 | Matrix Fibroblast |
| Gm18025 | 0.26 | 0.00 | 0.00 | 0.28 | 0.20 | 13 | Matrix Fibroblast |
| Gm6576 | 0.27 | 0.00 | 0.00 | 0.40 | 0.31 | 13 | Matrix Fibroblast |
| Hist1h2ap | 2.56 | 0.00 | 0.00 | 0.79 | 0.13 | 14 | Endothelial |
| Hist1h2ao | 2.56 | 0.00 | 0.00 | 0.79 | 0.13 | 14 | Endothelial |
| Hist1h2ag | 2.41 | 0.00 | 0.00 | 0.72 | 0.10 | 14 | Endothelial |
| Hist1h2ac | 2.31 | 0.00 | 0.00 | 0.69 | 0.09 | 14 | Endothelial |
| Hist1h2ad | 2.29 | 0.00 | 0.00 | 0.69 | 0.09 | 14 | Endothelial |
| Hist1h2ai | 2.26 | 0.00 | 0.00 | 0.67 | 0.08 | 14 | Endothelial |
| Hist1h2ah | 2.23 | 0.00 | 0.00 | 0.66 | 0.08 | 14 | Endothelial |
| Hist1h2af | 2.20 | 0.00 | 0.00 | 0.66 | 0.07 | 14 | Endothelial |
| Hist1h2an | 2.19 | 0.00 | 0.00 | 0.64 | 0.07 | 14 | Endothelial |
| Hist1h2ae | 2.15 | 0.00 | 0.00 | 0.61 | 0.07 | 14 | Endothelial |
| Hist1h2ab | 2.11 | 0.00 | 0.00 | 0.62 | 0.08 | 14 | Endothelial |
| Cenpf | 2.07 | 0.00 | 0.00 | 0.68 | 0.06 | 14 | Endothelial |
| Top2a | 2.05 | 0.00 | 0.00 | 0.79 | 0.09 | 14 | Endothelial |
| Mki67 | 2.05 | 0.00 | 0.00 | 0.81 | 0.10 | 14 | Endothelial |
| Hist1h2ak | 1.97 | 0.00 | 0.00 | 0.58 | 0.06 | 14 | Endothelial |
| Prc1 | 1.80 | 0.00 | 0.00 | 0.59 | 0.05 | 14 | Endothelial |
| Cenpe | 1.76 | 0.00 | 0.00 | 0.62 | 0.06 | 14 | Endothelial |
| Hmgb2 | 1.76 | 0.00 | 0.00 | 0.96 | 0.41 | 14 | Endothelial |
| H2afx | 1.68 | 0.00 | 0.00 | 0.62 | 0.08 | 14 | Endothelial |
| Nusap1 | 1.67 | 0.00 | 0.00 | 0.57 | 0.04 | 14 | Endothelial |
| Smc2 | 1.66 | 0.00 | 0.00 | 0.72 | 0.12 | 14 | Endothelial |
| Birc5 | 1.55 | 0.00 | 0.00 | 0.61 | 0.05 | 14 | Endothelial |
| Stmn1 | 1.54 | 0.00 | 0.00 | 0.83 | 0.22 | 14 | Endothelial |
| 2810417H13Rik | 1.53 | 0.00 | 0.00 | 0.62 | 0.07 | 14 | Endothelial |
| Tpx2 | 1.49 | 0.00 | 0.00 | 0.55 | 0.04 | 14 | Endothelial |
| Ube2c | 1.48 | 0.00 | 0.00 | 0.46 | 0.04 | 14 | Endothelial |
| Ccnb2 | 1.46 | 0.00 | 0.00 | 0.51 | 0.05 | 14 | Endothelial |
| Cdc20 | 1.44 | 0.00 | 0.00 | 0.46 | 0.04 | 14 | Endothelial |
| Ckap2 | 1.42 | 0.00 | 0.00 | 0.48 | 0.03 | 14 | Endothelial |
| Kif23 | 1.41 | 0.00 | 0.00 | 0.50 | 0.04 | 14 | Endothelial |
| Cenpa | 1.39 | 0.00 | 0.00 | 0.48 | 0.05 | 14 | Endothelial |
| Cks2 | 1.35 | 0.00 | 0.00 | 0.52 | 0.07 | 14 | Endothelial |
| Kif20b | 1.33 | 0.00 | 0.00 | 0.49 | 0.05 | 14 | Endothelial |
| Ckap2l | 1.33 | 0.00 | 0.00 | 0.48 | 0.03 | 14 | Endothelial |
| Gm10282 | 1.31 | 0.00 | 0.00 | 0.87 | 0.39 | 14 | Endothelial |
| Hmgn2 | 1.31 | 0.00 | 0.00 | 0.88 | 0.42 | 14 | Endothelial |
| Hmmr | 1.30 | 0.00 | 0.00 | 0.42 | 0.03 | 14 | Endothelial |
| Incenp | 1.29 | 0.00 | 0.00 | 0.52 | 0.07 | 14 | Endothelial |
| Spc25 | 1.27 | 0.00 | 0.00 | 0.46 | 0.04 | 14 | Endothelial |
| D17H6S56E-5 | 1.24 | 0.00 | 0.00 | 0.58 | 0.09 | 14 | Endothelial |
| Cdca8 | 1.24 | 0.00 | 0.00 | 0.46 | 0.04 | 14 | Endothelial |
| Smc4 | 1.22 | 0.00 | 0.00 | 0.83 | 0.33 | 14 | Endothelial |
| H2afz | 1.21 | 0.00 | 0.00 | 0.86 | 0.39 | 14 | Endothelial |
| Rrm2 | 1.21 | 0.00 | 0.00 | 0.42 | 0.04 | 14 | Endothelial |
| Sgol2 | 1.20 | 0.00 | 0.00 | 0.39 | 0.03 | 14 | Endothelial |
| Cdk1 | 1.18 | 0.00 | 0.00 | 0.47 | 0.04 | 14 | Endothelial |
| Pbk | 1.17 | 0.00 | 0.00 | 0.39 | 0.02 | 14 | Endothelial |
| Egfl7 | 1.16 | 0.00 | 0.00 | 0.90 | 0.26 | 14 | Endothelial |
| Casc5 | 1.16 | 0.00 | 0.00 | 0.41 | 0.04 | 14 | Endothelial |
| Tuba1b | 1.15 | 0.00 | 0.00 | 0.87 | 0.44 | 14 | Endothelial |
| Cd93 | 1.15 | 0.00 | 0.00 | 0.80 | 0.20 | 14 | Endothelial |
| Esco2 | 1.14 | 0.00 | 0.00 | 0.36 | 0.03 | 14 | Endothelial |
| Anln | 1.14 | 0.00 | 0.00 | 0.39 | 0.02 | 14 | Endothelial |
| Hist1h1b | 1.13 | 0.00 | 0.00 | 0.33 | 0.03 | 14 | Endothelial |
| Tacc3 | 1.12 | 0.00 | 0.00 | 0.43 | 0.05 | 14 | Endothelial |
| Hjurp | 1.11 | 0.00 | 0.00 | 0.55 | 0.13 | 14 | Endothelial |
| Diap3 | 1.11 | 0.00 | 0.00 | 0.43 | 0.05 | 14 | Endothelial |
| Lmnb1 | 1.09 | 0.00 | 0.00 | 0.47 | 0.07 | 14 | Endothelial |
| Fam64a | 1.09 | 0.00 | 0.00 | 0.34 | 0.01 | 14 | Endothelial |
| Nuf2 | 1.08 | 0.00 | 0.00 | 0.36 | 0.02 | 14 | Endothelial |
| Kif15 | 1.07 | 0.00 | 0.00 | 0.40 | 0.04 | 14 | Endothelial |
| Cep55 | 1.05 | 0.00 | 0.00 | 0.34 | 0.02 | 14 | Endothelial |
| Ccna2 | 1.04 | 0.00 | 0.00 | 0.39 | 0.03 | 14 | Endothelial |
| Slfn9 | 1.04 | 0.00 | 0.00 | 0.30 | 0.03 | 14 | Endothelial |
| Fbxo5 | 1.04 | 0.00 | 0.00 | 0.38 | 0.04 | 14 | Endothelial |
| Kif11 | 1.03 | 0.00 | 0.00 | 0.36 | 0.03 | 14 | Endothelial |
| Atad2 | 1.02 | 0.00 | 0.00 | 0.41 | 0.07 | 14 | Endothelial |
| Tmpo | 1.02 | 0.00 | 0.00 | 0.55 | 0.14 | 14 | Endothelial |
| Scn7a | 1.02 | 0.00 | 0.00 | 0.88 | 0.28 | 14 | Endothelial |
| Pcdh17 | 1.02 | 0.00 | 0.00 | 0.73 | 0.19 | 14 | Endothelial |
| Aspm | 1.01 | 0.00 | 0.00 | 0.30 | 0.02 | 14 | Endothelial |
| Hpgd | 1.00 | 0.00 | 0.00 | 0.76 | 0.25 | 14 | Endothelial |
| Cks1b | 1.00 | 0.00 | 0.00 | 0.46 | 0.08 | 14 | Endothelial |
| Tmem100 | 1.00 | 0.00 | 0.00 | 0.85 | 0.28 | 14 | Endothelial |
| Ncapg | 0.99 | 0.00 | 0.00 | 0.39 | 0.04 | 14 | Endothelial |
| RP23-45G16.5 | 0.98 | 0.00 | 0.00 | 0.38 | 0.04 | 14 | Endothelial |
| Hirip3 | 0.98 | 0.00 | 0.00 | 0.41 | 0.08 | 14 | Endothelial |
| Ccdc34 | 0.98 | 0.00 | 0.00 | 0.53 | 0.13 | 14 | Endothelial |
| Tk1 | 0.98 | 0.00 | 0.00 | 0.34 | 0.03 | 14 | Endothelial |
| Cenpq | 0.97 | 0.00 | 0.00 | 0.39 | 0.06 | 14 | Endothelial |
| Ndc80 | 0.97 | 0.00 | 0.00 | 0.37 | 0.03 | 14 | Endothelial |
| Ripply3 | 0.97 | 0.00 | 0.00 | 0.55 | 0.12 | 14 | Endothelial |
| Racgap1 | 0.96 | 0.00 | 0.00 | 0.36 | 0.03 | 14 | Endothelial |
| Knstrn | 0.96 | 0.00 | 0.00 | 0.34 | 0.03 | 14 | Endothelial |
| Ptprb | 0.94 | 0.00 | 0.00 | 0.78 | 0.20 | 14 | Endothelial |
| Cdh5 | 0.93 | 0.00 | 0.00 | 0.80 | 0.21 | 14 | Endothelial |
| Neil3 | 0.93 | 0.00 | 0.00 | 0.30 | 0.02 | 14 | Endothelial |
| Gpihbp1 | 0.93 | 0.00 | 0.00 | 0.63 | 0.15 | 14 | Endothelial |
| Clspn | 0.91 | 0.00 | 0.00 | 0.31 | 0.04 | 14 | Endothelial |
| Ncapd2 | 0.90 | 0.00 | 0.00 | 0.35 | 0.04 | 14 | Endothelial |
| Pecam1 | 0.89 | 0.00 | 0.00 | 0.77 | 0.23 | 14 | Endothelial |
| Mis18bp1 | 0.89 | 0.00 | 0.00 | 0.31 | 0.03 | 14 | Endothelial |
| Trim59 | 0.89 | 0.00 | 0.00 | 0.31 | 0.04 | 14 | Endothelial |
| Ccne2 | 0.88 | 0.00 | 0.00 | 0.26 | 0.03 | 14 | Endothelial |
| Calcrl | 0.87 | 0.00 | 0.00 | 0.80 | 0.24 | 14 | Endothelial |
| Spc24 | 0.87 | 0.00 | 0.00 | 0.34 | 0.04 | 14 | Endothelial |
| Rasgrp3 | 0.85 | 0.00 | 0.00 | 0.52 | 0.12 | 14 | Endothelial |
| C330027C09Rik | 0.85 | 0.00 | 0.00 | 0.27 | 0.02 | 14 | Endothelial |
| Plvap | 0.83 | 0.00 | 0.00 | 0.52 | 0.12 | 14 | Endothelial |
| Sgol1 | 0.82 | 0.00 | 0.00 | 0.28 | 0.02 | 14 | Endothelial |
| Clec14a | 0.81 | 0.00 | 0.00 | 0.63 | 0.16 | 14 | Endothelial |
| Tyms | 0.81 | 0.00 | 0.00 | 0.32 | 0.05 | 14 | Endothelial |
| Cenpw | 0.78 | 0.00 | 0.00 | 0.31 | 0.04 | 14 | Endothelial |
| Cdca3 | 0.78 | 0.00 | 0.00 | 0.29 | 0.03 | 14 | Endothelial |
| Aurkb | 0.77 | 0.00 | 0.00 | 0.27 | 0.02 | 14 | Endothelial |
| Kif22 | 0.74 | 0.00 | 0.00 | 0.26 | 0.02 | 14 | Endothelial |
| Bub1 | 0.74 | 0.00 | 0.00 | 0.25 | 0.02 | 14 | Endothelial |
| Lyve1 | 0.71 | 0.00 | 0.00 | 0.73 | 0.21 | 14 | Endothelial |
| Rad51ap1 | 0.71 | 0.00 | 0.00 | 0.27 | 0.03 | 14 | Endothelial |
| Hist2h2ab | 0.78 | 0.00 | 0.00 | 0.26 | 0.04 | 14 | Endothelial |
| Cyyr1 | 0.84 | 0.00 | 0.00 | 0.55 | 0.14 | 14 | Endothelial |
| Epas1 | 0.85 | 0.00 | 0.00 | 0.75 | 0.24 | 14 | Endothelial |
| Sox17 | 0.78 | 0.00 | 0.00 | 0.53 | 0.13 | 14 | Endothelial |
| Anp32e | 1.07 | 0.00 | 0.00 | 0.73 | 0.27 | 14 | Endothelial |
| Mest | 0.87 | 0.00 | 0.00 | 0.64 | 0.19 | 14 | Endothelial |
| BC028528 | 0.85 | 0.00 | 0.00 | 0.78 | 0.28 | 14 | Endothelial |
| Cit | 0.66 | 0.00 | 0.00 | 0.27 | 0.04 | 14 | Endothelial |
| Tubb6 | 0.80 | 0.00 | 0.00 | 0.36 | 0.07 | 14 | Endothelial |
| Sdpr | 0.92 | 0.00 | 0.00 | 0.94 | 0.51 | 14 | Endothelial |
| Cldn5 | 0.69 | 0.00 | 0.00 | 0.71 | 0.21 | 14 | Endothelial |
| Ramp2 | 0.67 | 0.00 | 0.00 | 0.80 | 0.27 | 14 | Endothelial |
| Ecscr | 0.72 | 0.00 | 0.00 | 0.59 | 0.16 | 14 | Endothelial |
| Gpr116 | 0.83 | 0.00 | 0.00 | 0.87 | 0.36 | 14 | Endothelial |
| Tspan13 | 0.78 | 0.00 | 0.00 | 0.72 | 0.24 | 14 | Endothelial |
| Plk4 | 0.62 | 0.00 | 0.00 | 0.26 | 0.04 | 14 | Endothelial |
| Erg | 0.78 | 0.00 | 0.00 | 0.50 | 0.12 | 14 | Endothelial |
| Tie1 | 0.72 | 0.00 | 0.00 | 0.46 | 0.10 | 14 | Endothelial |
| Tek | 0.73 | 0.00 | 0.00 | 0.49 | 0.12 | 14 | Endothelial |
| Ctla2a | 0.74 | 0.00 | 0.00 | 0.64 | 0.19 | 14 | Endothelial |
| Flt1 | 0.73 | 0.00 | 0.00 | 0.54 | 0.14 | 14 | Endothelial |
| Tspan7 | 0.72 | 0.00 | 0.00 | 0.66 | 0.20 | 14 | Endothelial |
| Myzap | 0.71 | 0.00 | 0.00 | 0.52 | 0.13 | 14 | Endothelial |
| 2810025M15Rik | 0.76 | 0.00 | 0.00 | 0.47 | 0.11 | 14 | Endothelial |
| Gmnn | 0.77 | 0.00 | 0.00 | 0.34 | 0.06 | 14 | Endothelial |
| Ncapd3 | 0.78 | 0.00 | 0.00 | 0.29 | 0.05 | 14 | Endothelial |
| Hist1h1e | 1.32 | 0.00 | 0.00 | 0.52 | 0.15 | 14 | Endothelial |
| Tubb5 | 1.04 | 0.00 | 0.00 | 0.82 | 0.43 | 14 | Endothelial |
| 2700094K13Rik | 0.94 | 0.00 | 0.00 | 0.53 | 0.15 | 14 | Endothelial |
| Ppap2a | 0.75 | 0.00 | 0.00 | 0.60 | 0.18 | 14 | Endothelial |
| Ptma | 0.67 | 0.00 | 0.00 | 0.99 | 0.91 | 14 | Endothelial |
| Ncaph | 0.61 | 0.00 | 0.00 | 0.26 | 0.04 | 14 | Endothelial |
| H2-Q7 | 0.83 | 0.00 | 0.00 | 0.34 | 0.07 | 14 | Endothelial |
| Nsg1 | 0.68 | 0.00 | 0.00 | 0.37 | 0.08 | 14 | Endothelial |
| Ckap5 | 0.90 | 0.00 | 0.00 | 0.43 | 0.11 | 14 | Endothelial |
| Myct1 | 0.71 | 0.00 | 0.00 | 0.49 | 0.12 | 14 | Endothelial |
| Kpna2 | 0.80 | 0.00 | 0.00 | 0.30 | 0.06 | 14 | Endothelial |
| Rad21 | 0.97 | 0.00 | 0.00 | 0.63 | 0.22 | 14 | Endothelial |
| Dbf4 | 0.65 | 0.00 | 0.00 | 0.28 | 0.05 | 14 | Endothelial |
| Kit | 0.63 | 0.00 | 0.00 | 0.37 | 0.08 | 14 | Endothelial |
| Kdr | 0.63 | 0.00 | 0.00 | 0.55 | 0.15 | 14 | Endothelial |
| Tnfsf10 | 0.65 | 0.00 | 0.00 | 0.38 | 0.08 | 14 | Endothelial |
| Ushbp1 | 0.65 | 0.00 | 0.00 | 0.38 | 0.08 | 14 | Endothelial |
| Gm10184 | 0.83 | 0.00 | 0.00 | 0.31 | 0.06 | 14 | Endothelial |
| Hist1h1d | 1.04 | 0.00 | 0.00 | 0.39 | 0.09 | 14 | Endothelial |
| Sema3g | 0.67 | 0.00 | 0.00 | 0.33 | 0.07 | 14 | Endothelial |
| Slc43a3 | 0.77 | 0.00 | 0.00 | 0.63 | 0.21 | 14 | Endothelial |
| Rangap1 | 0.71 | 0.00 | 0.00 | 0.34 | 0.07 | 14 | Endothelial |
| Icam2 | 0.66 | 0.00 | 0.00 | 0.54 | 0.15 | 14 | Endothelial |
| Palmd | 0.80 | 0.00 | 0.00 | 0.46 | 0.12 | 14 | Endothelial |
| Aplnr | 0.57 | 0.00 | 0.00 | 0.33 | 0.06 | 14 | Endothelial |
| Nucks1 | 0.92 | 0.00 | 0.00 | 0.69 | 0.27 | 14 | Endothelial |
| Emcn | 0.62 | 0.00 | 0.00 | 0.43 | 0.10 | 14 | Endothelial |
| Tspan18 | 0.66 | 0.00 | 0.00 | 0.48 | 0.13 | 14 | Endothelial |
| Hist2h2ac | 0.88 | 0.00 | 0.00 | 0.30 | 0.06 | 14 | Endothelial |
| Ece1 | 0.76 | 0.00 | 0.00 | 0.68 | 0.24 | 14 | Endothelial |
| Tmem88 | 0.66 | 0.00 | 0.00 | 0.41 | 0.10 | 14 | Endothelial |
| Gm10259 | 0.73 | 0.00 | 0.00 | 0.29 | 0.05 | 14 | Endothelial |
| Ska2 | 0.66 | 0.00 | 0.00 | 0.30 | 0.06 | 14 | Endothelial |
| Esam | 0.68 | 0.00 | 0.00 | 0.56 | 0.17 | 14 | Endothelial |
| Hn1 | 0.83 | 0.00 | 0.00 | 0.56 | 0.19 | 14 | Endothelial |
| Arhgef15 | 0.60 | 0.00 | 0.00 | 0.33 | 0.07 | 14 | Endothelial |
| Nostrin | 0.71 | 0.00 | 0.00 | 0.50 | 0.14 | 14 | Endothelial |
| S100a16 | 0.76 | 0.00 | 0.00 | 0.64 | 0.23 | 14 | Endothelial |
| Tuba1c | 0.93 | 0.00 | 0.00 | 0.70 | 0.31 | 14 | Endothelial |
| G2e3 | 0.73 | 0.00 | 0.00 | 0.29 | 0.06 | 14 | Endothelial |
| Acvrl1 | 0.63 | 0.00 | 0.00 | 0.55 | 0.17 | 14 | Endothelial |
| Clec1a | 0.59 | 0.00 | 0.00 | 0.53 | 0.16 | 14 | Endothelial |
| Podxl | 0.61 | 0.00 | 0.00 | 0.41 | 0.10 | 14 | Endothelial |
| Ldb2 | 0.69 | 0.00 | 0.00 | 0.52 | 0.15 | 14 | Endothelial |
| Prim1 | 0.65 | 0.00 | 0.00 | 0.26 | 0.05 | 14 | Endothelial |
| She | 0.51 | 0.00 | 0.00 | 0.28 | 0.05 | 14 | Endothelial |
| Eng | 0.68 | 0.00 | 0.00 | 0.41 | 0.11 | 14 | Endothelial |
| Dek | 0.97 | 0.00 | 0.00 | 0.83 | 0.49 | 14 | Endothelial |
| Armcx4 | 0.66 | 0.00 | 0.00 | 0.33 | 0.07 | 14 | Endothelial |
| Cd34 | 0.69 | 0.00 | 0.00 | 0.57 | 0.19 | 14 | Endothelial |
| Piezo2 | 0.67 | 0.00 | 0.00 | 0.41 | 0.11 | 14 | Endothelial |
| Sgk1 | 0.72 | 0.00 | 0.00 | 0.54 | 0.18 | 14 | Endothelial |
| Tm4sf1 | 0.58 | 0.00 | 0.00 | 0.61 | 0.20 | 14 | Endothelial |
| Ran | 0.83 | 0.00 | 0.00 | 0.76 | 0.37 | 14 | Endothelial |
| Rrm1 | 0.61 | 0.00 | 0.00 | 0.29 | 0.06 | 14 | Endothelial |
| Lig1 | 0.79 | 0.00 | 0.00 | 0.36 | 0.09 | 14 | Endothelial |
| Gata2 | 0.58 | 0.00 | 0.00 | 0.46 | 0.13 | 14 | Endothelial |
| Afap1l1 | 0.70 | 0.00 | 0.00 | 0.52 | 0.17 | 14 | Endothelial |
| Whsc1 | 0.79 | 0.00 | 0.00 | 0.44 | 0.13 | 14 | Endothelial |
| Stmn2 | 0.70 | 0.00 | 0.00 | 0.48 | 0.15 | 14 | Endothelial |
| Srgn | 0.62 | 0.00 | 0.00 | 0.75 | 0.32 | 14 | Endothelial |
| Ehd4 | 0.68 | 0.00 | 0.00 | 0.68 | 0.27 | 14 | Endothelial |
| Emp1 | 0.71 | 0.00 | 0.00 | 0.58 | 0.21 | 14 | Endothelial |
| Myef2 | 0.75 | 0.00 | 0.00 | 0.47 | 0.15 | 14 | Endothelial |
| Ddah1 | 0.50 | 0.00 | 0.00 | 0.27 | 0.06 | 14 | Endothelial |
| H1f0 | 0.98 | 0.00 | 0.00 | 0.62 | 0.26 | 14 | Endothelial |
| Itga6 | 0.64 | 0.00 | 0.00 | 0.43 | 0.13 | 14 | Endothelial |
| Hmgb1 | 0.74 | 0.00 | 0.00 | 0.88 | 0.60 | 14 | Endothelial |
| Gng11 | 0.68 | 0.00 | 0.00 | 0.68 | 0.28 | 14 | Endothelial |
| Cav1 | 0.61 | 0.00 | 0.00 | 0.73 | 0.30 | 14 | Endothelial |
| Sema3c | 0.78 | 0.00 | 0.00 | 0.53 | 0.19 | 14 | Endothelial |
| Fbln2 | 0.50 | 0.00 | 0.00 | 0.29 | 0.07 | 14 | Endothelial |
| Rasip1 | 0.54 | 0.00 | 0.00 | 0.41 | 0.11 | 14 | Endothelial |
| Gm9833 | 0.64 | 0.00 | 0.00 | 0.38 | 0.11 | 14 | Endothelial |
| Ddah2 | 0.74 | 0.00 | 0.00 | 0.62 | 0.25 | 14 | Endothelial |
| Id3 | 0.66 | 0.00 | 0.00 | 0.83 | 0.44 | 14 | Endothelial |
| Arap3 | 0.49 | 0.00 | 0.00 | 0.30 | 0.07 | 14 | Endothelial |
| H2-Q4 | 0.60 | 0.00 | 0.00 | 0.27 | 0.06 | 14 | Endothelial |
| Hist2h2aa2 | 0.98 | 0.00 | 0.00 | 0.41 | 0.13 | 14 | Endothelial |
| Kitl | 0.51 | 0.00 | 0.00 | 0.59 | 0.21 | 14 | Endothelial |
| Mcam | 0.53 | 0.00 | 0.00 | 0.43 | 0.12 | 14 | Endothelial |
| Ezh2 | 0.77 | 0.00 | 0.00 | 0.51 | 0.18 | 14 | Endothelial |
| Nav1 | 0.59 | 0.00 | 0.00 | 0.38 | 0.11 | 14 | Endothelial |
| Ptprg | 0.63 | 0.00 | 0.00 | 0.40 | 0.12 | 14 | Endothelial |
| Anp32a | 0.69 | 0.00 | 0.00 | 0.87 | 0.59 | 14 | Endothelial |
| Elk3 | 0.59 | 0.00 | 0.00 | 0.43 | 0.13 | 14 | Endothelial |
| Tubb4b | 0.67 | 0.00 | 0.00 | 0.51 | 0.18 | 14 | Endothelial |
| Lbr | 0.63 | 0.00 | 0.00 | 0.39 | 0.12 | 14 | Endothelial |
| Col4a2 | 0.64 | 0.00 | 0.00 | 0.80 | 0.42 | 14 | Endothelial |
| Mcm3 | 0.63 | 0.00 | 0.00 | 0.28 | 0.07 | 14 | Endothelial |
| Tbx3 | 0.59 | 0.00 | 0.00 | 0.55 | 0.20 | 14 | Endothelial |
| Eltd1 | 0.47 | 0.00 | 0.00 | 0.28 | 0.06 | 14 | Endothelial |
| Col4a1 | 0.66 | 0.00 | 0.00 | 0.85 | 0.48 | 14 | Endothelial |
| Tuba1a | 0.66 | 0.00 | 0.00 | 0.85 | 0.50 | 14 | Endothelial |
| Fendrr | 0.63 | 0.00 | 0.00 | 0.54 | 0.19 | 14 | Endothelial |
| Adcy4 | 0.53 | 0.00 | 0.00 | 0.32 | 0.08 | 14 | Endothelial |
| Usp1 | 0.75 | 0.00 | 0.00 | 0.44 | 0.15 | 14 | Endothelial |
| Ccnd1 | 0.73 | 0.00 | 0.00 | 0.51 | 0.18 | 14 | Endothelial |
| Nasp | 0.79 | 0.00 | 0.00 | 0.51 | 0.19 | 14 | Endothelial |
| Prx | 0.51 | 0.00 | 0.00 | 0.36 | 0.10 | 14 | Endothelial |
| Ly6e | 0.60 | 0.00 | 0.00 | 0.81 | 0.41 | 14 | Endothelial |
| Gimap1 | 0.53 | 0.00 | 0.00 | 0.43 | 0.13 | 14 | Endothelial |
| Hmgb3 | 0.65 | 0.00 | 0.00 | 0.43 | 0.14 | 14 | Endothelial |
| Dnajc9 | 0.64 | 0.00 | 0.00 | 0.38 | 0.11 | 14 | Endothelial |
| Tubb2a | 0.66 | 0.00 | 0.00 | 0.54 | 0.21 | 14 | Endothelial |
| Bst2 | 0.59 | 0.00 | 0.00 | 0.42 | 0.13 | 14 | Endothelial |
| Stard4 | 0.60 | 0.00 | 0.00 | 0.34 | 0.09 | 14 | Endothelial |
| Gm6104 | 0.63 | 0.00 | 0.00 | 0.39 | 0.12 | 14 | Endothelial |
| Pcdh1 | 0.46 | 0.00 | 0.00 | 0.28 | 0.07 | 14 | Endothelial |
| Cav2 | 0.55 | 0.00 | 0.00 | 0.52 | 0.18 | 14 | Endothelial |
| Hist1h1c | 0.86 | 0.00 | 0.00 | 0.36 | 0.11 | 14 | Endothelial |
| Klhl5 | 0.60 | 0.00 | 0.00 | 0.43 | 0.14 | 14 | Endothelial |
| Gimap4 | 0.52 | 0.00 | 0.00 | 0.50 | 0.17 | 14 | Endothelial |
| Cenpc1 | 0.63 | 0.00 | 0.00 | 0.31 | 0.08 | 14 | Endothelial |
| Nde1 | 0.53 | 0.00 | 0.00 | 0.26 | 0.06 | 14 | Endothelial |
| Slc29a1 | 0.53 | 0.00 | 0.00 | 0.30 | 0.08 | 14 | Endothelial |
| Fkbp1a | 0.64 | 0.00 | 0.00 | 0.75 | 0.40 | 14 | Endothelial |
| Foxf1 | 0.52 | 0.00 | 0.00 | 0.38 | 0.11 | 14 | Endothelial |
| Ace | 0.53 | 0.00 | 0.00 | 0.42 | 0.13 | 14 | Endothelial |
| S1pr1 | 0.56 | 0.00 | 0.00 | 0.51 | 0.18 | 14 | Endothelial |
| Dnmt1 | 0.66 | 0.00 | 0.00 | 0.34 | 0.10 | 14 | Endothelial |
| Pcna | 0.87 | 0.00 | 0.00 | 0.46 | 0.17 | 14 | Endothelial |
| Pcna-ps2 | 0.88 | 0.00 | 0.00 | 0.46 | 0.17 | 14 | Endothelial |
| Fli1 | 0.54 | 0.00 | 0.00 | 0.37 | 0.11 | 14 | Endothelial |
| Casz1 | 0.55 | 0.00 | 0.00 | 0.41 | 0.13 | 14 | Endothelial |
| Ccdc85b | 0.54 | 0.00 | 0.00 | 0.37 | 0.11 | 14 | Endothelial |
| Tmem2 | 0.51 | 0.00 | 0.00 | 0.43 | 0.14 | 14 | Endothelial |
| Calm2 | 0.64 | 0.00 | 0.00 | 0.88 | 0.63 | 14 | Endothelial |
| Tubb2b | 0.56 | 0.00 | 0.00 | 0.39 | 0.12 | 14 | Endothelial |
| Klf4 | 0.56 | 0.00 | 0.00 | 0.56 | 0.22 | 14 | Endothelial |
| Prcp | 0.53 | 0.00 | 0.00 | 0.32 | 0.09 | 14 | Endothelial |
| Rnf144a | 0.56 | 0.00 | 0.00 | 0.44 | 0.15 | 14 | Endothelial |
| Jup | 0.50 | 0.00 | 0.00 | 0.45 | 0.15 | 14 | Endothelial |
| Tpm4 | 0.60 | 0.00 | 0.00 | 0.82 | 0.48 | 14 | Endothelial |
| Ppia | 0.57 | 0.00 | 0.00 | 0.93 | 0.79 | 14 | Endothelial |
| Arhgap29 | 0.61 | 0.00 | 0.00 | 0.53 | 0.21 | 14 | Endothelial |
| 5830418K08Rik | 0.59 | 0.00 | 0.00 | 0.30 | 0.08 | 14 | Endothelial |
| Hnrnpa2b1 | 0.58 | 0.00 | 0.00 | 0.92 | 0.71 | 14 | Endothelial |
| Lxn | 0.50 | 0.00 | 0.00 | 0.37 | 0.11 | 14 | Endothelial |
| Aqp1 | 0.43 | 0.00 | 0.00 | 0.51 | 0.19 | 14 | Endothelial |
| Slco2a1 | 0.46 | 0.00 | 0.00 | 0.35 | 0.11 | 14 | Endothelial |
| 10-Sep | 0.53 | 0.00 | 0.00 | 0.31 | 0.09 | 14 | Endothelial |
| Id1 | 0.58 | 0.00 | 0.00 | 0.56 | 0.23 | 14 | Endothelial |
| Lsm2 | 0.51 | 0.00 | 0.00 | 0.30 | 0.08 | 14 | Endothelial |
| H2afv | 0.68 | 0.00 | 0.00 | 0.60 | 0.27 | 14 | Endothelial |
| Thbd | 0.52 | 0.00 | 0.00 | 0.59 | 0.24 | 14 | Endothelial |
| Ahr | 0.45 | 0.00 | 0.00 | 0.32 | 0.09 | 14 | Endothelial |
| Tipin | 0.57 | 0.00 | 0.00 | 0.32 | 0.10 | 14 | Endothelial |
| Cdkn2d | 0.53 | 0.00 | 0.00 | 0.25 | 0.06 | 14 | Endothelial |
| Edn1 | 0.53 | 0.00 | 0.00 | 0.32 | 0.09 | 14 | Endothelial |
| Gm10123 | 0.55 | 0.00 | 0.00 | 0.92 | 0.81 | 14 | Endothelial |
| Rgs12 | 0.48 | 0.00 | 0.00 | 0.37 | 0.12 | 14 | Endothelial |
| Pde4b | 0.54 | 0.00 | 0.00 | 0.40 | 0.13 | 14 | Endothelial |
| Cdk5rap2 | 0.54 | 0.00 | 0.00 | 0.30 | 0.09 | 14 | Endothelial |
| Crip2 | 0.54 | 0.00 | 0.00 | 0.76 | 0.40 | 14 | Endothelial |
| Rapgef5 | 0.50 | 0.00 | 0.00 | 0.33 | 0.10 | 14 | Endothelial |
| Ets1 | 0.49 | 0.00 | 0.00 | 0.57 | 0.24 | 14 | Endothelial |
| Arl4c | 0.57 | 0.00 | 0.00 | 0.32 | 0.10 | 14 | Endothelial |
| Bmpr2 | 0.50 | 0.00 | 0.00 | 0.31 | 0.09 | 14 | Endothelial |
| Ly6a | 0.34 | 0.00 | 0.00 | 0.50 | 0.19 | 14 | Endothelial |
| Cd36 | 0.51 | 0.00 | 0.00 | 0.68 | 0.32 | 14 | Endothelial |
| Lphn2 | 0.51 | 0.00 | 0.00 | 0.38 | 0.13 | 14 | Endothelial |
| Rdx | 0.63 | 0.00 | 0.00 | 0.74 | 0.44 | 14 | Endothelial |
| Gm12355 | 0.58 | 0.00 | 0.00 | 0.56 | 0.25 | 14 | Endothelial |
| Ptprm | 0.46 | 0.00 | 0.00 | 0.31 | 0.09 | 14 | Endothelial |
| Rgcc | 0.37 | 0.00 | 0.00 | 0.46 | 0.17 | 14 | Endothelial |
| Pea15a | 0.47 | 0.00 | 0.00 | 0.39 | 0.14 | 14 | Endothelial |
| Anp32b | 0.61 | 0.00 | 0.00 | 0.79 | 0.49 | 14 | Endothelial |
| Nes | 0.43 | 0.00 | 0.00 | 0.52 | 0.21 | 14 | Endothelial |
| Srsf3 | 0.60 | 0.00 | 0.00 | 0.75 | 0.44 | 14 | Endothelial |
| Arl6ip1 | 0.93 | 0.00 | 0.00 | 0.78 | 0.52 | 14 | Endothelial |
| Foxp1 | 0.53 | 0.00 | 0.00 | 0.76 | 0.44 | 14 | Endothelial |
| Ephb4 | 0.41 | 0.00 | 0.00 | 0.27 | 0.08 | 14 | Endothelial |
| Slc9a3r2 | 0.47 | 0.00 | 0.00 | 0.50 | 0.20 | 14 | Endothelial |
| Gm10068 | 0.61 | 0.00 | 0.00 | 0.60 | 0.29 | 14 | Endothelial |
| Ppp1r16b | 0.40 | 0.00 | 0.00 | 0.30 | 0.09 | 14 | Endothelial |
| Dut | 0.49 | 0.00 | 0.00 | 0.28 | 0.08 | 14 | Endothelial |
| Cbx3 | 0.59 | 0.00 | 0.00 | 0.64 | 0.33 | 14 | Endothelial |
| Tjp1 | 0.45 | 0.00 | 0.00 | 0.42 | 0.16 | 14 | Endothelial |
| Suz12 | 0.52 | 0.00 | 0.00 | 0.36 | 0.13 | 14 | Endothelial |
| Adam15 | 0.42 | 0.00 | 0.00 | 0.27 | 0.08 | 14 | Endothelial |
| Csrp2 | 0.46 | 0.00 | 0.00 | 0.43 | 0.16 | 14 | Endothelial |
| Slbp | 0.63 | 0.00 | 0.00 | 0.38 | 0.14 | 14 | Endothelial |
| Slk | 0.56 | 0.00 | 0.00 | 0.74 | 0.42 | 14 | Endothelial |
| Grap | 0.40 | 0.00 | 0.00 | 0.37 | 0.13 | 14 | Endothelial |
| 9430020K01Rik | 0.44 | 0.00 | 0.00 | 0.36 | 0.12 | 14 | Endothelial |
| Nin | 0.60 | 0.00 | 0.00 | 0.37 | 0.14 | 14 | Endothelial |
| 1810011O10Rik | 0.35 | 0.00 | 0.00 | 0.27 | 0.08 | 14 | Endothelial |
| 4931406P16Rik | 0.47 | 0.00 | 0.00 | 0.45 | 0.17 | 14 | Endothelial |
| Sptbn1 | 0.55 | 0.00 | 0.00 | 0.82 | 0.54 | 14 | Endothelial |
| Cntln | 0.58 | 0.00 | 0.00 | 0.33 | 0.11 | 14 | Endothelial |
| Hes1 | 0.56 | 0.00 | 0.00 | 0.59 | 0.28 | 14 | Endothelial |
| Orc6 | 0.54 | 0.00 | 0.00 | 0.30 | 0.10 | 14 | Endothelial |
| Tubb4b-ps1 | 0.45 | 0.00 | 0.00 | 0.27 | 0.08 | 14 | Endothelial |
| Snx3 | 0.57 | 0.00 | 0.00 | 0.62 | 0.32 | 14 | Endothelial |
| Efnb2 | 0.50 | 0.00 | 0.00 | 0.39 | 0.14 | 14 | Endothelial |
| Maoa | 0.43 | 0.00 | 0.00 | 0.26 | 0.08 | 14 | Endothelial |
| Acer2 | 0.38 | 0.00 | 0.00 | 0.30 | 0.10 | 14 | Endothelial |
| H2-D1 | 0.46 | 0.00 | 0.00 | 0.82 | 0.49 | 14 | Endothelial |
| Scn3b | 0.35 | 0.00 | 0.00 | 0.25 | 0.07 | 14 | Endothelial |
| Pitpnc1 | 0.38 | 0.00 | 0.00 | 0.35 | 0.12 | 14 | Endothelial |
| Snrpd1 | 0.53 | 0.00 | 0.00 | 0.47 | 0.20 | 14 | Endothelial |
| Cpd | 0.52 | 0.00 | 0.00 | 0.38 | 0.14 | 14 | Endothelial |
| Vim | 0.46 | 0.00 | 0.00 | 0.91 | 0.65 | 14 | Endothelial |
| Prkd2 | 0.43 | 0.00 | 0.00 | 0.29 | 0.09 | 14 | Endothelial |
| Prkce | 0.41 | 0.00 | 0.00 | 0.25 | 0.07 | 14 | Endothelial |
| Uaca | 0.45 | 0.00 | 0.00 | 0.37 | 0.13 | 14 | Endothelial |
| Banf1 | 0.57 | 0.00 | 0.00 | 0.57 | 0.28 | 14 | Endothelial |
| Yes1 | 0.45 | 0.00 | 0.00 | 0.29 | 0.10 | 14 | Endothelial |
| Gimap6 | 0.35 | 0.00 | 0.00 | 0.42 | 0.16 | 14 | Endothelial |
| Sae1 | 0.47 | 0.00 | 0.00 | 0.31 | 0.10 | 14 | Endothelial |
| B2m | 0.40 | 0.00 | 0.00 | 0.91 | 0.67 | 14 | Endothelial |
| Klf7 | 0.47 | 0.00 | 0.00 | 0.44 | 0.18 | 14 | Endothelial |
| Heg1 | 0.44 | 0.00 | 0.00 | 0.41 | 0.16 | 14 | Endothelial |
| Brd8 | 0.56 | 0.00 | 0.00 | 0.44 | 0.19 | 14 | Endothelial |
| Myl12b | 0.52 | 0.00 | 0.00 | 0.82 | 0.60 | 14 | Endothelial |
| Gcc2 | 0.54 | 0.00 | 0.00 | 0.52 | 0.24 | 14 | Endothelial |
| Rhoj | 0.40 | 0.00 | 0.00 | 0.45 | 0.18 | 14 | Endothelial |
| Cyth3 | 0.48 | 0.00 | 0.00 | 0.52 | 0.24 | 14 | Endothelial |
| Mllt4 | 0.51 | 0.00 | 0.00 | 0.47 | 0.20 | 14 | Endothelial |
| Gnai2 | 0.51 | 0.00 | 0.00 | 0.68 | 0.38 | 14 | Endothelial |
| Lrrc8c | 0.40 | 0.00 | 0.00 | 0.25 | 0.08 | 14 | Endothelial |
| Dtymk | 0.50 | 0.00 | 0.00 | 0.35 | 0.13 | 14 | Endothelial |
| Smc1a | 0.54 | 0.00 | 0.00 | 0.64 | 0.35 | 14 | Endothelial |
| Prdx4 | 0.49 | 0.00 | 0.00 | 0.40 | 0.16 | 14 | Endothelial |
| Carhsp1 | 0.40 | 0.00 | 0.00 | 0.36 | 0.13 | 14 | Endothelial |
| Rnf168 | 0.52 | 0.00 | 0.00 | 0.35 | 0.13 | 14 | Endothelial |
| Zfp503 | 0.45 | 0.00 | 0.00 | 0.38 | 0.15 | 14 | Endothelial |
| Fnbp1l | 0.49 | 0.00 | 0.00 | 0.53 | 0.25 | 14 | Endothelial |
| Arhgef12 | 0.47 | 0.00 | 0.00 | 0.51 | 0.23 | 14 | Endothelial |
| Cbx5 | 0.60 | 0.00 | 0.00 | 0.53 | 0.25 | 14 | Endothelial |
| Actn4 | 0.48 | 0.00 | 0.00 | 0.58 | 0.28 | 14 | Endothelial |
| Dnajc8 | 0.53 | 0.00 | 0.00 | 0.70 | 0.42 | 14 | Endothelial |
| Zeb1 | 0.40 | 0.00 | 0.00 | 0.31 | 0.11 | 14 | Endothelial |
| Fam111a | 0.46 | 0.00 | 0.00 | 0.38 | 0.15 | 14 | Endothelial |
| Ranbp1 | 0.54 | 0.00 | 0.00 | 0.57 | 0.28 | 14 | Endothelial |
| Cxx1a | 0.39 | 0.00 | 0.00 | 0.49 | 0.21 | 14 | Endothelial |
| Ubald2 | 0.47 | 0.00 | 0.00 | 0.36 | 0.14 | 14 | Endothelial |
| Fmo1 | 0.38 | 0.00 | 0.00 | 0.32 | 0.11 | 14 | Endothelial |
| Ywhah | 0.55 | 0.00 | 0.00 | 0.52 | 0.26 | 14 | Endothelial |
| Dlc1 | 0.44 | 0.00 | 0.00 | 0.41 | 0.17 | 14 | Endothelial |
| Meis2 | 0.39 | 0.00 | 0.00 | 0.32 | 0.11 | 14 | Endothelial |
| Stim2 | 0.40 | 0.00 | 0.00 | 0.27 | 0.09 | 14 | Endothelial |
| Hip1 | 0.43 | 0.00 | 0.00 | 0.38 | 0.15 | 14 | Endothelial |
| St3gal6 | 0.36 | 0.00 | 0.00 | 0.27 | 0.09 | 14 | Endothelial |
| Taok2 | 0.39 | 0.00 | 0.00 | 0.29 | 0.10 | 14 | Endothelial |
| Gimap5 | 0.34 | 0.00 | 0.00 | 0.31 | 0.11 | 14 | Endothelial |
| Gm6625 | 0.55 | 0.00 | 0.00 | 0.51 | 0.25 | 14 | Endothelial |
| Notch1 | 0.40 | 0.00 | 0.00 | 0.34 | 0.13 | 14 | Endothelial |
| Luzp1 | 0.47 | 0.00 | 0.00 | 0.44 | 0.19 | 14 | Endothelial |
| Xiap | 0.45 | 0.00 | 0.00 | 0.45 | 0.20 | 14 | Endothelial |
| Xpo1 | 0.39 | 0.00 | 0.00 | 0.27 | 0.09 | 14 | Endothelial |
| Marcks | 0.35 | 0.00 | 0.00 | 0.81 | 0.49 | 14 | Endothelial |
| Elovl5 | 0.49 | 0.00 | 0.00 | 0.37 | 0.15 | 14 | Endothelial |
| Cep57 | 0.41 | 0.00 | 0.00 | 0.29 | 0.11 | 14 | Endothelial |
| Cbx1 | 0.52 | 0.00 | 0.00 | 0.69 | 0.41 | 14 | Endothelial |
| Nfyb | 0.43 | 0.00 | 0.00 | 0.31 | 0.11 | 14 | Endothelial |
| Ddx39 | 0.40 | 0.00 | 0.00 | 0.26 | 0.09 | 14 | Endothelial |
| Bub3 | 0.45 | 0.00 | 0.00 | 0.35 | 0.14 | 14 | Endothelial |
| Ptgfrn | 0.40 | 0.00 | 0.00 | 0.27 | 0.09 | 14 | Endothelial |
| Cnn3 | 0.39 | 0.00 | 0.00 | 0.59 | 0.29 | 14 | Endothelial |
| Tpm3-rs7 | 0.40 | 0.00 | 0.00 | 0.88 | 0.69 | 14 | Endothelial |
| Pltp | 0.45 | 0.00 | 0.00 | 0.41 | 0.18 | 14 | Endothelial |
| Arhgap31 | 0.42 | 0.00 | 0.00 | 0.31 | 0.12 | 14 | Endothelial |
| Sema6a | 0.34 | 0.00 | 0.00 | 0.31 | 0.11 | 14 | Endothelial |
| Hoxa5 | 0.35 | 0.00 | 0.00 | 0.33 | 0.12 | 14 | Endothelial |
| Hdac7 | 0.38 | 0.00 | 0.00 | 0.34 | 0.13 | 14 | Endothelial |
| Ssrp1 | 0.46 | 0.00 | 0.00 | 0.45 | 0.21 | 14 | Endothelial |
| Ly6c1 | 0.27 | 0.00 | 0.00 | 0.43 | 0.18 | 14 | Endothelial |
| Aplp2 | 0.48 | 0.00 | 0.00 | 0.66 | 0.38 | 14 | Endothelial |
| Sypl | 0.46 | 0.00 | 0.00 | 0.52 | 0.25 | 14 | Endothelial |
| Nop56 | 0.49 | 0.00 | 0.00 | 0.43 | 0.19 | 14 | Endothelial |
| Wasf2 | 0.45 | 0.00 | 0.00 | 0.57 | 0.30 | 14 | Endothelial |
| Cd97 | 0.36 | 0.00 | 0.00 | 0.41 | 0.18 | 14 | Endothelial |
| Ctnnd1 | 0.43 | 0.00 | 0.00 | 0.37 | 0.16 | 14 | Endothelial |
| Actl6a | 0.37 | 0.00 | 0.00 | 0.29 | 0.10 | 14 | Endothelial |
| Snrk | 0.36 | 0.00 | 0.00 | 0.37 | 0.15 | 14 | Endothelial |
| Tcf4 | 0.40 | 0.00 | 0.00 | 0.70 | 0.40 | 14 | Endothelial |
| Nrp1 | 0.42 | 0.00 | 0.00 | 0.66 | 0.39 | 14 | Endothelial |
| Itgb1 | 0.38 | 0.00 | 0.00 | 0.85 | 0.62 | 14 | Endothelial |
| Ybx1 | 0.40 | 0.00 | 0.00 | 0.84 | 0.64 | 14 | Endothelial |
| Cxx1b | 0.32 | 0.00 | 0.00 | 0.47 | 0.22 | 14 | Endothelial |
| Cxx1c | 0.33 | 0.00 | 0.00 | 0.46 | 0.21 | 14 | Endothelial |
| Ptrf | 0.45 | 0.00 | 0.00 | 0.62 | 0.35 | 14 | Endothelial |
| Mapk3 | 0.37 | 0.00 | 0.00 | 0.33 | 0.13 | 14 | Endothelial |
| Frmd4a | 0.35 | 0.00 | 0.00 | 0.26 | 0.09 | 14 | Endothelial |
| Amotl1 | 0.37 | 0.00 | 0.00 | 0.37 | 0.15 | 14 | Endothelial |
| Ctnna1 | 0.41 | 0.00 | 0.00 | 0.63 | 0.36 | 14 | Endothelial |
| Hnrnpu | 0.42 | 0.00 | 0.00 | 0.83 | 0.63 | 14 | Endothelial |
| Baz1b | 0.48 | 0.00 | 0.00 | 0.51 | 0.26 | 14 | Endothelial |
| Srrt | 0.44 | 0.00 | 0.00 | 0.42 | 0.19 | 14 | Endothelial |
| Clic1 | 0.43 | 0.00 | 0.00 | 0.63 | 0.37 | 14 | Endothelial |
| Rell1 | 0.44 | 0.00 | 0.00 | 0.43 | 0.20 | 14 | Endothelial |
| Cd9 | 0.28 | 0.00 | 0.00 | 0.64 | 0.34 | 14 | Endothelial |
| Sptan1 | 0.44 | 0.00 | 0.00 | 0.37 | 0.16 | 14 | Endothelial |
| Itga1 | 0.34 | 0.00 | 0.00 | 0.26 | 0.09 | 14 | Endothelial |
| Kpnb1 | 0.43 | 0.00 | 0.00 | 0.40 | 0.18 | 14 | Endothelial |
| Guk1 | 0.42 | 0.00 | 0.00 | 0.35 | 0.15 | 14 | Endothelial |
| Hnrnpab | 0.50 | 0.00 | 0.00 | 0.57 | 0.32 | 14 | Endothelial |
| Smad7 | 0.33 | 0.00 | 0.00 | 0.29 | 0.11 | 14 | Endothelial |
| Ilf2 | 0.41 | 0.00 | 0.00 | 0.41 | 0.19 | 14 | Endothelial |
| Smc6 | 0.49 | 0.00 | 0.00 | 0.57 | 0.32 | 14 | Endothelial |
| Tpm3 | 0.37 | 0.00 | 0.00 | 0.88 | 0.69 | 14 | Endothelial |
| Kank3 | 0.34 | 0.00 | 0.00 | 0.34 | 0.14 | 14 | Endothelial |
| Scoc | 0.36 | 0.00 | 0.00 | 0.31 | 0.12 | 14 | Endothelial |
| Hnrnpa3 | 0.45 | 0.00 | 0.00 | 0.70 | 0.46 | 14 | Endothelial |
| Gm6793 | 0.45 | 0.00 | 0.00 | 0.53 | 0.29 | 14 | Endothelial |
| Cdk4 | 0.42 | 0.00 | 0.00 | 0.47 | 0.23 | 14 | Endothelial |
| Gkap1 | 0.38 | 0.00 | 0.00 | 0.30 | 0.12 | 14 | Endothelial |
| Add3 | 0.42 | 0.00 | 0.00 | 0.52 | 0.27 | 14 | Endothelial |
| Zfp608 | 0.36 | 0.00 | 0.00 | 0.33 | 0.14 | 14 | Endothelial |
| Tpst2 | 0.29 | 0.00 | 0.00 | 0.30 | 0.12 | 14 | Endothelial |
| Trib2 | 0.28 | 0.00 | 0.00 | 0.28 | 0.11 | 14 | Endothelial |
| Prex2 | 0.36 | 0.00 | 0.00 | 0.40 | 0.18 | 14 | Endothelial |
| Hnrnpf | 0.39 | 0.00 | 0.00 | 0.60 | 0.34 | 14 | Endothelial |
| Bnip2 | 0.42 | 0.00 | 0.00 | 0.55 | 0.29 | 14 | Endothelial |
| Fubp1 | 0.40 | 0.00 | 0.00 | 0.48 | 0.24 | 14 | Endothelial |
| Myo10 | 0.39 | 0.00 | 0.00 | 0.30 | 0.13 | 14 | Endothelial |
| Cd151 | 0.33 | 0.00 | 0.00 | 0.36 | 0.16 | 14 | Endothelial |
| Ablim1 | 0.42 | 0.00 | 0.00 | 0.48 | 0.24 | 14 | Endothelial |
| Snhg5 | 0.41 | 0.00 | 0.00 | 0.38 | 0.17 | 14 | Endothelial |
| Srsf2 | 0.48 | 0.00 | 0.00 | 0.63 | 0.38 | 14 | Endothelial |
| Dusp6 | 0.39 | 0.00 | 0.00 | 0.38 | 0.18 | 14 | Endothelial |
| Syne2 | 0.44 | 0.00 | 0.00 | 0.32 | 0.14 | 14 | Endothelial |
| Gbp7 | 0.35 | 0.00 | 0.00 | 0.33 | 0.14 | 14 | Endothelial |
| Ctcf | 0.44 | 0.00 | 0.00 | 0.45 | 0.23 | 14 | Endothelial |
| Jun | 0.27 | 0.00 | 0.00 | 0.64 | 0.36 | 14 | Endothelial |
| Rap1a | 0.38 | 0.00 | 0.00 | 0.55 | 0.30 | 14 | Endothelial |
| Plekha2 | 0.30 | 0.00 | 0.00 | 0.25 | 0.09 | 14 | Endothelial |
| Myo18a | 0.36 | 0.00 | 0.00 | 0.26 | 0.10 | 14 | Endothelial |
| Vps36 | 0.44 | 0.00 | 0.00 | 0.52 | 0.28 | 14 | Endothelial |
| Calm3 | 0.39 | 0.00 | 0.00 | 0.40 | 0.19 | 14 | Endothelial |
| Nfic | 0.37 | 0.00 | 0.00 | 0.47 | 0.24 | 14 | Endothelial |
| Gclm | 0.48 | 0.00 | 0.00 | 0.64 | 0.41 | 14 | Endothelial |
| Pds5b | 0.39 | 0.00 | 0.00 | 0.29 | 0.12 | 14 | Endothelial |
| H2-K1 | 0.44 | 0.00 | 0.00 | 0.56 | 0.32 | 14 | Endothelial |
| Rhoc | 0.32 | 0.00 | 0.00 | 0.42 | 0.20 | 14 | Endothelial |
| Ubb | 0.32 | 0.00 | 0.00 | 0.91 | 0.80 | 14 | Endothelial |
| Ccser2 | 0.38 | 0.00 | 0.00 | 0.33 | 0.14 | 14 | Endothelial |
| Gm5641 | 0.44 | 0.00 | 0.00 | 0.59 | 0.36 | 14 | Endothelial |
| Lyn | 0.27 | 0.00 | 0.00 | 0.31 | 0.13 | 14 | Endothelial |
| Rif1 | 0.38 | 0.00 | 0.00 | 0.35 | 0.16 | 14 | Endothelial |
| Smarca2 | 0.43 | 0.00 | 0.00 | 0.58 | 0.34 | 14 | Endothelial |
| Map7d1 | 0.35 | 0.00 | 0.00 | 0.38 | 0.18 | 14 | Endothelial |
| Dusp3 | 0.33 | 0.00 | 0.00 | 0.40 | 0.19 | 14 | Endothelial |
| Hspa14 | 0.32 | 0.00 | 0.00 | 0.26 | 0.10 | 14 | Endothelial |
| Ehd2 | 0.29 | 0.00 | 0.00 | 0.33 | 0.14 | 14 | Endothelial |
| Cd200 | 0.26 | 0.00 | 0.00 | 0.51 | 0.27 | 14 | Endothelial |
| Supt16 | 0.38 | 0.00 | 0.00 | 0.47 | 0.24 | 14 | Endothelial |
| Ccdc88a | 0.35 | 0.00 | 0.00 | 0.47 | 0.24 | 14 | Endothelial |
| Lgals9 | 0.29 | 0.00 | 0.00 | 0.32 | 0.14 | 14 | Endothelial |
| Gm12728 | 0.37 | 0.00 | 0.00 | 0.36 | 0.17 | 14 | Endothelial |
| Nmi | 0.28 | 0.00 | 0.00 | 0.27 | 0.11 | 14 | Endothelial |
| Mxra7 | 0.31 | 0.00 | 0.00 | 0.31 | 0.13 | 14 | Endothelial |
| Aph1a | 0.31 | 0.00 | 0.00 | 0.28 | 0.12 | 14 | Endothelial |
| Nsmce1 | 0.35 | 0.00 | 0.00 | 0.30 | 0.13 | 14 | Endothelial |
| Rbm3 | 0.42 | 0.00 | 0.00 | 0.63 | 0.40 | 14 | Endothelial |
| Dynll1 | 0.42 | 0.00 | 0.00 | 0.69 | 0.44 | 14 | Endothelial |
| Arl2bp | 0.34 | 0.00 | 0.00 | 0.30 | 0.13 | 14 | Endothelial |
| Elf2 | 0.37 | 0.00 | 0.00 | 0.37 | 0.17 | 14 | Endothelial |
| Ehd1 | 0.29 | 0.00 | 0.00 | 0.27 | 0.11 | 14 | Endothelial |
| Srsf4 | 0.38 | 0.00 | 0.00 | 0.29 | 0.13 | 14 | Endothelial |
| Vdac3 | 0.33 | 0.00 | 0.00 | 0.36 | 0.17 | 14 | Endothelial |
| Srsf7 | 0.36 | 0.00 | 0.00 | 0.33 | 0.15 | 14 | Endothelial |
| Xist | 0.46 | 0.00 | 0.00 | 0.74 | 0.50 | 14 | Endothelial |
| Rpa3 | 0.34 | 0.00 | 0.00 | 0.29 | 0.12 | 14 | Endothelial |
| Klf2 | 0.34 | 0.00 | 0.00 | 0.40 | 0.19 | 14 | Endothelial |
| Mrpl51 | 0.34 | 0.00 | 0.00 | 0.33 | 0.15 | 14 | Endothelial |
| Naa38 | 0.31 | 0.00 | 0.00 | 0.32 | 0.15 | 14 | Endothelial |
| Gm5160 | 0.42 | 0.00 | 0.00 | 0.41 | 0.21 | 14 | Endothelial |
| Acat1 | 0.38 | 0.00 | 0.00 | 0.37 | 0.18 | 14 | Endothelial |
| Ilf3 | 0.33 | 0.00 | 0.00 | 0.28 | 0.12 | 14 | Endothelial |
| 11-Sep | 0.33 | 0.00 | 0.00 | 0.44 | 0.22 | 14 | Endothelial |
| Ndufa8 | 0.36 | 0.00 | 0.00 | 0.46 | 0.25 | 14 | Endothelial |
| Fermt2 | 0.28 | 0.00 | 0.00 | 0.64 | 0.39 | 14 | Endothelial |
| Tmco1 | 0.36 | 0.00 | 0.00 | 0.52 | 0.29 | 14 | Endothelial |
| Cnot6 | 0.38 | 0.00 | 0.00 | 0.33 | 0.16 | 14 | Endothelial |
| Mapre1 | 0.37 | 0.00 | 0.00 | 0.51 | 0.29 | 14 | Endothelial |
| Larp7 | 0.35 | 0.00 | 0.00 | 0.35 | 0.17 | 14 | Endothelial |
| Smc3 | 0.41 | 0.00 | 0.00 | 0.60 | 0.37 | 14 | Endothelial |
| Itpripl2 | 0.32 | 0.00 | 0.00 | 0.35 | 0.17 | 14 | Endothelial |
| Nop58 | 0.38 | 0.00 | 0.00 | 0.47 | 0.26 | 14 | Endothelial |
| Rbbp4 | 0.36 | 0.00 | 0.00 | 0.39 | 0.20 | 14 | Endothelial |
| Galnt18 | 0.27 | 0.00 | 0.00 | 0.26 | 0.11 | 14 | Endothelial |
| Snrpb | 0.39 | 0.00 | 0.00 | 0.57 | 0.35 | 14 | Endothelial |
| Sh3glb1 | 0.36 | 0.00 | 0.00 | 0.63 | 0.39 | 14 | Endothelial |
| Nckap5 | 0.38 | 0.00 | 0.00 | 0.28 | 0.13 | 14 | Endothelial |
| Tjp2 | 0.30 | 0.00 | 0.00 | 0.34 | 0.16 | 14 | Endothelial |
| Acap2 | 0.28 | 0.00 | 0.00 | 0.39 | 0.19 | 14 | Endothelial |
| Vbp1 | 0.30 | 0.00 | 0.00 | 0.32 | 0.15 | 14 | Endothelial |
| Dot1l | 0.32 | 0.00 | 0.00 | 0.29 | 0.13 | 14 | Endothelial |
| G3bp1 | 0.30 | 0.00 | 0.00 | 0.35 | 0.17 | 14 | Endothelial |
| Zcrb1 | 0.38 | 0.00 | 0.00 | 0.50 | 0.28 | 14 | Endothelial |
| Mtf2 | 0.39 | 0.00 | 0.00 | 0.35 | 0.17 | 14 | Endothelial |
| Vdac3-ps1 | 0.27 | 0.00 | 0.00 | 0.32 | 0.15 | 14 | Endothelial |
| Rfc1 | 0.41 | 0.00 | 0.00 | 0.39 | 0.20 | 14 | Endothelial |
| Clic4 | 0.30 | 0.00 | 0.00 | 0.51 | 0.29 | 14 | Endothelial |
| Syncrip | 0.36 | 0.00 | 0.00 | 0.45 | 0.24 | 14 | Endothelial |
| Fam101b | 0.38 | 0.00 | 0.00 | 0.36 | 0.18 | 14 | Endothelial |
| Ccdc25 | 0.35 | 0.00 | 0.00 | 0.30 | 0.14 | 14 | Endothelial |
| Lsm5 | 0.29 | 0.00 | 0.00 | 0.27 | 0.12 | 14 | Endothelial |
| Dnaja1 | 0.37 | 0.00 | 0.00 | 0.72 | 0.51 | 14 | Endothelial |
| Gm10093 | 0.36 | 0.00 | 0.00 | 0.37 | 0.19 | 14 | Endothelial |
| Hmgn5 | 0.34 | 0.00 | 0.00 | 0.47 | 0.26 | 14 | Endothelial |
| Ssna1 | 0.29 | 0.00 | 0.00 | 0.29 | 0.13 | 14 | Endothelial |
| Rnps1 | 0.29 | 0.00 | 0.00 | 0.31 | 0.15 | 14 | Endothelial |
| 7-Sep | 0.30 | 0.00 | 0.00 | 0.78 | 0.58 | 14 | Endothelial |
| 2700029M09Rik | 0.30 | 0.00 | 0.00 | 0.28 | 0.12 | 14 | Endothelial |
| Rassf1 | 0.30 | 0.00 | 0.00 | 0.30 | 0.14 | 14 | Endothelial |
| Uchl5 | 0.26 | 0.00 | 0.00 | 0.26 | 0.11 | 14 | Endothelial |
| Ssb | 0.35 | 0.00 | 0.00 | 0.75 | 0.54 | 14 | Endothelial |
| Dynlt1f | 0.30 | 0.00 | 0.00 | 0.44 | 0.24 | 14 | Endothelial |
| Skil | 0.32 | 0.00 | 0.00 | 0.30 | 0.14 | 14 | Endothelial |
| Rnf7 | 0.29 | 0.00 | 0.00 | 0.44 | 0.24 | 14 | Endothelial |
| Map4k4 | 0.34 | 0.00 | 0.00 | 0.36 | 0.18 | 14 | Endothelial |
| Hdgf | 0.40 | 0.00 | 0.00 | 0.44 | 0.24 | 14 | Endothelial |
| Gm8991 | 0.37 | 0.00 | 0.00 | 0.41 | 0.22 | 14 | Endothelial |
| Anapc5 | 0.29 | 0.00 | 0.00 | 0.36 | 0.18 | 14 | Endothelial |
| Nap1l4 | 0.33 | 0.00 | 0.00 | 0.46 | 0.26 | 14 | Endothelial |
| Nudt21 | 0.28 | 0.00 | 0.00 | 0.27 | 0.12 | 14 | Endothelial |
| Serbp1 | 0.28 | 0.00 | 0.00 | 0.87 | 0.74 | 14 | Endothelial |
| Hnrnpd | 0.35 | 0.00 | 0.00 | 0.48 | 0.28 | 14 | Endothelial |
| Pa2g4 | 0.33 | 0.00 | 0.00 | 0.46 | 0.26 | 14 | Endothelial |
| Brd9 | 0.30 | 0.00 | 0.00 | 0.38 | 0.19 | 14 | Endothelial |
| Marcksl1 | 0.26 | 0.00 | 0.00 | 0.38 | 0.19 | 14 | Endothelial |
| Smchd1 | 0.31 | 0.00 | 0.00 | 0.32 | 0.16 | 14 | Endothelial |
| Mgll | 0.27 | 0.00 | 0.00 | 0.33 | 0.16 | 14 | Endothelial |
| Leprot | 0.29 | 0.00 | 0.00 | 0.55 | 0.32 | 14 | Endothelial |
| Gm9242 | 0.33 | 0.00 | 0.00 | 0.42 | 0.23 | 14 | Endothelial |
| 1700020I14Rik | 0.31 | 0.00 | 0.00 | 0.41 | 0.22 | 14 | Endothelial |
| Cyb5r3 | 0.34 | 0.00 | 0.00 | 0.60 | 0.38 | 14 | Endothelial |
| Rhoa | 0.33 | 0.00 | 0.00 | 0.66 | 0.46 | 14 | Endothelial |
| Tmem109 | 0.28 | 0.00 | 0.00 | 0.27 | 0.12 | 14 | Endothelial |
| Stag1 | 0.34 | 0.00 | 0.00 | 0.26 | 0.12 | 14 | Endothelial |
| Stk4 | 0.25 | 0.00 | 0.00 | 0.29 | 0.14 | 14 | Endothelial |
| Ywhae | 0.35 | 0.00 | 0.00 | 0.73 | 0.54 | 14 | Endothelial |
| Ctnnb1 | 0.33 | 0.00 | 0.00 | 0.56 | 0.34 | 14 | Endothelial |
| Fmr1 | 0.30 | 0.00 | 0.00 | 0.29 | 0.14 | 14 | Endothelial |
| Lyar | 0.26 | 0.00 | 0.00 | 0.31 | 0.15 | 14 | Endothelial |
| Hnrnpr | 0.31 | 0.00 | 0.00 | 0.44 | 0.24 | 14 | Endothelial |
| Cox7a2 | 0.34 | 0.00 | 0.00 | 0.68 | 0.48 | 14 | Endothelial |
| H3f3b | 0.30 | 0.00 | 0.00 | 0.95 | 0.89 | 14 | Endothelial |
| Ywhaq | 0.30 | 0.00 | 0.00 | 0.58 | 0.36 | 14 | Endothelial |
| Taf7 | 0.26 | 0.00 | 0.00 | 0.25 | 0.11 | 14 | Endothelial |
| Ywhab | 0.30 | 0.00 | 0.00 | 0.77 | 0.56 | 14 | Endothelial |
| Tardbp | 0.31 | 0.00 | 0.00 | 0.51 | 0.30 | 14 | Endothelial |
| Tead2 | 0.26 | 0.00 | 0.00 | 0.27 | 0.12 | 14 | Endothelial |
| Hdac1 | 0.34 | 0.00 | 0.00 | 0.36 | 0.19 | 14 | Endothelial |
| Eif4a3 | 0.28 | 0.00 | 0.00 | 0.36 | 0.19 | 14 | Endothelial |
| Dynlt1b | 0.27 | 0.00 | 0.00 | 0.44 | 0.25 | 14 | Endothelial |
| Srsf1 | 0.28 | 0.00 | 0.00 | 0.28 | 0.14 | 14 | Endothelial |
| Rab11a | 0.27 | 0.00 | 0.00 | 0.45 | 0.26 | 14 | Endothelial |
| Dynlt1c | 0.28 | 0.00 | 0.00 | 0.46 | 0.26 | 14 | Endothelial |
| Taf9 | 0.30 | 0.00 | 0.00 | 0.34 | 0.18 | 14 | Endothelial |
| Eif4g2 | 0.34 | 0.00 | 0.00 | 0.61 | 0.40 | 14 | Endothelial |
| Ccdc174 | 0.29 | 0.00 | 0.00 | 0.31 | 0.15 | 14 | Endothelial |
| Dram2 | 0.28 | 0.00 | 0.00 | 0.32 | 0.16 | 14 | Endothelial |
| B230219D22Rik | 0.32 | 0.00 | 0.00 | 0.47 | 0.27 | 14 | Endothelial |
| Sash1 | 0.25 | 0.00 | 0.00 | 0.27 | 0.13 | 14 | Endothelial |
| Pcmt1 | 0.25 | 0.00 | 0.00 | 0.36 | 0.19 | 14 | Endothelial |
| Dynlt1a | 0.27 | 0.00 | 0.00 | 0.40 | 0.22 | 14 | Endothelial |
| Gnb1 | 0.29 | 0.00 | 0.00 | 0.39 | 0.21 | 14 | Endothelial |
| Nono | 0.25 | 0.00 | 0.00 | 0.38 | 0.20 | 14 | Endothelial |
| Qk | 0.28 | 0.00 | 0.00 | 0.49 | 0.29 | 14 | Endothelial |
| U2af1 | 0.28 | 0.00 | 0.00 | 0.33 | 0.18 | 14 | Endothelial |
| Hist1h2bc | 0.52 | 0.00 | 0.00 | 0.27 | 0.13 | 14 | Endothelial |
| Atpif1 | 0.28 | 0.00 | 0.00 | 0.73 | 0.52 | 14 | Endothelial |
| Sumo2 | 0.31 | 0.00 | 0.00 | 0.50 | 0.30 | 14 | Endothelial |
| Gm9825 | 0.26 | 0.00 | 0.00 | 0.26 | 0.13 | 14 | Endothelial |
| Kat6a | 0.28 | 0.00 | 0.00 | 0.28 | 0.14 | 14 | Endothelial |
| Txn1 | 0.28 | 0.00 | 0.00 | 0.69 | 0.49 | 14 | Endothelial |
| Pdap1 | 0.32 | 0.00 | 0.00 | 0.65 | 0.45 | 14 | Endothelial |
| Mat2a | 0.30 | 0.00 | 0.00 | 0.46 | 0.27 | 14 | Endothelial |
| Dhx15 | 0.32 | 0.00 | 0.00 | 0.44 | 0.26 | 14 | Endothelial |
| Myh9 | 0.28 | 0.00 | 0.00 | 0.53 | 0.33 | 14 | Endothelial |
| Tnfaip1 | 0.26 | 0.00 | 0.00 | 0.31 | 0.16 | 14 | Endothelial |
| Dnm1l | 0.28 | 0.00 | 0.00 | 0.32 | 0.17 | 14 | Endothelial |
| Hint1 | 0.32 | 0.00 | 0.00 | 0.65 | 0.46 | 14 | Endothelial |
| Erh | 0.26 | 0.00 | 0.00 | 0.47 | 0.28 | 14 | Endothelial |
| Snrpe | 0.29 | 0.00 | 0.00 | 0.56 | 0.36 | 14 | Endothelial |
| Commd1 | 0.26 | 0.00 | 0.00 | 0.37 | 0.20 | 14 | Endothelial |
| Rock2 | 0.28 | 0.00 | 0.00 | 0.51 | 0.32 | 14 | Endothelial |
| Smarca4 | 0.27 | 0.00 | 0.00 | 0.32 | 0.17 | 14 | Endothelial |
| Hnrnpa0 | 0.25 | 0.00 | 0.00 | 0.38 | 0.21 | 14 | Endothelial |
| Fkbp3 | 0.28 | 0.00 | 0.00 | 0.49 | 0.30 | 14 | Endothelial |
| Hp1bp3 | 0.33 | 0.00 | 0.00 | 0.63 | 0.44 | 14 | Endothelial |
| Casp8ap2 | 0.34 | 0.00 | 0.00 | 0.32 | 0.17 | 14 | Endothelial |
| Bzw1 | 0.27 | 0.00 | 0.00 | 0.52 | 0.32 | 14 | Endothelial |
| Rbbp7 | 0.28 | 0.00 | 0.00 | 0.44 | 0.26 | 14 | Endothelial |
| Ing1 | 0.27 | 0.00 | 0.00 | 0.29 | 0.15 | 14 | Endothelial |
| Tra2b | 0.30 | 0.00 | 0.00 | 0.48 | 0.29 | 14 | Endothelial |
| Sfpq | 0.30 | 0.00 | 0.00 | 0.55 | 0.36 | 14 | Endothelial |
| Oxct1 | 0.25 | 0.00 | 0.00 | 0.28 | 0.15 | 14 | Endothelial |
| Wsb1 | 0.28 | 0.00 | 0.00 | 0.36 | 0.21 | 14 | Endothelial |
| B020018G12Rik | 0.25 | 0.00 | 0.00 | 0.42 | 0.25 | 14 | Endothelial |
| Slc6a6 | 0.29 | 0.00 | 0.00 | 0.42 | 0.25 | 14 | Endothelial |
| Hnrnpa1 | 0.26 | 0.00 | 0.00 | 0.43 | 0.26 | 14 | Endothelial |
| Kras | 0.27 | 0.00 | 0.00 | 0.33 | 0.18 | 14 | Endothelial |
| Thrap3 | 0.28 | 0.00 | 0.00 | 0.56 | 0.37 | 14 | Endothelial |
| Kif5b | 0.25 | 0.00 | 0.00 | 0.67 | 0.47 | 14 | Endothelial |
| Tsix | 0.30 | 0.00 | 0.00 | 0.32 | 0.18 | 14 | Endothelial |
| Zfp326 | 0.26 | 0.00 | 0.00 | 0.39 | 0.23 | 14 | Endothelial |
| Hnrnpdl | 0.26 | 0.00 | 0.00 | 0.56 | 0.36 | 14 | Endothelial |
| Zc3h7a | 0.26 | 0.00 | 0.00 | 0.45 | 0.28 | 14 | Endothelial |
| Nudt4 | 0.26 | 0.00 | 0.00 | 0.30 | 0.17 | 14 | Endothelial |
| Fus | 0.26 | 0.00 | 0.00 | 0.66 | 0.48 | 14 | Endothelial |
| Cep110 | 0.26 | 0.00 | 0.00 | 0.32 | 0.18 | 14 | Endothelial |
| Tcf21 | 2.15 | 0.00 | 0.00 | 0.93 | 0.15 | 15 | Matrix Fibroblast |
| Inmt | 2.06 | 0.00 | 0.00 | 0.56 | 0.04 | 15 | Matrix Fibroblast |
| Gsn | 1.80 | 0.00 | 0.00 | 0.87 | 0.23 | 15 | Matrix Fibroblast |
| Ifitm1 | 1.77 | 0.00 | 0.00 | 0.83 | 0.12 | 15 | Matrix Fibroblast |
| Mfap4 | 1.70 | 0.00 | 0.00 | 0.96 | 0.23 | 15 | Matrix Fibroblast |
| Ppp1r14a | 1.69 | 0.00 | 0.00 | 0.55 | 0.07 | 15 | Matrix Fibroblast |
| Fmo2 | 1.69 | 0.00 | 0.00 | 0.83 | 0.15 | 15 | Matrix Fibroblast |
| Gyg | 1.65 | 0.00 | 0.00 | 0.78 | 0.18 | 15 | Matrix Fibroblast |
| Plac9a | 1.62 | 0.00 | 0.00 | 0.90 | 0.29 | 15 | Matrix Fibroblast |
| Plac9b | 1.61 | 0.00 | 0.00 | 0.89 | 0.29 | 15 | Matrix Fibroblast |
| Gpx3 | 1.59 | 0.00 | 0.00 | 0.70 | 0.12 | 15 | Matrix Fibroblast |
| Adh1 | 1.55 | 0.00 | 0.00 | 0.82 | 0.15 | 15 | Matrix Fibroblast |
| Clec3b | 1.54 | 0.00 | 0.00 | 0.48 | 0.06 | 15 | Matrix Fibroblast |
| Col13a1 | 1.52 | 0.00 | 0.00 | 0.54 | 0.09 | 15 | Matrix Fibroblast |
| Nexn | 1.52 | 0.00 | 0.00 | 0.70 | 0.21 | 15 | Matrix Fibroblast |
| Olfml3 | 1.49 | 0.00 | 0.00 | 0.48 | 0.08 | 15 | Matrix Fibroblast |
| Mgp | 1.49 | 0.00 | 0.00 | 0.98 | 0.30 | 15 | Matrix Fibroblast |
| G0s2 | 1.46 | 0.00 | 0.00 | 0.57 | 0.12 | 15 | Matrix Fibroblast |
| Fhl1 | 1.45 | 0.00 | 0.00 | 0.86 | 0.28 | 15 | Matrix Fibroblast |
| Hba-a2 | 1.42 | 0.00 | 0.00 | 0.97 | 0.69 | 15 | Matrix Fibroblast |
| Lbh | 1.41 | 0.00 | 0.00 | 0.68 | 0.18 | 15 | Matrix Fibroblast |
| Hba-a1 | 1.40 | 0.00 | 0.00 | 0.98 | 0.70 | 15 | Matrix Fibroblast |
| Maf | 1.38 | 0.00 | 0.00 | 0.69 | 0.18 | 15 | Matrix Fibroblast |
| Hsd11b1 | 1.34 | 0.00 | 0.00 | 0.49 | 0.08 | 15 | Matrix Fibroblast |
| Limch1 | 1.32 | 0.00 | 0.00 | 0.90 | 0.29 | 15 | Matrix Fibroblast |
| Rbp1 | 1.26 | 0.00 | 0.00 | 0.65 | 0.18 | 15 | Matrix Fibroblast |
| Hbb-bt | 1.21 | 0.00 | 0.00 | 0.98 | 0.71 | 15 | Matrix Fibroblast |
| Hbb-bs | 1.21 | 0.00 | 0.00 | 0.99 | 0.80 | 15 | Matrix Fibroblast |
| Dpep1 | 1.17 | 0.00 | 0.00 | 0.37 | 0.06 | 15 | Matrix Fibroblast |
| Phlda1 | 1.48 | 0.00 | 0.00 | 0.63 | 0.19 | 15 | Matrix Fibroblast |
| Gm3940 | 1.32 | 0.00 | 0.00 | 0.78 | 0.35 | 15 | Matrix Fibroblast |
| Fn1 | 1.14 | 0.00 | 0.00 | 0.70 | 0.22 | 15 | Matrix Fibroblast |
| Serping1 | 1.24 | 0.00 | 0.00 | 0.48 | 0.11 | 15 | Matrix Fibroblast |
| Gm10923 | 1.37 | 0.00 | 0.00 | 0.61 | 0.21 | 15 | Matrix Fibroblast |
| 6030408B16Rik | 1.05 | 0.00 | 0.00 | 0.28 | 0.04 | 15 | Matrix Fibroblast |
| Apoe | 0.72 | 0.00 | 0.00 | 0.74 | 0.27 | 15 | Matrix Fibroblast |
| Gm10132 | 0.93 | 0.00 | 0.00 | 0.94 | 0.72 | 15 | Matrix Fibroblast |
| Gm10335 | 0.94 | 0.00 | 0.00 | 0.95 | 0.73 | 15 | Matrix Fibroblast |
| Rpl23a | 1.00 | 0.00 | 0.00 | 0.92 | 0.66 | 15 | Matrix Fibroblast |
| Rpl23a-ps3 | 0.91 | 0.00 | 0.00 | 0.94 | 0.74 | 15 | Matrix Fibroblast |
| mt-Td | 1.27 | 0.00 | 0.00 | 0.52 | 0.16 | 15 | Matrix Fibroblast |
| Col1a1 | 0.85 | 0.00 | 0.00 | 0.71 | 0.25 | 15 | Matrix Fibroblast |
| Atp1a2 | 1.20 | 0.00 | 0.00 | 0.40 | 0.09 | 15 | Matrix Fibroblast |
| Rps4x | 0.77 | 0.00 | 0.00 | 0.98 | 0.86 | 15 | Matrix Fibroblast |
| Cdh11 | 0.99 | 0.00 | 0.00 | 0.59 | 0.18 | 15 | Matrix Fibroblast |
| Pcolce2 | 1.14 | 0.00 | 0.00 | 0.38 | 0.09 | 15 | Matrix Fibroblast |
| Rpl10 | 0.90 | 0.00 | 0.00 | 0.93 | 0.71 | 15 | Matrix Fibroblast |
| Gm22758 | 0.88 | 0.00 | 0.00 | 0.93 | 0.68 | 15 | Matrix Fibroblast |
| RPL24 | 1.05 | 0.00 | 0.00 | 0.80 | 0.45 | 15 | Matrix Fibroblast |
| Gm7676 | 1.05 | 0.00 | 0.00 | 0.54 | 0.18 | 15 | Matrix Fibroblast |
| Rps27 | 0.74 | 0.00 | 0.00 | 0.97 | 0.81 | 15 | Matrix Fibroblast |
| Ogn | 1.01 | 0.00 | 0.00 | 0.46 | 0.13 | 15 | Matrix Fibroblast |
| Col1a2 | 0.47 | 0.00 | 0.00 | 0.92 | 0.38 | 15 | Matrix Fibroblast |
| Gm22759 | 0.87 | 0.00 | 0.00 | 0.89 | 0.64 | 15 | Matrix Fibroblast |
| Selenbp1 | 0.97 | 0.00 | 0.00 | 0.59 | 0.22 | 15 | Matrix Fibroblast |
| Rpl21 | 0.76 | 0.00 | 0.00 | 0.94 | 0.74 | 15 | Matrix Fibroblast |
| Rpl10-ps3 | 0.83 | 0.00 | 0.00 | 0.92 | 0.72 | 15 | Matrix Fibroblast |
| Col6a1 | 1.09 | 0.00 | 0.00 | 0.52 | 0.18 | 15 | Matrix Fibroblast |
| Nr2f2 | 1.20 | 0.00 | 0.00 | 0.54 | 0.20 | 15 | Matrix Fibroblast |
| Nap1l1 | 1.14 | 0.00 | 0.00 | 0.76 | 0.46 | 15 | Matrix Fibroblast |
| Rps8 | 0.71 | 0.00 | 0.00 | 0.96 | 0.81 | 15 | Matrix Fibroblast |
| 4-Sep | 0.98 | 0.00 | 0.00 | 0.59 | 0.22 | 15 | Matrix Fibroblast |
| Selenbp2 | 0.99 | 0.00 | 0.00 | 0.49 | 0.16 | 15 | Matrix Fibroblast |
| Gm26300 | 1.03 | 0.00 | 0.00 | 0.70 | 0.37 | 15 | Matrix Fibroblast |
| Meox2 | 0.99 | 0.00 | 0.00 | 0.32 | 0.07 | 15 | Matrix Fibroblast |
| Rpl37a | 0.67 | 0.00 | 0.00 | 0.95 | 0.83 | 15 | Matrix Fibroblast |
| Gm9846 | 0.59 | 0.00 | 0.00 | 0.98 | 0.90 | 15 | Matrix Fibroblast |
| Rps28 | 0.67 | 0.00 | 0.00 | 0.97 | 0.83 | 15 | Matrix Fibroblast |
| Fxyd1 | 0.99 | 0.00 | 0.00 | 0.33 | 0.09 | 15 | Matrix Fibroblast |
| Npnt | 0.83 | 0.00 | 0.00 | 0.60 | 0.23 | 15 | Matrix Fibroblast |
| Bgn | 0.93 | 0.00 | 0.00 | 0.58 | 0.23 | 15 | Matrix Fibroblast |
| Tns1 | 1.09 | 0.00 | 0.00 | 0.55 | 0.23 | 15 | Matrix Fibroblast |
| Rps29 | 0.57 | 0.00 | 0.00 | 0.98 | 0.90 | 15 | Matrix Fibroblast |
| Gm4204 | 1.03 | 0.00 | 0.00 | 0.76 | 0.50 | 15 | Matrix Fibroblast |
| Echdc2 | 0.93 | 0.00 | 0.00 | 0.26 | 0.06 | 15 | Matrix Fibroblast |
| Tmem176a | 1.09 | 0.00 | 0.00 | 0.50 | 0.20 | 15 | Matrix Fibroblast |
| Fblim1 | 1.02 | 0.00 | 0.00 | 0.45 | 0.16 | 15 | Matrix Fibroblast |
| Gm5428 | 0.66 | 0.00 | 0.00 | 0.93 | 0.79 | 15 | Matrix Fibroblast |
| Macf1 | 0.70 | 0.00 | 0.00 | 0.83 | 0.52 | 15 | Matrix Fibroblast |
| Rpl36-ps3 | 0.86 | 0.00 | 0.00 | 0.78 | 0.53 | 15 | Matrix Fibroblast |
| Rps17 | 0.67 | 0.00 | 0.00 | 0.93 | 0.77 | 15 | Matrix Fibroblast |
| Peg3 | 0.90 | 0.00 | 0.00 | 0.63 | 0.29 | 15 | Matrix Fibroblast |
| Rpl37 | 0.65 | 0.00 | 0.00 | 0.93 | 0.80 | 15 | Matrix Fibroblast |
| Rpl13-ps3 | 0.66 | 0.00 | 0.00 | 0.92 | 0.75 | 15 | Matrix Fibroblast |
| Rps23 | 0.62 | 0.00 | 0.00 | 0.93 | 0.79 | 15 | Matrix Fibroblast |
| Rps16 | 0.61 | 0.00 | 0.00 | 0.95 | 0.84 | 15 | Matrix Fibroblast |
| Dpt | 0.43 | 0.00 | 0.00 | 0.50 | 0.17 | 15 | Matrix Fibroblast |
| Rpl13 | 0.62 | 0.00 | 0.00 | 0.93 | 0.79 | 15 | Matrix Fibroblast |
| Rps15a | 0.59 | 0.00 | 0.00 | 0.94 | 0.81 | 15 | Matrix Fibroblast |
| Rpl31 | 0.76 | 0.00 | 0.00 | 0.83 | 0.58 | 15 | Matrix Fibroblast |
| Rpl35a | 0.60 | 0.00 | 0.00 | 0.92 | 0.79 | 15 | Matrix Fibroblast |
| Ndrg2 | 0.84 | 0.00 | 0.00 | 0.28 | 0.07 | 15 | Matrix Fibroblast |
| Cdo1 | 0.93 | 0.00 | 0.00 | 0.26 | 0.06 | 15 | Matrix Fibroblast |
| Mir682 | 0.54 | 0.00 | 0.00 | 0.95 | 0.85 | 15 | Matrix Fibroblast |
| Gas5 | 0.85 | 0.00 | 0.00 | 0.76 | 0.53 | 15 | Matrix Fibroblast |
| Rpl6 | 0.52 | 0.00 | 0.00 | 0.96 | 0.87 | 15 | Matrix Fibroblast |
| Ifitm2 | 0.86 | 0.00 | 0.00 | 0.61 | 0.31 | 15 | Matrix Fibroblast |
| Rpl10a-ps2 | 1.02 | 0.00 | 0.00 | 0.55 | 0.28 | 15 | Matrix Fibroblast |
| Col6a2 | 0.96 | 0.00 | 0.00 | 0.39 | 0.14 | 15 | Matrix Fibroblast |
| Plxdc2 | 0.81 | 0.00 | 0.00 | 0.37 | 0.12 | 15 | Matrix Fibroblast |
| Cp | 0.89 | 0.00 | 0.00 | 0.36 | 0.12 | 15 | Matrix Fibroblast |
| Rpl39 | 0.55 | 0.00 | 0.00 | 0.94 | 0.80 | 15 | Matrix Fibroblast |
| Gm5786 | 0.87 | 0.00 | 0.00 | 0.64 | 0.39 | 15 | Matrix Fibroblast |
| Rps2-ps10 | 0.82 | 0.00 | 0.00 | 0.68 | 0.43 | 15 | Matrix Fibroblast |
| Gm24276 | 0.97 | 0.00 | 0.00 | 0.47 | 0.22 | 15 | Matrix Fibroblast |
| Eln | 0.66 | 0.00 | 0.00 | 0.42 | 0.15 | 15 | Matrix Fibroblast |
| Rpl9 | 0.71 | 0.00 | 0.00 | 0.81 | 0.62 | 15 | Matrix Fibroblast |
| Enpep | 0.89 | 0.00 | 0.00 | 0.31 | 0.10 | 15 | Matrix Fibroblast |
| Rpl36 | 0.68 | 0.00 | 0.00 | 0.82 | 0.63 | 15 | Matrix Fibroblast |
| Tbx2 | 0.88 | 0.00 | 0.00 | 0.34 | 0.11 | 15 | Matrix Fibroblast |
| Gpc3 | 0.73 | 0.00 | 0.00 | 0.42 | 0.16 | 15 | Matrix Fibroblast |
| Rpl21-ps6 | 0.83 | 0.00 | 0.00 | 0.67 | 0.43 | 15 | Matrix Fibroblast |
| Rps25 | 0.53 | 0.00 | 0.00 | 0.92 | 0.80 | 15 | Matrix Fibroblast |
| Gm10076 | 0.43 | 0.00 | 0.00 | 0.98 | 0.92 | 15 | Matrix Fibroblast |
| Palld | 0.82 | 0.00 | 0.00 | 0.42 | 0.17 | 15 | Matrix Fibroblast |
| Gm25593 | 0.92 | 0.00 | 0.00 | 0.44 | 0.21 | 15 | Matrix Fibroblast |
| Ptms | 0.77 | 0.00 | 0.00 | 0.64 | 0.39 | 15 | Matrix Fibroblast |
| Rpl38 | 0.55 | 0.00 | 0.00 | 0.89 | 0.73 | 15 | Matrix Fibroblast |
| Rpl36a | 0.65 | 0.00 | 0.00 | 0.80 | 0.62 | 15 | Matrix Fibroblast |
| Lox | 0.71 | 0.00 | 0.00 | 0.36 | 0.13 | 15 | Matrix Fibroblast |
| Zyx | 0.89 | 0.00 | 0.00 | 0.43 | 0.19 | 15 | Matrix Fibroblast |
| Gm15013 | 0.81 | 0.00 | 0.00 | 0.60 | 0.36 | 15 | Matrix Fibroblast |
| Rps27a | 0.51 | 0.00 | 0.00 | 0.91 | 0.76 | 15 | Matrix Fibroblast |
| Rps2-ps6 | 0.79 | 0.00 | 0.00 | 0.63 | 0.40 | 15 | Matrix Fibroblast |
| Rps3a3 | 0.51 | 0.00 | 0.00 | 0.91 | 0.77 | 15 | Matrix Fibroblast |
| Rpl34 | 0.50 | 0.00 | 0.00 | 0.91 | 0.80 | 15 | Matrix Fibroblast |
| Gm7808 | 0.64 | 0.00 | 0.00 | 0.78 | 0.59 | 15 | Matrix Fibroblast |
| Cdkn1c | 0.75 | 0.00 | 0.00 | 0.42 | 0.18 | 15 | Matrix Fibroblast |
| Cd81 | 0.75 | 0.00 | 0.00 | 0.64 | 0.40 | 15 | Matrix Fibroblast |
| Rpl18a | 0.48 | 0.00 | 0.00 | 0.93 | 0.82 | 15 | Matrix Fibroblast |
| Loxl1 | 0.74 | 0.00 | 0.00 | 0.30 | 0.10 | 15 | Matrix Fibroblast |
| Gm6133 | 0.79 | 0.00 | 0.00 | 0.57 | 0.34 | 15 | Matrix Fibroblast |
| Vldlr | 0.91 | 0.00 | 0.00 | 0.30 | 0.11 | 15 | Matrix Fibroblast |
| Zfp36l1 | 0.83 | 0.00 | 0.00 | 0.66 | 0.45 | 15 | Matrix Fibroblast |
| Rps14 | 0.41 | 0.00 | 0.00 | 0.98 | 0.92 | 15 | Matrix Fibroblast |
| Rpl11 | 0.57 | 0.00 | 0.00 | 0.83 | 0.68 | 15 | Matrix Fibroblast |
| Mxra8 | 0.77 | 0.00 | 0.00 | 0.38 | 0.16 | 15 | Matrix Fibroblast |
| Gm5239 | 0.64 | 0.00 | 0.00 | 0.72 | 0.51 | 15 | Matrix Fibroblast |
| Gm6139 | 0.75 | 0.00 | 0.00 | 0.63 | 0.42 | 15 | Matrix Fibroblast |
| Rpl27a | 0.49 | 0.00 | 0.00 | 0.91 | 0.78 | 15 | Matrix Fibroblast |
| Gm11808 | 0.50 | 0.00 | 0.00 | 0.89 | 0.77 | 15 | Matrix Fibroblast |
| Gm26384 | 0.67 | 0.00 | 0.00 | 0.69 | 0.48 | 15 | Matrix Fibroblast |
| Rps26-ps1 | 0.41 | 0.00 | 0.00 | 0.95 | 0.87 | 15 | Matrix Fibroblast |
| Rps13 | 0.58 | 0.00 | 0.00 | 0.80 | 0.64 | 15 | Matrix Fibroblast |
| Rpl26 | 0.47 | 0.00 | 0.00 | 0.91 | 0.81 | 15 | Matrix Fibroblast |
| Rpl23 | 0.41 | 0.00 | 0.00 | 0.94 | 0.87 | 15 | Matrix Fibroblast |
| Gm22426 | 0.65 | 0.00 | 0.00 | 0.69 | 0.48 | 15 | Matrix Fibroblast |
| Rps18 | 0.55 | 0.00 | 0.00 | 0.80 | 0.65 | 15 | Matrix Fibroblast |
| Gm6576 | 0.80 | 0.00 | 0.00 | 0.52 | 0.31 | 15 | Matrix Fibroblast |
| Gm10263 | 0.47 | 0.00 | 0.00 | 0.90 | 0.76 | 15 | Matrix Fibroblast |
| Mylk | 0.63 | 0.00 | 0.00 | 0.52 | 0.27 | 15 | Matrix Fibroblast |
| Gm9493 | 0.50 | 0.00 | 0.00 | 0.85 | 0.74 | 15 | Matrix Fibroblast |
| Rps26 | 0.37 | 0.00 | 0.00 | 0.97 | 0.90 | 15 | Matrix Fibroblast |
| Gm5093 | 0.62 | 0.00 | 0.00 | 0.65 | 0.46 | 15 | Matrix Fibroblast |
| Sh3bgrl | 0.64 | 0.00 | 0.00 | 0.65 | 0.44 | 15 | Matrix Fibroblast |
| Itga8 | 0.68 | 0.00 | 0.00 | 0.26 | 0.09 | 15 | Matrix Fibroblast |
| Rps7 | 0.49 | 0.00 | 0.00 | 0.84 | 0.74 | 15 | Matrix Fibroblast |
| Rpl9-ps6 | 0.50 | 0.00 | 0.00 | 0.84 | 0.73 | 15 | Matrix Fibroblast |
| 0610007N19Rik | 0.81 | 0.00 | 0.00 | 0.33 | 0.14 | 15 | Matrix Fibroblast |
| Tmem254a | 0.89 | 0.00 | 0.00 | 0.38 | 0.19 | 15 | Matrix Fibroblast |
| Rpl27 | 0.50 | 0.00 | 0.00 | 0.84 | 0.69 | 15 | Matrix Fibroblast |
| Gm5244 | 0.78 | 0.00 | 0.00 | 0.47 | 0.28 | 15 | Matrix Fibroblast |
| Rpl28 | 0.63 | 0.00 | 0.00 | 0.69 | 0.53 | 15 | Matrix Fibroblast |
| Rps12 | 0.48 | 0.00 | 0.00 | 0.84 | 0.68 | 15 | Matrix Fibroblast |
| Gm10709 | 0.79 | 0.00 | 0.00 | 0.46 | 0.27 | 15 | Matrix Fibroblast |
| Tmem254b | 0.89 | 0.00 | 0.00 | 0.38 | 0.19 | 15 | Matrix Fibroblast |
| Camk2n1 | 0.85 | 0.00 | 0.00 | 0.32 | 0.14 | 15 | Matrix Fibroblast |
| Rpl41 | 0.32 | 0.00 | 0.00 | 0.98 | 0.95 | 15 | Matrix Fibroblast |
| Rpl35 | 0.43 | 0.00 | 0.00 | 0.89 | 0.80 | 15 | Matrix Fibroblast |
| Nenf | 0.75 | 0.00 | 0.00 | 0.37 | 0.18 | 15 | Matrix Fibroblast |
| Rps18-ps3 | 0.56 | 0.00 | 0.00 | 0.71 | 0.56 | 15 | Matrix Fibroblast |
| Ifitm3 | 0.58 | 0.00 | 0.00 | 0.61 | 0.39 | 15 | Matrix Fibroblast |
| Csrp1 | 0.83 | 0.00 | 0.00 | 0.34 | 0.16 | 15 | Matrix Fibroblast |
| Eif4ebp1 | 0.82 | 0.00 | 0.00 | 0.30 | 0.13 | 15 | Matrix Fibroblast |
| Rpl21-ps4 | 0.67 | 0.00 | 0.00 | 0.53 | 0.35 | 15 | Matrix Fibroblast |
| Rpl3 | 0.52 | 0.00 | 0.00 | 0.75 | 0.63 | 15 | Matrix Fibroblast |
| Rps3a2 | 0.37 | 0.00 | 0.00 | 0.93 | 0.82 | 15 | Matrix Fibroblast |
| Rpl17 | 0.55 | 0.00 | 0.00 | 0.70 | 0.53 | 15 | Matrix Fibroblast |
| Cox6c | 0.52 | 0.00 | 0.00 | 0.76 | 0.65 | 15 | Matrix Fibroblast |
| Gm7589 | 0.47 | 0.00 | 0.00 | 0.78 | 0.65 | 15 | Matrix Fibroblast |
| Rps19 | 0.34 | 0.00 | 0.00 | 0.93 | 0.82 | 15 | Matrix Fibroblast |
| Snai2 | 0.75 | 0.00 | 0.00 | 0.28 | 0.12 | 15 | Matrix Fibroblast |
| Tmem254c | 0.82 | 0.00 | 0.00 | 0.35 | 0.18 | 15 | Matrix Fibroblast |
| Gm8730 | 0.37 | 0.00 | 0.00 | 0.90 | 0.82 | 15 | Matrix Fibroblast |
| Socs2 | 0.69 | 0.00 | 0.00 | 0.52 | 0.33 | 15 | Matrix Fibroblast |
| Gm10036 | 0.42 | 0.00 | 0.00 | 0.84 | 0.74 | 15 | Matrix Fibroblast |
| Ech1 | 0.79 | 0.00 | 0.00 | 0.31 | 0.15 | 15 | Matrix Fibroblast |
| Rap2a | 0.78 | 0.00 | 0.00 | 0.32 | 0.15 | 15 | Matrix Fibroblast |
| Rplp0 | 0.35 | 0.00 | 0.00 | 0.91 | 0.83 | 15 | Matrix Fibroblast |
| Gm6472 | 0.38 | 0.00 | 0.00 | 0.87 | 0.80 | 15 | Matrix Fibroblast |
| Akap12 | 0.63 | 0.00 | 0.00 | 0.43 | 0.24 | 15 | Matrix Fibroblast |
| Tpm1 | 0.39 | 0.00 | 0.00 | 0.71 | 0.51 | 15 | Matrix Fibroblast |
| Gm24865 | 0.45 | 0.00 | 0.00 | 0.75 | 0.62 | 15 | Matrix Fibroblast |
| Gm7536 | 0.36 | 0.00 | 0.00 | 0.91 | 0.82 | 15 | Matrix Fibroblast |
| Rps21 | 0.37 | 0.00 | 0.00 | 0.86 | 0.75 | 15 | Matrix Fibroblast |
| mt-Co1 | 0.29 | 0.00 | 0.00 | 0.98 | 0.95 | 15 | Matrix Fibroblast |
| Rplp2 | 0.34 | 0.00 | 0.00 | 0.89 | 0.81 | 15 | Matrix Fibroblast |
| Rpl13a-ps1 | 0.45 | 0.00 | 0.00 | 0.69 | 0.55 | 15 | Matrix Fibroblast |
| Rpl15 | 0.46 | 0.00 | 0.00 | 0.70 | 0.58 | 15 | Matrix Fibroblast |
| Rps3 | 0.34 | 0.00 | 0.00 | 0.90 | 0.83 | 15 | Matrix Fibroblast |
| Gm2000 | 0.35 | 0.00 | 0.00 | 0.90 | 0.84 | 15 | Matrix Fibroblast |
| Rpl10a | 0.39 | 0.00 | 0.00 | 0.82 | 0.74 | 15 | Matrix Fibroblast |
| Tmem176b | 0.66 | 0.00 | 0.00 | 0.49 | 0.34 | 15 | Matrix Fibroblast |
| Limd1 | 0.73 | 0.00 | 0.00 | 0.32 | 0.18 | 15 | Matrix Fibroblast |
| Rpl19 | 0.37 | 0.00 | 0.00 | 0.84 | 0.75 | 15 | Matrix Fibroblast |
| Rps2 | 0.41 | 0.00 | 0.00 | 0.77 | 0.67 | 15 | Matrix Fibroblast |
| Rpl30 | 0.47 | 0.00 | 0.00 | 0.69 | 0.59 | 15 | Matrix Fibroblast |
| Eva1b | 0.64 | 0.00 | 0.00 | 0.35 | 0.20 | 15 | Matrix Fibroblast |
| Gm10288 | 0.34 | 0.00 | 0.00 | 0.86 | 0.78 | 15 | Matrix Fibroblast |
| Rpl27-ps3 | 0.35 | 0.00 | 0.00 | 0.85 | 0.76 | 15 | Matrix Fibroblast |
| Rarres2 | 0.56 | 0.00 | 0.00 | 0.27 | 0.13 | 15 | Matrix Fibroblast |
| Gm17511 | 0.40 | 0.00 | 0.00 | 0.73 | 0.61 | 15 | Matrix Fibroblast |
| Rpl14 | 0.31 | 0.00 | 0.00 | 0.90 | 0.84 | 15 | Matrix Fibroblast |
| Cnn2 | 0.58 | 0.00 | 0.00 | 0.53 | 0.39 | 15 | Matrix Fibroblast |
| Ppap2b | 0.62 | 0.00 | 0.00 | 0.39 | 0.24 | 15 | Matrix Fibroblast |
| Fibin | 0.53 | 0.00 | 0.00 | 0.26 | 0.12 | 15 | Matrix Fibroblast |
| Ypel3 | 0.60 | 0.00 | 0.00 | 0.35 | 0.20 | 15 | Matrix Fibroblast |
| Rps3a1 | 0.27 | 0.00 | 0.00 | 0.93 | 0.85 | 15 | Matrix Fibroblast |
| Gm17430 | 0.72 | 0.00 | 0.00 | 0.26 | 0.14 | 15 | Matrix Fibroblast |
| Rps12-ps3 | 0.31 | 0.00 | 0.00 | 0.88 | 0.79 | 15 | Matrix Fibroblast |
| Nupr1 | 0.49 | 0.00 | 0.00 | 0.33 | 0.18 | 15 | Matrix Fibroblast |
| Rpl7a | 0.57 | 0.00 | 0.00 | 0.51 | 0.39 | 15 | Matrix Fibroblast |
| Rpl7a-ps10 | 0.61 | 0.00 | 0.00 | 0.39 | 0.26 | 15 | Matrix Fibroblast |
| Rps15 | 0.33 | 0.00 | 0.00 | 0.84 | 0.78 | 15 | Matrix Fibroblast |
| Gm16519 | 0.62 | 0.00 | 0.00 | 0.27 | 0.15 | 15 | Matrix Fibroblast |
| Rpl32 | 0.27 | 0.00 | 0.00 | 0.91 | 0.84 | 15 | Matrix Fibroblast |
| Sec61g | 0.53 | 0.00 | 0.00 | 0.47 | 0.35 | 15 | Matrix Fibroblast |
| Nfib | 0.45 | 0.00 | 0.00 | 0.62 | 0.47 | 15 | Matrix Fibroblast |
| 2410006H16Rik | 0.63 | 0.00 | 0.00 | 0.39 | 0.26 | 15 | Matrix Fibroblast |
| Plin2 | 0.56 | 0.00 | 0.00 | 0.25 | 0.13 | 15 | Matrix Fibroblast |
| Gm10260 | 0.43 | 0.00 | 0.00 | 0.59 | 0.49 | 15 | Matrix Fibroblast |
| Hmgb1 | 0.38 | 0.00 | 0.00 | 0.71 | 0.61 | 15 | Matrix Fibroblast |
| Gm26669 | 0.71 | 0.00 | 0.00 | 0.35 | 0.22 | 15 | Matrix Fibroblast |
| Maged2 | 0.54 | 0.00 | 0.00 | 0.29 | 0.17 | 15 | Matrix Fibroblast |
| Prdx5 | 0.43 | 0.00 | 0.00 | 0.45 | 0.32 | 15 | Matrix Fibroblast |
| Rpl7a-ps5 | 0.50 | 0.00 | 0.00 | 0.47 | 0.37 | 15 | Matrix Fibroblast |
| Hadh | 0.60 | 0.00 | 0.00 | 0.28 | 0.17 | 15 | Matrix Fibroblast |
| Pbx1 | 0.63 | 0.00 | 0.00 | 0.31 | 0.20 | 15 | Matrix Fibroblast |
| Emd | 0.48 | 0.00 | 0.00 | 0.49 | 0.40 | 15 | Matrix Fibroblast |
| Rpl29 | 0.46 | 0.00 | 0.00 | 0.54 | 0.46 | 15 | Matrix Fibroblast |
| Gm20091 | 0.53 | 0.00 | 0.00 | 0.33 | 0.22 | 15 | Matrix Fibroblast |
| Gnas | 0.31 | 0.00 | 0.00 | 0.75 | 0.67 | 15 | Matrix Fibroblast |
| Cpm | 0.50 | 0.00 | 0.00 | 0.30 | 0.18 | 15 | Matrix Fibroblast |
| Rps10 | 0.29 | 0.00 | 0.00 | 0.76 | 0.70 | 15 | Matrix Fibroblast |
| Pdlim2 | 0.52 | 0.00 | 0.00 | 0.28 | 0.18 | 15 | Matrix Fibroblast |
| Eif3e | 0.52 | 0.00 | 0.00 | 0.47 | 0.40 | 15 | Matrix Fibroblast |
| Scp2 | 0.51 | 0.00 | 0.00 | 0.42 | 0.33 | 15 | Matrix Fibroblast |
| Gng11 | 0.41 | 0.00 | 0.00 | 0.41 | 0.29 | 15 | Matrix Fibroblast |
| Mesdc2 | 0.59 | 0.00 | 0.00 | 0.29 | 0.19 | 15 | Matrix Fibroblast |
| Tacc1 | 0.56 | 0.00 | 0.00 | 0.33 | 0.22 | 15 | Matrix Fibroblast |
| Tagln2 | 0.39 | 0.00 | 0.00 | 0.53 | 0.45 | 15 | Matrix Fibroblast |
| Gm10020 | 0.36 | 0.00 | 0.00 | 0.59 | 0.52 | 15 | Matrix Fibroblast |
| Atp5l | 0.42 | 0.00 | 0.00 | 0.51 | 0.44 | 15 | Matrix Fibroblast |
| Aldh2 | 0.41 | 0.00 | 0.00 | 0.43 | 0.33 | 15 | Matrix Fibroblast |
| Rpl22l1 | 0.32 | 0.00 | 0.00 | 0.61 | 0.56 | 15 | Matrix Fibroblast |
| Gm10221 | 0.37 | 0.00 | 0.00 | 0.51 | 0.46 | 15 | Matrix Fibroblast |
| Itgam | 0.39 | 0.00 | 0.00 | 0.36 | 0.28 | 15 | Matrix Fibroblast |
| Cirbp | 0.49 | 0.00 | 0.00 | 0.29 | 0.21 | 15 | Matrix Fibroblast |
| Fbn1 | 0.33 | 0.00 | 0.00 | 0.26 | 0.17 | 15 | Matrix Fibroblast |
| Nfia | 0.48 | 0.00 | 0.00 | 0.32 | 0.24 | 15 | Matrix Fibroblast |
| Rpl12 | 0.27 | 0.00 | 0.00 | 0.61 | 0.55 | 15 | Matrix Fibroblast |
| Cd63 | 0.35 | 0.00 | 0.00 | 0.38 | 0.30 | 15 | Matrix Fibroblast |
| Igfbp4 | 0.33 | 0.00 | 0.00 | 0.33 | 0.25 | 15 | Matrix Fibroblast |
| Sqstm1 | 0.42 | 0.00 | 0.00 | 0.37 | 0.32 | 15 | Matrix Fibroblast |
| Gm17669 | 0.46 | 0.00 | 0.00 | 0.28 | 0.22 | 15 | Matrix Fibroblast |
| Cox6a1 | 0.33 | 0.00 | 0.00 | 0.56 | 0.55 | 15 | Matrix Fibroblast |
| 1810058I24Rik | 0.49 | 0.00 | 0.00 | 0.27 | 0.20 | 15 | Matrix Fibroblast |
| Sepw1 | 0.45 | 0.00 | 0.00 | 0.26 | 0.20 | 15 | Matrix Fibroblast |
| D4Wsu53e | 0.28 | 0.00 | 0.00 | 0.61 | 0.60 | 15 | Matrix Fibroblast |
| Pfdn5 | 0.29 | 0.00 | 0.00 | 0.52 | 0.50 | 15 | Matrix Fibroblast |
| Morf4l1 | 0.30 | 0.00 | 0.00 | 0.57 | 0.55 | 15 | Matrix Fibroblast |
| Celf2 | 0.37 | 0.00 | 0.00 | 0.34 | 0.28 | 15 | Matrix Fibroblast |
| Ilk | 0.41 | 0.00 | 0.00 | 0.28 | 0.23 | 15 | Matrix Fibroblast |
| Gm5619 | 0.39 | 0.00 | 0.00 | 0.37 | 0.33 | 15 | Matrix Fibroblast |
| Uqcrh | 0.36 | 0.00 | 0.00 | 0.49 | 0.47 | 15 | Matrix Fibroblast |
| Sh3bp5 | 0.34 | 0.00 | 0.00 | 0.25 | 0.19 | 15 | Matrix Fibroblast |
| Orc5 | 0.26 | 0.00 | 0.00 | 0.58 | 0.56 | 15 | Matrix Fibroblast |
| Gpx4 | 0.32 | 0.00 | 0.00 | 0.42 | 0.38 | 15 | Matrix Fibroblast |
| Nisch | 0.38 | 0.00 | 0.00 | 0.35 | 0.30 | 15 | Matrix Fibroblast |
| Map1lc3b | 0.37 | 0.00 | 0.00 | 0.42 | 0.40 | 15 | Matrix Fibroblast |
| Atp5l-ps1 | 0.34 | 0.00 | 0.01 | 0.31 | 0.28 | 15 | Matrix Fibroblast |
| Gm10320 | 0.35 | 0.00 | 0.02 | 0.41 | 0.39 | 15 | Matrix Fibroblast |
| Wsb1 | 0.42 | 0.00 | 0.02 | 0.25 | 0.21 | 15 | Matrix Fibroblast |
| Eif3f | 0.30 | 0.00 | 0.04 | 0.39 | 0.36 | 15 | Matrix Fibroblast |
| Fbln2 | 2.52 | 0.00 | 0.00 | 0.74 | 0.06 | 16 | Endothelial |
| Vwf | 2.31 | 0.00 | 0.00 | 0.72 | 0.02 | 16 | Endothelial |
| Fbln5 | 2.20 | 0.00 | 0.00 | 0.83 | 0.16 | 16 | Endothelial |
| Ptprb | 2.11 | 0.00 | 0.00 | 0.95 | 0.20 | 16 | Endothelial |
| Fabp4 | 1.67 | 0.00 | 0.00 | 0.32 | 0.03 | 16 | Endothelial |
| Tm4sf1 | 1.65 | 0.00 | 0.00 | 0.81 | 0.20 | 16 | Endothelial |
| Prss23 | 1.52 | 0.00 | 0.00 | 0.48 | 0.06 | 16 | Endothelial |
| Car8 | 1.51 | 0.00 | 0.00 | 0.47 | 0.07 | 16 | Endothelial |
| Cpe | 1.48 | 0.00 | 0.00 | 0.37 | 0.02 | 16 | Endothelial |
| Stmn2 | 1.43 | 0.00 | 0.00 | 0.65 | 0.15 | 16 | Endothelial |
| Lyve1 | 1.41 | 0.00 | 0.00 | 0.81 | 0.21 | 16 | Endothelial |
| Edn1 | 1.39 | 0.00 | 0.00 | 0.54 | 0.09 | 16 | Endothelial |
| Heg1 | 1.35 | 0.00 | 0.00 | 0.65 | 0.15 | 16 | Endothelial |
| Gpr126 | 1.35 | 0.00 | 0.00 | 0.54 | 0.10 | 16 | Endothelial |
| Tmem2 | 1.32 | 0.00 | 0.00 | 0.64 | 0.14 | 16 | Endothelial |
| Bmx | 1.31 | 0.00 | 0.00 | 0.29 | 0.02 | 16 | Endothelial |
| Ackr3 | 1.29 | 0.00 | 0.00 | 0.38 | 0.04 | 16 | Endothelial |
| Jam2 | 1.19 | 0.00 | 0.00 | 0.43 | 0.06 | 16 | Endothelial |
| Thsd7a | 1.19 | 0.00 | 0.00 | 0.30 | 0.02 | 16 | Endothelial |
| Calcrl | 1.16 | 0.00 | 0.00 | 0.85 | 0.24 | 16 | Endothelial |
| Tek | 1.13 | 0.00 | 0.00 | 0.58 | 0.12 | 16 | Endothelial |
| Cthrc1 | 0.96 | 0.00 | 0.00 | 0.28 | 0.02 | 16 | Endothelial |
| Procr | 0.96 | 0.00 | 0.00 | 0.29 | 0.01 | 16 | Endothelial |
| Samd5 | 0.93 | 0.00 | 0.00 | 0.30 | 0.03 | 16 | Endothelial |
| Plvap | 1.26 | 0.00 | 0.00 | 0.57 | 0.12 | 16 | Endothelial |
| Flt1 | 1.18 | 0.00 | 0.00 | 0.64 | 0.15 | 16 | Endothelial |
| Pecam1 | 1.12 | 0.00 | 0.00 | 0.80 | 0.23 | 16 | Endothelial |
| Clec14a | 1.07 | 0.00 | 0.00 | 0.68 | 0.17 | 16 | Endothelial |
| Pgm5 | 0.85 | 0.00 | 0.00 | 0.25 | 0.03 | 16 | Endothelial |
| Ramp2 | 1.00 | 0.00 | 0.00 | 0.84 | 0.27 | 16 | Endothelial |
| Egfl7 | 0.94 | 0.00 | 0.00 | 0.87 | 0.27 | 16 | Endothelial |
| Cd59a | 1.14 | 0.00 | 0.00 | 0.43 | 0.08 | 16 | Endothelial |
| Mmrn2 | 0.87 | 0.00 | 0.00 | 0.30 | 0.04 | 16 | Endothelial |
| Sdpr | 1.15 | 0.00 | 0.00 | 0.95 | 0.51 | 16 | Endothelial |
| Ltbp4 | 1.16 | 0.00 | 0.00 | 0.47 | 0.10 | 16 | Endothelial |
| Fam101b | 1.40 | 0.00 | 0.00 | 0.60 | 0.18 | 16 | Endothelial |
| Fstl1 | 1.03 | 0.00 | 0.00 | 0.91 | 0.44 | 16 | Endothelial |
| Palmd | 1.00 | 0.00 | 0.00 | 0.52 | 0.12 | 16 | Endothelial |
| Fgl2 | 0.94 | 0.00 | 0.00 | 0.26 | 0.03 | 16 | Endothelial |
| Cd93 | 0.84 | 0.00 | 0.00 | 0.71 | 0.21 | 16 | Endothelial |
| Aplnr | 1.01 | 0.00 | 0.00 | 0.35 | 0.07 | 16 | Endothelial |
| Megf6 | 0.78 | 0.00 | 0.00 | 0.26 | 0.04 | 16 | Endothelial |
| Ece1 | 1.06 | 0.00 | 0.00 | 0.69 | 0.25 | 16 | Endothelial |
| Cxcl12 | 2.33 | 0.00 | 0.00 | 0.45 | 0.12 | 16 | Endothelial |
| Cd200 | 1.02 | 0.00 | 0.00 | 0.71 | 0.27 | 16 | Endothelial |
| Col5a2 | 1.21 | 0.00 | 0.00 | 0.56 | 0.17 | 16 | Endothelial |
| Tie1 | 0.85 | 0.00 | 0.00 | 0.45 | 0.11 | 16 | Endothelial |
| Jag1 | 0.95 | 0.00 | 0.00 | 0.27 | 0.05 | 16 | Endothelial |
| Cldn5 | 0.80 | 0.00 | 0.00 | 0.67 | 0.22 | 16 | Endothelial |
| S100a16 | 0.88 | 0.00 | 0.00 | 0.67 | 0.23 | 16 | Endothelial |
| Aqp1 | 0.91 | 0.00 | 0.00 | 0.60 | 0.19 | 16 | Endothelial |
| Ctla2a | 1.15 | 0.00 | 0.00 | 0.60 | 0.20 | 16 | Endothelial |
| Tspan7 | 0.81 | 0.00 | 0.00 | 0.64 | 0.21 | 16 | Endothelial |
| Ehd4 | 0.90 | 0.00 | 0.00 | 0.70 | 0.27 | 16 | Endothelial |
| Id3 | 0.92 | 0.00 | 0.00 | 0.85 | 0.45 | 16 | Endothelial |
| Gja4 | 1.07 | 0.00 | 0.00 | 0.31 | 0.07 | 16 | Endothelial |
| Adam15 | 0.75 | 0.00 | 0.00 | 0.35 | 0.08 | 16 | Endothelial |
| Adcy4 | 0.81 | 0.00 | 0.00 | 0.36 | 0.08 | 16 | Endothelial |
| Entpd1 | 0.91 | 0.00 | 0.00 | 0.37 | 0.09 | 16 | Endothelial |
| Tgm2 | 0.83 | 0.00 | 0.00 | 0.37 | 0.09 | 16 | Endothelial |
| Cdh5 | 0.59 | 0.00 | 0.00 | 0.68 | 0.22 | 16 | Endothelial |
| Csrp2 | 0.99 | 0.00 | 0.00 | 0.50 | 0.16 | 16 | Endothelial |
| Meis2 | 0.83 | 0.00 | 0.00 | 0.43 | 0.11 | 16 | Endothelial |
| Plk2 | 1.01 | 0.00 | 0.00 | 0.49 | 0.16 | 16 | Endothelial |
| Foxp1 | 0.89 | 0.00 | 0.00 | 0.81 | 0.44 | 16 | Endothelial |
| Ecscr | 0.67 | 0.00 | 0.00 | 0.53 | 0.16 | 16 | Endothelial |
| Igf2 | 1.39 | 0.00 | 0.00 | 0.31 | 0.07 | 16 | Endothelial |
| Ephb4 | 0.75 | 0.00 | 0.00 | 0.33 | 0.08 | 16 | Endothelial |
| Fgd5 | 0.68 | 0.00 | 0.00 | 0.29 | 0.06 | 16 | Endothelial |
| Srgn | 0.82 | 0.00 | 0.00 | 0.74 | 0.32 | 16 | Endothelial |
| Sema3g | 0.99 | 0.00 | 0.00 | 0.30 | 0.07 | 16 | Endothelial |
| Cd9 | 0.84 | 0.00 | 0.00 | 0.73 | 0.34 | 16 | Endothelial |
| Efnb2 | 1.06 | 0.00 | 0.00 | 0.45 | 0.14 | 16 | Endothelial |
| She | 0.70 | 0.00 | 0.00 | 0.27 | 0.06 | 16 | Endothelial |
| Epas1 | 0.67 | 0.00 | 0.00 | 0.66 | 0.25 | 16 | Endothelial |
| Slc43a3 | 0.84 | 0.00 | 0.00 | 0.57 | 0.21 | 16 | Endothelial |
| Foxo1 | 0.76 | 0.00 | 0.00 | 0.32 | 0.08 | 16 | Endothelial |
| Emcn | 0.75 | 0.00 | 0.00 | 0.39 | 0.11 | 16 | Endothelial |
| Acer2 | 0.80 | 0.00 | 0.00 | 0.36 | 0.10 | 16 | Endothelial |
| Erg | 0.78 | 0.00 | 0.00 | 0.43 | 0.13 | 16 | Endothelial |
| Cav1 | 0.73 | 0.00 | 0.00 | 0.70 | 0.31 | 16 | Endothelial |
| Plxnd1 | 0.76 | 0.00 | 0.00 | 0.30 | 0.07 | 16 | Endothelial |
| Eln | 0.91 | 0.00 | 0.00 | 0.46 | 0.15 | 16 | Endothelial |
| Rasip1 | 0.66 | 0.00 | 0.00 | 0.39 | 0.12 | 16 | Endothelial |
| Myct1 | 0.67 | 0.00 | 0.00 | 0.41 | 0.13 | 16 | Endothelial |
| Rhoj | 0.74 | 0.00 | 0.00 | 0.50 | 0.18 | 16 | Endothelial |
| Ripply3 | 0.95 | 0.00 | 0.00 | 0.39 | 0.12 | 16 | Endothelial |
| Ptrf | 0.85 | 0.00 | 0.00 | 0.68 | 0.35 | 16 | Endothelial |
| Hyal2 | 0.67 | 0.00 | 0.00 | 0.29 | 0.07 | 16 | Endothelial |
| Icam2 | 0.60 | 0.00 | 0.00 | 0.47 | 0.16 | 16 | Endothelial |
| Ace | 0.78 | 0.00 | 0.00 | 0.41 | 0.13 | 16 | Endothelial |
| Tmem88 | 0.73 | 0.00 | 0.00 | 0.36 | 0.10 | 16 | Endothelial |
| Ctla2b | 0.63 | 0.00 | 0.00 | 0.28 | 0.07 | 16 | Endothelial |
| Ifitm3 | 0.65 | 0.00 | 0.00 | 0.76 | 0.39 | 16 | Endothelial |
| Id1 | 0.80 | 0.00 | 0.00 | 0.56 | 0.23 | 16 | Endothelial |
| Esam | 0.61 | 0.00 | 0.00 | 0.50 | 0.18 | 16 | Endothelial |
| Rnf144a | 0.77 | 0.00 | 0.00 | 0.44 | 0.15 | 16 | Endothelial |
| Igfbp4 | 0.89 | 0.00 | 0.00 | 0.56 | 0.25 | 16 | Endothelial |
| Ceacam1 | 0.68 | 0.00 | 0.00 | 0.32 | 0.09 | 16 | Endothelial |
| Cbfa2t3 | 0.73 | 0.00 | 0.00 | 0.41 | 0.14 | 16 | Endothelial |
| Gata6 | 0.79 | 0.00 | 0.00 | 0.30 | 0.08 | 16 | Endothelial |
| Plcb4 | 0.79 | 0.00 | 0.00 | 0.40 | 0.14 | 16 | Endothelial |
| Slc9a3r2 | 0.66 | 0.00 | 0.00 | 0.52 | 0.20 | 16 | Endothelial |
| Gata2 | 0.60 | 0.00 | 0.00 | 0.41 | 0.14 | 16 | Endothelial |
| Prcp | 0.62 | 0.00 | 0.00 | 0.31 | 0.09 | 16 | Endothelial |
| Armcx1 | 0.71 | 0.00 | 0.00 | 0.31 | 0.09 | 16 | Endothelial |
| Myzap | 0.60 | 0.00 | 0.00 | 0.41 | 0.14 | 16 | Endothelial |
| Tspan6 | 0.59 | 0.00 | 0.00 | 0.29 | 0.08 | 16 | Endothelial |
| Dst | 0.74 | 0.00 | 0.00 | 0.38 | 0.13 | 16 | Endothelial |
| Smad7 | 0.74 | 0.00 | 0.00 | 0.35 | 0.11 | 16 | Endothelial |
| Prkcdbp | 0.69 | 0.00 | 0.00 | 0.34 | 0.11 | 16 | Endothelial |
| Tnfsf10 | 0.62 | 0.00 | 0.00 | 0.29 | 0.08 | 16 | Endothelial |
| Crim1 | 0.69 | 0.00 | 0.00 | 0.29 | 0.09 | 16 | Endothelial |
| Fkbp1a | 0.64 | 0.00 | 0.00 | 0.70 | 0.40 | 16 | Endothelial |
| Itgb1 | 0.65 | 0.00 | 0.00 | 0.84 | 0.63 | 16 | Endothelial |
| Cav2 | 0.59 | 0.00 | 0.00 | 0.47 | 0.19 | 16 | Endothelial |
| Utrn | 0.88 | 0.00 | 0.00 | 0.41 | 0.16 | 16 | Endothelial |
| Ednrb | 0.56 | 0.00 | 0.00 | 0.41 | 0.15 | 16 | Endothelial |
| Elk3 | 0.65 | 0.00 | 0.00 | 0.38 | 0.13 | 16 | Endothelial |
| Notch1 | 0.70 | 0.00 | 0.00 | 0.37 | 0.13 | 16 | Endothelial |
| 2900026A02Rik | 0.58 | 0.00 | 0.00 | 0.27 | 0.08 | 16 | Endothelial |
| Acvrl1 | 0.56 | 0.00 | 0.00 | 0.45 | 0.17 | 16 | Endothelial |
| Hsp90ab1 | 0.40 | 0.00 | 0.00 | 0.96 | 0.89 | 16 | Endothelial |
| Pcsk5 | 0.85 | 0.00 | 0.00 | 0.30 | 0.10 | 16 | Endothelial |
| Hoxa5 | 0.65 | 0.00 | 0.00 | 0.36 | 0.13 | 16 | Endothelial |
| Rab11a | 0.69 | 0.00 | 0.00 | 0.53 | 0.26 | 16 | Endothelial |
| 4931406P16Rik | 0.59 | 0.00 | 0.00 | 0.44 | 0.18 | 16 | Endothelial |
| Ccdc80 | 0.63 | 0.00 | 0.00 | 0.32 | 0.11 | 16 | Endothelial |
| Eng | 0.56 | 0.00 | 0.00 | 0.32 | 0.11 | 16 | Endothelial |
| Slk | 0.65 | 0.00 | 0.00 | 0.70 | 0.43 | 16 | Endothelial |
| Mcam | 0.50 | 0.00 | 0.00 | 0.36 | 0.13 | 16 | Endothelial |
| Klf4 | 0.57 | 0.00 | 0.00 | 0.51 | 0.23 | 16 | Endothelial |
| Cyyr1 | 0.47 | 0.00 | 0.00 | 0.39 | 0.15 | 16 | Endothelial |
| Ehd2 | 0.59 | 0.00 | 0.00 | 0.37 | 0.14 | 16 | Endothelial |
| Podxl | 0.50 | 0.00 | 0.00 | 0.31 | 0.11 | 16 | Endothelial |
| Luzp1 | 0.63 | 0.00 | 0.00 | 0.44 | 0.20 | 16 | Endothelial |
| Fam3c | 0.95 | 0.00 | 0.00 | 0.33 | 0.13 | 16 | Endothelial |
| Sox17 | 0.61 | 0.00 | 0.00 | 0.35 | 0.14 | 16 | Endothelial |
| Ldb2 | 0.59 | 0.00 | 0.00 | 0.39 | 0.16 | 16 | Endothelial |
| Gnaq | 0.57 | 0.00 | 0.00 | 0.45 | 0.21 | 16 | Endothelial |
| Apbb2 | 0.56 | 0.00 | 0.00 | 0.27 | 0.09 | 16 | Endothelial |
| Nrp1 | 0.51 | 0.00 | 0.00 | 0.67 | 0.39 | 16 | Endothelial |
| Aplp2 | 0.60 | 0.00 | 0.00 | 0.64 | 0.39 | 16 | Endothelial |
| Cd2ap | 0.68 | 0.00 | 0.00 | 0.59 | 0.34 | 16 | Endothelial |
| S1pr1 | 0.46 | 0.00 | 0.00 | 0.43 | 0.18 | 16 | Endothelial |
| Gng11 | 0.45 | 0.00 | 0.00 | 0.57 | 0.29 | 16 | Endothelial |
| Ushbp1 | 0.50 | 0.00 | 0.00 | 0.26 | 0.09 | 16 | Endothelial |
| Slfn5 | 0.46 | 0.00 | 0.00 | 0.27 | 0.10 | 16 | Endothelial |
| Fnbp1l | 0.50 | 0.00 | 0.00 | 0.50 | 0.25 | 16 | Endothelial |
| Mecom | 0.79 | 0.00 | 0.00 | 0.27 | 0.10 | 16 | Endothelial |
| Azin1 | 0.71 | 0.00 | 0.00 | 0.32 | 0.13 | 16 | Endothelial |
| Gimap1 | 0.41 | 0.00 | 0.00 | 0.34 | 0.14 | 16 | Endothelial |
| Cmip | 0.60 | 0.00 | 0.00 | 0.41 | 0.20 | 16 | Endothelial |
| Cd151 | 0.53 | 0.00 | 0.00 | 0.36 | 0.16 | 16 | Endothelial |
| Slc6a6 | 0.73 | 0.00 | 0.00 | 0.47 | 0.25 | 16 | Endothelial |
| Slc16a2 | 0.57 | 0.00 | 0.00 | 0.26 | 0.10 | 16 | Endothelial |
| Nav1 | 0.45 | 0.00 | 0.00 | 0.29 | 0.11 | 16 | Endothelial |
| Hmcn1 | 0.53 | 0.00 | 0.00 | 0.32 | 0.13 | 16 | Endothelial |
| Ppap2a | 0.42 | 0.00 | 0.00 | 0.41 | 0.19 | 16 | Endothelial |
| Mcfd2 | 0.50 | 0.00 | 0.00 | 0.29 | 0.12 | 16 | Endothelial |
| Cxx1a | 0.49 | 0.00 | 0.00 | 0.44 | 0.22 | 16 | Endothelial |
| Pea15a | 0.49 | 0.00 | 0.00 | 0.33 | 0.14 | 16 | Endothelial |
| Plec | 0.48 | 0.00 | 0.00 | 0.30 | 0.12 | 16 | Endothelial |
| Tspan18 | 0.38 | 0.00 | 0.00 | 0.33 | 0.13 | 16 | Endothelial |
| Nck1 | 0.46 | 0.00 | 0.00 | 0.31 | 0.13 | 16 | Endothelial |
| Emp1 | 0.42 | 0.00 | 0.00 | 0.45 | 0.22 | 16 | Endothelial |
| Itm2b | 0.33 | 0.00 | 0.00 | 0.89 | 0.76 | 16 | Endothelial |
| Ifitm2 | 0.47 | 0.00 | 0.00 | 0.55 | 0.32 | 16 | Endothelial |
| Tcf4 | 0.51 | 0.00 | 0.00 | 0.65 | 0.41 | 16 | Endothelial |
| Swap70 | 0.50 | 0.00 | 0.00 | 0.28 | 0.11 | 16 | Endothelial |
| Adam10 | 0.67 | 0.00 | 0.00 | 0.38 | 0.18 | 16 | Endothelial |
| Id2 | 0.47 | 0.00 | 0.00 | 0.57 | 0.33 | 16 | Endothelial |
| Rfk | 0.52 | 0.00 | 0.00 | 0.25 | 0.10 | 16 | Endothelial |
| Hip1 | 0.46 | 0.00 | 0.00 | 0.35 | 0.16 | 16 | Endothelial |
| Crip2 | 0.39 | 0.00 | 0.00 | 0.68 | 0.41 | 16 | Endothelial |
| Fam115a | 0.49 | 0.00 | 0.00 | 0.32 | 0.14 | 16 | Endothelial |
| Golm1 | 0.56 | 0.00 | 0.00 | 0.26 | 0.10 | 16 | Endothelial |
| Rdx | 0.47 | 0.00 | 0.00 | 0.67 | 0.45 | 16 | Endothelial |
| Agrn | 0.39 | 0.00 | 0.00 | 0.30 | 0.13 | 16 | Endothelial |
| Trib2 | 0.46 | 0.00 | 0.00 | 0.27 | 0.11 | 16 | Endothelial |
| Cxx1b | 0.45 | 0.00 | 0.00 | 0.43 | 0.22 | 16 | Endothelial |
| Klf2 | 0.51 | 0.00 | 0.00 | 0.40 | 0.20 | 16 | Endothelial |
| Actn4 | 0.50 | 0.00 | 0.00 | 0.50 | 0.29 | 16 | Endothelial |
| Anxa2 | 0.48 | 0.00 | 0.00 | 0.67 | 0.47 | 16 | Endothelial |
| Skap2 | 0.38 | 0.00 | 0.00 | 0.29 | 0.13 | 16 | Endothelial |
| Thbd | 0.48 | 0.00 | 0.00 | 0.46 | 0.25 | 16 | Endothelial |
| Cnn3 | 0.41 | 0.00 | 0.00 | 0.54 | 0.30 | 16 | Endothelial |
| Sema6a | 0.33 | 0.00 | 0.00 | 0.28 | 0.12 | 16 | Endothelial |
| Ghr | 0.44 | 0.00 | 0.00 | 0.32 | 0.15 | 16 | Endothelial |
| Anxa3 | 0.40 | 0.00 | 0.00 | 0.43 | 0.22 | 16 | Endothelial |
| Lima1 | 0.48 | 0.00 | 0.00 | 0.31 | 0.14 | 16 | Endothelial |
| Gimap5 | 0.32 | 0.00 | 0.00 | 0.27 | 0.11 | 16 | Endothelial |
| Atp2b1 | 0.49 | 0.00 | 0.00 | 0.76 | 0.58 | 16 | Endothelial |
| Tmem109 | 0.42 | 0.00 | 0.00 | 0.28 | 0.12 | 16 | Endothelial |
| Marcks | 0.28 | 0.00 | 0.00 | 0.75 | 0.50 | 16 | Endothelial |
| Rapgef5 | 0.45 | 0.00 | 0.00 | 0.25 | 0.11 | 16 | Endothelial |
| Map7d1 | 0.41 | 0.00 | 0.00 | 0.36 | 0.18 | 16 | Endothelial |
| Nfe2l1 | 0.49 | 0.00 | 0.00 | 0.42 | 0.23 | 16 | Endothelial |
| Cxx1c | 0.42 | 0.00 | 0.00 | 0.41 | 0.22 | 16 | Endothelial |
| Lphn2 | 0.34 | 0.00 | 0.00 | 0.30 | 0.13 | 16 | Endothelial |
| Hdac7 | 0.40 | 0.00 | 0.00 | 0.30 | 0.14 | 16 | Endothelial |
| Ccdc50 | 0.45 | 0.00 | 0.00 | 0.45 | 0.26 | 16 | Endothelial |
| Lgals9 | 0.37 | 0.00 | 0.00 | 0.30 | 0.14 | 16 | Endothelial |
| Ppap2b | 0.45 | 0.00 | 0.00 | 0.43 | 0.24 | 16 | Endothelial |
| Arhgap31 | 0.48 | 0.00 | 0.00 | 0.26 | 0.12 | 16 | Endothelial |
| Mmp2 | 0.39 | 0.00 | 0.00 | 0.29 | 0.13 | 16 | Endothelial |
| Golim4 | 0.45 | 0.00 | 0.00 | 0.38 | 0.20 | 16 | Endothelial |
| Rabgap1 | 0.45 | 0.00 | 0.00 | 0.28 | 0.13 | 16 | Endothelial |
| Tmod3 | 0.47 | 0.00 | 0.00 | 0.45 | 0.26 | 16 | Endothelial |
| Fli1 | 0.38 | 0.00 | 0.00 | 0.27 | 0.12 | 16 | Endothelial |
| Tsc22d1 | 0.51 | 0.00 | 0.00 | 0.53 | 0.33 | 16 | Endothelial |
| Sox4 | 0.50 | 0.00 | 0.00 | 0.57 | 0.36 | 16 | Endothelial |
| Ehd1 | 0.38 | 0.00 | 0.00 | 0.26 | 0.12 | 16 | Endothelial |
| Itga6 | 0.40 | 0.00 | 0.00 | 0.29 | 0.13 | 16 | Endothelial |
| Pdlim1 | 0.39 | 0.00 | 0.00 | 0.30 | 0.14 | 16 | Endothelial |
| Rhoc | 0.39 | 0.00 | 0.00 | 0.39 | 0.21 | 16 | Endothelial |
| Sat1 | 0.63 | 0.00 | 0.00 | 0.52 | 0.33 | 16 | Endothelial |
| Ttc28 | 0.38 | 0.00 | 0.00 | 0.27 | 0.12 | 16 | Endothelial |
| Ppfibp1 | 0.39 | 0.00 | 0.00 | 0.30 | 0.14 | 16 | Endothelial |
| Tead2 | 0.37 | 0.00 | 0.00 | 0.27 | 0.12 | 16 | Endothelial |
| Gbp7 | 0.38 | 0.00 | 0.00 | 0.29 | 0.14 | 16 | Endothelial |
| Alas1 | 0.43 | 0.00 | 0.00 | 0.26 | 0.12 | 16 | Endothelial |
| Kank3 | 0.36 | 0.00 | 0.00 | 0.30 | 0.14 | 16 | Endothelial |
| Ddah2 | 0.32 | 0.00 | 0.00 | 0.46 | 0.26 | 16 | Endothelial |
| Spag9 | 0.43 | 0.00 | 0.00 | 0.33 | 0.17 | 16 | Endothelial |
| Tmem100 | 0.32 | 0.00 | 0.00 | 0.51 | 0.30 | 16 | Endothelial |
| Gm7676 | 0.28 | 0.00 | 0.00 | 0.36 | 0.19 | 16 | Endothelial |
| Cdc42ep3 | 0.49 | 0.00 | 0.00 | 0.29 | 0.14 | 16 | Endothelial |
| Ppic | 0.38 | 0.00 | 0.00 | 0.58 | 0.38 | 16 | Endothelial |
| Ybx3 | 0.40 | 0.00 | 0.00 | 0.36 | 0.19 | 16 | Endothelial |
| Ccdc85b | 0.35 | 0.00 | 0.00 | 0.25 | 0.12 | 16 | Endothelial |
| Klhl5 | 0.40 | 0.00 | 0.00 | 0.29 | 0.14 | 16 | Endothelial |
| Itm2c | 0.41 | 0.00 | 0.00 | 0.39 | 0.22 | 16 | Endothelial |
| Uaca | 0.39 | 0.00 | 0.00 | 0.28 | 0.14 | 16 | Endothelial |
| Adamts1 | 0.36 | 0.00 | 0.00 | 0.39 | 0.21 | 16 | Endothelial |
| Casz1 | 0.39 | 0.00 | 0.00 | 0.28 | 0.14 | 16 | Endothelial |
| Pam | 0.34 | 0.00 | 0.00 | 0.33 | 0.17 | 16 | Endothelial |
| Vim | 0.25 | 0.00 | 0.00 | 0.86 | 0.66 | 16 | Endothelial |
| Gcc2 | 0.47 | 0.00 | 0.00 | 0.42 | 0.25 | 16 | Endothelial |
| Mgp | 0.59 | 0.00 | 0.00 | 0.50 | 0.31 | 16 | Endothelial |
| Bst2 | 0.32 | 0.00 | 0.00 | 0.28 | 0.14 | 16 | Endothelial |
| Msn | 0.34 | 0.00 | 0.00 | 0.62 | 0.43 | 16 | Endothelial |
| Tubb5 | 0.33 | 0.00 | 0.00 | 0.64 | 0.44 | 16 | Endothelial |
| Ctnna1 | 0.37 | 0.00 | 0.00 | 0.55 | 0.36 | 16 | Endothelial |
| Mapk3 | 0.35 | 0.00 | 0.00 | 0.27 | 0.14 | 16 | Endothelial |
| Pdia3 | 0.39 | 0.00 | 0.00 | 0.62 | 0.44 | 16 | Endothelial |
| Amotl1 | 0.30 | 0.00 | 0.00 | 0.30 | 0.16 | 16 | Endothelial |
| Sptbn1 | 0.32 | 0.00 | 0.00 | 0.72 | 0.54 | 16 | Endothelial |
| Laptm4a | 0.35 | 0.00 | 0.00 | 0.67 | 0.52 | 16 | Endothelial |
| Ttc3 | 0.40 | 0.00 | 0.00 | 0.46 | 0.29 | 16 | Endothelial |
| Marcksl1 | 0.39 | 0.00 | 0.00 | 0.35 | 0.19 | 16 | Endothelial |
| Arhgef2 | 0.33 | 0.00 | 0.00 | 0.36 | 0.20 | 16 | Endothelial |
| Nfic | 0.37 | 0.00 | 0.00 | 0.41 | 0.24 | 16 | Endothelial |
| F11r | 0.37 | 0.00 | 0.00 | 0.31 | 0.16 | 16 | Endothelial |
| Ralb | 0.34 | 0.00 | 0.00 | 0.34 | 0.19 | 16 | Endothelial |
| Cd81 | 0.30 | 0.00 | 0.00 | 0.59 | 0.40 | 16 | Endothelial |
| App | 0.25 | 0.00 | 0.00 | 0.77 | 0.59 | 16 | Endothelial |
| Dnaja1 | 0.37 | 0.00 | 0.00 | 0.67 | 0.51 | 16 | Endothelial |
| G3bp2 | 0.45 | 0.00 | 0.00 | 0.48 | 0.31 | 16 | Endothelial |
| Zmiz1 | 0.41 | 0.00 | 0.00 | 0.37 | 0.22 | 16 | Endothelial |
| Fbn1 | 0.34 | 0.00 | 0.00 | 0.32 | 0.17 | 16 | Endothelial |
| 2-Sep | 0.33 | 0.00 | 0.00 | 0.53 | 0.35 | 16 | Endothelial |
| H2-T23 | 0.32 | 0.00 | 0.00 | 0.25 | 0.13 | 16 | Endothelial |
| Sypl | 0.37 | 0.00 | 0.00 | 0.42 | 0.26 | 16 | Endothelial |
| Cd97 | 0.33 | 0.00 | 0.00 | 0.33 | 0.18 | 16 | Endothelial |
| Bnip2 | 0.36 | 0.00 | 0.00 | 0.47 | 0.30 | 16 | Endothelial |
| Serpinb6a | 0.35 | 0.00 | 0.00 | 0.32 | 0.18 | 16 | Endothelial |
| Phactr2 | 0.33 | 0.00 | 0.00 | 0.25 | 0.13 | 16 | Endothelial |
| Tenc1 | 0.32 | 0.00 | 0.00 | 0.27 | 0.14 | 16 | Endothelial |
| Ano6 | 0.38 | 0.00 | 0.00 | 0.31 | 0.18 | 16 | Endothelial |
| Calu | 0.31 | 0.00 | 0.00 | 0.40 | 0.24 | 16 | Endothelial |
| Arhgap29 | 0.31 | 0.00 | 0.00 | 0.37 | 0.22 | 16 | Endothelial |
| Ktn1 | 0.33 | 0.00 | 0.00 | 0.54 | 0.38 | 16 | Endothelial |
| Fabp5 | 0.27 | 0.00 | 0.00 | 0.25 | 0.13 | 16 | Endothelial |
| Clic1 | 0.27 | 0.00 | 0.00 | 0.54 | 0.38 | 16 | Endothelial |
| Cpd | 0.33 | 0.00 | 0.00 | 0.27 | 0.14 | 16 | Endothelial |
| Smarca2 | 0.36 | 0.00 | 0.00 | 0.50 | 0.34 | 16 | Endothelial |
| Pbx1 | 0.35 | 0.00 | 0.00 | 0.34 | 0.20 | 16 | Endothelial |
| Atp1b3 | 0.33 | 0.00 | 0.00 | 0.41 | 0.25 | 16 | Endothelial |
| Cnih1 | 0.31 | 0.00 | 0.00 | 0.32 | 0.19 | 16 | Endothelial |
| Arglu1 | 0.33 | 0.00 | 0.00 | 0.58 | 0.42 | 16 | Endothelial |
| Zbtb20 | 0.43 | 0.00 | 0.00 | 0.41 | 0.27 | 16 | Endothelial |
| Pltp | 0.35 | 0.00 | 0.00 | 0.31 | 0.18 | 16 | Endothelial |
| Clic4 | 0.32 | 0.00 | 0.00 | 0.45 | 0.29 | 16 | Endothelial |
| Xiap | 0.34 | 0.00 | 0.00 | 0.34 | 0.20 | 16 | Endothelial |
| S100a10 | 0.35 | 0.00 | 0.00 | 0.53 | 0.38 | 16 | Endothelial |
| Tjp1 | 0.31 | 0.00 | 0.00 | 0.29 | 0.16 | 16 | Endothelial |
| Capns1 | 0.34 | 0.00 | 0.00 | 0.39 | 0.24 | 16 | Endothelial |
| Ccser2 | 0.30 | 0.00 | 0.00 | 0.26 | 0.14 | 16 | Endothelial |
| Mest | 0.28 | 0.00 | 0.00 | 0.34 | 0.20 | 16 | Endothelial |
| Emp3 | 0.26 | 0.00 | 0.00 | 0.36 | 0.21 | 16 | Endothelial |
| Cdc42bpa | 0.45 | 0.00 | 0.00 | 0.26 | 0.15 | 16 | Endothelial |
| Rac1 | 0.29 | 0.00 | 0.00 | 0.45 | 0.30 | 16 | Endothelial |
| Snx3 | 0.32 | 0.00 | 0.00 | 0.48 | 0.33 | 16 | Endothelial |
| Cyb5r3 | 0.30 | 0.00 | 0.00 | 0.54 | 0.39 | 16 | Endothelial |
| 2700089E24Rik | 0.28 | 0.00 | 0.00 | 0.37 | 0.23 | 16 | Endothelial |
| Nr2f2 | 0.32 | 0.00 | 0.00 | 0.34 | 0.21 | 16 | Endothelial |
| Ndufa8 | 0.32 | 0.00 | 0.00 | 0.38 | 0.25 | 16 | Endothelial |
| Ppp1r12a | 0.37 | 0.00 | 0.00 | 0.38 | 0.25 | 16 | Endothelial |
| Gnai2 | 0.27 | 0.00 | 0.00 | 0.55 | 0.38 | 16 | Endothelial |
| S100a13 | 0.29 | 0.00 | 0.00 | 0.39 | 0.25 | 16 | Endothelial |
| Srpr | 0.29 | 0.00 | 0.00 | 0.37 | 0.24 | 16 | Endothelial |
| 11-Sep | 0.26 | 0.00 | 0.00 | 0.36 | 0.22 | 16 | Endothelial |
| Cflar | 0.26 | 0.00 | 0.00 | 0.31 | 0.19 | 16 | Endothelial |
| Gm26735 | 0.30 | 0.00 | 0.00 | 0.31 | 0.19 | 16 | Endothelial |
| Nras | 0.27 | 0.00 | 0.00 | 0.28 | 0.16 | 16 | Endothelial |
| Sptlc2 | 0.25 | 0.00 | 0.00 | 0.26 | 0.15 | 16 | Endothelial |
| Zc3h7a | 0.29 | 0.00 | 0.00 | 0.42 | 0.29 | 16 | Endothelial |
| Mllt4 | 0.28 | 0.00 | 0.00 | 0.33 | 0.21 | 16 | Endothelial |
| Rbm28 | 0.28 | 0.00 | 0.00 | 0.32 | 0.19 | 16 | Endothelial |
| Ctnnb1 | 0.29 | 0.00 | 0.00 | 0.48 | 0.34 | 16 | Endothelial |
| Amd1 | 0.67 | 0.00 | 0.00 | 0.29 | 0.18 | 16 | Endothelial |
| Tnfaip1 | 0.33 | 0.00 | 0.00 | 0.27 | 0.16 | 16 | Endothelial |
| Plcg1 | 0.26 | 0.00 | 0.00 | 0.27 | 0.16 | 16 | Endothelial |
| Iqgap1 | 0.29 | 0.00 | 0.00 | 0.59 | 0.45 | 16 | Endothelial |
| Chd3 | 0.33 | 0.00 | 0.00 | 0.29 | 0.18 | 16 | Endothelial |
| Calr | 0.32 | 0.00 | 0.00 | 0.56 | 0.43 | 16 | Endothelial |
| 1700020I14Rik | 0.27 | 0.00 | 0.00 | 0.34 | 0.23 | 16 | Endothelial |
| Leprot | 0.28 | 0.00 | 0.00 | 0.46 | 0.33 | 16 | Endothelial |
| Sh3glb1 | 0.26 | 0.00 | 0.00 | 0.54 | 0.40 | 16 | Endothelial |
| Akr1a1 | 0.25 | 0.00 | 0.00 | 0.61 | 0.49 | 16 | Endothelial |
| Hes1 | 0.33 | 0.00 | 0.00 | 0.42 | 0.29 | 16 | Endothelial |
| Xist | 0.26 | 0.00 | 0.00 | 0.64 | 0.51 | 16 | Endothelial |
| Tmem59 | 0.26 | 0.00 | 0.00 | 0.53 | 0.40 | 16 | Endothelial |
| Thoc2 | 0.31 | 0.00 | 0.00 | 0.45 | 0.33 | 16 | Endothelial |
| Atrx | 0.26 | 0.00 | 0.00 | 0.75 | 0.66 | 16 | Endothelial |
| Tmem30a | 0.26 | 0.00 | 0.00 | 0.40 | 0.28 | 16 | Endothelial |
| Atp6v0e | 0.25 | 0.00 | 0.00 | 0.31 | 0.20 | 16 | Endothelial |
| Pcmtd1 | 0.25 | 0.00 | 0.00 | 0.26 | 0.16 | 16 | Endothelial |
| Zrsr2 | 0.27 | 0.00 | 0.00 | 0.25 | 0.16 | 16 | Endothelial |
| Ddx3x | 0.31 | 0.00 | 0.00 | 0.41 | 0.30 | 16 | Endothelial |
| Amd2 | 0.65 | 0.00 | 0.00 | 0.27 | 0.18 | 16 | Endothelial |
| Ginm1 | 0.27 | 0.00 | 0.00 | 0.33 | 0.23 | 16 | Endothelial |
| Trip11 | 0.27 | 0.00 | 0.00 | 0.31 | 0.22 | 16 | Endothelial |
| Tia1 | 0.26 | 0.00 | 0.00 | 0.31 | 0.22 | 16 | Endothelial |
| Bsg | 0.33 | 0.00 | 0.00 | 0.45 | 0.36 | 16 | Endothelial |
| Sec14l3 | 3.43 | 0.00 | 0.00 | 0.99 | 0.10 | 17 | Ciliated Cell |
| Dynlrb2 | 3.08 | 0.00 | 0.00 | 0.94 | 0.01 | 17 | Ciliated Cell |
| AU040972 | 2.72 | 0.00 | 0.00 | 0.69 | 0.00 | 17 | Ciliated Cell |
| Ccdc153 | 2.70 | 0.00 | 0.00 | 0.82 | 0.00 | 17 | Ciliated Cell |
| Cbr2 | 2.62 | 0.00 | 0.00 | 0.98 | 0.15 | 17 | Ciliated Cell |
| Tppp3 | 2.62 | 0.00 | 0.00 | 0.94 | 0.11 | 17 | Ciliated Cell |
| Aldh1a1 | 2.50 | 0.00 | 0.00 | 0.88 | 0.03 | 17 | Ciliated Cell |
| Cdkn1c | 2.47 | 0.00 | 0.00 | 0.85 | 0.17 | 17 | Ciliated Cell |
| Ccdc39 | 2.39 | 0.00 | 0.00 | 0.83 | 0.02 | 17 | Ciliated Cell |
| Fhad1 | 2.38 | 0.00 | 0.00 | 0.81 | 0.01 | 17 | Ciliated Cell |
| Tmem212 | 2.34 | 0.00 | 0.00 | 0.80 | 0.00 | 17 | Ciliated Cell |
| 1700009P17Rik | 2.26 | 0.00 | 0.00 | 0.83 | 0.01 | 17 | Ciliated Cell |
| Fam161a | 2.24 | 0.00 | 0.00 | 0.79 | 0.01 | 17 | Ciliated Cell |
| Rsph1 | 2.24 | 0.00 | 0.00 | 0.77 | 0.00 | 17 | Ciliated Cell |
| 1700016K19Rik | 2.23 | 0.00 | 0.00 | 0.80 | 0.00 | 17 | Ciliated Cell |
| Spag17 | 2.22 | 0.00 | 0.00 | 0.81 | 0.00 | 17 | Ciliated Cell |
| Wdr66 | 2.21 | 0.00 | 0.00 | 0.79 | 0.01 | 17 | Ciliated Cell |
| Ccdc113 | 2.19 | 0.00 | 0.00 | 0.80 | 0.00 | 17 | Ciliated Cell |
| Spef2 | 2.19 | 0.00 | 0.00 | 0.75 | 0.01 | 17 | Ciliated Cell |
| Mt1 | 2.17 | 0.00 | 0.00 | 0.84 | 0.15 | 17 | Ciliated Cell |
| Chchd10 | 2.17 | 0.00 | 0.00 | 0.90 | 0.12 | 17 | Ciliated Cell |
| Fam183b | 2.16 | 0.00 | 0.00 | 0.77 | 0.00 | 17 | Ciliated Cell |
| Cyp2s1 | 2.16 | 0.00 | 0.00 | 0.79 | 0.01 | 17 | Ciliated Cell |
| Cd24a | 2.13 | 0.00 | 0.00 | 0.89 | 0.12 | 17 | Ciliated Cell |
| Foxj1 | 2.11 | 0.00 | 0.00 | 0.75 | 0.00 | 17 | Ciliated Cell |
| Hspa4l | 2.10 | 0.00 | 0.00 | 0.81 | 0.05 | 17 | Ciliated Cell |
| Wdr52 | 2.09 | 0.00 | 0.00 | 0.78 | 0.00 | 17 | Ciliated Cell |
| Ccdc181 | 2.08 | 0.00 | 0.00 | 0.75 | 0.02 | 17 | Ciliated Cell |
| Wdr96 | 2.06 | 0.00 | 0.00 | 0.71 | 0.00 | 17 | Ciliated Cell |
| Ccdc19 | 2.05 | 0.00 | 0.00 | 0.74 | 0.01 | 17 | Ciliated Cell |
| Tubb4b | 2.04 | 0.00 | 0.00 | 0.88 | 0.18 | 17 | Ciliated Cell |
| Enkur | 2.02 | 0.00 | 0.00 | 0.78 | 0.01 | 17 | Ciliated Cell |
| Traf3ip1 | 2.01 | 0.00 | 0.00 | 0.78 | 0.03 | 17 | Ciliated Cell |
| Spa17 | 2.01 | 0.00 | 0.00 | 0.75 | 0.02 | 17 | Ciliated Cell |
| Ccdc146 | 2.00 | 0.00 | 0.00 | 0.70 | 0.00 | 17 | Ciliated Cell |
| Mlf1 | 1.97 | 0.00 | 0.00 | 0.75 | 0.01 | 17 | Ciliated Cell |
| Cyp2f2 | 1.96 | 0.00 | 0.00 | 0.58 | 0.02 | 17 | Ciliated Cell |
| Lrriq1 | 1.94 | 0.00 | 0.00 | 0.68 | 0.00 | 17 | Ciliated Cell |
| Ccdc11 | 1.92 | 0.00 | 0.00 | 0.66 | 0.00 | 17 | Ciliated Cell |
| Sntn | 1.91 | 0.00 | 0.00 | 0.58 | 0.00 | 17 | Ciliated Cell |
| Iqcg | 1.90 | 0.00 | 0.00 | 0.71 | 0.01 | 17 | Ciliated Cell |
| Hsp90aa1 | 1.89 | 0.00 | 0.00 | 0.96 | 0.47 | 17 | Ciliated Cell |
| Elof1 | 1.87 | 0.00 | 0.00 | 0.82 | 0.10 | 17 | Ciliated Cell |
| 1110017D15Rik | 1.86 | 0.00 | 0.00 | 0.70 | 0.00 | 17 | Ciliated Cell |
| Anxa1 | 1.85 | 0.00 | 0.00 | 0.87 | 0.18 | 17 | Ciliated Cell |
| Riiad1 | 1.84 | 0.00 | 0.00 | 0.68 | 0.01 | 17 | Ciliated Cell |
| 3300002A11Rik | 1.84 | 0.00 | 0.00 | 0.64 | 0.00 | 17 | Ciliated Cell |
| BC051019 | 1.83 | 0.00 | 0.00 | 0.70 | 0.00 | 17 | Ciliated Cell |
| 1110004E09Rik | 1.83 | 0.00 | 0.00 | 0.77 | 0.07 | 17 | Ciliated Cell |
| Tcea3 | 1.81 | 0.00 | 0.00 | 0.77 | 0.06 | 17 | Ciliated Cell |
| Mt2 | 1.81 | 0.00 | 0.00 | 0.59 | 0.06 | 17 | Ciliated Cell |
| Dnah12 | 1.79 | 0.00 | 0.00 | 0.65 | 0.00 | 17 | Ciliated Cell |
| Ak7 | 1.77 | 0.00 | 0.00 | 0.67 | 0.00 | 17 | Ciliated Cell |
| Erich2 | 1.76 | 0.00 | 0.00 | 0.67 | 0.01 | 17 | Ciliated Cell |
| Nme5 | 1.76 | 0.00 | 0.00 | 0.68 | 0.00 | 17 | Ciliated Cell |
| Drc1 | 1.75 | 0.00 | 0.00 | 0.62 | 0.00 | 17 | Ciliated Cell |
| Igfbp5 | 1.75 | 0.00 | 0.00 | 0.87 | 0.09 | 17 | Ciliated Cell |
| 1600029I14Rik | 1.75 | 0.00 | 0.00 | 0.64 | 0.00 | 17 | Ciliated Cell |
| Wfdc2 | 1.75 | 0.00 | 0.00 | 0.91 | 0.15 | 17 | Ciliated Cell |
| Ezr | 1.74 | 0.00 | 0.00 | 0.90 | 0.21 | 17 | Ciliated Cell |
| Ccdc17 | 1.73 | 0.00 | 0.00 | 0.63 | 0.01 | 17 | Ciliated Cell |
| Krt8 | 1.71 | 0.00 | 0.00 | 0.73 | 0.07 | 17 | Ciliated Cell |
| Lrrc23 | 1.71 | 0.00 | 0.00 | 0.60 | 0.00 | 17 | Ciliated Cell |
| Tspan1 | 1.70 | 0.00 | 0.00 | 0.68 | 0.03 | 17 | Ciliated Cell |
| AK129341 | 1.69 | 0.00 | 0.00 | 0.66 | 0.01 | 17 | Ciliated Cell |
| Calml4 | 1.68 | 0.00 | 0.00 | 0.61 | 0.02 | 17 | Ciliated Cell |
| Cldn3 | 1.67 | 0.00 | 0.00 | 0.82 | 0.11 | 17 | Ciliated Cell |
| Hp | 1.65 | 0.00 | 0.00 | 0.68 | 0.10 | 17 | Ciliated Cell |
| Wdr60 | 1.65 | 0.00 | 0.00 | 0.73 | 0.07 | 17 | Ciliated Cell |
| Rsph4a | 1.62 | 0.00 | 0.00 | 0.61 | 0.00 | 17 | Ciliated Cell |
| 1700007K13Rik | 1.62 | 0.00 | 0.00 | 0.62 | 0.00 | 17 | Ciliated Cell |
| Tmem107 | 1.62 | 0.00 | 0.00 | 0.66 | 0.03 | 17 | Ciliated Cell |
| Ccdc41 | 1.61 | 0.00 | 0.00 | 0.81 | 0.16 | 17 | Ciliated Cell |
| Hspa2 | 1.60 | 0.00 | 0.00 | 0.61 | 0.01 | 17 | Ciliated Cell |
| Meig1 | 1.59 | 0.00 | 0.00 | 0.60 | 0.00 | 17 | Ciliated Cell |
| Cep290 | 1.59 | 0.00 | 0.00 | 0.72 | 0.08 | 17 | Ciliated Cell |
| 2610028H24Rik | 1.58 | 0.00 | 0.00 | 0.62 | 0.01 | 17 | Ciliated Cell |
| Ces1d | 1.58 | 0.00 | 0.00 | 0.83 | 0.16 | 17 | Ciliated Cell |
| E230008N13Rik | 1.57 | 0.00 | 0.00 | 0.59 | 0.01 | 17 | Ciliated Cell |
| Lrrc48 | 1.57 | 0.00 | 0.00 | 0.62 | 0.00 | 17 | Ciliated Cell |
| Ift74 | 1.55 | 0.00 | 0.00 | 0.71 | 0.09 | 17 | Ciliated Cell |
| Ccdc176 | 1.54 | 0.00 | 0.00 | 0.59 | 0.01 | 17 | Ciliated Cell |
| Osbpl6 | 1.54 | 0.00 | 0.00 | 0.67 | 0.04 | 17 | Ciliated Cell |
| Nek5 | 1.54 | 0.00 | 0.00 | 0.59 | 0.00 | 17 | Ciliated Cell |
| Dnah5 | 1.52 | 0.00 | 0.00 | 0.55 | 0.00 | 17 | Ciliated Cell |
| 1700094D03Rik | 1.52 | 0.00 | 0.00 | 0.55 | 0.02 | 17 | Ciliated Cell |
| Lrrc71 | 1.52 | 0.00 | 0.00 | 0.56 | 0.00 | 17 | Ciliated Cell |
| Efcab10 | 1.52 | 0.00 | 0.00 | 0.57 | 0.00 | 17 | Ciliated Cell |
| Mns1 | 1.51 | 0.00 | 0.00 | 0.61 | 0.03 | 17 | Ciliated Cell |
| Aldh3b1 | 1.51 | 0.00 | 0.00 | 0.58 | 0.02 | 17 | Ciliated Cell |
| Lrrc6 | 1.50 | 0.00 | 0.00 | 0.58 | 0.00 | 17 | Ciliated Cell |
| Ak9 | 1.50 | 0.00 | 0.00 | 0.55 | 0.00 | 17 | Ciliated Cell |
| Akap14 | 1.49 | 0.00 | 0.00 | 0.54 | 0.00 | 17 | Ciliated Cell |
| Ccdc78 | 1.49 | 0.00 | 0.00 | 0.54 | 0.00 | 17 | Ciliated Cell |
| Lrrc51 | 1.49 | 0.00 | 0.00 | 0.57 | 0.01 | 17 | Ciliated Cell |
| Ccdc104 | 1.48 | 0.00 | 0.00 | 0.76 | 0.17 | 17 | Ciliated Cell |
| Fam213a | 1.48 | 0.00 | 0.00 | 0.65 | 0.07 | 17 | Ciliated Cell |
| Mettl7a1 | 1.48 | 0.00 | 0.00 | 0.84 | 0.21 | 17 | Ciliated Cell |
| Kif21a | 1.48 | 0.00 | 0.00 | 0.59 | 0.05 | 17 | Ciliated Cell |
| 1700003M02Rik | 1.46 | 0.00 | 0.00 | 0.57 | 0.00 | 17 | Ciliated Cell |
| Tekt1 | 1.46 | 0.00 | 0.00 | 0.58 | 0.00 | 17 | Ciliated Cell |
| Rsph9 | 1.46 | 0.00 | 0.00 | 0.61 | 0.01 | 17 | Ciliated Cell |
| Cldn7 | 1.45 | 0.00 | 0.00 | 0.68 | 0.07 | 17 | Ciliated Cell |
| Gm973 | 1.45 | 0.00 | 0.00 | 0.53 | 0.00 | 17 | Ciliated Cell |
| Tubb4b-ps1 | 1.41 | 0.00 | 0.00 | 0.65 | 0.08 | 17 | Ciliated Cell |
| Smim22 | 1.41 | 0.00 | 0.00 | 0.60 | 0.03 | 17 | Ciliated Cell |
| A330021E22Rik | 1.41 | 0.00 | 0.00 | 0.58 | 0.01 | 17 | Ciliated Cell |
| Hdc | 1.41 | 0.00 | 0.00 | 0.62 | 0.03 | 17 | Ciliated Cell |
| Bphl | 1.40 | 0.00 | 0.00 | 0.62 | 0.04 | 17 | Ciliated Cell |
| Tctex1d4 | 1.40 | 0.00 | 0.00 | 0.54 | 0.00 | 17 | Ciliated Cell |
| Dcxr | 1.39 | 0.00 | 0.00 | 0.69 | 0.10 | 17 | Ciliated Cell |
| Nudc | 1.39 | 0.00 | 0.00 | 0.78 | 0.20 | 17 | Ciliated Cell |
| Odf3b | 1.38 | 0.00 | 0.00 | 0.54 | 0.00 | 17 | Ciliated Cell |
| Ppp1r36 | 1.37 | 0.00 | 0.00 | 0.52 | 0.00 | 17 | Ciliated Cell |
| Ift81 | 1.37 | 0.00 | 0.00 | 0.63 | 0.06 | 17 | Ciliated Cell |
| Ccdc37 | 1.36 | 0.00 | 0.00 | 0.50 | 0.00 | 17 | Ciliated Cell |
| Ttc25 | 1.36 | 0.00 | 0.00 | 0.52 | 0.00 | 17 | Ciliated Cell |
| Stk33 | 1.35 | 0.00 | 0.00 | 0.51 | 0.00 | 17 | Ciliated Cell |
| Ift88 | 1.35 | 0.00 | 0.00 | 0.59 | 0.04 | 17 | Ciliated Cell |
| Ascc1 | 1.34 | 0.00 | 0.00 | 0.60 | 0.04 | 17 | Ciliated Cell |
| Fam154b | 1.34 | 0.00 | 0.00 | 0.53 | 0.00 | 17 | Ciliated Cell |
| Myb | 1.34 | 0.00 | 0.00 | 0.51 | 0.01 | 17 | Ciliated Cell |
| Ccdc30 | 1.33 | 0.00 | 0.00 | 0.53 | 0.00 | 17 | Ciliated Cell |
| Cetn2 | 1.33 | 0.00 | 0.00 | 0.74 | 0.18 | 17 | Ciliated Cell |
| Fam216b | 1.32 | 0.00 | 0.00 | 0.40 | 0.00 | 17 | Ciliated Cell |
| Dnah6 | 1.32 | 0.00 | 0.00 | 0.47 | 0.00 | 17 | Ciliated Cell |
| Mycbp | 1.32 | 0.00 | 0.00 | 0.69 | 0.11 | 17 | Ciliated Cell |
| Wdr78 | 1.32 | 0.00 | 0.00 | 0.52 | 0.01 | 17 | Ciliated Cell |
| Strbp | 1.31 | 0.00 | 0.00 | 0.67 | 0.11 | 17 | Ciliated Cell |
| Smim5 | 1.31 | 0.00 | 0.00 | 0.53 | 0.01 | 17 | Ciliated Cell |
| Calm1 | 1.31 | 0.00 | 0.00 | 1.00 | 0.86 | 17 | Ciliated Cell |
| Gsta4 | 1.31 | 0.00 | 0.00 | 0.55 | 0.03 | 17 | Ciliated Cell |
| Efcab1 | 1.30 | 0.00 | 0.00 | 0.51 | 0.00 | 17 | Ciliated Cell |
| Fank1 | 1.30 | 0.00 | 0.00 | 0.50 | 0.00 | 17 | Ciliated Cell |
| Cdh1 | 1.29 | 0.00 | 0.00 | 0.61 | 0.07 | 17 | Ciliated Cell |
| Tm4sf1 | 1.29 | 0.00 | 0.00 | 0.87 | 0.20 | 17 | Ciliated Cell |
| Zc2hc1a | 1.29 | 0.00 | 0.00 | 0.59 | 0.05 | 17 | Ciliated Cell |
| Ccdc108 | 1.28 | 0.00 | 0.00 | 0.48 | 0.00 | 17 | Ciliated Cell |
| Dnali1 | 1.27 | 0.00 | 0.00 | 0.49 | 0.00 | 17 | Ciliated Cell |
| Capsl | 1.26 | 0.00 | 0.00 | 0.49 | 0.00 | 17 | Ciliated Cell |
| 2610015P09Rik | 1.26 | 0.00 | 0.00 | 0.55 | 0.04 | 17 | Ciliated Cell |
| Dpcd | 1.25 | 0.00 | 0.00 | 0.60 | 0.06 | 17 | Ciliated Cell |
| 1700024G13Rik | 1.25 | 0.00 | 0.00 | 0.39 | 0.00 | 17 | Ciliated Cell |
| 1700088E04Rik | 1.25 | 0.00 | 0.00 | 0.53 | 0.01 | 17 | Ciliated Cell |
| Cxcl17 | 1.25 | 0.00 | 0.00 | 0.39 | 0.01 | 17 | Ciliated Cell |
| Gpx8 | 1.25 | 0.00 | 0.00 | 0.67 | 0.13 | 17 | Ciliated Cell |
| Anxa8 | 1.24 | 0.00 | 0.00 | 0.46 | 0.01 | 17 | Ciliated Cell |
| Mcee | 1.24 | 0.00 | 0.00 | 0.62 | 0.08 | 17 | Ciliated Cell |
| Krt18 | 1.23 | 0.00 | 0.00 | 0.60 | 0.07 | 17 | Ciliated Cell |
| Ttll6 | 1.23 | 0.00 | 0.00 | 0.48 | 0.00 | 17 | Ciliated Cell |
| Il18r1 | 1.23 | 0.00 | 0.00 | 0.58 | 0.06 | 17 | Ciliated Cell |
| Cdkl2 | 1.23 | 0.00 | 0.00 | 0.49 | 0.02 | 17 | Ciliated Cell |
| 1110032A03Rik | 1.23 | 0.00 | 0.00 | 0.57 | 0.06 | 17 | Ciliated Cell |
| Kif27 | 1.23 | 0.00 | 0.00 | 0.48 | 0.00 | 17 | Ciliated Cell |
| Pifo | 1.22 | 0.00 | 0.00 | 0.48 | 0.00 | 17 | Ciliated Cell |
| Spint2 | 1.22 | 0.00 | 0.00 | 0.68 | 0.10 | 17 | Ciliated Cell |
| Ppil6 | 1.21 | 0.00 | 0.00 | 0.49 | 0.00 | 17 | Ciliated Cell |
| Cd55 | 1.20 | 0.00 | 0.00 | 0.60 | 0.07 | 17 | Ciliated Cell |
| Nphp1 | 1.19 | 0.00 | 0.00 | 0.54 | 0.05 | 17 | Ciliated Cell |
| 1700026D08Rik | 1.19 | 0.00 | 0.00 | 0.47 | 0.00 | 17 | Ciliated Cell |
| Spag6 | 1.19 | 0.00 | 0.00 | 0.47 | 0.00 | 17 | Ciliated Cell |
| Chchd6 | 1.18 | 0.00 | 0.00 | 0.51 | 0.03 | 17 | Ciliated Cell |
| Cspp1 | 1.18 | 0.00 | 0.00 | 0.69 | 0.15 | 17 | Ciliated Cell |
| Prr15l | 1.18 | 0.00 | 0.00 | 0.56 | 0.05 | 17 | Ciliated Cell |
| Pih1d2 | 1.18 | 0.00 | 0.00 | 0.48 | 0.00 | 17 | Ciliated Cell |
| Dnah3 | 1.18 | 0.00 | 0.00 | 0.42 | 0.00 | 17 | Ciliated Cell |
| Lrrc34 | 1.18 | 0.00 | 0.00 | 0.46 | 0.00 | 17 | Ciliated Cell |
| Gipc2 | 1.18 | 0.00 | 0.00 | 0.53 | 0.03 | 17 | Ciliated Cell |
| Mak | 1.17 | 0.00 | 0.00 | 0.46 | 0.00 | 17 | Ciliated Cell |
| 1700001C02Rik | 1.17 | 0.00 | 0.00 | 0.46 | 0.00 | 17 | Ciliated Cell |
| Dpy30 | 1.17 | 0.00 | 0.00 | 0.64 | 0.13 | 17 | Ciliated Cell |
| 1700026L06Rik | 1.17 | 0.00 | 0.00 | 0.44 | 0.00 | 17 | Ciliated Cell |
| Clic6 | 1.16 | 0.00 | 0.00 | 0.49 | 0.01 | 17 | Ciliated Cell |
| 1600029D21Rik | 1.15 | 0.00 | 0.00 | 0.48 | 0.02 | 17 | Ciliated Cell |
| Rsph10b | 1.15 | 0.00 | 0.00 | 0.43 | 0.00 | 17 | Ciliated Cell |
| Acsl3 | 1.15 | 0.00 | 0.00 | 0.52 | 0.04 | 17 | Ciliated Cell |
| Ccdc40 | 1.15 | 0.00 | 0.00 | 0.43 | 0.00 | 17 | Ciliated Cell |
| 1700007G11Rik | 1.15 | 0.00 | 0.00 | 0.45 | 0.00 | 17 | Ciliated Cell |
| Fam216a | 1.15 | 0.00 | 0.00 | 0.52 | 0.05 | 17 | Ciliated Cell |
| Ccdc114 | 1.14 | 0.00 | 0.00 | 0.45 | 0.01 | 17 | Ciliated Cell |
| Cetn4 | 1.14 | 0.00 | 0.00 | 0.46 | 0.01 | 17 | Ciliated Cell |
| Cdc14a | 1.14 | 0.00 | 0.00 | 0.46 | 0.03 | 17 | Ciliated Cell |
| Agr3 | 1.14 | 0.00 | 0.00 | 0.43 | 0.00 | 17 | Ciliated Cell |
| Krt19 | 1.14 | 0.00 | 0.00 | 0.56 | 0.06 | 17 | Ciliated Cell |
| Map1b | 1.14 | 0.00 | 0.00 | 0.60 | 0.09 | 17 | Ciliated Cell |
| Clic3 | 1.13 | 0.00 | 0.00 | 0.58 | 0.08 | 17 | Ciliated Cell |
| Ccdc173 | 1.13 | 0.00 | 0.00 | 0.45 | 0.02 | 17 | Ciliated Cell |
| Cdhr3 | 1.13 | 0.00 | 0.00 | 0.40 | 0.00 | 17 | Ciliated Cell |
| Tekt2 | 1.12 | 0.00 | 0.00 | 0.43 | 0.00 | 17 | Ciliated Cell |
| Ift43 | 1.12 | 0.00 | 0.00 | 0.61 | 0.09 | 17 | Ciliated Cell |
| Map9 | 1.12 | 0.00 | 0.00 | 0.46 | 0.02 | 17 | Ciliated Cell |
| Gm5918 | 1.12 | 0.00 | 0.00 | 0.45 | 0.01 | 17 | Ciliated Cell |
| Adgb | 1.12 | 0.00 | 0.00 | 0.43 | 0.00 | 17 | Ciliated Cell |
| Rsph3a | 1.12 | 0.00 | 0.00 | 0.53 | 0.06 | 17 | Ciliated Cell |
| Gm872 | 1.12 | 0.00 | 0.00 | 0.40 | 0.00 | 17 | Ciliated Cell |
| Iqca | 1.12 | 0.00 | 0.00 | 0.39 | 0.00 | 17 | Ciliated Cell |
| Myh14 | 1.11 | 0.00 | 0.00 | 0.61 | 0.09 | 17 | Ciliated Cell |
| Rfk | 1.11 | 0.00 | 0.00 | 0.57 | 0.09 | 17 | Ciliated Cell |
| Ttc21a | 1.10 | 0.00 | 0.00 | 0.42 | 0.00 | 17 | Ciliated Cell |
| Kif3a | 1.10 | 0.00 | 0.00 | 0.60 | 0.10 | 17 | Ciliated Cell |
| Ttll3 | 1.10 | 0.00 | 0.00 | 0.44 | 0.01 | 17 | Ciliated Cell |
| Net1 | 1.10 | 0.00 | 0.00 | 0.73 | 0.16 | 17 | Ciliated Cell |
| Cystm1 | 1.09 | 0.00 | 0.00 | 0.58 | 0.07 | 17 | Ciliated Cell |
| Stmnd1 | 1.09 | 0.00 | 0.00 | 0.36 | 0.00 | 17 | Ciliated Cell |
| Aldh1a7 | 1.09 | 0.00 | 0.00 | 0.45 | 0.04 | 17 | Ciliated Cell |
| S100a1 | 1.09 | 0.00 | 0.00 | 0.65 | 0.14 | 17 | Ciliated Cell |
| 4833427G06Rik | 1.09 | 0.00 | 0.00 | 0.43 | 0.00 | 17 | Ciliated Cell |
| Areg | 1.09 | 0.00 | 0.00 | 0.33 | 0.02 | 17 | Ciliated Cell |
| Dmkn | 1.09 | 0.00 | 0.00 | 0.42 | 0.01 | 17 | Ciliated Cell |
| Fam167a | 1.09 | 0.00 | 0.00 | 0.41 | 0.01 | 17 | Ciliated Cell |
| Rsph3b | 1.08 | 0.00 | 0.00 | 0.51 | 0.06 | 17 | Ciliated Cell |
| Lca5 | 1.08 | 0.00 | 0.00 | 0.44 | 0.03 | 17 | Ciliated Cell |
| Epcam | 1.08 | 0.00 | 0.00 | 0.61 | 0.09 | 17 | Ciliated Cell |
| Gm7094 | 1.07 | 0.00 | 0.00 | 0.42 | 0.01 | 17 | Ciliated Cell |
| Spata18 | 1.07 | 0.00 | 0.00 | 0.39 | 0.00 | 17 | Ciliated Cell |
| Gas8 | 1.06 | 0.00 | 0.00 | 0.49 | 0.03 | 17 | Ciliated Cell |
| Wdr16 | 1.06 | 0.00 | 0.00 | 0.41 | 0.00 | 17 | Ciliated Cell |
| Tmprss2 | 1.06 | 0.00 | 0.00 | 0.45 | 0.02 | 17 | Ciliated Cell |
| Mettl7a2 | 1.06 | 0.00 | 0.00 | 0.58 | 0.10 | 17 | Ciliated Cell |
| Vwa3a | 1.06 | 0.00 | 0.00 | 0.40 | 0.00 | 17 | Ciliated Cell |
| Pla2g16 | 1.06 | 0.00 | 0.00 | 0.50 | 0.04 | 17 | Ciliated Cell |
| Sema3e | 1.06 | 0.00 | 0.00 | 0.50 | 0.04 | 17 | Ciliated Cell |
| 6820408C15Rik | 1.05 | 0.00 | 0.00 | 0.41 | 0.00 | 17 | Ciliated Cell |
| Maats1 | 1.05 | 0.00 | 0.00 | 0.42 | 0.01 | 17 | Ciliated Cell |
| Trp53bp2 | 1.05 | 0.00 | 0.00 | 0.51 | 0.08 | 17 | Ciliated Cell |
| Aebp1 | 1.05 | 0.00 | 0.00 | 0.51 | 0.07 | 17 | Ciliated Cell |
| Dync2li1 | 1.05 | 0.00 | 0.00 | 0.46 | 0.03 | 17 | Ciliated Cell |
| Phtf1 | 1.04 | 0.00 | 0.00 | 0.48 | 0.04 | 17 | Ciliated Cell |
| 1700012B09Rik | 1.04 | 0.00 | 0.00 | 0.35 | 0.00 | 17 | Ciliated Cell |
| Slc23a1 | 1.03 | 0.00 | 0.00 | 0.39 | 0.01 | 17 | Ciliated Cell |
| 1500015O10Rik | 1.03 | 0.00 | 0.00 | 0.36 | 0.01 | 17 | Ciliated Cell |
| Prr18 | 1.03 | 0.00 | 0.00 | 0.42 | 0.02 | 17 | Ciliated Cell |
| Ubxn10 | 1.03 | 0.00 | 0.00 | 0.41 | 0.00 | 17 | Ciliated Cell |
| Arl3 | 1.03 | 0.00 | 0.00 | 0.60 | 0.12 | 17 | Ciliated Cell |
| Ttc18 | 1.02 | 0.00 | 0.00 | 0.39 | 0.00 | 17 | Ciliated Cell |
| Fam47e | 1.02 | 0.00 | 0.00 | 0.37 | 0.00 | 17 | Ciliated Cell |
| Spef1 | 1.02 | 0.00 | 0.00 | 0.47 | 0.03 | 17 | Ciliated Cell |
| Pdgfa | 1.02 | 0.00 | 0.00 | 0.59 | 0.10 | 17 | Ciliated Cell |
| Ankrd65 | 1.02 | 0.00 | 0.00 | 0.36 | 0.00 | 17 | Ciliated Cell |
| Gm867 | 1.01 | 0.00 | 0.00 | 0.35 | 0.00 | 17 | Ciliated Cell |
| Dcdc2b | 1.01 | 0.00 | 0.00 | 0.41 | 0.01 | 17 | Ciliated Cell |
| Ankrd42 | 1.01 | 0.00 | 0.00 | 0.42 | 0.01 | 17 | Ciliated Cell |
| Wdr63 | 1.01 | 0.00 | 0.00 | 0.39 | 0.00 | 17 | Ciliated Cell |
| Ift57 | 1.01 | 0.00 | 0.00 | 0.45 | 0.03 | 17 | Ciliated Cell |
| Sord | 1.01 | 0.00 | 0.00 | 0.43 | 0.03 | 17 | Ciliated Cell |
| Dync2h1 | 1.00 | 0.00 | 0.00 | 0.44 | 0.03 | 17 | Ciliated Cell |
| 4930451C15Rik | 1.00 | 0.00 | 0.00 | 0.36 | 0.00 | 17 | Ciliated Cell |
| Ttc29 | 0.99 | 0.00 | 0.00 | 0.37 | 0.00 | 17 | Ciliated Cell |
| Dusp18 | 0.99 | 0.00 | 0.00 | 0.42 | 0.01 | 17 | Ciliated Cell |
| Dusp14 | 0.99 | 0.00 | 0.00 | 0.43 | 0.02 | 17 | Ciliated Cell |
| Arhgef28 | 0.98 | 0.00 | 0.00 | 0.44 | 0.03 | 17 | Ciliated Cell |
| B9d1 | 0.98 | 0.00 | 0.00 | 0.43 | 0.02 | 17 | Ciliated Cell |
| Fsd1l | 0.97 | 0.00 | 0.00 | 0.42 | 0.02 | 17 | Ciliated Cell |
| Dzip1l | 0.97 | 0.00 | 0.00 | 0.43 | 0.03 | 17 | Ciliated Cell |
| Rabgap1l | 0.96 | 0.00 | 0.00 | 0.56 | 0.10 | 17 | Ciliated Cell |
| Fam84b | 0.96 | 0.00 | 0.00 | 0.43 | 0.05 | 17 | Ciliated Cell |
| Rabl2 | 0.95 | 0.00 | 0.00 | 0.43 | 0.02 | 17 | Ciliated Cell |
| Scnn1a | 0.95 | 0.00 | 0.00 | 0.52 | 0.06 | 17 | Ciliated Cell |
| Appl2 | 0.95 | 0.00 | 0.00 | 0.47 | 0.06 | 17 | Ciliated Cell |
| Ldhb | 0.95 | 0.00 | 0.00 | 0.50 | 0.07 | 17 | Ciliated Cell |
| Tob1 | 0.94 | 0.00 | 0.00 | 0.46 | 0.06 | 17 | Ciliated Cell |
| Slc25a17 | 0.94 | 0.00 | 0.00 | 0.53 | 0.08 | 17 | Ciliated Cell |
| Hydin | 0.94 | 0.00 | 0.00 | 0.33 | 0.00 | 17 | Ciliated Cell |
| Fam81a | 0.94 | 0.00 | 0.00 | 0.38 | 0.02 | 17 | Ciliated Cell |
| Syt5 | 0.94 | 0.00 | 0.00 | 0.33 | 0.00 | 17 | Ciliated Cell |
| Dcdc2a | 0.94 | 0.00 | 0.00 | 0.35 | 0.00 | 17 | Ciliated Cell |
| Elmod1 | 0.93 | 0.00 | 0.00 | 0.33 | 0.00 | 17 | Ciliated Cell |
| Nek1 | 0.93 | 0.00 | 0.00 | 0.44 | 0.05 | 17 | Ciliated Cell |
| Lmo7 | 0.93 | 0.00 | 0.00 | 0.64 | 0.12 | 17 | Ciliated Cell |
| Zbbx | 0.93 | 0.00 | 0.00 | 0.36 | 0.00 | 17 | Ciliated Cell |
| Zdhhc1 | 0.93 | 0.00 | 0.00 | 0.40 | 0.03 | 17 | Ciliated Cell |
| Rpgr | 0.92 | 0.00 | 0.00 | 0.40 | 0.02 | 17 | Ciliated Cell |
| Plcb3 | 0.92 | 0.00 | 0.00 | 0.45 | 0.06 | 17 | Ciliated Cell |
| Cyp2b10 | 0.91 | 0.00 | 0.00 | 0.32 | 0.00 | 17 | Ciliated Cell |
| Tjp3 | 0.91 | 0.00 | 0.00 | 0.37 | 0.02 | 17 | Ciliated Cell |
| 2410004P03Rik | 0.91 | 0.00 | 0.00 | 0.36 | 0.00 | 17 | Ciliated Cell |
| Gm166 | 0.91 | 0.00 | 0.00 | 0.39 | 0.01 | 17 | Ciliated Cell |
| 1700023L04Rik | 0.91 | 0.00 | 0.00 | 0.37 | 0.02 | 17 | Ciliated Cell |
| Tekt4 | 0.91 | 0.00 | 0.00 | 0.34 | 0.00 | 17 | Ciliated Cell |
| Gm1673 | 0.91 | 0.00 | 0.00 | 0.44 | 0.04 | 17 | Ciliated Cell |
| Aig1 | 0.91 | 0.00 | 0.00 | 0.50 | 0.06 | 17 | Ciliated Cell |
| 1700001L19Rik | 0.90 | 0.00 | 0.00 | 0.37 | 0.00 | 17 | Ciliated Cell |
| Eml2 | 0.90 | 0.00 | 0.00 | 0.40 | 0.03 | 17 | Ciliated Cell |
| Oscp1 | 0.90 | 0.00 | 0.00 | 0.39 | 0.01 | 17 | Ciliated Cell |
| Gsto1 | 0.90 | 0.00 | 0.00 | 0.46 | 0.05 | 17 | Ciliated Cell |
| Ccdc96 | 0.90 | 0.00 | 0.00 | 0.33 | 0.00 | 17 | Ciliated Cell |
| Cds1 | 0.90 | 0.00 | 0.00 | 0.39 | 0.02 | 17 | Ciliated Cell |
| Mdm1 | 0.90 | 0.00 | 0.00 | 0.38 | 0.03 | 17 | Ciliated Cell |
| Ttc39a | 0.90 | 0.00 | 0.00 | 0.35 | 0.00 | 17 | Ciliated Cell |
| Smim6 | 0.89 | 0.00 | 0.00 | 0.40 | 0.03 | 17 | Ciliated Cell |
| Dnah9 | 0.89 | 0.00 | 0.00 | 0.37 | 0.00 | 17 | Ciliated Cell |
| Mapk15 | 0.89 | 0.00 | 0.00 | 0.36 | 0.00 | 17 | Ciliated Cell |
| Ccdc170 | 0.88 | 0.00 | 0.00 | 0.33 | 0.00 | 17 | Ciliated Cell |
| Gm216 | 0.88 | 0.00 | 0.00 | 0.31 | 0.00 | 17 | Ciliated Cell |
| Ccdc65 | 0.88 | 0.00 | 0.00 | 0.35 | 0.00 | 17 | Ciliated Cell |
| Rpgrip1l | 0.88 | 0.00 | 0.00 | 0.37 | 0.02 | 17 | Ciliated Cell |
| Spata24 | 0.88 | 0.00 | 0.00 | 0.37 | 0.03 | 17 | Ciliated Cell |
| Spata17 | 0.87 | 0.00 | 0.00 | 0.33 | 0.00 | 17 | Ciliated Cell |
| Porcn | 0.87 | 0.00 | 0.00 | 0.38 | 0.02 | 17 | Ciliated Cell |
| Ttc12 | 0.87 | 0.00 | 0.00 | 0.35 | 0.01 | 17 | Ciliated Cell |
| Cyb561 | 0.86 | 0.00 | 0.00 | 0.42 | 0.03 | 17 | Ciliated Cell |
| Odf2l | 0.86 | 0.00 | 0.00 | 0.40 | 0.05 | 17 | Ciliated Cell |
| 1700028P14Rik | 0.86 | 0.00 | 0.00 | 0.33 | 0.00 | 17 | Ciliated Cell |
| Wwc1 | 0.85 | 0.00 | 0.00 | 0.44 | 0.06 | 17 | Ciliated Cell |
| 1700019L03Rik | 0.85 | 0.00 | 0.00 | 0.28 | 0.00 | 17 | Ciliated Cell |
| Hmgcs2 | 0.85 | 0.00 | 0.00 | 0.33 | 0.01 | 17 | Ciliated Cell |
| Ebp | 0.85 | 0.00 | 0.00 | 0.48 | 0.07 | 17 | Ciliated Cell |
| Tctex1d2 | 0.85 | 0.00 | 0.00 | 0.36 | 0.03 | 17 | Ciliated Cell |
| Pcbd1 | 0.85 | 0.00 | 0.00 | 0.38 | 0.02 | 17 | Ciliated Cell |
| Hs6st2 | 0.85 | 0.00 | 0.00 | 0.33 | 0.01 | 17 | Ciliated Cell |
| Rmdn1 | 0.84 | 0.00 | 0.00 | 0.44 | 0.05 | 17 | Ciliated Cell |
| Zmynd10 | 0.84 | 0.00 | 0.00 | 0.35 | 0.00 | 17 | Ciliated Cell |
| Fermt1 | 0.84 | 0.00 | 0.00 | 0.27 | 0.00 | 17 | Ciliated Cell |
| Sox2 | 0.83 | 0.00 | 0.00 | 0.31 | 0.00 | 17 | Ciliated Cell |
| Slc39a4 | 0.83 | 0.00 | 0.00 | 0.34 | 0.01 | 17 | Ciliated Cell |
| Dnaic2 | 0.83 | 0.00 | 0.00 | 0.32 | 0.00 | 17 | Ciliated Cell |
| Rnf128 | 0.83 | 0.00 | 0.00 | 0.33 | 0.01 | 17 | Ciliated Cell |
| Wdr65 | 0.83 | 0.00 | 0.00 | 0.33 | 0.00 | 17 | Ciliated Cell |
| Tex9 | 0.83 | 0.00 | 0.00 | 0.38 | 0.03 | 17 | Ciliated Cell |
| Stim1 | 0.83 | 0.00 | 0.00 | 0.42 | 0.06 | 17 | Ciliated Cell |
| Foxa2 | 0.82 | 0.00 | 0.00 | 0.38 | 0.04 | 17 | Ciliated Cell |
| Myo5c | 0.82 | 0.00 | 0.00 | 0.47 | 0.07 | 17 | Ciliated Cell |
| Bola1 | 0.82 | 0.00 | 0.00 | 0.44 | 0.06 | 17 | Ciliated Cell |
| Agbl2 | 0.82 | 0.00 | 0.00 | 0.32 | 0.00 | 17 | Ciliated Cell |
| Fbxo36 | 0.82 | 0.00 | 0.00 | 0.34 | 0.01 | 17 | Ciliated Cell |
| Cgn | 0.82 | 0.00 | 0.00 | 0.38 | 0.03 | 17 | Ciliated Cell |
| Ptn | 0.82 | 0.00 | 0.00 | 0.33 | 0.03 | 17 | Ciliated Cell |
| Slc23a2 | 0.82 | 0.00 | 0.00 | 0.29 | 0.03 | 17 | Ciliated Cell |
| Cobl | 0.81 | 0.00 | 0.00 | 0.34 | 0.02 | 17 | Ciliated Cell |
| Jag1 | 0.81 | 0.00 | 0.00 | 0.40 | 0.04 | 17 | Ciliated Cell |
| Endog | 0.80 | 0.00 | 0.00 | 0.31 | 0.02 | 17 | Ciliated Cell |
| Gstz1 | 0.80 | 0.00 | 0.00 | 0.38 | 0.04 | 17 | Ciliated Cell |
| Kcnmb2 | 0.80 | 0.00 | 0.00 | 0.32 | 0.00 | 17 | Ciliated Cell |
| Piwil4 | 0.80 | 0.00 | 0.00 | 0.28 | 0.01 | 17 | Ciliated Cell |
| Ccdc135 | 0.80 | 0.00 | 0.00 | 0.30 | 0.00 | 17 | Ciliated Cell |
| 1700040L02Rik | 0.79 | 0.00 | 0.00 | 0.31 | 0.00 | 17 | Ciliated Cell |
| Rabl5 | 0.79 | 0.00 | 0.00 | 0.35 | 0.02 | 17 | Ciliated Cell |
| 1700029J07Rik | 0.79 | 0.00 | 0.00 | 0.32 | 0.01 | 17 | Ciliated Cell |
| Iqub | 0.79 | 0.00 | 0.00 | 0.29 | 0.00 | 17 | Ciliated Cell |
| D430036J16Rik | 0.79 | 0.00 | 0.00 | 0.30 | 0.00 | 17 | Ciliated Cell |
| Fam179a | 0.79 | 0.00 | 0.00 | 0.31 | 0.00 | 17 | Ciliated Cell |
| Ccdc151 | 0.79 | 0.00 | 0.00 | 0.30 | 0.00 | 17 | Ciliated Cell |
| Ccdc81 | 0.78 | 0.00 | 0.00 | 0.28 | 0.00 | 17 | Ciliated Cell |
| Casc1 | 0.78 | 0.00 | 0.00 | 0.30 | 0.01 | 17 | Ciliated Cell |
| Wdr19 | 0.78 | 0.00 | 0.00 | 0.30 | 0.01 | 17 | Ciliated Cell |
| Kcnrg | 0.78 | 0.00 | 0.00 | 0.26 | 0.00 | 17 | Ciliated Cell |
| Fhdc1 | 0.78 | 0.00 | 0.00 | 0.34 | 0.01 | 17 | Ciliated Cell |
| Mr1 | 0.78 | 0.00 | 0.00 | 0.34 | 0.04 | 17 | Ciliated Cell |
| Irak1bp1 | 0.78 | 0.00 | 0.00 | 0.35 | 0.02 | 17 | Ciliated Cell |
| Usp2 | 0.77 | 0.00 | 0.00 | 0.34 | 0.01 | 17 | Ciliated Cell |
| D630024D03Rik | 0.77 | 0.00 | 0.00 | 0.28 | 0.00 | 17 | Ciliated Cell |
| Dzip1 | 0.77 | 0.00 | 0.00 | 0.33 | 0.03 | 17 | Ciliated Cell |
| Dnajb13 | 0.77 | 0.00 | 0.00 | 0.31 | 0.00 | 17 | Ciliated Cell |
| Dnal1 | 0.77 | 0.00 | 0.00 | 0.35 | 0.02 | 17 | Ciliated Cell |
| 9330101J02Rik | 0.76 | 0.00 | 0.00 | 0.29 | 0.00 | 17 | Ciliated Cell |
| Syt10 | 0.76 | 0.00 | 0.00 | 0.27 | 0.00 | 17 | Ciliated Cell |
| Gm16136 | 0.76 | 0.00 | 0.00 | 0.30 | 0.01 | 17 | Ciliated Cell |
| Ccdc162 | 0.76 | 0.00 | 0.00 | 0.28 | 0.00 | 17 | Ciliated Cell |
| 4930556M19Rik | 0.76 | 0.00 | 0.00 | 0.34 | 0.04 | 17 | Ciliated Cell |
| Igfbp2 | 0.75 | 0.00 | 0.00 | 0.33 | 0.02 | 17 | Ciliated Cell |
| Dalrd3 | 0.75 | 0.00 | 0.00 | 0.39 | 0.03 | 17 | Ciliated Cell |
| Gfra1 | 0.75 | 0.00 | 0.00 | 0.30 | 0.01 | 17 | Ciliated Cell |
| Fhit | 0.75 | 0.00 | 0.00 | 0.28 | 0.00 | 17 | Ciliated Cell |
| Rgs22 | 0.75 | 0.00 | 0.00 | 0.28 | 0.00 | 17 | Ciliated Cell |
| Cep164 | 0.74 | 0.00 | 0.00 | 0.39 | 0.05 | 17 | Ciliated Cell |
| Lrrc46 | 0.74 | 0.00 | 0.00 | 0.30 | 0.00 | 17 | Ciliated Cell |
| Dnah11 | 0.74 | 0.00 | 0.00 | 0.29 | 0.01 | 17 | Ciliated Cell |
| Ccdc147 | 0.74 | 0.00 | 0.00 | 0.27 | 0.00 | 17 | Ciliated Cell |
| Vpreb3 | 0.73 | 0.00 | 0.00 | 0.30 | 0.02 | 17 | Ciliated Cell |
| Ubxn11 | 0.73 | 0.00 | 0.00 | 0.32 | 0.01 | 17 | Ciliated Cell |
| Fam13a | 0.72 | 0.00 | 0.00 | 0.32 | 0.02 | 17 | Ciliated Cell |
| Plekhb1 | 0.72 | 0.00 | 0.00 | 0.31 | 0.02 | 17 | Ciliated Cell |
| Ttc30b | 0.72 | 0.00 | 0.00 | 0.32 | 0.01 | 17 | Ciliated Cell |
| Slc44a3 | 0.72 | 0.00 | 0.00 | 0.33 | 0.02 | 17 | Ciliated Cell |
| Mycbpap | 0.72 | 0.00 | 0.00 | 0.27 | 0.00 | 17 | Ciliated Cell |
| Ddr1 | 0.71 | 0.00 | 0.00 | 0.33 | 0.03 | 17 | Ciliated Cell |
| 4930562C15Rik | 0.71 | 0.00 | 0.00 | 0.28 | 0.00 | 17 | Ciliated Cell |
| Ift172 | 0.71 | 0.00 | 0.00 | 0.30 | 0.02 | 17 | Ciliated Cell |
| Fam149a | 0.71 | 0.00 | 0.00 | 0.30 | 0.02 | 17 | Ciliated Cell |
| Pla2g4a | 0.70 | 0.00 | 0.00 | 0.33 | 0.02 | 17 | Ciliated Cell |
| Lypd2 | 0.70 | 0.00 | 0.00 | 0.26 | 0.01 | 17 | Ciliated Cell |
| Tmem216 | 0.70 | 0.00 | 0.00 | 0.34 | 0.03 | 17 | Ciliated Cell |
| Lrrc45 | 0.70 | 0.00 | 0.00 | 0.36 | 0.03 | 17 | Ciliated Cell |
| Slc1a4 | 0.70 | 0.00 | 0.00 | 0.27 | 0.02 | 17 | Ciliated Cell |
| Cdkl4 | 0.69 | 0.00 | 0.00 | 0.27 | 0.00 | 17 | Ciliated Cell |
| Pir | 0.69 | 0.00 | 0.00 | 0.33 | 0.03 | 17 | Ciliated Cell |
| Dnaic1 | 0.69 | 0.00 | 0.00 | 0.25 | 0.01 | 17 | Ciliated Cell |
| Tex26 | 0.69 | 0.00 | 0.00 | 0.25 | 0.00 | 17 | Ciliated Cell |
| Efcab12 | 0.69 | 0.00 | 0.00 | 0.26 | 0.00 | 17 | Ciliated Cell |
| Susd2 | 0.69 | 0.00 | 0.00 | 0.25 | 0.00 | 17 | Ciliated Cell |
| Prom1 | 0.68 | 0.00 | 0.00 | 0.30 | 0.03 | 17 | Ciliated Cell |
| 4430402I18Rik | 0.68 | 0.00 | 0.00 | 0.27 | 0.01 | 17 | Ciliated Cell |
| Pacrg | 0.66 | 0.00 | 0.00 | 0.26 | 0.00 | 17 | Ciliated Cell |
| Eno4 | 0.66 | 0.00 | 0.00 | 0.27 | 0.00 | 17 | Ciliated Cell |
| Wdyhv1 | 0.66 | 0.00 | 0.00 | 0.33 | 0.03 | 17 | Ciliated Cell |
| Rnf186 | 0.66 | 0.00 | 0.00 | 0.27 | 0.01 | 17 | Ciliated Cell |
| Dnaaf3 | 0.66 | 0.00 | 0.00 | 0.30 | 0.00 | 17 | Ciliated Cell |
| Perp | 0.66 | 0.00 | 0.00 | 0.32 | 0.03 | 17 | Ciliated Cell |
| Ribc1 | 0.66 | 0.00 | 0.00 | 0.25 | 0.00 | 17 | Ciliated Cell |
| Gm16119 | 0.66 | 0.00 | 0.00 | 0.26 | 0.00 | 17 | Ciliated Cell |
| Armc3 | 0.66 | 0.00 | 0.00 | 0.25 | 0.01 | 17 | Ciliated Cell |
| Ctxn1 | 0.65 | 0.00 | 0.00 | 0.31 | 0.02 | 17 | Ciliated Cell |
| Cmbl | 0.65 | 0.00 | 0.00 | 0.27 | 0.01 | 17 | Ciliated Cell |
| Tacstd2 | 0.65 | 0.00 | 0.00 | 0.30 | 0.03 | 17 | Ciliated Cell |
| Tnn | 0.65 | 0.00 | 0.00 | 0.27 | 0.00 | 17 | Ciliated Cell |
| Ropn1l | 0.65 | 0.00 | 0.00 | 0.27 | 0.01 | 17 | Ciliated Cell |
| Dnaja4 | 0.64 | 0.00 | 0.00 | 0.27 | 0.01 | 17 | Ciliated Cell |
| Dnah10 | 0.63 | 0.00 | 0.00 | 0.26 | 0.00 | 17 | Ciliated Cell |
| Galnt3 | 0.63 | 0.00 | 0.00 | 0.29 | 0.02 | 17 | Ciliated Cell |
| Wdr35 | 0.63 | 0.00 | 0.00 | 0.29 | 0.02 | 17 | Ciliated Cell |
| Acox2 | 0.63 | 0.00 | 0.00 | 0.26 | 0.00 | 17 | Ciliated Cell |
| Spag8 | 0.61 | 0.00 | 0.00 | 0.26 | 0.00 | 17 | Ciliated Cell |
| Wdr31 | 0.60 | 0.00 | 0.00 | 0.25 | 0.01 | 17 | Ciliated Cell |
| Prkcz | 0.59 | 0.00 | 0.00 | 0.28 | 0.02 | 17 | Ciliated Cell |
| Stk30 | 0.59 | 0.00 | 0.00 | 0.27 | 0.01 | 17 | Ciliated Cell |
| 2310007B03Rik | 0.58 | 0.00 | 0.00 | 0.27 | 0.02 | 17 | Ciliated Cell |
| Arl6 | 0.58 | 0.00 | 0.00 | 0.27 | 0.02 | 17 | Ciliated Cell |
| Foxa1 | 0.58 | 0.00 | 0.00 | 0.26 | 0.02 | 17 | Ciliated Cell |
| Daf2 | 0.56 | 0.00 | 0.00 | 0.28 | 0.02 | 17 | Ciliated Cell |
| Dixdc1 | 0.56 | 0.00 | 0.00 | 0.28 | 0.02 | 17 | Ciliated Cell |
| Wdr34 | 0.55 | 0.00 | 0.00 | 0.25 | 0.01 | 17 | Ciliated Cell |
| Kcnj13 | 0.54 | 0.00 | 0.00 | 0.25 | 0.01 | 17 | Ciliated Cell |
| Pfn2 | 0.51 | 0.00 | 0.00 | 0.27 | 0.02 | 17 | Ciliated Cell |
| Ift27 | 0.73 | 0.00 | 0.00 | 0.41 | 0.06 | 17 | Ciliated Cell |
| Ank3 | 0.80 | 0.00 | 0.00 | 0.47 | 0.07 | 17 | Ciliated Cell |
| B230118H07Rik | 0.59 | 0.00 | 0.00 | 0.31 | 0.03 | 17 | Ciliated Cell |
| Maob | 0.74 | 0.00 | 0.00 | 0.30 | 0.03 | 17 | Ciliated Cell |
| Dnajb2 | 0.57 | 0.00 | 0.00 | 0.27 | 0.02 | 17 | Ciliated Cell |
| Spice1 | 0.70 | 0.00 | 0.00 | 0.35 | 0.04 | 17 | Ciliated Cell |
| Tsga10 | 0.65 | 0.00 | 0.00 | 0.28 | 0.03 | 17 | Ciliated Cell |
| Sdccag8 | 0.77 | 0.00 | 0.00 | 0.39 | 0.05 | 17 | Ciliated Cell |
| Hmgn3 | 0.77 | 0.00 | 0.00 | 0.37 | 0.05 | 17 | Ciliated Cell |
| Bok | 0.54 | 0.00 | 0.00 | 0.26 | 0.02 | 17 | Ciliated Cell |
| Slc9a3r1 | 0.58 | 0.00 | 0.00 | 0.33 | 0.04 | 17 | Ciliated Cell |
| Tmc4 | 0.63 | 0.00 | 0.00 | 0.32 | 0.04 | 17 | Ciliated Cell |
| Fam134b | 0.78 | 0.00 | 0.00 | 0.42 | 0.06 | 17 | Ciliated Cell |
| Gstm1 | 1.03 | 0.00 | 0.00 | 0.64 | 0.14 | 17 | Ciliated Cell |
| Fgfr1op | 0.66 | 0.00 | 0.00 | 0.36 | 0.05 | 17 | Ciliated Cell |
| Gtl3 | 1.04 | 0.00 | 0.00 | 0.66 | 0.16 | 17 | Ciliated Cell |
| Alcam | 1.02 | 0.00 | 0.00 | 0.81 | 0.22 | 17 | Ciliated Cell |
| Dnajc15 | 0.80 | 0.00 | 0.00 | 0.46 | 0.08 | 17 | Ciliated Cell |
| Nudt4 | 1.15 | 0.00 | 0.00 | 0.67 | 0.16 | 17 | Ciliated Cell |
| Sqle | 0.61 | 0.00 | 0.00 | 0.31 | 0.04 | 17 | Ciliated Cell |
| Dynlt1c | 1.19 | 0.00 | 0.00 | 0.79 | 0.26 | 17 | Ciliated Cell |
| Runx1 | 0.78 | 0.00 | 0.00 | 0.42 | 0.07 | 17 | Ciliated Cell |
| Srsf10 | 1.20 | 0.00 | 0.00 | 0.79 | 0.24 | 17 | Ciliated Cell |
| Sorl1 | 0.78 | 0.00 | 0.00 | 0.43 | 0.07 | 17 | Ciliated Cell |
| Hspb11 | 0.75 | 0.00 | 0.00 | 0.41 | 0.06 | 17 | Ciliated Cell |
| Med31 | 0.82 | 0.00 | 0.00 | 0.45 | 0.08 | 17 | Ciliated Cell |
| Cep19 | 0.61 | 0.00 | 0.00 | 0.33 | 0.04 | 17 | Ciliated Cell |
| Dynlt1b | 1.21 | 0.00 | 0.00 | 0.76 | 0.24 | 17 | Ciliated Cell |
| Irx3 | 0.75 | 0.00 | 0.00 | 0.43 | 0.07 | 17 | Ciliated Cell |
| Dynlt1f | 1.18 | 0.00 | 0.00 | 0.75 | 0.23 | 17 | Ciliated Cell |
| Katnal1 | 0.63 | 0.00 | 0.00 | 0.26 | 0.03 | 17 | Ciliated Cell |
| Gch1 | 0.56 | 0.00 | 0.00 | 0.26 | 0.03 | 17 | Ciliated Cell |
| Fam174b | 0.60 | 0.00 | 0.00 | 0.35 | 0.05 | 17 | Ciliated Cell |
| Acot1 | 0.80 | 0.00 | 0.00 | 0.43 | 0.07 | 17 | Ciliated Cell |
| Atp1b1 | 0.86 | 0.00 | 0.00 | 0.72 | 0.18 | 17 | Ciliated Cell |
| Fgf1 | 0.74 | 0.00 | 0.00 | 0.38 | 0.06 | 17 | Ciliated Cell |
| Cc2d2a | 0.71 | 0.00 | 0.00 | 0.35 | 0.05 | 17 | Ciliated Cell |
| Slc6a14 | 0.61 | 0.00 | 0.00 | 0.28 | 0.03 | 17 | Ciliated Cell |
| Mdm2 | 0.83 | 0.00 | 0.00 | 0.46 | 0.08 | 17 | Ciliated Cell |
| Kifap3 | 0.92 | 0.00 | 0.00 | 0.55 | 0.12 | 17 | Ciliated Cell |
| Ift46 | 0.57 | 0.00 | 0.00 | 0.33 | 0.04 | 17 | Ciliated Cell |
| Hes6 | 0.73 | 0.00 | 0.00 | 0.36 | 0.05 | 17 | Ciliated Cell |
| Lztfl1 | 0.80 | 0.00 | 0.00 | 0.47 | 0.09 | 17 | Ciliated Cell |
| Gdpd1 | 0.62 | 0.00 | 0.00 | 0.38 | 0.06 | 17 | Ciliated Cell |
| Ccp110 | 0.68 | 0.00 | 0.00 | 0.34 | 0.05 | 17 | Ciliated Cell |
| Cxadr | 0.68 | 0.00 | 0.00 | 0.37 | 0.06 | 17 | Ciliated Cell |
| Apoa1bp | 0.95 | 0.00 | 0.00 | 0.59 | 0.14 | 17 | Ciliated Cell |
| Cmtm8 | 0.83 | 0.00 | 0.00 | 0.55 | 0.12 | 17 | Ciliated Cell |
| Akip1 | 0.73 | 0.00 | 0.00 | 0.37 | 0.06 | 17 | Ciliated Cell |
| Alas1 | 0.97 | 0.00 | 0.00 | 0.53 | 0.12 | 17 | Ciliated Cell |
| Ppp1r14c | 0.69 | 0.00 | 0.00 | 0.53 | 0.11 | 17 | Ciliated Cell |
| Dynlt1a | 1.11 | 0.00 | 0.00 | 0.71 | 0.22 | 17 | Ciliated Cell |
| Akap9 | 1.18 | 0.00 | 0.00 | 0.87 | 0.35 | 17 | Ciliated Cell |
| Cpm | 0.97 | 0.00 | 0.00 | 0.67 | 0.17 | 17 | Ciliated Cell |
| Arhgap18 | 0.76 | 0.00 | 0.00 | 0.41 | 0.07 | 17 | Ciliated Cell |
| 2610301B20Rik | 0.53 | 0.00 | 0.00 | 0.29 | 0.04 | 17 | Ciliated Cell |
| Mal2 | 0.61 | 0.00 | 0.00 | 0.38 | 0.06 | 17 | Ciliated Cell |
| Rassf9 | 0.53 | 0.00 | 0.00 | 0.29 | 0.04 | 17 | Ciliated Cell |
| Baiap2l1 | 0.62 | 0.00 | 0.00 | 0.29 | 0.04 | 17 | Ciliated Cell |
| Hsph1 | 0.74 | 0.00 | 0.00 | 0.32 | 0.05 | 17 | Ciliated Cell |
| Sod2 | 0.86 | 0.00 | 0.00 | 0.56 | 0.13 | 17 | Ciliated Cell |
| Fbxw9 | 0.50 | 0.00 | 0.00 | 0.27 | 0.03 | 17 | Ciliated Cell |
| S100a11 | 1.16 | 0.00 | 0.00 | 0.91 | 0.48 | 17 | Ciliated Cell |
| Sclt1 | 0.67 | 0.00 | 0.00 | 0.37 | 0.06 | 17 | Ciliated Cell |
| Mgst3 | 0.78 | 0.00 | 0.00 | 0.47 | 0.09 | 17 | Ciliated Cell |
| Msi2 | 0.93 | 0.00 | 0.00 | 0.63 | 0.17 | 17 | Ciliated Cell |
| Grb14 | 0.61 | 0.00 | 0.00 | 0.36 | 0.06 | 17 | Ciliated Cell |
| Gga1 | 0.55 | 0.00 | 0.00 | 0.32 | 0.05 | 17 | Ciliated Cell |
| Ankrd32 | 0.87 | 0.00 | 0.00 | 0.48 | 0.10 | 17 | Ciliated Cell |
| Odf2 | 0.72 | 0.00 | 0.00 | 0.43 | 0.08 | 17 | Ciliated Cell |
| Fam53b | 0.58 | 0.00 | 0.00 | 0.35 | 0.05 | 17 | Ciliated Cell |
| Smco4 | 0.68 | 0.00 | 0.00 | 0.40 | 0.07 | 17 | Ciliated Cell |
| Oxct1 | 0.89 | 0.00 | 0.00 | 0.58 | 0.14 | 17 | Ciliated Cell |
| Cyp4b1 | 0.66 | 0.00 | 0.00 | 0.50 | 0.11 | 17 | Ciliated Cell |
| Rcc2 | 0.61 | 0.00 | 0.00 | 0.34 | 0.05 | 17 | Ciliated Cell |
| Odc1 | 0.63 | 0.00 | 0.00 | 0.35 | 0.06 | 17 | Ciliated Cell |
| Nudt14 | 0.61 | 0.00 | 0.00 | 0.37 | 0.06 | 17 | Ciliated Cell |
| Cdkn1a | 0.84 | 0.00 | 0.00 | 0.61 | 0.16 | 17 | Ciliated Cell |
| Tbcel | 0.69 | 0.00 | 0.00 | 0.36 | 0.06 | 17 | Ciliated Cell |
| Ssx2ip | 0.54 | 0.00 | 0.00 | 0.28 | 0.04 | 17 | Ciliated Cell |
| 1700047I17Rik2 | 0.89 | 0.00 | 0.00 | 0.64 | 0.18 | 17 | Ciliated Cell |
| Spop | 1.09 | 0.00 | 0.00 | 0.81 | 0.32 | 17 | Ciliated Cell |
| Ocln | 0.55 | 0.00 | 0.00 | 0.29 | 0.04 | 17 | Ciliated Cell |
| Hipk3 | 0.59 | 0.00 | 0.00 | 0.34 | 0.06 | 17 | Ciliated Cell |
| S100a6 | 1.09 | 0.00 | 0.00 | 0.81 | 0.31 | 17 | Ciliated Cell |
| Sned1 | 0.59 | 0.00 | 0.00 | 0.33 | 0.05 | 17 | Ciliated Cell |
| Acot13 | 0.66 | 0.00 | 0.00 | 0.40 | 0.08 | 17 | Ciliated Cell |
| Ndufaf3 | 0.59 | 0.00 | 0.00 | 0.31 | 0.05 | 17 | Ciliated Cell |
| Trim37 | 0.70 | 0.00 | 0.00 | 0.40 | 0.08 | 17 | Ciliated Cell |
| Fam177a | 0.93 | 0.00 | 0.00 | 0.70 | 0.23 | 17 | Ciliated Cell |
| Pcbd2 | 0.63 | 0.00 | 0.00 | 0.35 | 0.06 | 17 | Ciliated Cell |
| Ptprf | 0.68 | 0.00 | 0.00 | 0.60 | 0.15 | 17 | Ciliated Cell |
| Tagap1 | 0.55 | 0.00 | 0.00 | 0.30 | 0.05 | 17 | Ciliated Cell |
| Scgb3a2 | 2.17 | 0.00 | 0.00 | 0.37 | 0.07 | 17 | Ciliated Cell |
| Cluap1 | 0.47 | 0.00 | 0.00 | 0.27 | 0.04 | 17 | Ciliated Cell |
| Pick1 | 0.50 | 0.00 | 0.00 | 0.32 | 0.05 | 17 | Ciliated Cell |
| Cry2 | 0.47 | 0.00 | 0.00 | 0.26 | 0.04 | 17 | Ciliated Cell |
| Mrps17 | 0.79 | 0.00 | 0.00 | 0.53 | 0.13 | 17 | Ciliated Cell |
| Sfta2 | 0.92 | 0.00 | 0.00 | 0.58 | 0.15 | 17 | Ciliated Cell |
| Epb4.1l5 | 0.50 | 0.00 | 0.00 | 0.26 | 0.04 | 17 | Ciliated Cell |
| Avpi1 | 0.66 | 0.00 | 0.00 | 0.41 | 0.08 | 17 | Ciliated Cell |
| Hipk1 | 0.68 | 0.00 | 0.00 | 0.40 | 0.08 | 17 | Ciliated Cell |
| Plxnb2 | 0.56 | 0.00 | 0.00 | 0.32 | 0.05 | 17 | Ciliated Cell |
| Wdr6 | 0.47 | 0.00 | 0.00 | 0.29 | 0.04 | 17 | Ciliated Cell |
| Wif1 | 0.69 | 0.00 | 0.00 | 0.29 | 0.04 | 17 | Ciliated Cell |
| Cep135 | 0.63 | 0.00 | 0.00 | 0.33 | 0.06 | 17 | Ciliated Cell |
| BC005624 | 0.71 | 0.00 | 0.00 | 0.53 | 0.13 | 17 | Ciliated Cell |
| Tchp | 0.48 | 0.00 | 0.00 | 0.26 | 0.04 | 17 | Ciliated Cell |
| Anxa2 | 0.96 | 0.00 | 0.00 | 0.89 | 0.47 | 17 | Ciliated Cell |
| Lasp1 | 0.86 | 0.00 | 0.00 | 0.55 | 0.15 | 17 | Ciliated Cell |
| Tuba1a | 1.05 | 0.00 | 0.00 | 0.92 | 0.50 | 17 | Ciliated Cell |
| Vimp | 0.96 | 0.00 | 0.00 | 0.76 | 0.29 | 17 | Ciliated Cell |
| Megf6 | 0.51 | 0.00 | 0.00 | 0.26 | 0.04 | 17 | Ciliated Cell |
| 1810037I17Rik | 0.97 | 0.00 | 0.00 | 0.78 | 0.31 | 17 | Ciliated Cell |
| Nr2c2ap | 0.45 | 0.00 | 0.00 | 0.26 | 0.04 | 17 | Ciliated Cell |
| Map7 | 0.56 | 0.00 | 0.00 | 0.29 | 0.05 | 17 | Ciliated Cell |
| Cep89 | 0.52 | 0.00 | 0.00 | 0.29 | 0.05 | 17 | Ciliated Cell |
| Sccpdh | 0.57 | 0.00 | 0.00 | 0.31 | 0.05 | 17 | Ciliated Cell |
| Tmem254b | 0.82 | 0.00 | 0.00 | 0.62 | 0.19 | 17 | Ciliated Cell |
| Sdc1 | 0.74 | 0.00 | 0.00 | 0.48 | 0.11 | 17 | Ciliated Cell |
| Tmem254c | 0.80 | 0.00 | 0.00 | 0.61 | 0.18 | 17 | Ciliated Cell |
| Crim1 | 0.64 | 0.00 | 0.00 | 0.40 | 0.08 | 17 | Ciliated Cell |
| Ppl | 0.50 | 0.00 | 0.00 | 0.25 | 0.04 | 17 | Ciliated Cell |
| Hist1h2al | 0.74 | 0.00 | 0.00 | 0.42 | 0.09 | 17 | Ciliated Cell |
| Cbx6 | 0.65 | 0.00 | 0.00 | 0.44 | 0.10 | 17 | Ciliated Cell |
| Tmem254a | 0.80 | 0.00 | 0.00 | 0.62 | 0.19 | 17 | Ciliated Cell |
| Emb | 0.63 | 0.00 | 0.00 | 0.46 | 0.11 | 17 | Ciliated Cell |
| Chuk | 0.62 | 0.00 | 0.00 | 0.39 | 0.08 | 17 | Ciliated Cell |
| 4931406C07Rik | 0.66 | 0.00 | 0.00 | 0.32 | 0.06 | 17 | Ciliated Cell |
| Ruvbl1 | 0.59 | 0.00 | 0.00 | 0.35 | 0.07 | 17 | Ciliated Cell |
| Kif3b | 0.53 | 0.00 | 0.00 | 0.32 | 0.06 | 17 | Ciliated Cell |
| Nkx2-1 | 0.59 | 0.00 | 0.00 | 0.43 | 0.09 | 17 | Ciliated Cell |
| Cnp | 0.55 | 0.00 | 0.00 | 0.29 | 0.05 | 17 | Ciliated Cell |
| Plxnb1 | 0.45 | 0.00 | 0.00 | 0.26 | 0.04 | 17 | Ciliated Cell |
| Emc6 | 0.76 | 0.00 | 0.00 | 0.51 | 0.14 | 17 | Ciliated Cell |
| Ndufc2 | 0.92 | 0.00 | 0.00 | 0.82 | 0.38 | 17 | Ciliated Cell |
| Cit | 0.51 | 0.00 | 0.00 | 0.27 | 0.04 | 17 | Ciliated Cell |
| Atp13a3 | 0.66 | 0.00 | 0.00 | 0.41 | 0.09 | 17 | Ciliated Cell |
| Gramd2 | 0.46 | 0.00 | 0.00 | 0.28 | 0.05 | 17 | Ciliated Cell |
| Slc25a36 | 0.58 | 0.00 | 0.00 | 0.35 | 0.07 | 17 | Ciliated Cell |
| Dynll1 | 0.95 | 0.00 | 0.00 | 0.85 | 0.44 | 17 | Ciliated Cell |
| Cdc42ep3 | 0.67 | 0.00 | 0.00 | 0.53 | 0.14 | 17 | Ciliated Cell |
| Slc26a2 | 0.62 | 0.00 | 0.00 | 0.33 | 0.06 | 17 | Ciliated Cell |
| Fem1b | 0.51 | 0.00 | 0.00 | 0.28 | 0.05 | 17 | Ciliated Cell |
| Vapa | 0.80 | 0.00 | 0.00 | 0.65 | 0.22 | 17 | Ciliated Cell |
| Eif2b2 | 0.57 | 0.00 | 0.00 | 0.34 | 0.07 | 17 | Ciliated Cell |
| Ptgr1 | 0.53 | 0.00 | 0.00 | 0.29 | 0.05 | 17 | Ciliated Cell |
| Baiap2 | 0.56 | 0.00 | 0.00 | 0.36 | 0.08 | 17 | Ciliated Cell |
| Hist1h2bc | 0.75 | 0.00 | 0.00 | 0.49 | 0.13 | 17 | Ciliated Cell |
| Gbas | 0.70 | 0.00 | 0.00 | 0.44 | 0.11 | 17 | Ciliated Cell |
| Gas6 | 0.72 | 0.00 | 0.00 | 0.52 | 0.14 | 17 | Ciliated Cell |
| Por | 0.70 | 0.00 | 0.00 | 0.47 | 0.12 | 17 | Ciliated Cell |
| Dnal4 | 0.42 | 0.00 | 0.00 | 0.27 | 0.04 | 17 | Ciliated Cell |
| Atp1a1 | 0.71 | 0.00 | 0.00 | 0.51 | 0.14 | 17 | Ciliated Cell |
| Hist2h2aa2 | 0.73 | 0.00 | 0.00 | 0.49 | 0.13 | 17 | Ciliated Cell |
| Lgals3bp | 0.48 | 0.00 | 0.00 | 0.26 | 0.04 | 17 | Ciliated Cell |
| Cib1 | 0.58 | 0.00 | 0.00 | 0.46 | 0.12 | 17 | Ciliated Cell |
| Ide | 0.55 | 0.00 | 0.00 | 0.38 | 0.08 | 17 | Ciliated Cell |
| Csrp2 | 0.76 | 0.00 | 0.00 | 0.55 | 0.16 | 17 | Ciliated Cell |
| Tes | 0.60 | 0.00 | 0.00 | 0.40 | 0.09 | 17 | Ciliated Cell |
| Atpif1 | 0.81 | 0.00 | 0.00 | 0.90 | 0.52 | 17 | Ciliated Cell |
| Synj1 | 0.50 | 0.00 | 0.00 | 0.29 | 0.05 | 17 | Ciliated Cell |
| Ahi1 | 0.57 | 0.00 | 0.00 | 0.27 | 0.05 | 17 | Ciliated Cell |
| Chchd4 | 0.48 | 0.00 | 0.00 | 0.27 | 0.05 | 17 | Ciliated Cell |
| BC048507 | 0.61 | 0.00 | 0.00 | 0.46 | 0.12 | 17 | Ciliated Cell |
| Klhdc8b | 0.52 | 0.00 | 0.00 | 0.32 | 0.06 | 17 | Ciliated Cell |
| Pgrmc1 | 0.72 | 0.00 | 0.00 | 0.57 | 0.18 | 17 | Ciliated Cell |
| Tacc2 | 0.56 | 0.00 | 0.00 | 0.34 | 0.07 | 17 | Ciliated Cell |
| Ssbp4 | 0.43 | 0.00 | 0.00 | 0.29 | 0.06 | 17 | Ciliated Cell |
| Nceh1 | 0.55 | 0.00 | 0.00 | 0.31 | 0.06 | 17 | Ciliated Cell |
| Mgst1 | 0.97 | 0.00 | 0.00 | 0.57 | 0.19 | 17 | Ciliated Cell |
| Gde1 | 0.64 | 0.00 | 0.00 | 0.50 | 0.14 | 17 | Ciliated Cell |
| Clu | 0.65 | 0.00 | 0.00 | 0.28 | 0.05 | 17 | Ciliated Cell |
| Pisd | 0.67 | 0.00 | 0.00 | 0.42 | 0.11 | 17 | Ciliated Cell |
| Gnl2 | 0.65 | 0.00 | 0.00 | 0.46 | 0.12 | 17 | Ciliated Cell |
| Crip2 | 0.80 | 0.00 | 0.00 | 0.84 | 0.41 | 17 | Ciliated Cell |
| Chmp2b | 0.59 | 0.00 | 0.00 | 0.53 | 0.16 | 17 | Ciliated Cell |
| Pygl | 0.47 | 0.00 | 0.00 | 0.42 | 0.10 | 17 | Ciliated Cell |
| Cbr1 | 0.56 | 0.00 | 0.00 | 0.41 | 0.10 | 17 | Ciliated Cell |
| Rbm47 | 0.48 | 0.00 | 0.00 | 0.27 | 0.05 | 17 | Ciliated Cell |
| Trak1 | 0.45 | 0.00 | 0.00 | 0.28 | 0.05 | 17 | Ciliated Cell |
| Syne1 | 0.60 | 0.00 | 0.00 | 0.41 | 0.10 | 17 | Ciliated Cell |
| Ccdc112 | 0.41 | 0.00 | 0.00 | 0.25 | 0.04 | 17 | Ciliated Cell |
| Arl13b | 0.60 | 0.00 | 0.00 | 0.35 | 0.08 | 17 | Ciliated Cell |
| Csgalnact1 | 0.51 | 0.00 | 0.00 | 0.26 | 0.05 | 17 | Ciliated Cell |
| Med19 | 0.57 | 0.00 | 0.00 | 0.42 | 0.11 | 17 | Ciliated Cell |
| Mrpl39 | 0.47 | 0.00 | 0.00 | 0.34 | 0.08 | 17 | Ciliated Cell |
| Cadm1 | 0.53 | 0.00 | 0.00 | 0.49 | 0.14 | 17 | Ciliated Cell |
| Pttg1 | 0.63 | 0.00 | 0.00 | 0.48 | 0.14 | 17 | Ciliated Cell |
| Ccdc137 | 0.49 | 0.00 | 0.00 | 0.32 | 0.07 | 17 | Ciliated Cell |
| Casz1 | 0.54 | 0.00 | 0.00 | 0.48 | 0.13 | 17 | Ciliated Cell |
| Mprip | 0.74 | 0.00 | 0.00 | 0.59 | 0.20 | 17 | Ciliated Cell |
| Krt7 | 0.43 | 0.00 | 0.00 | 0.34 | 0.07 | 17 | Ciliated Cell |
| Txnrd1 | 0.67 | 0.00 | 0.00 | 0.41 | 0.11 | 17 | Ciliated Cell |
| Prkci | 0.60 | 0.00 | 0.00 | 0.46 | 0.13 | 17 | Ciliated Cell |
| Aldoa | 0.71 | 0.00 | 0.00 | 0.56 | 0.19 | 17 | Ciliated Cell |
| Ckb | 0.65 | 0.00 | 0.00 | 0.48 | 0.14 | 17 | Ciliated Cell |
| Chka | 0.56 | 0.00 | 0.00 | 0.39 | 0.10 | 17 | Ciliated Cell |
| Pebp1 | 0.76 | 0.00 | 0.00 | 0.64 | 0.25 | 17 | Ciliated Cell |
| Ccnc | 0.43 | 0.00 | 0.00 | 0.27 | 0.05 | 17 | Ciliated Cell |
| Swt1 | 0.62 | 0.00 | 0.00 | 0.50 | 0.15 | 17 | Ciliated Cell |
| Ccnd2 | 0.74 | 0.00 | 0.00 | 0.69 | 0.27 | 17 | Ciliated Cell |
| Cby1 | 0.48 | 0.00 | 0.00 | 0.25 | 0.05 | 17 | Ciliated Cell |
| Znhit1 | 0.56 | 0.00 | 0.00 | 0.48 | 0.14 | 17 | Ciliated Cell |
| Ino80b | 0.43 | 0.00 | 0.00 | 0.29 | 0.06 | 17 | Ciliated Cell |
| Cox14 | 0.65 | 0.00 | 0.00 | 0.60 | 0.21 | 17 | Ciliated Cell |
| Tusc3 | 0.60 | 0.00 | 0.00 | 0.41 | 0.11 | 17 | Ciliated Cell |
| Ahsa1 | 0.61 | 0.00 | 0.00 | 0.44 | 0.12 | 17 | Ciliated Cell |
| Rmdn3 | 0.45 | 0.00 | 0.00 | 0.27 | 0.05 | 17 | Ciliated Cell |
| Unc119b | 0.46 | 0.00 | 0.00 | 0.26 | 0.05 | 17 | Ciliated Cell |
| Ptges3 | 0.78 | 0.00 | 0.00 | 0.62 | 0.24 | 17 | Ciliated Cell |
| Cisd1 | 0.55 | 0.00 | 0.00 | 0.41 | 0.11 | 17 | Ciliated Cell |
| Mnat1 | 0.61 | 0.00 | 0.00 | 0.41 | 0.11 | 17 | Ciliated Cell |
| Sugt1 | 0.70 | 0.00 | 0.00 | 0.56 | 0.20 | 17 | Ciliated Cell |
| 4922501C03Rik | 0.44 | 0.00 | 0.00 | 0.30 | 0.06 | 17 | Ciliated Cell |
| Cldn18 | 0.29 | 0.00 | 0.00 | 0.56 | 0.17 | 17 | Ciliated Cell |
| Neat1 | 0.62 | 0.00 | 0.00 | 0.69 | 0.27 | 17 | Ciliated Cell |
| Fkbp4 | 0.55 | 0.00 | 0.00 | 0.48 | 0.14 | 17 | Ciliated Cell |
| Ift20 | 0.73 | 0.00 | 0.00 | 0.63 | 0.25 | 17 | Ciliated Cell |
| Golgb1 | 0.75 | 0.00 | 0.00 | 0.74 | 0.32 | 17 | Ciliated Cell |
| Rfc2 | 0.54 | 0.00 | 0.00 | 0.33 | 0.08 | 17 | Ciliated Cell |
| Larp4b | 0.59 | 0.00 | 0.00 | 0.49 | 0.15 | 17 | Ciliated Cell |
| Phf20l1 | 0.76 | 0.00 | 0.00 | 0.68 | 0.28 | 17 | Ciliated Cell |
| Prpf40b | 0.40 | 0.00 | 0.00 | 0.27 | 0.05 | 17 | Ciliated Cell |
| Suclg1 | 0.68 | 0.00 | 0.00 | 0.58 | 0.21 | 17 | Ciliated Cell |
| Pcm1 | 0.70 | 0.00 | 0.00 | 0.66 | 0.26 | 17 | Ciliated Cell |
| Scgb1a1 | 2.48 | 0.00 | 0.00 | 0.71 | 0.35 | 17 | Ciliated Cell |
| Uhrf1bp1l | 0.53 | 0.00 | 0.00 | 0.36 | 0.09 | 17 | Ciliated Cell |
| Sdc4 | 0.54 | 0.00 | 0.00 | 0.63 | 0.22 | 17 | Ciliated Cell |
| Chpt1 | 0.56 | 0.00 | 0.00 | 0.41 | 0.12 | 17 | Ciliated Cell |
| Ccdc34 | 0.66 | 0.00 | 0.00 | 0.44 | 0.13 | 17 | Ciliated Cell |
| Stip1 | 0.57 | 0.00 | 0.00 | 0.42 | 0.12 | 17 | Ciliated Cell |
| Ptpla | 0.40 | 0.00 | 0.00 | 0.31 | 0.07 | 17 | Ciliated Cell |
| 0610011F06Rik | 0.37 | 0.00 | 0.00 | 0.28 | 0.06 | 17 | Ciliated Cell |
| Wdr13 | 0.39 | 0.00 | 0.00 | 0.26 | 0.06 | 17 | Ciliated Cell |
| Mpc1 | 0.57 | 0.00 | 0.00 | 0.48 | 0.15 | 17 | Ciliated Cell |
| Krcc1 | 0.59 | 0.00 | 0.00 | 0.58 | 0.21 | 17 | Ciliated Cell |
| Hes1 | 0.70 | 0.00 | 0.00 | 0.67 | 0.29 | 17 | Ciliated Cell |
| Sec23b | 0.44 | 0.00 | 0.00 | 0.30 | 0.07 | 17 | Ciliated Cell |
| Lgals3 | 0.41 | 0.00 | 0.00 | 0.47 | 0.14 | 17 | Ciliated Cell |
| Smpdl3a | 0.50 | 0.00 | 0.00 | 0.38 | 0.10 | 17 | Ciliated Cell |
| Rgs3 | 0.46 | 0.00 | 0.00 | 0.28 | 0.06 | 17 | Ciliated Cell |
| Gclc | 0.54 | 0.00 | 0.00 | 0.33 | 0.08 | 17 | Ciliated Cell |
| 4930402H24Rik | 0.47 | 0.00 | 0.00 | 0.32 | 0.08 | 17 | Ciliated Cell |
| Otud7b | 0.43 | 0.00 | 0.00 | 0.35 | 0.09 | 17 | Ciliated Cell |
| 8430429K09Rik | 0.40 | 0.00 | 0.00 | 0.27 | 0.06 | 17 | Ciliated Cell |
| Tspan2 | 0.52 | 0.00 | 0.00 | 0.40 | 0.12 | 17 | Ciliated Cell |
| Mea1 | 0.42 | 0.00 | 0.00 | 0.32 | 0.08 | 17 | Ciliated Cell |
| Cpne3 | 0.55 | 0.00 | 0.00 | 0.52 | 0.18 | 17 | Ciliated Cell |
| Tuba1c | 0.73 | 0.00 | 0.00 | 0.70 | 0.32 | 17 | Ciliated Cell |
| Fam60a | 0.38 | 0.00 | 0.00 | 0.30 | 0.07 | 17 | Ciliated Cell |
| Mecom | 0.45 | 0.00 | 0.00 | 0.36 | 0.10 | 17 | Ciliated Cell |
| Adss | 0.61 | 0.00 | 0.00 | 0.47 | 0.16 | 17 | Ciliated Cell |
| Psmd9 | 0.44 | 0.00 | 0.00 | 0.31 | 0.08 | 17 | Ciliated Cell |
| Insig2 | 0.49 | 0.00 | 0.00 | 0.36 | 0.10 | 17 | Ciliated Cell |
| Atp8a1 | 0.59 | 0.00 | 0.00 | 0.46 | 0.15 | 17 | Ciliated Cell |
| Rrp1 | 0.65 | 0.00 | 0.00 | 0.76 | 0.36 | 17 | Ciliated Cell |
| Setd8 | 0.48 | 0.00 | 0.00 | 0.38 | 0.11 | 17 | Ciliated Cell |
| Arl4a | 0.48 | 0.00 | 0.00 | 0.38 | 0.11 | 17 | Ciliated Cell |
| Nagk | 0.43 | 0.00 | 0.00 | 0.30 | 0.08 | 17 | Ciliated Cell |
| Ppp5c | 0.42 | 0.00 | 0.00 | 0.29 | 0.07 | 17 | Ciliated Cell |
| Tuba1b | 0.72 | 0.00 | 0.00 | 0.81 | 0.45 | 17 | Ciliated Cell |
| Gsr | 0.30 | 0.00 | 0.00 | 0.33 | 0.09 | 17 | Ciliated Cell |
| Pcnt | 0.52 | 0.00 | 0.00 | 0.43 | 0.14 | 17 | Ciliated Cell |
| Usp46 | 0.42 | 0.00 | 0.00 | 0.30 | 0.08 | 17 | Ciliated Cell |
| Nrbp2 | 0.39 | 0.00 | 0.00 | 0.25 | 0.06 | 17 | Ciliated Cell |
| Asph | 0.36 | 0.00 | 0.00 | 0.26 | 0.06 | 17 | Ciliated Cell |
| Creld2 | 0.40 | 0.00 | 0.00 | 0.33 | 0.09 | 17 | Ciliated Cell |
| Bri3 | 0.59 | 0.00 | 0.00 | 0.54 | 0.20 | 17 | Ciliated Cell |
| Etfa | 0.57 | 0.00 | 0.00 | 0.47 | 0.16 | 17 | Ciliated Cell |
| Anxa4 | 0.43 | 0.00 | 0.00 | 0.34 | 0.09 | 17 | Ciliated Cell |
| Ppfibp2 | 0.46 | 0.00 | 0.00 | 0.25 | 0.06 | 17 | Ciliated Cell |
| Cep63 | 0.37 | 0.00 | 0.00 | 0.27 | 0.06 | 17 | Ciliated Cell |
| Rpap3 | 0.41 | 0.00 | 0.00 | 0.32 | 0.09 | 17 | Ciliated Cell |
| Rexo2 | 0.48 | 0.00 | 0.00 | 0.44 | 0.14 | 17 | Ciliated Cell |
| Hint2 | 0.44 | 0.00 | 0.00 | 0.36 | 0.10 | 17 | Ciliated Cell |
| Ift52 | 0.37 | 0.00 | 0.00 | 0.27 | 0.06 | 17 | Ciliated Cell |
| Gkap1 | 0.46 | 0.00 | 0.00 | 0.39 | 0.12 | 17 | Ciliated Cell |
| Naa35 | 0.43 | 0.00 | 0.00 | 0.35 | 0.10 | 17 | Ciliated Cell |
| Usp6nl | 0.37 | 0.00 | 0.00 | 0.26 | 0.06 | 17 | Ciliated Cell |
| Dynll2 | 0.49 | 0.00 | 0.00 | 0.38 | 0.12 | 17 | Ciliated Cell |
| Paip2 | 0.61 | 0.00 | 0.00 | 0.73 | 0.36 | 17 | Ciliated Cell |
| Cyb5 | 0.63 | 0.00 | 0.00 | 0.60 | 0.26 | 17 | Ciliated Cell |
| Tomm7 | 0.66 | 0.00 | 0.00 | 0.71 | 0.37 | 17 | Ciliated Cell |
| Fth1 | 0.36 | 0.00 | 0.00 | 0.97 | 0.81 | 17 | Ciliated Cell |
| Plp2 | 0.43 | 0.00 | 0.00 | 0.40 | 0.13 | 17 | Ciliated Cell |
| Bbip1 | 0.45 | 0.00 | 0.00 | 0.37 | 0.11 | 17 | Ciliated Cell |
| Bag1 | 0.60 | 0.00 | 0.00 | 0.57 | 0.23 | 17 | Ciliated Cell |
| Cyth2 | 0.41 | 0.00 | 0.00 | 0.33 | 0.09 | 17 | Ciliated Cell |
| Pltp | 0.47 | 0.00 | 0.00 | 0.50 | 0.18 | 17 | Ciliated Cell |
| Tns3 | 0.39 | 0.00 | 0.00 | 0.36 | 0.11 | 17 | Ciliated Cell |
| Gns | 0.45 | 0.00 | 0.00 | 0.39 | 0.12 | 17 | Ciliated Cell |
| Thap3 | 0.43 | 0.00 | 0.00 | 0.31 | 0.08 | 17 | Ciliated Cell |
| Hltf | 0.38 | 0.00 | 0.00 | 0.27 | 0.07 | 17 | Ciliated Cell |
| Tcp11l2 | 0.36 | 0.00 | 0.00 | 0.28 | 0.07 | 17 | Ciliated Cell |
| Lpgat1 | 0.45 | 0.00 | 0.00 | 0.31 | 0.09 | 17 | Ciliated Cell |
| Txnip | 0.56 | 0.00 | 0.00 | 0.60 | 0.25 | 17 | Ciliated Cell |
| Galk1 | 0.27 | 0.00 | 0.00 | 0.25 | 0.06 | 17 | Ciliated Cell |
| Nav2 | 0.43 | 0.00 | 0.00 | 0.30 | 0.08 | 17 | Ciliated Cell |
| Casp6 | 0.41 | 0.00 | 0.00 | 0.28 | 0.07 | 17 | Ciliated Cell |
| Sec14l1 | 0.50 | 0.00 | 0.00 | 0.38 | 0.12 | 17 | Ciliated Cell |
| Carkd | 0.48 | 0.00 | 0.00 | 0.44 | 0.15 | 17 | Ciliated Cell |
| Nmt2 | 0.46 | 0.00 | 0.00 | 0.33 | 0.10 | 17 | Ciliated Cell |
| Pigp | 0.40 | 0.00 | 0.00 | 0.34 | 0.10 | 17 | Ciliated Cell |
| Psmd10 | 0.33 | 0.00 | 0.00 | 0.28 | 0.07 | 17 | Ciliated Cell |
| Gm6169 | 0.39 | 0.00 | 0.00 | 0.38 | 0.12 | 17 | Ciliated Cell |
| Gtf2b | 0.44 | 0.00 | 0.00 | 0.34 | 0.10 | 17 | Ciliated Cell |
| H3f3b | 0.44 | 0.00 | 0.00 | 0.98 | 0.89 | 17 | Ciliated Cell |
| Scand1 | 0.47 | 0.00 | 0.00 | 0.44 | 0.16 | 17 | Ciliated Cell |
| Tubgcp3 | 0.35 | 0.00 | 0.00 | 0.28 | 0.08 | 17 | Ciliated Cell |
| Atp6v1d | 0.42 | 0.00 | 0.00 | 0.46 | 0.16 | 17 | Ciliated Cell |
| Rock1 | 0.58 | 0.00 | 0.00 | 0.70 | 0.34 | 17 | Ciliated Cell |
| Impad1 | 0.49 | 0.00 | 0.00 | 0.42 | 0.15 | 17 | Ciliated Cell |
| Cct7 | 0.50 | 0.00 | 0.00 | 0.51 | 0.20 | 17 | Ciliated Cell |
| Car8 | 0.31 | 0.00 | 0.00 | 0.28 | 0.08 | 17 | Ciliated Cell |
| Pcyt1a | 0.40 | 0.00 | 0.00 | 0.34 | 0.11 | 17 | Ciliated Cell |
| Tmem176a | 0.51 | 0.00 | 0.00 | 0.51 | 0.20 | 17 | Ciliated Cell |
| Arhgap5 | 0.48 | 0.00 | 0.00 | 0.48 | 0.19 | 17 | Ciliated Cell |
| Siva1 | 0.50 | 0.00 | 0.00 | 0.41 | 0.14 | 17 | Ciliated Cell |
| Eif1a | 0.49 | 0.00 | 0.00 | 0.41 | 0.14 | 17 | Ciliated Cell |
| Hopx | 0.25 | 0.00 | 0.00 | 0.33 | 0.10 | 17 | Ciliated Cell |
| Naga | 0.33 | 0.00 | 0.00 | 0.26 | 0.07 | 17 | Ciliated Cell |
| Dhrs3 | 0.34 | 0.00 | 0.00 | 0.25 | 0.07 | 17 | Ciliated Cell |
| Magi3 | 0.37 | 0.00 | 0.00 | 0.35 | 0.11 | 17 | Ciliated Cell |
| 2510002D24Rik | 0.39 | 0.00 | 0.00 | 0.29 | 0.08 | 17 | Ciliated Cell |
| Ercc6l2 | 0.34 | 0.00 | 0.00 | 0.28 | 0.08 | 17 | Ciliated Cell |
| Usp19 | 0.33 | 0.00 | 0.00 | 0.26 | 0.07 | 17 | Ciliated Cell |
| Nrbf2 | 0.34 | 0.00 | 0.00 | 0.29 | 0.08 | 17 | Ciliated Cell |
| Selenbp1 | 0.62 | 0.00 | 0.00 | 0.54 | 0.22 | 17 | Ciliated Cell |
| Ttf1 | 0.37 | 0.00 | 0.00 | 0.32 | 0.10 | 17 | Ciliated Cell |
| Zc3h14 | 0.32 | 0.00 | 0.00 | 0.28 | 0.08 | 17 | Ciliated Cell |
| Abr | 0.28 | 0.00 | 0.00 | 0.27 | 0.07 | 17 | Ciliated Cell |
| Fam98b | 0.36 | 0.00 | 0.00 | 0.32 | 0.10 | 17 | Ciliated Cell |
| Prdx5 | 0.45 | 0.00 | 0.00 | 0.68 | 0.32 | 17 | Ciliated Cell |
| Vamp8 | 0.52 | 0.00 | 0.00 | 0.69 | 0.34 | 17 | Ciliated Cell |
| Churc1 | 0.40 | 0.00 | 0.00 | 0.39 | 0.14 | 17 | Ciliated Cell |
| Ndufab1 | 0.45 | 0.00 | 0.00 | 0.48 | 0.19 | 17 | Ciliated Cell |
| Med1 | 0.36 | 0.00 | 0.00 | 0.30 | 0.09 | 17 | Ciliated Cell |
| Rnase4 | 0.43 | 0.00 | 0.00 | 0.48 | 0.18 | 17 | Ciliated Cell |
| Ube2b | 0.53 | 0.00 | 0.00 | 0.70 | 0.36 | 17 | Ciliated Cell |
| Hirip3 | 0.38 | 0.00 | 0.00 | 0.29 | 0.09 | 17 | Ciliated Cell |
| Glrx5 | 0.42 | 0.00 | 0.00 | 0.39 | 0.14 | 17 | Ciliated Cell |
| Laptm4b | 0.32 | 0.00 | 0.00 | 0.27 | 0.08 | 17 | Ciliated Cell |
| Clip1 | 0.48 | 0.00 | 0.00 | 0.50 | 0.20 | 17 | Ciliated Cell |
| Lrrfip2 | 0.42 | 0.00 | 0.00 | 0.39 | 0.14 | 17 | Ciliated Cell |
| Upf3b | 0.44 | 0.00 | 0.00 | 0.46 | 0.18 | 17 | Ciliated Cell |
| Lpin2 | 0.46 | 0.00 | 0.00 | 0.40 | 0.15 | 17 | Ciliated Cell |
| Arl4c | 0.42 | 0.00 | 0.00 | 0.32 | 0.10 | 17 | Ciliated Cell |
| Commd2 | 0.37 | 0.00 | 0.00 | 0.32 | 0.10 | 17 | Ciliated Cell |
| Nfe2l2 | 0.45 | 0.00 | 0.00 | 0.57 | 0.25 | 17 | Ciliated Cell |
| Flot1 | 0.31 | 0.00 | 0.00 | 0.25 | 0.07 | 17 | Ciliated Cell |
| Coa6 | 0.33 | 0.00 | 0.00 | 0.26 | 0.07 | 17 | Ciliated Cell |
| Dstn | 0.49 | 0.00 | 0.00 | 0.81 | 0.50 | 17 | Ciliated Cell |
| Dapk1 | 0.36 | 0.00 | 0.00 | 0.33 | 0.11 | 17 | Ciliated Cell |
| Gabarapl1 | 0.38 | 0.00 | 0.00 | 0.42 | 0.15 | 17 | Ciliated Cell |
| Actn1 | 0.47 | 0.00 | 0.00 | 0.52 | 0.22 | 17 | Ciliated Cell |
| Rnh1 | 0.40 | 0.00 | 0.00 | 0.41 | 0.15 | 17 | Ciliated Cell |
| Idh3a | 0.33 | 0.00 | 0.00 | 0.25 | 0.07 | 17 | Ciliated Cell |
| Fip1l1 | 0.41 | 0.00 | 0.00 | 0.46 | 0.18 | 17 | Ciliated Cell |
| Slc25a5 | 0.53 | 0.00 | 0.00 | 0.86 | 0.59 | 17 | Ciliated Cell |
| Sdhd | 0.41 | 0.00 | 0.00 | 0.44 | 0.17 | 17 | Ciliated Cell |
| Lin7c | 0.38 | 0.00 | 0.00 | 0.32 | 0.10 | 17 | Ciliated Cell |
| Tmem64 | 0.42 | 0.00 | 0.00 | 0.32 | 0.10 | 17 | Ciliated Cell |
| Stk11 | 0.36 | 0.00 | 0.00 | 0.30 | 0.10 | 17 | Ciliated Cell |
| Acadm | 0.33 | 0.00 | 0.00 | 0.38 | 0.14 | 17 | Ciliated Cell |
| Cryzl1 | 0.33 | 0.00 | 0.00 | 0.26 | 0.08 | 17 | Ciliated Cell |
| Iqgap1 | 0.51 | 0.00 | 0.00 | 0.77 | 0.45 | 17 | Ciliated Cell |
| Psmb6 | 0.49 | 0.00 | 0.00 | 0.65 | 0.33 | 17 | Ciliated Cell |
| Ttc14 | 0.46 | 0.00 | 0.00 | 0.48 | 0.20 | 17 | Ciliated Cell |
| Adipor2 | 0.33 | 0.00 | 0.00 | 0.34 | 0.11 | 17 | Ciliated Cell |
| Mrpl40 | 0.31 | 0.00 | 0.00 | 0.32 | 0.11 | 17 | Ciliated Cell |
| Suds3 | 0.33 | 0.00 | 0.00 | 0.26 | 0.08 | 17 | Ciliated Cell |
| Rnf181 | 0.39 | 0.00 | 0.00 | 0.35 | 0.12 | 17 | Ciliated Cell |
| Trp53inp1 | 0.25 | 0.00 | 0.00 | 0.26 | 0.08 | 17 | Ciliated Cell |
| Dhx40 | 0.32 | 0.00 | 0.00 | 0.34 | 0.11 | 17 | Ciliated Cell |
| Usmg5 | 0.54 | 0.00 | 0.00 | 0.71 | 0.40 | 17 | Ciliated Cell |
| Ccdc66 | 0.33 | 0.00 | 0.00 | 0.29 | 0.09 | 17 | Ciliated Cell |
| Il10rb | 0.31 | 0.00 | 0.00 | 0.28 | 0.09 | 17 | Ciliated Cell |
| Sh3d19 | 0.35 | 0.00 | 0.00 | 0.29 | 0.09 | 17 | Ciliated Cell |
| Pkp4 | 0.31 | 0.00 | 0.00 | 0.27 | 0.08 | 17 | Ciliated Cell |
| Arl6ip1 | 0.43 | 0.00 | 0.00 | 0.82 | 0.52 | 17 | Ciliated Cell |
| Btf3l4 | 0.36 | 0.00 | 0.00 | 0.36 | 0.13 | 17 | Ciliated Cell |
| Atxn10 | 0.44 | 0.00 | 0.00 | 0.46 | 0.19 | 17 | Ciliated Cell |
| Pdlim1 | 0.34 | 0.00 | 0.00 | 0.38 | 0.14 | 17 | Ciliated Cell |
| Iars2 | 0.33 | 0.00 | 0.00 | 0.29 | 0.09 | 17 | Ciliated Cell |
| Atp6v1a | 0.44 | 0.00 | 0.00 | 0.58 | 0.27 | 17 | Ciliated Cell |
| Selenbp2 | 0.53 | 0.00 | 0.00 | 0.42 | 0.16 | 17 | Ciliated Cell |
| Trip4 | 0.25 | 0.00 | 0.00 | 0.25 | 0.07 | 17 | Ciliated Cell |
| Tmem176b | 0.45 | 0.00 | 0.00 | 0.66 | 0.34 | 17 | Ciliated Cell |
| Nin | 0.30 | 0.00 | 0.00 | 0.38 | 0.14 | 17 | Ciliated Cell |
| Rnf5 | 0.32 | 0.00 | 0.00 | 0.30 | 0.10 | 17 | Ciliated Cell |
| Cep110 | 0.36 | 0.00 | 0.00 | 0.45 | 0.18 | 17 | Ciliated Cell |
| Slain2 | 0.38 | 0.00 | 0.00 | 0.29 | 0.09 | 17 | Ciliated Cell |
| Gstp1 | 0.45 | 0.00 | 0.00 | 0.46 | 0.19 | 17 | Ciliated Cell |
| Arid2 | 0.39 | 0.00 | 0.00 | 0.42 | 0.17 | 17 | Ciliated Cell |
| Dnajc1 | 0.40 | 0.00 | 0.00 | 0.42 | 0.17 | 17 | Ciliated Cell |
| Snapin | 0.34 | 0.00 | 0.00 | 0.33 | 0.11 | 17 | Ciliated Cell |
| AU019823 | 0.32 | 0.00 | 0.00 | 0.27 | 0.09 | 17 | Ciliated Cell |
| Tmem50b | 0.31 | 0.00 | 0.00 | 0.26 | 0.08 | 17 | Ciliated Cell |
| Swi5 | 0.52 | 0.00 | 0.00 | 0.55 | 0.26 | 17 | Ciliated Cell |
| Slc31a1 | 0.34 | 0.00 | 0.00 | 0.32 | 0.11 | 17 | Ciliated Cell |
| Prkx | 0.35 | 0.00 | 0.00 | 0.32 | 0.11 | 17 | Ciliated Cell |
| Plekha1 | 0.28 | 0.00 | 0.00 | 0.33 | 0.11 | 17 | Ciliated Cell |
| Nelfb | 0.33 | 0.00 | 0.00 | 0.37 | 0.14 | 17 | Ciliated Cell |
| Zfp608 | 0.30 | 0.00 | 0.00 | 0.38 | 0.14 | 17 | Ciliated Cell |
| Tpd52l2 | 0.40 | 0.00 | 0.00 | 0.43 | 0.18 | 17 | Ciliated Cell |
| Gabarapl2 | 0.38 | 0.00 | 0.00 | 0.50 | 0.22 | 17 | Ciliated Cell |
| Ppa2 | 0.30 | 0.00 | 0.00 | 0.30 | 0.10 | 17 | Ciliated Cell |
| Acaa1a | 0.33 | 0.00 | 0.00 | 0.41 | 0.16 | 17 | Ciliated Cell |
| Hibadh | 0.34 | 0.00 | 0.00 | 0.31 | 0.11 | 17 | Ciliated Cell |
| Sod1 | 0.53 | 0.00 | 0.00 | 0.75 | 0.47 | 17 | Ciliated Cell |
| Nhp2 | 0.36 | 0.00 | 0.00 | 0.32 | 0.11 | 17 | Ciliated Cell |
| Tpi1 | 0.35 | 0.00 | 0.00 | 0.39 | 0.15 | 17 | Ciliated Cell |
| Kdm1a | 0.35 | 0.00 | 0.00 | 0.37 | 0.14 | 17 | Ciliated Cell |
| Fam129b | 0.32 | 0.00 | 0.00 | 0.25 | 0.08 | 17 | Ciliated Cell |
| Chd4 | 0.48 | 0.00 | 0.00 | 0.78 | 0.48 | 17 | Ciliated Cell |
| Rev3l | 0.28 | 0.00 | 0.00 | 0.32 | 0.11 | 17 | Ciliated Cell |
| Ergic3 | 0.32 | 0.00 | 0.00 | 0.39 | 0.15 | 17 | Ciliated Cell |
| Zfp704 | 0.35 | 0.00 | 0.00 | 0.34 | 0.12 | 17 | Ciliated Cell |
| Dock9 | 0.31 | 0.00 | 0.00 | 0.28 | 0.09 | 17 | Ciliated Cell |
| Eci1 | 0.33 | 0.00 | 0.00 | 0.25 | 0.08 | 17 | Ciliated Cell |
| Atp2a2 | 0.39 | 0.00 | 0.00 | 0.35 | 0.13 | 17 | Ciliated Cell |
| Tagln2 | 0.46 | 0.00 | 0.00 | 0.76 | 0.44 | 17 | Ciliated Cell |
| Sf3b2 | 0.47 | 0.00 | 0.00 | 0.70 | 0.39 | 17 | Ciliated Cell |
| Pls3 | 0.49 | 0.00 | 0.00 | 0.45 | 0.20 | 17 | Ciliated Cell |
| Sms | 0.33 | 0.00 | 0.00 | 0.32 | 0.11 | 17 | Ciliated Cell |
| Fdx1 | 0.33 | 0.00 | 0.00 | 0.36 | 0.14 | 17 | Ciliated Cell |
| Tmx1 | 0.29 | 0.00 | 0.00 | 0.29 | 0.10 | 17 | Ciliated Cell |
| Sys1 | 0.31 | 0.00 | 0.00 | 0.37 | 0.14 | 17 | Ciliated Cell |
| Aco2 | 0.32 | 0.00 | 0.00 | 0.39 | 0.15 | 17 | Ciliated Cell |
| Eif4g3 | 0.39 | 0.00 | 0.00 | 0.56 | 0.27 | 17 | Ciliated Cell |
| Syne2 | 0.28 | 0.00 | 0.00 | 0.37 | 0.14 | 17 | Ciliated Cell |
| Prdx6 | 0.48 | 0.00 | 0.00 | 0.55 | 0.27 | 17 | Ciliated Cell |
| Cops5 | 0.28 | 0.00 | 0.00 | 0.36 | 0.14 | 17 | Ciliated Cell |
| Bnip3 | 0.26 | 0.00 | 0.00 | 0.26 | 0.08 | 17 | Ciliated Cell |
| Vamp5 | 0.27 | 0.00 | 0.00 | 0.28 | 0.09 | 17 | Ciliated Cell |
| Ubn2 | 0.35 | 0.00 | 0.00 | 0.39 | 0.16 | 17 | Ciliated Cell |
| Clcn3 | 0.29 | 0.00 | 0.00 | 0.28 | 0.09 | 17 | Ciliated Cell |
| Tspan3 | 0.29 | 0.00 | 0.00 | 0.39 | 0.15 | 17 | Ciliated Cell |
| Cct4 | 0.41 | 0.00 | 0.00 | 0.49 | 0.22 | 17 | Ciliated Cell |
| Ctnnb1 | 0.47 | 0.00 | 0.00 | 0.65 | 0.34 | 17 | Ciliated Cell |
| Aftph | 0.26 | 0.00 | 0.00 | 0.29 | 0.10 | 17 | Ciliated Cell |
| Nupr1 | 0.62 | 0.00 | 0.00 | 0.41 | 0.18 | 17 | Ciliated Cell |
| 1810009A15Rik | 0.28 | 0.00 | 0.00 | 0.35 | 0.13 | 17 | Ciliated Cell |
| 1110058L19Rik | 0.25 | 0.00 | 0.00 | 0.25 | 0.08 | 17 | Ciliated Cell |
| Gstp2 | 0.37 | 0.00 | 0.00 | 0.41 | 0.17 | 17 | Ciliated Cell |
| Prdx1 | 0.44 | 0.00 | 0.00 | 0.77 | 0.50 | 17 | Ciliated Cell |
| Cep57 | 0.35 | 0.00 | 0.00 | 0.30 | 0.11 | 17 | Ciliated Cell |
| Gclm | 0.41 | 0.00 | 0.00 | 0.72 | 0.41 | 17 | Ciliated Cell |
| Calm3 | 0.33 | 0.00 | 0.00 | 0.45 | 0.19 | 17 | Ciliated Cell |
| Tsc22d1 | 0.35 | 0.00 | 0.00 | 0.64 | 0.33 | 17 | Ciliated Cell |
| Pi4k2b | 0.26 | 0.00 | 0.00 | 0.28 | 0.10 | 17 | Ciliated Cell |
| Pole3 | 0.32 | 0.00 | 0.00 | 0.29 | 0.10 | 17 | Ciliated Cell |
| Crip1 | 0.31 | 0.00 | 0.00 | 0.80 | 0.47 | 17 | Ciliated Cell |
| Gm9803 | 0.29 | 0.00 | 0.00 | 0.27 | 0.09 | 17 | Ciliated Cell |
| Ak3 | 0.28 | 0.00 | 0.00 | 0.28 | 0.10 | 17 | Ciliated Cell |
| Qsox1 | 0.26 | 0.00 | 0.00 | 0.28 | 0.10 | 17 | Ciliated Cell |
| Bcap31 | 0.32 | 0.00 | 0.00 | 0.47 | 0.21 | 17 | Ciliated Cell |
| Phf3 | 0.36 | 0.00 | 0.00 | 0.52 | 0.24 | 17 | Ciliated Cell |
| Tspo | 0.34 | 0.00 | 0.00 | 0.53 | 0.25 | 17 | Ciliated Cell |
| Shfm1 | 0.41 | 0.00 | 0.00 | 0.86 | 0.64 | 17 | Ciliated Cell |
| Mtus1 | 0.29 | 0.00 | 0.00 | 0.41 | 0.17 | 17 | Ciliated Cell |
| Rpn2 | 0.34 | 0.00 | 0.00 | 0.46 | 0.20 | 17 | Ciliated Cell |
| Sri | 0.37 | 0.00 | 0.00 | 0.56 | 0.27 | 17 | Ciliated Cell |
| Emc4 | 0.28 | 0.00 | 0.00 | 0.35 | 0.14 | 17 | Ciliated Cell |
| Txn1 | 0.45 | 0.00 | 0.00 | 0.77 | 0.49 | 17 | Ciliated Cell |
| Mrps6 | 0.30 | 0.00 | 0.00 | 0.29 | 0.11 | 17 | Ciliated Cell |
| Gng5 | 0.42 | 0.00 | 0.00 | 0.73 | 0.45 | 17 | Ciliated Cell |
| BC031181 | 0.25 | 0.00 | 0.00 | 0.35 | 0.14 | 17 | Ciliated Cell |
| Ier5 | 0.26 | 0.00 | 0.00 | 0.25 | 0.08 | 17 | Ciliated Cell |
| Golga4 | 0.33 | 0.00 | 0.00 | 0.59 | 0.29 | 17 | Ciliated Cell |
| Acat1 | 0.35 | 0.00 | 0.00 | 0.41 | 0.18 | 17 | Ciliated Cell |
| Cntln | 0.27 | 0.00 | 0.00 | 0.31 | 0.12 | 17 | Ciliated Cell |
| Cstb | 0.25 | 0.00 | 0.00 | 0.55 | 0.27 | 17 | Ciliated Cell |
| Mdh1 | 0.28 | 0.00 | 0.00 | 0.43 | 0.19 | 17 | Ciliated Cell |
| Vps28 | 0.34 | 0.00 | 0.00 | 0.41 | 0.18 | 17 | Ciliated Cell |
| Hsbp1 | 0.36 | 0.00 | 0.00 | 0.59 | 0.31 | 17 | Ciliated Cell |
| Fnta | 0.27 | 0.00 | 0.00 | 0.31 | 0.12 | 17 | Ciliated Cell |
| Diablo | 0.26 | 0.00 | 0.00 | 0.27 | 0.10 | 17 | Ciliated Cell |
| Srpk2 | 0.27 | 0.00 | 0.00 | 0.38 | 0.16 | 17 | Ciliated Cell |
| Tmem179b | 0.25 | 0.00 | 0.00 | 0.30 | 0.11 | 17 | Ciliated Cell |
| Anapc13 | 0.31 | 0.00 | 0.00 | 0.46 | 0.21 | 17 | Ciliated Cell |
| Efr3a | 0.29 | 0.00 | 0.00 | 0.32 | 0.12 | 17 | Ciliated Cell |
| Dnaja2 | 0.32 | 0.00 | 0.00 | 0.52 | 0.25 | 17 | Ciliated Cell |
| Gm3150 | 0.33 | 0.00 | 0.00 | 0.38 | 0.16 | 17 | Ciliated Cell |
| Ddx26b | 0.29 | 0.00 | 0.00 | 0.34 | 0.14 | 17 | Ciliated Cell |
| Serf1 | 0.28 | 0.00 | 0.00 | 0.28 | 0.11 | 17 | Ciliated Cell |
| Ppp1cb | 0.30 | 0.00 | 0.00 | 0.43 | 0.19 | 17 | Ciliated Cell |
| Polr2i | 0.27 | 0.00 | 0.00 | 0.36 | 0.15 | 17 | Ciliated Cell |
| Mbip | 0.26 | 0.00 | 0.00 | 0.30 | 0.11 | 17 | Ciliated Cell |
| Man1a2 | 0.30 | 0.00 | 0.00 | 0.36 | 0.15 | 17 | Ciliated Cell |
| Rin2 | 0.27 | 0.00 | 0.00 | 0.36 | 0.15 | 17 | Ciliated Cell |
| Arid4b | 0.34 | 0.00 | 0.00 | 0.61 | 0.33 | 17 | Ciliated Cell |
| Upf2 | 0.27 | 0.00 | 0.00 | 0.35 | 0.15 | 17 | Ciliated Cell |
| Cab39 | 0.27 | 0.00 | 0.00 | 0.38 | 0.16 | 17 | Ciliated Cell |
| Ngfrap1 | 0.31 | 0.00 | 0.00 | 0.48 | 0.23 | 17 | Ciliated Cell |
| 9530068E07Rik | 0.28 | 0.00 | 0.00 | 0.39 | 0.17 | 17 | Ciliated Cell |
| Bsg | 0.38 | 0.00 | 0.00 | 0.62 | 0.36 | 17 | Ciliated Cell |
| Usp47 | 0.27 | 0.00 | 0.00 | 0.31 | 0.12 | 17 | Ciliated Cell |
| F11r | 0.25 | 0.00 | 0.00 | 0.38 | 0.16 | 17 | Ciliated Cell |
| Dctn3 | 0.25 | 0.00 | 0.00 | 0.39 | 0.17 | 17 | Ciliated Cell |
| Knop1 | 0.27 | 0.00 | 0.00 | 0.38 | 0.17 | 17 | Ciliated Cell |
| Pcmt1 | 0.28 | 0.00 | 0.00 | 0.42 | 0.19 | 17 | Ciliated Cell |
| Atp11a | 0.29 | 0.00 | 0.00 | 0.29 | 0.11 | 17 | Ciliated Cell |
| Rif1 | 0.26 | 0.00 | 0.00 | 0.38 | 0.16 | 17 | Ciliated Cell |
| Ndufb5 | 0.33 | 0.00 | 0.00 | 0.47 | 0.23 | 17 | Ciliated Cell |
| Hcfc1r1 | 0.28 | 0.00 | 0.00 | 0.34 | 0.14 | 17 | Ciliated Cell |
| Papola | 0.27 | 0.00 | 0.00 | 0.48 | 0.23 | 17 | Ciliated Cell |
| Tmbim6 | 0.33 | 0.00 | 0.00 | 0.66 | 0.37 | 17 | Ciliated Cell |
| Erdr1 | 0.25 | 0.00 | 0.00 | 0.25 | 0.09 | 17 | Ciliated Cell |
| Ptov1 | 0.27 | 0.00 | 0.00 | 0.34 | 0.14 | 17 | Ciliated Cell |
| Gorasp2 | 0.26 | 0.00 | 0.00 | 0.28 | 0.11 | 17 | Ciliated Cell |
| Skp1a | 0.37 | 0.00 | 0.00 | 0.67 | 0.40 | 17 | Ciliated Cell |
| 2-Mar | 0.27 | 0.00 | 0.00 | 0.32 | 0.13 | 17 | Ciliated Cell |
| Dnajb6 | 0.32 | 0.00 | 0.00 | 0.52 | 0.26 | 17 | Ciliated Cell |
| Eif3c | 0.37 | 0.00 | 0.00 | 0.74 | 0.47 | 17 | Ciliated Cell |
| Arhgef2 | 0.29 | 0.00 | 0.00 | 0.43 | 0.20 | 17 | Ciliated Cell |
| 4930523C07Rik | 0.25 | 0.00 | 0.00 | 0.40 | 0.18 | 17 | Ciliated Cell |
| Wbp11 | 0.27 | 0.00 | 0.00 | 0.48 | 0.24 | 17 | Ciliated Cell |
| Morf4l2 | 0.26 | 0.00 | 0.00 | 0.46 | 0.22 | 17 | Ciliated Cell |
| Gtf2i | 0.27 | 0.00 | 0.00 | 0.37 | 0.17 | 17 | Ciliated Cell |
| Hmgb3 | 0.26 | 0.00 | 0.00 | 0.33 | 0.14 | 17 | Ciliated Cell |
| Atg12 | 0.26 | 0.00 | 0.00 | 0.26 | 0.10 | 17 | Ciliated Cell |
| Dad1 | 0.35 | 0.00 | 0.00 | 0.62 | 0.36 | 17 | Ciliated Cell |
| Dhrs7 | 0.27 | 0.00 | 0.00 | 0.35 | 0.16 | 17 | Ciliated Cell |
| Spcs1 | 0.30 | 0.00 | 0.00 | 0.55 | 0.29 | 17 | Ciliated Cell |
| Hint1 | 0.33 | 0.00 | 0.00 | 0.73 | 0.46 | 17 | Ciliated Cell |
| Hprt | 0.25 | 0.00 | 0.00 | 0.31 | 0.13 | 17 | Ciliated Cell |
| Srp14 | 0.27 | 0.00 | 0.00 | 0.55 | 0.29 | 17 | Ciliated Cell |
| Cox6c | 0.34 | 0.00 | 0.00 | 0.85 | 0.65 | 17 | Ciliated Cell |
| Cetn3 | 0.29 | 0.00 | 0.00 | 0.57 | 0.31 | 17 | Ciliated Cell |
| Ywhae | 0.34 | 0.00 | 0.00 | 0.79 | 0.54 | 17 | Ciliated Cell |
| Wbp5 | 0.35 | 0.00 | 0.00 | 0.73 | 0.48 | 17 | Ciliated Cell |
| Mtpn | 0.26 | 0.00 | 0.00 | 0.51 | 0.27 | 17 | Ciliated Cell |
| Dync1i2 | 0.32 | 0.00 | 0.00 | 0.75 | 0.48 | 17 | Ciliated Cell |
| Acaa2 | 0.27 | 0.00 | 0.00 | 0.35 | 0.16 | 17 | Ciliated Cell |
| Azin1 | 0.26 | 0.00 | 0.00 | 0.30 | 0.13 | 17 | Ciliated Cell |
| Ldha | 0.25 | 0.00 | 0.00 | 0.54 | 0.29 | 17 | Ciliated Cell |
| Calm2 | 0.29 | 0.00 | 0.00 | 0.88 | 0.64 | 17 | Ciliated Cell |
| Ubb | 0.29 | 0.00 | 0.00 | 0.92 | 0.81 | 17 | Ciliated Cell |
| Usp34 | 0.27 | 0.00 | 0.00 | 0.47 | 0.25 | 17 | Ciliated Cell |
| Gpx4 | 0.31 | 0.00 | 0.00 | 0.61 | 0.37 | 17 | Ciliated Cell |
| mt-Co1 | 0.27 | 0.00 | 0.00 | 0.98 | 0.95 | 17 | Ciliated Cell |
| mt-Cytb | 0.29 | 0.00 | 0.00 | 0.93 | 0.82 | 17 | Ciliated Cell |
| Senp6 | 0.26 | 0.00 | 0.00 | 0.57 | 0.33 | 17 | Ciliated Cell |
| Ktn1 | 0.26 | 0.00 | 0.00 | 0.63 | 0.38 | 17 | Ciliated Cell |
| Gm10222 | 0.28 | 0.00 | 0.00 | 0.67 | 0.43 | 17 | Ciliated Cell |
| Atp5g3 | 0.26 | 0.00 | 0.00 | 0.60 | 0.36 | 17 | Ciliated Cell |
| mt-Nd4l | 0.25 | 0.00 | 0.00 | 0.61 | 0.38 | 17 | Ciliated Cell |
| mt-Nd4 | 0.28 | 0.00 | 0.00 | 0.89 | 0.75 | 17 | Ciliated Cell |
| Dbi | 0.26 | 0.00 | 0.00 | 0.56 | 0.37 | 17 | Ciliated Cell |
| mt-Nd5 | 0.28 | 0.00 | 0.00 | 0.88 | 0.75 | 17 | Ciliated Cell |
| Gucy1a3 | 3.27 | 0.00 | 0.00 | 0.94 | 0.08 | 18 | Matrix Fibroblast |
| Postn | 2.90 | 0.00 | 0.00 | 0.90 | 0.07 | 18 | Matrix Fibroblast |
| Gucy1b3 | 2.72 | 0.00 | 0.00 | 0.81 | 0.04 | 18 | Matrix Fibroblast |
| Higd1b | 2.59 | 0.00 | 0.00 | 0.70 | 0.01 | 18 | Matrix Fibroblast |
| Cox4i2 | 2.34 | 0.00 | 0.00 | 0.71 | 0.01 | 18 | Matrix Fibroblast |
| Pdgfrb | 2.19 | 0.00 | 0.00 | 0.71 | 0.03 | 18 | Matrix Fibroblast |
| Itm2a | 2.18 | 0.00 | 0.00 | 0.60 | 0.06 | 18 | Matrix Fibroblast |
| Pcdh18 | 2.04 | 0.00 | 0.00 | 0.56 | 0.03 | 18 | Matrix Fibroblast |
| Pde5a | 2.01 | 0.00 | 0.00 | 0.68 | 0.11 | 18 | Matrix Fibroblast |
| Rgs5 | 1.86 | 0.00 | 0.00 | 0.45 | 0.03 | 18 | Matrix Fibroblast |
| Itga1 | 1.86 | 0.00 | 0.00 | 0.60 | 0.09 | 18 | Matrix Fibroblast |
| Pdzd2 | 1.83 | 0.00 | 0.00 | 0.50 | 0.02 | 18 | Matrix Fibroblast |
| Cdh2 | 1.82 | 0.00 | 0.00 | 0.51 | 0.01 | 18 | Matrix Fibroblast |
| Sparcl1 | 1.76 | 0.00 | 0.00 | 0.94 | 0.38 | 18 | Matrix Fibroblast |
| Lipg | 1.70 | 0.00 | 0.00 | 0.40 | 0.01 | 18 | Matrix Fibroblast |
| Mfge8 | 1.70 | 0.00 | 0.00 | 0.69 | 0.15 | 18 | Matrix Fibroblast |
| Hbegf | 1.69 | 0.00 | 0.00 | 0.52 | 0.05 | 18 | Matrix Fibroblast |
| Ndufa4l2 | 1.69 | 0.00 | 0.00 | 0.47 | 0.01 | 18 | Matrix Fibroblast |
| Lhfp | 1.67 | 0.00 | 0.00 | 0.69 | 0.15 | 18 | Matrix Fibroblast |
| Mcam | 1.66 | 0.00 | 0.00 | 0.65 | 0.12 | 18 | Matrix Fibroblast |
| Maged2 | 1.66 | 0.00 | 0.00 | 0.66 | 0.16 | 18 | Matrix Fibroblast |
| Fam162b | 1.63 | 0.00 | 0.00 | 0.40 | 0.01 | 18 | Matrix Fibroblast |
| Ebf1 | 1.62 | 0.00 | 0.00 | 0.52 | 0.04 | 18 | Matrix Fibroblast |
| Trpc6 | 1.56 | 0.00 | 0.00 | 0.40 | 0.01 | 18 | Matrix Fibroblast |
| Gpr126 | 1.56 | 0.00 | 0.00 | 0.56 | 0.10 | 18 | Matrix Fibroblast |
| Kcnk3 | 1.52 | 0.00 | 0.00 | 0.49 | 0.04 | 18 | Matrix Fibroblast |
| Lmcd1 | 1.52 | 0.00 | 0.00 | 0.45 | 0.04 | 18 | Matrix Fibroblast |
| Notch3 | 1.51 | 0.00 | 0.00 | 0.44 | 0.01 | 18 | Matrix Fibroblast |
| Nkain4 | 1.46 | 0.00 | 0.00 | 0.44 | 0.04 | 18 | Matrix Fibroblast |
| Sdc2 | 1.45 | 0.00 | 0.00 | 0.59 | 0.11 | 18 | Matrix Fibroblast |
| Fam101a | 1.44 | 0.00 | 0.00 | 0.35 | 0.01 | 18 | Matrix Fibroblast |
| Heyl | 1.44 | 0.00 | 0.00 | 0.39 | 0.02 | 18 | Matrix Fibroblast |
| Klhl23 | 1.42 | 0.00 | 0.00 | 0.44 | 0.05 | 18 | Matrix Fibroblast |
| Emid1 | 1.41 | 0.00 | 0.00 | 0.40 | 0.04 | 18 | Matrix Fibroblast |
| Tbx5 | 1.40 | 0.00 | 0.00 | 0.46 | 0.07 | 18 | Matrix Fibroblast |
| Gap43 | 1.35 | 0.00 | 0.00 | 0.26 | 0.02 | 18 | Matrix Fibroblast |
| Parm1 | 1.34 | 0.00 | 0.00 | 0.31 | 0.01 | 18 | Matrix Fibroblast |
| S1pr3 | 1.27 | 0.00 | 0.00 | 0.37 | 0.04 | 18 | Matrix Fibroblast |
| Naalad2 | 1.26 | 0.00 | 0.00 | 0.30 | 0.02 | 18 | Matrix Fibroblast |
| Wisp1 | 1.24 | 0.00 | 0.00 | 0.31 | 0.03 | 18 | Matrix Fibroblast |
| Cspg4 | 1.10 | 0.00 | 0.00 | 0.27 | 0.01 | 18 | Matrix Fibroblast |
| Vsnl1 | 1.09 | 0.00 | 0.00 | 0.27 | 0.02 | 18 | Matrix Fibroblast |
| Angpt1 | 1.40 | 0.00 | 0.00 | 0.44 | 0.07 | 18 | Matrix Fibroblast |
| Crim1 | 1.38 | 0.00 | 0.00 | 0.47 | 0.08 | 18 | Matrix Fibroblast |
| Ltbp2 | 1.21 | 0.00 | 0.00 | 0.49 | 0.09 | 18 | Matrix Fibroblast |
| Rarres2 | 1.30 | 0.00 | 0.00 | 0.57 | 0.13 | 18 | Matrix Fibroblast |
| Col4a1 | 1.41 | 0.00 | 0.00 | 0.92 | 0.49 | 18 | Matrix Fibroblast |
| F2r | 1.50 | 0.00 | 0.00 | 0.55 | 0.14 | 18 | Matrix Fibroblast |
| Fermt2 | 1.56 | 0.00 | 0.00 | 0.83 | 0.39 | 18 | Matrix Fibroblast |
| Gjc1 | 1.17 | 0.00 | 0.00 | 0.39 | 0.07 | 18 | Matrix Fibroblast |
| Col4a2 | 1.28 | 0.00 | 0.00 | 0.87 | 0.42 | 18 | Matrix Fibroblast |
| 4-Sep | 1.30 | 0.00 | 0.00 | 0.70 | 0.23 | 18 | Matrix Fibroblast |
| 1190002N15Rik | 0.96 | 0.00 | 0.00 | 0.26 | 0.03 | 18 | Matrix Fibroblast |
| Ppp1r14a | 1.05 | 0.00 | 0.00 | 0.40 | 0.08 | 18 | Matrix Fibroblast |
| Ednra | 1.10 | 0.00 | 0.00 | 0.41 | 0.08 | 18 | Matrix Fibroblast |
| Plac9b | 0.94 | 0.00 | 0.00 | 0.82 | 0.29 | 18 | Matrix Fibroblast |
| P2ry14 | 0.93 | 0.00 | 0.00 | 0.40 | 0.08 | 18 | Matrix Fibroblast |
| Ehd3 | 0.96 | 0.00 | 0.00 | 0.26 | 0.04 | 18 | Matrix Fibroblast |
| Rasgrp2 | 1.06 | 0.00 | 0.00 | 0.42 | 0.09 | 18 | Matrix Fibroblast |
| Plac9a | 0.93 | 0.00 | 0.00 | 0.81 | 0.30 | 18 | Matrix Fibroblast |
| Cadm1 | 1.31 | 0.00 | 0.00 | 0.50 | 0.14 | 18 | Matrix Fibroblast |
| Ccrl2 | 0.91 | 0.00 | 0.00 | 0.29 | 0.05 | 18 | Matrix Fibroblast |
| Mir143hg | 0.93 | 0.00 | 0.00 | 0.27 | 0.04 | 18 | Matrix Fibroblast |
| Nid1 | 1.21 | 0.00 | 0.00 | 0.64 | 0.23 | 18 | Matrix Fibroblast |
| Mapt | 0.92 | 0.00 | 0.00 | 0.29 | 0.05 | 18 | Matrix Fibroblast |
| Sparc | 0.78 | 0.00 | 0.00 | 0.99 | 0.73 | 18 | Matrix Fibroblast |
| Meg3 | 1.06 | 0.00 | 0.00 | 0.64 | 0.21 | 18 | Matrix Fibroblast |
| Lamb1 | 1.16 | 0.00 | 0.00 | 0.51 | 0.15 | 18 | Matrix Fibroblast |
| Cald1 | 0.98 | 0.00 | 0.00 | 0.82 | 0.41 | 18 | Matrix Fibroblast |
| Btbd3 | 1.05 | 0.00 | 0.00 | 0.48 | 0.14 | 18 | Matrix Fibroblast |
| Myh11 | 0.81 | 0.00 | 0.00 | 0.42 | 0.10 | 18 | Matrix Fibroblast |
| 7-Sep | 1.14 | 0.00 | 0.00 | 0.86 | 0.58 | 18 | Matrix Fibroblast |
| Cd81 | 1.04 | 0.00 | 0.00 | 0.77 | 0.40 | 18 | Matrix Fibroblast |
| H19 | 1.15 | 0.00 | 0.00 | 0.32 | 0.07 | 18 | Matrix Fibroblast |
| Aspn | 0.52 | 0.00 | 0.00 | 0.39 | 0.10 | 18 | Matrix Fibroblast |
| Myl9 | 1.05 | 0.00 | 0.00 | 0.42 | 0.12 | 18 | Matrix Fibroblast |
| Nostrin | 1.01 | 0.00 | 0.00 | 0.48 | 0.14 | 18 | Matrix Fibroblast |
| Ism1 | 1.02 | 0.00 | 0.00 | 0.35 | 0.08 | 18 | Matrix Fibroblast |
| Nr2f2 | 1.05 | 0.00 | 0.00 | 0.56 | 0.20 | 18 | Matrix Fibroblast |
| Cst3 | 0.96 | 0.00 | 0.00 | 0.87 | 0.59 | 18 | Matrix Fibroblast |
| Tiparp | 1.07 | 0.00 | 0.00 | 0.35 | 0.09 | 18 | Matrix Fibroblast |
| Prr5l | 0.98 | 0.00 | 0.00 | 0.29 | 0.06 | 18 | Matrix Fibroblast |
| Marcks | 0.84 | 0.00 | 0.00 | 0.84 | 0.50 | 18 | Matrix Fibroblast |
| Gnas | 1.00 | 0.00 | 0.00 | 0.89 | 0.67 | 18 | Matrix Fibroblast |
| Zeb2 | 0.99 | 0.00 | 0.00 | 0.68 | 0.31 | 18 | Matrix Fibroblast |
| Lama4 | 0.97 | 0.00 | 0.00 | 0.43 | 0.13 | 18 | Matrix Fibroblast |
| Ndn | 0.89 | 0.00 | 0.00 | 0.26 | 0.05 | 18 | Matrix Fibroblast |
| Myo1b | 1.02 | 0.00 | 0.00 | 0.48 | 0.16 | 18 | Matrix Fibroblast |
| Igf2r | 0.98 | 0.00 | 0.00 | 0.37 | 0.10 | 18 | Matrix Fibroblast |
| Gng11 | 0.97 | 0.00 | 0.00 | 0.64 | 0.28 | 18 | Matrix Fibroblast |
| Col1a2 | 0.26 | 0.00 | 0.00 | 0.91 | 0.39 | 18 | Matrix Fibroblast |
| Car2 | 1.12 | 0.00 | 0.00 | 0.63 | 0.29 | 18 | Matrix Fibroblast |
| Egflam | 0.85 | 0.00 | 0.00 | 0.30 | 0.08 | 18 | Matrix Fibroblast |
| Filip1l | 0.96 | 0.00 | 0.00 | 0.67 | 0.32 | 18 | Matrix Fibroblast |
| Rasl11a | 0.88 | 0.00 | 0.00 | 0.26 | 0.06 | 18 | Matrix Fibroblast |
| Hsd11b1 | 0.95 | 0.00 | 0.00 | 0.33 | 0.09 | 18 | Matrix Fibroblast |
| Nes | 0.88 | 0.00 | 0.00 | 0.55 | 0.21 | 18 | Matrix Fibroblast |
| Pcolce | 0.87 | 0.00 | 0.00 | 0.31 | 0.08 | 18 | Matrix Fibroblast |
| Tbx4 | 0.77 | 0.00 | 0.00 | 0.27 | 0.06 | 18 | Matrix Fibroblast |
| Pten | 0.89 | 0.00 | 0.00 | 0.37 | 0.12 | 18 | Matrix Fibroblast |
| Ifitm1 | 0.42 | 0.00 | 0.00 | 0.43 | 0.14 | 18 | Matrix Fibroblast |
| Isca1 | 0.88 | 0.00 | 0.00 | 0.33 | 0.10 | 18 | Matrix Fibroblast |
| Itgb1 | 0.75 | 0.00 | 0.00 | 0.87 | 0.63 | 18 | Matrix Fibroblast |
| Tpm2 | 0.77 | 0.00 | 0.00 | 0.39 | 0.13 | 18 | Matrix Fibroblast |
| Cryab | 0.96 | 0.00 | 0.00 | 0.39 | 0.14 | 18 | Matrix Fibroblast |
| Col3a1 | 0.26 | 0.00 | 0.00 | 0.74 | 0.32 | 18 | Matrix Fibroblast |
| 11-Sep | 0.98 | 0.00 | 0.00 | 0.50 | 0.22 | 18 | Matrix Fibroblast |
| Loxl2 | 0.65 | 0.00 | 0.00 | 0.40 | 0.13 | 18 | Matrix Fibroblast |
| Cdkn1c | 0.88 | 0.00 | 0.00 | 0.45 | 0.18 | 18 | Matrix Fibroblast |
| Itga4 | 0.78 | 0.00 | 0.00 | 0.31 | 0.09 | 18 | Matrix Fibroblast |
| Ndrg2 | 0.68 | 0.00 | 0.00 | 0.28 | 0.08 | 18 | Matrix Fibroblast |
| Rhoj | 0.77 | 0.00 | 0.00 | 0.46 | 0.18 | 18 | Matrix Fibroblast |
| Mxra8 | 0.79 | 0.00 | 0.00 | 0.42 | 0.16 | 18 | Matrix Fibroblast |
| Epb4.1l2 | 0.88 | 0.00 | 0.00 | 0.45 | 0.19 | 18 | Matrix Fibroblast |
| Ccdc80 | 0.80 | 0.00 | 0.00 | 0.33 | 0.11 | 18 | Matrix Fibroblast |
| Dkk3 | 0.75 | 0.00 | 0.00 | 0.34 | 0.11 | 18 | Matrix Fibroblast |
| AK157302 | 0.79 | 0.00 | 0.00 | 0.27 | 0.08 | 18 | Matrix Fibroblast |
| Gnb4 | 0.84 | 0.00 | 0.00 | 0.35 | 0.13 | 18 | Matrix Fibroblast |
| Mylk | 0.68 | 0.00 | 0.00 | 0.56 | 0.27 | 18 | Matrix Fibroblast |
| Tbx2 | 0.80 | 0.00 | 0.00 | 0.33 | 0.12 | 18 | Matrix Fibroblast |
| Lims1 | 0.86 | 0.00 | 0.00 | 0.54 | 0.28 | 18 | Matrix Fibroblast |
| Eva1b | 0.81 | 0.00 | 0.00 | 0.44 | 0.20 | 18 | Matrix Fibroblast |
| Tgfb2 | 0.92 | 0.00 | 0.00 | 0.36 | 0.14 | 18 | Matrix Fibroblast |
| Cbfa2t3 | 0.75 | 0.00 | 0.00 | 0.37 | 0.14 | 18 | Matrix Fibroblast |
| Armcx3 | 0.74 | 0.00 | 0.00 | 0.30 | 0.11 | 18 | Matrix Fibroblast |
| Agpat5 | 0.84 | 0.00 | 0.00 | 0.28 | 0.09 | 18 | Matrix Fibroblast |
| Acta2 | 0.75 | 0.00 | 0.00 | 0.43 | 0.19 | 18 | Matrix Fibroblast |
| Fstl1 | 0.36 | 0.00 | 0.00 | 0.76 | 0.44 | 18 | Matrix Fibroblast |
| Fam162a | 0.82 | 0.00 | 0.00 | 0.35 | 0.14 | 18 | Matrix Fibroblast |
| Ifitm3 | 0.54 | 0.00 | 0.00 | 0.68 | 0.39 | 18 | Matrix Fibroblast |
| App | 0.55 | 0.00 | 0.00 | 0.80 | 0.59 | 18 | Matrix Fibroblast |
| Mgst3 | 0.77 | 0.00 | 0.00 | 0.27 | 0.10 | 18 | Matrix Fibroblast |
| Oaz2 | 0.74 | 0.00 | 0.00 | 0.52 | 0.28 | 18 | Matrix Fibroblast |
| Laptm4a | 0.61 | 0.00 | 0.00 | 0.72 | 0.52 | 18 | Matrix Fibroblast |
| Tns1 | 0.60 | 0.00 | 0.00 | 0.48 | 0.23 | 18 | Matrix Fibroblast |
| Mfap2 | 0.54 | 0.00 | 0.00 | 0.63 | 0.35 | 18 | Matrix Fibroblast |
| Ssr3 | 0.76 | 0.00 | 0.00 | 0.53 | 0.31 | 18 | Matrix Fibroblast |
| Mat2a | 0.78 | 0.00 | 0.00 | 0.50 | 0.27 | 18 | Matrix Fibroblast |
| Vasp | 0.72 | 0.00 | 0.00 | 0.40 | 0.19 | 18 | Matrix Fibroblast |
| G0s2 | 0.50 | 0.00 | 0.00 | 0.33 | 0.13 | 18 | Matrix Fibroblast |
| Zfhx3 | 0.80 | 0.00 | 0.00 | 0.37 | 0.17 | 18 | Matrix Fibroblast |
| Tagln | 0.59 | 0.00 | 0.00 | 0.35 | 0.15 | 18 | Matrix Fibroblast |
| RP23-103I12.13 | 0.52 | 0.00 | 0.00 | 0.27 | 0.10 | 18 | Matrix Fibroblast |
| Fam104a | 0.74 | 0.00 | 0.00 | 0.38 | 0.18 | 18 | Matrix Fibroblast |
| Serpinh1 | 0.68 | 0.00 | 0.00 | 0.61 | 0.39 | 18 | Matrix Fibroblast |
| Gpx3 | 0.60 | 0.00 | 0.00 | 0.32 | 0.14 | 18 | Matrix Fibroblast |
| Mest | 0.81 | 0.00 | 0.00 | 0.40 | 0.20 | 18 | Matrix Fibroblast |
| Col5a2 | 0.62 | 0.00 | 0.00 | 0.38 | 0.18 | 18 | Matrix Fibroblast |
| Rbmx | 0.76 | 0.00 | 0.00 | 0.27 | 0.11 | 18 | Matrix Fibroblast |
| Eif4e3 | 0.66 | 0.00 | 0.00 | 0.26 | 0.11 | 18 | Matrix Fibroblast |
| Nptn | 0.73 | 0.00 | 0.00 | 0.40 | 0.21 | 18 | Matrix Fibroblast |
| Bgn | 0.43 | 0.00 | 0.00 | 0.47 | 0.24 | 18 | Matrix Fibroblast |
| Lgals1 | 0.38 | 0.00 | 0.00 | 0.69 | 0.41 | 18 | Matrix Fibroblast |
| Nfia | 0.68 | 0.00 | 0.00 | 0.43 | 0.24 | 18 | Matrix Fibroblast |
| Tsc22d1 | 0.54 | 0.00 | 0.00 | 0.55 | 0.33 | 18 | Matrix Fibroblast |
| Tpm4 | 0.63 | 0.00 | 0.00 | 0.65 | 0.49 | 18 | Matrix Fibroblast |
| mt-Cytb | 0.37 | 0.00 | 0.00 | 0.90 | 0.82 | 18 | Matrix Fibroblast |
| Gpx8 | 0.58 | 0.00 | 0.00 | 0.31 | 0.14 | 18 | Matrix Fibroblast |
| Ginm1 | 0.65 | 0.00 | 0.00 | 0.41 | 0.23 | 18 | Matrix Fibroblast |
| Tuba1a | 0.47 | 0.00 | 0.00 | 0.70 | 0.51 | 18 | Matrix Fibroblast |
| Ppic | 0.52 | 0.00 | 0.00 | 0.56 | 0.38 | 18 | Matrix Fibroblast |
| Csnk1e | 0.55 | 0.00 | 0.00 | 0.28 | 0.13 | 18 | Matrix Fibroblast |
| Fbn1 | 0.55 | 0.00 | 0.00 | 0.34 | 0.17 | 18 | Matrix Fibroblast |
| Myh9 | 0.54 | 0.00 | 0.00 | 0.51 | 0.33 | 18 | Matrix Fibroblast |
| Rell1 | 0.62 | 0.00 | 0.00 | 0.37 | 0.20 | 18 | Matrix Fibroblast |
| Rap2a | 0.56 | 0.00 | 0.00 | 0.32 | 0.16 | 18 | Matrix Fibroblast |
| Ccnd2 | 0.57 | 0.00 | 0.00 | 0.46 | 0.28 | 18 | Matrix Fibroblast |
| Acaa2 | 0.61 | 0.00 | 0.00 | 0.31 | 0.16 | 18 | Matrix Fibroblast |
| Spred1 | 0.63 | 0.00 | 0.00 | 0.26 | 0.12 | 18 | Matrix Fibroblast |
| Pcdhga9 | 0.55 | 0.00 | 0.00 | 0.33 | 0.18 | 18 | Matrix Fibroblast |
| Rsu1 | 0.58 | 0.00 | 0.00 | 0.35 | 0.20 | 18 | Matrix Fibroblast |
| Slc25a4 | 0.45 | 0.00 | 0.00 | 0.65 | 0.50 | 18 | Matrix Fibroblast |
| Ilk | 0.56 | 0.00 | 0.00 | 0.39 | 0.23 | 18 | Matrix Fibroblast |
| mt-Atp6 | 0.29 | 0.00 | 0.00 | 0.91 | 0.85 | 18 | Matrix Fibroblast |
| mt-Nd4 | 0.33 | 0.00 | 0.00 | 0.86 | 0.75 | 18 | Matrix Fibroblast |
| Tln1 | 0.51 | 0.00 | 0.00 | 0.55 | 0.40 | 18 | Matrix Fibroblast |
| Ppfibp1 | 0.54 | 0.00 | 0.00 | 0.27 | 0.14 | 18 | Matrix Fibroblast |
| Cnn2 | 0.48 | 0.00 | 0.00 | 0.55 | 0.39 | 18 | Matrix Fibroblast |
| Prex2 | 0.54 | 0.00 | 0.00 | 0.33 | 0.18 | 18 | Matrix Fibroblast |
| mt-Nd5 | 0.31 | 0.00 | 0.00 | 0.85 | 0.76 | 18 | Matrix Fibroblast |
| Dnajc10 | 0.47 | 0.00 | 0.00 | 0.28 | 0.15 | 18 | Matrix Fibroblast |
| Pja2 | 0.50 | 0.00 | 0.00 | 0.27 | 0.14 | 18 | Matrix Fibroblast |
| Ckb | 0.47 | 0.00 | 0.00 | 0.27 | 0.14 | 18 | Matrix Fibroblast |
| Cd302 | 0.52 | 0.00 | 0.00 | 0.27 | 0.14 | 18 | Matrix Fibroblast |
| Gm7676 | 0.35 | 0.00 | 0.00 | 0.33 | 0.19 | 18 | Matrix Fibroblast |
| Cd63 | 0.41 | 0.00 | 0.00 | 0.45 | 0.30 | 18 | Matrix Fibroblast |
| Myl12a | 0.36 | 0.00 | 0.00 | 0.75 | 0.66 | 18 | Matrix Fibroblast |
| Camk2n1 | 0.50 | 0.00 | 0.00 | 0.26 | 0.14 | 18 | Matrix Fibroblast |
| Pam | 0.37 | 0.00 | 0.00 | 0.31 | 0.17 | 18 | Matrix Fibroblast |
| mt-Nd2 | 0.32 | 0.00 | 0.00 | 0.78 | 0.70 | 18 | Matrix Fibroblast |
| Dpysl2 | 0.46 | 0.00 | 0.00 | 0.29 | 0.18 | 18 | Matrix Fibroblast |
| Nedd4 | 0.30 | 0.00 | 0.00 | 0.66 | 0.53 | 18 | Matrix Fibroblast |
| Rbms1 | 0.45 | 0.00 | 0.00 | 0.50 | 0.39 | 18 | Matrix Fibroblast |
| Tpm1 | 0.49 | 0.00 | 0.00 | 0.65 | 0.52 | 18 | Matrix Fibroblast |
| Tacc1 | 0.46 | 0.00 | 0.00 | 0.34 | 0.23 | 18 | Matrix Fibroblast |
| Cdc42bpa | 0.45 | 0.00 | 0.00 | 0.26 | 0.15 | 18 | Matrix Fibroblast |
| Lamc1 | 0.44 | 0.00 | 0.00 | 0.27 | 0.16 | 18 | Matrix Fibroblast |
| Crip1 | 0.26 | 0.00 | 0.00 | 0.61 | 0.48 | 18 | Matrix Fibroblast |
| Crip2 | 0.26 | 0.00 | 0.00 | 0.57 | 0.41 | 18 | Matrix Fibroblast |
| Mxd4 | 0.38 | 0.00 | 0.00 | 0.50 | 0.38 | 18 | Matrix Fibroblast |
| Hsp90b1 | 0.29 | 0.00 | 0.00 | 0.82 | 0.76 | 18 | Matrix Fibroblast |
| Serinc1 | 0.39 | 0.00 | 0.00 | 0.41 | 0.29 | 18 | Matrix Fibroblast |
| Eid1 | 0.39 | 0.00 | 0.00 | 0.41 | 0.29 | 18 | Matrix Fibroblast |
| Lpp | 0.40 | 0.00 | 0.00 | 0.33 | 0.22 | 18 | Matrix Fibroblast |
| Dync1i2 | 0.34 | 0.00 | 0.00 | 0.59 | 0.48 | 18 | Matrix Fibroblast |
| Ktn1 | 0.45 | 0.00 | 0.00 | 0.48 | 0.38 | 18 | Matrix Fibroblast |
| Myadm | 0.38 | 0.00 | 0.00 | 0.30 | 0.20 | 18 | Matrix Fibroblast |
| Map1lc3a | 0.39 | 0.00 | 0.00 | 0.30 | 0.20 | 18 | Matrix Fibroblast |
| Thra | 0.40 | 0.00 | 0.00 | 0.27 | 0.17 | 18 | Matrix Fibroblast |
| Tcf4 | 0.30 | 0.00 | 0.00 | 0.53 | 0.41 | 18 | Matrix Fibroblast |
| mt-Nd3 | 0.32 | 0.00 | 0.00 | 0.58 | 0.48 | 18 | Matrix Fibroblast |
| Htatsf1 | 0.46 | 0.00 | 0.00 | 0.36 | 0.26 | 18 | Matrix Fibroblast |
| Brd3 | 0.42 | 0.00 | 0.00 | 0.35 | 0.25 | 18 | Matrix Fibroblast |
| Timp3 | 0.58 | 0.00 | 0.00 | 0.44 | 0.36 | 18 | Matrix Fibroblast |
| Leprot | 0.37 | 0.00 | 0.00 | 0.42 | 0.33 | 18 | Matrix Fibroblast |
| Mprip | 0.41 | 0.00 | 0.00 | 0.30 | 0.21 | 18 | Matrix Fibroblast |
| Phlda1 | 0.31 | 0.00 | 0.00 | 0.30 | 0.20 | 18 | Matrix Fibroblast |
| Ppib | 0.32 | 0.00 | 0.00 | 0.51 | 0.43 | 18 | Matrix Fibroblast |
| Ptms | 0.28 | 0.00 | 0.00 | 0.50 | 0.40 | 18 | Matrix Fibroblast |
| Itm2c | 0.33 | 0.00 | 0.00 | 0.31 | 0.23 | 18 | Matrix Fibroblast |
| Eif4g3 | 0.42 | 0.00 | 0.00 | 0.36 | 0.27 | 18 | Matrix Fibroblast |
| Ostc | 0.39 | 0.00 | 0.00 | 0.32 | 0.23 | 18 | Matrix Fibroblast |
| Sepw1 | 0.37 | 0.00 | 0.00 | 0.28 | 0.20 | 18 | Matrix Fibroblast |
| Pnn | 0.27 | 0.00 | 0.00 | 0.61 | 0.54 | 18 | Matrix Fibroblast |
| Calu | 0.31 | 0.00 | 0.00 | 0.33 | 0.24 | 18 | Matrix Fibroblast |
| Vezf1 | 0.35 | 0.00 | 0.00 | 0.32 | 0.24 | 18 | Matrix Fibroblast |
| Cnn3 | 0.25 | 0.00 | 0.00 | 0.41 | 0.30 | 18 | Matrix Fibroblast |
| Ube3a | 0.36 | 0.00 | 0.00 | 0.26 | 0.18 | 18 | Matrix Fibroblast |
| Id3 | 0.29 | 0.00 | 0.00 | 0.54 | 0.46 | 18 | Matrix Fibroblast |
| Rbpms | 0.31 | 0.00 | 0.00 | 0.27 | 0.19 | 18 | Matrix Fibroblast |
| Tmem30a | 0.40 | 0.00 | 0.00 | 0.36 | 0.28 | 18 | Matrix Fibroblast |
| Tmem50a | 0.31 | 0.00 | 0.00 | 0.52 | 0.45 | 18 | Matrix Fibroblast |
| Gm10222 | 0.28 | 0.00 | 0.00 | 0.51 | 0.43 | 18 | Matrix Fibroblast |
| Ccdc88a | 0.35 | 0.00 | 0.00 | 0.33 | 0.24 | 18 | Matrix Fibroblast |
| 2410006H16Rik | 0.36 | 0.00 | 0.00 | 0.34 | 0.26 | 18 | Matrix Fibroblast |
| Canx | 0.29 | 0.00 | 0.00 | 0.56 | 0.49 | 18 | Matrix Fibroblast |
| Pfdn1 | 0.38 | 0.00 | 0.00 | 0.34 | 0.26 | 18 | Matrix Fibroblast |
| Ddah2 | 0.25 | 0.00 | 0.00 | 0.35 | 0.26 | 18 | Matrix Fibroblast |
| Hspa5 | 0.28 | 0.00 | 0.00 | 0.61 | 0.55 | 18 | Matrix Fibroblast |
| Tmem176b | 0.26 | 0.00 | 0.00 | 0.42 | 0.34 | 18 | Matrix Fibroblast |
| Aldh2 | 0.30 | 0.00 | 0.00 | 0.40 | 0.33 | 18 | Matrix Fibroblast |
| Zbtb20 | 0.34 | 0.00 | 0.01 | 0.34 | 0.27 | 18 | Matrix Fibroblast |
| Jak1 | 0.28 | 0.00 | 0.01 | 0.33 | 0.25 | 18 | Matrix Fibroblast |
| Ppp1r12a | 0.27 | 0.00 | 0.01 | 0.33 | 0.25 | 18 | Matrix Fibroblast |
| Selk | 0.26 | 0.00 | 0.01 | 0.39 | 0.32 | 18 | Matrix Fibroblast |
| Flna | 0.31 | 0.00 | 0.01 | 0.30 | 0.23 | 18 | Matrix Fibroblast |
| Coro1b | 0.35 | 0.00 | 0.02 | 0.31 | 0.24 | 18 | Matrix Fibroblast |
| B230219D22Rik | 0.28 | 0.00 | 0.04 | 0.34 | 0.28 | 18 | Matrix Fibroblast |
| Ccl21a | 3.83 | 0.00 | 0.00 | 0.28 | 0.01 | 19 | Endothelial |
| Gm21541 | 3.72 | 0.00 | 0.00 | 0.29 | 0.01 | 19 | Endothelial |
| Gm13304 | 3.70 | 0.00 | 0.00 | 0.29 | 0.00 | 19 | Endothelial |
| Gm10591 | 2.51 | 0.00 | 0.00 | 0.26 | 0.00 | 19 | Endothelial |
| Mmrn1 | 1.60 | 0.00 | 0.00 | 0.30 | 0.00 | 19 | Endothelial |
| Igf2 | 1.55 | 0.00 | 0.00 | 0.29 | 0.07 | 19 | Endothelial |
| H19 | 1.58 | 0.00 | 0.00 | 0.28 | 0.07 | 19 | Endothelial |
| Arl4a | 0.81 | 0.00 | 0.00 | 0.35 | 0.11 | 19 | Endothelial |
| Lhfpl2 | 1.27 | 0.00 | 0.00 | 0.27 | 0.09 | 19 | Endothelial |
| Tubb4b-ps1 | 0.95 | 0.00 | 0.00 | 0.25 | 0.09 | 19 | Endothelial |
| Hba-a2 | 2.21 | 0.00 | 0.00 | 0.79 | 0.70 | 19 | Endothelial |
| Hba-a1 | 2.21 | 0.00 | 0.00 | 0.78 | 0.71 | 19 | Endothelial |
| Hbb-bt | 2.20 | 0.00 | 0.00 | 0.80 | 0.71 | 19 | Endothelial |
| Hbb-bs | 2.12 | 0.00 | 0.00 | 0.85 | 0.81 | 19 | Endothelial |
| Cd24a | 0.60 | 0.00 | 0.00 | 0.28 | 0.13 | 19 | Endothelial |
| Atg3 | 1.06 | 0.00 | 0.00 | 0.33 | 0.19 | 19 | Endothelial |
| Tia1 | 0.88 | 0.00 | 0.00 | 0.36 | 0.22 | 19 | Endothelial |
| Gng11 | 0.94 | 0.00 | 0.00 | 0.38 | 0.29 | 19 | Endothelial |
| Maf | 0.37 | 0.00 | 0.00 | 0.30 | 0.19 | 19 | Endothelial |
| Creg1 | 0.35 | 0.00 | 0.00 | 0.28 | 0.17 | 19 | Endothelial |
| Rab11a | 0.39 | 0.00 | 0.00 | 0.37 | 0.26 | 19 | Endothelial |
| Pabpc1 | 0.40 | 0.00 | 0.00 | 0.80 | 0.81 | 19 | Endothelial |
| Nudt4 | 0.37 | 0.00 | 0.04 | 0.25 | 0.17 | 19 | Endothelial |
| Igfbp3 | 3.21 | 0.00 | 0.00 | 0.81 | 0.06 | 20 | Matrix Fibroblast |
| Enpp2 | 2.50 | 0.00 | 0.00 | 0.72 | 0.06 | 20 | Matrix Fibroblast |
| Tgfbi | 2.47 | 0.00 | 0.00 | 0.88 | 0.18 | 20 | Matrix Fibroblast |
| Dnm3os | 2.38 | 0.00 | 0.00 | 0.69 | 0.05 | 20 | Matrix Fibroblast |
| Rgs2 | 2.35 | 0.00 | 0.00 | 0.89 | 0.27 | 20 | Matrix Fibroblast |
| Agt | 2.09 | 0.00 | 0.00 | 0.76 | 0.09 | 20 | Matrix Fibroblast |
| Fibin | 2.02 | 0.00 | 0.00 | 0.68 | 0.12 | 20 | Matrix Fibroblast |
| Meg3 | 1.87 | 0.00 | 0.00 | 0.88 | 0.21 | 20 | Matrix Fibroblast |
| Fam105a | 1.80 | 0.00 | 0.00 | 0.59 | 0.06 | 20 | Matrix Fibroblast |
| Penk | 1.72 | 0.00 | 0.00 | 0.34 | 0.01 | 20 | Matrix Fibroblast |
| Tgfbr3 | 1.69 | 0.00 | 0.00 | 0.56 | 0.08 | 20 | Matrix Fibroblast |
| Scara5 | 1.37 | 0.00 | 0.00 | 0.31 | 0.01 | 20 | Matrix Fibroblast |
| Crispld2 | 1.36 | 0.00 | 0.00 | 0.32 | 0.03 | 20 | Matrix Fibroblast |
| Kcnj8 | 1.29 | 0.00 | 0.00 | 0.28 | 0.02 | 20 | Matrix Fibroblast |
| H19 | 1.66 | 0.00 | 0.00 | 0.48 | 0.07 | 20 | Matrix Fibroblast |
| Egr1 | 2.07 | 0.00 | 0.00 | 0.62 | 0.13 | 20 | Matrix Fibroblast |
| Fos | 2.11 | 0.00 | 0.00 | 0.72 | 0.19 | 20 | Matrix Fibroblast |
| Jun | 1.86 | 0.00 | 0.00 | 0.87 | 0.36 | 20 | Matrix Fibroblast |
| Trp53inp1 | 1.54 | 0.00 | 0.00 | 0.45 | 0.07 | 20 | Matrix Fibroblast |
| Hspb1 | 2.23 | 0.00 | 0.00 | 0.59 | 0.13 | 20 | Matrix Fibroblast |
| Ctsl | 1.78 | 0.00 | 0.00 | 0.87 | 0.39 | 20 | Matrix Fibroblast |
| Gpx3 | 1.58 | 0.00 | 0.00 | 0.60 | 0.13 | 20 | Matrix Fibroblast |
| Lpin1 | 1.22 | 0.00 | 0.00 | 0.29 | 0.03 | 20 | Matrix Fibroblast |
| Rhou | 1.35 | 0.00 | 0.00 | 0.38 | 0.06 | 20 | Matrix Fibroblast |
| Fmo2 | 1.59 | 0.00 | 0.00 | 0.64 | 0.17 | 20 | Matrix Fibroblast |
| Pcsk5 | 1.46 | 0.00 | 0.00 | 0.47 | 0.10 | 20 | Matrix Fibroblast |
| Tsc22d3 | 1.60 | 0.00 | 0.00 | 0.69 | 0.24 | 20 | Matrix Fibroblast |
| Loxl2 | 1.53 | 0.00 | 0.00 | 0.53 | 0.13 | 20 | Matrix Fibroblast |
| Adamts1 | 1.60 | 0.00 | 0.00 | 0.64 | 0.21 | 20 | Matrix Fibroblast |
| Zbtb16 | 1.17 | 0.00 | 0.00 | 0.33 | 0.05 | 20 | Matrix Fibroblast |
| Klf9 | 1.43 | 0.00 | 0.00 | 0.61 | 0.19 | 20 | Matrix Fibroblast |
| Cdkn1a | 1.22 | 0.00 | 0.00 | 0.57 | 0.16 | 20 | Matrix Fibroblast |
| RP23-103I12.13 | 1.35 | 0.00 | 0.00 | 0.44 | 0.10 | 20 | Matrix Fibroblast |
| Ybx3 | 1.36 | 0.00 | 0.00 | 0.60 | 0.19 | 20 | Matrix Fibroblast |
| Npr3 | 1.48 | 0.00 | 0.00 | 0.50 | 0.13 | 20 | Matrix Fibroblast |
| Bmp2 | 1.15 | 0.00 | 0.00 | 0.26 | 0.04 | 20 | Matrix Fibroblast |
| Btg2 | 1.78 | 0.00 | 0.00 | 0.62 | 0.23 | 20 | Matrix Fibroblast |
| Glul | 1.37 | 0.00 | 0.00 | 0.62 | 0.22 | 20 | Matrix Fibroblast |
| Junb | 1.49 | 0.00 | 0.00 | 0.58 | 0.19 | 20 | Matrix Fibroblast |
| Gpr64 | 1.07 | 0.00 | 0.00 | 0.29 | 0.05 | 20 | Matrix Fibroblast |
| Bnip3 | 1.18 | 0.00 | 0.00 | 0.38 | 0.08 | 20 | Matrix Fibroblast |
| Socs2 | 1.30 | 0.00 | 0.00 | 0.73 | 0.33 | 20 | Matrix Fibroblast |
| Eln | 1.01 | 0.00 | 0.00 | 0.53 | 0.16 | 20 | Matrix Fibroblast |
| Tshz1 | 1.30 | 0.00 | 0.00 | 0.43 | 0.12 | 20 | Matrix Fibroblast |
| Hhip | 1.03 | 0.00 | 0.00 | 0.33 | 0.07 | 20 | Matrix Fibroblast |
| Il6ra | 1.12 | 0.00 | 0.00 | 0.40 | 0.10 | 20 | Matrix Fibroblast |
| Ndrg2 | 1.00 | 0.00 | 0.00 | 0.35 | 0.08 | 20 | Matrix Fibroblast |
| Marcks | 1.08 | 0.00 | 0.00 | 0.84 | 0.50 | 20 | Matrix Fibroblast |
| Fkbp5 | 1.16 | 0.00 | 0.00 | 0.36 | 0.09 | 20 | Matrix Fibroblast |
| D4Wsu53e | 1.14 | 0.00 | 0.00 | 0.84 | 0.60 | 20 | Matrix Fibroblast |
| Col1a2 | 0.49 | 0.00 | 0.00 | 0.91 | 0.39 | 20 | Matrix Fibroblast |
| Gm13889 | 1.16 | 0.00 | 0.00 | 0.30 | 0.07 | 20 | Matrix Fibroblast |
| Aldh2 | 1.27 | 0.00 | 0.00 | 0.67 | 0.33 | 20 | Matrix Fibroblast |
| Mgp | 1.09 | 0.00 | 0.00 | 0.74 | 0.31 | 20 | Matrix Fibroblast |
| Lox | 1.07 | 0.00 | 0.00 | 0.45 | 0.13 | 20 | Matrix Fibroblast |
| Cyr61 | 1.46 | 0.00 | 0.00 | 0.42 | 0.13 | 20 | Matrix Fibroblast |
| Ism1 | 1.09 | 0.00 | 0.00 | 0.35 | 0.09 | 20 | Matrix Fibroblast |
| Igf2 | 1.01 | 0.00 | 0.00 | 0.31 | 0.07 | 20 | Matrix Fibroblast |
| Pdzrn3 | 1.01 | 0.00 | 0.00 | 0.26 | 0.06 | 20 | Matrix Fibroblast |
| Zfp36 | 1.29 | 0.00 | 0.00 | 0.46 | 0.16 | 20 | Matrix Fibroblast |
| Net1 | 1.14 | 0.00 | 0.00 | 0.48 | 0.17 | 20 | Matrix Fibroblast |
| Arrdc3 | 1.12 | 0.00 | 0.00 | 0.38 | 0.11 | 20 | Matrix Fibroblast |
| Prdm6 | 0.93 | 0.00 | 0.00 | 0.28 | 0.06 | 20 | Matrix Fibroblast |
| Pros1 | 1.02 | 0.00 | 0.00 | 0.35 | 0.10 | 20 | Matrix Fibroblast |
| Rora | 1.15 | 0.00 | 0.00 | 0.35 | 0.10 | 20 | Matrix Fibroblast |
| Malat1 | 0.45 | 0.00 | 0.00 | 1.00 | 1.00 | 20 | Matrix Fibroblast |
| Atp2b1 | 0.99 | 0.00 | 0.00 | 0.80 | 0.58 | 20 | Matrix Fibroblast |
| Pdgfra | 0.86 | 0.00 | 0.00 | 0.26 | 0.06 | 20 | Matrix Fibroblast |
| Creb3l2 | 0.99 | 0.00 | 0.00 | 0.39 | 0.13 | 20 | Matrix Fibroblast |
| Tbx5 | 1.03 | 0.00 | 0.00 | 0.29 | 0.08 | 20 | Matrix Fibroblast |
| Nbl1 | 0.91 | 0.00 | 0.00 | 0.27 | 0.07 | 20 | Matrix Fibroblast |
| Socs3 | 1.06 | 0.00 | 0.00 | 0.27 | 0.07 | 20 | Matrix Fibroblast |
| Drap1 | 1.17 | 0.00 | 0.00 | 0.53 | 0.25 | 20 | Matrix Fibroblast |
| Apoe | 0.79 | 0.00 | 0.00 | 0.60 | 0.28 | 20 | Matrix Fibroblast |
| Zfp36l1 | 1.09 | 0.00 | 0.00 | 0.70 | 0.45 | 20 | Matrix Fibroblast |
| Dusp1 | 1.10 | 0.00 | 0.00 | 0.43 | 0.17 | 20 | Matrix Fibroblast |
| Col1a1 | 0.58 | 0.00 | 0.00 | 0.59 | 0.26 | 20 | Matrix Fibroblast |
| Gm26669 | 1.18 | 0.00 | 0.00 | 0.49 | 0.22 | 20 | Matrix Fibroblast |
| Thbs1 | 1.08 | 0.00 | 0.00 | 0.37 | 0.13 | 20 | Matrix Fibroblast |
| Col3a1 | 0.40 | 0.00 | 0.00 | 0.71 | 0.33 | 20 | Matrix Fibroblast |
| Ednrb | 0.71 | 0.00 | 0.00 | 0.41 | 0.15 | 20 | Matrix Fibroblast |
| Ier2 | 1.11 | 0.00 | 0.00 | 0.50 | 0.25 | 20 | Matrix Fibroblast |
| Mark3 | 1.08 | 0.00 | 0.00 | 0.40 | 0.16 | 20 | Matrix Fibroblast |
| Gm9847 | 0.96 | 0.00 | 0.00 | 0.39 | 0.16 | 20 | Matrix Fibroblast |
| Nrep | 0.71 | 0.00 | 0.00 | 0.73 | 0.43 | 20 | Matrix Fibroblast |
| Higd1a | 1.16 | 0.00 | 0.00 | 0.41 | 0.18 | 20 | Matrix Fibroblast |
| 1700025G04Rik | 0.94 | 0.00 | 0.00 | 0.29 | 0.10 | 20 | Matrix Fibroblast |
| Samd4 | 0.91 | 0.00 | 0.00 | 0.27 | 0.08 | 20 | Matrix Fibroblast |
| Insig2 | 0.95 | 0.00 | 0.00 | 0.29 | 0.10 | 20 | Matrix Fibroblast |
| Rhoj | 0.88 | 0.00 | 0.00 | 0.42 | 0.18 | 20 | Matrix Fibroblast |
| Mfap2 | 0.81 | 0.00 | 0.00 | 0.61 | 0.35 | 20 | Matrix Fibroblast |
| Pxdn | 0.79 | 0.00 | 0.00 | 0.29 | 0.11 | 20 | Matrix Fibroblast |
| Lamc1 | 0.80 | 0.00 | 0.00 | 0.36 | 0.16 | 20 | Matrix Fibroblast |
| Gm9790 | 1.02 | 0.00 | 0.00 | 0.36 | 0.17 | 20 | Matrix Fibroblast |
| Lama4 | 0.90 | 0.00 | 0.00 | 0.32 | 0.13 | 20 | Matrix Fibroblast |
| Fstl1 | 0.57 | 0.00 | 0.00 | 0.67 | 0.45 | 20 | Matrix Fibroblast |
| Chkb | 0.78 | 0.00 | 0.00 | 0.26 | 0.10 | 20 | Matrix Fibroblast |
| Mettl7a1 | 0.91 | 0.00 | 0.00 | 0.42 | 0.22 | 20 | Matrix Fibroblast |
| Nrip1 | 0.88 | 0.00 | 0.00 | 0.29 | 0.12 | 20 | Matrix Fibroblast |
| Igf1 | 0.89 | 0.00 | 0.00 | 0.32 | 0.14 | 20 | Matrix Fibroblast |
| Sfrs18 | 0.68 | 0.00 | 0.00 | 0.72 | 0.59 | 20 | Matrix Fibroblast |
| Col4a1 | 0.49 | 0.00 | 0.00 | 0.73 | 0.49 | 20 | Matrix Fibroblast |
| Hba-a2 | 0.41 | 0.00 | 0.00 | 0.86 | 0.70 | 20 | Matrix Fibroblast |
| Wnt5a | 0.67 | 0.00 | 0.00 | 0.28 | 0.10 | 20 | Matrix Fibroblast |
| Tcf25 | 0.67 | 0.00 | 0.00 | 0.67 | 0.51 | 20 | Matrix Fibroblast |
| Mxra8 | 0.69 | 0.00 | 0.00 | 0.36 | 0.16 | 20 | Matrix Fibroblast |
| Mmp2 | 0.71 | 0.00 | 0.00 | 0.32 | 0.14 | 20 | Matrix Fibroblast |
| Hba-a1 | 0.41 | 0.00 | 0.00 | 0.88 | 0.70 | 20 | Matrix Fibroblast |
| Clk1 | 0.70 | 0.00 | 0.00 | 0.54 | 0.35 | 20 | Matrix Fibroblast |
| Gabarapl1 | 0.79 | 0.00 | 0.00 | 0.33 | 0.16 | 20 | Matrix Fibroblast |
| Celf2 | 0.74 | 0.00 | 0.00 | 0.47 | 0.28 | 20 | Matrix Fibroblast |
| Id2 | 1.00 | 0.00 | 0.00 | 0.52 | 0.33 | 20 | Matrix Fibroblast |
| Hbb-bs | 0.30 | 0.00 | 0.00 | 0.94 | 0.81 | 20 | Matrix Fibroblast |
| Med13l | 0.78 | 0.00 | 0.00 | 0.28 | 0.12 | 20 | Matrix Fibroblast |
| Cd81 | 0.62 | 0.00 | 0.00 | 0.59 | 0.41 | 20 | Matrix Fibroblast |
| Clip1 | 0.77 | 0.00 | 0.00 | 0.39 | 0.21 | 20 | Matrix Fibroblast |
| Grb10 | 0.76 | 0.00 | 0.00 | 0.33 | 0.16 | 20 | Matrix Fibroblast |
| Mllt3 | 0.86 | 0.00 | 0.00 | 0.26 | 0.11 | 20 | Matrix Fibroblast |
| Mpzl1 | 0.74 | 0.00 | 0.00 | 0.26 | 0.11 | 20 | Matrix Fibroblast |
| Ghr | 0.74 | 0.00 | 0.00 | 0.31 | 0.15 | 20 | Matrix Fibroblast |
| Luc7l2 | 0.71 | 0.00 | 0.00 | 0.55 | 0.39 | 20 | Matrix Fibroblast |
| Rb1cc1 | 0.79 | 0.00 | 0.00 | 0.47 | 0.30 | 20 | Matrix Fibroblast |
| Btg1 | 0.67 | 0.00 | 0.00 | 0.46 | 0.28 | 20 | Matrix Fibroblast |
| Timp3 | 0.37 | 0.00 | 0.00 | 0.56 | 0.36 | 20 | Matrix Fibroblast |
| Hmgb1 | 0.48 | 0.00 | 0.00 | 0.74 | 0.61 | 20 | Matrix Fibroblast |
| Zcchc11 | 0.69 | 0.00 | 0.00 | 0.36 | 0.21 | 20 | Matrix Fibroblast |
| Pmp22 | 0.48 | 0.00 | 0.00 | 0.50 | 0.32 | 20 | Matrix Fibroblast |
| Col4a2 | 0.43 | 0.00 | 0.00 | 0.61 | 0.43 | 20 | Matrix Fibroblast |
| Ccnl2 | 0.68 | 0.00 | 0.00 | 0.40 | 0.24 | 20 | Matrix Fibroblast |
| Nexn | 0.65 | 0.00 | 0.00 | 0.39 | 0.22 | 20 | Matrix Fibroblast |
| Ubb | 0.37 | 0.00 | 0.00 | 0.87 | 0.81 | 20 | Matrix Fibroblast |
| Zfp503 | 0.74 | 0.00 | 0.00 | 0.30 | 0.15 | 20 | Matrix Fibroblast |
| Jund | 0.69 | 0.00 | 0.00 | 0.43 | 0.28 | 20 | Matrix Fibroblast |
| Map1lc3a | 0.64 | 0.00 | 0.00 | 0.35 | 0.20 | 20 | Matrix Fibroblast |
| Txnip | 0.69 | 0.00 | 0.00 | 0.41 | 0.25 | 20 | Matrix Fibroblast |
| Mef2a | 0.69 | 0.00 | 0.00 | 0.50 | 0.36 | 20 | Matrix Fibroblast |
| Serpine2 | 0.46 | 0.00 | 0.00 | 0.44 | 0.25 | 20 | Matrix Fibroblast |
| Pik3r1 | 0.70 | 0.00 | 0.00 | 0.26 | 0.13 | 20 | Matrix Fibroblast |
| Arglu1 | 0.58 | 0.00 | 0.00 | 0.55 | 0.42 | 20 | Matrix Fibroblast |
| Hbb-bt | 0.28 | 0.00 | 0.00 | 0.85 | 0.71 | 20 | Matrix Fibroblast |
| Ilk | 0.67 | 0.00 | 0.00 | 0.37 | 0.23 | 20 | Matrix Fibroblast |
| Map1lc3b | 0.57 | 0.00 | 0.00 | 0.53 | 0.40 | 20 | Matrix Fibroblast |
| Pnrc1 | 0.65 | 0.00 | 0.00 | 0.34 | 0.20 | 20 | Matrix Fibroblast |
| Tbx3 | 0.49 | 0.00 | 0.00 | 0.36 | 0.21 | 20 | Matrix Fibroblast |
| Camk2n1 | 0.63 | 0.00 | 0.00 | 0.27 | 0.15 | 20 | Matrix Fibroblast |
| Lhfp | 0.55 | 0.00 | 0.00 | 0.29 | 0.16 | 20 | Matrix Fibroblast |
| Rhoc | 0.53 | 0.00 | 0.00 | 0.35 | 0.21 | 20 | Matrix Fibroblast |
| Nfkbia | 0.52 | 0.00 | 0.00 | 0.51 | 0.37 | 20 | Matrix Fibroblast |
| mt-Cytb | 0.28 | 0.00 | 0.00 | 0.88 | 0.82 | 20 | Matrix Fibroblast |
| Peg3 | 0.37 | 0.00 | 0.00 | 0.47 | 0.30 | 20 | Matrix Fibroblast |
| Tmem59 | 0.54 | 0.00 | 0.00 | 0.52 | 0.41 | 20 | Matrix Fibroblast |
| Csrp1 | 0.60 | 0.00 | 0.00 | 0.29 | 0.17 | 20 | Matrix Fibroblast |
| Tmem50a | 0.48 | 0.00 | 0.00 | 0.57 | 0.45 | 20 | Matrix Fibroblast |
| Eif1 | 0.34 | 0.00 | 0.00 | 0.74 | 0.67 | 20 | Matrix Fibroblast |
| Ube2b | 0.57 | 0.00 | 0.00 | 0.48 | 0.36 | 20 | Matrix Fibroblast |
| Dnajc3 | 0.57 | 0.00 | 0.00 | 0.39 | 0.27 | 20 | Matrix Fibroblast |
| Morf4l1 | 0.43 | 0.00 | 0.00 | 0.64 | 0.55 | 20 | Matrix Fibroblast |
| Zeb2 | 0.50 | 0.00 | 0.00 | 0.46 | 0.31 | 20 | Matrix Fibroblast |
| Nupr1 | 0.44 | 0.00 | 0.00 | 0.32 | 0.18 | 20 | Matrix Fibroblast |
| Zmym5 | 0.63 | 0.00 | 0.00 | 0.32 | 0.20 | 20 | Matrix Fibroblast |
| Arid4b | 0.65 | 0.00 | 0.00 | 0.45 | 0.34 | 20 | Matrix Fibroblast |
| Cald1 | 0.38 | 0.00 | 0.00 | 0.56 | 0.41 | 20 | Matrix Fibroblast |
| Cd44 | 0.59 | 0.00 | 0.00 | 0.32 | 0.20 | 20 | Matrix Fibroblast |
| Csrp2 | 0.51 | 0.00 | 0.00 | 0.29 | 0.17 | 20 | Matrix Fibroblast |
| Smc5 | 0.71 | 0.00 | 0.00 | 0.28 | 0.16 | 20 | Matrix Fibroblast |
| Slc43a3 | 0.56 | 0.00 | 0.00 | 0.35 | 0.22 | 20 | Matrix Fibroblast |
| Srrm2 | 0.45 | 0.00 | 0.00 | 0.67 | 0.61 | 20 | Matrix Fibroblast |
| Luc7l3 | 0.53 | 0.00 | 0.00 | 0.62 | 0.57 | 20 | Matrix Fibroblast |
| Ginm1 | 0.65 | 0.00 | 0.00 | 0.35 | 0.23 | 20 | Matrix Fibroblast |
| Nfib | 0.41 | 0.00 | 0.00 | 0.60 | 0.47 | 20 | Matrix Fibroblast |
| Tagln | 0.47 | 0.00 | 0.00 | 0.27 | 0.15 | 20 | Matrix Fibroblast |
| Pnn | 0.47 | 0.00 | 0.00 | 0.61 | 0.54 | 20 | Matrix Fibroblast |
| Arid5b | 0.65 | 0.00 | 0.00 | 0.34 | 0.22 | 20 | Matrix Fibroblast |
| Zfr | 0.59 | 0.00 | 0.00 | 0.39 | 0.29 | 20 | Matrix Fibroblast |
| Vezf1 | 0.58 | 0.00 | 0.00 | 0.35 | 0.24 | 20 | Matrix Fibroblast |
| Gm26384 | 0.33 | 0.00 | 0.00 | 0.60 | 0.48 | 20 | Matrix Fibroblast |
| Fth1 | 0.31 | 0.00 | 0.00 | 0.85 | 0.81 | 20 | Matrix Fibroblast |
| Rbm39 | 0.35 | 0.00 | 0.00 | 0.74 | 0.72 | 20 | Matrix Fibroblast |
| Rbbp6 | 0.58 | 0.00 | 0.00 | 0.45 | 0.35 | 20 | Matrix Fibroblast |
| Slc38a2 | 0.58 | 0.00 | 0.00 | 0.29 | 0.19 | 20 | Matrix Fibroblast |
| Kmt2e | 0.53 | 0.00 | 0.00 | 0.50 | 0.43 | 20 | Matrix Fibroblast |
| Atrx | 0.45 | 0.00 | 0.00 | 0.68 | 0.66 | 20 | Matrix Fibroblast |
| Vimp | 0.56 | 0.00 | 0.00 | 0.39 | 0.30 | 20 | Matrix Fibroblast |
| Sec62 | 0.45 | 0.00 | 0.00 | 0.52 | 0.45 | 20 | Matrix Fibroblast |
| Rbpms | 0.50 | 0.00 | 0.00 | 0.28 | 0.19 | 20 | Matrix Fibroblast |
| Ccar1 | 0.54 | 0.00 | 0.00 | 0.41 | 0.33 | 20 | Matrix Fibroblast |
| Gnas | 0.28 | 0.00 | 0.00 | 0.72 | 0.67 | 20 | Matrix Fibroblast |
| Fendrr | 0.29 | 0.00 | 0.00 | 0.32 | 0.20 | 20 | Matrix Fibroblast |
| Lmna | 0.51 | 0.00 | 0.00 | 0.30 | 0.20 | 20 | Matrix Fibroblast |
| Laptm4a | 0.39 | 0.00 | 0.00 | 0.59 | 0.52 | 20 | Matrix Fibroblast |
| Ptms | 0.36 | 0.00 | 0.00 | 0.49 | 0.40 | 20 | Matrix Fibroblast |
| Maged1 | 0.45 | 0.00 | 0.00 | 0.28 | 0.18 | 20 | Matrix Fibroblast |
| Cir1 | 0.46 | 0.00 | 0.00 | 0.30 | 0.21 | 20 | Matrix Fibroblast |
| Mmp14 | 0.40 | 0.00 | 0.00 | 0.35 | 0.25 | 20 | Matrix Fibroblast |
| Cd63 | 0.37 | 0.00 | 0.00 | 0.39 | 0.30 | 20 | Matrix Fibroblast |
| Sf3b1 | 0.39 | 0.00 | 0.00 | 0.53 | 0.47 | 20 | Matrix Fibroblast |
| Selk | 0.45 | 0.00 | 0.00 | 0.40 | 0.32 | 20 | Matrix Fibroblast |
| Srsf5 | 0.32 | 0.00 | 0.00 | 0.56 | 0.51 | 20 | Matrix Fibroblast |
| Ppap2b | 0.42 | 0.00 | 0.00 | 0.34 | 0.24 | 20 | Matrix Fibroblast |
| Serinc1 | 0.42 | 0.00 | 0.00 | 0.38 | 0.29 | 20 | Matrix Fibroblast |
| Itgb1 | 0.25 | 0.00 | 0.00 | 0.69 | 0.63 | 20 | Matrix Fibroblast |
| Svil | 0.54 | 0.00 | 0.00 | 0.25 | 0.17 | 20 | Matrix Fibroblast |
| Cirbp | 0.45 | 0.00 | 0.00 | 0.30 | 0.22 | 20 | Matrix Fibroblast |
| Srsf11 | 0.34 | 0.00 | 0.00 | 0.54 | 0.49 | 20 | Matrix Fibroblast |
| Hp1bp3 | 0.38 | 0.00 | 0.00 | 0.50 | 0.45 | 20 | Matrix Fibroblast |
| Ifitm2 | 0.31 | 0.00 | 0.00 | 0.41 | 0.32 | 20 | Matrix Fibroblast |
| Akap9 | 0.46 | 0.00 | 0.00 | 0.43 | 0.36 | 20 | Matrix Fibroblast |
| Rbms3 | 0.41 | 0.00 | 0.00 | 0.25 | 0.17 | 20 | Matrix Fibroblast |
| Atf4 | 0.46 | 0.00 | 0.00 | 0.35 | 0.28 | 20 | Matrix Fibroblast |
| Sdc4 | 0.35 | 0.00 | 0.00 | 0.32 | 0.23 | 20 | Matrix Fibroblast |
| Tmem176b | 0.41 | 0.00 | 0.00 | 0.42 | 0.34 | 20 | Matrix Fibroblast |
| Ppic | 0.30 | 0.00 | 0.00 | 0.47 | 0.38 | 20 | Matrix Fibroblast |
| Mxd4 | 0.43 | 0.00 | 0.01 | 0.44 | 0.38 | 20 | Matrix Fibroblast |
| Eea1 | 0.50 | 0.00 | 0.01 | 0.28 | 0.21 | 20 | Matrix Fibroblast |
| Tsc22d1 | 0.41 | 0.00 | 0.01 | 0.41 | 0.34 | 20 | Matrix Fibroblast |
| Ik | 0.45 | 0.00 | 0.01 | 0.41 | 0.36 | 20 | Matrix Fibroblast |
| Ssr3 | 0.41 | 0.00 | 0.01 | 0.37 | 0.31 | 20 | Matrix Fibroblast |
| Gas5 | 0.31 | 0.00 | 0.02 | 0.57 | 0.54 | 20 | Matrix Fibroblast |
| Brd3 | 0.40 | 0.00 | 0.02 | 0.32 | 0.25 | 20 | Matrix Fibroblast |
| Wbp5 | 0.25 | 0.00 | 0.02 | 0.56 | 0.48 | 20 | Matrix Fibroblast |
| Dek | 0.34 | 0.00 | 0.03 | 0.54 | 0.50 | 20 | Matrix Fibroblast |
| Sfpq | 0.37 | 0.00 | 0.03 | 0.42 | 0.37 | 20 | Matrix Fibroblast |
| Bgn | 0.26 | 0.00 | 0.03 | 0.33 | 0.24 | 20 | Matrix Fibroblast |
| Sfr1 | 0.49 | 0.00 | 0.03 | 0.39 | 0.34 | 20 | Matrix Fibroblast |
| Prrc2c | 0.34 | 0.00 | 0.04 | 0.64 | 0.63 | 20 | Matrix Fibroblast |
| Pf4 | 3.38 | 0.00 | 0.00 | 0.69 | 0.02 | 21 | Macrophage |
| C1qc | 3.05 | 0.00 | 0.00 | 0.75 | 0.01 | 21 | Macrophage |
| C1qb | 3.04 | 0.00 | 0.00 | 0.77 | 0.01 | 21 | Macrophage |
| C1qa | 2.82 | 0.00 | 0.00 | 0.71 | 0.01 | 21 | Macrophage |
| Mrc1 | 2.68 | 0.00 | 0.00 | 0.67 | 0.02 | 21 | Macrophage |
| Ms4a7 | 2.40 | 0.00 | 0.00 | 0.56 | 0.01 | 21 | Macrophage |
| Cx3cr1 | 2.22 | 0.00 | 0.00 | 0.58 | 0.03 | 21 | Macrophage |
| Dab2 | 2.17 | 0.00 | 0.00 | 0.62 | 0.07 | 21 | Macrophage |
| F13a1 | 2.04 | 0.00 | 0.00 | 0.47 | 0.02 | 21 | Macrophage |
| Fcrls | 1.95 | 0.00 | 0.00 | 0.47 | 0.00 | 21 | Macrophage |
| Aif1 | 1.92 | 0.00 | 0.00 | 0.59 | 0.03 | 21 | Macrophage |
| Ccl12 | 1.86 | 0.00 | 0.00 | 0.26 | 0.00 | 21 | Macrophage |
| Stab1 | 1.85 | 0.00 | 0.00 | 0.49 | 0.04 | 21 | Macrophage |
| Csf1r | 1.84 | 0.00 | 0.00 | 0.58 | 0.04 | 21 | Macrophage |
| Fcer1g | 1.79 | 0.00 | 0.00 | 0.83 | 0.11 | 21 | Macrophage |
| Ctss | 1.77 | 0.00 | 0.00 | 0.87 | 0.14 | 21 | Macrophage |
| Hpgds | 1.73 | 0.00 | 0.00 | 0.44 | 0.01 | 21 | Macrophage |
| Laptm5 | 1.65 | 0.00 | 0.00 | 0.79 | 0.13 | 21 | Macrophage |
| Gatm | 1.61 | 0.00 | 0.00 | 0.41 | 0.01 | 21 | Macrophage |
| Ms4a6c | 1.58 | 0.00 | 0.00 | 0.59 | 0.06 | 21 | Macrophage |
| Tyrobp | 1.49 | 0.00 | 0.00 | 0.80 | 0.12 | 21 | Macrophage |
| C3ar1 | 1.45 | 0.00 | 0.00 | 0.39 | 0.01 | 21 | Macrophage |
| Fcgr3 | 1.45 | 0.00 | 0.00 | 0.48 | 0.04 | 21 | Macrophage |
| Clec4n | 1.38 | 0.00 | 0.00 | 0.36 | 0.02 | 21 | Macrophage |
| Ccr1 | 1.35 | 0.00 | 0.00 | 0.42 | 0.03 | 21 | Macrophage |
| Fcgr2b | 1.34 | 0.00 | 0.00 | 0.41 | 0.03 | 21 | Macrophage |
| Clec4a2 | 1.27 | 0.00 | 0.00 | 0.40 | 0.03 | 21 | Macrophage |
| Emr1 | 1.25 | 0.00 | 0.00 | 0.31 | 0.01 | 21 | Macrophage |
| Gpr65 | 1.23 | 0.00 | 0.00 | 0.31 | 0.02 | 21 | Macrophage |
| AI607873 | 1.17 | 0.00 | 0.00 | 0.35 | 0.03 | 21 | Macrophage |
| Trem2 | 1.07 | 0.00 | 0.00 | 0.28 | 0.01 | 21 | Macrophage |
| Ms4a6d | 1.00 | 0.00 | 0.00 | 0.28 | 0.02 | 21 | Macrophage |
| Ly86 | 1.51 | 0.00 | 0.00 | 0.52 | 0.06 | 21 | Macrophage |
| Fcgr1 | 0.93 | 0.00 | 0.00 | 0.26 | 0.02 | 21 | Macrophage |
| Lgmn | 2.02 | 0.00 | 0.00 | 0.65 | 0.12 | 21 | Macrophage |
| Apoe | 3.29 | 0.00 | 0.00 | 0.87 | 0.28 | 21 | Macrophage |
| Ms4a6b | 1.25 | 0.00 | 0.00 | 0.47 | 0.06 | 21 | Macrophage |
| Msr1 | 0.92 | 0.00 | 0.00 | 0.26 | 0.02 | 21 | Macrophage |
| Cyth4 | 1.10 | 0.00 | 0.00 | 0.38 | 0.04 | 21 | Macrophage |
| Maf | 2.10 | 0.00 | 0.00 | 0.76 | 0.19 | 21 | Macrophage |
| Unc93b1 | 1.25 | 0.00 | 0.00 | 0.49 | 0.07 | 21 | Macrophage |
| Pla2g7 | 1.32 | 0.00 | 0.00 | 0.42 | 0.05 | 21 | Macrophage |
| Cybb | 1.05 | 0.00 | 0.00 | 0.56 | 0.09 | 21 | Macrophage |
| Clec4a3 | 0.99 | 0.00 | 0.00 | 0.35 | 0.04 | 21 | Macrophage |
| Lst1 | 1.17 | 0.00 | 0.00 | 0.45 | 0.06 | 21 | Macrophage |
| Ctsb | 1.83 | 0.00 | 0.00 | 0.83 | 0.27 | 21 | Macrophage |
| Cd68 | 1.00 | 0.00 | 0.00 | 0.29 | 0.03 | 21 | Macrophage |
| AF251705 | 0.89 | 0.00 | 0.00 | 0.28 | 0.03 | 21 | Macrophage |
| Cxcl16 | 1.19 | 0.00 | 0.00 | 0.32 | 0.04 | 21 | Macrophage |
| Ctsc | 1.47 | 0.00 | 0.00 | 0.71 | 0.19 | 21 | Macrophage |
| Cd300a | 0.85 | 0.00 | 0.00 | 0.31 | 0.04 | 21 | Macrophage |
| Mafb | 1.16 | 0.00 | 0.00 | 0.26 | 0.03 | 21 | Macrophage |
| Cst3 | 1.48 | 0.00 | 0.00 | 0.96 | 0.59 | 21 | Macrophage |
| Evi2a | 0.95 | 0.00 | 0.00 | 0.28 | 0.03 | 21 | Macrophage |
| Pld4 | 0.92 | 0.00 | 0.00 | 0.37 | 0.05 | 21 | Macrophage |
| Spi1 | 0.93 | 0.00 | 0.00 | 0.40 | 0.06 | 21 | Macrophage |
| Myo5a | 1.19 | 0.00 | 0.00 | 0.40 | 0.07 | 21 | Macrophage |
| Hexb | 1.20 | 0.00 | 0.00 | 0.42 | 0.07 | 21 | Macrophage |
| Rgs10 | 1.16 | 0.00 | 0.00 | 0.49 | 0.10 | 21 | Macrophage |
| Mpeg1 | 1.05 | 0.00 | 0.00 | 0.42 | 0.07 | 21 | Macrophage |
| Arrb2 | 1.11 | 0.00 | 0.00 | 0.39 | 0.06 | 21 | Macrophage |
| Bcl2a1b | 1.02 | 0.00 | 0.00 | 0.41 | 0.07 | 21 | Macrophage |
| Cd14 | 1.17 | 0.00 | 0.00 | 0.26 | 0.03 | 21 | Macrophage |
| Trf | 1.18 | 0.00 | 0.00 | 0.47 | 0.10 | 21 | Macrophage |
| Bcl2a1d | 1.02 | 0.00 | 0.00 | 0.42 | 0.08 | 21 | Macrophage |
| Tmsb4x | 0.95 | 0.00 | 0.00 | 1.00 | 0.89 | 21 | Macrophage |
| Sepp1 | 1.79 | 0.00 | 0.00 | 0.81 | 0.41 | 21 | Macrophage |
| Sirpa | 1.01 | 0.00 | 0.00 | 0.36 | 0.07 | 21 | Macrophage |
| Grn | 1.24 | 0.00 | 0.00 | 0.52 | 0.13 | 21 | Macrophage |
| Cfp | 1.14 | 0.00 | 0.00 | 0.30 | 0.05 | 21 | Macrophage |
| Ifi204 | 0.86 | 0.00 | 0.00 | 0.32 | 0.05 | 21 | Macrophage |
| Cfh | 0.93 | 0.00 | 0.00 | 0.30 | 0.05 | 21 | Macrophage |
| Cd84 | 0.91 | 0.00 | 0.00 | 0.31 | 0.05 | 21 | Macrophage |
| Ptpn18 | 1.07 | 0.00 | 0.00 | 0.53 | 0.14 | 21 | Macrophage |
| Bcl2a1a | 0.87 | 0.00 | 0.00 | 0.36 | 0.06 | 21 | Macrophage |
| Fcgrt | 1.18 | 0.00 | 0.00 | 0.42 | 0.09 | 21 | Macrophage |
| Igsf6 | 0.73 | 0.00 | 0.00 | 0.28 | 0.04 | 21 | Macrophage |
| Lyz2 | 0.42 | 0.00 | 0.00 | 0.77 | 0.26 | 21 | Macrophage |
| B2m | 0.99 | 0.00 | 0.00 | 0.95 | 0.67 | 21 | Macrophage |
| Fyb | 1.02 | 0.00 | 0.00 | 0.58 | 0.18 | 21 | Macrophage |
| Psap | 0.95 | 0.00 | 0.00 | 0.79 | 0.36 | 21 | Macrophage |
| Ptplad2 | 0.85 | 0.00 | 0.00 | 0.28 | 0.05 | 21 | Macrophage |
| Efhd2 | 0.91 | 0.00 | 0.00 | 0.38 | 0.09 | 21 | Macrophage |
| Snx5 | 1.06 | 0.00 | 0.00 | 0.47 | 0.13 | 21 | Macrophage |
| Cyba | 0.98 | 0.00 | 0.00 | 0.72 | 0.30 | 21 | Macrophage |
| Mnda | 0.84 | 0.00 | 0.00 | 0.31 | 0.06 | 21 | Macrophage |
| Slfn2 | 0.83 | 0.00 | 0.00 | 0.49 | 0.13 | 21 | Macrophage |
| Irf8 | 0.90 | 0.00 | 0.00 | 0.31 | 0.06 | 21 | Macrophage |
| KCTD12 | 0.90 | 0.00 | 0.00 | 0.34 | 0.07 | 21 | Macrophage |
| Gm13139 | 0.99 | 0.00 | 0.00 | 0.33 | 0.07 | 21 | Macrophage |
| Cotl1 | 0.95 | 0.00 | 0.00 | 0.54 | 0.17 | 21 | Macrophage |
| Gm10116 | 1.10 | 0.00 | 0.00 | 0.88 | 0.62 | 21 | Macrophage |
| Gm13248 | 0.84 | 0.00 | 0.00 | 0.29 | 0.06 | 21 | Macrophage |
| Serinc3 | 1.01 | 0.00 | 0.00 | 0.72 | 0.32 | 21 | Macrophage |
| Gm13251 | 0.87 | 0.00 | 0.00 | 0.27 | 0.05 | 21 | Macrophage |
| Fermt3 | 0.69 | 0.00 | 0.00 | 0.30 | 0.06 | 21 | Macrophage |
| Cd53 | 0.78 | 0.00 | 0.00 | 0.38 | 0.09 | 21 | Macrophage |
| Man2b1 | 0.87 | 0.00 | 0.00 | 0.31 | 0.07 | 21 | Macrophage |
| Lcp1 | 0.92 | 0.00 | 0.00 | 0.61 | 0.22 | 21 | Macrophage |
| Tm6sf1 | 0.82 | 0.00 | 0.00 | 0.27 | 0.05 | 21 | Macrophage |
| Scamp2 | 1.01 | 0.00 | 0.00 | 0.47 | 0.15 | 21 | Macrophage |
| Arhgap30 | 0.71 | 0.00 | 0.00 | 0.40 | 0.10 | 21 | Macrophage |
| Ap1b1 | 0.81 | 0.00 | 0.00 | 0.25 | 0.05 | 21 | Macrophage |
| Clta | 1.01 | 0.00 | 0.00 | 0.77 | 0.43 | 21 | Macrophage |
| Ctsz | 0.89 | 0.00 | 0.00 | 0.47 | 0.15 | 21 | Macrophage |
| Mndal | 0.80 | 0.00 | 0.00 | 0.51 | 0.16 | 21 | Macrophage |
| Ctsa | 0.91 | 0.00 | 0.00 | 0.43 | 0.13 | 21 | Macrophage |
| Ptpn6 | 0.62 | 0.00 | 0.00 | 0.34 | 0.08 | 21 | Macrophage |
| Ftl1 | 1.07 | 0.00 | 0.00 | 0.81 | 0.57 | 21 | Macrophage |
| Alox5ap | 0.57 | 0.00 | 0.00 | 0.37 | 0.10 | 21 | Macrophage |
| Actb | 0.49 | 0.00 | 0.00 | 1.00 | 0.98 | 21 | Macrophage |
| AI662270 | 0.73 | 0.00 | 0.00 | 0.30 | 0.07 | 21 | Macrophage |
| Arhgdib | 0.71 | 0.00 | 0.00 | 0.49 | 0.16 | 21 | Macrophage |
| Hexa | 0.86 | 0.00 | 0.00 | 0.31 | 0.08 | 21 | Macrophage |
| Sat1 | 0.83 | 0.00 | 0.00 | 0.69 | 0.33 | 21 | Macrophage |
| Hcls1 | 0.70 | 0.00 | 0.00 | 0.30 | 0.07 | 21 | Macrophage |
| Mef2c | 1.01 | 0.00 | 0.00 | 0.47 | 0.16 | 21 | Macrophage |
| Blvrb | 0.90 | 0.00 | 0.00 | 0.30 | 0.08 | 21 | Macrophage |
| Snx6 | 1.22 | 0.00 | 0.00 | 0.59 | 0.27 | 21 | Macrophage |
| Abca1 | 0.99 | 0.00 | 0.00 | 0.26 | 0.06 | 21 | Macrophage |
| Ptprc | 0.61 | 0.00 | 0.00 | 0.46 | 0.15 | 21 | Macrophage |
| Fth1 | 0.68 | 0.00 | 0.00 | 0.93 | 0.81 | 21 | Macrophage |
| Pip4k2a | 0.70 | 0.00 | 0.00 | 0.26 | 0.06 | 21 | Macrophage |
| Ncf1 | 0.57 | 0.00 | 0.00 | 0.28 | 0.07 | 21 | Macrophage |
| Lilrb4 | 0.64 | 0.00 | 0.00 | 0.26 | 0.06 | 21 | Macrophage |
| Gusb | 0.73 | 0.00 | 0.00 | 0.30 | 0.08 | 21 | Macrophage |
| Cenpe | 1.03 | 0.00 | 0.00 | 0.29 | 0.08 | 21 | Macrophage |
| Csf2ra | 0.63 | 0.00 | 0.00 | 0.26 | 0.06 | 21 | Macrophage |
| Plek | 0.61 | 0.00 | 0.00 | 0.28 | 0.07 | 21 | Macrophage |
| Birc5 | 0.74 | 0.00 | 0.00 | 0.26 | 0.06 | 21 | Macrophage |
| Ifi30 | 0.73 | 0.00 | 0.00 | 0.35 | 0.11 | 21 | Macrophage |
| Incenp | 0.73 | 0.00 | 0.00 | 0.29 | 0.08 | 21 | Macrophage |
| Arpc1b | 0.72 | 0.00 | 0.00 | 0.69 | 0.37 | 21 | Macrophage |
| Ifi27l2a | 0.83 | 0.00 | 0.00 | 0.32 | 0.09 | 21 | Macrophage |
| Fam105a | 0.62 | 0.00 | 0.00 | 0.25 | 0.06 | 21 | Macrophage |
| Cd52 | 0.42 | 0.00 | 0.00 | 0.55 | 0.20 | 21 | Macrophage |
| Snx2 | 0.93 | 0.00 | 0.00 | 0.43 | 0.17 | 21 | Macrophage |
| Cltc | 0.90 | 0.00 | 0.00 | 0.45 | 0.18 | 21 | Macrophage |
| Rps29 | 0.56 | 0.00 | 0.00 | 0.98 | 0.91 | 21 | Macrophage |
| Pycard | 0.65 | 0.00 | 0.00 | 0.30 | 0.09 | 21 | Macrophage |
| Runx1 | 0.72 | 0.00 | 0.00 | 0.26 | 0.07 | 21 | Macrophage |
| Apbb1ip | 0.59 | 0.00 | 0.00 | 0.32 | 0.09 | 21 | Macrophage |
| Mki67 | 0.98 | 0.00 | 0.00 | 0.35 | 0.12 | 21 | Macrophage |
| Tgfbr1 | 0.83 | 0.00 | 0.00 | 0.27 | 0.08 | 21 | Macrophage |
| Gm9843 | 0.44 | 0.00 | 0.00 | 0.99 | 0.93 | 21 | Macrophage |
| Tpd52 | 0.67 | 0.00 | 0.00 | 0.32 | 0.10 | 21 | Macrophage |
| Coro1a | 0.55 | 0.00 | 0.00 | 0.32 | 0.10 | 21 | Macrophage |
| Il6ra | 0.61 | 0.00 | 0.00 | 0.32 | 0.10 | 21 | Macrophage |
| Tnfaip8 | 0.63 | 0.00 | 0.00 | 0.32 | 0.10 | 21 | Macrophage |
| Rnf130 | 0.65 | 0.00 | 0.00 | 0.25 | 0.07 | 21 | Macrophage |
| Ctsd | 0.82 | 0.00 | 0.00 | 0.51 | 0.25 | 21 | Macrophage |
| H2-D1 | 0.60 | 0.00 | 0.00 | 0.79 | 0.49 | 21 | Macrophage |
| Hist1h2ai | 0.91 | 0.00 | 0.00 | 0.30 | 0.10 | 21 | Macrophage |
| Top2a | 1.03 | 0.00 | 0.00 | 0.32 | 0.11 | 21 | Macrophage |
| Gpx1 | 0.62 | 0.00 | 0.00 | 0.70 | 0.43 | 21 | Macrophage |
| Gdi2 | 0.79 | 0.00 | 0.00 | 0.59 | 0.33 | 21 | Macrophage |
| Itm2b | 0.72 | 0.00 | 0.00 | 0.89 | 0.76 | 21 | Macrophage |
| Rnasel | 0.74 | 0.00 | 0.00 | 0.30 | 0.10 | 21 | Macrophage |
| H2-K1 | 0.68 | 0.00 | 0.00 | 0.62 | 0.32 | 21 | Macrophage |
| Erp29 | 0.70 | 0.00 | 0.00 | 0.52 | 0.26 | 21 | Macrophage |
| Igf1 | 0.79 | 0.00 | 0.00 | 0.36 | 0.14 | 21 | Macrophage |
| Capza2 | 0.78 | 0.00 | 0.00 | 0.60 | 0.35 | 21 | Macrophage |
| Atp6v0c | 0.77 | 0.00 | 0.00 | 0.50 | 0.25 | 21 | Macrophage |
| Sh3bgrl3 | 0.62 | 0.00 | 0.00 | 0.62 | 0.34 | 21 | Macrophage |
| Asah1 | 0.69 | 0.00 | 0.00 | 0.33 | 0.12 | 21 | Macrophage |
| Ifnar2 | 0.69 | 0.00 | 0.00 | 0.34 | 0.13 | 21 | Macrophage |
| Creg1 | 0.67 | 0.00 | 0.00 | 0.40 | 0.17 | 21 | Macrophage |
| Zfp36l2 | 0.57 | 0.00 | 0.00 | 0.31 | 0.11 | 21 | Macrophage |
| Mt1 | 0.88 | 0.00 | 0.00 | 0.38 | 0.16 | 21 | Macrophage |
| 9930111J21Rik2 | 0.68 | 0.00 | 0.00 | 0.32 | 0.12 | 21 | Macrophage |
| Fam49b | 0.60 | 0.00 | 0.00 | 0.30 | 0.11 | 21 | Macrophage |
| Snx30 | 0.62 | 0.00 | 0.00 | 0.29 | 0.10 | 21 | Macrophage |
| Ifi203 | 0.55 | 0.00 | 0.00 | 0.36 | 0.14 | 21 | Macrophage |
| Hist1h2af | 0.83 | 0.00 | 0.00 | 0.27 | 0.09 | 21 | Macrophage |
| Lrrc58 | 0.37 | 0.00 | 0.00 | 0.99 | 0.91 | 21 | Macrophage |
| Hist1h2ap | 1.09 | 0.00 | 0.00 | 0.35 | 0.15 | 21 | Macrophage |
| Ucp2 | 0.65 | 0.00 | 0.00 | 0.31 | 0.12 | 21 | Macrophage |
| Hist1h2ag | 0.97 | 0.00 | 0.00 | 0.30 | 0.12 | 21 | Macrophage |
| Arpc2 | 0.54 | 0.00 | 0.00 | 0.78 | 0.61 | 21 | Macrophage |
| Gm26445 | 0.75 | 0.00 | 0.00 | 0.36 | 0.15 | 21 | Macrophage |
| Fam111a | 0.67 | 0.00 | 0.00 | 0.36 | 0.15 | 21 | Macrophage |
| Hist1h2ao | 1.13 | 0.00 | 0.00 | 0.35 | 0.15 | 21 | Macrophage |
| Rnase4 | 0.84 | 0.00 | 0.00 | 0.40 | 0.18 | 21 | Macrophage |
| Hist1h2ac | 0.89 | 0.00 | 0.00 | 0.28 | 0.11 | 21 | Macrophage |
| Fau | 0.38 | 0.00 | 0.00 | 0.95 | 0.87 | 21 | Macrophage |
| D17H6S56E-5 | 0.58 | 0.00 | 0.00 | 0.29 | 0.11 | 21 | Macrophage |
| Cd74 | 1.06 | 0.00 | 0.00 | 0.32 | 0.13 | 21 | Macrophage |
| Hist1h2ab | 0.88 | 0.00 | 0.00 | 0.25 | 0.09 | 21 | Macrophage |
| Hist1h2ah | 0.79 | 0.00 | 0.00 | 0.26 | 0.10 | 21 | Macrophage |
| Arpc4 | 0.61 | 0.00 | 0.00 | 0.39 | 0.18 | 21 | Macrophage |
| Hist1h2ad | 0.89 | 0.00 | 0.00 | 0.28 | 0.11 | 21 | Macrophage |
| Rac2 | 0.38 | 0.00 | 0.00 | 0.30 | 0.11 | 21 | Macrophage |
| Hist1h2an | 0.79 | 0.00 | 0.00 | 0.25 | 0.09 | 21 | Macrophage |
| Lyn | 0.55 | 0.00 | 0.00 | 0.32 | 0.13 | 21 | Macrophage |
| Gm2a | 0.55 | 0.00 | 0.00 | 0.25 | 0.09 | 21 | Macrophage |
| Marcks | 0.55 | 0.00 | 0.00 | 0.75 | 0.50 | 21 | Macrophage |
| Pfn1 | 0.59 | 0.00 | 0.00 | 0.67 | 0.46 | 21 | Macrophage |
| Ehd4 | 0.43 | 0.00 | 0.00 | 0.55 | 0.28 | 21 | Macrophage |
| Ptgs1 | 0.65 | 0.00 | 0.00 | 0.27 | 0.10 | 21 | Macrophage |
| Dek | 0.65 | 0.00 | 0.00 | 0.68 | 0.50 | 21 | Macrophage |
| Rbpj | 0.56 | 0.00 | 0.00 | 0.27 | 0.10 | 21 | Macrophage |
| Psmb8 | 0.49 | 0.00 | 0.00 | 0.40 | 0.18 | 21 | Macrophage |
| Fam46a | 0.62 | 0.00 | 0.00 | 0.26 | 0.10 | 21 | Macrophage |
| Tpm3 | 0.45 | 0.00 | 0.00 | 0.84 | 0.70 | 21 | Macrophage |
| Mir692-2a | 0.80 | 0.00 | 0.00 | 0.39 | 0.19 | 21 | Macrophage |
| Gmfg | 0.44 | 0.00 | 0.00 | 0.33 | 0.14 | 21 | Macrophage |
| Gbp7 | 0.60 | 0.00 | 0.00 | 0.33 | 0.14 | 21 | Macrophage |
| Tpm3-rs7 | 0.46 | 0.00 | 0.00 | 0.83 | 0.69 | 21 | Macrophage |
| Gm22774 | 0.73 | 0.00 | 0.00 | 0.36 | 0.17 | 21 | Macrophage |
| Prkcd | 0.48 | 0.00 | 0.00 | 0.28 | 0.11 | 21 | Macrophage |
| Actr3 | 0.57 | 0.00 | 0.00 | 0.55 | 0.33 | 21 | Macrophage |
| Lamp1 | 0.58 | 0.00 | 0.00 | 0.63 | 0.42 | 21 | Macrophage |
| Ap2a2 | 0.66 | 0.00 | 0.00 | 0.29 | 0.12 | 21 | Macrophage |
| Gns | 0.65 | 0.00 | 0.00 | 0.30 | 0.13 | 21 | Macrophage |
| Gng2 | 0.51 | 0.00 | 0.00 | 0.29 | 0.12 | 21 | Macrophage |
| Ptbp3 | 0.49 | 0.00 | 0.00 | 0.59 | 0.36 | 21 | Macrophage |
| Gm22751 | 0.74 | 0.00 | 0.00 | 0.33 | 0.16 | 21 | Macrophage |
| Plin2 | 0.47 | 0.00 | 0.00 | 0.31 | 0.13 | 21 | Macrophage |
| Ifngr1 | 0.51 | 0.00 | 0.00 | 0.44 | 0.24 | 21 | Macrophage |
| Ap2s1 | 0.53 | 0.00 | 0.00 | 0.35 | 0.18 | 21 | Macrophage |
| Clic1 | 0.50 | 0.00 | 0.00 | 0.58 | 0.38 | 21 | Macrophage |
| Zeb2 | 0.49 | 0.00 | 0.00 | 0.53 | 0.31 | 21 | Macrophage |
| Fos | 0.65 | 0.00 | 0.00 | 0.37 | 0.19 | 21 | Macrophage |
| Bst2 | 0.49 | 0.00 | 0.00 | 0.30 | 0.14 | 21 | Macrophage |
| Coro1b | 0.58 | 0.00 | 0.00 | 0.43 | 0.24 | 21 | Macrophage |
| Rgs2 | 0.34 | 0.00 | 0.00 | 0.50 | 0.28 | 21 | Macrophage |
| Sft2d1 | 0.54 | 0.00 | 0.00 | 0.28 | 0.13 | 21 | Macrophage |
| H2afz | 0.58 | 0.00 | 0.00 | 0.57 | 0.40 | 21 | Macrophage |
| Serp1 | 0.48 | 0.00 | 0.00 | 0.41 | 0.23 | 21 | Macrophage |
| Rplp1 | 0.27 | 0.00 | 0.00 | 0.94 | 0.90 | 21 | Macrophage |
| Gm12166 | 0.52 | 0.00 | 0.00 | 0.28 | 0.13 | 21 | Macrophage |
| Actr2 | 0.48 | 0.00 | 0.00 | 0.46 | 0.27 | 21 | Macrophage |
| Nmt1 | 0.51 | 0.00 | 0.00 | 0.41 | 0.23 | 21 | Macrophage |
| Arpc3 | 0.48 | 0.00 | 0.00 | 0.44 | 0.26 | 21 | Macrophage |
| Smc2 | 0.46 | 0.00 | 0.00 | 0.28 | 0.14 | 21 | Macrophage |
| Gm10126 | 0.46 | 0.00 | 0.00 | 0.58 | 0.40 | 21 | Macrophage |
| Dusp6 | 0.58 | 0.00 | 0.00 | 0.34 | 0.18 | 21 | Macrophage |
| Zfp36 | 0.64 | 0.00 | 0.00 | 0.32 | 0.17 | 21 | Macrophage |
| Atp6v0b | 0.55 | 0.00 | 0.00 | 0.35 | 0.19 | 21 | Macrophage |
| Arpc5 | 0.41 | 0.00 | 0.00 | 0.71 | 0.56 | 21 | Macrophage |
| Emp3 | 0.45 | 0.00 | 0.00 | 0.38 | 0.21 | 21 | Macrophage |
| Gm4070 | 0.31 | 0.00 | 0.00 | 0.26 | 0.12 | 21 | Macrophage |
| Cd302 | 0.42 | 0.00 | 0.00 | 0.30 | 0.15 | 21 | Macrophage |
| Hmgb2 | 0.75 | 0.00 | 0.00 | 0.55 | 0.43 | 21 | Macrophage |
| Gas6 | 0.66 | 0.00 | 0.00 | 0.28 | 0.15 | 21 | Macrophage |
| Gm10073 | 0.26 | 0.00 | 0.00 | 0.92 | 0.86 | 21 | Macrophage |
| Tmem176b | 0.60 | 0.00 | 0.00 | 0.49 | 0.34 | 21 | Macrophage |
| M6pr | 0.46 | 0.00 | 0.00 | 0.39 | 0.23 | 21 | Macrophage |
| Ccdc50 | 0.58 | 0.00 | 0.00 | 0.42 | 0.26 | 21 | Macrophage |
| Atp2b1 | 0.43 | 0.00 | 0.00 | 0.73 | 0.59 | 21 | Macrophage |
| Cfl1 | 0.37 | 0.00 | 0.00 | 0.71 | 0.58 | 21 | Macrophage |
| Atox1 | 0.46 | 0.00 | 0.00 | 0.53 | 0.37 | 21 | Macrophage |
| Ppt1 | 0.38 | 0.00 | 0.00 | 0.26 | 0.12 | 21 | Macrophage |
| Sdcbp | 0.46 | 0.00 | 0.00 | 0.44 | 0.28 | 21 | Macrophage |
| Cyfip1 | 0.51 | 0.00 | 0.00 | 0.30 | 0.16 | 21 | Macrophage |
| Cd63 | 0.62 | 0.00 | 0.00 | 0.44 | 0.30 | 21 | Macrophage |
| Nucks1 | 0.47 | 0.00 | 0.00 | 0.44 | 0.28 | 21 | Macrophage |
| Tor1aip1 | 0.47 | 0.00 | 0.00 | 0.33 | 0.19 | 21 | Macrophage |
| Arl6ip1 | 0.38 | 0.00 | 0.00 | 0.68 | 0.52 | 21 | Macrophage |
| Rin2 | 0.46 | 0.00 | 0.00 | 0.29 | 0.16 | 21 | Macrophage |
| Tab2 | 0.48 | 0.00 | 0.00 | 0.38 | 0.23 | 21 | Macrophage |
| Ivns1abp | 0.40 | 0.00 | 0.00 | 0.38 | 0.22 | 21 | Macrophage |
| Sec11c | 0.34 | 0.00 | 0.00 | 0.32 | 0.18 | 21 | Macrophage |
| Ier3 | 0.83 | 0.00 | 0.00 | 0.25 | 0.14 | 21 | Macrophage |
| Akap13 | 0.38 | 0.00 | 0.00 | 0.42 | 0.27 | 21 | Macrophage |
| Atp6ap2 | 0.48 | 0.00 | 0.00 | 0.28 | 0.15 | 21 | Macrophage |
| Ccdc34 | 0.36 | 0.00 | 0.00 | 0.26 | 0.14 | 21 | Macrophage |
| BC005537 | 0.41 | 0.00 | 0.00 | 0.48 | 0.33 | 21 | Macrophage |
| Hprt | 0.34 | 0.00 | 0.00 | 0.26 | 0.14 | 21 | Macrophage |
| Egr1 | 0.46 | 0.00 | 0.00 | 0.26 | 0.14 | 21 | Macrophage |
| Celf2 | 0.33 | 0.00 | 0.00 | 0.44 | 0.28 | 21 | Macrophage |
| Smchd1 | 0.38 | 0.00 | 0.00 | 0.29 | 0.16 | 21 | Macrophage |
| Grb2 | 0.33 | 0.00 | 0.00 | 0.31 | 0.18 | 21 | Macrophage |
| Tmed5 | 0.33 | 0.00 | 0.00 | 0.30 | 0.17 | 21 | Macrophage |
| Mycbp2 | 0.38 | 0.00 | 0.00 | 0.38 | 0.23 | 21 | Macrophage |
| H2afy | 0.38 | 0.00 | 0.00 | 0.33 | 0.20 | 21 | Macrophage |
| Cstb | 0.42 | 0.00 | 0.00 | 0.41 | 0.28 | 21 | Macrophage |
| Snx3 | 0.38 | 0.00 | 0.00 | 0.47 | 0.33 | 21 | Macrophage |
| Junb | 0.52 | 0.00 | 0.00 | 0.32 | 0.20 | 21 | Macrophage |
| Akr1a1 | 0.34 | 0.00 | 0.00 | 0.64 | 0.50 | 21 | Macrophage |
| Fkbp2 | 0.35 | 0.00 | 0.00 | 0.31 | 0.18 | 21 | Macrophage |
| Arhgap17 | 0.43 | 0.00 | 0.00 | 0.30 | 0.18 | 21 | Macrophage |
| Stmn1 | 0.34 | 0.00 | 0.00 | 0.37 | 0.24 | 21 | Macrophage |
| Nars | 0.43 | 0.00 | 0.00 | 0.46 | 0.33 | 21 | Macrophage |
| Irf2bp2 | 0.44 | 0.00 | 0.00 | 0.27 | 0.16 | 21 | Macrophage |
| Rrbp1 | 0.29 | 0.00 | 0.00 | 0.68 | 0.55 | 21 | Macrophage |
| Ctage5 | 0.35 | 0.00 | 0.00 | 0.42 | 0.28 | 21 | Macrophage |
| Limd2 | 0.32 | 0.00 | 0.00 | 0.27 | 0.15 | 21 | Macrophage |
| Itgam | 0.34 | 0.00 | 0.00 | 0.41 | 0.28 | 21 | Macrophage |
| Gnai2 | 0.29 | 0.00 | 0.00 | 0.54 | 0.39 | 21 | Macrophage |
| Calr | 0.30 | 0.00 | 0.00 | 0.57 | 0.43 | 21 | Macrophage |
| Tmem256 | 0.34 | 0.00 | 0.00 | 0.46 | 0.33 | 21 | Macrophage |
| Gabarap | 0.37 | 0.00 | 0.00 | 0.56 | 0.44 | 21 | Macrophage |
| Taldo1 | 0.27 | 0.00 | 0.00 | 0.39 | 0.26 | 21 | Macrophage |
| Cdc42 | 0.25 | 0.00 | 0.00 | 0.68 | 0.54 | 21 | Macrophage |
| Capzb | 0.33 | 0.00 | 0.00 | 0.54 | 0.42 | 21 | Macrophage |
| Dbi | 0.35 | 0.00 | 0.00 | 0.50 | 0.37 | 21 | Macrophage |
| Klf6 | 0.29 | 0.00 | 0.00 | 0.49 | 0.36 | 21 | Macrophage |
| Pomp | 0.33 | 0.00 | 0.00 | 0.43 | 0.32 | 21 | Macrophage |
| Ctsl | 0.43 | 0.00 | 0.00 | 0.49 | 0.39 | 21 | Macrophage |
| Atp6v1f | 0.37 | 0.00 | 0.00 | 0.36 | 0.25 | 21 | Macrophage |
| Cox5a | 0.29 | 0.00 | 0.00 | 0.44 | 0.31 | 21 | Macrophage |
| Reep5 | 0.33 | 0.00 | 0.00 | 0.35 | 0.24 | 21 | Macrophage |
| Taok3 | 0.27 | 0.00 | 0.00 | 0.29 | 0.19 | 21 | Macrophage |
| 2810428I15Rik | 0.29 | 0.00 | 0.00 | 0.30 | 0.20 | 21 | Macrophage |
| Anp32b | 0.41 | 0.00 | 0.00 | 0.57 | 0.50 | 21 | Macrophage |
| Hspa8 | 0.26 | 0.00 | 0.00 | 0.76 | 0.67 | 21 | Macrophage |
| Igfbp4 | 0.31 | 0.00 | 0.00 | 0.37 | 0.25 | 21 | Macrophage |
| Ostf1 | 0.26 | 0.00 | 0.00 | 0.38 | 0.26 | 21 | Macrophage |
| Magt1 | 0.29 | 0.00 | 0.00 | 0.30 | 0.20 | 21 | Macrophage |
| Ik | 0.29 | 0.00 | 0.00 | 0.47 | 0.36 | 21 | Macrophage |
| Lamp2 | 0.36 | 0.00 | 0.00 | 0.37 | 0.26 | 21 | Macrophage |
| Brk1 | 0.29 | 0.00 | 0.00 | 0.37 | 0.26 | 21 | Macrophage |
| Psme2b | 0.27 | 0.00 | 0.00 | 0.41 | 0.30 | 21 | Macrophage |
| Tsc22d4 | 0.26 | 0.00 | 0.01 | 0.39 | 0.28 | 21 | Macrophage |
| Rnaset2b | 0.39 | 0.00 | 0.02 | 0.35 | 0.26 | 21 | Macrophage |
| Rtn4 | 0.30 | 0.00 | 0.02 | 0.33 | 0.24 | 21 | Macrophage |
| Ran | 0.28 | 0.00 | 0.02 | 0.48 | 0.38 | 21 | Macrophage |
| Rnaset2a | 0.39 | 0.00 | 0.02 | 0.35 | 0.26 | 21 | Macrophage |
| Gm10282 | 0.28 | 0.00 | 0.02 | 0.50 | 0.40 | 21 | Macrophage |
| Pak2 | 0.28 | 0.00 | 0.02 | 0.40 | 0.29 | 21 | Macrophage |
| Gm26924 | 0.35 | 0.00 | 0.03 | 0.65 | 0.57 | 21 | Macrophage |
| Etfb | 0.28 | 0.00 | 0.03 | 0.39 | 0.28 | 21 | Macrophage |
| S100a9 | 5.52 | 0.00 | 0.00 | 0.94 | 0.06 | 22 | Neutrophil |
| Stfa1 | 5.43 | 0.00 | 0.00 | 0.93 | 0.05 | 22 | Neutrophil |
| BC100530 | 5.43 | 0.00 | 0.00 | 0.93 | 0.04 | 22 | Neutrophil |
| S100a8 | 5.09 | 0.00 | 0.00 | 0.99 | 0.10 | 22 | Neutrophil |
| BC117090 | 4.86 | 0.00 | 0.00 | 0.89 | 0.02 | 22 | Neutrophil |
| Gm5483 | 4.83 | 0.00 | 0.00 | 0.97 | 0.03 | 22 | Neutrophil |
| Stfa2l1 | 4.83 | 0.00 | 0.00 | 0.92 | 0.02 | 22 | Neutrophil |
| Stfa2 | 4.73 | 0.00 | 0.00 | 0.88 | 0.02 | 22 | Neutrophil |
| 2010005H15Rik | 4.60 | 0.00 | 0.00 | 0.95 | 0.02 | 22 | Neutrophil |
| Retnlg | 3.89 | 0.00 | 0.00 | 0.60 | 0.01 | 22 | Neutrophil |
| Stfa3 | 3.66 | 0.00 | 0.00 | 0.66 | 0.02 | 22 | Neutrophil |
| 1100001G20Rik | 3.43 | 0.00 | 0.00 | 0.79 | 0.02 | 22 | Neutrophil |
| Csta | 3.34 | 0.00 | 0.00 | 0.65 | 0.00 | 22 | Neutrophil |
| Cxcr2 | 2.74 | 0.00 | 0.00 | 0.75 | 0.01 | 22 | Neutrophil |
| Gm5416 | 2.59 | 0.00 | 0.00 | 0.73 | 0.00 | 22 | Neutrophil |
| Il1b | 2.41 | 0.00 | 0.00 | 0.49 | 0.02 | 22 | Neutrophil |
| Mmp8 | 2.23 | 0.00 | 0.00 | 0.44 | 0.01 | 22 | Neutrophil |
| Gm4758 | 2.15 | 0.00 | 0.00 | 0.49 | 0.00 | 22 | Neutrophil |
| Slpi | 2.07 | 0.00 | 0.00 | 0.51 | 0.02 | 22 | Neutrophil |
| Tyrobp | 1.99 | 0.00 | 0.00 | 0.87 | 0.12 | 22 | Neutrophil |
| Alox5ap | 1.89 | 0.00 | 0.00 | 0.74 | 0.09 | 22 | Neutrophil |
| Pygl | 1.87 | 0.00 | 0.00 | 0.73 | 0.10 | 22 | Neutrophil |
| Msrb1 | 1.87 | 0.00 | 0.00 | 0.78 | 0.14 | 22 | Neutrophil |
| Csf3r | 1.83 | 0.00 | 0.00 | 0.48 | 0.01 | 22 | Neutrophil |
| Asprv1 | 1.79 | 0.00 | 0.00 | 0.40 | 0.00 | 22 | Neutrophil |
| Gda | 1.78 | 0.00 | 0.00 | 0.57 | 0.05 | 22 | Neutrophil |
| Pglyrp1 | 1.75 | 0.00 | 0.00 | 0.49 | 0.02 | 22 | Neutrophil |
| Mmp9 | 1.66 | 0.00 | 0.00 | 0.42 | 0.00 | 22 | Neutrophil |
| Cd300lf | 1.64 | 0.00 | 0.00 | 0.53 | 0.03 | 22 | Neutrophil |
| Gm15845 | 1.64 | 0.00 | 0.00 | 0.30 | 0.00 | 22 | Neutrophil |
| Il1r2 | 1.63 | 0.00 | 0.00 | 0.33 | 0.01 | 22 | Neutrophil |
| Clec4d | 1.56 | 0.00 | 0.00 | 0.38 | 0.01 | 22 | Neutrophil |
| Ccr1 | 1.47 | 0.00 | 0.00 | 0.43 | 0.03 | 22 | Neutrophil |
| C5ar1 | 1.40 | 0.00 | 0.00 | 0.41 | 0.02 | 22 | Neutrophil |
| F630028O10Rik | 1.40 | 0.00 | 0.00 | 0.29 | 0.01 | 22 | Neutrophil |
| Slfn1 | 1.39 | 0.00 | 0.00 | 0.36 | 0.02 | 22 | Neutrophil |
| Slfn4 | 1.39 | 0.00 | 0.00 | 0.28 | 0.01 | 22 | Neutrophil |
| AA467197 | 1.37 | 0.00 | 0.00 | 0.31 | 0.01 | 22 | Neutrophil |
| 1810033B17Rik | 1.34 | 0.00 | 0.00 | 0.42 | 0.03 | 22 | Neutrophil |
| Fpr2 | 1.14 | 0.00 | 0.00 | 0.33 | 0.02 | 22 | Neutrophil |
| Cd300ld | 1.11 | 0.00 | 0.00 | 0.29 | 0.02 | 22 | Neutrophil |
| Fcer1g | 1.39 | 0.00 | 0.00 | 0.73 | 0.11 | 22 | Neutrophil |
| Gp49a | 1.39 | 0.00 | 0.00 | 0.46 | 0.05 | 22 | Neutrophil |
| Ncf1 | 1.45 | 0.00 | 0.00 | 0.53 | 0.07 | 22 | Neutrophil |
| Spi1 | 1.39 | 0.00 | 0.00 | 0.52 | 0.06 | 22 | Neutrophil |
| Hp | 1.50 | 0.00 | 0.00 | 0.67 | 0.11 | 22 | Neutrophil |
| Lilrb4 | 1.40 | 0.00 | 0.00 | 0.49 | 0.06 | 22 | Neutrophil |
| Pilra | 1.13 | 0.00 | 0.00 | 0.33 | 0.03 | 22 | Neutrophil |
| Gsr | 1.57 | 0.00 | 0.00 | 0.57 | 0.09 | 22 | Neutrophil |
| Pla2g7 | 1.51 | 0.00 | 0.00 | 0.46 | 0.05 | 22 | Neutrophil |
| Igsf6 | 1.29 | 0.00 | 0.00 | 0.40 | 0.04 | 22 | Neutrophil |
| Lrg1 | 1.14 | 0.00 | 0.00 | 0.28 | 0.02 | 22 | Neutrophil |
| Slfn2 | 1.57 | 0.00 | 0.00 | 0.68 | 0.13 | 22 | Neutrophil |
| Pirb | 1.18 | 0.00 | 0.00 | 0.36 | 0.04 | 22 | Neutrophil |
| Clec5a | 1.12 | 0.00 | 0.00 | 0.25 | 0.02 | 22 | Neutrophil |
| Ifitm6 | 1.63 | 0.00 | 0.00 | 0.32 | 0.03 | 22 | Neutrophil |
| Mxd1 | 1.31 | 0.00 | 0.00 | 0.36 | 0.04 | 22 | Neutrophil |
| Tnfaip2 | 1.55 | 0.00 | 0.00 | 0.39 | 0.05 | 22 | Neutrophil |
| Lst1 | 1.26 | 0.00 | 0.00 | 0.46 | 0.06 | 22 | Neutrophil |
| Fcgr3 | 1.08 | 0.00 | 0.00 | 0.37 | 0.04 | 22 | Neutrophil |
| Lcn2 | 2.12 | 0.00 | 0.00 | 0.45 | 0.07 | 22 | Neutrophil |
| Cd52 | 1.31 | 0.00 | 0.00 | 0.79 | 0.20 | 22 | Neutrophil |
| A430104N18Rik | 1.17 | 0.00 | 0.00 | 0.41 | 0.06 | 22 | Neutrophil |
| Ncf2 | 0.99 | 0.00 | 0.00 | 0.33 | 0.04 | 22 | Neutrophil |
| Grina | 1.37 | 0.00 | 0.00 | 0.42 | 0.07 | 22 | Neutrophil |
| Arrb2 | 1.07 | 0.00 | 0.00 | 0.40 | 0.06 | 22 | Neutrophil |
| Wfdc17 | 1.54 | 0.00 | 0.00 | 0.30 | 0.04 | 22 | Neutrophil |
| Ccl6 | 1.37 | 0.00 | 0.00 | 0.49 | 0.09 | 22 | Neutrophil |
| Lcp1 | 1.35 | 0.00 | 0.00 | 0.73 | 0.22 | 22 | Neutrophil |
| Ncf4 | 0.93 | 0.00 | 0.00 | 0.28 | 0.03 | 22 | Neutrophil |
| Apobr | 0.95 | 0.00 | 0.00 | 0.25 | 0.03 | 22 | Neutrophil |
| Cd84 | 1.16 | 0.00 | 0.00 | 0.33 | 0.05 | 22 | Neutrophil |
| Sirpb1c | 1.03 | 0.00 | 0.00 | 0.27 | 0.03 | 22 | Neutrophil |
| Cd14 | 1.16 | 0.00 | 0.00 | 0.25 | 0.03 | 22 | Neutrophil |
| Srgn | 1.41 | 0.00 | 0.00 | 0.81 | 0.32 | 22 | Neutrophil |
| Coro1a | 1.04 | 0.00 | 0.00 | 0.46 | 0.10 | 22 | Neutrophil |
| Ptprc | 1.04 | 0.00 | 0.00 | 0.58 | 0.14 | 22 | Neutrophil |
| Rac2 | 1.07 | 0.00 | 0.00 | 0.48 | 0.11 | 22 | Neutrophil |
| Anxa1 | 1.62 | 0.00 | 0.00 | 0.60 | 0.18 | 22 | Neutrophil |
| Itgb2 | 0.90 | 0.00 | 0.00 | 0.29 | 0.04 | 22 | Neutrophil |
| Prdx5 | 1.22 | 0.00 | 0.00 | 0.76 | 0.32 | 22 | Neutrophil |
| S100a11 | 1.32 | 0.00 | 0.00 | 0.82 | 0.48 | 22 | Neutrophil |
| S100a6 | 0.90 | 0.00 | 0.00 | 0.77 | 0.31 | 22 | Neutrophil |
| Cyba | 0.99 | 0.00 | 0.00 | 0.73 | 0.30 | 22 | Neutrophil |
| Fxyd5 | 1.16 | 0.00 | 0.00 | 0.54 | 0.18 | 22 | Neutrophil |
| Fos | 1.10 | 0.00 | 0.00 | 0.57 | 0.19 | 22 | Neutrophil |
| Arhgdib | 1.03 | 0.00 | 0.00 | 0.51 | 0.16 | 22 | Neutrophil |
| Junb | 1.29 | 0.00 | 0.00 | 0.55 | 0.19 | 22 | Neutrophil |
| Csf2ra | 0.88 | 0.00 | 0.00 | 0.30 | 0.06 | 22 | Neutrophil |
| Apbb1ip | 0.89 | 0.00 | 0.00 | 0.38 | 0.09 | 22 | Neutrophil |
| Samsn1 | 0.86 | 0.00 | 0.00 | 0.31 | 0.07 | 22 | Neutrophil |
| Tpd52 | 1.05 | 0.00 | 0.00 | 0.39 | 0.10 | 22 | Neutrophil |
| Plek | 0.98 | 0.00 | 0.00 | 0.31 | 0.07 | 22 | Neutrophil |
| Ptpn6 | 0.91 | 0.00 | 0.00 | 0.34 | 0.08 | 22 | Neutrophil |
| Gmfg | 0.91 | 0.00 | 0.00 | 0.45 | 0.14 | 22 | Neutrophil |
| 4632428N05Rik | 0.87 | 0.00 | 0.00 | 0.28 | 0.06 | 22 | Neutrophil |
| Taldo1 | 0.92 | 0.00 | 0.00 | 0.62 | 0.26 | 22 | Neutrophil |
| Cdk2ap2 | 1.13 | 0.00 | 0.00 | 0.46 | 0.15 | 22 | Neutrophil |
| Btg2 | 1.04 | 0.00 | 0.00 | 0.57 | 0.23 | 22 | Neutrophil |
| Lyz2 | 0.36 | 0.00 | 0.00 | 0.68 | 0.26 | 22 | Neutrophil |
| Cd44 | 1.00 | 0.00 | 0.00 | 0.52 | 0.20 | 22 | Neutrophil |
| Itgam | 0.97 | 0.00 | 0.00 | 0.62 | 0.28 | 22 | Neutrophil |
| Stk17b | 0.80 | 0.00 | 0.00 | 0.28 | 0.06 | 22 | Neutrophil |
| Tmsb4x | 0.62 | 0.00 | 0.00 | 0.98 | 0.89 | 22 | Neutrophil |
| Fyb | 0.88 | 0.00 | 0.00 | 0.50 | 0.18 | 22 | Neutrophil |
| Laptm5 | 0.67 | 0.00 | 0.00 | 0.45 | 0.14 | 22 | Neutrophil |
| Zfp36 | 0.95 | 0.00 | 0.00 | 0.46 | 0.16 | 22 | Neutrophil |
| Hmgb2 | 0.75 | 0.00 | 0.00 | 0.77 | 0.42 | 22 | Neutrophil |
| Arpc5 | 0.79 | 0.00 | 0.00 | 0.84 | 0.56 | 22 | Neutrophil |
| Hcls1 | 0.75 | 0.00 | 0.00 | 0.29 | 0.07 | 22 | Neutrophil |
| Cd53 | 0.69 | 0.00 | 0.00 | 0.33 | 0.09 | 22 | Neutrophil |
| Arid3a | 0.75 | 0.00 | 0.00 | 0.34 | 0.10 | 22 | Neutrophil |
| Thbs1 | 0.92 | 0.00 | 0.00 | 0.39 | 0.13 | 22 | Neutrophil |
| Samhd1 | 0.96 | 0.00 | 0.00 | 0.48 | 0.19 | 22 | Neutrophil |
| Actb | 0.50 | 0.00 | 0.00 | 1.00 | 0.98 | 22 | Neutrophil |
| Arhgap30 | 0.76 | 0.00 | 0.00 | 0.33 | 0.10 | 22 | Neutrophil |
| Pgd | 0.76 | 0.00 | 0.00 | 0.34 | 0.11 | 22 | Neutrophil |
| Sat1 | 0.88 | 0.00 | 0.00 | 0.63 | 0.33 | 22 | Neutrophil |
| Hist2h2aa2 | 0.88 | 0.00 | 0.00 | 0.38 | 0.14 | 22 | Neutrophil |
| Btg1 | 1.00 | 0.00 | 0.00 | 0.57 | 0.28 | 22 | Neutrophil |
| Dhrs7 | 0.89 | 0.00 | 0.00 | 0.41 | 0.16 | 22 | Neutrophil |
| Mcl1 | 0.92 | 0.00 | 0.00 | 0.50 | 0.23 | 22 | Neutrophil |
| Adipor1 | 0.91 | 0.00 | 0.00 | 0.51 | 0.24 | 22 | Neutrophil |
| Rgs2 | 0.77 | 0.00 | 0.00 | 0.59 | 0.28 | 22 | Neutrophil |
| Lgals3 | 0.56 | 0.00 | 0.00 | 0.41 | 0.15 | 22 | Neutrophil |
| Cd24a | 0.76 | 0.00 | 0.00 | 0.37 | 0.13 | 22 | Neutrophil |
| G0s2 | 0.97 | 0.00 | 0.00 | 0.37 | 0.13 | 22 | Neutrophil |
| Cotl1 | 0.81 | 0.00 | 0.00 | 0.43 | 0.17 | 22 | Neutrophil |
| Nudt4 | 0.84 | 0.00 | 0.00 | 0.42 | 0.17 | 22 | Neutrophil |
| Dusp1 | 1.09 | 0.00 | 0.00 | 0.40 | 0.17 | 22 | Neutrophil |
| Bcl2a1d | 1.37 | 0.00 | 0.00 | 0.27 | 0.08 | 22 | Neutrophil |
| Prr13 | 0.76 | 0.00 | 0.00 | 0.35 | 0.13 | 22 | Neutrophil |
| Trim30a | 0.87 | 0.00 | 0.00 | 0.26 | 0.08 | 22 | Neutrophil |
| Hist1h2al | 0.85 | 0.00 | 0.00 | 0.29 | 0.10 | 22 | Neutrophil |
| Pkm | 0.85 | 0.00 | 0.00 | 0.47 | 0.22 | 22 | Neutrophil |
| Syne1 | 0.72 | 0.00 | 0.00 | 0.30 | 0.11 | 22 | Neutrophil |
| Lrrc58 | 0.51 | 0.00 | 0.00 | 0.98 | 0.91 | 22 | Neutrophil |
| Emb | 0.63 | 0.00 | 0.00 | 0.31 | 0.11 | 22 | Neutrophil |
| Fam49b | 0.60 | 0.00 | 0.00 | 0.30 | 0.11 | 22 | Neutrophil |
| Ostf1 | 0.78 | 0.00 | 0.00 | 0.50 | 0.26 | 22 | Neutrophil |
| 2810474O19Rik | 1.05 | 0.00 | 0.00 | 0.50 | 0.26 | 22 | Neutrophil |
| Sh3bgrl3 | 0.62 | 0.00 | 0.00 | 0.61 | 0.34 | 22 | Neutrophil |
| H2-D1 | 0.60 | 0.00 | 0.00 | 0.78 | 0.49 | 22 | Neutrophil |
| Trim12c | 0.78 | 0.00 | 0.00 | 0.27 | 0.09 | 22 | Neutrophil |
| Arpc1b | 0.63 | 0.00 | 0.00 | 0.62 | 0.37 | 22 | Neutrophil |
| Pim1 | 0.72 | 0.00 | 0.00 | 0.27 | 0.09 | 22 | Neutrophil |
| Egr1 | 0.88 | 0.00 | 0.00 | 0.33 | 0.14 | 22 | Neutrophil |
| Ier2 | 0.68 | 0.00 | 0.00 | 0.48 | 0.25 | 22 | Neutrophil |
| Gpcpd1 | 0.71 | 0.00 | 0.00 | 0.25 | 0.09 | 22 | Neutrophil |
| Vasp | 0.69 | 0.00 | 0.00 | 0.39 | 0.19 | 22 | Neutrophil |
| Rhog | 0.62 | 0.00 | 0.00 | 0.28 | 0.11 | 22 | Neutrophil |
| Cybb | 0.55 | 0.00 | 0.00 | 0.25 | 0.10 | 22 | Neutrophil |
| Ogfrl1 | 0.65 | 0.00 | 0.00 | 0.25 | 0.10 | 22 | Neutrophil |
| Lsp1 | 0.48 | 0.00 | 0.00 | 0.40 | 0.19 | 22 | Neutrophil |
| Txn1 | 0.51 | 0.00 | 0.00 | 0.70 | 0.50 | 22 | Neutrophil |
| Cap1 | 0.62 | 0.00 | 0.00 | 0.35 | 0.17 | 22 | Neutrophil |
| Fth1 | 0.92 | 0.00 | 0.00 | 0.90 | 0.81 | 22 | Neutrophil |
| Ifitm1 | 0.59 | 0.00 | 0.00 | 0.31 | 0.14 | 22 | Neutrophil |
| Prkcd | 0.56 | 0.00 | 0.00 | 0.26 | 0.11 | 22 | Neutrophil |
| Klf6 | 0.57 | 0.00 | 0.00 | 0.56 | 0.36 | 22 | Neutrophil |
| Zcchc6 | 0.64 | 0.00 | 0.00 | 0.43 | 0.25 | 22 | Neutrophil |
| Gnai2 | 0.44 | 0.00 | 0.00 | 0.59 | 0.39 | 22 | Neutrophil |
| B2m | 0.50 | 0.00 | 0.00 | 0.86 | 0.67 | 22 | Neutrophil |
| Jhdm1d | 0.57 | 0.00 | 0.00 | 0.32 | 0.16 | 22 | Neutrophil |
| Pfn1 | 0.45 | 0.00 | 0.00 | 0.65 | 0.46 | 22 | Neutrophil |
| Nfkbia | 0.78 | 0.00 | 0.00 | 0.56 | 0.37 | 22 | Neutrophil |
| Ier3 | 0.78 | 0.00 | 0.00 | 0.28 | 0.14 | 22 | Neutrophil |
| Neat1 | 0.46 | 0.00 | 0.00 | 0.47 | 0.27 | 22 | Neutrophil |
| Fam101b | 0.50 | 0.00 | 0.00 | 0.35 | 0.19 | 22 | Neutrophil |
| Plin2 | 0.73 | 0.00 | 0.00 | 0.28 | 0.13 | 22 | Neutrophil |
| Gpx1 | 0.50 | 0.00 | 0.00 | 0.61 | 0.43 | 22 | Neutrophil |
| Tkt | 0.48 | 0.00 | 0.00 | 0.29 | 0.15 | 22 | Neutrophil |
| 9930111J21Rik2 | 0.59 | 0.00 | 0.00 | 0.25 | 0.12 | 22 | Neutrophil |
| Lrrfip1 | 0.49 | 0.00 | 0.00 | 0.42 | 0.26 | 22 | Neutrophil |
| mt-Rnr1 | 0.50 | 0.00 | 0.00 | 0.64 | 0.46 | 22 | Neutrophil |
| Ctsd | 0.58 | 0.00 | 0.00 | 0.40 | 0.25 | 22 | Neutrophil |
| Fam111a | 0.48 | 0.00 | 0.00 | 0.30 | 0.15 | 22 | Neutrophil |
| Gpi1 | 0.49 | 0.00 | 0.00 | 0.27 | 0.14 | 22 | Neutrophil |
| Tspo | 0.43 | 0.00 | 0.00 | 0.42 | 0.26 | 22 | Neutrophil |
| Akap13 | 0.55 | 0.00 | 0.00 | 0.42 | 0.27 | 22 | Neutrophil |
| Mgst1 | 0.49 | 0.00 | 0.00 | 0.35 | 0.20 | 22 | Neutrophil |
| Shfm1 | 0.35 | 0.00 | 0.00 | 0.77 | 0.64 | 22 | Neutrophil |
| Arpc2 | 0.35 | 0.00 | 0.00 | 0.76 | 0.61 | 22 | Neutrophil |
| Actr2 | 0.42 | 0.00 | 0.00 | 0.43 | 0.28 | 22 | Neutrophil |
| Atp6v1g1 | 0.51 | 0.00 | 0.00 | 0.48 | 0.32 | 22 | Neutrophil |
| Fam32a | 0.50 | 0.00 | 0.00 | 0.39 | 0.24 | 22 | Neutrophil |
| Cmip | 0.47 | 0.00 | 0.00 | 0.34 | 0.20 | 22 | Neutrophil |
| Stk38 | 0.45 | 0.00 | 0.00 | 0.34 | 0.20 | 22 | Neutrophil |
| Lmo4 | 0.43 | 0.00 | 0.00 | 0.40 | 0.26 | 22 | Neutrophil |
| Arid4a | 0.58 | 0.00 | 0.00 | 0.43 | 0.29 | 22 | Neutrophil |
| Atp6v0c | 0.65 | 0.00 | 0.00 | 0.39 | 0.26 | 22 | Neutrophil |
| Tsc22d4 | 0.50 | 0.00 | 0.00 | 0.41 | 0.28 | 22 | Neutrophil |
| Irf2 | 0.50 | 0.00 | 0.00 | 0.38 | 0.24 | 22 | Neutrophil |
| Ptpn1 | 0.50 | 0.00 | 0.00 | 0.33 | 0.20 | 22 | Neutrophil |
| Mrpl33 | 0.47 | 0.00 | 0.00 | 0.43 | 0.30 | 22 | Neutrophil |
| Jund | 0.46 | 0.00 | 0.00 | 0.41 | 0.28 | 22 | Neutrophil |
| Serp1 | 0.47 | 0.00 | 0.00 | 0.36 | 0.23 | 22 | Neutrophil |
| Gm5506 | 0.39 | 0.00 | 0.00 | 0.26 | 0.14 | 22 | Neutrophil |
| Actg1 | 0.26 | 0.00 | 0.00 | 0.86 | 0.76 | 22 | Neutrophil |
| Iqgap1 | 0.39 | 0.00 | 0.00 | 0.58 | 0.45 | 22 | Neutrophil |
| Ptbp3 | 0.38 | 0.00 | 0.00 | 0.49 | 0.36 | 22 | Neutrophil |
| Ftl1 | 0.56 | 0.00 | 0.00 | 0.69 | 0.57 | 22 | Neutrophil |
| Actn1 | 0.41 | 0.00 | 0.00 | 0.35 | 0.23 | 22 | Neutrophil |
| Emd | 0.39 | 0.00 | 0.00 | 0.53 | 0.40 | 22 | Neutrophil |
| Grb2 | 0.42 | 0.00 | 0.00 | 0.29 | 0.18 | 22 | Neutrophil |
| Gm10282 | 0.41 | 0.00 | 0.00 | 0.52 | 0.40 | 22 | Neutrophil |
| Limd2 | 0.32 | 0.00 | 0.00 | 0.26 | 0.15 | 22 | Neutrophil |
| Msl1 | 0.41 | 0.00 | 0.00 | 0.30 | 0.19 | 22 | Neutrophil |
| Ckap4 | 0.32 | 0.00 | 0.00 | 0.29 | 0.18 | 22 | Neutrophil |
| Map1lc3b | 0.45 | 0.00 | 0.00 | 0.49 | 0.40 | 22 | Neutrophil |
| Eno1 | 0.37 | 0.00 | 0.00 | 0.26 | 0.16 | 22 | Neutrophil |
| Arpc3 | 0.34 | 0.00 | 0.00 | 0.37 | 0.26 | 22 | Neutrophil |
| Atg3 | 0.48 | 0.00 | 0.00 | 0.28 | 0.19 | 22 | Neutrophil |
| Myh9 | 0.29 | 0.00 | 0.00 | 0.46 | 0.33 | 22 | Neutrophil |
| Actr3 | 0.30 | 0.00 | 0.00 | 0.45 | 0.33 | 22 | Neutrophil |
| Ubc | 0.37 | 0.00 | 0.00 | 0.58 | 0.49 | 22 | Neutrophil |
| Gabarap | 0.39 | 0.00 | 0.00 | 0.53 | 0.44 | 22 | Neutrophil |
| Ypel3 | 0.42 | 0.00 | 0.00 | 0.30 | 0.21 | 22 | Neutrophil |
| Ubb | 0.27 | 0.00 | 0.00 | 0.86 | 0.81 | 22 | Neutrophil |
| Aldoa | 0.36 | 0.00 | 0.00 | 0.29 | 0.19 | 22 | Neutrophil |
| Gm17087 | 0.44 | 0.00 | 0.01 | 0.34 | 0.24 | 22 | Neutrophil |
| Arpc4 | 0.32 | 0.00 | 0.02 | 0.27 | 0.18 | 22 | Neutrophil |
| Atp6v1e1 | 0.42 | 0.00 | 0.03 | 0.34 | 0.25 | 22 | Neutrophil |
| Cd97 | 0.27 | 0.00 | 0.04 | 0.27 | 0.18 | 22 | Neutrophil |
| Top2a | 1.92 | 0.00 | 0.00 | 0.83 | 0.11 | 23 | Matrix Fibroblast |
| Prc1 | 1.79 | 0.00 | 0.00 | 0.71 | 0.06 | 23 | Matrix Fibroblast |
| Mki67 | 1.73 | 0.00 | 0.00 | 0.84 | 0.12 | 23 | Matrix Fibroblast |
| 2810417H13Rik | 1.64 | 0.00 | 0.00 | 0.70 | 0.08 | 23 | Matrix Fibroblast |
| Spc25 | 1.44 | 0.00 | 0.00 | 0.58 | 0.05 | 23 | Matrix Fibroblast |
| Tpx2 | 1.42 | 0.00 | 0.00 | 0.62 | 0.05 | 23 | Matrix Fibroblast |
| Hmmr | 1.39 | 0.00 | 0.00 | 0.54 | 0.04 | 23 | Matrix Fibroblast |
| Ckap2 | 1.28 | 0.00 | 0.00 | 0.53 | 0.04 | 23 | Matrix Fibroblast |
| RP23-45G16.5 | 1.26 | 0.00 | 0.00 | 0.57 | 0.05 | 23 | Matrix Fibroblast |
| Birc5 | 1.23 | 0.00 | 0.00 | 0.62 | 0.06 | 23 | Matrix Fibroblast |
| Cdk1 | 1.22 | 0.00 | 0.00 | 0.57 | 0.05 | 23 | Matrix Fibroblast |
| Spc24 | 1.18 | 0.00 | 0.00 | 0.57 | 0.05 | 23 | Matrix Fibroblast |
| Ckap2l | 1.14 | 0.00 | 0.00 | 0.53 | 0.04 | 23 | Matrix Fibroblast |
| Pbk | 1.07 | 0.00 | 0.00 | 0.45 | 0.03 | 23 | Matrix Fibroblast |
| Tk1 | 1.06 | 0.00 | 0.00 | 0.45 | 0.03 | 23 | Matrix Fibroblast |
| Nuf2 | 1.06 | 0.00 | 0.00 | 0.44 | 0.03 | 23 | Matrix Fibroblast |
| Cenpk | 0.99 | 0.00 | 0.00 | 0.47 | 0.03 | 23 | Matrix Fibroblast |
| Cdca3 | 1.08 | 0.00 | 0.00 | 0.46 | 0.03 | 23 | Matrix Fibroblast |
| Nusap1 | 1.36 | 0.00 | 0.00 | 0.57 | 0.06 | 23 | Matrix Fibroblast |
| Cenpm | 0.72 | 0.00 | 0.00 | 0.36 | 0.02 | 23 | Matrix Fibroblast |
| Sgol2 | 1.20 | 0.00 | 0.00 | 0.47 | 0.04 | 23 | Matrix Fibroblast |
| Anln | 1.00 | 0.00 | 0.00 | 0.43 | 0.03 | 23 | Matrix Fibroblast |
| Kif15 | 1.16 | 0.00 | 0.00 | 0.49 | 0.04 | 23 | Matrix Fibroblast |
| Kif20b | 1.53 | 0.00 | 0.00 | 0.58 | 0.06 | 23 | Matrix Fibroblast |
| Neil3 | 0.88 | 0.00 | 0.00 | 0.36 | 0.02 | 23 | Matrix Fibroblast |
| Hist1h2an | 1.83 | 0.00 | 0.00 | 0.67 | 0.09 | 23 | Matrix Fibroblast |
| Hist1h2ae | 1.86 | 0.00 | 0.00 | 0.66 | 0.09 | 23 | Matrix Fibroblast |
| Cenpf | 1.80 | 0.00 | 0.00 | 0.63 | 0.08 | 23 | Matrix Fibroblast |
| Hist1h2ak | 1.72 | 0.00 | 0.00 | 0.61 | 0.07 | 23 | Matrix Fibroblast |
| Hist1h2ad | 1.90 | 0.00 | 0.00 | 0.71 | 0.10 | 23 | Matrix Fibroblast |
| Cks2 | 1.34 | 0.00 | 0.00 | 0.63 | 0.08 | 23 | Matrix Fibroblast |
| Tacc3 | 1.13 | 0.00 | 0.00 | 0.53 | 0.05 | 23 | Matrix Fibroblast |
| Ncapg | 1.02 | 0.00 | 0.00 | 0.48 | 0.04 | 23 | Matrix Fibroblast |
| Fam64a | 0.85 | 0.00 | 0.00 | 0.32 | 0.02 | 23 | Matrix Fibroblast |
| Ccnb2 | 1.32 | 0.00 | 0.00 | 0.54 | 0.06 | 23 | Matrix Fibroblast |
| Ndc80 | 1.05 | 0.00 | 0.00 | 0.44 | 0.04 | 23 | Matrix Fibroblast |
| Diap3 | 1.19 | 0.00 | 0.00 | 0.54 | 0.06 | 23 | Matrix Fibroblast |
| Cenpe | 1.52 | 0.00 | 0.00 | 0.61 | 0.08 | 23 | Matrix Fibroblast |
| Smc2 | 1.58 | 0.00 | 0.00 | 0.79 | 0.13 | 23 | Matrix Fibroblast |
| Ube2c | 1.32 | 0.00 | 0.00 | 0.50 | 0.05 | 23 | Matrix Fibroblast |
| Kif11 | 0.94 | 0.00 | 0.00 | 0.44 | 0.04 | 23 | Matrix Fibroblast |
| Casc5 | 1.08 | 0.00 | 0.00 | 0.46 | 0.04 | 23 | Matrix Fibroblast |
| Racgap1 | 0.87 | 0.00 | 0.00 | 0.43 | 0.04 | 23 | Matrix Fibroblast |
| Esco2 | 0.96 | 0.00 | 0.00 | 0.40 | 0.03 | 23 | Matrix Fibroblast |
| Hist1h2ah | 1.86 | 0.00 | 0.00 | 0.66 | 0.09 | 23 | Matrix Fibroblast |
| Hist1h2ag | 1.98 | 0.00 | 0.00 | 0.72 | 0.12 | 23 | Matrix Fibroblast |
| Tubb6 | 1.20 | 0.00 | 0.00 | 0.59 | 0.07 | 23 | Matrix Fibroblast |
| Ska3 | 0.67 | 0.00 | 0.00 | 0.28 | 0.02 | 23 | Matrix Fibroblast |
| Hist1h2af | 1.79 | 0.00 | 0.00 | 0.63 | 0.09 | 23 | Matrix Fibroblast |
| Hist1h2ai | 1.91 | 0.00 | 0.00 | 0.66 | 0.10 | 23 | Matrix Fibroblast |
| Hist1h2ab | 1.83 | 0.00 | 0.00 | 0.63 | 0.09 | 23 | Matrix Fibroblast |
| Cenpp | 0.69 | 0.00 | 0.00 | 0.29 | 0.02 | 23 | Matrix Fibroblast |
| Hist1h2ac | 1.92 | 0.00 | 0.00 | 0.67 | 0.10 | 23 | Matrix Fibroblast |
| H2afx | 1.39 | 0.00 | 0.00 | 0.64 | 0.09 | 23 | Matrix Fibroblast |
| Ccna2 | 1.01 | 0.00 | 0.00 | 0.41 | 0.04 | 23 | Matrix Fibroblast |
| Melk | 0.68 | 0.00 | 0.00 | 0.28 | 0.02 | 23 | Matrix Fibroblast |
| Cdca8 | 1.06 | 0.00 | 0.00 | 0.47 | 0.05 | 23 | Matrix Fibroblast |
| Kif20a | 0.75 | 0.00 | 0.00 | 0.31 | 0.02 | 23 | Matrix Fibroblast |
| Cenpa | 1.33 | 0.00 | 0.00 | 0.52 | 0.06 | 23 | Matrix Fibroblast |
| Mxd3 | 0.65 | 0.00 | 0.00 | 0.29 | 0.02 | 23 | Matrix Fibroblast |
| Aspm | 0.93 | 0.00 | 0.00 | 0.36 | 0.03 | 23 | Matrix Fibroblast |
| Hist1h2ap | 2.07 | 0.00 | 0.00 | 0.76 | 0.15 | 23 | Matrix Fibroblast |
| Mis18bp1 | 1.00 | 0.00 | 0.00 | 0.37 | 0.03 | 23 | Matrix Fibroblast |
| Sgol1 | 0.79 | 0.00 | 0.00 | 0.34 | 0.03 | 23 | Matrix Fibroblast |
| Kif23 | 1.15 | 0.00 | 0.00 | 0.47 | 0.05 | 23 | Matrix Fibroblast |
| Hist1h2ao | 2.09 | 0.00 | 0.00 | 0.75 | 0.15 | 23 | Matrix Fibroblast |
| Knstrn | 0.88 | 0.00 | 0.00 | 0.39 | 0.04 | 23 | Matrix Fibroblast |
| Bub1 | 0.64 | 0.00 | 0.00 | 0.31 | 0.03 | 23 | Matrix Fibroblast |
| Ska1 | 0.65 | 0.00 | 0.00 | 0.28 | 0.02 | 23 | Matrix Fibroblast |
| Dlgap5 | 0.70 | 0.00 | 0.00 | 0.31 | 0.03 | 23 | Matrix Fibroblast |
| Tyms | 0.97 | 0.00 | 0.00 | 0.45 | 0.05 | 23 | Matrix Fibroblast |
| Incenp | 1.20 | 0.00 | 0.00 | 0.55 | 0.08 | 23 | Matrix Fibroblast |
| Cdc20 | 1.08 | 0.00 | 0.00 | 0.42 | 0.05 | 23 | Matrix Fibroblast |
| Prrx1 | 0.77 | 0.00 | 0.00 | 0.33 | 0.03 | 23 | Matrix Fibroblast |
| Cenph | 0.84 | 0.00 | 0.00 | 0.35 | 0.03 | 23 | Matrix Fibroblast |
| Ccnb1 | 0.72 | 0.00 | 0.00 | 0.29 | 0.02 | 23 | Matrix Fibroblast |
| Rad51ap1 | 0.71 | 0.00 | 0.00 | 0.34 | 0.03 | 23 | Matrix Fibroblast |
| Lmnb1 | 0.94 | 0.00 | 0.00 | 0.55 | 0.08 | 23 | Matrix Fibroblast |
| Ect2 | 0.67 | 0.00 | 0.00 | 0.28 | 0.02 | 23 | Matrix Fibroblast |
| Clspn | 0.88 | 0.00 | 0.00 | 0.37 | 0.04 | 23 | Matrix Fibroblast |
| Kif22 | 0.66 | 0.00 | 0.00 | 0.28 | 0.02 | 23 | Matrix Fibroblast |
| Fbxo5 | 0.82 | 0.00 | 0.00 | 0.39 | 0.05 | 23 | Matrix Fibroblast |
| Rrm2 | 0.87 | 0.00 | 0.00 | 0.39 | 0.05 | 23 | Matrix Fibroblast |
| C330027C09Rik | 0.74 | 0.00 | 0.00 | 0.31 | 0.03 | 23 | Matrix Fibroblast |
| Cdca2 | 0.58 | 0.00 | 0.00 | 0.25 | 0.02 | 23 | Matrix Fibroblast |
| Aurkb | 0.69 | 0.00 | 0.00 | 0.29 | 0.03 | 23 | Matrix Fibroblast |
| Hist1h1b | 1.00 | 0.00 | 0.00 | 0.34 | 0.04 | 23 | Matrix Fibroblast |
| Arhgap11a | 0.84 | 0.00 | 0.00 | 0.37 | 0.04 | 23 | Matrix Fibroblast |
| Spdl1 | 0.58 | 0.00 | 0.00 | 0.26 | 0.02 | 23 | Matrix Fibroblast |
| Gm10282 | 1.40 | 0.00 | 0.00 | 0.95 | 0.40 | 23 | Matrix Fibroblast |
| Gpc3 | 1.23 | 0.00 | 0.00 | 0.73 | 0.16 | 23 | Matrix Fibroblast |
| Rrm1 | 0.79 | 0.00 | 0.00 | 0.45 | 0.06 | 23 | Matrix Fibroblast |
| Hmgb2 | 1.48 | 0.00 | 0.00 | 0.97 | 0.42 | 23 | Matrix Fibroblast |
| Cep55 | 0.59 | 0.00 | 0.00 | 0.29 | 0.03 | 23 | Matrix Fibroblast |
| Hmgn2 | 1.39 | 0.00 | 0.00 | 0.96 | 0.43 | 23 | Matrix Fibroblast |
| Stmn1 | 1.16 | 0.00 | 0.00 | 0.83 | 0.23 | 23 | Matrix Fibroblast |
| Plk4 | 0.72 | 0.00 | 0.00 | 0.35 | 0.04 | 23 | Matrix Fibroblast |
| Kif4 | 0.62 | 0.00 | 0.00 | 0.26 | 0.03 | 23 | Matrix Fibroblast |
| H2afz | 1.32 | 0.00 | 0.00 | 0.94 | 0.40 | 23 | Matrix Fibroblast |
| Ncapd2 | 0.65 | 0.00 | 0.00 | 0.35 | 0.04 | 23 | Matrix Fibroblast |
| Tpm2 | 1.05 | 0.00 | 0.00 | 0.64 | 0.13 | 23 | Matrix Fibroblast |
| D17H6S56E-5 | 0.92 | 0.00 | 0.00 | 0.56 | 0.10 | 23 | Matrix Fibroblast |
| Cenpw | 0.70 | 0.00 | 0.00 | 0.37 | 0.05 | 23 | Matrix Fibroblast |
| Ccdc34 | 1.07 | 0.00 | 0.00 | 0.63 | 0.14 | 23 | Matrix Fibroblast |
| Cks1b | 0.91 | 0.00 | 0.00 | 0.52 | 0.09 | 23 | Matrix Fibroblast |
| Ncapg2 | 0.59 | 0.00 | 0.00 | 0.28 | 0.03 | 23 | Matrix Fibroblast |
| Hjurp | 1.06 | 0.00 | 0.00 | 0.64 | 0.14 | 23 | Matrix Fibroblast |
| Tuba1b | 1.24 | 0.00 | 0.00 | 0.96 | 0.45 | 23 | Matrix Fibroblast |
| Lgals1 | 1.25 | 0.00 | 0.00 | 0.97 | 0.41 | 23 | Matrix Fibroblast |
| Cenpq | 0.72 | 0.00 | 0.00 | 0.43 | 0.07 | 23 | Matrix Fibroblast |
| Slfn9 | 0.82 | 0.00 | 0.00 | 0.31 | 0.04 | 23 | Matrix Fibroblast |
| Mdk | 0.99 | 0.00 | 0.00 | 0.74 | 0.18 | 23 | Matrix Fibroblast |
| Tcf19 | 0.59 | 0.00 | 0.00 | 0.28 | 0.03 | 23 | Matrix Fibroblast |
| Col1a2 | 1.18 | 0.00 | 0.00 | 0.98 | 0.39 | 23 | Matrix Fibroblast |
| Atad2 | 0.90 | 0.00 | 0.00 | 0.45 | 0.08 | 23 | Matrix Fibroblast |
| Mns1 | 0.72 | 0.00 | 0.00 | 0.32 | 0.04 | 23 | Matrix Fibroblast |
| Slc27a6 | 0.68 | 0.00 | 0.00 | 0.32 | 0.04 | 23 | Matrix Fibroblast |
| Ska2 | 0.66 | 0.00 | 0.00 | 0.41 | 0.07 | 23 | Matrix Fibroblast |
| Itga8 | 0.85 | 0.00 | 0.00 | 0.49 | 0.09 | 23 | Matrix Fibroblast |
| Cdh11 | 1.00 | 0.00 | 0.00 | 0.72 | 0.19 | 23 | Matrix Fibroblast |
| Smc4 | 1.21 | 0.00 | 0.00 | 0.88 | 0.34 | 23 | Matrix Fibroblast |
| Col1a1 | 1.20 | 0.00 | 0.00 | 0.83 | 0.26 | 23 | Matrix Fibroblast |
| Prim1 | 0.70 | 0.00 | 0.00 | 0.35 | 0.05 | 23 | Matrix Fibroblast |
| Rangap1 | 0.77 | 0.00 | 0.00 | 0.43 | 0.08 | 23 | Matrix Fibroblast |
| Plac9a | 0.92 | 0.00 | 0.00 | 0.91 | 0.30 | 23 | Matrix Fibroblast |
| Hirip3 | 0.88 | 0.00 | 0.00 | 0.46 | 0.09 | 23 | Matrix Fibroblast |
| Plac9b | 0.89 | 0.00 | 0.00 | 0.91 | 0.30 | 23 | Matrix Fibroblast |
| Ccne2 | 0.78 | 0.00 | 0.00 | 0.28 | 0.03 | 23 | Matrix Fibroblast |
| Nexn | 1.00 | 0.00 | 0.00 | 0.75 | 0.21 | 23 | Matrix Fibroblast |
| Fbln1 | 0.65 | 0.00 | 0.00 | 0.35 | 0.05 | 23 | Matrix Fibroblast |
| Col3a1 | 0.93 | 0.00 | 0.00 | 0.93 | 0.33 | 23 | Matrix Fibroblast |
| Lig1 | 0.99 | 0.00 | 0.00 | 0.46 | 0.09 | 23 | Matrix Fibroblast |
| Cdkn2c | 0.63 | 0.00 | 0.00 | 0.31 | 0.04 | 23 | Matrix Fibroblast |
| Pola1 | 0.60 | 0.00 | 0.00 | 0.31 | 0.04 | 23 | Matrix Fibroblast |
| Colec12 | 0.83 | 0.00 | 0.00 | 0.49 | 0.10 | 23 | Matrix Fibroblast |
| Myh10 | 0.94 | 0.00 | 0.00 | 0.76 | 0.22 | 23 | Matrix Fibroblast |
| Nucks1 | 1.08 | 0.00 | 0.00 | 0.80 | 0.28 | 23 | Matrix Fibroblast |
| Dpt | 1.00 | 0.00 | 0.00 | 0.65 | 0.17 | 23 | Matrix Fibroblast |
| Fn1 | 0.81 | 0.00 | 0.00 | 0.79 | 0.23 | 23 | Matrix Fibroblast |
| Loxl1 | 0.72 | 0.00 | 0.00 | 0.51 | 0.11 | 23 | Matrix Fibroblast |
| 4930579G24Rik | 0.45 | 0.00 | 0.00 | 0.26 | 0.03 | 23 | Matrix Fibroblast |
| H2afv | 0.99 | 0.00 | 0.00 | 0.79 | 0.28 | 23 | Matrix Fibroblast |
| Hist2h2ab | 0.61 | 0.00 | 0.00 | 0.29 | 0.04 | 23 | Matrix Fibroblast |
| 2700094K13Rik | 0.81 | 0.00 | 0.00 | 0.61 | 0.16 | 23 | Matrix Fibroblast |
| Bgn | 0.84 | 0.00 | 0.00 | 0.77 | 0.24 | 23 | Matrix Fibroblast |
| Gm10259 | 0.70 | 0.00 | 0.00 | 0.35 | 0.06 | 23 | Matrix Fibroblast |
| Pcolce | 0.71 | 0.00 | 0.00 | 0.43 | 0.09 | 23 | Matrix Fibroblast |
| Dhfr | 0.57 | 0.00 | 0.00 | 0.29 | 0.04 | 23 | Matrix Fibroblast |
| Gmnn | 0.63 | 0.00 | 0.00 | 0.38 | 0.07 | 23 | Matrix Fibroblast |
| Tmpo | 0.81 | 0.00 | 0.00 | 0.57 | 0.15 | 23 | Matrix Fibroblast |
| Hist2h2ac | 0.79 | 0.00 | 0.00 | 0.35 | 0.06 | 23 | Matrix Fibroblast |
| Figf | 0.71 | 0.00 | 0.00 | 0.37 | 0.07 | 23 | Matrix Fibroblast |
| Mfap4 | 0.91 | 0.00 | 0.00 | 0.75 | 0.24 | 23 | Matrix Fibroblast |
| Tuba1c | 0.99 | 0.00 | 0.00 | 0.81 | 0.32 | 23 | Matrix Fibroblast |
| Adamts2 | 0.66 | 0.00 | 0.00 | 0.47 | 0.10 | 23 | Matrix Fibroblast |
| Dut | 0.66 | 0.00 | 0.00 | 0.42 | 0.09 | 23 | Matrix Fibroblast |
| Ncapd3 | 0.55 | 0.00 | 0.00 | 0.32 | 0.06 | 23 | Matrix Fibroblast |
| Tubb5 | 0.98 | 0.00 | 0.00 | 0.86 | 0.44 | 23 | Matrix Fibroblast |
| Meox2 | 0.61 | 0.00 | 0.00 | 0.39 | 0.08 | 23 | Matrix Fibroblast |
| Ran | 0.92 | 0.00 | 0.00 | 0.84 | 0.38 | 23 | Matrix Fibroblast |
| Tcf21 | 0.63 | 0.00 | 0.00 | 0.63 | 0.17 | 23 | Matrix Fibroblast |
| Plxdc2 | 0.62 | 0.00 | 0.00 | 0.51 | 0.12 | 23 | Matrix Fibroblast |
| Lbh | 0.80 | 0.00 | 0.00 | 0.64 | 0.19 | 23 | Matrix Fibroblast |
| Dnajc9 | 0.67 | 0.00 | 0.00 | 0.49 | 0.12 | 23 | Matrix Fibroblast |
| Adh1 | 0.93 | 0.00 | 0.00 | 0.59 | 0.17 | 23 | Matrix Fibroblast |
| Usp1 | 0.75 | 0.00 | 0.00 | 0.56 | 0.15 | 23 | Matrix Fibroblast |
| Ezh2 | 0.79 | 0.00 | 0.00 | 0.62 | 0.19 | 23 | Matrix Fibroblast |
| Dek | 0.90 | 0.00 | 0.00 | 0.91 | 0.50 | 23 | Matrix Fibroblast |
| Ckap5 | 0.76 | 0.00 | 0.00 | 0.46 | 0.11 | 23 | Matrix Fibroblast |
| Mgp | 0.98 | 0.00 | 0.00 | 0.82 | 0.31 | 23 | Matrix Fibroblast |
| Ncaph | 0.47 | 0.00 | 0.00 | 0.28 | 0.05 | 23 | Matrix Fibroblast |
| Fbn1 | 0.67 | 0.00 | 0.00 | 0.61 | 0.17 | 23 | Matrix Fibroblast |
| Hmgb1 | 0.85 | 0.00 | 0.00 | 0.94 | 0.61 | 23 | Matrix Fibroblast |
| Myl9 | 0.83 | 0.00 | 0.00 | 0.48 | 0.12 | 23 | Matrix Fibroblast |
| Cenpc1 | 0.70 | 0.00 | 0.00 | 0.39 | 0.09 | 23 | Matrix Fibroblast |
| Gm10184 | 0.71 | 0.00 | 0.00 | 0.34 | 0.07 | 23 | Matrix Fibroblast |
| Prdx4 | 0.75 | 0.00 | 0.00 | 0.56 | 0.16 | 23 | Matrix Fibroblast |
| Acta2 | 1.02 | 0.00 | 0.00 | 0.63 | 0.19 | 23 | Matrix Fibroblast |
| Whsc1 | 0.71 | 0.00 | 0.00 | 0.50 | 0.14 | 23 | Matrix Fibroblast |
| Cmc2 | 0.48 | 0.00 | 0.00 | 0.29 | 0.05 | 23 | Matrix Fibroblast |
| Trim59 | 0.59 | 0.00 | 0.00 | 0.26 | 0.04 | 23 | Matrix Fibroblast |
| Lox | 0.55 | 0.00 | 0.00 | 0.52 | 0.13 | 23 | Matrix Fibroblast |
| Cped1 | 0.62 | 0.00 | 0.00 | 0.42 | 0.10 | 23 | Matrix Fibroblast |
| Ugdh | 0.61 | 0.00 | 0.00 | 0.39 | 0.09 | 23 | Matrix Fibroblast |
| Ccdc80 | 0.70 | 0.00 | 0.00 | 0.45 | 0.11 | 23 | Matrix Fibroblast |
| Gpx7 | 0.63 | 0.00 | 0.00 | 0.41 | 0.10 | 23 | Matrix Fibroblast |
| Kpna2 | 0.68 | 0.00 | 0.00 | 0.31 | 0.06 | 23 | Matrix Fibroblast |
| Rbp1 | 0.64 | 0.00 | 0.00 | 0.63 | 0.19 | 23 | Matrix Fibroblast |
| Nkain4 | 0.52 | 0.00 | 0.00 | 0.26 | 0.05 | 23 | Matrix Fibroblast |
| 6030408B16Rik | 0.51 | 0.00 | 0.00 | 0.27 | 0.05 | 23 | Matrix Fibroblast |
| G2e3 | 0.57 | 0.00 | 0.00 | 0.32 | 0.06 | 23 | Matrix Fibroblast |
| Mfap5 | 0.81 | 0.00 | 0.00 | 0.32 | 0.06 | 23 | Matrix Fibroblast |
| Ckap4 | 0.75 | 0.00 | 0.00 | 0.57 | 0.18 | 23 | Matrix Fibroblast |
| Vcan | 0.68 | 0.00 | 0.00 | 0.44 | 0.11 | 23 | Matrix Fibroblast |
| Fhl1 | 0.67 | 0.00 | 0.00 | 0.78 | 0.29 | 23 | Matrix Fibroblast |
| Hist1h1d | 0.85 | 0.00 | 0.00 | 0.41 | 0.10 | 23 | Matrix Fibroblast |
| Dtymk | 0.59 | 0.00 | 0.00 | 0.49 | 0.13 | 23 | Matrix Fibroblast |
| Serpine2 | 0.51 | 0.00 | 0.00 | 0.74 | 0.25 | 23 | Matrix Fibroblast |
| Mmp2 | 0.63 | 0.00 | 0.00 | 0.49 | 0.13 | 23 | Matrix Fibroblast |
| Slc38a5 | 0.54 | 0.00 | 0.00 | 0.30 | 0.06 | 23 | Matrix Fibroblast |
| Rarres2 | 0.56 | 0.00 | 0.00 | 0.49 | 0.13 | 23 | Matrix Fibroblast |
| Fstl1 | 0.69 | 0.00 | 0.00 | 0.91 | 0.44 | 23 | Matrix Fibroblast |
| Cntln | 0.72 | 0.00 | 0.00 | 0.44 | 0.12 | 23 | Matrix Fibroblast |
| 4632419I22Rik | 0.51 | 0.00 | 0.00 | 0.35 | 0.08 | 23 | Matrix Fibroblast |
| Rnd3 | 0.47 | 0.00 | 0.00 | 0.35 | 0.07 | 23 | Matrix Fibroblast |
| Dbf4 | 0.56 | 0.00 | 0.00 | 0.28 | 0.05 | 23 | Matrix Fibroblast |
| Tubb4b | 0.79 | 0.00 | 0.00 | 0.57 | 0.19 | 23 | Matrix Fibroblast |
| Enpep | 0.59 | 0.00 | 0.00 | 0.41 | 0.10 | 23 | Matrix Fibroblast |
| Pcolce2 | 0.59 | 0.00 | 0.00 | 0.39 | 0.09 | 23 | Matrix Fibroblast |
| Spon1 | 0.54 | 0.00 | 0.00 | 0.33 | 0.07 | 23 | Matrix Fibroblast |
| Psip1 | 0.74 | 0.00 | 0.00 | 0.50 | 0.15 | 23 | Matrix Fibroblast |
| Gucy1b3 | 0.55 | 0.00 | 0.00 | 0.29 | 0.06 | 23 | Matrix Fibroblast |
| Cbx5 | 0.72 | 0.00 | 0.00 | 0.69 | 0.26 | 23 | Matrix Fibroblast |
| Fxyd1 | 0.49 | 0.00 | 0.00 | 0.38 | 0.09 | 23 | Matrix Fibroblast |
| Atad5 | 0.54 | 0.00 | 0.00 | 0.28 | 0.05 | 23 | Matrix Fibroblast |
| Serping1 | 0.51 | 0.00 | 0.00 | 0.46 | 0.12 | 23 | Matrix Fibroblast |
| Nasp | 0.69 | 0.00 | 0.00 | 0.59 | 0.20 | 23 | Matrix Fibroblast |
| Ptprd | 0.58 | 0.00 | 0.00 | 0.43 | 0.11 | 23 | Matrix Fibroblast |
| Tuba1a | 0.75 | 0.00 | 0.00 | 0.90 | 0.51 | 23 | Matrix Fibroblast |
| Fblim1 | 0.56 | 0.00 | 0.00 | 0.54 | 0.16 | 23 | Matrix Fibroblast |
| Ednra | 0.56 | 0.00 | 0.00 | 0.37 | 0.09 | 23 | Matrix Fibroblast |
| Iigp1 | 0.77 | 0.00 | 0.00 | 0.47 | 0.13 | 23 | Matrix Fibroblast |
| Col6a2 | 0.53 | 0.00 | 0.00 | 0.50 | 0.15 | 23 | Matrix Fibroblast |
| Nr2f2 | 0.62 | 0.00 | 0.00 | 0.61 | 0.21 | 23 | Matrix Fibroblast |
| Olfml3 | 0.56 | 0.00 | 0.00 | 0.36 | 0.08 | 23 | Matrix Fibroblast |
| 10-Sep | 0.55 | 0.00 | 0.00 | 0.37 | 0.09 | 23 | Matrix Fibroblast |
| Rad21 | 0.85 | 0.00 | 0.00 | 0.62 | 0.23 | 23 | Matrix Fibroblast |
| Nnat | 0.75 | 0.00 | 0.00 | 0.33 | 0.08 | 23 | Matrix Fibroblast |
| Gucy1a3 | 0.71 | 0.00 | 0.00 | 0.39 | 0.10 | 23 | Matrix Fibroblast |
| Cdk5rap2 | 0.57 | 0.00 | 0.00 | 0.37 | 0.09 | 23 | Matrix Fibroblast |
| Fbln5 | 0.58 | 0.00 | 0.00 | 0.54 | 0.17 | 23 | Matrix Fibroblast |
| Bicc1 | 0.55 | 0.00 | 0.00 | 0.31 | 0.07 | 23 | Matrix Fibroblast |
| Col13a1 | 0.53 | 0.00 | 0.00 | 0.38 | 0.10 | 23 | Matrix Fibroblast |
| Hist1h1e | 1.01 | 0.00 | 0.00 | 0.50 | 0.16 | 23 | Matrix Fibroblast |
| Ppp1r14a | 0.37 | 0.00 | 0.00 | 0.35 | 0.08 | 23 | Matrix Fibroblast |
| Hmgn1 | 0.72 | 0.00 | 0.00 | 0.85 | 0.45 | 23 | Matrix Fibroblast |
| Col6a3 | 0.51 | 0.00 | 0.00 | 0.38 | 0.10 | 23 | Matrix Fibroblast |
| Pcna-ps2 | 0.87 | 0.00 | 0.00 | 0.51 | 0.18 | 23 | Matrix Fibroblast |
| Ptma | 0.51 | 0.00 | 0.00 | 1.00 | 0.91 | 23 | Matrix Fibroblast |
| Pcna | 0.85 | 0.00 | 0.00 | 0.51 | 0.18 | 23 | Matrix Fibroblast |
| Anp32b | 0.67 | 0.00 | 0.00 | 0.88 | 0.50 | 23 | Matrix Fibroblast |
| Akap12 | 0.70 | 0.00 | 0.00 | 0.63 | 0.24 | 23 | Matrix Fibroblast |
| Rfc3 | 0.50 | 0.00 | 0.00 | 0.35 | 0.09 | 23 | Matrix Fibroblast |
| Fam111a | 0.52 | 0.00 | 0.00 | 0.49 | 0.15 | 23 | Matrix Fibroblast |
| 5830418K08Rik | 0.46 | 0.00 | 0.00 | 0.35 | 0.09 | 23 | Matrix Fibroblast |
| Anp32e | 0.69 | 0.00 | 0.00 | 0.69 | 0.28 | 23 | Matrix Fibroblast |
| Cox6b2 | 0.42 | 0.00 | 0.00 | 0.25 | 0.05 | 23 | Matrix Fibroblast |
| Sema5a | 0.41 | 0.00 | 0.00 | 0.26 | 0.06 | 23 | Matrix Fibroblast |
| Oxct1 | 0.56 | 0.00 | 0.00 | 0.47 | 0.15 | 23 | Matrix Fibroblast |
| Postn | 0.99 | 0.00 | 0.00 | 0.32 | 0.08 | 23 | Matrix Fibroblast |
| Dnmt1 | 0.53 | 0.00 | 0.00 | 0.38 | 0.10 | 23 | Matrix Fibroblast |
| Idh2 | 0.54 | 0.00 | 0.00 | 0.55 | 0.19 | 23 | Matrix Fibroblast |
| Gm10068 | 0.65 | 0.00 | 0.00 | 0.69 | 0.30 | 23 | Matrix Fibroblast |
| Mcm4 | 0.40 | 0.00 | 0.00 | 0.26 | 0.06 | 23 | Matrix Fibroblast |
| Hells | 0.50 | 0.00 | 0.00 | 0.26 | 0.06 | 23 | Matrix Fibroblast |
| Sparc | 0.59 | 0.00 | 0.00 | 0.99 | 0.73 | 23 | Matrix Fibroblast |
| Rap2a | 0.62 | 0.00 | 0.00 | 0.48 | 0.16 | 23 | Matrix Fibroblast |
| Gm10123 | 0.56 | 0.00 | 0.00 | 0.95 | 0.81 | 23 | Matrix Fibroblast |
| Prkcdbp | 0.52 | 0.00 | 0.00 | 0.39 | 0.11 | 23 | Matrix Fibroblast |
| Hsp90aa1 | 0.58 | 0.00 | 0.00 | 0.85 | 0.47 | 23 | Matrix Fibroblast |
| Col16a1 | 0.39 | 0.00 | 0.00 | 0.26 | 0.06 | 23 | Matrix Fibroblast |
| Cp | 0.49 | 0.00 | 0.00 | 0.42 | 0.12 | 23 | Matrix Fibroblast |
| Rcn3 | 0.47 | 0.00 | 0.00 | 0.43 | 0.13 | 23 | Matrix Fibroblast |
| Banf1 | 0.63 | 0.00 | 0.00 | 0.67 | 0.28 | 23 | Matrix Fibroblast |
| Nde1 | 0.47 | 0.00 | 0.00 | 0.28 | 0.06 | 23 | Matrix Fibroblast |
| Slit2 | 0.55 | 0.00 | 0.00 | 0.31 | 0.08 | 23 | Matrix Fibroblast |
| Hist2h2aa2 | 0.79 | 0.00 | 0.00 | 0.42 | 0.14 | 23 | Matrix Fibroblast |
| Ranbp1 | 0.65 | 0.00 | 0.00 | 0.67 | 0.29 | 23 | Matrix Fibroblast |
| Snai2 | 0.50 | 0.00 | 0.00 | 0.41 | 0.12 | 23 | Matrix Fibroblast |
| Nop56 | 0.60 | 0.00 | 0.00 | 0.54 | 0.20 | 23 | Matrix Fibroblast |
| Cbx3 | 0.64 | 0.00 | 0.00 | 0.73 | 0.34 | 23 | Matrix Fibroblast |
| Gyg | 0.62 | 0.00 | 0.00 | 0.53 | 0.19 | 23 | Matrix Fibroblast |
| Mcm7 | 0.35 | 0.00 | 0.00 | 0.30 | 0.07 | 23 | Matrix Fibroblast |
| Limch1 | 0.57 | 0.00 | 0.00 | 0.70 | 0.30 | 23 | Matrix Fibroblast |
| Sae1 | 0.45 | 0.00 | 0.00 | 0.38 | 0.11 | 23 | Matrix Fibroblast |
| Lmna | 0.57 | 0.00 | 0.00 | 0.54 | 0.20 | 23 | Matrix Fibroblast |
| Gsn | 0.44 | 0.00 | 0.00 | 0.64 | 0.25 | 23 | Matrix Fibroblast |
| Itm2a | 0.38 | 0.00 | 0.00 | 0.28 | 0.07 | 23 | Matrix Fibroblast |
| Snrpd1 | 0.56 | 0.00 | 0.00 | 0.55 | 0.20 | 23 | Matrix Fibroblast |
| Tsen15 | 0.38 | 0.00 | 0.00 | 0.27 | 0.06 | 23 | Matrix Fibroblast |
| Cald1 | 0.60 | 0.00 | 0.00 | 0.85 | 0.41 | 23 | Matrix Fibroblast |
| Smc6 | 0.61 | 0.00 | 0.00 | 0.72 | 0.33 | 23 | Matrix Fibroblast |
| Lsm5 | 0.41 | 0.00 | 0.00 | 0.39 | 0.12 | 23 | Matrix Fibroblast |
| Igfbp7 | 0.57 | 0.00 | 0.00 | 0.80 | 0.39 | 23 | Matrix Fibroblast |
| Sh3bgrl | 0.59 | 0.00 | 0.00 | 0.84 | 0.44 | 23 | Matrix Fibroblast |
| Col5a2 | 0.46 | 0.00 | 0.00 | 0.51 | 0.18 | 23 | Matrix Fibroblast |
| Gjc1 | 0.44 | 0.00 | 0.00 | 0.30 | 0.08 | 23 | Matrix Fibroblast |
| Fkbp9 | 0.43 | 0.00 | 0.00 | 0.44 | 0.14 | 23 | Matrix Fibroblast |
| Hist1h1c | 0.59 | 0.00 | 0.00 | 0.37 | 0.11 | 23 | Matrix Fibroblast |
| 1700025G04Rik | 0.43 | 0.00 | 0.00 | 0.35 | 0.10 | 23 | Matrix Fibroblast |
| Slbp | 0.51 | 0.00 | 0.00 | 0.44 | 0.15 | 23 | Matrix Fibroblast |
| Ltbp4 | 0.41 | 0.00 | 0.00 | 0.37 | 0.11 | 23 | Matrix Fibroblast |
| Etaa1 | 0.39 | 0.00 | 0.00 | 0.26 | 0.06 | 23 | Matrix Fibroblast |
| Ilf3 | 0.51 | 0.00 | 0.00 | 0.39 | 0.12 | 23 | Matrix Fibroblast |
| Cryab | 0.36 | 0.00 | 0.00 | 0.44 | 0.14 | 23 | Matrix Fibroblast |
| Gpm6b | 0.49 | 0.00 | 0.00 | 0.37 | 0.11 | 23 | Matrix Fibroblast |
| Lsm6 | 0.45 | 0.00 | 0.00 | 0.42 | 0.13 | 23 | Matrix Fibroblast |
| Hmgn5 | 0.61 | 0.00 | 0.00 | 0.63 | 0.26 | 23 | Matrix Fibroblast |
| Rab34 | 0.35 | 0.00 | 0.00 | 0.31 | 0.08 | 23 | Matrix Fibroblast |
| Ssrp1 | 0.55 | 0.00 | 0.00 | 0.55 | 0.21 | 23 | Matrix Fibroblast |
| 7-Sep | 0.60 | 0.00 | 0.00 | 0.91 | 0.58 | 23 | Matrix Fibroblast |
| Tssc4 | 0.41 | 0.00 | 0.00 | 0.32 | 0.09 | 23 | Matrix Fibroblast |
| Nfia | 0.55 | 0.00 | 0.00 | 0.59 | 0.24 | 23 | Matrix Fibroblast |
| Pls3 | 0.50 | 0.00 | 0.00 | 0.54 | 0.20 | 23 | Matrix Fibroblast |
| Ptms | 0.68 | 0.00 | 0.00 | 0.77 | 0.40 | 23 | Matrix Fibroblast |
| Palld | 0.36 | 0.00 | 0.00 | 0.51 | 0.17 | 23 | Matrix Fibroblast |
| 2700029M09Rik | 0.46 | 0.00 | 0.00 | 0.40 | 0.13 | 23 | Matrix Fibroblast |
| Adarb1 | 0.41 | 0.00 | 0.00 | 0.29 | 0.08 | 23 | Matrix Fibroblast |
| Hhip | 0.40 | 0.00 | 0.00 | 0.28 | 0.07 | 23 | Matrix Fibroblast |
| Maf | 0.44 | 0.00 | 0.00 | 0.53 | 0.19 | 23 | Matrix Fibroblast |
| Tubb4b-ps1 | 0.50 | 0.00 | 0.00 | 0.31 | 0.09 | 23 | Matrix Fibroblast |
| 0610007N19Rik | 0.39 | 0.00 | 0.00 | 0.44 | 0.15 | 23 | Matrix Fibroblast |
| Cdkn2d | 0.45 | 0.00 | 0.00 | 0.27 | 0.07 | 23 | Matrix Fibroblast |
| Ppia | 0.53 | 0.00 | 0.00 | 0.93 | 0.80 | 23 | Matrix Fibroblast |
| Crtap | 0.32 | 0.00 | 0.00 | 0.29 | 0.07 | 23 | Matrix Fibroblast |
| Emilin1 | 0.36 | 0.00 | 0.00 | 0.33 | 0.09 | 23 | Matrix Fibroblast |
| Ism1 | 0.39 | 0.00 | 0.00 | 0.32 | 0.09 | 23 | Matrix Fibroblast |
| Mrpl51 | 0.47 | 0.00 | 0.00 | 0.45 | 0.16 | 23 | Matrix Fibroblast |
| Twsg1 | 0.43 | 0.00 | 0.00 | 0.38 | 0.12 | 23 | Matrix Fibroblast |
| Snhg5 | 0.45 | 0.00 | 0.00 | 0.49 | 0.18 | 23 | Matrix Fibroblast |
| Rbbp7 | 0.49 | 0.00 | 0.00 | 0.62 | 0.26 | 23 | Matrix Fibroblast |
| Npnt | 0.64 | 0.00 | 0.00 | 0.58 | 0.24 | 23 | Matrix Fibroblast |
| Hivep3 | 0.44 | 0.00 | 0.00 | 0.28 | 0.07 | 23 | Matrix Fibroblast |
| Myof | 0.40 | 0.00 | 0.00 | 0.33 | 0.10 | 23 | Matrix Fibroblast |
| Gm4204 | 0.51 | 0.00 | 0.00 | 0.85 | 0.50 | 23 | Matrix Fibroblast |
| Smc1a | 0.52 | 0.00 | 0.00 | 0.73 | 0.35 | 23 | Matrix Fibroblast |
| Baz1b | 0.49 | 0.00 | 0.00 | 0.63 | 0.26 | 23 | Matrix Fibroblast |
| Csrp1 | 0.39 | 0.00 | 0.00 | 0.47 | 0.17 | 23 | Matrix Fibroblast |
| Sparcl1 | 0.39 | 0.00 | 0.00 | 0.81 | 0.39 | 23 | Matrix Fibroblast |
| Egflam | 0.41 | 0.00 | 0.00 | 0.29 | 0.08 | 23 | Matrix Fibroblast |
| Emp3 | 0.42 | 0.00 | 0.00 | 0.55 | 0.21 | 23 | Matrix Fibroblast |
| Bmpr1a | 0.41 | 0.00 | 0.00 | 0.32 | 0.09 | 23 | Matrix Fibroblast |
| Kdelc2 | 0.38 | 0.00 | 0.00 | 0.35 | 0.11 | 23 | Matrix Fibroblast |
| Tspan3 | 0.39 | 0.00 | 0.00 | 0.45 | 0.16 | 23 | Matrix Fibroblast |
| Igf1 | 0.31 | 0.00 | 0.00 | 0.42 | 0.14 | 23 | Matrix Fibroblast |
| Nap1l1 | 0.48 | 0.00 | 0.00 | 0.82 | 0.47 | 23 | Matrix Fibroblast |
| Isoc1 | 0.37 | 0.00 | 0.00 | 0.28 | 0.08 | 23 | Matrix Fibroblast |
| Angpt1 | 0.36 | 0.00 | 0.00 | 0.28 | 0.08 | 23 | Matrix Fibroblast |
| Gpx8 | 0.40 | 0.00 | 0.00 | 0.42 | 0.14 | 23 | Matrix Fibroblast |
| Aebp1 | 0.45 | 0.00 | 0.00 | 0.28 | 0.08 | 23 | Matrix Fibroblast |
| Mesdc2 | 0.43 | 0.00 | 0.00 | 0.50 | 0.19 | 23 | Matrix Fibroblast |
| Nfib | 0.51 | 0.00 | 0.00 | 0.84 | 0.47 | 23 | Matrix Fibroblast |
| Ptprs | 0.32 | 0.00 | 0.00 | 0.29 | 0.08 | 23 | Matrix Fibroblast |
| C1qtnf7 | 0.32 | 0.00 | 0.00 | 0.32 | 0.09 | 23 | Matrix Fibroblast |
| Lsm3 | 0.43 | 0.00 | 0.00 | 0.34 | 0.11 | 23 | Matrix Fibroblast |
| Nid1 | 0.44 | 0.00 | 0.00 | 0.57 | 0.23 | 23 | Matrix Fibroblast |
| Lyar | 0.40 | 0.00 | 0.00 | 0.43 | 0.15 | 23 | Matrix Fibroblast |
| Dkk3 | 0.36 | 0.00 | 0.00 | 0.37 | 0.12 | 23 | Matrix Fibroblast |
| Atp1a2 | 0.33 | 0.00 | 0.00 | 0.33 | 0.10 | 23 | Matrix Fibroblast |
| Gm5160 | 0.40 | 0.00 | 0.00 | 0.54 | 0.22 | 23 | Matrix Fibroblast |
| Lamb1 | 0.42 | 0.00 | 0.00 | 0.44 | 0.16 | 23 | Matrix Fibroblast |
| Nfix | 0.36 | 0.00 | 0.00 | 0.46 | 0.17 | 23 | Matrix Fibroblast |
| Mcm3 | 0.40 | 0.00 | 0.00 | 0.26 | 0.07 | 23 | Matrix Fibroblast |
| Tipin | 0.40 | 0.00 | 0.00 | 0.32 | 0.10 | 23 | Matrix Fibroblast |
| Smchd1 | 0.45 | 0.00 | 0.00 | 0.44 | 0.16 | 23 | Matrix Fibroblast |
| Xpo1 | 0.40 | 0.00 | 0.00 | 0.31 | 0.10 | 23 | Matrix Fibroblast |
| Eln | 0.37 | 0.00 | 0.00 | 0.44 | 0.16 | 23 | Matrix Fibroblast |
| Nop58 | 0.45 | 0.00 | 0.00 | 0.60 | 0.26 | 23 | Matrix Fibroblast |
| Pa2g4 | 0.41 | 0.00 | 0.00 | 0.61 | 0.26 | 23 | Matrix Fibroblast |
| Rbms3 | 0.33 | 0.00 | 0.00 | 0.46 | 0.17 | 23 | Matrix Fibroblast |
| Zeb2 | 0.43 | 0.00 | 0.00 | 0.68 | 0.31 | 23 | Matrix Fibroblast |
| Brd8 | 0.44 | 0.00 | 0.00 | 0.49 | 0.19 | 23 | Matrix Fibroblast |
| Larp7 | 0.43 | 0.00 | 0.00 | 0.45 | 0.17 | 23 | Matrix Fibroblast |
| Gm6625 | 0.47 | 0.00 | 0.00 | 0.57 | 0.25 | 23 | Matrix Fibroblast |
| 2310022B05Rik | 0.37 | 0.00 | 0.00 | 0.29 | 0.08 | 23 | Matrix Fibroblast |
| Mylk | 0.37 | 0.00 | 0.00 | 0.64 | 0.27 | 23 | Matrix Fibroblast |
| Pbdc1 | 0.43 | 0.00 | 0.00 | 0.37 | 0.13 | 23 | Matrix Fibroblast |
| Eny2 | 0.43 | 0.00 | 0.00 | 0.55 | 0.23 | 23 | Matrix Fibroblast |
| Hmgb3 | 0.40 | 0.00 | 0.00 | 0.41 | 0.15 | 23 | Matrix Fibroblast |
| Antxr1 | 0.32 | 0.00 | 0.00 | 0.36 | 0.12 | 23 | Matrix Fibroblast |
| Calm2 | 0.58 | 0.00 | 0.00 | 0.91 | 0.64 | 23 | Matrix Fibroblast |
| Fkbp2 | 0.36 | 0.00 | 0.00 | 0.47 | 0.18 | 23 | Matrix Fibroblast |
| Cdk4 | 0.42 | 0.00 | 0.00 | 0.56 | 0.24 | 23 | Matrix Fibroblast |
| Vkorc1 | 0.33 | 0.00 | 0.00 | 0.32 | 0.10 | 23 | Matrix Fibroblast |
| Rpa3 | 0.42 | 0.00 | 0.00 | 0.37 | 0.13 | 23 | Matrix Fibroblast |
| Tbx5 | 0.26 | 0.00 | 0.00 | 0.28 | 0.08 | 23 | Matrix Fibroblast |
| Lsm2 | 0.34 | 0.00 | 0.00 | 0.29 | 0.09 | 23 | Matrix Fibroblast |
| Stub1 | 0.49 | 0.00 | 0.00 | 0.52 | 0.22 | 23 | Matrix Fibroblast |
| Lhfp | 0.35 | 0.00 | 0.00 | 0.42 | 0.15 | 23 | Matrix Fibroblast |
| G0s2 | 0.27 | 0.00 | 0.00 | 0.39 | 0.13 | 23 | Matrix Fibroblast |
| Fam92a | 0.35 | 0.00 | 0.00 | 0.31 | 0.10 | 23 | Matrix Fibroblast |
| Phlda1 | 0.58 | 0.00 | 0.00 | 0.49 | 0.20 | 23 | Matrix Fibroblast |
| Rab13 | 0.29 | 0.00 | 0.00 | 0.30 | 0.09 | 23 | Matrix Fibroblast |
| Ilf2 | 0.41 | 0.00 | 0.00 | 0.48 | 0.19 | 23 | Matrix Fibroblast |
| Cbx1 | 0.45 | 0.00 | 0.00 | 0.78 | 0.41 | 23 | Matrix Fibroblast |
| Cep57 | 0.38 | 0.00 | 0.00 | 0.33 | 0.11 | 23 | Matrix Fibroblast |
| Ncaph2 | 0.33 | 0.00 | 0.00 | 0.28 | 0.09 | 23 | Matrix Fibroblast |
| Col6a1 | 0.33 | 0.00 | 0.00 | 0.49 | 0.19 | 23 | Matrix Fibroblast |
| Nptn | 0.39 | 0.00 | 0.00 | 0.51 | 0.21 | 23 | Matrix Fibroblast |
| Fmr1 | 0.38 | 0.00 | 0.00 | 0.39 | 0.15 | 23 | Matrix Fibroblast |
| Tsen34 | 0.31 | 0.00 | 0.00 | 0.38 | 0.14 | 23 | Matrix Fibroblast |
| Bub3 | 0.41 | 0.00 | 0.00 | 0.38 | 0.14 | 23 | Matrix Fibroblast |
| Nfic | 0.42 | 0.00 | 0.00 | 0.56 | 0.24 | 23 | Matrix Fibroblast |
| Ccdc82 | 0.34 | 0.00 | 0.00 | 0.25 | 0.07 | 23 | Matrix Fibroblast |
| Cpq | 0.34 | 0.00 | 0.00 | 0.28 | 0.09 | 23 | Matrix Fibroblast |
| Snrpf | 0.43 | 0.00 | 0.00 | 0.61 | 0.28 | 23 | Matrix Fibroblast |
| Ptges3 | 0.38 | 0.00 | 0.00 | 0.56 | 0.25 | 23 | Matrix Fibroblast |
| Rnaseh2b | 0.26 | 0.00 | 0.00 | 0.27 | 0.08 | 23 | Matrix Fibroblast |
| Ercc6l2 | 0.34 | 0.00 | 0.00 | 0.27 | 0.08 | 23 | Matrix Fibroblast |
| Tpm4 | 0.43 | 0.00 | 0.00 | 0.85 | 0.49 | 23 | Matrix Fibroblast |
| Gm9833 | 0.40 | 0.00 | 0.00 | 0.33 | 0.11 | 23 | Matrix Fibroblast |
| H1f0 | 0.55 | 0.00 | 0.00 | 0.58 | 0.27 | 23 | Matrix Fibroblast |
| Macf1 | 0.56 | 0.00 | 0.00 | 0.81 | 0.52 | 23 | Matrix Fibroblast |
| Tpm1 | 0.40 | 0.00 | 0.00 | 0.86 | 0.52 | 23 | Matrix Fibroblast |
| Nudt21 | 0.29 | 0.00 | 0.00 | 0.36 | 0.12 | 23 | Matrix Fibroblast |
| Hspa14 | 0.31 | 0.00 | 0.00 | 0.32 | 0.10 | 23 | Matrix Fibroblast |
| Kpnb1 | 0.42 | 0.00 | 0.00 | 0.46 | 0.19 | 23 | Matrix Fibroblast |
| Ccdc88a | 0.47 | 0.00 | 0.00 | 0.55 | 0.24 | 23 | Matrix Fibroblast |
| Smc3 | 0.48 | 0.00 | 0.00 | 0.70 | 0.38 | 23 | Matrix Fibroblast |
| Tns3 | 0.33 | 0.00 | 0.00 | 0.32 | 0.11 | 23 | Matrix Fibroblast |
| Acat1 | 0.41 | 0.00 | 0.00 | 0.44 | 0.18 | 23 | Matrix Fibroblast |
| Mpdz | 0.31 | 0.00 | 0.00 | 0.26 | 0.08 | 23 | Matrix Fibroblast |
| Anapc13 | 0.33 | 0.00 | 0.00 | 0.50 | 0.21 | 23 | Matrix Fibroblast |
| Spats2l | 0.30 | 0.00 | 0.00 | 0.28 | 0.09 | 23 | Matrix Fibroblast |
| Ift74 | 0.33 | 0.00 | 0.00 | 0.30 | 0.10 | 23 | Matrix Fibroblast |
| Ikbip | 0.34 | 0.00 | 0.00 | 0.26 | 0.08 | 23 | Matrix Fibroblast |
| Fkbp7 | 0.28 | 0.00 | 0.00 | 0.37 | 0.13 | 23 | Matrix Fibroblast |
| Pgp | 0.31 | 0.00 | 0.00 | 0.26 | 0.08 | 23 | Matrix Fibroblast |
| Gm6104 | 0.33 | 0.00 | 0.00 | 0.35 | 0.13 | 23 | Matrix Fibroblast |
| AI597468 | 0.38 | 0.00 | 0.00 | 0.27 | 0.08 | 23 | Matrix Fibroblast |
| Hsd11b1 | 0.35 | 0.00 | 0.00 | 0.29 | 0.09 | 23 | Matrix Fibroblast |
| Ankrd32 | 0.36 | 0.00 | 0.00 | 0.31 | 0.11 | 23 | Matrix Fibroblast |
| Morf4l2 | 0.37 | 0.00 | 0.00 | 0.51 | 0.22 | 23 | Matrix Fibroblast |
| Ndufaf2 | 0.32 | 0.00 | 0.00 | 0.34 | 0.12 | 23 | Matrix Fibroblast |
| Smim4 | 0.32 | 0.00 | 0.00 | 0.35 | 0.12 | 23 | Matrix Fibroblast |
| Mrpl13 | 0.36 | 0.00 | 0.00 | 0.33 | 0.12 | 23 | Matrix Fibroblast |
| Nudcd2 | 0.33 | 0.00 | 0.00 | 0.31 | 0.10 | 23 | Matrix Fibroblast |
| Zyx | 0.34 | 0.00 | 0.00 | 0.48 | 0.20 | 23 | Matrix Fibroblast |
| Apip | 0.28 | 0.00 | 0.00 | 0.26 | 0.08 | 23 | Matrix Fibroblast |
| Cdc16 | 0.29 | 0.00 | 0.00 | 0.41 | 0.16 | 23 | Matrix Fibroblast |
| Leo1 | 0.29 | 0.00 | 0.00 | 0.35 | 0.12 | 23 | Matrix Fibroblast |
| Ext1 | 0.29 | 0.00 | 0.00 | 0.26 | 0.08 | 23 | Matrix Fibroblast |
| Atpif1 | 0.45 | 0.00 | 0.00 | 0.86 | 0.53 | 23 | Matrix Fibroblast |
| Cd81 | 0.44 | 0.00 | 0.00 | 0.74 | 0.41 | 23 | Matrix Fibroblast |
| Kdelr2 | 0.37 | 0.00 | 0.00 | 0.45 | 0.19 | 23 | Matrix Fibroblast |
| Pmepa1 | 0.33 | 0.00 | 0.00 | 0.32 | 0.11 | 23 | Matrix Fibroblast |
| Pole3 | 0.31 | 0.00 | 0.00 | 0.31 | 0.11 | 23 | Matrix Fibroblast |
| Mfge8 | 0.45 | 0.00 | 0.00 | 0.40 | 0.16 | 23 | Matrix Fibroblast |
| Nedd4 | 0.41 | 0.00 | 0.00 | 0.87 | 0.53 | 23 | Matrix Fibroblast |
| Eif4e | 0.33 | 0.00 | 0.00 | 0.30 | 0.10 | 23 | Matrix Fibroblast |
| Slirp | 0.30 | 0.00 | 0.00 | 0.51 | 0.22 | 23 | Matrix Fibroblast |
| Hn1 | 0.44 | 0.00 | 0.00 | 0.46 | 0.20 | 23 | Matrix Fibroblast |
| Rfc1 | 0.46 | 0.00 | 0.00 | 0.47 | 0.20 | 23 | Matrix Fibroblast |
| Calm3 | 0.40 | 0.00 | 0.00 | 0.45 | 0.19 | 23 | Matrix Fibroblast |
| Ssr1 | 0.37 | 0.00 | 0.00 | 0.54 | 0.25 | 23 | Matrix Fibroblast |
| Pdzd11 | 0.31 | 0.00 | 0.00 | 0.28 | 0.09 | 23 | Matrix Fibroblast |
| G3bp1 | 0.36 | 0.00 | 0.00 | 0.42 | 0.17 | 23 | Matrix Fibroblast |
| Srrt | 0.33 | 0.00 | 0.00 | 0.47 | 0.20 | 23 | Matrix Fibroblast |
| Specc1l | 0.37 | 0.00 | 0.00 | 0.33 | 0.12 | 23 | Matrix Fibroblast |
| Tead1 | 0.34 | 0.00 | 0.00 | 0.36 | 0.14 | 23 | Matrix Fibroblast |
| Ywhaq | 0.38 | 0.00 | 0.00 | 0.70 | 0.37 | 23 | Matrix Fibroblast |
| Hint1 | 0.42 | 0.00 | 0.00 | 0.78 | 0.46 | 23 | Matrix Fibroblast |
| Smarcc1 | 0.36 | 0.00 | 0.00 | 0.41 | 0.17 | 23 | Matrix Fibroblast |
| Cpm | 0.51 | 0.00 | 0.00 | 0.42 | 0.18 | 23 | Matrix Fibroblast |
| Mrfap1 | 0.36 | 0.00 | 0.00 | 0.66 | 0.33 | 23 | Matrix Fibroblast |
| Ppic | 0.35 | 0.00 | 0.00 | 0.73 | 0.38 | 23 | Matrix Fibroblast |
| Fkbp3 | 0.43 | 0.00 | 0.00 | 0.59 | 0.30 | 23 | Matrix Fibroblast |
| Srsf2 | 0.40 | 0.00 | 0.00 | 0.71 | 0.38 | 23 | Matrix Fibroblast |
| Matr3 | 0.43 | 0.00 | 0.00 | 0.73 | 0.41 | 23 | Matrix Fibroblast |
| Pdlim2 | 0.35 | 0.00 | 0.00 | 0.42 | 0.18 | 23 | Matrix Fibroblast |
| Cbx6 | 0.28 | 0.00 | 0.00 | 0.30 | 0.11 | 23 | Matrix Fibroblast |
| Clic4 | 0.33 | 0.00 | 0.00 | 0.60 | 0.29 | 23 | Matrix Fibroblast |
| Pdap1 | 0.43 | 0.00 | 0.00 | 0.77 | 0.45 | 23 | Matrix Fibroblast |
| Cep290 | 0.32 | 0.00 | 0.00 | 0.28 | 0.10 | 23 | Matrix Fibroblast |
| Erh | 0.37 | 0.00 | 0.00 | 0.59 | 0.28 | 23 | Matrix Fibroblast |
| Myef2 | 0.38 | 0.00 | 0.00 | 0.38 | 0.16 | 23 | Matrix Fibroblast |
| Ctdspl2 | 0.34 | 0.00 | 0.00 | 0.31 | 0.11 | 23 | Matrix Fibroblast |
| Dkc1 | 0.33 | 0.00 | 0.00 | 0.31 | 0.12 | 23 | Matrix Fibroblast |
| Blmh | 0.34 | 0.00 | 0.00 | 0.41 | 0.17 | 23 | Matrix Fibroblast |
| Gclm | 0.39 | 0.00 | 0.00 | 0.74 | 0.42 | 23 | Matrix Fibroblast |
| Anapc11 | 0.33 | 0.00 | 0.00 | 0.41 | 0.17 | 23 | Matrix Fibroblast |
| Hnrnph1 | 0.40 | 0.00 | 0.00 | 0.69 | 0.38 | 23 | Matrix Fibroblast |
| Gm12728 | 0.32 | 0.00 | 0.00 | 0.41 | 0.17 | 23 | Matrix Fibroblast |
| Gm9242 | 0.39 | 0.00 | 0.00 | 0.50 | 0.24 | 23 | Matrix Fibroblast |
| Slc25a4 | 0.35 | 0.00 | 0.00 | 0.82 | 0.50 | 23 | Matrix Fibroblast |
| Mmp14 | 0.35 | 0.00 | 0.00 | 0.54 | 0.25 | 23 | Matrix Fibroblast |
| Htatsf1 | 0.31 | 0.00 | 0.00 | 0.55 | 0.26 | 23 | Matrix Fibroblast |
| Rnaseh2c | 0.27 | 0.00 | 0.00 | 0.29 | 0.11 | 23 | Matrix Fibroblast |
| Smarca5 | 0.35 | 0.00 | 0.00 | 0.60 | 0.30 | 23 | Matrix Fibroblast |
| Timp2 | 0.27 | 0.00 | 0.00 | 0.48 | 0.21 | 23 | Matrix Fibroblast |
| Nsmce1 | 0.25 | 0.00 | 0.00 | 0.35 | 0.14 | 23 | Matrix Fibroblast |
| Tgfb2 | 0.46 | 0.00 | 0.00 | 0.35 | 0.14 | 23 | Matrix Fibroblast |
| Nrep | 0.28 | 0.00 | 0.00 | 0.77 | 0.43 | 23 | Matrix Fibroblast |
| Gm8991 | 0.32 | 0.00 | 0.00 | 0.50 | 0.23 | 23 | Matrix Fibroblast |
| Hp1bp3 | 0.41 | 0.00 | 0.00 | 0.76 | 0.44 | 23 | Matrix Fibroblast |
| Fgfr1 | 0.29 | 0.00 | 0.00 | 0.29 | 0.11 | 23 | Matrix Fibroblast |
| Dnajc10 | 0.38 | 0.00 | 0.00 | 0.36 | 0.15 | 23 | Matrix Fibroblast |
| Hnrnpa3 | 0.39 | 0.00 | 0.00 | 0.78 | 0.46 | 23 | Matrix Fibroblast |
| Ube2e3 | 0.29 | 0.00 | 0.00 | 0.35 | 0.14 | 23 | Matrix Fibroblast |
| Tagln2 | 0.40 | 0.00 | 0.00 | 0.76 | 0.45 | 23 | Matrix Fibroblast |
| F2r | 0.33 | 0.00 | 0.00 | 0.36 | 0.14 | 23 | Matrix Fibroblast |
| Ndufa5 | 0.31 | 0.00 | 0.00 | 0.54 | 0.26 | 23 | Matrix Fibroblast |
| Rbm3 | 0.40 | 0.00 | 0.00 | 0.70 | 0.41 | 23 | Matrix Fibroblast |
| Casp8ap2 | 0.37 | 0.00 | 0.00 | 0.40 | 0.17 | 23 | Matrix Fibroblast |
| Srsf3 | 0.40 | 0.00 | 0.00 | 0.76 | 0.45 | 23 | Matrix Fibroblast |
| Rnf168 | 0.27 | 0.00 | 0.00 | 0.34 | 0.14 | 23 | Matrix Fibroblast |
| Epb4.1l2 | 0.34 | 0.00 | 0.00 | 0.43 | 0.19 | 23 | Matrix Fibroblast |
| Gm12355 | 0.33 | 0.00 | 0.00 | 0.54 | 0.26 | 23 | Matrix Fibroblast |
| Gm6793 | 0.37 | 0.00 | 0.00 | 0.58 | 0.30 | 23 | Matrix Fibroblast |
| Ctcf | 0.43 | 0.00 | 0.00 | 0.49 | 0.23 | 23 | Matrix Fibroblast |
| Rcn2 | 0.26 | 0.00 | 0.00 | 0.37 | 0.15 | 23 | Matrix Fibroblast |
| Tmsb10 | 0.41 | 0.00 | 0.00 | 0.98 | 0.86 | 23 | Matrix Fibroblast |
| Nudc | 0.32 | 0.00 | 0.00 | 0.45 | 0.21 | 23 | Matrix Fibroblast |
| Actn1 | 0.33 | 0.00 | 0.00 | 0.48 | 0.23 | 23 | Matrix Fibroblast |
| Cnep1r1 | 0.25 | 0.00 | 0.00 | 0.30 | 0.11 | 23 | Matrix Fibroblast |
| Nenf | 0.25 | 0.00 | 0.00 | 0.42 | 0.18 | 23 | Matrix Fibroblast |
| Hdgf | 0.29 | 0.00 | 0.00 | 0.52 | 0.25 | 23 | Matrix Fibroblast |
| Pros1 | 0.26 | 0.00 | 0.00 | 0.28 | 0.10 | 23 | Matrix Fibroblast |
| Hnrnpab | 0.35 | 0.00 | 0.00 | 0.62 | 0.33 | 23 | Matrix Fibroblast |
| Thoc7 | 0.35 | 0.00 | 0.00 | 0.61 | 0.32 | 23 | Matrix Fibroblast |
| Polr2m | 0.25 | 0.00 | 0.00 | 0.39 | 0.17 | 23 | Matrix Fibroblast |
| Gm10108 | 0.30 | 0.00 | 0.00 | 0.33 | 0.14 | 23 | Matrix Fibroblast |
| Snrpg | 0.30 | 0.00 | 0.00 | 0.72 | 0.39 | 23 | Matrix Fibroblast |
| Zranb2 | 0.31 | 0.00 | 0.00 | 0.53 | 0.25 | 23 | Matrix Fibroblast |
| Sumo2 | 0.33 | 0.00 | 0.00 | 0.59 | 0.30 | 23 | Matrix Fibroblast |
| Tsix | 0.34 | 0.00 | 0.00 | 0.41 | 0.18 | 23 | Matrix Fibroblast |
| A630007B06Rik | 0.34 | 0.00 | 0.00 | 0.43 | 0.20 | 23 | Matrix Fibroblast |
| Brd3 | 0.32 | 0.00 | 0.00 | 0.53 | 0.25 | 23 | Matrix Fibroblast |
| Cnn2 | 0.35 | 0.00 | 0.00 | 0.70 | 0.39 | 23 | Matrix Fibroblast |
| Magt1 | 0.27 | 0.00 | 0.00 | 0.44 | 0.20 | 23 | Matrix Fibroblast |
| Msl3 | 0.27 | 0.00 | 0.00 | 0.35 | 0.14 | 23 | Matrix Fibroblast |
| Rad50 | 0.30 | 0.00 | 0.00 | 0.43 | 0.19 | 23 | Matrix Fibroblast |
| Sec13 | 0.30 | 0.00 | 0.00 | 0.39 | 0.17 | 23 | Matrix Fibroblast |
| Serpinh1 | 0.34 | 0.00 | 0.00 | 0.69 | 0.39 | 23 | Matrix Fibroblast |
| Pbx1 | 0.32 | 0.00 | 0.00 | 0.43 | 0.20 | 23 | Matrix Fibroblast |
| Arl6ip1 | 0.65 | 0.00 | 0.00 | 0.77 | 0.52 | 23 | Matrix Fibroblast |
| Hsp90b1 | 0.35 | 0.00 | 0.00 | 0.94 | 0.76 | 23 | Matrix Fibroblast |
| Mxra8 | 0.28 | 0.00 | 0.00 | 0.38 | 0.16 | 23 | Matrix Fibroblast |
| Gm10053 | 0.34 | 0.00 | 0.00 | 0.46 | 0.22 | 23 | Matrix Fibroblast |
| Sarnp | 0.29 | 0.00 | 0.00 | 0.52 | 0.25 | 23 | Matrix Fibroblast |
| Gmps | 0.32 | 0.00 | 0.00 | 0.29 | 0.11 | 23 | Matrix Fibroblast |
| Anapc5 | 0.25 | 0.00 | 0.00 | 0.41 | 0.18 | 23 | Matrix Fibroblast |
| Mrpl18 | 0.28 | 0.00 | 0.00 | 0.45 | 0.21 | 23 | Matrix Fibroblast |
| Gm5641 | 0.33 | 0.00 | 0.00 | 0.66 | 0.37 | 23 | Matrix Fibroblast |
| Parp2 | 0.25 | 0.00 | 0.00 | 0.28 | 0.10 | 23 | Matrix Fibroblast |
| Atp5o | 0.31 | 0.00 | 0.00 | 0.65 | 0.35 | 23 | Matrix Fibroblast |
| Ywhah | 0.27 | 0.00 | 0.00 | 0.53 | 0.26 | 23 | Matrix Fibroblast |
| Igfbp4 | 0.30 | 0.00 | 0.00 | 0.51 | 0.25 | 23 | Matrix Fibroblast |
| Pds5b | 0.32 | 0.00 | 0.00 | 0.31 | 0.13 | 23 | Matrix Fibroblast |
| Naa50 | 0.31 | 0.00 | 0.00 | 0.32 | 0.13 | 23 | Matrix Fibroblast |
| Snrpd2 | 0.25 | 0.00 | 0.00 | 0.51 | 0.25 | 23 | Matrix Fibroblast |
| Uba2 | 0.25 | 0.00 | 0.00 | 0.29 | 0.11 | 23 | Matrix Fibroblast |
| Golim4 | 0.29 | 0.00 | 0.00 | 0.43 | 0.20 | 23 | Matrix Fibroblast |
| Rbbp4 | 0.27 | 0.00 | 0.00 | 0.43 | 0.20 | 23 | Matrix Fibroblast |
| Nhp2l1 | 0.29 | 0.00 | 0.00 | 0.42 | 0.19 | 23 | Matrix Fibroblast |
| Tagln | 0.36 | 0.00 | 0.00 | 0.34 | 0.15 | 23 | Matrix Fibroblast |
| Cox20 | 0.26 | 0.00 | 0.00 | 0.40 | 0.18 | 23 | Matrix Fibroblast |
| Wbp5 | 0.26 | 0.00 | 0.00 | 0.80 | 0.48 | 23 | Matrix Fibroblast |
| Purb | 0.32 | 0.00 | 0.00 | 0.66 | 0.37 | 23 | Matrix Fibroblast |
| Tcf12 | 0.32 | 0.00 | 0.00 | 0.38 | 0.17 | 23 | Matrix Fibroblast |
| Cycs | 0.31 | 0.00 | 0.00 | 0.44 | 0.22 | 23 | Matrix Fibroblast |
| Nfyb | 0.27 | 0.00 | 0.00 | 0.29 | 0.12 | 23 | Matrix Fibroblast |
| Syncrip | 0.28 | 0.00 | 0.00 | 0.49 | 0.25 | 23 | Matrix Fibroblast |
| Zfhx3 | 0.32 | 0.00 | 0.00 | 0.37 | 0.17 | 23 | Matrix Fibroblast |
| Mfap2 | 0.27 | 0.00 | 0.00 | 0.65 | 0.35 | 23 | Matrix Fibroblast |
| Siva1 | 0.31 | 0.00 | 0.00 | 0.33 | 0.15 | 23 | Matrix Fibroblast |
| Tmem256 | 0.27 | 0.00 | 0.00 | 0.61 | 0.33 | 23 | Matrix Fibroblast |
| Supt16 | 0.30 | 0.00 | 0.00 | 0.49 | 0.25 | 23 | Matrix Fibroblast |
| Pcdhga9 | 0.25 | 0.00 | 0.00 | 0.38 | 0.18 | 23 | Matrix Fibroblast |
| Srsf7 | 0.28 | 0.00 | 0.00 | 0.35 | 0.16 | 23 | Matrix Fibroblast |
| Hnrnpdl | 0.31 | 0.00 | 0.00 | 0.64 | 0.37 | 23 | Matrix Fibroblast |
| Hnrnpa1 | 0.30 | 0.00 | 0.00 | 0.52 | 0.27 | 23 | Matrix Fibroblast |
| Hadh | 0.28 | 0.00 | 0.00 | 0.37 | 0.17 | 23 | Matrix Fibroblast |
| Taf9 | 0.28 | 0.00 | 0.00 | 0.39 | 0.18 | 23 | Matrix Fibroblast |
| Hdac2 | 0.26 | 0.00 | 0.00 | 0.35 | 0.15 | 23 | Matrix Fibroblast |
| Mbnl2 | 0.29 | 0.00 | 0.00 | 0.62 | 0.35 | 23 | Matrix Fibroblast |
| Lamc1 | 0.26 | 0.00 | 0.00 | 0.35 | 0.16 | 23 | Matrix Fibroblast |
| Smc5 | 0.29 | 0.00 | 0.00 | 0.36 | 0.16 | 23 | Matrix Fibroblast |
| Gtf2h5 | 0.28 | 0.00 | 0.00 | 0.53 | 0.28 | 23 | Matrix Fibroblast |
| Cox7a2 | 0.32 | 0.00 | 0.00 | 0.76 | 0.48 | 23 | Matrix Fibroblast |
| Nin | 0.27 | 0.00 | 0.00 | 0.32 | 0.14 | 23 | Matrix Fibroblast |
| Cetn3 | 0.30 | 0.00 | 0.00 | 0.56 | 0.31 | 23 | Matrix Fibroblast |
| Xist | 0.38 | 0.00 | 0.00 | 0.76 | 0.51 | 23 | Matrix Fibroblast |
| Pdcd5 | 0.25 | 0.00 | 0.00 | 0.53 | 0.28 | 23 | Matrix Fibroblast |
| Tmed3 | 0.25 | 0.00 | 0.00 | 0.29 | 0.13 | 23 | Matrix Fibroblast |
| Tceb1 | 0.27 | 0.00 | 0.00 | 0.45 | 0.23 | 23 | Matrix Fibroblast |
| Pnrc2 | 0.29 | 0.00 | 0.00 | 0.39 | 0.19 | 23 | Matrix Fibroblast |
| Hnrnpu | 0.25 | 0.00 | 0.00 | 0.88 | 0.64 | 23 | Matrix Fibroblast |
| Pde5a | 0.33 | 0.00 | 0.00 | 0.28 | 0.12 | 23 | Matrix Fibroblast |
| Rif1 | 0.30 | 0.00 | 0.00 | 0.35 | 0.17 | 23 | Matrix Fibroblast |
| P4hb | 0.28 | 0.00 | 0.00 | 0.67 | 0.40 | 23 | Matrix Fibroblast |
| Tceb2 | 0.31 | 0.00 | 0.00 | 0.73 | 0.47 | 23 | Matrix Fibroblast |
| Suz12 | 0.32 | 0.00 | 0.00 | 0.29 | 0.13 | 23 | Matrix Fibroblast |
| Gm20091 | 0.27 | 0.00 | 0.00 | 0.43 | 0.22 | 23 | Matrix Fibroblast |
| Hnrnpa2b1 | 0.30 | 0.00 | 0.00 | 0.91 | 0.72 | 23 | Matrix Fibroblast |
| H3f3b | 0.31 | 0.00 | 0.00 | 0.98 | 0.89 | 23 | Matrix Fibroblast |
| Mat2a | 0.26 | 0.00 | 0.00 | 0.50 | 0.27 | 23 | Matrix Fibroblast |
| Cox7c | 0.28 | 0.00 | 0.00 | 0.87 | 0.63 | 23 | Matrix Fibroblast |
| Hnrnpa0 | 0.25 | 0.00 | 0.00 | 0.42 | 0.21 | 23 | Matrix Fibroblast |
| Hist1h2bc | 0.52 | 0.00 | 0.00 | 0.29 | 0.14 | 23 | Matrix Fibroblast |
| Srrm1 | 0.28 | 0.00 | 0.00 | 0.73 | 0.48 | 23 | Matrix Fibroblast |
| Sec61g | 0.32 | 0.00 | 0.00 | 0.57 | 0.35 | 23 | Matrix Fibroblast |
| Syne2 | 0.25 | 0.00 | 0.00 | 0.30 | 0.14 | 23 | Matrix Fibroblast |
| Fermt2 | 0.29 | 0.00 | 0.00 | 0.67 | 0.39 | 23 | Matrix Fibroblast |
| Gm5611 | 0.26 | 0.00 | 0.00 | 0.57 | 0.36 | 23 | Matrix Fibroblast |
| Morf4l1 | 0.26 | 0.00 | 0.00 | 0.77 | 0.55 | 23 | Matrix Fibroblast |
| Csrp2 | 0.33 | 0.00 | 0.00 | 0.32 | 0.17 | 23 | Matrix Fibroblast |
| Top2a | 2.16 | 0.00 | 0.00 | 0.90 | 0.11 | 24 | Matrix Fibroblast |
| Cenpf | 2.05 | 0.00 | 0.00 | 0.76 | 0.08 | 24 | Matrix Fibroblast |
| Mki67 | 2.00 | 0.00 | 0.00 | 0.95 | 0.12 | 24 | Matrix Fibroblast |
| Prc1 | 1.96 | 0.00 | 0.00 | 0.78 | 0.06 | 24 | Matrix Fibroblast |
| 2810417H13Rik | 1.94 | 0.00 | 0.00 | 0.87 | 0.08 | 24 | Matrix Fibroblast |
| Htra1 | 1.84 | 0.00 | 0.00 | 0.85 | 0.08 | 24 | Matrix Fibroblast |
| Cenpe | 1.80 | 0.00 | 0.00 | 0.72 | 0.08 | 24 | Matrix Fibroblast |
| Nusap1 | 1.73 | 0.00 | 0.00 | 0.74 | 0.05 | 24 | Matrix Fibroblast |
| Ube2c | 1.73 | 0.00 | 0.00 | 0.64 | 0.05 | 24 | Matrix Fibroblast |
| Tpx2 | 1.61 | 0.00 | 0.00 | 0.72 | 0.05 | 24 | Matrix Fibroblast |
| Hmmr | 1.61 | 0.00 | 0.00 | 0.66 | 0.04 | 24 | Matrix Fibroblast |
| Spc25 | 1.60 | 0.00 | 0.00 | 0.68 | 0.05 | 24 | Matrix Fibroblast |
| Cks2 | 1.57 | 0.00 | 0.00 | 0.71 | 0.08 | 24 | Matrix Fibroblast |
| Prss35 | 1.52 | 0.00 | 0.00 | 0.68 | 0.07 | 24 | Matrix Fibroblast |
| Ckap2l | 1.51 | 0.00 | 0.00 | 0.65 | 0.04 | 24 | Matrix Fibroblast |
| Birc5 | 1.51 | 0.00 | 0.00 | 0.71 | 0.06 | 24 | Matrix Fibroblast |
| Ccnb2 | 1.46 | 0.00 | 0.00 | 0.67 | 0.06 | 24 | Matrix Fibroblast |
| Diap3 | 1.43 | 0.00 | 0.00 | 0.68 | 0.06 | 24 | Matrix Fibroblast |
| Cdk1 | 1.43 | 0.00 | 0.00 | 0.64 | 0.05 | 24 | Matrix Fibroblast |
| Ncapg | 1.42 | 0.00 | 0.00 | 0.65 | 0.04 | 24 | Matrix Fibroblast |
| Kif23 | 1.39 | 0.00 | 0.00 | 0.61 | 0.05 | 24 | Matrix Fibroblast |
| Sgol2 | 1.37 | 0.00 | 0.00 | 0.57 | 0.04 | 24 | Matrix Fibroblast |
| Kif11 | 1.37 | 0.00 | 0.00 | 0.63 | 0.04 | 24 | Matrix Fibroblast |
| Ckap2 | 1.34 | 0.00 | 0.00 | 0.57 | 0.04 | 24 | Matrix Fibroblast |
| P2ry14 | 1.32 | 0.00 | 0.00 | 0.73 | 0.08 | 24 | Matrix Fibroblast |
| Anln | 1.31 | 0.00 | 0.00 | 0.58 | 0.03 | 24 | Matrix Fibroblast |
| RP23-45G16.5 | 1.30 | 0.00 | 0.00 | 0.60 | 0.05 | 24 | Matrix Fibroblast |
| Ccna2 | 1.25 | 0.00 | 0.00 | 0.56 | 0.04 | 24 | Matrix Fibroblast |
| Rrm2 | 1.25 | 0.00 | 0.00 | 0.55 | 0.05 | 24 | Matrix Fibroblast |
| Kif15 | 1.24 | 0.00 | 0.00 | 0.56 | 0.04 | 24 | Matrix Fibroblast |
| Casc5 | 1.22 | 0.00 | 0.00 | 0.53 | 0.04 | 24 | Matrix Fibroblast |
| Esco2 | 1.22 | 0.00 | 0.00 | 0.52 | 0.03 | 24 | Matrix Fibroblast |
| Egfem1 | 1.20 | 0.00 | 0.00 | 0.62 | 0.05 | 24 | Matrix Fibroblast |
| Pbk | 1.19 | 0.00 | 0.00 | 0.52 | 0.03 | 24 | Matrix Fibroblast |
| Spc24 | 1.18 | 0.00 | 0.00 | 0.59 | 0.05 | 24 | Matrix Fibroblast |
| Ndc80 | 1.17 | 0.00 | 0.00 | 0.51 | 0.04 | 24 | Matrix Fibroblast |
| Cdca3 | 1.17 | 0.00 | 0.00 | 0.52 | 0.03 | 24 | Matrix Fibroblast |
| Mis18bp1 | 1.13 | 0.00 | 0.00 | 0.47 | 0.03 | 24 | Matrix Fibroblast |
| Nuf2 | 1.10 | 0.00 | 0.00 | 0.52 | 0.03 | 24 | Matrix Fibroblast |
| Aspm | 1.07 | 0.00 | 0.00 | 0.46 | 0.03 | 24 | Matrix Fibroblast |
| Ccnb1 | 1.00 | 0.00 | 0.00 | 0.40 | 0.02 | 24 | Matrix Fibroblast |
| Bub1 | 0.98 | 0.00 | 0.00 | 0.42 | 0.02 | 24 | Matrix Fibroblast |
| Depdc1a | 0.94 | 0.00 | 0.00 | 0.39 | 0.02 | 24 | Matrix Fibroblast |
| Ect2 | 0.92 | 0.00 | 0.00 | 0.42 | 0.02 | 24 | Matrix Fibroblast |
| Mxd3 | 0.80 | 0.00 | 0.00 | 0.35 | 0.02 | 24 | Matrix Fibroblast |
| Shcbp1 | 0.71 | 0.00 | 0.00 | 0.31 | 0.01 | 24 | Matrix Fibroblast |
| Pdgfra | 1.26 | 0.00 | 0.00 | 0.61 | 0.06 | 24 | Matrix Fibroblast |
| Racgap1 | 1.01 | 0.00 | 0.00 | 0.49 | 0.04 | 24 | Matrix Fibroblast |
| Fam64a | 0.83 | 0.00 | 0.00 | 0.36 | 0.02 | 24 | Matrix Fibroblast |
| Neil3 | 0.83 | 0.00 | 0.00 | 0.39 | 0.02 | 24 | Matrix Fibroblast |
| Ska1 | 0.74 | 0.00 | 0.00 | 0.35 | 0.02 | 24 | Matrix Fibroblast |
| Incenp | 1.41 | 0.00 | 0.00 | 0.69 | 0.08 | 24 | Matrix Fibroblast |
| Prr11 | 0.87 | 0.00 | 0.00 | 0.35 | 0.02 | 24 | Matrix Fibroblast |
| Cdca8 | 1.15 | 0.00 | 0.00 | 0.55 | 0.05 | 24 | Matrix Fibroblast |
| Cdca2 | 0.79 | 0.00 | 0.00 | 0.35 | 0.02 | 24 | Matrix Fibroblast |
| C330027C09Rik | 0.96 | 0.00 | 0.00 | 0.42 | 0.03 | 24 | Matrix Fibroblast |
| Knstrn | 1.22 | 0.00 | 0.00 | 0.47 | 0.04 | 24 | Matrix Fibroblast |
| Tacc3 | 1.13 | 0.00 | 0.00 | 0.57 | 0.05 | 24 | Matrix Fibroblast |
| Cenpp | 0.70 | 0.00 | 0.00 | 0.33 | 0.02 | 24 | Matrix Fibroblast |
| Smc2 | 1.66 | 0.00 | 0.00 | 0.84 | 0.13 | 24 | Matrix Fibroblast |
| Cdc20 | 1.34 | 0.00 | 0.00 | 0.52 | 0.05 | 24 | Matrix Fibroblast |
| Agt | 1.25 | 0.00 | 0.00 | 0.76 | 0.10 | 24 | Matrix Fibroblast |
| Kif22 | 0.82 | 0.00 | 0.00 | 0.37 | 0.02 | 24 | Matrix Fibroblast |
| Tnc | 1.45 | 0.00 | 0.00 | 0.88 | 0.14 | 24 | Matrix Fibroblast |
| Kif20b | 1.45 | 0.00 | 0.00 | 0.60 | 0.06 | 24 | Matrix Fibroblast |
| Plk1 | 0.69 | 0.00 | 0.00 | 0.30 | 0.02 | 24 | Matrix Fibroblast |
| Cenpa | 1.56 | 0.00 | 0.00 | 0.59 | 0.06 | 24 | Matrix Fibroblast |
| Hist1h2ae | 1.66 | 0.00 | 0.00 | 0.68 | 0.09 | 24 | Matrix Fibroblast |
| Pdlim3 | 1.10 | 0.00 | 0.00 | 0.60 | 0.06 | 24 | Matrix Fibroblast |
| Melk | 0.76 | 0.00 | 0.00 | 0.30 | 0.02 | 24 | Matrix Fibroblast |
| Cd248 | 1.00 | 0.00 | 0.00 | 0.52 | 0.05 | 24 | Matrix Fibroblast |
| Hist1h2ad | 1.81 | 0.00 | 0.00 | 0.72 | 0.10 | 24 | Matrix Fibroblast |
| Cks1b | 1.29 | 0.00 | 0.00 | 0.70 | 0.09 | 24 | Matrix Fibroblast |
| Hist1h2ak | 1.52 | 0.00 | 0.00 | 0.62 | 0.07 | 24 | Matrix Fibroblast |
| Kif2c | 0.64 | 0.00 | 0.00 | 0.28 | 0.02 | 24 | Matrix Fibroblast |
| Aurkb | 0.81 | 0.00 | 0.00 | 0.37 | 0.03 | 24 | Matrix Fibroblast |
| Tenm4 | 0.93 | 0.00 | 0.00 | 0.43 | 0.04 | 24 | Matrix Fibroblast |
| Tubb6 | 1.15 | 0.00 | 0.00 | 0.62 | 0.07 | 24 | Matrix Fibroblast |
| Hist1h2ab | 1.65 | 0.00 | 0.00 | 0.67 | 0.09 | 24 | Matrix Fibroblast |
| Hist1h2ai | 1.69 | 0.00 | 0.00 | 0.69 | 0.10 | 24 | Matrix Fibroblast |
| Ncapd2 | 0.96 | 0.00 | 0.00 | 0.47 | 0.04 | 24 | Matrix Fibroblast |
| Hist1h2ac | 1.77 | 0.00 | 0.00 | 0.71 | 0.10 | 24 | Matrix Fibroblast |
| Hist1h2an | 1.72 | 0.00 | 0.00 | 0.66 | 0.09 | 24 | Matrix Fibroblast |
| Hist1h2ap | 2.15 | 0.00 | 0.00 | 0.81 | 0.15 | 24 | Matrix Fibroblast |
| Kifc1 | 0.63 | 0.00 | 0.00 | 0.29 | 0.02 | 24 | Matrix Fibroblast |
| Tk1 | 0.93 | 0.00 | 0.00 | 0.40 | 0.03 | 24 | Matrix Fibroblast |
| Hist1h2af | 1.66 | 0.00 | 0.00 | 0.66 | 0.09 | 24 | Matrix Fibroblast |
| Hist1h2ah | 1.67 | 0.00 | 0.00 | 0.67 | 0.09 | 24 | Matrix Fibroblast |
| Kif20a | 0.81 | 0.00 | 0.00 | 0.33 | 0.02 | 24 | Matrix Fibroblast |
| Sgol1 | 0.90 | 0.00 | 0.00 | 0.37 | 0.03 | 24 | Matrix Fibroblast |
| Tgfbi | 1.38 | 0.00 | 0.00 | 0.95 | 0.19 | 24 | Matrix Fibroblast |
| Fbxo5 | 1.04 | 0.00 | 0.00 | 0.47 | 0.04 | 24 | Matrix Fibroblast |
| Hist1h2ag | 1.87 | 0.00 | 0.00 | 0.73 | 0.12 | 24 | Matrix Fibroblast |
| Myh11 | 1.00 | 0.00 | 0.00 | 0.74 | 0.10 | 24 | Matrix Fibroblast |
| Dlgap5 | 0.90 | 0.00 | 0.00 | 0.35 | 0.03 | 24 | Matrix Fibroblast |
| Sdc2 | 1.24 | 0.00 | 0.00 | 0.77 | 0.12 | 24 | Matrix Fibroblast |
| Bub1b | 0.76 | 0.00 | 0.00 | 0.33 | 0.02 | 24 | Matrix Fibroblast |
| Hist1h2ao | 2.12 | 0.00 | 0.00 | 0.80 | 0.15 | 24 | Matrix Fibroblast |
| Arhgap11a | 1.04 | 0.00 | 0.00 | 0.44 | 0.04 | 24 | Matrix Fibroblast |
| Cenpm | 0.69 | 0.00 | 0.00 | 0.33 | 0.02 | 24 | Matrix Fibroblast |
| Cenpk | 0.89 | 0.00 | 0.00 | 0.40 | 0.04 | 24 | Matrix Fibroblast |
| Mns1 | 0.89 | 0.00 | 0.00 | 0.43 | 0.04 | 24 | Matrix Fibroblast |
| Hlf | 0.90 | 0.00 | 0.00 | 0.47 | 0.05 | 24 | Matrix Fibroblast |
| Robo2 | 1.04 | 0.00 | 0.00 | 0.64 | 0.09 | 24 | Matrix Fibroblast |
| Zwilch | 0.65 | 0.00 | 0.00 | 0.27 | 0.02 | 24 | Matrix Fibroblast |
| Fbln1 | 0.83 | 0.00 | 0.00 | 0.49 | 0.05 | 24 | Matrix Fibroblast |
| Wnt5a | 1.19 | 0.00 | 0.00 | 0.68 | 0.10 | 24 | Matrix Fibroblast |
| Mad2l1 | 0.69 | 0.00 | 0.00 | 0.32 | 0.02 | 24 | Matrix Fibroblast |
| Ska3 | 0.56 | 0.00 | 0.00 | 0.27 | 0.02 | 24 | Matrix Fibroblast |
| H2afx | 1.18 | 0.00 | 0.00 | 0.63 | 0.09 | 24 | Matrix Fibroblast |
| Olfml2b | 0.95 | 0.00 | 0.00 | 0.53 | 0.06 | 24 | Matrix Fibroblast |
| Vsnl1 | 0.66 | 0.00 | 0.00 | 0.28 | 0.02 | 24 | Matrix Fibroblast |
| Etv1 | 0.95 | 0.00 | 0.00 | 0.56 | 0.07 | 24 | Matrix Fibroblast |
| Ltbp2 | 0.94 | 0.00 | 0.00 | 0.65 | 0.09 | 24 | Matrix Fibroblast |
| Ccdc34 | 1.30 | 0.00 | 0.00 | 0.74 | 0.14 | 24 | Matrix Fibroblast |
| Pde5a | 1.11 | 0.00 | 0.00 | 0.70 | 0.11 | 24 | Matrix Fibroblast |
| Hist1h1b | 1.14 | 0.00 | 0.00 | 0.38 | 0.04 | 24 | Matrix Fibroblast |
| Plk4 | 0.80 | 0.00 | 0.00 | 0.41 | 0.04 | 24 | Matrix Fibroblast |
| Stmn1 | 1.52 | 0.00 | 0.00 | 0.90 | 0.23 | 24 | Matrix Fibroblast |
| Kif4 | 0.77 | 0.00 | 0.00 | 0.31 | 0.03 | 24 | Matrix Fibroblast |
| D17H6S56E-5 | 0.98 | 0.00 | 0.00 | 0.66 | 0.10 | 24 | Matrix Fibroblast |
| Fndc1 | 0.83 | 0.00 | 0.00 | 0.45 | 0.05 | 24 | Matrix Fibroblast |
| Spon2 | 0.92 | 0.00 | 0.00 | 0.51 | 0.06 | 24 | Matrix Fibroblast |
| Wnt11 | 0.67 | 0.00 | 0.00 | 0.30 | 0.02 | 24 | Matrix Fibroblast |
| Fam129a | 1.00 | 0.00 | 0.00 | 0.58 | 0.08 | 24 | Matrix Fibroblast |
| Chn1 | 0.71 | 0.00 | 0.00 | 0.35 | 0.03 | 24 | Matrix Fibroblast |
| Cenph | 0.79 | 0.00 | 0.00 | 0.35 | 0.03 | 24 | Matrix Fibroblast |
| Cdkn2c | 0.85 | 0.00 | 0.00 | 0.40 | 0.04 | 24 | Matrix Fibroblast |
| Cep55 | 0.72 | 0.00 | 0.00 | 0.31 | 0.03 | 24 | Matrix Fibroblast |
| Trip13 | 0.61 | 0.00 | 0.00 | 0.27 | 0.02 | 24 | Matrix Fibroblast |
| Spdl1 | 0.62 | 0.00 | 0.00 | 0.27 | 0.02 | 24 | Matrix Fibroblast |
| Car2 | 1.58 | 0.00 | 0.00 | 0.92 | 0.30 | 24 | Matrix Fibroblast |
| Aurka | 0.55 | 0.00 | 0.00 | 0.26 | 0.02 | 24 | Matrix Fibroblast |
| Gm10259 | 1.08 | 0.00 | 0.00 | 0.46 | 0.06 | 24 | Matrix Fibroblast |
| Hmgb2 | 1.63 | 0.00 | 0.00 | 0.99 | 0.42 | 24 | Matrix Fibroblast |
| Hirip3 | 1.11 | 0.00 | 0.00 | 0.56 | 0.09 | 24 | Matrix Fibroblast |
| C1qtnf7 | 1.05 | 0.00 | 0.00 | 0.58 | 0.09 | 24 | Matrix Fibroblast |
| Clspn | 0.86 | 0.00 | 0.00 | 0.38 | 0.04 | 24 | Matrix Fibroblast |
| Hjurp | 1.19 | 0.00 | 0.00 | 0.71 | 0.14 | 24 | Matrix Fibroblast |
| Hoxb2 | 0.72 | 0.00 | 0.00 | 0.38 | 0.04 | 24 | Matrix Fibroblast |
| Mdk | 1.14 | 0.00 | 0.00 | 0.83 | 0.18 | 24 | Matrix Fibroblast |
| Ddr2 | 0.74 | 0.00 | 0.00 | 0.43 | 0.05 | 24 | Matrix Fibroblast |
| Acta2 | 0.88 | 0.00 | 0.00 | 0.85 | 0.19 | 24 | Matrix Fibroblast |
| Ncaph | 0.71 | 0.00 | 0.00 | 0.38 | 0.04 | 24 | Matrix Fibroblast |
| Gm10184 | 1.07 | 0.00 | 0.00 | 0.46 | 0.06 | 24 | Matrix Fibroblast |
| Igf1 | 1.17 | 0.00 | 0.00 | 0.69 | 0.14 | 24 | Matrix Fibroblast |
| Pdgfrl | 0.68 | 0.00 | 0.00 | 0.35 | 0.04 | 24 | Matrix Fibroblast |
| Nrcam | 0.81 | 0.00 | 0.00 | 0.43 | 0.06 | 24 | Matrix Fibroblast |
| Nes | 1.20 | 0.00 | 0.00 | 0.85 | 0.21 | 24 | Matrix Fibroblast |
| Tpm2 | 0.99 | 0.00 | 0.00 | 0.68 | 0.13 | 24 | Matrix Fibroblast |
| Trim59 | 0.75 | 0.00 | 0.00 | 0.37 | 0.04 | 24 | Matrix Fibroblast |
| Kpna2 | 1.00 | 0.00 | 0.00 | 0.43 | 0.06 | 24 | Matrix Fibroblast |
| Gm10282 | 1.38 | 0.00 | 0.00 | 0.96 | 0.40 | 24 | Matrix Fibroblast |
| Cenpl | 0.56 | 0.00 | 0.00 | 0.27 | 0.02 | 24 | Matrix Fibroblast |
| Hmgn2 | 1.43 | 0.00 | 0.00 | 0.95 | 0.43 | 24 | Matrix Fibroblast |
| Lgals1 | 1.41 | 0.00 | 0.00 | 0.99 | 0.41 | 24 | Matrix Fibroblast |
| Tyms | 0.79 | 0.00 | 0.00 | 0.41 | 0.05 | 24 | Matrix Fibroblast |
| H2afz | 1.38 | 0.00 | 0.00 | 0.94 | 0.40 | 24 | Matrix Fibroblast |
| Loxl2 | 1.06 | 0.00 | 0.00 | 0.67 | 0.14 | 24 | Matrix Fibroblast |
| Rad51ap1 | 0.70 | 0.00 | 0.00 | 0.31 | 0.03 | 24 | Matrix Fibroblast |
| Tnfrsf19 | 0.64 | 0.00 | 0.00 | 0.35 | 0.04 | 24 | Matrix Fibroblast |
| Gmnn | 0.91 | 0.00 | 0.00 | 0.46 | 0.07 | 24 | Matrix Fibroblast |
| Myh10 | 1.17 | 0.00 | 0.00 | 0.83 | 0.22 | 24 | Matrix Fibroblast |
| Rrm1 | 0.82 | 0.00 | 0.00 | 0.44 | 0.06 | 24 | Matrix Fibroblast |
| Basp1 | 0.75 | 0.00 | 0.00 | 0.42 | 0.06 | 24 | Matrix Fibroblast |
| Hoxb5 | 0.75 | 0.00 | 0.00 | 0.40 | 0.05 | 24 | Matrix Fibroblast |
| Cenpw | 0.70 | 0.00 | 0.00 | 0.38 | 0.05 | 24 | Matrix Fibroblast |
| Cald1 | 1.21 | 0.00 | 0.00 | 0.97 | 0.41 | 24 | Matrix Fibroblast |
| Smc4 | 1.32 | 0.00 | 0.00 | 0.92 | 0.34 | 24 | Matrix Fibroblast |
| Ncapg2 | 0.71 | 0.00 | 0.00 | 0.29 | 0.03 | 24 | Matrix Fibroblast |
| Ska2 | 0.84 | 0.00 | 0.00 | 0.43 | 0.07 | 24 | Matrix Fibroblast |
| Rnf152 | 0.59 | 0.00 | 0.00 | 0.31 | 0.03 | 24 | Matrix Fibroblast |
| Robo1 | 0.61 | 0.00 | 0.00 | 0.33 | 0.04 | 24 | Matrix Fibroblast |
| 1500009L16Rik | 0.61 | 0.00 | 0.00 | 0.33 | 0.04 | 24 | Matrix Fibroblast |
| Cygb | 0.69 | 0.00 | 0.00 | 0.40 | 0.06 | 24 | Matrix Fibroblast |
| Igsf10 | 0.62 | 0.00 | 0.00 | 0.33 | 0.04 | 24 | Matrix Fibroblast |
| Fbn2 | 0.71 | 0.00 | 0.00 | 0.39 | 0.05 | 24 | Matrix Fibroblast |
| Gm13889 | 0.74 | 0.00 | 0.00 | 0.43 | 0.07 | 24 | Matrix Fibroblast |
| Stc1 | 0.85 | 0.00 | 0.00 | 0.27 | 0.03 | 24 | Matrix Fibroblast |
| H2afy2 | 0.76 | 0.00 | 0.00 | 0.47 | 0.08 | 24 | Matrix Fibroblast |
| Tcf19 | 0.63 | 0.00 | 0.00 | 0.28 | 0.03 | 24 | Matrix Fibroblast |
| Pdzrn3 | 0.74 | 0.00 | 0.00 | 0.39 | 0.06 | 24 | Matrix Fibroblast |
| D8Ertd82e | 0.59 | 0.00 | 0.00 | 0.29 | 0.03 | 24 | Matrix Fibroblast |
| Atad2 | 0.96 | 0.00 | 0.00 | 0.47 | 0.08 | 24 | Matrix Fibroblast |
| RP23-103I12.13 | 0.86 | 0.00 | 0.00 | 0.53 | 0.10 | 24 | Matrix Fibroblast |
| 6330403K07Rik | 0.64 | 0.00 | 0.00 | 0.34 | 0.04 | 24 | Matrix Fibroblast |
| Col5a1 | 0.72 | 0.00 | 0.00 | 0.51 | 0.09 | 24 | Matrix Fibroblast |
| Nucks1 | 1.11 | 0.00 | 0.00 | 0.85 | 0.28 | 24 | Matrix Fibroblast |
| Dkk3 | 0.84 | 0.00 | 0.00 | 0.58 | 0.12 | 24 | Matrix Fibroblast |
| Lmnb1 | 0.81 | 0.00 | 0.00 | 0.48 | 0.08 | 24 | Matrix Fibroblast |
| Tubb5 | 1.19 | 0.00 | 0.00 | 0.94 | 0.44 | 24 | Matrix Fibroblast |
| 2700094K13Rik | 0.94 | 0.00 | 0.00 | 0.67 | 0.16 | 24 | Matrix Fibroblast |
| Bora | 0.61 | 0.00 | 0.00 | 0.28 | 0.03 | 24 | Matrix Fibroblast |
| Nrep | 1.18 | 0.00 | 0.00 | 0.95 | 0.43 | 24 | Matrix Fibroblast |
| Tmpo | 1.00 | 0.00 | 0.00 | 0.64 | 0.15 | 24 | Matrix Fibroblast |
| Cdkn2d | 0.74 | 0.00 | 0.00 | 0.42 | 0.07 | 24 | Matrix Fibroblast |
| Prim1 | 0.74 | 0.00 | 0.00 | 0.36 | 0.05 | 24 | Matrix Fibroblast |
| Cenpq | 0.78 | 0.00 | 0.00 | 0.41 | 0.07 | 24 | Matrix Fibroblast |
| Gpc4 | 0.52 | 0.00 | 0.00 | 0.32 | 0.04 | 24 | Matrix Fibroblast |
| Ckap5 | 0.94 | 0.00 | 0.00 | 0.54 | 0.11 | 24 | Matrix Fibroblast |
| Tbx5 | 0.70 | 0.00 | 0.00 | 0.45 | 0.08 | 24 | Matrix Fibroblast |
| Col3a1 | 0.73 | 0.00 | 0.00 | 0.96 | 0.33 | 24 | Matrix Fibroblast |
| 4930579G24Rik | 0.50 | 0.00 | 0.00 | 0.28 | 0.03 | 24 | Matrix Fibroblast |
| Prdx4 | 0.96 | 0.00 | 0.00 | 0.65 | 0.16 | 24 | Matrix Fibroblast |
| Ezh2 | 0.97 | 0.00 | 0.00 | 0.70 | 0.19 | 24 | Matrix Fibroblast |
| Foxf2 | 0.53 | 0.00 | 0.00 | 0.30 | 0.04 | 24 | Matrix Fibroblast |
| C1qtnf2 | 0.62 | 0.00 | 0.00 | 0.30 | 0.04 | 24 | Matrix Fibroblast |
| Tuba1b | 1.18 | 0.00 | 0.00 | 0.91 | 0.45 | 24 | Matrix Fibroblast |
| Nnat | 0.84 | 0.00 | 0.00 | 0.44 | 0.08 | 24 | Matrix Fibroblast |
| Tshz2 | 0.77 | 0.00 | 0.00 | 0.49 | 0.09 | 24 | Matrix Fibroblast |
| Phf17 | 0.64 | 0.00 | 0.00 | 0.34 | 0.05 | 24 | Matrix Fibroblast |
| 4632419I22Rik | 0.76 | 0.00 | 0.00 | 0.43 | 0.08 | 24 | Matrix Fibroblast |
| Ptma | 0.80 | 0.00 | 0.00 | 1.00 | 0.91 | 24 | Matrix Fibroblast |
| G2e3 | 0.78 | 0.00 | 0.00 | 0.38 | 0.06 | 24 | Matrix Fibroblast |
| Dbf4 | 0.69 | 0.00 | 0.00 | 0.34 | 0.05 | 24 | Matrix Fibroblast |
| Uhrf1 | 0.51 | 0.00 | 0.00 | 0.25 | 0.03 | 24 | Matrix Fibroblast |
| Dnajc9 | 0.80 | 0.00 | 0.00 | 0.54 | 0.12 | 24 | Matrix Fibroblast |
| Leprel2 | 0.66 | 0.00 | 0.00 | 0.41 | 0.07 | 24 | Matrix Fibroblast |
| Rad21 | 0.99 | 0.00 | 0.00 | 0.75 | 0.23 | 24 | Matrix Fibroblast |
| Tmem132c | 0.65 | 0.00 | 0.00 | 0.34 | 0.05 | 24 | Matrix Fibroblast |
| Nasp | 1.00 | 0.00 | 0.00 | 0.68 | 0.20 | 24 | Matrix Fibroblast |
| Gm6104 | 0.73 | 0.00 | 0.00 | 0.55 | 0.12 | 24 | Matrix Fibroblast |
| Filip1l | 0.99 | 0.00 | 0.00 | 0.87 | 0.33 | 24 | Matrix Fibroblast |
| Ckb | 0.75 | 0.00 | 0.00 | 0.60 | 0.14 | 24 | Matrix Fibroblast |
| Calm2 | 1.01 | 0.00 | 0.00 | 0.99 | 0.64 | 24 | Matrix Fibroblast |
| Fbn1 | 0.77 | 0.00 | 0.00 | 0.67 | 0.17 | 24 | Matrix Fibroblast |
| Oxct1 | 0.85 | 0.00 | 0.00 | 0.59 | 0.15 | 24 | Matrix Fibroblast |
| Actg2 | 0.63 | 0.00 | 0.00 | 0.42 | 0.07 | 24 | Matrix Fibroblast |
| Anp32e | 1.01 | 0.00 | 0.00 | 0.81 | 0.28 | 24 | Matrix Fibroblast |
| Hmgb3 | 0.81 | 0.00 | 0.00 | 0.58 | 0.14 | 24 | Matrix Fibroblast |
| Tgfb2 | 0.84 | 0.00 | 0.00 | 0.58 | 0.14 | 24 | Matrix Fibroblast |
| Smtn | 0.70 | 0.00 | 0.00 | 0.42 | 0.08 | 24 | Matrix Fibroblast |
| Des | 0.54 | 0.00 | 0.00 | 0.35 | 0.06 | 24 | Matrix Fibroblast |
| 10-Sep | 0.71 | 0.00 | 0.00 | 0.45 | 0.09 | 24 | Matrix Fibroblast |
| Lig1 | 0.84 | 0.00 | 0.00 | 0.46 | 0.09 | 24 | Matrix Fibroblast |
| Enc1 | 0.52 | 0.00 | 0.00 | 0.27 | 0.04 | 24 | Matrix Fibroblast |
| Tagln | 0.64 | 0.00 | 0.00 | 0.61 | 0.15 | 24 | Matrix Fibroblast |
| Rangap1 | 0.66 | 0.00 | 0.00 | 0.41 | 0.08 | 24 | Matrix Fibroblast |
| H2afv | 0.99 | 0.00 | 0.00 | 0.76 | 0.28 | 24 | Matrix Fibroblast |
| Mylk | 0.76 | 0.00 | 0.00 | 0.82 | 0.27 | 24 | Matrix Fibroblast |
| Tuba1a | 0.98 | 0.00 | 0.00 | 0.91 | 0.51 | 24 | Matrix Fibroblast |
| Lhfpl2 | 0.62 | 0.00 | 0.00 | 0.42 | 0.09 | 24 | Matrix Fibroblast |
| Cmc2 | 0.57 | 0.00 | 0.00 | 0.31 | 0.05 | 24 | Matrix Fibroblast |
| Fstl1 | 0.79 | 0.00 | 0.00 | 0.95 | 0.44 | 24 | Matrix Fibroblast |
| Dut | 0.77 | 0.00 | 0.00 | 0.42 | 0.09 | 24 | Matrix Fibroblast |
| Pmf1 | 0.44 | 0.00 | 0.00 | 0.28 | 0.04 | 24 | Matrix Fibroblast |
| Col1a2 | 0.54 | 0.00 | 0.00 | 0.99 | 0.39 | 24 | Matrix Fibroblast |
| Rarres2 | 0.65 | 0.00 | 0.00 | 0.54 | 0.13 | 24 | Matrix Fibroblast |
| Mex3a | 0.63 | 0.00 | 0.00 | 0.41 | 0.08 | 24 | Matrix Fibroblast |
| Ran | 0.94 | 0.00 | 0.00 | 0.85 | 0.38 | 24 | Matrix Fibroblast |
| Cdk5rap2 | 0.60 | 0.00 | 0.00 | 0.43 | 0.09 | 24 | Matrix Fibroblast |
| Cdh11 | 0.58 | 0.00 | 0.00 | 0.68 | 0.19 | 24 | Matrix Fibroblast |
| Mis18a | 0.46 | 0.00 | 0.00 | 0.28 | 0.04 | 24 | Matrix Fibroblast |
| Mis12 | 0.64 | 0.00 | 0.00 | 0.36 | 0.07 | 24 | Matrix Fibroblast |
| Rfc4 | 0.50 | 0.00 | 0.00 | 0.25 | 0.04 | 24 | Matrix Fibroblast |
| Prr5l | 0.58 | 0.00 | 0.00 | 0.35 | 0.06 | 24 | Matrix Fibroblast |
| Pola1 | 0.55 | 0.00 | 0.00 | 0.28 | 0.04 | 24 | Matrix Fibroblast |
| Plac9a | 0.59 | 0.00 | 0.00 | 0.88 | 0.30 | 24 | Matrix Fibroblast |
| Myl9 | 0.61 | 0.00 | 0.00 | 0.50 | 0.12 | 24 | Matrix Fibroblast |
| Gpr126 | 0.60 | 0.00 | 0.00 | 0.46 | 0.10 | 24 | Matrix Fibroblast |
| Hmgb1 | 0.85 | 0.00 | 0.00 | 0.94 | 0.61 | 24 | Matrix Fibroblast |
| Bgn | 0.65 | 0.00 | 0.00 | 0.75 | 0.24 | 24 | Matrix Fibroblast |
| Tbx4 | 0.58 | 0.00 | 0.00 | 0.35 | 0.07 | 24 | Matrix Fibroblast |
| Nfib | 0.87 | 0.00 | 0.00 | 0.92 | 0.47 | 24 | Matrix Fibroblast |
| Lhfp | 0.55 | 0.00 | 0.00 | 0.58 | 0.15 | 24 | Matrix Fibroblast |
| Psip1 | 0.76 | 0.00 | 0.00 | 0.54 | 0.15 | 24 | Matrix Fibroblast |
| Tgfb3 | 0.52 | 0.00 | 0.00 | 0.31 | 0.05 | 24 | Matrix Fibroblast |
| Kcnk3 | 0.43 | 0.00 | 0.00 | 0.29 | 0.05 | 24 | Matrix Fibroblast |
| Tipin | 0.72 | 0.00 | 0.00 | 0.43 | 0.10 | 24 | Matrix Fibroblast |
| Plac9b | 0.56 | 0.00 | 0.00 | 0.86 | 0.30 | 24 | Matrix Fibroblast |
| Tshz1 | 0.77 | 0.00 | 0.00 | 0.47 | 0.12 | 24 | Matrix Fibroblast |
| Cit | 0.52 | 0.00 | 0.00 | 0.27 | 0.05 | 24 | Matrix Fibroblast |
| Tuba1c | 0.92 | 0.00 | 0.00 | 0.78 | 0.32 | 24 | Matrix Fibroblast |
| Lmna | 0.81 | 0.00 | 0.00 | 0.64 | 0.20 | 24 | Matrix Fibroblast |
| Gpr64 | 0.51 | 0.00 | 0.00 | 0.29 | 0.05 | 24 | Matrix Fibroblast |
| Pcna | 0.96 | 0.00 | 0.00 | 0.57 | 0.18 | 24 | Matrix Fibroblast |
| Nrm | 0.53 | 0.00 | 0.00 | 0.30 | 0.05 | 24 | Matrix Fibroblast |
| Efemp2 | 0.52 | 0.00 | 0.00 | 0.35 | 0.07 | 24 | Matrix Fibroblast |
| Hist1h1e | 1.03 | 0.00 | 0.00 | 0.55 | 0.16 | 24 | Matrix Fibroblast |
| Cbx3 | 0.80 | 0.00 | 0.00 | 0.83 | 0.34 | 24 | Matrix Fibroblast |
| Ncapd3 | 0.56 | 0.00 | 0.00 | 0.31 | 0.06 | 24 | Matrix Fibroblast |
| Golim4 | 0.77 | 0.00 | 0.00 | 0.63 | 0.20 | 24 | Matrix Fibroblast |
| Snai2 | 0.58 | 0.00 | 0.00 | 0.48 | 0.12 | 24 | Matrix Fibroblast |
| Serpine2 | 0.71 | 0.00 | 0.00 | 0.74 | 0.25 | 24 | Matrix Fibroblast |
| Calu | 0.71 | 0.00 | 0.00 | 0.70 | 0.24 | 24 | Matrix Fibroblast |
| Grb10 | 0.68 | 0.00 | 0.00 | 0.56 | 0.16 | 24 | Matrix Fibroblast |
| Larp7 | 0.75 | 0.00 | 0.00 | 0.57 | 0.17 | 24 | Matrix Fibroblast |
| Col16a1 | 0.55 | 0.00 | 0.00 | 0.30 | 0.06 | 24 | Matrix Fibroblast |
| Angptl2 | 0.44 | 0.00 | 0.00 | 0.26 | 0.04 | 24 | Matrix Fibroblast |
| Pcna-ps2 | 0.92 | 0.00 | 0.00 | 0.57 | 0.18 | 24 | Matrix Fibroblast |
| Cenpc1 | 0.69 | 0.00 | 0.00 | 0.38 | 0.09 | 24 | Matrix Fibroblast |
| Whsc1 | 0.66 | 0.00 | 0.00 | 0.50 | 0.14 | 24 | Matrix Fibroblast |
| Ranbp1 | 0.84 | 0.00 | 0.00 | 0.73 | 0.29 | 24 | Matrix Fibroblast |
| Cd34 | 0.58 | 0.00 | 0.00 | 0.64 | 0.20 | 24 | Matrix Fibroblast |
| Palld | 0.65 | 0.00 | 0.00 | 0.58 | 0.17 | 24 | Matrix Fibroblast |
| Rbms3 | 0.64 | 0.00 | 0.00 | 0.57 | 0.17 | 24 | Matrix Fibroblast |
| Ism1 | 0.49 | 0.00 | 0.00 | 0.39 | 0.09 | 24 | Matrix Fibroblast |
| Fzd1 | 0.44 | 0.00 | 0.00 | 0.29 | 0.06 | 24 | Matrix Fibroblast |
| Usp1 | 0.65 | 0.00 | 0.00 | 0.53 | 0.15 | 24 | Matrix Fibroblast |
| Banf1 | 0.71 | 0.00 | 0.00 | 0.73 | 0.28 | 24 | Matrix Fibroblast |
| B3galtl | 0.56 | 0.00 | 0.00 | 0.31 | 0.06 | 24 | Matrix Fibroblast |
| Dtymk | 0.65 | 0.00 | 0.00 | 0.48 | 0.13 | 24 | Matrix Fibroblast |
| Nexn | 0.67 | 0.00 | 0.00 | 0.66 | 0.22 | 24 | Matrix Fibroblast |
| Zeb2 | 0.72 | 0.00 | 0.00 | 0.79 | 0.31 | 24 | Matrix Fibroblast |
| Egflam | 0.54 | 0.00 | 0.00 | 0.36 | 0.08 | 24 | Matrix Fibroblast |
| Gm10068 | 0.75 | 0.00 | 0.00 | 0.75 | 0.30 | 24 | Matrix Fibroblast |
| Rasl11b | 0.41 | 0.00 | 0.00 | 0.28 | 0.05 | 24 | Matrix Fibroblast |
| Ckap4 | 0.72 | 0.00 | 0.00 | 0.56 | 0.18 | 24 | Matrix Fibroblast |
| Tubb4b | 0.75 | 0.00 | 0.00 | 0.57 | 0.19 | 24 | Matrix Fibroblast |
| Tpm1 | 0.74 | 0.00 | 0.00 | 0.92 | 0.52 | 24 | Matrix Fibroblast |
| H19 | 0.35 | 0.00 | 0.00 | 0.34 | 0.07 | 24 | Matrix Fibroblast |
| Klhl23 | 0.42 | 0.00 | 0.00 | 0.28 | 0.05 | 24 | Matrix Fibroblast |
| Rfc3 | 0.58 | 0.00 | 0.00 | 0.37 | 0.09 | 24 | Matrix Fibroblast |
| Rnaseh2b | 0.52 | 0.00 | 0.00 | 0.35 | 0.08 | 24 | Matrix Fibroblast |
| Cbx5 | 0.71 | 0.00 | 0.00 | 0.69 | 0.26 | 24 | Matrix Fibroblast |
| Gm9847 | 0.59 | 0.00 | 0.00 | 0.53 | 0.16 | 24 | Matrix Fibroblast |
| Iigp1 | 0.49 | 0.00 | 0.00 | 0.48 | 0.13 | 24 | Matrix Fibroblast |
| Net1 | 0.62 | 0.00 | 0.00 | 0.55 | 0.17 | 24 | Matrix Fibroblast |
| Smc1a | 0.77 | 0.00 | 0.00 | 0.78 | 0.35 | 24 | Matrix Fibroblast |
| Cpq | 0.45 | 0.00 | 0.00 | 0.37 | 0.08 | 24 | Matrix Fibroblast |
| Nfix | 0.63 | 0.00 | 0.00 | 0.54 | 0.17 | 24 | Matrix Fibroblast |
| Rcn3 | 0.52 | 0.00 | 0.00 | 0.47 | 0.13 | 24 | Matrix Fibroblast |
| Qpct | 0.38 | 0.00 | 0.00 | 0.25 | 0.05 | 24 | Matrix Fibroblast |
| Nde1 | 0.49 | 0.00 | 0.00 | 0.30 | 0.06 | 24 | Matrix Fibroblast |
| Gpm6b | 0.51 | 0.00 | 0.00 | 0.42 | 0.11 | 24 | Matrix Fibroblast |
| Pros1 | 0.47 | 0.00 | 0.00 | 0.40 | 0.10 | 24 | Matrix Fibroblast |
| Dek | 0.81 | 0.00 | 0.00 | 0.88 | 0.50 | 24 | Matrix Fibroblast |
| 11-Sep | 0.65 | 0.00 | 0.00 | 0.63 | 0.22 | 24 | Matrix Fibroblast |
| Socs2 | 0.64 | 0.00 | 0.00 | 0.80 | 0.34 | 24 | Matrix Fibroblast |
| Hist1h1d | 0.83 | 0.00 | 0.00 | 0.38 | 0.10 | 24 | Matrix Fibroblast |
| Tbx2 | 0.47 | 0.00 | 0.00 | 0.44 | 0.12 | 24 | Matrix Fibroblast |
| Snrpd1 | 0.65 | 0.00 | 0.00 | 0.58 | 0.20 | 24 | Matrix Fibroblast |
| Cdk4 | 0.69 | 0.00 | 0.00 | 0.64 | 0.24 | 24 | Matrix Fibroblast |
| Lamb1 | 0.48 | 0.00 | 0.00 | 0.53 | 0.16 | 24 | Matrix Fibroblast |
| 1700025G04Rik | 0.53 | 0.00 | 0.00 | 0.39 | 0.10 | 24 | Matrix Fibroblast |
| Asph | 0.45 | 0.00 | 0.00 | 0.29 | 0.06 | 24 | Matrix Fibroblast |
| Dnmt1 | 0.52 | 0.00 | 0.00 | 0.40 | 0.10 | 24 | Matrix Fibroblast |
| Rpn2 | 0.66 | 0.00 | 0.00 | 0.59 | 0.21 | 24 | Matrix Fibroblast |
| Actn1 | 0.69 | 0.00 | 0.00 | 0.61 | 0.23 | 24 | Matrix Fibroblast |
| Hsp90b1 | 0.62 | 0.00 | 0.00 | 0.98 | 0.76 | 24 | Matrix Fibroblast |
| Hist2h2ac | 0.51 | 0.00 | 0.00 | 0.29 | 0.06 | 24 | Matrix Fibroblast |
| Aebp1 | 0.45 | 0.00 | 0.00 | 0.33 | 0.08 | 24 | Matrix Fibroblast |
| Map1b | 0.57 | 0.00 | 0.00 | 0.39 | 0.10 | 24 | Matrix Fibroblast |
| Cntln | 0.65 | 0.00 | 0.00 | 0.42 | 0.12 | 24 | Matrix Fibroblast |
| Cdca7 | 0.48 | 0.00 | 0.00 | 0.27 | 0.06 | 24 | Matrix Fibroblast |
| Med30 | 0.53 | 0.00 | 0.00 | 0.35 | 0.09 | 24 | Matrix Fibroblast |
| Bub3 | 0.61 | 0.00 | 0.00 | 0.46 | 0.14 | 24 | Matrix Fibroblast |
| 5830418K08Rik | 0.61 | 0.00 | 0.00 | 0.35 | 0.09 | 24 | Matrix Fibroblast |
| Cd44 | 0.46 | 0.00 | 0.00 | 0.60 | 0.20 | 24 | Matrix Fibroblast |
| Hnrnpa2b1 | 0.61 | 0.00 | 0.00 | 0.95 | 0.72 | 24 | Matrix Fibroblast |
| Sae1 | 0.47 | 0.00 | 0.00 | 0.40 | 0.11 | 24 | Matrix Fibroblast |
| Col1a1 | 0.30 | 0.00 | 0.00 | 0.71 | 0.26 | 24 | Matrix Fibroblast |
| Cdca4 | 0.36 | 0.00 | 0.00 | 0.25 | 0.05 | 24 | Matrix Fibroblast |
| Tpm4 | 0.66 | 0.00 | 0.00 | 0.87 | 0.49 | 24 | Matrix Fibroblast |
| Haus8 | 0.46 | 0.00 | 0.00 | 0.25 | 0.05 | 24 | Matrix Fibroblast |
| Hmgn1 | 0.67 | 0.00 | 0.00 | 0.85 | 0.45 | 24 | Matrix Fibroblast |
| Mcm7 | 0.48 | 0.00 | 0.00 | 0.31 | 0.07 | 24 | Matrix Fibroblast |
| Hells | 0.44 | 0.00 | 0.00 | 0.27 | 0.06 | 24 | Matrix Fibroblast |
| Hn1 | 0.70 | 0.00 | 0.00 | 0.55 | 0.20 | 24 | Matrix Fibroblast |
| Mfap2 | 0.55 | 0.00 | 0.00 | 0.82 | 0.35 | 24 | Matrix Fibroblast |
| Rgs2 | 0.46 | 0.00 | 0.00 | 0.71 | 0.28 | 24 | Matrix Fibroblast |
| Creb3l2 | 0.54 | 0.00 | 0.00 | 0.45 | 0.13 | 24 | Matrix Fibroblast |
| Thbs1 | 0.51 | 0.00 | 0.00 | 0.44 | 0.13 | 24 | Matrix Fibroblast |
| Lsp1 | 0.68 | 0.00 | 0.00 | 0.56 | 0.19 | 24 | Matrix Fibroblast |
| Rpa3 | 0.57 | 0.00 | 0.00 | 0.43 | 0.13 | 24 | Matrix Fibroblast |
| Cep112 | 0.41 | 0.00 | 0.00 | 0.26 | 0.05 | 24 | Matrix Fibroblast |
| Myef2 | 0.56 | 0.00 | 0.00 | 0.49 | 0.16 | 24 | Matrix Fibroblast |
| Fut8 | 0.36 | 0.00 | 0.00 | 0.26 | 0.06 | 24 | Matrix Fibroblast |
| Mapre2 | 0.55 | 0.00 | 0.00 | 0.39 | 0.11 | 24 | Matrix Fibroblast |
| Prdm6 | 0.38 | 0.00 | 0.00 | 0.29 | 0.07 | 24 | Matrix Fibroblast |
| Papss1 | 0.37 | 0.00 | 0.00 | 0.30 | 0.07 | 24 | Matrix Fibroblast |
| Pdlim7 | 0.40 | 0.00 | 0.00 | 0.38 | 0.10 | 24 | Matrix Fibroblast |
| H3f3b | 0.53 | 0.00 | 0.00 | 0.99 | 0.89 | 24 | Matrix Fibroblast |
| Atad5 | 0.56 | 0.00 | 0.00 | 0.25 | 0.05 | 24 | Matrix Fibroblast |
| Lsm3 | 0.47 | 0.00 | 0.00 | 0.39 | 0.11 | 24 | Matrix Fibroblast |
| Mgp | 0.48 | 0.00 | 0.00 | 0.77 | 0.32 | 24 | Matrix Fibroblast |
| Mcm4 | 0.39 | 0.00 | 0.00 | 0.26 | 0.06 | 24 | Matrix Fibroblast |
| Atpif1 | 0.64 | 0.00 | 0.00 | 0.90 | 0.53 | 24 | Matrix Fibroblast |
| Pxdn | 0.41 | 0.00 | 0.00 | 0.39 | 0.11 | 24 | Matrix Fibroblast |
| Hoxb4 | 0.41 | 0.00 | 0.00 | 0.29 | 0.07 | 24 | Matrix Fibroblast |
| Gm10123 | 0.54 | 0.00 | 0.00 | 0.96 | 0.81 | 24 | Matrix Fibroblast |
| Itga9 | 0.48 | 0.00 | 0.00 | 0.36 | 0.09 | 24 | Matrix Fibroblast |
| Smc6 | 0.58 | 0.00 | 0.00 | 0.76 | 0.33 | 24 | Matrix Fibroblast |
| Slbp | 0.64 | 0.00 | 0.00 | 0.46 | 0.15 | 24 | Matrix Fibroblast |
| Cbx1 | 0.61 | 0.00 | 0.00 | 0.83 | 0.41 | 24 | Matrix Fibroblast |
| Gm9833 | 0.48 | 0.00 | 0.00 | 0.39 | 0.11 | 24 | Matrix Fibroblast |
| Rbbp7 | 0.67 | 0.00 | 0.00 | 0.63 | 0.26 | 24 | Matrix Fibroblast |
| Ccdc88a | 0.55 | 0.00 | 0.00 | 0.64 | 0.24 | 24 | Matrix Fibroblast |
| Dnajc10 | 0.52 | 0.00 | 0.00 | 0.47 | 0.15 | 24 | Matrix Fibroblast |
| Nudcd2 | 0.50 | 0.00 | 0.00 | 0.37 | 0.10 | 24 | Matrix Fibroblast |
| Tubb4b-ps1 | 0.48 | 0.00 | 0.00 | 0.33 | 0.09 | 24 | Matrix Fibroblast |
| Adk | 0.46 | 0.00 | 0.00 | 0.40 | 0.12 | 24 | Matrix Fibroblast |
| Bmp4 | 0.45 | 0.00 | 0.00 | 0.37 | 0.10 | 24 | Matrix Fibroblast |
| Rnaseh2c | 0.48 | 0.00 | 0.00 | 0.37 | 0.11 | 24 | Matrix Fibroblast |
| Serf1 | 0.56 | 0.00 | 0.00 | 0.38 | 0.11 | 24 | Matrix Fibroblast |
| Serpinh1 | 0.55 | 0.00 | 0.00 | 0.82 | 0.39 | 24 | Matrix Fibroblast |
| Rcn2 | 0.49 | 0.00 | 0.00 | 0.46 | 0.15 | 24 | Matrix Fibroblast |
| Ppia | 0.53 | 0.00 | 0.00 | 0.95 | 0.80 | 24 | Matrix Fibroblast |
| Col6a3 | 0.43 | 0.00 | 0.00 | 0.35 | 0.10 | 24 | Matrix Fibroblast |
| Gm6625 | 0.61 | 0.00 | 0.00 | 0.63 | 0.25 | 24 | Matrix Fibroblast |
| Mrpl51 | 0.52 | 0.00 | 0.00 | 0.47 | 0.16 | 24 | Matrix Fibroblast |
| Ssrp1 | 0.59 | 0.00 | 0.00 | 0.56 | 0.21 | 24 | Matrix Fibroblast |
| Hsp90aa1 | 0.52 | 0.00 | 0.00 | 0.85 | 0.47 | 24 | Matrix Fibroblast |
| Kpnb1 | 0.50 | 0.00 | 0.00 | 0.53 | 0.19 | 24 | Matrix Fibroblast |
| Pknox1 | 0.42 | 0.00 | 0.00 | 0.28 | 0.07 | 24 | Matrix Fibroblast |
| Mpdz | 0.43 | 0.00 | 0.00 | 0.30 | 0.08 | 24 | Matrix Fibroblast |
| Mpzl1 | 0.45 | 0.00 | 0.00 | 0.38 | 0.11 | 24 | Matrix Fibroblast |
| Adamts10 | 0.47 | 0.00 | 0.00 | 0.39 | 0.11 | 24 | Matrix Fibroblast |
| Gjc1 | 0.43 | 0.00 | 0.00 | 0.30 | 0.08 | 24 | Matrix Fibroblast |
| Pam | 0.49 | 0.00 | 0.00 | 0.50 | 0.17 | 24 | Matrix Fibroblast |
| Stub1 | 0.47 | 0.00 | 0.00 | 0.58 | 0.22 | 24 | Matrix Fibroblast |
| Samd4 | 0.45 | 0.00 | 0.00 | 0.32 | 0.09 | 24 | Matrix Fibroblast |
| Gnb4 | 0.51 | 0.00 | 0.00 | 0.41 | 0.13 | 24 | Matrix Fibroblast |
| Nptn | 0.51 | 0.00 | 0.00 | 0.56 | 0.21 | 24 | Matrix Fibroblast |
| Smarcc1 | 0.49 | 0.00 | 0.00 | 0.47 | 0.17 | 24 | Matrix Fibroblast |
| Terf1 | 0.41 | 0.00 | 0.00 | 0.25 | 0.06 | 24 | Matrix Fibroblast |
| Ryk | 0.45 | 0.00 | 0.00 | 0.41 | 0.13 | 24 | Matrix Fibroblast |
| Gpx7 | 0.47 | 0.00 | 0.00 | 0.34 | 0.10 | 24 | Matrix Fibroblast |
| Vdac1 | 0.51 | 0.00 | 0.00 | 0.54 | 0.21 | 24 | Matrix Fibroblast |
| Ncald | 0.46 | 0.00 | 0.00 | 0.34 | 0.10 | 24 | Matrix Fibroblast |
| Erh | 0.61 | 0.00 | 0.00 | 0.65 | 0.28 | 24 | Matrix Fibroblast |
| Cep290 | 0.52 | 0.00 | 0.00 | 0.34 | 0.10 | 24 | Matrix Fibroblast |
| Brd8 | 0.54 | 0.00 | 0.00 | 0.52 | 0.19 | 24 | Matrix Fibroblast |
| Rbbp4 | 0.54 | 0.00 | 0.00 | 0.53 | 0.20 | 24 | Matrix Fibroblast |
| Ptms | 0.69 | 0.00 | 0.00 | 0.77 | 0.40 | 24 | Matrix Fibroblast |
| Acat1 | 0.55 | 0.00 | 0.00 | 0.49 | 0.18 | 24 | Matrix Fibroblast |
| Plxna2 | 0.42 | 0.00 | 0.00 | 0.29 | 0.07 | 24 | Matrix Fibroblast |
| Igfbp5 | 0.44 | 0.00 | 0.00 | 0.36 | 0.11 | 24 | Matrix Fibroblast |
| Cep57 | 0.47 | 0.00 | 0.00 | 0.36 | 0.11 | 24 | Matrix Fibroblast |
| Selm | 0.34 | 0.00 | 0.00 | 0.32 | 0.09 | 24 | Matrix Fibroblast |
| Lsm2 | 0.40 | 0.00 | 0.00 | 0.32 | 0.09 | 24 | Matrix Fibroblast |
| Pbdc1 | 0.42 | 0.00 | 0.00 | 0.40 | 0.13 | 24 | Matrix Fibroblast |
| Mllt3 | 0.46 | 0.00 | 0.00 | 0.37 | 0.11 | 24 | Matrix Fibroblast |
| Ccdc41 | 0.46 | 0.00 | 0.00 | 0.48 | 0.17 | 24 | Matrix Fibroblast |
| Lcorl | 0.37 | 0.00 | 0.00 | 0.28 | 0.07 | 24 | Matrix Fibroblast |
| Ddah2 | 0.49 | 0.00 | 0.00 | 0.64 | 0.26 | 24 | Matrix Fibroblast |
| Hist2h2aa2 | 0.56 | 0.00 | 0.00 | 0.41 | 0.14 | 24 | Matrix Fibroblast |
| Srsf3 | 0.56 | 0.00 | 0.00 | 0.83 | 0.44 | 24 | Matrix Fibroblast |
| Lama4 | 0.40 | 0.00 | 0.00 | 0.41 | 0.13 | 24 | Matrix Fibroblast |
| Sgce | 0.45 | 0.00 | 0.00 | 0.34 | 0.10 | 24 | Matrix Fibroblast |
| Xpo1 | 0.39 | 0.00 | 0.00 | 0.33 | 0.10 | 24 | Matrix Fibroblast |
| Fam111a | 0.42 | 0.00 | 0.00 | 0.45 | 0.15 | 24 | Matrix Fibroblast |
| Anp32b | 0.58 | 0.00 | 0.00 | 0.84 | 0.50 | 24 | Matrix Fibroblast |
| Spry1 | 0.38 | 0.00 | 0.00 | 0.27 | 0.07 | 24 | Matrix Fibroblast |
| Baz1b | 0.56 | 0.00 | 0.00 | 0.61 | 0.26 | 24 | Matrix Fibroblast |
| Snhg5 | 0.48 | 0.00 | 0.00 | 0.48 | 0.18 | 24 | Matrix Fibroblast |
| Adamts2 | 0.48 | 0.00 | 0.00 | 0.35 | 0.10 | 24 | Matrix Fibroblast |
| Tnfaip8 | 0.41 | 0.00 | 0.00 | 0.35 | 0.10 | 24 | Matrix Fibroblast |
| Tspan3 | 0.41 | 0.00 | 0.00 | 0.45 | 0.16 | 24 | Matrix Fibroblast |
| Gng12 | 0.44 | 0.00 | 0.00 | 0.38 | 0.12 | 24 | Matrix Fibroblast |
| Ilk | 0.49 | 0.00 | 0.00 | 0.56 | 0.23 | 24 | Matrix Fibroblast |
| Nr2f2 | 0.39 | 0.00 | 0.00 | 0.55 | 0.21 | 24 | Matrix Fibroblast |
| Naa50 | 0.40 | 0.00 | 0.00 | 0.40 | 0.13 | 24 | Matrix Fibroblast |
| Cgrrf1 | 0.36 | 0.00 | 0.00 | 0.30 | 0.08 | 24 | Matrix Fibroblast |
| Pa2g4 | 0.51 | 0.00 | 0.00 | 0.61 | 0.26 | 24 | Matrix Fibroblast |
| Ptprs | 0.36 | 0.00 | 0.00 | 0.29 | 0.08 | 24 | Matrix Fibroblast |
| Ilf3 | 0.49 | 0.00 | 0.00 | 0.37 | 0.12 | 24 | Matrix Fibroblast |
| Celf2 | 0.44 | 0.00 | 0.00 | 0.65 | 0.28 | 24 | Matrix Fibroblast |
| Nop58 | 0.52 | 0.00 | 0.00 | 0.60 | 0.26 | 24 | Matrix Fibroblast |
| Ilf2 | 0.47 | 0.00 | 0.00 | 0.50 | 0.19 | 24 | Matrix Fibroblast |
| Prkrir | 0.36 | 0.00 | 0.00 | 0.34 | 0.11 | 24 | Matrix Fibroblast |
| 2700029M09Rik | 0.42 | 0.00 | 0.00 | 0.38 | 0.13 | 24 | Matrix Fibroblast |
| Gtf2a2 | 0.45 | 0.00 | 0.00 | 0.49 | 0.19 | 24 | Matrix Fibroblast |
| Fkbp10 | 0.30 | 0.00 | 0.00 | 0.28 | 0.08 | 24 | Matrix Fibroblast |
| Syncrip | 0.50 | 0.00 | 0.00 | 0.58 | 0.25 | 24 | Matrix Fibroblast |
| Supt16 | 0.53 | 0.00 | 0.00 | 0.58 | 0.25 | 24 | Matrix Fibroblast |
| Ghr | 0.35 | 0.00 | 0.00 | 0.43 | 0.15 | 24 | Matrix Fibroblast |
| Nsmce1 | 0.38 | 0.00 | 0.00 | 0.40 | 0.14 | 24 | Matrix Fibroblast |
| Zadh2 | 0.37 | 0.00 | 0.00 | 0.29 | 0.08 | 24 | Matrix Fibroblast |
| Gm6793 | 0.55 | 0.00 | 0.00 | 0.64 | 0.30 | 24 | Matrix Fibroblast |
| Ikbip | 0.43 | 0.00 | 0.00 | 0.28 | 0.08 | 24 | Matrix Fibroblast |
| Mustn1 | 0.36 | 0.00 | 0.00 | 0.28 | 0.08 | 24 | Matrix Fibroblast |
| Ddx1 | 0.46 | 0.00 | 0.00 | 0.45 | 0.17 | 24 | Matrix Fibroblast |
| Exosc8 | 0.31 | 0.00 | 0.00 | 0.29 | 0.08 | 24 | Matrix Fibroblast |
| Cnn3 | 0.46 | 0.00 | 0.00 | 0.67 | 0.30 | 24 | Matrix Fibroblast |
| Srsf7 | 0.41 | 0.00 | 0.00 | 0.43 | 0.16 | 24 | Matrix Fibroblast |
| Eif4e | 0.37 | 0.00 | 0.00 | 0.33 | 0.10 | 24 | Matrix Fibroblast |
| Minos1 | 0.55 | 0.00 | 0.00 | 0.78 | 0.44 | 24 | Matrix Fibroblast |
| Csrp2 | 0.35 | 0.00 | 0.00 | 0.46 | 0.17 | 24 | Matrix Fibroblast |
| Rab13 | 0.34 | 0.00 | 0.00 | 0.31 | 0.09 | 24 | Matrix Fibroblast |
| Hnrnpr | 0.48 | 0.00 | 0.00 | 0.57 | 0.25 | 24 | Matrix Fibroblast |
| Uchl5 | 0.36 | 0.00 | 0.00 | 0.35 | 0.12 | 24 | Matrix Fibroblast |
| Tead2 | 0.39 | 0.00 | 0.00 | 0.38 | 0.13 | 24 | Matrix Fibroblast |
| Hnrnpab | 0.55 | 0.00 | 0.00 | 0.67 | 0.33 | 24 | Matrix Fibroblast |
| Fip1l1 | 0.46 | 0.00 | 0.00 | 0.47 | 0.18 | 24 | Matrix Fibroblast |
| Pmepa1 | 0.36 | 0.00 | 0.00 | 0.34 | 0.11 | 24 | Matrix Fibroblast |
| Slc39a6 | 0.32 | 0.00 | 0.00 | 0.27 | 0.07 | 24 | Matrix Fibroblast |
| Rcn1 | 0.38 | 0.00 | 0.00 | 0.40 | 0.14 | 24 | Matrix Fibroblast |
| Nbl1 | 0.37 | 0.00 | 0.00 | 0.25 | 0.07 | 24 | Matrix Fibroblast |
| Cgnl1 | 0.34 | 0.00 | 0.00 | 0.36 | 0.12 | 24 | Matrix Fibroblast |
| Hnrnpa1 | 0.49 | 0.00 | 0.00 | 0.59 | 0.27 | 24 | Matrix Fibroblast |
| Nfic | 0.48 | 0.00 | 0.00 | 0.56 | 0.24 | 24 | Matrix Fibroblast |
| Nono | 0.45 | 0.00 | 0.00 | 0.51 | 0.21 | 24 | Matrix Fibroblast |
| Ndufaf2 | 0.49 | 0.00 | 0.00 | 0.36 | 0.12 | 24 | Matrix Fibroblast |
| Thoc7 | 0.48 | 0.00 | 0.00 | 0.67 | 0.32 | 24 | Matrix Fibroblast |
| Col5a2 | 0.31 | 0.00 | 0.00 | 0.48 | 0.18 | 24 | Matrix Fibroblast |
| Ppic | 0.47 | 0.00 | 0.00 | 0.76 | 0.38 | 24 | Matrix Fibroblast |
| Rbbp8 | 0.43 | 0.00 | 0.00 | 0.29 | 0.09 | 24 | Matrix Fibroblast |
| P4hb | 0.47 | 0.00 | 0.00 | 0.76 | 0.40 | 24 | Matrix Fibroblast |
| Nfia | 0.43 | 0.00 | 0.00 | 0.56 | 0.24 | 24 | Matrix Fibroblast |
| Txnl1 | 0.42 | 0.00 | 0.00 | 0.48 | 0.19 | 24 | Matrix Fibroblast |
| Idh2 | 0.47 | 0.00 | 0.00 | 0.48 | 0.19 | 24 | Matrix Fibroblast |
| Fkbp3 | 0.43 | 0.00 | 0.00 | 0.64 | 0.30 | 24 | Matrix Fibroblast |
| Adam19 | 0.38 | 0.00 | 0.00 | 0.30 | 0.10 | 24 | Matrix Fibroblast |
| Arl6ip1 | 0.71 | 0.00 | 0.00 | 0.80 | 0.52 | 24 | Matrix Fibroblast |
| Commd1 | 0.42 | 0.00 | 0.00 | 0.50 | 0.21 | 24 | Matrix Fibroblast |
| Mmp14 | 0.37 | 0.00 | 0.00 | 0.59 | 0.25 | 24 | Matrix Fibroblast |
| Tsen34 | 0.36 | 0.00 | 0.00 | 0.38 | 0.14 | 24 | Matrix Fibroblast |
| Eny2 | 0.37 | 0.00 | 0.00 | 0.55 | 0.23 | 24 | Matrix Fibroblast |
| Tcerg1 | 0.39 | 0.00 | 0.00 | 0.49 | 0.20 | 24 | Matrix Fibroblast |
| Hist1h1c | 0.52 | 0.00 | 0.00 | 0.33 | 0.11 | 24 | Matrix Fibroblast |
| Srsf2 | 0.48 | 0.00 | 0.00 | 0.73 | 0.38 | 24 | Matrix Fibroblast |
| Pds5b | 0.42 | 0.00 | 0.00 | 0.36 | 0.13 | 24 | Matrix Fibroblast |
| 2010111I01Rik | 0.34 | 0.00 | 0.00 | 0.29 | 0.09 | 24 | Matrix Fibroblast |
| Nme1 | 0.35 | 0.00 | 0.00 | 0.47 | 0.19 | 24 | Matrix Fibroblast |
| Ctdspl2 | 0.39 | 0.00 | 0.00 | 0.34 | 0.11 | 24 | Matrix Fibroblast |
| Snrnp40 | 0.29 | 0.00 | 0.00 | 0.28 | 0.08 | 24 | Matrix Fibroblast |
| Ctcf | 0.43 | 0.00 | 0.00 | 0.54 | 0.23 | 24 | Matrix Fibroblast |
| Rbmxl1 | 0.34 | 0.00 | 0.00 | 0.32 | 0.11 | 24 | Matrix Fibroblast |
| Smc3 | 0.47 | 0.00 | 0.00 | 0.72 | 0.38 | 24 | Matrix Fibroblast |
| Orc6 | 0.35 | 0.00 | 0.00 | 0.31 | 0.10 | 24 | Matrix Fibroblast |
| Purb | 0.45 | 0.00 | 0.00 | 0.72 | 0.37 | 24 | Matrix Fibroblast |
| Actl6a | 0.32 | 0.00 | 0.00 | 0.33 | 0.11 | 24 | Matrix Fibroblast |
| Gm5641 | 0.55 | 0.00 | 0.00 | 0.69 | 0.36 | 24 | Matrix Fibroblast |
| Gm23493 | 0.28 | 0.00 | 0.00 | 0.27 | 0.08 | 24 | Matrix Fibroblast |
| Fam92a | 0.33 | 0.00 | 0.00 | 0.30 | 0.10 | 24 | Matrix Fibroblast |
| 9530068E07Rik | 0.38 | 0.00 | 0.00 | 0.44 | 0.17 | 24 | Matrix Fibroblast |
| Cacybp | 0.32 | 0.00 | 0.00 | 0.50 | 0.21 | 24 | Matrix Fibroblast |
| Hspa14 | 0.31 | 0.00 | 0.00 | 0.31 | 0.11 | 24 | Matrix Fibroblast |
| Anapc11 | 0.34 | 0.00 | 0.00 | 0.43 | 0.17 | 24 | Matrix Fibroblast |
| Sumo2 | 0.43 | 0.00 | 0.00 | 0.62 | 0.30 | 24 | Matrix Fibroblast |
| Map4k3 | 0.37 | 0.00 | 0.00 | 0.25 | 0.08 | 24 | Matrix Fibroblast |
| Gpx8 | 0.35 | 0.00 | 0.00 | 0.39 | 0.14 | 24 | Matrix Fibroblast |
| Pten | 0.40 | 0.00 | 0.00 | 0.34 | 0.12 | 24 | Matrix Fibroblast |
| Itgb1 | 0.41 | 0.00 | 0.00 | 0.90 | 0.63 | 24 | Matrix Fibroblast |
| Mapre1 | 0.38 | 0.00 | 0.00 | 0.62 | 0.29 | 24 | Matrix Fibroblast |
| Smchd1 | 0.38 | 0.00 | 0.00 | 0.41 | 0.16 | 24 | Matrix Fibroblast |
| Naa38 | 0.33 | 0.00 | 0.00 | 0.39 | 0.15 | 24 | Matrix Fibroblast |
| Phlda1 | 0.32 | 0.00 | 0.00 | 0.49 | 0.20 | 24 | Matrix Fibroblast |
| H1f0 | 0.50 | 0.00 | 0.00 | 0.57 | 0.27 | 24 | Matrix Fibroblast |
| Cyth3 | 0.33 | 0.00 | 0.00 | 0.57 | 0.25 | 24 | Matrix Fibroblast |
| Hars | 0.36 | 0.00 | 0.00 | 0.30 | 0.10 | 24 | Matrix Fibroblast |
| Morf4l2 | 0.44 | 0.00 | 0.00 | 0.51 | 0.22 | 24 | Matrix Fibroblast |
| Gm12355 | 0.46 | 0.00 | 0.00 | 0.57 | 0.26 | 24 | Matrix Fibroblast |
| Fkbp2 | 0.43 | 0.00 | 0.00 | 0.44 | 0.18 | 24 | Matrix Fibroblast |
| Rgs10 | 0.27 | 0.00 | 0.00 | 0.30 | 0.10 | 24 | Matrix Fibroblast |
| Taf9 | 0.37 | 0.00 | 0.00 | 0.45 | 0.18 | 24 | Matrix Fibroblast |
| Nudc | 0.41 | 0.00 | 0.00 | 0.48 | 0.21 | 24 | Matrix Fibroblast |
| Eng | 0.28 | 0.00 | 0.00 | 0.33 | 0.11 | 24 | Matrix Fibroblast |
| Fmr1 | 0.40 | 0.00 | 0.00 | 0.38 | 0.15 | 24 | Matrix Fibroblast |
| Pdap1 | 0.49 | 0.00 | 0.00 | 0.79 | 0.45 | 24 | Matrix Fibroblast |
| Tmsb10 | 0.34 | 0.00 | 0.00 | 0.99 | 0.86 | 24 | Matrix Fibroblast |
| Fundc2 | 0.34 | 0.00 | 0.00 | 0.42 | 0.17 | 24 | Matrix Fibroblast |
| Phf5a | 0.37 | 0.00 | 0.00 | 0.46 | 0.19 | 24 | Matrix Fibroblast |
| Gm6563 | 0.44 | 0.00 | 0.00 | 0.66 | 0.33 | 24 | Matrix Fibroblast |
| Nudt21 | 0.39 | 0.00 | 0.00 | 0.34 | 0.12 | 24 | Matrix Fibroblast |
| Oat | 0.31 | 0.00 | 0.00 | 0.35 | 0.13 | 24 | Matrix Fibroblast |
| Mxra7 | 0.28 | 0.00 | 0.00 | 0.38 | 0.14 | 24 | Matrix Fibroblast |
| Med19 | 0.35 | 0.00 | 0.00 | 0.32 | 0.11 | 24 | Matrix Fibroblast |
| Parva | 0.32 | 0.00 | 0.00 | 0.41 | 0.16 | 24 | Matrix Fibroblast |
| Snrpe | 0.43 | 0.00 | 0.00 | 0.69 | 0.36 | 24 | Matrix Fibroblast |
| 2-Sep | 0.44 | 0.00 | 0.00 | 0.67 | 0.35 | 24 | Matrix Fibroblast |
| Tgfb1i1 | 0.31 | 0.00 | 0.00 | 0.36 | 0.13 | 24 | Matrix Fibroblast |
| Nr3c1 | 0.39 | 0.00 | 0.00 | 0.56 | 0.25 | 24 | Matrix Fibroblast |
| Anapc5 | 0.38 | 0.00 | 0.00 | 0.44 | 0.18 | 24 | Matrix Fibroblast |
| Dynlt1b | 0.39 | 0.00 | 0.00 | 0.55 | 0.25 | 24 | Matrix Fibroblast |
| Tead1 | 0.33 | 0.00 | 0.00 | 0.37 | 0.14 | 24 | Matrix Fibroblast |
| Lsm6 | 0.37 | 0.00 | 0.00 | 0.35 | 0.13 | 24 | Matrix Fibroblast |
| Pole3 | 0.27 | 0.00 | 0.00 | 0.31 | 0.11 | 24 | Matrix Fibroblast |
| Vezf1 | 0.36 | 0.00 | 0.00 | 0.53 | 0.24 | 24 | Matrix Fibroblast |
| Picalm | 0.36 | 0.00 | 0.00 | 0.49 | 0.21 | 24 | Matrix Fibroblast |
| Set | 0.46 | 0.00 | 0.00 | 0.65 | 0.34 | 24 | Matrix Fibroblast |
| Hnrnpa3 | 0.47 | 0.00 | 0.00 | 0.78 | 0.46 | 24 | Matrix Fibroblast |
| Hnrnph1 | 0.41 | 0.00 | 0.00 | 0.73 | 0.38 | 24 | Matrix Fibroblast |
| Prnp | 0.39 | 0.00 | 0.00 | 0.33 | 0.12 | 24 | Matrix Fibroblast |
| Hmcn1 | 0.25 | 0.00 | 0.00 | 0.37 | 0.14 | 24 | Matrix Fibroblast |
| Brd3 | 0.42 | 0.00 | 0.00 | 0.54 | 0.25 | 24 | Matrix Fibroblast |
| Lyar | 0.38 | 0.00 | 0.00 | 0.39 | 0.15 | 24 | Matrix Fibroblast |
| Gm8991 | 0.44 | 0.00 | 0.00 | 0.51 | 0.23 | 24 | Matrix Fibroblast |
| Nutf2 | 0.29 | 0.00 | 0.00 | 0.35 | 0.13 | 24 | Matrix Fibroblast |
| Pagr1a | 0.31 | 0.00 | 0.00 | 0.26 | 0.09 | 24 | Matrix Fibroblast |
| Lap3 | 0.31 | 0.00 | 0.00 | 0.29 | 0.10 | 24 | Matrix Fibroblast |
| Ddx39b | 0.36 | 0.00 | 0.00 | 0.47 | 0.21 | 24 | Matrix Fibroblast |
| Ing1 | 0.33 | 0.00 | 0.00 | 0.39 | 0.15 | 24 | Matrix Fibroblast |
| Cox20 | 0.36 | 0.00 | 0.00 | 0.43 | 0.18 | 24 | Matrix Fibroblast |
| Pdia6 | 0.34 | 0.00 | 0.00 | 0.63 | 0.30 | 24 | Matrix Fibroblast |
| Ddost | 0.37 | 0.00 | 0.00 | 0.45 | 0.19 | 24 | Matrix Fibroblast |
| Rbm3 | 0.44 | 0.00 | 0.00 | 0.73 | 0.41 | 24 | Matrix Fibroblast |
| Sh3pxd2a | 0.34 | 0.00 | 0.00 | 0.29 | 0.10 | 24 | Matrix Fibroblast |
| Mrpl18 | 0.33 | 0.00 | 0.00 | 0.48 | 0.21 | 24 | Matrix Fibroblast |
| Ddx39 | 0.28 | 0.00 | 0.00 | 0.28 | 0.09 | 24 | Matrix Fibroblast |
| Txndc12 | 0.27 | 0.00 | 0.00 | 0.26 | 0.09 | 24 | Matrix Fibroblast |
| Dynlt1a | 0.33 | 0.00 | 0.00 | 0.52 | 0.23 | 24 | Matrix Fibroblast |
| Matr3 | 0.47 | 0.00 | 0.00 | 0.71 | 0.41 | 24 | Matrix Fibroblast |
| Sptssa | 0.31 | 0.00 | 0.00 | 0.47 | 0.20 | 24 | Matrix Fibroblast |
| Ndufa11 | 0.37 | 0.00 | 0.00 | 0.57 | 0.27 | 24 | Matrix Fibroblast |
| Cnot6 | 0.28 | 0.00 | 0.00 | 0.40 | 0.16 | 24 | Matrix Fibroblast |
| Kdelr2 | 0.38 | 0.00 | 0.00 | 0.45 | 0.19 | 24 | Matrix Fibroblast |
| Ttc3 | 0.38 | 0.00 | 0.00 | 0.60 | 0.29 | 24 | Matrix Fibroblast |
| Snrpd3 | 0.43 | 0.00 | 0.00 | 0.62 | 0.32 | 24 | Matrix Fibroblast |
| Marcks | 0.28 | 0.00 | 0.00 | 0.88 | 0.50 | 24 | Matrix Fibroblast |
| Fam115a | 0.36 | 0.00 | 0.00 | 0.36 | 0.14 | 24 | Matrix Fibroblast |
| Hnrnpdl | 0.36 | 0.00 | 0.00 | 0.71 | 0.37 | 24 | Matrix Fibroblast |
| Gtf2e2 | 0.29 | 0.00 | 0.00 | 0.28 | 0.09 | 24 | Matrix Fibroblast |
| 7-Sep | 0.38 | 0.00 | 0.00 | 0.87 | 0.59 | 24 | Matrix Fibroblast |
| Hint1 | 0.45 | 0.00 | 0.00 | 0.76 | 0.46 | 24 | Matrix Fibroblast |
| Nfyb | 0.31 | 0.00 | 0.00 | 0.32 | 0.12 | 24 | Matrix Fibroblast |
| Bzw2 | 0.28 | 0.00 | 0.00 | 0.38 | 0.15 | 24 | Matrix Fibroblast |
| Fgfr1 | 0.30 | 0.00 | 0.00 | 0.30 | 0.11 | 24 | Matrix Fibroblast |
| Nop56 | 0.34 | 0.00 | 0.00 | 0.46 | 0.20 | 24 | Matrix Fibroblast |
| Dpy30 | 0.32 | 0.00 | 0.00 | 0.37 | 0.14 | 24 | Matrix Fibroblast |
| Gm4204 | 0.36 | 0.00 | 0.00 | 0.80 | 0.50 | 24 | Matrix Fibroblast |
| Anxa6 | 0.31 | 0.00 | 0.00 | 0.47 | 0.20 | 24 | Matrix Fibroblast |
| Cfdp1 | 0.32 | 0.00 | 0.00 | 0.72 | 0.38 | 24 | Matrix Fibroblast |
| Maged1 | 0.26 | 0.00 | 0.00 | 0.45 | 0.18 | 24 | Matrix Fibroblast |
| Nedd4 | 0.38 | 0.00 | 0.00 | 0.85 | 0.53 | 24 | Matrix Fibroblast |
| Zfp131 | 0.30 | 0.00 | 0.00 | 0.33 | 0.12 | 24 | Matrix Fibroblast |
| Cct8 | 0.27 | 0.00 | 0.00 | 0.48 | 0.21 | 24 | Matrix Fibroblast |
| Epb4.1l2 | 0.29 | 0.00 | 0.00 | 0.45 | 0.19 | 24 | Matrix Fibroblast |
| Tra2b | 0.35 | 0.00 | 0.00 | 0.60 | 0.29 | 24 | Matrix Fibroblast |
| Tssc4 | 0.30 | 0.00 | 0.00 | 0.26 | 0.09 | 24 | Matrix Fibroblast |
| Entpd1 | 0.29 | 0.00 | 0.00 | 0.27 | 0.09 | 24 | Matrix Fibroblast |
| G3bp1 | 0.30 | 0.00 | 0.00 | 0.42 | 0.17 | 24 | Matrix Fibroblast |
| Hnrnpd | 0.39 | 0.00 | 0.00 | 0.57 | 0.28 | 24 | Matrix Fibroblast |
| Nutf2-ps1 | 0.28 | 0.00 | 0.00 | 0.30 | 0.11 | 24 | Matrix Fibroblast |
| Sec13 | 0.29 | 0.00 | 0.00 | 0.42 | 0.17 | 24 | Matrix Fibroblast |
| Snrpa1 | 0.27 | 0.00 | 0.00 | 0.37 | 0.14 | 24 | Matrix Fibroblast |
| Rnps1 | 0.27 | 0.00 | 0.00 | 0.38 | 0.15 | 24 | Matrix Fibroblast |
| Emp3 | 0.26 | 0.00 | 0.00 | 0.49 | 0.21 | 24 | Matrix Fibroblast |
| Anxa5 | 0.39 | 0.00 | 0.00 | 0.73 | 0.40 | 24 | Matrix Fibroblast |
| B230219D22Rik | 0.35 | 0.00 | 0.00 | 0.58 | 0.27 | 24 | Matrix Fibroblast |
| Snrpd2 | 0.38 | 0.00 | 0.00 | 0.53 | 0.25 | 24 | Matrix Fibroblast |
| Ccdc127 | 0.37 | 0.00 | 0.00 | 0.30 | 0.11 | 24 | Matrix Fibroblast |
| Mesdc2 | 0.38 | 0.00 | 0.00 | 0.43 | 0.19 | 24 | Matrix Fibroblast |
| Lgmn | 0.25 | 0.00 | 0.00 | 0.34 | 0.13 | 24 | Matrix Fibroblast |
| Fam98b | 0.26 | 0.00 | 0.00 | 0.29 | 0.10 | 24 | Matrix Fibroblast |
| Lman1 | 0.27 | 0.00 | 0.00 | 0.43 | 0.18 | 24 | Matrix Fibroblast |
| Rnf168 | 0.36 | 0.00 | 0.00 | 0.34 | 0.14 | 24 | Matrix Fibroblast |
| Scpep1 | 0.27 | 0.00 | 0.00 | 0.26 | 0.09 | 24 | Matrix Fibroblast |
| Cdc123 | 0.31 | 0.00 | 0.00 | 0.33 | 0.13 | 24 | Matrix Fibroblast |
| Ankrd32 | 0.37 | 0.00 | 0.00 | 0.29 | 0.11 | 24 | Matrix Fibroblast |
| Gm5160 | 0.33 | 0.00 | 0.00 | 0.48 | 0.22 | 24 | Matrix Fibroblast |
| Stip1 | 0.30 | 0.00 | 0.00 | 0.34 | 0.13 | 24 | Matrix Fibroblast |
| Srrt | 0.31 | 0.00 | 0.00 | 0.46 | 0.20 | 24 | Matrix Fibroblast |
| Ssb | 0.42 | 0.00 | 0.00 | 0.84 | 0.55 | 24 | Matrix Fibroblast |
| Tjp2 | 0.30 | 0.00 | 0.00 | 0.40 | 0.17 | 24 | Matrix Fibroblast |
| Fkbp7 | 0.28 | 0.00 | 0.00 | 0.34 | 0.13 | 24 | Matrix Fibroblast |
| Snrpf | 0.38 | 0.00 | 0.00 | 0.57 | 0.28 | 24 | Matrix Fibroblast |
| Leo1 | 0.33 | 0.00 | 0.00 | 0.33 | 0.13 | 24 | Matrix Fibroblast |
| Ctgf | 0.25 | 0.00 | 0.00 | 0.25 | 0.09 | 24 | Matrix Fibroblast |
| Gsk3b | 0.32 | 0.00 | 0.00 | 0.57 | 0.28 | 24 | Matrix Fibroblast |
| Ngfrap1 | 0.27 | 0.00 | 0.00 | 0.52 | 0.24 | 24 | Matrix Fibroblast |
| Ubald2 | 0.29 | 0.00 | 0.00 | 0.37 | 0.15 | 24 | Matrix Fibroblast |
| Cnep1r1 | 0.30 | 0.00 | 0.00 | 0.30 | 0.11 | 24 | Matrix Fibroblast |
| Acp1 | 0.33 | 0.00 | 0.00 | 0.35 | 0.14 | 24 | Matrix Fibroblast |
| Hnrnph3 | 0.35 | 0.00 | 0.00 | 0.41 | 0.18 | 24 | Matrix Fibroblast |
| Cnih1 | 0.34 | 0.00 | 0.00 | 0.43 | 0.19 | 24 | Matrix Fibroblast |
| Dynlt1f | 0.29 | 0.00 | 0.00 | 0.52 | 0.24 | 24 | Matrix Fibroblast |
| Dynlt1c | 0.33 | 0.00 | 0.00 | 0.56 | 0.27 | 24 | Matrix Fibroblast |
| Vbp1 | 0.34 | 0.00 | 0.00 | 0.37 | 0.15 | 24 | Matrix Fibroblast |
| Smarca4 | 0.35 | 0.00 | 0.00 | 0.40 | 0.17 | 24 | Matrix Fibroblast |
| Pnrc2 | 0.39 | 0.00 | 0.00 | 0.43 | 0.19 | 24 | Matrix Fibroblast |
| Stag1 | 0.29 | 0.00 | 0.00 | 0.32 | 0.12 | 24 | Matrix Fibroblast |
| Lsm5 | 0.31 | 0.00 | 0.00 | 0.31 | 0.12 | 24 | Matrix Fibroblast |
| Zfp260 | 0.30 | 0.00 | 0.00 | 0.33 | 0.13 | 24 | Matrix Fibroblast |
| Lsm4 | 0.34 | 0.00 | 0.00 | 0.45 | 0.21 | 24 | Matrix Fibroblast |
| Adprh | 0.25 | 0.00 | 0.00 | 0.31 | 0.12 | 24 | Matrix Fibroblast |
| Phldb2 | 0.31 | 0.00 | 0.00 | 0.59 | 0.29 | 24 | Matrix Fibroblast |
| Gm10093 | 0.26 | 0.00 | 0.00 | 0.44 | 0.19 | 24 | Matrix Fibroblast |
| Atp2b1 | 0.36 | 0.00 | 0.00 | 0.88 | 0.58 | 24 | Matrix Fibroblast |
| Sox4 | 0.28 | 0.00 | 0.00 | 0.69 | 0.37 | 24 | Matrix Fibroblast |
| Pdia4 | 0.33 | 0.00 | 0.00 | 0.46 | 0.21 | 24 | Matrix Fibroblast |
| Spin1 | 0.26 | 0.00 | 0.00 | 0.29 | 0.11 | 24 | Matrix Fibroblast |
| Srsf1 | 0.29 | 0.00 | 0.00 | 0.34 | 0.14 | 24 | Matrix Fibroblast |
| Pcnt | 0.35 | 0.00 | 0.00 | 0.34 | 0.14 | 24 | Matrix Fibroblast |
| Cd81 | 0.30 | 0.00 | 0.00 | 0.74 | 0.41 | 24 | Matrix Fibroblast |
| Thrap3 | 0.35 | 0.00 | 0.00 | 0.69 | 0.37 | 24 | Matrix Fibroblast |
| Topors | 0.29 | 0.00 | 0.00 | 0.30 | 0.12 | 24 | Matrix Fibroblast |
| Ptges3 | 0.35 | 0.00 | 0.00 | 0.50 | 0.25 | 24 | Matrix Fibroblast |
| Gm9242 | 0.39 | 0.00 | 0.00 | 0.49 | 0.24 | 24 | Matrix Fibroblast |
| Impad1 | 0.27 | 0.00 | 0.00 | 0.36 | 0.15 | 24 | Matrix Fibroblast |
| Lamc1 | 0.28 | 0.00 | 0.00 | 0.37 | 0.16 | 24 | Matrix Fibroblast |
| Mlf2 | 0.27 | 0.00 | 0.00 | 0.34 | 0.14 | 24 | Matrix Fibroblast |
| Twsg1 | 0.26 | 0.00 | 0.00 | 0.30 | 0.12 | 24 | Matrix Fibroblast |
| Ssr1 | 0.29 | 0.00 | 0.00 | 0.51 | 0.25 | 24 | Matrix Fibroblast |
| Nudt4 | 0.30 | 0.00 | 0.00 | 0.39 | 0.17 | 24 | Matrix Fibroblast |
| Nap1l1 | 0.29 | 0.00 | 0.00 | 0.77 | 0.47 | 24 | Matrix Fibroblast |
| Magt1 | 0.27 | 0.00 | 0.00 | 0.44 | 0.20 | 24 | Matrix Fibroblast |
| Smc5 | 0.26 | 0.00 | 0.00 | 0.38 | 0.16 | 24 | Matrix Fibroblast |
| Ywhah | 0.30 | 0.00 | 0.00 | 0.53 | 0.26 | 24 | Matrix Fibroblast |
| Mrfap1 | 0.33 | 0.00 | 0.00 | 0.62 | 0.33 | 24 | Matrix Fibroblast |
| Papola | 0.29 | 0.00 | 0.00 | 0.49 | 0.24 | 24 | Matrix Fibroblast |
| Psmd2 | 0.27 | 0.00 | 0.00 | 0.36 | 0.15 | 24 | Matrix Fibroblast |
| Hspa5 | 0.33 | 0.00 | 0.00 | 0.83 | 0.55 | 24 | Matrix Fibroblast |
| Cfl2 | 0.29 | 0.00 | 0.00 | 0.51 | 0.25 | 24 | Matrix Fibroblast |
| Dhx15 | 0.27 | 0.00 | 0.00 | 0.54 | 0.27 | 24 | Matrix Fibroblast |
| Vkorc1 | 0.32 | 0.00 | 0.00 | 0.27 | 0.10 | 24 | Matrix Fibroblast |
| Scoc | 0.35 | 0.00 | 0.00 | 0.31 | 0.13 | 24 | Matrix Fibroblast |
| 1110004F10Rik | 0.31 | 0.00 | 0.00 | 0.55 | 0.29 | 24 | Matrix Fibroblast |
| Wbp11 | 0.28 | 0.00 | 0.00 | 0.49 | 0.24 | 24 | Matrix Fibroblast |
| Gm10053 | 0.26 | 0.00 | 0.00 | 0.46 | 0.22 | 24 | Matrix Fibroblast |
| Hnrnpc | 0.30 | 0.00 | 0.00 | 0.75 | 0.44 | 24 | Matrix Fibroblast |
| Magoh | 0.27 | 0.00 | 0.00 | 0.37 | 0.16 | 24 | Matrix Fibroblast |
| Pcm1 | 0.29 | 0.00 | 0.00 | 0.54 | 0.27 | 24 | Matrix Fibroblast |
| Hp1bp3 | 0.37 | 0.00 | 0.00 | 0.73 | 0.44 | 24 | Matrix Fibroblast |
| Ift74 | 0.25 | 0.00 | 0.00 | 0.26 | 0.10 | 24 | Matrix Fibroblast |
| Snrpb | 0.31 | 0.00 | 0.00 | 0.65 | 0.35 | 24 | Matrix Fibroblast |
| Wbp5 | 0.26 | 0.00 | 0.00 | 0.79 | 0.48 | 24 | Matrix Fibroblast |
| Rfc1 | 0.33 | 0.00 | 0.00 | 0.43 | 0.21 | 24 | Matrix Fibroblast |
| Nap1l4 | 0.27 | 0.00 | 0.00 | 0.53 | 0.26 | 24 | Matrix Fibroblast |
| Prdx2 | 0.30 | 0.00 | 0.00 | 0.62 | 0.33 | 24 | Matrix Fibroblast |
| Mrpl42 | 0.26 | 0.00 | 0.00 | 0.32 | 0.13 | 24 | Matrix Fibroblast |
| Hnrnpa0 | 0.35 | 0.00 | 0.00 | 0.44 | 0.21 | 24 | Matrix Fibroblast |
| Eif4a1 | 0.31 | 0.00 | 0.00 | 0.65 | 0.36 | 24 | Matrix Fibroblast |
| Gm26735 | 0.30 | 0.00 | 0.00 | 0.41 | 0.19 | 24 | Matrix Fibroblast |
| Pebp1 | 0.26 | 0.00 | 0.00 | 0.51 | 0.25 | 24 | Matrix Fibroblast |
| Lrrcc1 | 0.32 | 0.00 | 0.00 | 0.29 | 0.12 | 24 | Matrix Fibroblast |
| Jmy | 0.32 | 0.00 | 0.00 | 0.29 | 0.12 | 24 | Matrix Fibroblast |
| Arf4 | 0.30 | 0.00 | 0.00 | 0.53 | 0.27 | 24 | Matrix Fibroblast |
| Tial1 | 0.27 | 0.00 | 0.00 | 0.46 | 0.22 | 24 | Matrix Fibroblast |
| Ncl | 0.29 | 0.00 | 0.00 | 0.97 | 0.78 | 24 | Matrix Fibroblast |
| Cnih4 | 0.26 | 0.00 | 0.00 | 0.34 | 0.14 | 24 | Matrix Fibroblast |
| Myo1b | 0.27 | 0.00 | 0.00 | 0.38 | 0.17 | 24 | Matrix Fibroblast |
| Suz12 | 0.31 | 0.00 | 0.00 | 0.31 | 0.13 | 24 | Matrix Fibroblast |
| Ssr2 | 0.29 | 0.00 | 0.00 | 0.58 | 0.31 | 24 | Matrix Fibroblast |
| Srsf10 | 0.29 | 0.00 | 0.00 | 0.50 | 0.25 | 24 | Matrix Fibroblast |
| Calr | 0.40 | 0.00 | 0.00 | 0.69 | 0.43 | 24 | Matrix Fibroblast |
| Hdac2 | 0.26 | 0.00 | 0.00 | 0.35 | 0.15 | 24 | Matrix Fibroblast |
| Bcas2 | 0.25 | 0.00 | 0.00 | 0.44 | 0.21 | 24 | Matrix Fibroblast |
| Mtus1 | 0.28 | 0.00 | 0.00 | 0.38 | 0.17 | 24 | Matrix Fibroblast |
| Asf1a | 0.28 | 0.00 | 0.00 | 0.27 | 0.11 | 24 | Matrix Fibroblast |
| Khdrbs1 | 0.33 | 0.00 | 0.00 | 0.57 | 0.31 | 24 | Matrix Fibroblast |
| Rsrc1 | 0.25 | 0.00 | 0.00 | 0.33 | 0.14 | 24 | Matrix Fibroblast |
| Hnrnpu | 0.33 | 0.00 | 0.00 | 0.89 | 0.64 | 24 | Matrix Fibroblast |
| Uqcr10 | 0.30 | 0.00 | 0.00 | 0.63 | 0.36 | 24 | Matrix Fibroblast |
| Eif4g2 | 0.29 | 0.00 | 0.00 | 0.70 | 0.41 | 24 | Matrix Fibroblast |
| Tceb2 | 0.30 | 0.00 | 0.00 | 0.76 | 0.47 | 24 | Matrix Fibroblast |
| Dynll1 | 0.31 | 0.00 | 0.00 | 0.73 | 0.45 | 24 | Matrix Fibroblast |
| Pbrm1 | 0.30 | 0.00 | 0.00 | 0.56 | 0.30 | 24 | Matrix Fibroblast |
| Vps36 | 0.31 | 0.00 | 0.00 | 0.54 | 0.29 | 24 | Matrix Fibroblast |
| Dhx9 | 0.26 | 0.00 | 0.00 | 0.40 | 0.19 | 24 | Matrix Fibroblast |
| Eprs | 0.25 | 0.00 | 0.00 | 0.46 | 0.22 | 24 | Matrix Fibroblast |
| Pcnp | 0.27 | 0.00 | 0.00 | 0.42 | 0.20 | 24 | Matrix Fibroblast |
| Serbp1 | 0.28 | 0.00 | 0.00 | 0.93 | 0.75 | 24 | Matrix Fibroblast |
| Ube2n | 0.26 | 0.00 | 0.00 | 0.34 | 0.15 | 24 | Matrix Fibroblast |
| Sarnp | 0.30 | 0.00 | 0.00 | 0.49 | 0.25 | 24 | Matrix Fibroblast |
| Ccar1 | 0.28 | 0.00 | 0.00 | 0.60 | 0.33 | 24 | Matrix Fibroblast |
| Siva1 | 0.30 | 0.00 | 0.00 | 0.33 | 0.15 | 24 | Matrix Fibroblast |
| Tardbp | 0.29 | 0.00 | 0.00 | 0.56 | 0.31 | 24 | Matrix Fibroblast |
| Phip | 0.27 | 0.00 | 0.00 | 0.35 | 0.16 | 24 | Matrix Fibroblast |
| Ppp2r2d | 0.25 | 0.00 | 0.00 | 0.26 | 0.11 | 24 | Matrix Fibroblast |
| Lpp | 0.27 | 0.00 | 0.00 | 0.43 | 0.22 | 24 | Matrix Fibroblast |
| Cox7a2 | 0.27 | 0.00 | 0.00 | 0.77 | 0.48 | 24 | Matrix Fibroblast |
| Cux1 | 0.27 | 0.00 | 0.00 | 0.35 | 0.17 | 24 | Matrix Fibroblast |
| Canx | 0.26 | 0.00 | 0.00 | 0.78 | 0.49 | 24 | Matrix Fibroblast |
| Hnrnpm | 0.26 | 0.00 | 0.00 | 0.77 | 0.49 | 24 | Matrix Fibroblast |
| Gclm | 0.34 | 0.00 | 0.00 | 0.66 | 0.42 | 24 | Matrix Fibroblast |
| Sf3b2 | 0.25 | 0.00 | 0.00 | 0.67 | 0.40 | 24 | Matrix Fibroblast |
| Npm1 | 0.31 | 0.00 | 0.00 | 0.81 | 0.58 | 24 | Matrix Fibroblast |
| Glg1 | 0.25 | 0.00 | 0.00 | 0.39 | 0.19 | 24 | Matrix Fibroblast |
| Ostc | 0.25 | 0.00 | 0.00 | 0.45 | 0.23 | 24 | Matrix Fibroblast |
| Ywhaq | 0.30 | 0.00 | 0.00 | 0.61 | 0.37 | 24 | Matrix Fibroblast |
| Ubb | 0.26 | 0.00 | 0.00 | 0.95 | 0.81 | 24 | Matrix Fibroblast |
| Chd4 | 0.26 | 0.00 | 0.00 | 0.76 | 0.48 | 24 | Matrix Fibroblast |
| Gm4076 | 0.25 | 0.00 | 0.00 | 0.41 | 0.21 | 24 | Matrix Fibroblast |
| Hdgfrp2 | 0.26 | 0.00 | 0.00 | 0.26 | 0.12 | 24 | Matrix Fibroblast |
| Prdx1 | 0.27 | 0.00 | 0.00 | 0.75 | 0.50 | 24 | Matrix Fibroblast |
| Smarca5 | 0.31 | 0.00 | 0.00 | 0.52 | 0.31 | 24 | Matrix Fibroblast |
| Pdia3 | 0.26 | 0.00 | 0.00 | 0.69 | 0.44 | 24 | Matrix Fibroblast |
| Psmc3 | 0.27 | 0.00 | 0.00 | 0.50 | 0.29 | 24 | Matrix Fibroblast |
| Tcp1 | 0.25 | 0.00 | 0.00 | 0.43 | 0.24 | 24 | Matrix Fibroblast |
| Cox6c | 0.29 | 0.00 | 0.00 | 0.84 | 0.65 | 24 | Matrix Fibroblast |
| Zbtb20 | 0.26 | 0.00 | 0.00 | 0.48 | 0.27 | 24 | Matrix Fibroblast |
| mt-Nd5 | 0.27 | 0.00 | 0.00 | 0.93 | 0.76 | 24 | Matrix Fibroblast |
| mt-Nd2 | 0.26 | 0.00 | 0.00 | 0.88 | 0.70 | 24 | Matrix Fibroblast |
| Hist1h2ao | 2.51 | 0.00 | 0.00 | 0.92 | 0.15 | 25 | T Cell |
| Hist1h2ap | 2.49 | 0.00 | 0.00 | 0.92 | 0.15 | 25 | T Cell |
| Hist1h2ag | 2.34 | 0.00 | 0.00 | 0.87 | 0.11 | 25 | T Cell |
| Top2a | 2.27 | 0.00 | 0.00 | 0.94 | 0.11 | 25 | T Cell |
| Hist1h2ac | 2.24 | 0.00 | 0.00 | 0.87 | 0.10 | 25 | T Cell |
| Hist1h2ad | 2.22 | 0.00 | 0.00 | 0.85 | 0.10 | 25 | T Cell |
| Hist1h2ah | 2.20 | 0.00 | 0.00 | 0.85 | 0.09 | 25 | T Cell |
| Trbc2 | 2.17 | 0.00 | 0.00 | 0.86 | 0.04 | 25 | T Cell |
| Hist1h2ai | 2.16 | 0.00 | 0.00 | 0.83 | 0.10 | 25 | T Cell |
| Hist1h2an | 2.16 | 0.00 | 0.00 | 0.81 | 0.09 | 25 | T Cell |
| Mki67 | 2.09 | 0.00 | 0.00 | 0.95 | 0.12 | 25 | T Cell |
| Hist1h2af | 2.09 | 0.00 | 0.00 | 0.83 | 0.09 | 25 | T Cell |
| 2810417H13Rik | 2.08 | 0.00 | 0.00 | 0.86 | 0.08 | 25 | T Cell |
| Hist1h2ab | 2.03 | 0.00 | 0.00 | 0.84 | 0.09 | 25 | T Cell |
| Hist1h2ae | 2.02 | 0.00 | 0.00 | 0.81 | 0.09 | 25 | T Cell |
| Ms4a4b | 2.00 | 0.00 | 0.00 | 0.67 | 0.05 | 25 | T Cell |
| Hist1h2ak | 1.95 | 0.00 | 0.00 | 0.78 | 0.07 | 25 | T Cell |
| Trbc1 | 1.94 | 0.00 | 0.00 | 0.78 | 0.03 | 25 | T Cell |
| Nusap1 | 1.83 | 0.00 | 0.00 | 0.73 | 0.06 | 25 | T Cell |
| Birc5 | 1.77 | 0.00 | 0.00 | 0.81 | 0.06 | 25 | T Cell |
| H2afx | 1.73 | 0.00 | 0.00 | 0.80 | 0.09 | 25 | T Cell |
| Ube2c | 1.60 | 0.00 | 0.00 | 0.62 | 0.05 | 25 | T Cell |
| Cenpe | 1.55 | 0.00 | 0.00 | 0.73 | 0.08 | 25 | T Cell |
| Incenp | 1.54 | 0.00 | 0.00 | 0.73 | 0.08 | 25 | T Cell |
| Hmmr | 1.52 | 0.00 | 0.00 | 0.61 | 0.04 | 25 | T Cell |
| Hist1h1b | 1.50 | 0.00 | 0.00 | 0.49 | 0.04 | 25 | T Cell |
| Cd3g | 1.49 | 0.00 | 0.00 | 0.65 | 0.03 | 25 | T Cell |
| Cdca8 | 1.44 | 0.00 | 0.00 | 0.66 | 0.05 | 25 | T Cell |
| Rrm2 | 1.41 | 0.00 | 0.00 | 0.58 | 0.05 | 25 | T Cell |
| RP23-45G16.5 | 1.40 | 0.00 | 0.00 | 0.64 | 0.05 | 25 | T Cell |
| Cd3d | 1.39 | 0.00 | 0.00 | 0.58 | 0.03 | 25 | T Cell |
| Prc1 | 1.38 | 0.00 | 0.00 | 0.67 | 0.06 | 25 | T Cell |
| Nkg7 | 1.37 | 0.00 | 0.00 | 0.38 | 0.02 | 25 | T Cell |
| Ccnb2 | 1.37 | 0.00 | 0.00 | 0.67 | 0.06 | 25 | T Cell |
| Thy1 | 1.36 | 0.00 | 0.00 | 0.57 | 0.02 | 25 | T Cell |
| Ncapg | 1.35 | 0.00 | 0.00 | 0.61 | 0.04 | 25 | T Cell |
| Ptprcap | 1.35 | 0.00 | 0.00 | 0.77 | 0.07 | 25 | T Cell |
| Tpx2 | 1.26 | 0.00 | 0.00 | 0.61 | 0.05 | 25 | T Cell |
| Kif11 | 1.22 | 0.00 | 0.00 | 0.55 | 0.04 | 25 | T Cell |
| Esco2 | 1.22 | 0.00 | 0.00 | 0.49 | 0.03 | 25 | T Cell |
| Kif15 | 1.20 | 0.00 | 0.00 | 0.54 | 0.04 | 25 | T Cell |
| Ndc80 | 1.20 | 0.00 | 0.00 | 0.53 | 0.04 | 25 | T Cell |
| Skap1 | 1.18 | 0.00 | 0.00 | 0.53 | 0.02 | 25 | T Cell |
| Selplg | 1.16 | 0.00 | 0.00 | 0.61 | 0.05 | 25 | T Cell |
| Cdca3 | 1.14 | 0.00 | 0.00 | 0.52 | 0.03 | 25 | T Cell |
| Asf1b | 1.02 | 0.00 | 0.00 | 0.45 | 0.02 | 25 | T Cell |
| Bcl11b | 1.00 | 0.00 | 0.00 | 0.38 | 0.02 | 25 | T Cell |
| Itgb7 | 0.97 | 0.00 | 0.00 | 0.49 | 0.03 | 25 | T Cell |
| Lck | 0.97 | 0.00 | 0.00 | 0.43 | 0.02 | 25 | T Cell |
| Spn | 0.94 | 0.00 | 0.00 | 0.45 | 0.03 | 25 | T Cell |
| Aurkb | 0.88 | 0.00 | 0.00 | 0.41 | 0.03 | 25 | T Cell |
| Icos | 0.87 | 0.00 | 0.00 | 0.28 | 0.01 | 25 | T Cell |
| Ctsw | 0.77 | 0.00 | 0.00 | 0.29 | 0.01 | 25 | T Cell |
| Hist1h3c | 0.75 | 0.00 | 0.00 | 0.32 | 0.01 | 25 | T Cell |
| Cenpf | 1.57 | 0.00 | 0.00 | 0.70 | 0.08 | 25 | T Cell |
| Fbxo5 | 1.16 | 0.00 | 0.00 | 0.54 | 0.04 | 25 | T Cell |
| Cd3e | 0.73 | 0.00 | 0.00 | 0.31 | 0.01 | 25 | T Cell |
| Trac | 0.75 | 0.00 | 0.00 | 0.25 | 0.01 | 25 | T Cell |
| 1-Sep | 1.05 | 0.00 | 0.00 | 0.50 | 0.04 | 25 | T Cell |
| Lat | 0.75 | 0.00 | 0.00 | 0.33 | 0.02 | 25 | T Cell |
| Mis18bp1 | 1.09 | 0.00 | 0.00 | 0.46 | 0.03 | 25 | T Cell |
| Klrd1 | 1.35 | 0.00 | 0.00 | 0.30 | 0.01 | 25 | T Cell |
| Ccna2 | 1.04 | 0.00 | 0.00 | 0.48 | 0.04 | 25 | T Cell |
| Casc5 | 1.17 | 0.00 | 0.00 | 0.53 | 0.04 | 25 | T Cell |
| Sgol1 | 0.90 | 0.00 | 0.00 | 0.42 | 0.03 | 25 | T Cell |
| Cks2 | 1.36 | 0.00 | 0.00 | 0.69 | 0.08 | 25 | T Cell |
| Smc2 | 1.60 | 0.00 | 0.00 | 0.87 | 0.13 | 25 | T Cell |
| Rac2 | 1.27 | 0.00 | 0.00 | 0.81 | 0.11 | 25 | T Cell |
| Clspn | 1.22 | 0.00 | 0.00 | 0.49 | 0.04 | 25 | T Cell |
| Tacc3 | 1.22 | 0.00 | 0.00 | 0.56 | 0.05 | 25 | T Cell |
| Cenpa | 1.33 | 0.00 | 0.00 | 0.60 | 0.06 | 25 | T Cell |
| Ccnb1 | 0.82 | 0.00 | 0.00 | 0.36 | 0.02 | 25 | T Cell |
| AW112010 | 1.57 | 0.00 | 0.00 | 0.48 | 0.04 | 25 | T Cell |
| Cdc20 | 1.18 | 0.00 | 0.00 | 0.51 | 0.05 | 25 | T Cell |
| Cep55 | 0.78 | 0.00 | 0.00 | 0.39 | 0.03 | 25 | T Cell |
| Ptpn7 | 0.81 | 0.00 | 0.00 | 0.38 | 0.03 | 25 | T Cell |
| Cd2 | 1.05 | 0.00 | 0.00 | 0.49 | 0.04 | 25 | T Cell |
| Racgap1 | 0.99 | 0.00 | 0.00 | 0.45 | 0.04 | 25 | T Cell |
| Dlgap5 | 0.89 | 0.00 | 0.00 | 0.36 | 0.03 | 25 | T Cell |
| Ska1 | 0.73 | 0.00 | 0.00 | 0.32 | 0.02 | 25 | T Cell |
| Aspm | 1.01 | 0.00 | 0.00 | 0.40 | 0.03 | 25 | T Cell |
| Spc24 | 0.98 | 0.00 | 0.00 | 0.49 | 0.05 | 25 | T Cell |
| Bub1 | 0.76 | 0.00 | 0.00 | 0.35 | 0.03 | 25 | T Cell |
| Ptprc | 1.39 | 0.00 | 0.00 | 0.87 | 0.14 | 25 | T Cell |
| Sgol2 | 1.04 | 0.00 | 0.00 | 0.45 | 0.04 | 25 | T Cell |
| Nuf2 | 0.85 | 0.00 | 0.00 | 0.40 | 0.03 | 25 | T Cell |
| Kif23 | 1.01 | 0.00 | 0.00 | 0.52 | 0.05 | 25 | T Cell |
| Spc25 | 1.05 | 0.00 | 0.00 | 0.49 | 0.05 | 25 | T Cell |
| D17H6S56E-5 | 1.43 | 0.00 | 0.00 | 0.71 | 0.10 | 25 | T Cell |
| Ikzf2 | 0.80 | 0.00 | 0.00 | 0.31 | 0.02 | 25 | T Cell |
| Cenph | 0.90 | 0.00 | 0.00 | 0.40 | 0.03 | 25 | T Cell |
| Kif22 | 0.85 | 0.00 | 0.00 | 0.34 | 0.02 | 25 | T Cell |
| Rad51 | 0.71 | 0.00 | 0.00 | 0.34 | 0.03 | 25 | T Cell |
| Cd52 | 1.49 | 0.00 | 0.00 | 0.96 | 0.20 | 25 | T Cell |
| Ncapd2 | 1.01 | 0.00 | 0.00 | 0.45 | 0.04 | 25 | T Cell |
| Cks1b | 1.18 | 0.00 | 0.00 | 0.65 | 0.09 | 25 | T Cell |
| Plk1 | 0.60 | 0.00 | 0.00 | 0.27 | 0.02 | 25 | T Cell |
| Cd247 | 0.64 | 0.00 | 0.00 | 0.26 | 0.01 | 25 | T Cell |
| Kif20b | 1.19 | 0.00 | 0.00 | 0.54 | 0.07 | 25 | T Cell |
| Ckap2l | 0.93 | 0.00 | 0.00 | 0.45 | 0.05 | 25 | T Cell |
| Lmnb1 | 1.16 | 0.00 | 0.00 | 0.61 | 0.08 | 25 | T Cell |
| Arhgap11a | 0.94 | 0.00 | 0.00 | 0.43 | 0.04 | 25 | T Cell |
| Cenpl | 0.68 | 0.00 | 0.00 | 0.32 | 0.02 | 25 | T Cell |
| Cdk1 | 1.01 | 0.00 | 0.00 | 0.47 | 0.05 | 25 | T Cell |
| Bin2 | 0.85 | 0.00 | 0.00 | 0.44 | 0.04 | 25 | T Cell |
| Mad2l1 | 0.70 | 0.00 | 0.00 | 0.32 | 0.02 | 25 | T Cell |
| Hmgb2 | 2.04 | 0.00 | 0.00 | 1.00 | 0.42 | 25 | T Cell |
| Tbc1d10c | 0.65 | 0.00 | 0.00 | 0.35 | 0.03 | 25 | T Cell |
| Cd27 | 0.67 | 0.00 | 0.00 | 0.30 | 0.02 | 25 | T Cell |
| Ms4a6b | 1.09 | 0.00 | 0.00 | 0.50 | 0.06 | 25 | T Cell |
| Hcst | 0.72 | 0.00 | 0.00 | 0.39 | 0.04 | 25 | T Cell |
| Hist1h4i | 0.59 | 0.00 | 0.00 | 0.30 | 0.02 | 25 | T Cell |
| Pycard | 1.17 | 0.00 | 0.00 | 0.59 | 0.09 | 25 | T Cell |
| Cdca2 | 0.57 | 0.00 | 0.00 | 0.27 | 0.02 | 25 | T Cell |
| Diap3 | 0.82 | 0.00 | 0.00 | 0.48 | 0.06 | 25 | T Cell |
| Arhgdib | 1.11 | 0.00 | 0.00 | 0.80 | 0.16 | 25 | T Cell |
| Dnajc9 | 1.16 | 0.00 | 0.00 | 0.67 | 0.12 | 25 | T Cell |
| Depdc1a | 0.66 | 0.00 | 0.00 | 0.26 | 0.02 | 25 | T Cell |
| Knstrn | 0.85 | 0.00 | 0.00 | 0.37 | 0.04 | 25 | T Cell |
| C330027C09Rik | 0.66 | 0.00 | 0.00 | 0.32 | 0.03 | 25 | T Cell |
| Phgdh | 0.61 | 0.00 | 0.00 | 0.29 | 0.02 | 25 | T Cell |
| Gm10184 | 1.06 | 0.00 | 0.00 | 0.48 | 0.06 | 25 | T Cell |
| Trim59 | 0.82 | 0.00 | 0.00 | 0.39 | 0.04 | 25 | T Cell |
| Gmnn | 0.98 | 0.00 | 0.00 | 0.50 | 0.07 | 25 | T Cell |
| Gata3 | 0.75 | 0.00 | 0.00 | 0.28 | 0.02 | 25 | T Cell |
| Rrm1 | 0.88 | 0.00 | 0.00 | 0.48 | 0.06 | 25 | T Cell |
| Gm10259 | 0.92 | 0.00 | 0.00 | 0.46 | 0.06 | 25 | T Cell |
| Tyms | 0.93 | 0.00 | 0.00 | 0.44 | 0.05 | 25 | T Cell |
| Iqgap2 | 0.75 | 0.00 | 0.00 | 0.37 | 0.04 | 25 | T Cell |
| Cd48 | 0.78 | 0.00 | 0.00 | 0.41 | 0.05 | 25 | T Cell |
| 6-Sep | 0.82 | 0.00 | 0.00 | 0.49 | 0.07 | 25 | T Cell |
| Ikzf3 | 0.58 | 0.00 | 0.00 | 0.27 | 0.02 | 25 | T Cell |
| Was | 0.71 | 0.00 | 0.00 | 0.43 | 0.05 | 25 | T Cell |
| Kpna2 | 0.92 | 0.00 | 0.00 | 0.45 | 0.06 | 25 | T Cell |
| Ezh2 | 1.23 | 0.00 | 0.00 | 0.78 | 0.19 | 25 | T Cell |
| Dut | 1.01 | 0.00 | 0.00 | 0.54 | 0.09 | 25 | T Cell |
| Rad51ap1 | 0.73 | 0.00 | 0.00 | 0.33 | 0.03 | 25 | T Cell |
| Kif4 | 0.67 | 0.00 | 0.00 | 0.28 | 0.03 | 25 | T Cell |
| Dnmt1 | 1.03 | 0.00 | 0.00 | 0.59 | 0.10 | 25 | T Cell |
| Stmn1 | 1.22 | 0.00 | 0.00 | 0.87 | 0.23 | 25 | T Cell |
| Bub1b | 0.67 | 0.00 | 0.00 | 0.27 | 0.02 | 25 | T Cell |
| Il2rg | 0.84 | 0.00 | 0.00 | 0.58 | 0.10 | 25 | T Cell |
| Kif20a | 0.60 | 0.00 | 0.00 | 0.26 | 0.02 | 25 | T Cell |
| Mns1 | 0.80 | 0.00 | 0.00 | 0.36 | 0.04 | 25 | T Cell |
| Smc4 | 1.52 | 0.00 | 0.00 | 0.94 | 0.34 | 25 | T Cell |
| Prim1 | 0.81 | 0.00 | 0.00 | 0.41 | 0.05 | 25 | T Cell |
| Il7r | 0.76 | 0.00 | 0.00 | 0.28 | 0.03 | 25 | T Cell |
| Dbf4 | 0.78 | 0.00 | 0.00 | 0.40 | 0.05 | 25 | T Cell |
| Epsti1 | 0.83 | 0.00 | 0.00 | 0.35 | 0.04 | 25 | T Cell |
| Coro1a | 0.90 | 0.00 | 0.00 | 0.57 | 0.10 | 25 | T Cell |
| AI662270 | 0.81 | 0.00 | 0.00 | 0.48 | 0.07 | 25 | T Cell |
| Rbm38 | 0.59 | 0.00 | 0.00 | 0.30 | 0.03 | 25 | T Cell |
| Psmb8 | 1.02 | 0.00 | 0.00 | 0.78 | 0.18 | 25 | T Cell |
| H2afz | 1.41 | 0.00 | 0.00 | 0.95 | 0.40 | 25 | T Cell |
| Spdl1 | 0.64 | 0.00 | 0.00 | 0.25 | 0.02 | 25 | T Cell |
| Lcp1 | 1.11 | 0.00 | 0.00 | 0.85 | 0.22 | 25 | T Cell |
| Hist1h1e | 1.55 | 0.00 | 0.00 | 0.70 | 0.16 | 25 | T Cell |
| Fgl2 | 0.79 | 0.00 | 0.00 | 0.33 | 0.04 | 25 | T Cell |
| Anp32e | 1.24 | 0.00 | 0.00 | 0.90 | 0.28 | 25 | T Cell |
| Tk1 | 0.57 | 0.00 | 0.00 | 0.31 | 0.03 | 25 | T Cell |
| Ikzf1 | 0.58 | 0.00 | 0.00 | 0.32 | 0.04 | 25 | T Cell |
| Pbk | 0.55 | 0.00 | 0.00 | 0.30 | 0.03 | 25 | T Cell |
| Fam107b | 0.81 | 0.00 | 0.00 | 0.45 | 0.07 | 25 | T Cell |
| Laptm5 | 0.73 | 0.00 | 0.00 | 0.68 | 0.13 | 25 | T Cell |
| Hist2h2ac | 0.89 | 0.00 | 0.00 | 0.42 | 0.06 | 25 | T Cell |
| Cenpw | 0.65 | 0.00 | 0.00 | 0.37 | 0.05 | 25 | T Cell |
| Hist1h1d | 1.30 | 0.00 | 0.00 | 0.53 | 0.10 | 25 | T Cell |
| Cenpk | 0.72 | 0.00 | 0.00 | 0.31 | 0.04 | 25 | T Cell |
| Lig1 | 1.08 | 0.00 | 0.00 | 0.52 | 0.09 | 25 | T Cell |
| Ncapg2 | 0.53 | 0.00 | 0.00 | 0.28 | 0.03 | 25 | T Cell |
| Mcm5 | 0.65 | 0.00 | 0.00 | 0.31 | 0.04 | 25 | T Cell |
| Atad2 | 0.92 | 0.00 | 0.00 | 0.47 | 0.08 | 25 | T Cell |
| Ckap2 | 0.62 | 0.00 | 0.00 | 0.35 | 0.04 | 25 | T Cell |
| Arhgap9 | 0.59 | 0.00 | 0.00 | 0.32 | 0.04 | 25 | T Cell |
| Rfc4 | 0.52 | 0.00 | 0.00 | 0.31 | 0.04 | 25 | T Cell |
| Cdc25b | 0.72 | 0.00 | 0.00 | 0.34 | 0.04 | 25 | T Cell |
| Def6 | 0.52 | 0.00 | 0.00 | 0.26 | 0.03 | 25 | T Cell |
| Ms4a4c | 0.68 | 0.00 | 0.00 | 0.29 | 0.03 | 25 | T Cell |
| Hist2h2aa2 | 1.05 | 0.00 | 0.00 | 0.60 | 0.14 | 25 | T Cell |
| Ccr7 | 0.97 | 0.00 | 0.00 | 0.25 | 0.03 | 25 | T Cell |
| Sell | 0.79 | 0.00 | 0.00 | 0.31 | 0.04 | 25 | T Cell |
| Coro2a | 0.87 | 0.00 | 0.00 | 0.41 | 0.06 | 25 | T Cell |
| Pcna-ps2 | 1.32 | 0.00 | 0.00 | 0.67 | 0.18 | 25 | T Cell |
| Arhgap15 | 0.66 | 0.00 | 0.00 | 0.35 | 0.05 | 25 | T Cell |
| Cd53 | 0.72 | 0.00 | 0.00 | 0.51 | 0.09 | 25 | T Cell |
| Gimap9 | 0.82 | 0.00 | 0.00 | 0.43 | 0.07 | 25 | T Cell |
| Cotl1 | 0.97 | 0.00 | 0.00 | 0.69 | 0.17 | 25 | T Cell |
| Cytip | 0.81 | 0.00 | 0.00 | 0.50 | 0.09 | 25 | T Cell |
| Ptma | 0.84 | 0.00 | 0.00 | 1.00 | 0.91 | 25 | T Cell |
| Lbr | 0.88 | 0.00 | 0.00 | 0.56 | 0.12 | 25 | T Cell |
| Mthfd2 | 0.55 | 0.00 | 0.00 | 0.26 | 0.03 | 25 | T Cell |
| Tubb5 | 1.10 | 0.00 | 0.00 | 0.94 | 0.44 | 25 | T Cell |
| Itgb2 | 0.51 | 0.00 | 0.00 | 0.33 | 0.04 | 25 | T Cell |
| Plk4 | 0.60 | 0.00 | 0.00 | 0.31 | 0.04 | 25 | T Cell |
| Lef1 | 0.65 | 0.00 | 0.00 | 0.30 | 0.04 | 25 | T Cell |
| Camk4 | 0.64 | 0.00 | 0.00 | 0.29 | 0.04 | 25 | T Cell |
| Fen1 | 0.57 | 0.00 | 0.00 | 0.30 | 0.04 | 25 | T Cell |
| Pcna | 1.28 | 0.00 | 0.00 | 0.64 | 0.18 | 25 | T Cell |
| Mcm6 | 0.73 | 0.00 | 0.00 | 0.41 | 0.07 | 25 | T Cell |
| Traf3ip3 | 0.52 | 0.00 | 0.00 | 0.28 | 0.03 | 25 | T Cell |
| Hist2h2ab | 0.72 | 0.00 | 0.00 | 0.30 | 0.04 | 25 | T Cell |
| Nucks1 | 1.03 | 0.00 | 0.00 | 0.83 | 0.28 | 25 | T Cell |
| Tuba1b | 1.08 | 0.00 | 0.00 | 0.94 | 0.45 | 25 | T Cell |
| Slbp | 1.04 | 0.00 | 0.00 | 0.59 | 0.15 | 25 | T Cell |
| Cdc7 | 0.50 | 0.00 | 0.00 | 0.25 | 0.03 | 25 | T Cell |
| Ran | 1.04 | 0.00 | 0.00 | 0.91 | 0.38 | 25 | T Cell |
| Mcm3 | 0.65 | 0.00 | 0.00 | 0.41 | 0.07 | 25 | T Cell |
| Apobec3 | 0.75 | 0.00 | 0.00 | 0.43 | 0.08 | 25 | T Cell |
| Stk10 | 0.50 | 0.00 | 0.00 | 0.32 | 0.05 | 25 | T Cell |
| S1pr4 | 0.61 | 0.00 | 0.00 | 0.29 | 0.04 | 25 | T Cell |
| Cdkn2d | 0.77 | 0.00 | 0.00 | 0.39 | 0.07 | 25 | T Cell |
| Ddx39 | 0.67 | 0.00 | 0.00 | 0.46 | 0.09 | 25 | T Cell |
| Gm10282 | 1.00 | 0.00 | 0.00 | 0.91 | 0.40 | 25 | T Cell |
| Lcp2 | 0.56 | 0.00 | 0.00 | 0.26 | 0.03 | 25 | T Cell |
| Fermt3 | 0.56 | 0.00 | 0.00 | 0.37 | 0.06 | 25 | T Cell |
| Pola1 | 0.65 | 0.00 | 0.00 | 0.30 | 0.04 | 25 | T Cell |
| Psmb9 | 0.71 | 0.00 | 0.00 | 0.45 | 0.09 | 25 | T Cell |
| Nrm | 0.58 | 0.00 | 0.00 | 0.34 | 0.05 | 25 | T Cell |
| Ltb | 0.47 | 0.00 | 0.00 | 0.27 | 0.03 | 25 | T Cell |
| Limd2 | 0.83 | 0.00 | 0.00 | 0.60 | 0.15 | 25 | T Cell |
| S100a4 | 0.95 | 0.00 | 0.00 | 0.38 | 0.07 | 25 | T Cell |
| Ccdc34 | 0.71 | 0.00 | 0.00 | 0.58 | 0.14 | 25 | T Cell |
| Anp32b | 1.01 | 0.00 | 0.00 | 0.94 | 0.50 | 25 | T Cell |
| Hmgn2 | 0.99 | 0.00 | 0.00 | 0.91 | 0.43 | 25 | T Cell |
| Dek | 1.09 | 0.00 | 0.00 | 0.94 | 0.50 | 25 | T Cell |
| Sh3bgrl3 | 0.92 | 0.00 | 0.00 | 0.87 | 0.34 | 25 | T Cell |
| H2-K1 | 1.02 | 0.00 | 0.00 | 0.86 | 0.32 | 25 | T Cell |
| Gmfg | 0.72 | 0.00 | 0.00 | 0.57 | 0.14 | 25 | T Cell |
| Rangap1 | 0.68 | 0.00 | 0.00 | 0.41 | 0.08 | 25 | T Cell |
| Mcm7 | 0.73 | 0.00 | 0.00 | 0.38 | 0.07 | 25 | T Cell |
| Apbb1ip | 0.58 | 0.00 | 0.00 | 0.46 | 0.09 | 25 | T Cell |
| Ckap5 | 0.75 | 0.00 | 0.00 | 0.50 | 0.11 | 25 | T Cell |
| Tcf7 | 0.91 | 0.00 | 0.00 | 0.35 | 0.06 | 25 | T Cell |
| Hells | 0.65 | 0.00 | 0.00 | 0.34 | 0.06 | 25 | T Cell |
| Sh3kbp1 | 0.78 | 0.00 | 0.00 | 0.48 | 0.10 | 25 | T Cell |
| Ptpn18 | 0.64 | 0.00 | 0.00 | 0.59 | 0.14 | 25 | T Cell |
| Hist1h1c | 1.00 | 0.00 | 0.00 | 0.48 | 0.11 | 25 | T Cell |
| Lrmp | 0.49 | 0.00 | 0.00 | 0.26 | 0.03 | 25 | T Cell |
| Hn1 | 0.81 | 0.00 | 0.00 | 0.67 | 0.20 | 25 | T Cell |
| Ifi47 | 0.62 | 0.00 | 0.00 | 0.32 | 0.05 | 25 | T Cell |
| Haus3 | 0.56 | 0.00 | 0.00 | 0.29 | 0.04 | 25 | T Cell |
| Psat1 | 0.49 | 0.00 | 0.00 | 0.26 | 0.03 | 25 | T Cell |
| Ncaph | 0.55 | 0.00 | 0.00 | 0.29 | 0.05 | 25 | T Cell |
| Me2 | 0.60 | 0.00 | 0.00 | 0.38 | 0.07 | 25 | T Cell |
| Actb | 0.74 | 0.00 | 0.00 | 1.00 | 0.98 | 25 | T Cell |
| Hcls1 | 0.48 | 0.00 | 0.00 | 0.40 | 0.07 | 25 | T Cell |
| Fnbp1 | 0.69 | 0.00 | 0.00 | 0.42 | 0.09 | 25 | T Cell |
| Cit | 0.56 | 0.00 | 0.00 | 0.29 | 0.05 | 25 | T Cell |
| Stk17b | 0.53 | 0.00 | 0.00 | 0.35 | 0.06 | 25 | T Cell |
| Tmpo | 0.87 | 0.00 | 0.00 | 0.57 | 0.15 | 25 | T Cell |
| Arl6ip1 | 1.31 | 0.00 | 0.00 | 0.93 | 0.52 | 25 | T Cell |
| Eri1 | 0.49 | 0.00 | 0.00 | 0.33 | 0.06 | 25 | T Cell |
| Myo1g | 0.44 | 0.00 | 0.00 | 0.26 | 0.04 | 25 | T Cell |
| Fyb | 0.76 | 0.00 | 0.00 | 0.64 | 0.18 | 25 | T Cell |
| Pmf1 | 0.49 | 0.00 | 0.00 | 0.28 | 0.04 | 25 | T Cell |
| Gimap4 | 0.82 | 0.00 | 0.00 | 0.63 | 0.17 | 25 | T Cell |
| Tuba1c | 0.94 | 0.00 | 0.00 | 0.80 | 0.32 | 25 | T Cell |
| Sp110 | 0.50 | 0.00 | 0.00 | 0.38 | 0.07 | 25 | T Cell |
| 2700094K13Rik | 0.76 | 0.00 | 0.00 | 0.58 | 0.16 | 25 | T Cell |
| Emp3 | 0.75 | 0.00 | 0.00 | 0.69 | 0.21 | 25 | T Cell |
| Shisa5 | 0.83 | 0.00 | 0.00 | 0.66 | 0.20 | 25 | T Cell |
| Rad21 | 0.88 | 0.00 | 0.00 | 0.70 | 0.23 | 25 | T Cell |
| Sp100 | 0.70 | 0.00 | 0.00 | 0.40 | 0.08 | 25 | T Cell |
| Tes | 0.64 | 0.00 | 0.00 | 0.43 | 0.10 | 25 | T Cell |
| A430104N18Rik | 0.51 | 0.00 | 0.00 | 0.35 | 0.06 | 25 | T Cell |
| Lrrc58 | 0.76 | 0.00 | 0.00 | 0.99 | 0.91 | 25 | T Cell |
| Fam111a | 0.67 | 0.00 | 0.00 | 0.56 | 0.15 | 25 | T Cell |
| B2m | 0.79 | 0.00 | 0.00 | 0.98 | 0.67 | 25 | T Cell |
| Lsp1 | 0.54 | 0.00 | 0.00 | 0.67 | 0.19 | 25 | T Cell |
| Smchd1 | 0.73 | 0.00 | 0.00 | 0.57 | 0.16 | 25 | T Cell |
| Hmha1 | 0.51 | 0.00 | 0.00 | 0.39 | 0.08 | 25 | T Cell |
| Actr3 | 0.88 | 0.00 | 0.00 | 0.81 | 0.33 | 25 | T Cell |
| Rpsa-ps10 | 0.75 | 0.00 | 0.00 | 0.96 | 0.68 | 25 | T Cell |
| Pim1 | 0.70 | 0.00 | 0.00 | 0.41 | 0.09 | 25 | T Cell |
| Nasp | 0.84 | 0.00 | 0.00 | 0.62 | 0.20 | 25 | T Cell |
| Arhgap30 | 0.53 | 0.00 | 0.00 | 0.44 | 0.10 | 25 | T Cell |
| Exosc8 | 0.54 | 0.00 | 0.00 | 0.38 | 0.08 | 25 | T Cell |
| Hjurp | 0.69 | 0.00 | 0.00 | 0.53 | 0.14 | 25 | T Cell |
| Lgals1 | 0.75 | 0.00 | 0.00 | 0.93 | 0.41 | 25 | T Cell |
| Ncapd3 | 0.61 | 0.00 | 0.00 | 0.30 | 0.06 | 25 | T Cell |
| Gm10123 | 0.68 | 0.00 | 0.00 | 0.96 | 0.81 | 25 | T Cell |
| Fxyd5 | 0.64 | 0.00 | 0.00 | 0.60 | 0.18 | 25 | T Cell |
| Hirip3 | 0.58 | 0.00 | 0.00 | 0.39 | 0.09 | 25 | T Cell |
| Rpa1 | 0.45 | 0.00 | 0.00 | 0.29 | 0.05 | 25 | T Cell |
| Tmsb4x | 0.67 | 0.00 | 0.00 | 0.99 | 0.89 | 25 | T Cell |
| Ppil1 | 0.52 | 0.00 | 0.00 | 0.29 | 0.05 | 25 | T Cell |
| Pfn1 | 0.84 | 0.00 | 0.00 | 0.87 | 0.46 | 25 | T Cell |
| Fkbp3 | 0.74 | 0.00 | 0.00 | 0.78 | 0.30 | 25 | T Cell |
| Srpk1 | 0.64 | 0.00 | 0.00 | 0.47 | 0.12 | 25 | T Cell |
| Itgal | 0.31 | 0.00 | 0.00 | 0.25 | 0.04 | 25 | T Cell |
| Cmc2 | 0.55 | 0.00 | 0.00 | 0.28 | 0.05 | 25 | T Cell |
| Atad5 | 0.57 | 0.00 | 0.00 | 0.29 | 0.05 | 25 | T Cell |
| Ska2 | 0.55 | 0.00 | 0.00 | 0.32 | 0.07 | 25 | T Cell |
| Gm5506 | 0.60 | 0.00 | 0.00 | 0.51 | 0.14 | 25 | T Cell |
| Tipin | 0.55 | 0.00 | 0.00 | 0.41 | 0.10 | 25 | T Cell |
| Ncbp1 | 0.47 | 0.00 | 0.00 | 0.29 | 0.06 | 25 | T Cell |
| Dnajc15 | 0.55 | 0.00 | 0.00 | 0.36 | 0.08 | 25 | T Cell |
| Pyhin1 | 0.66 | 0.00 | 0.00 | 0.27 | 0.05 | 25 | T Cell |
| Crip1 | 0.80 | 0.00 | 0.00 | 0.92 | 0.47 | 25 | T Cell |
| Tubb4b | 0.61 | 0.00 | 0.00 | 0.59 | 0.19 | 25 | T Cell |
| Mcm4 | 0.54 | 0.00 | 0.00 | 0.29 | 0.06 | 25 | T Cell |
| Usp1 | 0.68 | 0.00 | 0.00 | 0.53 | 0.15 | 25 | T Cell |
| 2700029M09Rik | 0.71 | 0.00 | 0.00 | 0.46 | 0.13 | 25 | T Cell |
| Sfxn1 | 0.44 | 0.00 | 0.00 | 0.33 | 0.07 | 25 | T Cell |
| Ccr2 | 0.64 | 0.00 | 0.00 | 0.34 | 0.07 | 25 | T Cell |
| Esyt1 | 0.47 | 0.00 | 0.00 | 0.32 | 0.07 | 25 | T Cell |
| Ldha | 0.77 | 0.00 | 0.00 | 0.72 | 0.29 | 25 | T Cell |
| Slfn2 | 0.52 | 0.00 | 0.00 | 0.49 | 0.13 | 25 | T Cell |
| Cdca4 | 0.48 | 0.00 | 0.00 | 0.27 | 0.05 | 25 | T Cell |
| Pgk1 | 0.59 | 0.00 | 0.00 | 0.43 | 0.11 | 25 | T Cell |
| Rnaseh2b | 0.50 | 0.00 | 0.00 | 0.36 | 0.08 | 25 | T Cell |
| Snrpd1 | 0.65 | 0.00 | 0.00 | 0.61 | 0.20 | 25 | T Cell |
| Eno1 | 0.66 | 0.00 | 0.00 | 0.53 | 0.16 | 25 | T Cell |
| Sec11c | 0.67 | 0.00 | 0.00 | 0.56 | 0.18 | 25 | T Cell |
| S100a10 | 1.07 | 0.00 | 0.00 | 0.80 | 0.38 | 25 | T Cell |
| Acot7 | 0.40 | 0.00 | 0.00 | 0.31 | 0.06 | 25 | T Cell |
| Gimap1 | 0.50 | 0.00 | 0.00 | 0.50 | 0.14 | 25 | T Cell |
| Dgka | 0.61 | 0.00 | 0.00 | 0.27 | 0.05 | 25 | T Cell |
| Emb | 0.67 | 0.00 | 0.00 | 0.42 | 0.11 | 25 | T Cell |
| Cdk2ap2 | 0.57 | 0.00 | 0.00 | 0.52 | 0.15 | 25 | T Cell |
| Banf1 | 0.74 | 0.00 | 0.00 | 0.71 | 0.28 | 25 | T Cell |
| Ppia | 0.67 | 0.00 | 0.00 | 0.97 | 0.80 | 25 | T Cell |
| Uchl5 | 0.59 | 0.00 | 0.00 | 0.43 | 0.11 | 25 | T Cell |
| Nudt21 | 0.53 | 0.00 | 0.00 | 0.45 | 0.12 | 25 | T Cell |
| Mbnl1 | 0.71 | 0.00 | 0.00 | 0.68 | 0.27 | 25 | T Cell |
| Topbp1 | 0.46 | 0.00 | 0.00 | 0.27 | 0.05 | 25 | T Cell |
| Rpsa | 0.67 | 0.00 | 0.00 | 0.94 | 0.62 | 25 | T Cell |
| Vars | 0.46 | 0.00 | 0.00 | 0.34 | 0.08 | 25 | T Cell |
| Rbm3 | 0.75 | 0.00 | 0.00 | 0.83 | 0.41 | 25 | T Cell |
| Rplp0 | 0.56 | 0.00 | 0.00 | 0.99 | 0.83 | 25 | T Cell |
| H2-Q7 | 0.69 | 0.00 | 0.00 | 0.32 | 0.07 | 25 | T Cell |
| H2-D1 | 0.75 | 0.00 | 0.00 | 0.91 | 0.49 | 25 | T Cell |
| Leprotl1 | 0.60 | 0.00 | 0.00 | 0.49 | 0.15 | 25 | T Cell |
| Cenpc1 | 0.55 | 0.00 | 0.00 | 0.36 | 0.09 | 25 | T Cell |
| G2e3 | 0.46 | 0.00 | 0.00 | 0.30 | 0.06 | 25 | T Cell |
| Ets1 | 0.59 | 0.00 | 0.00 | 0.69 | 0.25 | 25 | T Cell |
| Dtymk | 0.57 | 0.00 | 0.00 | 0.46 | 0.13 | 25 | T Cell |
| Hat1 | 0.49 | 0.00 | 0.00 | 0.32 | 0.07 | 25 | T Cell |
| Gm8730 | 0.55 | 0.00 | 0.00 | 0.98 | 0.82 | 25 | T Cell |
| Rps26 | 0.51 | 0.00 | 0.00 | 0.99 | 0.90 | 25 | T Cell |
| Gm17087 | 0.68 | 0.00 | 0.00 | 0.63 | 0.24 | 25 | T Cell |
| Tpm3-rs7 | 0.62 | 0.00 | 0.00 | 0.97 | 0.69 | 25 | T Cell |
| Cep57 | 0.51 | 0.00 | 0.00 | 0.40 | 0.11 | 25 | T Cell |
| Ssrp1 | 0.63 | 0.00 | 0.00 | 0.59 | 0.21 | 25 | T Cell |
| Snrpe | 0.66 | 0.00 | 0.00 | 0.78 | 0.36 | 25 | T Cell |
| Lsm3 | 0.44 | 0.00 | 0.00 | 0.40 | 0.11 | 25 | T Cell |
| Nfyb | 0.54 | 0.00 | 0.00 | 0.42 | 0.12 | 25 | T Cell |
| Pabpc1 | 0.57 | 0.00 | 0.00 | 0.99 | 0.81 | 25 | T Cell |
| Pgk1-rs7 | 0.53 | 0.00 | 0.00 | 0.38 | 0.10 | 25 | T Cell |
| Ifi203 | 0.57 | 0.00 | 0.00 | 0.48 | 0.14 | 25 | T Cell |
| Ppp1r18 | 0.44 | 0.00 | 0.00 | 0.29 | 0.06 | 25 | T Cell |
| Gm6625 | 0.64 | 0.00 | 0.00 | 0.65 | 0.25 | 25 | T Cell |
| Bub3 | 0.54 | 0.00 | 0.00 | 0.46 | 0.14 | 25 | T Cell |
| Ctcf | 0.60 | 0.00 | 0.00 | 0.63 | 0.23 | 25 | T Cell |
| Cep110 | 0.59 | 0.00 | 0.00 | 0.54 | 0.19 | 25 | T Cell |
| Hprt | 0.62 | 0.00 | 0.00 | 0.44 | 0.14 | 25 | T Cell |
| Hnrnpf | 0.63 | 0.00 | 0.00 | 0.77 | 0.34 | 25 | T Cell |
| Rpa3 | 0.53 | 0.00 | 0.00 | 0.43 | 0.13 | 25 | T Cell |
| Gm10068 | 0.64 | 0.00 | 0.00 | 0.71 | 0.30 | 25 | T Cell |
| H2afv | 0.81 | 0.00 | 0.00 | 0.67 | 0.28 | 25 | T Cell |
| Mif | 0.63 | 0.00 | 0.00 | 0.54 | 0.19 | 25 | T Cell |
| Fam49b | 0.50 | 0.00 | 0.00 | 0.39 | 0.11 | 25 | T Cell |
| Arrb2 | 0.37 | 0.00 | 0.00 | 0.29 | 0.07 | 25 | T Cell |
| Gm8991 | 0.72 | 0.00 | 0.00 | 0.59 | 0.23 | 25 | T Cell |
| Itga4 | 0.62 | 0.00 | 0.00 | 0.36 | 0.10 | 25 | T Cell |
| Gm5641 | 0.76 | 0.00 | 0.00 | 0.75 | 0.36 | 25 | T Cell |
| Cenpq | 0.49 | 0.00 | 0.00 | 0.29 | 0.07 | 25 | T Cell |
| Gm6793 | 0.65 | 0.00 | 0.00 | 0.69 | 0.30 | 25 | T Cell |
| Pbdc1 | 0.52 | 0.00 | 0.00 | 0.43 | 0.13 | 25 | T Cell |
| Mrpl18 | 0.64 | 0.00 | 0.00 | 0.56 | 0.21 | 25 | T Cell |
| Tpm3 | 0.58 | 0.00 | 0.00 | 0.97 | 0.70 | 25 | T Cell |
| Elf1 | 0.59 | 0.00 | 0.00 | 0.57 | 0.20 | 25 | T Cell |
| Hnrnpa2b1 | 0.59 | 0.00 | 0.00 | 0.96 | 0.72 | 25 | T Cell |
| Samsn1 | 0.50 | 0.00 | 0.00 | 0.29 | 0.07 | 25 | T Cell |
| Wipf1 | 0.47 | 0.00 | 0.00 | 0.36 | 0.10 | 25 | T Cell |
| Rps11 | 0.53 | 0.00 | 0.00 | 0.96 | 0.76 | 25 | T Cell |
| Smc1a | 0.62 | 0.00 | 0.00 | 0.76 | 0.35 | 25 | T Cell |
| Serbp1 | 0.60 | 0.00 | 0.00 | 0.96 | 0.75 | 25 | T Cell |
| Lsm5 | 0.50 | 0.00 | 0.00 | 0.40 | 0.12 | 25 | T Cell |
| Cbx3 | 0.67 | 0.00 | 0.00 | 0.72 | 0.34 | 25 | T Cell |
| Calm3 | 0.60 | 0.00 | 0.00 | 0.54 | 0.19 | 25 | T Cell |
| Pkm | 0.62 | 0.00 | 0.00 | 0.57 | 0.22 | 25 | T Cell |
| Lsm4 | 0.56 | 0.00 | 0.00 | 0.56 | 0.20 | 25 | T Cell |
| Rps26-ps1 | 0.50 | 0.00 | 0.00 | 0.99 | 0.87 | 25 | T Cell |
| Anp32a | 0.57 | 0.00 | 0.00 | 0.95 | 0.60 | 25 | T Cell |
| Srsf7 | 0.54 | 0.00 | 0.00 | 0.47 | 0.16 | 25 | T Cell |
| Rplp1 | 0.46 | 0.00 | 0.00 | 0.99 | 0.90 | 25 | T Cell |
| Mndal | 0.47 | 0.00 | 0.00 | 0.50 | 0.16 | 25 | T Cell |
| Gapdh | 0.49 | 0.00 | 0.00 | 0.44 | 0.14 | 25 | T Cell |
| Rbbp4 | 0.51 | 0.00 | 0.00 | 0.56 | 0.20 | 25 | T Cell |
| Ifi27l2a | 0.82 | 0.00 | 0.00 | 0.34 | 0.09 | 25 | T Cell |
| Uchl3 | 0.37 | 0.00 | 0.00 | 0.28 | 0.07 | 25 | T Cell |
| Rad50 | 0.50 | 0.00 | 0.00 | 0.54 | 0.19 | 25 | T Cell |
| Psip1 | 0.53 | 0.00 | 0.00 | 0.46 | 0.15 | 25 | T Cell |
| Gm9242 | 0.68 | 0.00 | 0.00 | 0.58 | 0.24 | 25 | T Cell |
| Hmga1 | 0.40 | 0.00 | 0.00 | 0.30 | 0.08 | 25 | T Cell |
| Nubp1 | 0.43 | 0.00 | 0.00 | 0.26 | 0.06 | 25 | T Cell |
| Zc3hav1 | 0.41 | 0.00 | 0.00 | 0.30 | 0.08 | 25 | T Cell |
| Nop58 | 0.65 | 0.00 | 0.00 | 0.62 | 0.26 | 25 | T Cell |
| Cbx5 | 0.59 | 0.00 | 0.00 | 0.64 | 0.26 | 25 | T Cell |
| Whsc1 | 0.53 | 0.00 | 0.00 | 0.42 | 0.14 | 25 | T Cell |
| Capza1 | 0.44 | 0.00 | 0.00 | 0.40 | 0.12 | 25 | T Cell |
| 11-Sep | 0.55 | 0.00 | 0.00 | 0.58 | 0.22 | 25 | T Cell |
| Pa2g4 | 0.69 | 0.00 | 0.00 | 0.61 | 0.26 | 25 | T Cell |
| Gm12355 | 0.54 | 0.00 | 0.00 | 0.63 | 0.26 | 25 | T Cell |
| Hmgb1 | 0.63 | 0.00 | 0.00 | 0.89 | 0.61 | 25 | T Cell |
| Snrpf | 0.58 | 0.00 | 0.00 | 0.65 | 0.28 | 25 | T Cell |
| Tmem71 | 0.42 | 0.00 | 0.00 | 0.26 | 0.06 | 25 | T Cell |
| Mdh1 | 0.51 | 0.00 | 0.00 | 0.51 | 0.19 | 25 | T Cell |
| Gm12728 | 0.47 | 0.00 | 0.00 | 0.48 | 0.17 | 25 | T Cell |
| Suz12 | 0.43 | 0.00 | 0.00 | 0.41 | 0.13 | 25 | T Cell |
| Atp1b3 | 0.61 | 0.00 | 0.00 | 0.61 | 0.25 | 25 | T Cell |
| Jak1 | 0.53 | 0.00 | 0.00 | 0.63 | 0.25 | 25 | T Cell |
| Dnajc2 | 0.50 | 0.00 | 0.00 | 0.53 | 0.20 | 25 | T Cell |
| Gm10275 | 0.49 | 0.00 | 0.00 | 0.93 | 0.66 | 25 | T Cell |
| Rpl4 | 0.45 | 0.00 | 0.00 | 0.97 | 0.83 | 25 | T Cell |
| Bzw2 | 0.53 | 0.00 | 0.00 | 0.43 | 0.15 | 25 | T Cell |
| Clic1 | 0.58 | 0.00 | 0.00 | 0.76 | 0.38 | 25 | T Cell |
| Supt16 | 0.59 | 0.00 | 0.00 | 0.59 | 0.25 | 25 | T Cell |
| Gm8186 | 0.59 | 0.00 | 0.00 | 0.75 | 0.39 | 25 | T Cell |
| Rpl14 | 0.44 | 0.00 | 0.00 | 0.98 | 0.84 | 25 | T Cell |
| Crlf3 | 0.45 | 0.00 | 0.00 | 0.41 | 0.14 | 25 | T Cell |
| Ubash3b | 0.37 | 0.00 | 0.00 | 0.27 | 0.07 | 25 | T Cell |
| Hmgn5 | 0.57 | 0.00 | 0.00 | 0.62 | 0.26 | 25 | T Cell |
| Srgn | 0.42 | 0.00 | 0.00 | 0.77 | 0.33 | 25 | T Cell |
| Srsf3 | 0.57 | 0.00 | 0.00 | 0.85 | 0.44 | 25 | T Cell |
| Rbbp7 | 0.58 | 0.00 | 0.00 | 0.60 | 0.26 | 25 | T Cell |
| Hspa14 | 0.38 | 0.00 | 0.00 | 0.35 | 0.10 | 25 | T Cell |
| Ranbp1 | 0.61 | 0.00 | 0.00 | 0.64 | 0.29 | 25 | T Cell |
| Rfc2 | 0.35 | 0.00 | 0.00 | 0.30 | 0.08 | 25 | T Cell |
| Rps3 | 0.44 | 0.00 | 0.00 | 0.99 | 0.83 | 25 | T Cell |
| Cdc42se2 | 0.45 | 0.00 | 0.00 | 0.41 | 0.14 | 25 | T Cell |
| Npm1 | 0.60 | 0.00 | 0.00 | 0.88 | 0.58 | 25 | T Cell |
| Gimap6 | 0.47 | 0.00 | 0.00 | 0.46 | 0.16 | 25 | T Cell |
| Rpl32 | 0.43 | 0.00 | 0.00 | 0.98 | 0.84 | 25 | T Cell |
| Wdr89 | 0.39 | 0.00 | 0.00 | 0.99 | 0.89 | 25 | T Cell |
| Tap2 | 0.35 | 0.00 | 0.00 | 0.27 | 0.07 | 25 | T Cell |
| Stk4 | 0.40 | 0.00 | 0.00 | 0.42 | 0.14 | 25 | T Cell |
| Psme2b | 0.45 | 0.00 | 0.00 | 0.67 | 0.30 | 25 | T Cell |
| Hist1h2al | 0.44 | 0.00 | 0.00 | 0.33 | 0.10 | 25 | T Cell |
| Stip1 | 0.47 | 0.00 | 0.00 | 0.38 | 0.13 | 25 | T Cell |
| G3bp1 | 0.41 | 0.00 | 0.00 | 0.48 | 0.17 | 25 | T Cell |
| Snrpg | 0.55 | 0.00 | 0.00 | 0.74 | 0.39 | 25 | T Cell |
| Ncaph2 | 0.41 | 0.00 | 0.00 | 0.30 | 0.09 | 25 | T Cell |
| Rpl13a | 0.37 | 0.00 | 0.00 | 0.99 | 0.87 | 25 | T Cell |
| Isy1 | 0.35 | 0.00 | 0.00 | 0.28 | 0.08 | 25 | T Cell |
| Srrt | 0.42 | 0.00 | 0.00 | 0.51 | 0.20 | 25 | T Cell |
| Nhp2 | 0.43 | 0.00 | 0.00 | 0.36 | 0.12 | 25 | T Cell |
| Hars | 0.32 | 0.00 | 0.00 | 0.32 | 0.10 | 25 | T Cell |
| Exosc3 | 0.35 | 0.00 | 0.00 | 0.26 | 0.07 | 25 | T Cell |
| Nup50 | 0.30 | 0.00 | 0.00 | 0.26 | 0.07 | 25 | T Cell |
| Emg1 | 0.39 | 0.00 | 0.00 | 0.43 | 0.16 | 25 | T Cell |
| Gltp | 0.36 | 0.00 | 0.00 | 0.33 | 0.10 | 25 | T Cell |
| Dkc1 | 0.35 | 0.00 | 0.00 | 0.36 | 0.12 | 25 | T Cell |
| Cap1 | 0.44 | 0.00 | 0.00 | 0.46 | 0.17 | 25 | T Cell |
| Ezr | 0.36 | 0.00 | 0.00 | 0.55 | 0.22 | 25 | T Cell |
| Ccm2 | 0.38 | 0.00 | 0.00 | 0.35 | 0.12 | 25 | T Cell |
| Snrpd3 | 0.49 | 0.00 | 0.00 | 0.67 | 0.32 | 25 | T Cell |
| Abracl | 0.41 | 0.00 | 0.00 | 0.56 | 0.23 | 25 | T Cell |
| Ncl | 0.49 | 0.00 | 0.00 | 0.95 | 0.78 | 25 | T Cell |
| Rps3a1 | 0.38 | 0.00 | 0.00 | 0.98 | 0.85 | 25 | T Cell |
| Rpl12 | 0.46 | 0.00 | 0.00 | 0.87 | 0.55 | 25 | T Cell |
| Rbbp8 | 0.37 | 0.00 | 0.00 | 0.29 | 0.09 | 25 | T Cell |
| Vrk1 | 0.38 | 0.00 | 0.00 | 0.27 | 0.08 | 25 | T Cell |
| Ola1 | 0.37 | 0.00 | 0.00 | 0.41 | 0.15 | 25 | T Cell |
| Gtf2f1 | 0.45 | 0.00 | 0.00 | 0.52 | 0.21 | 25 | T Cell |
| Gm10073 | 0.41 | 0.00 | 0.00 | 0.96 | 0.86 | 25 | T Cell |
| Tmsb10 | 0.45 | 0.00 | 0.00 | 0.98 | 0.86 | 25 | T Cell |
| Rps5 | 0.41 | 0.00 | 0.00 | 0.96 | 0.78 | 25 | T Cell |
| Hnrnpa3 | 0.53 | 0.00 | 0.00 | 0.80 | 0.46 | 25 | T Cell |
| Impdh2 | 0.36 | 0.00 | 0.00 | 0.31 | 0.10 | 25 | T Cell |
| Med30 | 0.34 | 0.00 | 0.00 | 0.29 | 0.09 | 25 | T Cell |
| Lsm6 | 0.40 | 0.00 | 0.00 | 0.38 | 0.13 | 25 | T Cell |
| Psmd14 | 0.43 | 0.00 | 0.00 | 0.46 | 0.18 | 25 | T Cell |
| Slc25a5 | 0.51 | 0.00 | 0.00 | 0.84 | 0.59 | 25 | T Cell |
| Hnrnpu | 0.49 | 0.00 | 0.00 | 0.91 | 0.64 | 25 | T Cell |
| Nop56 | 0.48 | 0.00 | 0.00 | 0.49 | 0.20 | 25 | T Cell |
| Sae1 | 0.34 | 0.00 | 0.00 | 0.34 | 0.11 | 25 | T Cell |
| Pole4 | 0.37 | 0.00 | 0.00 | 0.30 | 0.09 | 25 | T Cell |
| 5830418K08Rik | 0.41 | 0.00 | 0.00 | 0.29 | 0.09 | 25 | T Cell |
| Rfc3 | 0.39 | 0.00 | 0.00 | 0.29 | 0.09 | 25 | T Cell |
| Setd8 | 0.32 | 0.00 | 0.00 | 0.35 | 0.12 | 25 | T Cell |
| Tpi1 | 0.43 | 0.00 | 0.00 | 0.40 | 0.15 | 25 | T Cell |
| Atp11b | 0.38 | 0.00 | 0.00 | 0.26 | 0.07 | 25 | T Cell |
| Nme1 | 0.47 | 0.00 | 0.00 | 0.47 | 0.19 | 25 | T Cell |
| Hint1 | 0.47 | 0.00 | 0.00 | 0.81 | 0.46 | 25 | T Cell |
| Rapgef6 | 0.42 | 0.00 | 0.00 | 0.38 | 0.14 | 25 | T Cell |
| Naa10 | 0.41 | 0.00 | 0.00 | 0.29 | 0.09 | 25 | T Cell |
| Eef1b2 | 0.45 | 0.00 | 0.00 | 0.91 | 0.65 | 25 | T Cell |
| Eif2s1 | 0.39 | 0.00 | 0.00 | 0.37 | 0.13 | 25 | T Cell |
| Rcsd1 | 0.25 | 0.00 | 0.00 | 0.28 | 0.08 | 25 | T Cell |
| Nans | 0.32 | 0.00 | 0.00 | 0.29 | 0.09 | 25 | T Cell |
| Rgs10 | 0.33 | 0.00 | 0.00 | 0.31 | 0.10 | 25 | T Cell |
| Rps20 | 0.42 | 0.00 | 0.00 | 0.95 | 0.79 | 25 | T Cell |
| Cdc42se1 | 0.42 | 0.00 | 0.00 | 0.48 | 0.19 | 25 | T Cell |
| H2-T22 | 0.34 | 0.00 | 0.00 | 0.28 | 0.08 | 25 | T Cell |
| Smc6 | 0.42 | 0.00 | 0.00 | 0.67 | 0.33 | 25 | T Cell |
| Psme2 | 0.36 | 0.00 | 0.00 | 0.55 | 0.24 | 25 | T Cell |
| Nmi | 0.28 | 0.00 | 0.00 | 0.34 | 0.11 | 25 | T Cell |
| Nudc | 0.39 | 0.00 | 0.00 | 0.50 | 0.21 | 25 | T Cell |
| Kars | 0.32 | 0.00 | 0.00 | 0.26 | 0.07 | 25 | T Cell |
| Tfam | 0.28 | 0.00 | 0.00 | 0.27 | 0.08 | 25 | T Cell |
| Gclm | 0.51 | 0.00 | 0.00 | 0.74 | 0.42 | 25 | T Cell |
| Syncrip | 0.44 | 0.00 | 0.00 | 0.55 | 0.25 | 25 | T Cell |
| Ppp1ca | 0.48 | 0.00 | 0.00 | 0.66 | 0.33 | 25 | T Cell |
| Arhgef1 | 0.34 | 0.00 | 0.00 | 0.34 | 0.12 | 25 | T Cell |
| Naa38 | 0.35 | 0.00 | 0.00 | 0.40 | 0.15 | 25 | T Cell |
| Gm5160 | 0.43 | 0.00 | 0.00 | 0.50 | 0.22 | 25 | T Cell |
| Gnb2l1 | 0.43 | 0.00 | 0.00 | 0.92 | 0.65 | 25 | T Cell |
| Tnfaip8 | 0.39 | 0.00 | 0.00 | 0.32 | 0.11 | 25 | T Cell |
| Gm4204 | 0.40 | 0.00 | 0.00 | 0.83 | 0.50 | 25 | T Cell |
| Gm9396 | 0.43 | 0.00 | 0.00 | 0.85 | 0.54 | 25 | T Cell |
| Kpnb1 | 0.40 | 0.00 | 0.00 | 0.46 | 0.19 | 25 | T Cell |
| Prelid1 | 0.45 | 0.00 | 0.00 | 0.60 | 0.29 | 25 | T Cell |
| Arpc2 | 0.41 | 0.00 | 0.00 | 0.92 | 0.61 | 25 | T Cell |
| Snrpa1 | 0.29 | 0.00 | 0.00 | 0.38 | 0.14 | 25 | T Cell |
| Sf3b5 | 0.40 | 0.00 | 0.00 | 0.51 | 0.22 | 25 | T Cell |
| Sf3a3 | 0.33 | 0.00 | 0.00 | 0.34 | 0.12 | 25 | T Cell |
| Hmga1-rs1 | 0.28 | 0.00 | 0.00 | 0.25 | 0.08 | 25 | T Cell |
| Larp7 | 0.33 | 0.00 | 0.00 | 0.43 | 0.17 | 25 | T Cell |
| Rps15a | 0.34 | 0.00 | 0.00 | 0.96 | 0.81 | 25 | T Cell |
| Hnrnpd | 0.43 | 0.00 | 0.00 | 0.60 | 0.28 | 25 | T Cell |
| Rps16 | 0.34 | 0.00 | 0.00 | 0.97 | 0.84 | 25 | T Cell |
| Gm4070 | 0.37 | 0.00 | 0.00 | 0.34 | 0.12 | 25 | T Cell |
| Cfl1 | 0.48 | 0.00 | 0.00 | 0.86 | 0.58 | 25 | T Cell |
| Lrrfip1 | 0.38 | 0.00 | 0.00 | 0.56 | 0.25 | 25 | T Cell |
| Bzw1 | 0.44 | 0.00 | 0.00 | 0.65 | 0.32 | 25 | T Cell |
| Hnrnpc | 0.44 | 0.00 | 0.00 | 0.78 | 0.44 | 25 | T Cell |
| Mrpl40 | 0.29 | 0.00 | 0.00 | 0.32 | 0.11 | 25 | T Cell |
| Tubb4b-ps1 | 0.29 | 0.00 | 0.00 | 0.27 | 0.09 | 25 | T Cell |
| Odf2 | 0.34 | 0.00 | 0.00 | 0.27 | 0.09 | 25 | T Cell |
| Eif4a3 | 0.40 | 0.00 | 0.00 | 0.45 | 0.19 | 25 | T Cell |
| Tcof1 | 0.31 | 0.00 | 0.00 | 0.32 | 0.11 | 25 | T Cell |
| Ywhah | 0.47 | 0.00 | 0.00 | 0.55 | 0.26 | 25 | T Cell |
| Tsc22d4 | 0.37 | 0.00 | 0.00 | 0.59 | 0.28 | 25 | T Cell |
| Slmo2 | 0.32 | 0.00 | 0.00 | 0.28 | 0.09 | 25 | T Cell |
| Nt5c | 0.27 | 0.00 | 0.00 | 0.41 | 0.16 | 25 | T Cell |
| Hsp90aa1 | 0.41 | 0.00 | 0.00 | 0.79 | 0.47 | 25 | T Cell |
| Gvin1 | 0.38 | 0.00 | 0.00 | 0.33 | 0.12 | 25 | T Cell |
| Mettl23 | 0.28 | 0.00 | 0.00 | 0.32 | 0.11 | 25 | T Cell |
| Rps6 | 0.34 | 0.00 | 0.00 | 0.98 | 0.82 | 25 | T Cell |
| Nap1l1 | 0.33 | 0.00 | 0.00 | 0.80 | 0.47 | 25 | T Cell |
| Snrnp40 | 0.30 | 0.00 | 0.00 | 0.26 | 0.08 | 25 | T Cell |
| Hnrnpab | 0.52 | 0.00 | 0.00 | 0.63 | 0.33 | 25 | T Cell |
| Gm10093 | 0.39 | 0.00 | 0.00 | 0.46 | 0.19 | 25 | T Cell |
| Stat1 | 0.39 | 0.00 | 0.00 | 0.30 | 0.10 | 25 | T Cell |
| Ide | 0.33 | 0.00 | 0.00 | 0.27 | 0.09 | 25 | T Cell |
| Cox20 | 0.27 | 0.00 | 0.00 | 0.45 | 0.18 | 25 | T Cell |
| Celf2 | 0.32 | 0.00 | 0.00 | 0.60 | 0.28 | 25 | T Cell |
| Set | 0.48 | 0.00 | 0.00 | 0.65 | 0.34 | 25 | T Cell |
| Casp3 | 0.38 | 0.00 | 0.00 | 0.25 | 0.08 | 25 | T Cell |
| Ppp2r5c | 0.37 | 0.00 | 0.00 | 0.48 | 0.21 | 25 | T Cell |
| Cox5a | 0.37 | 0.00 | 0.00 | 0.63 | 0.31 | 25 | T Cell |
| Stra13 | 0.30 | 0.00 | 0.00 | 0.26 | 0.08 | 25 | T Cell |
| Ptbp3 | 0.34 | 0.00 | 0.00 | 0.72 | 0.36 | 25 | T Cell |
| Tmed5 | 0.33 | 0.00 | 0.00 | 0.42 | 0.17 | 25 | T Cell |
| Sptssa | 0.32 | 0.00 | 0.00 | 0.47 | 0.20 | 25 | T Cell |
| Parp2 | 0.33 | 0.00 | 0.00 | 0.30 | 0.10 | 25 | T Cell |
| Rps3a2 | 0.34 | 0.00 | 0.00 | 0.95 | 0.82 | 25 | T Cell |
| Tcp1 | 0.32 | 0.00 | 0.00 | 0.53 | 0.24 | 25 | T Cell |
| Rplp2 | 0.33 | 0.00 | 0.00 | 0.97 | 0.81 | 25 | T Cell |
| Smarca5 | 0.38 | 0.00 | 0.00 | 0.62 | 0.30 | 25 | T Cell |
| Rnaseh2c | 0.32 | 0.00 | 0.00 | 0.30 | 0.11 | 25 | T Cell |
| Psmb2 | 0.30 | 0.00 | 0.00 | 0.56 | 0.25 | 25 | T Cell |
| Ing1 | 0.32 | 0.00 | 0.00 | 0.39 | 0.15 | 25 | T Cell |
| Rps2 | 0.44 | 0.00 | 0.00 | 0.88 | 0.67 | 25 | T Cell |
| Mrpl42 | 0.38 | 0.00 | 0.00 | 0.35 | 0.13 | 25 | T Cell |
| Ak2 | 0.30 | 0.00 | 0.00 | 0.39 | 0.15 | 25 | T Cell |
| Arpc5l | 0.32 | 0.00 | 0.00 | 0.41 | 0.17 | 25 | T Cell |
| Dhx9 | 0.37 | 0.00 | 0.00 | 0.44 | 0.19 | 25 | T Cell |
| Ahsa1 | 0.28 | 0.00 | 0.00 | 0.34 | 0.13 | 25 | T Cell |
| Naa50 | 0.37 | 0.00 | 0.00 | 0.34 | 0.13 | 25 | T Cell |
| Cd47 | 0.37 | 0.00 | 0.00 | 0.77 | 0.43 | 25 | T Cell |
| Npm3 | 0.26 | 0.00 | 0.00 | 0.28 | 0.10 | 25 | T Cell |
| Eef1d | 0.33 | 0.00 | 0.00 | 0.54 | 0.26 | 25 | T Cell |
| Nsmce2 | 0.30 | 0.00 | 0.00 | 0.33 | 0.12 | 25 | T Cell |
| Xrn2 | 0.39 | 0.00 | 0.00 | 0.60 | 0.30 | 25 | T Cell |
| Arl4c | 0.38 | 0.00 | 0.00 | 0.29 | 0.10 | 25 | T Cell |
| Smarcc1 | 0.30 | 0.00 | 0.00 | 0.40 | 0.17 | 25 | T Cell |
| Eif5a | 0.41 | 0.00 | 0.00 | 0.64 | 0.34 | 25 | T Cell |
| Rpl5 | 0.34 | 0.00 | 0.00 | 0.94 | 0.74 | 25 | T Cell |
| Pdcd4 | 0.40 | 0.00 | 0.00 | 0.49 | 0.23 | 25 | T Cell |
| Aebp2 | 0.29 | 0.00 | 0.00 | 0.30 | 0.11 | 25 | T Cell |
| Hdac1 | 0.36 | 0.00 | 0.00 | 0.44 | 0.19 | 25 | T Cell |
| Wdr1 | 0.31 | 0.00 | 0.00 | 0.40 | 0.17 | 25 | T Cell |
| Thrap3 | 0.37 | 0.00 | 0.00 | 0.69 | 0.37 | 25 | T Cell |
| Tma7 | 0.35 | 0.00 | 0.00 | 0.62 | 0.32 | 25 | T Cell |
| Rpl3 | 0.37 | 0.00 | 0.00 | 0.86 | 0.63 | 25 | T Cell |
| Cdc5l | 0.32 | 0.00 | 0.00 | 0.49 | 0.22 | 25 | T Cell |
| Actr2 | 0.33 | 0.00 | 0.00 | 0.57 | 0.27 | 25 | T Cell |
| Erh | 0.33 | 0.00 | 0.00 | 0.58 | 0.28 | 25 | T Cell |
| Utp3 | 0.39 | 0.00 | 0.00 | 0.48 | 0.22 | 25 | T Cell |
| Hspa8 | 0.42 | 0.00 | 0.00 | 0.89 | 0.67 | 25 | T Cell |
| Nap1l4 | 0.33 | 0.00 | 0.00 | 0.55 | 0.26 | 25 | T Cell |
| Bola2 | 0.31 | 0.00 | 0.00 | 0.39 | 0.16 | 25 | T Cell |
| Sf3b2 | 0.36 | 0.00 | 0.00 | 0.72 | 0.39 | 25 | T Cell |
| Smc3 | 0.35 | 0.00 | 0.00 | 0.68 | 0.38 | 25 | T Cell |
| AC121131.2 | 0.36 | 0.00 | 0.00 | 0.72 | 0.40 | 25 | T Cell |
| Gspt1 | 0.31 | 0.00 | 0.00 | 0.37 | 0.15 | 25 | T Cell |
| Cnot6l | 0.27 | 0.00 | 0.00 | 0.34 | 0.13 | 25 | T Cell |
| Uba2 | 0.27 | 0.00 | 0.00 | 0.30 | 0.11 | 25 | T Cell |
| Ppm1g | 0.34 | 0.00 | 0.00 | 0.34 | 0.13 | 25 | T Cell |
| Rab8a | 0.36 | 0.00 | 0.00 | 0.34 | 0.14 | 25 | T Cell |
| Gm6104 | 0.28 | 0.00 | 0.00 | 0.32 | 0.13 | 25 | T Cell |
| Orc6 | 0.33 | 0.00 | 0.00 | 0.28 | 0.10 | 25 | T Cell |
| Khdrbs1 | 0.31 | 0.00 | 0.00 | 0.62 | 0.31 | 25 | T Cell |
| Nsmce1 | 0.37 | 0.00 | 0.00 | 0.34 | 0.14 | 25 | T Cell |
| Rnps1 | 0.27 | 0.00 | 0.00 | 0.37 | 0.15 | 25 | T Cell |
| Ptpn2 | 0.28 | 0.00 | 0.00 | 0.33 | 0.13 | 25 | T Cell |
| AC167036.1 | 0.30 | 0.00 | 0.00 | 0.61 | 0.30 | 25 | T Cell |
| Agpat5 | 0.27 | 0.00 | 0.00 | 0.27 | 0.10 | 25 | T Cell |
| Arpc1b | 0.31 | 0.00 | 0.00 | 0.68 | 0.37 | 25 | T Cell |
| Mdh2 | 0.30 | 0.00 | 0.00 | 0.48 | 0.22 | 25 | T Cell |
| Mtf2 | 0.33 | 0.00 | 0.00 | 0.40 | 0.18 | 25 | T Cell |
| Rps18 | 0.34 | 0.00 | 0.00 | 0.89 | 0.65 | 25 | T Cell |
| Rps10-ps1 | 0.31 | 0.00 | 0.00 | 0.94 | 0.77 | 25 | T Cell |
| Nop10 | 0.31 | 0.00 | 0.00 | 0.56 | 0.28 | 25 | T Cell |
| AC123611.1 | 0.34 | 0.00 | 0.00 | 0.70 | 0.39 | 25 | T Cell |
| Pak2 | 0.31 | 0.00 | 0.00 | 0.58 | 0.29 | 25 | T Cell |
| Actg1 | 0.37 | 0.00 | 0.00 | 0.92 | 0.76 | 25 | T Cell |
| G3bp2 | 0.35 | 0.00 | 0.00 | 0.60 | 0.31 | 25 | T Cell |
| Fip1l1 | 0.27 | 0.00 | 0.00 | 0.41 | 0.18 | 25 | T Cell |
| Taf15 | 0.30 | 0.00 | 0.00 | 0.27 | 0.10 | 25 | T Cell |
| Trmt112 | 0.27 | 0.00 | 0.00 | 0.35 | 0.14 | 25 | T Cell |
| Cct2 | 0.30 | 0.00 | 0.00 | 0.54 | 0.27 | 25 | T Cell |
| Psma2 | 0.38 | 0.00 | 0.00 | 0.72 | 0.44 | 25 | T Cell |
| Gm5611 | 0.43 | 0.00 | 0.00 | 0.62 | 0.36 | 25 | T Cell |
| Vps28 | 0.27 | 0.00 | 0.00 | 0.41 | 0.18 | 25 | T Cell |
| Srsf2 | 0.38 | 0.00 | 0.00 | 0.67 | 0.38 | 25 | T Cell |
| Baz1b | 0.27 | 0.00 | 0.00 | 0.54 | 0.27 | 25 | T Cell |
| Hnrnpm | 0.30 | 0.00 | 0.00 | 0.82 | 0.49 | 25 | T Cell |
| Srp9 | 0.29 | 0.00 | 0.00 | 0.56 | 0.28 | 25 | T Cell |
| Hnrnpul1 | 0.28 | 0.00 | 0.00 | 0.30 | 0.12 | 25 | T Cell |
| Rpl8 | 0.31 | 0.00 | 0.00 | 0.95 | 0.79 | 25 | T Cell |
| Ifngr1 | 0.35 | 0.00 | 0.00 | 0.49 | 0.24 | 25 | T Cell |
| Lyar | 0.35 | 0.00 | 0.00 | 0.36 | 0.16 | 25 | T Cell |
| Twistnb | 0.26 | 0.00 | 0.00 | 0.33 | 0.13 | 25 | T Cell |
| Rfc1 | 0.33 | 0.00 | 0.00 | 0.43 | 0.21 | 25 | T Cell |
| Eef1a1 | 0.27 | 0.00 | 0.00 | 0.97 | 0.87 | 25 | T Cell |
| Gm9825 | 0.26 | 0.00 | 0.00 | 0.32 | 0.13 | 25 | T Cell |
| Rps13 | 0.34 | 0.00 | 0.00 | 0.88 | 0.64 | 25 | T Cell |
| Rpl19 | 0.28 | 0.00 | 0.00 | 0.95 | 0.75 | 25 | T Cell |
| Rps3a3 | 0.31 | 0.00 | 0.00 | 0.95 | 0.78 | 25 | T Cell |
| Sumo2 | 0.33 | 0.00 | 0.00 | 0.58 | 0.30 | 25 | T Cell |
| Ddx21 | 0.38 | 0.00 | 0.00 | 0.51 | 0.26 | 25 | T Cell |
| Nolc1 | 0.34 | 0.00 | 0.00 | 0.39 | 0.18 | 25 | T Cell |
| Atp5a1 | 0.34 | 0.00 | 0.00 | 0.70 | 0.41 | 25 | T Cell |
| Calm2 | 0.35 | 0.00 | 0.00 | 0.90 | 0.64 | 25 | T Cell |
| Rpl6 | 0.27 | 0.00 | 0.00 | 0.99 | 0.87 | 25 | T Cell |
| Oaz1 | 0.37 | 0.00 | 0.00 | 0.86 | 0.62 | 25 | T Cell |
| Cdc123 | 0.28 | 0.00 | 0.00 | 0.31 | 0.13 | 25 | T Cell |
| Gm10269 | 0.30 | 0.00 | 0.00 | 0.95 | 0.79 | 25 | T Cell |
| Cct7 | 0.28 | 0.00 | 0.00 | 0.45 | 0.21 | 25 | T Cell |
| Gm2000 | 0.28 | 0.00 | 0.00 | 0.97 | 0.84 | 25 | T Cell |
| Hspa4 | 0.29 | 0.00 | 0.00 | 0.48 | 0.23 | 25 | T Cell |
| Eml4 | 0.35 | 0.00 | 0.00 | 0.38 | 0.18 | 25 | T Cell |
| Spcs2 | 0.31 | 0.00 | 0.00 | 0.58 | 0.31 | 25 | T Cell |
| Cdc42 | 0.35 | 0.00 | 0.00 | 0.82 | 0.54 | 25 | T Cell |
| Rpl22 | 0.32 | 0.00 | 0.00 | 0.94 | 0.79 | 25 | T Cell |
| Rexo2 | 0.28 | 0.00 | 0.00 | 0.34 | 0.15 | 25 | T Cell |
| Rpl18a | 0.26 | 0.00 | 0.00 | 0.99 | 0.82 | 25 | T Cell |
| Rpl39 | 0.27 | 0.00 | 0.00 | 0.95 | 0.80 | 25 | T Cell |
| 0610009D07Rik | 0.25 | 0.00 | 0.00 | 0.45 | 0.22 | 25 | T Cell |
| Hspd1 | 0.33 | 0.00 | 0.00 | 0.50 | 0.25 | 25 | T Cell |
| Ube2i | 0.27 | 0.00 | 0.00 | 0.55 | 0.29 | 25 | T Cell |
| Orc5 | 0.37 | 0.00 | 0.00 | 0.80 | 0.56 | 25 | T Cell |
| Capzb | 0.30 | 0.00 | 0.00 | 0.71 | 0.42 | 25 | T Cell |
| Fgfr1op2 | 0.32 | 0.00 | 0.00 | 0.53 | 0.28 | 25 | T Cell |
| Gm6472 | 0.29 | 0.00 | 0.00 | 0.95 | 0.80 | 25 | T Cell |
| Efr3a | 0.27 | 0.00 | 0.00 | 0.30 | 0.13 | 25 | T Cell |
| Capg | 0.27 | 0.00 | 0.00 | 0.27 | 0.11 | 25 | T Cell |
| Ngdn | 0.31 | 0.00 | 0.00 | 0.27 | 0.11 | 25 | T Cell |
| Dnaja2 | 0.29 | 0.00 | 0.00 | 0.50 | 0.25 | 25 | T Cell |
| Gm10260 | 0.33 | 0.00 | 0.00 | 0.75 | 0.49 | 25 | T Cell |
| Rpl10a | 0.29 | 0.00 | 0.00 | 0.93 | 0.74 | 25 | T Cell |
| Cct5 | 0.26 | 0.00 | 0.00 | 0.47 | 0.23 | 25 | T Cell |
| Gm11273 | 0.30 | 0.00 | 0.00 | 0.65 | 0.38 | 25 | T Cell |
| Psma1 | 0.27 | 0.00 | 0.00 | 0.50 | 0.26 | 25 | T Cell |
| Usp3 | 0.27 | 0.00 | 0.00 | 0.26 | 0.11 | 25 | T Cell |
| Gm8225 | 0.27 | 0.00 | 0.00 | 0.34 | 0.15 | 25 | T Cell |
| Gm4968 | 0.28 | 0.00 | 0.00 | 0.35 | 0.16 | 25 | T Cell |
| Mapre1 | 0.26 | 0.00 | 0.00 | 0.56 | 0.29 | 25 | T Cell |
| Rif1 | 0.27 | 0.00 | 0.00 | 0.36 | 0.17 | 25 | T Cell |
| Stk38 | 0.30 | 0.00 | 0.00 | 0.41 | 0.20 | 25 | T Cell |
| Bcas2 | 0.27 | 0.00 | 0.00 | 0.43 | 0.21 | 25 | T Cell |
| Tra2b | 0.29 | 0.00 | 0.00 | 0.55 | 0.29 | 25 | T Cell |
| Ube2l3 | 0.27 | 0.00 | 0.00 | 0.50 | 0.26 | 25 | T Cell |
| Trp53 | 0.26 | 0.00 | 0.00 | 0.35 | 0.16 | 25 | T Cell |
| Gm17511 | 0.31 | 0.00 | 0.00 | 0.87 | 0.61 | 25 | T Cell |
| Sf3b1 | 0.28 | 0.00 | 0.00 | 0.75 | 0.46 | 25 | T Cell |
| Yeats4 | 0.28 | 0.00 | 0.00 | 0.39 | 0.19 | 25 | T Cell |
| Tpr | 0.28 | 0.00 | 0.00 | 0.78 | 0.50 | 25 | T Cell |
| Cox8a | 0.30 | 0.00 | 0.00 | 0.83 | 0.56 | 25 | T Cell |
| Rpl22l1 | 0.33 | 0.00 | 0.00 | 0.79 | 0.56 | 25 | T Cell |
| Rpl23a | 0.26 | 0.00 | 0.00 | 0.90 | 0.66 | 25 | T Cell |
| Rps7 | 0.28 | 0.00 | 0.00 | 0.92 | 0.74 | 25 | T Cell |
| Cox5b | 0.26 | 0.00 | 0.00 | 0.67 | 0.39 | 25 | T Cell |
| Rps17 | 0.26 | 0.00 | 0.00 | 0.92 | 0.77 | 25 | T Cell |
| Ctdspl2 | 0.34 | 0.00 | 0.00 | 0.27 | 0.11 | 25 | T Cell |
| Rps18-ps3 | 0.33 | 0.00 | 0.00 | 0.80 | 0.56 | 25 | T Cell |
| Gm11808 | 0.26 | 0.00 | 0.00 | 0.94 | 0.77 | 25 | T Cell |
| Cast | 0.29 | 0.00 | 0.00 | 0.40 | 0.20 | 25 | T Cell |
| Rsl1d1 | 0.29 | 0.00 | 0.00 | 0.51 | 0.27 | 25 | T Cell |
| Rpl13a-ps1 | 0.28 | 0.00 | 0.00 | 0.81 | 0.55 | 25 | T Cell |
| Rps10 | 0.28 | 0.00 | 0.00 | 0.88 | 0.70 | 25 | T Cell |
| Cdkn1b | 0.28 | 0.00 | 0.00 | 0.42 | 0.21 | 25 | T Cell |
| Gm6139 | 0.28 | 0.00 | 0.00 | 0.70 | 0.42 | 25 | T Cell |
| Arpc5 | 0.29 | 0.00 | 0.00 | 0.82 | 0.56 | 25 | T Cell |
| Ilf2 | 0.25 | 0.00 | 0.00 | 0.39 | 0.19 | 25 | T Cell |
| Rps2-ps6 | 0.31 | 0.00 | 0.00 | 0.65 | 0.41 | 25 | T Cell |
| Atp5o | 0.28 | 0.00 | 0.00 | 0.61 | 0.35 | 25 | T Cell |
| Tagln2 | 0.33 | 0.00 | 0.00 | 0.69 | 0.45 | 25 | T Cell |
| Rpl18 | 0.28 | 0.00 | 0.00 | 0.85 | 0.64 | 25 | T Cell |
| Atp5j | 0.26 | 0.00 | 0.00 | 0.73 | 0.46 | 25 | T Cell |
| Id2 | 0.73 | 0.00 | 0.00 | 0.49 | 0.33 | 25 | T Cell |
| Rpl36al | 0.26 | 0.00 | 0.00 | 0.79 | 0.52 | 25 | T Cell |
| Atp5b | 0.28 | 0.00 | 0.00 | 0.69 | 0.44 | 25 | T Cell |
| Ndufa1 | 0.25 | 0.00 | 0.00 | 0.55 | 0.31 | 25 | T Cell |
| Rps12-ps3 | 0.26 | 0.00 | 0.00 | 0.93 | 0.79 | 25 | T Cell |
| Rpl23a-ps1 | 0.25 | 0.00 | 0.00 | 0.33 | 0.16 | 25 | T Cell |
| Eef1g | 0.27 | 0.00 | 0.00 | 0.73 | 0.49 | 25 | T Cell |
| Rpl17 | 0.28 | 0.00 | 0.00 | 0.78 | 0.54 | 25 | T Cell |
| Gm6576 | 0.28 | 0.00 | 0.00 | 0.52 | 0.31 | 25 | T Cell |
| Atp5f1 | 0.26 | 0.00 | 0.00 | 0.74 | 0.47 | 25 | T Cell |
| Rps2-ps10 | 0.27 | 0.00 | 0.00 | 0.65 | 0.43 | 25 | T Cell |
| Paics | 0.26 | 0.00 | 0.00 | 0.40 | 0.23 | 25 | T Cell |
| Igkc | 2.44 | 0.00 | 0.00 | 0.88 | 0.05 | 26 | Fibroblast |
| Hist1h2ag | 2.43 | 0.00 | 0.00 | 0.89 | 0.11 | 26 | Fibroblast |
| Ighm | 2.40 | 0.00 | 0.00 | 0.94 | 0.07 | 26 | Fibroblast |
| Hist1h2ac | 2.32 | 0.00 | 0.00 | 0.87 | 0.10 | 26 | Fibroblast |
| Hist1h2ad | 2.29 | 0.00 | 0.00 | 0.85 | 0.10 | 26 | Fibroblast |
| Hist1h2ah | 2.25 | 0.00 | 0.00 | 0.87 | 0.09 | 26 | Fibroblast |
| Hist1h2an | 2.18 | 0.00 | 0.00 | 0.85 | 0.09 | 26 | Fibroblast |
| Ms4a1 | 2.17 | 0.00 | 0.00 | 0.91 | 0.05 | 26 | Fibroblast |
| Hist1h2ai | 2.17 | 0.00 | 0.00 | 0.84 | 0.10 | 26 | Fibroblast |
| Hist1h2af | 2.17 | 0.00 | 0.00 | 0.84 | 0.09 | 26 | Fibroblast |
| Top2a | 2.05 | 0.00 | 0.00 | 0.94 | 0.11 | 26 | Fibroblast |
| Hist1h2ae | 2.03 | 0.00 | 0.00 | 0.80 | 0.09 | 26 | Fibroblast |
| Ly6d | 2.02 | 0.00 | 0.00 | 0.82 | 0.04 | 26 | Fibroblast |
| Hist1h2ak | 2.01 | 0.00 | 0.00 | 0.79 | 0.07 | 26 | Fibroblast |
| Mki67 | 2.00 | 0.00 | 0.00 | 0.92 | 0.12 | 26 | Fibroblast |
| 2810417H13Rik | 1.99 | 0.00 | 0.00 | 0.78 | 0.08 | 26 | Fibroblast |
| Hist1h2ab | 1.95 | 0.00 | 0.00 | 0.77 | 0.09 | 26 | Fibroblast |
| Iglc2 | 1.91 | 0.00 | 0.00 | 0.64 | 0.03 | 26 | Fibroblast |
| Cd79a | 1.77 | 0.00 | 0.00 | 0.81 | 0.04 | 26 | Fibroblast |
| Nusap1 | 1.69 | 0.00 | 0.00 | 0.70 | 0.06 | 26 | Fibroblast |
| Cd79b | 1.69 | 0.00 | 0.00 | 0.81 | 0.05 | 26 | Fibroblast |
| Mzb1 | 1.29 | 0.00 | 0.00 | 0.52 | 0.02 | 26 | Fibroblast |
| Siglecg | 1.26 | 0.00 | 0.00 | 0.50 | 0.02 | 26 | Fibroblast |
| Bank1 | 1.24 | 0.00 | 0.00 | 0.46 | 0.03 | 26 | Fibroblast |
| Fcrla | 1.18 | 0.00 | 0.00 | 0.45 | 0.02 | 26 | Fibroblast |
| Fam129c | 1.10 | 0.00 | 0.00 | 0.43 | 0.02 | 26 | Fibroblast |
| Cd19 | 1.09 | 0.00 | 0.00 | 0.44 | 0.02 | 26 | Fibroblast |
| Pax5 | 1.09 | 0.00 | 0.00 | 0.44 | 0.02 | 26 | Fibroblast |
| Tnfrsf13c | 1.07 | 0.00 | 0.00 | 0.40 | 0.02 | 26 | Fibroblast |
| Gm19980 | 0.95 | 0.00 | 0.00 | 0.34 | 0.01 | 26 | Fibroblast |
| Spib | 0.85 | 0.00 | 0.00 | 0.33 | 0.01 | 26 | Fibroblast |
| Pou2af1 | 0.82 | 0.00 | 0.00 | 0.31 | 0.01 | 26 | Fibroblast |
| Hist1h2ap | 2.58 | 0.00 | 0.00 | 0.94 | 0.15 | 26 | Fibroblast |
| Cd37 | 1.50 | 0.00 | 0.00 | 0.71 | 0.07 | 26 | Fibroblast |
| Hist1h2ao | 2.56 | 0.00 | 0.00 | 0.91 | 0.15 | 26 | Fibroblast |
| Blnk | 0.78 | 0.00 | 0.00 | 0.31 | 0.01 | 26 | Fibroblast |
| Ncapg | 1.22 | 0.00 | 0.00 | 0.51 | 0.04 | 26 | Fibroblast |
| Ptprcap | 1.21 | 0.00 | 0.00 | 0.66 | 0.07 | 26 | Fibroblast |
| H2afx | 1.64 | 0.00 | 0.00 | 0.72 | 0.09 | 26 | Fibroblast |
| Faim3 | 0.87 | 0.00 | 0.00 | 0.26 | 0.01 | 26 | Fibroblast |
| Prc1 | 1.27 | 0.00 | 0.00 | 0.61 | 0.07 | 26 | Fibroblast |
| Incenp | 1.41 | 0.00 | 0.00 | 0.66 | 0.08 | 26 | Fibroblast |
| Birc5 | 1.21 | 0.00 | 0.00 | 0.59 | 0.06 | 26 | Fibroblast |
| Phgdh | 0.86 | 0.00 | 0.00 | 0.35 | 0.02 | 26 | Fibroblast |
| Esco2 | 1.20 | 0.00 | 0.00 | 0.41 | 0.03 | 26 | Fibroblast |
| Ly86 | 1.11 | 0.00 | 0.00 | 0.59 | 0.07 | 26 | Fibroblast |
| Hist1h1b | 1.40 | 0.00 | 0.00 | 0.43 | 0.04 | 26 | Fibroblast |
| Cdca3 | 0.98 | 0.00 | 0.00 | 0.42 | 0.04 | 26 | Fibroblast |
| Cdca8 | 1.13 | 0.00 | 0.00 | 0.49 | 0.05 | 26 | Fibroblast |
| Sgol1 | 0.99 | 0.00 | 0.00 | 0.37 | 0.03 | 26 | Fibroblast |
| Hmmr | 1.22 | 0.00 | 0.00 | 0.46 | 0.04 | 26 | Fibroblast |
| Ube2c | 1.46 | 0.00 | 0.00 | 0.49 | 0.05 | 26 | Fibroblast |
| C130026I21Rik | 0.94 | 0.00 | 0.00 | 0.37 | 0.03 | 26 | Fibroblast |
| Cd74 | 1.32 | 0.00 | 0.00 | 0.75 | 0.12 | 26 | Fibroblast |
| Ncapd2 | 1.04 | 0.00 | 0.00 | 0.44 | 0.04 | 26 | Fibroblast |
| Aurkb | 0.88 | 0.00 | 0.00 | 0.34 | 0.03 | 26 | Fibroblast |
| Cd2 | 1.10 | 0.00 | 0.00 | 0.44 | 0.04 | 26 | Fibroblast |
| Kif11 | 1.02 | 0.00 | 0.00 | 0.41 | 0.04 | 26 | Fibroblast |
| Cenpe | 1.37 | 0.00 | 0.00 | 0.58 | 0.08 | 26 | Fibroblast |
| Bcl11a | 0.96 | 0.00 | 0.00 | 0.36 | 0.03 | 26 | Fibroblast |
| Cdca2 | 0.80 | 0.00 | 0.00 | 0.29 | 0.02 | 26 | Fibroblast |
| Aspm | 1.04 | 0.00 | 0.00 | 0.36 | 0.03 | 26 | Fibroblast |
| Smc2 | 1.30 | 0.00 | 0.00 | 0.75 | 0.13 | 26 | Fibroblast |
| Rac2 | 1.17 | 0.00 | 0.00 | 0.69 | 0.11 | 26 | Fibroblast |
| H2-Ab1 | 1.31 | 0.00 | 0.00 | 0.61 | 0.09 | 26 | Fibroblast |
| Sp140 | 1.15 | 0.00 | 0.00 | 0.54 | 0.07 | 26 | Fibroblast |
| Gm15987 | 0.70 | 0.00 | 0.00 | 0.25 | 0.02 | 26 | Fibroblast |
| Kif15 | 0.99 | 0.00 | 0.00 | 0.42 | 0.04 | 26 | Fibroblast |
| Clspn | 1.09 | 0.00 | 0.00 | 0.41 | 0.04 | 26 | Fibroblast |
| Cenpa | 1.19 | 0.00 | 0.00 | 0.51 | 0.06 | 26 | Fibroblast |
| Plac8 | 1.14 | 0.00 | 0.00 | 0.83 | 0.17 | 26 | Fibroblast |
| Ccna2 | 0.93 | 0.00 | 0.00 | 0.38 | 0.04 | 26 | Fibroblast |
| Cenpf | 1.36 | 0.00 | 0.00 | 0.55 | 0.08 | 26 | Fibroblast |
| Cks2 | 1.10 | 0.00 | 0.00 | 0.56 | 0.08 | 26 | Fibroblast |
| Hmgb2 | 1.85 | 0.00 | 0.00 | 1.00 | 0.42 | 26 | Fibroblast |
| Prkcb | 0.75 | 0.00 | 0.00 | 0.30 | 0.02 | 26 | Fibroblast |
| Cd52 | 1.30 | 0.00 | 0.00 | 0.90 | 0.20 | 26 | Fibroblast |
| H2-Aa | 1.22 | 0.00 | 0.00 | 0.55 | 0.08 | 26 | Fibroblast |
| Kif20b | 1.08 | 0.00 | 0.00 | 0.49 | 0.07 | 26 | Fibroblast |
| Tacc3 | 1.01 | 0.00 | 0.00 | 0.44 | 0.06 | 26 | Fibroblast |
| Tnfrsf13b | 0.88 | 0.00 | 0.00 | 0.40 | 0.04 | 26 | Fibroblast |
| Kif23 | 1.05 | 0.00 | 0.00 | 0.44 | 0.06 | 26 | Fibroblast |
| RP23-45G16.5 | 1.08 | 0.00 | 0.00 | 0.41 | 0.05 | 26 | Fibroblast |
| Rad51 | 0.81 | 0.00 | 0.00 | 0.29 | 0.03 | 26 | Fibroblast |
| Vars | 0.96 | 0.00 | 0.00 | 0.52 | 0.08 | 26 | Fibroblast |
| Casc5 | 0.97 | 0.00 | 0.00 | 0.39 | 0.05 | 26 | Fibroblast |
| Spc24 | 0.91 | 0.00 | 0.00 | 0.41 | 0.05 | 26 | Fibroblast |
| Ndc80 | 0.93 | 0.00 | 0.00 | 0.36 | 0.04 | 26 | Fibroblast |
| H2-Eb1 | 1.09 | 0.00 | 0.00 | 0.48 | 0.07 | 26 | Fibroblast |
| Ccnb2 | 1.05 | 0.00 | 0.00 | 0.44 | 0.06 | 26 | Fibroblast |
| Rrm2 | 1.07 | 0.00 | 0.00 | 0.39 | 0.05 | 26 | Fibroblast |
| Sgol2 | 0.94 | 0.00 | 0.00 | 0.36 | 0.04 | 26 | Fibroblast |
| Uchl3 | 1.02 | 0.00 | 0.00 | 0.46 | 0.07 | 26 | Fibroblast |
| Hist1h4i | 0.72 | 0.00 | 0.00 | 0.27 | 0.02 | 26 | Fibroblast |
| H2-DMb2 | 0.80 | 0.00 | 0.00 | 0.25 | 0.02 | 26 | Fibroblast |
| Cenph | 0.81 | 0.00 | 0.00 | 0.33 | 0.04 | 26 | Fibroblast |
| A530032D15Rik | 0.72 | 0.00 | 0.00 | 0.26 | 0.02 | 26 | Fibroblast |
| Hist2h2ab | 0.91 | 0.00 | 0.00 | 0.35 | 0.04 | 26 | Fibroblast |
| Tpx2 | 0.92 | 0.00 | 0.00 | 0.41 | 0.05 | 26 | Fibroblast |
| Fbxo5 | 0.90 | 0.00 | 0.00 | 0.37 | 0.05 | 26 | Fibroblast |
| Lrmp | 0.78 | 0.00 | 0.00 | 0.32 | 0.03 | 26 | Fibroblast |
| Cks1b | 1.03 | 0.00 | 0.00 | 0.54 | 0.09 | 26 | Fibroblast |
| Btk | 0.81 | 0.00 | 0.00 | 0.33 | 0.04 | 26 | Fibroblast |
| Ikzf3 | 0.73 | 0.00 | 0.00 | 0.25 | 0.02 | 26 | Fibroblast |
| Dnajc9 | 1.13 | 0.00 | 0.00 | 0.60 | 0.12 | 26 | Fibroblast |
| Kif22 | 0.67 | 0.00 | 0.00 | 0.26 | 0.03 | 26 | Fibroblast |
| Fam111a | 1.17 | 0.00 | 0.00 | 0.68 | 0.15 | 26 | Fibroblast |
| D17H6S56E-5 | 1.29 | 0.00 | 0.00 | 0.56 | 0.10 | 26 | Fibroblast |
| Cdc20 | 0.95 | 0.00 | 0.00 | 0.37 | 0.05 | 26 | Fibroblast |
| Hist1h1d | 1.38 | 0.00 | 0.00 | 0.54 | 0.10 | 26 | Fibroblast |
| Ckap2l | 0.86 | 0.00 | 0.00 | 0.36 | 0.05 | 26 | Fibroblast |
| Syk | 0.85 | 0.00 | 0.00 | 0.41 | 0.06 | 26 | Fibroblast |
| Dlgap5 | 0.74 | 0.00 | 0.00 | 0.26 | 0.03 | 26 | Fibroblast |
| Nuf2 | 0.71 | 0.00 | 0.00 | 0.29 | 0.03 | 26 | Fibroblast |
| Hist1h1e | 1.56 | 0.00 | 0.00 | 0.66 | 0.16 | 26 | Fibroblast |
| Arhgdib | 1.00 | 0.00 | 0.00 | 0.70 | 0.16 | 26 | Fibroblast |
| Uhrf1 | 0.77 | 0.00 | 0.00 | 0.27 | 0.03 | 26 | Fibroblast |
| Mis18bp1 | 0.76 | 0.00 | 0.00 | 0.29 | 0.03 | 26 | Fibroblast |
| Smc4 | 1.17 | 0.00 | 0.00 | 0.90 | 0.34 | 26 | Fibroblast |
| Lmnb1 | 0.96 | 0.00 | 0.00 | 0.47 | 0.09 | 26 | Fibroblast |
| Gmnn | 0.86 | 0.00 | 0.00 | 0.42 | 0.07 | 26 | Fibroblast |
| Arhgap11a | 0.75 | 0.00 | 0.00 | 0.32 | 0.04 | 26 | Fibroblast |
| Ebf1 | 0.76 | 0.00 | 0.00 | 0.33 | 0.05 | 26 | Fibroblast |
| Ifi30 | 1.03 | 0.00 | 0.00 | 0.52 | 0.11 | 26 | Fibroblast |
| Apobec3 | 0.89 | 0.00 | 0.00 | 0.44 | 0.08 | 26 | Fibroblast |
| Uchl4 | 0.89 | 0.00 | 0.00 | 0.37 | 0.06 | 26 | Fibroblast |
| Ptma | 0.88 | 0.00 | 0.00 | 1.00 | 0.91 | 26 | Fibroblast |
| Cep55 | 0.64 | 0.00 | 0.00 | 0.25 | 0.03 | 26 | Fibroblast |
| Lig1 | 1.11 | 0.00 | 0.00 | 0.47 | 0.09 | 26 | Fibroblast |
| Cdk1 | 0.81 | 0.00 | 0.00 | 0.35 | 0.05 | 26 | Fibroblast |
| Cnp | 0.67 | 0.00 | 0.00 | 0.35 | 0.05 | 26 | Fibroblast |
| Ifi203 | 0.89 | 0.00 | 0.00 | 0.60 | 0.14 | 26 | Fibroblast |
| Hells | 0.85 | 0.00 | 0.00 | 0.36 | 0.06 | 26 | Fibroblast |
| Cd48 | 0.69 | 0.00 | 0.00 | 0.33 | 0.05 | 26 | Fibroblast |
| Hirip3 | 1.01 | 0.00 | 0.00 | 0.45 | 0.09 | 26 | Fibroblast |
| Mndal | 0.90 | 0.00 | 0.00 | 0.64 | 0.16 | 26 | Fibroblast |
| Sub1 | 1.16 | 0.00 | 0.00 | 0.93 | 0.51 | 26 | Fibroblast |
| Wdr76 | 0.66 | 0.00 | 0.00 | 0.27 | 0.04 | 26 | Fibroblast |
| Rad51ap1 | 0.66 | 0.00 | 0.00 | 0.27 | 0.03 | 26 | Fibroblast |
| AI662270 | 0.72 | 0.00 | 0.00 | 0.40 | 0.07 | 26 | Fibroblast |
| Ezh2 | 0.97 | 0.00 | 0.00 | 0.67 | 0.19 | 26 | Fibroblast |
| Pim1 | 0.99 | 0.00 | 0.00 | 0.46 | 0.09 | 26 | Fibroblast |
| Parp1 | 0.83 | 0.00 | 0.00 | 0.48 | 0.10 | 26 | Fibroblast |
| Rassf4 | 0.74 | 0.00 | 0.00 | 0.29 | 0.04 | 26 | Fibroblast |
| Cmah | 0.71 | 0.00 | 0.00 | 0.28 | 0.04 | 26 | Fibroblast |
| Coro1a | 0.75 | 0.00 | 0.00 | 0.49 | 0.10 | 26 | Fibroblast |
| Ptpn6 | 0.71 | 0.00 | 0.00 | 0.44 | 0.08 | 26 | Fibroblast |
| Anp32e | 1.12 | 0.00 | 0.00 | 0.78 | 0.28 | 26 | Fibroblast |
| Rilpl2 | 0.76 | 0.00 | 0.00 | 0.37 | 0.07 | 26 | Fibroblast |
| Ran | 1.04 | 0.00 | 0.00 | 0.87 | 0.38 | 26 | Fibroblast |
| Tbc1d10c | 0.56 | 0.00 | 0.00 | 0.25 | 0.03 | 26 | Fibroblast |
| Ptprc | 0.63 | 0.00 | 0.00 | 0.62 | 0.15 | 26 | Fibroblast |
| Tec | 0.67 | 0.00 | 0.00 | 0.26 | 0.03 | 26 | Fibroblast |
| Ifi27l2a | 0.81 | 0.00 | 0.00 | 0.45 | 0.09 | 26 | Fibroblast |
| Ncf4 | 0.57 | 0.00 | 0.00 | 0.26 | 0.04 | 26 | Fibroblast |
| Dut | 0.94 | 0.00 | 0.00 | 0.42 | 0.09 | 26 | Fibroblast |
| Gm10184 | 0.78 | 0.00 | 0.00 | 0.36 | 0.07 | 26 | Fibroblast |
| Hist2h2ac | 0.79 | 0.00 | 0.00 | 0.35 | 0.06 | 26 | Fibroblast |
| Dnmt1 | 0.88 | 0.00 | 0.00 | 0.46 | 0.10 | 26 | Fibroblast |
| Cenpw | 0.65 | 0.00 | 0.00 | 0.31 | 0.05 | 26 | Fibroblast |
| Nucks1 | 1.00 | 0.00 | 0.00 | 0.77 | 0.28 | 26 | Fibroblast |
| Anp32b | 0.94 | 0.00 | 0.00 | 0.93 | 0.50 | 26 | Fibroblast |
| Sec11c | 0.93 | 0.00 | 0.00 | 0.61 | 0.18 | 26 | Fibroblast |
| Arhgap15 | 0.72 | 0.00 | 0.00 | 0.31 | 0.05 | 26 | Fibroblast |
| Knstrn | 0.67 | 0.00 | 0.00 | 0.27 | 0.04 | 26 | Fibroblast |
| Kpna2 | 0.84 | 0.00 | 0.00 | 0.34 | 0.06 | 26 | Fibroblast |
| 1-Sep | 0.57 | 0.00 | 0.00 | 0.27 | 0.04 | 26 | Fibroblast |
| A430104N18Rik | 0.63 | 0.00 | 0.00 | 0.36 | 0.06 | 26 | Fibroblast |
| Sp110 | 0.72 | 0.00 | 0.00 | 0.37 | 0.07 | 26 | Fibroblast |
| Gm10259 | 0.79 | 0.00 | 0.00 | 0.33 | 0.06 | 26 | Fibroblast |
| Pola1 | 0.63 | 0.00 | 0.00 | 0.28 | 0.04 | 26 | Fibroblast |
| Cenpk | 0.62 | 0.00 | 0.00 | 0.25 | 0.04 | 26 | Fibroblast |
| Nrm | 0.67 | 0.00 | 0.00 | 0.31 | 0.05 | 26 | Fibroblast |
| Lsp1 | 0.86 | 0.00 | 0.00 | 0.65 | 0.19 | 26 | Fibroblast |
| Ralgps2 | 0.75 | 0.00 | 0.00 | 0.37 | 0.07 | 26 | Fibroblast |
| Prim1 | 0.86 | 0.00 | 0.00 | 0.31 | 0.05 | 26 | Fibroblast |
| Snx5 | 0.85 | 0.00 | 0.00 | 0.50 | 0.13 | 26 | Fibroblast |
| Dek | 1.06 | 0.00 | 0.00 | 0.89 | 0.50 | 26 | Fibroblast |
| Tuba1b | 0.93 | 0.00 | 0.00 | 0.89 | 0.45 | 26 | Fibroblast |
| Pcna-ps2 | 1.16 | 0.00 | 0.00 | 0.57 | 0.18 | 26 | Fibroblast |
| Diap3 | 0.71 | 0.00 | 0.00 | 0.33 | 0.06 | 26 | Fibroblast |
| Tcof1 | 0.73 | 0.00 | 0.00 | 0.45 | 0.11 | 26 | Fibroblast |
| Hist2h2aa2 | 0.94 | 0.00 | 0.00 | 0.50 | 0.14 | 26 | Fibroblast |
| Spc25 | 0.67 | 0.00 | 0.00 | 0.29 | 0.05 | 26 | Fibroblast |
| Hist1h1c | 1.03 | 0.00 | 0.00 | 0.45 | 0.11 | 26 | Fibroblast |
| Ncf1 | 0.65 | 0.00 | 0.00 | 0.35 | 0.07 | 26 | Fibroblast |
| Sell | 0.65 | 0.00 | 0.00 | 0.25 | 0.04 | 26 | Fibroblast |
| Arid3a | 0.85 | 0.00 | 0.00 | 0.43 | 0.10 | 26 | Fibroblast |
| Racgap1 | 0.62 | 0.00 | 0.00 | 0.25 | 0.04 | 26 | Fibroblast |
| Trim59 | 0.66 | 0.00 | 0.00 | 0.27 | 0.04 | 26 | Fibroblast |
| H2afz | 0.91 | 0.00 | 0.00 | 0.83 | 0.40 | 26 | Fibroblast |
| Gm4204 | 0.85 | 0.00 | 0.00 | 0.91 | 0.50 | 26 | Fibroblast |
| Sfxn1 | 0.62 | 0.00 | 0.00 | 0.34 | 0.07 | 26 | Fibroblast |
| 2700094K13Rik | 0.77 | 0.00 | 0.00 | 0.55 | 0.16 | 26 | Fibroblast |
| Pcna | 1.05 | 0.00 | 0.00 | 0.56 | 0.18 | 26 | Fibroblast |
| Cd24a | 0.95 | 0.00 | 0.00 | 0.48 | 0.13 | 26 | Fibroblast |
| Fam65b | 0.52 | 0.00 | 0.00 | 0.26 | 0.04 | 26 | Fibroblast |
| Stmn1 | 0.98 | 0.00 | 0.00 | 0.65 | 0.23 | 26 | Fibroblast |
| Ass1 | 0.57 | 0.00 | 0.00 | 0.26 | 0.04 | 26 | Fibroblast |
| Capg | 0.70 | 0.00 | 0.00 | 0.42 | 0.10 | 26 | Fibroblast |
| Ncl | 0.82 | 0.00 | 0.00 | 0.96 | 0.78 | 26 | Fibroblast |
| Cd53 | 0.66 | 0.00 | 0.00 | 0.40 | 0.09 | 26 | Fibroblast |
| Tubb5 | 0.86 | 0.00 | 0.00 | 0.86 | 0.44 | 26 | Fibroblast |
| Dbf4 | 0.60 | 0.00 | 0.00 | 0.28 | 0.05 | 26 | Fibroblast |
| Snx2 | 0.78 | 0.00 | 0.00 | 0.53 | 0.17 | 26 | Fibroblast |
| Was | 0.45 | 0.00 | 0.00 | 0.29 | 0.06 | 26 | Fibroblast |
| Slbp | 0.89 | 0.00 | 0.00 | 0.49 | 0.15 | 26 | Fibroblast |
| Nap1l1 | 0.78 | 0.00 | 0.00 | 0.87 | 0.47 | 26 | Fibroblast |
| Pyhin1 | 0.53 | 0.00 | 0.00 | 0.26 | 0.05 | 26 | Fibroblast |
| Man1a | 0.60 | 0.00 | 0.00 | 0.26 | 0.05 | 26 | Fibroblast |
| Rps5 | 0.63 | 0.00 | 0.00 | 0.97 | 0.78 | 26 | Fibroblast |
| Ckap2 | 0.62 | 0.00 | 0.00 | 0.25 | 0.05 | 26 | Fibroblast |
| Lbr | 0.64 | 0.00 | 0.00 | 0.45 | 0.12 | 26 | Fibroblast |
| Rrm1 | 0.69 | 0.00 | 0.00 | 0.31 | 0.07 | 26 | Fibroblast |
| Ptpn18 | 0.63 | 0.00 | 0.00 | 0.49 | 0.14 | 26 | Fibroblast |
| Sh3bgrl3 | 0.71 | 0.00 | 0.00 | 0.79 | 0.34 | 26 | Fibroblast |
| Cep110 | 0.73 | 0.00 | 0.00 | 0.57 | 0.19 | 26 | Fibroblast |
| Cbx5 | 0.82 | 0.00 | 0.00 | 0.66 | 0.26 | 26 | Fibroblast |
| Rnase6 | 0.39 | 0.00 | 0.00 | 0.27 | 0.05 | 26 | Fibroblast |
| Rps26 | 0.57 | 0.00 | 0.00 | 0.99 | 0.90 | 26 | Fibroblast |
| Psmb9 | 0.67 | 0.00 | 0.00 | 0.36 | 0.09 | 26 | Fibroblast |
| Gm10123 | 0.61 | 0.00 | 0.00 | 0.97 | 0.81 | 26 | Fibroblast |
| Atad5 | 0.56 | 0.00 | 0.00 | 0.27 | 0.05 | 26 | Fibroblast |
| Serbp1 | 0.67 | 0.00 | 0.00 | 0.97 | 0.75 | 26 | Fibroblast |
| Hn1 | 0.74 | 0.00 | 0.00 | 0.57 | 0.20 | 26 | Fibroblast |
| 6-Sep | 0.53 | 0.00 | 0.00 | 0.31 | 0.07 | 26 | Fibroblast |
| Rpl4 | 0.58 | 0.00 | 0.00 | 0.98 | 0.83 | 26 | Fibroblast |
| Mef2c | 0.71 | 0.00 | 0.00 | 0.51 | 0.16 | 26 | Fibroblast |
| Npm1 | 0.84 | 0.00 | 0.00 | 0.89 | 0.58 | 26 | Fibroblast |
| Ppil1 | 0.53 | 0.00 | 0.00 | 0.27 | 0.06 | 26 | Fibroblast |
| Usp1 | 0.76 | 0.00 | 0.00 | 0.49 | 0.15 | 26 | Fibroblast |
| Unc93b1 | 0.59 | 0.00 | 0.00 | 0.32 | 0.07 | 26 | Fibroblast |
| Serp1 | 0.67 | 0.00 | 0.00 | 0.62 | 0.23 | 26 | Fibroblast |
| Tyms | 0.68 | 0.00 | 0.00 | 0.27 | 0.06 | 26 | Fibroblast |
| Rps26-ps1 | 0.59 | 0.00 | 0.00 | 0.98 | 0.87 | 26 | Fibroblast |
| Whsc1 | 0.77 | 0.00 | 0.00 | 0.45 | 0.14 | 26 | Fibroblast |
| Atic | 0.53 | 0.00 | 0.00 | 0.28 | 0.06 | 26 | Fibroblast |
| Gimap4 | 0.64 | 0.00 | 0.00 | 0.54 | 0.18 | 26 | Fibroblast |
| Crip1 | 0.91 | 0.00 | 0.00 | 0.87 | 0.48 | 26 | Fibroblast |
| Nme1 | 0.71 | 0.00 | 0.00 | 0.54 | 0.19 | 26 | Fibroblast |
| Rpia | 0.56 | 0.00 | 0.00 | 0.29 | 0.07 | 26 | Fibroblast |
| Hmga1 | 0.63 | 0.00 | 0.00 | 0.32 | 0.08 | 26 | Fibroblast |
| Mcm6 | 0.63 | 0.00 | 0.00 | 0.30 | 0.07 | 26 | Fibroblast |
| Arhgap30 | 0.49 | 0.00 | 0.00 | 0.39 | 0.10 | 26 | Fibroblast |
| Gmfg | 0.68 | 0.00 | 0.00 | 0.46 | 0.14 | 26 | Fibroblast |
| Rps3a1 | 0.53 | 0.00 | 0.00 | 0.97 | 0.85 | 26 | Fibroblast |
| Mcm7 | 0.70 | 0.00 | 0.00 | 0.31 | 0.07 | 26 | Fibroblast |
| Sh3bp5 | 0.68 | 0.00 | 0.00 | 0.56 | 0.19 | 26 | Fibroblast |
| Hmgn2 | 0.72 | 0.00 | 0.00 | 0.82 | 0.43 | 26 | Fibroblast |
| Tuba1c | 0.72 | 0.00 | 0.00 | 0.72 | 0.32 | 26 | Fibroblast |
| Psmb8 | 0.63 | 0.00 | 0.00 | 0.54 | 0.18 | 26 | Fibroblast |
| Pld4 | 0.51 | 0.00 | 0.00 | 0.26 | 0.06 | 26 | Fibroblast |
| Nol7 | 0.79 | 0.00 | 0.00 | 0.73 | 0.35 | 26 | Fibroblast |
| Ppia | 0.58 | 0.00 | 0.00 | 0.96 | 0.80 | 26 | Fibroblast |
| Dctpp1 | 0.55 | 0.00 | 0.00 | 0.26 | 0.05 | 26 | Fibroblast |
| Ezr | 0.67 | 0.00 | 0.00 | 0.59 | 0.22 | 26 | Fibroblast |
| Ranbp1 | 0.80 | 0.00 | 0.00 | 0.65 | 0.29 | 26 | Fibroblast |
| Chd1 | 0.71 | 0.00 | 0.00 | 0.50 | 0.17 | 26 | Fibroblast |
| Ncapd3 | 0.57 | 0.00 | 0.00 | 0.26 | 0.06 | 26 | Fibroblast |
| Ddx39 | 0.58 | 0.00 | 0.00 | 0.34 | 0.09 | 26 | Fibroblast |
| Hcls1 | 0.45 | 0.00 | 0.00 | 0.31 | 0.07 | 26 | Fibroblast |
| Pa2g4 | 0.77 | 0.00 | 0.00 | 0.62 | 0.26 | 26 | Fibroblast |
| Dnajc2 | 0.75 | 0.00 | 0.00 | 0.53 | 0.20 | 26 | Fibroblast |
| Nubp1 | 0.50 | 0.00 | 0.00 | 0.26 | 0.06 | 26 | Fibroblast |
| Coro2a | 0.52 | 0.00 | 0.00 | 0.28 | 0.07 | 26 | Fibroblast |
| Rpl14 | 0.50 | 0.00 | 0.00 | 0.98 | 0.84 | 26 | Fibroblast |
| Nasp | 0.78 | 0.00 | 0.00 | 0.53 | 0.20 | 26 | Fibroblast |
| Atad2 | 0.63 | 0.00 | 0.00 | 0.31 | 0.08 | 26 | Fibroblast |
| Apbb1ip | 0.44 | 0.00 | 0.00 | 0.36 | 0.09 | 26 | Fibroblast |
| Lcp1 | 0.59 | 0.00 | 0.00 | 0.59 | 0.23 | 26 | Fibroblast |
| Rps11 | 0.58 | 0.00 | 0.00 | 0.93 | 0.76 | 26 | Fibroblast |
| Smchd1 | 0.63 | 0.00 | 0.00 | 0.47 | 0.16 | 26 | Fibroblast |
| Csk | 0.47 | 0.00 | 0.00 | 0.27 | 0.06 | 26 | Fibroblast |
| Rpsa-ps10 | 0.61 | 0.00 | 0.00 | 0.94 | 0.68 | 26 | Fibroblast |
| Pycard | 0.56 | 0.00 | 0.00 | 0.34 | 0.09 | 26 | Fibroblast |
| Gm10282 | 0.71 | 0.00 | 0.00 | 0.77 | 0.40 | 26 | Fibroblast |
| Strbp | 0.66 | 0.00 | 0.00 | 0.39 | 0.12 | 26 | Fibroblast |
| Ssrp1 | 0.66 | 0.00 | 0.00 | 0.55 | 0.21 | 26 | Fibroblast |
| Gnl3 | 0.62 | 0.00 | 0.00 | 0.39 | 0.12 | 26 | Fibroblast |
| Rps3a2 | 0.49 | 0.00 | 0.00 | 0.96 | 0.82 | 26 | Fibroblast |
| Actb | 0.42 | 0.00 | 0.00 | 1.00 | 0.98 | 26 | Fibroblast |
| Fam107b | 0.50 | 0.00 | 0.00 | 0.29 | 0.07 | 26 | Fibroblast |
| Fermt3 | 0.50 | 0.00 | 0.00 | 0.27 | 0.06 | 26 | Fibroblast |
| Vrk1 | 0.51 | 0.00 | 0.00 | 0.30 | 0.08 | 26 | Fibroblast |
| Itga4 | 0.42 | 0.00 | 0.00 | 0.35 | 0.10 | 26 | Fibroblast |
| Ckap5 | 0.64 | 0.00 | 0.00 | 0.38 | 0.11 | 26 | Fibroblast |
| Arl6ip1 | 0.88 | 0.00 | 0.00 | 0.84 | 0.52 | 26 | Fibroblast |
| Set | 0.77 | 0.00 | 0.00 | 0.69 | 0.34 | 26 | Fibroblast |
| H2-K1 | 0.64 | 0.00 | 0.00 | 0.70 | 0.32 | 26 | Fibroblast |
| Dkc1 | 0.58 | 0.00 | 0.00 | 0.38 | 0.12 | 26 | Fibroblast |
| Uchl5 | 0.53 | 0.00 | 0.00 | 0.38 | 0.12 | 26 | Fibroblast |
| Rad21 | 0.64 | 0.00 | 0.00 | 0.57 | 0.23 | 26 | Fibroblast |
| Nop58 | 0.82 | 0.00 | 0.00 | 0.59 | 0.26 | 26 | Fibroblast |
| Fam49b | 0.45 | 0.00 | 0.00 | 0.36 | 0.11 | 26 | Fibroblast |
| Tmpo | 0.70 | 0.00 | 0.00 | 0.44 | 0.15 | 26 | Fibroblast |
| Cd55 | 0.63 | 0.00 | 0.00 | 0.29 | 0.07 | 26 | Fibroblast |
| Pabpc1 | 0.50 | 0.00 | 0.00 | 0.97 | 0.81 | 26 | Fibroblast |
| Lrrc58 | 0.47 | 0.00 | 0.00 | 1.00 | 0.91 | 26 | Fibroblast |
| Pkig | 0.54 | 0.00 | 0.00 | 0.31 | 0.09 | 26 | Fibroblast |
| Hsp90aa1 | 0.65 | 0.00 | 0.00 | 0.80 | 0.47 | 26 | Fibroblast |
| Nop56 | 0.74 | 0.00 | 0.00 | 0.50 | 0.20 | 26 | Fibroblast |
| Elf1 | 0.57 | 0.00 | 0.00 | 0.53 | 0.20 | 26 | Fibroblast |
| H2afv | 0.63 | 0.00 | 0.00 | 0.64 | 0.28 | 26 | Fibroblast |
| Srsf3 | 0.66 | 0.00 | 0.00 | 0.79 | 0.45 | 26 | Fibroblast |
| Rbm3 | 0.65 | 0.00 | 0.00 | 0.74 | 0.41 | 26 | Fibroblast |
| Rangap1 | 0.56 | 0.00 | 0.00 | 0.29 | 0.08 | 26 | Fibroblast |
| Cotl1 | 0.49 | 0.00 | 0.00 | 0.49 | 0.18 | 26 | Fibroblast |
| Ly6e | 0.51 | 0.00 | 0.00 | 0.84 | 0.42 | 26 | Fibroblast |
| Mcm3 | 0.52 | 0.00 | 0.00 | 0.27 | 0.07 | 26 | Fibroblast |
| Snrpd1 | 0.58 | 0.00 | 0.00 | 0.51 | 0.20 | 26 | Fibroblast |
| Pfn1 | 0.63 | 0.00 | 0.00 | 0.77 | 0.46 | 26 | Fibroblast |
| Tubb4b | 0.57 | 0.00 | 0.00 | 0.49 | 0.19 | 26 | Fibroblast |
| Swap70 | 0.62 | 0.00 | 0.00 | 0.35 | 0.11 | 26 | Fibroblast |
| Gm13139 | 0.45 | 0.00 | 0.00 | 0.27 | 0.07 | 26 | Fibroblast |
| Hnrnpa2b1 | 0.56 | 0.00 | 0.00 | 0.93 | 0.72 | 26 | Fibroblast |
| Hmgb1 | 0.60 | 0.00 | 0.00 | 0.88 | 0.61 | 26 | Fibroblast |
| Gm5641 | 0.68 | 0.00 | 0.00 | 0.70 | 0.37 | 26 | Fibroblast |
| Ak2 | 0.51 | 0.00 | 0.00 | 0.43 | 0.15 | 26 | Fibroblast |
| Rpl13a | 0.41 | 0.00 | 0.00 | 0.97 | 0.87 | 26 | Fibroblast |
| Gar1 | 0.45 | 0.00 | 0.00 | 0.26 | 0.07 | 26 | Fibroblast |
| Hnrnpu | 0.66 | 0.00 | 0.00 | 0.88 | 0.64 | 26 | Fibroblast |
| 4930523C07Rik | 0.58 | 0.00 | 0.00 | 0.48 | 0.19 | 26 | Fibroblast |
| Rabgap1l | 0.53 | 0.00 | 0.00 | 0.34 | 0.11 | 26 | Fibroblast |
| Brd8 | 0.61 | 0.00 | 0.00 | 0.49 | 0.19 | 26 | Fibroblast |
| Gm12355 | 0.61 | 0.00 | 0.00 | 0.58 | 0.26 | 26 | Fibroblast |
| Mycbp2 | 0.66 | 0.00 | 0.00 | 0.54 | 0.23 | 26 | Fibroblast |
| Ska2 | 0.52 | 0.00 | 0.00 | 0.25 | 0.07 | 26 | Fibroblast |
| Eef1b2 | 0.50 | 0.00 | 0.00 | 0.89 | 0.65 | 26 | Fibroblast |
| Snrpe | 0.55 | 0.00 | 0.00 | 0.70 | 0.36 | 26 | Fibroblast |
| Mif | 0.64 | 0.00 | 0.00 | 0.46 | 0.19 | 26 | Fibroblast |
| Zfp706 | 0.63 | 0.00 | 0.00 | 0.64 | 0.32 | 26 | Fibroblast |
| Supt16 | 0.63 | 0.00 | 0.00 | 0.56 | 0.25 | 26 | Fibroblast |
| Hmga1-rs1 | 0.48 | 0.00 | 0.00 | 0.27 | 0.08 | 26 | Fibroblast |
| Rpl32 | 0.45 | 0.00 | 0.00 | 0.95 | 0.84 | 26 | Fibroblast |
| Emp3 | 0.63 | 0.00 | 0.00 | 0.51 | 0.21 | 26 | Fibroblast |
| Slfn2 | 0.41 | 0.00 | 0.00 | 0.39 | 0.13 | 26 | Fibroblast |
| Gm7729 | 0.53 | 0.00 | 0.00 | 0.36 | 0.12 | 26 | Fibroblast |
| Wdr89 | 0.39 | 0.00 | 0.00 | 0.98 | 0.89 | 26 | Fibroblast |
| Hat1 | 0.43 | 0.00 | 0.00 | 0.26 | 0.07 | 26 | Fibroblast |
| Exosc3 | 0.54 | 0.00 | 0.00 | 0.26 | 0.07 | 26 | Fibroblast |
| Snx30 | 0.48 | 0.00 | 0.00 | 0.33 | 0.11 | 26 | Fibroblast |
| Calm1 | 0.42 | 0.00 | 0.00 | 0.99 | 0.87 | 26 | Fibroblast |
| Hjurp | 0.51 | 0.00 | 0.00 | 0.40 | 0.14 | 26 | Fibroblast |
| Tnfaip8 | 0.51 | 0.00 | 0.00 | 0.33 | 0.11 | 26 | Fibroblast |
| Gtf2f1 | 0.56 | 0.00 | 0.00 | 0.51 | 0.21 | 26 | Fibroblast |
| Gm6625 | 0.62 | 0.00 | 0.00 | 0.55 | 0.25 | 26 | Fibroblast |
| Erp29 | 0.54 | 0.00 | 0.00 | 0.57 | 0.26 | 26 | Fibroblast |
| Ctcf | 0.62 | 0.00 | 0.00 | 0.53 | 0.23 | 26 | Fibroblast |
| Ptbp3 | 0.60 | 0.00 | 0.00 | 0.70 | 0.36 | 26 | Fibroblast |
| Sms | 0.42 | 0.00 | 0.00 | 0.35 | 0.12 | 26 | Fibroblast |
| Rps16 | 0.42 | 0.00 | 0.00 | 0.98 | 0.84 | 26 | Fibroblast |
| Tipin | 0.60 | 0.00 | 0.00 | 0.31 | 0.10 | 26 | Fibroblast |
| Snrpa1 | 0.53 | 0.00 | 0.00 | 0.39 | 0.14 | 26 | Fibroblast |
| Gm8730 | 0.44 | 0.00 | 0.00 | 0.95 | 0.82 | 26 | Fibroblast |
| Eef1g | 0.61 | 0.00 | 0.00 | 0.77 | 0.49 | 26 | Fibroblast |
| Eif5a | 0.58 | 0.00 | 0.00 | 0.64 | 0.34 | 26 | Fibroblast |
| Snn | 0.43 | 0.00 | 0.00 | 0.27 | 0.08 | 26 | Fibroblast |
| Arhgap17 | 0.52 | 0.00 | 0.00 | 0.44 | 0.18 | 26 | Fibroblast |
| Rps3a3 | 0.43 | 0.00 | 0.00 | 0.93 | 0.78 | 26 | Fibroblast |
| Arpc5l | 0.52 | 0.00 | 0.00 | 0.42 | 0.17 | 26 | Fibroblast |
| Rps12-ps3 | 0.45 | 0.00 | 0.00 | 0.95 | 0.79 | 26 | Fibroblast |
| Klf2 | 0.56 | 0.00 | 0.00 | 0.48 | 0.20 | 26 | Fibroblast |
| Rps3 | 0.42 | 0.00 | 0.00 | 0.95 | 0.83 | 26 | Fibroblast |
| Limd2 | 0.48 | 0.00 | 0.00 | 0.41 | 0.15 | 26 | Fibroblast |
| H2afy | 0.59 | 0.00 | 0.00 | 0.46 | 0.20 | 26 | Fibroblast |
| Impdh2 | 0.41 | 0.00 | 0.00 | 0.31 | 0.10 | 26 | Fibroblast |
| Snrpf | 0.51 | 0.00 | 0.00 | 0.59 | 0.28 | 26 | Fibroblast |
| Crlf3 | 0.41 | 0.00 | 0.00 | 0.38 | 0.14 | 26 | Fibroblast |
| Rps9 | 0.36 | 0.00 | 0.00 | 0.98 | 0.91 | 26 | Fibroblast |
| Gm10275 | 0.45 | 0.00 | 0.00 | 0.89 | 0.66 | 26 | Fibroblast |
| H2-D1 | 0.43 | 0.00 | 0.00 | 0.85 | 0.49 | 26 | Fibroblast |
| Gm10260 | 0.52 | 0.00 | 0.00 | 0.79 | 0.49 | 26 | Fibroblast |
| Smarca5 | 0.58 | 0.00 | 0.00 | 0.62 | 0.30 | 26 | Fibroblast |
| Gm10269 | 0.40 | 0.00 | 0.00 | 0.96 | 0.79 | 26 | Fibroblast |
| Rbbp7 | 0.55 | 0.00 | 0.00 | 0.55 | 0.26 | 26 | Fibroblast |
| Sp100 | 0.37 | 0.00 | 0.00 | 0.28 | 0.08 | 26 | Fibroblast |
| Ccdc34 | 0.47 | 0.00 | 0.00 | 0.38 | 0.14 | 26 | Fibroblast |
| Cep57 | 0.50 | 0.00 | 0.00 | 0.32 | 0.11 | 26 | Fibroblast |
| Rpf2 | 0.49 | 0.00 | 0.00 | 0.28 | 0.09 | 26 | Fibroblast |
| Exosc8 | 0.39 | 0.00 | 0.00 | 0.27 | 0.08 | 26 | Fibroblast |
| Asnsd1 | 0.46 | 0.00 | 0.00 | 0.32 | 0.11 | 26 | Fibroblast |
| Tmsb4x | 0.32 | 0.00 | 0.00 | 0.99 | 0.89 | 26 | Fibroblast |
| Gimap1 | 0.40 | 0.00 | 0.00 | 0.39 | 0.14 | 26 | Fibroblast |
| Gm5611 | 0.64 | 0.00 | 0.00 | 0.63 | 0.36 | 26 | Fibroblast |
| Rplp1 | 0.35 | 0.00 | 0.00 | 0.99 | 0.90 | 26 | Fibroblast |
| Rps2 | 0.49 | 0.00 | 0.00 | 0.88 | 0.67 | 26 | Fibroblast |
| Rasgrp2 | 0.49 | 0.00 | 0.00 | 0.30 | 0.10 | 26 | Fibroblast |
| Gnb2l1 | 0.47 | 0.00 | 0.00 | 0.91 | 0.65 | 26 | Fibroblast |
| Gm6793 | 0.67 | 0.00 | 0.00 | 0.58 | 0.30 | 26 | Fibroblast |
| Rplp0 | 0.41 | 0.00 | 0.00 | 0.96 | 0.83 | 26 | Fibroblast |
| Grap | 0.45 | 0.00 | 0.00 | 0.37 | 0.13 | 26 | Fibroblast |
| Rpl8 | 0.41 | 0.00 | 0.00 | 0.95 | 0.79 | 26 | Fibroblast |
| Gimap9 | 0.38 | 0.00 | 0.00 | 0.25 | 0.07 | 26 | Fibroblast |
| Baz1a | 0.51 | 0.00 | 0.00 | 0.35 | 0.13 | 26 | Fibroblast |
| Tpm3 | 0.49 | 0.00 | 0.00 | 0.92 | 0.70 | 26 | Fibroblast |
| Erdr1 | 0.57 | 0.00 | 0.00 | 0.29 | 0.09 | 26 | Fibroblast |
| Fbl | 0.44 | 0.00 | 0.00 | 0.26 | 0.08 | 26 | Fibroblast |
| Yeats4 | 0.54 | 0.00 | 0.00 | 0.44 | 0.19 | 26 | Fibroblast |
| Nudt21 | 0.42 | 0.00 | 0.00 | 0.34 | 0.12 | 26 | Fibroblast |
| Lsm4 | 0.52 | 0.00 | 0.00 | 0.47 | 0.21 | 26 | Fibroblast |
| Actr3 | 0.47 | 0.00 | 0.00 | 0.64 | 0.33 | 26 | Fibroblast |
| G3bp1 | 0.54 | 0.00 | 0.00 | 0.42 | 0.17 | 26 | Fibroblast |
| Gm8186 | 0.52 | 0.00 | 0.00 | 0.68 | 0.39 | 26 | Fibroblast |
| Hnrnpf | 0.49 | 0.00 | 0.00 | 0.66 | 0.34 | 26 | Fibroblast |
| Rpsa | 0.48 | 0.00 | 0.00 | 0.86 | 0.62 | 26 | Fibroblast |
| mt-Co1 | 0.34 | 0.00 | 0.00 | 0.98 | 0.95 | 26 | Fibroblast |
| Gm8991 | 0.56 | 0.00 | 0.00 | 0.49 | 0.23 | 26 | Fibroblast |
| Cyba | 0.40 | 0.00 | 0.00 | 0.61 | 0.30 | 26 | Fibroblast |
| Emg1 | 0.46 | 0.00 | 0.00 | 0.39 | 0.16 | 26 | Fibroblast |
| Chchd10 | 0.45 | 0.00 | 0.00 | 0.36 | 0.13 | 26 | Fibroblast |
| 5830418K08Rik | 0.40 | 0.00 | 0.00 | 0.27 | 0.09 | 26 | Fibroblast |
| Smc1a | 0.51 | 0.00 | 0.00 | 0.66 | 0.35 | 26 | Fibroblast |
| Fnbp1 | 0.32 | 0.00 | 0.00 | 0.27 | 0.09 | 26 | Fibroblast |
| Utp3 | 0.51 | 0.00 | 0.00 | 0.49 | 0.22 | 26 | Fibroblast |
| Rps6 | 0.36 | 0.00 | 0.00 | 0.96 | 0.82 | 26 | Fibroblast |
| Fkbp3 | 0.55 | 0.00 | 0.00 | 0.59 | 0.30 | 26 | Fibroblast |
| Naa50 | 0.50 | 0.00 | 0.00 | 0.34 | 0.13 | 26 | Fibroblast |
| Hspa4 | 0.52 | 0.00 | 0.00 | 0.51 | 0.23 | 26 | Fibroblast |
| Nhp2 | 0.44 | 0.00 | 0.00 | 0.32 | 0.12 | 26 | Fibroblast |
| Tcf3 | 0.42 | 0.00 | 0.00 | 0.25 | 0.08 | 26 | Fibroblast |
| 9930111J21Rik2 | 0.41 | 0.00 | 0.00 | 0.33 | 0.12 | 26 | Fibroblast |
| Nsmce1 | 0.47 | 0.00 | 0.00 | 0.35 | 0.14 | 26 | Fibroblast |
| Gm9242 | 0.59 | 0.00 | 0.00 | 0.49 | 0.24 | 26 | Fibroblast |
| Ddx27 | 0.45 | 0.00 | 0.00 | 0.33 | 0.12 | 26 | Fibroblast |
| Eml4 | 0.49 | 0.00 | 0.00 | 0.42 | 0.18 | 26 | Fibroblast |
| Gm9843 | 0.31 | 0.00 | 0.00 | 0.99 | 0.93 | 26 | Fibroblast |
| Cytip | 0.39 | 0.00 | 0.00 | 0.29 | 0.10 | 26 | Fibroblast |
| Dhx9 | 0.47 | 0.00 | 0.00 | 0.44 | 0.19 | 26 | Fibroblast |
| Rpl10a | 0.43 | 0.00 | 0.00 | 0.93 | 0.74 | 26 | Fibroblast |
| 2700029M09Rik | 0.46 | 0.00 | 0.00 | 0.34 | 0.13 | 26 | Fibroblast |
| mt-Co3 | 0.36 | 0.00 | 0.00 | 0.98 | 0.89 | 26 | Fibroblast |
| S100a10 | 0.56 | 0.00 | 0.00 | 0.67 | 0.38 | 26 | Fibroblast |
| Mybbp1a | 0.48 | 0.00 | 0.00 | 0.34 | 0.13 | 26 | Fibroblast |
| Gclm | 0.52 | 0.00 | 0.00 | 0.69 | 0.42 | 26 | Fibroblast |
| Rsl1d1 | 0.49 | 0.00 | 0.00 | 0.55 | 0.27 | 26 | Fibroblast |
| Rps18 | 0.44 | 0.00 | 0.00 | 0.87 | 0.65 | 26 | Fibroblast |
| Stk38 | 0.57 | 0.00 | 0.00 | 0.44 | 0.20 | 26 | Fibroblast |
| Lsm5 | 0.52 | 0.00 | 0.00 | 0.31 | 0.12 | 26 | Fibroblast |
| Hist1h2al | 0.38 | 0.00 | 0.00 | 0.29 | 0.10 | 26 | Fibroblast |
| Gspt1 | 0.54 | 0.00 | 0.00 | 0.37 | 0.15 | 26 | Fibroblast |
| Gm10073 | 0.34 | 0.00 | 0.00 | 0.98 | 0.86 | 26 | Fibroblast |
| Gm4070 | 0.42 | 0.00 | 0.00 | 0.33 | 0.12 | 26 | Fibroblast |
| Rpl5 | 0.37 | 0.00 | 0.00 | 0.91 | 0.74 | 26 | Fibroblast |
| Mak16 | 0.46 | 0.00 | 0.00 | 0.32 | 0.12 | 26 | Fibroblast |
| Taf1d | 0.49 | 0.00 | 0.00 | 0.48 | 0.23 | 26 | Fibroblast |
| Rps20 | 0.39 | 0.00 | 0.00 | 0.95 | 0.79 | 26 | Fibroblast |
| Rpl22 | 0.39 | 0.00 | 0.00 | 0.92 | 0.79 | 26 | Fibroblast |
| Wdr43 | 0.42 | 0.00 | 0.00 | 0.34 | 0.13 | 26 | Fibroblast |
| mt-Co2 | 0.37 | 0.00 | 0.00 | 0.95 | 0.77 | 26 | Fibroblast |
| Tpm3-rs7 | 0.44 | 0.00 | 0.00 | 0.90 | 0.69 | 26 | Fibroblast |
| Rad50 | 0.45 | 0.00 | 0.00 | 0.44 | 0.19 | 26 | Fibroblast |
| Banf1 | 0.46 | 0.00 | 0.00 | 0.56 | 0.29 | 26 | Fibroblast |
| Gm2000 | 0.33 | 0.00 | 0.00 | 0.97 | 0.84 | 26 | Fibroblast |
| Orc5 | 0.44 | 0.00 | 0.00 | 0.83 | 0.55 | 26 | Fibroblast |
| Lsm2 | 0.45 | 0.00 | 0.00 | 0.26 | 0.09 | 26 | Fibroblast |
| Trnt1 | 0.45 | 0.00 | 0.00 | 0.32 | 0.12 | 26 | Fibroblast |
| Rps12 | 0.40 | 0.00 | 0.00 | 0.90 | 0.69 | 26 | Fibroblast |
| Grb2 | 0.43 | 0.00 | 0.00 | 0.41 | 0.18 | 26 | Fibroblast |
| Hnrnpd | 0.51 | 0.00 | 0.00 | 0.55 | 0.29 | 26 | Fibroblast |
| Pbdc1 | 0.46 | 0.00 | 0.00 | 0.33 | 0.13 | 26 | Fibroblast |
| Ndufs8 | 0.42 | 0.00 | 0.00 | 0.40 | 0.17 | 26 | Fibroblast |
| Vbp1 | 0.40 | 0.00 | 0.00 | 0.37 | 0.15 | 26 | Fibroblast |
| Ccm2 | 0.35 | 0.00 | 0.00 | 0.31 | 0.12 | 26 | Fibroblast |
| Plekha2 | 0.40 | 0.00 | 0.00 | 0.28 | 0.10 | 26 | Fibroblast |
| Lyn | 0.31 | 0.00 | 0.00 | 0.34 | 0.13 | 26 | Fibroblast |
| Smarca4 | 0.44 | 0.00 | 0.00 | 0.40 | 0.17 | 26 | Fibroblast |
| Cpsf2 | 0.47 | 0.00 | 0.00 | 0.32 | 0.13 | 26 | Fibroblast |
| Pgls | 0.43 | 0.00 | 0.00 | 0.39 | 0.17 | 26 | Fibroblast |
| Mrpl18 | 0.49 | 0.00 | 0.00 | 0.44 | 0.21 | 26 | Fibroblast |
| Npm3 | 0.34 | 0.00 | 0.00 | 0.27 | 0.10 | 26 | Fibroblast |
| Parp2 | 0.35 | 0.00 | 0.00 | 0.29 | 0.10 | 26 | Fibroblast |
| Naa10 | 0.30 | 0.00 | 0.00 | 0.26 | 0.09 | 26 | Fibroblast |
| Rps18-ps3 | 0.45 | 0.00 | 0.00 | 0.79 | 0.56 | 26 | Fibroblast |
| Rps10-ps1 | 0.35 | 0.00 | 0.00 | 0.93 | 0.77 | 26 | Fibroblast |
| Nudc | 0.42 | 0.00 | 0.00 | 0.44 | 0.21 | 26 | Fibroblast |
| Rpl18a | 0.32 | 0.00 | 0.00 | 0.98 | 0.82 | 26 | Fibroblast |
| Hspa14 | 0.40 | 0.00 | 0.00 | 0.29 | 0.11 | 26 | Fibroblast |
| Rpl6 | 0.31 | 0.00 | 0.00 | 0.97 | 0.87 | 26 | Fibroblast |
| Rbm17 | 0.47 | 0.00 | 0.00 | 0.49 | 0.24 | 26 | Fibroblast |
| Gm9396 | 0.40 | 0.00 | 0.00 | 0.79 | 0.54 | 26 | Fibroblast |
| Gvin1 | 0.35 | 0.00 | 0.00 | 0.31 | 0.12 | 26 | Fibroblast |
| Dnajc7 | 0.40 | 0.00 | 0.00 | 0.46 | 0.21 | 26 | Fibroblast |
| Rpl22l1 | 0.44 | 0.00 | 0.00 | 0.80 | 0.56 | 26 | Fibroblast |
| Rplp2 | 0.33 | 0.00 | 0.00 | 0.94 | 0.81 | 26 | Fibroblast |
| Rpl18 | 0.40 | 0.00 | 0.00 | 0.85 | 0.64 | 26 | Fibroblast |
| Gadd45gip1 | 0.41 | 0.00 | 0.00 | 0.36 | 0.16 | 26 | Fibroblast |
| Mrpl40 | 0.45 | 0.00 | 0.00 | 0.29 | 0.11 | 26 | Fibroblast |
| Clic1 | 0.42 | 0.00 | 0.00 | 0.66 | 0.38 | 26 | Fibroblast |
| Cenpc1 | 0.37 | 0.00 | 0.00 | 0.25 | 0.09 | 26 | Fibroblast |
| Mtf2 | 0.41 | 0.00 | 0.00 | 0.40 | 0.18 | 26 | Fibroblast |
| Iws1 | 0.37 | 0.00 | 0.00 | 0.38 | 0.16 | 26 | Fibroblast |
| C1qbp | 0.45 | 0.00 | 0.00 | 0.31 | 0.12 | 26 | Fibroblast |
| Eif5 | 0.45 | 0.00 | 0.00 | 0.63 | 0.37 | 26 | Fibroblast |
| Ccnd2 | 0.47 | 0.00 | 0.00 | 0.54 | 0.28 | 26 | Fibroblast |
| Fau | 0.29 | 0.00 | 0.00 | 0.98 | 0.87 | 26 | Fibroblast |
| Rpl3 | 0.41 | 0.00 | 0.00 | 0.84 | 0.63 | 26 | Fibroblast |
| Dtymk | 0.46 | 0.00 | 0.00 | 0.33 | 0.14 | 26 | Fibroblast |
| Srsf2 | 0.49 | 0.00 | 0.00 | 0.65 | 0.38 | 26 | Fibroblast |
| Slmo2 | 0.36 | 0.00 | 0.00 | 0.26 | 0.09 | 26 | Fibroblast |
| Rpa3 | 0.49 | 0.00 | 0.00 | 0.31 | 0.13 | 26 | Fibroblast |
| Pgk1-rs7 | 0.31 | 0.00 | 0.00 | 0.27 | 0.10 | 26 | Fibroblast |
| Hprt | 0.36 | 0.00 | 0.00 | 0.33 | 0.14 | 26 | Fibroblast |
| Pgk1 | 0.32 | 0.00 | 0.00 | 0.30 | 0.12 | 26 | Fibroblast |
| Ctsz | 0.44 | 0.00 | 0.00 | 0.36 | 0.15 | 26 | Fibroblast |
| Rps15a | 0.30 | 0.00 | 0.00 | 0.95 | 0.81 | 26 | Fibroblast |
| Lsm6 | 0.44 | 0.00 | 0.00 | 0.33 | 0.13 | 26 | Fibroblast |
| Mdh1 | 0.36 | 0.00 | 0.00 | 0.41 | 0.19 | 26 | Fibroblast |
| Hnrnpc | 0.40 | 0.00 | 0.00 | 0.72 | 0.44 | 26 | Fibroblast |
| Rps13 | 0.37 | 0.00 | 0.00 | 0.84 | 0.64 | 26 | Fibroblast |
| Ddx21 | 0.46 | 0.00 | 0.00 | 0.51 | 0.26 | 26 | Fibroblast |
| Il2rg | 0.30 | 0.00 | 0.00 | 0.27 | 0.10 | 26 | Fibroblast |
| Ppp1cc | 0.33 | 0.00 | 0.00 | 0.27 | 0.10 | 26 | Fibroblast |
| Bola2 | 0.38 | 0.00 | 0.00 | 0.37 | 0.16 | 26 | Fibroblast |
| Slc25a5 | 0.38 | 0.00 | 0.00 | 0.82 | 0.59 | 26 | Fibroblast |
| Smc3 | 0.39 | 0.00 | 0.00 | 0.66 | 0.38 | 26 | Fibroblast |
| Snrpg | 0.45 | 0.00 | 0.00 | 0.64 | 0.39 | 26 | Fibroblast |
| Gm10068 | 0.38 | 0.00 | 0.00 | 0.57 | 0.30 | 26 | Fibroblast |
| Hnrnpa1 | 0.44 | 0.00 | 0.00 | 0.51 | 0.27 | 26 | Fibroblast |
| Lman2 | 0.40 | 0.00 | 0.00 | 0.33 | 0.14 | 26 | Fibroblast |
| Gpx1 | 0.36 | 0.00 | 0.00 | 0.72 | 0.43 | 26 | Fibroblast |
| Gdi2 | 0.40 | 0.00 | 0.00 | 0.59 | 0.33 | 26 | Fibroblast |
| Xist | 0.44 | 0.00 | 0.00 | 0.77 | 0.51 | 26 | Fibroblast |
| Trp53 | 0.31 | 0.00 | 0.00 | 0.37 | 0.16 | 26 | Fibroblast |
| Lsm3 | 0.31 | 0.00 | 0.00 | 0.28 | 0.11 | 26 | Fibroblast |
| Setd8 | 0.36 | 0.00 | 0.00 | 0.29 | 0.12 | 26 | Fibroblast |
| Bin1 | 0.43 | 0.00 | 0.00 | 0.28 | 0.11 | 26 | Fibroblast |
| Lyar | 0.46 | 0.00 | 0.00 | 0.35 | 0.16 | 26 | Fibroblast |
| Gm5506 | 0.35 | 0.00 | 0.00 | 0.34 | 0.14 | 26 | Fibroblast |
| Gabpb2 | 0.41 | 0.00 | 0.00 | 0.26 | 0.10 | 26 | Fibroblast |
| Stk4 | 0.33 | 0.00 | 0.00 | 0.34 | 0.14 | 26 | Fibroblast |
| Shisa5 | 0.32 | 0.00 | 0.00 | 0.43 | 0.21 | 26 | Fibroblast |
| Naa15 | 0.36 | 0.00 | 0.00 | 0.45 | 0.22 | 26 | Fibroblast |
| Hspd1 | 0.49 | 0.00 | 0.00 | 0.47 | 0.25 | 26 | Fibroblast |
| Mrp63 | 0.37 | 0.00 | 0.00 | 0.34 | 0.15 | 26 | Fibroblast |
| Bub3 | 0.43 | 0.00 | 0.00 | 0.32 | 0.14 | 26 | Fibroblast |
| Suz12 | 0.45 | 0.00 | 0.00 | 0.31 | 0.13 | 26 | Fibroblast |
| Fh1 | 0.32 | 0.00 | 0.00 | 0.29 | 0.12 | 26 | Fibroblast |
| Smc6 | 0.36 | 0.00 | 0.00 | 0.59 | 0.33 | 26 | Fibroblast |
| Gm6169 | 0.43 | 0.00 | 0.00 | 0.30 | 0.13 | 26 | Fibroblast |
| Cbx3 | 0.39 | 0.00 | 0.00 | 0.61 | 0.34 | 26 | Fibroblast |
| Eif2s1 | 0.35 | 0.00 | 0.00 | 0.31 | 0.13 | 26 | Fibroblast |
| Taf15 | 0.34 | 0.00 | 0.00 | 0.26 | 0.10 | 26 | Fibroblast |
| Pdia4 | 0.48 | 0.00 | 0.00 | 0.42 | 0.21 | 26 | Fibroblast |
| Itsn2 | 0.39 | 0.00 | 0.00 | 0.48 | 0.25 | 26 | Fibroblast |
| Prpf38b | 0.39 | 0.00 | 0.00 | 0.77 | 0.50 | 26 | Fibroblast |
| Larp7 | 0.31 | 0.00 | 0.00 | 0.38 | 0.17 | 26 | Fibroblast |
| Magoh | 0.36 | 0.00 | 0.00 | 0.36 | 0.16 | 26 | Fibroblast |
| Hnrnpab | 0.42 | 0.00 | 0.00 | 0.57 | 0.33 | 26 | Fibroblast |
| Llph | 0.41 | 0.00 | 0.00 | 0.59 | 0.34 | 26 | Fibroblast |
| Srsf7 | 0.34 | 0.00 | 0.00 | 0.35 | 0.16 | 26 | Fibroblast |
| Nhp2l1 | 0.36 | 0.00 | 0.00 | 0.39 | 0.19 | 26 | Fibroblast |
| Rps19bp1 | 0.35 | 0.00 | 0.00 | 0.28 | 0.12 | 26 | Fibroblast |
| Hint1 | 0.42 | 0.00 | 0.00 | 0.71 | 0.46 | 26 | Fibroblast |
| Polr1d | 0.36 | 0.00 | 0.00 | 0.52 | 0.28 | 26 | Fibroblast |
| Rab8a | 0.35 | 0.00 | 0.00 | 0.31 | 0.14 | 26 | Fibroblast |
| Rpl12 | 0.34 | 0.00 | 0.00 | 0.77 | 0.55 | 26 | Fibroblast |
| Srsf4 | 0.38 | 0.00 | 0.00 | 0.30 | 0.13 | 26 | Fibroblast |
| Gbp7 | 0.36 | 0.00 | 0.00 | 0.32 | 0.14 | 26 | Fibroblast |
| Rpl35 | 0.29 | 0.00 | 0.00 | 0.94 | 0.80 | 26 | Fibroblast |
| Sf3a3 | 0.35 | 0.00 | 0.00 | 0.28 | 0.12 | 26 | Fibroblast |
| mt-Nd1 | 0.27 | 0.00 | 0.00 | 0.98 | 0.91 | 26 | Fibroblast |
| Nono | 0.35 | 0.00 | 0.00 | 0.42 | 0.21 | 26 | Fibroblast |
| Taf9 | 0.34 | 0.00 | 0.00 | 0.38 | 0.18 | 26 | Fibroblast |
| mt-Nd5 | 0.38 | 0.00 | 0.00 | 0.92 | 0.76 | 26 | Fibroblast |
| Arhgef1 | 0.34 | 0.00 | 0.00 | 0.28 | 0.12 | 26 | Fibroblast |
| Gimap6 | 0.29 | 0.00 | 0.00 | 0.36 | 0.16 | 26 | Fibroblast |
| Tmem123 | 0.32 | 0.00 | 0.00 | 0.29 | 0.12 | 26 | Fibroblast |
| Pdlim1 | 0.45 | 0.00 | 0.00 | 0.31 | 0.14 | 26 | Fibroblast |
| Eno1 | 0.40 | 0.00 | 0.00 | 0.34 | 0.16 | 26 | Fibroblast |
| Gm10222 | 0.40 | 0.00 | 0.00 | 0.67 | 0.43 | 26 | Fibroblast |
| Hsp90ab1 | 0.31 | 0.00 | 0.00 | 0.98 | 0.89 | 26 | Fibroblast |
| AC121131.2 | 0.37 | 0.00 | 0.00 | 0.64 | 0.40 | 26 | Fibroblast |
| Ucp2 | 0.33 | 0.00 | 0.00 | 0.29 | 0.12 | 26 | Fibroblast |
| Tomm70a | 0.30 | 0.00 | 0.00 | 0.30 | 0.13 | 26 | Fibroblast |
| D19Ertd737e | 0.35 | 0.00 | 0.00 | 0.34 | 0.16 | 26 | Fibroblast |
| 1810009A15Rik | 0.30 | 0.00 | 0.00 | 0.31 | 0.14 | 26 | Fibroblast |
| mt-Nd4l | 0.38 | 0.00 | 0.00 | 0.62 | 0.38 | 26 | Fibroblast |
| mt-Atp6 | 0.29 | 0.00 | 0.00 | 0.93 | 0.85 | 26 | Fibroblast |
| Cib1 | 0.35 | 0.00 | 0.00 | 0.28 | 0.12 | 26 | Fibroblast |
| Rpl13 | 0.29 | 0.00 | 0.00 | 0.92 | 0.79 | 26 | Fibroblast |
| Eef1d | 0.40 | 0.00 | 0.00 | 0.47 | 0.26 | 26 | Fibroblast |
| Rps19 | 0.25 | 0.00 | 0.00 | 0.94 | 0.83 | 26 | Fibroblast |
| Ywhah | 0.37 | 0.00 | 0.00 | 0.48 | 0.26 | 26 | Fibroblast |
| Hnrnpm | 0.35 | 0.00 | 0.00 | 0.72 | 0.49 | 26 | Fibroblast |
| Gm8225 | 0.36 | 0.00 | 0.00 | 0.32 | 0.15 | 26 | Fibroblast |
| Ppp4c | 0.37 | 0.00 | 0.00 | 0.33 | 0.16 | 26 | Fibroblast |
| Psmb1 | 0.36 | 0.00 | 0.00 | 0.71 | 0.47 | 26 | Fibroblast |
| Adss | 0.29 | 0.00 | 0.00 | 0.35 | 0.17 | 26 | Fibroblast |
| Sae1 | 0.30 | 0.00 | 0.00 | 0.26 | 0.11 | 26 | Fibroblast |
| Phip | 0.34 | 0.00 | 0.00 | 0.34 | 0.16 | 26 | Fibroblast |
| Gm5160 | 0.39 | 0.00 | 0.00 | 0.41 | 0.22 | 26 | Fibroblast |
| Stip1 | 0.38 | 0.00 | 0.00 | 0.29 | 0.13 | 26 | Fibroblast |
| Gm5428 | 0.25 | 0.00 | 0.00 | 0.94 | 0.79 | 26 | Fibroblast |
| H3f3a | 0.39 | 0.00 | 0.00 | 0.80 | 0.60 | 26 | Fibroblast |
| Sf1 | 0.31 | 0.00 | 0.00 | 0.28 | 0.12 | 26 | Fibroblast |
| Syncrip | 0.40 | 0.00 | 0.00 | 0.45 | 0.25 | 26 | Fibroblast |
| Rfc1 | 0.33 | 0.00 | 0.00 | 0.40 | 0.21 | 26 | Fibroblast |
| Rpl39 | 0.26 | 0.00 | 0.00 | 0.94 | 0.80 | 26 | Fibroblast |
| Prelid1 | 0.31 | 0.00 | 0.00 | 0.52 | 0.29 | 26 | Fibroblast |
| Cdc42se2 | 0.36 | 0.00 | 0.00 | 0.30 | 0.14 | 26 | Fibroblast |
| Nfyb | 0.25 | 0.00 | 0.00 | 0.27 | 0.12 | 26 | Fibroblast |
| Cacybp | 0.35 | 0.00 | 0.00 | 0.40 | 0.21 | 26 | Fibroblast |
| Gm6472 | 0.25 | 0.00 | 0.00 | 0.93 | 0.80 | 26 | Fibroblast |
| Rps25 | 0.28 | 0.00 | 0.00 | 0.93 | 0.80 | 26 | Fibroblast |
| Mrpl54 | 0.30 | 0.00 | 0.00 | 0.31 | 0.14 | 26 | Fibroblast |
| Hspe1 | 0.27 | 0.00 | 0.00 | 0.51 | 0.29 | 26 | Fibroblast |
| Gapdh | 0.35 | 0.00 | 0.00 | 0.30 | 0.14 | 26 | Fibroblast |
| Zcchc11 | 0.34 | 0.00 | 0.00 | 0.40 | 0.21 | 26 | Fibroblast |
| Psma1 | 0.32 | 0.00 | 0.00 | 0.47 | 0.26 | 26 | Fibroblast |
| Srrt | 0.32 | 0.00 | 0.00 | 0.39 | 0.20 | 26 | Fibroblast |
| Plp2 | 0.38 | 0.00 | 0.00 | 0.29 | 0.13 | 26 | Fibroblast |
| 11-Sep | 0.30 | 0.00 | 0.00 | 0.43 | 0.23 | 26 | Fibroblast |
| Trmt112 | 0.25 | 0.00 | 0.00 | 0.31 | 0.14 | 26 | Fibroblast |
| Caprin1 | 0.39 | 0.00 | 0.00 | 0.44 | 0.25 | 26 | Fibroblast |
| Cct3 | 0.31 | 0.00 | 0.00 | 0.34 | 0.17 | 26 | Fibroblast |
| Thrap3 | 0.37 | 0.00 | 0.00 | 0.59 | 0.37 | 26 | Fibroblast |
| Mrpl15 | 0.28 | 0.00 | 0.00 | 0.30 | 0.14 | 26 | Fibroblast |
| Sf3b5 | 0.35 | 0.00 | 0.00 | 0.41 | 0.22 | 26 | Fibroblast |
| Rpl28 | 0.29 | 0.00 | 0.00 | 0.76 | 0.53 | 26 | Fibroblast |
| Tsix | 0.27 | 0.00 | 0.00 | 0.36 | 0.18 | 26 | Fibroblast |
| Gm9825 | 0.28 | 0.00 | 0.00 | 0.28 | 0.13 | 26 | Fibroblast |
| Mtpn | 0.31 | 0.00 | 0.00 | 0.48 | 0.27 | 26 | Fibroblast |
| Nop10 | 0.32 | 0.00 | 0.00 | 0.48 | 0.28 | 26 | Fibroblast |
| Tmed5 | 0.35 | 0.00 | 0.00 | 0.34 | 0.17 | 26 | Fibroblast |
| Mrpl21 | 0.28 | 0.00 | 0.00 | 0.31 | 0.15 | 26 | Fibroblast |
| Nek7 | 0.36 | 0.00 | 0.00 | 0.26 | 0.12 | 26 | Fibroblast |
| Srsf6 | 0.25 | 0.00 | 0.00 | 0.37 | 0.19 | 26 | Fibroblast |
| Rpl13a-ps1 | 0.34 | 0.00 | 0.00 | 0.75 | 0.55 | 26 | Fibroblast |
| Cct8 | 0.31 | 0.00 | 0.00 | 0.39 | 0.21 | 26 | Fibroblast |
| Rps10 | 0.28 | 0.00 | 0.00 | 0.87 | 0.70 | 26 | Fibroblast |
| Calm3 | 0.29 | 0.00 | 0.00 | 0.37 | 0.20 | 26 | Fibroblast |
| Tpr | 0.32 | 0.00 | 0.00 | 0.71 | 0.50 | 26 | Fibroblast |
| Tcp1 | 0.31 | 0.00 | 0.00 | 0.43 | 0.24 | 26 | Fibroblast |
| AC167036.1 | 0.33 | 0.00 | 0.00 | 0.51 | 0.30 | 26 | Fibroblast |
| Tra2b | 0.29 | 0.00 | 0.00 | 0.52 | 0.29 | 26 | Fibroblast |
| Ptpn1 | 0.29 | 0.00 | 0.00 | 0.39 | 0.20 | 26 | Fibroblast |
| Ppm1g | 0.30 | 0.00 | 0.00 | 0.29 | 0.13 | 26 | Fibroblast |
| Lrrfip1 | 0.26 | 0.00 | 0.00 | 0.46 | 0.26 | 26 | Fibroblast |
| Twistnb | 0.29 | 0.00 | 0.00 | 0.29 | 0.13 | 26 | Fibroblast |
| Gm18025 | 0.32 | 0.00 | 0.00 | 0.38 | 0.20 | 26 | Fibroblast |
| Ccdc41 | 0.28 | 0.00 | 0.00 | 0.34 | 0.17 | 26 | Fibroblast |
| Smarcc1 | 0.33 | 0.00 | 0.00 | 0.33 | 0.17 | 26 | Fibroblast |
| Baz1b | 0.31 | 0.00 | 0.00 | 0.47 | 0.27 | 26 | Fibroblast |
| AC123611.1 | 0.33 | 0.00 | 0.00 | 0.61 | 0.39 | 26 | Fibroblast |
| Rpl36al | 0.34 | 0.00 | 0.00 | 0.71 | 0.52 | 26 | Fibroblast |
| Cbx1 | 0.32 | 0.00 | 0.00 | 0.64 | 0.42 | 26 | Fibroblast |
| U2surp | 0.29 | 0.00 | 0.00 | 0.54 | 0.33 | 26 | Fibroblast |
| Arl6ip4 | 0.32 | 0.00 | 0.00 | 0.36 | 0.19 | 26 | Fibroblast |
| Cox5a | 0.26 | 0.00 | 0.00 | 0.53 | 0.31 | 26 | Fibroblast |
| Kmt2a | 0.30 | 0.00 | 0.00 | 0.39 | 0.21 | 26 | Fibroblast |
| Mapk1 | 0.26 | 0.00 | 0.00 | 0.34 | 0.17 | 26 | Fibroblast |
| Sf3b2 | 0.27 | 0.00 | 0.00 | 0.62 | 0.40 | 26 | Fibroblast |
| Tpi1 | 0.26 | 0.00 | 0.00 | 0.31 | 0.15 | 26 | Fibroblast |
| Snrpd3 | 0.33 | 0.00 | 0.00 | 0.52 | 0.32 | 26 | Fibroblast |
| Tcerg1 | 0.28 | 0.00 | 0.00 | 0.37 | 0.20 | 26 | Fibroblast |
| Rnps1 | 0.32 | 0.00 | 0.00 | 0.30 | 0.15 | 26 | Fibroblast |
| Ppp1ca | 0.29 | 0.00 | 0.00 | 0.54 | 0.33 | 26 | Fibroblast |
| Snhg5 | 0.29 | 0.00 | 0.00 | 0.34 | 0.18 | 26 | Fibroblast |
| H3f3b | 0.26 | 0.00 | 0.00 | 0.96 | 0.89 | 26 | Fibroblast |
| Ube2a | 0.42 | 0.00 | 0.00 | 0.34 | 0.18 | 26 | Fibroblast |
| Atp5a1 | 0.30 | 0.00 | 0.00 | 0.61 | 0.41 | 26 | Fibroblast |
| Mat2a | 0.27 | 0.00 | 0.00 | 0.46 | 0.27 | 26 | Fibroblast |
| Rps17 | 0.25 | 0.00 | 0.00 | 0.90 | 0.77 | 26 | Fibroblast |
| Gm11273 | 0.26 | 0.00 | 0.00 | 0.60 | 0.38 | 26 | Fibroblast |
| Gm6563 | 0.31 | 0.00 | 0.00 | 0.53 | 0.34 | 26 | Fibroblast |
| 2310036O22Rik | 0.29 | 0.00 | 0.00 | 0.36 | 0.19 | 26 | Fibroblast |
| Pqbp1 | 0.25 | 0.00 | 0.00 | 0.26 | 0.12 | 26 | Fibroblast |
| Sumo2 | 0.30 | 0.00 | 0.00 | 0.49 | 0.30 | 26 | Fibroblast |
| Slc25a3 | 0.27 | 0.00 | 0.00 | 0.64 | 0.42 | 26 | Fibroblast |
| S100a6 | 0.47 | 0.00 | 0.00 | 0.49 | 0.31 | 26 | Fibroblast |
| Tardbp | 0.28 | 0.00 | 0.00 | 0.51 | 0.31 | 26 | Fibroblast |
| Stx7 | 0.33 | 0.00 | 0.00 | 0.29 | 0.15 | 26 | Fibroblast |
| Ndufb7 | 0.27 | 0.00 | 0.00 | 0.47 | 0.28 | 26 | Fibroblast |
| Stag2 | 0.32 | 0.00 | 0.00 | 0.40 | 0.23 | 26 | Fibroblast |
| Eif4a3 | 0.33 | 0.00 | 0.00 | 0.35 | 0.19 | 26 | Fibroblast |
| Cct7 | 0.30 | 0.00 | 0.00 | 0.38 | 0.21 | 26 | Fibroblast |
| Psme1 | 0.25 | 0.00 | 0.00 | 0.52 | 0.32 | 26 | Fibroblast |
| Mcl1 | 0.25 | 0.00 | 0.00 | 0.40 | 0.23 | 26 | Fibroblast |
| Hnrnpa3 | 0.32 | 0.00 | 0.00 | 0.64 | 0.46 | 26 | Fibroblast |
| Sarnp | 0.25 | 0.00 | 0.00 | 0.43 | 0.25 | 26 | Fibroblast |
| Erh | 0.31 | 0.00 | 0.00 | 0.46 | 0.29 | 26 | Fibroblast |
| Mrpl52 | 0.25 | 0.00 | 0.00 | 0.52 | 0.33 | 26 | Fibroblast |
| Ppdpf | 0.25 | 0.00 | 0.00 | 0.33 | 0.18 | 26 | Fibroblast |
| Cox20 | 0.27 | 0.00 | 0.00 | 0.33 | 0.18 | 26 | Fibroblast |
| Mtdh | 0.28 | 0.00 | 0.00 | 0.67 | 0.48 | 26 | Fibroblast |
| Snrpd2 | 0.29 | 0.00 | 0.00 | 0.41 | 0.25 | 26 | Fibroblast |
| Dnajb11 | 0.26 | 0.00 | 0.00 | 0.29 | 0.16 | 26 | Fibroblast |
| Oaz1 | 0.26 | 0.00 | 0.00 | 0.80 | 0.62 | 26 | Fibroblast |
| Ldha | 0.25 | 0.00 | 0.00 | 0.47 | 0.29 | 26 | Fibroblast |
| Atp5b | 0.27 | 0.00 | 0.00 | 0.63 | 0.44 | 26 | Fibroblast |
| Kpnb1 | 0.28 | 0.00 | 0.00 | 0.33 | 0.19 | 26 | Fibroblast |
| Gm12728 | 0.29 | 0.00 | 0.00 | 0.31 | 0.17 | 26 | Fibroblast |
| Rps2-ps6 | 0.26 | 0.00 | 0.00 | 0.58 | 0.41 | 26 | Fibroblast |
| Eif2s2 | 0.27 | 0.00 | 0.01 | 0.64 | 0.48 | 26 | Fibroblast |
| Gm6139 | 0.27 | 0.00 | 0.02 | 0.59 | 0.42 | 26 | Fibroblast |
| Manf | 0.31 | 0.00 | 0.02 | 0.33 | 0.20 | 26 | Fibroblast |
| Wbp11 | 0.28 | 0.00 | 0.02 | 0.39 | 0.24 | 26 | Fibroblast |
| Casp8ap2 | 0.29 | 0.00 | 0.03 | 0.30 | 0.17 | 26 | Fibroblast |
| Cct6a | 0.26 | 0.00 | 0.03 | 0.38 | 0.24 | 26 | Fibroblast |
| Tagln2 | 0.33 | 0.00 | 0.03 | 0.62 | 0.45 | 26 | Fibroblast |
| Scgb3a2 | 4.97 | 0.00 | 0.00 | 0.96 | 0.07 | 27 | Mixed Epithelial |
| Cyp2f2 | 3.52 | 0.00 | 0.00 | 0.95 | 0.02 | 27 | Mixed Epithelial |
| Cbr2 | 3.16 | 0.00 | 0.00 | 0.99 | 0.16 | 27 | Mixed Epithelial |
| Hp | 3.02 | 0.00 | 0.00 | 0.89 | 0.11 | 27 | Mixed Epithelial |
| Aldh1a1 | 2.56 | 0.00 | 0.00 | 0.88 | 0.04 | 27 | Mixed Epithelial |
| Aldh1a7 | 2.06 | 0.00 | 0.00 | 0.80 | 0.04 | 27 | Mixed Epithelial |
| Gsto1 | 2.01 | 0.00 | 0.00 | 0.66 | 0.06 | 27 | Mixed Epithelial |
| Gsta4 | 1.97 | 0.00 | 0.00 | 0.83 | 0.04 | 27 | Mixed Epithelial |
| Sec14l3 | 1.86 | 0.00 | 0.00 | 0.91 | 0.11 | 27 | Mixed Epithelial |
| B430010I23Rik | 1.83 | 0.00 | 0.00 | 0.50 | 0.02 | 27 | Mixed Epithelial |
| Lypd2 | 1.78 | 0.00 | 0.00 | 0.51 | 0.01 | 27 | Mixed Epithelial |
| Cldn10 | 1.76 | 0.00 | 0.00 | 0.71 | 0.02 | 27 | Mixed Epithelial |
| Krt15 | 1.39 | 0.00 | 0.00 | 0.27 | 0.00 | 27 | Mixed Epithelial |
| Gsta3 | 1.37 | 0.00 | 0.00 | 0.60 | 0.01 | 27 | Mixed Epithelial |
| Aox3 | 1.09 | 0.00 | 0.00 | 0.41 | 0.01 | 27 | Mixed Epithelial |
| 5330417C22Rik | 0.89 | 0.00 | 0.00 | 0.44 | 0.02 | 27 | Mixed Epithelial |
| Slc16a11 | 0.82 | 0.00 | 0.00 | 0.34 | 0.00 | 27 | Mixed Epithelial |
| Ace2 | 0.81 | 0.00 | 0.00 | 0.36 | 0.01 | 27 | Mixed Epithelial |
| Cckar | 0.80 | 0.00 | 0.00 | 0.36 | 0.01 | 27 | Mixed Epithelial |
| Gabrp | 0.73 | 0.00 | 0.00 | 0.32 | 0.00 | 27 | Mixed Epithelial |
| Gm15883 | 0.69 | 0.00 | 0.00 | 0.32 | 0.00 | 27 | Mixed Epithelial |
| Gpx2 | 0.67 | 0.00 | 0.00 | 0.29 | 0.01 | 27 | Mixed Epithelial |
| Sox2 | 0.62 | 0.00 | 0.00 | 0.27 | 0.01 | 27 | Mixed Epithelial |
| Lrrc26 | 0.62 | 0.00 | 0.00 | 0.27 | 0.00 | 27 | Mixed Epithelial |
| AU021092 | 1.15 | 0.00 | 0.00 | 0.56 | 0.04 | 27 | Mixed Epithelial |
| Homer2 | 0.65 | 0.00 | 0.00 | 0.29 | 0.01 | 27 | Mixed Epithelial |
| Krt19 | 1.24 | 0.00 | 0.00 | 0.66 | 0.06 | 27 | Mixed Epithelial |
| Wfdc2 | 2.46 | 0.00 | 0.00 | 0.95 | 0.16 | 27 | Mixed Epithelial |
| Ehf | 0.84 | 0.00 | 0.00 | 0.34 | 0.02 | 27 | Mixed Epithelial |
| Tspan1 | 1.01 | 0.00 | 0.00 | 0.51 | 0.04 | 27 | Mixed Epithelial |
| Mgst1 | 2.43 | 0.00 | 0.00 | 0.93 | 0.19 | 27 | Mixed Epithelial |
| Cgn | 0.87 | 0.00 | 0.00 | 0.48 | 0.04 | 27 | Mixed Epithelial |
| Ces1d | 1.84 | 0.00 | 0.00 | 0.90 | 0.17 | 27 | Mixed Epithelial |
| Chia | 1.83 | 0.00 | 0.00 | 0.35 | 0.02 | 27 | Mixed Epithelial |
| Ldhb | 1.22 | 0.00 | 0.00 | 0.61 | 0.08 | 27 | Mixed Epithelial |
| Scgb1a1 | 5.32 | 0.00 | 0.00 | 0.99 | 0.35 | 27 | Mixed Epithelial |
| Acsl1 | 0.77 | 0.00 | 0.00 | 0.36 | 0.03 | 27 | Mixed Epithelial |
| Sftpd | 1.34 | 0.00 | 0.00 | 0.90 | 0.16 | 27 | Mixed Epithelial |
| Kcnk2 | 0.61 | 0.00 | 0.00 | 0.25 | 0.01 | 27 | Mixed Epithelial |
| Por | 1.44 | 0.00 | 0.00 | 0.74 | 0.12 | 27 | Mixed Epithelial |
| Atp1b1 | 1.50 | 0.00 | 0.00 | 0.90 | 0.18 | 27 | Mixed Epithelial |
| Cldn7 | 1.04 | 0.00 | 0.00 | 0.60 | 0.08 | 27 | Mixed Epithelial |
| Dcxr | 1.17 | 0.00 | 0.00 | 0.69 | 0.11 | 27 | Mixed Epithelial |
| Gstt3 | 0.68 | 0.00 | 0.00 | 0.32 | 0.03 | 27 | Mixed Epithelial |
| Muc1 | 0.97 | 0.00 | 0.00 | 0.63 | 0.09 | 27 | Mixed Epithelial |
| Fam46c | 0.63 | 0.00 | 0.00 | 0.30 | 0.02 | 27 | Mixed Epithelial |
| Selenbp2 | 1.64 | 0.00 | 0.00 | 0.81 | 0.17 | 27 | Mixed Epithelial |
| Selenbp1 | 1.78 | 0.00 | 0.00 | 0.90 | 0.22 | 27 | Mixed Epithelial |
| Krt18 | 0.99 | 0.00 | 0.00 | 0.57 | 0.08 | 27 | Mixed Epithelial |
| Gm16136 | 0.57 | 0.00 | 0.00 | 0.26 | 0.02 | 27 | Mixed Epithelial |
| Retnla | 3.43 | 0.00 | 0.00 | 0.61 | 0.10 | 27 | Mixed Epithelial |
| Sfta2 | 1.17 | 0.00 | 0.00 | 0.80 | 0.15 | 27 | Mixed Epithelial |
| Dapk2 | 0.55 | 0.00 | 0.00 | 0.27 | 0.02 | 27 | Mixed Epithelial |
| Alas1 | 1.24 | 0.00 | 0.00 | 0.66 | 0.12 | 27 | Mixed Epithelial |
| Kcnj15 | 0.73 | 0.00 | 0.00 | 0.38 | 0.04 | 27 | Mixed Epithelial |
| Cisd1 | 1.06 | 0.00 | 0.00 | 0.64 | 0.11 | 27 | Mixed Epithelial |
| Epcam | 0.97 | 0.00 | 0.00 | 0.60 | 0.10 | 27 | Mixed Epithelial |
| Krt8 | 0.98 | 0.00 | 0.00 | 0.53 | 0.08 | 27 | Mixed Epithelial |
| Gstk1 | 0.91 | 0.00 | 0.00 | 0.51 | 0.08 | 27 | Mixed Epithelial |
| Cldn3 | 0.89 | 0.00 | 0.00 | 0.67 | 0.12 | 27 | Mixed Epithelial |
| Mettl7a1 | 1.35 | 0.00 | 0.00 | 0.83 | 0.22 | 27 | Mixed Epithelial |
| Mettl7a2 | 1.02 | 0.00 | 0.00 | 0.61 | 0.11 | 27 | Mixed Epithelial |
| Rasl11a | 0.92 | 0.00 | 0.00 | 0.45 | 0.06 | 27 | Mixed Epithelial |
| Tacstd2 | 0.70 | 0.00 | 0.00 | 0.31 | 0.03 | 27 | Mixed Epithelial |
| Ptgr1 | 0.92 | 0.00 | 0.00 | 0.41 | 0.05 | 27 | Mixed Epithelial |
| Spint2 | 0.88 | 0.00 | 0.00 | 0.62 | 0.11 | 27 | Mixed Epithelial |
| Nupr1 | 1.26 | 0.00 | 0.00 | 0.76 | 0.18 | 27 | Mixed Epithelial |
| Prdx6 | 1.56 | 0.00 | 0.00 | 0.87 | 0.27 | 27 | Mixed Epithelial |
| Cd55 | 0.80 | 0.00 | 0.00 | 0.48 | 0.07 | 27 | Mixed Epithelial |
| Clic3 | 0.87 | 0.00 | 0.00 | 0.54 | 0.09 | 27 | Mixed Epithelial |
| Scnn1b | 0.49 | 0.00 | 0.00 | 0.27 | 0.03 | 27 | Mixed Epithelial |
| Pir | 0.61 | 0.00 | 0.00 | 0.31 | 0.03 | 27 | Mixed Epithelial |
| Ptpn13 | 0.57 | 0.00 | 0.00 | 0.31 | 0.03 | 27 | Mixed Epithelial |
| Kcnk1 | 0.55 | 0.00 | 0.00 | 0.26 | 0.03 | 27 | Mixed Epithelial |
| Rab25 | 0.49 | 0.00 | 0.00 | 0.25 | 0.02 | 27 | Mixed Epithelial |
| Trf | 0.99 | 0.00 | 0.00 | 0.54 | 0.10 | 27 | Mixed Epithelial |
| Fam174b | 0.59 | 0.00 | 0.00 | 0.38 | 0.05 | 27 | Mixed Epithelial |
| 8430408G22Rik | 0.67 | 0.00 | 0.00 | 0.33 | 0.04 | 27 | Mixed Epithelial |
| Cdh1 | 0.74 | 0.00 | 0.00 | 0.45 | 0.08 | 27 | Mixed Epithelial |
| Pdzk1ip1 | 0.48 | 0.00 | 0.00 | 0.26 | 0.03 | 27 | Mixed Epithelial |
| Mal2 | 0.63 | 0.00 | 0.00 | 0.39 | 0.06 | 27 | Mixed Epithelial |
| Chchd10 | 0.65 | 0.00 | 0.00 | 0.60 | 0.13 | 27 | Mixed Epithelial |
| Chi3l1 | 1.05 | 0.00 | 0.00 | 0.52 | 0.11 | 27 | Mixed Epithelial |
| Prr15l | 0.60 | 0.00 | 0.00 | 0.37 | 0.06 | 27 | Mixed Epithelial |
| Klf5 | 0.53 | 0.00 | 0.00 | 0.26 | 0.03 | 27 | Mixed Epithelial |
| Acss1 | 0.52 | 0.00 | 0.00 | 0.28 | 0.04 | 27 | Mixed Epithelial |
| Il18r1 | 0.61 | 0.00 | 0.00 | 0.40 | 0.07 | 27 | Mixed Epithelial |
| 1810010H24Rik | 0.50 | 0.00 | 0.00 | 0.26 | 0.03 | 27 | Mixed Epithelial |
| Cxadr | 0.54 | 0.00 | 0.00 | 0.36 | 0.06 | 27 | Mixed Epithelial |
| Mal | 0.61 | 0.00 | 0.00 | 0.31 | 0.05 | 27 | Mixed Epithelial |
| Myo5c | 0.68 | 0.00 | 0.00 | 0.40 | 0.08 | 27 | Mixed Epithelial |
| Tmc4 | 0.50 | 0.00 | 0.00 | 0.28 | 0.04 | 27 | Mixed Epithelial |
| Alcam | 0.74 | 0.00 | 0.00 | 0.77 | 0.23 | 27 | Mixed Epithelial |
| Atp6v1c2 | 0.46 | 0.00 | 0.00 | 0.32 | 0.05 | 27 | Mixed Epithelial |
| Wwc1 | 0.54 | 0.00 | 0.00 | 0.36 | 0.06 | 27 | Mixed Epithelial |
| Gstm2 | 0.51 | 0.00 | 0.00 | 0.36 | 0.06 | 27 | Mixed Epithelial |
| Ascc2 | 0.45 | 0.00 | 0.00 | 0.28 | 0.04 | 27 | Mixed Epithelial |
| Col23a1 | 0.64 | 0.00 | 0.00 | 0.30 | 0.05 | 27 | Mixed Epithelial |
| 2200002D01Rik | 0.60 | 0.00 | 0.00 | 0.39 | 0.07 | 27 | Mixed Epithelial |
| Cyb5 | 0.97 | 0.00 | 0.00 | 0.73 | 0.26 | 27 | Mixed Epithelial |
| Carkd | 0.82 | 0.00 | 0.00 | 0.57 | 0.16 | 27 | Mixed Epithelial |
| Emb | 0.57 | 0.00 | 0.00 | 0.49 | 0.11 | 27 | Mixed Epithelial |
| Atp11a | 0.81 | 0.00 | 0.00 | 0.48 | 0.11 | 27 | Mixed Epithelial |
| C3 | 0.37 | 0.00 | 0.00 | 0.28 | 0.04 | 27 | Mixed Epithelial |
| Cd24a | 0.51 | 0.00 | 0.00 | 0.53 | 0.13 | 27 | Mixed Epithelial |
| Krt7 | 0.54 | 0.00 | 0.00 | 0.39 | 0.08 | 27 | Mixed Epithelial |
| Gclc | 0.59 | 0.00 | 0.00 | 0.40 | 0.09 | 27 | Mixed Epithelial |
| Pon3 | 0.55 | 0.00 | 0.00 | 0.44 | 0.10 | 27 | Mixed Epithelial |
| Cd9 | 0.83 | 0.00 | 0.00 | 0.84 | 0.35 | 27 | Mixed Epithelial |
| Smim6 | 0.47 | 0.00 | 0.00 | 0.26 | 0.04 | 27 | Mixed Epithelial |
| Gstm1 | 0.70 | 0.00 | 0.00 | 0.56 | 0.15 | 27 | Mixed Epithelial |
| Smim22 | 0.44 | 0.00 | 0.00 | 0.26 | 0.04 | 27 | Mixed Epithelial |
| Bsg | 0.83 | 0.00 | 0.00 | 0.84 | 0.36 | 27 | Mixed Epithelial |
| Ssx2ip | 0.56 | 0.00 | 0.00 | 0.26 | 0.04 | 27 | Mixed Epithelial |
| Pgap2 | 0.51 | 0.00 | 0.00 | 0.43 | 0.10 | 27 | Mixed Epithelial |
| Wfdc1 | 0.49 | 0.00 | 0.00 | 0.32 | 0.06 | 27 | Mixed Epithelial |
| Scnn1a | 0.56 | 0.00 | 0.00 | 0.35 | 0.07 | 27 | Mixed Epithelial |
| Cp | 0.91 | 0.00 | 0.00 | 0.47 | 0.12 | 27 | Mixed Epithelial |
| Fmo2 | 0.52 | 0.00 | 0.00 | 0.57 | 0.17 | 27 | Mixed Epithelial |
| Npc2 | 0.54 | 0.00 | 0.00 | 0.93 | 0.54 | 27 | Mixed Epithelial |
| Sdc4 | 0.59 | 0.00 | 0.00 | 0.69 | 0.23 | 27 | Mixed Epithelial |
| Gprc5a | 0.41 | 0.00 | 0.00 | 0.34 | 0.07 | 27 | Mixed Epithelial |
| Fdx1 | 0.65 | 0.00 | 0.00 | 0.49 | 0.14 | 27 | Mixed Epithelial |
| Sdc1 | 0.58 | 0.00 | 0.00 | 0.45 | 0.12 | 27 | Mixed Epithelial |
| Car8 | 0.39 | 0.00 | 0.00 | 0.36 | 0.08 | 27 | Mixed Epithelial |
| Chka | 0.61 | 0.00 | 0.00 | 0.41 | 0.10 | 27 | Mixed Epithelial |
| Prdx6b | 0.52 | 0.00 | 0.00 | 0.29 | 0.06 | 27 | Mixed Epithelial |
| Slc12a2 | 0.59 | 0.00 | 0.00 | 0.50 | 0.15 | 27 | Mixed Epithelial |
| Tmem176a | 0.63 | 0.00 | 0.00 | 0.60 | 0.20 | 27 | Mixed Epithelial |
| Tcea3 | 0.40 | 0.00 | 0.00 | 0.34 | 0.08 | 27 | Mixed Epithelial |
| Acsl4 | 0.54 | 0.00 | 0.00 | 0.43 | 0.12 | 27 | Mixed Epithelial |
| Ide | 0.47 | 0.00 | 0.00 | 0.36 | 0.09 | 27 | Mixed Epithelial |
| Cystm1 | 0.45 | 0.00 | 0.00 | 0.34 | 0.08 | 27 | Mixed Epithelial |
| Pgrmc1 | 0.64 | 0.00 | 0.00 | 0.54 | 0.19 | 27 | Mixed Epithelial |
| Atp8a1 | 0.63 | 0.00 | 0.00 | 0.50 | 0.15 | 27 | Mixed Epithelial |
| Anxa1 | 0.87 | 0.00 | 0.00 | 0.56 | 0.19 | 27 | Mixed Epithelial |
| Gdpd1 | 0.44 | 0.00 | 0.00 | 0.29 | 0.06 | 27 | Mixed Epithelial |
| Me1 | 0.48 | 0.00 | 0.00 | 0.27 | 0.06 | 27 | Mixed Epithelial |
| Fgf1 | 0.43 | 0.00 | 0.00 | 0.29 | 0.06 | 27 | Mixed Epithelial |
| Avpi1 | 0.37 | 0.00 | 0.00 | 0.35 | 0.09 | 27 | Mixed Epithelial |
| Tpd52 | 0.46 | 0.00 | 0.00 | 0.39 | 0.11 | 27 | Mixed Epithelial |
| Cyp4b1 | 0.32 | 0.00 | 0.00 | 0.41 | 0.11 | 27 | Mixed Epithelial |
| Larp4b | 0.56 | 0.00 | 0.00 | 0.48 | 0.16 | 27 | Mixed Epithelial |
| Slc26a2 | 0.46 | 0.00 | 0.00 | 0.29 | 0.07 | 27 | Mixed Epithelial |
| Spink5 | 0.80 | 0.00 | 0.00 | 0.32 | 0.08 | 27 | Mixed Epithelial |
| Hes1 | 0.83 | 0.00 | 0.00 | 0.66 | 0.29 | 27 | Mixed Epithelial |
| Myh14 | 0.33 | 0.00 | 0.00 | 0.36 | 0.10 | 27 | Mixed Epithelial |
| Phyh | 0.38 | 0.00 | 0.00 | 0.27 | 0.06 | 27 | Mixed Epithelial |
| Mecom | 0.33 | 0.00 | 0.00 | 0.36 | 0.10 | 27 | Mixed Epithelial |
| Ccnd1 | 0.38 | 0.00 | 0.00 | 0.54 | 0.19 | 27 | Mixed Epithelial |
| Neat1 | 0.41 | 0.00 | 0.00 | 0.67 | 0.27 | 27 | Mixed Epithelial |
| S100a11 | 0.59 | 0.00 | 0.00 | 0.85 | 0.49 | 27 | Mixed Epithelial |
| Gfpt1 | 0.35 | 0.00 | 0.00 | 0.27 | 0.07 | 27 | Mixed Epithelial |
| Wbp5 | 0.59 | 0.00 | 0.00 | 0.83 | 0.48 | 27 | Mixed Epithelial |
| Runx1 | 0.34 | 0.00 | 0.00 | 0.28 | 0.07 | 27 | Mixed Epithelial |
| Ppap2b | 0.49 | 0.00 | 0.00 | 0.59 | 0.24 | 27 | Mixed Epithelial |
| Nkx2-1 | 0.37 | 0.00 | 0.00 | 0.34 | 0.10 | 27 | Mixed Epithelial |
| Pgrmc2 | 0.34 | 0.00 | 0.00 | 0.31 | 0.08 | 27 | Mixed Epithelial |
| Ndufaf7 | 0.31 | 0.00 | 0.00 | 0.28 | 0.07 | 27 | Mixed Epithelial |
| Wwp1 | 0.36 | 0.00 | 0.00 | 0.41 | 0.14 | 27 | Mixed Epithelial |
| Idh2 | 0.44 | 0.00 | 0.00 | 0.51 | 0.19 | 27 | Mixed Epithelial |
| Uqcr10 | 0.51 | 0.00 | 0.00 | 0.73 | 0.36 | 27 | Mixed Epithelial |
| Cgnl1 | 0.50 | 0.00 | 0.00 | 0.37 | 0.12 | 27 | Mixed Epithelial |
| Tef | 0.29 | 0.00 | 0.00 | 0.28 | 0.08 | 27 | Mixed Epithelial |
| Vamp8 | 0.55 | 0.00 | 0.00 | 0.72 | 0.34 | 27 | Mixed Epithelial |
| Ifrd1 | 0.40 | 0.00 | 0.00 | 0.44 | 0.16 | 27 | Mixed Epithelial |
| Il6st | 0.34 | 0.00 | 0.00 | 0.31 | 0.09 | 27 | Mixed Epithelial |
| Cltb | 0.32 | 0.00 | 0.00 | 0.35 | 0.11 | 27 | Mixed Epithelial |
| Mtus1 | 0.45 | 0.00 | 0.00 | 0.45 | 0.17 | 27 | Mixed Epithelial |
| Oat | 0.41 | 0.00 | 0.00 | 0.37 | 0.13 | 27 | Mixed Epithelial |
| Fth1 | 0.31 | 0.00 | 0.00 | 0.96 | 0.81 | 27 | Mixed Epithelial |
| Rfk | 0.35 | 0.00 | 0.00 | 0.32 | 0.10 | 27 | Mixed Epithelial |
| Mbip | 0.35 | 0.00 | 0.00 | 0.34 | 0.12 | 27 | Mixed Epithelial |
| mt-Nd5 | 0.52 | 0.00 | 0.00 | 0.93 | 0.76 | 27 | Mixed Epithelial |
| Vmp1 | 0.29 | 0.00 | 0.00 | 0.40 | 0.14 | 27 | Mixed Epithelial |
| Ndufc2 | 0.42 | 0.00 | 0.00 | 0.74 | 0.39 | 27 | Mixed Epithelial |
| Ptplb | 0.28 | 0.00 | 0.00 | 0.26 | 0.08 | 27 | Mixed Epithelial |
| Fkbp4 | 0.37 | 0.00 | 0.00 | 0.40 | 0.15 | 27 | Mixed Epithelial |
| Sat1 | 0.35 | 0.00 | 0.00 | 0.67 | 0.33 | 27 | Mixed Epithelial |
| Tmem109 | 0.32 | 0.00 | 0.00 | 0.35 | 0.13 | 27 | Mixed Epithelial |
| Tmem50b | 0.33 | 0.00 | 0.00 | 0.26 | 0.08 | 27 | Mixed Epithelial |
| Gde1 | 0.38 | 0.00 | 0.00 | 0.39 | 0.15 | 27 | Mixed Epithelial |
| Nfe2l2 | 0.41 | 0.00 | 0.00 | 0.55 | 0.25 | 27 | Mixed Epithelial |
| Prrg3 | 0.34 | 0.00 | 0.00 | 0.25 | 0.08 | 27 | Mixed Epithelial |
| Vamp5 | 0.28 | 0.00 | 0.00 | 0.29 | 0.10 | 27 | Mixed Epithelial |
| Lrrfip2 | 0.32 | 0.00 | 0.00 | 0.37 | 0.14 | 27 | Mixed Epithelial |
| mt-Co1 | 0.34 | 0.00 | 0.00 | 0.99 | 0.95 | 27 | Mixed Epithelial |
| Pls3 | 0.41 | 0.00 | 0.00 | 0.47 | 0.20 | 27 | Mixed Epithelial |
| Paics | 0.34 | 0.00 | 0.00 | 0.52 | 0.23 | 27 | Mixed Epithelial |
| mt-Cytb | 0.35 | 0.00 | 0.00 | 0.96 | 0.82 | 27 | Mixed Epithelial |
| Emc6 | 0.28 | 0.00 | 0.00 | 0.36 | 0.14 | 27 | Mixed Epithelial |
| Prdx1 | 0.41 | 0.00 | 0.00 | 0.78 | 0.50 | 27 | Mixed Epithelial |
| Bag1 | 0.38 | 0.00 | 0.00 | 0.51 | 0.24 | 27 | Mixed Epithelial |
| Prkci | 0.33 | 0.00 | 0.00 | 0.34 | 0.13 | 27 | Mixed Epithelial |
| Pcyt1a | 0.28 | 0.00 | 0.00 | 0.30 | 0.11 | 27 | Mixed Epithelial |
| Commd2 | 0.31 | 0.00 | 0.00 | 0.28 | 0.10 | 27 | Mixed Epithelial |
| Sod2 | 0.32 | 0.00 | 0.00 | 0.34 | 0.14 | 27 | Mixed Epithelial |
| Idh1 | 0.37 | 0.00 | 0.00 | 0.27 | 0.10 | 27 | Mixed Epithelial |
| Sod1 | 0.46 | 0.00 | 0.00 | 0.77 | 0.47 | 27 | Mixed Epithelial |
| mt-Nd4l | 0.34 | 0.00 | 0.00 | 0.71 | 0.38 | 27 | Mixed Epithelial |
| Mdh1 | 0.29 | 0.00 | 0.00 | 0.43 | 0.19 | 27 | Mixed Epithelial |
| Aldh2 | 0.28 | 0.00 | 0.00 | 0.64 | 0.33 | 27 | Mixed Epithelial |
| Dstn | 0.39 | 0.00 | 0.00 | 0.80 | 0.51 | 27 | Mixed Epithelial |
| Ccnd2 | 0.34 | 0.00 | 0.00 | 0.56 | 0.28 | 27 | Mixed Epithelial |
| mt-Nd6 | 0.40 | 0.00 | 0.00 | 0.52 | 0.26 | 27 | Mixed Epithelial |
| Gm10222 | 0.34 | 0.00 | 0.00 | 0.72 | 0.43 | 27 | Mixed Epithelial |
| Dynll2 | 0.28 | 0.00 | 0.00 | 0.29 | 0.12 | 27 | Mixed Epithelial |
| Vimp | 0.27 | 0.00 | 0.00 | 0.56 | 0.30 | 27 | Mixed Epithelial |
| Qsox1 | 0.25 | 0.00 | 0.00 | 0.25 | 0.10 | 27 | Mixed Epithelial |
| Gstp1 | 0.26 | 0.00 | 0.00 | 0.40 | 0.20 | 27 | Mixed Epithelial |
| Ppp3ca | 0.32 | 0.00 | 0.00 | 0.52 | 0.28 | 27 | Mixed Epithelial |
| Hdgf | 0.27 | 0.00 | 0.00 | 0.47 | 0.25 | 27 | Mixed Epithelial |
| Gpx4 | 0.28 | 0.00 | 0.00 | 0.62 | 0.38 | 27 | Mixed Epithelial |
| Mcpt8 | 4.49 | 0.00 | 0.00 | 0.61 | 0.00 | 28 | Myeloid Cell |
| Cpa3 | 3.98 | 0.00 | 0.00 | 0.59 | 0.01 | 28 | Myeloid Cell |
| Prss34 | 3.13 | 0.00 | 0.00 | 0.30 | 0.00 | 28 | Myeloid Cell |
| Ccl4 | 3.01 | 0.00 | 0.00 | 0.42 | 0.01 | 28 | Myeloid Cell |
| Gp49a | 2.98 | 0.00 | 0.00 | 0.89 | 0.05 | 28 | Myeloid Cell |
| Ccl3 | 2.97 | 0.00 | 0.00 | 0.61 | 0.01 | 28 | Myeloid Cell |
| Ccl9 | 2.96 | 0.00 | 0.00 | 0.69 | 0.03 | 28 | Myeloid Cell |
| Hdc | 2.70 | 0.00 | 0.00 | 0.80 | 0.04 | 28 | Myeloid Cell |
| Alox5ap | 2.66 | 0.00 | 0.00 | 0.89 | 0.09 | 28 | Myeloid Cell |
| Lilrb4 | 2.63 | 0.00 | 0.00 | 0.80 | 0.06 | 28 | Myeloid Cell |
| Cd200r3 | 2.50 | 0.00 | 0.00 | 0.74 | 0.00 | 28 | Myeloid Cell |
| Cyp11a1 | 2.28 | 0.00 | 0.00 | 0.66 | 0.00 | 28 | Myeloid Cell |
| Il1rl1 | 2.18 | 0.00 | 0.00 | 0.46 | 0.01 | 28 | Myeloid Cell |
| Csf2rb | 2.03 | 0.00 | 0.00 | 0.62 | 0.04 | 28 | Myeloid Cell |
| Cxcr2 | 1.93 | 0.00 | 0.00 | 0.45 | 0.02 | 28 | Myeloid Cell |
| Lat2 | 1.86 | 0.00 | 0.00 | 0.59 | 0.03 | 28 | Myeloid Cell |
| Ifitm7 | 1.86 | 0.00 | 0.00 | 0.47 | 0.03 | 28 | Myeloid Cell |
| Osm | 1.82 | 0.00 | 0.00 | 0.36 | 0.01 | 28 | Myeloid Cell |
| Emilin2 | 1.68 | 0.00 | 0.00 | 0.45 | 0.02 | 28 | Myeloid Cell |
| Gpr183 | 1.66 | 0.00 | 0.00 | 0.42 | 0.02 | 28 | Myeloid Cell |
| Il4 | 1.54 | 0.00 | 0.00 | 0.31 | 0.00 | 28 | Myeloid Cell |
| Fcer1a | 1.51 | 0.00 | 0.00 | 0.32 | 0.00 | 28 | Myeloid Cell |
| Sytl3 | 1.48 | 0.00 | 0.00 | 0.38 | 0.00 | 28 | Myeloid Cell |
| Ms4a2 | 1.37 | 0.00 | 0.00 | 0.29 | 0.00 | 28 | Myeloid Cell |
| Csf2rb2 | 1.36 | 0.00 | 0.00 | 0.34 | 0.01 | 28 | Myeloid Cell |
| Il18rap | 1.35 | 0.00 | 0.00 | 0.36 | 0.01 | 28 | Myeloid Cell |
| Rgs18 | 1.21 | 0.00 | 0.00 | 0.34 | 0.01 | 28 | Myeloid Cell |
| Rab44 | 1.20 | 0.00 | 0.00 | 0.31 | 0.01 | 28 | Myeloid Cell |
| P2rx1 | 1.06 | 0.00 | 0.00 | 0.26 | 0.01 | 28 | Myeloid Cell |
| Tyrobp | 2.13 | 0.00 | 0.00 | 0.93 | 0.12 | 28 | Myeloid Cell |
| Cyp4f18 | 1.53 | 0.00 | 0.00 | 0.41 | 0.02 | 28 | Myeloid Cell |
| Rgs1 | 1.62 | 0.00 | 0.00 | 0.32 | 0.01 | 28 | Myeloid Cell |
| Hgf | 1.46 | 0.00 | 0.00 | 0.29 | 0.01 | 28 | Myeloid Cell |
| Csf1 | 1.84 | 0.00 | 0.00 | 0.46 | 0.03 | 28 | Myeloid Cell |
| Fcer1g | 1.74 | 0.00 | 0.00 | 0.85 | 0.11 | 28 | Myeloid Cell |
| Itgb7 | 1.37 | 0.00 | 0.00 | 0.43 | 0.03 | 28 | Myeloid Cell |
| Ifitm1 | 3.39 | 0.00 | 0.00 | 0.79 | 0.14 | 28 | Myeloid Cell |
| Tbc1d4 | 1.59 | 0.00 | 0.00 | 0.43 | 0.04 | 28 | Myeloid Cell |
| Gata2 | 2.02 | 0.00 | 0.00 | 0.79 | 0.14 | 28 | Myeloid Cell |
| Ccl6 | 2.75 | 0.00 | 0.00 | 0.67 | 0.10 | 28 | Myeloid Cell |
| Ccr2 | 1.64 | 0.00 | 0.00 | 0.61 | 0.07 | 28 | Myeloid Cell |
| Fxyd5 | 1.96 | 0.00 | 0.00 | 0.86 | 0.18 | 28 | Myeloid Cell |
| Tmem71 | 1.84 | 0.00 | 0.00 | 0.53 | 0.06 | 28 | Myeloid Cell |
| Laptm5 | 1.56 | 0.00 | 0.00 | 0.79 | 0.14 | 28 | Myeloid Cell |
| Fcgr3 | 1.41 | 0.00 | 0.00 | 0.45 | 0.04 | 28 | Myeloid Cell |
| Ikzf2 | 1.10 | 0.00 | 0.00 | 0.29 | 0.02 | 28 | Myeloid Cell |
| Srgn | 2.11 | 0.00 | 0.00 | 0.98 | 0.33 | 28 | Myeloid Cell |
| Cd53 | 1.47 | 0.00 | 0.00 | 0.60 | 0.09 | 28 | Myeloid Cell |
| Plek | 1.37 | 0.00 | 0.00 | 0.52 | 0.07 | 28 | Myeloid Cell |
| Rac2 | 1.51 | 0.00 | 0.00 | 0.65 | 0.11 | 28 | Myeloid Cell |
| Fyb | 1.70 | 0.00 | 0.00 | 0.78 | 0.18 | 28 | Myeloid Cell |
| Gm7676 | 2.41 | 0.00 | 0.00 | 0.74 | 0.19 | 28 | Myeloid Cell |
| Selplg | 1.32 | 0.00 | 0.00 | 0.43 | 0.05 | 28 | Myeloid Cell |
| Serpinb1a | 2.09 | 0.00 | 0.00 | 0.39 | 0.05 | 28 | Myeloid Cell |
| Tec | 1.33 | 0.00 | 0.00 | 0.32 | 0.03 | 28 | Myeloid Cell |
| Arhgdib | 1.38 | 0.00 | 0.00 | 0.72 | 0.16 | 28 | Myeloid Cell |
| Arhgap15 | 1.21 | 0.00 | 0.00 | 0.38 | 0.05 | 28 | Myeloid Cell |
| Ccr1 | 1.08 | 0.00 | 0.00 | 0.32 | 0.04 | 28 | Myeloid Cell |
| Ptplad2 | 1.30 | 0.00 | 0.00 | 0.37 | 0.05 | 28 | Myeloid Cell |
| Lgals3 | 1.31 | 0.00 | 0.00 | 0.65 | 0.15 | 28 | Myeloid Cell |
| Plac8 | 1.24 | 0.00 | 0.00 | 0.70 | 0.17 | 28 | Myeloid Cell |
| Btk | 1.13 | 0.00 | 0.00 | 0.31 | 0.04 | 28 | Myeloid Cell |
| Spn | 0.92 | 0.00 | 0.00 | 0.26 | 0.03 | 28 | Myeloid Cell |
| Samsn1 | 1.19 | 0.00 | 0.00 | 0.40 | 0.07 | 28 | Myeloid Cell |
| Lcp1 | 1.38 | 0.00 | 0.00 | 0.72 | 0.23 | 28 | Myeloid Cell |
| Ncf2 | 1.01 | 0.00 | 0.00 | 0.30 | 0.04 | 28 | Myeloid Cell |
| Tnfaip8 | 1.45 | 0.00 | 0.00 | 0.47 | 0.10 | 28 | Myeloid Cell |
| Cd63 | 1.32 | 0.00 | 0.00 | 0.76 | 0.30 | 28 | Myeloid Cell |
| A430104N18Rik | 1.07 | 0.00 | 0.00 | 0.37 | 0.06 | 28 | Myeloid Cell |
| Hcst | 1.05 | 0.00 | 0.00 | 0.28 | 0.04 | 28 | Myeloid Cell |
| Gmfg | 1.21 | 0.00 | 0.00 | 0.54 | 0.14 | 28 | Myeloid Cell |
| Sh3bgrl3 | 1.26 | 0.00 | 0.00 | 0.79 | 0.34 | 28 | Myeloid Cell |
| Ptpre | 1.07 | 0.00 | 0.00 | 0.35 | 0.06 | 28 | Myeloid Cell |
| Taldo1 | 1.32 | 0.00 | 0.00 | 0.70 | 0.26 | 28 | Myeloid Cell |
| Hk2 | 1.13 | 0.00 | 0.00 | 0.31 | 0.05 | 28 | Myeloid Cell |
| Gm6169 | 1.30 | 0.00 | 0.00 | 0.48 | 0.13 | 28 | Myeloid Cell |
| Ncf1 | 1.20 | 0.00 | 0.00 | 0.36 | 0.07 | 28 | Myeloid Cell |
| Hmha1 | 1.04 | 0.00 | 0.00 | 0.39 | 0.08 | 28 | Myeloid Cell |
| Klhl6 | 0.99 | 0.00 | 0.00 | 0.27 | 0.04 | 28 | Myeloid Cell |
| Coro2a | 1.16 | 0.00 | 0.00 | 0.35 | 0.07 | 28 | Myeloid Cell |
| Prr13 | 1.14 | 0.00 | 0.00 | 0.49 | 0.13 | 28 | Myeloid Cell |
| Lgals1 | 1.00 | 0.00 | 0.00 | 0.89 | 0.42 | 28 | Myeloid Cell |
| Plp2 | 1.29 | 0.00 | 0.00 | 0.47 | 0.13 | 28 | Myeloid Cell |
| Actb | 0.81 | 0.00 | 0.00 | 1.00 | 0.98 | 28 | Myeloid Cell |
| Syne1 | 1.32 | 0.00 | 0.00 | 0.43 | 0.11 | 28 | Myeloid Cell |
| Lst1 | 0.93 | 0.00 | 0.00 | 0.35 | 0.07 | 28 | Myeloid Cell |
| Crlf3 | 1.18 | 0.00 | 0.00 | 0.49 | 0.14 | 28 | Myeloid Cell |
| Cd37 | 0.78 | 0.00 | 0.00 | 0.37 | 0.08 | 28 | Myeloid Cell |
| Cotl1 | 1.16 | 0.00 | 0.00 | 0.55 | 0.18 | 28 | Myeloid Cell |
| Emb | 1.14 | 0.00 | 0.00 | 0.43 | 0.11 | 28 | Myeloid Cell |
| Was | 0.93 | 0.00 | 0.00 | 0.30 | 0.06 | 28 | Myeloid Cell |
| Tmsb4x | 1.10 | 0.00 | 0.00 | 0.98 | 0.89 | 28 | Myeloid Cell |
| Arpc5 | 1.00 | 0.00 | 0.00 | 0.88 | 0.56 | 28 | Myeloid Cell |
| Lrrc58 | 0.80 | 0.00 | 0.00 | 0.98 | 0.91 | 28 | Myeloid Cell |
| Cd52 | 0.79 | 0.00 | 0.00 | 0.64 | 0.20 | 28 | Myeloid Cell |
| Apbb1ip | 0.95 | 0.00 | 0.00 | 0.39 | 0.09 | 28 | Myeloid Cell |
| Igsf6 | 1.12 | 0.00 | 0.00 | 0.25 | 0.04 | 28 | Myeloid Cell |
| Cd84 | 1.00 | 0.00 | 0.00 | 0.27 | 0.05 | 28 | Myeloid Cell |
| Arhgap30 | 0.94 | 0.00 | 0.00 | 0.40 | 0.10 | 28 | Myeloid Cell |
| Fam46a | 1.20 | 0.00 | 0.00 | 0.38 | 0.10 | 28 | Myeloid Cell |
| Coro1a | 0.92 | 0.00 | 0.00 | 0.39 | 0.10 | 28 | Myeloid Cell |
| Pla2g7 | 0.88 | 0.00 | 0.00 | 0.29 | 0.06 | 28 | Myeloid Cell |
| Rgs2 | 1.27 | 0.00 | 0.00 | 0.66 | 0.28 | 28 | Myeloid Cell |
| Tmem64 | 1.08 | 0.00 | 0.00 | 0.38 | 0.11 | 28 | Myeloid Cell |
| Fermt3 | 0.85 | 0.00 | 0.00 | 0.29 | 0.06 | 28 | Myeloid Cell |
| Anxa1 | 1.02 | 0.00 | 0.00 | 0.52 | 0.19 | 28 | Myeloid Cell |
| Runx1 | 1.01 | 0.00 | 0.00 | 0.30 | 0.07 | 28 | Myeloid Cell |
| Man2b1 | 1.00 | 0.00 | 0.00 | 0.30 | 0.07 | 28 | Myeloid Cell |
| Arrb2 | 1.00 | 0.00 | 0.00 | 0.28 | 0.07 | 28 | Myeloid Cell |
| Plgrkt | 1.05 | 0.00 | 0.00 | 0.37 | 0.11 | 28 | Myeloid Cell |
| Ctsd | 1.01 | 0.00 | 0.00 | 0.57 | 0.25 | 28 | Myeloid Cell |
| Gsr | 0.95 | 0.00 | 0.00 | 0.33 | 0.09 | 28 | Myeloid Cell |
| Fam105a | 0.90 | 0.00 | 0.00 | 0.27 | 0.06 | 28 | Myeloid Cell |
| Rplp1 | 0.58 | 0.00 | 0.00 | 0.99 | 0.90 | 28 | Myeloid Cell |
| Ptpn18 | 0.95 | 0.00 | 0.00 | 0.43 | 0.14 | 28 | Myeloid Cell |
| Glipr2 | 0.80 | 0.00 | 0.00 | 0.26 | 0.06 | 28 | Myeloid Cell |
| Capg | 0.90 | 0.00 | 0.00 | 0.35 | 0.11 | 28 | Myeloid Cell |
| S100a10 | 0.95 | 0.00 | 0.00 | 0.70 | 0.38 | 28 | Myeloid Cell |
| Mpc2 | 1.24 | 0.00 | 0.00 | 0.52 | 0.24 | 28 | Myeloid Cell |
| Rfc2 | 0.79 | 0.00 | 0.00 | 0.30 | 0.08 | 28 | Myeloid Cell |
| Klf6 | 1.06 | 0.00 | 0.00 | 0.67 | 0.36 | 28 | Myeloid Cell |
| Casp3 | 0.94 | 0.00 | 0.00 | 0.29 | 0.08 | 28 | Myeloid Cell |
| Gm2a | 0.75 | 0.00 | 0.00 | 0.32 | 0.09 | 28 | Myeloid Cell |
| Egr1 | 1.30 | 0.00 | 0.00 | 0.39 | 0.14 | 28 | Myeloid Cell |
| H2-T23 | 0.86 | 0.00 | 0.00 | 0.38 | 0.13 | 28 | Myeloid Cell |
| Stk17b | 0.84 | 0.00 | 0.00 | 0.26 | 0.06 | 28 | Myeloid Cell |
| Gm9843 | 0.55 | 0.00 | 0.00 | 0.98 | 0.93 | 28 | Myeloid Cell |
| Arpc1b | 0.83 | 0.00 | 0.00 | 0.68 | 0.37 | 28 | Myeloid Cell |
| Il18r1 | 1.12 | 0.00 | 0.00 | 0.27 | 0.07 | 28 | Myeloid Cell |
| Limd2 | 0.92 | 0.00 | 0.00 | 0.42 | 0.15 | 28 | Myeloid Cell |
| Rps9 | 0.50 | 0.00 | 0.00 | 0.97 | 0.91 | 28 | Myeloid Cell |
| Os9 | 1.03 | 0.00 | 0.00 | 0.48 | 0.21 | 28 | Myeloid Cell |
| Ptprc | 0.64 | 0.00 | 0.00 | 0.43 | 0.15 | 28 | Myeloid Cell |
| Grina | 0.84 | 0.00 | 0.00 | 0.26 | 0.07 | 28 | Myeloid Cell |
| Fau | 0.56 | 0.00 | 0.00 | 0.95 | 0.87 | 28 | Myeloid Cell |
| Neat1 | 0.78 | 0.00 | 0.00 | 0.59 | 0.27 | 28 | Myeloid Cell |
| Zfp36 | 0.96 | 0.00 | 0.00 | 0.42 | 0.17 | 28 | Myeloid Cell |
| Leprotl1 | 0.89 | 0.00 | 0.00 | 0.40 | 0.15 | 28 | Myeloid Cell |
| Gpx1 | 0.74 | 0.00 | 0.00 | 0.71 | 0.43 | 28 | Myeloid Cell |
| Rps23 | 0.67 | 0.00 | 0.00 | 0.91 | 0.79 | 28 | Myeloid Cell |
| Slfn2 | 0.87 | 0.00 | 0.00 | 0.38 | 0.13 | 28 | Myeloid Cell |
| Itm2b | 0.62 | 0.00 | 0.00 | 0.92 | 0.76 | 28 | Myeloid Cell |
| Gm10073 | 0.56 | 0.00 | 0.00 | 0.95 | 0.86 | 28 | Myeloid Cell |
| Junb | 1.17 | 0.00 | 0.00 | 0.46 | 0.20 | 28 | Myeloid Cell |
| Sec11c | 0.85 | 0.00 | 0.00 | 0.44 | 0.18 | 28 | Myeloid Cell |
| Ptprcap | 0.62 | 0.00 | 0.00 | 0.27 | 0.07 | 28 | Myeloid Cell |
| Crip1 | 0.65 | 0.00 | 0.00 | 0.79 | 0.48 | 28 | Myeloid Cell |
| Pfn1 | 0.80 | 0.00 | 0.00 | 0.71 | 0.46 | 28 | Myeloid Cell |
| Sh3kbp1 | 0.87 | 0.00 | 0.00 | 0.32 | 0.11 | 28 | Myeloid Cell |
| Sfxn1 | 0.73 | 0.00 | 0.00 | 0.25 | 0.07 | 28 | Myeloid Cell |
| Creg1 | 0.83 | 0.00 | 0.00 | 0.42 | 0.17 | 28 | Myeloid Cell |
| Rnf130 | 0.86 | 0.00 | 0.00 | 0.25 | 0.07 | 28 | Myeloid Cell |
| Pabpc1 | 0.56 | 0.00 | 0.00 | 0.94 | 0.81 | 28 | Myeloid Cell |
| Pygl | 0.81 | 0.00 | 0.00 | 0.32 | 0.11 | 28 | Myeloid Cell |
| Taok3 | 0.81 | 0.00 | 0.00 | 0.43 | 0.19 | 28 | Myeloid Cell |
| Aldoa | 0.93 | 0.00 | 0.00 | 0.43 | 0.19 | 28 | Myeloid Cell |
| Arpc2 | 0.65 | 0.00 | 0.00 | 0.81 | 0.61 | 28 | Myeloid Cell |
| Rps11 | 0.58 | 0.00 | 0.00 | 0.88 | 0.76 | 28 | Myeloid Cell |
| Ier3 | 1.31 | 0.00 | 0.00 | 0.34 | 0.14 | 28 | Myeloid Cell |
| Stk4 | 0.69 | 0.00 | 0.00 | 0.36 | 0.14 | 28 | Myeloid Cell |
| Pim1 | 0.94 | 0.00 | 0.00 | 0.28 | 0.09 | 28 | Myeloid Cell |
| Cd9 | 0.70 | 0.00 | 0.00 | 0.60 | 0.35 | 28 | Myeloid Cell |
| Cd47 | 0.70 | 0.00 | 0.00 | 0.65 | 0.43 | 28 | Myeloid Cell |
| Prkcd | 0.72 | 0.00 | 0.00 | 0.30 | 0.11 | 28 | Myeloid Cell |
| Ostf1 | 0.67 | 0.00 | 0.00 | 0.48 | 0.26 | 28 | Myeloid Cell |
| Rpl38 | 0.54 | 0.00 | 0.00 | 0.87 | 0.73 | 28 | Myeloid Cell |
| Tspo | 0.70 | 0.00 | 0.00 | 0.48 | 0.26 | 28 | Myeloid Cell |
| Rnasel | 0.61 | 0.00 | 0.00 | 0.28 | 0.10 | 28 | Myeloid Cell |
| Clic1 | 0.66 | 0.00 | 0.00 | 0.59 | 0.38 | 28 | Myeloid Cell |
| H2-D1 | 0.47 | 0.00 | 0.00 | 0.75 | 0.50 | 28 | Myeloid Cell |
| Vasp | 0.71 | 0.00 | 0.00 | 0.39 | 0.19 | 28 | Myeloid Cell |
| Ldha | 0.77 | 0.00 | 0.00 | 0.49 | 0.29 | 28 | Myeloid Cell |
| Lbr | 0.77 | 0.00 | 0.00 | 0.30 | 0.12 | 28 | Myeloid Cell |
| Cd97 | 0.67 | 0.00 | 0.00 | 0.38 | 0.18 | 28 | Myeloid Cell |
| Ncor1 | 0.67 | 0.00 | 0.00 | 0.61 | 0.40 | 28 | Myeloid Cell |
| Herc1 | 0.89 | 0.00 | 0.00 | 0.29 | 0.13 | 28 | Myeloid Cell |
| Trim12c | 0.68 | 0.00 | 0.00 | 0.25 | 0.09 | 28 | Myeloid Cell |
| Actg1 | 0.61 | 0.00 | 0.00 | 0.81 | 0.76 | 28 | Myeloid Cell |
| Fam111a | 0.60 | 0.00 | 0.00 | 0.35 | 0.15 | 28 | Myeloid Cell |
| Fos | 0.73 | 0.00 | 0.00 | 0.39 | 0.19 | 28 | Myeloid Cell |
| Arid3a | 0.58 | 0.00 | 0.00 | 0.26 | 0.10 | 28 | Myeloid Cell |
| Tagln2 | 0.59 | 0.00 | 0.00 | 0.65 | 0.45 | 28 | Myeloid Cell |
| Tln1 | 0.67 | 0.00 | 0.00 | 0.58 | 0.40 | 28 | Myeloid Cell |
| Arl4c | 0.87 | 0.00 | 0.00 | 0.25 | 0.10 | 28 | Myeloid Cell |
| Myh9 | 0.74 | 0.00 | 0.00 | 0.53 | 0.33 | 28 | Myeloid Cell |
| Rps24 | 0.33 | 0.00 | 0.00 | 0.95 | 0.88 | 28 | Myeloid Cell |
| Zyx | 0.65 | 0.00 | 0.00 | 0.37 | 0.20 | 28 | Myeloid Cell |
| Prkacb | 0.66 | 0.00 | 0.00 | 0.25 | 0.11 | 28 | Myeloid Cell |
| Gm11808 | 0.40 | 0.00 | 0.00 | 0.86 | 0.77 | 28 | Myeloid Cell |
| Anp32a | 0.41 | 0.00 | 0.00 | 0.78 | 0.60 | 28 | Myeloid Cell |
| Phf20l1 | 0.70 | 0.00 | 0.00 | 0.45 | 0.29 | 28 | Myeloid Cell |
| Cnn2 | 0.65 | 0.00 | 0.00 | 0.55 | 0.39 | 28 | Myeloid Cell |
| Ier2 | 0.79 | 0.00 | 0.00 | 0.41 | 0.25 | 28 | Myeloid Cell |
| Emp3 | 0.62 | 0.00 | 0.00 | 0.38 | 0.21 | 28 | Myeloid Cell |
| Rpl39 | 0.35 | 0.00 | 0.00 | 0.89 | 0.80 | 28 | Myeloid Cell |
| Rpl32 | 0.32 | 0.00 | 0.00 | 0.92 | 0.84 | 28 | Myeloid Cell |
| Sptssa | 0.79 | 0.00 | 0.00 | 0.35 | 0.20 | 28 | Myeloid Cell |
| Gm9846 | 0.26 | 0.00 | 0.00 | 0.98 | 0.90 | 28 | Myeloid Cell |
| Aprt | 0.62 | 0.00 | 0.00 | 0.32 | 0.17 | 28 | Myeloid Cell |
| Actr2 | 0.57 | 0.00 | 0.00 | 0.44 | 0.28 | 28 | Myeloid Cell |
| Iqgap1 | 0.56 | 0.00 | 0.00 | 0.61 | 0.45 | 28 | Myeloid Cell |
| Cfl1 | 0.42 | 0.00 | 0.00 | 0.72 | 0.58 | 28 | Myeloid Cell |
| Gm10116 | 0.34 | 0.00 | 0.00 | 0.76 | 0.63 | 28 | Myeloid Cell |
| Akap13 | 0.63 | 0.00 | 0.00 | 0.43 | 0.27 | 28 | Myeloid Cell |
| Ftl1 | 0.41 | 0.00 | 0.00 | 0.70 | 0.57 | 28 | Myeloid Cell |
| D4Wsu53e | 0.38 | 0.00 | 0.00 | 0.71 | 0.60 | 28 | Myeloid Cell |
| Jak1 | 0.46 | 0.00 | 0.00 | 0.42 | 0.26 | 28 | Myeloid Cell |
| Gng2 | 0.52 | 0.00 | 0.00 | 0.25 | 0.12 | 28 | Myeloid Cell |
| Cap1 | 0.52 | 0.00 | 0.00 | 0.31 | 0.17 | 28 | Myeloid Cell |
| Srpk1 | 0.67 | 0.00 | 0.00 | 0.25 | 0.13 | 28 | Myeloid Cell |
| 2410006H16Rik | 0.64 | 0.00 | 0.00 | 0.41 | 0.26 | 28 | Myeloid Cell |
| Cetn2 | 0.57 | 0.00 | 0.00 | 0.33 | 0.19 | 28 | Myeloid Cell |
| Ywhaz | 0.45 | 0.00 | 0.00 | 0.67 | 0.51 | 28 | Myeloid Cell |
| Wdr89 | 0.27 | 0.00 | 0.00 | 0.95 | 0.89 | 28 | Myeloid Cell |
| Tmem50a | 0.45 | 0.00 | 0.00 | 0.58 | 0.45 | 28 | Myeloid Cell |
| Rnh1 | 0.55 | 0.00 | 0.00 | 0.29 | 0.16 | 28 | Myeloid Cell |
| Samhd1 | 0.52 | 0.00 | 0.00 | 0.35 | 0.20 | 28 | Myeloid Cell |
| Gpx4 | 0.50 | 0.00 | 0.00 | 0.52 | 0.38 | 28 | Myeloid Cell |
| Cyba | 0.32 | 0.00 | 0.00 | 0.47 | 0.30 | 28 | Myeloid Cell |
| Gm17087 | 0.54 | 0.00 | 0.00 | 0.38 | 0.24 | 28 | Myeloid Cell |
| Rpl17 | 0.37 | 0.00 | 0.00 | 0.68 | 0.54 | 28 | Myeloid Cell |
| Clint1 | 0.64 | 0.00 | 0.00 | 0.25 | 0.14 | 28 | Myeloid Cell |
| Hmgb2 | 0.39 | 0.00 | 0.00 | 0.58 | 0.43 | 28 | Myeloid Cell |
| Rpl10 | 0.32 | 0.00 | 0.00 | 0.81 | 0.71 | 28 | Myeloid Cell |
| Nt5c | 0.50 | 0.00 | 0.00 | 0.29 | 0.16 | 28 | Myeloid Cell |
| Capzb | 0.47 | 0.00 | 0.00 | 0.55 | 0.42 | 28 | Myeloid Cell |
| Gltscr2 | 0.43 | 0.00 | 0.00 | 0.60 | 0.48 | 28 | Myeloid Cell |
| Mrpl33 | 0.56 | 0.00 | 0.00 | 0.42 | 0.30 | 28 | Myeloid Cell |
| Myl12b | 0.37 | 0.00 | 0.00 | 0.73 | 0.60 | 28 | Myeloid Cell |
| Serp1 | 0.49 | 0.00 | 0.00 | 0.37 | 0.23 | 28 | Myeloid Cell |
| Dusp6 | 0.57 | 0.00 | 0.00 | 0.31 | 0.18 | 28 | Myeloid Cell |
| Tax1bp1 | 0.43 | 0.00 | 0.00 | 0.71 | 0.64 | 28 | Myeloid Cell |
| Rpl18a | 0.30 | 0.00 | 0.00 | 0.86 | 0.82 | 28 | Myeloid Cell |
| Itm2c | 0.53 | 0.00 | 0.00 | 0.35 | 0.23 | 28 | Myeloid Cell |
| Lrrfip1 | 0.48 | 0.00 | 0.01 | 0.39 | 0.26 | 28 | Myeloid Cell |
| Mir682 | 0.28 | 0.00 | 0.01 | 0.89 | 0.85 | 28 | Myeloid Cell |
| Rps10-ps1 | 0.31 | 0.00 | 0.01 | 0.86 | 0.77 | 28 | Myeloid Cell |
| Gm5244 | 0.59 | 0.00 | 0.01 | 0.40 | 0.29 | 28 | Myeloid Cell |
| Rpl37a | 0.26 | 0.00 | 0.01 | 0.89 | 0.83 | 28 | Myeloid Cell |
| Ifitm2 | 0.42 | 0.00 | 0.01 | 0.46 | 0.32 | 28 | Myeloid Cell |
| Sub1 | 0.43 | 0.00 | 0.01 | 0.61 | 0.51 | 28 | Myeloid Cell |
| Ndufa6 | 0.43 | 0.00 | 0.01 | 0.54 | 0.43 | 28 | Myeloid Cell |
| Cbfa2t3 | 0.33 | 0.00 | 0.01 | 0.27 | 0.15 | 28 | Myeloid Cell |
| Cdc42se2 | 0.40 | 0.00 | 0.01 | 0.25 | 0.14 | 28 | Myeloid Cell |
| Txnip | 0.53 | 0.00 | 0.01 | 0.38 | 0.26 | 28 | Myeloid Cell |
| Rpl27-ps3 | 0.32 | 0.00 | 0.02 | 0.82 | 0.76 | 28 | Myeloid Cell |
| Rpl34 | 0.25 | 0.00 | 0.02 | 0.87 | 0.80 | 28 | Myeloid Cell |
| Tsc22d4 | 0.49 | 0.00 | 0.03 | 0.40 | 0.28 | 28 | Myeloid Cell |
| Adipor1 | 0.47 | 0.00 | 0.04 | 0.36 | 0.24 | 28 | Myeloid Cell |
| Gm17511 | 0.44 | 0.00 | 0.04 | 0.66 | 0.61 | 28 | Myeloid Cell |
| mt-Rnr1 | 0.32 | 0.00 | 0.05 | 0.58 | 0.47 | 28 | Myeloid Cell |
| Fgfr1op2 | 0.50 | 0.00 | 0.05 | 0.40 | 0.28 | 28 | Myeloid Cell |
| Lamp3 | 1.63 | 0.00 | 0.00 | 0.85 | 0.12 | 29 | AT2 Cell |
| Cxcl15 | 1.62 | 0.00 | 0.00 | 0.91 | 0.15 | 29 | AT2 Cell |
| Hc | 1.39 | 0.00 | 0.00 | 0.82 | 0.12 | 29 | AT2 Cell |
| Egfl6 | 1.30 | 0.00 | 0.00 | 0.69 | 0.08 | 29 | AT2 Cell |
| Sftpd | 1.50 | 0.00 | 0.00 | 0.96 | 0.16 | 29 | AT2 Cell |
| Chi3l1 | 1.36 | 0.00 | 0.00 | 0.78 | 0.11 | 29 | AT2 Cell |
| Sftpb | 1.58 | 0.00 | 0.00 | 0.98 | 0.18 | 29 | AT2 Cell |
| Wfdc2 | 1.53 | 0.00 | 0.00 | 0.91 | 0.17 | 29 | AT2 Cell |
| Slc34a2 | 1.47 | 0.00 | 0.00 | 0.91 | 0.16 | 29 | AT2 Cell |
| S100g | 1.33 | 0.00 | 0.00 | 0.80 | 0.12 | 29 | AT2 Cell |
| Sftpa1 | 1.75 | 0.00 | 0.00 | 0.97 | 0.21 | 29 | AT2 Cell |
| Bex4 | 1.31 | 0.00 | 0.00 | 0.81 | 0.13 | 29 | AT2 Cell |
| Bex2 | 1.53 | 0.00 | 0.00 | 0.70 | 0.10 | 29 | AT2 Cell |
| Sfta2 | 1.30 | 0.00 | 0.00 | 0.86 | 0.16 | 29 | AT2 Cell |
| Bex1 | 1.21 | 0.00 | 0.00 | 0.68 | 0.10 | 29 | AT2 Cell |
| Spc24 | 1.19 | 0.00 | 0.00 | 0.45 | 0.05 | 29 | AT2 Cell |
| Muc1 | 1.10 | 0.00 | 0.00 | 0.63 | 0.09 | 29 | AT2 Cell |
| Dram1 | 1.25 | 0.00 | 0.00 | 0.69 | 0.12 | 29 | AT2 Cell |
| Prc1 | 1.36 | 0.00 | 0.00 | 0.51 | 0.07 | 29 | AT2 Cell |
| Etv5 | 0.98 | 0.00 | 0.00 | 0.50 | 0.06 | 29 | AT2 Cell |
| D17H6S56E-5 | 1.85 | 0.00 | 0.00 | 0.61 | 0.10 | 29 | AT2 Cell |
| Tinag | 0.77 | 0.00 | 0.00 | 0.37 | 0.04 | 29 | AT2 Cell |
| Lpcat1 | 1.20 | 0.00 | 0.00 | 0.85 | 0.18 | 29 | AT2 Cell |
| Lgi3 | 0.98 | 0.00 | 0.00 | 0.56 | 0.08 | 29 | AT2 Cell |
| Mki67 | 1.43 | 0.00 | 0.00 | 0.67 | 0.12 | 29 | AT2 Cell |
| Pbk | 0.88 | 0.00 | 0.00 | 0.34 | 0.03 | 29 | AT2 Cell |
| Top2a | 1.42 | 0.00 | 0.00 | 0.63 | 0.11 | 29 | AT2 Cell |
| Cenpf | 1.45 | 0.00 | 0.00 | 0.52 | 0.08 | 29 | AT2 Cell |
| Cdc20 | 1.18 | 0.00 | 0.00 | 0.40 | 0.05 | 29 | AT2 Cell |
| Ppp1r14c | 1.04 | 0.00 | 0.00 | 0.63 | 0.11 | 29 | AT2 Cell |
| Birc5 | 1.04 | 0.00 | 0.00 | 0.45 | 0.06 | 29 | AT2 Cell |
| Epcam | 1.06 | 0.00 | 0.00 | 0.57 | 0.10 | 29 | AT2 Cell |
| Cbr2 | 0.88 | 0.00 | 0.00 | 0.78 | 0.17 | 29 | AT2 Cell |
| RP23-45G16.5 | 0.93 | 0.00 | 0.00 | 0.39 | 0.05 | 29 | AT2 Cell |
| Rbpjl | 0.79 | 0.00 | 0.00 | 0.45 | 0.06 | 29 | AT2 Cell |
| Cd74 | 0.81 | 0.00 | 0.00 | 0.65 | 0.13 | 29 | AT2 Cell |
| Tex15 | 0.67 | 0.00 | 0.00 | 0.29 | 0.03 | 29 | AT2 Cell |
| Spc25 | 0.98 | 0.00 | 0.00 | 0.39 | 0.05 | 29 | AT2 Cell |
| Lyz1 | 1.34 | 0.00 | 0.00 | 0.76 | 0.19 | 29 | AT2 Cell |
| Nusap1 | 1.00 | 0.00 | 0.00 | 0.41 | 0.06 | 29 | AT2 Cell |
| Il33 | 0.66 | 0.00 | 0.00 | 0.33 | 0.04 | 29 | AT2 Cell |
| Cldn18 | 0.68 | 0.00 | 0.00 | 0.77 | 0.18 | 29 | AT2 Cell |
| Scd1 | 0.96 | 0.00 | 0.00 | 0.52 | 0.09 | 29 | AT2 Cell |
| Pla2g1b | 0.69 | 0.00 | 0.00 | 0.37 | 0.05 | 29 | AT2 Cell |
| Napsa | 1.08 | 0.00 | 0.00 | 0.79 | 0.20 | 29 | AT2 Cell |
| Sdc4 | 1.11 | 0.00 | 0.00 | 0.82 | 0.23 | 29 | AT2 Cell |
| Irx3 | 0.91 | 0.00 | 0.00 | 0.46 | 0.08 | 29 | AT2 Cell |
| Ckap2 | 0.85 | 0.00 | 0.00 | 0.35 | 0.05 | 29 | AT2 Cell |
| Spink5 | 1.17 | 0.00 | 0.00 | 0.46 | 0.08 | 29 | AT2 Cell |
| Ces1d | 0.99 | 0.00 | 0.00 | 0.72 | 0.17 | 29 | AT2 Cell |
| Sftpc | 1.87 | 0.00 | 0.00 | 0.97 | 0.63 | 29 | AT2 Cell |
| Ptprf | 0.89 | 0.00 | 0.00 | 0.69 | 0.16 | 29 | AT2 Cell |
| Kif23 | 0.81 | 0.00 | 0.00 | 0.39 | 0.06 | 29 | AT2 Cell |
| Fabp5 | 0.96 | 0.00 | 0.00 | 0.61 | 0.13 | 29 | AT2 Cell |
| Lyz2 | 1.18 | 0.00 | 0.00 | 0.84 | 0.27 | 29 | AT2 Cell |
| H2-Aa | 0.53 | 0.00 | 0.00 | 0.48 | 0.08 | 29 | AT2 Cell |
| Kif11 | 0.70 | 0.00 | 0.00 | 0.32 | 0.04 | 29 | AT2 Cell |
| Hmmr | 0.76 | 0.00 | 0.00 | 0.34 | 0.05 | 29 | AT2 Cell |
| Abca3 | 0.87 | 0.00 | 0.00 | 0.58 | 0.12 | 29 | AT2 Cell |
| Lcn2 | 0.76 | 0.00 | 0.00 | 0.43 | 0.07 | 29 | AT2 Cell |
| Kif20b | 0.86 | 0.00 | 0.00 | 0.41 | 0.07 | 29 | AT2 Cell |
| Nkx2-1 | 0.75 | 0.00 | 0.00 | 0.52 | 0.10 | 29 | AT2 Cell |
| Ager | 0.28 | 0.00 | 0.00 | 0.73 | 0.17 | 29 | AT2 Cell |
| Car8 | 0.77 | 0.00 | 0.00 | 0.45 | 0.08 | 29 | AT2 Cell |
| Aldh1a7 | 0.77 | 0.00 | 0.00 | 0.34 | 0.05 | 29 | AT2 Cell |
| Ube2c | 1.23 | 0.00 | 0.00 | 0.35 | 0.05 | 29 | AT2 Cell |
| Cdh1 | 0.76 | 0.00 | 0.00 | 0.45 | 0.08 | 29 | AT2 Cell |
| Wbp5 | 1.17 | 0.00 | 0.00 | 0.96 | 0.48 | 29 | AT2 Cell |
| Alcam | 0.87 | 0.00 | 0.00 | 0.81 | 0.23 | 29 | AT2 Cell |
| 2810417H13Rik | 0.90 | 0.00 | 0.00 | 0.45 | 0.08 | 29 | AT2 Cell |
| Hist1h2ag | 1.24 | 0.00 | 0.00 | 0.53 | 0.12 | 29 | AT2 Cell |
| Cdca3 | 0.72 | 0.00 | 0.00 | 0.29 | 0.04 | 29 | AT2 Cell |
| Acsl4 | 0.98 | 0.00 | 0.00 | 0.54 | 0.12 | 29 | AT2 Cell |
| Sgol2 | 0.75 | 0.00 | 0.00 | 0.31 | 0.04 | 29 | AT2 Cell |
| Tc2n | 0.77 | 0.00 | 0.00 | 0.30 | 0.04 | 29 | AT2 Cell |
| Stmn1 | 1.30 | 0.00 | 0.00 | 0.73 | 0.23 | 29 | AT2 Cell |
| Trf | 0.84 | 0.00 | 0.00 | 0.49 | 0.10 | 29 | AT2 Cell |
| Apoc1 | 0.62 | 0.00 | 0.00 | 0.31 | 0.04 | 29 | AT2 Cell |
| Cdca8 | 0.74 | 0.00 | 0.00 | 0.34 | 0.05 | 29 | AT2 Cell |
| Hist1h2ak | 0.95 | 0.00 | 0.00 | 0.41 | 0.08 | 29 | AT2 Cell |
| Hist1h2ad | 1.13 | 0.00 | 0.00 | 0.49 | 0.11 | 29 | AT2 Cell |
| Npc2 | 0.89 | 0.00 | 0.00 | 0.98 | 0.54 | 29 | AT2 Cell |
| Knstrn | 0.74 | 0.00 | 0.00 | 0.29 | 0.04 | 29 | AT2 Cell |
| Cks2 | 1.08 | 0.00 | 0.00 | 0.42 | 0.08 | 29 | AT2 Cell |
| Mal | 0.66 | 0.00 | 0.00 | 0.32 | 0.05 | 29 | AT2 Cell |
| Atp6v1c2 | 0.61 | 0.00 | 0.00 | 0.33 | 0.05 | 29 | AT2 Cell |
| Hist1h2ah | 1.07 | 0.00 | 0.00 | 0.46 | 0.10 | 29 | AT2 Cell |
| Ccnb2 | 0.87 | 0.00 | 0.00 | 0.35 | 0.06 | 29 | AT2 Cell |
| Hist1h2ac | 1.13 | 0.00 | 0.00 | 0.48 | 0.11 | 29 | AT2 Cell |
| Hist1h2ab | 0.97 | 0.00 | 0.00 | 0.45 | 0.09 | 29 | AT2 Cell |
| Hist1h2ap | 1.40 | 0.00 | 0.00 | 0.57 | 0.15 | 29 | AT2 Cell |
| Atp1b1 | 0.67 | 0.00 | 0.00 | 0.70 | 0.19 | 29 | AT2 Cell |
| Hist1h2ai | 1.01 | 0.00 | 0.00 | 0.46 | 0.10 | 29 | AT2 Cell |
| Spint2 | 0.68 | 0.00 | 0.00 | 0.51 | 0.11 | 29 | AT2 Cell |
| Hist1h2an | 1.11 | 0.00 | 0.00 | 0.43 | 0.09 | 29 | AT2 Cell |
| Ngfrap1 | 0.99 | 0.00 | 0.00 | 0.71 | 0.23 | 29 | AT2 Cell |
| Ctsh | 0.81 | 0.00 | 0.00 | 0.71 | 0.21 | 29 | AT2 Cell |
| Hist1h2af | 1.09 | 0.00 | 0.00 | 0.43 | 0.09 | 29 | AT2 Cell |
| Smc2 | 1.02 | 0.00 | 0.00 | 0.54 | 0.14 | 29 | AT2 Cell |
| Cenpe | 0.87 | 0.00 | 0.00 | 0.40 | 0.08 | 29 | AT2 Cell |
| Cdk1 | 0.84 | 0.00 | 0.00 | 0.32 | 0.05 | 29 | AT2 Cell |
| Cldn7 | 0.64 | 0.00 | 0.00 | 0.41 | 0.08 | 29 | AT2 Cell |
| Ctsc | 0.85 | 0.00 | 0.00 | 0.67 | 0.19 | 29 | AT2 Cell |
| Sdc1 | 0.74 | 0.00 | 0.00 | 0.51 | 0.12 | 29 | AT2 Cell |
| Retnla | 2.55 | 0.00 | 0.00 | 0.45 | 0.11 | 29 | AT2 Cell |
| Pi4k2b | 0.64 | 0.00 | 0.00 | 0.46 | 0.10 | 29 | AT2 Cell |
| Hist1h2ao | 1.41 | 0.00 | 0.00 | 0.55 | 0.15 | 29 | AT2 Cell |
| Irx5 | 0.54 | 0.00 | 0.00 | 0.25 | 0.04 | 29 | AT2 Cell |
| Myo5c | 0.57 | 0.00 | 0.00 | 0.39 | 0.08 | 29 | AT2 Cell |
| Irx1 | 0.61 | 0.00 | 0.00 | 0.30 | 0.05 | 29 | AT2 Cell |
| Dlk1 | 0.65 | 0.00 | 0.00 | 0.37 | 0.07 | 29 | AT2 Cell |
| Ccna2 | 0.57 | 0.00 | 0.00 | 0.26 | 0.04 | 29 | AT2 Cell |
| Hist1h2ae | 0.97 | 0.00 | 0.00 | 0.41 | 0.09 | 29 | AT2 Cell |
| Kif15 | 0.68 | 0.00 | 0.00 | 0.29 | 0.05 | 29 | AT2 Cell |
| Slc39a8 | 0.57 | 0.00 | 0.00 | 0.38 | 0.07 | 29 | AT2 Cell |
| Mgst1 | 0.75 | 0.00 | 0.00 | 0.66 | 0.20 | 29 | AT2 Cell |
| Cadm1 | 0.63 | 0.00 | 0.00 | 0.54 | 0.14 | 29 | AT2 Cell |
| Cks1b | 0.78 | 0.00 | 0.00 | 0.41 | 0.10 | 29 | AT2 Cell |
| Tfrc | 0.57 | 0.00 | 0.00 | 0.31 | 0.06 | 29 | AT2 Cell |
| Tpx2 | 0.63 | 0.00 | 0.00 | 0.30 | 0.06 | 29 | AT2 Cell |
| Cldn3 | 0.55 | 0.00 | 0.00 | 0.49 | 0.12 | 29 | AT2 Cell |
| Mecom | 0.63 | 0.00 | 0.00 | 0.43 | 0.10 | 29 | AT2 Cell |
| Casc5 | 0.57 | 0.00 | 0.00 | 0.27 | 0.05 | 29 | AT2 Cell |
| Ssx2ip | 0.56 | 0.00 | 0.00 | 0.26 | 0.04 | 29 | AT2 Cell |
| Ank3 | 0.61 | 0.00 | 0.00 | 0.36 | 0.08 | 29 | AT2 Cell |
| Incenp | 0.70 | 0.00 | 0.00 | 0.37 | 0.08 | 29 | AT2 Cell |
| Rrm2 | 0.68 | 0.00 | 0.00 | 0.27 | 0.05 | 29 | AT2 Cell |
| Slc12a2 | 0.63 | 0.00 | 0.00 | 0.53 | 0.15 | 29 | AT2 Cell |
| H2afx | 0.74 | 0.00 | 0.00 | 0.40 | 0.10 | 29 | AT2 Cell |
| Fasn | 0.61 | 0.00 | 0.00 | 0.35 | 0.08 | 29 | AT2 Cell |
| Fbxo5 | 0.53 | 0.00 | 0.00 | 0.26 | 0.05 | 29 | AT2 Cell |
| Cmtm8 | 0.64 | 0.00 | 0.00 | 0.47 | 0.13 | 29 | AT2 Cell |
| Oat | 0.65 | 0.00 | 0.00 | 0.47 | 0.13 | 29 | AT2 Cell |
| Mbip | 0.57 | 0.00 | 0.00 | 0.45 | 0.11 | 29 | AT2 Cell |
| Tacc3 | 0.59 | 0.00 | 0.00 | 0.29 | 0.06 | 29 | AT2 Cell |
| Mt1 | 0.54 | 0.00 | 0.00 | 0.54 | 0.16 | 29 | AT2 Cell |
| 2700094K13Rik | 0.68 | 0.00 | 0.00 | 0.54 | 0.16 | 29 | AT2 Cell |
| Ldhb | 0.55 | 0.00 | 0.00 | 0.35 | 0.08 | 29 | AT2 Cell |
| Diap3 | 0.62 | 0.00 | 0.00 | 0.30 | 0.06 | 29 | AT2 Cell |
| Atp8a1 | 0.72 | 0.00 | 0.00 | 0.51 | 0.15 | 29 | AT2 Cell |
| Avpi1 | 0.51 | 0.00 | 0.00 | 0.37 | 0.09 | 29 | AT2 Cell |
| Snx7 | 0.49 | 0.00 | 0.00 | 0.32 | 0.07 | 29 | AT2 Cell |
| Ppp1r9a | 0.61 | 0.00 | 0.00 | 0.50 | 0.14 | 29 | AT2 Cell |
| Fgfr2 | 0.67 | 0.00 | 0.00 | 0.32 | 0.07 | 29 | AT2 Cell |
| Snx25 | 0.66 | 0.00 | 0.00 | 0.29 | 0.06 | 29 | AT2 Cell |
| Nucb2 | 0.67 | 0.00 | 0.00 | 0.52 | 0.16 | 29 | AT2 Cell |
| Gclc | 0.72 | 0.00 | 0.00 | 0.35 | 0.09 | 29 | AT2 Cell |
| Id2 | 0.78 | 0.00 | 0.00 | 0.75 | 0.33 | 29 | AT2 Cell |
| Rab27b | 0.53 | 0.00 | 0.00 | 0.31 | 0.07 | 29 | AT2 Cell |
| Krt8 | 0.43 | 0.00 | 0.00 | 0.34 | 0.08 | 29 | AT2 Cell |
| Mt2 | 0.39 | 0.00 | 0.00 | 0.31 | 0.07 | 29 | AT2 Cell |
| Hmgb2 | 0.92 | 0.00 | 0.00 | 0.79 | 0.43 | 29 | AT2 Cell |
| Hirip3 | 0.66 | 0.00 | 0.00 | 0.35 | 0.09 | 29 | AT2 Cell |
| Rnase4 | 0.65 | 0.00 | 0.00 | 0.54 | 0.18 | 29 | AT2 Cell |
| Ccdc34 | 0.59 | 0.00 | 0.00 | 0.46 | 0.14 | 29 | AT2 Cell |
| Mid1ip1 | 0.66 | 0.00 | 0.00 | 0.46 | 0.14 | 29 | AT2 Cell |
| Cxadr | 0.49 | 0.00 | 0.00 | 0.28 | 0.06 | 29 | AT2 Cell |
| Ptplad1 | 0.48 | 0.00 | 0.00 | 0.36 | 0.09 | 29 | AT2 Cell |
| Cebpa | 0.39 | 0.00 | 0.00 | 0.26 | 0.05 | 29 | AT2 Cell |
| Tcea3 | 0.45 | 0.00 | 0.00 | 0.32 | 0.08 | 29 | AT2 Cell |
| Rai14 | 0.55 | 0.00 | 0.00 | 0.39 | 0.11 | 29 | AT2 Cell |
| Gm10282 | 0.83 | 0.00 | 0.00 | 0.76 | 0.40 | 29 | AT2 Cell |
| Hmgn2 | 0.78 | 0.00 | 0.00 | 0.81 | 0.43 | 29 | AT2 Cell |
| Me1 | 0.61 | 0.00 | 0.00 | 0.25 | 0.06 | 29 | AT2 Cell |
| St6galnac2 | 0.45 | 0.00 | 0.00 | 0.25 | 0.06 | 29 | AT2 Cell |
| Ptgs1 | 0.50 | 0.00 | 0.00 | 0.37 | 0.10 | 29 | AT2 Cell |
| Adk | 0.45 | 0.00 | 0.00 | 0.40 | 0.12 | 29 | AT2 Cell |
| Gas6 | 0.49 | 0.00 | 0.00 | 0.46 | 0.15 | 29 | AT2 Cell |
| Cenpa | 0.56 | 0.00 | 0.00 | 0.28 | 0.07 | 29 | AT2 Cell |
| Nucks1 | 0.71 | 0.00 | 0.00 | 0.65 | 0.28 | 29 | AT2 Cell |
| Rangap1 | 0.51 | 0.00 | 0.00 | 0.30 | 0.08 | 29 | AT2 Cell |
| Abcd3 | 0.62 | 0.00 | 0.00 | 0.51 | 0.19 | 29 | AT2 Cell |
| Brd7 | 0.62 | 0.00 | 0.00 | 0.62 | 0.26 | 29 | AT2 Cell |
| Prnp | 0.53 | 0.00 | 0.00 | 0.39 | 0.12 | 29 | AT2 Cell |
| H2afz | 0.87 | 0.00 | 0.00 | 0.70 | 0.40 | 29 | AT2 Cell |
| Atad2 | 0.63 | 0.00 | 0.00 | 0.30 | 0.08 | 29 | AT2 Cell |
| Tmpo | 0.64 | 0.00 | 0.00 | 0.44 | 0.15 | 29 | AT2 Cell |
| Mthfd1 | 0.39 | 0.00 | 0.00 | 0.31 | 0.08 | 29 | AT2 Cell |
| H1f0 | 0.64 | 0.00 | 0.00 | 0.62 | 0.27 | 29 | AT2 Cell |
| Acot7 | 0.46 | 0.00 | 0.00 | 0.26 | 0.06 | 29 | AT2 Cell |
| Wwc1 | 0.44 | 0.00 | 0.00 | 0.26 | 0.06 | 29 | AT2 Cell |
| Zdhhc3 | 0.58 | 0.00 | 0.00 | 0.45 | 0.16 | 29 | AT2 Cell |
| Ptma | 0.50 | 0.00 | 0.00 | 0.98 | 0.91 | 29 | AT2 Cell |
| Tspan8 | 0.42 | 0.00 | 0.00 | 0.39 | 0.12 | 29 | AT2 Cell |
| Acsl5 | 0.44 | 0.00 | 0.00 | 0.36 | 0.11 | 29 | AT2 Cell |
| Rad21 | 0.59 | 0.00 | 0.00 | 0.56 | 0.23 | 29 | AT2 Cell |
| Atp11a | 0.55 | 0.00 | 0.00 | 0.36 | 0.11 | 29 | AT2 Cell |
| Smc4 | 0.70 | 0.00 | 0.00 | 0.68 | 0.35 | 29 | AT2 Cell |
| Cat | 0.46 | 0.00 | 0.00 | 0.39 | 0.12 | 29 | AT2 Cell |
| Dcxr | 0.33 | 0.00 | 0.00 | 0.36 | 0.11 | 29 | AT2 Cell |
| Neat1 | 0.40 | 0.00 | 0.00 | 0.65 | 0.27 | 29 | AT2 Cell |
| Scd2 | 0.54 | 0.00 | 0.00 | 0.54 | 0.21 | 29 | AT2 Cell |
| Pkp4 | 0.42 | 0.00 | 0.00 | 0.30 | 0.08 | 29 | AT2 Cell |
| Tubb4b | 0.58 | 0.00 | 0.00 | 0.48 | 0.19 | 29 | AT2 Cell |
| Hmgn1 | 0.57 | 0.00 | 0.00 | 0.80 | 0.45 | 29 | AT2 Cell |
| Cenpq | 0.31 | 0.00 | 0.00 | 0.26 | 0.07 | 29 | AT2 Cell |
| Taok3 | 0.43 | 0.00 | 0.00 | 0.50 | 0.19 | 29 | AT2 Cell |
| Dbi | 0.59 | 0.00 | 0.00 | 0.73 | 0.37 | 29 | AT2 Cell |
| Gmnn | 0.52 | 0.00 | 0.00 | 0.26 | 0.07 | 29 | AT2 Cell |
| Chchd10 | 0.29 | 0.00 | 0.00 | 0.41 | 0.13 | 29 | AT2 Cell |
| Whsc1 | 0.50 | 0.00 | 0.00 | 0.40 | 0.14 | 29 | AT2 Cell |
| Arl6ip1 | 0.87 | 0.00 | 0.00 | 0.83 | 0.52 | 29 | AT2 Cell |
| Serpinb6b | 0.34 | 0.00 | 0.00 | 0.47 | 0.17 | 29 | AT2 Cell |
| Dynll2 | 0.49 | 0.00 | 0.00 | 0.36 | 0.12 | 29 | AT2 Cell |
| Tuba1b | 0.65 | 0.00 | 0.00 | 0.78 | 0.45 | 29 | AT2 Cell |
| Anxa4 | 0.41 | 0.00 | 0.00 | 0.31 | 0.10 | 29 | AT2 Cell |
| Fkbp2 | 0.53 | 0.00 | 0.00 | 0.46 | 0.18 | 29 | AT2 Cell |
| Scp2 | 0.48 | 0.00 | 0.00 | 0.69 | 0.33 | 29 | AT2 Cell |
| Exosc7 | 0.42 | 0.00 | 0.00 | 0.38 | 0.13 | 29 | AT2 Cell |
| H2afv | 0.57 | 0.00 | 0.00 | 0.60 | 0.28 | 29 | AT2 Cell |
| Cystm1 | 0.31 | 0.00 | 0.00 | 0.28 | 0.08 | 29 | AT2 Cell |
| Lig1 | 0.51 | 0.00 | 0.00 | 0.30 | 0.10 | 29 | AT2 Cell |
| Chka | 0.34 | 0.00 | 0.00 | 0.32 | 0.10 | 29 | AT2 Cell |
| Dpp4 | 0.34 | 0.00 | 0.00 | 0.37 | 0.13 | 29 | AT2 Cell |
| Cbx6 | 0.38 | 0.00 | 0.00 | 0.32 | 0.11 | 29 | AT2 Cell |
| Phldb2 | 0.40 | 0.00 | 0.00 | 0.64 | 0.29 | 29 | AT2 Cell |
| Dek | 0.71 | 0.00 | 0.00 | 0.77 | 0.50 | 29 | AT2 Cell |
| Gja1 | 0.39 | 0.00 | 0.00 | 0.29 | 0.09 | 29 | AT2 Cell |
| Prrg3 | 0.33 | 0.00 | 0.00 | 0.26 | 0.08 | 29 | AT2 Cell |
| Tgoln1 | 0.46 | 0.00 | 0.00 | 0.46 | 0.19 | 29 | AT2 Cell |
| Trim2 | 0.36 | 0.00 | 0.00 | 0.26 | 0.08 | 29 | AT2 Cell |
| Ezh2 | 0.47 | 0.00 | 0.00 | 0.46 | 0.19 | 29 | AT2 Cell |
| Hmga1-rs1 | 0.34 | 0.00 | 0.00 | 0.25 | 0.08 | 29 | AT2 Cell |
| Iah1 | 0.41 | 0.00 | 0.00 | 0.35 | 0.12 | 29 | AT2 Cell |
| Sh3d19 | 0.33 | 0.00 | 0.00 | 0.29 | 0.09 | 29 | AT2 Cell |
| Fkbp4 | 0.53 | 0.00 | 0.00 | 0.39 | 0.15 | 29 | AT2 Cell |
| Uhrf1bp1l | 0.33 | 0.00 | 0.00 | 0.30 | 0.10 | 29 | AT2 Cell |
| Apoa1bp | 0.35 | 0.00 | 0.00 | 0.39 | 0.15 | 29 | AT2 Cell |
| Tubb4b-ps1 | 0.35 | 0.00 | 0.00 | 0.28 | 0.09 | 29 | AT2 Cell |
| Pon3 | 0.35 | 0.00 | 0.00 | 0.30 | 0.10 | 29 | AT2 Cell |
| Pcna | 0.58 | 0.00 | 0.00 | 0.42 | 0.18 | 29 | AT2 Cell |
| Rnaseh2c | 0.37 | 0.00 | 0.00 | 0.31 | 0.11 | 29 | AT2 Cell |
| Lmnb1 | 0.34 | 0.00 | 0.00 | 0.27 | 0.09 | 29 | AT2 Cell |
| Hmga1 | 0.32 | 0.00 | 0.00 | 0.25 | 0.08 | 29 | AT2 Cell |
| Dnajc9 | 0.39 | 0.00 | 0.00 | 0.34 | 0.12 | 29 | AT2 Cell |
| Tmem243 | 0.31 | 0.00 | 0.00 | 0.25 | 0.08 | 29 | AT2 Cell |
| Tmem50b | 0.38 | 0.00 | 0.00 | 0.25 | 0.08 | 29 | AT2 Cell |
| Pcna-ps2 | 0.57 | 0.00 | 0.00 | 0.41 | 0.18 | 29 | AT2 Cell |
| Etfb | 0.44 | 0.00 | 0.00 | 0.58 | 0.28 | 29 | AT2 Cell |
| Usp1 | 0.46 | 0.00 | 0.00 | 0.38 | 0.16 | 29 | AT2 Cell |
| Krt18 | 0.27 | 0.00 | 0.00 | 0.25 | 0.08 | 29 | AT2 Cell |
| Gde1 | 0.33 | 0.00 | 0.00 | 0.38 | 0.15 | 29 | AT2 Cell |
| Soat1 | 0.32 | 0.00 | 0.00 | 0.30 | 0.11 | 29 | AT2 Cell |
| Iqgap1 | 0.45 | 0.00 | 0.00 | 0.77 | 0.45 | 29 | AT2 Cell |
| Atp13a3 | 0.28 | 0.00 | 0.00 | 0.28 | 0.10 | 29 | AT2 Cell |
| Hist1h1e | 0.75 | 0.00 | 0.00 | 0.39 | 0.17 | 29 | AT2 Cell |
| Dut | 0.41 | 0.00 | 0.00 | 0.26 | 0.09 | 29 | AT2 Cell |
| H2afj | 0.35 | 0.00 | 0.00 | 0.64 | 0.32 | 29 | AT2 Cell |
| Mtch1 | 0.44 | 0.00 | 0.00 | 0.36 | 0.15 | 29 | AT2 Cell |
| Oxct1 | 0.40 | 0.00 | 0.00 | 0.37 | 0.15 | 29 | AT2 Cell |
| Nudc | 0.33 | 0.00 | 0.00 | 0.47 | 0.21 | 29 | AT2 Cell |
| Gm26924 | 0.64 | 0.00 | 0.00 | 0.84 | 0.57 | 29 | AT2 Cell |
| Ran | 0.46 | 0.00 | 0.00 | 0.69 | 0.38 | 29 | AT2 Cell |
| Hmgn5 | 0.36 | 0.00 | 0.00 | 0.53 | 0.26 | 29 | AT2 Cell |
| Cd63 | 0.29 | 0.00 | 0.00 | 0.62 | 0.30 | 29 | AT2 Cell |
| Tmem30a | 0.39 | 0.00 | 0.00 | 0.57 | 0.28 | 29 | AT2 Cell |
| H3f3b | 0.32 | 0.00 | 0.00 | 0.99 | 0.89 | 29 | AT2 Cell |
| Ckap5 | 0.37 | 0.00 | 0.00 | 0.30 | 0.12 | 29 | AT2 Cell |
| Cpne3 | 0.41 | 0.00 | 0.00 | 0.41 | 0.18 | 29 | AT2 Cell |
| Sae1 | 0.28 | 0.00 | 0.00 | 0.30 | 0.11 | 29 | AT2 Cell |
| Arfip1 | 0.30 | 0.00 | 0.00 | 0.32 | 0.13 | 29 | AT2 Cell |
| Nckap5 | 0.29 | 0.00 | 0.00 | 0.33 | 0.13 | 29 | AT2 Cell |
| Mettl7a1 | 0.28 | 0.00 | 0.00 | 0.48 | 0.22 | 29 | AT2 Cell |
| Siva1 | 0.42 | 0.00 | 0.00 | 0.35 | 0.15 | 29 | AT2 Cell |
| Anp32b | 0.41 | 0.00 | 0.00 | 0.81 | 0.50 | 29 | AT2 Cell |
| Syne2 | 0.32 | 0.00 | 0.00 | 0.35 | 0.14 | 29 | AT2 Cell |
| Itga9 | 0.30 | 0.00 | 0.00 | 0.26 | 0.10 | 29 | AT2 Cell |
| Cbx5 | 0.31 | 0.00 | 0.00 | 0.53 | 0.26 | 29 | AT2 Cell |
| Ndufc2 | 0.34 | 0.00 | 0.00 | 0.70 | 0.39 | 29 | AT2 Cell |
| Azin1 | 0.27 | 0.00 | 0.00 | 0.33 | 0.13 | 29 | AT2 Cell |
| Dstn | 0.38 | 0.00 | 0.00 | 0.79 | 0.51 | 29 | AT2 Cell |
| Prdx4 | 0.31 | 0.00 | 0.00 | 0.37 | 0.16 | 29 | AT2 Cell |
| Baz1b | 0.26 | 0.00 | 0.00 | 0.54 | 0.27 | 29 | AT2 Cell |
| Cited2 | 0.31 | 0.00 | 0.00 | 0.27 | 0.10 | 29 | AT2 Cell |
| Nfix | 0.27 | 0.00 | 0.00 | 0.39 | 0.17 | 29 | AT2 Cell |
| Gm6104 | 0.30 | 0.00 | 0.00 | 0.31 | 0.13 | 29 | AT2 Cell |
| Enpep | 0.29 | 0.00 | 0.00 | 0.27 | 0.10 | 29 | AT2 Cell |
| Wls | 0.30 | 0.00 | 0.00 | 0.47 | 0.23 | 29 | AT2 Cell |
| Hist1h1d | 0.51 | 0.00 | 0.00 | 0.26 | 0.10 | 29 | AT2 Cell |
| Atp2a2 | 0.33 | 0.00 | 0.00 | 0.31 | 0.13 | 29 | AT2 Cell |
| Tmem245 | 0.29 | 0.00 | 0.00 | 0.26 | 0.10 | 29 | AT2 Cell |
| Anapc13 | 0.31 | 0.00 | 0.00 | 0.45 | 0.21 | 29 | AT2 Cell |
| Bsg | 0.33 | 0.00 | 0.00 | 0.63 | 0.36 | 29 | AT2 Cell |
| Hdgf | 0.31 | 0.00 | 0.00 | 0.50 | 0.25 | 29 | AT2 Cell |
| Hmgb3 | 0.32 | 0.00 | 0.00 | 0.34 | 0.15 | 29 | AT2 Cell |
| Fam111a | 0.27 | 0.00 | 0.00 | 0.35 | 0.15 | 29 | AT2 Cell |
| Hjurp | 0.29 | 0.00 | 0.00 | 0.34 | 0.15 | 29 | AT2 Cell |
| Tspo | 0.26 | 0.00 | 0.00 | 0.51 | 0.26 | 29 | AT2 Cell |
| Suz12 | 0.26 | 0.00 | 0.00 | 0.31 | 0.13 | 29 | AT2 Cell |
| Nmt2 | 0.31 | 0.00 | 0.00 | 0.26 | 0.10 | 29 | AT2 Cell |
| Emb | 0.25 | 0.00 | 0.00 | 0.28 | 0.11 | 29 | AT2 Cell |
| Ctnnb1 | 0.29 | 0.00 | 0.00 | 0.63 | 0.35 | 29 | AT2 Cell |
| Sod1 | 0.31 | 0.00 | 0.00 | 0.75 | 0.47 | 29 | AT2 Cell |
| Hsp90aa1 | 0.27 | 0.00 | 0.00 | 0.75 | 0.48 | 29 | AT2 Cell |
| Cep57 | 0.31 | 0.00 | 0.00 | 0.27 | 0.11 | 29 | AT2 Cell |
| Tuba1c | 0.41 | 0.00 | 0.00 | 0.57 | 0.32 | 29 | AT2 Cell |
| Stard3nl | 0.29 | 0.00 | 0.00 | 0.28 | 0.12 | 29 | AT2 Cell |
| Tubb5 | 0.43 | 0.00 | 0.00 | 0.70 | 0.44 | 29 | AT2 Cell |
| Tmbim6 | 0.26 | 0.00 | 0.00 | 0.67 | 0.38 | 29 | AT2 Cell |
| Usp33 | 0.32 | 0.00 | 0.00 | 0.27 | 0.11 | 29 | AT2 Cell |
| Dapk1 | 0.25 | 0.00 | 0.00 | 0.27 | 0.11 | 29 | AT2 Cell |
| Gm9790 | 0.34 | 0.00 | 0.00 | 0.36 | 0.17 | 29 | AT2 Cell |
| 1810037I17Rik | 0.32 | 0.00 | 0.00 | 0.58 | 0.32 | 29 | AT2 Cell |
| Smchd1 | 0.30 | 0.00 | 0.00 | 0.35 | 0.16 | 29 | AT2 Cell |
| Pigyl | 0.30 | 0.00 | 0.00 | 0.25 | 0.11 | 29 | AT2 Cell |
| Txn1 | 0.31 | 0.00 | 0.00 | 0.78 | 0.50 | 29 | AT2 Cell |
| Idh2 | 0.26 | 0.00 | 0.00 | 0.40 | 0.19 | 29 | AT2 Cell |
| Ckap4 | 0.25 | 0.00 | 0.00 | 0.37 | 0.18 | 29 | AT2 Cell |
| Mrpl14 | 0.30 | 0.00 | 0.00 | 0.33 | 0.15 | 29 | AT2 Cell |
| Vamp8 | 0.26 | 0.00 | 0.00 | 0.62 | 0.35 | 29 | AT2 Cell |
| Atp5e | 0.29 | 0.00 | 0.00 | 0.79 | 0.50 | 29 | AT2 Cell |
| Arf6 | 0.26 | 0.00 | 0.00 | 0.39 | 0.19 | 29 | AT2 Cell |
| Bub3 | 0.29 | 0.00 | 0.00 | 0.31 | 0.14 | 29 | AT2 Cell |
| Pfdn1 | 0.28 | 0.00 | 0.00 | 0.48 | 0.26 | 29 | AT2 Cell |
| Dtymk | 0.31 | 0.00 | 0.00 | 0.30 | 0.14 | 29 | AT2 Cell |
| Slbp | 0.33 | 0.00 | 0.00 | 0.31 | 0.15 | 29 | AT2 Cell |
| Bag1 | 0.34 | 0.00 | 0.00 | 0.44 | 0.24 | 29 | AT2 Cell |
| Dmtf1 | 0.26 | 0.00 | 0.00 | 0.26 | 0.11 | 29 | AT2 Cell |
| Atxn10 | 0.27 | 0.00 | 0.00 | 0.39 | 0.19 | 29 | AT2 Cell |
| Usp47 | 0.30 | 0.00 | 0.00 | 0.27 | 0.12 | 29 | AT2 Cell |
| Prdx1 | 0.29 | 0.00 | 0.00 | 0.74 | 0.50 | 29 | AT2 Cell |
| Tead1 | 0.30 | 0.00 | 0.00 | 0.29 | 0.14 | 29 | AT2 Cell |
| Gm5641 | 0.30 | 0.00 | 0.00 | 0.60 | 0.37 | 29 | AT2 Cell |
| Lrrcc1 | 0.25 | 0.00 | 0.00 | 0.26 | 0.12 | 29 | AT2 Cell |
| Ssrp1 | 0.28 | 0.00 | 0.00 | 0.40 | 0.21 | 29 | AT2 Cell |
| Calm2 | 0.33 | 0.00 | 0.00 | 0.85 | 0.64 | 29 | AT2 Cell |
| Larp7 | 0.26 | 0.00 | 0.00 | 0.34 | 0.17 | 29 | AT2 Cell |
| Rif1 | 0.28 | 0.00 | 0.00 | 0.32 | 0.17 | 29 | AT2 Cell |
| Mrpl52 | 0.25 | 0.00 | 0.00 | 0.54 | 0.33 | 29 | AT2 Cell |
| Ranbp1 | 0.29 | 0.00 | 0.00 | 0.49 | 0.29 | 29 | AT2 Cell |
| Hnrnpab | 0.27 | 0.00 | 0.01 | 0.53 | 0.33 | 29 | AT2 Cell |
| Higd1a | 0.25 | 0.00 | 0.01 | 0.34 | 0.18 | 29 | AT2 Cell |
| Pcnt | 0.29 | 0.00 | 0.01 | 0.28 | 0.14 | 29 | AT2 Cell |
| H2afy | 0.26 | 0.00 | 0.02 | 0.36 | 0.20 | 29 | AT2 Cell |
| Mfap4 | 1.08 | 0.00 | 0.00 | 0.94 | 0.25 | 30 | Myofibroblast |
| Tcf21 | 0.88 | 0.00 | 0.00 | 0.82 | 0.17 | 30 | Myofibroblast |
| Dpt | 1.06 | 0.00 | 0.00 | 0.72 | 0.18 | 30 | Myofibroblast |
| Adh1 | 0.94 | 0.00 | 0.00 | 0.71 | 0.17 | 30 | Myofibroblast |
| Adamts17 | 0.66 | 0.00 | 0.00 | 0.30 | 0.04 | 30 | Myofibroblast |
| Maf | 0.88 | 0.00 | 0.00 | 0.74 | 0.19 | 30 | Myofibroblast |
| Cp | 0.90 | 0.00 | 0.00 | 0.58 | 0.13 | 30 | Myofibroblast |
| Ogn | 0.79 | 0.00 | 0.00 | 0.60 | 0.13 | 30 | Myofibroblast |
| Limch1 | 0.88 | 0.00 | 0.00 | 0.89 | 0.31 | 30 | Myofibroblast |
| Col13a1 | 0.76 | 0.00 | 0.00 | 0.49 | 0.10 | 30 | Myofibroblast |
| Meox2 | 0.74 | 0.00 | 0.00 | 0.42 | 0.08 | 30 | Myofibroblast |
| G0s2 | 0.79 | 0.00 | 0.00 | 0.57 | 0.13 | 30 | Myofibroblast |
| Cdh11 | 0.80 | 0.00 | 0.00 | 0.70 | 0.19 | 30 | Myofibroblast |
| Macf1 | 0.96 | 0.00 | 0.00 | 0.98 | 0.52 | 30 | Myofibroblast |
| Fhl1 | 0.79 | 0.00 | 0.00 | 0.85 | 0.29 | 30 | Myofibroblast |
| Fn1 | 0.84 | 0.00 | 0.00 | 0.76 | 0.23 | 30 | Myofibroblast |
| Rbp1 | 0.81 | 0.00 | 0.00 | 0.68 | 0.20 | 30 | Myofibroblast |
| Fmo2 | 0.96 | 0.00 | 0.00 | 0.61 | 0.17 | 30 | Myofibroblast |
| Lbh | 0.66 | 0.00 | 0.00 | 0.68 | 0.19 | 30 | Myofibroblast |
| Akap12 | 0.90 | 0.00 | 0.00 | 0.73 | 0.24 | 30 | Myofibroblast |
| Gyg | 0.74 | 0.00 | 0.00 | 0.65 | 0.19 | 30 | Myofibroblast |
| 6030408B16Rik | 0.71 | 0.00 | 0.00 | 0.28 | 0.05 | 30 | Myofibroblast |
| Pcolce2 | 0.59 | 0.00 | 0.00 | 0.42 | 0.09 | 30 | Myofibroblast |
| Ablim3 | 0.53 | 0.00 | 0.00 | 0.29 | 0.05 | 30 | Myofibroblast |
| Lyve1 | 1.05 | 0.00 | 0.00 | 0.68 | 0.22 | 30 | Myofibroblast |
| Mgp | 0.55 | 0.00 | 0.00 | 0.88 | 0.32 | 30 | Myofibroblast |
| Ptprd | 0.78 | 0.00 | 0.00 | 0.47 | 0.12 | 30 | Myofibroblast |
| Npnt | 0.81 | 0.00 | 0.00 | 0.72 | 0.24 | 30 | Myofibroblast |
| Figf | 0.50 | 0.00 | 0.00 | 0.35 | 0.07 | 30 | Myofibroblast |
| Nebl | 0.69 | 0.00 | 0.00 | 0.43 | 0.10 | 30 | Myofibroblast |
| 4-Sep | 0.65 | 0.00 | 0.00 | 0.71 | 0.23 | 30 | Myofibroblast |
| Ramp2 | 0.65 | 0.00 | 0.00 | 0.80 | 0.28 | 30 | Myofibroblast |
| Plac9b | 0.67 | 0.00 | 0.00 | 0.81 | 0.30 | 30 | Myofibroblast |
| Inmt | 0.63 | 0.00 | 0.00 | 0.29 | 0.06 | 30 | Myofibroblast |
| Clec3b | 0.53 | 0.00 | 0.00 | 0.33 | 0.07 | 30 | Myofibroblast |
| Spon1 | 0.57 | 0.00 | 0.00 | 0.33 | 0.07 | 30 | Myofibroblast |
| Gpihbp1 | 0.62 | 0.00 | 0.00 | 0.56 | 0.16 | 30 | Myofibroblast |
| Calcrl | 0.50 | 0.00 | 0.00 | 0.75 | 0.25 | 30 | Myofibroblast |
| Slc38a5 | 0.58 | 0.00 | 0.00 | 0.29 | 0.06 | 30 | Myofibroblast |
| Plac9a | 0.62 | 0.00 | 0.00 | 0.79 | 0.30 | 30 | Myofibroblast |
| Col1a2 | 0.49 | 0.00 | 0.00 | 0.93 | 0.39 | 30 | Myofibroblast |
| Fgf7 | 0.52 | 0.00 | 0.00 | 0.26 | 0.05 | 30 | Myofibroblast |
| Enpep | 0.61 | 0.00 | 0.00 | 0.40 | 0.10 | 30 | Myofibroblast |
| Tmem100 | 0.43 | 0.00 | 0.00 | 0.82 | 0.30 | 30 | Myofibroblast |
| Cldn5 | 0.58 | 0.00 | 0.00 | 0.67 | 0.22 | 30 | Myofibroblast |
| Gsn | 0.64 | 0.00 | 0.00 | 0.67 | 0.25 | 30 | Myofibroblast |
| Rap2a | 0.69 | 0.00 | 0.00 | 0.51 | 0.16 | 30 | Myofibroblast |
| Nr2f2 | 0.62 | 0.00 | 0.00 | 0.62 | 0.21 | 30 | Myofibroblast |
| Serping1 | 0.48 | 0.00 | 0.00 | 0.46 | 0.12 | 30 | Myofibroblast |
| Cped1 | 0.57 | 0.00 | 0.00 | 0.38 | 0.10 | 30 | Myofibroblast |
| Myh10 | 0.61 | 0.00 | 0.00 | 0.65 | 0.23 | 30 | Myofibroblast |
| Epas1 | 0.57 | 0.00 | 0.00 | 0.70 | 0.25 | 30 | Myofibroblast |
| Gpx3 | 0.66 | 0.00 | 0.00 | 0.47 | 0.14 | 30 | Myofibroblast |
| Ifitm1 | 0.45 | 0.00 | 0.00 | 0.48 | 0.14 | 30 | Myofibroblast |
| Egfl7 | 0.40 | 0.00 | 0.00 | 0.78 | 0.28 | 30 | Myofibroblast |
| Atp1a2 | 0.55 | 0.00 | 0.00 | 0.38 | 0.10 | 30 | Myofibroblast |
| Fblim1 | 0.65 | 0.00 | 0.00 | 0.51 | 0.16 | 30 | Myofibroblast |
| Acvrl1 | 0.50 | 0.00 | 0.00 | 0.55 | 0.18 | 30 | Myofibroblast |
| Scn7a | 0.48 | 0.00 | 0.00 | 0.73 | 0.30 | 30 | Myofibroblast |
| Plxdc2 | 0.71 | 0.00 | 0.00 | 0.42 | 0.13 | 30 | Myofibroblast |
| Sema6a | 0.53 | 0.00 | 0.00 | 0.41 | 0.12 | 30 | Myofibroblast |
| Ednra | 0.48 | 0.00 | 0.00 | 0.34 | 0.09 | 30 | Myofibroblast |
| Col1a1 | 0.41 | 0.00 | 0.00 | 0.70 | 0.26 | 30 | Myofibroblast |
| Ptprb | 0.48 | 0.00 | 0.00 | 0.62 | 0.22 | 30 | Myofibroblast |
| Sh3bgrl | 0.65 | 0.00 | 0.00 | 0.84 | 0.45 | 30 | Myofibroblast |
| Sned1 | 0.46 | 0.00 | 0.00 | 0.26 | 0.06 | 30 | Myofibroblast |
| Ly6a | 0.55 | 0.00 | 0.00 | 0.57 | 0.19 | 30 | Myofibroblast |
| Cd93 | 0.44 | 0.00 | 0.00 | 0.63 | 0.22 | 30 | Myofibroblast |
| Thbd | 0.62 | 0.00 | 0.00 | 0.64 | 0.25 | 30 | Myofibroblast |
| Tns1 | 0.54 | 0.00 | 0.00 | 0.62 | 0.23 | 30 | Myofibroblast |
| Eva1b | 0.58 | 0.00 | 0.00 | 0.55 | 0.20 | 30 | Myofibroblast |
| Hip1 | 0.62 | 0.00 | 0.00 | 0.48 | 0.16 | 30 | Myofibroblast |
| Slit2 | 0.59 | 0.00 | 0.00 | 0.30 | 0.08 | 30 | Myofibroblast |
| Gpc3 | 0.58 | 0.00 | 0.00 | 0.49 | 0.17 | 30 | Myofibroblast |
| Ppap2b | 0.65 | 0.00 | 0.00 | 0.62 | 0.24 | 30 | Myofibroblast |
| Hsd11b1 | 0.44 | 0.00 | 0.00 | 0.34 | 0.09 | 30 | Myofibroblast |
| Tspan7 | 0.51 | 0.00 | 0.00 | 0.59 | 0.22 | 30 | Myofibroblast |
| Dpep1 | 0.60 | 0.00 | 0.00 | 0.27 | 0.07 | 30 | Myofibroblast |
| Peg3 | 0.76 | 0.00 | 0.00 | 0.68 | 0.30 | 30 | Myofibroblast |
| Pecam1 | 0.31 | 0.00 | 0.00 | 0.67 | 0.24 | 30 | Myofibroblast |
| Sparcl1 | 0.47 | 0.00 | 0.00 | 0.82 | 0.39 | 30 | Myofibroblast |
| Mylk | 0.52 | 0.00 | 0.00 | 0.67 | 0.27 | 30 | Myofibroblast |
| Kitl | 0.48 | 0.00 | 0.00 | 0.58 | 0.23 | 30 | Myofibroblast |
| Colec12 | 0.39 | 0.00 | 0.00 | 0.36 | 0.10 | 30 | Myofibroblast |
| Clec1a | 0.51 | 0.00 | 0.00 | 0.49 | 0.17 | 30 | Myofibroblast |
| Bgn | 0.43 | 0.00 | 0.00 | 0.64 | 0.24 | 30 | Myofibroblast |
| BC028528 | 0.42 | 0.00 | 0.00 | 0.70 | 0.29 | 30 | Myofibroblast |
| Tspan13 | 0.36 | 0.00 | 0.00 | 0.64 | 0.25 | 30 | Myofibroblast |
| Gng11 | 0.47 | 0.00 | 0.00 | 0.69 | 0.29 | 30 | Myofibroblast |
| Slc29a1 | 0.41 | 0.00 | 0.00 | 0.30 | 0.08 | 30 | Myofibroblast |
| Nexn | 0.41 | 0.00 | 0.00 | 0.57 | 0.22 | 30 | Myofibroblast |
| Adamts1 | 0.67 | 0.00 | 0.00 | 0.55 | 0.21 | 30 | Myofibroblast |
| Vcan | 0.62 | 0.00 | 0.00 | 0.35 | 0.11 | 30 | Myofibroblast |
| Igfbp4 | 0.65 | 0.00 | 0.00 | 0.61 | 0.25 | 30 | Myofibroblast |
| Palld | 0.47 | 0.00 | 0.00 | 0.49 | 0.18 | 30 | Myofibroblast |
| Scn3b | 0.55 | 0.00 | 0.00 | 0.28 | 0.08 | 30 | Myofibroblast |
| Serpine2 | 0.41 | 0.00 | 0.00 | 0.62 | 0.26 | 30 | Myofibroblast |
| Ppap2a | 0.40 | 0.00 | 0.00 | 0.52 | 0.19 | 30 | Myofibroblast |
| Agpat4 | 0.54 | 0.00 | 0.00 | 0.39 | 0.13 | 30 | Myofibroblast |
| Fgfr1 | 0.45 | 0.00 | 0.00 | 0.34 | 0.11 | 30 | Myofibroblast |
| Ace | 0.44 | 0.00 | 0.00 | 0.40 | 0.14 | 30 | Myofibroblast |
| Tbx2 | 0.49 | 0.00 | 0.00 | 0.36 | 0.12 | 30 | Myofibroblast |
| Foxf1 | 0.44 | 0.00 | 0.00 | 0.37 | 0.12 | 30 | Myofibroblast |
| Eltd1 | 0.37 | 0.00 | 0.00 | 0.26 | 0.07 | 30 | Myofibroblast |
| Phlda1 | 0.49 | 0.00 | 0.00 | 0.52 | 0.21 | 30 | Myofibroblast |
| Pmp22 | 0.39 | 0.00 | 0.00 | 0.69 | 0.32 | 30 | Myofibroblast |
| Gimap5 | 0.49 | 0.00 | 0.00 | 0.35 | 0.12 | 30 | Myofibroblast |
| Meis2 | 0.47 | 0.00 | 0.00 | 0.36 | 0.12 | 30 | Myofibroblast |
| Heg1 | 0.38 | 0.00 | 0.00 | 0.44 | 0.16 | 30 | Myofibroblast |
| Selenbp1 | 0.38 | 0.00 | 0.00 | 0.56 | 0.23 | 30 | Myofibroblast |
| S100a16 | 0.33 | 0.00 | 0.00 | 0.59 | 0.24 | 30 | Myofibroblast |
| Ehd4 | 0.42 | 0.00 | 0.00 | 0.63 | 0.28 | 30 | Myofibroblast |
| Pcdh17 | 0.41 | 0.00 | 0.00 | 0.53 | 0.21 | 30 | Myofibroblast |
| Itga8 | 0.56 | 0.00 | 0.00 | 0.30 | 0.09 | 30 | Myofibroblast |
| Hivep3 | 0.44 | 0.00 | 0.00 | 0.26 | 0.08 | 30 | Myofibroblast |
| 1810011O10Rik | 0.34 | 0.00 | 0.00 | 0.28 | 0.08 | 30 | Myofibroblast |
| Emilin1 | 0.41 | 0.00 | 0.00 | 0.30 | 0.09 | 30 | Myofibroblast |
| Rgs12 | 0.35 | 0.00 | 0.00 | 0.36 | 0.12 | 30 | Myofibroblast |
| Slc9a3r2 | 0.41 | 0.00 | 0.00 | 0.52 | 0.21 | 30 | Myofibroblast |
| Nfib | 0.48 | 0.00 | 0.00 | 0.82 | 0.47 | 30 | Myofibroblast |
| Nid1 | 0.49 | 0.00 | 0.00 | 0.55 | 0.23 | 30 | Myofibroblast |
| Cxx1b | 0.51 | 0.00 | 0.00 | 0.53 | 0.22 | 30 | Myofibroblast |
| Ppp1r14a | 0.32 | 0.00 | 0.00 | 0.28 | 0.08 | 30 | Myofibroblast |
| Aqp1 | 0.32 | 0.00 | 0.00 | 0.51 | 0.20 | 30 | Myofibroblast |
| Prex2 | 0.43 | 0.00 | 0.00 | 0.48 | 0.19 | 30 | Myofibroblast |
| Ednrb | 0.58 | 0.00 | 0.00 | 0.40 | 0.15 | 30 | Myofibroblast |
| Mxra8 | 0.46 | 0.00 | 0.00 | 0.43 | 0.17 | 30 | Myofibroblast |
| Mmp2 | 0.48 | 0.00 | 0.00 | 0.38 | 0.14 | 30 | Myofibroblast |
| Olfml3 | 0.44 | 0.00 | 0.00 | 0.28 | 0.09 | 30 | Myofibroblast |
| S1pr1 | 0.39 | 0.00 | 0.00 | 0.49 | 0.19 | 30 | Myofibroblast |
| Id1 | 0.39 | 0.00 | 0.00 | 0.56 | 0.24 | 30 | Myofibroblast |
| Emcn | 0.27 | 0.00 | 0.00 | 0.34 | 0.11 | 30 | Myofibroblast |
| Cd200 | 0.46 | 0.00 | 0.00 | 0.60 | 0.27 | 30 | Myofibroblast |
| Ldb2 | 0.44 | 0.00 | 0.00 | 0.43 | 0.17 | 30 | Myofibroblast |
| Kank3 | 0.28 | 0.00 | 0.00 | 0.40 | 0.14 | 30 | Myofibroblast |
| Ripply3 | 0.38 | 0.00 | 0.00 | 0.37 | 0.13 | 30 | Myofibroblast |
| Tcf4 | 0.52 | 0.00 | 0.00 | 0.74 | 0.41 | 30 | Myofibroblast |
| Cxx1c | 0.57 | 0.00 | 0.00 | 0.51 | 0.22 | 30 | Myofibroblast |
| Tie1 | 0.37 | 0.00 | 0.00 | 0.34 | 0.12 | 30 | Myofibroblast |
| Hspb1 | 0.49 | 0.00 | 0.00 | 0.37 | 0.14 | 30 | Myofibroblast |
| Csrp1 | 0.43 | 0.00 | 0.00 | 0.42 | 0.17 | 30 | Myofibroblast |
| Isoc1 | 0.49 | 0.00 | 0.00 | 0.26 | 0.08 | 30 | Myofibroblast |
| Emp1 | 0.35 | 0.00 | 0.00 | 0.54 | 0.22 | 30 | Myofibroblast |
| Gpr116 | 0.28 | 0.00 | 0.00 | 0.77 | 0.38 | 30 | Myofibroblast |
| Ccdc80 | 0.28 | 0.00 | 0.00 | 0.34 | 0.12 | 30 | Myofibroblast |
| Ism1 | 0.45 | 0.00 | 0.00 | 0.28 | 0.09 | 30 | Myofibroblast |
| Cdc42ep1 | 0.49 | 0.00 | 0.00 | 0.26 | 0.08 | 30 | Myofibroblast |
| Igfbp7 | 0.43 | 0.00 | 0.00 | 0.73 | 0.39 | 30 | Myofibroblast |
| Col4a1 | 0.38 | 0.00 | 0.00 | 0.83 | 0.50 | 30 | Myofibroblast |
| Marcks | 0.37 | 0.00 | 0.00 | 0.84 | 0.50 | 30 | Myofibroblast |
| Tmem204 | 0.44 | 0.00 | 0.00 | 0.40 | 0.16 | 30 | Myofibroblast |
| Mest | 0.28 | 0.00 | 0.00 | 0.50 | 0.20 | 30 | Myofibroblast |
| Ecscr | 0.28 | 0.00 | 0.00 | 0.44 | 0.17 | 30 | Myofibroblast |
| Gata2 | 0.31 | 0.00 | 0.00 | 0.38 | 0.14 | 30 | Myofibroblast |
| Tmem88 | 0.29 | 0.00 | 0.00 | 0.32 | 0.11 | 30 | Myofibroblast |
| C230081A13Rik | 0.36 | 0.00 | 0.00 | 0.30 | 0.10 | 30 | Myofibroblast |
| Ctsl | 0.37 | 0.00 | 0.00 | 0.75 | 0.39 | 30 | Myofibroblast |
| Rnf144a | 0.31 | 0.00 | 0.00 | 0.41 | 0.16 | 30 | Myofibroblast |
| Ppp1r16b | 0.31 | 0.00 | 0.00 | 0.29 | 0.10 | 30 | Myofibroblast |
| Ly6c1 | 0.37 | 0.00 | 0.00 | 0.47 | 0.19 | 30 | Myofibroblast |
| Slc43a3 | 0.45 | 0.00 | 0.00 | 0.51 | 0.22 | 30 | Myofibroblast |
| Fendrr | 0.27 | 0.00 | 0.00 | 0.50 | 0.20 | 30 | Myofibroblast |
| Cav2 | 0.33 | 0.00 | 0.00 | 0.48 | 0.19 | 30 | Myofibroblast |
| Adarb1 | 0.37 | 0.00 | 0.00 | 0.25 | 0.08 | 30 | Myofibroblast |
| Cpm | 0.55 | 0.00 | 0.00 | 0.43 | 0.18 | 30 | Myofibroblast |
| Myzap | 0.41 | 0.00 | 0.00 | 0.38 | 0.15 | 30 | Myofibroblast |
| Esam | 0.39 | 0.00 | 0.00 | 0.44 | 0.18 | 30 | Myofibroblast |
| Mcam | 0.41 | 0.00 | 0.00 | 0.35 | 0.13 | 30 | Myofibroblast |
| Ext1 | 0.45 | 0.00 | 0.00 | 0.25 | 0.08 | 30 | Myofibroblast |
| Car4 | 0.42 | 0.00 | 0.00 | 0.25 | 0.08 | 30 | Myofibroblast |
| Specc1l | 0.40 | 0.00 | 0.00 | 0.33 | 0.12 | 30 | Myofibroblast |
| Cd34 | 0.38 | 0.00 | 0.00 | 0.48 | 0.20 | 30 | Myofibroblast |
| Nrep | 0.49 | 0.00 | 0.00 | 0.73 | 0.43 | 30 | Myofibroblast |
| Mpv17 | 0.33 | 0.00 | 0.00 | 0.28 | 0.10 | 30 | Myofibroblast |
| Rasgrp3 | 0.48 | 0.00 | 0.00 | 0.34 | 0.13 | 30 | Myofibroblast |
| Btbd3 | 0.44 | 0.00 | 0.00 | 0.37 | 0.15 | 30 | Myofibroblast |
| Kit | 0.29 | 0.00 | 0.00 | 0.26 | 0.08 | 30 | Myofibroblast |
| Sdpr | 0.35 | 0.00 | 0.00 | 0.85 | 0.52 | 30 | Myofibroblast |
| Igf2r | 0.42 | 0.00 | 0.00 | 0.30 | 0.11 | 30 | Myofibroblast |
| Plcb4 | 0.32 | 0.00 | 0.00 | 0.37 | 0.14 | 30 | Myofibroblast |
| Myo6 | 0.32 | 0.00 | 0.00 | 0.36 | 0.14 | 30 | Myofibroblast |
| Mfap2 | 0.36 | 0.00 | 0.00 | 0.71 | 0.35 | 30 | Myofibroblast |
| Tmem47 | 0.29 | 0.00 | 0.00 | 0.29 | 0.10 | 30 | Myofibroblast |
| Ccnd1 | 0.45 | 0.00 | 0.00 | 0.44 | 0.19 | 30 | Myofibroblast |
| Snai2 | 0.37 | 0.00 | 0.00 | 0.33 | 0.12 | 30 | Myofibroblast |
| Pltp | 0.46 | 0.00 | 0.00 | 0.42 | 0.18 | 30 | Myofibroblast |
| Ivd | 0.36 | 0.00 | 0.00 | 0.26 | 0.09 | 30 | Myofibroblast |
| Plk2 | 0.47 | 0.00 | 0.00 | 0.39 | 0.16 | 30 | Myofibroblast |
| Lama4 | 0.47 | 0.00 | 0.00 | 0.34 | 0.13 | 30 | Myofibroblast |
| Klf2 | 0.33 | 0.00 | 0.00 | 0.47 | 0.20 | 30 | Myofibroblast |
| Npr3 | 0.47 | 0.00 | 0.00 | 0.33 | 0.13 | 30 | Myofibroblast |
| Klf4 | 0.38 | 0.00 | 0.00 | 0.51 | 0.24 | 30 | Myofibroblast |
| Prx | 0.33 | 0.00 | 0.00 | 0.29 | 0.11 | 30 | Myofibroblast |
| Fas | 0.26 | 0.00 | 0.00 | 0.28 | 0.10 | 30 | Myofibroblast |
| Clec14a | 0.36 | 0.00 | 0.00 | 0.42 | 0.18 | 30 | Myofibroblast |
| Col6a2 | 0.32 | 0.00 | 0.00 | 0.37 | 0.15 | 30 | Myofibroblast |
| Cxx1a | 0.50 | 0.00 | 0.00 | 0.48 | 0.22 | 30 | Myofibroblast |
| Col4a2 | 0.32 | 0.00 | 0.00 | 0.76 | 0.43 | 30 | Myofibroblast |
| Ptrf | 0.37 | 0.00 | 0.00 | 0.65 | 0.35 | 30 | Myofibroblast |
| Nfia | 0.40 | 0.00 | 0.00 | 0.51 | 0.24 | 30 | Myofibroblast |
| Tns3 | 0.43 | 0.00 | 0.00 | 0.29 | 0.11 | 30 | Myofibroblast |
| Gstm1 | 0.40 | 0.00 | 0.00 | 0.36 | 0.15 | 30 | Myofibroblast |
| Pcolce | 0.31 | 0.00 | 0.00 | 0.25 | 0.09 | 30 | Myofibroblast |
| Plcg1 | 0.35 | 0.00 | 0.00 | 0.38 | 0.16 | 30 | Myofibroblast |
| Ifitm2 | 0.40 | 0.00 | 0.00 | 0.60 | 0.32 | 30 | Myofibroblast |
| Fxyd1 | 0.31 | 0.00 | 0.00 | 0.26 | 0.09 | 30 | Myofibroblast |
| 9430020K01Rik | 0.26 | 0.00 | 0.00 | 0.33 | 0.13 | 30 | Myofibroblast |
| Lpl | 0.35 | 0.00 | 0.00 | 0.35 | 0.15 | 30 | Myofibroblast |
| Nrp1 | 0.36 | 0.00 | 0.00 | 0.70 | 0.39 | 30 | Myofibroblast |
| Xist | 0.39 | 0.00 | 0.00 | 0.79 | 0.51 | 30 | Myofibroblast |
| Smarca2 | 0.36 | 0.00 | 0.00 | 0.63 | 0.35 | 30 | Myofibroblast |
| Mdk | 0.33 | 0.00 | 0.00 | 0.42 | 0.19 | 30 | Myofibroblast |
| Adamts10 | 0.26 | 0.00 | 0.00 | 0.30 | 0.12 | 30 | Myofibroblast |
| Epb4.1l2 | 0.32 | 0.00 | 0.00 | 0.42 | 0.19 | 30 | Myofibroblast |
| Trim12c | 0.29 | 0.00 | 0.00 | 0.26 | 0.10 | 30 | Myofibroblast |
| Blmh | 0.45 | 0.00 | 0.00 | 0.38 | 0.17 | 30 | Myofibroblast |
| Erg | 0.34 | 0.00 | 0.00 | 0.33 | 0.14 | 30 | Myofibroblast |
| Taok2 | 0.32 | 0.00 | 0.00 | 0.27 | 0.10 | 30 | Myofibroblast |
| Cnep1r1 | 0.31 | 0.00 | 0.00 | 0.29 | 0.12 | 30 | Myofibroblast |
| Casz1 | 0.33 | 0.00 | 0.00 | 0.33 | 0.14 | 30 | Myofibroblast |
| Vldlr | 0.30 | 0.00 | 0.00 | 0.29 | 0.11 | 30 | Myofibroblast |
| Acadsb | 0.33 | 0.00 | 0.00 | 0.28 | 0.11 | 30 | Myofibroblast |
| Arhgap29 | 0.33 | 0.00 | 0.00 | 0.46 | 0.22 | 30 | Myofibroblast |
| Zyx | 0.27 | 0.00 | 0.00 | 0.43 | 0.20 | 30 | Myofibroblast |
| Mgll | 0.28 | 0.00 | 0.00 | 0.37 | 0.16 | 30 | Myofibroblast |
| Selenbp2 | 0.32 | 0.00 | 0.00 | 0.38 | 0.17 | 30 | Myofibroblast |
| Tenc1 | 0.47 | 0.00 | 0.01 | 0.33 | 0.15 | 30 | Myofibroblast |
| Tmem176a | 0.34 | 0.00 | 0.01 | 0.42 | 0.21 | 30 | Myofibroblast |
| Col6a1 | 0.32 | 0.00 | 0.01 | 0.41 | 0.19 | 30 | Myofibroblast |
| Vim | 0.28 | 0.00 | 0.01 | 0.94 | 0.66 | 30 | Myofibroblast |
| Ccdc88a | 0.30 | 0.00 | 0.01 | 0.50 | 0.25 | 30 | Myofibroblast |
| Nenf | 0.35 | 0.00 | 0.01 | 0.39 | 0.18 | 30 | Myofibroblast |
| Nisch | 0.39 | 0.00 | 0.01 | 0.56 | 0.30 | 30 | Myofibroblast |
| Efnb1 | 0.29 | 0.00 | 0.01 | 0.34 | 0.15 | 30 | Myofibroblast |
| Syne1 | 0.32 | 0.00 | 0.01 | 0.27 | 0.11 | 30 | Myofibroblast |
| Sox4 | 0.41 | 0.00 | 0.01 | 0.62 | 0.37 | 30 | Myofibroblast |
| Fbn1 | 0.26 | 0.00 | 0.01 | 0.38 | 0.17 | 30 | Myofibroblast |
| Rasgrp2 | 0.36 | 0.00 | 0.01 | 0.25 | 0.10 | 30 | Myofibroblast |
| Ltbp4 | 0.34 | 0.00 | 0.01 | 0.27 | 0.11 | 30 | Myofibroblast |
| Fbln5 | 0.38 | 0.00 | 0.01 | 0.37 | 0.17 | 30 | Myofibroblast |
| Efnb2 | 0.44 | 0.00 | 0.01 | 0.33 | 0.15 | 30 | Myofibroblast |
| Polr2i | 0.34 | 0.00 | 0.02 | 0.34 | 0.15 | 30 | Myofibroblast |
| Oaz2 | 0.31 | 0.00 | 0.02 | 0.54 | 0.29 | 30 | Myofibroblast |
| Nav1 | 0.34 | 0.00 | 0.02 | 0.28 | 0.12 | 30 | Myofibroblast |
| Zbtb20 | 0.48 | 0.00 | 0.02 | 0.50 | 0.27 | 30 | Myofibroblast |
| Tgfb1i1 | 0.28 | 0.00 | 0.02 | 0.31 | 0.13 | 30 | Myofibroblast |
| Upf3b | 0.46 | 0.00 | 0.02 | 0.37 | 0.18 | 30 | Myofibroblast |
| Fkbp9 | 0.36 | 0.00 | 0.02 | 0.32 | 0.14 | 30 | Myofibroblast |
| Clic4 | 0.33 | 0.00 | 0.02 | 0.54 | 0.29 | 30 | Myofibroblast |
| Ttc28 | 0.31 | 0.00 | 0.03 | 0.29 | 0.12 | 30 | Myofibroblast |
| Ppp4c | 0.32 | 0.00 | 0.03 | 0.34 | 0.16 | 30 | Myofibroblast |
| Jun | 0.31 | 0.00 | 0.03 | 0.64 | 0.37 | 30 | Myofibroblast |
| Myo1b | 0.27 | 0.00 | 0.03 | 0.36 | 0.17 | 30 | Myofibroblast |
| Mbnl2 | 0.33 | 0.00 | 0.03 | 0.62 | 0.35 | 30 | Myofibroblast |
| Ybx1 | 0.29 | 0.00 | 0.03 | 0.87 | 0.64 | 30 | Myofibroblast |
| Rdx | 0.36 | 0.00 | 0.04 | 0.69 | 0.45 | 30 | Myofibroblast |
| Pbx1 | 0.32 | 0.00 | 0.04 | 0.40 | 0.20 | 30 | Myofibroblast |
| Ptprg | 0.25 | 0.00 | 0.04 | 0.29 | 0.13 | 30 | Myofibroblast |
| Sepp1 | 0.30 | 0.00 | 0.04 | 0.67 | 0.41 | 30 | Myofibroblast |
| Dnajc8 | 0.29 | 0.00 | 0.04 | 0.71 | 0.43 | 30 | Myofibroblast |
| Timp2 | 0.30 | 0.00 | 0.04 | 0.42 | 0.21 | 30 | Myofibroblast |
| Ralbp1 | 0.26 | 0.00 | 0.05 | 0.60 | 0.33 | 30 | Myofibroblast |
| N6amt1 | 0.28 | 0.00 | 0.05 | 0.29 | 0.13 | 30 | Myofibroblast |
| Sptbn1 | 0.34 | 0.00 | 0.05 | 0.78 | 0.55 | 30 | Myofibroblast |
| Slc34a2 | 1.62 | 0.00 | 0.00 | 0.99 | 0.16 | 31 | AT2 Cell |
| Cxcl15 | 1.58 | 0.00 | 0.00 | 0.93 | 0.15 | 31 | AT2 Cell |
| Sftpb | 1.72 | 0.00 | 0.00 | 0.98 | 0.18 | 31 | AT2 Cell |
| Hc | 1.42 | 0.00 | 0.00 | 0.81 | 0.12 | 31 | AT2 Cell |
| Il33 | 1.14 | 0.00 | 0.00 | 0.45 | 0.04 | 31 | AT2 Cell |
| Sftpa1 | 1.69 | 0.00 | 0.00 | 0.99 | 0.22 | 31 | AT2 Cell |
| Lamp3 | 1.38 | 0.00 | 0.00 | 0.76 | 0.12 | 31 | AT2 Cell |
| Sftpd | 1.45 | 0.00 | 0.00 | 0.90 | 0.17 | 31 | AT2 Cell |
| Sfta2 | 1.19 | 0.00 | 0.00 | 0.90 | 0.16 | 31 | AT2 Cell |
| Lpcat1 | 1.37 | 0.00 | 0.00 | 0.93 | 0.18 | 31 | AT2 Cell |
| S100g | 1.28 | 0.00 | 0.00 | 0.78 | 0.13 | 31 | AT2 Cell |
| Chi3l1 | 1.28 | 0.00 | 0.00 | 0.71 | 0.11 | 31 | AT2 Cell |
| Bex2 | 1.39 | 0.00 | 0.00 | 0.66 | 0.11 | 31 | AT2 Cell |
| Bex1 | 1.23 | 0.00 | 0.00 | 0.65 | 0.10 | 31 | AT2 Cell |
| Muc1 | 1.12 | 0.00 | 0.00 | 0.60 | 0.09 | 31 | AT2 Cell |
| Bex4 | 1.27 | 0.00 | 0.00 | 0.71 | 0.13 | 31 | AT2 Cell |
| Lgi3 | 0.94 | 0.00 | 0.00 | 0.54 | 0.08 | 31 | AT2 Cell |
| Egfl6 | 0.97 | 0.00 | 0.00 | 0.55 | 0.09 | 31 | AT2 Cell |
| Dram1 | 1.13 | 0.00 | 0.00 | 0.65 | 0.12 | 31 | AT2 Cell |
| Cbr2 | 0.92 | 0.00 | 0.00 | 0.79 | 0.17 | 31 | AT2 Cell |
| Wfdc2 | 0.98 | 0.00 | 0.00 | 0.80 | 0.17 | 31 | AT2 Cell |
| Lyz1 | 1.35 | 0.00 | 0.00 | 0.81 | 0.19 | 31 | AT2 Cell |
| Napsa | 1.08 | 0.00 | 0.00 | 0.85 | 0.20 | 31 | AT2 Cell |
| Scd1 | 1.05 | 0.00 | 0.00 | 0.54 | 0.09 | 31 | AT2 Cell |
| Slco4c1 | 0.73 | 0.00 | 0.00 | 0.29 | 0.03 | 31 | AT2 Cell |
| Rbpjl | 0.91 | 0.00 | 0.00 | 0.45 | 0.06 | 31 | AT2 Cell |
| Ppp1r14c | 1.00 | 0.00 | 0.00 | 0.61 | 0.11 | 31 | AT2 Cell |
| Sftpc | 1.72 | 0.00 | 0.00 | 1.00 | 0.63 | 31 | AT2 Cell |
| Cldn18 | 0.71 | 0.00 | 0.00 | 0.78 | 0.18 | 31 | AT2 Cell |
| Etv5 | 0.80 | 0.00 | 0.00 | 0.44 | 0.07 | 31 | AT2 Cell |
| Pla2g1b | 0.67 | 0.00 | 0.00 | 0.38 | 0.05 | 31 | AT2 Cell |
| Fabp5 | 0.99 | 0.00 | 0.00 | 0.61 | 0.13 | 31 | AT2 Cell |
| Ctsh | 0.98 | 0.00 | 0.00 | 0.78 | 0.21 | 31 | AT2 Cell |
| Kcnc3 | 0.53 | 0.00 | 0.00 | 0.29 | 0.04 | 31 | AT2 Cell |
| Lyz2 | 0.92 | 0.00 | 0.00 | 0.85 | 0.27 | 31 | AT2 Cell |
| Abca3 | 0.95 | 0.00 | 0.00 | 0.56 | 0.12 | 31 | AT2 Cell |
| Spink5 | 0.92 | 0.00 | 0.00 | 0.44 | 0.08 | 31 | AT2 Cell |
| Npc2 | 1.07 | 0.00 | 0.00 | 0.99 | 0.54 | 31 | AT2 Cell |
| Snhg11 | 0.75 | 0.00 | 0.00 | 0.29 | 0.04 | 31 | AT2 Cell |
| Nkx2-1 | 0.75 | 0.00 | 0.00 | 0.49 | 0.10 | 31 | AT2 Cell |
| Ptprf | 0.80 | 0.00 | 0.00 | 0.63 | 0.16 | 31 | AT2 Cell |
| Atp6v1c2 | 0.61 | 0.00 | 0.00 | 0.33 | 0.05 | 31 | AT2 Cell |
| Cldn3 | 0.67 | 0.00 | 0.00 | 0.54 | 0.12 | 31 | AT2 Cell |
| Ces1d | 0.73 | 0.00 | 0.00 | 0.64 | 0.17 | 31 | AT2 Cell |
| Lcn2 | 0.88 | 0.00 | 0.00 | 0.38 | 0.07 | 31 | AT2 Cell |
| Car8 | 0.87 | 0.00 | 0.00 | 0.39 | 0.08 | 31 | AT2 Cell |
| Acsl4 | 0.77 | 0.00 | 0.00 | 0.49 | 0.12 | 31 | AT2 Cell |
| Atp1b1 | 0.71 | 0.00 | 0.00 | 0.65 | 0.19 | 31 | AT2 Cell |
| Irx3 | 0.63 | 0.00 | 0.00 | 0.38 | 0.08 | 31 | AT2 Cell |
| Chchd10 | 0.62 | 0.00 | 0.00 | 0.53 | 0.13 | 31 | AT2 Cell |
| Cd74 | 0.61 | 0.00 | 0.00 | 0.50 | 0.13 | 31 | AT2 Cell |
| Scn7a | 0.57 | 0.00 | 0.00 | 0.81 | 0.30 | 31 | AT2 Cell |
| Neat1 | 0.68 | 0.00 | 0.00 | 0.76 | 0.27 | 31 | AT2 Cell |
| Sdc4 | 0.63 | 0.00 | 0.00 | 0.70 | 0.23 | 31 | AT2 Cell |
| Prrg3 | 0.68 | 0.00 | 0.00 | 0.36 | 0.08 | 31 | AT2 Cell |
| Dcxr | 0.64 | 0.00 | 0.00 | 0.44 | 0.11 | 31 | AT2 Cell |
| Rab27b | 0.65 | 0.00 | 0.00 | 0.34 | 0.07 | 31 | AT2 Cell |
| Fasn | 0.78 | 0.00 | 0.00 | 0.35 | 0.08 | 31 | AT2 Cell |
| Meg3 | 0.77 | 0.00 | 0.00 | 0.66 | 0.22 | 31 | AT2 Cell |
| Retnla | 1.32 | 0.00 | 0.00 | 0.41 | 0.11 | 31 | AT2 Cell |
| Alcam | 0.76 | 0.00 | 0.00 | 0.66 | 0.23 | 31 | AT2 Cell |
| Ank3 | 0.74 | 0.00 | 0.00 | 0.35 | 0.08 | 31 | AT2 Cell |
| Ctsc | 0.69 | 0.00 | 0.00 | 0.60 | 0.20 | 31 | AT2 Cell |
| Clec1a | 0.44 | 0.00 | 0.00 | 0.58 | 0.17 | 31 | AT2 Cell |
| Atp8a1 | 0.66 | 0.00 | 0.00 | 0.53 | 0.16 | 31 | AT2 Cell |
| Fgfr2 | 0.66 | 0.00 | 0.00 | 0.33 | 0.07 | 31 | AT2 Cell |
| Per3 | 0.74 | 0.00 | 0.00 | 0.26 | 0.05 | 31 | AT2 Cell |
| Gpr116 | 0.57 | 0.00 | 0.00 | 0.88 | 0.38 | 31 | AT2 Cell |
| Atp11a | 0.61 | 0.00 | 0.00 | 0.43 | 0.11 | 31 | AT2 Cell |
| Pi4k2b | 0.60 | 0.00 | 0.00 | 0.40 | 0.10 | 31 | AT2 Cell |
| Sdc1 | 0.41 | 0.00 | 0.00 | 0.45 | 0.12 | 31 | AT2 Cell |
| Epcam | 0.64 | 0.00 | 0.00 | 0.39 | 0.10 | 31 | AT2 Cell |
| Cat | 0.62 | 0.00 | 0.00 | 0.44 | 0.13 | 31 | AT2 Cell |
| Hpgd | 0.40 | 0.00 | 0.00 | 0.73 | 0.26 | 31 | AT2 Cell |
| Fendrr | 0.49 | 0.00 | 0.00 | 0.61 | 0.20 | 31 | AT2 Cell |
| Ptprb | 0.31 | 0.00 | 0.00 | 0.66 | 0.22 | 31 | AT2 Cell |
| Trf | 0.56 | 0.00 | 0.00 | 0.39 | 0.10 | 31 | AT2 Cell |
| Tmem100 | 0.38 | 0.00 | 0.00 | 0.80 | 0.30 | 31 | AT2 Cell |
| Gas6 | 0.64 | 0.00 | 0.00 | 0.48 | 0.15 | 31 | AT2 Cell |
| Mecom | 0.48 | 0.00 | 0.00 | 0.39 | 0.11 | 31 | AT2 Cell |
| Rnase4 | 0.79 | 0.00 | 0.00 | 0.54 | 0.19 | 31 | AT2 Cell |
| Wbp5 | 0.77 | 0.00 | 0.00 | 0.84 | 0.48 | 31 | AT2 Cell |
| Slc39a8 | 0.47 | 0.00 | 0.00 | 0.31 | 0.08 | 31 | AT2 Cell |
| Arhgef15 | 0.48 | 0.00 | 0.00 | 0.31 | 0.08 | 31 | AT2 Cell |
| Mgst1 | 0.47 | 0.00 | 0.00 | 0.58 | 0.20 | 31 | AT2 Cell |
| Ppp1r9a | 0.54 | 0.00 | 0.00 | 0.46 | 0.14 | 31 | AT2 Cell |
| Scd2 | 0.55 | 0.00 | 0.00 | 0.59 | 0.21 | 31 | AT2 Cell |
| Chka | 0.55 | 0.00 | 0.00 | 0.38 | 0.10 | 31 | AT2 Cell |
| H2-Aa | 0.33 | 0.00 | 0.00 | 0.33 | 0.08 | 31 | AT2 Cell |
| Ptgs1 | 0.41 | 0.00 | 0.00 | 0.38 | 0.11 | 31 | AT2 Cell |
| Nucb2 | 0.52 | 0.00 | 0.00 | 0.49 | 0.16 | 31 | AT2 Cell |
| Ngfrap1 | 0.54 | 0.00 | 0.00 | 0.61 | 0.24 | 31 | AT2 Cell |
| Nckap5 | 0.60 | 0.00 | 0.00 | 0.41 | 0.13 | 31 | AT2 Cell |
| Dpp4 | 0.44 | 0.00 | 0.00 | 0.43 | 0.13 | 31 | AT2 Cell |
| Brd7 | 0.69 | 0.00 | 0.00 | 0.63 | 0.26 | 31 | AT2 Cell |
| Arhgap31 | 0.42 | 0.00 | 0.00 | 0.40 | 0.12 | 31 | AT2 Cell |
| Myo5c | 0.44 | 0.00 | 0.00 | 0.30 | 0.08 | 31 | AT2 Cell |
| Abcd3 | 0.60 | 0.00 | 0.00 | 0.50 | 0.19 | 31 | AT2 Cell |
| Pecam1 | 0.42 | 0.00 | 0.00 | 0.64 | 0.24 | 31 | AT2 Cell |
| Kit | 0.40 | 0.00 | 0.00 | 0.31 | 0.08 | 31 | AT2 Cell |
| Cd93 | 0.28 | 0.00 | 0.00 | 0.61 | 0.22 | 31 | AT2 Cell |
| Mt1 | 0.42 | 0.00 | 0.00 | 0.46 | 0.16 | 31 | AT2 Cell |
| Zdhhc3 | 0.50 | 0.00 | 0.00 | 0.45 | 0.16 | 31 | AT2 Cell |
| Pon3 | 0.42 | 0.00 | 0.00 | 0.34 | 0.10 | 31 | AT2 Cell |
| Arap3 | 0.39 | 0.00 | 0.00 | 0.29 | 0.08 | 31 | AT2 Cell |
| Cadm1 | 0.43 | 0.00 | 0.00 | 0.43 | 0.14 | 31 | AT2 Cell |
| Id2 | 0.46 | 0.00 | 0.00 | 0.71 | 0.33 | 31 | AT2 Cell |
| Serpinb6b | 0.40 | 0.00 | 0.00 | 0.48 | 0.17 | 31 | AT2 Cell |
| Tspan8 | 0.32 | 0.00 | 0.00 | 0.39 | 0.12 | 31 | AT2 Cell |
| Mbip | 0.50 | 0.00 | 0.00 | 0.36 | 0.12 | 31 | AT2 Cell |
| Mid1ip1 | 0.54 | 0.00 | 0.00 | 0.41 | 0.14 | 31 | AT2 Cell |
| Flt1 | 0.36 | 0.00 | 0.00 | 0.46 | 0.16 | 31 | AT2 Cell |
| Cldn5 | 0.31 | 0.00 | 0.00 | 0.59 | 0.23 | 31 | AT2 Cell |
| Mapk3 | 0.35 | 0.00 | 0.00 | 0.41 | 0.14 | 31 | AT2 Cell |
| Prnp | 0.46 | 0.00 | 0.00 | 0.38 | 0.12 | 31 | AT2 Cell |
| Cd36 | 0.41 | 0.00 | 0.00 | 0.74 | 0.33 | 31 | AT2 Cell |
| Ctla2a | 0.50 | 0.00 | 0.00 | 0.53 | 0.21 | 31 | AT2 Cell |
| S100a16 | 0.41 | 0.00 | 0.00 | 0.59 | 0.24 | 31 | AT2 Cell |
| Stard4 | 0.39 | 0.00 | 0.00 | 0.33 | 0.10 | 31 | AT2 Cell |
| Acsl5 | 0.39 | 0.00 | 0.00 | 0.35 | 0.11 | 31 | AT2 Cell |
| Cldn7 | 0.32 | 0.00 | 0.00 | 0.29 | 0.08 | 31 | AT2 Cell |
| Sgk1 | 0.32 | 0.00 | 0.00 | 0.51 | 0.19 | 31 | AT2 Cell |
| Egfl7 | 0.25 | 0.00 | 0.00 | 0.70 | 0.28 | 31 | AT2 Cell |
| Kdr | 0.33 | 0.00 | 0.00 | 0.45 | 0.17 | 31 | AT2 Cell |
| Cmtm8 | 0.37 | 0.00 | 0.00 | 0.38 | 0.13 | 31 | AT2 Cell |
| Soat1 | 0.36 | 0.00 | 0.00 | 0.34 | 0.11 | 31 | AT2 Cell |
| Tmem243 | 0.52 | 0.00 | 0.00 | 0.28 | 0.08 | 31 | AT2 Cell |
| Tgoln1 | 0.46 | 0.00 | 0.00 | 0.48 | 0.19 | 31 | AT2 Cell |
| Pgs1 | 0.41 | 0.00 | 0.00 | 0.26 | 0.08 | 31 | AT2 Cell |
| Npc1 | 0.48 | 0.00 | 0.00 | 0.29 | 0.09 | 31 | AT2 Cell |
| Yae1d1 | 0.44 | 0.00 | 0.00 | 0.29 | 0.09 | 31 | AT2 Cell |
| Atp1a1 | 0.34 | 0.00 | 0.00 | 0.41 | 0.15 | 31 | AT2 Cell |
| Aqp1 | 0.26 | 0.00 | 0.00 | 0.51 | 0.20 | 31 | AT2 Cell |
| Secisbp2l | 0.47 | 0.00 | 0.00 | 0.46 | 0.18 | 31 | AT2 Cell |
| Myzap | 0.26 | 0.00 | 0.00 | 0.41 | 0.15 | 31 | AT2 Cell |
| Ptgfrn | 0.45 | 0.00 | 0.00 | 0.30 | 0.10 | 31 | AT2 Cell |
| Erg | 0.33 | 0.00 | 0.00 | 0.39 | 0.14 | 31 | AT2 Cell |
| Ccnd1 | 0.43 | 0.00 | 0.00 | 0.48 | 0.19 | 31 | AT2 Cell |
| Gclc | 0.64 | 0.00 | 0.00 | 0.28 | 0.09 | 31 | AT2 Cell |
| Bcam | 0.29 | 0.00 | 0.00 | 0.30 | 0.10 | 31 | AT2 Cell |
| Scn3b | 0.29 | 0.00 | 0.00 | 0.26 | 0.08 | 31 | AT2 Cell |
| Taok3 | 0.40 | 0.00 | 0.00 | 0.46 | 0.19 | 31 | AT2 Cell |
| Cdh1 | 0.35 | 0.00 | 0.00 | 0.26 | 0.08 | 31 | AT2 Cell |
| H2afj | 0.51 | 0.00 | 0.00 | 0.65 | 0.32 | 31 | AT2 Cell |
| Elovl1 | 0.43 | 0.00 | 0.00 | 0.40 | 0.15 | 31 | AT2 Cell |
| Plcg1 | 0.55 | 0.00 | 0.00 | 0.41 | 0.16 | 31 | AT2 Cell |
| Mettl7a1 | 0.38 | 0.00 | 0.00 | 0.53 | 0.22 | 31 | AT2 Cell |
| Afap1l1 | 0.33 | 0.00 | 0.00 | 0.45 | 0.18 | 31 | AT2 Cell |
| Casz1 | 0.39 | 0.00 | 0.00 | 0.38 | 0.14 | 31 | AT2 Cell |
| Ptplad1 | 0.32 | 0.00 | 0.00 | 0.29 | 0.09 | 31 | AT2 Cell |
| Nrp1 | 0.42 | 0.00 | 0.00 | 0.71 | 0.39 | 31 | AT2 Cell |
| Edn1 | 0.31 | 0.00 | 0.00 | 0.30 | 0.10 | 31 | AT2 Cell |
| Acvrl1 | 0.28 | 0.00 | 0.00 | 0.45 | 0.18 | 31 | AT2 Cell |
| Tbx3 | 0.31 | 0.00 | 0.00 | 0.50 | 0.21 | 31 | AT2 Cell |
| Podxl | 0.28 | 0.00 | 0.00 | 0.33 | 0.11 | 31 | AT2 Cell |
| Slco2a1 | 0.32 | 0.00 | 0.00 | 0.33 | 0.11 | 31 | AT2 Cell |
| Acly | 0.46 | 0.00 | 0.00 | 0.30 | 0.10 | 31 | AT2 Cell |
| Ldb2 | 0.38 | 0.00 | 0.00 | 0.43 | 0.17 | 31 | AT2 Cell |
| Ecscr | 0.31 | 0.00 | 0.00 | 0.44 | 0.17 | 31 | AT2 Cell |
| Sdpr | 0.32 | 0.00 | 0.00 | 0.88 | 0.52 | 31 | AT2 Cell |
| Rassf3 | 0.39 | 0.00 | 0.00 | 0.26 | 0.09 | 31 | AT2 Cell |
| Oraov1 | 0.35 | 0.00 | 0.00 | 0.28 | 0.09 | 31 | AT2 Cell |
| Tnfaip1 | 0.34 | 0.00 | 0.00 | 0.41 | 0.16 | 31 | AT2 Cell |
| Atp13a3 | 0.26 | 0.00 | 0.00 | 0.29 | 0.10 | 31 | AT2 Cell |
| Wwc2 | 0.36 | 0.00 | 0.00 | 0.35 | 0.13 | 31 | AT2 Cell |
| Gde1 | 0.37 | 0.00 | 0.00 | 0.38 | 0.15 | 31 | AT2 Cell |
| Id1 | 0.36 | 0.00 | 0.00 | 0.53 | 0.24 | 31 | AT2 Cell |
| 1500011K16Rik | 0.36 | 0.00 | 0.00 | 0.35 | 0.13 | 31 | AT2 Cell |
| Mest | 0.30 | 0.00 | 0.00 | 0.46 | 0.20 | 31 | AT2 Cell |
| Ano6 | 0.37 | 0.00 | 0.00 | 0.43 | 0.18 | 31 | AT2 Cell |
| Lpin2 | 0.41 | 0.00 | 0.00 | 0.38 | 0.15 | 31 | AT2 Cell |
| BC028528 | 0.30 | 0.00 | 0.00 | 0.61 | 0.29 | 31 | AT2 Cell |
| Sptlc2 | 0.29 | 0.00 | 0.00 | 0.38 | 0.15 | 31 | AT2 Cell |
| Rasgef1a | 0.33 | 0.00 | 0.00 | 0.26 | 0.09 | 31 | AT2 Cell |
| Ccz1 | 0.34 | 0.00 | 0.00 | 0.34 | 0.13 | 31 | AT2 Cell |
| Stmn2 | 0.30 | 0.00 | 0.00 | 0.39 | 0.16 | 31 | AT2 Cell |
| Nkd1 | 0.43 | 0.00 | 0.00 | 0.30 | 0.11 | 31 | AT2 Cell |
| Esam | 0.30 | 0.00 | 0.01 | 0.44 | 0.18 | 31 | AT2 Cell |
| 2810025M15Rik | 0.34 | 0.00 | 0.01 | 0.33 | 0.13 | 31 | AT2 Cell |
| Ushbp1 | 0.30 | 0.00 | 0.01 | 0.26 | 0.09 | 31 | AT2 Cell |
| Vamp8 | 0.42 | 0.00 | 0.01 | 0.64 | 0.35 | 31 | AT2 Cell |
| Dock9 | 0.32 | 0.00 | 0.01 | 0.28 | 0.10 | 31 | AT2 Cell |
| Sepp1 | 0.30 | 0.00 | 0.01 | 0.73 | 0.41 | 31 | AT2 Cell |
| Lphn2 | 0.38 | 0.00 | 0.01 | 0.34 | 0.14 | 31 | AT2 Cell |
| Pvrl3 | 0.45 | 0.00 | 0.01 | 0.28 | 0.10 | 31 | AT2 Cell |
| Apc | 0.32 | 0.00 | 0.01 | 0.50 | 0.24 | 31 | AT2 Cell |
| Tmem245 | 0.42 | 0.00 | 0.01 | 0.28 | 0.10 | 31 | AT2 Cell |
| Rnaset2a | 0.36 | 0.00 | 0.02 | 0.53 | 0.26 | 31 | AT2 Cell |
| Rai14 | 0.31 | 0.00 | 0.02 | 0.29 | 0.11 | 31 | AT2 Cell |
| Ctnnd1 | 0.31 | 0.00 | 0.02 | 0.39 | 0.16 | 31 | AT2 Cell |
| Piezo2 | 0.35 | 0.00 | 0.02 | 0.30 | 0.12 | 31 | AT2 Cell |
| Sypl | 0.28 | 0.00 | 0.03 | 0.54 | 0.26 | 31 | AT2 Cell |
| App | 0.30 | 0.00 | 0.03 | 0.86 | 0.60 | 31 | AT2 Cell |
| Tmbim6 | 0.41 | 0.00 | 0.03 | 0.69 | 0.38 | 31 | AT2 Cell |
| Pitpnc1 | 0.37 | 0.00 | 0.03 | 0.31 | 0.13 | 31 | AT2 Cell |
| Heg1 | 0.39 | 0.00 | 0.03 | 0.38 | 0.16 | 31 | AT2 Cell |
| Mtch1 | 0.37 | 0.00 | 0.03 | 0.35 | 0.15 | 31 | AT2 Cell |
| Uhrf1bp1l | 0.37 | 0.00 | 0.03 | 0.26 | 0.10 | 31 | AT2 Cell |
| Exosc7 | 0.41 | 0.00 | 0.04 | 0.33 | 0.13 | 31 | AT2 Cell |
| Tie1 | 0.28 | 0.00 | 0.04 | 0.30 | 0.12 | 31 | AT2 Cell |
| Tmem30a | 0.33 | 0.00 | 0.04 | 0.54 | 0.28 | 31 | AT2 Cell |
| Myh14 | 0.32 | 0.00 | 0.04 | 0.26 | 0.10 | 31 | AT2 Cell |

Table S8: Marker genes for individual clusters of cells from combined human and mouse cells

| Genes | Average Log Fold Change | Unadjusted P-value | Adjusted P-Value | Proportion of cells in Group 1 | Proportion of cells in Group 2 | Cluster No | Sub-Type |
| --- | --- | --- | --- | --- | --- | --- | --- |
| LYVE1 | 2.03 | 0.00 | 0.00 | 0.68 | 0.12 | 0 | Endothelial |
| CD93 | 1.95 | 0.00 | 0.00 | 0.77 | 0.11 | 0 | Endothelial |
| GPIHBP1 | 1.95 | 0.00 | 0.00 | 0.63 | 0.06 | 0 | Endothelial |
| TMEM100 | 1.81 | 0.00 | 0.00 | 0.85 | 0.19 | 0 | Endothelial |
| CALCRL | 1.78 | 0.00 | 0.00 | 0.80 | 0.15 | 0 | Endothelial |
| PTPRB | 1.70 | 0.00 | 0.00 | 0.73 | 0.11 | 0 | Endothelial |
| C1orf54 | 1.70 | 0.00 | 0.00 | 0.79 | 0.20 | 0 | Endothelial |
| SOX17 | 1.70 | 0.00 | 0.00 | 0.52 | 0.06 | 0 | Endothelial |
| SCN7A | 1.68 | 0.00 | 0.00 | 0.80 | 0.19 | 0 | Endothelial |
| PCDH17 | 1.67 | 0.00 | 0.00 | 0.66 | 0.12 | 0 | Endothelial |
| CLEC14A | 1.67 | 0.00 | 0.00 | 0.64 | 0.10 | 0 | Endothelial |
| EGFL7 | 1.66 | 0.00 | 0.00 | 0.88 | 0.17 | 0 | Endothelial |
| HPGD | 1.62 | 0.00 | 0.00 | 0.70 | 0.18 | 0 | Endothelial |
| CDH5 | 1.61 | 0.00 | 0.00 | 0.77 | 0.13 | 0 | Endothelial |
| PLVAP | 1.59 | 0.00 | 0.00 | 0.46 | 0.05 | 0 | Endothelial |
| EPAS1 | 1.57 | 0.00 | 0.00 | 0.75 | 0.18 | 0 | Endothelial |
| TSPAN7 | 1.56 | 0.00 | 0.00 | 0.65 | 0.13 | 0 | Endothelial |
| RAMP2 | 1.56 | 0.00 | 0.00 | 0.85 | 0.19 | 0 | Endothelial |
| CYYR1 | 1.55 | 0.00 | 0.00 | 0.51 | 0.08 | 0 | Endothelial |
| TEK | 1.54 | 0.00 | 0.00 | 0.48 | 0.05 | 0 | Endothelial |
| RASGRP3 | 1.50 | 0.00 | 0.00 | 0.46 | 0.06 | 0 | Endothelial |
| PECAM1 | 1.48 | 0.00 | 0.00 | 0.75 | 0.15 | 0 | Endothelial |
| LDB2 | 1.46 | 0.00 | 0.00 | 0.54 | 0.10 | 0 | Endothelial |
| S100A16 | 1.41 | 0.00 | 0.00 | 0.66 | 0.18 | 0 | Endothelial |
| CD200 | 1.40 | 0.00 | 0.00 | 0.57 | 0.21 | 0 | Endothelial |
| CLEC1A | 1.38 | 0.00 | 0.00 | 0.55 | 0.09 | 0 | Endothelial |
| EDN1 | 1.36 | 0.00 | 0.00 | 0.34 | 0.05 | 0 | Endothelial |
| SLC43A3 | 1.36 | 0.00 | 0.00 | 0.54 | 0.12 | 0 | Endothelial |
| CD36 | 1.35 | 0.00 | 0.00 | 0.75 | 0.25 | 0 | Endothelial |
| MYZAP | 1.34 | 0.00 | 0.00 | 0.48 | 0.08 | 0 | Endothelial |
| STMN2 | 1.34 | 0.00 | 0.00 | 0.46 | 0.09 | 0 | Endothelial |
| PALMD | 1.34 | 0.00 | 0.00 | 0.40 | 0.07 | 0 | Endothelial |
| AQP1 | 1.32 | 0.00 | 0.00 | 0.56 | 0.14 | 0 | Endothelial |
| TM4SF1 | 1.32 | 0.00 | 0.00 | 0.61 | 0.15 | 0 | Endothelial |
| SEMA3C | 1.32 | 0.00 | 0.00 | 0.47 | 0.14 | 0 | Endothelial |
| CLDN5 | 1.31 | 0.00 | 0.00 | 0.73 | 0.14 | 0 | Endothelial |
| TSPAN18 | 1.31 | 0.00 | 0.00 | 0.44 | 0.07 | 0 | Endothelial |
| KIT | 1.31 | 0.00 | 0.00 | 0.33 | 0.03 | 0 | Endothelial |
| RASIP1 | 1.30 | 0.00 | 0.00 | 0.42 | 0.06 | 0 | Endothelial |
| EFNB2 | 1.28 | 0.00 | 0.00 | 0.43 | 0.09 | 0 | Endothelial |
| GATA2 | 1.27 | 0.00 | 0.00 | 0.44 | 0.08 | 0 | Endothelial |
| ACER2 | 1.25 | 0.00 | 0.00 | 0.33 | 0.04 | 0 | Endothelial |
| THBD | 1.25 | 0.00 | 0.00 | 0.61 | 0.18 | 0 | Endothelial |
| RIPPLY3 | 1.24 | 0.00 | 0.00 | 0.40 | 0.06 | 0 | Endothelial |
| EHD4 | 1.24 | 0.00 | 0.00 | 0.65 | 0.19 | 0 | Endothelial |
| NOSTRIN | 1.24 | 0.00 | 0.00 | 0.45 | 0.09 | 0 | Endothelial |
| ECSCR | 1.24 | 0.00 | 0.00 | 0.51 | 0.09 | 0 | Endothelial |
| TIE1 | 1.23 | 0.00 | 0.00 | 0.40 | 0.06 | 0 | Endothelial |
| AFAP1L1 | 1.21 | 0.00 | 0.00 | 0.48 | 0.11 | 0 | Endothelial |
| TSPAN13 | 1.21 | 0.00 | 0.00 | 0.66 | 0.16 | 0 | Endothelial |
| ACVRL1 | 1.19 | 0.00 | 0.00 | 0.52 | 0.12 | 0 | Endothelial |
| SGK1 | 1.19 | 0.00 | 0.00 | 0.47 | 0.13 | 0 | Endothelial |
| ITGA6 | 1.18 | 0.00 | 0.00 | 0.38 | 0.08 | 0 | Endothelial |
| FLT1 | 1.18 | 0.00 | 0.00 | 0.45 | 0.09 | 0 | Endothelial |
| ESAM | 1.16 | 0.00 | 0.00 | 0.54 | 0.14 | 0 | Endothelial |
| CAV1 | 1.16 | 0.00 | 0.00 | 0.78 | 0.29 | 0 | Endothelial |
| MYCT1 | 1.15 | 0.00 | 0.00 | 0.41 | 0.07 | 0 | Endothelial |
| SEMA3G | 1.15 | 0.00 | 0.00 | 0.27 | 0.03 | 0 | Endothelial |
| ECE1 | 1.15 | 0.00 | 0.00 | 0.58 | 0.18 | 0 | Endothelial |
| RNF144A | 1.15 | 0.00 | 0.00 | 0.40 | 0.10 | 0 | Endothelial |
| USHBP1 | 1.15 | 0.00 | 0.00 | 0.31 | 0.04 | 0 | Endothelial |
| SLC9A3R2 | 1.14 | 0.00 | 0.00 | 0.55 | 0.17 | 0 | Endothelial |
| CXCL12 | 1.14 | 0.00 | 0.00 | 0.31 | 0.10 | 0 | Endothelial |
| KLF4 | 1.10 | 0.00 | 0.00 | 0.51 | 0.18 | 0 | Endothelial |
| APLNR | 1.09 | 0.00 | 0.00 | 0.28 | 0.04 | 0 | Endothelial |
| PODXL | 1.07 | 0.00 | 0.00 | 0.33 | 0.07 | 0 | Endothelial |
| ICAM2 | 1.06 | 0.00 | 0.00 | 0.49 | 0.11 | 0 | Endothelial |
| KLHL5 | 1.06 | 0.00 | 0.00 | 0.37 | 0.09 | 0 | Endothelial |
| FMO1 | 1.05 | 0.00 | 0.00 | 0.33 | 0.07 | 0 | Endothelial |
| SLCO2A1 | 1.05 | 0.00 | 0.00 | 0.34 | 0.07 | 0 | Endothelial |
| TMEM88 | 1.04 | 0.00 | 0.00 | 0.32 | 0.06 | 0 | Endothelial |
| GJA4 | 1.03 | 0.00 | 0.00 | 0.25 | 0.04 | 0 | Endothelial |
| TUBB2A | 1.03 | 0.00 | 0.00 | 0.45 | 0.17 | 0 | Endothelial |
| EMCN | 1.02 | 0.00 | 0.00 | 0.37 | 0.07 | 0 | Endothelial |
| CCND1 | 1.00 | 0.00 | 0.00 | 0.43 | 0.15 | 0 | Endothelial |
| PDE4B | 0.99 | 0.00 | 0.00 | 0.34 | 0.10 | 0 | Endothelial |
| PPP1R16B | 0.98 | 0.00 | 0.00 | 0.28 | 0.05 | 0 | Endothelial |
| S1PR1 | 0.98 | 0.00 | 0.00 | 0.46 | 0.14 | 0 | Endothelial |
| ADCY4 | 0.97 | 0.00 | 0.00 | 0.28 | 0.05 | 0 | Endothelial |
| ETS1 | 0.97 | 0.00 | 0.00 | 0.54 | 0.18 | 0 | Endothelial |
| KIAA0355 | 0.97 | 0.00 | 0.00 | 0.41 | 0.13 | 0 | Endothelial |
| SLC6A6 | 0.96 | 0.00 | 0.00 | 0.45 | 0.19 | 0 | Endothelial |
| ARHGEF15 | 0.94 | 0.00 | 0.00 | 0.26 | 0.04 | 0 | Endothelial |
| RHOJ | 0.94 | 0.00 | 0.00 | 0.42 | 0.13 | 0 | Endothelial |
| TJP1 | 0.93 | 0.00 | 0.00 | 0.39 | 0.13 | 0 | Endothelial |
| TUBB2B | 0.93 | 0.00 | 0.00 | 0.31 | 0.09 | 0 | Endothelial |
| TBX3 | 0.92 | 0.00 | 0.00 | 0.48 | 0.16 | 0 | Endothelial |
| HOXA5 | 0.92 | 0.00 | 0.00 | 0.32 | 0.10 | 0 | Endothelial |
| LXN | 0.91 | 0.00 | 0.00 | 0.31 | 0.09 | 0 | Endothelial |
| ADAM15 | 0.90 | 0.00 | 0.00 | 0.25 | 0.05 | 0 | Endothelial |
| TNFSF10 | 0.90 | 0.00 | 0.00 | 0.29 | 0.07 | 0 | Endothelial |
| MCAM | 0.89 | 0.00 | 0.00 | 0.36 | 0.09 | 0 | Endothelial |
| GRAP | 0.89 | 0.00 | 0.00 | 0.34 | 0.08 | 0 | Endothelial |
| ELK3 | 0.89 | 0.00 | 0.00 | 0.34 | 0.10 | 0 | Endothelial |
| KANK3 | 0.89 | 0.00 | 0.00 | 0.34 | 0.10 | 0 | Endothelial |
| FKBP1A | 0.89 | 0.00 | 0.00 | 0.71 | 0.38 | 0 | Endothelial |
| CAV2 | 0.88 | 0.00 | 0.00 | 0.48 | 0.17 | 0 | Endothelial |
| CRIP2 | 0.88 | 0.00 | 0.00 | 0.72 | 0.38 | 0 | Endothelial |
| KLF2 | 0.88 | 0.00 | 0.00 | 0.40 | 0.16 | 0 | Endothelial |
| CD34 | 0.88 | 0.00 | 0.00 | 0.45 | 0.16 | 0 | Endothelial |
| PEG3 | 0.88 | 0.00 | 0.00 | 0.49 | 0.22 | 0 | Endothelial |
| PIEZO2 | 0.88 | 0.00 | 0.00 | 0.29 | 0.08 | 0 | Endothelial |
| PTPRM | 0.88 | 0.00 | 0.00 | 0.25 | 0.07 | 0 | Endothelial |
| AHR | 0.87 | 0.00 | 0.00 | 0.27 | 0.06 | 0 | Endothelial |
| GNG11 | 0.86 | 0.00 | 0.00 | 0.60 | 0.26 | 0 | Endothelial |
| GIMAP1 | 0.86 | 0.00 | 0.00 | 0.35 | 0.09 | 0 | Endothelial |
| PLCB4 | 0.86 | 0.00 | 0.00 | 0.28 | 0.09 | 0 | Endothelial |
| TCF4 | 0.86 | 0.00 | 0.00 | 0.67 | 0.37 | 0 | Endothelial |
| PTPRG | 0.86 | 0.00 | 0.00 | 0.29 | 0.09 | 0 | Endothelial |
| RGS12 | 0.86 | 0.00 | 0.00 | 0.30 | 0.08 | 0 | Endothelial |
| LY6E | 0.85 | 0.00 | 0.00 | 0.77 | 0.36 | 0 | Endothelial |
| HEG1 | 0.85 | 0.00 | 0.00 | 0.36 | 0.12 | 0 | Endothelial |
| SLFN5 | 0.85 | 0.00 | 0.00 | 0.28 | 0.07 | 0 | Endothelial |
| ARHGAP29 | 0.85 | 0.00 | 0.00 | 0.45 | 0.19 | 0 | Endothelial |
| SNRK | 0.85 | 0.00 | 0.00 | 0.35 | 0.12 | 0 | Endothelial |
| SLK | 0.84 | 0.00 | 0.00 | 0.63 | 0.36 | 0 | Endothelial |
| DDAH2 | 0.84 | 0.00 | 0.00 | 0.51 | 0.24 | 0 | Endothelial |
| MARCKS | 0.84 | 0.00 | 0.00 | 0.75 | 0.45 | 0 | Endothelial |
| FOXF1 | 0.84 | 0.00 | 0.00 | 0.31 | 0.10 | 0 | Endothelial |
| JUP | 0.83 | 0.00 | 0.00 | 0.36 | 0.12 | 0 | Endothelial |
| MEIS2 | 0.83 | 0.00 | 0.00 | 0.27 | 0.09 | 0 | Endothelial |
| NOTCH1 | 0.82 | 0.00 | 0.00 | 0.29 | 0.09 | 0 | Endothelial |
| SEMA6A | 0.82 | 0.00 | 0.00 | 0.30 | 0.08 | 0 | Endothelial |
| SRGN | 0.82 | 0.00 | 0.00 | 0.67 | 0.25 | 0 | Endothelial |
| FNBP1L | 0.81 | 0.00 | 0.00 | 0.44 | 0.21 | 0 | Endothelial |
| RAPGEF5 | 0.81 | 0.00 | 0.00 | 0.28 | 0.08 | 0 | Endothelial |
| KDR | 0.80 | 0.00 | 0.00 | 0.45 | 0.11 | 0 | Endothelial |
| TCN2 | 0.80 | 0.00 | 0.00 | 0.27 | 0.08 | 0 | Endothelial |
| FLI1 | 0.80 | 0.00 | 0.00 | 0.29 | 0.08 | 0 | Endothelial |
| PPP1R2 | 0.79 | 0.00 | 0.00 | 0.45 | 0.20 | 0 | Endothelial |
| ENG | 0.79 | 0.00 | 0.00 | 0.30 | 0.10 | 0 | Endothelial |
| SPTBN1 | 0.79 | 0.00 | 0.00 | 0.78 | 0.49 | 0 | Endothelial |
| KLF7 | 0.79 | 0.00 | 0.00 | 0.37 | 0.14 | 0 | Endothelial |
| COL4A1 | 0.78 | 0.00 | 0.00 | 0.76 | 0.46 | 0 | Endothelial |
| GIMAP6 | 0.78 | 0.00 | 0.00 | 0.38 | 0.11 | 0 | Endothelial |
| COL4A2 | 0.77 | 0.00 | 0.00 | 0.70 | 0.40 | 0 | Endothelial |
| NAV1 | 0.77 | 0.00 | 0.00 | 0.26 | 0.08 | 0 | Endothelial |
| BST2 | 0.75 | 0.00 | 0.00 | 0.34 | 0.12 | 0 | Endothelial |
| CYTH3 | 0.75 | 0.00 | 0.00 | 0.44 | 0.19 | 0 | Endothelial |
| ANP32A | 0.74 | 0.00 | 0.00 | 0.78 | 0.54 | 0 | Endothelial |
| GIMAP4 | 0.73 | 0.00 | 0.00 | 0.42 | 0.13 | 0 | Endothelial |
| CLIC5 | 0.72 | 0.00 | 0.00 | 0.53 | 0.17 | 0 | Endothelial |
| ITM2B | 0.71 | 0.00 | 0.00 | 0.90 | 0.75 | 0 | Endothelial |
| SPARCL1 | 0.68 | 0.00 | 0.00 | 0.63 | 0.37 | 0 | Endothelial |
| GNAI2 | 0.65 | 0.00 | 0.00 | 0.60 | 0.34 | 0 | Endothelial |
| JUN | 0.63 | 0.00 | 0.00 | 0.62 | 0.33 | 0 | Endothelial |
| NRP1 | 0.63 | 0.00 | 0.00 | 0.60 | 0.35 | 0 | Endothelial |
| KITLG | 0.56 | 0.00 | 0.00 | 0.45 | 0.17 | 0 | Endothelial |
| VIM | 0.42 | 0.00 | 0.00 | 0.89 | 0.65 | 0 | Endothelial |
| ARHGEF12 | 0.78 | 0.00 | 0.00 | 0.42 | 0.20 | 0 | Endothelial |
| PREX2 | 0.78 | 0.00 | 0.00 | 0.34 | 0.14 | 0 | Endothelial |
| GCC2 | 0.78 | 0.00 | 0.00 | 0.42 | 0.21 | 0 | Endothelial |
| CNN3 | 0.58 | 0.00 | 0.00 | 0.54 | 0.29 | 0 | Endothelial |
| CCDC85B | 0.71 | 0.00 | 0.00 | 0.30 | 0.11 | 0 | Endothelial |
| STMN1 | 0.75 | 0.00 | 0.00 | 0.42 | 0.21 | 0 | Endothelial |
| PLTP | 0.75 | 0.00 | 0.00 | 0.34 | 0.14 | 0 | Endothelial |
| CD9 | 0.47 | 0.00 | 0.00 | 0.58 | 0.32 | 0 | Endothelial |
| GALNT18 | 0.70 | 0.00 | 0.00 | 0.25 | 0.08 | 0 | Endothelial |
| CYB5R3 | 0.72 | 0.00 | 0.00 | 0.57 | 0.36 | 0 | Endothelial |
| HIP1 | 0.73 | 0.00 | 0.00 | 0.31 | 0.12 | 0 | Endothelial |
| DNAJC8 | 0.65 | 0.00 | 0.00 | 0.60 | 0.39 | 0 | Endothelial |
| SMARCA2 | 0.74 | 0.00 | 0.00 | 0.51 | 0.30 | 0 | Endothelial |
| ZEB1 | 0.73 | 0.00 | 0.00 | 0.25 | 0.09 | 0 | Endothelial |
| RAB11A | 0.68 | 0.00 | 0.00 | 0.45 | 0.24 | 0 | Endothelial |
| CTNNA1 | 0.66 | 0.00 | 0.00 | 0.54 | 0.33 | 0 | Endothelial |
| MYL12B | 0.51 | 0.00 | 0.00 | 0.75 | 0.60 | 0 | Endothelial |
| DLC1 | 0.69 | 0.00 | 0.00 | 0.34 | 0.15 | 0 | Endothelial |
| RGCC | 0.45 | 0.00 | 0.00 | 0.37 | 0.16 | 0 | Endothelial |
| ITGB1 | 0.43 | 0.00 | 0.00 | 0.79 | 0.61 | 0 | Endothelial |
| CARHSP1 | 0.69 | 0.00 | 0.00 | 0.30 | 0.13 | 0 | Endothelial |
| UACA | 0.70 | 0.00 | 0.00 | 0.29 | 0.12 | 0 | Endothelial |
| LUZP1 | 0.71 | 0.00 | 0.00 | 0.35 | 0.17 | 0 | Endothelial |
| SNX3 | 0.62 | 0.00 | 0.00 | 0.52 | 0.32 | 0 | Endothelial |
| PEA15 | 0.65 | 0.00 | 0.00 | 0.29 | 0.12 | 0 | Endothelial |
| PLCG1 | 0.68 | 0.00 | 0.00 | 0.30 | 0.13 | 0 | Endothelial |
| NES | 0.46 | 0.00 | 0.00 | 0.39 | 0.18 | 0 | Endothelial |
| RHOC | 0.62 | 0.00 | 0.00 | 0.40 | 0.21 | 0 | Endothelial |
| EFNB1 | 0.68 | 0.00 | 0.00 | 0.29 | 0.12 | 0 | Endothelial |
| HDAC7 | 0.64 | 0.00 | 0.00 | 0.27 | 0.11 | 0 | Endothelial |
| SASH1 | 0.68 | 0.00 | 0.00 | 0.26 | 0.11 | 0 | Endothelial |
| BTBD3 | 0.67 | 0.00 | 0.00 | 0.28 | 0.12 | 0 | Endothelial |
| ABLIM1 | 0.65 | 0.00 | 0.00 | 0.39 | 0.20 | 0 | Endothelial |
| PLK2 | 0.69 | 0.00 | 0.00 | 0.31 | 0.14 | 0 | Endothelial |
| MGLL | 0.64 | 0.00 | 0.00 | 0.31 | 0.15 | 0 | Endothelial |
| RAP1A | 0.54 | 0.00 | 0.00 | 0.46 | 0.27 | 0 | Endothelial |
| HES1 | 0.61 | 0.00 | 0.00 | 0.45 | 0.26 | 0 | Endothelial |
| CLIC1 | 0.52 | 0.00 | 0.00 | 0.55 | 0.36 | 0 | Endothelial |
| ACTN4 | 0.60 | 0.00 | 0.00 | 0.45 | 0.27 | 0 | Endothelial |
| WASF2 | 0.59 | 0.00 | 0.00 | 0.46 | 0.28 | 0 | Endothelial |
| TAX1BP1 | 0.55 | 0.00 | 0.00 | 0.71 | 0.58 | 0 | Endothelial |
| FERMT2 | 0.44 | 0.00 | 0.00 | 0.56 | 0.36 | 0 | Endothelial |
| CD151 | 0.57 | 0.00 | 0.00 | 0.33 | 0.17 | 0 | Endothelial |
| ELOVL5 | 0.61 | 0.00 | 0.00 | 0.27 | 0.13 | 0 | Endothelial |
| BNIP2 | 0.56 | 0.00 | 0.00 | 0.43 | 0.26 | 0 | Endothelial |
| XIAP | 0.62 | 0.00 | 0.00 | 0.32 | 0.17 | 0 | Endothelial |
| ARL2BP | 0.64 | 0.00 | 0.00 | 0.26 | 0.12 | 0 | Endothelial |
| APLP2 | 0.52 | 0.00 | 0.00 | 0.55 | 0.38 | 0 | Endothelial |
| ANXA3 | 0.44 | 0.00 | 0.00 | 0.37 | 0.19 | 0 | Endothelial |
| TMEM204 | 0.50 | 0.00 | 0.00 | 0.29 | 0.14 | 0 | Endothelial |
| CTNND1 | 0.58 | 0.00 | 0.00 | 0.29 | 0.15 | 0 | Endothelial |
| DUSP6 | 0.54 | 0.00 | 0.00 | 0.31 | 0.16 | 0 | Endothelial |
| CBFA2T3 | 0.49 | 0.00 | 0.00 | 0.26 | 0.12 | 0 | Endothelial |
| GUK1 | 0.64 | 0.00 | 0.00 | 0.32 | 0.17 | 0 | Endothelial |
| MAPK3 | 0.59 | 0.00 | 0.00 | 0.25 | 0.12 | 0 | Endothelial |
| SH3GLB1 | 0.51 | 0.00 | 0.00 | 0.52 | 0.35 | 0 | Endothelial |
| LEPROT | 0.54 | 0.00 | 0.00 | 0.48 | 0.31 | 0 | Endothelial |
| ITPRIPL2 | 0.62 | 0.00 | 0.00 | 0.29 | 0.14 | 0 | Endothelial |
| ZNF503 | 0.59 | 0.00 | 0.00 | 0.28 | 0.14 | 0 | Endothelial |
| ACAP2 | 0.58 | 0.00 | 0.00 | 0.31 | 0.16 | 0 | Endothelial |
| ATP1B3 | 0.52 | 0.00 | 0.00 | 0.40 | 0.23 | 0 | Endothelial |
| MAP7D1 | 0.58 | 0.00 | 0.00 | 0.31 | 0.16 | 0 | Endothelial |
| SPTAN1 | 0.58 | 0.00 | 0.00 | 0.29 | 0.15 | 0 | Endothelial |
| SYPL1 | 0.54 | 0.00 | 0.00 | 0.39 | 0.24 | 0 | Endothelial |
| ADD3 | 0.48 | 0.00 | 0.00 | 0.41 | 0.25 | 0 | Endothelial |
| MXRA7 | 0.50 | 0.00 | 0.00 | 0.26 | 0.12 | 0 | Endothelial |
| SERINC3 | 0.61 | 0.00 | 0.00 | 0.43 | 0.28 | 0 | Endothelial |
| CSRP2 | 0.45 | 0.00 | 0.00 | 0.28 | 0.14 | 0 | Endothelial |
| IER2 | 0.47 | 0.00 | 0.00 | 0.39 | 0.23 | 0 | Endothelial |
| MXD4 | 0.55 | 0.00 | 0.00 | 0.47 | 0.34 | 0 | Endothelial |
| CLIC4 | 0.49 | 0.00 | 0.00 | 0.42 | 0.28 | 0 | Endothelial |
| TPM4 | 0.37 | 0.00 | 0.00 | 0.63 | 0.49 | 0 | Endothelial |
| DNAJA1 | 0.38 | 0.00 | 0.00 | 0.62 | 0.49 | 0 | Endothelial |
| AMOTL1 | 0.49 | 0.00 | 0.00 | 0.26 | 0.14 | 0 | Endothelial |
| GLUL | 0.48 | 0.00 | 0.00 | 0.33 | 0.20 | 0 | Endothelial |
| IVNS1ABP | 0.74 | 0.00 | 0.00 | 0.32 | 0.19 | 0 | Endothelial |
| NFKBIA | 0.46 | 0.00 | 0.00 | 0.48 | 0.34 | 0 | Endothelial |
| DRAM2 | 0.52 | 0.00 | 0.00 | 0.26 | 0.15 | 0 | Endothelial |
| TPM3 | 0.25 | 0.00 | 0.00 | 0.77 | 0.65 | 0 | Endothelial |
| MSN | 0.33 | 0.00 | 0.00 | 0.57 | 0.42 | 0 | Endothelial |
| ADAMTS1 | 0.47 | 0.00 | 0.00 | 0.31 | 0.18 | 0 | Endothelial |
| EDNRB | 0.37 | 0.00 | 0.00 | 0.27 | 0.14 | 0 | Endothelial |
| MEF2A | 0.40 | 0.00 | 0.00 | 0.46 | 0.32 | 0 | Endothelial |
| CSNK1A1 | 0.35 | 0.00 | 0.00 | 0.62 | 0.49 | 0 | Endothelial |
| HMG20B | 0.46 | 0.00 | 0.00 | 0.44 | 0.31 | 0 | Endothelial |
| ZMIZ1 | 0.46 | 0.00 | 0.00 | 0.31 | 0.19 | 0 | Endothelial |
| CTNNB1 | 0.42 | 0.00 | 0.00 | 0.46 | 0.33 | 0 | Endothelial |
| TOP1 | 0.42 | 0.00 | 0.00 | 0.53 | 0.41 | 0 | Endothelial |
| MARCKSL1 | 0.47 | 0.00 | 0.00 | 0.33 | 0.21 | 0 | Endothelial |
| SOX4 | 0.35 | 0.00 | 0.00 | 0.52 | 0.38 | 0 | Endothelial |
| EHD2 | 0.43 | 0.00 | 0.00 | 0.26 | 0.15 | 0 | Endothelial |
| IGFBP4 | 0.28 | 0.00 | 0.00 | 0.39 | 0.25 | 0 | Endothelial |
| RHOA | 0.33 | 0.00 | 0.00 | 0.57 | 0.47 | 0 | Endothelial |
| S100A13 | 0.41 | 0.00 | 0.00 | 0.37 | 0.25 | 0 | Endothelial |
| RELL1 | 0.41 | 0.00 | 0.00 | 0.28 | 0.17 | 0 | Endothelial |
| ARGLU1 | 0.38 | 0.00 | 0.00 | 0.51 | 0.40 | 0 | Endothelial |
| QKI | 0.38 | 0.00 | 0.00 | 0.39 | 0.27 | 0 | Endothelial |
| CD2AP | 0.38 | 0.00 | 0.00 | 0.42 | 0.30 | 0 | Endothelial |
| ATP2B1 | 0.32 | 0.00 | 0.00 | 0.65 | 0.54 | 0 | Endothelial |
| DYNLL1 | 0.33 | 0.00 | 0.00 | 0.56 | 0.45 | 0 | Endothelial |
| RALB | 0.37 | 0.00 | 0.00 | 0.28 | 0.18 | 0 | Endothelial |
| ARHGEF2 | 0.40 | 0.00 | 0.00 | 0.28 | 0.18 | 0 | Endothelial |
| SNRPB | 0.36 | 0.00 | 0.00 | 0.45 | 0.34 | 0 | Endothelial |
| RBPMS | 0.39 | 0.00 | 0.00 | 0.27 | 0.17 | 0 | Endothelial |
| NDUFA8 | 0.40 | 0.00 | 0.00 | 0.34 | 0.23 | 0 | Endothelial |
| FUBP1 | 0.41 | 0.00 | 0.00 | 0.32 | 0.22 | 0 | Endothelial |
| RNF7 | 0.40 | 0.00 | 0.00 | 0.34 | 0.24 | 0 | Endothelial |
| HOOK3 | 0.40 | 0.00 | 0.00 | 0.31 | 0.20 | 0 | Endothelial |
| H1F0 | 0.43 | 0.00 | 0.00 | 0.35 | 0.25 | 0 | Endothelial |
| PHLDB2 | 0.36 | 0.00 | 0.00 | 0.37 | 0.26 | 0 | Endothelial |
| ANP32E | 0.36 | 0.00 | 0.00 | 0.35 | 0.24 | 0 | Endothelial |
| YWHAB | 0.28 | 0.00 | 0.00 | 0.65 | 0.56 | 0 | Endothelial |
| GNB1 | 0.39 | 0.00 | 0.00 | 0.30 | 0.20 | 0 | Endothelial |
| PHF20L1 | 0.39 | 0.00 | 0.00 | 0.35 | 0.25 | 0 | Endothelial |
| TXNIP | 0.37 | 0.00 | 0.00 | 0.36 | 0.25 | 0 | Endothelial |
| ITSN2 | 0.32 | 0.00 | 0.00 | 0.32 | 0.22 | 0 | Endothelial |
| OAZ2 | 0.41 | 0.00 | 0.00 | 0.39 | 0.29 | 0 | Endothelial |
| FCHO2 | 0.35 | 0.00 | 0.00 | 0.25 | 0.16 | 0 | Endothelial |
| RAB6A | 0.38 | 0.00 | 0.00 | 0.28 | 0.20 | 0 | Endothelial |
| ARPC3 | 0.33 | 0.00 | 0.00 | 0.37 | 0.27 | 0 | Endothelial |
| NAP1L4 | 0.33 | 0.00 | 0.00 | 0.33 | 0.24 | 0 | Endothelial |
| ZNF326 | 0.35 | 0.00 | 0.00 | 0.29 | 0.21 | 0 | Endothelial |
| DYNC1I2 | 0.29 | 0.00 | 0.00 | 0.55 | 0.47 | 0 | Endothelial |
| ARF1 | 0.30 | 0.00 | 0.00 | 0.51 | 0.43 | 0 | Endothelial |
| MAP4K4 | 0.38 | 0.00 | 0.00 | 0.25 | 0.17 | 0 | Endothelial |
| MOCS2 | 0.34 | 0.00 | 0.00 | 0.25 | 0.17 | 0 | Endothelial |
| HNRNPF | 0.28 | 0.00 | 0.00 | 0.42 | 0.33 | 0 | Endothelial |
| AKR1A1 | 0.26 | 0.00 | 0.00 | 0.55 | 0.47 | 0 | Endothelial |
| EIF4G2 | 0.31 | 0.00 | 0.00 | 0.49 | 0.41 | 0 | Endothelial |
| ZC3H7A | 0.31 | 0.00 | 0.00 | 0.34 | 0.25 | 0 | Endothelial |
| CFLAR | 0.31 | 0.00 | 0.00 | 0.26 | 0.18 | 0 | Endothelial |
| C5orf24 | 0.33 | 0.00 | 0.00 | 0.34 | 0.25 | 0 | Endothelial |
| G3BP2 | 0.30 | 0.00 | 0.00 | 0.36 | 0.27 | 0 | Endothelial |
| ZCRB1 | 0.33 | 0.00 | 0.00 | 0.35 | 0.27 | 0 | Endothelial |
| TMCO1 | 0.32 | 0.00 | 0.00 | 0.37 | 0.28 | 0 | Endothelial |
| GNB2 | 0.30 | 0.00 | 0.00 | 0.44 | 0.37 | 0 | Endothelial |
| KTN1 | 0.30 | 0.00 | 0.00 | 0.45 | 0.38 | 0 | Endothelial |
| ZBTB20 | 0.31 | 0.00 | 0.00 | 0.34 | 0.26 | 0 | Endothelial |
| CBX1 | 0.29 | 0.00 | 0.00 | 0.46 | 0.39 | 0 | Endothelial |
| SEPT2 | 0.26 | 0.00 | 0.00 | 0.43 | 0.36 | 0 | Endothelial |
| SDCBP | 0.25 | 0.00 | 0.00 | 0.35 | 0.28 | 0 | Endothelial |
| SERINC1 | 0.26 | 0.00 | 0.00 | 0.35 | 0.28 | 0 | Endothelial |
| CCDC50 | 0.29 | 0.00 | 0.00 | 0.31 | 0.24 | 0 | Endothelial |
| PRKAR1A | 0.26 | 0.00 | 0.00 | 0.39 | 0.32 | 0 | Endothelial |
| VPS36 | 0.28 | 0.00 | 0.00 | 0.33 | 0.26 | 0 | Endothelial |
| RAD21 | 0.27 | 0.00 | 0.00 | 0.27 | 0.21 | 0 | Endothelial |
| STRN3 | 0.27 | 0.00 | 0.00 | 0.25 | 0.19 | 0 | Endothelial |
| RSU1 | 0.26 | 0.00 | 0.00 | 0.26 | 0.20 | 0 | Endothelial |
| BAZ2B | 0.25 | 0.00 | 0.00 | 0.32 | 0.27 | 0 | Endothelial |
| MFAP4 | 3.12 | 0.00 | 0.00 | 0.97 | 0.14 | 1 | Matrix Fibroblast |
| ADH1A | 3.02 | 0.00 | 0.00 | 0.85 | 0.04 | 1 | Matrix Fibroblast |
| DPT | 2.74 | 0.00 | 0.00 | 0.76 | 0.08 | 1 | Matrix Fibroblast |
| TCF21 | 2.70 | 0.00 | 0.00 | 0.87 | 0.07 | 1 | Matrix Fibroblast |
| LIMCH1 | 2.46 | 0.00 | 0.00 | 0.94 | 0.19 | 1 | Matrix Fibroblast |
| GYG1 | 2.36 | 0.00 | 0.00 | 0.79 | 0.10 | 1 | Matrix Fibroblast |
| CDH11 | 2.28 | 0.00 | 0.00 | 0.79 | 0.10 | 1 | Matrix Fibroblast |
| FMO2 | 2.27 | 0.00 | 0.00 | 0.60 | 0.07 | 1 | Matrix Fibroblast |
| FHL1 | 2.21 | 0.00 | 0.00 | 0.91 | 0.21 | 1 | Matrix Fibroblast |
| MAF | 2.20 | 0.00 | 0.00 | 0.76 | 0.08 | 1 | Matrix Fibroblast |
| MACF1 | 2.05 | 0.00 | 0.00 | 0.93 | 0.44 | 1 | Matrix Fibroblast |
| NPNT | 2.02 | 0.00 | 0.00 | 0.77 | 0.15 | 1 | Matrix Fibroblast |
| G0S2 | 1.99 | 0.00 | 0.00 | 0.61 | 0.06 | 1 | Matrix Fibroblast |
| INMT | 1.98 | 0.00 | 0.00 | 0.26 | 0.03 | 1 | Matrix Fibroblast |
| LBH | 1.95 | 0.00 | 0.00 | 0.75 | 0.12 | 1 | Matrix Fibroblast |
| ENPEP | 1.94 | 0.00 | 0.00 | 0.53 | 0.04 | 1 | Matrix Fibroblast |
| NEXN | 1.93 | 0.00 | 0.00 | 0.73 | 0.13 | 1 | Matrix Fibroblast |
| PCOLCE2 | 1.92 | 0.00 | 0.00 | 0.50 | 0.02 | 1 | Matrix Fibroblast |
| PLAC9 | 1.90 | 0.00 | 0.00 | 0.88 | 0.21 | 1 | Matrix Fibroblast |
| COL13A1 | 1.88 | 0.00 | 0.00 | 0.52 | 0.03 | 1 | Matrix Fibroblast |
| GSN | 1.88 | 0.00 | 0.00 | 0.74 | 0.17 | 1 | Matrix Fibroblast |
| OGN | 1.83 | 0.00 | 0.00 | 0.60 | 0.06 | 1 | Matrix Fibroblast |
| PLXDC2 | 1.82 | 0.00 | 0.00 | 0.56 | 0.06 | 1 | Matrix Fibroblast |
| MEOX2 | 1.78 | 0.00 | 0.00 | 0.48 | 0.03 | 1 | Matrix Fibroblast |
| GPX3 | 1.73 | 0.00 | 0.00 | 0.50 | 0.09 | 1 | Matrix Fibroblast |
| CPM | 1.70 | 0.00 | 0.00 | 0.56 | 0.10 | 1 | Matrix Fibroblast |
| COLEC12 | 1.65 | 0.00 | 0.00 | 0.48 | 0.05 | 1 | Matrix Fibroblast |
| PTPRD | 1.63 | 0.00 | 0.00 | 0.48 | 0.05 | 1 | Matrix Fibroblast |
| GPC3 | 1.63 | 0.00 | 0.00 | 0.66 | 0.10 | 1 | Matrix Fibroblast |
| ATP1A2 | 1.59 | 0.00 | 0.00 | 0.44 | 0.04 | 1 | Matrix Fibroblast |
| HSD11B1 | 1.58 | 0.00 | 0.00 | 0.38 | 0.03 | 1 | Matrix Fibroblast |
| SLIT2 | 1.58 | 0.00 | 0.00 | 0.40 | 0.03 | 1 | Matrix Fibroblast |
| CLEC3B | 1.58 | 0.00 | 0.00 | 0.38 | 0.02 | 1 | Matrix Fibroblast |
| OLFML3 | 1.51 | 0.00 | 0.00 | 0.42 | 0.05 | 1 | Matrix Fibroblast |
| PHLDA1 | 1.50 | 0.00 | 0.00 | 0.58 | 0.14 | 1 | Matrix Fibroblast |
| SERPING1 | 1.50 | 0.00 | 0.00 | 0.53 | 0.08 | 1 | Matrix Fibroblast |
| ITGA8 | 1.50 | 0.00 | 0.00 | 0.44 | 0.04 | 1 | Matrix Fibroblast |
| PALLD | 1.48 | 0.00 | 0.00 | 0.58 | 0.11 | 1 | Matrix Fibroblast |
| MGP | 1.47 | 0.00 | 0.00 | 0.92 | 0.24 | 1 | Matrix Fibroblast |
| CPED1 | 1.46 | 0.00 | 0.00 | 0.41 | 0.04 | 1 | Matrix Fibroblast |
| SLC38A5 | 1.44 | 0.00 | 0.00 | 0.35 | 0.01 | 1 | Matrix Fibroblast |
| SH3BGRL | 1.42 | 0.00 | 0.00 | 0.83 | 0.40 | 1 | Matrix Fibroblast |
| SLC27A6 | 1.39 | 0.00 | 0.00 | 0.29 | 0.00 | 1 | Matrix Fibroblast |
| AKAP12 | 1.38 | 0.00 | 0.00 | 0.64 | 0.18 | 1 | Matrix Fibroblast |
| VCAM1 | 1.37 | 0.00 | 0.00 | 0.32 | 0.02 | 1 | Matrix Fibroblast |
| SNAI2 | 1.36 | 0.00 | 0.00 | 0.42 | 0.07 | 1 | Matrix Fibroblast |
| SPON1 | 1.35 | 0.00 | 0.00 | 0.36 | 0.03 | 1 | Matrix Fibroblast |
| MXRA8 | 1.34 | 0.00 | 0.00 | 0.53 | 0.11 | 1 | Matrix Fibroblast |
| RAP2A | 1.34 | 0.00 | 0.00 | 0.47 | 0.10 | 1 | Matrix Fibroblast |
| WNT2 | 1.34 | 0.00 | 0.00 | 0.32 | 0.02 | 1 | Matrix Fibroblast |
| PPP1R14A | 1.33 | 0.00 | 0.00 | 0.40 | 0.06 | 1 | Matrix Fibroblast |
| FBLIM1 | 1.31 | 0.00 | 0.00 | 0.51 | 0.11 | 1 | Matrix Fibroblast |
| MDK | 1.31 | 0.00 | 0.00 | 0.58 | 0.16 | 1 | Matrix Fibroblast |
| COL1A2 | 1.31 | 0.00 | 0.00 | 0.97 | 0.32 | 1 | Matrix Fibroblast |
| NR2F2 | 1.30 | 0.00 | 0.00 | 0.56 | 0.15 | 1 | Matrix Fibroblast |
| DPEP1 | 1.29 | 0.00 | 0.00 | 0.26 | 0.02 | 1 | Matrix Fibroblast |
| CDO1 | 1.28 | 0.00 | 0.00 | 0.28 | 0.03 | 1 | Matrix Fibroblast |
| MYH10 | 1.28 | 0.00 | 0.00 | 0.61 | 0.16 | 1 | Matrix Fibroblast |
| ADARB1 | 1.27 | 0.00 | 0.00 | 0.32 | 0.03 | 1 | Matrix Fibroblast |
| CACNA1D | 1.27 | 0.00 | 0.00 | 0.29 | 0.02 | 1 | Matrix Fibroblast |
| SPECC1L-ADORA2A | 1.27 | 0.00 | 0.00 | 0.37 | 0.06 | 1 | Matrix Fibroblast |
| VLDLR | 1.27 | 0.00 | 0.00 | 0.36 | 0.06 | 1 | Matrix Fibroblast |
| SERPINE2 | 1.27 | 0.00 | 0.00 | 0.70 | 0.18 | 1 | Matrix Fibroblast |
| TNS1 | 1.27 | 0.00 | 0.00 | 0.57 | 0.17 | 1 | Matrix Fibroblast |
| SEPT4 | 1.24 | 0.00 | 0.00 | 0.63 | 0.16 | 1 | Matrix Fibroblast |
| ABLIM3 | 1.23 | 0.00 | 0.00 | 0.28 | 0.02 | 1 | Matrix Fibroblast |
| HIVEP3 | 1.18 | 0.00 | 0.00 | 0.30 | 0.04 | 1 | Matrix Fibroblast |
| ANGPT1 | 1.16 | 0.00 | 0.00 | 0.31 | 0.05 | 1 | Matrix Fibroblast |
| VCAN | 1.12 | 0.00 | 0.00 | 0.40 | 0.09 | 1 | Matrix Fibroblast |
| EDNRA | 1.12 | 0.00 | 0.00 | 0.33 | 0.06 | 1 | Matrix Fibroblast |
| TNS3 | 1.12 | 0.00 | 0.00 | 0.34 | 0.07 | 1 | Matrix Fibroblast |
| TMEM119 | 1.10 | 0.00 | 0.00 | 0.26 | 0.02 | 1 | Matrix Fibroblast |
| FXYD1 | 1.09 | 0.00 | 0.00 | 0.34 | 0.06 | 1 | Matrix Fibroblast |
| FIBIN | 1.09 | 0.00 | 0.00 | 0.37 | 0.07 | 1 | Matrix Fibroblast |
| CDKN1C | 1.09 | 0.00 | 0.00 | 0.44 | 0.13 | 1 | Matrix Fibroblast |
| MYLK | 1.08 | 0.00 | 0.00 | 0.69 | 0.23 | 1 | Matrix Fibroblast |
| BGN | 1.08 | 0.00 | 0.00 | 0.67 | 0.20 | 1 | Matrix Fibroblast |
| ZYX | 1.08 | 0.00 | 0.00 | 0.50 | 0.16 | 1 | Matrix Fibroblast |
| TRIL | 1.07 | 0.00 | 0.00 | 0.28 | 0.04 | 1 | Matrix Fibroblast |
| SELENBP1 | 1.06 | 0.00 | 0.00 | 0.55 | 0.15 | 1 | Matrix Fibroblast |
| COL6A1 | 1.03 | 0.00 | 0.00 | 0.53 | 0.17 | 1 | Matrix Fibroblast |
| LOXL1 | 1.02 | 0.00 | 0.00 | 0.37 | 0.08 | 1 | Matrix Fibroblast |
| GPM6B | 1.02 | 0.00 | 0.00 | 0.35 | 0.09 | 1 | Matrix Fibroblast |
| NREP | 1.01 | 0.00 | 0.00 | 0.76 | 0.38 | 1 | Matrix Fibroblast |
| PEG3 | 0.99 | 0.00 | 0.00 | 0.54 | 0.22 | 1 | Matrix Fibroblast |
| ACADSB | 0.97 | 0.00 | 0.00 | 0.31 | 0.07 | 1 | Matrix Fibroblast |
| LIMD1 | 0.97 | 0.00 | 0.00 | 0.38 | 0.13 | 1 | Matrix Fibroblast |
| FBN1 | 0.96 | 0.00 | 0.00 | 0.44 | 0.13 | 1 | Matrix Fibroblast |
| TMEM176A | 0.96 | 0.00 | 0.00 | 0.42 | 0.16 | 1 | Matrix Fibroblast |
| NAP1L1 | 0.95 | 0.00 | 0.00 | 0.72 | 0.44 | 1 | Matrix Fibroblast |
| COL6A2 | 0.95 | 0.00 | 0.00 | 0.45 | 0.14 | 1 | Matrix Fibroblast |
| COL1A1 | 0.94 | 0.00 | 0.00 | 0.77 | 0.21 | 1 | Matrix Fibroblast |
| H6PD | 0.93 | 0.00 | 0.00 | 0.26 | 0.06 | 1 | Matrix Fibroblast |
| TMEM254 | 0.92 | 0.00 | 0.00 | 0.38 | 0.14 | 1 | Matrix Fibroblast |
| CSRP1 | 0.92 | 0.00 | 0.00 | 0.45 | 0.15 | 1 | Matrix Fibroblast |
| AGPAT4 | 0.92 | 0.00 | 0.00 | 0.33 | 0.09 | 1 | Matrix Fibroblast |
| PBX1 | 0.91 | 0.00 | 0.00 | 0.41 | 0.17 | 1 | Matrix Fibroblast |
| PDLIM2 | 0.90 | 0.00 | 0.00 | 0.41 | 0.15 | 1 | Matrix Fibroblast |
| CCDC80 | 0.90 | 0.00 | 0.00 | 0.34 | 0.09 | 1 | Matrix Fibroblast |
| LAMB1 | 0.88 | 0.00 | 0.00 | 0.37 | 0.12 | 1 | Matrix Fibroblast |
| CNN2 | 0.87 | 0.00 | 0.00 | 0.66 | 0.34 | 1 | Matrix Fibroblast |
| ISM1 | 0.87 | 0.00 | 0.00 | 0.26 | 0.06 | 1 | Matrix Fibroblast |
| IGFBP4 | 0.86 | 0.00 | 0.00 | 0.56 | 0.23 | 1 | Matrix Fibroblast |
| EIF4EBP1 | 0.86 | 0.00 | 0.00 | 0.32 | 0.10 | 1 | Matrix Fibroblast |
| NFIB | 0.85 | 0.00 | 0.00 | 0.77 | 0.43 | 1 | Matrix Fibroblast |
| FKBP9 | 0.84 | 0.00 | 0.00 | 0.33 | 0.11 | 1 | Matrix Fibroblast |
| ROBO2 | 0.84 | 0.00 | 0.00 | 0.27 | 0.06 | 1 | Matrix Fibroblast |
| PREX2 | 0.84 | 0.00 | 0.00 | 0.40 | 0.13 | 1 | Matrix Fibroblast |
| TBX2 | 0.83 | 0.00 | 0.00 | 0.33 | 0.11 | 1 | Matrix Fibroblast |
| COL6A3 | 0.83 | 0.00 | 0.00 | 0.33 | 0.10 | 1 | Matrix Fibroblast |
| COL3A1 | 0.82 | 0.00 | 0.00 | 0.88 | 0.27 | 1 | Matrix Fibroblast |
| ADAMTS1 | 0.82 | 0.00 | 0.00 | 0.42 | 0.16 | 1 | Matrix Fibroblast |
| PCOLCE | 0.82 | 0.00 | 0.00 | 0.30 | 0.08 | 1 | Matrix Fibroblast |
| LAMA4 | 0.81 | 0.00 | 0.00 | 0.32 | 0.11 | 1 | Matrix Fibroblast |
| PCDHGA9 | 0.79 | 0.00 | 0.00 | 0.36 | 0.13 | 1 | Matrix Fibroblast |
| FBLN5 | 0.79 | 0.00 | 0.00 | 0.45 | 0.14 | 1 | Matrix Fibroblast |
| CD81 | 0.72 | 0.00 | 0.00 | 0.65 | 0.37 | 1 | Matrix Fibroblast |
| DKK3 | 0.72 | 0.00 | 0.00 | 0.32 | 0.11 | 1 | Matrix Fibroblast |
| ADAMTS2 | 0.67 | 0.00 | 0.00 | 0.27 | 0.08 | 1 | Matrix Fibroblast |
| NEDD4 | 0.64 | 0.00 | 0.00 | 0.72 | 0.46 | 1 | Matrix Fibroblast |
| RARRES2 | 0.64 | 0.00 | 0.00 | 0.38 | 0.13 | 1 | Matrix Fibroblast |
| APOE | 0.63 | 0.00 | 0.00 | 0.56 | 0.23 | 1 | Matrix Fibroblast |
| TPM1 | 0.61 | 0.00 | 0.00 | 0.82 | 0.48 | 1 | Matrix Fibroblast |
| RPS4X | 0.53 | 0.00 | 0.00 | 0.97 | 0.86 | 1 | Matrix Fibroblast |
| SPARC | 0.51 | 0.00 | 0.00 | 0.96 | 0.73 | 1 | Matrix Fibroblast |
| SPARCL1 | 0.50 | 0.00 | 0.00 | 0.73 | 0.36 | 1 | Matrix Fibroblast |
| LGALS1 | 0.50 | 0.00 | 0.00 | 0.77 | 0.42 | 1 | Matrix Fibroblast |
| TMSB10 | 0.48 | 0.00 | 0.00 | 0.96 | 0.87 | 1 | Matrix Fibroblast |
| KDELC2 | 0.80 | 0.00 | 0.00 | 0.26 | 0.08 | 1 | Matrix Fibroblast |
| LIMS3 | 0.73 | 0.00 | 0.00 | 0.48 | 0.23 | 1 | Matrix Fibroblast |
| NPR3 | 0.81 | 0.00 | 0.00 | 0.28 | 0.09 | 1 | Matrix Fibroblast |
| MAGED1 | 0.74 | 0.00 | 0.00 | 0.37 | 0.15 | 1 | Matrix Fibroblast |
| COL5A2 | 0.62 | 0.00 | 0.00 | 0.38 | 0.16 | 1 | Matrix Fibroblast |
| NISCH | 0.73 | 0.00 | 0.00 | 0.48 | 0.24 | 1 | Matrix Fibroblast |
| COX6C | 0.54 | 0.00 | 0.00 | 0.81 | 0.65 | 1 | Matrix Fibroblast |
| IGFBP7 | 0.28 | 0.00 | 0.00 | 0.69 | 0.39 | 1 | Matrix Fibroblast |
| SH3BP5 | 0.63 | 0.00 | 0.00 | 0.39 | 0.17 | 1 | Matrix Fibroblast |
| PRDX5 | 0.51 | 0.00 | 0.00 | 0.54 | 0.30 | 1 | Matrix Fibroblast |
| RBMS3 | 0.70 | 0.00 | 0.00 | 0.35 | 0.15 | 1 | Matrix Fibroblast |
| LRP1 | 0.68 | 0.00 | 0.00 | 0.28 | 0.10 | 1 | Matrix Fibroblast |
| MBNL2 | 0.67 | 0.00 | 0.00 | 0.52 | 0.30 | 1 | Matrix Fibroblast |
| CRYAB | 0.54 | 0.00 | 0.00 | 0.33 | 0.13 | 1 | Matrix Fibroblast |
| LMO4 | 0.64 | 0.00 | 0.00 | 0.45 | 0.23 | 1 | Matrix Fibroblast |
| N6AMT1 | 0.72 | 0.00 | 0.00 | 0.26 | 0.09 | 1 | Matrix Fibroblast |
| NENF | 0.74 | 0.00 | 0.00 | 0.37 | 0.17 | 1 | Matrix Fibroblast |
| FGFR1 | 0.72 | 0.00 | 0.00 | 0.25 | 0.09 | 1 | Matrix Fibroblast |
| SOCS2 | 0.57 | 0.00 | 0.00 | 0.53 | 0.29 | 1 | Matrix Fibroblast |
| MFAP2 | 0.50 | 0.00 | 0.00 | 0.58 | 0.32 | 1 | Matrix Fibroblast |
| TAGLN2 | 0.53 | 0.00 | 0.00 | 0.66 | 0.44 | 1 | Matrix Fibroblast |
| TACC1 | 0.69 | 0.00 | 0.00 | 0.41 | 0.21 | 1 | Matrix Fibroblast |
| NID1 | 0.59 | 0.00 | 0.00 | 0.42 | 0.20 | 1 | Matrix Fibroblast |
| LTBP4 | 0.61 | 0.00 | 0.00 | 0.26 | 0.10 | 1 | Matrix Fibroblast |
| ZFP36L1 | 0.64 | 0.00 | 0.00 | 0.64 | 0.44 | 1 | Matrix Fibroblast |
| CELF2 | 0.57 | 0.00 | 0.00 | 0.45 | 0.24 | 1 | Matrix Fibroblast |
| PLTP | 0.58 | 0.00 | 0.00 | 0.34 | 0.15 | 1 | Matrix Fibroblast |
| EVA1B | 0.61 | 0.00 | 0.00 | 0.36 | 0.18 | 1 | Matrix Fibroblast |
| IGF1 | 0.43 | 0.00 | 0.00 | 0.29 | 0.11 | 1 | Matrix Fibroblast |
| CD302 | 0.56 | 0.00 | 0.00 | 0.29 | 0.12 | 1 | Matrix Fibroblast |
| S100A1 | 0.61 | 0.00 | 0.00 | 0.26 | 0.11 | 1 | Matrix Fibroblast |
| CBX5 | 0.61 | 0.00 | 0.00 | 0.40 | 0.21 | 1 | Matrix Fibroblast |
| GNG11 | 0.41 | 0.00 | 0.00 | 0.51 | 0.29 | 1 | Matrix Fibroblast |
| RCN3 | 0.52 | 0.00 | 0.00 | 0.27 | 0.11 | 1 | Matrix Fibroblast |
| SCP2 | 0.56 | 0.00 | 0.00 | 0.49 | 0.31 | 1 | Matrix Fibroblast |
| GNAS | 0.34 | 0.00 | 0.00 | 0.79 | 0.64 | 1 | Matrix Fibroblast |
| ALDH2 | 0.53 | 0.00 | 0.00 | 0.48 | 0.30 | 1 | Matrix Fibroblast |
| IFRD1 | 0.66 | 0.00 | 0.00 | 0.27 | 0.13 | 1 | Matrix Fibroblast |
| ZMYND11 | 0.54 | 0.00 | 0.00 | 0.47 | 0.28 | 1 | Matrix Fibroblast |
| FKBP7 | 0.54 | 0.00 | 0.00 | 0.27 | 0.12 | 1 | Matrix Fibroblast |
| SLC25A4 | 0.43 | 0.00 | 0.00 | 0.63 | 0.45 | 1 | Matrix Fibroblast |
| TIMP2 | 0.52 | 0.00 | 0.00 | 0.37 | 0.20 | 1 | Matrix Fibroblast |
| ECH1 | 0.61 | 0.00 | 0.00 | 0.31 | 0.15 | 1 | Matrix Fibroblast |
| TSPAN3 | 0.56 | 0.00 | 0.00 | 0.30 | 0.15 | 1 | Matrix Fibroblast |
| TMEM176B | 0.55 | 0.00 | 0.00 | 0.48 | 0.30 | 1 | Matrix Fibroblast |
| GTF2H5 | 0.54 | 0.00 | 0.00 | 0.42 | 0.26 | 1 | Matrix Fibroblast |
| NFIA | 0.56 | 0.00 | 0.00 | 0.39 | 0.22 | 1 | Matrix Fibroblast |
| ELN | 0.33 | 0.00 | 0.00 | 0.33 | 0.16 | 1 | Matrix Fibroblast |
| HMGN1 | 0.46 | 0.00 | 0.00 | 0.60 | 0.44 | 1 | Matrix Fibroblast |
| SERPINH1 | 0.38 | 0.00 | 0.00 | 0.59 | 0.39 | 1 | Matrix Fibroblast |
| BLMH | 0.53 | 0.00 | 0.00 | 0.29 | 0.15 | 1 | Matrix Fibroblast |
| THRA | 0.54 | 0.00 | 0.00 | 0.29 | 0.15 | 1 | Matrix Fibroblast |
| LIMA1 | 0.53 | 0.00 | 0.00 | 0.27 | 0.13 | 1 | Matrix Fibroblast |
| YWHAQ | 0.49 | 0.00 | 0.00 | 0.52 | 0.37 | 1 | Matrix Fibroblast |
| HMGN5 | 0.66 | 0.00 | 0.00 | 0.36 | 0.21 | 1 | Matrix Fibroblast |
| SQSTM1 | 0.47 | 0.00 | 0.00 | 0.45 | 0.30 | 1 | Matrix Fibroblast |
| EID1 | 0.45 | 0.00 | 0.00 | 0.46 | 0.31 | 1 | Matrix Fibroblast |
| RAD50 | 0.54 | 0.00 | 0.00 | 0.29 | 0.16 | 1 | Matrix Fibroblast |
| HSPB1 | 0.37 | 0.00 | 0.00 | 0.29 | 0.16 | 1 | Matrix Fibroblast |
| MAGED2 | 0.44 | 0.00 | 0.00 | 0.33 | 0.19 | 1 | Matrix Fibroblast |
| SUB1 | 0.34 | 0.00 | 0.00 | 0.64 | 0.50 | 1 | Matrix Fibroblast |
| FLNA | 0.38 | 0.00 | 0.00 | 0.38 | 0.23 | 1 | Matrix Fibroblast |
| COX7C | 0.34 | 0.00 | 0.00 | 0.74 | 0.63 | 1 | Matrix Fibroblast |
| RALB | 0.48 | 0.00 | 0.00 | 0.31 | 0.18 | 1 | Matrix Fibroblast |
| CD63 | 0.27 | 0.00 | 0.00 | 0.50 | 0.32 | 1 | Matrix Fibroblast |
| SEPT7 | 0.32 | 0.00 | 0.00 | 0.70 | 0.58 | 1 | Matrix Fibroblast |
| LAMC1 | 0.46 | 0.00 | 0.00 | 0.27 | 0.14 | 1 | Matrix Fibroblast |
| RPL23A | 0.33 | 0.00 | 0.00 | 0.80 | 0.68 | 1 | Matrix Fibroblast |
| MYO1B | 0.49 | 0.00 | 0.00 | 0.26 | 0.15 | 1 | Matrix Fibroblast |
| TGFB1I1 | 0.41 | 0.00 | 0.00 | 0.26 | 0.14 | 1 | Matrix Fibroblast |
| PPIC | 0.34 | 0.00 | 0.00 | 0.51 | 0.37 | 1 | Matrix Fibroblast |
| IDH2 | 0.46 | 0.00 | 0.00 | 0.30 | 0.18 | 1 | Matrix Fibroblast |
| COX6A1 | 0.34 | 0.00 | 0.00 | 0.66 | 0.55 | 1 | Matrix Fibroblast |
| PDCD4 | 0.34 | 0.00 | 0.00 | 0.32 | 0.19 | 1 | Matrix Fibroblast |
| WSB1 | 0.41 | 0.00 | 0.00 | 0.35 | 0.22 | 1 | Matrix Fibroblast |
| CLIC4 | 0.35 | 0.00 | 0.00 | 0.41 | 0.28 | 1 | Matrix Fibroblast |
| MMP14 | 0.35 | 0.00 | 0.00 | 0.36 | 0.23 | 1 | Matrix Fibroblast |
| PLS3 | 0.39 | 0.00 | 0.00 | 0.32 | 0.20 | 1 | Matrix Fibroblast |
| NFIX | 0.42 | 0.00 | 0.00 | 0.26 | 0.15 | 1 | Matrix Fibroblast |
| COX8A | 0.30 | 0.00 | 0.00 | 0.65 | 0.55 | 1 | Matrix Fibroblast |
| TCF12 | 0.43 | 0.00 | 0.00 | 0.25 | 0.15 | 1 | Matrix Fibroblast |
| TMEM204 | 0.34 | 0.00 | 0.00 | 0.26 | 0.15 | 1 | Matrix Fibroblast |
| SEC61B | 0.34 | 0.00 | 0.00 | 0.58 | 0.47 | 1 | Matrix Fibroblast |
| SFR1 | 0.38 | 0.00 | 0.00 | 0.41 | 0.29 | 1 | Matrix Fibroblast |
| NKTR | 0.38 | 0.00 | 0.00 | 0.46 | 0.34 | 1 | Matrix Fibroblast |
| CNPY2 | 0.40 | 0.00 | 0.00 | 0.30 | 0.19 | 1 | Matrix Fibroblast |
| KIF1B | 0.41 | 0.00 | 0.00 | 0.31 | 0.20 | 1 | Matrix Fibroblast |
| CKAP4 | 0.37 | 0.00 | 0.00 | 0.27 | 0.17 | 1 | Matrix Fibroblast |
| MPC2 | 0.37 | 0.00 | 0.00 | 0.33 | 0.22 | 1 | Matrix Fibroblast |
| SLC25A5 | 0.29 | 0.00 | 0.00 | 0.66 | 0.57 | 1 | Matrix Fibroblast |
| HSPA5 | 0.30 | 0.00 | 0.00 | 0.63 | 0.54 | 1 | Matrix Fibroblast |
| NDUFB9 | 0.34 | 0.00 | 0.00 | 0.47 | 0.36 | 1 | Matrix Fibroblast |
| DRAP1 | 0.50 | 0.00 | 0.00 | 0.34 | 0.24 | 1 | Matrix Fibroblast |
| SMIM15 | 0.38 | 0.00 | 0.00 | 0.28 | 0.17 | 1 | Matrix Fibroblast |
| ENY2 | 0.40 | 0.00 | 0.00 | 0.33 | 0.23 | 1 | Matrix Fibroblast |
| ATXN7L3B | 0.32 | 0.00 | 0.00 | 0.48 | 0.36 | 1 | Matrix Fibroblast |
| TBL1X | 0.36 | 0.00 | 0.00 | 0.32 | 0.22 | 1 | Matrix Fibroblast |
| DNAJA1 | 0.30 | 0.00 | 0.00 | 0.59 | 0.50 | 1 | Matrix Fibroblast |
| NDUFA4 | 0.27 | 0.00 | 0.00 | 0.65 | 0.56 | 1 | Matrix Fibroblast |
| TSC22D1 | 0.27 | 0.00 | 0.00 | 0.46 | 0.34 | 1 | Matrix Fibroblast |
| SEC61G | 0.34 | 0.00 | 0.00 | 0.46 | 0.36 | 1 | Matrix Fibroblast |
| ANXA6 | 0.32 | 0.00 | 0.00 | 0.31 | 0.21 | 1 | Matrix Fibroblast |
| MPRIP | 0.37 | 0.00 | 0.00 | 0.28 | 0.19 | 1 | Matrix Fibroblast |
| CCND2 | 0.25 | 0.00 | 0.00 | 0.37 | 0.26 | 1 | Matrix Fibroblast |
| MORF4L1 | 0.27 | 0.00 | 0.00 | 0.64 | 0.55 | 1 | Matrix Fibroblast |
| MRFAP1 | 0.31 | 0.00 | 0.00 | 0.44 | 0.34 | 1 | Matrix Fibroblast |
| SSR1 | 0.33 | 0.00 | 0.00 | 0.33 | 0.23 | 1 | Matrix Fibroblast |
| EIF3E | 0.31 | 0.00 | 0.00 | 0.52 | 0.42 | 1 | Matrix Fibroblast |
| MAP1LC3B | 0.34 | 0.00 | 0.00 | 0.47 | 0.37 | 1 | Matrix Fibroblast |
| NDUFC1 | 0.32 | 0.00 | 0.00 | 0.39 | 0.30 | 1 | Matrix Fibroblast |
| EIF4A2 | 0.27 | 0.00 | 0.00 | 0.44 | 0.34 | 1 | Matrix Fibroblast |
| LUC7L2 | 0.33 | 0.00 | 0.00 | 0.45 | 0.36 | 1 | Matrix Fibroblast |
| RPL24 | 0.31 | 0.00 | 0.00 | 0.58 | 0.48 | 1 | Matrix Fibroblast |
| SWI5 | 0.33 | 0.00 | 0.00 | 0.32 | 0.23 | 1 | Matrix Fibroblast |
| HSBP1 | 0.30 | 0.00 | 0.00 | 0.41 | 0.32 | 1 | Matrix Fibroblast |
| NDUFB11 | 0.30 | 0.00 | 0.00 | 0.43 | 0.34 | 1 | Matrix Fibroblast |
| UQCRB | 0.26 | 0.00 | 0.00 | 0.55 | 0.46 | 1 | Matrix Fibroblast |
| HTATSF1 | 0.36 | 0.00 | 0.00 | 0.32 | 0.24 | 1 | Matrix Fibroblast |
| RBBP7 | 0.32 | 0.00 | 0.00 | 0.32 | 0.24 | 1 | Matrix Fibroblast |
| IER3IP1 | 0.30 | 0.00 | 0.00 | 0.38 | 0.29 | 1 | Matrix Fibroblast |
| NDUFB3 | 0.31 | 0.00 | 0.00 | 0.39 | 0.30 | 1 | Matrix Fibroblast |
| LAMP2 | 0.32 | 0.00 | 0.00 | 0.33 | 0.24 | 1 | Matrix Fibroblast |
| ROMO1 | 0.26 | 0.00 | 0.00 | 0.39 | 0.30 | 1 | Matrix Fibroblast |
| NR3C1 | 0.31 | 0.00 | 0.00 | 0.33 | 0.25 | 1 | Matrix Fibroblast |
| NDUFA7 | 0.27 | 0.00 | 0.00 | 0.42 | 0.34 | 1 | Matrix Fibroblast |
| RPL17-C18orf32 | 0.29 | 0.00 | 0.00 | 0.38 | 0.30 | 1 | Matrix Fibroblast |
| NDUFB6 | 0.28 | 0.00 | 0.00 | 0.30 | 0.23 | 1 | Matrix Fibroblast |
| ANAPC13 | 0.30 | 0.00 | 0.00 | 0.27 | 0.20 | 1 | Matrix Fibroblast |
| NDUFA5 | 0.28 | 0.00 | 0.00 | 0.33 | 0.26 | 1 | Matrix Fibroblast |
| ZBTB20 | 0.26 | 0.00 | 0.00 | 0.34 | 0.26 | 1 | Matrix Fibroblast |
| PDCD5 | 0.28 | 0.00 | 0.00 | 0.34 | 0.27 | 1 | Matrix Fibroblast |
| ILK | 0.29 | 0.00 | 0.00 | 0.30 | 0.23 | 1 | Matrix Fibroblast |
| H2AFV | 0.27 | 0.00 | 0.00 | 0.33 | 0.25 | 1 | Matrix Fibroblast |
| COMMD1 | 0.26 | 0.00 | 0.00 | 0.26 | 0.19 | 1 | Matrix Fibroblast |
| ING4 | 0.26 | 0.00 | 0.00 | 0.27 | 0.20 | 1 | Matrix Fibroblast |
| PSMC5 | 0.26 | 0.00 | 0.00 | 0.29 | 0.23 | 1 | Matrix Fibroblast |
| HBA2 | 0.39 | 0.00 | 0.00 | 0.68 | 0.64 | 1 | Matrix Fibroblast |
| ARID5B | 0.30 | 0.00 | 0.00 | 0.28 | 0.22 | 1 | Matrix Fibroblast |
| AAMP | 0.26 | 0.00 | 0.00 | 0.27 | 0.21 | 1 | Matrix Fibroblast |
| C19orf70 | 0.26 | 0.00 | 0.00 | 0.29 | 0.23 | 1 | Matrix Fibroblast |
| PSMC1 | 0.26 | 0.00 | 0.00 | 0.37 | 0.30 | 1 | Matrix Fibroblast |
| NDUFB4 | 0.27 | 0.00 | 0.00 | 0.32 | 0.25 | 1 | Matrix Fibroblast |
| THOC2 | 0.26 | 0.00 | 0.00 | 0.38 | 0.31 | 1 | Matrix Fibroblast |
| TBRG1 | 0.26 | 0.00 | 0.00 | 0.34 | 0.28 | 1 | Matrix Fibroblast |
| PPP1R2 | 0.33 | 0.00 | 0.00 | 0.28 | 0.24 | 1 | Matrix Fibroblast |
| SFTPC | 4.04 | 0.00 | 0.00 | 0.97 | 0.54 | 2 | AT2 Cell |
| SFTPA2 | 3.82 | 0.00 | 0.00 | 0.97 | 0.11 | 2 | AT2 Cell |
| SFTPB | 3.54 | 0.00 | 0.00 | 0.97 | 0.08 | 2 | AT2 Cell |
| SLC34A2 | 3.34 | 0.00 | 0.00 | 0.94 | 0.06 | 2 | AT2 Cell |
| SFTPD | 3.12 | 0.00 | 0.00 | 0.95 | 0.06 | 2 | AT2 Cell |
| S100G | 2.85 | 0.00 | 0.00 | 0.81 | 0.04 | 2 | AT2 Cell |
| WFDC2 | 2.74 | 0.00 | 0.00 | 0.89 | 0.06 | 2 | AT2 Cell |
| SFTA2 | 2.54 | 0.00 | 0.00 | 0.88 | 0.05 | 2 | AT2 Cell |
| LPCAT1 | 2.44 | 0.00 | 0.00 | 0.85 | 0.10 | 2 | AT2 Cell |
| BEX2 | 2.44 | 0.00 | 0.00 | 0.65 | 0.03 | 2 | AT2 Cell |
| PPP1R14C | 2.33 | 0.00 | 0.00 | 0.70 | 0.03 | 2 | AT2 Cell |
| NAPSA | 2.32 | 0.00 | 0.00 | 0.87 | 0.12 | 2 | AT2 Cell |
| SPINK5 | 2.31 | 0.00 | 0.00 | 0.51 | 0.02 | 2 | AT2 Cell |
| NPC2 | 2.27 | 0.00 | 0.00 | 0.99 | 0.49 | 2 | AT2 Cell |
| DRAM1 | 2.22 | 0.00 | 0.00 | 0.73 | 0.05 | 2 | AT2 Cell |
| ABCA3 | 2.01 | 0.00 | 0.00 | 0.66 | 0.05 | 2 | AT2 Cell |
| CTSH | 1.93 | 0.00 | 0.00 | 0.76 | 0.14 | 2 | AT2 Cell |
| ATP1B1 | 1.91 | 0.00 | 0.00 | 0.79 | 0.11 | 2 | AT2 Cell |
| LGI3 | 1.89 | 0.00 | 0.00 | 0.54 | 0.02 | 2 | AT2 Cell |
| RBPJL | 1.78 | 0.00 | 0.00 | 0.45 | 0.02 | 2 | AT2 Cell |
| DLK1 | 1.74 | 0.00 | 0.00 | 0.34 | 0.03 | 2 | AT2 Cell |
| FABP5 | 1.70 | 0.00 | 0.00 | 0.59 | 0.09 | 2 | AT2 Cell |
| EGFL6 | 1.70 | 0.00 | 0.00 | 0.60 | 0.05 | 2 | AT2 Cell |
| ALCAM | 1.70 | 0.00 | 0.00 | 0.79 | 0.14 | 2 | AT2 Cell |
| SDC4 | 1.69 | 0.00 | 0.00 | 0.73 | 0.14 | 2 | AT2 Cell |
| PI4K2B | 1.66 | 0.00 | 0.00 | 0.50 | 0.04 | 2 | AT2 Cell |
| CLDN3 | 1.62 | 0.00 | 0.00 | 0.57 | 0.05 | 2 | AT2 Cell |
| DCXR | 1.62 | 0.00 | 0.00 | 0.51 | 0.06 | 2 | AT2 Cell |
| PTPRF | 1.61 | 0.00 | 0.00 | 0.63 | 0.09 | 2 | AT2 Cell |
| MGST1 | 1.59 | 0.00 | 0.00 | 0.67 | 0.12 | 2 | AT2 Cell |
| ANK3 | 1.52 | 0.00 | 0.00 | 0.42 | 0.03 | 2 | AT2 Cell |
| ACSL4 | 1.48 | 0.00 | 0.00 | 0.49 | 0.07 | 2 | AT2 Cell |
| ETV5 | 1.47 | 0.00 | 0.00 | 0.40 | 0.03 | 2 | AT2 Cell |
| ATP6V1C2 | 1.45 | 0.00 | 0.00 | 0.36 | 0.01 | 2 | AT2 Cell |
| FASN | 1.44 | 0.00 | 0.00 | 0.40 | 0.03 | 2 | AT2 Cell |
| MID1IP1 | 1.44 | 0.00 | 0.00 | 0.49 | 0.10 | 2 | AT2 Cell |
| PLA2G1B | 1.43 | 0.00 | 0.00 | 0.36 | 0.01 | 2 | AT2 Cell |
| RNASE4 | 1.43 | 0.00 | 0.00 | 0.57 | 0.12 | 2 | AT2 Cell |
| PPP1R9A | 1.42 | 0.00 | 0.00 | 0.51 | 0.09 | 2 | AT2 Cell |
| IRX3 | 1.41 | 0.00 | 0.00 | 0.39 | 0.03 | 2 | AT2 Cell |
| NKX2-1 | 1.40 | 0.00 | 0.00 | 0.48 | 0.04 | 2 | AT2 Cell |
| ABCD3 | 1.40 | 0.00 | 0.00 | 0.58 | 0.13 | 2 | AT2 Cell |
| MYO5C | 1.39 | 0.00 | 0.00 | 0.39 | 0.03 | 2 | AT2 Cell |
| EPCAM | 1.39 | 0.00 | 0.00 | 0.46 | 0.04 | 2 | AT2 Cell |
| IL33 | 1.37 | 0.00 | 0.00 | 0.28 | 0.02 | 2 | AT2 Cell |
| CHCHD10 | 1.37 | 0.00 | 0.00 | 0.51 | 0.08 | 2 | AT2 Cell |
| ATP8A1 | 1.35 | 0.00 | 0.00 | 0.54 | 0.09 | 2 | AT2 Cell |
| FGFR2 | 1.30 | 0.00 | 0.00 | 0.36 | 0.03 | 2 | AT2 Cell |
| ZDHHC3 | 1.28 | 0.00 | 0.00 | 0.51 | 0.11 | 2 | AT2 Cell |
| TGOLN2 | 1.27 | 0.00 | 0.00 | 0.53 | 0.15 | 2 | AT2 Cell |
| GCLC | 1.24 | 0.00 | 0.00 | 0.34 | 0.05 | 2 | AT2 Cell |
| EXOSC7 | 1.23 | 0.00 | 0.00 | 0.44 | 0.09 | 2 | AT2 Cell |
| MT1M | 1.22 | 0.00 | 0.00 | 0.26 | 0.03 | 2 | AT2 Cell |
| CLDN18 | 1.21 | 0.00 | 0.00 | 0.80 | 0.09 | 2 | AT2 Cell |
| ERRFI1 | 1.21 | 0.00 | 0.00 | 0.34 | 0.06 | 2 | AT2 Cell |
| SPINT2 | 1.19 | 0.00 | 0.00 | 0.44 | 0.07 | 2 | AT2 Cell |
| NUPR1 | 1.18 | 0.00 | 0.00 | 0.46 | 0.18 | 2 | AT2 Cell |
| PRR15L | 1.18 | 0.00 | 0.00 | 0.30 | 0.02 | 2 | AT2 Cell |
| TFCP2L1 | 1.16 | 0.00 | 0.00 | 0.26 | 0.01 | 2 | AT2 Cell |
| TMEM243 | 1.15 | 0.00 | 0.00 | 0.33 | 0.06 | 2 | AT2 Cell |
| SCD | 1.14 | 0.00 | 0.00 | 0.56 | 0.16 | 2 | AT2 Cell |
| ATP11A | 1.13 | 0.00 | 0.00 | 0.39 | 0.07 | 2 | AT2 Cell |
| SOAT1 | 1.12 | 0.00 | 0.00 | 0.38 | 0.07 | 2 | AT2 Cell |
| BRD7 | 1.11 | 0.00 | 0.00 | 0.60 | 0.21 | 2 | AT2 Cell |
| CDH1 | 1.10 | 0.00 | 0.00 | 0.32 | 0.03 | 2 | AT2 Cell |
| NUCB2 | 1.09 | 0.00 | 0.00 | 0.47 | 0.13 | 2 | AT2 Cell |
| CEBPA | 1.09 | 0.00 | 0.00 | 0.28 | 0.02 | 2 | AT2 Cell |
| H2AFJ | 1.07 | 0.00 | 0.00 | 0.67 | 0.28 | 2 | AT2 Cell |
| CTSC | 1.07 | 0.00 | 0.00 | 0.56 | 0.16 | 2 | AT2 Cell |
| IRX1 | 1.06 | 0.00 | 0.00 | 0.26 | 0.02 | 2 | AT2 Cell |
| CMTM8 | 1.05 | 0.00 | 0.00 | 0.40 | 0.08 | 2 | AT2 Cell |
| GDE1 | 1.05 | 0.00 | 0.00 | 0.44 | 0.11 | 2 | AT2 Cell |
| LPIN2 | 1.05 | 0.00 | 0.00 | 0.40 | 0.10 | 2 | AT2 Cell |
| SLC12A2 | 1.04 | 0.00 | 0.00 | 0.45 | 0.11 | 2 | AT2 Cell |
| CCZ1B | 1.04 | 0.00 | 0.00 | 0.32 | 0.10 | 2 | AT2 Cell |
| CLDN7 | 1.04 | 0.00 | 0.00 | 0.33 | 0.04 | 2 | AT2 Cell |
| MECOM | 1.04 | 0.00 | 0.00 | 0.35 | 0.07 | 2 | AT2 Cell |
| GAS6 | 1.04 | 0.00 | 0.00 | 0.45 | 0.12 | 2 | AT2 Cell |
| SNX25 | 1.02 | 0.00 | 0.00 | 0.26 | 0.03 | 2 | AT2 Cell |
| MBIP | 1.01 | 0.00 | 0.00 | 0.38 | 0.08 | 2 | AT2 Cell |
| RAB27B | 0.96 | 0.00 | 0.00 | 0.28 | 0.04 | 2 | AT2 Cell |
| UHRF1BP1L | 0.96 | 0.00 | 0.00 | 0.31 | 0.06 | 2 | AT2 Cell |
| TAOK3 | 0.95 | 0.00 | 0.00 | 0.47 | 0.15 | 2 | AT2 Cell |
| RAI14 | 0.94 | 0.00 | 0.00 | 0.34 | 0.08 | 2 | AT2 Cell |
| CHKA | 0.93 | 0.00 | 0.00 | 0.32 | 0.06 | 2 | AT2 Cell |
| AVPI1 | 0.90 | 0.00 | 0.00 | 0.29 | 0.05 | 2 | AT2 Cell |
| VAMP8 | 0.89 | 0.00 | 0.00 | 0.66 | 0.29 | 2 | AT2 Cell |
| SWT1 | 0.88 | 0.00 | 0.00 | 0.37 | 0.11 | 2 | AT2 Cell |
| SLC39A8 | 0.84 | 0.00 | 0.00 | 0.29 | 0.05 | 2 | AT2 Cell |
| NPC1 | 0.84 | 0.00 | 0.00 | 0.28 | 0.06 | 2 | AT2 Cell |
| SECISBP2L | 0.83 | 0.00 | 0.00 | 0.41 | 0.15 | 2 | AT2 Cell |
| BRI3 | 0.83 | 0.00 | 0.00 | 0.46 | 0.18 | 2 | AT2 Cell |
| PON3 | 0.82 | 0.00 | 0.00 | 0.30 | 0.07 | 2 | AT2 Cell |
| CADM1 | 0.82 | 0.00 | 0.00 | 0.41 | 0.11 | 2 | AT2 Cell |
| MYH14 | 0.82 | 0.00 | 0.00 | 0.32 | 0.06 | 2 | AT2 Cell |
| TMEM245 | 0.78 | 0.00 | 0.00 | 0.29 | 0.08 | 2 | AT2 Cell |
| DPP4 | 0.75 | 0.00 | 0.00 | 0.36 | 0.10 | 2 | AT2 Cell |
| KRT18 | 0.74 | 0.00 | 0.00 | 0.27 | 0.06 | 2 | AT2 Cell |
| ARL6IP1 | 0.74 | 0.00 | 0.00 | 0.76 | 0.46 | 2 | AT2 Cell |
| CYSTM1 | 0.74 | 0.00 | 0.00 | 0.28 | 0.07 | 2 | AT2 Cell |
| EMB | 0.67 | 0.00 | 0.00 | 0.30 | 0.08 | 2 | AT2 Cell |
| IQGAP1 | 0.66 | 0.00 | 0.00 | 0.73 | 0.39 | 2 | AT2 Cell |
| AGER | 0.63 | 0.00 | 0.00 | 0.71 | 0.10 | 2 | AT2 Cell |
| CLIC3 | 0.53 | 0.00 | 0.00 | 0.27 | 0.06 | 2 | AT2 Cell |
| ACSL5 | 0.72 | 0.00 | 0.00 | 0.28 | 0.08 | 2 | AT2 Cell |
| CD36 | 0.45 | 0.00 | 0.00 | 0.65 | 0.29 | 2 | AT2 Cell |
| ITGA9 | 0.78 | 0.00 | 0.00 | 0.27 | 0.07 | 2 | AT2 Cell |
| PRNP | 0.80 | 0.00 | 0.00 | 0.34 | 0.11 | 2 | AT2 Cell |
| CPM | 0.55 | 0.00 | 0.00 | 0.40 | 0.14 | 2 | AT2 Cell |
| ATP13A3 | 0.65 | 0.00 | 0.00 | 0.25 | 0.07 | 2 | AT2 Cell |
| CD74 | 0.27 | 0.00 | 0.00 | 0.37 | 0.13 | 2 | AT2 Cell |
| LDHB | 0.69 | 0.00 | 0.00 | 0.29 | 0.09 | 2 | AT2 Cell |
| EZR | 0.57 | 0.00 | 0.00 | 0.44 | 0.18 | 2 | AT2 Cell |
| SELENBP1 | 0.73 | 0.00 | 0.00 | 0.43 | 0.18 | 2 | AT2 Cell |
| PCNT | 0.75 | 0.00 | 0.00 | 0.29 | 0.10 | 2 | AT2 Cell |
| DAPK1 | 0.65 | 0.00 | 0.00 | 0.26 | 0.08 | 2 | AT2 Cell |
| ADK | 0.67 | 0.00 | 0.00 | 0.29 | 0.09 | 2 | AT2 Cell |
| APP | 0.44 | 0.00 | 0.00 | 0.84 | 0.58 | 2 | AT2 Cell |
| ELOVL1 | 0.71 | 0.00 | 0.00 | 0.33 | 0.13 | 2 | AT2 Cell |
| ATP6V1A | 0.58 | 0.00 | 0.00 | 0.47 | 0.24 | 2 | AT2 Cell |
| SCP2 | 0.64 | 0.00 | 0.00 | 0.54 | 0.31 | 2 | AT2 Cell |
| IAH1 | 0.65 | 0.00 | 0.00 | 0.29 | 0.11 | 2 | AT2 Cell |
| DBI | 0.56 | 0.00 | 0.00 | 0.58 | 0.35 | 2 | AT2 Cell |
| CITED2 | 0.61 | 0.00 | 0.00 | 0.25 | 0.09 | 2 | AT2 Cell |
| FKBP4 | 0.61 | 0.00 | 0.00 | 0.30 | 0.12 | 2 | AT2 Cell |
| TMBIM6 | 0.50 | 0.00 | 0.00 | 0.61 | 0.38 | 2 | AT2 Cell |
| LMO7 | 0.28 | 0.00 | 0.00 | 0.28 | 0.10 | 2 | AT2 Cell |
| SOCS2 | 0.54 | 0.00 | 0.00 | 0.52 | 0.30 | 2 | AT2 Cell |
| TNFAIP1 | 0.55 | 0.00 | 0.00 | 0.32 | 0.14 | 2 | AT2 Cell |
| PRKCI | 0.55 | 0.00 | 0.00 | 0.26 | 0.10 | 2 | AT2 Cell |
| CTNNB1 | 0.50 | 0.00 | 0.00 | 0.54 | 0.32 | 2 | AT2 Cell |
| NFE2L2 | 0.52 | 0.00 | 0.00 | 0.43 | 0.23 | 2 | AT2 Cell |
| BSG | 0.45 | 0.00 | 0.00 | 0.57 | 0.37 | 2 | AT2 Cell |
| RNASET2 | 0.48 | 0.00 | 0.00 | 0.42 | 0.23 | 2 | AT2 Cell |
| ARF6 | 0.49 | 0.00 | 0.00 | 0.34 | 0.17 | 2 | AT2 Cell |
| NDUFC2-KCTD14 | 0.41 | 0.00 | 0.00 | 0.53 | 0.33 | 2 | AT2 Cell |
| EMC6 | 0.48 | 0.00 | 0.00 | 0.27 | 0.12 | 2 | AT2 Cell |
| HERPUD1 | 0.56 | 0.00 | 0.00 | 0.28 | 0.14 | 2 | AT2 Cell |
| CSTB | 0.37 | 0.00 | 0.00 | 0.44 | 0.27 | 2 | AT2 Cell |
| GOLGA4 | 0.44 | 0.00 | 0.00 | 0.45 | 0.27 | 2 | AT2 Cell |
| DSTN | 0.30 | 0.00 | 0.00 | 0.71 | 0.51 | 2 | AT2 Cell |
| PHLDB2 | 0.39 | 0.00 | 0.00 | 0.45 | 0.26 | 2 | AT2 Cell |
| ATP6V1G1 | 0.44 | 0.00 | 0.00 | 0.50 | 0.33 | 2 | AT2 Cell |
| WLS | 0.39 | 0.00 | 0.00 | 0.37 | 0.20 | 2 | AT2 Cell |
| IFT20 | 0.44 | 0.00 | 0.00 | 0.39 | 0.23 | 2 | AT2 Cell |
| SIVA1 | 0.51 | 0.00 | 0.00 | 0.28 | 0.15 | 2 | AT2 Cell |
| TMEM30A | 0.44 | 0.00 | 0.00 | 0.42 | 0.26 | 2 | AT2 Cell |
| SNX4 | 0.42 | 0.00 | 0.00 | 0.39 | 0.23 | 2 | AT2 Cell |
| GPX4 | 0.41 | 0.00 | 0.00 | 0.53 | 0.37 | 2 | AT2 Cell |
| IRF2BP2 | 0.42 | 0.00 | 0.00 | 0.27 | 0.14 | 2 | AT2 Cell |
| AKAP9 | 0.36 | 0.00 | 0.00 | 0.50 | 0.33 | 2 | AT2 Cell |
| PRDX6 | 0.49 | 0.00 | 0.00 | 0.42 | 0.28 | 2 | AT2 Cell |
| SOD1 | 0.34 | 0.00 | 0.00 | 0.62 | 0.46 | 2 | AT2 Cell |
| CAMK2N1 | 0.37 | 0.00 | 0.00 | 0.26 | 0.13 | 2 | AT2 Cell |
| ATP1A1 | 0.37 | 0.00 | 0.00 | 0.26 | 0.13 | 2 | AT2 Cell |
| PIK3CA | 0.40 | 0.00 | 0.00 | 0.30 | 0.17 | 2 | AT2 Cell |
| ETFB | 0.38 | 0.00 | 0.00 | 0.41 | 0.26 | 2 | AT2 Cell |
| EIF4H | 0.51 | 0.00 | 0.00 | 0.40 | 0.26 | 2 | AT2 Cell |
| MTCH1 | 0.47 | 0.00 | 0.00 | 0.28 | 0.16 | 2 | AT2 Cell |
| USE1 | 0.34 | 0.00 | 0.00 | 0.39 | 0.24 | 2 | AT2 Cell |
| CREG1 | 0.32 | 0.00 | 0.00 | 0.28 | 0.16 | 2 | AT2 Cell |
| TSPO | 0.28 | 0.00 | 0.00 | 0.39 | 0.25 | 2 | AT2 Cell |
| POLR2E | 0.32 | 0.00 | 0.00 | 0.27 | 0.16 | 2 | AT2 Cell |
| HIGD1A | 0.31 | 0.00 | 0.00 | 0.28 | 0.17 | 2 | AT2 Cell |
| HDGF | 0.29 | 0.00 | 0.00 | 0.34 | 0.22 | 2 | AT2 Cell |
| ACADL | 0.27 | 0.00 | 0.00 | 0.31 | 0.19 | 2 | AT2 Cell |
| TCEAL8 | 0.34 | 0.00 | 0.00 | 0.30 | 0.20 | 2 | AT2 Cell |
| SCGB1A1 | 3.18 | 0.00 | 0.00 | 0.43 | 0.31 | 2 | AT2 Cell |
| MAP1LC3A | 0.27 | 0.00 | 0.00 | 0.30 | 0.20 | 2 | AT2 Cell |
| NEMF | 0.27 | 0.00 | 0.00 | 0.31 | 0.21 | 2 | AT2 Cell |
| ATXN10 | 0.28 | 0.00 | 0.00 | 0.28 | 0.19 | 2 | AT2 Cell |
| VAPA | 0.27 | 0.00 | 0.00 | 0.33 | 0.23 | 2 | AT2 Cell |
| RGCC | 0.54 | 0.00 | 0.00 | 0.26 | 0.19 | 2 | AT2 Cell |
| TGFBI | 3.06 | 0.00 | 0.00 | 0.93 | 0.13 | 3 | Myofibroblast |
| ASPN | 2.88 | 0.00 | 0.00 | 0.46 | 0.07 | 3 | Myofibroblast |
| TAGLN | 2.27 | 0.00 | 0.00 | 0.79 | 0.12 | 3 | Myofibroblast |
| TNC | 2.27 | 0.00 | 0.00 | 0.75 | 0.08 | 3 | Myofibroblast |
| HTRA1 | 2.24 | 0.00 | 0.00 | 0.54 | 0.04 | 3 | Myofibroblast |
| AGT | 2.19 | 0.00 | 0.00 | 0.62 | 0.03 | 3 | Myofibroblast |
| ACTA2 | 2.17 | 0.00 | 0.00 | 0.85 | 0.18 | 3 | Myofibroblast |
| P2RY14 | 2.16 | 0.00 | 0.00 | 0.59 | 0.03 | 3 | Myofibroblast |
| LOXL2 | 2.14 | 0.00 | 0.00 | 0.64 | 0.08 | 3 | Myofibroblast |
| ACTG2 | 2.06 | 0.00 | 0.00 | 0.54 | 0.04 | 3 | Myofibroblast |
| MYH11 | 2.04 | 0.00 | 0.00 | 0.73 | 0.06 | 3 | Myofibroblast |
| IGF1 | 2.03 | 0.00 | 0.00 | 0.65 | 0.08 | 3 | Myofibroblast |
| PRSS35 | 2.02 | 0.00 | 0.00 | 0.52 | 0.03 | 3 | Myofibroblast |
| WNT5A | 2.02 | 0.00 | 0.00 | 0.58 | 0.05 | 3 | Myofibroblast |
| LTBP2 | 1.93 | 0.00 | 0.00 | 0.59 | 0.05 | 3 | Myofibroblast |
| PDLIM3 | 1.80 | 0.00 | 0.00 | 0.54 | 0.03 | 3 | Myofibroblast |
| DES | 1.80 | 0.00 | 0.00 | 0.44 | 0.03 | 3 | Myofibroblast |
| NET1 | 1.79 | 0.00 | 0.00 | 0.59 | 0.11 | 3 | Myofibroblast |
| FILIP1L | 1.73 | 0.00 | 0.00 | 0.85 | 0.28 | 3 | Myofibroblast |
| SPON2 | 1.71 | 0.00 | 0.00 | 0.48 | 0.03 | 3 | Myofibroblast |
| CALD1 | 1.64 | 0.00 | 0.00 | 0.94 | 0.39 | 3 | Myofibroblast |
| SERPINE2 | 1.60 | 0.00 | 0.00 | 0.74 | 0.20 | 3 | Myofibroblast |
| MUSTN1 | 1.60 | 0.00 | 0.00 | 0.41 | 0.04 | 3 | Myofibroblast |
| FSTL1 | 1.59 | 0.00 | 0.00 | 0.92 | 0.41 | 3 | Myofibroblast |
| OLFML2B | 1.58 | 0.00 | 0.00 | 0.42 | 0.03 | 3 | Myofibroblast |
| CKB | 1.54 | 0.00 | 0.00 | 0.56 | 0.09 | 3 | Myofibroblast |
| TPM2 | 1.51 | 0.00 | 0.00 | 0.65 | 0.13 | 3 | Myofibroblast |
| COL3A1 | 1.49 | 0.00 | 0.00 | 0.97 | 0.28 | 3 | Myofibroblast |
| PDE5A | 1.48 | 0.00 | 0.00 | 0.51 | 0.07 | 3 | Myofibroblast |
| THBS1 | 1.47 | 0.00 | 0.00 | 0.49 | 0.09 | 3 | Myofibroblast |
| NES | 1.42 | 0.00 | 0.00 | 0.61 | 0.17 | 3 | Myofibroblast |
| MYL9 | 1.41 | 0.00 | 0.00 | 0.61 | 0.12 | 3 | Myofibroblast |
| LGALS1 | 1.41 | 0.00 | 0.00 | 0.96 | 0.41 | 3 | Myofibroblast |
| CD248 | 1.39 | 0.00 | 0.00 | 0.33 | 0.02 | 3 | Myofibroblast |
| NREP | 1.37 | 0.00 | 0.00 | 0.86 | 0.39 | 3 | Myofibroblast |
| CNN1 | 1.36 | 0.00 | 0.00 | 0.25 | 0.02 | 3 | Myofibroblast |
| FNDC1 | 1.32 | 0.00 | 0.00 | 0.33 | 0.02 | 3 | Myofibroblast |
| ELN | 1.29 | 0.00 | 0.00 | 0.53 | 0.15 | 3 | Myofibroblast |
| HLF | 1.29 | 0.00 | 0.00 | 0.28 | 0.02 | 3 | Myofibroblast |
| PDGFRA | 1.27 | 0.00 | 0.00 | 0.30 | 0.03 | 3 | Myofibroblast |
| MYLK | 1.26 | 0.00 | 0.00 | 0.74 | 0.25 | 3 | Myofibroblast |
| COL5A1 | 1.25 | 0.00 | 0.00 | 0.44 | 0.07 | 3 | Myofibroblast |
| MAP1B | 1.22 | 0.00 | 0.00 | 0.39 | 0.07 | 3 | Myofibroblast |
| TGFB2 | 1.22 | 0.00 | 0.00 | 0.43 | 0.10 | 3 | Myofibroblast |
| IGFBP5 | 1.21 | 0.00 | 0.00 | 0.39 | 0.07 | 3 | Myofibroblast |
| DKK3 | 1.20 | 0.00 | 0.00 | 0.49 | 0.10 | 3 | Myofibroblast |
| TPM1 | 1.20 | 0.00 | 0.00 | 0.91 | 0.48 | 3 | Myofibroblast |
| MDK | 1.19 | 0.00 | 0.00 | 0.65 | 0.18 | 3 | Myofibroblast |
| LPAR1 | 1.19 | 0.00 | 0.00 | 0.29 | 0.04 | 3 | Myofibroblast |
| ROBO2 | 1.19 | 0.00 | 0.00 | 0.36 | 0.06 | 3 | Myofibroblast |
| PAMR1 | 1.18 | 0.00 | 0.00 | 0.26 | 0.03 | 3 | Myofibroblast |
| C1QTNF7 | 1.17 | 0.00 | 0.00 | 0.35 | 0.06 | 3 | Myofibroblast |
| MYH10 | 1.17 | 0.00 | 0.00 | 0.63 | 0.18 | 3 | Myofibroblast |
| BGN | 1.12 | 0.00 | 0.00 | 0.72 | 0.21 | 3 | Myofibroblast |
| RCN3 | 1.10 | 0.00 | 0.00 | 0.44 | 0.10 | 3 | Myofibroblast |
| ANGPTL2 | 1.10 | 0.00 | 0.00 | 0.28 | 0.03 | 3 | Myofibroblast |
| CTGF | 1.09 | 0.00 | 0.00 | 0.32 | 0.07 | 3 | Myofibroblast |
| COL27A1 | 1.09 | 0.00 | 0.00 | 0.29 | 0.04 | 3 | Myofibroblast |
| TGFB3 | 1.08 | 0.00 | 0.00 | 0.28 | 0.04 | 3 | Myofibroblast |
| COL1A2 | 1.08 | 0.00 | 0.00 | 0.98 | 0.35 | 3 | Myofibroblast |
| LMCD1 | 1.07 | 0.00 | 0.00 | 0.31 | 0.05 | 3 | Myofibroblast |
| ADAMTS2 | 1.06 | 0.00 | 0.00 | 0.34 | 0.08 | 3 | Myofibroblast |
| MFAP2 | 1.06 | 0.00 | 0.00 | 0.76 | 0.31 | 3 | Myofibroblast |
| FAM129A | 1.06 | 0.00 | 0.00 | 0.32 | 0.06 | 3 | Myofibroblast |
| PALLD | 1.02 | 0.00 | 0.00 | 0.53 | 0.14 | 3 | Myofibroblast |
| LMNA | 1.01 | 0.00 | 0.00 | 0.51 | 0.20 | 3 | Myofibroblast |
| SAMD4A | 1.01 | 0.00 | 0.00 | 0.29 | 0.06 | 3 | Myofibroblast |
| TSHZ2 | 1.01 | 0.00 | 0.00 | 0.32 | 0.07 | 3 | Myofibroblast |
| DDR2 | 1.00 | 0.00 | 0.00 | 0.27 | 0.04 | 3 | Myofibroblast |
| LSP1 | 0.99 | 0.00 | 0.00 | 0.48 | 0.16 | 3 | Myofibroblast |
| FBN1 | 0.99 | 0.00 | 0.00 | 0.49 | 0.14 | 3 | Myofibroblast |
| CREB3L2 | 0.99 | 0.00 | 0.00 | 0.38 | 0.11 | 3 | Myofibroblast |
| PDZRN3 | 0.99 | 0.00 | 0.00 | 0.27 | 0.04 | 3 | Myofibroblast |
| MGP | 0.98 | 0.00 | 0.00 | 0.72 | 0.30 | 3 | Myofibroblast |
| CYGB | 0.97 | 0.00 | 0.00 | 0.28 | 0.05 | 3 | Myofibroblast |
| LHFPL2 | 0.95 | 0.00 | 0.00 | 0.31 | 0.06 | 3 | Myofibroblast |
| POSTN | 0.95 | 0.00 | 0.00 | 0.31 | 0.07 | 3 | Myofibroblast |
| C11orf96 | 0.94 | 0.00 | 0.00 | 0.30 | 0.06 | 3 | Myofibroblast |
| RGS2 | 0.94 | 0.00 | 0.00 | 0.57 | 0.25 | 3 | Myofibroblast |
| CYR61 | 0.94 | 0.00 | 0.00 | 0.34 | 0.10 | 3 | Myofibroblast |
| PDLIM7 | 0.93 | 0.00 | 0.00 | 0.36 | 0.10 | 3 | Myofibroblast |
| GRB10 | 0.93 | 0.00 | 0.00 | 0.40 | 0.12 | 3 | Myofibroblast |
| PRNP | 0.93 | 0.00 | 0.00 | 0.36 | 0.12 | 3 | Myofibroblast |
| ITGA9 | 0.91 | 0.00 | 0.00 | 0.32 | 0.07 | 3 | Myofibroblast |
| COL1A1 | 0.91 | 0.00 | 0.00 | 0.80 | 0.23 | 3 | Myofibroblast |
| SOCS2 | 0.90 | 0.00 | 0.00 | 0.63 | 0.29 | 3 | Myofibroblast |
| RBMS3 | 0.89 | 0.00 | 0.00 | 0.45 | 0.15 | 3 | Myofibroblast |
| PMEPA1 | 0.89 | 0.00 | 0.00 | 0.33 | 0.09 | 3 | Myofibroblast |
| GOLIM4 | 0.88 | 0.00 | 0.00 | 0.48 | 0.18 | 3 | Myofibroblast |
| CNN3 | 0.88 | 0.00 | 0.00 | 0.67 | 0.29 | 3 | Myofibroblast |
| ETV1 | 0.87 | 0.00 | 0.00 | 0.26 | 0.06 | 3 | Myofibroblast |
| CALU | 0.86 | 0.00 | 0.00 | 0.55 | 0.22 | 3 | Myofibroblast |
| COL5A2 | 0.85 | 0.00 | 0.00 | 0.46 | 0.16 | 3 | Myofibroblast |
| MXRA7 | 0.84 | 0.00 | 0.00 | 0.38 | 0.12 | 3 | Myofibroblast |
| ISM1 | 0.82 | 0.00 | 0.00 | 0.28 | 0.06 | 3 | Myofibroblast |
| PRDM6 | 0.80 | 0.00 | 0.00 | 0.25 | 0.05 | 3 | Myofibroblast |
| SMTN | 0.79 | 0.00 | 0.00 | 0.29 | 0.07 | 3 | Myofibroblast |
| NFIB | 0.79 | 0.00 | 0.00 | 0.78 | 0.44 | 3 | Myofibroblast |
| SOX4 | 0.78 | 0.00 | 0.00 | 0.71 | 0.37 | 3 | Myofibroblast |
| NEXN | 0.78 | 0.00 | 0.00 | 0.55 | 0.18 | 3 | Myofibroblast |
| COL6A3 | 0.78 | 0.00 | 0.00 | 0.38 | 0.10 | 3 | Myofibroblast |
| SERPINH1 | 0.77 | 0.00 | 0.00 | 0.73 | 0.38 | 3 | Myofibroblast |
| RARRES2 | 0.77 | 0.00 | 0.00 | 0.48 | 0.13 | 3 | Myofibroblast |
| ANXA5 | 0.76 | 0.00 | 0.00 | 0.71 | 0.40 | 3 | Myofibroblast |
| SPARC | 0.62 | 0.00 | 0.00 | 0.98 | 0.74 | 3 | Myofibroblast |
| PLAC9 | 0.53 | 0.00 | 0.00 | 0.65 | 0.27 | 3 | Myofibroblast |
| CD44 | 0.68 | 0.00 | 0.00 | 0.47 | 0.18 | 3 | Myofibroblast |
| TUBA1A | 0.67 | 0.00 | 0.00 | 0.78 | 0.50 | 3 | Myofibroblast |
| HMCN1 | 0.96 | 0.00 | 0.00 | 0.35 | 0.11 | 3 | Myofibroblast |
| PROS1 | 0.78 | 0.00 | 0.00 | 0.28 | 0.08 | 3 | Myofibroblast |
| SEMA3C | 0.60 | 0.00 | 0.00 | 0.46 | 0.17 | 3 | Myofibroblast |
| TGFB1I1 | 0.72 | 0.00 | 0.00 | 0.38 | 0.13 | 3 | Myofibroblast |
| CALM2 | 0.57 | 0.00 | 0.00 | 0.86 | 0.64 | 3 | Myofibroblast |
| MMP14 | 0.75 | 0.00 | 0.00 | 0.51 | 0.22 | 3 | Myofibroblast |
| FLNA | 0.71 | 0.00 | 0.00 | 0.51 | 0.22 | 3 | Myofibroblast |
| LOXL1 | 0.75 | 0.00 | 0.00 | 0.31 | 0.10 | 3 | Myofibroblast |
| FKBP7 | 0.74 | 0.00 | 0.00 | 0.35 | 0.12 | 3 | Myofibroblast |
| GHR | 0.81 | 0.00 | 0.00 | 0.33 | 0.12 | 3 | Myofibroblast |
| CSRP2 | 0.66 | 0.00 | 0.00 | 0.38 | 0.14 | 3 | Myofibroblast |
| ILK | 0.67 | 0.00 | 0.00 | 0.47 | 0.21 | 3 | Myofibroblast |
| ITGB1 | 0.52 | 0.00 | 0.00 | 0.85 | 0.62 | 3 | Myofibroblast |
| CELF2 | 0.70 | 0.00 | 0.00 | 0.51 | 0.24 | 3 | Myofibroblast |
| ACTN1 | 0.71 | 0.00 | 0.00 | 0.46 | 0.21 | 3 | Myofibroblast |
| RORA | 0.81 | 0.00 | 0.00 | 0.26 | 0.08 | 3 | Myofibroblast |
| TIMP2 | 0.67 | 0.00 | 0.00 | 0.44 | 0.20 | 3 | Myofibroblast |
| COL6A2 | 0.58 | 0.00 | 0.00 | 0.40 | 0.16 | 3 | Myofibroblast |
| TSPAN3 | 0.64 | 0.00 | 0.00 | 0.37 | 0.15 | 3 | Myofibroblast |
| PXDN | 0.66 | 0.00 | 0.00 | 0.29 | 0.10 | 3 | Myofibroblast |
| CD63 | 0.56 | 0.00 | 0.00 | 0.60 | 0.32 | 3 | Myofibroblast |
| TPM4 | 0.55 | 0.00 | 0.00 | 0.73 | 0.48 | 3 | Myofibroblast |
| PTMA | 0.35 | 0.00 | 0.00 | 0.97 | 0.91 | 3 | Myofibroblast |
| GPM6B | 0.61 | 0.00 | 0.00 | 0.31 | 0.11 | 3 | Myofibroblast |
| LAMA4 | 0.66 | 0.00 | 0.00 | 0.32 | 0.12 | 3 | Myofibroblast |
| COL6A1 | 0.52 | 0.00 | 0.00 | 0.45 | 0.20 | 3 | Myofibroblast |
| PPIC | 0.56 | 0.00 | 0.00 | 0.62 | 0.36 | 3 | Myofibroblast |
| FNDC3B | 0.66 | 0.00 | 0.00 | 0.28 | 0.10 | 3 | Myofibroblast |
| MYL6 | 0.55 | 0.00 | 0.00 | 0.86 | 0.71 | 3 | Myofibroblast |
| RCN2 | 0.65 | 0.00 | 0.00 | 0.35 | 0.15 | 3 | Myofibroblast |
| SH3PXD2A | 0.63 | 0.00 | 0.00 | 0.27 | 0.09 | 3 | Myofibroblast |
| MLLT3 | 0.70 | 0.00 | 0.00 | 0.27 | 0.09 | 3 | Myofibroblast |
| LAMB1 | 0.62 | 0.00 | 0.00 | 0.34 | 0.14 | 3 | Myofibroblast |
| MXRA8 | 0.51 | 0.00 | 0.00 | 0.37 | 0.15 | 3 | Myofibroblast |
| RYK | 0.66 | 0.00 | 0.00 | 0.28 | 0.10 | 3 | Myofibroblast |
| C9orf3 | 0.64 | 0.00 | 0.00 | 0.26 | 0.09 | 3 | Myofibroblast |
| ADAMTS10 | 0.65 | 0.00 | 0.00 | 0.26 | 0.09 | 3 | Myofibroblast |
| DSTN | 0.56 | 0.00 | 0.00 | 0.74 | 0.51 | 3 | Myofibroblast |
| FGFR1 | 0.59 | 0.00 | 0.00 | 0.27 | 0.10 | 3 | Myofibroblast |
| ANTXR1 | 0.57 | 0.00 | 0.00 | 0.30 | 0.11 | 3 | Myofibroblast |
| HSP90B1 | 0.47 | 0.00 | 0.00 | 0.87 | 0.74 | 3 | Myofibroblast |
| P4HB | 0.56 | 0.00 | 0.00 | 0.63 | 0.40 | 3 | Myofibroblast |
| MAGED2 | 0.48 | 0.00 | 0.00 | 0.40 | 0.19 | 3 | Myofibroblast |
| NFIX | 0.60 | 0.00 | 0.00 | 0.34 | 0.15 | 3 | Myofibroblast |
| ARF4 | 0.54 | 0.00 | 0.00 | 0.50 | 0.27 | 3 | Myofibroblast |
| GPX8 | 0.60 | 0.00 | 0.00 | 0.29 | 0.12 | 3 | Myofibroblast |
| CD81 | 0.46 | 0.00 | 0.00 | 0.62 | 0.38 | 3 | Myofibroblast |
| TTC3 | 0.57 | 0.00 | 0.00 | 0.49 | 0.27 | 3 | Myofibroblast |
| PARVA | 0.54 | 0.00 | 0.00 | 0.32 | 0.14 | 3 | Myofibroblast |
| AMOTL1 | 0.61 | 0.00 | 0.00 | 0.31 | 0.14 | 3 | Myofibroblast |
| SNAI2 | 0.56 | 0.00 | 0.00 | 0.27 | 0.11 | 3 | Myofibroblast |
| MORF4L2 | 0.54 | 0.00 | 0.00 | 0.42 | 0.22 | 3 | Myofibroblast |
| NUPR1 | 0.36 | 0.00 | 0.00 | 0.39 | 0.19 | 3 | Myofibroblast |
| MAGED1 | 0.53 | 0.00 | 0.00 | 0.35 | 0.17 | 3 | Myofibroblast |
| VCAN | 0.48 | 0.00 | 0.00 | 0.28 | 0.12 | 3 | Myofibroblast |
| CDK4 | 0.51 | 0.00 | 0.00 | 0.42 | 0.22 | 3 | Myofibroblast |
| TWSG1 | 0.53 | 0.00 | 0.00 | 0.25 | 0.11 | 3 | Myofibroblast |
| CKAP4 | 0.52 | 0.00 | 0.00 | 0.34 | 0.16 | 3 | Myofibroblast |
| TSC22D1 | 0.46 | 0.00 | 0.00 | 0.54 | 0.33 | 3 | Myofibroblast |
| GNG12 | 0.48 | 0.00 | 0.00 | 0.26 | 0.11 | 3 | Myofibroblast |
| NFIC | 0.45 | 0.00 | 0.00 | 0.43 | 0.23 | 3 | Myofibroblast |
| PICALM | 0.57 | 0.00 | 0.00 | 0.36 | 0.19 | 3 | Myofibroblast |
| PLS3 | 0.48 | 0.00 | 0.00 | 0.38 | 0.20 | 3 | Myofibroblast |
| MINOS1 | 0.45 | 0.00 | 0.00 | 0.62 | 0.43 | 3 | Myofibroblast |
| RRBP1 | 0.46 | 0.00 | 0.00 | 0.71 | 0.53 | 3 | Myofibroblast |
| ANXA6 | 0.46 | 0.00 | 0.00 | 0.39 | 0.21 | 3 | Myofibroblast |
| PRDX4 | 0.47 | 0.00 | 0.00 | 0.30 | 0.15 | 3 | Myofibroblast |
| SVIL | 0.51 | 0.00 | 0.00 | 0.31 | 0.15 | 3 | Myofibroblast |
| TBX2 | 0.41 | 0.00 | 0.00 | 0.28 | 0.12 | 3 | Myofibroblast |
| NFIA | 0.46 | 0.00 | 0.00 | 0.41 | 0.23 | 3 | Myofibroblast |
| NDUFA11 | 0.44 | 0.00 | 0.00 | 0.47 | 0.28 | 3 | Myofibroblast |
| FKBP9 | 0.44 | 0.00 | 0.00 | 0.27 | 0.13 | 3 | Myofibroblast |
| VIM | 0.29 | 0.00 | 0.00 | 0.89 | 0.67 | 3 | Myofibroblast |
| LAMC1 | 0.45 | 0.00 | 0.00 | 0.30 | 0.15 | 3 | Myofibroblast |
| HMGN1 | 0.38 | 0.00 | 0.00 | 0.64 | 0.44 | 3 | Myofibroblast |
| PHLDB2 | 0.47 | 0.00 | 0.00 | 0.44 | 0.27 | 3 | Myofibroblast |
| TJP2 | 0.54 | 0.00 | 0.00 | 0.29 | 0.15 | 3 | Myofibroblast |
| CSRP1 | 0.56 | 0.00 | 0.00 | 0.33 | 0.18 | 3 | Myofibroblast |
| ITM2C | 0.44 | 0.00 | 0.00 | 0.39 | 0.23 | 3 | Myofibroblast |
| ARID5B | 0.46 | 0.00 | 0.00 | 0.37 | 0.21 | 3 | Myofibroblast |
| MARCKSL1 | 0.36 | 0.00 | 0.00 | 0.39 | 0.22 | 3 | Myofibroblast |
| PRDX2 | 0.38 | 0.00 | 0.00 | 0.52 | 0.34 | 3 | Myofibroblast |
| DST | 0.44 | 0.00 | 0.00 | 0.29 | 0.15 | 3 | Myofibroblast |
| CDC42EP3 | 0.48 | 0.00 | 0.00 | 0.26 | 0.12 | 3 | Myofibroblast |
| SEPT2 | 0.37 | 0.00 | 0.00 | 0.53 | 0.35 | 3 | Myofibroblast |
| DDAH2 | 0.34 | 0.00 | 0.00 | 0.45 | 0.27 | 3 | Myofibroblast |
| NEDD4 | 0.38 | 0.00 | 0.00 | 0.65 | 0.48 | 3 | Myofibroblast |
| SEPT11 | 0.41 | 0.00 | 0.00 | 0.37 | 0.21 | 3 | Myofibroblast |
| NPTN | 0.42 | 0.00 | 0.00 | 0.33 | 0.19 | 3 | Myofibroblast |
| LAPTM4A | 0.30 | 0.00 | 0.00 | 0.72 | 0.54 | 3 | Myofibroblast |
| SLMAP | 0.45 | 0.00 | 0.00 | 0.27 | 0.14 | 3 | Myofibroblast |
| SSR2 | 0.35 | 0.00 | 0.00 | 0.50 | 0.33 | 3 | Myofibroblast |
| SLC25A4 | 0.34 | 0.00 | 0.00 | 0.62 | 0.46 | 3 | Myofibroblast |
| DYNLRB1 | 0.37 | 0.00 | 0.00 | 0.51 | 0.34 | 3 | Myofibroblast |
| CCND2 | 0.50 | 0.00 | 0.00 | 0.41 | 0.26 | 3 | Myofibroblast |
| KDELR2 | 0.38 | 0.00 | 0.00 | 0.34 | 0.20 | 3 | Myofibroblast |
| NR2F2 | 0.26 | 0.00 | 0.00 | 0.35 | 0.20 | 3 | Myofibroblast |
| C5orf15 | 0.37 | 0.00 | 0.00 | 0.29 | 0.16 | 3 | Myofibroblast |
| TMED9 | 0.35 | 0.00 | 0.00 | 0.42 | 0.27 | 3 | Myofibroblast |
| CRYAB | 0.30 | 0.00 | 0.00 | 0.28 | 0.15 | 3 | Myofibroblast |
| RBBP7 | 0.34 | 0.00 | 0.00 | 0.38 | 0.23 | 3 | Myofibroblast |
| SERF2 | 0.28 | 0.00 | 0.00 | 0.74 | 0.58 | 3 | Myofibroblast |
| RSU1 | 0.35 | 0.00 | 0.00 | 0.33 | 0.19 | 3 | Myofibroblast |
| RTN4 | 0.36 | 0.00 | 0.00 | 0.40 | 0.25 | 3 | Myofibroblast |
| ATP2B1 | 0.35 | 0.00 | 0.00 | 0.66 | 0.55 | 3 | Myofibroblast |
| MRFAP1 | 0.30 | 0.00 | 0.00 | 0.50 | 0.34 | 3 | Myofibroblast |
| HSPA5 | 0.31 | 0.00 | 0.00 | 0.69 | 0.53 | 3 | Myofibroblast |
| LMAN1 | 0.35 | 0.00 | 0.00 | 0.30 | 0.17 | 3 | Myofibroblast |
| THOC7 | 0.34 | 0.00 | 0.00 | 0.44 | 0.29 | 3 | Myofibroblast |
| MAP1LC3A | 0.38 | 0.00 | 0.00 | 0.32 | 0.20 | 3 | Myofibroblast |
| MYH9 | 0.29 | 0.00 | 0.00 | 0.48 | 0.33 | 3 | Myofibroblast |
| TIA1 | 0.31 | 0.00 | 0.00 | 0.33 | 0.20 | 3 | Myofibroblast |
| CBX1 | 0.30 | 0.00 | 0.00 | 0.54 | 0.38 | 3 | Myofibroblast |
| NDUFS5 | 0.46 | 0.00 | 0.00 | 0.38 | 0.26 | 3 | Myofibroblast |
| COX20 | 0.30 | 0.00 | 0.00 | 0.29 | 0.17 | 3 | Myofibroblast |
| ZFP36L1 | 0.29 | 0.00 | 0.00 | 0.60 | 0.45 | 3 | Myofibroblast |
| RANBP1 | 0.30 | 0.00 | 0.00 | 0.43 | 0.29 | 3 | Myofibroblast |
| DDOST | 0.34 | 0.00 | 0.00 | 0.31 | 0.19 | 3 | Myofibroblast |
| PDAP1 | 0.29 | 0.00 | 0.00 | 0.57 | 0.43 | 3 | Myofibroblast |
| PDIA6 | 0.31 | 0.00 | 0.00 | 0.45 | 0.31 | 3 | Myofibroblast |
| GSTP1 | 0.27 | 0.00 | 0.00 | 0.36 | 0.23 | 3 | Myofibroblast |
| IDH2 | 0.31 | 0.00 | 0.00 | 0.30 | 0.19 | 3 | Myofibroblast |
| PDLIM1 | 0.30 | 0.00 | 0.00 | 0.26 | 0.15 | 3 | Myofibroblast |
| MYO1B | 0.30 | 0.00 | 0.00 | 0.26 | 0.15 | 3 | Myofibroblast |
| EPRS | 0.34 | 0.00 | 0.00 | 0.32 | 0.20 | 3 | Myofibroblast |
| HNRNPA1 | 0.43 | 0.00 | 0.00 | 0.42 | 0.30 | 3 | Myofibroblast |
| RPN2 | 0.31 | 0.00 | 0.00 | 0.32 | 0.20 | 3 | Myofibroblast |
| LARP7 | 0.31 | 0.00 | 0.00 | 0.27 | 0.16 | 3 | Myofibroblast |
| PPIB | 0.26 | 0.00 | 0.00 | 0.59 | 0.45 | 3 | Myofibroblast |
| SPCS1 | 0.26 | 0.00 | 0.00 | 0.43 | 0.29 | 3 | Myofibroblast |
| HNRNPR | 0.28 | 0.00 | 0.00 | 0.37 | 0.25 | 3 | Myofibroblast |
| NENF | 0.31 | 0.00 | 0.00 | 0.30 | 0.19 | 3 | Myofibroblast |
| HDLBP | 0.31 | 0.00 | 0.00 | 0.32 | 0.20 | 3 | Myofibroblast |
| SEC13 | 0.27 | 0.00 | 0.00 | 0.28 | 0.17 | 3 | Myofibroblast |
| YBX3 | 0.32 | 0.00 | 0.00 | 0.29 | 0.18 | 3 | Myofibroblast |
| MIF | 0.26 | 0.00 | 0.00 | 0.34 | 0.22 | 3 | Myofibroblast |
| TMEM59 | 0.27 | 0.00 | 0.00 | 0.55 | 0.41 | 3 | Myofibroblast |
| COPB2 | 0.31 | 0.00 | 0.00 | 0.27 | 0.17 | 3 | Myofibroblast |
| FUNDC2 | 0.27 | 0.00 | 0.00 | 0.27 | 0.16 | 3 | Myofibroblast |
| ROMO1 | 0.26 | 0.00 | 0.00 | 0.43 | 0.30 | 3 | Myofibroblast |
| PSMD7 | 0.25 | 0.00 | 0.00 | 0.40 | 0.27 | 3 | Myofibroblast |
| VEZF1 | 0.32 | 0.00 | 0.00 | 0.32 | 0.21 | 3 | Myofibroblast |
| CALR | 0.30 | 0.00 | 0.00 | 0.56 | 0.44 | 3 | Myofibroblast |
| SAR1A | 0.29 | 0.00 | 0.00 | 0.31 | 0.20 | 3 | Myofibroblast |
| GLG1 | 0.28 | 0.00 | 0.00 | 0.29 | 0.18 | 3 | Myofibroblast |
| CNPY2 | 0.27 | 0.00 | 0.00 | 0.31 | 0.20 | 3 | Myofibroblast |
| TXNL1 | 0.26 | 0.00 | 0.00 | 0.29 | 0.18 | 3 | Myofibroblast |
| SFR1 | 0.28 | 0.00 | 0.00 | 0.41 | 0.29 | 3 | Myofibroblast |
| DDX1 | 0.28 | 0.00 | 0.00 | 0.25 | 0.16 | 3 | Myofibroblast |
| RBPMS | 0.28 | 0.00 | 0.00 | 0.28 | 0.18 | 3 | Myofibroblast |
| SUMO2 | 0.25 | 0.00 | 0.00 | 0.45 | 0.33 | 3 | Myofibroblast |
| ILF2 | 0.29 | 0.00 | 0.00 | 0.29 | 0.19 | 3 | Myofibroblast |
| RHEB | 0.27 | 0.00 | 0.00 | 0.29 | 0.19 | 3 | Myofibroblast |
| CCNI | 0.38 | 0.00 | 0.00 | 0.29 | 0.19 | 3 | Myofibroblast |
| NARS | 0.25 | 0.00 | 0.00 | 0.42 | 0.30 | 3 | Myofibroblast |
| CFL2 | 0.25 | 0.00 | 0.00 | 0.34 | 0.23 | 3 | Myofibroblast |
| TMEM167A | 0.26 | 0.00 | 0.00 | 0.31 | 0.21 | 3 | Myofibroblast |
| BCCIP | 0.27 | 0.00 | 0.00 | 0.29 | 0.19 | 3 | Myofibroblast |
| GINM1 | 0.27 | 0.00 | 0.00 | 0.33 | 0.22 | 3 | Myofibroblast |
| PURA | 0.30 | 0.00 | 0.00 | 0.29 | 0.19 | 3 | Myofibroblast |
| CLIP1 | 0.30 | 0.00 | 0.00 | 0.27 | 0.18 | 3 | Myofibroblast |
| PURB | 0.26 | 0.00 | 0.00 | 0.43 | 0.33 | 3 | Myofibroblast |
| CTSS | 3.35 | 0.00 | 0.00 | 0.90 | 0.06 | 4 | Macrophage |
| S100A8 | 3.18 | 0.00 | 0.00 | 0.38 | 0.08 | 4 | Macrophage |
| TYROBP | 2.89 | 0.00 | 0.00 | 0.85 | 0.05 | 4 | Macrophage |
| FCER1G | 2.85 | 0.00 | 0.00 | 0.81 | 0.04 | 4 | Macrophage |
| CYBB | 2.71 | 0.00 | 0.00 | 0.70 | 0.03 | 4 | Macrophage |
| CCR2 | 2.57 | 0.00 | 0.00 | 0.52 | 0.03 | 4 | Macrophage |
| MPEG1 | 2.45 | 0.00 | 0.00 | 0.63 | 0.01 | 4 | Macrophage |
| ALOX5AP | 2.42 | 0.00 | 0.00 | 0.70 | 0.03 | 4 | Macrophage |
| PLAC8 | 2.34 | 0.00 | 0.00 | 0.58 | 0.12 | 4 | Macrophage |
| LGALS3 | 2.32 | 0.00 | 0.00 | 0.75 | 0.10 | 4 | Macrophage |
| LST1 | 2.29 | 0.00 | 0.00 | 0.54 | 0.02 | 4 | Macrophage |
| SPI1 | 2.10 | 0.00 | 0.00 | 0.53 | 0.01 | 4 | Macrophage |
| PLA2G7 | 2.07 | 0.00 | 0.00 | 0.42 | 0.01 | 4 | Macrophage |
| MSRB1 | 2.04 | 0.00 | 0.00 | 0.62 | 0.09 | 4 | Macrophage |
| LAPTM5 | 2.03 | 0.00 | 0.00 | 0.74 | 0.07 | 4 | Macrophage |
| MNDA | 1.92 | 0.00 | 0.00 | 0.41 | 0.02 | 4 | Macrophage |
| APOE | 1.85 | 0.00 | 0.00 | 0.54 | 0.24 | 4 | Macrophage |
| CYBA | 1.85 | 0.00 | 0.00 | 0.83 | 0.26 | 4 | Macrophage |
| IGSF6 | 1.84 | 0.00 | 0.00 | 0.40 | 0.01 | 4 | Macrophage |
| AIF1 | 1.82 | 0.00 | 0.00 | 0.29 | 0.01 | 4 | Macrophage |
| PLBD1 | 1.82 | 0.00 | 0.00 | 0.37 | 0.01 | 4 | Macrophage |
| GDA | 1.76 | 0.00 | 0.00 | 0.35 | 0.01 | 4 | Macrophage |
| S100A4 | 1.76 | 0.00 | 0.00 | 0.47 | 0.06 | 4 | Macrophage |
| PLEK | 1.72 | 0.00 | 0.00 | 0.42 | 0.03 | 4 | Macrophage |
| PYGL | 1.65 | 0.00 | 0.00 | 0.44 | 0.06 | 4 | Macrophage |
| CST3 | 1.65 | 0.00 | 0.00 | 0.86 | 0.56 | 4 | Macrophage |
| APBB1IP | 1.65 | 0.00 | 0.00 | 0.48 | 0.05 | 4 | Macrophage |
| PLD4 | 1.64 | 0.00 | 0.00 | 0.39 | 0.02 | 4 | Macrophage |
| SAMSN1 | 1.64 | 0.00 | 0.00 | 0.37 | 0.03 | 4 | Macrophage |
| TNFAIP2 | 1.61 | 0.00 | 0.00 | 0.32 | 0.02 | 4 | Macrophage |
| CD52 | 1.61 | 0.00 | 0.00 | 0.79 | 0.13 | 4 | Macrophage |
| PTPRC | 1.60 | 0.00 | 0.00 | 0.64 | 0.09 | 4 | Macrophage |
| CD300LF | 1.60 | 0.00 | 0.00 | 0.31 | 0.01 | 4 | Macrophage |
| GSR | 1.59 | 0.00 | 0.00 | 0.37 | 0.05 | 4 | Macrophage |
| RNASE6 | 1.58 | 0.00 | 0.00 | 0.34 | 0.02 | 4 | Macrophage |
| CORO1A | 1.57 | 0.00 | 0.00 | 0.49 | 0.06 | 4 | Macrophage |
| SAMHD1 | 1.56 | 0.00 | 0.00 | 0.61 | 0.15 | 4 | Macrophage |
| LCP1 | 1.54 | 0.00 | 0.00 | 0.66 | 0.16 | 4 | Macrophage |
| CTSB | 1.53 | 0.00 | 0.00 | 0.62 | 0.25 | 4 | Macrophage |
| CSF1R | 1.52 | 0.00 | 0.00 | 0.31 | 0.01 | 4 | Macrophage |
| PRDX5 | 1.51 | 0.00 | 0.00 | 0.69 | 0.29 | 4 | Macrophage |
| ARHGAP30 | 1.51 | 0.00 | 0.00 | 0.49 | 0.06 | 4 | Macrophage |
| GM2A | 1.47 | 0.00 | 0.00 | 0.39 | 0.05 | 4 | Macrophage |
| GPX1 | 1.47 | 0.00 | 0.00 | 0.84 | 0.40 | 4 | Macrophage |
| CSF2RA | 1.46 | 0.00 | 0.00 | 0.34 | 0.03 | 4 | Macrophage |
| CEBPB | 1.45 | 0.00 | 0.00 | 0.36 | 0.07 | 4 | Macrophage |
| CFP | 1.44 | 0.00 | 0.00 | 0.31 | 0.02 | 4 | Macrophage |
| COTL1 | 1.44 | 0.00 | 0.00 | 0.56 | 0.13 | 4 | Macrophage |
| GNGT2 | 1.44 | 0.00 | 0.00 | 0.33 | 0.08 | 4 | Macrophage |
| CD44 | 1.43 | 0.00 | 0.00 | 0.57 | 0.17 | 4 | Macrophage |
| PTPN6 | 1.43 | 0.00 | 0.00 | 0.42 | 0.05 | 4 | Macrophage |
| NCF2 | 1.43 | 0.00 | 0.00 | 0.33 | 0.01 | 4 | Macrophage |
| FOS | 1.42 | 0.00 | 0.00 | 0.48 | 0.14 | 4 | Macrophage |
| ITGAL | 1.42 | 0.00 | 0.00 | 0.27 | 0.02 | 4 | Macrophage |
| TPD52 | 1.40 | 0.00 | 0.00 | 0.38 | 0.07 | 4 | Macrophage |
| NCF1 | 1.39 | 0.00 | 0.00 | 0.36 | 0.03 | 4 | Macrophage |
| HCLS1 | 1.39 | 0.00 | 0.00 | 0.39 | 0.04 | 4 | Macrophage |
| SNX10 | 1.39 | 0.00 | 0.00 | 0.31 | 0.02 | 4 | Macrophage |
| TALDO1 | 1.37 | 0.00 | 0.00 | 0.65 | 0.23 | 4 | Macrophage |
| ARHGDIB | 1.37 | 0.00 | 0.00 | 0.60 | 0.12 | 4 | Macrophage |
| CCR1 | 1.36 | 0.00 | 0.00 | 0.27 | 0.01 | 4 | Macrophage |
| IFITM2 | 1.33 | 0.00 | 0.00 | 0.31 | 0.06 | 4 | Macrophage |
| LY86 | 1.33 | 0.00 | 0.00 | 0.34 | 0.03 | 4 | Macrophage |
| CYTIP | 1.31 | 0.00 | 0.00 | 0.37 | 0.06 | 4 | Macrophage |
| CTSC | 1.29 | 0.00 | 0.00 | 0.53 | 0.17 | 4 | Macrophage |
| MARCH1 | 1.28 | 0.00 | 0.00 | 0.26 | 0.01 | 4 | Macrophage |
| FAM49B | 1.28 | 0.00 | 0.00 | 0.40 | 0.07 | 4 | Macrophage |
| FGR | 1.28 | 0.00 | 0.00 | 0.25 | 0.01 | 4 | Macrophage |
| ARPC1B | 1.27 | 0.00 | 0.00 | 0.73 | 0.33 | 4 | Macrophage |
| FAM111A | 1.26 | 0.00 | 0.00 | 0.43 | 0.11 | 4 | Macrophage |
| HCK | 1.26 | 0.00 | 0.00 | 0.27 | 0.02 | 4 | Macrophage |
| CD84 | 1.26 | 0.00 | 0.00 | 0.28 | 0.02 | 4 | Macrophage |
| IL6R | 1.25 | 0.00 | 0.00 | 0.34 | 0.05 | 4 | Macrophage |
| CORO2A | 1.24 | 0.00 | 0.00 | 0.29 | 0.03 | 4 | Macrophage |
| FTH1 | 1.24 | 0.00 | 0.00 | 0.93 | 0.81 | 4 | Macrophage |
| PLIN2 | 1.23 | 0.00 | 0.00 | 0.37 | 0.10 | 4 | Macrophage |
| ITGA4 | 1.22 | 0.00 | 0.00 | 0.33 | 0.07 | 4 | Macrophage |
| CASP1 | 1.21 | 0.00 | 0.00 | 0.25 | 0.02 | 4 | Macrophage |
| CD302 | 1.20 | 0.00 | 0.00 | 0.41 | 0.12 | 4 | Macrophage |
| CDK2AP2 | 1.20 | 0.00 | 0.00 | 0.39 | 0.12 | 4 | Macrophage |
| PRKCD | 1.20 | 0.00 | 0.00 | 0.37 | 0.08 | 4 | Macrophage |
| PTPRE | 1.17 | 0.00 | 0.00 | 0.30 | 0.04 | 4 | Macrophage |
| LSP1 | 1.17 | 0.00 | 0.00 | 0.56 | 0.15 | 4 | Macrophage |
| EFHD2 | 1.17 | 0.00 | 0.00 | 0.32 | 0.06 | 4 | Macrophage |
| EMB | 1.17 | 0.00 | 0.00 | 0.35 | 0.08 | 4 | Macrophage |
| UNC93B1 | 1.16 | 0.00 | 0.00 | 0.31 | 0.04 | 4 | Macrophage |
| PTPN18 | 1.14 | 0.00 | 0.00 | 0.42 | 0.10 | 4 | Macrophage |
| TMSB4X | 1.12 | 0.00 | 0.00 | 0.99 | 0.89 | 4 | Macrophage |
| CD53 | 1.11 | 0.00 | 0.00 | 0.38 | 0.06 | 4 | Macrophage |
| PYCARD | 1.11 | 0.00 | 0.00 | 0.34 | 0.07 | 4 | Macrophage |
| APRT | 1.10 | 0.00 | 0.00 | 0.45 | 0.16 | 4 | Macrophage |
| BAZ1A | 1.09 | 0.00 | 0.00 | 0.35 | 0.11 | 4 | Macrophage |
| ITGAM | 1.09 | 0.00 | 0.00 | 0.53 | 0.23 | 4 | Macrophage |
| PTPN1 | 1.09 | 0.00 | 0.00 | 0.44 | 0.17 | 4 | Macrophage |
| CTSZ | 1.08 | 0.00 | 0.00 | 0.39 | 0.13 | 4 | Macrophage |
| PSMB8 | 1.07 | 0.00 | 0.00 | 0.48 | 0.16 | 4 | Macrophage |
| SMPDL3A | 1.05 | 0.00 | 0.00 | 0.31 | 0.09 | 4 | Macrophage |
| RGS2 | 1.02 | 0.00 | 0.00 | 0.60 | 0.24 | 4 | Macrophage |
| SH3BGRL3 | 1.01 | 0.00 | 0.00 | 0.64 | 0.33 | 4 | Macrophage |
| ARID3A | 1.01 | 0.00 | 0.00 | 0.30 | 0.08 | 4 | Macrophage |
| FERMT3 | 1.01 | 0.00 | 0.00 | 0.28 | 0.04 | 4 | Macrophage |
| CAPG | 0.98 | 0.00 | 0.00 | 0.31 | 0.08 | 4 | Macrophage |
| ACTR3 | 0.96 | 0.00 | 0.00 | 0.61 | 0.30 | 4 | Macrophage |
| LYN | 0.96 | 0.00 | 0.00 | 0.34 | 0.10 | 4 | Macrophage |
| ACTR2 | 0.96 | 0.00 | 0.00 | 0.53 | 0.25 | 4 | Macrophage |
| ARPC5 | 0.96 | 0.00 | 0.00 | 0.83 | 0.55 | 4 | Macrophage |
| ARPC2 | 0.92 | 0.00 | 0.00 | 0.84 | 0.60 | 4 | Macrophage |
| RAC2 | 0.81 | 0.00 | 0.00 | 0.35 | 0.08 | 4 | Macrophage |
| B2M | 0.79 | 0.00 | 0.00 | 0.89 | 0.68 | 4 | Macrophage |
| S100A6 | 0.78 | 0.00 | 0.00 | 0.71 | 0.32 | 4 | Macrophage |
| FAU | 0.69 | 0.00 | 0.00 | 0.96 | 0.87 | 4 | Macrophage |
| GMFG | 0.86 | 0.00 | 0.00 | 0.38 | 0.13 | 4 | Macrophage |
| ZFP36L2 | 0.95 | 0.00 | 0.00 | 0.36 | 0.12 | 4 | Macrophage |
| IFNGR1 | 1.00 | 0.00 | 0.00 | 0.47 | 0.21 | 4 | Macrophage |
| CTSA | 0.95 | 0.00 | 0.00 | 0.33 | 0.11 | 4 | Macrophage |
| EMP3 | 0.99 | 0.00 | 0.00 | 0.48 | 0.22 | 4 | Macrophage |
| CREG1 | 0.98 | 0.00 | 0.00 | 0.39 | 0.15 | 4 | Macrophage |
| LRRC58 | 0.53 | 0.00 | 0.00 | 0.94 | 0.85 | 4 | Macrophage |
| CAPZA2 | 0.86 | 0.00 | 0.00 | 0.59 | 0.34 | 4 | Macrophage |
| AKAP13 | 0.91 | 0.00 | 0.00 | 0.50 | 0.24 | 4 | Macrophage |
| CTSD | 1.29 | 0.00 | 0.00 | 0.46 | 0.23 | 4 | Macrophage |
| ATP6V0C | 0.98 | 0.00 | 0.00 | 0.44 | 0.21 | 4 | Macrophage |
| RPS27A | 0.47 | 0.00 | 0.00 | 0.90 | 0.77 | 4 | Macrophage |
| JUNB | 0.93 | 0.00 | 0.00 | 0.42 | 0.19 | 4 | Macrophage |
| LLPH | 0.90 | 0.00 | 0.00 | 0.53 | 0.29 | 4 | Macrophage |
| LRP1 | 0.84 | 0.00 | 0.00 | 0.30 | 0.10 | 4 | Macrophage |
| TPM3 | 0.55 | 0.00 | 0.00 | 0.84 | 0.65 | 4 | Macrophage |
| WDR89 | 0.44 | 0.00 | 0.00 | 0.92 | 0.81 | 4 | Macrophage |
| ZFP36 | 0.94 | 0.00 | 0.00 | 0.35 | 0.14 | 4 | Macrophage |
| PSMB10 | 0.81 | 0.00 | 0.00 | 0.33 | 0.12 | 4 | Macrophage |
| S100A11 | 0.85 | 0.00 | 0.00 | 0.67 | 0.48 | 4 | Macrophage |
| CLTA | 0.75 | 0.00 | 0.00 | 0.63 | 0.41 | 4 | Macrophage |
| BTG1 | 0.89 | 0.00 | 0.00 | 0.52 | 0.28 | 4 | Macrophage |
| CORO1B | 0.82 | 0.00 | 0.00 | 0.45 | 0.22 | 4 | Macrophage |
| PPT1 | 0.85 | 0.00 | 0.00 | 0.31 | 0.11 | 4 | Macrophage |
| TKT | 0.84 | 0.00 | 0.00 | 0.33 | 0.13 | 4 | Macrophage |
| STAT1 | 0.91 | 0.00 | 0.00 | 0.26 | 0.09 | 4 | Macrophage |
| SERP1 | 0.80 | 0.00 | 0.00 | 0.45 | 0.22 | 4 | Macrophage |
| SEC11C | 0.79 | 0.00 | 0.00 | 0.35 | 0.15 | 4 | Macrophage |
| IQGAP1 | 0.70 | 0.00 | 0.00 | 0.64 | 0.41 | 4 | Macrophage |
| CRIP1 | 0.78 | 0.00 | 0.00 | 0.66 | 0.46 | 4 | Macrophage |
| PFN1 | 0.67 | 0.00 | 0.00 | 0.67 | 0.47 | 4 | Macrophage |
| ARPC4 | 0.77 | 0.00 | 0.00 | 0.37 | 0.17 | 4 | Macrophage |
| CCDC88A | 0.86 | 0.00 | 0.00 | 0.42 | 0.21 | 4 | Macrophage |
| ANXA1 | 0.85 | 0.00 | 0.00 | 0.39 | 0.18 | 4 | Macrophage |
| SRGN | 0.73 | 0.00 | 0.00 | 0.55 | 0.30 | 4 | Macrophage |
| ENO1 | 0.79 | 0.00 | 0.00 | 0.37 | 0.17 | 4 | Macrophage |
| PSMA7 | 0.65 | 0.00 | 0.00 | 0.71 | 0.56 | 4 | Macrophage |
| PTBP3 | 0.71 | 0.00 | 0.00 | 0.53 | 0.31 | 4 | Macrophage |
| RHOG | 0.75 | 0.00 | 0.00 | 0.27 | 0.10 | 4 | Macrophage |
| CSTB | 1.29 | 0.00 | 0.00 | 0.46 | 0.27 | 4 | Macrophage |
| GNS | 0.82 | 0.00 | 0.00 | 0.27 | 0.10 | 4 | Macrophage |
| PSME2 | 0.79 | 0.00 | 0.00 | 0.44 | 0.24 | 4 | Macrophage |
| UCP2 | 0.73 | 0.00 | 0.00 | 0.27 | 0.10 | 4 | Macrophage |
| PKM | 0.80 | 0.00 | 0.00 | 0.41 | 0.22 | 4 | Macrophage |
| GNAI2 | 0.60 | 0.00 | 0.00 | 0.58 | 0.37 | 4 | Macrophage |
| ATOX1 | 0.75 | 0.00 | 0.00 | 0.54 | 0.36 | 4 | Macrophage |
| ATP6V1B2 | 0.76 | 0.00 | 0.00 | 0.28 | 0.11 | 4 | Macrophage |
| REEP5 | 0.75 | 0.00 | 0.00 | 0.42 | 0.23 | 4 | Macrophage |
| OSTF1 | 0.66 | 0.00 | 0.00 | 0.43 | 0.24 | 4 | Macrophage |
| STK38 | 0.72 | 0.00 | 0.00 | 0.34 | 0.17 | 4 | Macrophage |
| POMP | 0.67 | 0.00 | 0.00 | 0.50 | 0.32 | 4 | Macrophage |
| TMED5 | 0.69 | 0.00 | 0.00 | 0.32 | 0.14 | 4 | Macrophage |
| LY6E | 0.53 | 0.00 | 0.00 | 0.62 | 0.41 | 4 | Macrophage |
| ATP6V1E1 | 0.79 | 0.00 | 0.00 | 0.42 | 0.24 | 4 | Macrophage |
| AHNAK | 0.59 | 0.00 | 0.00 | 0.60 | 0.42 | 4 | Macrophage |
| MRPL33 | 0.72 | 0.00 | 0.00 | 0.46 | 0.28 | 4 | Macrophage |
| RPLP2 | 0.32 | 0.00 | 0.00 | 0.91 | 0.82 | 4 | Macrophage |
| ATP6V0B | 0.68 | 0.00 | 0.00 | 0.36 | 0.19 | 4 | Macrophage |
| M6PR | 0.72 | 0.00 | 0.00 | 0.39 | 0.22 | 4 | Macrophage |
| LRRFIP1 | 0.64 | 0.00 | 0.00 | 0.42 | 0.24 | 4 | Macrophage |
| MYCBP2 | 0.66 | 0.00 | 0.00 | 0.38 | 0.21 | 4 | Macrophage |
| DBI | 0.64 | 0.00 | 0.00 | 0.53 | 0.36 | 4 | Macrophage |
| GABARAP | 0.60 | 0.00 | 0.00 | 0.58 | 0.43 | 4 | Macrophage |
| CAPZB | 0.57 | 0.00 | 0.00 | 0.58 | 0.42 | 4 | Macrophage |
| ARPC3 | 0.63 | 0.00 | 0.00 | 0.44 | 0.27 | 4 | Macrophage |
| TSPO | 0.69 | 0.00 | 0.00 | 0.41 | 0.25 | 4 | Macrophage |
| ERP29 | 0.64 | 0.00 | 0.00 | 0.42 | 0.25 | 4 | Macrophage |
| CD47 | 0.57 | 0.00 | 0.00 | 0.57 | 0.40 | 4 | Macrophage |
| DUSP6 | 0.70 | 0.00 | 0.00 | 0.33 | 0.17 | 4 | Macrophage |
| RAP1B | 0.68 | 0.00 | 0.00 | 0.39 | 0.23 | 4 | Macrophage |
| HSPA8 | 0.45 | 0.00 | 0.00 | 0.77 | 0.67 | 4 | Macrophage |
| COX4I1 | 0.40 | 0.00 | 0.00 | 0.81 | 0.74 | 4 | Macrophage |
| ANXA2 | 0.55 | 0.00 | 0.00 | 0.62 | 0.48 | 4 | Macrophage |
| SKAP2 | 0.61 | 0.00 | 0.00 | 0.27 | 0.13 | 4 | Macrophage |
| ACTG1 | 0.33 | 0.00 | 0.00 | 0.86 | 0.77 | 4 | Macrophage |
| CFL1 | 0.44 | 0.00 | 0.00 | 0.71 | 0.60 | 4 | Macrophage |
| GRB2 | 0.58 | 0.00 | 0.00 | 0.31 | 0.16 | 4 | Macrophage |
| SPPL2A | 0.62 | 0.00 | 0.00 | 0.27 | 0.13 | 4 | Macrophage |
| ABRACL | 0.62 | 0.00 | 0.00 | 0.37 | 0.21 | 4 | Macrophage |
| PSME1 | 0.56 | 0.00 | 0.00 | 0.48 | 0.32 | 4 | Macrophage |
| GLUD2 | 0.63 | 0.00 | 0.00 | 0.28 | 0.14 | 4 | Macrophage |
| LAMTOR4 | 0.62 | 0.00 | 0.00 | 0.27 | 0.13 | 4 | Macrophage |
| HMGB2 | 0.55 | 0.00 | 0.00 | 0.53 | 0.37 | 4 | Macrophage |
| C6orf62 | 0.62 | 0.00 | 0.00 | 0.46 | 0.30 | 4 | Macrophage |
| TOR1AIP1 | 0.60 | 0.00 | 0.00 | 0.31 | 0.16 | 4 | Macrophage |
| MSN | 0.44 | 0.00 | 0.00 | 0.60 | 0.43 | 4 | Macrophage |
| ANP32B | 0.59 | 0.00 | 0.00 | 0.60 | 0.47 | 4 | Macrophage |
| VAMP8 | 0.45 | 0.00 | 0.00 | 0.49 | 0.32 | 4 | Macrophage |
| KLF6 | 0.49 | 0.00 | 0.00 | 0.50 | 0.34 | 4 | Macrophage |
| ASAH1 | 0.58 | 0.00 | 0.00 | 0.26 | 0.13 | 4 | Macrophage |
| KLF4 | 0.47 | 0.00 | 0.00 | 0.38 | 0.22 | 4 | Macrophage |
| RPL37A | 0.27 | 0.00 | 0.00 | 0.90 | 0.84 | 4 | Macrophage |
| BOLA2 | 0.60 | 0.00 | 0.00 | 0.26 | 0.14 | 4 | Macrophage |
| SLK | 0.43 | 0.00 | 0.00 | 0.55 | 0.39 | 4 | Macrophage |
| LMO4 | 0.60 | 0.00 | 0.00 | 0.39 | 0.25 | 4 | Macrophage |
| TAOK3 | 0.54 | 0.00 | 0.00 | 0.31 | 0.17 | 4 | Macrophage |
| CELF2 | 0.49 | 0.00 | 0.00 | 0.41 | 0.26 | 4 | Macrophage |
| CLIC1 | 0.46 | 0.00 | 0.00 | 0.52 | 0.38 | 4 | Macrophage |
| TAPBP | 0.57 | 0.00 | 0.00 | 0.27 | 0.14 | 4 | Macrophage |
| FAM32A | 0.60 | 0.00 | 0.00 | 0.37 | 0.23 | 4 | Macrophage |
| ARHGAP17 | 0.57 | 0.00 | 0.00 | 0.28 | 0.15 | 4 | Macrophage |
| MGST1 | 0.51 | 0.00 | 0.00 | 0.31 | 0.17 | 4 | Macrophage |
| KLF13 | 0.54 | 0.00 | 0.00 | 0.27 | 0.15 | 4 | Macrophage |
| H2AFY | 0.54 | 0.00 | 0.00 | 0.32 | 0.20 | 4 | Macrophage |
| CMIP | 0.51 | 0.00 | 0.00 | 0.30 | 0.18 | 4 | Macrophage |
| ARID4A | 0.54 | 0.00 | 0.00 | 0.39 | 0.26 | 4 | Macrophage |
| PAK2 | 0.48 | 0.00 | 0.00 | 0.40 | 0.27 | 4 | Macrophage |
| LAMP1 | 0.49 | 0.00 | 0.00 | 0.53 | 0.42 | 4 | Macrophage |
| SUB1 | 0.37 | 0.00 | 0.00 | 0.62 | 0.51 | 4 | Macrophage |
| BTG2 | 0.63 | 0.00 | 0.00 | 0.33 | 0.21 | 4 | Macrophage |
| COX6B1 | 0.37 | 0.00 | 0.00 | 0.66 | 0.58 | 4 | Macrophage |
| LAMTOR2 | 0.52 | 0.00 | 0.00 | 0.26 | 0.15 | 4 | Macrophage |
| CMPK1 | 0.54 | 0.00 | 0.00 | 0.32 | 0.21 | 4 | Macrophage |
| PRPF40A | 0.49 | 0.00 | 0.00 | 0.42 | 0.30 | 4 | Macrophage |
| ZNHIT1 | 0.50 | 0.00 | 0.00 | 0.25 | 0.15 | 4 | Macrophage |
| CCDC12 | 0.50 | 0.00 | 0.00 | 0.29 | 0.18 | 4 | Macrophage |
| COX7B | 0.42 | 0.00 | 0.00 | 0.56 | 0.47 | 4 | Macrophage |
| FLNA | 0.45 | 0.00 | 0.00 | 0.36 | 0.24 | 4 | Macrophage |
| NME1 | 0.49 | 0.00 | 0.00 | 0.29 | 0.18 | 4 | Macrophage |
| YWHAZ | 0.41 | 0.00 | 0.00 | 0.59 | 0.49 | 4 | Macrophage |
| TSC22D4 | 0.48 | 0.00 | 0.00 | 0.36 | 0.25 | 4 | Macrophage |
| PTP4A2 | 0.40 | 0.00 | 0.00 | 0.52 | 0.40 | 4 | Macrophage |
| ALDOA | 0.51 | 0.00 | 0.00 | 0.33 | 0.22 | 4 | Macrophage |
| SLC25A5 | 0.34 | 0.00 | 0.00 | 0.67 | 0.58 | 4 | Macrophage |
| IRF2 | 0.50 | 0.00 | 0.00 | 0.33 | 0.22 | 4 | Macrophage |
| ATP6V0E1 | 0.47 | 0.00 | 0.00 | 0.33 | 0.22 | 4 | Macrophage |
| RAP1A | 0.47 | 0.00 | 0.00 | 0.41 | 0.29 | 4 | Macrophage |
| S100A10 | 0.41 | 0.00 | 0.00 | 0.51 | 0.40 | 4 | Macrophage |
| EIF3A | 0.39 | 0.00 | 0.00 | 0.66 | 0.58 | 4 | Macrophage |
| COX17 | 0.52 | 0.00 | 0.00 | 0.35 | 0.24 | 4 | Macrophage |
| CHMP4B | 0.52 | 0.00 | 0.00 | 0.30 | 0.20 | 4 | Macrophage |
| DAZAP2 | 0.44 | 0.00 | 0.00 | 0.32 | 0.21 | 4 | Macrophage |
| ZBTB7A | 0.46 | 0.00 | 0.00 | 0.28 | 0.18 | 4 | Macrophage |
| H2AFZ | 0.43 | 0.00 | 0.00 | 0.50 | 0.40 | 4 | Macrophage |
| RBM25 | 0.35 | 0.00 | 0.00 | 0.73 | 0.66 | 4 | Macrophage |
| NSA2 | 0.45 | 0.00 | 0.00 | 0.40 | 0.29 | 4 | Macrophage |
| RNH1 | 0.50 | 0.00 | 0.00 | 0.26 | 0.17 | 4 | Macrophage |
| H3F3A | 0.30 | 0.00 | 0.00 | 0.69 | 0.62 | 4 | Macrophage |
| TMEM14C | 0.48 | 0.00 | 0.00 | 0.32 | 0.22 | 4 | Macrophage |
| SERF2 | 0.32 | 0.00 | 0.00 | 0.66 | 0.59 | 4 | Macrophage |
| ATG3 | 0.43 | 0.00 | 0.00 | 0.27 | 0.17 | 4 | Macrophage |
| CDC42SE1 | 0.46 | 0.00 | 0.00 | 0.27 | 0.18 | 4 | Macrophage |
| EIF5B | 0.37 | 0.00 | 0.00 | 0.58 | 0.50 | 4 | Macrophage |
| SDCBP | 0.48 | 0.00 | 0.00 | 0.38 | 0.28 | 4 | Macrophage |
| TMEM256-PLSCR3 | 0.41 | 0.00 | 0.00 | 0.40 | 0.30 | 4 | Macrophage |
| ETFB | 0.47 | 0.00 | 0.00 | 0.37 | 0.27 | 4 | Macrophage |
| EIF5A | 0.39 | 0.00 | 0.00 | 0.42 | 0.33 | 4 | Macrophage |
| ATP6V1F | 0.41 | 0.00 | 0.00 | 0.34 | 0.25 | 4 | Macrophage |
| VASP | 0.42 | 0.00 | 0.00 | 0.28 | 0.19 | 4 | Macrophage |
| RNF103-CHMP3 | 0.45 | 0.00 | 0.00 | 0.27 | 0.18 | 4 | Macrophage |
| DDX21 | 0.40 | 0.00 | 0.00 | 0.34 | 0.25 | 4 | Macrophage |
| GDI2 | 0.39 | 0.00 | 0.00 | 0.41 | 0.33 | 4 | Macrophage |
| DEK | 0.39 | 0.00 | 0.00 | 0.54 | 0.47 | 4 | Macrophage |
| PRDX1 | 0.41 | 0.00 | 0.00 | 0.57 | 0.51 | 4 | Macrophage |
| MBNL1 | 0.33 | 0.00 | 0.00 | 0.34 | 0.25 | 4 | Macrophage |
| NFE2L2 | 0.44 | 0.00 | 0.00 | 0.33 | 0.25 | 4 | Macrophage |
| NFKBIA | 0.42 | 0.00 | 0.00 | 0.44 | 0.36 | 4 | Macrophage |
| UBL3 | 0.39 | 0.00 | 0.00 | 0.26 | 0.18 | 4 | Macrophage |
| WDR26 | 0.40 | 0.00 | 0.00 | 0.32 | 0.24 | 4 | Macrophage |
| UQCRQ | 0.35 | 0.00 | 0.00 | 0.44 | 0.36 | 4 | Macrophage |
| RHOA | 0.28 | 0.00 | 0.00 | 0.55 | 0.48 | 4 | Macrophage |
| HNRNPM | 0.33 | 0.00 | 0.00 | 0.55 | 0.48 | 4 | Macrophage |
| UTP3 | 0.39 | 0.00 | 0.00 | 0.27 | 0.19 | 4 | Macrophage |
| SH3GLB1 | 0.31 | 0.00 | 0.00 | 0.46 | 0.38 | 4 | Macrophage |
| CHMP2A | 0.35 | 0.00 | 0.00 | 0.40 | 0.32 | 4 | Macrophage |
| MTPN | 0.38 | 0.00 | 0.00 | 0.33 | 0.25 | 4 | Macrophage |
| RRBP1 | 0.28 | 0.00 | 0.00 | 0.60 | 0.54 | 4 | Macrophage |
| RBM3 | 0.32 | 0.00 | 0.00 | 0.47 | 0.41 | 4 | Macrophage |
| ARPP19 | 0.38 | 0.00 | 0.00 | 0.36 | 0.29 | 4 | Macrophage |
| SEC61B | 0.30 | 0.00 | 0.00 | 0.54 | 0.48 | 4 | Macrophage |
| MTDH | 0.32 | 0.00 | 0.00 | 0.54 | 0.48 | 4 | Macrophage |
| ATP1B3 | 0.32 | 0.00 | 0.00 | 0.34 | 0.25 | 4 | Macrophage |
| CAST | 0.38 | 0.00 | 0.00 | 0.27 | 0.19 | 4 | Macrophage |
| ATP6V0D1 | 0.35 | 0.00 | 0.00 | 0.25 | 0.18 | 4 | Macrophage |
| DNAJC2 | 0.36 | 0.00 | 0.00 | 0.26 | 0.18 | 4 | Macrophage |
| CCNL1 | 0.32 | 0.00 | 0.00 | 0.39 | 0.31 | 4 | Macrophage |
| ABI1 | 0.35 | 0.00 | 0.00 | 0.28 | 0.20 | 4 | Macrophage |
| CDC5L | 0.39 | 0.00 | 0.00 | 0.28 | 0.20 | 4 | Macrophage |
| ADIPOR1 | 0.36 | 0.00 | 0.00 | 0.29 | 0.22 | 4 | Macrophage |
| BRK1 | 0.31 | 0.00 | 0.00 | 0.36 | 0.29 | 4 | Macrophage |
| COX8A | 0.27 | 0.00 | 0.00 | 0.60 | 0.56 | 4 | Macrophage |
| GAPDH | 0.43 | 0.00 | 0.00 | 0.26 | 0.19 | 4 | Macrophage |
| ATP6V1G1 | 0.36 | 0.00 | 0.00 | 0.40 | 0.34 | 4 | Macrophage |
| SRP9 | 0.34 | 0.00 | 0.00 | 0.37 | 0.30 | 4 | Macrophage |
| CFLAR | 0.30 | 0.00 | 0.00 | 0.26 | 0.19 | 4 | Macrophage |
| NDUFA1 | 0.30 | 0.00 | 0.00 | 0.39 | 0.33 | 4 | Macrophage |
| NDUFB7 | 0.33 | 0.00 | 0.00 | 0.36 | 0.30 | 4 | Macrophage |
| ROCK1 | 0.30 | 0.00 | 0.00 | 0.40 | 0.33 | 4 | Macrophage |
| TMA7 | 0.26 | 0.00 | 0.00 | 0.41 | 0.35 | 4 | Macrophage |
| NDUFA3 | 0.27 | 0.00 | 0.00 | 0.47 | 0.42 | 4 | Macrophage |
| EIF2S2 | 0.29 | 0.00 | 0.00 | 0.51 | 0.47 | 4 | Macrophage |
| TOMM22 | 0.27 | 0.00 | 0.00 | 0.32 | 0.26 | 4 | Macrophage |
| PSMA4 | 0.28 | 0.00 | 0.00 | 0.39 | 0.33 | 4 | Macrophage |
| FKBP2 | 0.29 | 0.00 | 0.00 | 0.25 | 0.19 | 4 | Macrophage |
| PPIG | 0.27 | 0.00 | 0.00 | 0.44 | 0.39 | 4 | Macrophage |
| NDUFB3 | 0.27 | 0.00 | 0.00 | 0.36 | 0.31 | 4 | Macrophage |
| SNRPB2 | 0.26 | 0.00 | 0.00 | 0.31 | 0.25 | 4 | Macrophage |
| ABCF1 | 0.28 | 0.00 | 0.00 | 0.33 | 0.27 | 4 | Macrophage |
| SMDT1 | 0.27 | 0.00 | 0.00 | 0.34 | 0.29 | 4 | Macrophage |
| NARS | 0.29 | 0.00 | 0.00 | 0.36 | 0.31 | 4 | Macrophage |
| CYCS | 0.29 | 0.00 | 0.00 | 0.27 | 0.22 | 4 | Macrophage |
| CALM3 | 0.29 | 0.00 | 0.00 | 0.25 | 0.20 | 4 | Macrophage |
| PRELID1 | 0.25 | 0.00 | 0.00 | 0.34 | 0.30 | 4 | Macrophage |
| C19orf53 | 0.28 | 0.00 | 0.00 | 0.33 | 0.29 | 4 | Macrophage |
| YWHAH | 0.25 | 0.00 | 0.00 | 0.30 | 0.26 | 4 | Macrophage |
| CWC15 | 0.26 | 0.00 | 0.00 | 0.34 | 0.29 | 4 | Macrophage |
| MIER1 | 0.26 | 0.00 | 0.00 | 0.27 | 0.22 | 4 | Macrophage |
| CCDC50 | 0.26 | 0.00 | 0.00 | 0.30 | 0.25 | 4 | Macrophage |
| ESD | 0.31 | 0.00 | 0.00 | 0.27 | 0.22 | 4 | Macrophage |
| POLR1D | 0.26 | 0.00 | 0.00 | 0.32 | 0.28 | 4 | Macrophage |
| HSPA4 | 0.27 | 0.00 | 0.00 | 0.26 | 0.22 | 4 | Macrophage |
| EIF3J | 0.26 | 0.00 | 0.00 | 0.35 | 0.31 | 4 | Macrophage |
| GGNBP2 | 0.26 | 0.00 | 0.00 | 0.30 | 0.27 | 4 | Macrophage |
| PSMB4 | 0.26 | 0.00 | 0.00 | 0.29 | 0.25 | 4 | Macrophage |
| SSU72 | 0.25 | 0.00 | 0.00 | 0.26 | 0.22 | 4 | Macrophage |
| EDNRB | 2.25 | 0.00 | 0.00 | 0.76 | 0.12 | 5 | Endothelial |
| KDR | 2.19 | 0.00 | 0.00 | 0.75 | 0.12 | 5 | Endothelial |
| KITLG | 2.13 | 0.00 | 0.00 | 0.74 | 0.18 | 5 | Endothelial |
| CYP4B1 | 2.00 | 0.00 | 0.00 | 0.54 | 0.07 | 5 | Endothelial |
| APLN | 1.97 | 0.00 | 0.00 | 0.46 | 0.02 | 5 | Endothelial |
| CLDN5 | 1.92 | 0.00 | 0.00 | 0.89 | 0.20 | 5 | Endothelial |
| SCN7A | 1.77 | 0.00 | 0.00 | 0.80 | 0.26 | 5 | Endothelial |
| CLEC1A | 1.69 | 0.00 | 0.00 | 0.68 | 0.13 | 5 | Endothelial |
| TSPAN13 | 1.64 | 0.00 | 0.00 | 0.73 | 0.21 | 5 | Endothelial |
| ICAM2 | 1.64 | 0.00 | 0.00 | 0.71 | 0.14 | 5 | Endothelial |
| TBX3 | 1.60 | 0.00 | 0.00 | 0.71 | 0.18 | 5 | Endothelial |
| HPGD | 1.56 | 0.00 | 0.00 | 0.82 | 0.23 | 5 | Endothelial |
| RAMP2 | 1.56 | 0.00 | 0.00 | 0.91 | 0.26 | 5 | Endothelial |
| ECSCR | 1.54 | 0.00 | 0.00 | 0.59 | 0.13 | 5 | Endothelial |
| EMCN | 1.52 | 0.00 | 0.00 | 0.50 | 0.09 | 5 | Endothelial |
| PMP22 | 1.51 | 0.00 | 0.00 | 0.67 | 0.32 | 5 | Endothelial |
| CDH5 | 1.51 | 0.00 | 0.00 | 0.84 | 0.20 | 5 | Endothelial |
| EGFL7 | 1.50 | 0.00 | 0.00 | 0.93 | 0.25 | 5 | Endothelial |
| CD34 | 1.40 | 0.00 | 0.00 | 0.65 | 0.18 | 5 | Endothelial |
| SEMA6A | 1.40 | 0.00 | 0.00 | 0.47 | 0.10 | 5 | Endothelial |
| ESAM | 1.35 | 0.00 | 0.00 | 0.68 | 0.17 | 5 | Endothelial |
| ACVRL1 | 1.34 | 0.00 | 0.00 | 0.64 | 0.15 | 5 | Endothelial |
| PECAM1 | 1.34 | 0.00 | 0.00 | 0.81 | 0.22 | 5 | Endothelial |
| IGFBP7 | 1.33 | 0.00 | 0.00 | 0.77 | 0.41 | 5 | Endothelial |
| TSPAN12 | 1.32 | 0.00 | 0.00 | 0.34 | 0.05 | 5 | Endothelial |
| TMEM100 | 1.31 | 0.00 | 0.00 | 0.83 | 0.26 | 5 | Endothelial |
| PCDH17 | 1.31 | 0.00 | 0.00 | 0.69 | 0.18 | 5 | Endothelial |
| EHD4 | 1.29 | 0.00 | 0.00 | 0.68 | 0.24 | 5 | Endothelial |
| TNFSF10 | 1.28 | 0.00 | 0.00 | 0.40 | 0.08 | 5 | Endothelial |
| MYCT1 | 1.28 | 0.00 | 0.00 | 0.49 | 0.11 | 5 | Endothelial |
| SLC9A3R2 | 1.26 | 0.00 | 0.00 | 0.66 | 0.20 | 5 | Endothelial |
| RGS12 | 1.25 | 0.00 | 0.00 | 0.44 | 0.10 | 5 | Endothelial |
| NHLRC2 | 1.24 | 0.00 | 0.00 | 0.33 | 0.06 | 5 | Endothelial |
| S1PR1 | 1.24 | 0.00 | 0.00 | 0.56 | 0.16 | 5 | Endothelial |
| NRP1 | 1.22 | 0.00 | 0.00 | 0.74 | 0.37 | 5 | Endothelial |
| STMN2 | 1.21 | 0.00 | 0.00 | 0.50 | 0.12 | 5 | Endothelial |
| FLT1 | 1.20 | 0.00 | 0.00 | 0.54 | 0.13 | 5 | Endothelial |
| TMEM204 | 1.20 | 0.00 | 0.00 | 0.51 | 0.14 | 5 | Endothelial |
| CAV2 | 1.19 | 0.00 | 0.00 | 0.59 | 0.20 | 5 | Endothelial |
| MYZAP | 1.19 | 0.00 | 0.00 | 0.54 | 0.12 | 5 | Endothelial |
| CYTH3 | 1.18 | 0.00 | 0.00 | 0.54 | 0.21 | 5 | Endothelial |
| PCDH1 | 1.16 | 0.00 | 0.00 | 0.31 | 0.06 | 5 | Endothelial |
| FMO1 | 1.16 | 0.00 | 0.00 | 0.39 | 0.09 | 5 | Endothelial |
| FKBP1A | 1.16 | 0.00 | 0.00 | 0.82 | 0.41 | 5 | Endothelial |
| TSPAN7 | 1.14 | 0.00 | 0.00 | 0.65 | 0.19 | 5 | Endothelial |
| FOXF1 | 1.14 | 0.00 | 0.00 | 0.46 | 0.11 | 5 | Endothelial |
| PIEZO2 | 1.13 | 0.00 | 0.00 | 0.37 | 0.10 | 5 | Endothelial |
| AHR | 1.10 | 0.00 | 0.00 | 0.36 | 0.08 | 5 | Endothelial |
| CCND1 | 1.10 | 0.00 | 0.00 | 0.54 | 0.17 | 5 | Endothelial |
| TSPAN18 | 1.10 | 0.00 | 0.00 | 0.43 | 0.12 | 5 | Endothelial |
| AFAP1L1 | 1.07 | 0.00 | 0.00 | 0.52 | 0.15 | 5 | Endothelial |
| KLF4 | 1.07 | 0.00 | 0.00 | 0.60 | 0.21 | 5 | Endothelial |
| EPAS1 | 1.06 | 0.00 | 0.00 | 0.73 | 0.25 | 5 | Endothelial |
| NOSTRIN | 1.05 | 0.00 | 0.00 | 0.47 | 0.13 | 5 | Endothelial |
| RAPGEF5 | 1.04 | 0.00 | 0.00 | 0.39 | 0.09 | 5 | Endothelial |
| KLF7 | 1.03 | 0.00 | 0.00 | 0.48 | 0.16 | 5 | Endothelial |
| RIPPLY3 | 1.03 | 0.00 | 0.00 | 0.40 | 0.10 | 5 | Endothelial |
| CALCRL | 1.02 | 0.00 | 0.00 | 0.73 | 0.23 | 5 | Endothelial |
| STXBP6 | 1.02 | 0.00 | 0.00 | 0.28 | 0.04 | 5 | Endothelial |
| JUN | 1.01 | 0.00 | 0.00 | 0.78 | 0.35 | 5 | Endothelial |
| THBD | 1.01 | 0.00 | 0.00 | 0.59 | 0.23 | 5 | Endothelial |
| ARHGAP29 | 1.01 | 0.00 | 0.00 | 0.57 | 0.21 | 5 | Endothelial |
| TRIB2 | 1.01 | 0.00 | 0.00 | 0.36 | 0.09 | 5 | Endothelial |
| GATA2 | 0.99 | 0.00 | 0.00 | 0.47 | 0.12 | 5 | Endothelial |
| SLCO2A1 | 0.99 | 0.00 | 0.00 | 0.39 | 0.10 | 5 | Endothelial |
| LY6E | 0.98 | 0.00 | 0.00 | 0.85 | 0.40 | 5 | Endothelial |
| ROBO4 | 0.97 | 0.00 | 0.00 | 0.29 | 0.05 | 5 | Endothelial |
| RCSD1 | 0.97 | 0.00 | 0.00 | 0.34 | 0.07 | 5 | Endothelial |
| PODXL | 0.95 | 0.00 | 0.00 | 0.37 | 0.09 | 5 | Endothelial |
| GNG11 | 0.95 | 0.00 | 0.00 | 0.66 | 0.29 | 5 | Endothelial |
| GIMAP4 | 0.93 | 0.00 | 0.00 | 0.53 | 0.16 | 5 | Endothelial |
| ARHGEF3 | 0.93 | 0.00 | 0.00 | 0.29 | 0.06 | 5 | Endothelial |
| GIMAP6 | 0.91 | 0.00 | 0.00 | 0.45 | 0.14 | 5 | Endothelial |
| S100A16 | 0.89 | 0.00 | 0.00 | 0.62 | 0.23 | 5 | Endothelial |
| APP | 0.86 | 0.00 | 0.00 | 0.87 | 0.59 | 5 | Endothelial |
| CLIC5 | 0.82 | 0.00 | 0.00 | 0.57 | 0.20 | 5 | Endothelial |
| MARCKS | 0.97 | 0.00 | 0.00 | 0.82 | 0.48 | 5 | Endothelial |
| C1orf54 | 0.86 | 0.00 | 0.00 | 0.65 | 0.27 | 5 | Endothelial |
| TIE1 | 0.85 | 0.00 | 0.00 | 0.38 | 0.10 | 5 | Endothelial |
| TIMP3 | 0.99 | 0.00 | 0.00 | 0.72 | 0.36 | 5 | Endothelial |
| COL4A2 | 0.86 | 0.00 | 0.00 | 0.77 | 0.43 | 5 | Endothelial |
| RASIP1 | 0.88 | 0.00 | 0.00 | 0.39 | 0.11 | 5 | Endothelial |
| AQP1 | 0.81 | 0.00 | 0.00 | 0.53 | 0.19 | 5 | Endothelial |
| HSPA12B | 0.80 | 0.00 | 0.00 | 0.26 | 0.05 | 5 | Endothelial |
| SCN3B | 0.97 | 0.00 | 0.00 | 0.27 | 0.06 | 5 | Endothelial |
| CD36 | 0.83 | 0.00 | 0.00 | 0.68 | 0.31 | 5 | Endothelial |
| ITM2B | 0.79 | 0.00 | 0.00 | 0.90 | 0.76 | 5 | Endothelial |
| CD200 | 0.99 | 0.00 | 0.00 | 0.59 | 0.25 | 5 | Endothelial |
| SPTBN1 | 0.78 | 0.00 | 0.00 | 0.83 | 0.52 | 5 | Endothelial |
| CCDC85B | 0.95 | 0.00 | 0.00 | 0.40 | 0.13 | 5 | Endothelial |
| ARHGEF12 | 0.93 | 0.00 | 0.00 | 0.53 | 0.21 | 5 | Endothelial |
| BST2 | 1.06 | 0.00 | 0.00 | 0.41 | 0.14 | 5 | Endothelial |
| LYVE1 | 0.93 | 0.00 | 0.00 | 0.51 | 0.19 | 5 | Endothelial |
| SGK1 | 0.88 | 0.00 | 0.00 | 0.47 | 0.17 | 5 | Endothelial |
| BTBD3 | 1.02 | 0.00 | 0.00 | 0.39 | 0.13 | 5 | Endothelial |
| JUP | 0.90 | 0.00 | 0.00 | 0.42 | 0.14 | 5 | Endothelial |
| KIAA0355 | 0.87 | 0.00 | 0.00 | 0.45 | 0.16 | 5 | Endothelial |
| IMPDH1 | 0.92 | 0.00 | 0.00 | 0.27 | 0.07 | 5 | Endothelial |
| COL4A1 | 0.74 | 0.00 | 0.00 | 0.79 | 0.49 | 5 | Endothelial |
| KANK3 | 0.83 | 0.00 | 0.00 | 0.38 | 0.12 | 5 | Endothelial |
| ARHGEF15 | 0.78 | 0.00 | 0.00 | 0.26 | 0.06 | 5 | Endothelial |
| PPP1R16B | 0.84 | 0.00 | 0.00 | 0.29 | 0.08 | 5 | Endothelial |
| CAV1 | 0.83 | 0.00 | 0.00 | 0.68 | 0.35 | 5 | Endothelial |
| TMEM47 | 1.07 | 0.00 | 0.00 | 0.31 | 0.10 | 5 | Endothelial |
| BMPR2 | 0.86 | 0.00 | 0.00 | 0.31 | 0.09 | 5 | Endothelial |
| CYYR1 | 0.71 | 0.00 | 0.00 | 0.39 | 0.13 | 5 | Endothelial |
| CLEC14A | 0.63 | 0.00 | 0.00 | 0.46 | 0.17 | 5 | Endothelial |
| CARHSP1 | 0.87 | 0.00 | 0.00 | 0.38 | 0.14 | 5 | Endothelial |
| TJP1 | 0.82 | 0.00 | 0.00 | 0.42 | 0.16 | 5 | Endothelial |
| USHBP1 | 0.76 | 0.00 | 0.00 | 0.28 | 0.08 | 5 | Endothelial |
| PEA15 | 0.83 | 0.00 | 0.00 | 0.38 | 0.14 | 5 | Endothelial |
| ANXA3 | 0.72 | 0.00 | 0.00 | 0.49 | 0.21 | 5 | Endothelial |
| SLFN5 | 0.77 | 0.00 | 0.00 | 0.30 | 0.09 | 5 | Endothelial |
| CRIP2 | 0.66 | 0.00 | 0.00 | 0.71 | 0.42 | 5 | Endothelial |
| MGLL | 0.87 | 0.00 | 0.00 | 0.41 | 0.16 | 5 | Endothelial |
| TPST2 | 0.79 | 0.00 | 0.00 | 0.32 | 0.11 | 5 | Endothelial |
| HES1 | 0.88 | 0.00 | 0.00 | 0.56 | 0.28 | 5 | Endothelial |
| NAV1 | 0.86 | 0.00 | 0.00 | 0.30 | 0.10 | 5 | Endothelial |
| RHOJ | 0.78 | 0.00 | 0.00 | 0.41 | 0.16 | 5 | Endothelial |
| SEPT4 | 0.91 | 0.00 | 0.00 | 0.47 | 0.21 | 5 | Endothelial |
| SNRK | 0.81 | 0.00 | 0.00 | 0.38 | 0.15 | 5 | Endothelial |
| CNN3 | 0.65 | 0.00 | 0.00 | 0.61 | 0.31 | 5 | Endothelial |
| VAT1 | 0.75 | 0.00 | 0.00 | 0.28 | 0.09 | 5 | Endothelial |
| ETS1 | 0.64 | 0.00 | 0.00 | 0.50 | 0.22 | 5 | Endothelial |
| PLEKHA1 | 0.80 | 0.00 | 0.00 | 0.31 | 0.11 | 5 | Endothelial |
| VIM | 0.49 | 0.00 | 0.00 | 0.90 | 0.68 | 5 | Endothelial |
| LDB2 | 0.63 | 0.00 | 0.00 | 0.41 | 0.16 | 5 | Endothelial |
| SCARB1 | 0.78 | 0.00 | 0.00 | 0.27 | 0.08 | 5 | Endothelial |
| GALNT18 | 0.73 | 0.00 | 0.00 | 0.30 | 0.10 | 5 | Endothelial |
| ADD3 | 0.74 | 0.00 | 0.00 | 0.52 | 0.26 | 5 | Endothelial |
| B2M | 1.07 | 0.00 | 0.00 | 0.88 | 0.69 | 5 | Endothelial |
| KLF2 | 0.73 | 0.00 | 0.00 | 0.43 | 0.18 | 5 | Endothelial |
| RHOC | 0.71 | 0.00 | 0.00 | 0.47 | 0.23 | 5 | Endothelial |
| RAB11A | 0.72 | 0.00 | 0.00 | 0.50 | 0.26 | 5 | Endothelial |
| NES | 0.63 | 0.00 | 0.00 | 0.45 | 0.20 | 5 | Endothelial |
| SMARCA2 | 0.73 | 0.00 | 0.00 | 0.56 | 0.32 | 5 | Endothelial |
| TBX2 | 0.73 | 0.00 | 0.00 | 0.32 | 0.13 | 5 | Endothelial |
| SEMA3C | 0.67 | 0.00 | 0.00 | 0.41 | 0.18 | 5 | Endothelial |
| BCAM | 0.79 | 0.00 | 0.00 | 0.28 | 0.10 | 5 | Endothelial |
| CD151 | 0.69 | 0.00 | 0.00 | 0.40 | 0.18 | 5 | Endothelial |
| ENG | 0.66 | 0.00 | 0.00 | 0.31 | 0.12 | 5 | Endothelial |
| SOX17 | 0.52 | 0.00 | 0.00 | 0.33 | 0.13 | 5 | Endothelial |
| KLHL5 | 0.66 | 0.00 | 0.00 | 0.33 | 0.13 | 5 | Endothelial |
| TCF4 | 0.59 | 0.00 | 0.00 | 0.65 | 0.40 | 5 | Endothelial |
| FERMT2 | 0.61 | 0.00 | 0.00 | 0.61 | 0.38 | 5 | Endothelial |
| ECE1 | 0.62 | 0.00 | 0.00 | 0.47 | 0.24 | 5 | Endothelial |
| RASGRP3 | 0.64 | 0.00 | 0.00 | 0.30 | 0.11 | 5 | Endothelial |
| ACBD5 | 0.76 | 0.00 | 0.00 | 0.31 | 0.12 | 5 | Endothelial |
| DDAH2 | 0.61 | 0.00 | 0.00 | 0.51 | 0.27 | 5 | Endothelial |
| PTPRG | 0.72 | 0.00 | 0.00 | 0.29 | 0.11 | 5 | Endothelial |
| TUBB2A | 0.71 | 0.00 | 0.00 | 0.41 | 0.20 | 5 | Endothelial |
| PDE4B | 0.68 | 0.00 | 0.00 | 0.32 | 0.13 | 5 | Endothelial |
| GRAP | 0.63 | 0.00 | 0.00 | 0.30 | 0.11 | 5 | Endothelial |
| GPIHBP1 | 0.65 | 0.00 | 0.00 | 0.35 | 0.15 | 5 | Endothelial |
| RAP1A | 0.65 | 0.00 | 0.00 | 0.51 | 0.29 | 5 | Endothelial |
| ZEB1 | 0.64 | 0.00 | 0.00 | 0.28 | 0.11 | 5 | Endothelial |
| SYN3 | 0.81 | 0.00 | 0.00 | 0.31 | 0.13 | 5 | Endothelial |
| TM4SF1 | 0.56 | 0.00 | 0.00 | 0.43 | 0.21 | 5 | Endothelial |
| GIMAP1 | 0.56 | 0.00 | 0.00 | 0.31 | 0.13 | 5 | Endothelial |
| LYN | 0.63 | 0.00 | 0.00 | 0.28 | 0.11 | 5 | Endothelial |
| SNX3 | 0.58 | 0.00 | 0.00 | 0.56 | 0.34 | 5 | Endothelial |
| RELL1 | 0.72 | 0.00 | 0.00 | 0.36 | 0.17 | 5 | Endothelial |
| CMTM3 | 0.64 | 0.00 | 0.00 | 0.32 | 0.14 | 5 | Endothelial |
| CTNND1 | 0.64 | 0.00 | 0.00 | 0.34 | 0.16 | 5 | Endothelial |
| QKI | 0.60 | 0.00 | 0.00 | 0.49 | 0.28 | 5 | Endothelial |
| TUBA1A | 0.47 | 0.00 | 0.00 | 0.73 | 0.51 | 5 | Endothelial |
| ETS2 | 0.60 | 0.00 | 0.00 | 0.29 | 0.12 | 5 | Endothelial |
| MGST3 | 0.66 | 0.00 | 0.00 | 0.29 | 0.13 | 5 | Endothelial |
| CLIC4 | 0.64 | 0.00 | 0.00 | 0.49 | 0.29 | 5 | Endothelial |
| GNAI2 | 0.53 | 0.00 | 0.00 | 0.59 | 0.37 | 5 | Endothelial |
| GUK1 | 0.72 | 0.00 | 0.00 | 0.37 | 0.18 | 5 | Endothelial |
| SRGN | 0.34 | 0.00 | 0.00 | 0.56 | 0.31 | 5 | Endothelial |
| MEF2A | 0.59 | 0.00 | 0.00 | 0.53 | 0.33 | 5 | Endothelial |
| ACAP2 | 0.64 | 0.00 | 0.00 | 0.35 | 0.18 | 5 | Endothelial |
| CDKN1A | 1.00 | 0.00 | 0.00 | 0.28 | 0.13 | 5 | Endothelial |
| GCC2 | 0.62 | 0.00 | 0.00 | 0.42 | 0.23 | 5 | Endothelial |
| SASH1 | 0.58 | 0.00 | 0.00 | 0.28 | 0.12 | 5 | Endothelial |
| MXRA7 | 0.61 | 0.00 | 0.00 | 0.29 | 0.14 | 5 | Endothelial |
| CD93 | 0.42 | 0.00 | 0.00 | 0.42 | 0.21 | 5 | Endothelial |
| IER2 | 0.55 | 0.00 | 0.00 | 0.44 | 0.25 | 5 | Endothelial |
| LAPTM4A | 0.46 | 0.00 | 0.00 | 0.70 | 0.54 | 5 | Endothelial |
| ELK3 | 0.53 | 0.00 | 0.00 | 0.29 | 0.13 | 5 | Endothelial |
| SERPINH1 | 0.42 | 0.00 | 0.00 | 0.61 | 0.41 | 5 | Endothelial |
| DYNLL1 | 0.48 | 0.00 | 0.00 | 0.63 | 0.46 | 5 | Endothelial |
| VAMP5 | 0.76 | 0.00 | 0.00 | 0.26 | 0.12 | 5 | Endothelial |
| DLC1 | 0.55 | 0.00 | 0.00 | 0.34 | 0.17 | 5 | Endothelial |
| HIP1 | 0.54 | 0.00 | 0.00 | 0.29 | 0.14 | 5 | Endothelial |
| MAP7D1 | 0.58 | 0.00 | 0.00 | 0.33 | 0.17 | 5 | Endothelial |
| SERINC3 | 0.56 | 0.00 | 0.00 | 0.47 | 0.29 | 5 | Endothelial |
| CLTB | 0.57 | 0.00 | 0.00 | 0.26 | 0.12 | 5 | Endothelial |
| ABI1 | 0.57 | 0.00 | 0.00 | 0.36 | 0.20 | 5 | Endothelial |
| MYO6 | 0.49 | 0.00 | 0.00 | 0.27 | 0.13 | 5 | Endothelial |
| ITGB1 | 0.35 | 0.00 | 0.00 | 0.80 | 0.63 | 5 | Endothelial |
| MCAM | 0.48 | 0.00 | 0.00 | 0.27 | 0.13 | 5 | Endothelial |
| SLC43A3 | 0.55 | 0.00 | 0.00 | 0.34 | 0.18 | 5 | Endothelial |
| NID1 | 0.55 | 0.00 | 0.00 | 0.39 | 0.22 | 5 | Endothelial |
| NOTCH1 | 0.48 | 0.00 | 0.00 | 0.25 | 0.12 | 5 | Endothelial |
| SKIL | 0.56 | 0.00 | 0.00 | 0.27 | 0.14 | 5 | Endothelial |
| WASF2 | 0.49 | 0.00 | 0.00 | 0.47 | 0.30 | 5 | Endothelial |
| DUSP6 | 0.45 | 0.00 | 0.00 | 0.33 | 0.18 | 5 | Endothelial |
| REEP3 | 0.52 | 0.00 | 0.00 | 0.33 | 0.18 | 5 | Endothelial |
| PLCG1 | 0.54 | 0.00 | 0.00 | 0.29 | 0.15 | 5 | Endothelial |
| CLIC1 | 0.43 | 0.00 | 0.00 | 0.55 | 0.38 | 5 | Endothelial |
| CYB5R3 | 0.52 | 0.00 | 0.00 | 0.54 | 0.39 | 5 | Endothelial |
| MYL12B | 0.35 | 0.00 | 0.00 | 0.73 | 0.62 | 5 | Endothelial |
| SPTAN1 | 0.46 | 0.00 | 0.00 | 0.30 | 0.16 | 5 | Endothelial |
| EVA1B | 0.49 | 0.00 | 0.00 | 0.34 | 0.19 | 5 | Endothelial |
| LUZP1 | 0.47 | 0.00 | 0.00 | 0.34 | 0.19 | 5 | Endothelial |
| HDAC7 | 0.49 | 0.00 | 0.00 | 0.26 | 0.13 | 5 | Endothelial |
| DNAJC8 | 0.42 | 0.00 | 0.00 | 0.57 | 0.42 | 5 | Endothelial |
| SH3BP5 | 0.44 | 0.00 | 0.00 | 0.34 | 0.19 | 5 | Endothelial |
| GMFG | 0.33 | 0.00 | 0.00 | 0.28 | 0.14 | 5 | Endothelial |
| CTNNA1 | 0.40 | 0.00 | 0.00 | 0.52 | 0.35 | 5 | Endothelial |
| EFNB1 | 0.43 | 0.00 | 0.00 | 0.27 | 0.14 | 5 | Endothelial |
| EFNB2 | 0.38 | 0.00 | 0.00 | 0.27 | 0.14 | 5 | Endothelial |
| PON2 | 0.44 | 0.00 | 0.00 | 0.27 | 0.14 | 5 | Endothelial |
| PTTG1IP | 0.48 | 0.00 | 0.00 | 0.30 | 0.17 | 5 | Endothelial |
| MSN | 0.31 | 0.00 | 0.00 | 0.60 | 0.43 | 5 | Endothelial |
| MTUS1 | 0.48 | 0.00 | 0.00 | 0.28 | 0.15 | 5 | Endothelial |
| PLK2 | 0.45 | 0.00 | 0.00 | 0.29 | 0.16 | 5 | Endothelial |
| RHOA | 0.36 | 0.00 | 0.00 | 0.61 | 0.48 | 5 | Endothelial |
| SYPL1 | 0.42 | 0.00 | 0.00 | 0.40 | 0.26 | 5 | Endothelial |
| LEPROTL1 | 0.42 | 0.00 | 0.00 | 0.27 | 0.15 | 5 | Endothelial |
| PPP2R5A | 0.46 | 0.00 | 0.00 | 0.25 | 0.14 | 5 | Endothelial |
| SERINC1 | 0.47 | 0.00 | 0.00 | 0.42 | 0.28 | 5 | Endothelial |
| DAAM1 | 0.44 | 0.00 | 0.00 | 0.28 | 0.16 | 5 | Endothelial |
| FNBP1L | 0.46 | 0.00 | 0.00 | 0.37 | 0.24 | 5 | Endothelial |
| ITPRIPL2 | 0.46 | 0.00 | 0.00 | 0.28 | 0.16 | 5 | Endothelial |
| BNIP2 | 0.40 | 0.00 | 0.00 | 0.42 | 0.28 | 5 | Endothelial |
| CD47 | 0.44 | 0.00 | 0.00 | 0.53 | 0.41 | 5 | Endothelial |
| TJP2 | 0.37 | 0.00 | 0.00 | 0.27 | 0.15 | 5 | Endothelial |
| MOCS2 | 0.48 | 0.00 | 0.00 | 0.30 | 0.18 | 5 | Endothelial |
| PPP1R2 | 0.43 | 0.00 | 0.00 | 0.36 | 0.24 | 5 | Endothelial |
| NFIC | 0.45 | 0.00 | 0.00 | 0.37 | 0.24 | 5 | Endothelial |
| MSI2 | 0.46 | 0.00 | 0.00 | 0.26 | 0.15 | 5 | Endothelial |
| KLC1 | 0.40 | 0.00 | 0.00 | 0.25 | 0.14 | 5 | Endothelial |
| HPCAL1 | 0.39 | 0.00 | 0.00 | 0.26 | 0.15 | 5 | Endothelial |
| ZFHX3 | 0.49 | 0.00 | 0.00 | 0.28 | 0.17 | 5 | Endothelial |
| VAMP3 | 0.39 | 0.00 | 0.00 | 0.25 | 0.15 | 5 | Endothelial |
| RBMS1 | 0.40 | 0.00 | 0.00 | 0.50 | 0.38 | 5 | Endothelial |
| RALA | 0.41 | 0.00 | 0.00 | 0.36 | 0.24 | 5 | Endothelial |
| DUSP1 | 0.34 | 0.00 | 0.00 | 0.28 | 0.17 | 5 | Endothelial |
| NFKBIA | 0.35 | 0.00 | 0.00 | 0.49 | 0.36 | 5 | Endothelial |
| DPYSL2 | 0.39 | 0.00 | 0.00 | 0.29 | 0.18 | 5 | Endothelial |
| ZMIZ1 | 0.38 | 0.00 | 0.00 | 0.31 | 0.20 | 5 | Endothelial |
| S100A13 | 0.42 | 0.00 | 0.00 | 0.38 | 0.27 | 5 | Endothelial |
| RPN1 | 0.36 | 0.00 | 0.00 | 0.32 | 0.20 | 5 | Endothelial |
| ARF1 | 0.31 | 0.00 | 0.00 | 0.55 | 0.44 | 5 | Endothelial |
| VPS36 | 0.40 | 0.00 | 0.00 | 0.38 | 0.26 | 5 | Endothelial |
| MXD4 | 0.41 | 0.00 | 0.00 | 0.46 | 0.35 | 5 | Endothelial |
| RAP1B | 0.33 | 0.00 | 0.00 | 0.35 | 0.24 | 5 | Endothelial |
| SDCBP | 0.32 | 0.00 | 0.00 | 0.41 | 0.28 | 5 | Endothelial |
| CCND3 | 0.38 | 0.00 | 0.00 | 0.28 | 0.18 | 5 | Endothelial |
| ARPC3 | 0.32 | 0.00 | 0.00 | 0.40 | 0.28 | 5 | Endothelial |
| FCHO2 | 0.41 | 0.00 | 0.00 | 0.27 | 0.17 | 5 | Endothelial |
| ITSN2 | 0.37 | 0.00 | 0.00 | 0.34 | 0.23 | 5 | Endothelial |
| APLP2 | 0.31 | 0.00 | 0.00 | 0.51 | 0.40 | 5 | Endothelial |
| RTN4 | 0.42 | 0.00 | 0.00 | 0.36 | 0.26 | 5 | Endothelial |
| ZCRB1 | 0.35 | 0.00 | 0.00 | 0.38 | 0.27 | 5 | Endothelial |
| TMOD3 | 0.36 | 0.00 | 0.00 | 0.36 | 0.26 | 5 | Endothelial |
| LEPROT | 0.33 | 0.00 | 0.00 | 0.44 | 0.33 | 5 | Endothelial |
| TXNIP | 0.32 | 0.00 | 0.00 | 0.37 | 0.26 | 5 | Endothelial |
| CHMP5 | 0.36 | 0.00 | 0.00 | 0.35 | 0.25 | 5 | Endothelial |
| GNB2 | 0.34 | 0.00 | 0.00 | 0.47 | 0.37 | 5 | Endothelial |
| LAMP1 | 0.27 | 0.00 | 0.00 | 0.53 | 0.43 | 5 | Endothelial |
| C5orf24 | 0.33 | 0.00 | 0.00 | 0.36 | 0.26 | 5 | Endothelial |
| TMEM50A | 0.27 | 0.00 | 0.00 | 0.54 | 0.44 | 5 | Endothelial |
| PPP1CA | 0.29 | 0.00 | 0.00 | 0.44 | 0.33 | 5 | Endothelial |
| TMEM176A | 0.32 | 0.00 | 0.00 | 0.28 | 0.19 | 5 | Endothelial |
| SBDS | 0.30 | 0.00 | 0.00 | 0.33 | 0.24 | 5 | Endothelial |
| SRPRA | 0.37 | 0.00 | 0.00 | 0.31 | 0.23 | 5 | Endothelial |
| GABARAPL2 | 0.29 | 0.00 | 0.00 | 0.33 | 0.25 | 5 | Endothelial |
| HOOK3 | 0.33 | 0.00 | 0.00 | 0.30 | 0.22 | 5 | Endothelial |
| ROCK2 | 0.32 | 0.00 | 0.00 | 0.38 | 0.29 | 5 | Endothelial |
| RAB6A | 0.27 | 0.00 | 0.00 | 0.29 | 0.21 | 5 | Endothelial |
| MARCKSL1 | 0.30 | 0.00 | 0.00 | 0.31 | 0.23 | 5 | Endothelial |
| RAB7A | 0.26 | 0.00 | 0.00 | 0.35 | 0.27 | 5 | Endothelial |
| RAB2A | 0.26 | 0.00 | 0.00 | 0.36 | 0.28 | 5 | Endothelial |
| IRF2 | 0.26 | 0.00 | 0.00 | 0.30 | 0.22 | 5 | Endothelial |
| TMCO1 | 0.25 | 0.00 | 0.00 | 0.37 | 0.29 | 5 | Endothelial |
| RAB10 | 0.26 | 0.00 | 0.00 | 0.31 | 0.24 | 5 | Endothelial |
| GNB1 | 0.26 | 0.00 | 0.00 | 0.28 | 0.22 | 5 | Endothelial |
| XIAP | 0.26 | 0.00 | 0.00 | 0.26 | 0.19 | 5 | Endothelial |
| KIF1B | 0.29 | 0.00 | 0.00 | 0.27 | 0.21 | 5 | Endothelial |
| PDCD10 | 0.26 | 0.00 | 0.00 | 0.26 | 0.20 | 5 | Endothelial |
| IGFBP4 | 0.26 | 0.00 | 0.00 | 0.34 | 0.27 | 5 | Endothelial |
| AKAP5 | 3.64 | 0.00 | 0.00 | 0.94 | 0.04 | 6 | AT1 Cell |
| AGER | 3.45 | 0.00 | 0.00 | 0.99 | 0.13 | 6 | AT1 Cell |
| CLDN18 | 2.99 | 0.00 | 0.00 | 0.98 | 0.13 | 6 | AT1 Cell |
| RTKN2 | 2.88 | 0.00 | 0.00 | 0.77 | 0.02 | 6 | AT1 Cell |
| SEC14L3 | 2.71 | 0.00 | 0.00 | 0.86 | 0.05 | 6 | AT1 Cell |
| LMO7 | 2.65 | 0.00 | 0.00 | 0.89 | 0.08 | 6 | AT1 Cell |
| KRT7 | 2.53 | 0.00 | 0.00 | 0.79 | 0.03 | 6 | AT1 Cell |
| AQP5 | 2.52 | 0.00 | 0.00 | 0.72 | 0.03 | 6 | AT1 Cell |
| COL4A3 | 2.42 | 0.00 | 0.00 | 0.66 | 0.01 | 6 | AT1 Cell |
| SPOCK2 | 2.41 | 0.00 | 0.00 | 0.67 | 0.02 | 6 | AT1 Cell |
| GPRC5A | 2.36 | 0.00 | 0.00 | 0.74 | 0.03 | 6 | AT1 Cell |
| VEGFA | 2.33 | 0.00 | 0.00 | 0.84 | 0.12 | 6 | AT1 Cell |
| CLIC3 | 2.28 | 0.00 | 0.00 | 0.74 | 0.05 | 6 | AT1 Cell |
| MSLN | 2.26 | 0.00 | 0.00 | 0.61 | 0.01 | 6 | AT1 Cell |
| MYH14 | 2.23 | 0.00 | 0.00 | 0.74 | 0.05 | 6 | AT1 Cell |
| CLIC5 | 2.21 | 0.00 | 0.00 | 0.92 | 0.19 | 6 | AT1 Cell |
| COL4A4 | 2.19 | 0.00 | 0.00 | 0.58 | 0.01 | 6 | AT1 Cell |
| HOPX | 2.19 | 0.00 | 0.00 | 0.74 | 0.07 | 6 | AT1 Cell |
| C19orf33 | 2.17 | 0.00 | 0.00 | 0.67 | 0.04 | 6 | AT1 Cell |
| KRT19 | 2.11 | 0.00 | 0.00 | 0.60 | 0.02 | 6 | AT1 Cell |
| SCNN1G | 2.08 | 0.00 | 0.00 | 0.57 | 0.01 | 6 | AT1 Cell |
| S100A14 | 2.06 | 0.00 | 0.00 | 0.52 | 0.01 | 6 | AT1 Cell |
| SCNN1A | 2.04 | 0.00 | 0.00 | 0.63 | 0.03 | 6 | AT1 Cell |
| PDGFA | 2.03 | 0.00 | 0.00 | 0.64 | 0.07 | 6 | AT1 Cell |
| SEMA3E | 2.01 | 0.00 | 0.00 | 0.49 | 0.01 | 6 | AT1 Cell |
| TINAGL1 | 2.01 | 0.00 | 0.00 | 0.67 | 0.10 | 6 | AT1 Cell |
| PDPN | 1.98 | 0.00 | 0.00 | 0.55 | 0.03 | 6 | AT1 Cell |
| CDKN2B | 1.96 | 0.00 | 0.00 | 0.56 | 0.02 | 6 | AT1 Cell |
| NDNF | 1.91 | 0.00 | 0.00 | 0.49 | 0.03 | 6 | AT1 Cell |
| KRT8 | 1.88 | 0.00 | 0.00 | 0.57 | 0.04 | 6 | AT1 Cell |
| S100A6 | 1.86 | 0.00 | 0.00 | 0.93 | 0.32 | 6 | AT1 Cell |
| TIMP3 | 1.86 | 0.00 | 0.00 | 0.91 | 0.35 | 6 | AT1 Cell |
| SLC39A8 | 1.84 | 0.00 | 0.00 | 0.57 | 0.05 | 6 | AT1 Cell |
| SEMA3A | 1.84 | 0.00 | 0.00 | 0.45 | 0.01 | 6 | AT1 Cell |
| TSPAN8 | 1.81 | 0.00 | 0.00 | 0.68 | 0.09 | 6 | AT1 Cell |
| PRDX6 | 1.81 | 0.00 | 0.00 | 0.83 | 0.26 | 6 | AT1 Cell |
| GIPC2 | 1.77 | 0.00 | 0.00 | 0.43 | 0.01 | 6 | AT1 Cell |
| ICAM1 | 1.73 | 0.00 | 0.00 | 0.63 | 0.07 | 6 | AT1 Cell |
| ANO1 | 1.73 | 0.00 | 0.00 | 0.48 | 0.03 | 6 | AT1 Cell |
| TMEM37 | 1.69 | 0.00 | 0.00 | 0.45 | 0.03 | 6 | AT1 Cell |
| ATP1B1 | 1.67 | 0.00 | 0.00 | 0.70 | 0.16 | 6 | AT1 Cell |
| ITGB6 | 1.66 | 0.00 | 0.00 | 0.35 | 0.01 | 6 | AT1 Cell |
| ANXA1 | 1.65 | 0.00 | 0.00 | 0.71 | 0.17 | 6 | AT1 Cell |
| FAM174B | 1.64 | 0.00 | 0.00 | 0.43 | 0.03 | 6 | AT1 Cell |
| F3 | 1.64 | 0.00 | 0.00 | 0.41 | 0.02 | 6 | AT1 Cell |
| RAB11FIP1 | 1.63 | 0.00 | 0.00 | 0.48 | 0.03 | 6 | AT1 Cell |
| FLRT3 | 1.63 | 0.00 | 0.00 | 0.39 | 0.02 | 6 | AT1 Cell |
| NDST1 | 1.62 | 0.00 | 0.00 | 0.52 | 0.09 | 6 | AT1 Cell |
| KRT18 | 1.62 | 0.00 | 0.00 | 0.53 | 0.05 | 6 | AT1 Cell |
| RGCC | 1.60 | 0.00 | 0.00 | 0.65 | 0.17 | 6 | AT1 Cell |
| ALCAM | 1.60 | 0.00 | 0.00 | 0.84 | 0.18 | 6 | AT1 Cell |
| MTHFD1 | 1.60 | 0.00 | 0.00 | 0.51 | 0.06 | 6 | AT1 Cell |
| MAL2 | 1.59 | 0.00 | 0.00 | 0.48 | 0.03 | 6 | AT1 Cell |
| CLDN3 | 1.58 | 0.00 | 0.00 | 0.58 | 0.08 | 6 | AT1 Cell |
| PTPRF | 1.57 | 0.00 | 0.00 | 0.70 | 0.12 | 6 | AT1 Cell |
| ANXA3 | 1.56 | 0.00 | 0.00 | 0.76 | 0.19 | 6 | AT1 Cell |
| SYN3 | 1.56 | 0.00 | 0.00 | 0.59 | 0.12 | 6 | AT1 Cell |
| CYSTM1 | 1.55 | 0.00 | 0.00 | 0.53 | 0.07 | 6 | AT1 Cell |
| AGRN | 1.54 | 0.00 | 0.00 | 0.60 | 0.10 | 6 | AT1 Cell |
| CRYAB | 1.52 | 0.00 | 0.00 | 0.65 | 0.13 | 6 | AT1 Cell |
| AHNAK | 1.52 | 0.00 | 0.00 | 0.90 | 0.41 | 6 | AT1 Cell |
| SCD | 1.47 | 0.00 | 0.00 | 0.68 | 0.18 | 6 | AT1 Cell |
| LAMC2 | 1.46 | 0.00 | 0.00 | 0.40 | 0.02 | 6 | AT1 Cell |
| CADM1 | 1.46 | 0.00 | 0.00 | 0.58 | 0.12 | 6 | AT1 Cell |
| NKX2-1 | 1.45 | 0.00 | 0.00 | 0.55 | 0.07 | 6 | AT1 Cell |
| EZR | 1.45 | 0.00 | 0.00 | 0.70 | 0.18 | 6 | AT1 Cell |
| SELENBP1 | 1.44 | 0.00 | 0.00 | 0.55 | 0.19 | 6 | AT1 Cell |
| DAG1 | 1.43 | 0.00 | 0.00 | 0.45 | 0.07 | 6 | AT1 Cell |
| CTSH | 1.43 | 0.00 | 0.00 | 0.73 | 0.18 | 6 | AT1 Cell |
| IL18R1 | 1.41 | 0.00 | 0.00 | 0.44 | 0.04 | 6 | AT1 Cell |
| EPCAM | 1.40 | 0.00 | 0.00 | 0.52 | 0.07 | 6 | AT1 Cell |
| TSPAN15 | 1.38 | 0.00 | 0.00 | 0.46 | 0.06 | 6 | AT1 Cell |
| CLDN7 | 1.37 | 0.00 | 0.00 | 0.44 | 0.05 | 6 | AT1 Cell |
| WWC1 | 1.36 | 0.00 | 0.00 | 0.42 | 0.03 | 6 | AT1 Cell |
| TP53BP2 | 1.36 | 0.00 | 0.00 | 0.42 | 0.06 | 6 | AT1 Cell |
| SCNN1B | 1.35 | 0.00 | 0.00 | 0.33 | 0.01 | 6 | AT1 Cell |
| CXADR | 1.34 | 0.00 | 0.00 | 0.38 | 0.04 | 6 | AT1 Cell |
| LLGL2 | 1.34 | 0.00 | 0.00 | 0.36 | 0.02 | 6 | AT1 Cell |
| WWP1 | 1.33 | 0.00 | 0.00 | 0.53 | 0.10 | 6 | AT1 Cell |
| EPS8L2 | 1.32 | 0.00 | 0.00 | 0.34 | 0.01 | 6 | AT1 Cell |
| CTGF | 1.29 | 0.00 | 0.00 | 0.44 | 0.08 | 6 | AT1 Cell |
| TEAD1 | 1.28 | 0.00 | 0.00 | 0.50 | 0.11 | 6 | AT1 Cell |
| TACSTD2 | 1.28 | 0.00 | 0.00 | 0.28 | 0.01 | 6 | AT1 Cell |
| FAM189A2 | 1.28 | 0.00 | 0.00 | 0.35 | 0.02 | 6 | AT1 Cell |
| MMP11 | 1.28 | 0.00 | 0.00 | 0.34 | 0.03 | 6 | AT1 Cell |
| PARD6B | 1.27 | 0.00 | 0.00 | 0.31 | 0.01 | 6 | AT1 Cell |
| SPINT2 | 1.26 | 0.00 | 0.00 | 0.51 | 0.09 | 6 | AT1 Cell |
| CD9 | 1.26 | 0.00 | 0.00 | 0.85 | 0.33 | 6 | AT1 Cell |
| OSBPL6 | 1.25 | 0.00 | 0.00 | 0.31 | 0.03 | 6 | AT1 Cell |
| MPP5 | 1.25 | 0.00 | 0.00 | 0.46 | 0.09 | 6 | AT1 Cell |
| PPP3CA | 1.24 | 0.00 | 0.00 | 0.70 | 0.24 | 6 | AT1 Cell |
| PKP2 | 1.24 | 0.00 | 0.00 | 0.32 | 0.02 | 6 | AT1 Cell |
| NAV2 | 1.24 | 0.00 | 0.00 | 0.39 | 0.06 | 6 | AT1 Cell |
| DENND3 | 1.23 | 0.00 | 0.00 | 0.40 | 0.05 | 6 | AT1 Cell |
| SH3D21 | 1.20 | 0.00 | 0.00 | 0.28 | 0.02 | 6 | AT1 Cell |
| PDE7A | 1.17 | 0.00 | 0.00 | 0.37 | 0.05 | 6 | AT1 Cell |
| WDR45 | 1.17 | 0.00 | 0.00 | 0.41 | 0.07 | 6 | AT1 Cell |
| GJA1 | 1.15 | 0.00 | 0.00 | 0.45 | 0.07 | 6 | AT1 Cell |
| MAGI3 | 1.15 | 0.00 | 0.00 | 0.41 | 0.09 | 6 | AT1 Cell |
| HS2ST1 | 1.15 | 0.00 | 0.00 | 0.31 | 0.04 | 6 | AT1 Cell |
| SLC1A5 | 1.15 | 0.00 | 0.00 | 0.43 | 0.09 | 6 | AT1 Cell |
| SLCO3A1 | 1.12 | 0.00 | 0.00 | 0.36 | 0.06 | 6 | AT1 Cell |
| ARHGEF26 | 1.12 | 0.00 | 0.00 | 0.34 | 0.05 | 6 | AT1 Cell |
| IGFBP6 | 1.12 | 0.00 | 0.00 | 0.27 | 0.03 | 6 | AT1 Cell |
| ERRFI1 | 1.11 | 0.00 | 0.00 | 0.40 | 0.07 | 6 | AT1 Cell |
| PRKCI | 1.11 | 0.00 | 0.00 | 0.46 | 0.10 | 6 | AT1 Cell |
| IER3 | 1.11 | 0.00 | 0.00 | 0.46 | 0.12 | 6 | AT1 Cell |
| CTTNBP2NL | 1.11 | 0.00 | 0.00 | 0.39 | 0.09 | 6 | AT1 Cell |
| GDE1 | 1.10 | 0.00 | 0.00 | 0.50 | 0.12 | 6 | AT1 Cell |
| SPHK1 | 1.10 | 0.00 | 0.00 | 0.26 | 0.02 | 6 | AT1 Cell |
| BCAM | 1.08 | 0.00 | 0.00 | 0.41 | 0.10 | 6 | AT1 Cell |
| FBLN5 | 1.07 | 0.00 | 0.00 | 0.58 | 0.16 | 6 | AT1 Cell |
| CHCHD10 | 1.07 | 0.00 | 0.00 | 0.48 | 0.11 | 6 | AT1 Cell |
| TSPAN2 | 1.06 | 0.00 | 0.00 | 0.43 | 0.10 | 6 | AT1 Cell |
| CDH1 | 1.06 | 0.00 | 0.00 | 0.36 | 0.05 | 6 | AT1 Cell |
| VAMP8 | 1.06 | 0.00 | 0.00 | 0.74 | 0.31 | 6 | AT1 Cell |
| CHKA | 1.06 | 0.00 | 0.00 | 0.41 | 0.07 | 6 | AT1 Cell |
| COL4A5 | 1.05 | 0.00 | 0.00 | 0.25 | 0.03 | 6 | AT1 Cell |
| KLF6 | 1.05 | 0.00 | 0.00 | 0.72 | 0.33 | 6 | AT1 Cell |
| FGF1 | 1.04 | 0.00 | 0.00 | 0.29 | 0.04 | 6 | AT1 Cell |
| TMEM63B | 1.04 | 0.00 | 0.00 | 0.33 | 0.05 | 6 | AT1 Cell |
| LAMB3 | 1.02 | 0.00 | 0.00 | 0.26 | 0.02 | 6 | AT1 Cell |
| GATA6 | 1.02 | 0.00 | 0.00 | 0.36 | 0.07 | 6 | AT1 Cell |
| MAL | 1.01 | 0.00 | 0.00 | 0.29 | 0.03 | 6 | AT1 Cell |
| FADS3 | 1.00 | 0.00 | 0.00 | 0.26 | 0.03 | 6 | AT1 Cell |
| CRB3 | 1.00 | 0.00 | 0.00 | 0.28 | 0.02 | 6 | AT1 Cell |
| C1orf116 | 0.99 | 0.00 | 0.00 | 0.26 | 0.02 | 6 | AT1 Cell |
| NRBP2 | 0.99 | 0.00 | 0.00 | 0.29 | 0.04 | 6 | AT1 Cell |
| NPNT | 0.97 | 0.00 | 0.00 | 0.75 | 0.21 | 6 | AT1 Cell |
| NCEH1 | 0.93 | 0.00 | 0.00 | 0.29 | 0.05 | 6 | AT1 Cell |
| IRX3 | 0.91 | 0.00 | 0.00 | 0.31 | 0.06 | 6 | AT1 Cell |
| SFTA2 | 0.90 | 0.00 | 0.00 | 0.50 | 0.13 | 6 | AT1 Cell |
| APP | 0.85 | 0.00 | 0.00 | 0.92 | 0.59 | 6 | AT1 Cell |
| MYO5C | 0.81 | 0.00 | 0.00 | 0.30 | 0.06 | 6 | AT1 Cell |
| LIMCH1 | 0.75 | 0.00 | 0.00 | 0.82 | 0.26 | 6 | AT1 Cell |
| ANXA2 | 0.97 | 0.00 | 0.00 | 0.84 | 0.48 | 6 | AT1 Cell |
| IGFBP7 | 0.93 | 0.00 | 0.00 | 0.83 | 0.41 | 6 | AT1 Cell |
| SAMHD1 | 0.88 | 0.00 | 0.00 | 0.54 | 0.17 | 6 | AT1 Cell |
| QKI | 0.95 | 0.00 | 0.00 | 0.65 | 0.27 | 6 | AT1 Cell |
| CYR61 | 0.99 | 0.00 | 0.00 | 0.40 | 0.11 | 6 | AT1 Cell |
| ATP8A1 | 0.89 | 0.00 | 0.00 | 0.45 | 0.13 | 6 | AT1 Cell |
| LGALS3 | 0.68 | 0.00 | 0.00 | 0.49 | 0.15 | 6 | AT1 Cell |
| S100A11 | 0.89 | 0.00 | 0.00 | 0.82 | 0.48 | 6 | AT1 Cell |
| CUX1 | 0.97 | 0.00 | 0.00 | 0.46 | 0.15 | 6 | AT1 Cell |
| TACC2 | 0.85 | 0.00 | 0.00 | 0.29 | 0.06 | 6 | AT1 Cell |
| FAM177A1 | 0.90 | 0.00 | 0.00 | 0.50 | 0.17 | 6 | AT1 Cell |
| BSG | 0.94 | 0.00 | 0.00 | 0.73 | 0.37 | 6 | AT1 Cell |
| PMP22 | 0.89 | 0.00 | 0.00 | 0.70 | 0.32 | 6 | AT1 Cell |
| SPARC | 0.71 | 0.00 | 0.00 | 0.98 | 0.75 | 6 | AT1 Cell |
| MSN | 0.84 | 0.00 | 0.00 | 0.80 | 0.43 | 6 | AT1 Cell |
| TEF | 0.77 | 0.00 | 0.00 | 0.27 | 0.06 | 6 | AT1 Cell |
| WNK1 | 0.92 | 0.00 | 0.00 | 0.46 | 0.15 | 6 | AT1 Cell |
| HDGF | 0.91 | 0.00 | 0.00 | 0.54 | 0.21 | 6 | AT1 Cell |
| CTDSPL | 0.88 | 0.00 | 0.00 | 0.34 | 0.09 | 6 | AT1 Cell |
| TFDP2 | 0.88 | 0.00 | 0.00 | 0.31 | 0.08 | 6 | AT1 Cell |
| SWT1 | 0.89 | 0.00 | 0.00 | 0.40 | 0.12 | 6 | AT1 Cell |
| CRIP2 | 0.77 | 0.00 | 0.00 | 0.78 | 0.42 | 6 | AT1 Cell |
| HSPG2 | 0.83 | 0.00 | 0.00 | 0.27 | 0.06 | 6 | AT1 Cell |
| WWC2 | 0.89 | 0.00 | 0.00 | 0.38 | 0.11 | 6 | AT1 Cell |
| LPIN2 | 0.93 | 0.00 | 0.00 | 0.40 | 0.12 | 6 | AT1 Cell |
| C16orf70 | 0.76 | 0.00 | 0.00 | 0.26 | 0.06 | 6 | AT1 Cell |
| CHPT1 | 0.85 | 0.00 | 0.00 | 0.35 | 0.10 | 6 | AT1 Cell |
| MBIP | 0.80 | 0.00 | 0.00 | 0.36 | 0.10 | 6 | AT1 Cell |
| FDX1 | 0.91 | 0.00 | 0.00 | 0.40 | 0.13 | 6 | AT1 Cell |
| DSTN | 0.67 | 0.00 | 0.00 | 0.82 | 0.52 | 6 | AT1 Cell |
| PGRMC1 | 0.92 | 0.00 | 0.00 | 0.48 | 0.18 | 6 | AT1 Cell |
| PTGS1 | 0.83 | 0.00 | 0.00 | 0.32 | 0.08 | 6 | AT1 Cell |
| TNS1 | 0.82 | 0.00 | 0.00 | 0.53 | 0.21 | 6 | AT1 Cell |
| DDX47 | 0.85 | 0.00 | 0.00 | 0.39 | 0.12 | 6 | AT1 Cell |
| AMOTL2 | 0.74 | 0.00 | 0.00 | 0.31 | 0.08 | 6 | AT1 Cell |
| WFDC2 | 0.53 | 0.00 | 0.00 | 0.44 | 0.14 | 6 | AT1 Cell |
| WLS | 0.86 | 0.00 | 0.00 | 0.51 | 0.20 | 6 | AT1 Cell |
| ARL4C | 0.75 | 0.00 | 0.00 | 0.30 | 0.08 | 6 | AT1 Cell |
| MYO1C | 0.79 | 0.00 | 0.00 | 0.32 | 0.09 | 6 | AT1 Cell |
| ASAH1 | 0.82 | 0.00 | 0.00 | 0.39 | 0.13 | 6 | AT1 Cell |
| CERS2 | 0.84 | 0.00 | 0.00 | 0.40 | 0.14 | 6 | AT1 Cell |
| LAMA5 | 0.78 | 0.00 | 0.00 | 0.29 | 0.08 | 6 | AT1 Cell |
| GPX4 | 0.80 | 0.00 | 0.00 | 0.67 | 0.37 | 6 | AT1 Cell |
| IQGAP1 | 0.69 | 0.00 | 0.00 | 0.73 | 0.42 | 6 | AT1 Cell |
| TJP2 | 0.76 | 0.00 | 0.00 | 0.41 | 0.15 | 6 | AT1 Cell |
| PHACTR1 | 0.81 | 0.00 | 0.00 | 0.25 | 0.06 | 6 | AT1 Cell |
| BTG3 | 0.79 | 0.00 | 0.00 | 0.34 | 0.11 | 6 | AT1 Cell |
| CNN2 | 0.66 | 0.00 | 0.00 | 0.71 | 0.37 | 6 | AT1 Cell |
| ANXA4 | 0.76 | 0.00 | 0.00 | 0.30 | 0.09 | 6 | AT1 Cell |
| KLF9 | 0.75 | 0.00 | 0.00 | 0.42 | 0.16 | 6 | AT1 Cell |
| UTRN | 0.77 | 0.00 | 0.00 | 0.41 | 0.15 | 6 | AT1 Cell |
| TMSB4X | 0.29 | 0.00 | 0.00 | 0.99 | 0.90 | 6 | AT1 Cell |
| EFNA1 | 0.75 | 0.00 | 0.00 | 0.28 | 0.08 | 6 | AT1 Cell |
| SDC4 | 0.62 | 0.00 | 0.00 | 0.48 | 0.19 | 6 | AT1 Cell |
| TMEM245 | 0.69 | 0.00 | 0.00 | 0.30 | 0.09 | 6 | AT1 Cell |
| RNASE4 | 0.68 | 0.00 | 0.00 | 0.43 | 0.16 | 6 | AT1 Cell |
| MRPL14 | 0.76 | 0.00 | 0.00 | 0.38 | 0.14 | 6 | AT1 Cell |
| ARRDC1 | 0.62 | 0.00 | 0.00 | 0.29 | 0.09 | 6 | AT1 Cell |
| GALNT18 | 0.71 | 0.00 | 0.00 | 0.31 | 0.10 | 6 | AT1 Cell |
| CREB3L2 | 0.80 | 0.00 | 0.00 | 0.35 | 0.12 | 6 | AT1 Cell |
| AQP1 | 0.63 | 0.00 | 0.00 | 0.48 | 0.19 | 6 | AT1 Cell |
| SFTPD | 0.71 | 0.00 | 0.00 | 0.40 | 0.15 | 6 | AT1 Cell |
| NCKAP5 | 0.68 | 0.00 | 0.00 | 0.34 | 0.12 | 6 | AT1 Cell |
| CDKN1A | 0.47 | 0.00 | 0.00 | 0.36 | 0.12 | 6 | AT1 Cell |
| DPYSL2 | 0.80 | 0.00 | 0.00 | 0.42 | 0.18 | 6 | AT1 Cell |
| RABAC1 | 0.72 | 0.00 | 0.00 | 0.57 | 0.29 | 6 | AT1 Cell |
| EPN2 | 0.71 | 0.00 | 0.00 | 0.32 | 0.11 | 6 | AT1 Cell |
| NAPSA | 0.76 | 0.00 | 0.00 | 0.45 | 0.19 | 6 | AT1 Cell |
| SOD1 | 0.69 | 0.00 | 0.00 | 0.71 | 0.47 | 6 | AT1 Cell |
| GADD45B | 0.70 | 0.00 | 0.00 | 0.28 | 0.09 | 6 | AT1 Cell |
| RAI14 | 0.60 | 0.00 | 0.00 | 0.30 | 0.10 | 6 | AT1 Cell |
| ARHGAP5 | 0.74 | 0.00 | 0.00 | 0.40 | 0.16 | 6 | AT1 Cell |
| CTNNA1 | 0.64 | 0.00 | 0.00 | 0.64 | 0.35 | 6 | AT1 Cell |
| LRPAP1 | 0.86 | 0.00 | 0.00 | 0.39 | 0.16 | 6 | AT1 Cell |
| HES1 | 0.62 | 0.00 | 0.00 | 0.57 | 0.28 | 6 | AT1 Cell |
| FNBP1L | 0.67 | 0.00 | 0.00 | 0.50 | 0.23 | 6 | AT1 Cell |
| RABGAP1L | 0.65 | 0.00 | 0.00 | 0.26 | 0.08 | 6 | AT1 Cell |
| GSN | 0.29 | 0.00 | 0.00 | 0.53 | 0.24 | 6 | AT1 Cell |
| PCNT | 0.61 | 0.00 | 0.00 | 0.31 | 0.11 | 6 | AT1 Cell |
| NBEAL1 | 0.64 | 0.00 | 0.00 | 0.33 | 0.12 | 6 | AT1 Cell |
| CAPN2 | 0.69 | 0.00 | 0.00 | 0.37 | 0.16 | 6 | AT1 Cell |
| ARL6IP1 | 0.58 | 0.00 | 0.00 | 0.73 | 0.49 | 6 | AT1 Cell |
| GNAS | 0.47 | 0.00 | 0.00 | 0.84 | 0.65 | 6 | AT1 Cell |
| PPP1R9A | 0.55 | 0.00 | 0.00 | 0.33 | 0.13 | 6 | AT1 Cell |
| RB1CC1 | 0.63 | 0.00 | 0.00 | 0.51 | 0.26 | 6 | AT1 Cell |
| TPPP3 | 0.67 | 0.00 | 0.00 | 0.33 | 0.13 | 6 | AT1 Cell |
| ELOVL5 | 0.58 | 0.00 | 0.00 | 0.35 | 0.14 | 6 | AT1 Cell |
| LPCAT1 | 0.49 | 0.00 | 0.00 | 0.40 | 0.17 | 6 | AT1 Cell |
| MECOM | 0.55 | 0.00 | 0.00 | 0.26 | 0.09 | 6 | AT1 Cell |
| RALBP1 | 0.57 | 0.00 | 0.00 | 0.55 | 0.30 | 6 | AT1 Cell |
| MYO1B | 0.65 | 0.00 | 0.00 | 0.36 | 0.15 | 6 | AT1 Cell |
| PGAP2 | 0.65 | 0.00 | 0.00 | 0.25 | 0.09 | 6 | AT1 Cell |
| PDLIM2 | 0.61 | 0.00 | 0.00 | 0.39 | 0.18 | 6 | AT1 Cell |
| ATP6V1A | 0.56 | 0.00 | 0.00 | 0.49 | 0.25 | 6 | AT1 Cell |
| SPTLC2 | 0.60 | 0.00 | 0.00 | 0.33 | 0.13 | 6 | AT1 Cell |
| CHP1 | 0.55 | 0.00 | 0.00 | 0.32 | 0.13 | 6 | AT1 Cell |
| ZFYVE21 | 0.63 | 0.00 | 0.00 | 0.26 | 0.09 | 6 | AT1 Cell |
| KIAA0040 | 0.60 | 0.00 | 0.00 | 0.37 | 0.17 | 6 | AT1 Cell |
| SLC38A2 | 0.59 | 0.00 | 0.00 | 0.39 | 0.18 | 6 | AT1 Cell |
| VAPA | 0.59 | 0.00 | 0.00 | 0.45 | 0.23 | 6 | AT1 Cell |
| TPM3 | 0.44 | 0.00 | 0.00 | 0.87 | 0.66 | 6 | AT1 Cell |
| ARHGEF2 | 0.59 | 0.00 | 0.00 | 0.40 | 0.19 | 6 | AT1 Cell |
| RBMS1 | 0.56 | 0.00 | 0.00 | 0.61 | 0.37 | 6 | AT1 Cell |
| ECE1 | 0.48 | 0.00 | 0.00 | 0.48 | 0.24 | 6 | AT1 Cell |
| RNF181 | 0.60 | 0.00 | 0.00 | 0.30 | 0.13 | 6 | AT1 Cell |
| EHD2 | 0.62 | 0.00 | 0.00 | 0.35 | 0.16 | 6 | AT1 Cell |
| LAS1L | 0.83 | 0.00 | 0.00 | 0.29 | 0.12 | 6 | AT1 Cell |
| NPC2 | 0.58 | 0.00 | 0.00 | 0.72 | 0.54 | 6 | AT1 Cell |
| BTG2 | 0.49 | 0.00 | 0.00 | 0.43 | 0.21 | 6 | AT1 Cell |
| ACTN4 | 0.56 | 0.00 | 0.00 | 0.51 | 0.29 | 6 | AT1 Cell |
| MPRIP | 0.54 | 0.00 | 0.00 | 0.40 | 0.19 | 6 | AT1 Cell |
| ZDHHC3 | 0.48 | 0.00 | 0.00 | 0.33 | 0.15 | 6 | AT1 Cell |
| DDB1 | 0.55 | 0.00 | 0.00 | 0.36 | 0.17 | 6 | AT1 Cell |
| TGFB2 | 0.52 | 0.00 | 0.00 | 0.31 | 0.13 | 6 | AT1 Cell |
| CRLF3 | 0.46 | 0.00 | 0.00 | 0.29 | 0.12 | 6 | AT1 Cell |
| KANK2 | 0.53 | 0.00 | 0.00 | 0.30 | 0.13 | 6 | AT1 Cell |
| CALM2 | 0.45 | 0.00 | 0.00 | 0.83 | 0.65 | 6 | AT1 Cell |
| SPTBN1 | 0.43 | 0.00 | 0.00 | 0.77 | 0.53 | 6 | AT1 Cell |
| SSR2 | 0.54 | 0.00 | 0.00 | 0.56 | 0.33 | 6 | AT1 Cell |
| CREBRF | 0.50 | 0.00 | 0.00 | 0.26 | 0.11 | 6 | AT1 Cell |
| TM9SF3 | 0.52 | 0.00 | 0.00 | 0.54 | 0.32 | 6 | AT1 Cell |
| SYPL1 | 0.48 | 0.00 | 0.00 | 0.47 | 0.25 | 6 | AT1 Cell |
| DRAP1 | 0.50 | 0.00 | 0.00 | 0.44 | 0.24 | 6 | AT1 Cell |
| AKAP13 | 0.43 | 0.00 | 0.00 | 0.47 | 0.25 | 6 | AT1 Cell |
| TAOK3 | 0.43 | 0.00 | 0.00 | 0.37 | 0.18 | 6 | AT1 Cell |
| ROCK1 | 0.50 | 0.00 | 0.00 | 0.54 | 0.33 | 6 | AT1 Cell |
| CDKN1B | 0.50 | 0.00 | 0.00 | 0.38 | 0.19 | 6 | AT1 Cell |
| AKAP9 | 0.48 | 0.00 | 0.00 | 0.55 | 0.34 | 6 | AT1 Cell |
| SLC12A2 | 0.46 | 0.00 | 0.00 | 0.31 | 0.14 | 6 | AT1 Cell |
| SRSF11 | 0.56 | 0.00 | 0.00 | 0.66 | 0.48 | 6 | AT1 Cell |
| CD2AP | 0.42 | 0.00 | 0.00 | 0.54 | 0.31 | 6 | AT1 Cell |
| COL4A2 | 0.39 | 0.00 | 0.00 | 0.67 | 0.44 | 6 | AT1 Cell |
| ZFHX3 | 0.50 | 0.00 | 0.00 | 0.34 | 0.16 | 6 | AT1 Cell |
| ABCD3 | 0.42 | 0.00 | 0.00 | 0.35 | 0.17 | 6 | AT1 Cell |
| PHLDB2 | 0.44 | 0.00 | 0.00 | 0.49 | 0.27 | 6 | AT1 Cell |
| DNAJC3 | 0.49 | 0.00 | 0.00 | 0.44 | 0.25 | 6 | AT1 Cell |
| TJP1 | 0.43 | 0.00 | 0.00 | 0.35 | 0.17 | 6 | AT1 Cell |
| CAB39 | 0.45 | 0.00 | 0.00 | 0.31 | 0.14 | 6 | AT1 Cell |
| LRRFIP1 | 0.44 | 0.00 | 0.00 | 0.45 | 0.25 | 6 | AT1 Cell |
| MYO9A | 0.53 | 0.00 | 0.00 | 0.27 | 0.12 | 6 | AT1 Cell |
| DRAM1 | 0.29 | 0.00 | 0.00 | 0.27 | 0.12 | 6 | AT1 Cell |
| TMBIM6 | 0.42 | 0.00 | 0.00 | 0.61 | 0.39 | 6 | AT1 Cell |
| OSBPL9 | 0.45 | 0.00 | 0.00 | 0.27 | 0.12 | 6 | AT1 Cell |
| LAMP1 | 0.42 | 0.00 | 0.00 | 0.64 | 0.42 | 6 | AT1 Cell |
| SRRM2 | 0.40 | 0.00 | 0.00 | 0.75 | 0.58 | 6 | AT1 Cell |
| SEC14L1 | 0.43 | 0.00 | 0.00 | 0.26 | 0.11 | 6 | AT1 Cell |
| YPEL3 | 0.37 | 0.00 | 0.00 | 0.37 | 0.19 | 6 | AT1 Cell |
| WWTR1 | 0.48 | 0.00 | 0.00 | 0.29 | 0.14 | 6 | AT1 Cell |
| ACTN1 | 0.46 | 0.00 | 0.00 | 0.41 | 0.23 | 6 | AT1 Cell |
| COL4A3BP | 0.41 | 0.00 | 0.00 | 0.29 | 0.14 | 6 | AT1 Cell |
| SYNE2 | 0.47 | 0.00 | 0.00 | 0.28 | 0.13 | 6 | AT1 Cell |
| JUND | 0.43 | 0.00 | 0.00 | 0.45 | 0.26 | 6 | AT1 Cell |
| RHOA | 0.45 | 0.00 | 0.00 | 0.67 | 0.48 | 6 | AT1 Cell |
| SFTPB | 0.27 | 0.00 | 0.00 | 0.35 | 0.17 | 6 | AT1 Cell |
| KRAS | 0.47 | 0.00 | 0.00 | 0.34 | 0.17 | 6 | AT1 Cell |
| BRI3 | 0.42 | 0.00 | 0.00 | 0.37 | 0.20 | 6 | AT1 Cell |
| SLC34A2 | 0.42 | 0.00 | 0.00 | 0.31 | 0.15 | 6 | AT1 Cell |
| STX7 | 0.50 | 0.00 | 0.00 | 0.28 | 0.14 | 6 | AT1 Cell |
| TBRG1 | 0.43 | 0.00 | 0.00 | 0.46 | 0.27 | 6 | AT1 Cell |
| TSC22D3 | 0.38 | 0.00 | 0.00 | 0.40 | 0.22 | 6 | AT1 Cell |
| CAV2 | 0.34 | 0.00 | 0.00 | 0.40 | 0.22 | 6 | AT1 Cell |
| S100G | 0.42 | 0.00 | 0.00 | 0.26 | 0.12 | 6 | AT1 Cell |
| SVIL | 0.46 | 0.00 | 0.00 | 0.31 | 0.16 | 6 | AT1 Cell |
| HPCAL1 | 0.37 | 0.00 | 0.00 | 0.29 | 0.15 | 6 | AT1 Cell |
| ACADL | 0.36 | 0.00 | 0.00 | 0.36 | 0.20 | 6 | AT1 Cell |
| TBL1X | 0.39 | 0.00 | 0.00 | 0.39 | 0.22 | 6 | AT1 Cell |
| APLP2 | 0.41 | 0.00 | 0.00 | 0.58 | 0.40 | 6 | AT1 Cell |
| HMGN1 | 0.34 | 0.00 | 0.00 | 0.64 | 0.45 | 6 | AT1 Cell |
| BAG1 | 0.44 | 0.00 | 0.00 | 0.39 | 0.23 | 6 | AT1 Cell |
| MBNL1 | 0.35 | 0.00 | 0.00 | 0.42 | 0.25 | 6 | AT1 Cell |
| ATP2B1 | 0.29 | 0.00 | 0.00 | 0.75 | 0.55 | 6 | AT1 Cell |
| SNX4 | 0.37 | 0.00 | 0.00 | 0.40 | 0.24 | 6 | AT1 Cell |
| PARVA | 0.42 | 0.00 | 0.00 | 0.29 | 0.15 | 6 | AT1 Cell |
| SECISBP2L | 0.36 | 0.00 | 0.00 | 0.31 | 0.17 | 6 | AT1 Cell |
| KIF5B | 0.34 | 0.00 | 0.00 | 0.64 | 0.47 | 6 | AT1 Cell |
| LARP4B | 0.33 | 0.00 | 0.00 | 0.26 | 0.13 | 6 | AT1 Cell |
| TMEM59 | 0.38 | 0.00 | 0.00 | 0.59 | 0.42 | 6 | AT1 Cell |
| PPP2R5A | 0.34 | 0.00 | 0.00 | 0.27 | 0.14 | 6 | AT1 Cell |
| PHF3 | 0.35 | 0.00 | 0.00 | 0.37 | 0.22 | 6 | AT1 Cell |
| ADIPOR1 | 0.34 | 0.00 | 0.00 | 0.37 | 0.22 | 6 | AT1 Cell |
| ATP1A1 | 0.33 | 0.00 | 0.00 | 0.27 | 0.14 | 6 | AT1 Cell |
| PKM | 0.37 | 0.00 | 0.00 | 0.39 | 0.23 | 6 | AT1 Cell |
| ANKRD12 | 0.42 | 0.00 | 0.00 | 0.41 | 0.26 | 6 | AT1 Cell |
| ARID4B | 0.36 | 0.00 | 0.00 | 0.48 | 0.32 | 6 | AT1 Cell |
| GAS6 | 0.34 | 0.00 | 0.00 | 0.29 | 0.15 | 6 | AT1 Cell |
| USP34 | 0.35 | 0.00 | 0.00 | 0.38 | 0.23 | 6 | AT1 Cell |
| DUSP1 | 0.33 | 0.00 | 0.00 | 0.30 | 0.17 | 6 | AT1 Cell |
| PTOV1 | 0.40 | 0.00 | 0.00 | 0.27 | 0.14 | 6 | AT1 Cell |
| TRA2A | 0.39 | 0.00 | 0.00 | 0.35 | 0.20 | 6 | AT1 Cell |
| SPTAN1 | 0.39 | 0.00 | 0.00 | 0.29 | 0.16 | 6 | AT1 Cell |
| LRRFIP2 | 0.36 | 0.00 | 0.00 | 0.25 | 0.13 | 6 | AT1 Cell |
| ZFP91 | 0.34 | 0.00 | 0.00 | 0.42 | 0.27 | 6 | AT1 Cell |
| WASL | 0.37 | 0.00 | 0.00 | 0.29 | 0.17 | 6 | AT1 Cell |
| PPIC | 0.33 | 0.00 | 0.00 | 0.56 | 0.38 | 6 | AT1 Cell |
| P4HB | 0.36 | 0.00 | 0.00 | 0.57 | 0.41 | 6 | AT1 Cell |
| JUP | 0.26 | 0.00 | 0.00 | 0.29 | 0.16 | 6 | AT1 Cell |
| PLS3 | 0.36 | 0.00 | 0.00 | 0.35 | 0.21 | 6 | AT1 Cell |
| BRD7 | 0.31 | 0.00 | 0.00 | 0.40 | 0.25 | 6 | AT1 Cell |
| EIF1 | 0.27 | 0.00 | 0.00 | 0.78 | 0.69 | 6 | AT1 Cell |
| MLF2 | 0.39 | 0.00 | 0.00 | 0.27 | 0.15 | 6 | AT1 Cell |
| ADAM10 | 0.34 | 0.00 | 0.00 | 0.30 | 0.17 | 6 | AT1 Cell |
| TM2D2 | 0.35 | 0.00 | 0.00 | 0.27 | 0.15 | 6 | AT1 Cell |
| LAMC1 | 0.34 | 0.00 | 0.00 | 0.27 | 0.16 | 6 | AT1 Cell |
| ATP6V1E1 | 0.29 | 0.00 | 0.00 | 0.39 | 0.25 | 6 | AT1 Cell |
| SIVA1 | 0.30 | 0.00 | 0.00 | 0.27 | 0.16 | 6 | AT1 Cell |
| UBL3 | 0.29 | 0.00 | 0.00 | 0.30 | 0.18 | 6 | AT1 Cell |
| ANXA5 | 0.35 | 0.00 | 0.00 | 0.56 | 0.42 | 6 | AT1 Cell |
| LAMP2 | 0.28 | 0.00 | 0.00 | 0.39 | 0.25 | 6 | AT1 Cell |
| FOS | 0.29 | 0.00 | 0.00 | 0.29 | 0.17 | 6 | AT1 Cell |
| RAB5A | 0.30 | 0.00 | 0.00 | 0.26 | 0.15 | 6 | AT1 Cell |
| RPL22L1 | 0.27 | 0.00 | 0.00 | 0.65 | 0.53 | 6 | AT1 Cell |
| TMED2 | 0.31 | 0.00 | 0.00 | 0.39 | 0.26 | 6 | AT1 Cell |
| CFL2 | 0.28 | 0.00 | 0.00 | 0.36 | 0.23 | 6 | AT1 Cell |
| EMD | 0.28 | 0.00 | 0.00 | 0.52 | 0.37 | 6 | AT1 Cell |
| KTN1 | 0.29 | 0.00 | 0.00 | 0.52 | 0.38 | 6 | AT1 Cell |
| BCAP31 | 0.33 | 0.00 | 0.00 | 0.33 | 0.21 | 6 | AT1 Cell |
| ATP6V1G1 | 0.27 | 0.00 | 0.00 | 0.47 | 0.34 | 6 | AT1 Cell |
| ORC5 | 0.26 | 0.00 | 0.00 | 0.64 | 0.51 | 6 | AT1 Cell |
| ATXN10 | 0.32 | 0.00 | 0.00 | 0.30 | 0.19 | 6 | AT1 Cell |
| ZC3H7A | 0.29 | 0.00 | 0.00 | 0.39 | 0.26 | 6 | AT1 Cell |
| OS9 | 0.34 | 0.00 | 0.00 | 0.32 | 0.21 | 6 | AT1 Cell |
| TMED10 | 0.28 | 0.00 | 0.00 | 0.48 | 0.35 | 6 | AT1 Cell |
| CHMP2B | 0.29 | 0.00 | 0.00 | 0.26 | 0.16 | 6 | AT1 Cell |
| MMP14 | 0.27 | 0.00 | 0.00 | 0.36 | 0.25 | 6 | AT1 Cell |
| TMEM176B | 0.31 | 0.00 | 0.00 | 0.44 | 0.32 | 6 | AT1 Cell |
| H2AFJ | 0.27 | 0.00 | 0.00 | 0.43 | 0.32 | 6 | AT1 Cell |
| TUBB4B | 0.36 | 0.00 | 0.00 | 0.26 | 0.18 | 6 | AT1 Cell |
| RAB6A | 0.26 | 0.00 | 0.00 | 0.29 | 0.21 | 6 | AT1 Cell |
| IGKC | 3.87 | 0.00 | 0.00 | 0.86 | 0.01 | 7 | B Cell |
| MS4A1 | 3.58 | 0.00 | 0.00 | 0.86 | 0.01 | 7 | B Cell |
| IGHM | 3.50 | 0.00 | 0.00 | 0.89 | 0.02 | 7 | B Cell |
| LY6D | 3.18 | 0.00 | 0.00 | 0.79 | 0.01 | 7 | B Cell |
| CD79A | 3.14 | 0.00 | 0.00 | 0.81 | 0.01 | 7 | B Cell |
| IGLL5 | 2.99 | 0.00 | 0.00 | 0.59 | 0.01 | 7 | B Cell |
| CD79B | 2.98 | 0.00 | 0.00 | 0.78 | 0.02 | 7 | B Cell |
| CD37 | 2.57 | 0.00 | 0.00 | 0.72 | 0.04 | 7 | B Cell |
| CD74 | 2.25 | 0.00 | 0.00 | 0.81 | 0.12 | 7 | B Cell |
| VPREB3 | 2.12 | 0.00 | 0.00 | 0.30 | 0.00 | 7 | B Cell |
| MZB1 | 2.09 | 0.00 | 0.00 | 0.44 | 0.00 | 7 | B Cell |
| HLA-DQB1 | 2.07 | 0.00 | 0.00 | 0.60 | 0.06 | 7 | B Cell |
| BANK1 | 2.03 | 0.00 | 0.00 | 0.40 | 0.01 | 7 | B Cell |
| SIGLEC10 | 1.96 | 0.00 | 0.00 | 0.41 | 0.01 | 7 | B Cell |
| PTPRCAP | 1.95 | 0.00 | 0.00 | 0.57 | 0.04 | 7 | B Cell |
| CD19 | 1.93 | 0.00 | 0.00 | 0.41 | 0.00 | 7 | B Cell |
| FCRLA | 1.92 | 0.00 | 0.00 | 0.40 | 0.00 | 7 | B Cell |
| CD52 | 1.84 | 0.00 | 0.00 | 0.87 | 0.16 | 7 | B Cell |
| PAX5 | 1.82 | 0.00 | 0.00 | 0.36 | 0.00 | 7 | B Cell |
| TNFRSF13C | 1.75 | 0.00 | 0.00 | 0.32 | 0.00 | 7 | B Cell |
| FAM129C | 1.70 | 0.00 | 0.00 | 0.32 | 0.01 | 7 | B Cell |
| SP110 | 1.60 | 0.00 | 0.00 | 0.40 | 0.06 | 7 | B Cell |
| PLAC8 | 1.59 | 0.00 | 0.00 | 0.74 | 0.14 | 7 | B Cell |
| SPIB | 1.56 | 0.00 | 0.00 | 0.28 | 0.00 | 7 | B Cell |
| LY86 | 1.55 | 0.00 | 0.00 | 0.43 | 0.05 | 7 | B Cell |
| EBF1 | 1.50 | 0.00 | 0.00 | 0.32 | 0.04 | 7 | B Cell |
| SYK | 1.50 | 0.00 | 0.00 | 0.35 | 0.04 | 7 | B Cell |
| BCL11A | 1.44 | 0.00 | 0.00 | 0.26 | 0.02 | 7 | B Cell |
| RALGPS2 | 1.42 | 0.00 | 0.00 | 0.34 | 0.05 | 7 | B Cell |
| RAC2 | 1.41 | 0.00 | 0.00 | 0.51 | 0.09 | 7 | B Cell |
| TNFRSF13B | 1.39 | 0.00 | 0.00 | 0.31 | 0.03 | 7 | B Cell |
| RNASE6 | 1.35 | 0.00 | 0.00 | 0.34 | 0.04 | 7 | B Cell |
| ARHGDIB | 1.33 | 0.00 | 0.00 | 0.61 | 0.15 | 7 | B Cell |
| APOBEC3H | 1.32 | 0.00 | 0.00 | 0.33 | 0.06 | 7 | B Cell |
| PTPN6 | 1.31 | 0.00 | 0.00 | 0.39 | 0.07 | 7 | B Cell |
| UNC93B1 | 1.26 | 0.00 | 0.00 | 0.32 | 0.06 | 7 | B Cell |
| CD53 | 1.22 | 0.00 | 0.00 | 0.40 | 0.08 | 7 | B Cell |
| ARHGAP30 | 1.21 | 0.00 | 0.00 | 0.40 | 0.09 | 7 | B Cell |
| LSP1 | 1.16 | 0.00 | 0.00 | 0.58 | 0.17 | 7 | B Cell |
| PTPRC | 1.00 | 0.00 | 0.00 | 0.51 | 0.13 | 7 | B Cell |
| CORO1A | 1.08 | 0.00 | 0.00 | 0.39 | 0.09 | 7 | B Cell |
| CD55 | 1.34 | 0.00 | 0.00 | 0.29 | 0.06 | 7 | B Cell |
| NCF1 | 1.09 | 0.00 | 0.00 | 0.29 | 0.05 | 7 | B Cell |
| VARS | 1.17 | 0.00 | 0.00 | 0.30 | 0.06 | 7 | B Cell |
| WDR89 | 0.69 | 0.00 | 0.00 | 0.94 | 0.82 | 7 | B Cell |
| LY6E | 0.85 | 0.00 | 0.00 | 0.79 | 0.42 | 7 | B Cell |
| PARP1 | 1.19 | 0.00 | 0.00 | 0.34 | 0.09 | 7 | B Cell |
| HCLS1 | 0.95 | 0.00 | 0.00 | 0.29 | 0.07 | 7 | B Cell |
| FAM111A | 1.09 | 0.00 | 0.00 | 0.40 | 0.13 | 7 | B Cell |
| MT-CO1 | 0.46 | 0.00 | 0.00 | 0.99 | 0.95 | 7 | B Cell |
| SUB1 | 1.07 | 0.00 | 0.00 | 0.72 | 0.51 | 7 | B Cell |
| PTPN18 | 0.95 | 0.00 | 0.00 | 0.39 | 0.12 | 7 | B Cell |
| LIMD2 | 1.08 | 0.00 | 0.00 | 0.40 | 0.14 | 7 | B Cell |
| SNX5 | 1.12 | 0.00 | 0.00 | 0.36 | 0.12 | 7 | B Cell |
| FAU | 0.55 | 0.00 | 0.00 | 0.95 | 0.87 | 7 | B Cell |
| SERP1 | 1.08 | 0.00 | 0.00 | 0.51 | 0.23 | 7 | B Cell |
| PSMB8 | 0.95 | 0.00 | 0.00 | 0.46 | 0.18 | 7 | B Cell |
| LRRC58 | 0.54 | 0.00 | 0.00 | 0.97 | 0.85 | 7 | B Cell |
| RPL18A | 0.58 | 0.00 | 0.00 | 0.95 | 0.83 | 7 | B Cell |
| RPLP1 | 0.50 | 0.00 | 0.00 | 0.97 | 0.91 | 7 | B Cell |
| SP140 | 0.92 | 0.00 | 0.00 | 0.27 | 0.07 | 7 | B Cell |
| RPL10A | 0.63 | 0.00 | 0.00 | 0.90 | 0.76 | 7 | B Cell |
| SH3BP5 | 1.02 | 0.00 | 0.00 | 0.46 | 0.19 | 7 | B Cell |
| SH3BGRL3 | 0.89 | 0.00 | 0.00 | 0.63 | 0.35 | 7 | B Cell |
| RPS20 | 0.60 | 0.00 | 0.00 | 0.91 | 0.80 | 7 | B Cell |
| MEF2C | 1.11 | 0.00 | 0.00 | 0.42 | 0.17 | 7 | B Cell |
| EEF1B2 | 0.67 | 0.00 | 0.00 | 0.80 | 0.65 | 7 | B Cell |
| GIMAP4 | 0.93 | 0.00 | 0.00 | 0.42 | 0.17 | 7 | B Cell |
| SNX2 | 1.05 | 0.00 | 0.00 | 0.39 | 0.16 | 7 | B Cell |
| CRIP1 | 0.95 | 0.00 | 0.00 | 0.72 | 0.47 | 7 | B Cell |
| PIM1 | 1.04 | 0.00 | 0.00 | 0.26 | 0.08 | 7 | B Cell |
| RPL8 | 0.55 | 0.00 | 0.00 | 0.90 | 0.81 | 7 | B Cell |
| NOL7 | 0.91 | 0.00 | 0.00 | 0.59 | 0.34 | 7 | B Cell |
| RABGAP1L | 0.99 | 0.00 | 0.00 | 0.27 | 0.08 | 7 | B Cell |
| GIMAP6 | 0.92 | 0.00 | 0.00 | 0.38 | 0.15 | 7 | B Cell |
| EEF1G | 0.83 | 0.00 | 0.00 | 0.67 | 0.46 | 7 | B Cell |
| TMSB4X | 0.33 | 0.00 | 0.00 | 0.98 | 0.90 | 7 | B Cell |
| GIMAP1 | 0.87 | 0.00 | 0.00 | 0.35 | 0.13 | 7 | B Cell |
| SERBP1 | 0.63 | 0.00 | 0.00 | 0.83 | 0.73 | 7 | B Cell |
| RPL6 | 0.46 | 0.00 | 0.00 | 0.95 | 0.88 | 7 | B Cell |
| ITGA4 | 0.80 | 0.00 | 0.00 | 0.28 | 0.09 | 7 | B Cell |
| SEC11C | 0.94 | 0.00 | 0.00 | 0.38 | 0.16 | 7 | B Cell |
| CAPG | 0.92 | 0.00 | 0.00 | 0.28 | 0.09 | 7 | B Cell |
| LCP1 | 0.79 | 0.00 | 0.00 | 0.45 | 0.20 | 7 | B Cell |
| RPLP2 | 0.48 | 0.00 | 0.00 | 0.91 | 0.83 | 7 | B Cell |
| NAP1L1 | 0.78 | 0.00 | 0.00 | 0.66 | 0.47 | 7 | B Cell |
| STRBP | 0.93 | 0.00 | 0.00 | 0.27 | 0.09 | 7 | B Cell |
| SHISA5 | 0.94 | 0.00 | 0.00 | 0.43 | 0.20 | 7 | B Cell |
| GMFG | 0.88 | 0.00 | 0.00 | 0.35 | 0.14 | 7 | B Cell |
| TCOF1 | 0.90 | 0.00 | 0.00 | 0.28 | 0.10 | 7 | B Cell |
| ARID3A | 0.92 | 0.00 | 0.00 | 0.27 | 0.09 | 7 | B Cell |
| IL2RG | 0.82 | 0.00 | 0.00 | 0.27 | 0.09 | 7 | B Cell |
| MT-CO3 | 0.42 | 0.00 | 0.00 | 0.95 | 0.89 | 7 | B Cell |
| NAPSA | 0.39 | 0.00 | 0.00 | 0.47 | 0.19 | 7 | B Cell |
| APBB1IP | 0.66 | 0.00 | 0.00 | 0.26 | 0.08 | 7 | B Cell |
| GRAP | 0.83 | 0.00 | 0.00 | 0.31 | 0.12 | 7 | B Cell |
| ARPC5L | 0.97 | 0.00 | 0.00 | 0.35 | 0.15 | 7 | B Cell |
| TPM3 | 0.59 | 0.00 | 0.00 | 0.81 | 0.67 | 7 | B Cell |
| RPS4X | 0.39 | 0.00 | 0.00 | 0.95 | 0.88 | 7 | B Cell |
| MYCBP2 | 0.95 | 0.00 | 0.00 | 0.43 | 0.22 | 7 | B Cell |
| CYTIP | 0.70 | 0.00 | 0.00 | 0.25 | 0.08 | 7 | B Cell |
| EZR | 0.81 | 0.00 | 0.00 | 0.41 | 0.20 | 7 | B Cell |
| HSP90AB1 | 0.46 | 0.00 | 0.00 | 0.92 | 0.89 | 7 | B Cell |
| KLF2 | 0.78 | 0.00 | 0.00 | 0.41 | 0.19 | 7 | B Cell |
| DNAJC9 | 0.91 | 0.00 | 0.00 | 0.27 | 0.10 | 7 | B Cell |
| SWAP70 | 0.90 | 0.00 | 0.00 | 0.27 | 0.10 | 7 | B Cell |
| MT-ND1 | 0.37 | 0.00 | 0.00 | 0.95 | 0.92 | 7 | B Cell |
| NME1 | 0.90 | 0.00 | 0.00 | 0.38 | 0.18 | 7 | B Cell |
| HIST1H2AB | 1.56 | 0.00 | 0.00 | 0.27 | 0.11 | 7 | B Cell |
| CYBA | 0.60 | 0.00 | 0.00 | 0.53 | 0.31 | 7 | B Cell |
| COTL1 | 0.74 | 0.00 | 0.00 | 0.36 | 0.17 | 7 | B Cell |
| H3F3A | 0.57 | 0.00 | 0.00 | 0.76 | 0.62 | 7 | B Cell |
| MT-ND5 | 0.48 | 0.00 | 0.00 | 0.84 | 0.75 | 7 | B Cell |
| MT-CO2 | 0.42 | 0.00 | 0.00 | 0.87 | 0.79 | 7 | B Cell |
| RPS13 | 0.50 | 0.00 | 0.00 | 0.79 | 0.66 | 7 | B Cell |
| BIN1 | 0.84 | 0.00 | 0.00 | 0.26 | 0.10 | 7 | B Cell |
| ZNF706 | 0.77 | 0.00 | 0.00 | 0.50 | 0.32 | 7 | B Cell |
| NOP58 | 0.89 | 0.00 | 0.00 | 0.42 | 0.23 | 7 | B Cell |
| ARHGAP17 | 0.82 | 0.00 | 0.00 | 0.33 | 0.16 | 7 | B Cell |
| DNAJC2 | 0.77 | 0.00 | 0.00 | 0.36 | 0.18 | 7 | B Cell |
| CCM2 | 0.85 | 0.00 | 0.00 | 0.25 | 0.11 | 7 | B Cell |
| CRLF3 | 0.75 | 0.00 | 0.00 | 0.28 | 0.12 | 7 | B Cell |
| ACTR3 | 0.69 | 0.00 | 0.00 | 0.51 | 0.32 | 7 | B Cell |
| HNRNPA2B1 | 0.50 | 0.00 | 0.00 | 0.79 | 0.72 | 7 | B Cell |
| NOP56 | 0.84 | 0.00 | 0.00 | 0.36 | 0.19 | 7 | B Cell |
| LYN | 0.70 | 0.00 | 0.00 | 0.27 | 0.12 | 7 | B Cell |
| ITSN2 | 0.77 | 0.00 | 0.00 | 0.41 | 0.23 | 7 | B Cell |
| ORC5 | 0.55 | 0.00 | 0.00 | 0.66 | 0.51 | 7 | B Cell |
| PTBP3 | 0.65 | 0.00 | 0.00 | 0.50 | 0.32 | 7 | B Cell |
| ARPC5 | 0.50 | 0.00 | 0.00 | 0.70 | 0.57 | 7 | B Cell |
| ETS1 | 0.65 | 0.00 | 0.00 | 0.42 | 0.23 | 7 | B Cell |
| RPL22L1 | 0.52 | 0.00 | 0.00 | 0.67 | 0.53 | 7 | B Cell |
| ERP29 | 0.70 | 0.00 | 0.00 | 0.42 | 0.26 | 7 | B Cell |
| LLPH | 0.66 | 0.00 | 0.00 | 0.48 | 0.31 | 7 | B Cell |
| ANP32B | 0.60 | 0.00 | 0.00 | 0.61 | 0.48 | 7 | B Cell |
| KIAA0040 | 0.73 | 0.00 | 0.00 | 0.32 | 0.17 | 7 | B Cell |
| MT-ATP6 | 0.31 | 0.00 | 0.00 | 0.90 | 0.85 | 7 | B Cell |
| PFN1 | 0.55 | 0.00 | 0.00 | 0.62 | 0.49 | 7 | B Cell |
| PDCD4 | 0.71 | 0.00 | 0.00 | 0.36 | 0.20 | 7 | B Cell |
| DDX5 | 0.43 | 0.00 | 0.00 | 0.78 | 0.71 | 7 | B Cell |
| CIB1 | 0.78 | 0.00 | 0.00 | 0.26 | 0.13 | 7 | B Cell |
| SRGN | 0.30 | 0.00 | 0.00 | 0.53 | 0.31 | 7 | B Cell |
| DNAJC7 | 0.74 | 0.00 | 0.00 | 0.35 | 0.20 | 7 | B Cell |
| HNRNPU | 0.47 | 0.00 | 0.00 | 0.70 | 0.62 | 7 | B Cell |
| RPS27A | 0.32 | 0.00 | 0.00 | 0.86 | 0.78 | 7 | B Cell |
| DDX21 | 0.70 | 0.00 | 0.00 | 0.40 | 0.26 | 7 | B Cell |
| ARPC2 | 0.37 | 0.00 | 0.00 | 0.73 | 0.61 | 7 | B Cell |
| LSM4 | 0.64 | 0.00 | 0.00 | 0.34 | 0.20 | 7 | B Cell |
| SMCHD1 | 0.72 | 0.00 | 0.00 | 0.27 | 0.14 | 7 | B Cell |
| PTMA | 0.32 | 0.00 | 0.00 | 0.93 | 0.92 | 7 | B Cell |
| STK38 | 0.64 | 0.00 | 0.00 | 0.31 | 0.18 | 7 | B Cell |
| RPL7 | 0.26 | 0.00 | 0.00 | 0.84 | 0.77 | 7 | B Cell |
| RAN | 0.61 | 0.00 | 0.00 | 0.50 | 0.39 | 7 | B Cell |
| RBM3 | 0.54 | 0.00 | 0.00 | 0.52 | 0.41 | 7 | B Cell |
| CCND2 | 0.66 | 0.00 | 0.00 | 0.41 | 0.27 | 7 | B Cell |
| HNRNPCL1 | 0.52 | 0.00 | 0.00 | 0.52 | 0.39 | 7 | B Cell |
| HSPE1 | 0.64 | 0.00 | 0.00 | 0.42 | 0.30 | 7 | B Cell |
| GPX1 | 0.38 | 0.00 | 0.00 | 0.57 | 0.44 | 7 | B Cell |
| SLC25A5 | 0.40 | 0.00 | 0.00 | 0.67 | 0.58 | 7 | B Cell |
| SMIM14 | 0.74 | 0.00 | 0.00 | 0.31 | 0.19 | 7 | B Cell |
| EML4 | 0.66 | 0.00 | 0.00 | 0.28 | 0.16 | 7 | B Cell |
| ARPC1B | 0.45 | 0.00 | 0.00 | 0.50 | 0.36 | 7 | B Cell |
| YEATS4 | 0.62 | 0.00 | 0.00 | 0.29 | 0.17 | 7 | B Cell |
| BTG1 | 0.66 | 0.00 | 0.00 | 0.42 | 0.30 | 7 | B Cell |
| ANP32E | 0.63 | 0.00 | 0.00 | 0.38 | 0.25 | 7 | B Cell |
| SF3B1 | 0.48 | 0.00 | 0.00 | 0.55 | 0.44 | 7 | B Cell |
| EIF5A | 0.55 | 0.00 | 0.00 | 0.45 | 0.33 | 7 | B Cell |
| GDI2 | 0.59 | 0.00 | 0.00 | 0.44 | 0.33 | 7 | B Cell |
| IVNS1ABP | 0.49 | 0.00 | 0.00 | 0.33 | 0.20 | 7 | B Cell |
| HMGB2 | 0.97 | 0.00 | 0.00 | 0.47 | 0.38 | 7 | B Cell |
| OAZ1 | 0.39 | 0.00 | 0.00 | 0.69 | 0.63 | 7 | B Cell |
| GRB2 | 0.58 | 0.00 | 0.00 | 0.29 | 0.17 | 7 | B Cell |
| EMG1 | 0.55 | 0.00 | 0.00 | 0.25 | 0.15 | 7 | B Cell |
| S100A10 | 0.47 | 0.00 | 0.00 | 0.52 | 0.41 | 7 | B Cell |
| CDC42SE1 | 0.56 | 0.00 | 0.00 | 0.29 | 0.18 | 7 | B Cell |
| GTF2F1 | 0.59 | 0.00 | 0.00 | 0.31 | 0.20 | 7 | B Cell |
| ACIN1 | 0.39 | 0.00 | 0.00 | 0.60 | 0.51 | 7 | B Cell |
| PGLS | 0.58 | 0.00 | 0.00 | 0.27 | 0.17 | 7 | B Cell |
| MSN | 0.41 | 0.00 | 0.00 | 0.55 | 0.44 | 7 | B Cell |
| CLTA | 0.43 | 0.00 | 0.00 | 0.52 | 0.43 | 7 | B Cell |
| PCNA | 0.63 | 0.00 | 0.00 | 0.26 | 0.16 | 7 | B Cell |
| HNRNPF | 0.48 | 0.00 | 0.00 | 0.44 | 0.34 | 7 | B Cell |
| FKBP3 | 0.52 | 0.00 | 0.00 | 0.39 | 0.29 | 7 | B Cell |
| PA2G4 | 0.61 | 0.00 | 0.00 | 0.36 | 0.26 | 7 | B Cell |
| DEK | 0.51 | 0.00 | 0.00 | 0.54 | 0.47 | 7 | B Cell |
| ZNF106 | 0.54 | 0.00 | 0.00 | 0.29 | 0.19 | 7 | B Cell |
| SNRPF | 0.51 | 0.00 | 0.00 | 0.38 | 0.28 | 7 | B Cell |
| EIF5 | 0.52 | 0.00 | 0.00 | 0.45 | 0.37 | 7 | B Cell |
| HNRNPM | 0.41 | 0.00 | 0.00 | 0.57 | 0.49 | 7 | B Cell |
| RSL1D1 | 0.55 | 0.00 | 0.00 | 0.37 | 0.27 | 7 | B Cell |
| NT5C | 0.52 | 0.00 | 0.00 | 0.26 | 0.16 | 7 | B Cell |
| NACA | 0.33 | 0.00 | 0.00 | 0.66 | 0.61 | 7 | B Cell |
| RRP1 | 0.47 | 0.00 | 0.00 | 0.43 | 0.34 | 7 | B Cell |
| XRN2 | 0.52 | 0.00 | 0.00 | 0.39 | 0.29 | 7 | B Cell |
| RANBP1 | 0.56 | 0.00 | 0.00 | 0.39 | 0.30 | 7 | B Cell |
| POLR1D | 0.48 | 0.00 | 0.00 | 0.37 | 0.28 | 7 | B Cell |
| PTPN1 | 0.48 | 0.00 | 0.00 | 0.29 | 0.19 | 7 | B Cell |
| TAF1D | 0.52 | 0.00 | 0.00 | 0.31 | 0.22 | 7 | B Cell |
| SNRPE | 0.44 | 0.00 | 0.00 | 0.45 | 0.37 | 7 | B Cell |
| NUCKS1 | 0.50 | 0.00 | 0.00 | 0.38 | 0.29 | 7 | B Cell |
| SAMHD1 | 0.38 | 0.00 | 0.00 | 0.29 | 0.19 | 7 | B Cell |
| AKAP13 | 0.52 | 0.00 | 0.00 | 0.36 | 0.26 | 7 | B Cell |
| CEBPZ | 0.51 | 0.00 | 0.00 | 0.26 | 0.17 | 7 | B Cell |
| EIF3CL | 0.39 | 0.00 | 0.00 | 0.51 | 0.43 | 7 | B Cell |
| RPL36A | 0.29 | 0.00 | 0.00 | 0.69 | 0.64 | 7 | B Cell |
| SAFB2 | 0.51 | 0.00 | 0.00 | 0.31 | 0.22 | 7 | B Cell |
| ARL6IP1 | 0.41 | 0.00 | 0.00 | 0.57 | 0.50 | 7 | B Cell |
| HSPD1 | 0.50 | 0.00 | 0.00 | 0.34 | 0.26 | 7 | B Cell |
| SRRM2 | 0.32 | 0.00 | 0.00 | 0.63 | 0.59 | 7 | B Cell |
| PDIA4 | 0.52 | 0.00 | 0.00 | 0.29 | 0.21 | 7 | B Cell |
| MIF | 0.49 | 0.00 | 0.00 | 0.31 | 0.23 | 7 | B Cell |
| DDX46 | 0.43 | 0.00 | 0.00 | 0.38 | 0.30 | 7 | B Cell |
| CLIC1 | 0.35 | 0.00 | 0.00 | 0.47 | 0.39 | 7 | B Cell |
| DHX9 | 0.47 | 0.00 | 0.00 | 0.26 | 0.18 | 7 | B Cell |
| NAA15 | 0.44 | 0.00 | 0.00 | 0.29 | 0.20 | 7 | B Cell |
| EMP3 | 0.42 | 0.00 | 0.00 | 0.33 | 0.24 | 7 | B Cell |
| ACTR2 | 0.39 | 0.00 | 0.00 | 0.36 | 0.27 | 7 | B Cell |
| FGFR1OP2 | 0.40 | 0.00 | 0.00 | 0.35 | 0.26 | 7 | B Cell |
| SUPT16H | 0.45 | 0.00 | 0.00 | 0.31 | 0.23 | 7 | B Cell |
| MAT2A | 0.47 | 0.00 | 0.00 | 0.33 | 0.25 | 7 | B Cell |
| CTCF | 0.47 | 0.00 | 0.00 | 0.29 | 0.21 | 7 | B Cell |
| MBNL1 | 0.38 | 0.00 | 0.00 | 0.33 | 0.25 | 7 | B Cell |
| YWHAZ | 0.28 | 0.00 | 0.00 | 0.56 | 0.50 | 7 | B Cell |
| PSME1 | 0.40 | 0.00 | 0.00 | 0.41 | 0.34 | 7 | B Cell |
| SMARCA5 | 0.40 | 0.00 | 0.00 | 0.36 | 0.28 | 7 | B Cell |
| NASP | 0.47 | 0.00 | 0.00 | 0.26 | 0.19 | 7 | B Cell |
| SLTM | 0.41 | 0.00 | 0.00 | 0.45 | 0.38 | 7 | B Cell |
| PSMB1 | 0.36 | 0.00 | 0.00 | 0.52 | 0.48 | 7 | B Cell |
| GCLM | 0.35 | 0.00 | 0.00 | 0.44 | 0.38 | 7 | B Cell |
| HSPA4 | 0.41 | 0.00 | 0.00 | 0.29 | 0.22 | 7 | B Cell |
| CCT2 | 0.43 | 0.00 | 0.00 | 0.33 | 0.26 | 7 | B Cell |
| SERINC3 | 0.38 | 0.00 | 0.00 | 0.37 | 0.30 | 7 | B Cell |
| TCP1 | 0.40 | 0.00 | 0.00 | 0.30 | 0.23 | 7 | B Cell |
| ADD3 | 0.38 | 0.00 | 0.00 | 0.35 | 0.27 | 7 | B Cell |
| MTPN | 0.38 | 0.00 | 0.00 | 0.33 | 0.26 | 7 | B Cell |
| IRF2 | 0.43 | 0.00 | 0.00 | 0.30 | 0.22 | 7 | B Cell |
| SSRP1 | 0.41 | 0.00 | 0.00 | 0.27 | 0.20 | 7 | B Cell |
| TMOD3 | 0.42 | 0.00 | 0.00 | 0.33 | 0.26 | 7 | B Cell |
| KMT2A | 0.43 | 0.00 | 0.00 | 0.26 | 0.19 | 7 | B Cell |
| MTDH | 0.35 | 0.00 | 0.00 | 0.52 | 0.48 | 7 | B Cell |
| TSC22D4 | 0.40 | 0.00 | 0.00 | 0.32 | 0.26 | 7 | B Cell |
| CAPZB | 0.29 | 0.00 | 0.00 | 0.49 | 0.44 | 7 | B Cell |
| THRAP3 | 0.36 | 0.00 | 0.00 | 0.41 | 0.36 | 7 | B Cell |
| CCT5 | 0.42 | 0.00 | 0.00 | 0.29 | 0.23 | 7 | B Cell |
| ENO1 | 0.36 | 0.00 | 0.00 | 0.25 | 0.19 | 7 | B Cell |
| SNRPD3 | 0.38 | 0.00 | 0.00 | 0.37 | 0.31 | 7 | B Cell |
| CNBP | 0.34 | 0.00 | 0.00 | 0.50 | 0.47 | 7 | B Cell |
| MT-ND4L | 0.34 | 0.00 | 0.00 | 0.43 | 0.39 | 7 | B Cell |
| PNN | 0.27 | 0.00 | 0.00 | 0.55 | 0.51 | 7 | B Cell |
| DDX6 | 0.39 | 0.00 | 0.00 | 0.34 | 0.29 | 7 | B Cell |
| KMT2E | 0.35 | 0.00 | 0.00 | 0.47 | 0.43 | 7 | B Cell |
| PPP1CA | 0.34 | 0.00 | 0.00 | 0.38 | 0.33 | 7 | B Cell |
| RBM17 | 0.39 | 0.00 | 0.00 | 0.30 | 0.25 | 7 | B Cell |
| HNRNPDL | 0.34 | 0.00 | 0.00 | 0.42 | 0.38 | 7 | B Cell |
| TRA2B | 0.39 | 0.00 | 0.00 | 0.35 | 0.30 | 7 | B Cell |
| H2AFY | 0.39 | 0.00 | 0.00 | 0.26 | 0.21 | 7 | B Cell |
| SBNO1 | 0.43 | 0.00 | 0.00 | 0.26 | 0.20 | 7 | B Cell |
| NOP10 | 0.38 | 0.00 | 0.00 | 0.34 | 0.29 | 7 | B Cell |
| SNRPD1 | 0.37 | 0.00 | 0.00 | 0.26 | 0.21 | 7 | B Cell |
| SRSF8 | 0.31 | 0.00 | 0.00 | 0.40 | 0.36 | 7 | B Cell |
| SF3B2 | 0.34 | 0.00 | 0.00 | 0.42 | 0.38 | 7 | B Cell |
| JUND | 0.39 | 0.00 | 0.00 | 0.32 | 0.27 | 7 | B Cell |
| UTP3 | 0.36 | 0.00 | 0.00 | 0.25 | 0.20 | 7 | B Cell |
| HNRNPD | 0.37 | 0.00 | 0.00 | 0.33 | 0.28 | 7 | B Cell |
| RSBN1L | 0.37 | 0.00 | 0.00 | 0.29 | 0.24 | 7 | B Cell |
| EIF3K | 0.32 | 0.00 | 0.00 | 0.40 | 0.37 | 7 | B Cell |
| CCDC50 | 0.34 | 0.00 | 0.00 | 0.30 | 0.25 | 7 | B Cell |
| SRSF3 | 0.34 | 0.00 | 0.00 | 0.32 | 0.28 | 7 | B Cell |
| EIF3E | 0.26 | 0.00 | 0.00 | 0.46 | 0.43 | 7 | B Cell |
| PRPF4B | 0.27 | 0.00 | 0.00 | 0.46 | 0.43 | 7 | B Cell |
| BTG2 | 0.32 | 0.00 | 0.00 | 0.26 | 0.22 | 7 | B Cell |
| SEPT11 | 0.32 | 0.00 | 0.01 | 0.27 | 0.23 | 7 | B Cell |
| CCT8 | 0.33 | 0.00 | 0.01 | 0.25 | 0.21 | 7 | B Cell |
| RPL17-C18orf32 | 0.27 | 0.00 | 0.01 | 0.35 | 0.31 | 7 | B Cell |
| SCAF11 | 0.33 | 0.00 | 0.01 | 0.31 | 0.27 | 7 | B Cell |
| DUSP11 | 0.30 | 0.00 | 0.01 | 0.29 | 0.24 | 7 | B Cell |
| HNRNPA3 | 0.26 | 0.00 | 0.01 | 0.47 | 0.47 | 7 | B Cell |
| MT-ATP8 | 0.33 | 0.00 | 0.02 | 0.29 | 0.25 | 7 | B Cell |
| ASH1L | 0.34 | 0.00 | 0.03 | 0.32 | 0.28 | 7 | B Cell |
| PSME2 | 0.28 | 0.00 | 0.04 | 0.29 | 0.26 | 7 | B Cell |
| TRBC1 | 2.96 | 0.00 | 0.00 | 0.70 | 0.01 | 8 | T Cell |
| CCL5 | 2.85 | 0.00 | 0.00 | 0.26 | 0.01 | 8 | T Cell |
| CD3D | 2.62 | 0.00 | 0.00 | 0.62 | 0.01 | 8 | T Cell |
| CD3G | 2.59 | 0.00 | 0.00 | 0.65 | 0.01 | 8 | T Cell |
| NKG7 | 2.23 | 0.00 | 0.00 | 0.37 | 0.01 | 8 | T Cell |
| TCF7 | 2.07 | 0.00 | 0.00 | 0.46 | 0.04 | 8 | T Cell |
| KLRD1 | 2.04 | 0.00 | 0.00 | 0.29 | 0.00 | 8 | T Cell |
| PTPRCAP | 2.00 | 0.00 | 0.00 | 0.60 | 0.04 | 8 | T Cell |
| LCK | 1.98 | 0.00 | 0.00 | 0.44 | 0.01 | 8 | T Cell |
| PTPRC | 1.96 | 0.00 | 0.00 | 0.75 | 0.12 | 8 | T Cell |
| IL7R | 1.95 | 0.00 | 0.00 | 0.34 | 0.02 | 8 | T Cell |
| RAC2 | 1.88 | 0.00 | 0.00 | 0.65 | 0.08 | 8 | T Cell |
| CD52 | 1.86 | 0.00 | 0.00 | 0.86 | 0.16 | 8 | T Cell |
| CCR7 | 1.84 | 0.00 | 0.00 | 0.29 | 0.01 | 8 | T Cell |
| SELPLG | 1.75 | 0.00 | 0.00 | 0.45 | 0.04 | 8 | T Cell |
| HCST | 1.72 | 0.00 | 0.00 | 0.40 | 0.03 | 8 | T Cell |
| LEF1 | 1.71 | 0.00 | 0.00 | 0.33 | 0.03 | 8 | T Cell |
| CD27 | 1.66 | 0.00 | 0.00 | 0.32 | 0.01 | 8 | T Cell |
| SEPT1 | 1.65 | 0.00 | 0.00 | 0.40 | 0.03 | 8 | T Cell |
| SHISA5 | 1.65 | 0.00 | 0.00 | 0.66 | 0.19 | 8 | T Cell |
| THY1 | 1.65 | 0.00 | 0.00 | 0.37 | 0.03 | 8 | T Cell |
| SKAP1 | 1.59 | 0.00 | 0.00 | 0.33 | 0.01 | 8 | T Cell |
| BCL11B | 1.59 | 0.00 | 0.00 | 0.28 | 0.01 | 8 | T Cell |
| LIMD2 | 1.58 | 0.00 | 0.00 | 0.57 | 0.13 | 8 | T Cell |
| CD3E | 1.54 | 0.00 | 0.00 | 0.29 | 0.00 | 8 | T Cell |
| GIMAP4 | 1.53 | 0.00 | 0.00 | 0.61 | 0.16 | 8 | T Cell |
| ARHGDIB | 1.52 | 0.00 | 0.00 | 0.68 | 0.15 | 8 | T Cell |
| SP140 | 1.52 | 0.00 | 0.00 | 0.42 | 0.06 | 8 | T Cell |
| EPSTI1 | 1.52 | 0.00 | 0.00 | 0.33 | 0.03 | 8 | T Cell |
| SELL | 1.50 | 0.00 | 0.00 | 0.30 | 0.03 | 8 | T Cell |
| CD53 | 1.48 | 0.00 | 0.00 | 0.46 | 0.07 | 8 | T Cell |
| IL2RG | 1.46 | 0.00 | 0.00 | 0.45 | 0.08 | 8 | T Cell |
| SATB1 | 1.42 | 0.00 | 0.00 | 0.27 | 0.03 | 8 | T Cell |
| CYTIP | 1.37 | 0.00 | 0.00 | 0.41 | 0.08 | 8 | T Cell |
| LAPTM5 | 1.36 | 0.00 | 0.00 | 0.60 | 0.12 | 8 | T Cell |
| BIN2 | 1.36 | 0.00 | 0.00 | 0.33 | 0.04 | 8 | T Cell |
| DGKA | 1.35 | 0.00 | 0.00 | 0.29 | 0.04 | 8 | T Cell |
| PTPN18 | 1.35 | 0.00 | 0.00 | 0.50 | 0.12 | 8 | T Cell |
| ITGB7 | 1.32 | 0.00 | 0.00 | 0.30 | 0.02 | 8 | T Cell |
| TBC1D10C | 1.30 | 0.00 | 0.00 | 0.28 | 0.02 | 8 | T Cell |
| CORO1A | 1.22 | 0.00 | 0.00 | 0.41 | 0.09 | 8 | T Cell |
| ARHGAP30 | 1.19 | 0.00 | 0.00 | 0.41 | 0.09 | 8 | T Cell |
| CD48 | 1.10 | 0.00 | 0.00 | 0.27 | 0.04 | 8 | T Cell |
| WDR89 | 1.00 | 0.00 | 0.00 | 0.97 | 0.82 | 8 | T Cell |
| RPLP1 | 0.92 | 0.00 | 0.00 | 0.99 | 0.90 | 8 | T Cell |
| B2M | 0.78 | 0.00 | 0.00 | 0.97 | 0.69 | 8 | T Cell |
| GIMAP6 | 1.30 | 0.00 | 0.00 | 0.50 | 0.14 | 8 | T Cell |
| PSMB8 | 1.20 | 0.00 | 0.00 | 0.55 | 0.18 | 8 | T Cell |
| COTL1 | 1.24 | 0.00 | 0.00 | 0.52 | 0.16 | 8 | T Cell |
| APBB1IP | 1.11 | 0.00 | 0.00 | 0.37 | 0.08 | 8 | T Cell |
| STK17B | 1.24 | 0.00 | 0.00 | 0.29 | 0.05 | 8 | T Cell |
| SH3KBP1 | 1.37 | 0.00 | 0.00 | 0.40 | 0.10 | 8 | T Cell |
| TMSB10 | 0.75 | 0.00 | 0.00 | 0.99 | 0.88 | 8 | T Cell |
| TMSB4X | 0.63 | 0.00 | 0.00 | 1.00 | 0.90 | 8 | T Cell |
| WAS | 1.07 | 0.00 | 0.00 | 0.27 | 0.05 | 8 | T Cell |
| PDCD4 | 1.40 | 0.00 | 0.00 | 0.55 | 0.20 | 8 | T Cell |
| PYCARD | 1.25 | 0.00 | 0.00 | 0.37 | 0.08 | 8 | T Cell |
| SEPT6 | 1.22 | 0.00 | 0.00 | 0.33 | 0.07 | 8 | T Cell |
| RPLP2 | 0.77 | 0.00 | 0.00 | 0.96 | 0.82 | 8 | T Cell |
| RPL18A | 0.80 | 0.00 | 0.00 | 0.97 | 0.83 | 8 | T Cell |
| EEF1B2 | 0.95 | 0.00 | 0.00 | 0.86 | 0.65 | 8 | T Cell |
| RPS20 | 0.80 | 0.00 | 0.00 | 0.93 | 0.80 | 8 | T Cell |
| LRRC58 | 0.75 | 0.00 | 0.00 | 0.96 | 0.85 | 8 | T Cell |
| RPL8 | 0.73 | 0.00 | 0.00 | 0.94 | 0.81 | 8 | T Cell |
| TMEM71 | 1.15 | 0.00 | 0.00 | 0.26 | 0.05 | 8 | T Cell |
| LCP1 | 1.02 | 0.00 | 0.00 | 0.54 | 0.20 | 8 | T Cell |
| RPL10A | 0.73 | 0.00 | 0.00 | 0.90 | 0.76 | 8 | T Cell |
| ARHGAP15 | 1.10 | 0.00 | 0.00 | 0.26 | 0.05 | 8 | T Cell |
| CD37 | 0.79 | 0.00 | 0.00 | 0.29 | 0.06 | 8 | T Cell |
| ETS1 | 1.15 | 0.00 | 0.00 | 0.56 | 0.23 | 8 | T Cell |
| FAU | 0.62 | 0.00 | 0.00 | 0.96 | 0.87 | 8 | T Cell |
| RPS4X | 0.59 | 0.00 | 0.00 | 0.97 | 0.88 | 8 | T Cell |
| RPL6 | 0.60 | 0.00 | 0.00 | 0.95 | 0.88 | 8 | T Cell |
| GMFG | 1.05 | 0.00 | 0.00 | 0.42 | 0.14 | 8 | T Cell |
| LSP1 | 0.81 | 0.00 | 0.00 | 0.50 | 0.18 | 8 | T Cell |
| EMB | 1.04 | 0.00 | 0.00 | 0.33 | 0.09 | 8 | T Cell |
| JAK1 | 1.15 | 0.00 | 0.00 | 0.54 | 0.25 | 8 | T Cell |
| MBNL1 | 1.11 | 0.00 | 0.00 | 0.53 | 0.25 | 8 | T Cell |
| ACTR3 | 0.97 | 0.00 | 0.00 | 0.60 | 0.32 | 8 | T Cell |
| RAPGEF6 | 1.20 | 0.00 | 0.00 | 0.35 | 0.11 | 8 | T Cell |
| HCLS1 | 0.87 | 0.00 | 0.00 | 0.27 | 0.07 | 8 | T Cell |
| RPS13 | 0.73 | 0.00 | 0.00 | 0.84 | 0.66 | 8 | T Cell |
| GIMAP1 | 0.95 | 0.00 | 0.00 | 0.37 | 0.13 | 8 | T Cell |
| CRLF3 | 1.14 | 0.00 | 0.00 | 0.35 | 0.12 | 8 | T Cell |
| SH3BGRL3 | 0.86 | 0.00 | 0.00 | 0.62 | 0.35 | 8 | T Cell |
| DNAJC15 | 1.06 | 0.00 | 0.00 | 0.27 | 0.08 | 8 | T Cell |
| SRPK1 | 1.05 | 0.00 | 0.00 | 0.32 | 0.11 | 8 | T Cell |
| ARPC2 | 0.67 | 0.00 | 0.00 | 0.80 | 0.61 | 8 | T Cell |
| PFN1 | 0.85 | 0.00 | 0.00 | 0.69 | 0.48 | 8 | T Cell |
| RGS10 | 1.05 | 0.00 | 0.00 | 0.28 | 0.09 | 8 | T Cell |
| ATP1B3 | 0.97 | 0.00 | 0.00 | 0.50 | 0.25 | 8 | T Cell |
| ITGA4 | 0.96 | 0.00 | 0.00 | 0.28 | 0.09 | 8 | T Cell |
| LEPROTL1 | 0.99 | 0.00 | 0.00 | 0.36 | 0.15 | 8 | T Cell |
| ABLIM1 | 0.93 | 0.00 | 0.00 | 0.46 | 0.23 | 8 | T Cell |
| ORC5 | 0.74 | 0.00 | 0.00 | 0.70 | 0.51 | 8 | T Cell |
| RPL23A | 0.53 | 0.00 | 0.00 | 0.84 | 0.69 | 8 | T Cell |
| FKBP3 | 0.97 | 0.00 | 0.00 | 0.51 | 0.28 | 8 | T Cell |
| PSMB9 | 0.85 | 0.00 | 0.00 | 0.27 | 0.09 | 8 | T Cell |
| RPL7 | 0.47 | 0.00 | 0.00 | 0.88 | 0.77 | 8 | T Cell |
| RPL22L1 | 0.70 | 0.00 | 0.00 | 0.70 | 0.53 | 8 | T Cell |
| CDK2AP2 | 0.86 | 0.00 | 0.00 | 0.34 | 0.14 | 8 | T Cell |
| SERBP1 | 0.58 | 0.00 | 0.00 | 0.82 | 0.73 | 8 | T Cell |
| RPL13A | 0.40 | 0.00 | 0.00 | 0.77 | 0.58 | 8 | T Cell |
| SRGN | 0.57 | 0.00 | 0.00 | 0.59 | 0.31 | 8 | T Cell |
| S100A10 | 0.81 | 0.00 | 0.00 | 0.60 | 0.40 | 8 | T Cell |
| HMGB2 | 0.96 | 0.00 | 0.00 | 0.56 | 0.38 | 8 | T Cell |
| ARHGEF1 | 0.90 | 0.00 | 0.00 | 0.28 | 0.11 | 8 | T Cell |
| RPS27A | 0.42 | 0.00 | 0.00 | 0.87 | 0.78 | 8 | T Cell |
| ARPC1B | 0.63 | 0.00 | 0.00 | 0.56 | 0.36 | 8 | T Cell |
| DNAJC9 | 0.82 | 0.00 | 0.00 | 0.25 | 0.10 | 8 | T Cell |
| BZW2 | 0.91 | 0.00 | 0.00 | 0.30 | 0.14 | 8 | T Cell |
| HNRNPA2B1 | 0.45 | 0.00 | 0.00 | 0.81 | 0.72 | 8 | T Cell |
| STK4 | 0.87 | 0.00 | 0.00 | 0.30 | 0.13 | 8 | T Cell |
| RPL36A | 0.54 | 0.00 | 0.00 | 0.75 | 0.64 | 8 | T Cell |
| MT-ND5 | 0.46 | 0.00 | 0.00 | 0.83 | 0.75 | 8 | T Cell |
| TPM3 | 0.45 | 0.00 | 0.00 | 0.78 | 0.67 | 8 | T Cell |
| ANP32A | 0.48 | 0.00 | 0.00 | 0.72 | 0.57 | 8 | T Cell |
| RPL37A | 0.36 | 0.00 | 0.00 | 0.91 | 0.85 | 8 | T Cell |
| EEF1G | 0.64 | 0.00 | 0.00 | 0.60 | 0.46 | 8 | T Cell |
| MT-ATP6 | 0.35 | 0.00 | 0.00 | 0.90 | 0.85 | 8 | T Cell |
| CCND2 | 0.71 | 0.00 | 0.00 | 0.45 | 0.27 | 8 | T Cell |
| LDHA | 0.70 | 0.00 | 0.00 | 0.47 | 0.31 | 8 | T Cell |
| MT-ND1 | 0.31 | 0.00 | 0.00 | 0.95 | 0.92 | 8 | T Cell |
| SAMHD1 | 0.70 | 0.00 | 0.00 | 0.36 | 0.19 | 8 | T Cell |
| NT5C | 0.74 | 0.00 | 0.00 | 0.32 | 0.16 | 8 | T Cell |
| ARPC5 | 0.53 | 0.00 | 0.00 | 0.67 | 0.57 | 8 | T Cell |
| SF3B1 | 0.65 | 0.00 | 0.00 | 0.59 | 0.44 | 8 | T Cell |
| CRIP1 | 0.42 | 0.00 | 0.00 | 0.65 | 0.47 | 8 | T Cell |
| CD47 | 0.61 | 0.00 | 0.00 | 0.57 | 0.41 | 8 | T Cell |
| PSME1 | 0.65 | 0.00 | 0.00 | 0.49 | 0.33 | 8 | T Cell |
| NACA | 0.46 | 0.00 | 0.00 | 0.71 | 0.61 | 8 | T Cell |
| HNRNPF | 0.64 | 0.00 | 0.00 | 0.49 | 0.34 | 8 | T Cell |
| RAB8A | 0.69 | 0.00 | 0.00 | 0.27 | 0.13 | 8 | T Cell |
| SEC11C | 0.62 | 0.00 | 0.00 | 0.31 | 0.16 | 8 | T Cell |
| ABRACL | 0.67 | 0.00 | 0.00 | 0.37 | 0.22 | 8 | T Cell |
| KLF2 | 0.64 | 0.00 | 0.00 | 0.35 | 0.19 | 8 | T Cell |
| ARL6IP1 | 0.57 | 0.00 | 0.00 | 0.62 | 0.49 | 8 | T Cell |
| STK38 | 0.77 | 0.00 | 0.00 | 0.32 | 0.18 | 8 | T Cell |
| DDX5 | 0.44 | 0.00 | 0.00 | 0.78 | 0.71 | 8 | T Cell |
| OAZ1 | 0.44 | 0.00 | 0.00 | 0.70 | 0.63 | 8 | T Cell |
| PSME2 | 0.61 | 0.00 | 0.00 | 0.40 | 0.25 | 8 | T Cell |
| CFL1 | 0.47 | 0.00 | 0.00 | 0.70 | 0.60 | 8 | T Cell |
| EMG1 | 0.67 | 0.00 | 0.00 | 0.28 | 0.14 | 8 | T Cell |
| ENO1 | 0.60 | 0.00 | 0.00 | 0.32 | 0.18 | 8 | T Cell |
| XRN2 | 0.71 | 0.00 | 0.00 | 0.43 | 0.29 | 8 | T Cell |
| HSPA8 | 0.40 | 0.00 | 0.00 | 0.73 | 0.68 | 8 | T Cell |
| SNRPF | 0.62 | 0.00 | 0.00 | 0.42 | 0.28 | 8 | T Cell |
| NME1 | 0.63 | 0.00 | 0.00 | 0.32 | 0.19 | 8 | T Cell |
| MT-CO3 | 0.25 | 0.00 | 0.00 | 0.92 | 0.89 | 8 | T Cell |
| CDC42SE1 | 0.67 | 0.00 | 0.00 | 0.32 | 0.18 | 8 | T Cell |
| TSC22D4 | 0.60 | 0.00 | 0.00 | 0.40 | 0.25 | 8 | T Cell |
| EIF3E | 0.52 | 0.00 | 0.00 | 0.55 | 0.43 | 8 | T Cell |
| EIF3H | 0.59 | 0.00 | 0.00 | 0.48 | 0.37 | 8 | T Cell |
| RNASET2 | 0.59 | 0.00 | 0.00 | 0.38 | 0.25 | 8 | T Cell |
| SLC25A5 | 0.44 | 0.00 | 0.00 | 0.65 | 0.58 | 8 | T Cell |
| SNRPD3 | 0.60 | 0.00 | 0.00 | 0.43 | 0.31 | 8 | T Cell |
| NOP10 | 0.61 | 0.00 | 0.00 | 0.41 | 0.29 | 8 | T Cell |
| SMCHD1 | 0.66 | 0.00 | 0.00 | 0.26 | 0.14 | 8 | T Cell |
| RBM3 | 0.53 | 0.00 | 0.00 | 0.51 | 0.41 | 8 | T Cell |
| FGFR1OP2 | 0.60 | 0.00 | 0.00 | 0.38 | 0.26 | 8 | T Cell |
| NOP58 | 0.60 | 0.00 | 0.00 | 0.36 | 0.24 | 8 | T Cell |
| EEF2 | 0.41 | 0.00 | 0.00 | 0.67 | 0.59 | 8 | T Cell |
| HSP90AB1 | 0.28 | 0.00 | 0.00 | 0.89 | 0.89 | 8 | T Cell |
| EMP3 | 0.49 | 0.00 | 0.00 | 0.37 | 0.24 | 8 | T Cell |
| AKAP13 | 0.59 | 0.00 | 0.00 | 0.38 | 0.26 | 8 | T Cell |
| ADD3 | 0.54 | 0.00 | 0.00 | 0.40 | 0.27 | 8 | T Cell |
| TP53 | 0.59 | 0.00 | 0.00 | 0.26 | 0.15 | 8 | T Cell |
| H2AFZ | 0.63 | 0.00 | 0.00 | 0.49 | 0.41 | 8 | T Cell |
| DUSP11 | 0.59 | 0.00 | 0.00 | 0.36 | 0.24 | 8 | T Cell |
| EZR | 0.48 | 0.00 | 0.00 | 0.33 | 0.21 | 8 | T Cell |
| SRRM2 | 0.41 | 0.00 | 0.00 | 0.65 | 0.59 | 8 | T Cell |
| PPP1CA | 0.52 | 0.00 | 0.00 | 0.44 | 0.33 | 8 | T Cell |
| DDX21 | 0.61 | 0.00 | 0.00 | 0.37 | 0.26 | 8 | T Cell |
| CNN2 | 0.41 | 0.00 | 0.00 | 0.51 | 0.38 | 8 | T Cell |
| EIF3M | 0.54 | 0.00 | 0.00 | 0.36 | 0.25 | 8 | T Cell |
| HNRNPCL1 | 0.46 | 0.00 | 0.00 | 0.50 | 0.40 | 8 | T Cell |
| NOP56 | 0.60 | 0.00 | 0.00 | 0.30 | 0.19 | 8 | T Cell |
| HNRNPU | 0.37 | 0.00 | 0.00 | 0.68 | 0.62 | 8 | T Cell |
| IFNGR1 | 0.62 | 0.00 | 0.00 | 0.34 | 0.23 | 8 | T Cell |
| RPL17-C18orf32 | 0.58 | 0.00 | 0.00 | 0.41 | 0.30 | 8 | T Cell |
| LSM4 | 0.56 | 0.00 | 0.00 | 0.31 | 0.20 | 8 | T Cell |
| SUPT4H1 | 0.57 | 0.00 | 0.00 | 0.27 | 0.17 | 8 | T Cell |
| MDH1 | 0.54 | 0.00 | 0.00 | 0.29 | 0.18 | 8 | T Cell |
| CDKN1B | 0.65 | 0.00 | 0.00 | 0.29 | 0.19 | 8 | T Cell |
| ANP32E | 0.55 | 0.00 | 0.00 | 0.36 | 0.26 | 8 | T Cell |
| MSN | 0.40 | 0.00 | 0.00 | 0.55 | 0.44 | 8 | T Cell |
| PCNA | 0.63 | 0.00 | 0.00 | 0.26 | 0.16 | 8 | T Cell |
| G3BP2 | 0.46 | 0.00 | 0.00 | 0.39 | 0.29 | 8 | T Cell |
| THRAP3 | 0.52 | 0.00 | 0.00 | 0.44 | 0.36 | 8 | T Cell |
| COX7A2L | 0.47 | 0.00 | 0.00 | 0.50 | 0.42 | 8 | T Cell |
| HSPE1 | 0.54 | 0.00 | 0.00 | 0.39 | 0.30 | 8 | T Cell |
| PRRC2C | 0.36 | 0.00 | 0.00 | 0.66 | 0.61 | 8 | T Cell |
| POLR1D | 0.51 | 0.00 | 0.00 | 0.37 | 0.28 | 8 | T Cell |
| ACIN1 | 0.41 | 0.00 | 0.00 | 0.59 | 0.51 | 8 | T Cell |
| MIF | 0.48 | 0.00 | 0.00 | 0.33 | 0.23 | 8 | T Cell |
| PPP2R5C | 0.50 | 0.00 | 0.00 | 0.29 | 0.19 | 8 | T Cell |
| RAN | 0.46 | 0.00 | 0.00 | 0.46 | 0.39 | 8 | T Cell |
| NCOR1 | 0.47 | 0.00 | 0.00 | 0.47 | 0.40 | 8 | T Cell |
| DNAJC2 | 0.52 | 0.00 | 0.00 | 0.27 | 0.19 | 8 | T Cell |
| LRRFIP1 | 0.47 | 0.00 | 0.00 | 0.35 | 0.25 | 8 | T Cell |
| ANP32B | 0.41 | 0.00 | 0.00 | 0.53 | 0.48 | 8 | T Cell |
| PKM | 0.50 | 0.00 | 0.00 | 0.33 | 0.24 | 8 | T Cell |
| PSMB1 | 0.40 | 0.00 | 0.00 | 0.53 | 0.48 | 8 | T Cell |
| GCLM | 0.41 | 0.00 | 0.00 | 0.46 | 0.38 | 8 | T Cell |
| DDX46 | 0.46 | 0.00 | 0.00 | 0.38 | 0.30 | 8 | T Cell |
| LLPH | 0.40 | 0.00 | 0.00 | 0.41 | 0.31 | 8 | T Cell |
| TSPO | 0.45 | 0.00 | 0.00 | 0.35 | 0.27 | 8 | T Cell |
| EIF3I | 0.46 | 0.00 | 0.00 | 0.33 | 0.24 | 8 | T Cell |
| ACTR2 | 0.41 | 0.00 | 0.00 | 0.36 | 0.27 | 8 | T Cell |
| PAK2 | 0.45 | 0.00 | 0.00 | 0.36 | 0.28 | 8 | T Cell |
| UQCRH | 0.37 | 0.00 | 0.00 | 0.54 | 0.48 | 8 | T Cell |
| EIF3K | 0.44 | 0.00 | 0.00 | 0.43 | 0.36 | 8 | T Cell |
| MYH9 | 0.40 | 0.00 | 0.00 | 0.42 | 0.34 | 8 | T Cell |
| SNRPE | 0.39 | 0.00 | 0.00 | 0.44 | 0.37 | 8 | T Cell |
| PSMD14 | 0.45 | 0.00 | 0.00 | 0.25 | 0.17 | 8 | T Cell |
| RSL1D1 | 0.49 | 0.00 | 0.00 | 0.35 | 0.27 | 8 | T Cell |
| EIF5A | 0.39 | 0.00 | 0.00 | 0.41 | 0.33 | 8 | T Cell |
| TOP2B | 0.47 | 0.00 | 0.00 | 0.33 | 0.25 | 8 | T Cell |
| BTF3 | 0.31 | 0.00 | 0.00 | 0.59 | 0.54 | 8 | T Cell |
| NOL7 | 0.37 | 0.00 | 0.00 | 0.43 | 0.35 | 8 | T Cell |
| EIF3F | 0.43 | 0.00 | 0.00 | 0.44 | 0.38 | 8 | T Cell |
| HNRNPM | 0.34 | 0.00 | 0.00 | 0.53 | 0.49 | 8 | T Cell |
| CNBP | 0.38 | 0.00 | 0.00 | 0.51 | 0.47 | 8 | T Cell |
| EIF3CL | 0.34 | 0.00 | 0.00 | 0.49 | 0.43 | 8 | T Cell |
| OSTF1 | 0.39 | 0.00 | 0.00 | 0.33 | 0.25 | 8 | T Cell |
| CCT2 | 0.45 | 0.00 | 0.00 | 0.33 | 0.26 | 8 | T Cell |
| TMEM50A | 0.37 | 0.00 | 0.00 | 0.50 | 0.44 | 8 | T Cell |
| SUB1 | 0.29 | 0.00 | 0.00 | 0.56 | 0.52 | 8 | T Cell |
| RSBN1L | 0.50 | 0.00 | 0.00 | 0.31 | 0.24 | 8 | T Cell |
| PNN | 0.31 | 0.00 | 0.00 | 0.56 | 0.51 | 8 | T Cell |
| ITSN2 | 0.42 | 0.00 | 0.00 | 0.31 | 0.23 | 8 | T Cell |
| PTBP3 | 0.36 | 0.00 | 0.00 | 0.40 | 0.33 | 8 | T Cell |
| S1PR1 | 0.36 | 0.00 | 0.00 | 0.26 | 0.19 | 8 | T Cell |
| SF3B2 | 0.36 | 0.00 | 0.00 | 0.44 | 0.38 | 8 | T Cell |
| H2AFY | 0.47 | 0.00 | 0.00 | 0.27 | 0.20 | 8 | T Cell |
| NAP1L4 | 0.44 | 0.00 | 0.00 | 0.32 | 0.25 | 8 | T Cell |
| CCT8 | 0.41 | 0.00 | 0.00 | 0.28 | 0.21 | 8 | T Cell |
| KHDRBS1 | 0.39 | 0.00 | 0.00 | 0.37 | 0.31 | 8 | T Cell |
| MRPL18 | 0.38 | 0.00 | 0.00 | 0.26 | 0.20 | 8 | T Cell |
| CAPZB | 0.32 | 0.00 | 0.00 | 0.48 | 0.44 | 8 | T Cell |
| RRP1 | 0.39 | 0.00 | 0.00 | 0.40 | 0.34 | 8 | T Cell |
| SSRP1 | 0.40 | 0.00 | 0.00 | 0.26 | 0.20 | 8 | T Cell |
| PA2G4 | 0.42 | 0.00 | 0.00 | 0.32 | 0.26 | 8 | T Cell |
| CELF2 | 0.30 | 0.00 | 0.00 | 0.34 | 0.27 | 8 | T Cell |
| PTP4A2 | 0.39 | 0.00 | 0.00 | 0.46 | 0.41 | 8 | T Cell |
| BANF1 | 0.37 | 0.00 | 0.00 | 0.35 | 0.29 | 8 | T Cell |
| NASP | 0.40 | 0.00 | 0.00 | 0.25 | 0.19 | 8 | T Cell |
| VASP | 0.42 | 0.00 | 0.00 | 0.26 | 0.20 | 8 | T Cell |
| CTSD | 0.30 | 0.00 | 0.00 | 0.31 | 0.25 | 8 | T Cell |
| UQCRC2 | 0.40 | 0.00 | 0.00 | 0.28 | 0.22 | 8 | T Cell |
| MYCBP2 | 0.39 | 0.00 | 0.00 | 0.28 | 0.22 | 8 | T Cell |
| YWHAZ | 0.26 | 0.00 | 0.00 | 0.53 | 0.50 | 8 | T Cell |
| TMA7 | 0.33 | 0.00 | 0.00 | 0.40 | 0.35 | 8 | T Cell |
| NDUFV3 | 0.36 | 0.00 | 0.00 | 0.29 | 0.23 | 8 | T Cell |
| SPCS2 | 0.35 | 0.00 | 0.00 | 0.38 | 0.33 | 8 | T Cell |
| TRA2B | 0.38 | 0.00 | 0.00 | 0.35 | 0.30 | 8 | T Cell |
| BTG1 | 0.42 | 0.00 | 0.00 | 0.35 | 0.31 | 8 | T Cell |
| SUPT16H | 0.37 | 0.00 | 0.00 | 0.28 | 0.23 | 8 | T Cell |
| PRELID1 | 0.34 | 0.00 | 0.00 | 0.35 | 0.30 | 8 | T Cell |
| DNAJA2 | 0.38 | 0.00 | 0.00 | 0.29 | 0.24 | 8 | T Cell |
| TOMM20 | 0.35 | 0.00 | 0.00 | 0.31 | 0.26 | 8 | T Cell |
| CCT4 | 0.39 | 0.00 | 0.00 | 0.27 | 0.22 | 8 | T Cell |
| ATP6V1F | 0.35 | 0.00 | 0.00 | 0.31 | 0.26 | 8 | T Cell |
| TMEM234 | 0.34 | 0.00 | 0.00 | 0.37 | 0.33 | 8 | T Cell |
| HSPD1 | 0.39 | 0.00 | 0.00 | 0.31 | 0.26 | 8 | T Cell |
| CCNL1 | 0.32 | 0.00 | 0.00 | 0.37 | 0.32 | 8 | T Cell |
| CSNK2A3 | 0.35 | 0.00 | 0.00 | 0.33 | 0.28 | 8 | T Cell |
| NSA2 | 0.31 | 0.00 | 0.00 | 0.35 | 0.30 | 8 | T Cell |
| CDC37 | 0.33 | 0.00 | 0.00 | 0.34 | 0.29 | 8 | T Cell |
| PHF20L1 | 0.36 | 0.00 | 0.00 | 0.31 | 0.26 | 8 | T Cell |
| CTCF | 0.33 | 0.00 | 0.00 | 0.26 | 0.21 | 8 | T Cell |
| SRI | 0.35 | 0.00 | 0.00 | 0.32 | 0.28 | 8 | T Cell |
| MT-ATP8 | 0.35 | 0.00 | 0.00 | 0.30 | 0.25 | 8 | T Cell |
| BZW1 | 0.29 | 0.00 | 0.00 | 0.37 | 0.33 | 8 | T Cell |
| CCT5 | 0.29 | 0.00 | 0.00 | 0.28 | 0.23 | 8 | T Cell |
| SLTM | 0.29 | 0.00 | 0.01 | 0.42 | 0.39 | 8 | T Cell |
| VPS28 | 0.34 | 0.00 | 0.01 | 0.25 | 0.21 | 8 | T Cell |
| NONO | 0.34 | 0.00 | 0.01 | 0.26 | 0.22 | 8 | T Cell |
| MIER1 | 0.35 | 0.00 | 0.02 | 0.27 | 0.22 | 8 | T Cell |
| ITGAM | 0.26 | 0.00 | 0.02 | 0.30 | 0.25 | 8 | T Cell |
| SRSF5 | 0.27 | 0.00 | 0.03 | 0.51 | 0.51 | 8 | T Cell |
| CDC5L | 0.32 | 0.00 | 0.04 | 0.25 | 0.21 | 8 | T Cell |
| FBLN2 | 2.47 | 0.00 | 0.00 | 0.61 | 0.05 | 9 | Endothelial |
| VWF | 2.41 | 0.00 | 0.00 | 0.64 | 0.02 | 9 | Endothelial |
| PTPRB | 2.07 | 0.00 | 0.00 | 0.90 | 0.19 | 9 | Endothelial |
| FBLN5 | 1.97 | 0.00 | 0.00 | 0.66 | 0.17 | 9 | Endothelial |
| TM4SF1 | 1.72 | 0.00 | 0.00 | 0.80 | 0.21 | 9 | Endothelial |
| EDN1 | 1.53 | 0.00 | 0.00 | 0.53 | 0.09 | 9 | Endothelial |
| STMN2 | 1.49 | 0.00 | 0.00 | 0.58 | 0.13 | 9 | Endothelial |
| GJA4 | 1.48 | 0.00 | 0.00 | 0.40 | 0.07 | 9 | Endothelial |
| PRSS23 | 1.44 | 0.00 | 0.00 | 0.41 | 0.07 | 9 | Endothelial |
| CPE | 1.40 | 0.00 | 0.00 | 0.27 | 0.03 | 9 | Endothelial |
| ACKR3 | 1.39 | 0.00 | 0.00 | 0.33 | 0.04 | 9 | Endothelial |
| HEG1 | 1.39 | 0.00 | 0.00 | 0.63 | 0.15 | 9 | Endothelial |
| JAM2 | 1.35 | 0.00 | 0.00 | 0.42 | 0.06 | 9 | Endothelial |
| PLVAP | 1.35 | 0.00 | 0.00 | 0.53 | 0.11 | 9 | Endothelial |
| CLEC14A | 1.30 | 0.00 | 0.00 | 0.71 | 0.17 | 9 | Endothelial |
| CALCRL | 1.28 | 0.00 | 0.00 | 0.85 | 0.24 | 9 | Endothelial |
| TEK | 1.27 | 0.00 | 0.00 | 0.56 | 0.11 | 9 | Endothelial |
| LYVE1 | 1.24 | 0.00 | 0.00 | 0.70 | 0.20 | 9 | Endothelial |
| MMRN2 | 1.23 | 0.00 | 0.00 | 0.36 | 0.04 | 9 | Endothelial |
| FLT1 | 1.19 | 0.00 | 0.00 | 0.58 | 0.14 | 9 | Endothelial |
| PECAM1 | 1.14 | 0.00 | 0.00 | 0.79 | 0.24 | 9 | Endothelial |
| CD93 | 1.05 | 0.00 | 0.00 | 0.73 | 0.21 | 9 | Endothelial |
| RAMP2 | 1.03 | 0.00 | 0.00 | 0.84 | 0.28 | 9 | Endothelial |
| EGFL7 | 0.93 | 0.00 | 0.00 | 0.85 | 0.27 | 9 | Endothelial |
| CXCL12 | 2.52 | 0.00 | 0.00 | 0.49 | 0.12 | 9 | Endothelial |
| APLNR | 1.26 | 0.00 | 0.00 | 0.37 | 0.07 | 9 | Endothelial |
| PALMD | 1.10 | 0.00 | 0.00 | 0.50 | 0.12 | 9 | Endothelial |
| ADAM15 | 1.04 | 0.00 | 0.00 | 0.39 | 0.08 | 9 | Endothelial |
| ADCY4 | 1.06 | 0.00 | 0.00 | 0.39 | 0.08 | 9 | Endothelial |
| TSPAN7 | 0.92 | 0.00 | 0.00 | 0.66 | 0.20 | 9 | Endothelial |
| CLDN5 | 0.87 | 0.00 | 0.00 | 0.69 | 0.23 | 9 | Endothelial |
| EFNB2 | 1.32 | 0.00 | 0.00 | 0.49 | 0.14 | 9 | Endothelial |
| S100A16 | 0.93 | 0.00 | 0.00 | 0.68 | 0.24 | 9 | Endothelial |
| AQP1 | 1.05 | 0.00 | 0.00 | 0.60 | 0.20 | 9 | Endothelial |
| CDH5 | 0.73 | 0.00 | 0.00 | 0.70 | 0.23 | 9 | Endothelial |
| SEMA3G | 1.20 | 0.00 | 0.00 | 0.32 | 0.06 | 9 | Endothelial |
| LTBP4 | 1.12 | 0.00 | 0.00 | 0.42 | 0.11 | 9 | Endothelial |
| ECE1 | 1.01 | 0.00 | 0.00 | 0.63 | 0.24 | 9 | Endothelial |
| SRGN | 0.87 | 0.00 | 0.00 | 0.74 | 0.31 | 9 | Endothelial |
| TIE1 | 0.90 | 0.00 | 0.00 | 0.42 | 0.11 | 9 | Endothelial |
| HYAL2 | 1.06 | 0.00 | 0.00 | 0.35 | 0.09 | 9 | Endothelial |
| SHE | 0.85 | 0.00 | 0.00 | 0.27 | 0.05 | 9 | Endothelial |
| CD9 | 0.86 | 0.00 | 0.00 | 0.73 | 0.35 | 9 | Endothelial |
| CD200 | 0.88 | 0.00 | 0.00 | 0.62 | 0.26 | 9 | Endothelial |
| SOX18 | 0.79 | 0.00 | 0.00 | 0.25 | 0.05 | 9 | Endothelial |
| EPAS1 | 0.76 | 0.00 | 0.00 | 0.67 | 0.27 | 9 | Endothelial |
| ENTPD1 | 0.94 | 0.00 | 0.00 | 0.33 | 0.08 | 9 | Endothelial |
| EHD4 | 0.84 | 0.00 | 0.00 | 0.62 | 0.26 | 9 | Endothelial |
| IGF2 | 1.38 | 0.00 | 0.00 | 0.27 | 0.06 | 9 | Endothelial |
| MEIS2 | 0.85 | 0.00 | 0.00 | 0.38 | 0.11 | 9 | Endothelial |
| MYCT1 | 0.78 | 0.00 | 0.00 | 0.41 | 0.12 | 9 | Endothelial |
| FSTL1 | 0.77 | 0.00 | 0.00 | 0.79 | 0.45 | 9 | Endothelial |
| SOX17 | 0.92 | 0.00 | 0.00 | 0.42 | 0.13 | 9 | Endothelial |
| EPHB4 | 0.81 | 0.00 | 0.00 | 0.30 | 0.07 | 9 | Endothelial |
| ECSCR | 0.70 | 0.00 | 0.00 | 0.46 | 0.15 | 9 | Endothelial |
| ICAM2 | 0.73 | 0.00 | 0.00 | 0.48 | 0.17 | 9 | Endothelial |
| PLK2 | 0.99 | 0.00 | 0.00 | 0.45 | 0.16 | 9 | Endothelial |
| CAV1 | 0.74 | 0.00 | 0.00 | 0.74 | 0.36 | 9 | Endothelial |
| ACER2 | 0.92 | 0.00 | 0.00 | 0.32 | 0.09 | 9 | Endothelial |
| RASIP1 | 0.77 | 0.00 | 0.00 | 0.38 | 0.12 | 9 | Endothelial |
| PODXL | 0.79 | 0.00 | 0.00 | 0.36 | 0.10 | 9 | Endothelial |
| TMEM88 | 0.83 | 0.00 | 0.00 | 0.35 | 0.10 | 9 | Endothelial |
| SLC43A3 | 0.89 | 0.00 | 0.00 | 0.49 | 0.18 | 9 | Endothelial |
| TGM2 | 0.83 | 0.00 | 0.00 | 0.34 | 0.10 | 9 | Endothelial |
| PCSK5 | 1.04 | 0.00 | 0.00 | 0.31 | 0.09 | 9 | Endothelial |
| SLC9A3R2 | 0.90 | 0.00 | 0.00 | 0.54 | 0.22 | 9 | Endothelial |
| CYYR1 | 0.73 | 0.00 | 0.00 | 0.42 | 0.14 | 9 | Endothelial |
| ESAM | 0.65 | 0.00 | 0.00 | 0.51 | 0.20 | 9 | Endothelial |
| RNF144A | 0.88 | 0.00 | 0.00 | 0.40 | 0.14 | 9 | Endothelial |
| GATA2 | 0.69 | 0.00 | 0.00 | 0.40 | 0.13 | 9 | Endothelial |
| CSRP2 | 0.89 | 0.00 | 0.00 | 0.41 | 0.15 | 9 | Endothelial |
| ACVRL1 | 0.68 | 0.00 | 0.00 | 0.47 | 0.18 | 9 | Endothelial |
| TNFSF10 | 0.74 | 0.00 | 0.00 | 0.32 | 0.10 | 9 | Endothelial |
| NOTCH1 | 0.79 | 0.00 | 0.00 | 0.36 | 0.12 | 9 | Endothelial |
| LDB2 | 0.76 | 0.00 | 0.00 | 0.44 | 0.17 | 9 | Endothelial |
| PLXND1 | 0.77 | 0.00 | 0.00 | 0.27 | 0.08 | 9 | Endothelial |
| RHOJ | 0.73 | 0.00 | 0.00 | 0.44 | 0.17 | 9 | Endothelial |
| RIPPLY3 | 0.97 | 0.00 | 0.00 | 0.34 | 0.11 | 9 | Endothelial |
| PRCP | 0.77 | 0.00 | 0.00 | 0.32 | 0.10 | 9 | Endothelial |
| FOXO1 | 0.74 | 0.00 | 0.00 | 0.25 | 0.07 | 9 | Endothelial |
| IGFBP4 | 0.90 | 0.00 | 0.00 | 0.56 | 0.27 | 9 | Endothelial |
| FKBP1A | 0.69 | 0.00 | 0.00 | 0.72 | 0.43 | 9 | Endothelial |
| HOXA5 | 0.77 | 0.00 | 0.00 | 0.37 | 0.13 | 9 | Endothelial |
| SLC6A6 | 0.97 | 0.00 | 0.00 | 0.50 | 0.23 | 9 | Endothelial |
| KLF2 | 0.72 | 0.00 | 0.00 | 0.47 | 0.19 | 9 | Endothelial |
| GATA6 | 0.84 | 0.00 | 0.00 | 0.26 | 0.08 | 9 | Endothelial |
| CAV2 | 0.62 | 0.00 | 0.00 | 0.50 | 0.22 | 9 | Endothelial |
| COL5A2 | 1.01 | 0.00 | 0.00 | 0.42 | 0.18 | 9 | Endothelial |
| EMCN | 0.58 | 0.00 | 0.00 | 0.34 | 0.11 | 9 | Endothelial |
| CRIM1 | 0.77 | 0.00 | 0.00 | 0.26 | 0.08 | 9 | Endothelial |
| ELK3 | 0.70 | 0.00 | 0.00 | 0.37 | 0.14 | 9 | Endothelial |
| RAB11A | 0.70 | 0.00 | 0.00 | 0.55 | 0.27 | 9 | Endothelial |
| SMAD7 | 0.77 | 0.00 | 0.00 | 0.31 | 0.11 | 9 | Endothelial |
| TMEM100 | 0.62 | 0.00 | 0.00 | 0.60 | 0.29 | 9 | Endothelial |
| ENG | 0.77 | 0.00 | 0.00 | 0.35 | 0.13 | 9 | Endothelial |
| PLCB4 | 0.80 | 0.00 | 0.00 | 0.32 | 0.12 | 9 | Endothelial |
| KLF4 | 0.61 | 0.00 | 0.00 | 0.49 | 0.23 | 9 | Endothelial |
| CBFA2T3 | 0.73 | 0.00 | 0.00 | 0.35 | 0.13 | 9 | Endothelial |
| THBD | 0.67 | 0.00 | 0.00 | 0.51 | 0.25 | 9 | Endothelial |
| MECOM | 0.89 | 0.00 | 0.00 | 0.27 | 0.09 | 9 | Endothelial |
| CRIP2 | 0.54 | 0.00 | 0.00 | 0.71 | 0.43 | 9 | Endothelial |
| MYZAP | 0.58 | 0.00 | 0.00 | 0.36 | 0.14 | 9 | Endothelial |
| UTRN | 0.79 | 0.00 | 0.00 | 0.37 | 0.16 | 9 | Endothelial |
| KIAA0355 | 0.63 | 0.00 | 0.00 | 0.40 | 0.17 | 9 | Endothelial |
| MCAM | 0.57 | 0.00 | 0.00 | 0.34 | 0.13 | 9 | Endothelial |
| ITGB1 | 0.55 | 0.00 | 0.00 | 0.83 | 0.64 | 9 | Endothelial |
| GNG11 | 0.49 | 0.00 | 0.00 | 0.60 | 0.31 | 9 | Endothelial |
| SLK | 0.62 | 0.00 | 0.00 | 0.65 | 0.40 | 9 | Endothelial |
| ARMCX1 | 0.72 | 0.00 | 0.00 | 0.27 | 0.10 | 9 | Endothelial |
| ITGA6 | 0.63 | 0.00 | 0.00 | 0.32 | 0.12 | 9 | Endothelial |
| LUZP1 | 0.62 | 0.00 | 0.00 | 0.41 | 0.19 | 9 | Endothelial |
| S1PR1 | 0.49 | 0.00 | 0.00 | 0.41 | 0.18 | 9 | Endothelial |
| TSPAN18 | 0.51 | 0.00 | 0.00 | 0.33 | 0.13 | 9 | Endothelial |
| SWAP70 | 0.59 | 0.00 | 0.00 | 0.27 | 0.10 | 9 | Endothelial |
| TSPAN13 | 0.35 | 0.00 | 0.00 | 0.50 | 0.24 | 9 | Endothelial |
| ELN | 0.62 | 0.00 | 0.00 | 0.39 | 0.18 | 9 | Endothelial |
| GIMAP1 | 0.45 | 0.00 | 0.00 | 0.32 | 0.13 | 9 | Endothelial |
| CD151 | 0.56 | 0.00 | 0.00 | 0.39 | 0.19 | 9 | Endothelial |
| EHD2 | 0.56 | 0.00 | 0.00 | 0.35 | 0.16 | 9 | Endothelial |
| NRP1 | 0.54 | 0.00 | 0.00 | 0.61 | 0.39 | 9 | Endothelial |
| APLP2 | 0.56 | 0.00 | 0.00 | 0.62 | 0.40 | 9 | Endothelial |
| HIP1 | 0.54 | 0.00 | 0.00 | 0.33 | 0.15 | 9 | Endothelial |
| FNBP1L | 0.50 | 0.00 | 0.00 | 0.45 | 0.24 | 9 | Endothelial |
| KANK3 | 0.49 | 0.00 | 0.00 | 0.31 | 0.13 | 9 | Endothelial |
| HMCN1 | 0.56 | 0.00 | 0.00 | 0.30 | 0.13 | 9 | Endothelial |
| RAPGEF5 | 0.51 | 0.00 | 0.00 | 0.26 | 0.11 | 9 | Endothelial |
| ITM2B | 0.31 | 0.00 | 0.00 | 0.90 | 0.77 | 9 | Endothelial |
| PEA15 | 0.55 | 0.00 | 0.00 | 0.32 | 0.15 | 9 | Endothelial |
| BST2 | 0.63 | 0.00 | 0.00 | 0.33 | 0.15 | 9 | Endothelial |
| NAV1 | 0.54 | 0.00 | 0.00 | 0.25 | 0.10 | 9 | Endothelial |
| ANXA2 | 0.47 | 0.00 | 0.00 | 0.68 | 0.49 | 9 | Endothelial |
| TRIB2 | 0.52 | 0.00 | 0.00 | 0.25 | 0.10 | 9 | Endothelial |
| TCF4 | 0.47 | 0.00 | 0.00 | 0.64 | 0.41 | 9 | Endothelial |
| FLI1 | 0.47 | 0.00 | 0.00 | 0.27 | 0.11 | 9 | Endothelial |
| SPTBN1 | 0.39 | 0.00 | 0.00 | 0.76 | 0.53 | 9 | Endothelial |
| AZIN1 | 0.67 | 0.00 | 0.00 | 0.27 | 0.12 | 9 | Endothelial |
| ACTN4 | 0.49 | 0.00 | 0.00 | 0.50 | 0.29 | 9 | Endothelial |
| CCDC85B | 0.51 | 0.00 | 0.00 | 0.30 | 0.14 | 9 | Endothelial |
| CD2AP | 0.61 | 0.00 | 0.00 | 0.50 | 0.31 | 9 | Endothelial |
| AGRN | 0.44 | 0.00 | 0.00 | 0.28 | 0.12 | 9 | Endothelial |
| SEMA6A | 0.41 | 0.00 | 0.00 | 0.27 | 0.12 | 9 | Endothelial |
| NCK1 | 0.50 | 0.00 | 0.00 | 0.28 | 0.13 | 9 | Endothelial |
| MAPK3 | 0.46 | 0.00 | 0.00 | 0.29 | 0.13 | 9 | Endothelial |
| C1orf54 | 0.36 | 0.00 | 0.00 | 0.52 | 0.29 | 9 | Endothelial |
| UACA | 0.51 | 0.00 | 0.00 | 0.30 | 0.14 | 9 | Endothelial |
| SOX4 | 0.55 | 0.00 | 0.00 | 0.60 | 0.40 | 9 | Endothelial |
| PLEC | 0.47 | 0.00 | 0.00 | 0.27 | 0.12 | 9 | Endothelial |
| HPGD | 0.35 | 0.00 | 0.00 | 0.47 | 0.26 | 9 | Endothelial |
| DDAH2 | 0.41 | 0.00 | 0.00 | 0.48 | 0.28 | 9 | Endothelial |
| RHOC | 0.50 | 0.00 | 0.00 | 0.42 | 0.24 | 9 | Endothelial |
| PDLIM1 | 0.51 | 0.00 | 0.00 | 0.31 | 0.15 | 9 | Endothelial |
| HDAC7 | 0.46 | 0.00 | 0.00 | 0.29 | 0.14 | 9 | Endothelial |
| AMD1 | 0.89 | 0.00 | 0.00 | 0.33 | 0.17 | 9 | Endothelial |
| HSP90AB1 | 0.27 | 0.00 | 0.00 | 0.94 | 0.89 | 9 | Endothelial |
| DST | 0.54 | 0.00 | 0.00 | 0.31 | 0.16 | 9 | Endothelial |
| VAMP5 | 0.49 | 0.00 | 0.00 | 0.27 | 0.13 | 9 | Endothelial |
| MARCKS | 0.32 | 0.00 | 0.00 | 0.70 | 0.49 | 9 | Endothelial |
| ANXA3 | 0.34 | 0.00 | 0.00 | 0.40 | 0.22 | 9 | Endothelial |
| TMEM109 | 0.44 | 0.00 | 0.00 | 0.27 | 0.13 | 9 | Endothelial |
| MARCKSL1 | 0.50 | 0.00 | 0.00 | 0.39 | 0.23 | 9 | Endothelial |
| CMIP | 0.49 | 0.00 | 0.00 | 0.33 | 0.18 | 9 | Endothelial |
| SKAP2 | 0.40 | 0.00 | 0.00 | 0.27 | 0.13 | 9 | Endothelial |
| MAP7D1 | 0.40 | 0.00 | 0.00 | 0.33 | 0.18 | 9 | Endothelial |
| RASGRP3 | 0.38 | 0.00 | 0.00 | 0.25 | 0.12 | 9 | Endothelial |
| TMOD3 | 0.46 | 0.00 | 0.00 | 0.42 | 0.26 | 9 | Endothelial |
| FAM3C | 0.81 | 0.00 | 0.00 | 0.26 | 0.14 | 9 | Endothelial |
| EDNRB | 0.28 | 0.00 | 0.00 | 0.30 | 0.16 | 9 | Endothelial |
| CCDC50 | 0.42 | 0.00 | 0.00 | 0.41 | 0.25 | 9 | Endothelial |
| ZMIZ1 | 0.45 | 0.00 | 0.00 | 0.35 | 0.20 | 9 | Endothelial |
| ADAM10 | 0.53 | 0.00 | 0.00 | 0.32 | 0.18 | 9 | Endothelial |
| CTNNA1 | 0.37 | 0.00 | 0.00 | 0.53 | 0.36 | 9 | Endothelial |
| CLIC1 | 0.33 | 0.00 | 0.00 | 0.56 | 0.39 | 9 | Endothelial |
| KLHL5 | 0.47 | 0.00 | 0.00 | 0.27 | 0.14 | 9 | Endothelial |
| TSC22D1 | 0.41 | 0.00 | 0.00 | 0.51 | 0.35 | 9 | Endothelial |
| ATP2B1 | 0.42 | 0.00 | 0.00 | 0.69 | 0.55 | 9 | Endothelial |
| ARHGAP29 | 0.34 | 0.00 | 0.00 | 0.39 | 0.23 | 9 | Endothelial |
| YBX3 | 0.40 | 0.00 | 0.00 | 0.33 | 0.19 | 9 | Endothelial |
| SPAG9 | 0.45 | 0.00 | 0.00 | 0.29 | 0.16 | 9 | Endothelial |
| SNX3 | 0.38 | 0.00 | 0.00 | 0.51 | 0.35 | 9 | Endothelial |
| TJP1 | 0.39 | 0.00 | 0.00 | 0.31 | 0.17 | 9 | Endothelial |
| RALB | 0.35 | 0.00 | 0.00 | 0.34 | 0.19 | 9 | Endothelial |
| PHACTR2 | 0.38 | 0.00 | 0.00 | 0.26 | 0.14 | 9 | Endothelial |
| ATP1B3 | 0.39 | 0.00 | 0.00 | 0.41 | 0.26 | 9 | Endothelial |
| LIMA1 | 0.41 | 0.00 | 0.00 | 0.27 | 0.15 | 9 | Endothelial |
| CNN3 | 0.32 | 0.00 | 0.00 | 0.51 | 0.32 | 9 | Endothelial |
| GPIHBP1 | 0.30 | 0.00 | 0.00 | 0.29 | 0.16 | 9 | Endothelial |
| CLIC4 | 0.36 | 0.00 | 0.00 | 0.45 | 0.30 | 9 | Endothelial |
| BNIP2 | 0.38 | 0.00 | 0.00 | 0.44 | 0.28 | 9 | Endothelial |
| NFE2L1 | 0.40 | 0.00 | 0.00 | 0.35 | 0.21 | 9 | Endothelial |
| PPFIBP1 | 0.40 | 0.00 | 0.00 | 0.26 | 0.14 | 9 | Endothelial |
| GCC2 | 0.46 | 0.00 | 0.00 | 0.38 | 0.24 | 9 | Endothelial |
| PREX2 | 0.42 | 0.00 | 0.00 | 0.29 | 0.17 | 9 | Endothelial |
| S100A10 | 0.36 | 0.00 | 0.00 | 0.56 | 0.41 | 9 | Endothelial |
| CCSER2 | 0.36 | 0.00 | 0.00 | 0.26 | 0.14 | 9 | Endothelial |
| AMOTL1 | 0.32 | 0.00 | 0.00 | 0.27 | 0.15 | 9 | Endothelial |
| CARHSP1 | 0.37 | 0.00 | 0.00 | 0.27 | 0.15 | 9 | Endothelial |
| XIAP | 0.37 | 0.00 | 0.00 | 0.32 | 0.19 | 9 | Endothelial |
| PLTP | 0.40 | 0.00 | 0.00 | 0.30 | 0.17 | 9 | Endothelial |
| GNAI2 | 0.30 | 0.00 | 0.00 | 0.55 | 0.38 | 9 | Endothelial |
| SYPL1 | 0.35 | 0.00 | 0.00 | 0.40 | 0.26 | 9 | Endothelial |
| KTN1 | 0.30 | 0.00 | 0.00 | 0.54 | 0.39 | 9 | Endothelial |
| PDIA3 | 0.33 | 0.00 | 0.00 | 0.60 | 0.46 | 9 | Endothelial |
| ADAMTS1 | 0.38 | 0.00 | 0.00 | 0.32 | 0.20 | 9 | Endothelial |
| STMN1 | 0.49 | 0.00 | 0.00 | 0.36 | 0.24 | 9 | Endothelial |
| G3BP2 | 0.39 | 0.00 | 0.00 | 0.42 | 0.29 | 9 | Endothelial |
| AFAP1L1 | 0.37 | 0.00 | 0.00 | 0.29 | 0.17 | 9 | Endothelial |
| SEPT2 | 0.30 | 0.00 | 0.00 | 0.51 | 0.37 | 9 | Endothelial |
| PTTG1IP | 0.30 | 0.00 | 0.00 | 0.30 | 0.18 | 9 | Endothelial |
| MSN | 0.26 | 0.00 | 0.00 | 0.59 | 0.44 | 9 | Endothelial |
| DNAJA1 | 0.32 | 0.00 | 0.00 | 0.64 | 0.51 | 9 | Endothelial |
| SPTAN1 | 0.33 | 0.00 | 0.00 | 0.28 | 0.17 | 9 | Endothelial |
| CFLAR | 0.30 | 0.00 | 0.00 | 0.31 | 0.19 | 9 | Endothelial |
| ARGLU1 | 0.32 | 0.00 | 0.00 | 0.55 | 0.41 | 9 | Endothelial |
| CD36 | 0.27 | 0.00 | 0.00 | 0.49 | 0.33 | 9 | Endothelial |
| CYB5R3 | 0.29 | 0.00 | 0.00 | 0.53 | 0.39 | 9 | Endothelial |
| DYNLL1 | 0.30 | 0.00 | 0.00 | 0.60 | 0.47 | 9 | Endothelial |
| NRAS | 0.33 | 0.00 | 0.00 | 0.26 | 0.15 | 9 | Endothelial |
| ARHGEF2 | 0.31 | 0.00 | 0.00 | 0.31 | 0.19 | 9 | Endothelial |
| NFIC | 0.36 | 0.00 | 0.00 | 0.37 | 0.25 | 9 | Endothelial |
| TXNIP | 0.27 | 0.00 | 0.00 | 0.40 | 0.27 | 9 | Endothelial |
| CTNNB1 | 0.32 | 0.00 | 0.00 | 0.48 | 0.34 | 9 | Endothelial |
| NFKBIA | 0.25 | 0.00 | 0.00 | 0.50 | 0.36 | 9 | Endothelial |
| ELOVL5 | 0.30 | 0.00 | 0.00 | 0.25 | 0.15 | 9 | Endothelial |
| VAMP3 | 0.33 | 0.00 | 0.00 | 0.25 | 0.15 | 9 | Endothelial |
| GLUL | 0.65 | 0.00 | 0.00 | 0.32 | 0.22 | 9 | Endothelial |
| PLCG1 | 0.31 | 0.00 | 0.00 | 0.26 | 0.15 | 9 | Endothelial |
| NDUFA8 | 0.34 | 0.00 | 0.00 | 0.36 | 0.25 | 9 | Endothelial |
| SEPT11 | 0.32 | 0.00 | 0.00 | 0.34 | 0.23 | 9 | Endothelial |
| IER2 | 0.27 | 0.00 | 0.00 | 0.38 | 0.26 | 9 | Endothelial |
| S100A13 | 0.30 | 0.00 | 0.00 | 0.39 | 0.27 | 9 | Endothelial |
| TNFAIP1 | 0.32 | 0.00 | 0.00 | 0.26 | 0.16 | 9 | Endothelial |
| PPIC | 0.27 | 0.00 | 0.00 | 0.52 | 0.38 | 9 | Endothelial |
| SMARCA2 | 0.32 | 0.00 | 0.00 | 0.45 | 0.33 | 9 | Endothelial |
| NUDT4 | 0.35 | 0.00 | 0.00 | 0.26 | 0.16 | 9 | Endothelial |
| LEPROT | 0.28 | 0.00 | 0.00 | 0.46 | 0.34 | 9 | Endothelial |
| CNIH1 | 0.28 | 0.00 | 0.00 | 0.29 | 0.19 | 9 | Endothelial |
| TSPO | 0.26 | 0.00 | 0.00 | 0.38 | 0.27 | 9 | Endothelial |
| ANO6 | 0.33 | 0.00 | 0.00 | 0.26 | 0.17 | 9 | Endothelial |
| EMP3 | 0.28 | 0.00 | 0.00 | 0.35 | 0.24 | 9 | Endothelial |
| NAP1L4 | 0.29 | 0.00 | 0.00 | 0.36 | 0.25 | 9 | Endothelial |
| WASF2 | 0.26 | 0.00 | 0.00 | 0.42 | 0.30 | 9 | Endothelial |
| HOOK3 | 0.29 | 0.00 | 0.00 | 0.32 | 0.22 | 9 | Endothelial |
| ZC3H7A | 0.31 | 0.00 | 0.00 | 0.37 | 0.26 | 9 | Endothelial |
| TTC3 | 0.29 | 0.00 | 0.00 | 0.40 | 0.29 | 9 | Endothelial |
| ZBTB20 | 0.34 | 0.00 | 0.00 | 0.37 | 0.27 | 9 | Endothelial |
| GOLIM4 | 0.30 | 0.00 | 0.00 | 0.30 | 0.20 | 9 | Endothelial |
| HES1 | 0.31 | 0.00 | 0.00 | 0.40 | 0.29 | 9 | Endothelial |
| GUK1 | 0.36 | 0.00 | 0.00 | 0.28 | 0.19 | 9 | Endothelial |
| ITM2C | 0.26 | 0.00 | 0.00 | 0.33 | 0.24 | 9 | Endothelial |
| SRPRA | 0.25 | 0.00 | 0.00 | 0.32 | 0.23 | 9 | Endothelial |
| PPIA | 0.29 | 0.00 | 0.00 | 0.30 | 0.21 | 9 | Endothelial |
| IVNS1ABP | 0.33 | 0.00 | 0.00 | 0.28 | 0.21 | 9 | Endothelial |
| DDX3X | 0.26 | 0.00 | 0.00 | 0.38 | 0.29 | 9 | Endothelial |
| TIA1 | 0.25 | 0.00 | 0.00 | 0.28 | 0.21 | 9 | Endothelial |
| MGP | 0.26 | 0.00 | 0.00 | 0.43 | 0.34 | 9 | Endothelial |
| COL1A1 | 2.65 | 0.00 | 0.00 | 0.97 | 0.28 | 10 | Myofibroblast |
| DCN | 2.62 | 0.00 | 0.00 | 0.70 | 0.05 | 10 | Myofibroblast |
| COL3A1 | 2.55 | 0.00 | 0.00 | 0.98 | 0.34 | 10 | Myofibroblast |
| COL1A2 | 2.36 | 0.00 | 0.00 | 0.99 | 0.40 | 10 | Myofibroblast |
| MFAP5 | 2.17 | 0.00 | 0.00 | 0.53 | 0.05 | 10 | Myofibroblast |
| COL14A1 | 2.05 | 0.00 | 0.00 | 0.59 | 0.05 | 10 | Myofibroblast |
| GPC3 | 1.98 | 0.00 | 0.00 | 0.80 | 0.17 | 10 | Myofibroblast |
| CCDC80 | 1.66 | 0.00 | 0.00 | 0.60 | 0.12 | 10 | Myofibroblast |
| AEBP1 | 1.56 | 0.00 | 0.00 | 0.48 | 0.07 | 10 | Myofibroblast |
| CYGB | 1.49 | 0.00 | 0.00 | 0.47 | 0.06 | 10 | Myofibroblast |
| OGN | 1.43 | 0.00 | 0.00 | 0.62 | 0.13 | 10 | Myofibroblast |
| PCOLCE | 1.41 | 0.00 | 0.00 | 0.58 | 0.10 | 10 | Myofibroblast |
| C1R | 1.37 | 0.00 | 0.00 | 0.40 | 0.04 | 10 | Myofibroblast |
| SERPINF1 | 1.36 | 0.00 | 0.00 | 0.31 | 0.02 | 10 | Myofibroblast |
| SERPING1 | 1.31 | 0.00 | 0.00 | 0.63 | 0.13 | 10 | Myofibroblast |
| COL5A1 | 1.30 | 0.00 | 0.00 | 0.54 | 0.10 | 10 | Myofibroblast |
| THBS2 | 1.16 | 0.00 | 0.00 | 0.29 | 0.02 | 10 | Myofibroblast |
| RCN3 | 1.33 | 0.00 | 0.00 | 0.58 | 0.12 | 10 | Myofibroblast |
| MGP | 1.77 | 0.00 | 0.00 | 0.89 | 0.32 | 10 | Myofibroblast |
| GAS1 | 1.16 | 0.00 | 0.00 | 0.32 | 0.04 | 10 | Myofibroblast |
| FBLN1 | 1.47 | 0.00 | 0.00 | 0.43 | 0.07 | 10 | Myofibroblast |
| ADAMTS2 | 1.25 | 0.00 | 0.00 | 0.50 | 0.09 | 10 | Myofibroblast |
| ASPN | 0.83 | 0.00 | 0.00 | 0.52 | 0.10 | 10 | Myofibroblast |
| BGN | 1.26 | 0.00 | 0.00 | 0.81 | 0.25 | 10 | Myofibroblast |
| COL6A2 | 1.30 | 0.00 | 0.00 | 0.67 | 0.17 | 10 | Myofibroblast |
| LUM | 1.36 | 0.00 | 0.00 | 0.31 | 0.04 | 10 | Myofibroblast |
| SPARC | 1.36 | 0.00 | 0.00 | 0.99 | 0.76 | 10 | Myofibroblast |
| PLAC9 | 1.17 | 0.00 | 0.00 | 0.88 | 0.29 | 10 | Myofibroblast |
| LTBP4 | 1.12 | 0.00 | 0.00 | 0.54 | 0.11 | 10 | Myofibroblast |
| CPE | 1.17 | 0.00 | 0.00 | 0.28 | 0.03 | 10 | Myofibroblast |
| PRRX1 | 1.01 | 0.00 | 0.00 | 0.26 | 0.03 | 10 | Myofibroblast |
| COL6A1 | 1.18 | 0.00 | 0.00 | 0.71 | 0.21 | 10 | Myofibroblast |
| IGFBP5 | 1.99 | 0.00 | 0.00 | 0.46 | 0.10 | 10 | Myofibroblast |
| RARRES2 | 1.36 | 0.00 | 0.00 | 0.60 | 0.16 | 10 | Myofibroblast |
| COL5A2 | 1.15 | 0.00 | 0.00 | 0.63 | 0.18 | 10 | Myofibroblast |
| IGFBP6 | 1.32 | 0.00 | 0.00 | 0.26 | 0.03 | 10 | Myofibroblast |
| BICC1 | 1.05 | 0.00 | 0.00 | 0.35 | 0.06 | 10 | Myofibroblast |
| ELN | 1.25 | 0.00 | 0.00 | 0.60 | 0.17 | 10 | Myofibroblast |
| LGALS1 | 1.03 | 0.00 | 0.00 | 0.92 | 0.46 | 10 | Myofibroblast |
| FBN1 | 1.12 | 0.00 | 0.00 | 0.58 | 0.17 | 10 | Myofibroblast |
| LOXL1 | 0.99 | 0.00 | 0.00 | 0.46 | 0.11 | 10 | Myofibroblast |
| FSTL1 | 1.12 | 0.00 | 0.00 | 0.86 | 0.45 | 10 | Myofibroblast |
| POSTN | 1.59 | 0.00 | 0.00 | 0.39 | 0.09 | 10 | Myofibroblast |
| COL6A3 | 0.95 | 0.00 | 0.00 | 0.47 | 0.12 | 10 | Myofibroblast |
| MMP23B | 0.76 | 0.00 | 0.00 | 0.26 | 0.04 | 10 | Myofibroblast |
| TIMP1 | 1.60 | 0.00 | 0.00 | 0.36 | 0.08 | 10 | Myofibroblast |
| IGF2 | 1.34 | 0.00 | 0.00 | 0.31 | 0.06 | 10 | Myofibroblast |
| GSN | 1.29 | 0.00 | 0.00 | 0.64 | 0.24 | 10 | Myofibroblast |
| IGFBP3 | 1.16 | 0.00 | 0.00 | 0.29 | 0.06 | 10 | Myofibroblast |
| IGFBP7 | 0.86 | 0.00 | 0.00 | 0.85 | 0.42 | 10 | Myofibroblast |
| SERPINH1 | 0.81 | 0.00 | 0.00 | 0.82 | 0.41 | 10 | Myofibroblast |
| CDH11 | 0.67 | 0.00 | 0.00 | 0.58 | 0.19 | 10 | Myofibroblast |
| CD63 | 1.03 | 0.00 | 0.00 | 0.71 | 0.34 | 10 | Myofibroblast |
| IGFBP4 | 0.92 | 0.00 | 0.00 | 0.64 | 0.27 | 10 | Myofibroblast |
| DPT | 1.00 | 0.00 | 0.00 | 0.50 | 0.17 | 10 | Myofibroblast |
| VCAN | 0.83 | 0.00 | 0.00 | 0.43 | 0.13 | 10 | Myofibroblast |
| VKORC1 | 0.78 | 0.00 | 0.00 | 0.39 | 0.11 | 10 | Myofibroblast |
| COL16A1 | 0.76 | 0.00 | 0.00 | 0.26 | 0.06 | 10 | Myofibroblast |
| TIMP2 | 0.85 | 0.00 | 0.00 | 0.56 | 0.21 | 10 | Myofibroblast |
| AKAP12 | 1.33 | 0.00 | 0.00 | 0.55 | 0.24 | 10 | Myofibroblast |
| NDN | 0.71 | 0.00 | 0.00 | 0.30 | 0.08 | 10 | Myofibroblast |
| S100A6 | 0.54 | 0.00 | 0.00 | 0.75 | 0.35 | 10 | Myofibroblast |
| CLEC3B | 0.86 | 0.00 | 0.00 | 0.27 | 0.07 | 10 | Myofibroblast |
| TUBB6 | 0.64 | 0.00 | 0.00 | 0.30 | 0.08 | 10 | Myofibroblast |
| LMNA | 1.07 | 0.00 | 0.00 | 0.54 | 0.22 | 10 | Myofibroblast |
| FXYD1 | 0.65 | 0.00 | 0.00 | 0.34 | 0.10 | 10 | Myofibroblast |
| TMED3 | 0.69 | 0.00 | 0.00 | 0.39 | 0.12 | 10 | Myofibroblast |
| PRSS35 | 0.70 | 0.00 | 0.00 | 0.30 | 0.08 | 10 | Myofibroblast |
| SEMA5A | 0.73 | 0.00 | 0.00 | 0.25 | 0.06 | 10 | Myofibroblast |
| LTBP1 | 0.75 | 0.00 | 0.00 | 0.25 | 0.06 | 10 | Myofibroblast |
| GPX8 | 0.75 | 0.00 | 0.00 | 0.39 | 0.13 | 10 | Myofibroblast |
| NFIB | 0.69 | 0.00 | 0.00 | 0.81 | 0.47 | 10 | Myofibroblast |
| RAB34 | 0.65 | 0.00 | 0.00 | 0.33 | 0.10 | 10 | Myofibroblast |
| CKAP4 | 0.73 | 0.00 | 0.00 | 0.46 | 0.17 | 10 | Myofibroblast |
| GPX3 | 0.69 | 0.00 | 0.00 | 0.41 | 0.14 | 10 | Myofibroblast |
| ITM2A | 0.85 | 0.00 | 0.00 | 0.28 | 0.08 | 10 | Myofibroblast |
| MMP14 | 0.77 | 0.00 | 0.00 | 0.56 | 0.25 | 10 | Myofibroblast |
| PEG3 | 0.94 | 0.00 | 0.00 | 0.56 | 0.26 | 10 | Myofibroblast |
| OLFML2B | 0.57 | 0.00 | 0.00 | 0.26 | 0.07 | 10 | Myofibroblast |
| PBX1 | 0.71 | 0.00 | 0.00 | 0.49 | 0.19 | 10 | Myofibroblast |
| COLEC12 | 0.60 | 0.00 | 0.00 | 0.34 | 0.11 | 10 | Myofibroblast |
| FAM114A1 | 0.65 | 0.00 | 0.00 | 0.31 | 0.09 | 10 | Myofibroblast |
| MAGED2 | 0.64 | 0.00 | 0.00 | 0.51 | 0.20 | 10 | Myofibroblast |
| MDK | 0.59 | 0.00 | 0.00 | 0.54 | 0.22 | 10 | Myofibroblast |
| NR2F2 | 0.65 | 0.00 | 0.00 | 0.49 | 0.20 | 10 | Myofibroblast |
| KDELR2 | 0.74 | 0.00 | 0.00 | 0.48 | 0.21 | 10 | Myofibroblast |
| EFEMP2 | 0.64 | 0.00 | 0.00 | 0.29 | 0.09 | 10 | Myofibroblast |
| PLXDC2 | 0.56 | 0.00 | 0.00 | 0.37 | 0.13 | 10 | Myofibroblast |
| CRTAP | 0.58 | 0.00 | 0.00 | 0.28 | 0.08 | 10 | Myofibroblast |
| FGFR1 | 0.56 | 0.00 | 0.00 | 0.33 | 0.11 | 10 | Myofibroblast |
| LRP1 | 0.58 | 0.00 | 0.00 | 0.35 | 0.12 | 10 | Myofibroblast |
| TPM2 | 0.46 | 0.00 | 0.00 | 0.45 | 0.17 | 10 | Myofibroblast |
| FKBP7 | 0.60 | 0.00 | 0.00 | 0.38 | 0.14 | 10 | Myofibroblast |
| MFAP2 | 0.68 | 0.00 | 0.00 | 0.66 | 0.35 | 10 | Myofibroblast |
| LOXL2 | 0.44 | 0.00 | 0.00 | 0.36 | 0.13 | 10 | Myofibroblast |
| IKBIP | 0.54 | 0.00 | 0.00 | 0.27 | 0.09 | 10 | Myofibroblast |
| PMEPA1 | 0.66 | 0.00 | 0.00 | 0.31 | 0.11 | 10 | Myofibroblast |
| FKBP9 | 0.59 | 0.00 | 0.00 | 0.36 | 0.14 | 10 | Myofibroblast |
| NID1 | 0.58 | 0.00 | 0.00 | 0.51 | 0.23 | 10 | Myofibroblast |
| S100A11 | 0.58 | 0.00 | 0.00 | 0.77 | 0.49 | 10 | Myofibroblast |
| RNASE4 | 0.64 | 0.00 | 0.00 | 0.42 | 0.17 | 10 | Myofibroblast |
| NDRG2 | 0.59 | 0.00 | 0.00 | 0.27 | 0.09 | 10 | Myofibroblast |
| ANTXR1 | 0.61 | 0.00 | 0.00 | 0.34 | 0.13 | 10 | Myofibroblast |
| GPX7 | 0.61 | 0.00 | 0.00 | 0.28 | 0.10 | 10 | Myofibroblast |
| CBX6 | 0.54 | 0.00 | 0.00 | 0.31 | 0.11 | 10 | Myofibroblast |
| FKBP10 | 0.52 | 0.00 | 0.00 | 0.26 | 0.09 | 10 | Myofibroblast |
| CPED1 | 0.61 | 0.00 | 0.00 | 0.27 | 0.09 | 10 | Myofibroblast |
| EID1 | 0.59 | 0.00 | 0.00 | 0.60 | 0.32 | 10 | Myofibroblast |
| NFIA | 0.61 | 0.00 | 0.00 | 0.50 | 0.24 | 10 | Myofibroblast |
| LHFPL2 | 0.71 | 0.00 | 0.00 | 0.25 | 0.08 | 10 | Myofibroblast |
| RRBP1 | 0.61 | 0.00 | 0.00 | 0.77 | 0.54 | 10 | Myofibroblast |
| ARF4 | 0.60 | 0.00 | 0.00 | 0.55 | 0.29 | 10 | Myofibroblast |
| EDNRA | 0.51 | 0.00 | 0.00 | 0.28 | 0.10 | 10 | Myofibroblast |
| IGF1 | 0.71 | 0.00 | 0.00 | 0.34 | 0.13 | 10 | Myofibroblast |
| S100A10 | 0.68 | 0.00 | 0.00 | 0.66 | 0.40 | 10 | Myofibroblast |
| TWSG1 | 0.47 | 0.00 | 0.00 | 0.31 | 0.12 | 10 | Myofibroblast |
| CST3 | 0.40 | 0.00 | 0.00 | 0.82 | 0.58 | 10 | Myofibroblast |
| LAPTM4A | 0.46 | 0.00 | 0.00 | 0.79 | 0.55 | 10 | Myofibroblast |
| PMP22 | 0.38 | 0.00 | 0.00 | 0.62 | 0.33 | 10 | Myofibroblast |
| LAMB1 | 0.54 | 0.00 | 0.00 | 0.36 | 0.15 | 10 | Myofibroblast |
| OLFML3 | 0.50 | 0.00 | 0.00 | 0.27 | 0.10 | 10 | Myofibroblast |
| CALU | 0.53 | 0.00 | 0.00 | 0.50 | 0.25 | 10 | Myofibroblast |
| CDKN1C | 0.82 | 0.00 | 0.00 | 0.37 | 0.17 | 10 | Myofibroblast |
| APOE | 0.34 | 0.00 | 0.00 | 0.53 | 0.27 | 10 | Myofibroblast |
| MYH10 | 0.41 | 0.00 | 0.00 | 0.48 | 0.22 | 10 | Myofibroblast |
| TUBA1A | 0.50 | 0.00 | 0.00 | 0.79 | 0.52 | 10 | Myofibroblast |
| DKK3 | 0.43 | 0.00 | 0.00 | 0.33 | 0.13 | 10 | Myofibroblast |
| FHL1 | 0.36 | 0.00 | 0.00 | 0.60 | 0.30 | 10 | Myofibroblast |
| FBLN5 | 0.35 | 0.00 | 0.00 | 0.41 | 0.18 | 10 | Myofibroblast |
| CD302 | 0.49 | 0.00 | 0.00 | 0.35 | 0.14 | 10 | Myofibroblast |
| RBMS3 | 0.48 | 0.00 | 0.00 | 0.40 | 0.17 | 10 | Myofibroblast |
| FIBIN | 0.51 | 0.00 | 0.00 | 0.29 | 0.11 | 10 | Myofibroblast |
| MORF4L2 | 0.51 | 0.00 | 0.00 | 0.47 | 0.23 | 10 | Myofibroblast |
| PPIC | 0.46 | 0.00 | 0.00 | 0.65 | 0.38 | 10 | Myofibroblast |
| NENF | 0.51 | 0.00 | 0.00 | 0.42 | 0.20 | 10 | Myofibroblast |
| NUPR1 | 0.39 | 0.00 | 0.00 | 0.44 | 0.20 | 10 | Myofibroblast |
| DPYSL3 | 0.42 | 0.00 | 0.00 | 0.25 | 0.09 | 10 | Myofibroblast |
| CYR61 | 0.77 | 0.00 | 0.00 | 0.29 | 0.12 | 10 | Myofibroblast |
| SPARCL1 | 0.37 | 0.00 | 0.00 | 0.70 | 0.41 | 10 | Myofibroblast |
| CALD1 | 0.26 | 0.00 | 0.00 | 0.74 | 0.44 | 10 | Myofibroblast |
| EGR1 | 0.72 | 0.00 | 0.00 | 0.28 | 0.11 | 10 | Myofibroblast |
| NFIX | 0.51 | 0.00 | 0.00 | 0.36 | 0.16 | 10 | Myofibroblast |
| SEC61G | 0.48 | 0.00 | 0.00 | 0.61 | 0.37 | 10 | Myofibroblast |
| SPATS2L | 0.49 | 0.00 | 0.00 | 0.26 | 0.10 | 10 | Myofibroblast |
| MYL9 | 0.30 | 0.00 | 0.00 | 0.38 | 0.17 | 10 | Myofibroblast |
| MAGED1 | 0.53 | 0.00 | 0.00 | 0.38 | 0.18 | 10 | Myofibroblast |
| ANXA5 | 0.43 | 0.00 | 0.00 | 0.67 | 0.43 | 10 | Myofibroblast |
| FCGRT | 0.39 | 0.00 | 0.00 | 0.31 | 0.13 | 10 | Myofibroblast |
| PPIB | 0.43 | 0.00 | 0.00 | 0.69 | 0.46 | 10 | Myofibroblast |
| EMP3 | 0.40 | 0.00 | 0.00 | 0.47 | 0.24 | 10 | Myofibroblast |
| PTPRD | 0.44 | 0.00 | 0.00 | 0.28 | 0.11 | 10 | Myofibroblast |
| PRDX4 | 0.47 | 0.00 | 0.00 | 0.34 | 0.16 | 10 | Myofibroblast |
| YBX3 | 0.44 | 0.00 | 0.00 | 0.39 | 0.19 | 10 | Myofibroblast |
| VIM | 0.49 | 0.00 | 0.00 | 0.88 | 0.69 | 10 | Myofibroblast |
| HDLBP | 0.43 | 0.00 | 0.00 | 0.41 | 0.21 | 10 | Myofibroblast |
| MTCH1 | 0.43 | 0.00 | 0.00 | 0.36 | 0.17 | 10 | Myofibroblast |
| GOLIM4 | 0.42 | 0.00 | 0.00 | 0.40 | 0.20 | 10 | Myofibroblast |
| SURF4 | 0.37 | 0.00 | 0.00 | 0.32 | 0.15 | 10 | Myofibroblast |
| MXRA8 | 0.34 | 0.00 | 0.00 | 0.36 | 0.17 | 10 | Myofibroblast |
| SSR4 | 0.39 | 0.00 | 0.00 | 0.48 | 0.27 | 10 | Myofibroblast |
| P4HB | 0.39 | 0.00 | 0.00 | 0.64 | 0.41 | 10 | Myofibroblast |
| ANXA2 | 0.42 | 0.00 | 0.00 | 0.70 | 0.49 | 10 | Myofibroblast |
| TSPAN3 | 0.43 | 0.00 | 0.00 | 0.34 | 0.17 | 10 | Myofibroblast |
| CRYAB | 0.37 | 0.00 | 0.00 | 0.33 | 0.16 | 10 | Myofibroblast |
| OST4 | 0.42 | 0.00 | 0.00 | 0.49 | 0.28 | 10 | Myofibroblast |
| LAMC1 | 0.38 | 0.00 | 0.00 | 0.33 | 0.16 | 10 | Myofibroblast |
| CD81 | 0.37 | 0.00 | 0.00 | 0.63 | 0.40 | 10 | Myofibroblast |
| LAMA4 | 0.45 | 0.00 | 0.00 | 0.29 | 0.13 | 10 | Myofibroblast |
| TPPP3 | 0.66 | 0.00 | 0.00 | 0.29 | 0.14 | 10 | Myofibroblast |
| RBPJ | 0.37 | 0.00 | 0.00 | 0.26 | 0.12 | 10 | Myofibroblast |
| ERLEC1 | 0.39 | 0.00 | 0.00 | 0.32 | 0.15 | 10 | Myofibroblast |
| TUBA1B | 0.41 | 0.00 | 0.00 | 0.68 | 0.46 | 10 | Myofibroblast |
| GSTP1 | 0.41 | 0.00 | 0.00 | 0.43 | 0.24 | 10 | Myofibroblast |
| FMO2 | 0.32 | 0.00 | 0.00 | 0.30 | 0.15 | 10 | Myofibroblast |
| RPL29 | 0.71 | 0.00 | 0.00 | 0.35 | 0.19 | 10 | Myofibroblast |
| NBEAL1 | 0.47 | 0.00 | 0.00 | 0.27 | 0.13 | 10 | Myofibroblast |
| GRB10 | 0.39 | 0.00 | 0.00 | 0.30 | 0.15 | 10 | Myofibroblast |
| DDOST | 0.37 | 0.00 | 0.00 | 0.37 | 0.19 | 10 | Myofibroblast |
| TCEAL8 | 0.38 | 0.00 | 0.00 | 0.39 | 0.21 | 10 | Myofibroblast |
| CELF2 | 0.32 | 0.00 | 0.00 | 0.47 | 0.27 | 10 | Myofibroblast |
| PHPT1 | 0.37 | 0.00 | 0.00 | 0.27 | 0.13 | 10 | Myofibroblast |
| ILK | 0.34 | 0.00 | 0.00 | 0.43 | 0.24 | 10 | Myofibroblast |
| MRFAP1 | 0.36 | 0.00 | 0.00 | 0.56 | 0.35 | 10 | Myofibroblast |
| SVIL | 0.39 | 0.00 | 0.00 | 0.32 | 0.16 | 10 | Myofibroblast |
| PDLIM2 | 0.35 | 0.00 | 0.00 | 0.35 | 0.18 | 10 | Myofibroblast |
| KDELR1 | 0.36 | 0.00 | 0.00 | 0.32 | 0.16 | 10 | Myofibroblast |
| PLS3 | 0.36 | 0.00 | 0.00 | 0.39 | 0.22 | 10 | Myofibroblast |
| NUCB1 | 0.30 | 0.00 | 0.00 | 0.31 | 0.16 | 10 | Myofibroblast |
| PDLIM7 | 0.31 | 0.00 | 0.00 | 0.26 | 0.12 | 10 | Myofibroblast |
| GAS6 | 0.69 | 0.00 | 0.00 | 0.30 | 0.16 | 10 | Myofibroblast |
| SEC61A1 | 0.34 | 0.00 | 0.00 | 0.31 | 0.16 | 10 | Myofibroblast |
| CALR | 0.33 | 0.00 | 0.00 | 0.66 | 0.45 | 10 | Myofibroblast |
| GPM6B | 0.40 | 0.00 | 0.00 | 0.26 | 0.13 | 10 | Myofibroblast |
| RPL24 | 0.41 | 0.00 | 0.00 | 0.67 | 0.49 | 10 | Myofibroblast |
| ZFP36L1 | 0.32 | 0.00 | 0.00 | 0.67 | 0.46 | 10 | Myofibroblast |
| RPL7A | 0.53 | 0.00 | 0.00 | 0.60 | 0.43 | 10 | Myofibroblast |
| THBS1 | 0.25 | 0.00 | 0.00 | 0.26 | 0.13 | 10 | Myofibroblast |
| SAR1A | 0.32 | 0.00 | 0.00 | 0.37 | 0.21 | 10 | Myofibroblast |
| TMED9 | 0.35 | 0.00 | 0.00 | 0.46 | 0.28 | 10 | Myofibroblast |
| RPN2 | 0.30 | 0.00 | 0.00 | 0.38 | 0.21 | 10 | Myofibroblast |
| SERF2 | 0.28 | 0.00 | 0.00 | 0.77 | 0.60 | 10 | Myofibroblast |
| CNPY2 | 0.31 | 0.00 | 0.00 | 0.37 | 0.20 | 10 | Myofibroblast |
| CBX5 | 0.42 | 0.00 | 0.00 | 0.40 | 0.24 | 10 | Myofibroblast |
| EEF1D | 0.44 | 0.00 | 0.00 | 0.47 | 0.30 | 10 | Myofibroblast |
| YIPF5 | 0.30 | 0.00 | 0.00 | 0.27 | 0.14 | 10 | Myofibroblast |
| COPB2 | 0.35 | 0.00 | 0.00 | 0.32 | 0.18 | 10 | Myofibroblast |
| ATRAID | 0.29 | 0.00 | 0.00 | 0.34 | 0.19 | 10 | Myofibroblast |
| COMT | 0.30 | 0.00 | 0.00 | 0.30 | 0.16 | 10 | Myofibroblast |
| HMCN1 | 0.34 | 0.00 | 0.00 | 0.27 | 0.13 | 10 | Myofibroblast |
| KANK2 | 0.33 | 0.00 | 0.00 | 0.26 | 0.13 | 10 | Myofibroblast |
| LRPAP1 | 0.32 | 0.00 | 0.00 | 0.31 | 0.17 | 10 | Myofibroblast |
| SSR2 | 0.35 | 0.00 | 0.00 | 0.52 | 0.34 | 10 | Myofibroblast |
| SFR1 | 0.40 | 0.00 | 0.00 | 0.47 | 0.30 | 10 | Myofibroblast |
| PDIA6 | 0.31 | 0.00 | 0.00 | 0.51 | 0.32 | 10 | Myofibroblast |
| REXO2 | 0.30 | 0.00 | 0.00 | 0.28 | 0.15 | 10 | Myofibroblast |
| NFIC | 0.34 | 0.00 | 0.00 | 0.41 | 0.25 | 10 | Myofibroblast |
| CAV1 | 0.31 | 0.00 | 0.00 | 0.58 | 0.37 | 10 | Myofibroblast |
| TTC3 | 0.25 | 0.00 | 0.00 | 0.48 | 0.29 | 10 | Myofibroblast |
| GAPDH | 0.41 | 0.00 | 0.00 | 0.33 | 0.19 | 10 | Myofibroblast |
| DAAM1 | 0.37 | 0.00 | 0.00 | 0.30 | 0.16 | 10 | Myofibroblast |
| NUCKS1 | 0.37 | 0.00 | 0.00 | 0.45 | 0.29 | 10 | Myofibroblast |
| CD164 | 0.31 | 0.00 | 0.00 | 0.29 | 0.16 | 10 | Myofibroblast |
| CAMK2N1 | 0.30 | 0.00 | 0.00 | 0.27 | 0.14 | 10 | Myofibroblast |
| HNRNPA1 | 0.58 | 0.00 | 0.00 | 0.45 | 0.31 | 10 | Myofibroblast |
| SEPT7 | 0.26 | 0.00 | 0.00 | 0.77 | 0.59 | 10 | Myofibroblast |
| RTN4 | 0.26 | 0.00 | 0.00 | 0.43 | 0.26 | 10 | Myofibroblast |
| BSG | 0.30 | 0.00 | 0.00 | 0.58 | 0.39 | 10 | Myofibroblast |
| GNAS | 0.32 | 0.00 | 0.00 | 0.80 | 0.66 | 10 | Myofibroblast |
| HSPB1 | 0.45 | 0.00 | 0.00 | 0.30 | 0.17 | 10 | Myofibroblast |
| HCFC1R1 | 0.29 | 0.00 | 0.00 | 0.27 | 0.15 | 10 | Myofibroblast |
| AP2M1 | 0.33 | 0.00 | 0.00 | 0.30 | 0.17 | 10 | Myofibroblast |
| ERGIC3 | 0.25 | 0.00 | 0.00 | 0.30 | 0.17 | 10 | Myofibroblast |
| CCNI | 0.48 | 0.00 | 0.00 | 0.32 | 0.20 | 10 | Myofibroblast |
| ZFP36L2 | 0.49 | 0.00 | 0.00 | 0.25 | 0.14 | 10 | Myofibroblast |
| TMED2 | 0.30 | 0.00 | 0.00 | 0.41 | 0.26 | 10 | Myofibroblast |
| COPE | 0.25 | 0.00 | 0.00 | 0.40 | 0.25 | 10 | Myofibroblast |
| PARVA | 0.26 | 0.00 | 0.00 | 0.28 | 0.16 | 10 | Myofibroblast |
| PDIA3 | 0.27 | 0.00 | 0.00 | 0.63 | 0.46 | 10 | Myofibroblast |
| RPL36A | 0.27 | 0.00 | 0.00 | 0.78 | 0.64 | 10 | Myofibroblast |
| BFAR | 0.28 | 0.00 | 0.00 | 0.25 | 0.14 | 10 | Myofibroblast |
| RPL13A | 0.65 | 0.00 | 0.00 | 0.70 | 0.59 | 10 | Myofibroblast |
| KRTCAP2 | 0.26 | 0.00 | 0.00 | 0.45 | 0.30 | 10 | Myofibroblast |
| HSPA5 | 0.29 | 0.00 | 0.00 | 0.70 | 0.55 | 10 | Myofibroblast |
| TRMT112 | 0.27 | 0.00 | 0.00 | 0.29 | 0.18 | 10 | Myofibroblast |
| LIMA1 | 0.28 | 0.00 | 0.00 | 0.26 | 0.15 | 10 | Myofibroblast |
| CNN3 | 0.28 | 0.00 | 0.00 | 0.47 | 0.33 | 10 | Myofibroblast |
| MIF | 0.27 | 0.00 | 0.00 | 0.35 | 0.23 | 10 | Myofibroblast |
| CIRBP | 0.33 | 0.00 | 0.00 | 0.38 | 0.26 | 10 | Myofibroblast |
| C12orf57 | 0.29 | 0.00 | 0.00 | 0.27 | 0.16 | 10 | Myofibroblast |
| NDUFS5 | 0.29 | 0.00 | 0.00 | 0.39 | 0.27 | 10 | Myofibroblast |
| HSP90B1 | 0.40 | 0.00 | 0.00 | 0.83 | 0.75 | 10 | Myofibroblast |
| CSRP2 | 0.27 | 0.00 | 0.00 | 0.26 | 0.16 | 10 | Myofibroblast |
| CRIP1 | 0.59 | 0.00 | 0.00 | 0.60 | 0.47 | 10 | Myofibroblast |
| SUMO2 | 0.26 | 0.00 | 0.00 | 0.47 | 0.34 | 10 | Myofibroblast |
| RPL23A | 0.25 | 0.00 | 0.00 | 0.79 | 0.69 | 10 | Myofibroblast |
| ACTA2 | 2.55 | 0.00 | 0.00 | 0.96 | 0.23 | 11 | Myofibroblast |
| MYH11 | 2.42 | 0.00 | 0.00 | 0.79 | 0.11 | 11 | Myofibroblast |
| MYL9 | 2.32 | 0.00 | 0.00 | 0.86 | 0.16 | 11 | Myofibroblast |
| IGFBP5 | 2.31 | 0.00 | 0.00 | 0.64 | 0.09 | 11 | Myofibroblast |
| TPM2 | 2.24 | 0.00 | 0.00 | 0.86 | 0.16 | 11 | Myofibroblast |
| ACTC1 | 2.19 | 0.00 | 0.00 | 0.36 | 0.02 | 11 | Myofibroblast |
| TAGLN | 2.18 | 0.00 | 0.00 | 0.92 | 0.17 | 11 | Myofibroblast |
| PI15 | 2.11 | 0.00 | 0.00 | 0.44 | 0.03 | 11 | Myofibroblast |
| GPC6 | 1.92 | 0.00 | 0.00 | 0.41 | 0.01 | 11 | Myofibroblast |
| ACTG2 | 1.80 | 0.00 | 0.00 | 0.57 | 0.08 | 11 | Myofibroblast |
| ELN | 1.66 | 0.00 | 0.00 | 0.75 | 0.17 | 11 | Myofibroblast |
| SH3BGR | 1.62 | 0.00 | 0.00 | 0.29 | 0.01 | 11 | Myofibroblast |
| CALD1 | 1.61 | 0.00 | 0.00 | 0.95 | 0.43 | 11 | Myofibroblast |
| CNN1 | 1.49 | 0.00 | 0.00 | 0.38 | 0.03 | 11 | Myofibroblast |
| MAP1B | 1.46 | 0.00 | 0.00 | 0.52 | 0.09 | 11 | Myofibroblast |
| THSD4 | 1.46 | 0.00 | 0.00 | 0.31 | 0.02 | 11 | Myofibroblast |
| AOC3 | 1.45 | 0.00 | 0.00 | 0.38 | 0.03 | 11 | Myofibroblast |
| SOD3 | 1.39 | 0.00 | 0.00 | 0.40 | 0.06 | 11 | Myofibroblast |
| CPE | 1.33 | 0.00 | 0.00 | 0.33 | 0.03 | 11 | Myofibroblast |
| NOTCH3 | 1.30 | 0.00 | 0.00 | 0.36 | 0.03 | 11 | Myofibroblast |
| CRISPLD2 | 1.22 | 0.00 | 0.00 | 0.30 | 0.03 | 11 | Myofibroblast |
| NTRK3 | 1.22 | 0.00 | 0.00 | 0.28 | 0.01 | 11 | Myofibroblast |
| HEYL | 1.12 | 0.00 | 0.00 | 0.28 | 0.03 | 11 | Myofibroblast |
| NDRG2 | 1.28 | 0.00 | 0.00 | 0.47 | 0.08 | 11 | Myofibroblast |
| PDGFRB | 1.21 | 0.00 | 0.00 | 0.40 | 0.06 | 11 | Myofibroblast |
| IGFBP7 | 1.33 | 0.00 | 0.00 | 0.92 | 0.42 | 11 | Myofibroblast |
| TPM1 | 1.59 | 0.00 | 0.00 | 0.95 | 0.51 | 11 | Myofibroblast |
| MUSTN1 | 1.52 | 0.00 | 0.00 | 0.38 | 0.07 | 11 | Myofibroblast |
| LGALS1 | 1.04 | 0.00 | 0.00 | 0.95 | 0.46 | 11 | Myofibroblast |
| POSTN | 1.21 | 0.00 | 0.00 | 0.44 | 0.09 | 11 | Myofibroblast |
| ITGA8 | 1.50 | 0.00 | 0.00 | 0.43 | 0.09 | 11 | Myofibroblast |
| FAM129A | 1.24 | 0.00 | 0.00 | 0.39 | 0.08 | 11 | Myofibroblast |
| MYL6 | 1.03 | 0.00 | 0.00 | 0.93 | 0.72 | 11 | Myofibroblast |
| FXYD1 | 1.04 | 0.00 | 0.00 | 0.42 | 0.10 | 11 | Myofibroblast |
| COL1A2 | 0.60 | 0.00 | 0.00 | 0.95 | 0.40 | 11 | Myofibroblast |
| CRIP1 | 1.16 | 0.00 | 0.00 | 0.84 | 0.47 | 11 | Myofibroblast |
| CSRP2 | 1.34 | 0.00 | 0.00 | 0.51 | 0.15 | 11 | Myofibroblast |
| FSTL1 | 1.09 | 0.00 | 0.00 | 0.84 | 0.45 | 11 | Myofibroblast |
| CXCL12 | 0.69 | 0.00 | 0.00 | 0.47 | 0.13 | 11 | Myofibroblast |
| DSTN | 1.10 | 0.00 | 0.00 | 0.84 | 0.53 | 11 | Myofibroblast |
| MGP | 0.91 | 0.00 | 0.00 | 0.79 | 0.33 | 11 | Myofibroblast |
| MYLK | 1.15 | 0.00 | 0.00 | 0.70 | 0.29 | 11 | Myofibroblast |
| FLNA | 1.06 | 0.00 | 0.00 | 0.61 | 0.24 | 11 | Myofibroblast |
| FBLIM1 | 1.24 | 0.00 | 0.00 | 0.49 | 0.16 | 11 | Myofibroblast |
| TINAGL1 | 0.99 | 0.00 | 0.00 | 0.42 | 0.12 | 11 | Myofibroblast |
| SEMA5A | 0.97 | 0.00 | 0.00 | 0.28 | 0.06 | 11 | Myofibroblast |
| PLS3 | 1.00 | 0.00 | 0.00 | 0.55 | 0.21 | 11 | Myofibroblast |
| ACTN1 | 0.97 | 0.00 | 0.00 | 0.57 | 0.23 | 11 | Myofibroblast |
| RASL12 | 0.79 | 0.00 | 0.00 | 0.26 | 0.06 | 11 | Myofibroblast |
| CRYAB | 0.88 | 0.00 | 0.00 | 0.46 | 0.16 | 11 | Myofibroblast |
| DES | 0.98 | 0.00 | 0.00 | 0.27 | 0.06 | 11 | Myofibroblast |
| PLAC9 | 0.56 | 0.00 | 0.00 | 0.72 | 0.30 | 11 | Myofibroblast |
| COL14A1 | 1.02 | 0.00 | 0.00 | 0.26 | 0.06 | 11 | Myofibroblast |
| RARRES2 | 0.67 | 0.00 | 0.00 | 0.48 | 0.16 | 11 | Myofibroblast |
| CCDC80 | 0.86 | 0.00 | 0.00 | 0.39 | 0.12 | 11 | Myofibroblast |
| COL1A1 | 0.54 | 0.00 | 0.00 | 0.68 | 0.28 | 11 | Myofibroblast |
| PTP4A3 | 0.92 | 0.00 | 0.00 | 0.30 | 0.08 | 11 | Myofibroblast |
| PRDM6 | 0.78 | 0.00 | 0.00 | 0.27 | 0.07 | 11 | Myofibroblast |
| CAV1 | 0.71 | 0.00 | 0.00 | 0.75 | 0.36 | 11 | Myofibroblast |
| HSPB1 | 1.12 | 0.00 | 0.00 | 0.45 | 0.17 | 11 | Myofibroblast |
| LOXL1 | 0.90 | 0.00 | 0.00 | 0.36 | 0.11 | 11 | Myofibroblast |
| PDLIM7 | 0.83 | 0.00 | 0.00 | 0.37 | 0.12 | 11 | Myofibroblast |
| STOM | 0.71 | 0.00 | 0.00 | 0.31 | 0.09 | 11 | Myofibroblast |
| SLMAP | 0.93 | 0.00 | 0.00 | 0.40 | 0.14 | 11 | Myofibroblast |
| NEXN | 0.66 | 0.00 | 0.00 | 0.53 | 0.21 | 11 | Myofibroblast |
| MEF2C | 0.79 | 0.00 | 0.00 | 0.45 | 0.18 | 11 | Myofibroblast |
| FILIP1L | 0.79 | 0.00 | 0.00 | 0.66 | 0.33 | 11 | Myofibroblast |
| SPARC | 0.49 | 0.00 | 0.00 | 0.98 | 0.76 | 11 | Myofibroblast |
| OLFML2B | 0.70 | 0.00 | 0.00 | 0.25 | 0.07 | 11 | Myofibroblast |
| ILK | 0.76 | 0.00 | 0.00 | 0.52 | 0.23 | 11 | Myofibroblast |
| DNAJB4 | 1.00 | 0.00 | 0.00 | 0.31 | 0.10 | 11 | Myofibroblast |
| PPP1R14A | 0.68 | 0.00 | 0.00 | 0.33 | 0.11 | 11 | Myofibroblast |
| C1orf198 | 0.77 | 0.00 | 0.00 | 0.28 | 0.08 | 11 | Myofibroblast |
| PPP1R12A | 0.86 | 0.00 | 0.00 | 0.55 | 0.26 | 11 | Myofibroblast |
| BGN | 0.68 | 0.00 | 0.00 | 0.57 | 0.26 | 11 | Myofibroblast |
| HCFC1R1 | 0.86 | 0.00 | 0.00 | 0.38 | 0.14 | 11 | Myofibroblast |
| ADAMTS2 | 0.84 | 0.00 | 0.00 | 0.31 | 0.10 | 11 | Myofibroblast |
| SMTN | 0.81 | 0.00 | 0.00 | 0.28 | 0.09 | 11 | Myofibroblast |
| CD63 | 0.61 | 0.00 | 0.00 | 0.65 | 0.34 | 11 | Myofibroblast |
| VIM | 0.57 | 0.00 | 0.00 | 0.90 | 0.69 | 11 | Myofibroblast |
| CTGF | 0.85 | 0.00 | 0.00 | 0.29 | 0.09 | 11 | Myofibroblast |
| RBPMS | 0.81 | 0.00 | 0.00 | 0.42 | 0.18 | 11 | Myofibroblast |
| MAGED2 | 0.70 | 0.00 | 0.00 | 0.47 | 0.21 | 11 | Myofibroblast |
| PDLIM1 | 0.67 | 0.00 | 0.00 | 0.39 | 0.15 | 11 | Myofibroblast |
| TGFB2 | 0.93 | 0.00 | 0.00 | 0.35 | 0.13 | 11 | Myofibroblast |
| PPP1CB | 0.72 | 0.00 | 0.00 | 0.45 | 0.20 | 11 | Myofibroblast |
| NDN | 0.61 | 0.00 | 0.00 | 0.25 | 0.08 | 11 | Myofibroblast |
| MCAM | 0.66 | 0.00 | 0.00 | 0.36 | 0.13 | 11 | Myofibroblast |
| MSRB3 | 0.67 | 0.00 | 0.00 | 0.28 | 0.09 | 11 | Myofibroblast |
| TM4SF1 | 0.53 | 0.00 | 0.00 | 0.49 | 0.22 | 11 | Myofibroblast |
| FHL1 | 0.32 | 0.00 | 0.00 | 0.62 | 0.30 | 11 | Myofibroblast |
| RPLP1 | 0.37 | 0.00 | 0.00 | 0.96 | 0.91 | 11 | Myofibroblast |
| MFGE8 | 0.62 | 0.00 | 0.00 | 0.39 | 0.17 | 11 | Myofibroblast |
| ZFHX3 | 0.71 | 0.00 | 0.00 | 0.39 | 0.17 | 11 | Myofibroblast |
| MYH10 | 0.65 | 0.00 | 0.00 | 0.48 | 0.22 | 11 | Myofibroblast |
| CBX6 | 0.65 | 0.00 | 0.00 | 0.29 | 0.11 | 11 | Myofibroblast |
| NUPR1 | 0.62 | 0.00 | 0.00 | 0.45 | 0.20 | 11 | Myofibroblast |
| LMNA | 0.66 | 0.00 | 0.00 | 0.46 | 0.23 | 11 | Myofibroblast |
| ITGA4 | 0.70 | 0.00 | 0.00 | 0.26 | 0.09 | 11 | Myofibroblast |
| TGFB1I1 | 0.64 | 0.00 | 0.00 | 0.36 | 0.15 | 11 | Myofibroblast |
| MGST3 | 0.75 | 0.00 | 0.00 | 0.32 | 0.13 | 11 | Myofibroblast |
| MT-CYB | 0.37 | 0.00 | 0.00 | 0.95 | 0.83 | 11 | Myofibroblast |
| CKB | 0.67 | 0.00 | 0.00 | 0.33 | 0.14 | 11 | Myofibroblast |
| MT-ND4 | 0.50 | 0.00 | 0.00 | 0.89 | 0.76 | 11 | Myofibroblast |
| CST3 | 0.42 | 0.00 | 0.00 | 0.81 | 0.59 | 11 | Myofibroblast |
| MIF | 0.61 | 0.00 | 0.00 | 0.45 | 0.23 | 11 | Myofibroblast |
| ANTXR1 | 0.62 | 0.00 | 0.00 | 0.31 | 0.13 | 11 | Myofibroblast |
| EID1 | 0.54 | 0.00 | 0.00 | 0.57 | 0.32 | 11 | Myofibroblast |
| MT-CO3 | 0.33 | 0.00 | 0.00 | 0.96 | 0.89 | 11 | Myofibroblast |
| EEF1D | 0.69 | 0.00 | 0.00 | 0.51 | 0.30 | 11 | Myofibroblast |
| RPL10A | 0.43 | 0.00 | 0.00 | 0.88 | 0.76 | 11 | Myofibroblast |
| RCN3 | 0.59 | 0.00 | 0.00 | 0.30 | 0.13 | 11 | Myofibroblast |
| SH3BGRL | 0.38 | 0.00 | 0.00 | 0.72 | 0.45 | 11 | Myofibroblast |
| GSN | 0.32 | 0.00 | 0.00 | 0.48 | 0.25 | 11 | Myofibroblast |
| MT-ND3 | 0.65 | 0.00 | 0.00 | 0.68 | 0.51 | 11 | Myofibroblast |
| RAB34 | 0.50 | 0.00 | 0.00 | 0.26 | 0.10 | 11 | Myofibroblast |
| HNRNPA1 | 0.79 | 0.00 | 0.00 | 0.49 | 0.31 | 11 | Myofibroblast |
| NENF | 0.50 | 0.00 | 0.00 | 0.40 | 0.20 | 11 | Myofibroblast |
| SERPINE2 | 0.44 | 0.00 | 0.00 | 0.47 | 0.25 | 11 | Myofibroblast |
| CD81 | 0.55 | 0.00 | 0.00 | 0.61 | 0.40 | 11 | Myofibroblast |
| NFIA | 0.62 | 0.00 | 0.00 | 0.44 | 0.24 | 11 | Myofibroblast |
| CNN3 | 0.47 | 0.00 | 0.00 | 0.55 | 0.32 | 11 | Myofibroblast |
| FBN1 | 0.52 | 0.00 | 0.00 | 0.35 | 0.17 | 11 | Myofibroblast |
| TPPP3 | 0.67 | 0.00 | 0.00 | 0.30 | 0.14 | 11 | Myofibroblast |
| SLC25A4 | 0.64 | 0.00 | 0.00 | 0.63 | 0.47 | 11 | Myofibroblast |
| GAPDH | 0.85 | 0.00 | 0.00 | 0.36 | 0.19 | 11 | Myofibroblast |
| NUDT4 | 0.50 | 0.00 | 0.00 | 0.33 | 0.16 | 11 | Myofibroblast |
| VCL | 0.58 | 0.00 | 0.00 | 0.30 | 0.14 | 11 | Myofibroblast |
| EIF3L | 0.52 | 0.00 | 0.00 | 0.33 | 0.16 | 11 | Myofibroblast |
| TSPAN3 | 0.50 | 0.00 | 0.00 | 0.34 | 0.17 | 11 | Myofibroblast |
| PBXIP1 | 0.56 | 0.00 | 0.00 | 0.30 | 0.14 | 11 | Myofibroblast |
| APOE | 0.33 | 0.00 | 0.00 | 0.47 | 0.27 | 11 | Myofibroblast |
| RPS4X | 0.35 | 0.00 | 0.00 | 0.95 | 0.88 | 11 | Myofibroblast |
| RPL29 | 1.05 | 0.00 | 0.00 | 0.35 | 0.19 | 11 | Myofibroblast |
| NACA | 0.40 | 0.00 | 0.00 | 0.76 | 0.61 | 11 | Myofibroblast |
| IMPDH2 | 0.47 | 0.00 | 0.00 | 0.26 | 0.11 | 11 | Myofibroblast |
| RHOJ | 0.52 | 0.00 | 0.00 | 0.35 | 0.18 | 11 | Myofibroblast |
| EEF2 | 0.48 | 0.00 | 0.00 | 0.73 | 0.59 | 11 | Myofibroblast |
| GSTP1 | 0.56 | 0.00 | 0.00 | 0.42 | 0.24 | 11 | Myofibroblast |
| CDC42EP3 | 0.69 | 0.00 | 0.00 | 0.28 | 0.13 | 11 | Myofibroblast |
| RBMX | 0.51 | 0.00 | 0.00 | 0.28 | 0.14 | 11 | Myofibroblast |
| SERPINH1 | 0.42 | 0.00 | 0.00 | 0.63 | 0.41 | 11 | Myofibroblast |
| SEPT7 | 0.41 | 0.00 | 0.00 | 0.76 | 0.59 | 11 | Myofibroblast |
| EIF4A2 | 0.47 | 0.00 | 0.00 | 0.53 | 0.35 | 11 | Myofibroblast |
| TUBA1A | 0.41 | 0.00 | 0.00 | 0.70 | 0.52 | 11 | Myofibroblast |
| NDUFS5 | 0.49 | 0.00 | 0.00 | 0.43 | 0.27 | 11 | Myofibroblast |
| CYR61 | 0.45 | 0.00 | 0.00 | 0.26 | 0.12 | 11 | Myofibroblast |
| NR2F2 | 0.48 | 0.00 | 0.00 | 0.38 | 0.21 | 11 | Myofibroblast |
| KANK2 | 0.45 | 0.00 | 0.00 | 0.27 | 0.13 | 11 | Myofibroblast |
| DAAM1 | 0.61 | 0.00 | 0.00 | 0.31 | 0.16 | 11 | Myofibroblast |
| CFL2 | 0.50 | 0.00 | 0.00 | 0.40 | 0.23 | 11 | Myofibroblast |
| RPL8 | 0.34 | 0.00 | 0.00 | 0.89 | 0.81 | 11 | Myofibroblast |
| MT-ND2 | 0.31 | 0.00 | 0.00 | 0.84 | 0.71 | 11 | Myofibroblast |
| TSC22D1 | 0.46 | 0.00 | 0.00 | 0.53 | 0.35 | 11 | Myofibroblast |
| RPL7 | 0.52 | 0.00 | 0.00 | 0.85 | 0.77 | 11 | Myofibroblast |
| MT-CO2 | 0.31 | 0.00 | 0.00 | 0.88 | 0.79 | 11 | Myofibroblast |
| RPL23A | 0.49 | 0.00 | 0.00 | 0.80 | 0.69 | 11 | Myofibroblast |
| CCNI | 0.70 | 0.00 | 0.00 | 0.34 | 0.20 | 11 | Myofibroblast |
| SERF2 | 0.32 | 0.00 | 0.00 | 0.75 | 0.60 | 11 | Myofibroblast |
| ZBTB38 | 0.53 | 0.00 | 0.00 | 0.27 | 0.13 | 11 | Myofibroblast |
| RSU1 | 0.46 | 0.00 | 0.00 | 0.36 | 0.20 | 11 | Myofibroblast |
| PPIA | 0.51 | 0.00 | 0.00 | 0.36 | 0.21 | 11 | Myofibroblast |
| THRA | 0.46 | 0.00 | 0.00 | 0.31 | 0.17 | 11 | Myofibroblast |
| ITGB1 | 0.32 | 0.00 | 0.00 | 0.83 | 0.64 | 11 | Myofibroblast |
| TIMP2 | 0.43 | 0.00 | 0.00 | 0.38 | 0.22 | 11 | Myofibroblast |
| TUBA1B | 0.40 | 0.00 | 0.00 | 0.63 | 0.47 | 11 | Myofibroblast |
| CSRP1 | 0.43 | 0.00 | 0.00 | 0.35 | 0.19 | 11 | Myofibroblast |
| ACADM | 0.50 | 0.00 | 0.00 | 0.26 | 0.13 | 11 | Myofibroblast |
| CAV2 | 0.31 | 0.00 | 0.00 | 0.39 | 0.22 | 11 | Myofibroblast |
| RHOC | 0.35 | 0.00 | 0.00 | 0.41 | 0.24 | 11 | Myofibroblast |
| NBEAL1 | 0.40 | 0.00 | 0.00 | 0.25 | 0.13 | 11 | Myofibroblast |
| DST | 0.39 | 0.00 | 0.00 | 0.29 | 0.16 | 11 | Myofibroblast |
| ANXA5 | 0.34 | 0.00 | 0.00 | 0.60 | 0.43 | 11 | Myofibroblast |
| BTBD3 | 0.56 | 0.00 | 0.00 | 0.26 | 0.14 | 11 | Myofibroblast |
| RTN4 | 0.37 | 0.00 | 0.00 | 0.43 | 0.26 | 11 | Myofibroblast |
| SNRPD2 | 0.36 | 0.00 | 0.00 | 0.44 | 0.28 | 11 | Myofibroblast |
| COX7C | 0.31 | 0.00 | 0.00 | 0.77 | 0.64 | 11 | Myofibroblast |
| TNS1 | 0.32 | 0.00 | 0.00 | 0.38 | 0.23 | 11 | Myofibroblast |
| LOXL2 | 0.36 | 0.00 | 0.00 | 0.26 | 0.13 | 11 | Myofibroblast |
| CALU | 0.40 | 0.00 | 0.00 | 0.40 | 0.25 | 11 | Myofibroblast |
| CIRBP | 0.62 | 0.00 | 0.00 | 0.38 | 0.26 | 11 | Myofibroblast |
| MRFAP1 | 0.39 | 0.00 | 0.00 | 0.51 | 0.35 | 11 | Myofibroblast |
| C12orf57 | 0.39 | 0.00 | 0.00 | 0.29 | 0.16 | 11 | Myofibroblast |
| MT-ND4L | 0.35 | 0.00 | 0.00 | 0.55 | 0.39 | 11 | Myofibroblast |
| MT-ND5 | 0.29 | 0.00 | 0.00 | 0.88 | 0.75 | 11 | Myofibroblast |
| TPM4 | 0.37 | 0.00 | 0.00 | 0.67 | 0.51 | 11 | Myofibroblast |
| S100A11 | 0.33 | 0.00 | 0.00 | 0.66 | 0.50 | 11 | Myofibroblast |
| NET1 | 0.52 | 0.00 | 0.00 | 0.28 | 0.16 | 11 | Myofibroblast |
| RPL7A | 0.64 | 0.00 | 0.00 | 0.54 | 0.44 | 11 | Myofibroblast |
| ALDOA | 0.41 | 0.00 | 0.00 | 0.36 | 0.23 | 11 | Myofibroblast |
| CALM2 | 0.31 | 0.00 | 0.00 | 0.77 | 0.66 | 11 | Myofibroblast |
| COL6A1 | 0.28 | 0.00 | 0.00 | 0.38 | 0.22 | 11 | Myofibroblast |
| OGN | 0.43 | 0.00 | 0.00 | 0.25 | 0.14 | 11 | Myofibroblast |
| RHEB | 0.38 | 0.00 | 0.00 | 0.33 | 0.20 | 11 | Myofibroblast |
| SUMO2 | 0.39 | 0.00 | 0.00 | 0.48 | 0.34 | 11 | Myofibroblast |
| UQCRB | 0.32 | 0.00 | 0.00 | 0.61 | 0.47 | 11 | Myofibroblast |
| COL6A2 | 0.27 | 0.00 | 0.00 | 0.32 | 0.18 | 11 | Myofibroblast |
| ECH1 | 0.38 | 0.00 | 0.00 | 0.30 | 0.17 | 11 | Myofibroblast |
| COMMD6 | 0.48 | 0.00 | 0.00 | 0.28 | 0.16 | 11 | Myofibroblast |
| AP2M1 | 0.37 | 0.00 | 0.00 | 0.30 | 0.17 | 11 | Myofibroblast |
| TOMM7 | 0.32 | 0.00 | 0.00 | 0.55 | 0.40 | 11 | Myofibroblast |
| COX5B | 0.33 | 0.00 | 0.00 | 0.54 | 0.40 | 11 | Myofibroblast |
| PTMA | 0.28 | 0.00 | 0.00 | 0.96 | 0.92 | 11 | Myofibroblast |
| IDH2 | 0.41 | 0.00 | 0.00 | 0.32 | 0.20 | 11 | Myofibroblast |
| EIF1AX | 0.34 | 0.00 | 0.00 | 0.35 | 0.22 | 11 | Myofibroblast |
| MPRIP | 0.36 | 0.00 | 0.00 | 0.33 | 0.20 | 11 | Myofibroblast |
| MAP1LC3A | 0.33 | 0.00 | 0.00 | 0.33 | 0.21 | 11 | Myofibroblast |
| ADAMTS1 | 0.38 | 0.00 | 0.00 | 0.32 | 0.20 | 11 | Myofibroblast |
| ESD | 0.35 | 0.00 | 0.00 | 0.35 | 0.23 | 11 | Myofibroblast |
| DYNLL1 | 0.27 | 0.00 | 0.00 | 0.62 | 0.47 | 11 | Myofibroblast |
| EIF4B | 0.33 | 0.00 | 0.00 | 0.31 | 0.19 | 11 | Myofibroblast |
| ANXA6 | 0.31 | 0.00 | 0.00 | 0.35 | 0.22 | 11 | Myofibroblast |
| HMGN1 | 0.31 | 0.00 | 0.00 | 0.60 | 0.46 | 11 | Myofibroblast |
| OAZ2 | 0.33 | 0.00 | 0.00 | 0.44 | 0.30 | 11 | Myofibroblast |
| RPL13A | 0.81 | 0.00 | 0.00 | 0.66 | 0.59 | 11 | Myofibroblast |
| SPARCL1 | 0.42 | 0.00 | 0.00 | 0.55 | 0.41 | 11 | Myofibroblast |
| NUCKS1 | 0.28 | 0.00 | 0.00 | 0.43 | 0.29 | 11 | Myofibroblast |
| GOLIM4 | 0.31 | 0.00 | 0.00 | 0.33 | 0.20 | 11 | Myofibroblast |
| UQCR11 | 0.28 | 0.00 | 0.00 | 0.46 | 0.32 | 11 | Myofibroblast |
| RCN2 | 0.26 | 0.00 | 0.00 | 0.28 | 0.17 | 11 | Myofibroblast |
| ZYX | 0.28 | 0.00 | 0.00 | 0.33 | 0.21 | 11 | Myofibroblast |
| RBMS3 | 0.32 | 0.00 | 0.00 | 0.29 | 0.18 | 11 | Myofibroblast |
| C11orf58 | 0.27 | 0.00 | 0.00 | 0.44 | 0.31 | 11 | Myofibroblast |
| PGRMC1 | 0.33 | 0.00 | 0.00 | 0.31 | 0.19 | 11 | Myofibroblast |
| PARK7 | 0.29 | 0.00 | 0.00 | 0.41 | 0.28 | 11 | Myofibroblast |
| ENO1 | 0.26 | 0.00 | 0.00 | 0.30 | 0.19 | 11 | Myofibroblast |
| CCT3 | 0.30 | 0.00 | 0.00 | 0.29 | 0.18 | 11 | Myofibroblast |
| CUTA | 0.28 | 0.00 | 0.00 | 0.32 | 0.20 | 11 | Myofibroblast |
| MORF4L2 | 0.31 | 0.00 | 0.00 | 0.35 | 0.24 | 11 | Myofibroblast |
| EDF1 | 0.28 | 0.00 | 0.00 | 0.46 | 0.34 | 11 | Myofibroblast |
| RPS27A | 0.35 | 0.00 | 0.00 | 0.83 | 0.78 | 11 | Myofibroblast |
| NDUFV3 | 0.41 | 0.00 | 0.00 | 0.33 | 0.23 | 11 | Myofibroblast |
| EIF3E | 0.27 | 0.00 | 0.00 | 0.54 | 0.43 | 11 | Myofibroblast |
| MMP14 | 0.28 | 0.00 | 0.00 | 0.36 | 0.25 | 11 | Myofibroblast |
| SIVA1 | 0.25 | 0.00 | 0.00 | 0.25 | 0.16 | 11 | Myofibroblast |
| NDUFB4 | 0.25 | 0.00 | 0.00 | 0.37 | 0.26 | 11 | Myofibroblast |
| FBLN5 | 0.41 | 0.00 | 0.00 | 0.27 | 0.18 | 11 | Myofibroblast |
| CNN2 | 0.29 | 0.00 | 0.00 | 0.49 | 0.39 | 11 | Myofibroblast |
| ZFP36L1 | 0.26 | 0.00 | 0.00 | 0.58 | 0.46 | 11 | Myofibroblast |
| KLF9 | 0.32 | 0.00 | 0.00 | 0.25 | 0.17 | 11 | Myofibroblast |
| ISCU | 0.26 | 0.00 | 0.00 | 0.26 | 0.17 | 11 | Myofibroblast |
| TCEAL8 | 0.29 | 0.00 | 0.00 | 0.30 | 0.21 | 11 | Myofibroblast |
| RPS13 | 0.29 | 0.00 | 0.00 | 0.73 | 0.67 | 11 | Myofibroblast |
| NDUFA4 | 0.30 | 0.00 | 0.00 | 0.63 | 0.57 | 11 | Myofibroblast |
| HSPD1 | 0.28 | 0.00 | 0.00 | 0.35 | 0.26 | 11 | Myofibroblast |
| PPDPF | 0.32 | 0.00 | 0.00 | 0.31 | 0.23 | 11 | Myofibroblast |
| EIF2A | 0.26 | 0.00 | 0.00 | 0.26 | 0.18 | 11 | Myofibroblast |
| SRP14 | 0.30 | 0.00 | 0.00 | 0.41 | 0.34 | 11 | Myofibroblast |
| RPL24 | 0.38 | 0.00 | 0.00 | 0.54 | 0.50 | 11 | Myofibroblast |
| HTATSF1 | 0.26 | 0.00 | 0.00 | 0.33 | 0.25 | 11 | Myofibroblast |
| TUBA1C | 0.27 | 0.00 | 0.00 | 0.37 | 0.29 | 11 | Myofibroblast |
| HIGD1B | 3.13 | 0.00 | 0.00 | 0.87 | 0.02 | 12 | Pericyte |
| COX4I2 | 2.83 | 0.00 | 0.00 | 0.84 | 0.02 | 12 | Pericyte |
| POSTN | 2.69 | 0.00 | 0.00 | 0.70 | 0.09 | 12 | Pericyte |
| FAM162B | 2.57 | 0.00 | 0.00 | 0.60 | 0.02 | 12 | Pericyte |
| NDUFA4L2 | 2.49 | 0.00 | 0.00 | 0.67 | 0.02 | 12 | Pericyte |
| PDGFRB | 2.48 | 0.00 | 0.00 | 0.80 | 0.05 | 12 | Pericyte |
| PCDH18 | 2.25 | 0.00 | 0.00 | 0.60 | 0.03 | 12 | Pericyte |
| ITM2A | 2.07 | 0.00 | 0.00 | 0.52 | 0.07 | 12 | Pericyte |
| PDZD2 | 1.99 | 0.00 | 0.00 | 0.49 | 0.02 | 12 | Pericyte |
| PDE5A | 1.99 | 0.00 | 0.00 | 0.61 | 0.11 | 12 | Pericyte |
| ITGA1 | 1.88 | 0.00 | 0.00 | 0.67 | 0.10 | 12 | Pericyte |
| LIPG | 1.83 | 0.00 | 0.00 | 0.33 | 0.01 | 12 | Pericyte |
| RGS5 | 1.80 | 0.00 | 0.00 | 0.44 | 0.05 | 12 | Pericyte |
| TRPC6 | 1.79 | 0.00 | 0.00 | 0.46 | 0.01 | 12 | Pericyte |
| CHN1 | 1.78 | 0.00 | 0.00 | 0.41 | 0.04 | 12 | Pericyte |
| CDH2 | 1.76 | 0.00 | 0.00 | 0.35 | 0.01 | 12 | Pericyte |
| KCNK3 | 1.73 | 0.00 | 0.00 | 0.54 | 0.05 | 12 | Pericyte |
| HBEGF | 1.73 | 0.00 | 0.00 | 0.42 | 0.05 | 12 | Pericyte |
| MFGE8 | 1.72 | 0.00 | 0.00 | 0.73 | 0.16 | 12 | Pericyte |
| NOTCH3 | 1.71 | 0.00 | 0.00 | 0.52 | 0.03 | 12 | Pericyte |
| EBF1 | 1.70 | 0.00 | 0.00 | 0.52 | 0.04 | 12 | Pericyte |
| KLHL23 | 1.63 | 0.00 | 0.00 | 0.50 | 0.05 | 12 | Pericyte |
| LMCD1 | 1.57 | 0.00 | 0.00 | 0.54 | 0.07 | 12 | Pericyte |
| TBX5 | 1.54 | 0.00 | 0.00 | 0.49 | 0.07 | 12 | Pericyte |
| HEYL | 1.53 | 0.00 | 0.00 | 0.42 | 0.03 | 12 | Pericyte |
| GJC1 | 1.38 | 0.00 | 0.00 | 0.46 | 0.07 | 12 | Pericyte |
| S1PR3 | 1.37 | 0.00 | 0.00 | 0.34 | 0.04 | 12 | Pericyte |
| PAG1 | 1.36 | 0.00 | 0.00 | 0.36 | 0.03 | 12 | Pericyte |
| CSPG4 | 1.32 | 0.00 | 0.00 | 0.31 | 0.02 | 12 | Pericyte |
| VSNL1 | 1.30 | 0.00 | 0.00 | 0.30 | 0.02 | 12 | Pericyte |
| NR2F1 | 1.23 | 0.00 | 0.00 | 0.33 | 0.03 | 12 | Pericyte |
| FOXS1 | 1.18 | 0.00 | 0.00 | 0.29 | 0.01 | 12 | Pericyte |
| MAGED2 | 1.65 | 0.00 | 0.00 | 0.74 | 0.20 | 12 | Pericyte |
| NKAIN4 | 1.45 | 0.00 | 0.00 | 0.34 | 0.04 | 12 | Pericyte |
| MCAM | 1.59 | 0.00 | 0.00 | 0.61 | 0.13 | 12 | Pericyte |
| EMID1 | 1.49 | 0.00 | 0.00 | 0.34 | 0.04 | 12 | Pericyte |
| C3orf58 | 1.11 | 0.00 | 0.00 | 0.29 | 0.03 | 12 | Pericyte |
| RERG | 1.15 | 0.00 | 0.00 | 0.34 | 0.05 | 12 | Pericyte |
| PPP1R14A | 1.22 | 0.00 | 0.00 | 0.52 | 0.10 | 12 | Pericyte |
| COL4A1 | 1.36 | 0.00 | 0.00 | 0.95 | 0.50 | 12 | Pericyte |
| NCK2 | 1.00 | 0.00 | 0.00 | 0.29 | 0.03 | 12 | Pericyte |
| P2RY14 | 1.06 | 0.00 | 0.00 | 0.46 | 0.08 | 12 | Pericyte |
| CADM1 | 1.40 | 0.00 | 0.00 | 0.59 | 0.14 | 12 | Pericyte |
| RASL12 | 1.22 | 0.00 | 0.00 | 0.36 | 0.06 | 12 | Pericyte |
| CRIM1 | 1.39 | 0.00 | 0.00 | 0.41 | 0.08 | 12 | Pericyte |
| COL4A2 | 1.24 | 0.00 | 0.00 | 0.90 | 0.44 | 12 | Pericyte |
| SPARCL1 | 1.60 | 0.00 | 0.00 | 0.87 | 0.41 | 12 | Pericyte |
| ARHGAP42 | 1.02 | 0.00 | 0.00 | 0.27 | 0.03 | 12 | Pericyte |
| RARRES2 | 1.13 | 0.00 | 0.00 | 0.61 | 0.16 | 12 | Pericyte |
| EDNRA | 1.17 | 0.00 | 0.00 | 0.45 | 0.09 | 12 | Pericyte |
| LAMB1 | 1.24 | 0.00 | 0.00 | 0.55 | 0.15 | 12 | Pericyte |
| NID1 | 1.20 | 0.00 | 0.00 | 0.68 | 0.23 | 12 | Pericyte |
| FERMT2 | 1.43 | 0.00 | 0.00 | 0.82 | 0.39 | 12 | Pericyte |
| AXL | 0.96 | 0.00 | 0.00 | 0.30 | 0.05 | 12 | Pericyte |
| MYO1B | 1.17 | 0.00 | 0.00 | 0.56 | 0.16 | 12 | Pericyte |
| ARHGEF17 | 0.94 | 0.00 | 0.00 | 0.28 | 0.04 | 12 | Pericyte |
| ANGPT1 | 1.33 | 0.00 | 0.00 | 0.38 | 0.08 | 12 | Pericyte |
| SEPT4 | 1.25 | 0.00 | 0.00 | 0.67 | 0.22 | 12 | Pericyte |
| NR2F2 | 1.08 | 0.00 | 0.00 | 0.62 | 0.20 | 12 | Pericyte |
| MAPT | 0.99 | 0.00 | 0.00 | 0.27 | 0.04 | 12 | Pericyte |
| PLAC9 | 0.82 | 0.00 | 0.00 | 0.80 | 0.30 | 12 | Pericyte |
| PCOLCE | 1.15 | 0.00 | 0.00 | 0.43 | 0.10 | 12 | Pericyte |
| LTBP2 | 1.06 | 0.00 | 0.00 | 0.42 | 0.10 | 12 | Pericyte |
| GPX3 | 1.29 | 0.00 | 0.00 | 0.50 | 0.14 | 12 | Pericyte |
| NDN | 1.05 | 0.00 | 0.00 | 0.35 | 0.08 | 12 | Pericyte |
| SPARC | 0.65 | 0.00 | 0.00 | 0.99 | 0.76 | 12 | Pericyte |
| NDRG2 | 0.99 | 0.00 | 0.00 | 0.37 | 0.08 | 12 | Pericyte |
| CALD1 | 0.84 | 0.00 | 0.00 | 0.84 | 0.43 | 12 | Pericyte |
| CCRL2 | 1.00 | 0.00 | 0.00 | 0.26 | 0.05 | 12 | Pericyte |
| SEPT7 | 1.06 | 0.00 | 0.00 | 0.88 | 0.59 | 12 | Pericyte |
| RASGRP2 | 1.07 | 0.00 | 0.00 | 0.37 | 0.09 | 12 | Pericyte |
| TBX2 | 1.03 | 0.00 | 0.00 | 0.46 | 0.13 | 12 | Pericyte |
| NT5DC2 | 0.90 | 0.00 | 0.00 | 0.25 | 0.05 | 12 | Pericyte |
| PTEN | 0.99 | 0.00 | 0.00 | 0.42 | 0.12 | 12 | Pericyte |
| MYL9 | 0.96 | 0.00 | 0.00 | 0.52 | 0.17 | 12 | Pericyte |
| SEPT11 | 1.05 | 0.00 | 0.00 | 0.58 | 0.22 | 12 | Pericyte |
| ISM1 | 1.04 | 0.00 | 0.00 | 0.33 | 0.08 | 12 | Pericyte |
| CD248 | 0.72 | 0.00 | 0.00 | 0.26 | 0.05 | 12 | Pericyte |
| FZD1 | 0.84 | 0.00 | 0.00 | 0.27 | 0.06 | 12 | Pericyte |
| ISCA1 | 0.95 | 0.00 | 0.00 | 0.33 | 0.08 | 12 | Pericyte |
| MARCKS | 0.79 | 0.00 | 0.00 | 0.87 | 0.49 | 12 | Pericyte |
| GNG11 | 0.85 | 0.00 | 0.00 | 0.71 | 0.31 | 12 | Pericyte |
| TPPP3 | 1.27 | 0.00 | 0.00 | 0.41 | 0.13 | 12 | Pericyte |
| BGN | 1.02 | 0.00 | 0.00 | 0.61 | 0.26 | 12 | Pericyte |
| CD81 | 0.95 | 0.00 | 0.00 | 0.74 | 0.40 | 12 | Pericyte |
| TPM2 | 0.75 | 0.00 | 0.00 | 0.49 | 0.17 | 12 | Pericyte |
| PTP4A3 | 0.79 | 0.00 | 0.00 | 0.31 | 0.08 | 12 | Pericyte |
| MXRA8 | 0.83 | 0.00 | 0.00 | 0.47 | 0.17 | 12 | Pericyte |
| TBX4 | 0.88 | 0.00 | 0.00 | 0.25 | 0.06 | 12 | Pericyte |
| GJA4 | 0.73 | 0.00 | 0.00 | 0.30 | 0.08 | 12 | Pericyte |
| TIPARP | 1.09 | 0.00 | 0.00 | 0.29 | 0.08 | 12 | Pericyte |
| LAMA4 | 0.95 | 0.00 | 0.00 | 0.40 | 0.13 | 12 | Pericyte |
| NOSTRIN | 0.98 | 0.00 | 0.00 | 0.41 | 0.14 | 12 | Pericyte |
| CST3 | 0.81 | 0.00 | 0.00 | 0.84 | 0.59 | 12 | Pericyte |
| STOM | 0.99 | 0.00 | 0.00 | 0.30 | 0.09 | 12 | Pericyte |
| OAZ2 | 0.84 | 0.00 | 0.00 | 0.62 | 0.30 | 12 | Pericyte |
| HSD11B1 | 0.98 | 0.00 | 0.00 | 0.28 | 0.08 | 12 | Pericyte |
| TACC1 | 0.92 | 0.00 | 0.00 | 0.52 | 0.23 | 12 | Pericyte |
| DCN | 1.06 | 0.00 | 0.00 | 0.25 | 0.07 | 12 | Pericyte |
| MYH11 | 0.35 | 0.00 | 0.00 | 0.40 | 0.13 | 12 | Pericyte |
| LAPTM4A | 0.64 | 0.00 | 0.00 | 0.82 | 0.55 | 12 | Pericyte |
| CYGB | 0.93 | 0.00 | 0.00 | 0.25 | 0.07 | 12 | Pericyte |
| PTK2 | 0.72 | 0.00 | 0.00 | 0.30 | 0.09 | 12 | Pericyte |
| IGF2R | 0.96 | 0.00 | 0.00 | 0.30 | 0.09 | 12 | Pericyte |
| NES | 0.77 | 0.00 | 0.00 | 0.50 | 0.21 | 12 | Pericyte |
| C11orf96 | 0.71 | 0.00 | 0.00 | 0.27 | 0.08 | 12 | Pericyte |
| RBMX | 0.85 | 0.00 | 0.00 | 0.36 | 0.14 | 12 | Pericyte |
| ARMCX3 | 0.78 | 0.00 | 0.00 | 0.34 | 0.12 | 12 | Pericyte |
| SMTN | 0.71 | 0.00 | 0.00 | 0.28 | 0.09 | 12 | Pericyte |
| DKK3 | 0.74 | 0.00 | 0.00 | 0.36 | 0.13 | 12 | Pericyte |
| MGST3 | 0.76 | 0.00 | 0.00 | 0.35 | 0.13 | 12 | Pericyte |
| GPM6B | 0.62 | 0.00 | 0.00 | 0.34 | 0.13 | 12 | Pericyte |
| CDKN1C | 1.05 | 0.00 | 0.00 | 0.39 | 0.17 | 12 | Pericyte |
| GNB4 | 0.83 | 0.00 | 0.00 | 0.34 | 0.13 | 12 | Pericyte |
| BTBD3 | 0.77 | 0.00 | 0.00 | 0.35 | 0.14 | 12 | Pericyte |
| CRYAB | 0.82 | 0.00 | 0.00 | 0.38 | 0.16 | 12 | Pericyte |
| FAM162A | 0.81 | 0.00 | 0.00 | 0.35 | 0.15 | 12 | Pericyte |
| ITGB1 | 0.57 | 0.00 | 0.00 | 0.86 | 0.64 | 12 | Pericyte |
| EVA1B | 0.76 | 0.00 | 0.00 | 0.43 | 0.20 | 12 | Pericyte |
| TNS1 | 0.61 | 0.00 | 0.00 | 0.48 | 0.23 | 12 | Pericyte |
| G0S2 | 0.48 | 0.00 | 0.00 | 0.35 | 0.13 | 12 | Pericyte |
| GNAS | 0.85 | 0.00 | 0.00 | 0.85 | 0.66 | 12 | Pericyte |
| SGCE | 0.70 | 0.00 | 0.00 | 0.29 | 0.11 | 12 | Pericyte |
| CD63 | 0.60 | 0.00 | 0.00 | 0.58 | 0.34 | 12 | Pericyte |
| EFEMP2 | 0.62 | 0.00 | 0.00 | 0.26 | 0.09 | 12 | Pericyte |
| TFPI | 0.64 | 0.00 | 0.00 | 0.29 | 0.11 | 12 | Pericyte |
| TUBA1A | 0.52 | 0.00 | 0.00 | 0.76 | 0.52 | 12 | Pericyte |
| ZFHX3 | 0.74 | 0.00 | 0.00 | 0.37 | 0.17 | 12 | Pericyte |
| SERPING1 | 0.55 | 0.00 | 0.00 | 0.34 | 0.14 | 12 | Pericyte |
| ITM2C | 0.65 | 0.00 | 0.00 | 0.45 | 0.24 | 12 | Pericyte |
| ITGA4 | 0.55 | 0.00 | 0.00 | 0.25 | 0.09 | 12 | Pericyte |
| TGFB1I1 | 0.59 | 0.00 | 0.00 | 0.35 | 0.16 | 12 | Pericyte |
| VASP | 0.64 | 0.00 | 0.00 | 0.40 | 0.20 | 12 | Pericyte |
| EGFL6 | 1.19 | 0.00 | 0.00 | 0.27 | 0.11 | 12 | Pericyte |
| MT-ND4 | 0.45 | 0.00 | 0.00 | 0.89 | 0.76 | 12 | Pericyte |
| MT-CYB | 0.35 | 0.00 | 0.00 | 0.93 | 0.83 | 12 | Pericyte |
| CCDC80 | 0.68 | 0.00 | 0.00 | 0.30 | 0.12 | 12 | Pericyte |
| DPYSL2 | 0.58 | 0.00 | 0.00 | 0.38 | 0.18 | 12 | Pericyte |
| TSC22D1 | 0.53 | 0.00 | 0.00 | 0.58 | 0.35 | 12 | Pericyte |
| LGALS1 | 0.37 | 0.00 | 0.00 | 0.74 | 0.46 | 12 | Pericyte |
| MYLK | 0.37 | 0.00 | 0.00 | 0.54 | 0.29 | 12 | Pericyte |
| COL6A2 | 0.58 | 0.00 | 0.00 | 0.37 | 0.18 | 12 | Pericyte |
| FOXF1 | 0.48 | 0.00 | 0.00 | 0.30 | 0.13 | 12 | Pericyte |
| FILIP1L | 0.52 | 0.00 | 0.00 | 0.55 | 0.33 | 12 | Pericyte |
| EID1 | 0.52 | 0.00 | 0.00 | 0.53 | 0.33 | 12 | Pericyte |
| MT-ATP6 | 0.31 | 0.00 | 0.00 | 0.94 | 0.85 | 12 | Pericyte |
| PPIC | 0.49 | 0.00 | 0.00 | 0.59 | 0.38 | 12 | Pericyte |
| COL6A3 | 0.54 | 0.00 | 0.00 | 0.29 | 0.13 | 12 | Pericyte |
| MT-ND3 | 0.54 | 0.00 | 0.00 | 0.68 | 0.51 | 12 | Pericyte |
| COL5A2 | 0.49 | 0.00 | 0.00 | 0.37 | 0.19 | 12 | Pericyte |
| LOXL2 | 0.38 | 0.00 | 0.00 | 0.29 | 0.13 | 12 | Pericyte |
| SERPINH1 | 0.53 | 0.00 | 0.00 | 0.63 | 0.42 | 12 | Pericyte |
| GINM1 | 0.62 | 0.00 | 0.00 | 0.41 | 0.23 | 12 | Pericyte |
| MYH9 | 0.45 | 0.00 | 0.00 | 0.54 | 0.34 | 12 | Pericyte |
| RPL29 | 0.90 | 0.00 | 0.00 | 0.35 | 0.20 | 12 | Pericyte |
| LIMS3 | 0.73 | 0.00 | 0.00 | 0.42 | 0.26 | 12 | Pericyte |
| CBFA2T3 | 0.57 | 0.00 | 0.00 | 0.29 | 0.14 | 12 | Pericyte |
| CIRBP | 0.54 | 0.00 | 0.00 | 0.43 | 0.26 | 12 | Pericyte |
| PIK3R1 | 0.42 | 0.00 | 0.00 | 0.26 | 0.12 | 12 | Pericyte |
| RPL7A | 0.58 | 0.00 | 0.00 | 0.58 | 0.44 | 12 | Pericyte |
| TMEM59 | 0.42 | 0.00 | 0.00 | 0.61 | 0.43 | 12 | Pericyte |
| RHOC | 0.47 | 0.00 | 0.00 | 0.42 | 0.24 | 12 | Pericyte |
| ILK | 0.48 | 0.00 | 0.00 | 0.42 | 0.24 | 12 | Pericyte |
| GAPDH | 0.77 | 0.00 | 0.00 | 0.34 | 0.19 | 12 | Pericyte |
| FAM104B | 0.65 | 0.00 | 0.00 | 0.31 | 0.17 | 12 | Pericyte |
| CALM2 | 0.43 | 0.00 | 0.00 | 0.75 | 0.66 | 12 | Pericyte |
| CAMK2N1 | 0.53 | 0.00 | 0.00 | 0.29 | 0.14 | 12 | Pericyte |
| MAP1LC3A | 0.47 | 0.00 | 0.00 | 0.37 | 0.21 | 12 | Pericyte |
| RHOB | 0.46 | 0.00 | 0.00 | 0.26 | 0.12 | 12 | Pericyte |
| CRIP2 | 0.32 | 0.00 | 0.00 | 0.65 | 0.43 | 12 | Pericyte |
| EIF4A2 | 0.45 | 0.00 | 0.00 | 0.52 | 0.35 | 12 | Pericyte |
| NPTN | 0.66 | 0.00 | 0.00 | 0.34 | 0.20 | 12 | Pericyte |
| PHLDA1 | 0.41 | 0.00 | 0.00 | 0.37 | 0.20 | 12 | Pericyte |
| ACAA2 | 0.63 | 0.00 | 0.00 | 0.29 | 0.16 | 12 | Pericyte |
| TGFB2 | 0.63 | 0.00 | 0.00 | 0.27 | 0.14 | 12 | Pericyte |
| MT-ND2 | 0.30 | 0.00 | 0.00 | 0.83 | 0.71 | 12 | Pericyte |
| TPM4 | 0.45 | 0.00 | 0.00 | 0.67 | 0.51 | 12 | Pericyte |
| RHOJ | 0.59 | 0.00 | 0.00 | 0.32 | 0.18 | 12 | Pericyte |
| TSPAN3 | 0.42 | 0.00 | 0.00 | 0.31 | 0.17 | 12 | Pericyte |
| SSR3 | 0.61 | 0.00 | 0.00 | 0.44 | 0.30 | 12 | Pericyte |
| TXNIP | 0.71 | 0.00 | 0.00 | 0.41 | 0.27 | 12 | Pericyte |
| GPX8 | 0.49 | 0.00 | 0.00 | 0.26 | 0.13 | 12 | Pericyte |
| CD302 | 0.48 | 0.00 | 0.00 | 0.28 | 0.15 | 12 | Pericyte |
| AP2M1 | 0.50 | 0.00 | 0.00 | 0.30 | 0.17 | 12 | Pericyte |
| FCGRT | 0.37 | 0.00 | 0.00 | 0.26 | 0.13 | 12 | Pericyte |
| ARL3 | 0.41 | 0.00 | 0.00 | 0.25 | 0.13 | 12 | Pericyte |
| PPP1R12A | 0.36 | 0.00 | 0.00 | 0.43 | 0.26 | 12 | Pericyte |
| RCN2 | 0.38 | 0.00 | 0.00 | 0.31 | 0.17 | 12 | Pericyte |
| ZBTB38 | 0.43 | 0.00 | 0.00 | 0.26 | 0.14 | 12 | Pericyte |
| NFIA | 0.54 | 0.00 | 0.00 | 0.39 | 0.24 | 12 | Pericyte |
| PSIP1 | 0.40 | 0.00 | 0.00 | 0.29 | 0.16 | 12 | Pericyte |
| ANXA6 | 0.46 | 0.00 | 0.00 | 0.36 | 0.22 | 12 | Pericyte |
| CD151 | 0.38 | 0.00 | 0.00 | 0.34 | 0.19 | 12 | Pericyte |
| MAT2A | 0.61 | 0.00 | 0.00 | 0.38 | 0.25 | 12 | Pericyte |
| TMEM204 | 0.28 | 0.00 | 0.00 | 0.30 | 0.16 | 12 | Pericyte |
| KANK2 | 0.39 | 0.00 | 0.00 | 0.25 | 0.13 | 12 | Pericyte |
| PPIA | 0.48 | 0.00 | 0.00 | 0.35 | 0.21 | 12 | Pericyte |
| MFAP2 | 0.34 | 0.00 | 0.00 | 0.53 | 0.35 | 12 | Pericyte |
| TCF4 | 0.26 | 0.00 | 0.00 | 0.61 | 0.42 | 12 | Pericyte |
| SUMO2 | 0.42 | 0.00 | 0.00 | 0.48 | 0.34 | 12 | Pericyte |
| COMMD6 | 0.43 | 0.00 | 0.00 | 0.29 | 0.16 | 12 | Pericyte |
| CCNI | 0.49 | 0.00 | 0.00 | 0.33 | 0.20 | 12 | Pericyte |
| SLC12A2 | 0.39 | 0.00 | 0.00 | 0.27 | 0.15 | 12 | Pericyte |
| MT-CO2 | 0.32 | 0.00 | 0.00 | 0.84 | 0.79 | 12 | Pericyte |
| RELL1 | 0.55 | 0.00 | 0.00 | 0.30 | 0.18 | 12 | Pericyte |
| PJA2 | 0.44 | 0.00 | 0.00 | 0.26 | 0.14 | 12 | Pericyte |
| EEF1D | 0.49 | 0.00 | 0.00 | 0.43 | 0.30 | 12 | Pericyte |
| NUCB1 | 0.39 | 0.00 | 0.00 | 0.28 | 0.16 | 12 | Pericyte |
| ATRAID | 0.36 | 0.00 | 0.00 | 0.32 | 0.19 | 12 | Pericyte |
| CDC42BPA | 0.42 | 0.00 | 0.00 | 0.26 | 0.14 | 12 | Pericyte |
| PPFIBP1 | 0.46 | 0.00 | 0.00 | 0.25 | 0.14 | 12 | Pericyte |
| PPDPF | 0.44 | 0.00 | 0.00 | 0.35 | 0.23 | 12 | Pericyte |
| APP | 0.29 | 0.00 | 0.00 | 0.77 | 0.60 | 12 | Pericyte |
| UACA | 0.44 | 0.00 | 0.00 | 0.26 | 0.15 | 12 | Pericyte |
| PREX2 | 0.46 | 0.00 | 0.00 | 0.29 | 0.17 | 12 | Pericyte |
| C12orf57 | 0.36 | 0.00 | 0.00 | 0.28 | 0.16 | 12 | Pericyte |
| RSU1 | 0.45 | 0.00 | 0.00 | 0.33 | 0.21 | 12 | Pericyte |
| RPL24 | 0.33 | 0.00 | 0.00 | 0.61 | 0.49 | 12 | Pericyte |
| GSTP1 | 0.38 | 0.00 | 0.00 | 0.37 | 0.24 | 12 | Pericyte |
| HNRNPA1 | 0.57 | 0.00 | 0.00 | 0.42 | 0.31 | 12 | Pericyte |
| LEPROT | 0.33 | 0.00 | 0.00 | 0.48 | 0.34 | 12 | Pericyte |
| RAP2A | 0.40 | 0.00 | 0.00 | 0.26 | 0.15 | 12 | Pericyte |
| RAB11B | 0.33 | 0.00 | 0.00 | 0.34 | 0.22 | 12 | Pericyte |
| TIMP3 | 0.49 | 0.00 | 0.00 | 0.51 | 0.38 | 12 | Pericyte |
| KTN1 | 0.40 | 0.00 | 0.00 | 0.52 | 0.39 | 12 | Pericyte |
| COL6A1 | 0.29 | 0.00 | 0.00 | 0.35 | 0.22 | 12 | Pericyte |
| PPIB | 0.30 | 0.00 | 0.00 | 0.59 | 0.46 | 12 | Pericyte |
| CCND2 | 0.38 | 0.00 | 0.00 | 0.40 | 0.28 | 12 | Pericyte |
| TRMT112 | 0.30 | 0.00 | 0.00 | 0.29 | 0.18 | 12 | Pericyte |
| MEF2C | 0.29 | 0.00 | 0.00 | 0.30 | 0.18 | 12 | Pericyte |
| RABAC1 | 0.29 | 0.00 | 0.00 | 0.43 | 0.30 | 12 | Pericyte |
| COMT | 0.31 | 0.00 | 0.00 | 0.26 | 0.16 | 12 | Pericyte |
| GOLIM4 | 0.31 | 0.00 | 0.00 | 0.32 | 0.20 | 12 | Pericyte |
| FKBP8 | 0.30 | 0.00 | 0.00 | 0.25 | 0.15 | 12 | Pericyte |
| TSC22D3 | 0.36 | 0.00 | 0.00 | 0.34 | 0.23 | 12 | Pericyte |
| UQCRC1 | 0.27 | 0.00 | 0.00 | 0.30 | 0.19 | 12 | Pericyte |
| NDUFS5 | 0.26 | 0.00 | 0.00 | 0.39 | 0.27 | 12 | Pericyte |
| ECH1 | 0.26 | 0.00 | 0.00 | 0.28 | 0.17 | 12 | Pericyte |
| BRD3 | 0.40 | 0.00 | 0.00 | 0.35 | 0.24 | 12 | Pericyte |
| RBMS1 | 0.31 | 0.00 | 0.00 | 0.51 | 0.38 | 12 | Pericyte |
| CUTA | 0.31 | 0.00 | 0.00 | 0.31 | 0.20 | 12 | Pericyte |
| ALDH2 | 0.35 | 0.00 | 0.00 | 0.44 | 0.32 | 12 | Pericyte |
| DRAP1 | 0.39 | 0.00 | 0.00 | 0.36 | 0.25 | 12 | Pericyte |
| SRP14 | 0.41 | 0.00 | 0.00 | 0.43 | 0.34 | 12 | Pericyte |
| DYNLL1 | 0.25 | 0.00 | 0.00 | 0.59 | 0.47 | 12 | Pericyte |
| CD164 | 0.28 | 0.00 | 0.00 | 0.25 | 0.16 | 12 | Pericyte |
| KIAA0040 | 0.27 | 0.00 | 0.00 | 0.27 | 0.18 | 12 | Pericyte |
| EIF3L | 0.32 | 0.00 | 0.00 | 0.25 | 0.16 | 12 | Pericyte |
| RNPS1 | 0.28 | 0.00 | 0.00 | 0.25 | 0.16 | 12 | Pericyte |
| PPP2R1A | 0.28 | 0.00 | 0.00 | 0.27 | 0.17 | 12 | Pericyte |
| DYNC1I2 | 0.26 | 0.00 | 0.00 | 0.61 | 0.48 | 12 | Pericyte |
| CALM3 | 0.26 | 0.00 | 0.00 | 0.30 | 0.21 | 12 | Pericyte |
| SERINC1 | 0.29 | 0.00 | 0.00 | 0.39 | 0.29 | 12 | Pericyte |
| MIF | 0.33 | 0.00 | 0.00 | 0.32 | 0.23 | 12 | Pericyte |
| PRKAR1A | 0.27 | 0.00 | 0.00 | 0.43 | 0.33 | 12 | Pericyte |
| RPL23A | 0.25 | 0.00 | 0.00 | 0.74 | 0.69 | 12 | Pericyte |
| HNRNPA0 | 0.28 | 0.00 | 0.00 | 0.34 | 0.24 | 12 | Pericyte |
| C5orf15 | 0.28 | 0.00 | 0.00 | 0.26 | 0.17 | 12 | Pericyte |
| MPRIP | 0.28 | 0.00 | 0.00 | 0.29 | 0.20 | 12 | Pericyte |
| RAB18 | 0.26 | 0.00 | 0.00 | 0.27 | 0.19 | 12 | Pericyte |
| PFDN1 | 0.32 | 0.00 | 0.00 | 0.35 | 0.26 | 12 | Pericyte |
| EIF4G3 | 0.37 | 0.00 | 0.02 | 0.32 | 0.25 | 12 | Pericyte |
| ALOX5AP | 1.93 | 0.00 | 0.00 | 0.43 | 0.09 | 13 | Myeloid |
| SAMSN1 | 1.42 | 0.00 | 0.00 | 0.31 | 0.06 | 13 | Myeloid |
| LAPTM5 | 1.19 | 0.00 | 0.00 | 0.48 | 0.13 | 13 | Myeloid |
| CCR2 | 1.20 | 0.00 | 0.00 | 0.33 | 0.07 | 13 | Myeloid |
| SELPLG | 1.10 | 0.00 | 0.00 | 0.26 | 0.05 | 13 | Myeloid |
| TYROBP | 1.29 | 0.00 | 0.00 | 0.41 | 0.12 | 13 | Myeloid |
| RAC2 | 1.08 | 0.00 | 0.00 | 0.36 | 0.10 | 13 | Myeloid |
| FCER1G | 1.00 | 0.00 | 0.00 | 0.38 | 0.11 | 13 | Myeloid |
| PLEK | 1.27 | 0.00 | 0.00 | 0.27 | 0.07 | 13 | Myeloid |
| SRGN | 1.48 | 0.00 | 0.00 | 0.60 | 0.32 | 13 | Myeloid |
| ARHGDIB | 1.01 | 0.00 | 0.00 | 0.45 | 0.17 | 13 | Myeloid |
| CD53 | 0.99 | 0.00 | 0.00 | 0.30 | 0.09 | 13 | Myeloid |
| CD52 | 0.75 | 0.00 | 0.00 | 0.48 | 0.19 | 13 | Myeloid |
| ARHGAP30 | 0.91 | 0.00 | 0.00 | 0.30 | 0.10 | 13 | Myeloid |
| LCP1 | 0.95 | 0.00 | 0.00 | 0.43 | 0.21 | 13 | Myeloid |
| TNFAIP8 | 1.02 | 0.00 | 0.00 | 0.27 | 0.10 | 13 | Myeloid |
| PTPRC | 0.64 | 0.00 | 0.00 | 0.35 | 0.14 | 13 | Myeloid |
| PLAC8 | 0.60 | 0.00 | 0.00 | 0.38 | 0.16 | 13 | Myeloid |
| APBB1IP | 0.72 | 0.00 | 0.00 | 0.25 | 0.09 | 13 | Myeloid |
| GATA2 | 1.32 | 0.00 | 0.00 | 0.31 | 0.14 | 13 | Myeloid |
| COTL1 | 0.84 | 0.00 | 0.00 | 0.36 | 0.17 | 13 | Myeloid |
| SEC11C | 0.82 | 0.00 | 0.00 | 0.34 | 0.16 | 13 | Myeloid |
| PTPN18 | 0.77 | 0.00 | 0.00 | 0.30 | 0.13 | 13 | Myeloid |
| SH3BGRL3 | 0.81 | 0.00 | 0.00 | 0.54 | 0.36 | 13 | Myeloid |
| GMFG | 0.79 | 0.00 | 0.00 | 0.32 | 0.15 | 13 | Myeloid |
| TALDO1 | 0.89 | 0.00 | 0.00 | 0.44 | 0.27 | 13 | Myeloid |
| GPX1 | 0.89 | 0.00 | 0.00 | 0.60 | 0.44 | 13 | Myeloid |
| LGALS3 | 0.67 | 0.00 | 0.00 | 0.33 | 0.16 | 13 | Myeloid |
| CREG1 | 0.84 | 0.00 | 0.00 | 0.33 | 0.17 | 13 | Myeloid |
| LGALS1 | 0.54 | 0.00 | 0.00 | 0.63 | 0.46 | 13 | Myeloid |
| TMSB4X | 0.63 | 0.00 | 0.00 | 0.87 | 0.90 | 13 | Myeloid |
| CRLF3 | 0.76 | 0.00 | 0.00 | 0.25 | 0.13 | 13 | Myeloid |
| LRRC58 | 0.55 | 0.00 | 0.00 | 0.85 | 0.85 | 13 | Myeloid |
| LIMD2 | 0.61 | 0.00 | 0.00 | 0.26 | 0.15 | 13 | Myeloid |
| ATG3 | 0.95 | 0.00 | 0.00 | 0.28 | 0.18 | 13 | Myeloid |
| ARPC1B | 0.53 | 0.00 | 0.00 | 0.48 | 0.37 | 13 | Myeloid |
| ARPC5 | 0.54 | 0.00 | 0.00 | 0.60 | 0.58 | 13 | Myeloid |
| CRIP1 | 0.46 | 0.00 | 0.00 | 0.58 | 0.48 | 13 | Myeloid |
| S100A10 | 0.61 | 0.00 | 0.00 | 0.49 | 0.41 | 13 | Myeloid |
| TAOK3 | 0.55 | 0.00 | 0.00 | 0.29 | 0.19 | 13 | Myeloid |
| ARPC2 | 0.41 | 0.00 | 0.00 | 0.64 | 0.62 | 13 | Myeloid |
| RPLP1 | 0.30 | 0.00 | 0.00 | 0.84 | 0.91 | 13 | Myeloid |
| CTSD | 0.56 | 0.00 | 0.00 | 0.34 | 0.25 | 13 | Myeloid |
| LDHA | 0.59 | 0.00 | 0.00 | 0.39 | 0.32 | 13 | Myeloid |
| WDR89 | 0.28 | 0.00 | 0.00 | 0.82 | 0.82 | 13 | Myeloid |
| HBA2 | 2.03 | 0.00 | 0.00 | 0.67 | 0.65 | 13 | Myeloid |
| JUNB | 0.67 | 0.00 | 0.00 | 0.29 | 0.21 | 13 | Myeloid |
| CD47 | 0.43 | 0.00 | 0.00 | 0.48 | 0.41 | 13 | Myeloid |
| PFN1 | 0.46 | 0.00 | 0.00 | 0.52 | 0.49 | 13 | Myeloid |
| KLF6 | 0.77 | 0.00 | 0.00 | 0.41 | 0.35 | 13 | Myeloid |
| NCOR1 | 0.44 | 0.00 | 0.00 | 0.46 | 0.40 | 13 | Myeloid |
| CYBA | 0.27 | 0.00 | 0.00 | 0.40 | 0.32 | 13 | Myeloid |
| TSPO | 0.46 | 0.00 | 0.00 | 0.34 | 0.27 | 13 | Myeloid |
| ALDOA | 0.54 | 0.00 | 0.00 | 0.30 | 0.23 | 13 | Myeloid |
| ADIPOR1 | 0.50 | 0.00 | 0.00 | 0.30 | 0.22 | 13 | Myeloid |
| MBNL1 | 0.52 | 0.00 | 0.00 | 0.32 | 0.26 | 13 | Myeloid |
| SAMHD1 | 0.36 | 0.00 | 0.03 | 0.26 | 0.19 | 13 | Myeloid |
| HMGB2 | 0.48 | 0.00 | 0.03 | 0.43 | 0.38 | 13 | Myeloid |
| ANXA1 | 0.44 | 0.00 | 0.04 | 0.27 | 0.20 | 13 | Myeloid |
| AKAP13 | 0.46 | 0.00 | 0.04 | 0.33 | 0.27 | 13 | Myeloid |
| LRRFIP1 | 0.40 | 0.00 | 0.05 | 0.32 | 0.26 | 13 | Myeloid |
| LUM | 3.55 | 0.00 | 0.00 | 0.84 | 0.03 | 14 | Matrix Fibroblast |
| A2M | 2.95 | 0.00 | 0.00 | 0.66 | 0.05 | 14 | Matrix Fibroblast |
| DCN | 2.67 | 0.00 | 0.00 | 0.87 | 0.06 | 14 | Matrix Fibroblast |
| FBLN1 | 2.60 | 0.00 | 0.00 | 0.85 | 0.07 | 14 | Matrix Fibroblast |
| RARRES2 | 2.37 | 0.00 | 0.00 | 0.88 | 0.16 | 14 | Matrix Fibroblast |
| RPL29 | 2.23 | 0.00 | 0.00 | 0.97 | 0.19 | 14 | Matrix Fibroblast |
| RARRES1 | 2.03 | 0.00 | 0.00 | 0.48 | 0.01 | 14 | Matrix Fibroblast |
| C7 | 2.02 | 0.00 | 0.00 | 0.70 | 0.02 | 14 | Matrix Fibroblast |
| COL6A3 | 1.96 | 0.00 | 0.00 | 0.80 | 0.12 | 14 | Matrix Fibroblast |
| TIMP1 | 1.89 | 0.00 | 0.00 | 0.76 | 0.08 | 14 | Matrix Fibroblast |
| PLEKHH2 | 1.78 | 0.00 | 0.00 | 0.66 | 0.06 | 14 | Matrix Fibroblast |
| GPC3 | 1.72 | 0.00 | 0.00 | 0.86 | 0.17 | 14 | Matrix Fibroblast |
| NR2F1 | 1.71 | 0.00 | 0.00 | 0.59 | 0.03 | 14 | Matrix Fibroblast |
| C1R | 1.71 | 0.00 | 0.00 | 0.56 | 0.04 | 14 | Matrix Fibroblast |
| SLC40A1 | 1.69 | 0.00 | 0.00 | 0.61 | 0.03 | 14 | Matrix Fibroblast |
| HSPA1A | 1.67 | 0.00 | 0.00 | 0.61 | 0.05 | 14 | Matrix Fibroblast |
| WNT2 | 1.66 | 0.00 | 0.00 | 0.64 | 0.05 | 14 | Matrix Fibroblast |
| SOD3 | 1.65 | 0.00 | 0.00 | 0.62 | 0.06 | 14 | Matrix Fibroblast |
| INMT | 1.64 | 0.00 | 0.00 | 0.66 | 0.05 | 14 | Matrix Fibroblast |
| SNRPN | 1.59 | 0.00 | 0.00 | 0.61 | 0.04 | 14 | Matrix Fibroblast |
| UBL5 | 1.57 | 0.00 | 0.00 | 0.72 | 0.06 | 14 | Matrix Fibroblast |
| AOC3 | 1.56 | 0.00 | 0.00 | 0.57 | 0.03 | 14 | Matrix Fibroblast |
| ABCA8 | 1.52 | 0.00 | 0.00 | 0.48 | 0.01 | 14 | Matrix Fibroblast |
| CCBE1 | 1.44 | 0.00 | 0.00 | 0.46 | 0.01 | 14 | Matrix Fibroblast |
| EPHX1 | 1.33 | 0.00 | 0.00 | 0.47 | 0.04 | 14 | Matrix Fibroblast |
| METTL7A | 1.33 | 0.00 | 0.00 | 0.42 | 0.02 | 14 | Matrix Fibroblast |
| ABCA6 | 1.32 | 0.00 | 0.00 | 0.38 | 0.01 | 14 | Matrix Fibroblast |
| RSPO2 | 1.31 | 0.00 | 0.00 | 0.43 | 0.01 | 14 | Matrix Fibroblast |
| GYPC | 1.30 | 0.00 | 0.00 | 0.56 | 0.06 | 14 | Matrix Fibroblast |
| CCL2 | 1.29 | 0.00 | 0.00 | 0.32 | 0.02 | 14 | Matrix Fibroblast |
| COX7A1 | 1.20 | 0.00 | 0.00 | 0.45 | 0.03 | 14 | Matrix Fibroblast |
| FGFR4 | 1.19 | 0.00 | 0.00 | 0.46 | 0.03 | 14 | Matrix Fibroblast |
| GDF10 | 1.19 | 0.00 | 0.00 | 0.35 | 0.01 | 14 | Matrix Fibroblast |
| EIF2S3 | 1.10 | 0.00 | 0.00 | 0.36 | 0.02 | 14 | Matrix Fibroblast |
| PYURF | 1.09 | 0.00 | 0.00 | 0.44 | 0.03 | 14 | Matrix Fibroblast |
| CLEC11A | 1.07 | 0.00 | 0.00 | 0.36 | 0.02 | 14 | Matrix Fibroblast |
| MYL6B | 1.07 | 0.00 | 0.00 | 0.38 | 0.02 | 14 | Matrix Fibroblast |
| SAT2 | 1.05 | 0.00 | 0.00 | 0.43 | 0.04 | 14 | Matrix Fibroblast |
| QPRT | 0.99 | 0.00 | 0.00 | 0.31 | 0.01 | 14 | Matrix Fibroblast |
| MOXD1 | 0.91 | 0.00 | 0.00 | 0.26 | 0.01 | 14 | Matrix Fibroblast |
| ITGAE | 0.81 | 0.00 | 0.00 | 0.26 | 0.01 | 14 | Matrix Fibroblast |
| CCNB1IP1 | 0.77 | 0.00 | 0.00 | 0.26 | 0.01 | 14 | Matrix Fibroblast |
| ENAH | 1.21 | 0.00 | 0.00 | 0.48 | 0.05 | 14 | Matrix Fibroblast |
| TMEM98 | 1.28 | 0.00 | 0.00 | 0.57 | 0.07 | 14 | Matrix Fibroblast |
| HNRNPA1 | 1.92 | 0.00 | 0.00 | 0.96 | 0.30 | 14 | Matrix Fibroblast |
| LITAF | 1.72 | 0.00 | 0.00 | 0.70 | 0.12 | 14 | Matrix Fibroblast |
| MDFI | 1.04 | 0.00 | 0.00 | 0.39 | 0.04 | 14 | Matrix Fibroblast |
| CSRP1 | 1.81 | 0.00 | 0.00 | 0.83 | 0.19 | 14 | Matrix Fibroblast |
| IFITM2 | 1.15 | 0.00 | 0.00 | 0.60 | 0.08 | 14 | Matrix Fibroblast |
| CYBRD1 | 0.85 | 0.00 | 0.00 | 0.28 | 0.02 | 14 | Matrix Fibroblast |
| XRCC6 | 1.11 | 0.00 | 0.00 | 0.51 | 0.06 | 14 | Matrix Fibroblast |
| CCNI | 1.55 | 0.00 | 0.00 | 0.86 | 0.19 | 14 | Matrix Fibroblast |
| CD59 | 1.22 | 0.00 | 0.00 | 0.54 | 0.07 | 14 | Matrix Fibroblast |
| PPP1R14A | 1.46 | 0.00 | 0.00 | 0.66 | 0.10 | 14 | Matrix Fibroblast |
| TCF21 | 1.43 | 0.00 | 0.00 | 0.85 | 0.17 | 14 | Matrix Fibroblast |
| HMGN3 | 1.27 | 0.00 | 0.00 | 0.54 | 0.07 | 14 | Matrix Fibroblast |
| HSPB6 | 0.85 | 0.00 | 0.00 | 0.27 | 0.02 | 14 | Matrix Fibroblast |
| OLFML3 | 1.39 | 0.00 | 0.00 | 0.63 | 0.09 | 14 | Matrix Fibroblast |
| ZNF428 | 1.03 | 0.00 | 0.00 | 0.44 | 0.05 | 14 | Matrix Fibroblast |
| CIRBP | 1.71 | 0.00 | 0.00 | 0.90 | 0.25 | 14 | Matrix Fibroblast |
| LDHB | 1.35 | 0.00 | 0.00 | 0.66 | 0.11 | 14 | Matrix Fibroblast |
| BAG2 | 0.93 | 0.00 | 0.00 | 0.33 | 0.03 | 14 | Matrix Fibroblast |
| ZFP36L2 | 1.65 | 0.00 | 0.00 | 0.71 | 0.14 | 14 | Matrix Fibroblast |
| RPL7A | 1.80 | 0.00 | 0.00 | 0.98 | 0.43 | 14 | Matrix Fibroblast |
| DST | 1.50 | 0.00 | 0.00 | 0.75 | 0.15 | 14 | Matrix Fibroblast |
| MGST3 | 1.35 | 0.00 | 0.00 | 0.70 | 0.13 | 14 | Matrix Fibroblast |
| RPL13A | 2.12 | 0.00 | 0.00 | 1.00 | 0.59 | 14 | Matrix Fibroblast |
| COL6A2 | 1.49 | 0.00 | 0.00 | 0.79 | 0.18 | 14 | Matrix Fibroblast |
| MMP23B | 0.93 | 0.00 | 0.00 | 0.40 | 0.04 | 14 | Matrix Fibroblast |
| MEOX2 | 1.11 | 0.00 | 0.00 | 0.59 | 0.09 | 14 | Matrix Fibroblast |
| ZSCAN18 | 0.81 | 0.00 | 0.00 | 0.29 | 0.02 | 14 | Matrix Fibroblast |
| PCOLCE | 1.23 | 0.00 | 0.00 | 0.63 | 0.10 | 14 | Matrix Fibroblast |
| LRRC17 | 1.14 | 0.00 | 0.00 | 0.47 | 0.06 | 14 | Matrix Fibroblast |
| FAT3 | 0.82 | 0.00 | 0.00 | 0.25 | 0.02 | 14 | Matrix Fibroblast |
| PTGER1 | 0.79 | 0.00 | 0.00 | 0.26 | 0.02 | 14 | Matrix Fibroblast |
| MSRB2 | 0.97 | 0.00 | 0.00 | 0.39 | 0.04 | 14 | Matrix Fibroblast |
| PPP1R3C | 1.29 | 0.00 | 0.00 | 0.40 | 0.05 | 14 | Matrix Fibroblast |
| NBEAL1 | 1.31 | 0.00 | 0.00 | 0.66 | 0.13 | 14 | Matrix Fibroblast |
| VCAN | 1.16 | 0.00 | 0.00 | 0.69 | 0.13 | 14 | Matrix Fibroblast |
| DEGS1 | 0.76 | 0.00 | 0.00 | 0.28 | 0.02 | 14 | Matrix Fibroblast |
| NME4 | 0.87 | 0.00 | 0.00 | 0.33 | 0.03 | 14 | Matrix Fibroblast |
| IFI27L2 | 0.96 | 0.00 | 0.00 | 0.32 | 0.03 | 14 | Matrix Fibroblast |
| LBH | 1.36 | 0.00 | 0.00 | 0.83 | 0.20 | 14 | Matrix Fibroblast |
| PPIA | 1.25 | 0.00 | 0.00 | 0.84 | 0.21 | 14 | Matrix Fibroblast |
| SNCA | 0.84 | 0.00 | 0.00 | 0.27 | 0.02 | 14 | Matrix Fibroblast |
| COMMD6 | 1.23 | 0.00 | 0.00 | 0.74 | 0.16 | 14 | Matrix Fibroblast |
| RBM42 | 0.81 | 0.00 | 0.00 | 0.27 | 0.02 | 14 | Matrix Fibroblast |
| EIF3L | 1.27 | 0.00 | 0.00 | 0.72 | 0.16 | 14 | Matrix Fibroblast |
| TMEM230 | 1.08 | 0.00 | 0.00 | 0.49 | 0.07 | 14 | Matrix Fibroblast |
| NDN | 1.03 | 0.00 | 0.00 | 0.50 | 0.08 | 14 | Matrix Fibroblast |
| HSPB1 | 1.11 | 0.00 | 0.00 | 0.76 | 0.17 | 14 | Matrix Fibroblast |
| PPDPF | 1.28 | 0.00 | 0.00 | 0.83 | 0.22 | 14 | Matrix Fibroblast |
| COL6A1 | 1.35 | 0.00 | 0.00 | 0.82 | 0.22 | 14 | Matrix Fibroblast |
| RPL7 | 1.35 | 0.00 | 0.00 | 1.00 | 0.77 | 14 | Matrix Fibroblast |
| FMOD | 0.78 | 0.00 | 0.00 | 0.26 | 0.02 | 14 | Matrix Fibroblast |
| CD82 | 1.05 | 0.00 | 0.00 | 0.38 | 0.04 | 14 | Matrix Fibroblast |
| LGALS3BP | 0.98 | 0.00 | 0.00 | 0.41 | 0.05 | 14 | Matrix Fibroblast |
| EMILIN1 | 1.14 | 0.00 | 0.00 | 0.58 | 0.10 | 14 | Matrix Fibroblast |
| LURAP1L | 1.00 | 0.00 | 0.00 | 0.41 | 0.05 | 14 | Matrix Fibroblast |
| GAPDH | 1.16 | 0.00 | 0.00 | 0.79 | 0.19 | 14 | Matrix Fibroblast |
| FBXO17 | 0.81 | 0.00 | 0.00 | 0.29 | 0.03 | 14 | Matrix Fibroblast |
| EEF1D | 1.30 | 0.00 | 0.00 | 0.91 | 0.29 | 14 | Matrix Fibroblast |
| EFEMP2 | 1.06 | 0.00 | 0.00 | 0.54 | 0.09 | 14 | Matrix Fibroblast |
| FGF7 | 1.03 | 0.00 | 0.00 | 0.39 | 0.05 | 14 | Matrix Fibroblast |
| SRP14 | 1.31 | 0.00 | 0.00 | 0.92 | 0.33 | 14 | Matrix Fibroblast |
| RPS27A | 1.24 | 0.00 | 0.00 | 1.00 | 0.78 | 14 | Matrix Fibroblast |
| RPL24 | 1.33 | 0.00 | 0.00 | 0.98 | 0.49 | 14 | Matrix Fibroblast |
| COL1A1 | 1.18 | 0.00 | 0.00 | 0.90 | 0.28 | 14 | Matrix Fibroblast |
| ELN | 1.14 | 0.00 | 0.00 | 0.75 | 0.18 | 14 | Matrix Fibroblast |
| MIF | 1.23 | 0.00 | 0.00 | 0.81 | 0.22 | 14 | Matrix Fibroblast |
| PFN2 | 0.86 | 0.00 | 0.00 | 0.32 | 0.04 | 14 | Matrix Fibroblast |
| NDFIP1 | 1.06 | 0.00 | 0.00 | 0.56 | 0.10 | 14 | Matrix Fibroblast |
| RPL23A | 1.29 | 0.00 | 0.00 | 0.99 | 0.69 | 14 | Matrix Fibroblast |
| MFAP4 | 1.08 | 0.00 | 0.00 | 0.88 | 0.26 | 14 | Matrix Fibroblast |
| XRCC5 | 0.94 | 0.00 | 0.00 | 0.40 | 0.06 | 14 | Matrix Fibroblast |
| SERPING1 | 0.96 | 0.00 | 0.00 | 0.66 | 0.14 | 14 | Matrix Fibroblast |
| LTBP4 | 1.10 | 0.00 | 0.00 | 0.59 | 0.12 | 14 | Matrix Fibroblast |
| CAMK2N1 | 1.11 | 0.00 | 0.00 | 0.64 | 0.14 | 14 | Matrix Fibroblast |
| TFPI | 0.98 | 0.00 | 0.00 | 0.58 | 0.11 | 14 | Matrix Fibroblast |
| DYNC2LI1 | 0.81 | 0.00 | 0.00 | 0.29 | 0.03 | 14 | Matrix Fibroblast |
| LMO4 | 1.38 | 0.00 | 0.00 | 0.81 | 0.25 | 14 | Matrix Fibroblast |
| RPS4X | 1.02 | 0.00 | 0.00 | 1.00 | 0.88 | 14 | Matrix Fibroblast |
| TOMM7 | 1.27 | 0.00 | 0.00 | 0.93 | 0.39 | 14 | Matrix Fibroblast |
| FCGRT | 1.05 | 0.00 | 0.00 | 0.61 | 0.13 | 14 | Matrix Fibroblast |
| GSTP1 | 1.18 | 0.00 | 0.00 | 0.80 | 0.24 | 14 | Matrix Fibroblast |
| RPS13 | 1.12 | 0.00 | 0.00 | 1.00 | 0.66 | 14 | Matrix Fibroblast |
| CD63 | 1.31 | 0.00 | 0.00 | 0.90 | 0.34 | 14 | Matrix Fibroblast |
| KRT10 | 0.88 | 0.00 | 0.00 | 0.39 | 0.06 | 14 | Matrix Fibroblast |
| G0S2 | 1.20 | 0.00 | 0.00 | 0.62 | 0.13 | 14 | Matrix Fibroblast |
| TRMT112 | 1.07 | 0.00 | 0.00 | 0.69 | 0.17 | 14 | Matrix Fibroblast |
| MT-ND3 | 1.30 | 0.00 | 0.00 | 0.95 | 0.51 | 14 | Matrix Fibroblast |
| GCSH | 0.91 | 0.00 | 0.00 | 0.39 | 0.06 | 14 | Matrix Fibroblast |
| S100A6 | 1.13 | 0.00 | 0.00 | 0.92 | 0.35 | 14 | Matrix Fibroblast |
| ALDH1A1 | 0.95 | 0.00 | 0.00 | 0.33 | 0.04 | 14 | Matrix Fibroblast |
| MDK | 1.05 | 0.00 | 0.00 | 0.80 | 0.22 | 14 | Matrix Fibroblast |
| PDGFRB | 0.76 | 0.00 | 0.00 | 0.42 | 0.07 | 14 | Matrix Fibroblast |
| TGM2 | 1.40 | 0.00 | 0.00 | 0.51 | 0.11 | 14 | Matrix Fibroblast |
| DUT | 0.96 | 0.00 | 0.00 | 0.52 | 0.10 | 14 | Matrix Fibroblast |
| ADI1 | 0.83 | 0.00 | 0.00 | 0.33 | 0.04 | 14 | Matrix Fibroblast |
| MZT2B | 0.89 | 0.00 | 0.00 | 0.46 | 0.08 | 14 | Matrix Fibroblast |
| SHISA3 | 0.82 | 0.00 | 0.00 | 0.25 | 0.03 | 14 | Matrix Fibroblast |
| ANGPT1 | 0.80 | 0.00 | 0.00 | 0.45 | 0.08 | 14 | Matrix Fibroblast |
| RSL24D1 | 0.98 | 0.00 | 0.00 | 0.55 | 0.12 | 14 | Matrix Fibroblast |
| VAMP2 | 0.94 | 0.00 | 0.00 | 0.48 | 0.09 | 14 | Matrix Fibroblast |
| MYL9 | 0.87 | 0.00 | 0.00 | 0.68 | 0.17 | 14 | Matrix Fibroblast |
| BEX4 | 0.71 | 0.00 | 0.00 | 0.26 | 0.03 | 14 | Matrix Fibroblast |
| RBMX | 1.02 | 0.00 | 0.00 | 0.58 | 0.13 | 14 | Matrix Fibroblast |
| TERF2IP | 0.86 | 0.00 | 0.00 | 0.44 | 0.08 | 14 | Matrix Fibroblast |
| EIF3E | 1.23 | 0.00 | 0.00 | 0.90 | 0.43 | 14 | Matrix Fibroblast |
| GSTM3 | 0.78 | 0.00 | 0.00 | 0.36 | 0.06 | 14 | Matrix Fibroblast |
| MT-ND4 | 0.95 | 0.00 | 0.00 | 0.99 | 0.76 | 14 | Matrix Fibroblast |
| RPLP1 | 0.79 | 0.00 | 0.00 | 1.00 | 0.91 | 14 | Matrix Fibroblast |
| DKK3 | 0.90 | 0.00 | 0.00 | 0.58 | 0.13 | 14 | Matrix Fibroblast |
| MAOB | 0.75 | 0.00 | 0.00 | 0.27 | 0.03 | 14 | Matrix Fibroblast |
| PRMT2 | 0.90 | 0.00 | 0.00 | 0.36 | 0.06 | 14 | Matrix Fibroblast |
| BTG1 | 1.05 | 0.00 | 0.00 | 0.86 | 0.30 | 14 | Matrix Fibroblast |
| RPLP2 | 0.89 | 0.00 | 0.00 | 1.00 | 0.83 | 14 | Matrix Fibroblast |
| RPL8 | 0.83 | 0.00 | 0.00 | 1.00 | 0.81 | 14 | Matrix Fibroblast |
| RAB13 | 1.02 | 0.00 | 0.00 | 0.52 | 0.12 | 14 | Matrix Fibroblast |
| OST4 | 1.04 | 0.00 | 0.00 | 0.80 | 0.28 | 14 | Matrix Fibroblast |
| PTK7 | 0.76 | 0.00 | 0.00 | 0.30 | 0.04 | 14 | Matrix Fibroblast |
| TUFM | 0.89 | 0.00 | 0.00 | 0.41 | 0.08 | 14 | Matrix Fibroblast |
| SUMO2 | 1.02 | 0.00 | 0.00 | 0.86 | 0.34 | 14 | Matrix Fibroblast |
| C12orf57 | 0.95 | 0.00 | 0.00 | 0.62 | 0.16 | 14 | Matrix Fibroblast |
| GRHPR | 0.70 | 0.00 | 0.00 | 0.30 | 0.04 | 14 | Matrix Fibroblast |
| UXT | 0.87 | 0.00 | 0.00 | 0.37 | 0.06 | 14 | Matrix Fibroblast |
| PRMT1 | 0.85 | 0.00 | 0.00 | 0.53 | 0.12 | 14 | Matrix Fibroblast |
| VSTM4 | 0.69 | 0.00 | 0.00 | 0.28 | 0.04 | 14 | Matrix Fibroblast |
| RPL10A | 0.85 | 0.00 | 0.00 | 0.99 | 0.76 | 14 | Matrix Fibroblast |
| RAB34 | 0.84 | 0.00 | 0.00 | 0.47 | 0.10 | 14 | Matrix Fibroblast |
| CHID1 | 0.65 | 0.00 | 0.00 | 0.27 | 0.04 | 14 | Matrix Fibroblast |
| ALKBH7 | 0.75 | 0.00 | 0.00 | 0.34 | 0.06 | 14 | Matrix Fibroblast |
| ENPP2 | 0.62 | 0.00 | 0.00 | 0.34 | 0.06 | 14 | Matrix Fibroblast |
| RPL37A | 0.74 | 0.00 | 0.00 | 1.00 | 0.85 | 14 | Matrix Fibroblast |
| TPM2 | 0.70 | 0.00 | 0.00 | 0.64 | 0.17 | 14 | Matrix Fibroblast |
| SLIT2 | 0.79 | 0.00 | 0.00 | 0.41 | 0.08 | 14 | Matrix Fibroblast |
| HNRNPA0 | 0.92 | 0.00 | 0.00 | 0.72 | 0.24 | 14 | Matrix Fibroblast |
| FBXO7 | 0.67 | 0.00 | 0.00 | 0.26 | 0.04 | 14 | Matrix Fibroblast |
| EID1 | 0.90 | 0.00 | 0.00 | 0.84 | 0.32 | 14 | Matrix Fibroblast |
| RASL12 | 0.76 | 0.00 | 0.00 | 0.34 | 0.06 | 14 | Matrix Fibroblast |
| EIF4A2 | 1.02 | 0.00 | 0.00 | 0.82 | 0.34 | 14 | Matrix Fibroblast |
| PNRC1 | 0.89 | 0.00 | 0.00 | 0.70 | 0.22 | 14 | Matrix Fibroblast |
| POSTN | 0.54 | 0.00 | 0.00 | 0.45 | 0.09 | 14 | Matrix Fibroblast |
| SLIT3 | 0.66 | 0.00 | 0.00 | 0.26 | 0.04 | 14 | Matrix Fibroblast |
| RPS20 | 0.73 | 0.00 | 0.00 | 0.99 | 0.81 | 14 | Matrix Fibroblast |
| FHL1 | 0.75 | 0.00 | 0.00 | 0.84 | 0.30 | 14 | Matrix Fibroblast |
| PIEZO2 | 0.75 | 0.00 | 0.00 | 0.49 | 0.11 | 14 | Matrix Fibroblast |
| MAFB | 0.74 | 0.00 | 0.00 | 0.26 | 0.04 | 14 | Matrix Fibroblast |
| FTH1 | 0.67 | 0.00 | 0.00 | 0.99 | 0.82 | 14 | Matrix Fibroblast |
| SNRPD2 | 0.91 | 0.00 | 0.00 | 0.78 | 0.28 | 14 | Matrix Fibroblast |
| PFDN5 | 0.87 | 0.00 | 0.00 | 0.95 | 0.52 | 14 | Matrix Fibroblast |
| NUCKS1 | 0.95 | 0.00 | 0.00 | 0.77 | 0.29 | 14 | Matrix Fibroblast |
| RPL18A | 0.74 | 0.00 | 0.00 | 1.00 | 0.84 | 14 | Matrix Fibroblast |
| GNAI1 | 0.82 | 0.00 | 0.00 | 0.38 | 0.08 | 14 | Matrix Fibroblast |
| FXYD1 | 0.74 | 0.00 | 0.00 | 0.45 | 0.10 | 14 | Matrix Fibroblast |
| COLEC12 | 0.69 | 0.00 | 0.00 | 0.47 | 0.11 | 14 | Matrix Fibroblast |
| ACTG1 | 0.82 | 0.00 | 0.00 | 0.98 | 0.78 | 14 | Matrix Fibroblast |
| RGS3 | 1.25 | 0.00 | 0.00 | 0.37 | 0.08 | 14 | Matrix Fibroblast |
| SLC38A5 | 0.66 | 0.00 | 0.00 | 0.33 | 0.06 | 14 | Matrix Fibroblast |
| MPST | 0.68 | 0.00 | 0.00 | 0.27 | 0.04 | 14 | Matrix Fibroblast |
| RUNX1T1 | 0.68 | 0.00 | 0.00 | 0.33 | 0.06 | 14 | Matrix Fibroblast |
| VAMP5 | 0.72 | 0.00 | 0.00 | 0.50 | 0.13 | 14 | Matrix Fibroblast |
| MAGED2 | 0.78 | 0.00 | 0.00 | 0.66 | 0.21 | 14 | Matrix Fibroblast |
| FAM43A | 0.70 | 0.00 | 0.00 | 0.26 | 0.04 | 14 | Matrix Fibroblast |
| PPA1 | 0.69 | 0.00 | 0.00 | 0.37 | 0.08 | 14 | Matrix Fibroblast |
| CD164 | 0.77 | 0.00 | 0.00 | 0.55 | 0.16 | 14 | Matrix Fibroblast |
| CITED2 | 0.84 | 0.00 | 0.00 | 0.45 | 0.11 | 14 | Matrix Fibroblast |
| CHST2 | 0.78 | 0.00 | 0.00 | 0.34 | 0.07 | 14 | Matrix Fibroblast |
| NUPR1 | 0.59 | 0.00 | 0.00 | 0.67 | 0.20 | 14 | Matrix Fibroblast |
| WSB1 | 0.85 | 0.00 | 0.00 | 0.69 | 0.24 | 14 | Matrix Fibroblast |
| TMSB4X | 0.72 | 0.00 | 0.00 | 1.00 | 0.90 | 14 | Matrix Fibroblast |
| PARK7 | 0.83 | 0.00 | 0.00 | 0.75 | 0.28 | 14 | Matrix Fibroblast |
| CDH11 | 0.45 | 0.00 | 0.00 | 0.67 | 0.19 | 14 | Matrix Fibroblast |
| TMA7 | 0.85 | 0.00 | 0.00 | 0.82 | 0.35 | 14 | Matrix Fibroblast |
| SSR2 | 0.84 | 0.00 | 0.00 | 0.80 | 0.34 | 14 | Matrix Fibroblast |
| MTCH1 | 0.76 | 0.00 | 0.00 | 0.57 | 0.17 | 14 | Matrix Fibroblast |
| EI24 | 0.69 | 0.00 | 0.00 | 0.35 | 0.07 | 14 | Matrix Fibroblast |
| CEBPB | 0.74 | 0.00 | 0.00 | 0.42 | 0.10 | 14 | Matrix Fibroblast |
| RPA2 | 0.65 | 0.00 | 0.00 | 0.30 | 0.06 | 14 | Matrix Fibroblast |
| UQCRB | 0.87 | 0.00 | 0.00 | 0.90 | 0.47 | 14 | Matrix Fibroblast |
| PSMB7 | 0.76 | 0.00 | 0.00 | 0.48 | 0.13 | 14 | Matrix Fibroblast |
| TMEM43 | 0.65 | 0.00 | 0.00 | 0.31 | 0.06 | 14 | Matrix Fibroblast |
| ARL6IP5 | 0.73 | 0.00 | 0.00 | 0.50 | 0.14 | 14 | Matrix Fibroblast |
| IMPDH2 | 0.78 | 0.00 | 0.00 | 0.44 | 0.11 | 14 | Matrix Fibroblast |
| NACA | 0.75 | 0.00 | 0.00 | 0.96 | 0.61 | 14 | Matrix Fibroblast |
| NDUFAF3 | 0.68 | 0.00 | 0.00 | 0.35 | 0.07 | 14 | Matrix Fibroblast |
| PSMF1 | 0.62 | 0.00 | 0.00 | 0.28 | 0.05 | 14 | Matrix Fibroblast |
| EDF1 | 0.78 | 0.00 | 0.00 | 0.81 | 0.34 | 14 | Matrix Fibroblast |
| FILIP1 | 0.69 | 0.00 | 0.00 | 0.33 | 0.07 | 14 | Matrix Fibroblast |
| SPINT2 | 0.65 | 0.00 | 0.00 | 0.44 | 0.11 | 14 | Matrix Fibroblast |
| POLR2L | 0.78 | 0.00 | 0.00 | 0.61 | 0.19 | 14 | Matrix Fibroblast |
| HEBP1 | 0.53 | 0.00 | 0.00 | 0.27 | 0.05 | 14 | Matrix Fibroblast |
| LAMA4 | 0.72 | 0.00 | 0.00 | 0.49 | 0.13 | 14 | Matrix Fibroblast |
| ZYX | 0.85 | 0.00 | 0.00 | 0.61 | 0.20 | 14 | Matrix Fibroblast |
| EEF2 | 0.81 | 0.00 | 0.00 | 0.94 | 0.59 | 14 | Matrix Fibroblast |
| APOE | 0.72 | 0.00 | 0.00 | 0.70 | 0.27 | 14 | Matrix Fibroblast |
| LSM3 | 0.66 | 0.00 | 0.00 | 0.46 | 0.12 | 14 | Matrix Fibroblast |
| PHPT1 | 0.74 | 0.00 | 0.00 | 0.47 | 0.13 | 14 | Matrix Fibroblast |
| TRIP10 | 0.65 | 0.00 | 0.00 | 0.31 | 0.06 | 14 | Matrix Fibroblast |
| LAPTM4A | 0.79 | 0.00 | 0.00 | 0.91 | 0.55 | 14 | Matrix Fibroblast |
| MRC2 | 0.66 | 0.00 | 0.00 | 0.28 | 0.05 | 14 | Matrix Fibroblast |
| TMBIM4 | 0.72 | 0.00 | 0.00 | 0.52 | 0.15 | 14 | Matrix Fibroblast |
| SERPINB6 | 0.57 | 0.00 | 0.00 | 0.29 | 0.06 | 14 | Matrix Fibroblast |
| QSOX1 | 0.72 | 0.00 | 0.00 | 0.40 | 0.10 | 14 | Matrix Fibroblast |
| TMEM205 | 0.66 | 0.00 | 0.00 | 0.34 | 0.08 | 14 | Matrix Fibroblast |
| SRSF9 | 0.63 | 0.00 | 0.00 | 0.46 | 0.12 | 14 | Matrix Fibroblast |
| ERGIC3 | 0.73 | 0.00 | 0.00 | 0.54 | 0.17 | 14 | Matrix Fibroblast |
| H3F3A | 0.69 | 0.00 | 0.00 | 0.94 | 0.62 | 14 | Matrix Fibroblast |
| OGN | 0.39 | 0.00 | 0.00 | 0.50 | 0.13 | 14 | Matrix Fibroblast |
| VPS28 | 0.70 | 0.00 | 0.00 | 0.61 | 0.21 | 14 | Matrix Fibroblast |
| PLD3 | 0.65 | 0.00 | 0.00 | 0.35 | 0.08 | 14 | Matrix Fibroblast |
| GPX3 | 0.68 | 0.00 | 0.00 | 0.49 | 0.14 | 14 | Matrix Fibroblast |
| GSTK1 | 0.68 | 0.00 | 0.00 | 0.38 | 0.09 | 14 | Matrix Fibroblast |
| GYG1 | 0.56 | 0.00 | 0.00 | 0.61 | 0.19 | 14 | Matrix Fibroblast |
| PMF1 | 0.61 | 0.00 | 0.00 | 0.26 | 0.05 | 14 | Matrix Fibroblast |
| MT-CO3 | 0.53 | 0.00 | 0.00 | 1.00 | 0.89 | 14 | Matrix Fibroblast |
| AAMDC | 0.61 | 0.00 | 0.00 | 0.33 | 0.07 | 14 | Matrix Fibroblast |
| POLR2G | 0.72 | 0.00 | 0.00 | 0.49 | 0.14 | 14 | Matrix Fibroblast |
| AP2M1 | 0.72 | 0.00 | 0.00 | 0.54 | 0.17 | 14 | Matrix Fibroblast |
| PLSCR4 | 0.57 | 0.00 | 0.00 | 0.30 | 0.06 | 14 | Matrix Fibroblast |
| CALM3 | 0.66 | 0.00 | 0.00 | 0.61 | 0.20 | 14 | Matrix Fibroblast |
| PMP22 | 0.84 | 0.00 | 0.00 | 0.76 | 0.33 | 14 | Matrix Fibroblast |
| EXOC7 | 0.62 | 0.00 | 0.00 | 0.29 | 0.06 | 14 | Matrix Fibroblast |
| HDDC2 | 0.59 | 0.00 | 0.00 | 0.34 | 0.08 | 14 | Matrix Fibroblast |
| CCNG1 | 0.60 | 0.00 | 0.00 | 0.33 | 0.07 | 14 | Matrix Fibroblast |
| LMNA | 0.94 | 0.00 | 0.00 | 0.61 | 0.23 | 14 | Matrix Fibroblast |
| BTF3 | 0.69 | 0.00 | 0.00 | 0.93 | 0.54 | 14 | Matrix Fibroblast |
| EDNRA | 0.63 | 0.00 | 0.00 | 0.39 | 0.10 | 14 | Matrix Fibroblast |
| TSPAN4 | 0.65 | 0.00 | 0.00 | 0.38 | 0.09 | 14 | Matrix Fibroblast |
| ITGA8 | 0.42 | 0.00 | 0.00 | 0.40 | 0.10 | 14 | Matrix Fibroblast |
| COL13A1 | 0.51 | 0.00 | 0.00 | 0.39 | 0.10 | 14 | Matrix Fibroblast |
| SERF2 | 0.67 | 0.00 | 0.00 | 0.96 | 0.60 | 14 | Matrix Fibroblast |
| PGAM1 | 0.72 | 0.00 | 0.00 | 0.46 | 0.13 | 14 | Matrix Fibroblast |
| TMEM9 | 0.57 | 0.00 | 0.00 | 0.32 | 0.07 | 14 | Matrix Fibroblast |
| RND3 | 0.78 | 0.00 | 0.00 | 0.35 | 0.08 | 14 | Matrix Fibroblast |
| ENSA | 0.58 | 0.00 | 0.00 | 0.48 | 0.14 | 14 | Matrix Fibroblast |
| SLC44A1 | 0.64 | 0.00 | 0.00 | 0.36 | 0.09 | 14 | Matrix Fibroblast |
| APEX1 | 0.67 | 0.00 | 0.00 | 0.40 | 0.11 | 14 | Matrix Fibroblast |
| PLAGL1 | 0.57 | 0.00 | 0.00 | 0.29 | 0.06 | 14 | Matrix Fibroblast |
| FOXO3 | 0.63 | 0.00 | 0.00 | 0.32 | 0.08 | 14 | Matrix Fibroblast |
| RPL6 | 0.51 | 0.00 | 0.00 | 1.00 | 0.88 | 14 | Matrix Fibroblast |
| B2M | 0.63 | 0.00 | 0.00 | 1.00 | 0.70 | 14 | Matrix Fibroblast |
| CDO1 | 0.48 | 0.00 | 0.00 | 0.29 | 0.06 | 14 | Matrix Fibroblast |
| EIF4A3 | 0.59 | 0.00 | 0.00 | 0.29 | 0.07 | 14 | Matrix Fibroblast |
| GABPB1 | 0.58 | 0.00 | 0.00 | 0.25 | 0.05 | 14 | Matrix Fibroblast |
| MARCKSL1 | 0.71 | 0.00 | 0.00 | 0.62 | 0.23 | 14 | Matrix Fibroblast |
| SOX4 | 0.78 | 0.00 | 0.00 | 0.84 | 0.40 | 14 | Matrix Fibroblast |
| GUK1 | 0.63 | 0.00 | 0.00 | 0.56 | 0.19 | 14 | Matrix Fibroblast |
| FNIP1 | 0.71 | 0.00 | 0.00 | 0.32 | 0.08 | 14 | Matrix Fibroblast |
| NDRG2 | 0.59 | 0.00 | 0.00 | 0.35 | 0.09 | 14 | Matrix Fibroblast |
| FKBP8 | 0.66 | 0.00 | 0.00 | 0.48 | 0.15 | 14 | Matrix Fibroblast |
| MYL6 | 0.59 | 0.00 | 0.00 | 0.96 | 0.72 | 14 | Matrix Fibroblast |
| HSD17B11 | 0.53 | 0.00 | 0.00 | 0.35 | 0.09 | 14 | Matrix Fibroblast |
| NDUFS5 | 0.66 | 0.00 | 0.00 | 0.67 | 0.26 | 14 | Matrix Fibroblast |
| ALDOA | 0.66 | 0.00 | 0.00 | 0.61 | 0.22 | 14 | Matrix Fibroblast |
| ARID1B | 0.59 | 0.00 | 0.00 | 0.35 | 0.09 | 14 | Matrix Fibroblast |
| GSTO1 | 0.49 | 0.00 | 0.00 | 0.32 | 0.08 | 14 | Matrix Fibroblast |
| PKIG | 0.60 | 0.00 | 0.00 | 0.38 | 0.10 | 14 | Matrix Fibroblast |
| RGCC | 0.55 | 0.00 | 0.00 | 0.56 | 0.19 | 14 | Matrix Fibroblast |
| KANK2 | 0.65 | 0.00 | 0.00 | 0.44 | 0.13 | 14 | Matrix Fibroblast |
| PIH1D1 | 0.56 | 0.00 | 0.00 | 0.29 | 0.07 | 14 | Matrix Fibroblast |
| CUTA | 0.67 | 0.00 | 0.00 | 0.56 | 0.20 | 14 | Matrix Fibroblast |
| GABARAPL2 | 0.61 | 0.00 | 0.00 | 0.65 | 0.25 | 14 | Matrix Fibroblast |
| CARHSP1 | 0.60 | 0.00 | 0.00 | 0.48 | 0.15 | 14 | Matrix Fibroblast |
| UBE2E3 | 0.63 | 0.00 | 0.00 | 0.47 | 0.15 | 14 | Matrix Fibroblast |
| EBPL | 0.53 | 0.00 | 0.00 | 0.25 | 0.05 | 14 | Matrix Fibroblast |
| BMP5 | 0.62 | 0.00 | 0.00 | 0.26 | 0.06 | 14 | Matrix Fibroblast |
| PLXDC2 | 0.38 | 0.00 | 0.00 | 0.45 | 0.13 | 14 | Matrix Fibroblast |
| COL5A1 | 0.58 | 0.00 | 0.00 | 0.38 | 0.10 | 14 | Matrix Fibroblast |
| UQCR11 | 0.64 | 0.00 | 0.00 | 0.74 | 0.32 | 14 | Matrix Fibroblast |
| CAMLG | 0.64 | 0.00 | 0.00 | 0.35 | 0.10 | 14 | Matrix Fibroblast |
| PET100 | 0.63 | 0.00 | 0.00 | 0.47 | 0.15 | 14 | Matrix Fibroblast |
| FBLN5 | 0.35 | 0.00 | 0.00 | 0.55 | 0.18 | 14 | Matrix Fibroblast |
| FBL | 0.50 | 0.00 | 0.00 | 0.35 | 0.09 | 14 | Matrix Fibroblast |
| 6-Mar | 0.58 | 0.00 | 0.00 | 0.31 | 0.08 | 14 | Matrix Fibroblast |
| IL6ST | 0.62 | 0.00 | 0.00 | 0.36 | 0.10 | 14 | Matrix Fibroblast |
| TAF7 | 0.55 | 0.00 | 0.00 | 0.43 | 0.13 | 14 | Matrix Fibroblast |
| ACADVL | 0.60 | 0.00 | 0.00 | 0.37 | 0.10 | 14 | Matrix Fibroblast |
| KDELR1 | 0.61 | 0.00 | 0.00 | 0.48 | 0.16 | 14 | Matrix Fibroblast |
| ENO1 | 0.73 | 0.00 | 0.00 | 0.53 | 0.19 | 14 | Matrix Fibroblast |
| CLU | 0.60 | 0.00 | 0.00 | 0.29 | 0.07 | 14 | Matrix Fibroblast |
| SMPDL3A | 0.66 | 0.00 | 0.00 | 0.38 | 0.11 | 14 | Matrix Fibroblast |
| N4BP2L2 | 0.61 | 0.00 | 0.00 | 0.60 | 0.23 | 14 | Matrix Fibroblast |
| DGUOK | 0.53 | 0.00 | 0.00 | 0.37 | 0.10 | 14 | Matrix Fibroblast |
| SRSF7 | 0.67 | 0.00 | 0.00 | 0.50 | 0.17 | 14 | Matrix Fibroblast |
| NENF | 0.55 | 0.00 | 0.00 | 0.56 | 0.20 | 14 | Matrix Fibroblast |
| BAD | 0.56 | 0.00 | 0.00 | 0.34 | 0.09 | 14 | Matrix Fibroblast |
| SMS | 0.58 | 0.00 | 0.00 | 0.38 | 0.11 | 14 | Matrix Fibroblast |
| NUCB1 | 0.53 | 0.00 | 0.00 | 0.48 | 0.16 | 14 | Matrix Fibroblast |
| ATRAID | 0.56 | 0.00 | 0.00 | 0.52 | 0.18 | 14 | Matrix Fibroblast |
| FZD1 | 0.48 | 0.00 | 0.00 | 0.26 | 0.06 | 14 | Matrix Fibroblast |
| GLO1 | 0.60 | 0.00 | 0.00 | 0.46 | 0.15 | 14 | Matrix Fibroblast |
| PRDX6 | 0.50 | 0.00 | 0.00 | 0.69 | 0.29 | 14 | Matrix Fibroblast |
| DECR1 | 0.50 | 0.00 | 0.00 | 0.32 | 0.08 | 14 | Matrix Fibroblast |
| ST13 | 0.63 | 0.00 | 0.00 | 0.75 | 0.35 | 14 | Matrix Fibroblast |
| LRP1 | 0.48 | 0.00 | 0.00 | 0.41 | 0.12 | 14 | Matrix Fibroblast |
| LOXL1 | 0.52 | 0.00 | 0.00 | 0.39 | 0.12 | 14 | Matrix Fibroblast |
| ATP6V0E1 | 0.57 | 0.00 | 0.00 | 0.59 | 0.23 | 14 | Matrix Fibroblast |
| RPL36A | 0.62 | 0.00 | 0.00 | 0.96 | 0.64 | 14 | Matrix Fibroblast |
| FSCN1 | 0.48 | 0.00 | 0.00 | 0.31 | 0.08 | 14 | Matrix Fibroblast |
| ITGB1BP1 | 0.50 | 0.00 | 0.00 | 0.29 | 0.07 | 14 | Matrix Fibroblast |
| EIF1AX | 0.55 | 0.00 | 0.00 | 0.58 | 0.22 | 14 | Matrix Fibroblast |
| MT-CO2 | 0.53 | 0.00 | 0.00 | 0.98 | 0.79 | 14 | Matrix Fibroblast |
| SRSF3 | 0.61 | 0.00 | 0.00 | 0.66 | 0.28 | 14 | Matrix Fibroblast |
| CAV1 | 0.57 | 0.00 | 0.00 | 0.78 | 0.37 | 14 | Matrix Fibroblast |
| RBPJ | 0.53 | 0.00 | 0.00 | 0.38 | 0.12 | 14 | Matrix Fibroblast |
| FHOD1 | 0.46 | 0.00 | 0.00 | 0.28 | 0.07 | 14 | Matrix Fibroblast |
| MAST4 | 0.54 | 0.00 | 0.00 | 0.27 | 0.06 | 14 | Matrix Fibroblast |
| ETV1 | 0.48 | 0.00 | 0.00 | 0.29 | 0.07 | 14 | Matrix Fibroblast |
| BRK1 | 0.57 | 0.00 | 0.00 | 0.68 | 0.29 | 14 | Matrix Fibroblast |
| CMTM6 | 0.53 | 0.00 | 0.00 | 0.35 | 0.10 | 14 | Matrix Fibroblast |
| ANXA6 | 0.55 | 0.00 | 0.00 | 0.57 | 0.22 | 14 | Matrix Fibroblast |
| NDUFB2 | 0.59 | 0.00 | 0.00 | 0.59 | 0.24 | 14 | Matrix Fibroblast |
| HADHB | 0.53 | 0.00 | 0.00 | 0.35 | 0.10 | 14 | Matrix Fibroblast |
| TERF1 | 0.47 | 0.00 | 0.00 | 0.26 | 0.06 | 14 | Matrix Fibroblast |
| SSR4 | 0.61 | 0.00 | 0.00 | 0.63 | 0.27 | 14 | Matrix Fibroblast |
| COL14A1 | 0.40 | 0.00 | 0.00 | 0.26 | 0.06 | 14 | Matrix Fibroblast |
| TAGLN2 | 0.62 | 0.00 | 0.00 | 0.83 | 0.47 | 14 | Matrix Fibroblast |
| ESD | 0.60 | 0.00 | 0.00 | 0.56 | 0.22 | 14 | Matrix Fibroblast |
| DDT | 0.56 | 0.00 | 0.00 | 0.32 | 0.09 | 14 | Matrix Fibroblast |
| TMEM258 | 0.57 | 0.00 | 0.00 | 0.75 | 0.35 | 14 | Matrix Fibroblast |
| ECH1 | 0.54 | 0.00 | 0.00 | 0.48 | 0.17 | 14 | Matrix Fibroblast |
| APH1A | 0.53 | 0.00 | 0.00 | 0.41 | 0.13 | 14 | Matrix Fibroblast |
| RALY | 0.52 | 0.00 | 0.00 | 0.41 | 0.14 | 14 | Matrix Fibroblast |
| S100A4 | 0.38 | 0.00 | 0.00 | 0.35 | 0.10 | 14 | Matrix Fibroblast |
| GULP1 | 0.54 | 0.00 | 0.00 | 0.30 | 0.08 | 14 | Matrix Fibroblast |
| LAMTOR5 | 0.54 | 0.00 | 0.00 | 0.52 | 0.20 | 14 | Matrix Fibroblast |
| MXRA8 | 0.44 | 0.00 | 0.00 | 0.49 | 0.17 | 14 | Matrix Fibroblast |
| EPC1 | 0.48 | 0.00 | 0.00 | 0.34 | 0.10 | 14 | Matrix Fibroblast |
| ADH5 | 0.52 | 0.00 | 0.00 | 0.46 | 0.16 | 14 | Matrix Fibroblast |
| PSMG2 | 0.47 | 0.00 | 0.00 | 0.27 | 0.07 | 14 | Matrix Fibroblast |
| DSTN | 0.57 | 0.00 | 0.00 | 0.86 | 0.53 | 14 | Matrix Fibroblast |
| AES | 0.55 | 0.00 | 0.00 | 0.44 | 0.16 | 14 | Matrix Fibroblast |
| SIVA1 | 0.51 | 0.00 | 0.00 | 0.45 | 0.16 | 14 | Matrix Fibroblast |
| PGRMC1 | 0.48 | 0.00 | 0.00 | 0.51 | 0.19 | 14 | Matrix Fibroblast |
| LGALS1 | 0.38 | 0.00 | 0.00 | 0.91 | 0.46 | 14 | Matrix Fibroblast |
| SMIM7 | 0.50 | 0.00 | 0.00 | 0.37 | 0.12 | 14 | Matrix Fibroblast |
| UBC | 0.60 | 0.00 | 0.00 | 0.87 | 0.50 | 14 | Matrix Fibroblast |
| DNPEP | 0.51 | 0.00 | 0.00 | 0.30 | 0.08 | 14 | Matrix Fibroblast |
| PRNP | 0.42 | 0.00 | 0.00 | 0.41 | 0.14 | 14 | Matrix Fibroblast |
| EIF4B | 0.53 | 0.00 | 0.00 | 0.51 | 0.19 | 14 | Matrix Fibroblast |
| BMP4 | 0.52 | 0.00 | 0.00 | 0.34 | 0.10 | 14 | Matrix Fibroblast |
| EIF1 | 0.52 | 0.00 | 0.00 | 0.96 | 0.69 | 14 | Matrix Fibroblast |
| METTL9 | 0.47 | 0.00 | 0.00 | 0.40 | 0.13 | 14 | Matrix Fibroblast |
| PRSS23 | 0.58 | 0.00 | 0.00 | 0.29 | 0.08 | 14 | Matrix Fibroblast |
| PPIB | 0.57 | 0.00 | 0.00 | 0.81 | 0.46 | 14 | Matrix Fibroblast |
| RNH1 | 0.51 | 0.00 | 0.00 | 0.47 | 0.17 | 14 | Matrix Fibroblast |
| SCARB2 | 0.47 | 0.00 | 0.00 | 0.29 | 0.08 | 14 | Matrix Fibroblast |
| VDAC2 | 0.50 | 0.00 | 0.00 | 0.59 | 0.25 | 14 | Matrix Fibroblast |
| TSPYL1 | 0.47 | 0.00 | 0.00 | 0.34 | 0.10 | 14 | Matrix Fibroblast |
| S100A10 | 0.59 | 0.00 | 0.00 | 0.78 | 0.41 | 14 | Matrix Fibroblast |
| LRPAP1 | 0.45 | 0.00 | 0.00 | 0.47 | 0.17 | 14 | Matrix Fibroblast |
| CHD9 | 0.43 | 0.00 | 0.00 | 0.38 | 0.13 | 14 | Matrix Fibroblast |
| RAB5C | 0.48 | 0.00 | 0.00 | 0.41 | 0.14 | 14 | Matrix Fibroblast |
| NHP2 | 0.51 | 0.00 | 0.00 | 0.38 | 0.13 | 14 | Matrix Fibroblast |
| STOM | 0.49 | 0.00 | 0.00 | 0.31 | 0.09 | 14 | Matrix Fibroblast |
| SEC11A | 0.48 | 0.00 | 0.00 | 0.52 | 0.20 | 14 | Matrix Fibroblast |
| RAD23A | 0.47 | 0.00 | 0.00 | 0.41 | 0.14 | 14 | Matrix Fibroblast |
| SNX17 | 0.45 | 0.00 | 0.00 | 0.34 | 0.11 | 14 | Matrix Fibroblast |
| DPM1 | 0.46 | 0.00 | 0.00 | 0.27 | 0.08 | 14 | Matrix Fibroblast |
| MT-ND2 | 0.49 | 0.00 | 0.00 | 0.96 | 0.71 | 14 | Matrix Fibroblast |
| LSM2 | 0.43 | 0.00 | 0.00 | 0.32 | 0.10 | 14 | Matrix Fibroblast |
| EIF3D | 0.44 | 0.00 | 0.00 | 0.51 | 0.20 | 14 | Matrix Fibroblast |
| RGS2 | 0.38 | 0.00 | 0.00 | 0.63 | 0.27 | 14 | Matrix Fibroblast |
| PSIP1 | 0.48 | 0.00 | 0.00 | 0.43 | 0.15 | 14 | Matrix Fibroblast |
| MT-CO1 | 0.35 | 0.00 | 0.00 | 1.00 | 0.95 | 14 | Matrix Fibroblast |
| LAMTOR4 | 0.53 | 0.00 | 0.00 | 0.40 | 0.14 | 14 | Matrix Fibroblast |
| SLC1A5 | 0.45 | 0.00 | 0.00 | 0.33 | 0.10 | 14 | Matrix Fibroblast |
| CYGB | 0.38 | 0.00 | 0.00 | 0.26 | 0.07 | 14 | Matrix Fibroblast |
| COPZ1 | 0.48 | 0.00 | 0.00 | 0.38 | 0.13 | 14 | Matrix Fibroblast |
| TMEM59 | 0.51 | 0.00 | 0.00 | 0.78 | 0.42 | 14 | Matrix Fibroblast |
| RHEB | 0.48 | 0.00 | 0.00 | 0.49 | 0.19 | 14 | Matrix Fibroblast |
| TNFRSF1A | 0.46 | 0.00 | 0.00 | 0.39 | 0.14 | 14 | Matrix Fibroblast |
| FKBP2 | 0.48 | 0.00 | 0.00 | 0.49 | 0.20 | 14 | Matrix Fibroblast |
| PDLIM2 | 0.50 | 0.00 | 0.00 | 0.47 | 0.18 | 14 | Matrix Fibroblast |
| MYH10 | 0.37 | 0.00 | 0.00 | 0.56 | 0.22 | 14 | Matrix Fibroblast |
| MRPS7 | 0.46 | 0.00 | 0.00 | 0.27 | 0.08 | 14 | Matrix Fibroblast |
| LMAN2 | 0.44 | 0.00 | 0.00 | 0.42 | 0.15 | 14 | Matrix Fibroblast |
| CNOT7 | 0.47 | 0.00 | 0.00 | 0.29 | 0.09 | 14 | Matrix Fibroblast |
| SNAI2 | 0.37 | 0.00 | 0.00 | 0.36 | 0.12 | 14 | Matrix Fibroblast |
| NDUFA4 | 0.51 | 0.00 | 0.00 | 0.88 | 0.57 | 14 | Matrix Fibroblast |
| STOML2 | 0.46 | 0.00 | 0.00 | 0.31 | 0.10 | 14 | Matrix Fibroblast |
| ARL6IP4 | 0.42 | 0.00 | 0.00 | 0.51 | 0.20 | 14 | Matrix Fibroblast |
| MAGEH1 | 0.42 | 0.00 | 0.00 | 0.26 | 0.08 | 14 | Matrix Fibroblast |
| ATP6AP2 | 0.45 | 0.00 | 0.00 | 0.44 | 0.17 | 14 | Matrix Fibroblast |
| PTMA | 0.34 | 0.00 | 0.00 | 1.00 | 0.92 | 14 | Matrix Fibroblast |
| RBFOX2 | 0.45 | 0.00 | 0.00 | 0.29 | 0.09 | 14 | Matrix Fibroblast |
| SF3B5 | 0.42 | 0.00 | 0.00 | 0.55 | 0.23 | 14 | Matrix Fibroblast |
| TMED4 | 0.48 | 0.00 | 0.00 | 0.30 | 0.09 | 14 | Matrix Fibroblast |
| LSM5 | 0.42 | 0.00 | 0.00 | 0.36 | 0.13 | 14 | Matrix Fibroblast |
| CYB5B | 0.44 | 0.00 | 0.00 | 0.29 | 0.09 | 14 | Matrix Fibroblast |
| TSPAN3 | 0.42 | 0.00 | 0.00 | 0.44 | 0.17 | 14 | Matrix Fibroblast |
| ISCU | 0.47 | 0.00 | 0.00 | 0.44 | 0.17 | 14 | Matrix Fibroblast |
| EIF3I | 0.47 | 0.00 | 0.00 | 0.55 | 0.24 | 14 | Matrix Fibroblast |
| KDELR2 | 0.49 | 0.00 | 0.00 | 0.50 | 0.21 | 14 | Matrix Fibroblast |
| AP2S1 | 0.43 | 0.00 | 0.00 | 0.48 | 0.19 | 14 | Matrix Fibroblast |
| MEAF6 | 0.46 | 0.00 | 0.00 | 0.27 | 0.08 | 14 | Matrix Fibroblast |
| TM2D3 | 0.41 | 0.00 | 0.00 | 0.25 | 0.08 | 14 | Matrix Fibroblast |
| DRAP1 | 0.41 | 0.00 | 0.00 | 0.56 | 0.25 | 14 | Matrix Fibroblast |
| TMEM147 | 0.46 | 0.00 | 0.00 | 0.36 | 0.13 | 14 | Matrix Fibroblast |
| FGFR1 | 0.38 | 0.00 | 0.00 | 0.34 | 0.11 | 14 | Matrix Fibroblast |
| MRPL51 | 0.43 | 0.00 | 0.00 | 0.42 | 0.16 | 14 | Matrix Fibroblast |
| TRAPPC1 | 0.44 | 0.00 | 0.00 | 0.36 | 0.13 | 14 | Matrix Fibroblast |
| FLOT1 | 0.44 | 0.00 | 0.00 | 0.28 | 0.09 | 14 | Matrix Fibroblast |
| SRP9 | 0.42 | 0.00 | 0.00 | 0.63 | 0.30 | 14 | Matrix Fibroblast |
| LAMTOR1 | 0.43 | 0.00 | 0.00 | 0.39 | 0.15 | 14 | Matrix Fibroblast |
| SPAG7 | 0.46 | 0.00 | 0.00 | 0.36 | 0.13 | 14 | Matrix Fibroblast |
| PPP1R15A | 0.47 | 0.00 | 0.00 | 0.26 | 0.08 | 14 | Matrix Fibroblast |
| JTB | 0.44 | 0.00 | 0.00 | 0.48 | 0.20 | 14 | Matrix Fibroblast |
| ZNF22 | 0.39 | 0.00 | 0.00 | 0.35 | 0.12 | 14 | Matrix Fibroblast |
| NELFE | 0.43 | 0.00 | 0.00 | 0.26 | 0.08 | 14 | Matrix Fibroblast |
| RNF146 | 0.44 | 0.00 | 0.00 | 0.33 | 0.12 | 14 | Matrix Fibroblast |
| NAP1L1 | 0.36 | 0.00 | 0.00 | 0.83 | 0.47 | 14 | Matrix Fibroblast |
| SAP18 | 0.48 | 0.00 | 0.00 | 0.64 | 0.32 | 14 | Matrix Fibroblast |
| COL5A2 | 0.35 | 0.00 | 0.00 | 0.47 | 0.19 | 14 | Matrix Fibroblast |
| CRTAP | 0.41 | 0.00 | 0.00 | 0.27 | 0.09 | 14 | Matrix Fibroblast |
| PDLIM3 | 0.44 | 0.00 | 0.00 | 0.27 | 0.08 | 14 | Matrix Fibroblast |
| MRFAP1 | 0.46 | 0.00 | 0.00 | 0.68 | 0.35 | 14 | Matrix Fibroblast |
| JUNB | 0.81 | 0.00 | 0.00 | 0.47 | 0.21 | 14 | Matrix Fibroblast |
| CD302 | 0.28 | 0.00 | 0.00 | 0.40 | 0.14 | 14 | Matrix Fibroblast |
| KLHDC3 | 0.42 | 0.00 | 0.00 | 0.29 | 0.10 | 14 | Matrix Fibroblast |
| EIF3G | 0.44 | 0.00 | 0.00 | 0.48 | 0.21 | 14 | Matrix Fibroblast |
| HSD17B12 | 0.42 | 0.00 | 0.00 | 0.33 | 0.12 | 14 | Matrix Fibroblast |
| EMC4 | 0.41 | 0.00 | 0.00 | 0.39 | 0.15 | 14 | Matrix Fibroblast |
| NOSIP | 0.44 | 0.00 | 0.00 | 0.29 | 0.10 | 14 | Matrix Fibroblast |
| TMED2 | 0.39 | 0.00 | 0.00 | 0.57 | 0.26 | 14 | Matrix Fibroblast |
| POLR2K | 0.38 | 0.00 | 0.00 | 0.38 | 0.14 | 14 | Matrix Fibroblast |
| CLNS1A | 0.41 | 0.00 | 0.00 | 0.29 | 0.10 | 14 | Matrix Fibroblast |
| GADD45B | 0.62 | 0.00 | 0.00 | 0.29 | 0.10 | 14 | Matrix Fibroblast |
| ITM2C | 0.41 | 0.00 | 0.00 | 0.54 | 0.24 | 14 | Matrix Fibroblast |
| CSNK2B | 0.41 | 0.00 | 0.00 | 0.43 | 0.18 | 14 | Matrix Fibroblast |
| EMC10 | 0.38 | 0.00 | 0.00 | 0.31 | 0.11 | 14 | Matrix Fibroblast |
| EGFL6 | 0.28 | 0.00 | 0.00 | 0.32 | 0.11 | 14 | Matrix Fibroblast |
| HERPUD1 | 0.38 | 0.00 | 0.00 | 0.39 | 0.15 | 14 | Matrix Fibroblast |
| EMP3 | 0.34 | 0.00 | 0.00 | 0.54 | 0.24 | 14 | Matrix Fibroblast |
| PNKD | 0.40 | 0.00 | 0.00 | 0.30 | 0.11 | 14 | Matrix Fibroblast |
| NEDD8 | 0.38 | 0.00 | 0.00 | 0.64 | 0.32 | 14 | Matrix Fibroblast |
| TECR | 0.41 | 0.00 | 0.00 | 0.39 | 0.16 | 14 | Matrix Fibroblast |
| TSC22D3 | 0.33 | 0.00 | 0.00 | 0.51 | 0.23 | 14 | Matrix Fibroblast |
| MEIS1 | 0.34 | 0.00 | 0.00 | 0.27 | 0.09 | 14 | Matrix Fibroblast |
| BLOC1S1 | 0.40 | 0.00 | 0.00 | 0.26 | 0.08 | 14 | Matrix Fibroblast |
| NDUFS4 | 0.44 | 0.00 | 0.00 | 0.47 | 0.21 | 14 | Matrix Fibroblast |
| COPS6 | 0.38 | 0.00 | 0.00 | 0.45 | 0.19 | 14 | Matrix Fibroblast |
| MRPS34 | 0.39 | 0.00 | 0.00 | 0.28 | 0.10 | 14 | Matrix Fibroblast |
| CNBP | 0.37 | 0.00 | 0.00 | 0.81 | 0.47 | 14 | Matrix Fibroblast |
| HNRNPDL | 0.38 | 0.00 | 0.00 | 0.72 | 0.38 | 14 | Matrix Fibroblast |
| ANXA1 | 0.34 | 0.00 | 0.00 | 0.47 | 0.20 | 14 | Matrix Fibroblast |
| HSBP1 | 0.41 | 0.00 | 0.00 | 0.63 | 0.32 | 14 | Matrix Fibroblast |
| C11orf58 | 0.36 | 0.00 | 0.00 | 0.63 | 0.30 | 14 | Matrix Fibroblast |
| PGK1 | 0.42 | 0.00 | 0.00 | 0.35 | 0.13 | 14 | Matrix Fibroblast |
| TMSB10 | 0.29 | 0.00 | 0.00 | 1.00 | 0.88 | 14 | Matrix Fibroblast |
| MRPS6 | 0.39 | 0.00 | 0.00 | 0.32 | 0.12 | 14 | Matrix Fibroblast |
| NFIA | 0.35 | 0.00 | 0.00 | 0.53 | 0.24 | 14 | Matrix Fibroblast |
| MRPS21 | 0.39 | 0.00 | 0.00 | 0.48 | 0.21 | 14 | Matrix Fibroblast |
| ANAPC16 | 0.41 | 0.00 | 0.00 | 0.44 | 0.19 | 14 | Matrix Fibroblast |
| SRSF4 | 0.35 | 0.00 | 0.00 | 0.36 | 0.14 | 14 | Matrix Fibroblast |
| LAMC1 | 0.41 | 0.00 | 0.00 | 0.39 | 0.16 | 14 | Matrix Fibroblast |
| TACC1 | 0.29 | 0.00 | 0.00 | 0.53 | 0.23 | 14 | Matrix Fibroblast |
| STAT3 | 0.40 | 0.00 | 0.00 | 0.40 | 0.16 | 14 | Matrix Fibroblast |
| CUEDC2 | 0.39 | 0.00 | 0.00 | 0.30 | 0.11 | 14 | Matrix Fibroblast |
| PPP2R1A | 0.42 | 0.00 | 0.00 | 0.41 | 0.17 | 14 | Matrix Fibroblast |
| MMADHC | 0.34 | 0.00 | 0.00 | 0.31 | 0.11 | 14 | Matrix Fibroblast |
| SNX3 | 0.36 | 0.00 | 0.00 | 0.69 | 0.35 | 14 | Matrix Fibroblast |
| SUPT4H1 | 0.37 | 0.00 | 0.00 | 0.40 | 0.17 | 14 | Matrix Fibroblast |
| RCN2 | 0.32 | 0.00 | 0.00 | 0.40 | 0.17 | 14 | Matrix Fibroblast |
| HEXIM1 | 0.48 | 0.00 | 0.00 | 0.31 | 0.12 | 14 | Matrix Fibroblast |
| MORF4L2 | 0.42 | 0.00 | 0.00 | 0.50 | 0.23 | 14 | Matrix Fibroblast |
| APRT | 0.32 | 0.00 | 0.00 | 0.44 | 0.19 | 14 | Matrix Fibroblast |
| CALM2 | 0.40 | 0.00 | 0.00 | 0.92 | 0.66 | 14 | Matrix Fibroblast |
| STMN1 | 0.28 | 0.00 | 0.00 | 0.54 | 0.24 | 14 | Matrix Fibroblast |
| SEC62 | 0.36 | 0.00 | 0.00 | 0.80 | 0.46 | 14 | Matrix Fibroblast |
| GADD45GIP1 | 0.29 | 0.00 | 0.00 | 0.43 | 0.18 | 14 | Matrix Fibroblast |
| NDUFA11 | 0.36 | 0.00 | 0.00 | 0.60 | 0.30 | 14 | Matrix Fibroblast |
| VIM | 0.41 | 0.00 | 0.00 | 0.94 | 0.69 | 14 | Matrix Fibroblast |
| SUMO1 | 0.34 | 0.00 | 0.00 | 0.56 | 0.27 | 14 | Matrix Fibroblast |
| PLS3 | 0.49 | 0.00 | 0.00 | 0.47 | 0.22 | 14 | Matrix Fibroblast |
| MLF2 | 0.39 | 0.00 | 0.00 | 0.37 | 0.15 | 14 | Matrix Fibroblast |
| ZFP36L1 | 0.41 | 0.00 | 0.00 | 0.79 | 0.46 | 14 | Matrix Fibroblast |
| CPQ | 0.29 | 0.00 | 0.00 | 0.27 | 0.10 | 14 | Matrix Fibroblast |
| C1orf52 | 0.33 | 0.00 | 0.00 | 0.27 | 0.09 | 14 | Matrix Fibroblast |
| TBX2 | 0.30 | 0.00 | 0.00 | 0.35 | 0.14 | 14 | Matrix Fibroblast |
| TMED9 | 0.39 | 0.00 | 0.00 | 0.57 | 0.28 | 14 | Matrix Fibroblast |
| VKORC1 | 0.34 | 0.00 | 0.00 | 0.32 | 0.12 | 14 | Matrix Fibroblast |
| SDHC | 0.37 | 0.00 | 0.00 | 0.35 | 0.14 | 14 | Matrix Fibroblast |
| ZFYVE21 | 0.32 | 0.00 | 0.00 | 0.28 | 0.10 | 14 | Matrix Fibroblast |
| TMEM14C | 0.34 | 0.00 | 0.00 | 0.48 | 0.22 | 14 | Matrix Fibroblast |
| TMEM219 | 0.33 | 0.00 | 0.00 | 0.27 | 0.10 | 14 | Matrix Fibroblast |
| CCDC80 | 0.41 | 0.00 | 0.00 | 0.32 | 0.13 | 14 | Matrix Fibroblast |
| RPS19BP1 | 0.33 | 0.00 | 0.00 | 0.33 | 0.13 | 14 | Matrix Fibroblast |
| C1QBP | 0.30 | 0.00 | 0.00 | 0.35 | 0.14 | 14 | Matrix Fibroblast |
| MT-CYB | 0.31 | 0.00 | 0.00 | 0.98 | 0.83 | 14 | Matrix Fibroblast |
| CD151 | 0.31 | 0.00 | 0.00 | 0.44 | 0.19 | 14 | Matrix Fibroblast |
| YWHAQ | 0.35 | 0.00 | 0.00 | 0.70 | 0.38 | 14 | Matrix Fibroblast |
| MRPL43 | 0.32 | 0.00 | 0.00 | 0.34 | 0.14 | 14 | Matrix Fibroblast |
| TXNIP | 0.31 | 0.00 | 0.00 | 0.56 | 0.27 | 14 | Matrix Fibroblast |
| NDUFV1 | 0.35 | 0.00 | 0.00 | 0.34 | 0.14 | 14 | Matrix Fibroblast |
| DARS | 0.39 | 0.00 | 0.00 | 0.27 | 0.10 | 14 | Matrix Fibroblast |
| HNRNPUL1 | 0.36 | 0.00 | 0.00 | 0.31 | 0.12 | 14 | Matrix Fibroblast |
| MEA1 | 0.30 | 0.00 | 0.00 | 0.25 | 0.09 | 14 | Matrix Fibroblast |
| FAM114A1 | 0.31 | 0.00 | 0.00 | 0.26 | 0.10 | 14 | Matrix Fibroblast |
| ATP6V1G1 | 0.31 | 0.00 | 0.00 | 0.65 | 0.34 | 14 | Matrix Fibroblast |
| PRDX3 | 0.32 | 0.00 | 0.00 | 0.30 | 0.12 | 14 | Matrix Fibroblast |
| RNF5 | 0.33 | 0.00 | 0.00 | 0.28 | 0.11 | 14 | Matrix Fibroblast |
| HMGB1 | 0.34 | 0.00 | 0.00 | 0.91 | 0.62 | 14 | Matrix Fibroblast |
| MAF1 | 0.30 | 0.00 | 0.00 | 0.40 | 0.18 | 14 | Matrix Fibroblast |
| SPCS2 | 0.30 | 0.00 | 0.00 | 0.63 | 0.33 | 14 | Matrix Fibroblast |
| SLC39A1 | 0.40 | 0.00 | 0.00 | 0.29 | 0.11 | 14 | Matrix Fibroblast |
| TBCA | 0.36 | 0.00 | 0.00 | 0.64 | 0.35 | 14 | Matrix Fibroblast |
| ANXA5 | 0.31 | 0.00 | 0.00 | 0.73 | 0.43 | 14 | Matrix Fibroblast |
| UQCRQ | 0.32 | 0.00 | 0.00 | 0.66 | 0.36 | 14 | Matrix Fibroblast |
| TMEM109 | 0.31 | 0.00 | 0.00 | 0.32 | 0.13 | 14 | Matrix Fibroblast |
| NDUFB4 | 0.33 | 0.00 | 0.00 | 0.52 | 0.26 | 14 | Matrix Fibroblast |
| CAMTA1 | 0.27 | 0.00 | 0.00 | 0.26 | 0.10 | 14 | Matrix Fibroblast |
| ENG | 0.40 | 0.00 | 0.00 | 0.32 | 0.13 | 14 | Matrix Fibroblast |
| DCTN3 | 0.34 | 0.00 | 0.00 | 0.40 | 0.18 | 14 | Matrix Fibroblast |
| COA3 | 0.33 | 0.00 | 0.00 | 0.32 | 0.14 | 14 | Matrix Fibroblast |
| POLR2I | 0.28 | 0.00 | 0.00 | 0.37 | 0.16 | 14 | Matrix Fibroblast |
| MRPL55 | 0.31 | 0.00 | 0.00 | 0.26 | 0.10 | 14 | Matrix Fibroblast |
| ILF3 | 0.39 | 0.00 | 0.00 | 0.30 | 0.12 | 14 | Matrix Fibroblast |
| POLR2J | 0.31 | 0.00 | 0.00 | 0.34 | 0.14 | 14 | Matrix Fibroblast |
| ECHS1 | 0.27 | 0.00 | 0.00 | 0.36 | 0.16 | 14 | Matrix Fibroblast |
| TMEM123 | 0.32 | 0.00 | 0.00 | 0.33 | 0.14 | 14 | Matrix Fibroblast |
| LSM7 | 0.30 | 0.00 | 0.00 | 0.38 | 0.17 | 14 | Matrix Fibroblast |
| SDHD | 0.30 | 0.00 | 0.00 | 0.39 | 0.18 | 14 | Matrix Fibroblast |
| PHACTR2 | 0.32 | 0.00 | 0.00 | 0.32 | 0.14 | 14 | Matrix Fibroblast |
| NRBP1 | 0.27 | 0.00 | 0.00 | 0.33 | 0.14 | 14 | Matrix Fibroblast |
| VAPA | 0.31 | 0.00 | 0.00 | 0.48 | 0.24 | 14 | Matrix Fibroblast |
| PDIA6 | 0.33 | 0.00 | 0.00 | 0.59 | 0.32 | 14 | Matrix Fibroblast |
| ARMCX3 | 0.31 | 0.00 | 0.00 | 0.30 | 0.12 | 14 | Matrix Fibroblast |
| POLR2E | 0.29 | 0.00 | 0.00 | 0.38 | 0.17 | 14 | Matrix Fibroblast |
| JAGN1 | 0.26 | 0.00 | 0.00 | 0.25 | 0.10 | 14 | Matrix Fibroblast |
| COMT | 0.27 | 0.00 | 0.00 | 0.36 | 0.16 | 14 | Matrix Fibroblast |
| SGCE | 0.33 | 0.00 | 0.00 | 0.27 | 0.11 | 14 | Matrix Fibroblast |
| ANXA4 | 0.27 | 0.00 | 0.00 | 0.25 | 0.10 | 14 | Matrix Fibroblast |
| EIF6 | 0.31 | 0.00 | 0.00 | 0.31 | 0.13 | 14 | Matrix Fibroblast |
| PHIP | 0.29 | 0.00 | 0.00 | 0.36 | 0.16 | 14 | Matrix Fibroblast |
| MED4 | 0.25 | 0.00 | 0.00 | 0.28 | 0.11 | 14 | Matrix Fibroblast |
| ZNHIT1 | 0.29 | 0.00 | 0.00 | 0.35 | 0.16 | 14 | Matrix Fibroblast |
| AUP1 | 0.28 | 0.00 | 0.00 | 0.28 | 0.12 | 14 | Matrix Fibroblast |
| NAA38 | 0.34 | 0.00 | 0.00 | 0.34 | 0.16 | 14 | Matrix Fibroblast |
| CAPZB | 0.30 | 0.00 | 0.00 | 0.72 | 0.44 | 14 | Matrix Fibroblast |
| MRPL41 | 0.28 | 0.00 | 0.00 | 0.28 | 0.12 | 14 | Matrix Fibroblast |
| PTOV1 | 0.28 | 0.00 | 0.00 | 0.33 | 0.15 | 14 | Matrix Fibroblast |
| CIB1 | 0.29 | 0.00 | 0.00 | 0.31 | 0.13 | 14 | Matrix Fibroblast |
| EIF3F | 0.28 | 0.00 | 0.00 | 0.67 | 0.38 | 14 | Matrix Fibroblast |
| DDX18 | 0.25 | 0.00 | 0.00 | 0.34 | 0.15 | 14 | Matrix Fibroblast |
| UBE2N | 0.29 | 0.00 | 0.00 | 0.35 | 0.16 | 14 | Matrix Fibroblast |
| TMBIM6 | 0.29 | 0.00 | 0.00 | 0.68 | 0.40 | 14 | Matrix Fibroblast |
| LRP10 | 0.30 | 0.00 | 0.00 | 0.27 | 0.12 | 14 | Matrix Fibroblast |
| UBE2E1 | 0.27 | 0.00 | 0.00 | 0.28 | 0.12 | 14 | Matrix Fibroblast |
| PSMB5 | 0.27 | 0.00 | 0.00 | 0.47 | 0.24 | 14 | Matrix Fibroblast |
| UBE2V2 | 0.25 | 0.00 | 0.00 | 0.27 | 0.11 | 14 | Matrix Fibroblast |
| PRDX4 | 0.29 | 0.00 | 0.00 | 0.34 | 0.16 | 14 | Matrix Fibroblast |
| C5orf15 | 0.27 | 0.00 | 0.00 | 0.36 | 0.17 | 14 | Matrix Fibroblast |
| ZNF503 | 0.31 | 0.00 | 0.00 | 0.34 | 0.16 | 14 | Matrix Fibroblast |
| ARF4 | 0.28 | 0.00 | 0.00 | 0.53 | 0.29 | 14 | Matrix Fibroblast |
| PPP1CC | 0.25 | 0.00 | 0.00 | 0.26 | 0.11 | 14 | Matrix Fibroblast |
| HNRNPH3 | 0.27 | 0.00 | 0.00 | 0.38 | 0.19 | 14 | Matrix Fibroblast |
| SCOC | 0.27 | 0.00 | 0.00 | 0.29 | 0.13 | 14 | Matrix Fibroblast |
| RNPS1 | 0.27 | 0.00 | 0.00 | 0.33 | 0.16 | 14 | Matrix Fibroblast |
| BBX | 0.25 | 0.00 | 0.00 | 0.29 | 0.13 | 14 | Matrix Fibroblast |
| LDHA | 0.31 | 0.00 | 0.00 | 0.55 | 0.31 | 14 | Matrix Fibroblast |
| PKN2 | 0.32 | 0.00 | 0.00 | 0.36 | 0.19 | 14 | Matrix Fibroblast |
| JUN | 0.30 | 0.00 | 0.00 | 0.64 | 0.38 | 14 | Matrix Fibroblast |
| TM2D2 | 0.26 | 0.00 | 0.00 | 0.32 | 0.15 | 14 | Matrix Fibroblast |
| CCT3 | 0.28 | 0.00 | 0.00 | 0.36 | 0.18 | 14 | Matrix Fibroblast |
| TUBA1B | 0.33 | 0.00 | 0.00 | 0.72 | 0.47 | 14 | Matrix Fibroblast |
| SOD2 | 0.25 | 0.00 | 0.00 | 0.27 | 0.13 | 14 | Matrix Fibroblast |
| PFN1 | 0.27 | 0.00 | 0.00 | 0.77 | 0.49 | 14 | Matrix Fibroblast |
| GIMAP7 | 2.21 | 0.00 | 0.00 | 0.75 | 0.02 | 15 | Endothelial |
| TMEM255B | 1.11 | 0.00 | 0.00 | 0.33 | 0.01 | 15 | Endothelial |
| IL7R | 2.21 | 0.00 | 0.00 | 0.72 | 0.03 | 15 | Endothelial |
| IGFBP2 | 1.84 | 0.00 | 0.00 | 0.68 | 0.03 | 15 | Endothelial |
| HSPA1A | 2.37 | 0.00 | 0.00 | 0.83 | 0.05 | 15 | Endothelial |
| A2M | 2.05 | 0.00 | 0.00 | 0.83 | 0.05 | 15 | Endothelial |
| IL33 | 1.79 | 0.00 | 0.00 | 0.70 | 0.05 | 15 | Endothelial |
| APOL3 | 1.20 | 0.00 | 0.00 | 0.35 | 0.01 | 15 | Endothelial |
| TNFSF10 | 2.40 | 0.00 | 0.00 | 0.90 | 0.10 | 15 | Endothelial |
| RAMP3 | 1.80 | 0.00 | 0.00 | 0.54 | 0.04 | 15 | Endothelial |
| VWF | 1.51 | 0.00 | 0.00 | 0.59 | 0.04 | 15 | Endothelial |
| MLF1 | 1.18 | 0.00 | 0.00 | 0.41 | 0.02 | 15 | Endothelial |
| THEM6 | 0.96 | 0.00 | 0.00 | 0.27 | 0.01 | 15 | Endothelial |
| SPEF2 | 1.03 | 0.00 | 0.00 | 0.26 | 0.01 | 15 | Endothelial |
| FAM167B | 1.13 | 0.00 | 0.00 | 0.35 | 0.02 | 15 | Endothelial |
| UBL5 | 1.84 | 0.00 | 0.00 | 0.69 | 0.07 | 15 | Endothelial |
| ARHGAP18 | 1.71 | 0.00 | 0.00 | 0.70 | 0.07 | 15 | Endothelial |
| BST2 | 2.40 | 0.00 | 0.00 | 0.93 | 0.16 | 15 | Endothelial |
| RGS5 | 1.47 | 0.00 | 0.00 | 0.62 | 0.06 | 15 | Endothelial |
| DTL | 1.23 | 0.00 | 0.00 | 0.35 | 0.02 | 15 | Endothelial |
| LEPR | 1.22 | 0.00 | 0.00 | 0.37 | 0.02 | 15 | Endothelial |
| EDN1 | 2.34 | 0.00 | 0.00 | 0.78 | 0.10 | 15 | Endothelial |
| IFITM2 | 1.60 | 0.00 | 0.00 | 0.74 | 0.08 | 15 | Endothelial |
| SERPINB6 | 1.58 | 0.00 | 0.00 | 0.59 | 0.06 | 15 | Endothelial |
| VAMP5 | 2.00 | 0.00 | 0.00 | 0.85 | 0.13 | 15 | Endothelial |
| RGCC | 2.50 | 0.00 | 0.00 | 0.95 | 0.20 | 15 | Endothelial |
| EGLN3 | 1.25 | 0.00 | 0.00 | 0.38 | 0.03 | 15 | Endothelial |
| THSD1 | 1.52 | 0.00 | 0.00 | 0.58 | 0.06 | 15 | Endothelial |
| ZNF428 | 1.33 | 0.00 | 0.00 | 0.52 | 0.05 | 15 | Endothelial |
| TIMP1 | 1.07 | 0.00 | 0.00 | 0.68 | 0.08 | 15 | Endothelial |
| RPL29 | 1.76 | 0.00 | 0.00 | 0.95 | 0.20 | 15 | Endothelial |
| CCDC85B | 1.70 | 0.00 | 0.00 | 0.83 | 0.14 | 15 | Endothelial |
| PYURF | 1.21 | 0.00 | 0.00 | 0.43 | 0.04 | 15 | Endothelial |
| BCAM | 1.56 | 0.00 | 0.00 | 0.74 | 0.11 | 15 | Endothelial |
| NPDC1 | 1.42 | 0.00 | 0.00 | 0.59 | 0.07 | 15 | Endothelial |
| MGST2 | 1.44 | 0.00 | 0.00 | 0.46 | 0.04 | 15 | Endothelial |
| CXorf36 | 0.98 | 0.00 | 0.00 | 0.27 | 0.02 | 15 | Endothelial |
| RNF144B | 0.84 | 0.00 | 0.00 | 0.26 | 0.01 | 15 | Endothelial |
| SVIP | 1.28 | 0.00 | 0.00 | 0.35 | 0.03 | 15 | Endothelial |
| IFI27L2 | 1.23 | 0.00 | 0.00 | 0.38 | 0.03 | 15 | Endothelial |
| ISOC2 | 0.98 | 0.00 | 0.00 | 0.35 | 0.03 | 15 | Endothelial |
| PROCR | 1.06 | 0.00 | 0.00 | 0.35 | 0.03 | 15 | Endothelial |
| PRCP | 1.59 | 0.00 | 0.00 | 0.69 | 0.11 | 15 | Endothelial |
| RAMP2 | 1.90 | 0.00 | 0.00 | 1.00 | 0.30 | 15 | Endothelial |
| CTNNAL1 | 1.29 | 0.00 | 0.00 | 0.47 | 0.05 | 15 | Endothelial |
| COX7A1 | 1.18 | 0.00 | 0.00 | 0.41 | 0.04 | 15 | Endothelial |
| TNFRSF14 | 0.82 | 0.00 | 0.00 | 0.26 | 0.02 | 15 | Endothelial |
| DEGS1 | 1.01 | 0.00 | 0.00 | 0.33 | 0.03 | 15 | Endothelial |
| GATA3 | 1.10 | 0.00 | 0.00 | 0.32 | 0.02 | 15 | Endothelial |
| CAV1 | 2.06 | 0.00 | 0.00 | 1.00 | 0.37 | 15 | Endothelial |
| PDE3B | 1.20 | 0.00 | 0.00 | 0.35 | 0.03 | 15 | Endothelial |
| SOX7 | 1.32 | 0.00 | 0.00 | 0.51 | 0.06 | 15 | Endothelial |
| FCGRT | 1.60 | 0.00 | 0.00 | 0.73 | 0.13 | 15 | Endothelial |
| STOM | 1.29 | 0.00 | 0.00 | 0.62 | 0.09 | 15 | Endothelial |
| GNG11 | 1.68 | 0.00 | 0.00 | 0.98 | 0.32 | 15 | Endothelial |
| MAP3K11 | 1.24 | 0.00 | 0.00 | 0.46 | 0.05 | 15 | Endothelial |
| POSTN | 1.09 | 0.00 | 0.00 | 0.64 | 0.10 | 15 | Endothelial |
| SOX18 | 1.19 | 0.00 | 0.00 | 0.47 | 0.06 | 15 | Endothelial |
| CAV2 | 1.57 | 0.00 | 0.00 | 0.89 | 0.22 | 15 | Endothelial |
| CCDC130 | 0.97 | 0.00 | 0.00 | 0.33 | 0.03 | 15 | Endothelial |
| ITM2A | 1.16 | 0.00 | 0.00 | 0.57 | 0.08 | 15 | Endothelial |
| IGFBP4 | 1.63 | 0.00 | 0.00 | 0.94 | 0.28 | 15 | Endothelial |
| GAPDH | 1.34 | 0.00 | 0.00 | 0.85 | 0.19 | 15 | Endothelial |
| ROBO4 | 1.27 | 0.00 | 0.00 | 0.49 | 0.06 | 15 | Endothelial |
| EPAS1 | 1.54 | 0.00 | 0.00 | 0.96 | 0.28 | 15 | Endothelial |
| FDPS | 1.36 | 0.00 | 0.00 | 0.54 | 0.08 | 15 | Endothelial |
| B2M | 2.22 | 0.00 | 0.00 | 1.00 | 0.70 | 15 | Endothelial |
| ENG | 1.41 | 0.00 | 0.00 | 0.69 | 0.13 | 15 | Endothelial |
| BEX4 | 1.06 | 0.00 | 0.00 | 0.33 | 0.03 | 15 | Endothelial |
| CD59 | 1.07 | 0.00 | 0.00 | 0.52 | 0.07 | 15 | Endothelial |
| NTHL1 | 0.85 | 0.00 | 0.00 | 0.27 | 0.02 | 15 | Endothelial |
| PCDH12 | 0.97 | 0.00 | 0.00 | 0.33 | 0.03 | 15 | Endothelial |
| MARCKSL1 | 1.78 | 0.00 | 0.00 | 0.82 | 0.23 | 15 | Endothelial |
| MT-ND3 | 1.58 | 0.00 | 0.00 | 0.99 | 0.51 | 15 | Endothelial |
| GMFG | 1.32 | 0.00 | 0.00 | 0.72 | 0.15 | 15 | Endothelial |
| SRPX | 1.05 | 0.00 | 0.00 | 0.33 | 0.04 | 15 | Endothelial |
| MGST3 | 1.20 | 0.00 | 0.00 | 0.68 | 0.14 | 15 | Endothelial |
| TMEM123 | 1.37 | 0.00 | 0.00 | 0.68 | 0.14 | 15 | Endothelial |
| PRMT1 | 1.39 | 0.00 | 0.00 | 0.63 | 0.13 | 15 | Endothelial |
| TMSB4X | 1.42 | 0.00 | 0.00 | 1.00 | 0.90 | 15 | Endothelial |
| SLC9A3R2 | 1.41 | 0.00 | 0.00 | 0.85 | 0.23 | 15 | Endothelial |
| NME3 | 0.91 | 0.00 | 0.00 | 0.27 | 0.03 | 15 | Endothelial |
| TSPAN4 | 1.17 | 0.00 | 0.00 | 0.56 | 0.10 | 15 | Endothelial |
| FRY | 1.01 | 0.00 | 0.00 | 0.36 | 0.04 | 15 | Endothelial |
| CLEC14A | 1.30 | 0.00 | 0.00 | 0.80 | 0.19 | 15 | Endothelial |
| KIAA1217 | 1.03 | 0.00 | 0.00 | 0.44 | 0.06 | 15 | Endothelial |
| ESAM | 1.27 | 0.00 | 0.00 | 0.82 | 0.21 | 15 | Endothelial |
| STXBP6 | 1.05 | 0.00 | 0.00 | 0.42 | 0.06 | 15 | Endothelial |
| POLR2L | 1.46 | 0.00 | 0.00 | 0.74 | 0.20 | 15 | Endothelial |
| LRRC32 | 1.00 | 0.00 | 0.00 | 0.30 | 0.03 | 15 | Endothelial |
| TIE1 | 1.17 | 0.00 | 0.00 | 0.62 | 0.12 | 15 | Endothelial |
| TMEM88 | 1.25 | 0.00 | 0.00 | 0.58 | 0.11 | 15 | Endothelial |
| APLNR | 1.09 | 0.00 | 0.00 | 0.48 | 0.08 | 15 | Endothelial |
| PRKCH | 0.97 | 0.00 | 0.00 | 0.38 | 0.05 | 15 | Endothelial |
| CD46 | 0.78 | 0.00 | 0.00 | 0.27 | 0.03 | 15 | Endothelial |
| EBPL | 0.94 | 0.00 | 0.00 | 0.40 | 0.05 | 15 | Endothelial |
| S100A10 | 1.24 | 0.00 | 0.00 | 0.98 | 0.41 | 15 | Endothelial |
| MRI1 | 1.15 | 0.00 | 0.00 | 0.37 | 0.05 | 15 | Endothelial |
| HYAL2 | 1.16 | 0.00 | 0.00 | 0.53 | 0.10 | 15 | Endothelial |
| HIST1H2BG | 0.89 | 0.00 | 0.00 | 0.27 | 0.03 | 15 | Endothelial |
| LIFR | 1.20 | 0.00 | 0.00 | 0.46 | 0.08 | 15 | Endothelial |
| CLDN5 | 1.19 | 0.00 | 0.00 | 0.86 | 0.24 | 15 | Endothelial |
| JAM2 | 1.19 | 0.00 | 0.00 | 0.44 | 0.07 | 15 | Endothelial |
| ARHGAP29 | 1.14 | 0.00 | 0.00 | 0.83 | 0.23 | 15 | Endothelial |
| TFPI | 1.32 | 0.00 | 0.00 | 0.57 | 0.12 | 15 | Endothelial |
| ISG15 | 1.21 | 0.00 | 0.00 | 0.40 | 0.06 | 15 | Endothelial |
| KCTD12 | 1.28 | 0.00 | 0.00 | 0.48 | 0.09 | 15 | Endothelial |
| FKBP1A | 1.24 | 0.00 | 0.00 | 0.95 | 0.44 | 15 | Endothelial |
| CCND1 | 1.40 | 0.00 | 0.00 | 0.73 | 0.20 | 15 | Endothelial |
| CD93 | 0.93 | 0.00 | 0.00 | 0.86 | 0.22 | 15 | Endothelial |
| RHOC | 1.22 | 0.00 | 0.00 | 0.82 | 0.24 | 15 | Endothelial |
| RPL13A | 1.17 | 0.00 | 0.00 | 0.98 | 0.59 | 15 | Endothelial |
| HNRNPA1 | 1.07 | 0.00 | 0.00 | 0.90 | 0.31 | 15 | Endothelial |
| STMN1 | 1.15 | 0.00 | 0.00 | 0.82 | 0.25 | 15 | Endothelial |
| TM4SF1 | 1.13 | 0.00 | 0.00 | 0.80 | 0.22 | 15 | Endothelial |
| COMMD6 | 1.09 | 0.00 | 0.00 | 0.68 | 0.17 | 15 | Endothelial |
| TMSB10 | 1.07 | 0.00 | 0.00 | 1.00 | 0.88 | 15 | Endothelial |
| GUK1 | 1.11 | 0.00 | 0.00 | 0.73 | 0.20 | 15 | Endothelial |
| PTMA | 0.86 | 0.00 | 0.00 | 1.00 | 0.92 | 15 | Endothelial |
| CIRBP | 1.10 | 0.00 | 0.00 | 0.84 | 0.26 | 15 | Endothelial |
| KRT10 | 1.05 | 0.00 | 0.00 | 0.40 | 0.06 | 15 | Endothelial |
| PPIA | 1.29 | 0.00 | 0.00 | 0.73 | 0.22 | 15 | Endothelial |
| FOXF1 | 1.13 | 0.00 | 0.00 | 0.61 | 0.13 | 15 | Endothelial |
| XRCC5 | 1.03 | 0.00 | 0.00 | 0.38 | 0.06 | 15 | Endothelial |
| HPGD | 0.86 | 0.00 | 0.00 | 0.90 | 0.27 | 15 | Endothelial |
| FSCN1 | 0.98 | 0.00 | 0.00 | 0.46 | 0.08 | 15 | Endothelial |
| MT-ND4 | 1.01 | 0.00 | 0.00 | 0.99 | 0.77 | 15 | Endothelial |
| HMGN3 | 1.04 | 0.00 | 0.00 | 0.43 | 0.08 | 15 | Endothelial |
| TMEM173 | 0.95 | 0.00 | 0.00 | 0.32 | 0.04 | 15 | Endothelial |
| SPHK1 | 0.79 | 0.00 | 0.00 | 0.26 | 0.03 | 15 | Endothelial |
| OCIAD2 | 0.94 | 0.00 | 0.00 | 0.32 | 0.04 | 15 | Endothelial |
| SCARF1 | 0.80 | 0.00 | 0.00 | 0.30 | 0.04 | 15 | Endothelial |
| GSTP1 | 1.01 | 0.00 | 0.00 | 0.80 | 0.24 | 15 | Endothelial |
| CALCRL | 0.99 | 0.00 | 0.00 | 0.85 | 0.26 | 15 | Endothelial |
| KIT | 0.85 | 0.00 | 0.00 | 0.46 | 0.08 | 15 | Endothelial |
| CHST12 | 0.99 | 0.00 | 0.00 | 0.33 | 0.05 | 15 | Endothelial |
| UBE2L6 | 0.96 | 0.00 | 0.00 | 0.36 | 0.06 | 15 | Endothelial |
| S100A16 | 1.08 | 0.00 | 0.00 | 0.82 | 0.26 | 15 | Endothelial |
| H3F3A | 1.00 | 0.00 | 0.00 | 0.96 | 0.62 | 15 | Endothelial |
| LXN | 1.36 | 0.00 | 0.00 | 0.54 | 0.13 | 15 | Endothelial |
| SRP14 | 1.05 | 0.00 | 0.00 | 0.88 | 0.34 | 15 | Endothelial |
| MYL6 | 0.99 | 0.00 | 0.00 | 0.98 | 0.72 | 15 | Endothelial |
| NDUFAF3 | 0.94 | 0.00 | 0.00 | 0.42 | 0.08 | 15 | Endothelial |
| MMRN2 | 0.74 | 0.00 | 0.00 | 0.33 | 0.05 | 15 | Endothelial |
| CD74 | 1.00 | 0.00 | 0.00 | 0.62 | 0.15 | 15 | Endothelial |
| LEPROTL1 | 1.00 | 0.00 | 0.00 | 0.62 | 0.16 | 15 | Endothelial |
| GIMAP4 | 0.86 | 0.00 | 0.00 | 0.69 | 0.18 | 15 | Endothelial |
| VAMP2 | 0.86 | 0.00 | 0.00 | 0.47 | 0.10 | 15 | Endothelial |
| CARHSP1 | 1.11 | 0.00 | 0.00 | 0.59 | 0.15 | 15 | Endothelial |
| HSPB8 | 0.88 | 0.00 | 0.00 | 0.27 | 0.04 | 15 | Endothelial |
| NAA38 | 1.15 | 0.00 | 0.00 | 0.59 | 0.16 | 15 | Endothelial |
| HOXA5 | 1.00 | 0.00 | 0.00 | 0.57 | 0.14 | 15 | Endothelial |
| PSMB9 | 0.97 | 0.00 | 0.00 | 0.47 | 0.10 | 15 | Endothelial |
| VAMP3 | 0.96 | 0.00 | 0.00 | 0.59 | 0.15 | 15 | Endothelial |
| MT-CO2 | 0.84 | 0.00 | 0.00 | 1.00 | 0.79 | 15 | Endothelial |
| FAM13C | 0.83 | 0.00 | 0.00 | 0.32 | 0.05 | 15 | Endothelial |
| ABHD17A | 0.84 | 0.00 | 0.00 | 0.38 | 0.07 | 15 | Endothelial |
| TSPAN5 | 0.76 | 0.00 | 0.00 | 0.27 | 0.04 | 15 | Endothelial |
| TXNIP | 1.06 | 0.00 | 0.00 | 0.78 | 0.27 | 15 | Endothelial |
| EGFL7 | 0.75 | 0.00 | 0.00 | 0.90 | 0.29 | 15 | Endothelial |
| WSB1 | 1.05 | 0.00 | 0.00 | 0.73 | 0.24 | 15 | Endothelial |
| EFNA1 | 0.99 | 0.00 | 0.00 | 0.42 | 0.09 | 15 | Endothelial |
| PPDPF | 1.03 | 0.00 | 0.00 | 0.70 | 0.23 | 15 | Endothelial |
| MYLIP | 0.80 | 0.00 | 0.00 | 0.43 | 0.09 | 15 | Endothelial |
| LDB2 | 0.78 | 0.00 | 0.00 | 0.65 | 0.18 | 15 | Endothelial |
| GSTO1 | 0.77 | 0.00 | 0.00 | 0.41 | 0.08 | 15 | Endothelial |
| NUDT14 | 0.87 | 0.00 | 0.00 | 0.36 | 0.07 | 15 | Endothelial |
| SDF2L1 | 0.77 | 0.00 | 0.00 | 0.30 | 0.05 | 15 | Endothelial |
| CTNND1 | 1.01 | 0.00 | 0.00 | 0.59 | 0.17 | 15 | Endothelial |
| SNX3 | 0.95 | 0.00 | 0.00 | 0.86 | 0.36 | 15 | Endothelial |
| HSPB1 | 0.93 | 0.00 | 0.00 | 0.63 | 0.17 | 15 | Endothelial |
| DYNLL1 | 1.02 | 0.00 | 0.00 | 0.90 | 0.47 | 15 | Endothelial |
| PRPSAP1 | 1.03 | 0.00 | 0.00 | 0.40 | 0.08 | 15 | Endothelial |
| SERF2 | 0.89 | 0.00 | 0.00 | 0.95 | 0.60 | 15 | Endothelial |
| MIF | 0.88 | 0.00 | 0.00 | 0.72 | 0.23 | 15 | Endothelial |
| SPSB3 | 0.73 | 0.00 | 0.00 | 0.31 | 0.05 | 15 | Endothelial |
| MT-ND2 | 0.83 | 0.00 | 0.00 | 0.99 | 0.72 | 15 | Endothelial |
| CD320 | 0.82 | 0.00 | 0.00 | 0.26 | 0.04 | 15 | Endothelial |
| AAMDC | 0.90 | 0.00 | 0.00 | 0.38 | 0.08 | 15 | Endothelial |
| SNRPN | 0.79 | 0.00 | 0.00 | 0.30 | 0.05 | 15 | Endothelial |
| APH1A | 0.96 | 0.00 | 0.00 | 0.52 | 0.14 | 15 | Endothelial |
| TRMT112 | 0.97 | 0.00 | 0.00 | 0.61 | 0.18 | 15 | Endothelial |
| GPIHBP1 | 0.80 | 0.00 | 0.00 | 0.61 | 0.16 | 15 | Endothelial |
| EMCN | 0.73 | 0.00 | 0.00 | 0.51 | 0.12 | 15 | Endothelial |
| TSPAN15 | 0.76 | 0.00 | 0.00 | 0.38 | 0.08 | 15 | Endothelial |
| HSPA12B | 0.83 | 0.00 | 0.00 | 0.35 | 0.07 | 15 | Endothelial |
| CDC42EP1 | 0.88 | 0.00 | 0.00 | 0.42 | 0.10 | 15 | Endothelial |
| RPL7A | 0.91 | 0.00 | 0.00 | 0.90 | 0.44 | 15 | Endothelial |
| JAM3 | 0.76 | 0.00 | 0.00 | 0.26 | 0.04 | 15 | Endothelial |
| TGFBR2 | 0.93 | 0.00 | 0.00 | 0.49 | 0.12 | 15 | Endothelial |
| ETS2 | 1.08 | 0.00 | 0.00 | 0.49 | 0.13 | 15 | Endothelial |
| DACH1 | 0.81 | 0.00 | 0.00 | 0.31 | 0.06 | 15 | Endothelial |
| PECAM1 | 0.75 | 0.00 | 0.00 | 0.78 | 0.26 | 15 | Endothelial |
| FABP5 | 1.47 | 0.00 | 0.00 | 0.53 | 0.15 | 15 | Endothelial |
| SH3BP5 | 0.79 | 0.00 | 0.00 | 0.65 | 0.20 | 15 | Endothelial |
| HSPG2 | 0.72 | 0.00 | 0.00 | 0.36 | 0.07 | 15 | Endothelial |
| MAOA | 0.84 | 0.00 | 0.00 | 0.38 | 0.08 | 15 | Endothelial |
| DUT | 0.86 | 0.00 | 0.00 | 0.44 | 0.11 | 15 | Endothelial |
| TMA7 | 0.94 | 0.00 | 0.00 | 0.80 | 0.35 | 15 | Endothelial |
| TMEM230 | 0.88 | 0.00 | 0.00 | 0.37 | 0.08 | 15 | Endothelial |
| TMEM219 | 0.86 | 0.00 | 0.00 | 0.42 | 0.10 | 15 | Endothelial |
| CYYR1 | 0.77 | 0.00 | 0.00 | 0.56 | 0.15 | 15 | Endothelial |
| CDH5 | 0.70 | 0.00 | 0.00 | 0.75 | 0.24 | 15 | Endothelial |
| STARD3 | 0.82 | 0.00 | 0.00 | 0.32 | 0.06 | 15 | Endothelial |
| CAMTA1 | 0.82 | 0.00 | 0.00 | 0.42 | 0.10 | 15 | Endothelial |
| ACVRL1 | 0.66 | 0.00 | 0.00 | 0.64 | 0.19 | 15 | Endothelial |
| RPS27A | 0.66 | 0.00 | 0.00 | 0.99 | 0.78 | 15 | Endothelial |
| BRK1 | 0.88 | 0.00 | 0.00 | 0.77 | 0.29 | 15 | Endothelial |
| CD63 | 0.85 | 0.00 | 0.00 | 0.83 | 0.35 | 15 | Endothelial |
| RPL24 | 0.83 | 0.00 | 0.00 | 0.94 | 0.50 | 15 | Endothelial |
| PARVB | 0.72 | 0.00 | 0.00 | 0.33 | 0.07 | 15 | Endothelial |
| MAGEH1 | 0.99 | 0.00 | 0.00 | 0.36 | 0.08 | 15 | Endothelial |
| RAB13 | 0.82 | 0.00 | 0.00 | 0.47 | 0.12 | 15 | Endothelial |
| VAT1 | 0.72 | 0.00 | 0.00 | 0.42 | 0.10 | 15 | Endothelial |
| PIM3 | 0.95 | 0.00 | 0.00 | 0.36 | 0.08 | 15 | Endothelial |
| LAP3 | 0.88 | 0.00 | 0.00 | 0.43 | 0.11 | 15 | Endothelial |
| GJA4 | 0.72 | 0.00 | 0.00 | 0.37 | 0.08 | 15 | Endothelial |
| LMCD1 | 0.73 | 0.00 | 0.00 | 0.36 | 0.08 | 15 | Endothelial |
| GPR146 | 0.80 | 0.00 | 0.00 | 0.30 | 0.06 | 15 | Endothelial |
| CCNI | 0.72 | 0.00 | 0.00 | 0.63 | 0.20 | 15 | Endothelial |
| GLUL | 0.99 | 0.00 | 0.00 | 0.63 | 0.22 | 15 | Endothelial |
| BLOC1S1 | 0.74 | 0.00 | 0.00 | 0.37 | 0.09 | 15 | Endothelial |
| RHOB | 0.96 | 0.00 | 0.00 | 0.46 | 0.13 | 15 | Endothelial |
| WARS | 0.93 | 0.00 | 0.00 | 0.40 | 0.10 | 15 | Endothelial |
| AP2M1 | 0.89 | 0.00 | 0.00 | 0.54 | 0.17 | 15 | Endothelial |
| LAMTOR4 | 0.84 | 0.00 | 0.00 | 0.51 | 0.15 | 15 | Endothelial |
| RAB5C | 0.83 | 0.00 | 0.00 | 0.49 | 0.14 | 15 | Endothelial |
| MAGED2 | 0.78 | 0.00 | 0.00 | 0.63 | 0.21 | 15 | Endothelial |
| PLXNC1 | 0.64 | 0.00 | 0.00 | 0.27 | 0.05 | 15 | Endothelial |
| CD40 | 0.66 | 0.00 | 0.00 | 0.30 | 0.06 | 15 | Endothelial |
| UQCR11 | 0.86 | 0.00 | 0.00 | 0.75 | 0.32 | 15 | Endothelial |
| HERPUD1 | 0.94 | 0.00 | 0.00 | 0.49 | 0.15 | 15 | Endothelial |
| PIN4 | 0.73 | 0.00 | 0.00 | 0.46 | 0.13 | 15 | Endothelial |
| SEC14L1 | 0.77 | 0.00 | 0.00 | 0.44 | 0.12 | 15 | Endothelial |
| PLAT | 1.27 | 0.00 | 0.00 | 0.30 | 0.06 | 15 | Endothelial |
| WDR82 | 0.69 | 0.00 | 0.00 | 0.28 | 0.06 | 15 | Endothelial |
| EEF1D | 0.87 | 0.00 | 0.00 | 0.70 | 0.30 | 15 | Endothelial |
| HEY1 | 0.77 | 0.00 | 0.00 | 0.27 | 0.05 | 15 | Endothelial |
| GABPB1 | 0.69 | 0.00 | 0.00 | 0.27 | 0.05 | 15 | Endothelial |
| HMGB1 | 0.72 | 0.00 | 0.00 | 0.94 | 0.62 | 15 | Endothelial |
| MT-CYB | 0.58 | 0.00 | 0.00 | 1.00 | 0.83 | 15 | Endothelial |
| ZNF22 | 0.73 | 0.00 | 0.00 | 0.44 | 0.12 | 15 | Endothelial |
| SLC44A2 | 0.68 | 0.00 | 0.00 | 0.32 | 0.07 | 15 | Endothelial |
| UBC | 0.82 | 0.00 | 0.00 | 0.88 | 0.51 | 15 | Endothelial |
| MYCT1 | 0.57 | 0.00 | 0.00 | 0.48 | 0.13 | 15 | Endothelial |
| SUMO2 | 0.88 | 0.00 | 0.00 | 0.75 | 0.34 | 15 | Endothelial |
| SREK1IP1 | 0.70 | 0.00 | 0.00 | 0.42 | 0.12 | 15 | Endothelial |
| HNRNPA0 | 0.76 | 0.00 | 0.00 | 0.64 | 0.24 | 15 | Endothelial |
| PET100 | 0.91 | 0.00 | 0.00 | 0.48 | 0.15 | 15 | Endothelial |
| PPA1 | 0.79 | 0.00 | 0.00 | 0.33 | 0.08 | 15 | Endothelial |
| EDF1 | 0.70 | 0.00 | 0.00 | 0.78 | 0.34 | 15 | Endothelial |
| SMIM7 | 0.80 | 0.00 | 0.00 | 0.42 | 0.12 | 15 | Endothelial |
| PALMD | 0.66 | 0.00 | 0.00 | 0.46 | 0.13 | 15 | Endothelial |
| PAK4 | 0.63 | 0.00 | 0.00 | 0.26 | 0.05 | 15 | Endothelial |
| TNIP1 | 0.63 | 0.00 | 0.00 | 0.26 | 0.05 | 15 | Endothelial |
| DPYSL3 | 0.75 | 0.00 | 0.00 | 0.37 | 0.10 | 15 | Endothelial |
| PSMB3 | 0.70 | 0.00 | 0.00 | 0.67 | 0.27 | 15 | Endothelial |
| S100A6 | 0.51 | 0.00 | 0.00 | 0.83 | 0.35 | 15 | Endothelial |
| XRCC6 | 0.78 | 0.00 | 0.00 | 0.30 | 0.07 | 15 | Endothelial |
| ARHGEF15 | 0.56 | 0.00 | 0.00 | 0.33 | 0.08 | 15 | Endothelial |
| OST4 | 0.81 | 0.00 | 0.00 | 0.67 | 0.29 | 15 | Endothelial |
| SGK1 | 0.97 | 0.00 | 0.00 | 0.54 | 0.19 | 15 | Endothelial |
| BMPR2 | 0.70 | 0.00 | 0.00 | 0.40 | 0.11 | 15 | Endothelial |
| MDK | 0.56 | 0.00 | 0.00 | 0.64 | 0.22 | 15 | Endothelial |
| DUSP1 | 0.90 | 0.00 | 0.00 | 0.51 | 0.17 | 15 | Endothelial |
| TSPAN12 | 0.60 | 0.00 | 0.00 | 0.31 | 0.07 | 15 | Endothelial |
| TMEM100 | 0.56 | 0.00 | 0.00 | 0.77 | 0.30 | 15 | Endothelial |
| N4BP2L2 | 0.69 | 0.00 | 0.00 | 0.62 | 0.23 | 15 | Endothelial |
| ADAR | 0.72 | 0.00 | 0.00 | 0.30 | 0.07 | 15 | Endothelial |
| PFN1 | 0.69 | 0.00 | 0.00 | 0.86 | 0.49 | 15 | Endothelial |
| SYNGR2 | 0.72 | 0.00 | 0.00 | 0.37 | 0.10 | 15 | Endothelial |
| TMBIM4 | 0.73 | 0.00 | 0.00 | 0.48 | 0.16 | 15 | Endothelial |
| DUSP6 | 0.68 | 0.00 | 0.00 | 0.54 | 0.18 | 15 | Endothelial |
| C12orf57 | 0.74 | 0.00 | 0.00 | 0.49 | 0.17 | 15 | Endothelial |
| UACA | 0.74 | 0.00 | 0.00 | 0.47 | 0.15 | 15 | Endothelial |
| PHACTR2 | 0.69 | 0.00 | 0.00 | 0.46 | 0.14 | 15 | Endothelial |
| RPLP2 | 0.49 | 0.00 | 0.00 | 0.99 | 0.83 | 15 | Endothelial |
| MEF2C | 0.57 | 0.00 | 0.00 | 0.54 | 0.18 | 15 | Endothelial |
| CD34 | 0.70 | 0.00 | 0.00 | 0.58 | 0.21 | 15 | Endothelial |
| NUMB | 0.68 | 0.00 | 0.00 | 0.35 | 0.09 | 15 | Endothelial |
| TLE4 | 1.15 | 0.00 | 0.00 | 0.42 | 0.13 | 15 | Endothelial |
| ZNF503 | 0.84 | 0.00 | 0.00 | 0.48 | 0.16 | 15 | Endothelial |
| RPL23A | 0.53 | 0.00 | 0.00 | 0.98 | 0.69 | 15 | Endothelial |
| TMEM258 | 0.73 | 0.00 | 0.00 | 0.74 | 0.35 | 15 | Endothelial |
| RALB | 0.80 | 0.00 | 0.00 | 0.53 | 0.20 | 15 | Endothelial |
| SCAND1 | 0.51 | 0.00 | 0.00 | 0.52 | 0.17 | 15 | Endothelial |
| PKIG | 0.60 | 0.00 | 0.00 | 0.37 | 0.10 | 15 | Endothelial |
| PODXL | 0.62 | 0.00 | 0.00 | 0.40 | 0.11 | 15 | Endothelial |
| NDUFB2 | 0.85 | 0.00 | 0.00 | 0.58 | 0.24 | 15 | Endothelial |
| SOX4 | 0.85 | 0.00 | 0.00 | 0.75 | 0.40 | 15 | Endothelial |
| TPRKB | 0.63 | 0.00 | 0.00 | 0.26 | 0.06 | 15 | Endothelial |
| NUDT4 | 0.81 | 0.00 | 0.00 | 0.47 | 0.16 | 15 | Endothelial |
| TUFM | 0.65 | 0.00 | 0.00 | 0.31 | 0.08 | 15 | Endothelial |
| CMTM3 | 0.75 | 0.00 | 0.00 | 0.44 | 0.15 | 15 | Endothelial |
| ICAM2 | 0.55 | 0.00 | 0.00 | 0.52 | 0.18 | 15 | Endothelial |
| PEA15 | 0.62 | 0.00 | 0.00 | 0.46 | 0.15 | 15 | Endothelial |
| TSPAN7 | 0.52 | 0.00 | 0.00 | 0.59 | 0.22 | 15 | Endothelial |
| TSC22D1 | 0.80 | 0.00 | 0.00 | 0.72 | 0.35 | 15 | Endothelial |
| ITM2B | 0.53 | 0.00 | 0.00 | 0.98 | 0.77 | 15 | Endothelial |
| GPX3 | 0.31 | 0.00 | 0.00 | 0.46 | 0.15 | 15 | Endothelial |
| RGS3 | 0.61 | 0.00 | 0.00 | 0.31 | 0.08 | 15 | Endothelial |
| RPS19BP1 | 0.63 | 0.00 | 0.00 | 0.41 | 0.13 | 15 | Endothelial |
| PLK2 | 0.68 | 0.00 | 0.00 | 0.48 | 0.17 | 15 | Endothelial |
| SNRPD2 | 0.69 | 0.00 | 0.00 | 0.64 | 0.28 | 15 | Endothelial |
| PIH1D1 | 0.79 | 0.00 | 0.00 | 0.27 | 0.07 | 15 | Endothelial |
| ANAPC16 | 0.67 | 0.00 | 0.00 | 0.52 | 0.20 | 15 | Endothelial |
| TMEM140 | 0.59 | 0.00 | 0.00 | 0.30 | 0.08 | 15 | Endothelial |
| CD36 | 0.43 | 0.00 | 0.00 | 0.79 | 0.33 | 15 | Endothelial |
| RAB11A | 0.65 | 0.00 | 0.00 | 0.64 | 0.28 | 15 | Endothelial |
| RBMX | 0.65 | 0.00 | 0.00 | 0.42 | 0.14 | 15 | Endothelial |
| CD2BP2 | 0.61 | 0.00 | 0.00 | 0.27 | 0.07 | 15 | Endothelial |
| MRPS7 | 0.56 | 0.00 | 0.00 | 0.30 | 0.08 | 15 | Endothelial |
| FOXN3 | 0.63 | 0.00 | 0.00 | 0.32 | 0.09 | 15 | Endothelial |
| VPS28 | 0.76 | 0.00 | 0.00 | 0.52 | 0.21 | 15 | Endothelial |
| RPS13 | 0.52 | 0.00 | 0.00 | 0.94 | 0.67 | 15 | Endothelial |
| C1orf43 | 0.69 | 0.00 | 0.00 | 0.32 | 0.09 | 15 | Endothelial |
| DHRS3 | 0.69 | 0.00 | 0.00 | 0.27 | 0.07 | 15 | Endothelial |
| YIPF3 | 0.67 | 0.00 | 0.00 | 0.42 | 0.14 | 15 | Endothelial |
| GABARAPL2 | 0.77 | 0.00 | 0.00 | 0.58 | 0.25 | 15 | Endothelial |
| MRPL51 | 0.76 | 0.00 | 0.00 | 0.46 | 0.17 | 15 | Endothelial |
| CNRIP1 | 0.49 | 0.00 | 0.00 | 0.26 | 0.06 | 15 | Endothelial |
| CDC37 | 0.62 | 0.00 | 0.00 | 0.65 | 0.29 | 15 | Endothelial |
| MRPS34 | 0.68 | 0.00 | 0.00 | 0.33 | 0.10 | 15 | Endothelial |
| PSME1 | 0.70 | 0.00 | 0.00 | 0.69 | 0.34 | 15 | Endothelial |
| TBX3 | 0.68 | 0.00 | 0.00 | 0.56 | 0.22 | 15 | Endothelial |
| SSR4 | 0.59 | 0.00 | 0.00 | 0.63 | 0.27 | 15 | Endothelial |
| COMMD2 | 0.58 | 0.00 | 0.00 | 0.35 | 0.10 | 15 | Endothelial |
| HMBOX1 | 0.68 | 0.00 | 0.00 | 0.28 | 0.08 | 15 | Endothelial |
| RPL7 | 0.45 | 0.00 | 0.00 | 0.98 | 0.77 | 15 | Endothelial |
| SPARCL1 | 0.49 | 0.00 | 0.00 | 0.82 | 0.42 | 15 | Endothelial |
| RGL2 | 0.69 | 0.00 | 0.00 | 0.32 | 0.09 | 15 | Endothelial |
| DOCK6 | 0.54 | 0.00 | 0.00 | 0.27 | 0.07 | 15 | Endothelial |
| ATP6V0E1 | 0.64 | 0.00 | 0.00 | 0.57 | 0.23 | 15 | Endothelial |
| SRSF3 | 0.64 | 0.00 | 0.00 | 0.63 | 0.28 | 15 | Endothelial |
| UXT | 0.62 | 0.00 | 0.00 | 0.26 | 0.07 | 15 | Endothelial |
| CLIC1 | 0.59 | 0.00 | 0.00 | 0.77 | 0.39 | 15 | Endothelial |
| CFL1 | 0.50 | 0.00 | 0.00 | 0.93 | 0.61 | 15 | Endothelial |
| MYZAP | 0.47 | 0.00 | 0.00 | 0.44 | 0.15 | 15 | Endothelial |
| NPR3 | 0.54 | 0.00 | 0.00 | 0.37 | 0.12 | 15 | Endothelial |
| ERICH1 | 0.57 | 0.00 | 0.00 | 0.26 | 0.07 | 15 | Endothelial |
| ARMCX3 | 0.64 | 0.00 | 0.00 | 0.38 | 0.13 | 15 | Endothelial |
| RPIA | 0.47 | 0.00 | 0.00 | 0.26 | 0.07 | 15 | Endothelial |
| NEDD8 | 0.60 | 0.00 | 0.00 | 0.68 | 0.32 | 15 | Endothelial |
| C9orf16 | 0.61 | 0.00 | 0.00 | 0.28 | 0.08 | 15 | Endothelial |
| DDIT4 | 0.63 | 0.00 | 0.00 | 0.27 | 0.07 | 15 | Endothelial |
| WWTR1 | 0.68 | 0.00 | 0.00 | 0.42 | 0.15 | 15 | Endothelial |
| S100A4 | 0.60 | 0.00 | 0.00 | 0.33 | 0.10 | 15 | Endothelial |
| CLEC3B | 0.50 | 0.00 | 0.00 | 0.27 | 0.07 | 15 | Endothelial |
| LPAR6 | 0.60 | 0.00 | 0.00 | 0.31 | 0.09 | 15 | Endothelial |
| LAMTOR1 | 0.68 | 0.00 | 0.00 | 0.41 | 0.15 | 15 | Endothelial |
| BTF3 | 0.55 | 0.00 | 0.00 | 0.86 | 0.54 | 15 | Endothelial |
| MZT2B | 0.75 | 0.00 | 0.00 | 0.30 | 0.09 | 15 | Endothelial |
| RALA | 0.67 | 0.00 | 0.00 | 0.56 | 0.24 | 15 | Endothelial |
| PARK7 | 0.62 | 0.00 | 0.00 | 0.62 | 0.28 | 15 | Endothelial |
| DGKZ | 0.55 | 0.00 | 0.00 | 0.28 | 0.08 | 15 | Endothelial |
| ZNHIT1 | 0.62 | 0.00 | 0.00 | 0.43 | 0.16 | 15 | Endothelial |
| LSM3 | 0.61 | 0.00 | 0.00 | 0.37 | 0.13 | 15 | Endothelial |
| TPST2 | 0.46 | 0.00 | 0.00 | 0.37 | 0.12 | 15 | Endothelial |
| RALY | 0.62 | 0.00 | 0.00 | 0.40 | 0.14 | 15 | Endothelial |
| FIS1 | 0.66 | 0.00 | 0.00 | 0.63 | 0.30 | 15 | Endothelial |
| NDRG2 | 0.49 | 0.00 | 0.00 | 0.31 | 0.09 | 15 | Endothelial |
| NDUFS5 | 0.53 | 0.00 | 0.00 | 0.61 | 0.27 | 15 | Endothelial |
| ARPC3 | 0.60 | 0.00 | 0.00 | 0.63 | 0.29 | 15 | Endothelial |
| MRPL22 | 0.64 | 0.00 | 0.00 | 0.26 | 0.07 | 15 | Endothelial |
| RTN4 | 0.64 | 0.00 | 0.00 | 0.59 | 0.27 | 15 | Endothelial |
| PSMB7 | 0.72 | 0.00 | 0.00 | 0.37 | 0.13 | 15 | Endothelial |
| LAPTM4A | 0.60 | 0.00 | 0.00 | 0.85 | 0.55 | 15 | Endothelial |
| PTTG1IP | 0.57 | 0.00 | 0.00 | 0.47 | 0.18 | 15 | Endothelial |
| ITGB1BP1 | 0.47 | 0.00 | 0.00 | 0.27 | 0.08 | 15 | Endothelial |
| DDX17 | 0.63 | 0.00 | 0.00 | 0.57 | 0.26 | 15 | Endothelial |
| ADCY4 | 0.64 | 0.00 | 0.00 | 0.30 | 0.09 | 15 | Endothelial |
| PNKD | 0.58 | 0.00 | 0.00 | 0.33 | 0.11 | 15 | Endothelial |
| SAP30BP | 0.42 | 0.00 | 0.00 | 0.26 | 0.07 | 15 | Endothelial |
| NDFIP1 | 0.60 | 0.00 | 0.00 | 0.33 | 0.11 | 15 | Endothelial |
| TMEM204 | 0.54 | 0.00 | 0.00 | 0.44 | 0.17 | 15 | Endothelial |
| DECR1 | 0.54 | 0.00 | 0.00 | 0.28 | 0.08 | 15 | Endothelial |
| FNBP1 | 0.50 | 0.00 | 0.00 | 0.31 | 0.09 | 15 | Endothelial |
| AES | 0.76 | 0.00 | 0.00 | 0.41 | 0.16 | 15 | Endothelial |
| TGM2 | 0.58 | 0.00 | 0.00 | 0.33 | 0.11 | 15 | Endothelial |
| UQCRQ | 0.53 | 0.00 | 0.00 | 0.72 | 0.36 | 15 | Endothelial |
| TIMM22 | 0.57 | 0.00 | 0.00 | 0.27 | 0.08 | 15 | Endothelial |
| PTBP1 | 0.55 | 0.00 | 0.00 | 0.28 | 0.09 | 15 | Endothelial |
| S100A13 | 0.74 | 0.00 | 0.00 | 0.57 | 0.27 | 15 | Endothelial |
| GIMAP1 | 0.33 | 0.00 | 0.00 | 0.41 | 0.14 | 15 | Endothelial |
| JUN | 0.67 | 0.00 | 0.00 | 0.70 | 0.38 | 15 | Endothelial |
| RNPS1 | 0.56 | 0.00 | 0.00 | 0.42 | 0.16 | 15 | Endothelial |
| ACTN4 | 0.48 | 0.00 | 0.00 | 0.64 | 0.30 | 15 | Endothelial |
| BRI3 | 0.56 | 0.00 | 0.00 | 0.51 | 0.21 | 15 | Endothelial |
| SDCBP | 0.52 | 0.00 | 0.00 | 0.63 | 0.29 | 15 | Endothelial |
| NDUFA11 | 0.68 | 0.00 | 0.00 | 0.59 | 0.30 | 15 | Endothelial |
| LAMTOR2 | 0.64 | 0.00 | 0.00 | 0.42 | 0.16 | 15 | Endothelial |
| SRSF7 | 0.55 | 0.00 | 0.00 | 0.44 | 0.18 | 15 | Endothelial |
| PSMB8 | 0.43 | 0.00 | 0.00 | 0.48 | 0.19 | 15 | Endothelial |
| DNAJC15 | 0.51 | 0.00 | 0.00 | 0.28 | 0.09 | 15 | Endothelial |
| ACADVL | 0.59 | 0.00 | 0.00 | 0.31 | 0.10 | 15 | Endothelial |
| NOSIP | 0.57 | 0.00 | 0.00 | 0.30 | 0.10 | 15 | Endothelial |
| PLXND1 | 0.54 | 0.00 | 0.00 | 0.27 | 0.08 | 15 | Endothelial |
| ALDOA | 0.54 | 0.00 | 0.00 | 0.52 | 0.23 | 15 | Endothelial |
| EIF1 | 0.46 | 0.00 | 0.00 | 0.94 | 0.69 | 15 | Endothelial |
| PPP2R5A | 0.50 | 0.00 | 0.00 | 0.38 | 0.14 | 15 | Endothelial |
| TMEM109 | 0.63 | 0.00 | 0.00 | 0.36 | 0.14 | 15 | Endothelial |
| XAF1 | 0.52 | 0.00 | 0.00 | 0.27 | 0.08 | 15 | Endothelial |
| SKAP2 | 0.60 | 0.00 | 0.00 | 0.37 | 0.14 | 15 | Endothelial |
| NECAP2 | 0.56 | 0.00 | 0.00 | 0.31 | 0.10 | 15 | Endothelial |
| PHPT1 | 0.53 | 0.00 | 0.00 | 0.36 | 0.13 | 15 | Endothelial |
| SOX17 | 0.30 | 0.00 | 0.00 | 0.40 | 0.14 | 15 | Endothelial |
| COPZ1 | 0.49 | 0.00 | 0.00 | 0.36 | 0.13 | 15 | Endothelial |
| SAP18 | 0.49 | 0.00 | 0.00 | 0.63 | 0.32 | 15 | Endothelial |
| MT-ATP6 | 0.35 | 0.00 | 0.00 | 0.98 | 0.85 | 15 | Endothelial |
| MEIS2 | 0.61 | 0.00 | 0.00 | 0.33 | 0.12 | 15 | Endothelial |
| RNF7 | 0.52 | 0.00 | 0.00 | 0.54 | 0.25 | 15 | Endothelial |
| TMEM165 | 0.56 | 0.00 | 0.00 | 0.33 | 0.12 | 15 | Endothelial |
| CFLAR | 0.55 | 0.00 | 0.00 | 0.46 | 0.20 | 15 | Endothelial |
| SEC11A | 0.46 | 0.00 | 0.00 | 0.48 | 0.21 | 15 | Endothelial |
| BTG1 | 0.53 | 0.00 | 0.00 | 0.61 | 0.31 | 15 | Endothelial |
| FKBP2 | 0.58 | 0.00 | 0.00 | 0.46 | 0.20 | 15 | Endothelial |
| CD151 | 0.54 | 0.00 | 0.00 | 0.46 | 0.20 | 15 | Endothelial |
| PSMB10 | 0.48 | 0.00 | 0.00 | 0.37 | 0.14 | 15 | Endothelial |
| SPTAN1 | 0.37 | 0.00 | 0.00 | 0.43 | 0.17 | 15 | Endothelial |
| AP2S1 | 0.58 | 0.00 | 0.00 | 0.44 | 0.20 | 15 | Endothelial |
| THBD | 0.36 | 0.00 | 0.00 | 0.58 | 0.26 | 15 | Endothelial |
| NCKAP5 | 0.56 | 0.00 | 0.00 | 0.35 | 0.13 | 15 | Endothelial |
| OFD1 | 0.38 | 0.00 | 0.00 | 0.26 | 0.08 | 15 | Endothelial |
| TERF2IP | 0.56 | 0.00 | 0.00 | 0.26 | 0.08 | 15 | Endothelial |
| NDUFB4 | 0.52 | 0.00 | 0.00 | 0.54 | 0.26 | 15 | Endothelial |
| BSG | 0.40 | 0.00 | 0.00 | 0.74 | 0.39 | 15 | Endothelial |
| TJP1 | 0.30 | 0.00 | 0.00 | 0.44 | 0.18 | 15 | Endothelial |
| DAZAP2 | 0.54 | 0.00 | 0.00 | 0.49 | 0.22 | 15 | Endothelial |
| TMBIM6 | 0.44 | 0.00 | 0.00 | 0.74 | 0.41 | 15 | Endothelial |
| HPCAL1 | 0.54 | 0.00 | 0.00 | 0.38 | 0.15 | 15 | Endothelial |
| METTL9 | 0.44 | 0.00 | 0.00 | 0.36 | 0.14 | 15 | Endothelial |
| IL6ST | 0.61 | 0.00 | 0.00 | 0.28 | 0.10 | 15 | Endothelial |
| LSM2 | 0.51 | 0.00 | 0.00 | 0.28 | 0.10 | 15 | Endothelial |
| HSPE1 | 0.56 | 0.00 | 0.00 | 0.59 | 0.30 | 15 | Endothelial |
| PON2 | 0.54 | 0.00 | 0.00 | 0.37 | 0.15 | 15 | Endothelial |
| SCARB2 | 0.48 | 0.00 | 0.00 | 0.26 | 0.09 | 15 | Endothelial |
| RBCK1 | 0.42 | 0.00 | 0.00 | 0.27 | 0.09 | 15 | Endothelial |
| ELK3 | 0.37 | 0.00 | 0.00 | 0.37 | 0.14 | 15 | Endothelial |
| TNFRSF1A | 0.51 | 0.00 | 0.00 | 0.36 | 0.14 | 15 | Endothelial |
| CRIP2 | 0.31 | 0.00 | 0.00 | 0.82 | 0.44 | 15 | Endothelial |
| H2AFZ | 0.48 | 0.00 | 0.00 | 0.69 | 0.41 | 15 | Endothelial |
| POLR2C | 0.46 | 0.00 | 0.00 | 0.36 | 0.14 | 15 | Endothelial |
| FLI1 | 0.45 | 0.00 | 0.00 | 0.32 | 0.12 | 15 | Endothelial |
| VIM | 0.46 | 0.00 | 0.00 | 0.95 | 0.69 | 15 | Endothelial |
| LSM7 | 0.45 | 0.00 | 0.00 | 0.41 | 0.17 | 15 | Endothelial |
| SRSF9 | 0.39 | 0.00 | 0.00 | 0.33 | 0.13 | 15 | Endothelial |
| GLO1 | 0.53 | 0.00 | 0.00 | 0.37 | 0.15 | 15 | Endothelial |
| GSTK1 | 0.57 | 0.00 | 0.00 | 0.27 | 0.10 | 15 | Endothelial |
| UBE2D2 | 0.47 | 0.00 | 0.00 | 0.44 | 0.20 | 15 | Endothelial |
| GADD45GIP1 | 0.54 | 0.00 | 0.00 | 0.41 | 0.18 | 15 | Endothelial |
| COMMD7 | 0.35 | 0.00 | 0.00 | 0.30 | 0.11 | 15 | Endothelial |
| EMP3 | 0.37 | 0.00 | 0.00 | 0.52 | 0.24 | 15 | Endothelial |
| SNRPC | 0.47 | 0.00 | 0.00 | 0.41 | 0.18 | 15 | Endothelial |
| VAPA | 0.50 | 0.00 | 0.00 | 0.49 | 0.24 | 15 | Endothelial |
| PTPRM | 0.50 | 0.00 | 0.00 | 0.27 | 0.10 | 15 | Endothelial |
| RAD23A | 0.53 | 0.00 | 0.00 | 0.35 | 0.14 | 15 | Endothelial |
| MAPK3 | 0.41 | 0.00 | 0.00 | 0.35 | 0.14 | 15 | Endothelial |
| RBPJ | 0.41 | 0.00 | 0.00 | 0.31 | 0.12 | 15 | Endothelial |
| NACA | 0.42 | 0.00 | 0.00 | 0.86 | 0.61 | 15 | Endothelial |
| BANF1 | 0.35 | 0.00 | 0.01 | 0.59 | 0.29 | 15 | Endothelial |
| POLR2K | 0.34 | 0.00 | 0.01 | 0.36 | 0.15 | 15 | Endothelial |
| ITGA6 | 0.31 | 0.00 | 0.01 | 0.33 | 0.13 | 15 | Endothelial |
| ARID1B | 0.45 | 0.00 | 0.01 | 0.26 | 0.09 | 15 | Endothelial |
| RNF13 | 0.42 | 0.00 | 0.01 | 0.28 | 0.11 | 15 | Endothelial |
| ENSA | 0.42 | 0.00 | 0.01 | 0.35 | 0.14 | 15 | Endothelial |
| TMEM50B | 0.37 | 0.00 | 0.01 | 0.26 | 0.09 | 15 | Endothelial |
| JTB | 0.42 | 0.00 | 0.01 | 0.44 | 0.20 | 15 | Endothelial |
| EFEMP2 | 0.47 | 0.00 | 0.01 | 0.26 | 0.09 | 15 | Endothelial |
| PDCD6 | 0.49 | 0.00 | 0.01 | 0.38 | 0.17 | 15 | Endothelial |
| EIF3G | 0.44 | 0.00 | 0.01 | 0.44 | 0.21 | 15 | Endothelial |
| APRT | 0.41 | 0.00 | 0.01 | 0.42 | 0.19 | 15 | Endothelial |
| HSBP1 | 0.42 | 0.00 | 0.01 | 0.61 | 0.33 | 15 | Endothelial |
| NDUFA1 | 0.40 | 0.00 | 0.01 | 0.62 | 0.33 | 15 | Endothelial |
| EID1 | 0.35 | 0.00 | 0.01 | 0.63 | 0.33 | 15 | Endothelial |
| CTDSP1 | 0.46 | 0.00 | 0.01 | 0.26 | 0.10 | 15 | Endothelial |
| COL4A2 | 0.32 | 0.00 | 0.01 | 0.77 | 0.45 | 15 | Endothelial |
| NOP10 | 0.44 | 0.00 | 0.01 | 0.54 | 0.29 | 15 | Endothelial |
| TIMM8B | 0.39 | 0.00 | 0.01 | 0.43 | 0.20 | 15 | Endothelial |
| NUCB1 | 0.43 | 0.00 | 0.01 | 0.37 | 0.16 | 15 | Endothelial |
| ZFP36L1 | 0.36 | 0.00 | 0.01 | 0.75 | 0.47 | 15 | Endothelial |
| PNRC1 | 0.38 | 0.00 | 0.01 | 0.47 | 0.23 | 15 | Endothelial |
| TOMM7 | 0.45 | 0.00 | 0.01 | 0.68 | 0.40 | 15 | Endothelial |
| ECHS1 | 0.40 | 0.00 | 0.01 | 0.37 | 0.16 | 15 | Endothelial |
| SRP9 | 0.49 | 0.00 | 0.01 | 0.56 | 0.31 | 15 | Endothelial |
| NDUFS7 | 0.42 | 0.00 | 0.02 | 0.41 | 0.19 | 15 | Endothelial |
| CIB1 | 0.42 | 0.00 | 0.02 | 0.32 | 0.13 | 15 | Endothelial |
| PGAM1 | 0.50 | 0.00 | 0.02 | 0.32 | 0.14 | 15 | Endothelial |
| ADRM1 | 0.50 | 0.00 | 0.02 | 0.32 | 0.14 | 15 | Endothelial |
| NAA10 | 0.43 | 0.00 | 0.02 | 0.27 | 0.11 | 15 | Endothelial |
| SH3BGRL3 | 0.25 | 0.00 | 0.02 | 0.67 | 0.36 | 15 | Endothelial |
| ZEB1 | 0.32 | 0.00 | 0.02 | 0.30 | 0.12 | 15 | Endothelial |
| SSR2 | 0.42 | 0.00 | 0.03 | 0.61 | 0.34 | 15 | Endothelial |
| MYO6 | 0.51 | 0.00 | 0.03 | 0.32 | 0.14 | 15 | Endothelial |
| SIVA1 | 0.42 | 0.00 | 0.03 | 0.36 | 0.16 | 15 | Endothelial |
| CCDC124 | 0.44 | 0.00 | 0.03 | 0.28 | 0.12 | 15 | Endothelial |
| RAB11B | 0.41 | 0.00 | 0.03 | 0.44 | 0.22 | 15 | Endothelial |
| FKBP8 | 0.47 | 0.00 | 0.04 | 0.35 | 0.16 | 15 | Endothelial |
| SEC61G | 0.35 | 0.00 | 0.04 | 0.67 | 0.38 | 15 | Endothelial |
| POMP | 0.32 | 0.00 | 0.04 | 0.62 | 0.34 | 15 | Endothelial |
| RABAC1 | 0.36 | 0.00 | 0.04 | 0.57 | 0.30 | 15 | Endothelial |
| DARS | 0.45 | 0.00 | 0.04 | 0.26 | 0.10 | 15 | Endothelial |
| PSME2 | 0.47 | 0.00 | 0.05 | 0.48 | 0.26 | 15 | Endothelial |
| POLR2G | 0.45 | 0.00 | 0.05 | 0.33 | 0.15 | 15 | Endothelial |
| IER2 | 0.45 | 0.00 | 0.05 | 0.48 | 0.26 | 15 | Endothelial |
| DCTN6 | 0.38 | 0.00 | 0.05 | 0.30 | 0.12 | 15 | Endothelial |
| GZMB | 3.89 | 0.00 | 0.00 | 0.61 | 0.00 | 16 | NK Cell |
| PRF1 | 3.52 | 0.00 | 0.00 | 0.71 | 0.00 | 16 | NK Cell |
| KLRB1 | 3.34 | 0.00 | 0.00 | 0.84 | 0.01 | 16 | NK Cell |
| XCL2 | 3.16 | 0.00 | 0.00 | 0.47 | 0.00 | 16 | NK Cell |
| CD7 | 3.04 | 0.00 | 0.00 | 0.95 | 0.01 | 16 | NK Cell |
| CTSW | 2.99 | 0.00 | 0.00 | 0.84 | 0.02 | 16 | NK Cell |
| CST7 | 2.86 | 0.00 | 0.00 | 0.74 | 0.01 | 16 | NK Cell |
| ITGB2 | 2.49 | 0.00 | 0.00 | 0.71 | 0.01 | 16 | NK Cell |
| GZMM | 2.45 | 0.00 | 0.00 | 0.76 | 0.01 | 16 | NK Cell |
| TBX21 | 1.73 | 0.00 | 0.00 | 0.42 | 0.00 | 16 | NK Cell |
| SAMD3 | 1.59 | 0.00 | 0.00 | 0.45 | 0.01 | 16 | NK Cell |
| CD247 | 2.53 | 0.00 | 0.00 | 0.76 | 0.02 | 16 | NK Cell |
| HSH2D | 1.35 | 0.00 | 0.00 | 0.26 | 0.00 | 16 | NK Cell |
| GZMA | 2.35 | 0.00 | 0.00 | 0.63 | 0.01 | 16 | NK Cell |
| NKG7 | 3.72 | 0.00 | 0.00 | 0.84 | 0.02 | 16 | NK Cell |
| MATK | 2.01 | 0.00 | 0.00 | 0.55 | 0.01 | 16 | NK Cell |
| KLRD1 | 2.53 | 0.00 | 0.00 | 0.66 | 0.02 | 16 | NK Cell |
| TRGC1 | 1.28 | 0.00 | 0.00 | 0.29 | 0.00 | 16 | NK Cell |
| CD160 | 1.96 | 0.00 | 0.00 | 0.40 | 0.01 | 16 | NK Cell |
| AOAH | 1.66 | 0.00 | 0.00 | 0.37 | 0.01 | 16 | NK Cell |
| CD96 | 1.26 | 0.00 | 0.00 | 0.32 | 0.01 | 16 | NK Cell |
| MYBL1 | 1.25 | 0.00 | 0.00 | 0.32 | 0.01 | 16 | NK Cell |
| SYTL1 | 1.49 | 0.00 | 0.00 | 0.32 | 0.01 | 16 | NK Cell |
| IL2RB | 1.54 | 0.00 | 0.00 | 0.34 | 0.01 | 16 | NK Cell |
| ZAP70 | 1.32 | 0.00 | 0.00 | 0.40 | 0.01 | 16 | NK Cell |
| LCK | 1.89 | 0.00 | 0.00 | 0.66 | 0.03 | 16 | NK Cell |
| AAK1 | 1.91 | 0.00 | 0.00 | 0.61 | 0.02 | 16 | NK Cell |
| TXK | 1.54 | 0.00 | 0.00 | 0.37 | 0.01 | 16 | NK Cell |
| GIMAP7 | 2.02 | 0.00 | 0.00 | 0.63 | 0.03 | 16 | NK Cell |
| CCL4 | 3.56 | 0.00 | 0.00 | 0.42 | 0.01 | 16 | NK Cell |
| TSTD1 | 1.35 | 0.00 | 0.00 | 0.34 | 0.01 | 16 | NK Cell |
| DOK2 | 1.68 | 0.00 | 0.00 | 0.50 | 0.02 | 16 | NK Cell |
| HCST | 2.43 | 0.00 | 0.00 | 0.74 | 0.04 | 16 | NK Cell |
| TRBC1 | 2.15 | 0.00 | 0.00 | 0.71 | 0.04 | 16 | NK Cell |
| ACAP1 | 1.61 | 0.00 | 0.00 | 0.45 | 0.02 | 16 | NK Cell |
| HSPA1A | 2.99 | 0.00 | 0.00 | 0.79 | 0.05 | 16 | NK Cell |
| HAVCR2 | 1.91 | 0.00 | 0.00 | 0.40 | 0.01 | 16 | NK Cell |
| FCGR3A | 2.45 | 0.00 | 0.00 | 0.53 | 0.02 | 16 | NK Cell |
| CCL5 | 2.11 | 0.00 | 0.00 | 0.45 | 0.02 | 16 | NK Cell |
| S1PR5 | 1.36 | 0.00 | 0.00 | 0.29 | 0.01 | 16 | NK Cell |
| SH2D2A | 1.45 | 0.00 | 0.00 | 0.29 | 0.01 | 16 | NK Cell |
| CD3D | 1.96 | 0.00 | 0.00 | 0.55 | 0.03 | 16 | NK Cell |
| CD48 | 1.80 | 0.00 | 0.00 | 0.63 | 0.05 | 16 | NK Cell |
| PTPN4 | 1.33 | 0.00 | 0.00 | 0.37 | 0.02 | 16 | NK Cell |
| RUNX3 | 1.26 | 0.00 | 0.00 | 0.34 | 0.02 | 16 | NK Cell |
| CLEC2D | 1.54 | 0.00 | 0.00 | 0.45 | 0.03 | 16 | NK Cell |
| BIN2 | 1.80 | 0.00 | 0.00 | 0.58 | 0.05 | 16 | NK Cell |
| ADA | 1.08 | 0.00 | 0.00 | 0.29 | 0.01 | 16 | NK Cell |
| SH2D1A | 0.91 | 0.00 | 0.00 | 0.26 | 0.01 | 16 | NK Cell |
| CORO1A | 2.10 | 0.00 | 0.00 | 0.79 | 0.10 | 16 | NK Cell |
| IFITM2 | 1.87 | 0.00 | 0.00 | 0.74 | 0.08 | 16 | NK Cell |
| TBC1D10C | 1.37 | 0.00 | 0.00 | 0.45 | 0.03 | 16 | NK Cell |
| CD3E | 1.27 | 0.00 | 0.00 | 0.32 | 0.02 | 16 | NK Cell |
| UBL5 | 1.73 | 0.00 | 0.00 | 0.63 | 0.07 | 16 | NK Cell |
| RHOF | 1.14 | 0.00 | 0.00 | 0.29 | 0.02 | 16 | NK Cell |
| AKNA | 1.50 | 0.00 | 0.00 | 0.42 | 0.03 | 16 | NK Cell |
| GADD45B | 1.93 | 0.00 | 0.00 | 0.74 | 0.10 | 16 | NK Cell |
| MYO1F | 1.58 | 0.00 | 0.00 | 0.42 | 0.03 | 16 | NK Cell |
| IL16 | 1.41 | 0.00 | 0.00 | 0.40 | 0.03 | 16 | NK Cell |
| RAC2 | 1.64 | 0.00 | 0.00 | 0.76 | 0.11 | 16 | NK Cell |
| EVL | 1.81 | 0.00 | 0.00 | 0.68 | 0.09 | 16 | NK Cell |
| TTC38 | 1.35 | 0.00 | 0.00 | 0.34 | 0.02 | 16 | NK Cell |
| DUSP2 | 1.27 | 0.00 | 0.00 | 0.32 | 0.02 | 16 | NK Cell |
| GAPDH | 2.20 | 0.00 | 0.00 | 0.92 | 0.20 | 16 | NK Cell |
| RHOH | 1.29 | 0.00 | 0.00 | 0.29 | 0.02 | 16 | NK Cell |
| BID | 0.98 | 0.00 | 0.00 | 0.34 | 0.02 | 16 | NK Cell |
| CD3G | 1.23 | 0.00 | 0.00 | 0.42 | 0.03 | 16 | NK Cell |
| PSMB9 | 1.58 | 0.00 | 0.00 | 0.68 | 0.10 | 16 | NK Cell |
| ARHGDIB | 1.58 | 0.00 | 0.00 | 0.90 | 0.17 | 16 | NK Cell |
| RPL29 | 2.04 | 0.00 | 0.00 | 0.90 | 0.20 | 16 | NK Cell |
| FMNL1 | 1.30 | 0.00 | 0.00 | 0.37 | 0.03 | 16 | NK Cell |
| RNF213 | 1.42 | 0.00 | 0.00 | 0.47 | 0.05 | 16 | NK Cell |
| C1orf56 | 1.39 | 0.00 | 0.00 | 0.26 | 0.02 | 16 | NK Cell |
| ZFP36L2 | 1.84 | 0.00 | 0.00 | 0.76 | 0.14 | 16 | NK Cell |
| RASAL3 | 0.98 | 0.00 | 0.00 | 0.26 | 0.02 | 16 | NK Cell |
| TGFB1 | 1.32 | 0.00 | 0.00 | 0.42 | 0.04 | 16 | NK Cell |
| EIF2S3 | 1.11 | 0.00 | 0.00 | 0.32 | 0.03 | 16 | NK Cell |
| GATA3 | 1.15 | 0.00 | 0.00 | 0.32 | 0.03 | 16 | NK Cell |
| IL2RG | 1.49 | 0.00 | 0.00 | 0.63 | 0.09 | 16 | NK Cell |
| SPON2 | 1.94 | 0.00 | 0.00 | 0.55 | 0.08 | 16 | NK Cell |
| FCER1G | 1.22 | 0.00 | 0.00 | 0.74 | 0.12 | 16 | NK Cell |
| ARHGAP4 | 1.08 | 0.00 | 0.00 | 0.32 | 0.03 | 16 | NK Cell |
| DNAJB1 | 1.62 | 0.00 | 0.00 | 0.50 | 0.07 | 16 | NK Cell |
| PTPRC | 1.22 | 0.00 | 0.00 | 0.79 | 0.14 | 16 | NK Cell |
| CD27 | 1.45 | 0.00 | 0.00 | 0.29 | 0.02 | 16 | NK Cell |
| UXT | 1.26 | 0.00 | 0.00 | 0.50 | 0.07 | 16 | NK Cell |
| C19orf66 | 1.47 | 0.00 | 0.00 | 0.42 | 0.05 | 16 | NK Cell |
| DEF6 | 1.15 | 0.00 | 0.00 | 0.32 | 0.03 | 16 | NK Cell |
| CD74 | 1.04 | 0.00 | 0.00 | 0.79 | 0.15 | 16 | NK Cell |
| JUNB | 1.67 | 0.00 | 0.00 | 0.84 | 0.21 | 16 | NK Cell |
| FOSB | 1.37 | 0.00 | 0.00 | 0.34 | 0.03 | 16 | NK Cell |
| CD53 | 1.51 | 0.00 | 0.00 | 0.58 | 0.09 | 16 | NK Cell |
| B2M | 2.48 | 0.00 | 0.00 | 1.00 | 0.70 | 16 | NK Cell |
| IRF1 | 1.60 | 0.00 | 0.00 | 0.55 | 0.09 | 16 | NK Cell |
| EVI2A | 1.05 | 0.00 | 0.00 | 0.34 | 0.03 | 16 | NK Cell |
| TYROBP | 1.05 | 0.00 | 0.00 | 0.71 | 0.13 | 16 | NK Cell |
| PPIA | 1.65 | 0.00 | 0.00 | 0.82 | 0.22 | 16 | NK Cell |
| LEF1 | 1.09 | 0.00 | 0.00 | 0.37 | 0.04 | 16 | NK Cell |
| FYN | 1.26 | 0.00 | 0.00 | 0.58 | 0.10 | 16 | NK Cell |
| ARL4C | 1.38 | 0.00 | 0.00 | 0.55 | 0.09 | 16 | NK Cell |
| HEXIM1 | 1.60 | 0.00 | 0.00 | 0.61 | 0.12 | 16 | NK Cell |
| GPSM3 | 1.18 | 0.00 | 0.00 | 0.45 | 0.06 | 16 | NK Cell |
| LIMD2 | 1.44 | 0.00 | 0.00 | 0.71 | 0.15 | 16 | NK Cell |
| RPL13A | 1.90 | 0.00 | 0.00 | 1.00 | 0.59 | 16 | NK Cell |
| PCSK7 | 1.22 | 0.00 | 0.00 | 0.37 | 0.04 | 16 | NK Cell |
| PTPN7 | 0.99 | 0.00 | 0.00 | 0.29 | 0.03 | 16 | NK Cell |
| PPP1R18 | 1.17 | 0.00 | 0.00 | 0.47 | 0.07 | 16 | NK Cell |
| TAGAP | 1.24 | 0.00 | 0.00 | 0.34 | 0.04 | 16 | NK Cell |
| RASSF5 | 1.05 | 0.00 | 0.00 | 0.34 | 0.04 | 16 | NK Cell |
| TUFM | 1.20 | 0.00 | 0.00 | 0.50 | 0.08 | 16 | NK Cell |
| CYBA | 1.53 | 0.00 | 0.00 | 0.92 | 0.32 | 16 | NK Cell |
| WIPF1 | 1.16 | 0.00 | 0.00 | 0.55 | 0.10 | 16 | NK Cell |
| XRCC5 | 1.27 | 0.00 | 0.00 | 0.42 | 0.06 | 16 | NK Cell |
| IER2 | 1.88 | 0.00 | 0.00 | 0.82 | 0.26 | 16 | NK Cell |
| SIAH2 | 1.01 | 0.00 | 0.00 | 0.26 | 0.02 | 16 | NK Cell |
| TRAF3IP3 | 1.20 | 0.00 | 0.00 | 0.32 | 0.03 | 16 | NK Cell |
| VAMP2 | 1.39 | 0.00 | 0.00 | 0.53 | 0.10 | 16 | NK Cell |
| GYPC | 1.27 | 0.00 | 0.00 | 0.45 | 0.07 | 16 | NK Cell |
| TMSB4X | 1.40 | 0.00 | 0.00 | 1.00 | 0.90 | 16 | NK Cell |
| CHST12 | 1.39 | 0.00 | 0.00 | 0.37 | 0.05 | 16 | NK Cell |
| CMC1 | 1.93 | 0.00 | 0.00 | 0.53 | 0.10 | 16 | NK Cell |
| 1-Sep | 1.08 | 0.00 | 0.00 | 0.34 | 0.04 | 16 | NK Cell |
| PLAC8 | 0.82 | 0.00 | 0.00 | 0.74 | 0.17 | 16 | NK Cell |
| APMAP | 1.29 | 0.00 | 0.00 | 0.40 | 0.06 | 16 | NK Cell |
| LAPTM5 | 0.98 | 0.00 | 0.00 | 0.68 | 0.14 | 16 | NK Cell |
| ARPC3 | 1.38 | 0.00 | 0.00 | 0.84 | 0.29 | 16 | NK Cell |
| PRKCH | 1.21 | 0.00 | 0.00 | 0.37 | 0.05 | 16 | NK Cell |
| RNF126 | 1.08 | 0.00 | 0.00 | 0.32 | 0.04 | 16 | NK Cell |
| EEF1D | 1.39 | 0.00 | 0.00 | 0.87 | 0.30 | 16 | NK Cell |
| PYURF | 1.13 | 0.00 | 0.00 | 0.32 | 0.04 | 16 | NK Cell |
| DPP7 | 1.05 | 0.00 | 0.00 | 0.26 | 0.03 | 16 | NK Cell |
| UCP2 | 1.49 | 0.00 | 0.00 | 0.55 | 0.12 | 16 | NK Cell |
| SH3BP1 | 0.84 | 0.00 | 0.00 | 0.26 | 0.03 | 16 | NK Cell |
| BTG1 | 1.52 | 0.00 | 0.00 | 0.84 | 0.31 | 16 | NK Cell |
| STK17B | 0.95 | 0.00 | 0.00 | 0.40 | 0.06 | 16 | NK Cell |
| CD37 | 1.06 | 0.00 | 0.00 | 0.45 | 0.07 | 16 | NK Cell |
| MT-CO2 | 1.08 | 0.00 | 0.00 | 1.00 | 0.79 | 16 | NK Cell |
| JUN | 1.61 | 0.00 | 0.00 | 0.90 | 0.38 | 16 | NK Cell |
| LDHB | 1.41 | 0.00 | 0.00 | 0.53 | 0.12 | 16 | NK Cell |
| PFN1 | 1.61 | 0.00 | 0.00 | 0.90 | 0.49 | 16 | NK Cell |
| GBP2 | 1.15 | 0.00 | 0.00 | 0.37 | 0.06 | 16 | NK Cell |
| 9-Sep | 1.24 | 0.00 | 0.00 | 0.50 | 0.10 | 16 | NK Cell |
| HNRNPA1 | 1.36 | 0.00 | 0.00 | 0.82 | 0.31 | 16 | NK Cell |
| PSMF1 | 1.00 | 0.00 | 0.00 | 0.34 | 0.05 | 16 | NK Cell |
| S100A4 | 1.31 | 0.00 | 0.00 | 0.50 | 0.10 | 16 | NK Cell |
| EMP3 | 1.36 | 0.00 | 0.00 | 0.74 | 0.24 | 16 | NK Cell |
| PRMT2 | 1.37 | 0.00 | 0.00 | 0.37 | 0.06 | 16 | NK Cell |
| HOPX | 1.37 | 0.00 | 0.00 | 0.50 | 0.11 | 16 | NK Cell |
| RPL7A | 1.20 | 0.00 | 0.00 | 0.92 | 0.44 | 16 | NK Cell |
| SLC9A3R1 | 0.91 | 0.00 | 0.00 | 0.29 | 0.04 | 16 | NK Cell |
| C12orf75 | 0.87 | 0.00 | 0.00 | 0.29 | 0.04 | 16 | NK Cell |
| RPL23A | 1.25 | 0.00 | 0.00 | 0.97 | 0.69 | 16 | NK Cell |
| GTF3A | 1.11 | 0.00 | 0.00 | 0.45 | 0.09 | 16 | NK Cell |
| SIGIRR | 1.16 | 0.00 | 0.00 | 0.29 | 0.04 | 16 | NK Cell |
| RPS27A | 1.01 | 0.00 | 0.00 | 1.00 | 0.78 | 16 | NK Cell |
| DENND2D | 1.11 | 0.00 | 0.00 | 0.29 | 0.04 | 16 | NK Cell |
| RPLP2 | 1.03 | 0.00 | 0.00 | 1.00 | 0.83 | 16 | NK Cell |
| LSP1 | 0.85 | 0.00 | 0.00 | 0.71 | 0.19 | 16 | NK Cell |
| CDKN2D | 0.94 | 0.00 | 0.00 | 0.34 | 0.06 | 16 | NK Cell |
| LYST | 1.14 | 0.00 | 0.00 | 0.29 | 0.04 | 16 | NK Cell |
| NFKB2 | 1.05 | 0.00 | 0.00 | 0.26 | 0.04 | 16 | NK Cell |
| COMMD6 | 1.09 | 0.00 | 0.00 | 0.61 | 0.17 | 16 | NK Cell |
| SRGN | 0.96 | 0.00 | 0.00 | 0.90 | 0.32 | 16 | NK Cell |
| ARHGEF1 | 1.13 | 0.00 | 0.00 | 0.50 | 0.12 | 16 | NK Cell |
| EIF1 | 1.05 | 0.00 | 0.00 | 1.00 | 0.69 | 16 | NK Cell |
| TTC19 | 0.84 | 0.00 | 0.00 | 0.26 | 0.04 | 16 | NK Cell |
| ARHGAP9 | 1.01 | 0.00 | 0.00 | 0.26 | 0.04 | 16 | NK Cell |
| TMC6 | 0.81 | 0.00 | 0.00 | 0.26 | 0.04 | 16 | NK Cell |
| PPP1R15A | 1.17 | 0.00 | 0.00 | 0.40 | 0.08 | 16 | NK Cell |
| RBCK1 | 1.03 | 0.00 | 0.00 | 0.42 | 0.09 | 16 | NK Cell |
| MIF | 1.12 | 0.00 | 0.00 | 0.68 | 0.23 | 16 | NK Cell |
| PGAM1 | 1.07 | 0.00 | 0.00 | 0.53 | 0.14 | 16 | NK Cell |
| PPDPF | 1.10 | 0.00 | 0.00 | 0.68 | 0.23 | 16 | NK Cell |
| TCF7 | 0.91 | 0.00 | 0.00 | 0.34 | 0.06 | 16 | NK Cell |
| SRP14 | 0.87 | 0.00 | 0.00 | 0.84 | 0.34 | 16 | NK Cell |
| PSME1 | 1.12 | 0.00 | 0.00 | 0.79 | 0.34 | 16 | NK Cell |
| MT-ND4 | 0.78 | 0.00 | 0.00 | 1.00 | 0.77 | 16 | NK Cell |
| INTS6 | 0.95 | 0.00 | 0.00 | 0.26 | 0.04 | 16 | NK Cell |
| PSMB10 | 1.02 | 0.00 | 0.00 | 0.53 | 0.14 | 16 | NK Cell |
| MDM4 | 0.85 | 0.00 | 0.00 | 0.29 | 0.05 | 16 | NK Cell |
| SH3BGRL3 | 1.00 | 0.00 | 0.00 | 0.82 | 0.36 | 16 | NK Cell |
| CCNI | 0.99 | 0.00 | 0.00 | 0.63 | 0.20 | 16 | NK Cell |
| WDR74 | 0.83 | 0.00 | 0.00 | 0.29 | 0.05 | 16 | NK Cell |
| FAM49B | 1.19 | 0.00 | 0.00 | 0.45 | 0.11 | 16 | NK Cell |
| CYTIP | 0.97 | 0.00 | 0.00 | 0.42 | 0.09 | 16 | NK Cell |
| CIB1 | 1.00 | 0.00 | 0.00 | 0.50 | 0.13 | 16 | NK Cell |
| GMFG | 0.94 | 0.00 | 0.00 | 0.55 | 0.15 | 16 | NK Cell |
| CLIC3 | 1.13 | 0.00 | 0.00 | 0.40 | 0.08 | 16 | NK Cell |
| CLIC1 | 1.10 | 0.00 | 0.00 | 0.84 | 0.39 | 16 | NK Cell |
| NOC2L | 0.87 | 0.00 | 0.00 | 0.29 | 0.05 | 16 | NK Cell |
| HCLS1 | 0.88 | 0.00 | 0.00 | 0.37 | 0.07 | 16 | NK Cell |
| CIRBP | 1.01 | 0.00 | 0.00 | 0.71 | 0.26 | 16 | NK Cell |
| LITAF | 1.06 | 0.00 | 0.00 | 0.47 | 0.13 | 16 | NK Cell |
| ARL6IP5 | 1.07 | 0.00 | 0.00 | 0.50 | 0.14 | 16 | NK Cell |
| 6-Sep | 0.87 | 0.00 | 0.00 | 0.37 | 0.08 | 16 | NK Cell |
| TERF2IP | 1.00 | 0.00 | 0.00 | 0.37 | 0.08 | 16 | NK Cell |
| PSMB8 | 0.92 | 0.00 | 0.00 | 0.61 | 0.19 | 16 | NK Cell |
| AES | 1.08 | 0.00 | 0.00 | 0.53 | 0.16 | 16 | NK Cell |
| MT-ND3 | 0.93 | 0.00 | 0.00 | 0.90 | 0.51 | 16 | NK Cell |
| NBEAL1 | 1.16 | 0.00 | 0.00 | 0.47 | 0.13 | 16 | NK Cell |
| RPL7 | 0.84 | 0.00 | 0.00 | 0.97 | 0.77 | 16 | NK Cell |
| TMA7 | 1.00 | 0.00 | 0.00 | 0.76 | 0.35 | 16 | NK Cell |
| GLIPR2 | 0.86 | 0.00 | 0.00 | 0.34 | 0.08 | 16 | NK Cell |
| RDH14 | 0.82 | 0.00 | 0.00 | 0.26 | 0.05 | 16 | NK Cell |
| NDUFB2 | 0.86 | 0.00 | 0.00 | 0.66 | 0.24 | 16 | NK Cell |
| SRSF7 | 1.19 | 0.00 | 0.00 | 0.53 | 0.18 | 16 | NK Cell |
| STK4 | 1.01 | 0.00 | 0.00 | 0.47 | 0.14 | 16 | NK Cell |
| RPL8 | 0.64 | 0.00 | 0.00 | 1.00 | 0.81 | 16 | NK Cell |
| C9orf16 | 0.96 | 0.00 | 0.00 | 0.34 | 0.08 | 16 | NK Cell |
| ALDOA | 0.83 | 0.00 | 0.00 | 0.63 | 0.23 | 16 | NK Cell |
| TXNIP | 1.07 | 0.00 | 0.00 | 0.68 | 0.27 | 16 | NK Cell |
| FDPS | 1.13 | 0.00 | 0.00 | 0.34 | 0.08 | 16 | NK Cell |
| MT-CO1 | 0.58 | 0.00 | 0.00 | 1.00 | 0.95 | 16 | NK Cell |
| CYTH1 | 0.91 | 0.00 | 0.00 | 0.32 | 0.07 | 16 | NK Cell |
| PLEK | 0.92 | 0.00 | 0.00 | 0.32 | 0.07 | 16 | NK Cell |
| CCDC107 | 0.83 | 0.00 | 0.00 | 0.40 | 0.11 | 16 | NK Cell |
| RHOG | 0.95 | 0.00 | 0.00 | 0.42 | 0.12 | 16 | NK Cell |
| PGK1 | 1.00 | 0.00 | 0.00 | 0.45 | 0.14 | 16 | NK Cell |
| FAM107B | 0.75 | 0.00 | 0.00 | 0.32 | 0.07 | 16 | NK Cell |
| FNBP1 | 0.92 | 0.00 | 0.00 | 0.37 | 0.09 | 16 | NK Cell |
| TNIP1 | 0.80 | 0.00 | 0.00 | 0.26 | 0.05 | 16 | NK Cell |
| DUSP1 | 1.12 | 0.00 | 0.00 | 0.53 | 0.17 | 16 | NK Cell |
| APRT | 0.81 | 0.00 | 0.00 | 0.55 | 0.19 | 16 | NK Cell |
| SERF2 | 0.79 | 0.00 | 0.00 | 0.95 | 0.60 | 16 | NK Cell |
| PSMG2 | 0.76 | 0.00 | 0.00 | 0.32 | 0.07 | 16 | NK Cell |
| MFSD10 | 0.79 | 0.00 | 0.00 | 0.29 | 0.06 | 16 | NK Cell |
| CFL1 | 0.72 | 0.00 | 0.00 | 0.92 | 0.61 | 16 | NK Cell |
| GSTK1 | 0.89 | 0.00 | 0.00 | 0.37 | 0.10 | 16 | NK Cell |
| PSME2 | 0.93 | 0.00 | 0.00 | 0.63 | 0.26 | 16 | NK Cell |
| CAND1 | 0.91 | 0.00 | 0.00 | 0.29 | 0.06 | 16 | NK Cell |
| RPS13 | 0.99 | 0.00 | 0.00 | 0.95 | 0.67 | 16 | NK Cell |
| SSBP4 | 0.67 | 0.00 | 0.00 | 0.29 | 0.06 | 16 | NK Cell |
| MYL6 | 0.61 | 0.00 | 0.00 | 0.97 | 0.73 | 16 | NK Cell |
| SSR4 | 0.82 | 0.00 | 0.00 | 0.66 | 0.27 | 16 | NK Cell |
| RPLP1 | 0.72 | 0.00 | 0.00 | 1.00 | 0.91 | 16 | NK Cell |
| RBMX | 0.85 | 0.00 | 0.00 | 0.45 | 0.14 | 16 | NK Cell |
| GNG2 | 0.81 | 0.00 | 0.00 | 0.42 | 0.12 | 16 | NK Cell |
| HMGN3 | 0.92 | 0.00 | 0.00 | 0.32 | 0.08 | 16 | NK Cell |
| CUTA | 0.85 | 0.00 | 0.00 | 0.55 | 0.21 | 16 | NK Cell |
| RPL24 | 0.87 | 0.00 | 0.00 | 0.87 | 0.50 | 16 | NK Cell |
| UBE2L6 | 0.87 | 0.00 | 0.00 | 0.26 | 0.06 | 16 | NK Cell |
| SORL1 | 0.91 | 0.00 | 0.00 | 0.29 | 0.07 | 16 | NK Cell |
| TWF2 | 0.67 | 0.00 | 0.00 | 0.37 | 0.10 | 16 | NK Cell |
| TMEM230 | 0.87 | 0.00 | 0.00 | 0.32 | 0.08 | 16 | NK Cell |
| CKLF | 0.75 | 0.00 | 0.00 | 0.26 | 0.06 | 16 | NK Cell |
| TRMT112 | 0.71 | 0.00 | 0.00 | 0.53 | 0.18 | 16 | NK Cell |
| RPL18A | 0.80 | 0.00 | 0.00 | 0.97 | 0.84 | 16 | NK Cell |
| CDC42SE2 | 0.84 | 0.00 | 0.00 | 0.42 | 0.13 | 16 | NK Cell |
| CD55 | 0.78 | 0.00 | 0.00 | 0.29 | 0.07 | 16 | NK Cell |
| EPC1 | 0.87 | 0.00 | 0.00 | 0.37 | 0.11 | 16 | NK Cell |
| MT-CO3 | 0.51 | 0.00 | 0.00 | 1.00 | 0.89 | 16 | NK Cell |
| RCSD1 | 0.75 | 0.00 | 0.00 | 0.34 | 0.09 | 16 | NK Cell |
| POLR2L | 0.90 | 0.00 | 0.00 | 0.53 | 0.20 | 16 | NK Cell |
| OST4 | 0.84 | 0.00 | 0.00 | 0.66 | 0.29 | 16 | NK Cell |
| HSPE1 | 0.85 | 0.00 | 0.00 | 0.66 | 0.30 | 16 | NK Cell |
| SLC38A2 | 0.96 | 0.00 | 0.00 | 0.50 | 0.19 | 16 | NK Cell |
| UBC | 1.05 | 0.00 | 0.00 | 0.82 | 0.51 | 16 | NK Cell |
| EIF3L | 0.77 | 0.00 | 0.00 | 0.47 | 0.17 | 16 | NK Cell |
| LCP1 | 0.47 | 0.00 | 0.00 | 0.61 | 0.21 | 16 | NK Cell |
| PSMB3 | 0.77 | 0.00 | 0.00 | 0.63 | 0.27 | 16 | NK Cell |
| TANK | 0.77 | 0.00 | 0.00 | 0.26 | 0.06 | 16 | NK Cell |
| ARHGAP30 | 0.52 | 0.00 | 0.00 | 0.37 | 0.10 | 16 | NK Cell |
| TAPBP | 0.84 | 0.00 | 0.00 | 0.45 | 0.16 | 16 | NK Cell |
| PFDN5 | 0.60 | 0.00 | 0.00 | 0.90 | 0.53 | 16 | NK Cell |
| ENO1 | 0.92 | 0.00 | 0.00 | 0.50 | 0.19 | 16 | NK Cell |
| NOSIP | 0.71 | 0.00 | 0.00 | 0.34 | 0.10 | 16 | NK Cell |
| EIF3G | 0.90 | 0.00 | 0.00 | 0.53 | 0.21 | 16 | NK Cell |
| PTPN6 | 0.73 | 0.00 | 0.00 | 0.32 | 0.08 | 16 | NK Cell |
| CD44 | 0.61 | 0.00 | 0.01 | 0.55 | 0.21 | 16 | NK Cell |
| PIM1 | 0.68 | 0.00 | 0.01 | 0.32 | 0.09 | 16 | NK Cell |
| TIMP1 | 0.38 | 0.00 | 0.01 | 0.32 | 0.09 | 16 | NK Cell |
| EFHD2 | 0.70 | 0.00 | 0.01 | 0.32 | 0.09 | 16 | NK Cell |
| EIF4A2 | 0.84 | 0.00 | 0.01 | 0.68 | 0.35 | 16 | NK Cell |
| FERMT3 | 0.53 | 0.00 | 0.01 | 0.26 | 0.06 | 16 | NK Cell |
| PIP4K2A | 0.71 | 0.00 | 0.01 | 0.26 | 0.07 | 16 | NK Cell |
| NACA | 0.64 | 0.00 | 0.01 | 0.87 | 0.61 | 16 | NK Cell |
| EXOSC8 | 0.86 | 0.00 | 0.01 | 0.29 | 0.08 | 16 | NK Cell |
| ARPC1B | 0.63 | 0.00 | 0.01 | 0.71 | 0.37 | 16 | NK Cell |
| RPS4X | 0.61 | 0.00 | 0.02 | 1.00 | 0.88 | 16 | NK Cell |
| FBL | 0.93 | 0.00 | 0.02 | 0.32 | 0.10 | 16 | NK Cell |
| BAZ1A | 0.82 | 0.00 | 0.02 | 0.40 | 0.13 | 16 | NK Cell |
| XRCC6 | 0.61 | 0.00 | 0.02 | 0.26 | 0.07 | 16 | NK Cell |
| LDHA | 0.84 | 0.00 | 0.02 | 0.63 | 0.32 | 16 | NK Cell |
| RBM8A | 0.75 | 0.00 | 0.02 | 0.63 | 0.30 | 16 | NK Cell |
| ADAR | 0.74 | 0.00 | 0.02 | 0.26 | 0.07 | 16 | NK Cell |
| HNRNPA0 | 0.72 | 0.00 | 0.02 | 0.55 | 0.25 | 16 | NK Cell |
| COPE | 0.70 | 0.00 | 0.04 | 0.55 | 0.25 | 16 | NK Cell |
| ABHD17A | 0.64 | 0.00 | 0.04 | 0.26 | 0.07 | 16 | NK Cell |
| KARS | 0.84 | 0.00 | 0.04 | 0.29 | 0.09 | 16 | NK Cell |
| BTF3 | 0.72 | 0.00 | 0.04 | 0.79 | 0.55 | 16 | NK Cell |
| RNF166 | 0.63 | 0.00 | 0.05 | 0.26 | 0.07 | 16 | NK Cell |
| AIP | 0.70 | 0.00 | 0.05 | 0.26 | 0.07 | 16 | NK Cell |

Table S9: Genes differentially expressed in Young and Old Matrix Fibroblasts

| **Gene** | **Log2 FoldChange** | **Unadjusted p-value** | **Adjusted p-value** | **High in** |
| --- | --- | --- | --- | --- |
| IGFBP7 | -2.32944 | 1.76E-16 | 6.37E-14 | Early |
| HES1 | -2.0363 | 5.05E-14 | 9.16E-12 | Early |
| RGS3 | -1.98867 | 5.12E-12 | 5.31E-10 | Early |
| TAGLN | -1.85646 | 6.78E-12 | 6.15E-10 | Early |
| C11orf96 | -1.75965 | 3.53E-08 | 9.43E-07 | Early |
| EGFL6 | -1.66064 | 2.19E-08 | 6.37E-07 | Early |
| TGM2 | -1.61017 | 1.47E-13 | 2.13E-11 | Early |
| TPPP3 | -1.53759 | 8.42E-06 | 0.00013 | Early |
| C1R | -1.41221 | 2.44E-11 | 1.77E-09 | Early |
| GPX3 | -1.39671 | 1.69E-10 | 1.12E-08 | Early |
| PDGFRB | -1.38072 | 8.95E-10 | 4.06E-08 | Early |
| BGN | -1.32151 | 3.73E-10 | 2.08E-08 | Early |
| ELN | -1.27578 | 3.63E-15 | 8.78E-13 | Early |
| LITAF | -1.24824 | 1.86E-12 | 2.25E-10 | Early |
| TIMP1 | -1.23053 | 2.72E-10 | 1.65E-08 | Early |
| LMNA | -1.22694 | 1.89E-09 | 7.99E-08 | Early |
| RARRES1 | -1.21227 | 6.42E-06 | 0.000104 | Early |
| CRIP2 | -1.15468 | 1.90E-07 | 3.82E-06 | Early |
| CNN3 | -1.14133 | 2.88E-07 | 5.65E-06 | Early |
| PMP22 | -1.13909 | 8.31E-12 | 6.70E-10 | Early |
| ZFP36 | -1.12008 | 0.003101 | 0.018012 | Early |
| IGFBP4 | -1.11364 | 0.000134 | 0.001453 | Early |
| CD59 | -1.1021 | 1.98E-09 | 7.99E-08 | Early |
| TPM2 | -1.07868 | 4.54E-10 | 2.20E-08 | Early |
| PLAC9 | -1.06543 | 6.56E-07 | 1.19E-05 | Early |
| TPM4 | -1.04635 | 1.19E-08 | 3.77E-07 | Early |
| ZYX | -1.01911 | 8.15E-09 | 2.96E-07 | Early |
| CCDC80 | -1.01612 | 0.000365 | 0.003195 | Early |
| TPM1 | -0.98081 | 1.24E-07 | 2.73E-06 | Early |
| PDLIM3 | -0.97266 | 0.000146 | 0.001518 | Early |
| TUBA1A | -0.94876 | 1.43E-08 | 4.31E-07 | Early |
| COL4A1 | -0.94321 | 3.05E-05 | 0.000381 | Early |
| CTGF | -0.94149 | 0.020365 | 0.075432 | Early |
| YBX3 | -0.93912 | 9.98E-05 | 0.00115 | Early |
| ENG | -0.89199 | 0.000332 | 0.002942 | Early |
| ZFP36L2 | -0.89135 | 1.55E-07 | 3.21E-06 | Early |
| MYL9 | -0.88525 | 3.64E-08 | 9.43E-07 | Early |
| TUBA1B | -0.85148 | 5.25E-08 | 1.27E-06 | Early |
| KDELR2 | -0.84422 | 1.03E-05 | 0.000152 | Early |
| TXNIP | -0.84004 | 1.62E-05 | 0.000222 | Early |
| RHOC | -0.81135 | 3.11E-05 | 0.000383 | Early |
| ANXA2 | -0.79852 | 6.59E-06 | 0.000104 | Early |
| HNRNPM | -0.79017 | 2.39E-05 | 0.00031 | Early |
| SERPING1 | -0.78543 | 6.55E-07 | 1.19E-05 | Early |
| ZFP36L1 | -0.78119 | 1.51E-07 | 3.21E-06 | Early |
| TSC22D1 | -0.77537 | 1.30E-05 | 0.000185 | Early |
| APOE | -0.75428 | 2.49E-05 | 0.000317 | Early |
| FMO2 | -0.73943 | 0.003523 | 0.019377 | Early |
| IFITM2 | -0.73789 | 1.35E-05 | 0.000188 | Early |
| NDRG2 | -0.71935 | 0.000546 | 0.004608 | Early |
| ITGB1 | -0.71754 | 5.34E-05 | 0.000636 | Early |
| CEBPB | -0.69796 | 0.001135 | 0.008494 | Early |
| COL4A2 | -0.6912 | 0.001327 | 0.009404 | Early |
| GPX4 | -0.68504 | 1.01E-05 | 0.000152 | Early |
| ARF1 | -0.67965 | 1.09E-05 | 0.000158 | Early |
| ARID5B | -0.67637 | 0.001334 | 0.009404 | Early |
| CD63 | -0.67523 | 8.88E-09 | 3.07E-07 | Early |
| CALM2 | -0.67503 | 4.53E-10 | 2.20E-08 | Early |
| TAGLN2 | -0.66315 | 1.16E-07 | 2.64E-06 | Early |
| ARF4 | -0.65749 | 0.000211 | 0.002015 | Early |
| INMT | -0.65208 | 0.00084 | 0.006435 | Early |
| TUBB4B | -0.6494 | 0.003954 | 0.021109 | Early |
| EDNRA | -0.647 | 0.002747 | 0.016744 | Early |
| ACTA2 | -0.63586 | 0.003701 | 0.0202 | Early |
| TNC | -0.63571 | 0.014358 | 0.057591 | Early |
| DPYSL2 | -0.63193 | 0.005437 | 0.026672 | Early |
| SPARC | -0.62971 | 4.46E-08 | 1.12E-06 | Early |
| CALU | -0.62885 | 0.001723 | 0.011249 | Early |
| PLS3 | -0.62687 | 0.001882 | 0.011983 | Early |
| GABARAP | -0.62009 | 0.001348 | 0.009413 | Early |
| MRFAP1 | -0.61928 | 1.65E-05 | 0.000222 | Early |
| VIM | -0.61924 | 9.89E-09 | 3.26E-07 | Early |
| OAZ2 | -0.61866 | 0.001939 | 0.012242 | Early |
| SRSF11 | -0.61306 | 0.000325 | 0.002915 | Early |
| LGALS3 | -0.60514 | 0.003318 | 0.01882 | Early |
| RNH1 | -0.60094 | 0.001303 | 0.009404 | Early |
| FHL1 | -0.59926 | 3.44E-06 | 5.94E-05 | Early |
| TNFRSF1A | -0.5988 | 0.003059 | 0.017929 | Early |
| DYNLL1 | -0.59679 | 0.000124 | 0.001366 | Early |
| COL1A1 | -0.59292 | 5.09E-07 | 9.72E-06 | Early |
| DNAJB6 | -0.58755 | 0.003766 | 0.020403 | Early |
| MYH10 | -0.58524 | 0.003317 | 0.01882 | Early |
| BZW1 | -0.58436 | 0.003225 | 0.018581 | Early |
| AOC3 | -0.5798 | 0.000709 | 0.005852 | Early |
| MORF4L2 | -0.57608 | 0.001735 | 0.011249 | Early |
| GABARAPL2 | -0.57497 | 0.000178 | 0.001751 | Early |
| LGALS1 | -0.57334 | 1.12E-07 | 2.63E-06 | Early |
| SPCS2 | -0.57222 | 0.00049 | 0.004184 | Early |
| MDFI | -0.57016 | 0.008299 | 0.037604 | Early |
| PNRC1 | -0.56973 | 0.000111 | 0.001236 | Early |
| Hs.SEPT2 | -0.56815 | 0.001309 | 0.009404 | Early |
| G0S2 | -0.56519 | 0.008273 | 0.037604 | Early |
| RPS27L | -0.55841 | 0.000842 | 0.006435 | Early |
| TIMP3 | -0.55406 | 0.001682 | 0.011104 | Early |
| YWHAQ | -0.55379 | 0.000138 | 0.001468 | Early |
| NDUFS5 | -0.5475 | 0.000192 | 0.001862 | Early |
| ANXA5 | -0.54611 | 0.000104 | 0.001177 | Early |
| CALD1 | -0.54219 | 5.13E-05 | 0.000621 | Early |
| NAP1L1 | -0.53864 | 6.62E-06 | 0.000104 | Early |
| Hs.SEPT11 | -0.53603 | 0.005985 | 0.028776 | Early |
| TACC1 | -0.53508 | 0.003372 | 0.018976 | Early |
| COL5A2 | -0.53468 | 0.01006 | 0.043217 | Early |
| MTCH1 | -0.53432 | 0.001653 | 0.011009 | Early |
| CARHSP1 | -0.53339 | 0.004333 | 0.022386 | Early |
| SOD3 | -0.52552 | 0.002627 | 0.016165 | Early |
| SF3B2 | -0.52503 | 0.006204 | 0.029632 | Early |
| EIF1B | -0.52112 | 0.006617 | 0.031398 | Early |
| SEC11A | -0.51765 | 0.002925 | 0.017553 | Early |
| SEC61B | -0.51712 | 0.000997 | 0.007542 | Early |
| TBX2 | -0.51581 | 0.015875 | 0.061702 | Early |
| HSPB1 | -0.51386 | 0.000145 | 0.001518 | Early |
| KIF5B | -0.50934 | 0.012122 | 0.050004 | Early |
| CSDE1 | -0.50819 | 0.002588 | 0.01606 | Early |
| TMED9 | -0.50057 | 0.004239 | 0.022143 | Early |
| NDUFS6 | -0.49675 | 0.009337 | 0.041083 | Early |
| SLC25A3 | -0.49396 | 0.000278 | 0.002525 | Early |
| S100A13 | -0.49001 | 0.018448 | 0.070863 | Early |
| KANK2 | -0.48986 | 0.011696 | 0.04852 | Early |
| SERPINH1 | -0.48578 | 0.004191 | 0.022123 | Early |
| FBLN5 | -0.48495 | 0.009904 | 0.042799 | Early |
| MFGE8 | -0.48076 | 0.024787 | 0.088412 | Early |
| RBM3 | -0.4738 | 0.01043 | 0.044281 | Early |
| ARPC5 | -0.47105 | 0.002768 | 0.016744 | Early |
| RPN2 | -0.47002 | 0.020478 | 0.075467 | Early |
| EIF4H | -0.46734 | 0.008878 | 0.039543 | Early |
| PRDX6 | -0.4666 | 0.001187 | 0.008792 | Early |
| MYL6 | -0.46088 | 3.26E-08 | 9.09E-07 | Early |
| GDI2 | -0.45425 | 0.015836 | 0.061702 | Early |
| MXRA8 | -0.45303 | 0.014476 | 0.057745 | Early |
| GNAS | -0.44941 | 0.007093 | 0.033221 | Early |
| PSMB5 | -0.44353 | 0.0137 | 0.055708 | Early |
| CAV1 | -0.44144 | 0.001638 | 0.011009 | Early |
| FTH1 | -0.43957 | 4.06E-09 | 1.55E-07 | Early |
| KRT10 | -0.43938 | 0.024725 | 0.088412 | Early |
| HNRNPDL | -0.43903 | 0.0021 | 0.01314 | Early |
| GPX1 | -0.43719 | 0.012529 | 0.05139 | Early |
| CSRP1 | -0.4347 | 0.00076 | 0.006017 | Early |
| TMBIM6 | -0.43397 | 0.003062 | 0.017929 | Early |
| HNRNPA3 | -0.4339 | 0.015177 | 0.059883 | Early |
| CSNK2B | -0.43013 | 0.018858 | 0.072057 | Early |
| ESD | -0.42708 | 0.009859 | 0.042799 | Early |
| SCP2 | -0.42113 | 0.028794 | 0.099544 | Early |
| MORF4L1 | -0.41907 | 0.000482 | 0.004165 | Early |
| DKK3 | -0.41707 | 0.019368 | 0.072855 | Early |
| CFL1 | -0.41632 | 0.000596 | 0.00497 | Early |
| SH3BGRL | -0.41207 | 0.007413 | 0.034497 | Early |
| NDUFA4 | -0.41154 | 0.000166 | 0.001677 | Early |
| UBC | -0.41108 | 0.000732 | 0.005969 | Early |
| S100A11 | -0.41022 | 0.021602 | 0.079207 | Early |
| TMED2 | -0.40968 | 0.013735 | 0.055708 | Early |
| MGP | -0.40669 | 0.010881 | 0.045661 | Early |
| CST3 | -0.40508 | 0.019363 | 0.072855 | Early |
| COL6A1 | -0.39744 | 0.001467 | 0.010051 | Early |
| P4HB | -0.39601 | 0.015893 | 0.061702 | Early |
| CAPZB | -0.39509 | 0.00509 | 0.025403 | Early |
| ALDOA | -0.39333 | 0.009519 | 0.041632 | Early |
| ACTG1 | -0.39223 | 2.19E-05 | 0.000289 | Early |
| C12orf57 | -0.38414 | 0.014872 | 0.059002 | Early |
| LBH | -0.38187 | 0.008339 | 0.037604 | Early |
| PTMA | -0.37831 | 4.06E-06 | 6.85E-05 | Early |
| RAN | -0.37737 | 0.022276 | 0.081267 | Early |
| SUB1 | -0.36821 | 0.005647 | 0.027331 | Early |
| SRP9 | -0.36753 | 0.014284 | 0.057591 | Early |
| COL1A2 | -0.36717 | 0.000273 | 0.002511 | Early |
| HMGB1 | -0.3665 | 0.000829 | 0.006435 | Early |
| TMEM59 | -0.35658 | 0.004205 | 0.022123 | Early |
| EID1 | -0.35165 | 0.001229 | 0.009016 | Early |
| COL3A1 | -0.34911 | 0.010161 | 0.043394 | Early |
| CCNI | -0.34676 | 0.000762 | 0.006017 | Early |
| UBL5 | -0.34671 | 0.009296 | 0.041083 | Early |
| HNRNPA0 | -0.34656 | 0.010825 | 0.045661 | Early |
| GSTP1 | -0.33657 | 0.005093 | 0.025403 | Early |
| MFAP4 | -0.32692 | 0.005135 | 0.025403 | Early |
| UQCRQ | -0.31804 | 0.026824 | 0.094996 | Early |
| BSG | -0.31785 | 0.027575 | 0.096309 | Early |
| LAPTM4A | -0.31742 | 0.001401 | 0.009684 | Early |
| HSP90AB1 | -0.30996 | 0.001622 | 0.011007 | Early |
| SOD1 | -0.3008 | 0.019587 | 0.0733 | Early |
| DSTN | -0.29783 | 0.006904 | 0.032547 | Early |
| NUCKS1 | -0.28211 | 0.024843 | 0.088412 | Early |
| CHCHD2 | -0.27445 | 0.007708 | 0.035641 | Early |
| EIF4A2 | -0.26852 | 0.019286 | 0.072855 | Early |
| HNRNPA1 | -0.26191 | 0.000231 | 0.00218 | Early |
| BTF3 | -0.24315 | 0.005569 | 0.027137 | Early |
| EEF2 | -0.23726 | 0.005055 | 0.025403 | Early |
| H3F3B | -0.17482 | 0.028502 | 0.099008 | Early |
| RPL8 | -0.1464 | 0.003827 | 0.020583 | Early |
| RPL23A | 0.127451 | 0.016545 | 0.06389 | Late |
| MT-CO3 | 0.163465 | 0.02017 | 0.075096 | Late |
| RPL29 | 0.169345 | 0.002965 | 0.017643 | Late |
| RPLP2 | 0.177702 | 0.000756 | 0.006017 | Late |
| RPL18A | 0.216409 | 9.67E-07 | 1.71E-05 | Late |
| MT-CO1 | 0.263336 | 8.68E-05 | 0.001017 | Late |
| EEF1B2 | 0.27718 | 0.027593 | 0.096309 | Late |
| PFN1 | 0.291909 | 0.027396 | 0.096309 | Late |
| LDHB | 0.349191 | 0.008439 | 0.037819 | Late |
| LMO4 | 0.353492 | 0.011561 | 0.048239 | Late |
| OLFML3 | 0.396225 | 0.024824 | 0.088412 | Late |
| BTG1 | 0.412775 | 0.000172 | 0.001707 | Late |
| PSME2 | 0.414944 | 0.022841 | 0.082914 | Late |
| HSPE1 | 0.470328 | 0.004613 | 0.023582 | Late |
| SH3BGRL3 | 0.478596 | 0.003433 | 0.019171 | Late |
| LY6E | 0.480952 | 0.004348 | 0.022386 | Late |
| TECR | 0.500961 | 0.007873 | 0.036174 | Late |
| EPHX1 | 0.523167 | 0.005144 | 0.025403 | Late |
| CCBE1 | 0.594546 | 0.001787 | 0.011484 | Late |
| CYBA | 0.624535 | 0.000166 | 0.001677 | Late |
| SMPDL3A | 0.655026 | 0.003516 | 0.019377 | Late |
| B2M | 0.842561 | 9.93E-19 | 7.21E-16 | Late |
| S100A4 | 0.937335 | 0.000272 | 0.002511 | Late |

Table S10: Pathways activated in early maturing fibroblasts

| Name | Source | pValue | -LOG  (pvalue) | Genes in Input | Genes in Annotation | Gene Names |
| --- | --- | --- | --- | --- | --- | --- |
| Smooth Muscle Contraction | [REACTOME](http://www.ncbi.nlm.nih.gov/biosystems/1269870) | 3.182E-10 | 31.54934711 | [9](https://toppgene.cchmc.org/showQueryTerms.jsp?userdata_id=d07d352b-06a4-47dc-9922-b8e696c1ea84&feature=pt&row=0) | [36](https://toppgene.cchmc.org/showTermDetail.jsp?userdata_id=d07d352b-06a4-47dc-9922-b8e696c1ea84&category=Pathway&id=1269870) | TPM1, TPM2, TPM4, MYL6, CALD1, CALM2, ANXA2, ACTA2, MYL9 |
| Scavenging by Class A Receptors | [REACTOME](http://www.ncbi.nlm.nih.gov/biosystems/1269899) | 3.387E-09 | 28.13734487 | [7](https://toppgene.cchmc.org/showQueryTerms.jsp?userdata_id=d07d352b-06a4-47dc-9922-b8e696c1ea84&feature=pt&row=1) | [21](https://toppgene.cchmc.org/showTermDetail.jsp?userdata_id=d07d352b-06a4-47dc-9922-b8e696c1ea84&category=Pathway&id=1269899) | COL3A1, COL4A1, COL4A2, APOE, FTH1, COL1A1, COL1A2 |
| Extracellular matrix organization | [REACTOME](http://www.ncbi.nlm.nih.gov/biosystems/1270244) | 2.048E-08 | 25.54120904 | [18](https://toppgene.cchmc.org/showQueryTerms.jsp?userdata_id=d07d352b-06a4-47dc-9922-b8e696c1ea84&feature=pt&row=3) | [298](https://toppgene.cchmc.org/showTermDetail.jsp?userdata_id=d07d352b-06a4-47dc-9922-b8e696c1ea84&category=Pathway&id=1270244) | COL3A1, COL4A1, COL4A2, COL5A2, COL6A1, FBLN5, SPARC, TNC, SERPINH1, ITGB1, BGN, MFAP4, TIMP1, P4HB, BSG, ELN, COL1A1, COL1A2 |
| Platelet degranulation | [REACTOME](http://www.ncbi.nlm.nih.gov/biosystems/1269367) | 6.536E-08 | 23.86701678 | [12](https://toppgene.cchmc.org/showQueryTerms.jsp?userdata_id=d07d352b-06a4-47dc-9922-b8e696c1ea84&feature=pt&row=11) | [132](https://toppgene.cchmc.org/showTermDetail.jsp?userdata_id=d07d352b-06a4-47dc-9922-b8e696c1ea84&category=Pathway&id=1269367) | SPARC, CALM2, CALU, CFL1, ANXA5, TIMP1, TIMP3, SERPING1, CD63, TAGLN2, ALDOA, SOD1 |
| Response to elevated platelet cytosolic Ca2+ | [REACTOME](http://www.ncbi.nlm.nih.gov/biosystems/1269365) | 9.884E-08 | 23.27032975 | [12](https://toppgene.cchmc.org/showQueryTerms.jsp?userdata_id=d07d352b-06a4-47dc-9922-b8e696c1ea84&feature=pt&row=12) | [137](https://toppgene.cchmc.org/showTermDetail.jsp?userdata_id=d07d352b-06a4-47dc-9922-b8e696c1ea84&category=Pathway&id=1269365) | SPARC, CALM2, CALU, CFL1, ANXA5, TIMP1, TIMP3, SERPING1, CD63, TAGLN2, ALDOA, SOD1 |
| Binding and Uptake of Ligands by Scavenger Receptors | [REACTOME](http://www.ncbi.nlm.nih.gov/biosystems/1269897) | 1.151E-07 | 23.05060883 | [8](https://toppgene.cchmc.org/showQueryTerms.jsp?userdata_id=d07d352b-06a4-47dc-9922-b8e696c1ea84&feature=pt&row=14) | [49](https://toppgene.cchmc.org/showTermDetail.jsp?userdata_id=d07d352b-06a4-47dc-9922-b8e696c1ea84&category=Pathway&id=1269897) | COL3A1, COL4A1, COL4A2, SPARC, APOE, FTH1, COL1A1, COL1A2 |
| Collagen biosynthesis and modifying enzymes | [REACTOME](http://www.ncbi.nlm.nih.gov/biosystems/1270246) | 1.548E-07 | 22.62309119 | [9](https://toppgene.cchmc.org/showQueryTerms.jsp?userdata_id=d07d352b-06a4-47dc-9922-b8e696c1ea84&feature=pt&row=15) | [70](https://toppgene.cchmc.org/showTermDetail.jsp?userdata_id=d07d352b-06a4-47dc-9922-b8e696c1ea84&category=Pathway&id=1270246) | COL3A1, COL4A1, COL4A2, COL5A2, COL6A1, SERPINH1, P4HB, COL1A1, COL1A2 |
| Detoxification of Reactive Oxygen Species | [REACTOME](http://www.ncbi.nlm.nih.gov/biosystems/1270420) | 3.093E-07 | 21.62448983 | [7](https://toppgene.cchmc.org/showQueryTerms.jsp?userdata_id=d07d352b-06a4-47dc-9922-b8e696c1ea84&feature=pt&row=16) | [38](https://toppgene.cchmc.org/showTermDetail.jsp?userdata_id=d07d352b-06a4-47dc-9922-b8e696c1ea84&category=Pathway&id=1270420) | GPX1, GPX3, PRDX6, GSTP1, P4HB, SOD1, SOD3 |
| Collagen chain trimerization | [REACTOME](http://www.ncbi.nlm.nih.gov/biosystems/1470926) | 1.407E-06 | 19.43894624 | [7](https://toppgene.cchmc.org/showQueryTerms.jsp?userdata_id=d07d352b-06a4-47dc-9922-b8e696c1ea84&feature=pt&row=19) | [47](https://toppgene.cchmc.org/showTermDetail.jsp?userdata_id=d07d352b-06a4-47dc-9922-b8e696c1ea84&category=Pathway&id=1470926) | COL3A1, COL4A1, COL4A2, COL5A2, COL6A1, COL1A1, COL1A2 |
| Semaphorin interactions | [REACTOME](http://www.ncbi.nlm.nih.gov/biosystems/1270304) | 1.553E-06 | 19.29651074 | [8](https://toppgene.cchmc.org/showQueryTerms.jsp?userdata_id=d07d352b-06a4-47dc-9922-b8e696c1ea84&feature=pt&row=20) | [68](https://toppgene.cchmc.org/showTermDetail.jsp?userdata_id=d07d352b-06a4-47dc-9922-b8e696c1ea84&category=Pathway&id=1270304) | DPYSL2, MYH10, MYL6, CFL1, ITGB1, RHOC, MYL9, HSP90AB1 |
| Collagen formation | [REACTOME](http://www.ncbi.nlm.nih.gov/biosystems/1270245) | 1.807E-06 | 19.07797206 | [9](https://toppgene.cchmc.org/showQueryTerms.jsp?userdata_id=d07d352b-06a4-47dc-9922-b8e696c1ea84&feature=pt&row=22) | [93](https://toppgene.cchmc.org/showTermDetail.jsp?userdata_id=d07d352b-06a4-47dc-9922-b8e696c1ea84&category=Pathway&id=1270245) | COL3A1, COL4A1, COL4A2, COL5A2, COL6A1, SERPINH1, P4HB, COL1A1, COL1A2 |
| Vesicle-mediated transport | [REACTOME](http://www.ncbi.nlm.nih.gov/biosystems/1269876) | 4.405E-06 | 17.79242655 | [23](https://toppgene.cchmc.org/showQueryTerms.jsp?userdata_id=d07d352b-06a4-47dc-9922-b8e696c1ea84&feature=pt&row=40) | [660](https://toppgene.cchmc.org/showTermDetail.jsp?userdata_id=d07d352b-06a4-47dc-9922-b8e696c1ea84&category=Pathway&id=1269876) | COL3A1, COL4A1, COL4A2, KDELR2, SPARC, CALM2, GABARAP, GABARAPL2, APOE, GDI2, ARPC5, ARF1, ARF4, UBC, FTH1, CD59, TMED9, DYNLL1, TMED2, KIF5B, YWHAQ, COL1A1, COL1A2 |
| Assembly of collagen fibrils and other multimeric structures | [REACTOME](http://www.ncbi.nlm.nih.gov/biosystems/1270247) | 7.573E-06 | 17.01070364 | [7](https://toppgene.cchmc.org/showQueryTerms.jsp?userdata_id=d07d352b-06a4-47dc-9922-b8e696c1ea84&feature=pt&row=44) | [60](https://toppgene.cchmc.org/showTermDetail.jsp?userdata_id=d07d352b-06a4-47dc-9922-b8e696c1ea84&category=Pathway&id=1270247) | COL3A1, COL4A1, COL4A2, COL5A2, COL6A1, COL1A1, COL1A2 |
| Neutrophil degranulation | [REACTOME](http://www.ncbi.nlm.nih.gov/biosystems/1457780) | 0.00002818 | 15.11496886 | [18](https://toppgene.cchmc.org/showQueryTerms.jsp?userdata_id=d07d352b-06a4-47dc-9922-b8e696c1ea84&feature=pt&row=47) | [492](https://toppgene.cchmc.org/showTermDetail.jsp?userdata_id=d07d352b-06a4-47dc-9922-b8e696c1ea84&category=Pathway&id=1457780) | SPARC, CALM2, CALU, CFL1, ANXA5, TIMP1, TIMP3, SERPING1, CD63, TAGLN2, ALDOA, SOD1, COL3A1, COL4A1, COL4A2, APOE, COL1A1, COL1A2 |

Table S11: Cellular Classification Accuracy

|  | **Newborn Human Only** | **Human Mouse Combined** | **Overlap** | **Accuracy** |
| --- | --- | --- | --- | --- |
| Mesenchymal | 3553 | 1476 | 1431 | 96.95 |
| Endothelial | 1119 | 641 | 625 | 97.50 |
| Immune | 618 | 186 | 182 | 97.85 |
| Epithelial | 209 | 24 | 18 | 75.00 |
| TOTAL | 5499 | 2327 | 2256 | 96.95 |

Figure S1: Schematic of the steps involved in the process of single-cell sequencing of the human lungs, further integrating the data from human cells to the mouse single cell sequencing data, and finally estimating ages of human cells using the mouse cells.

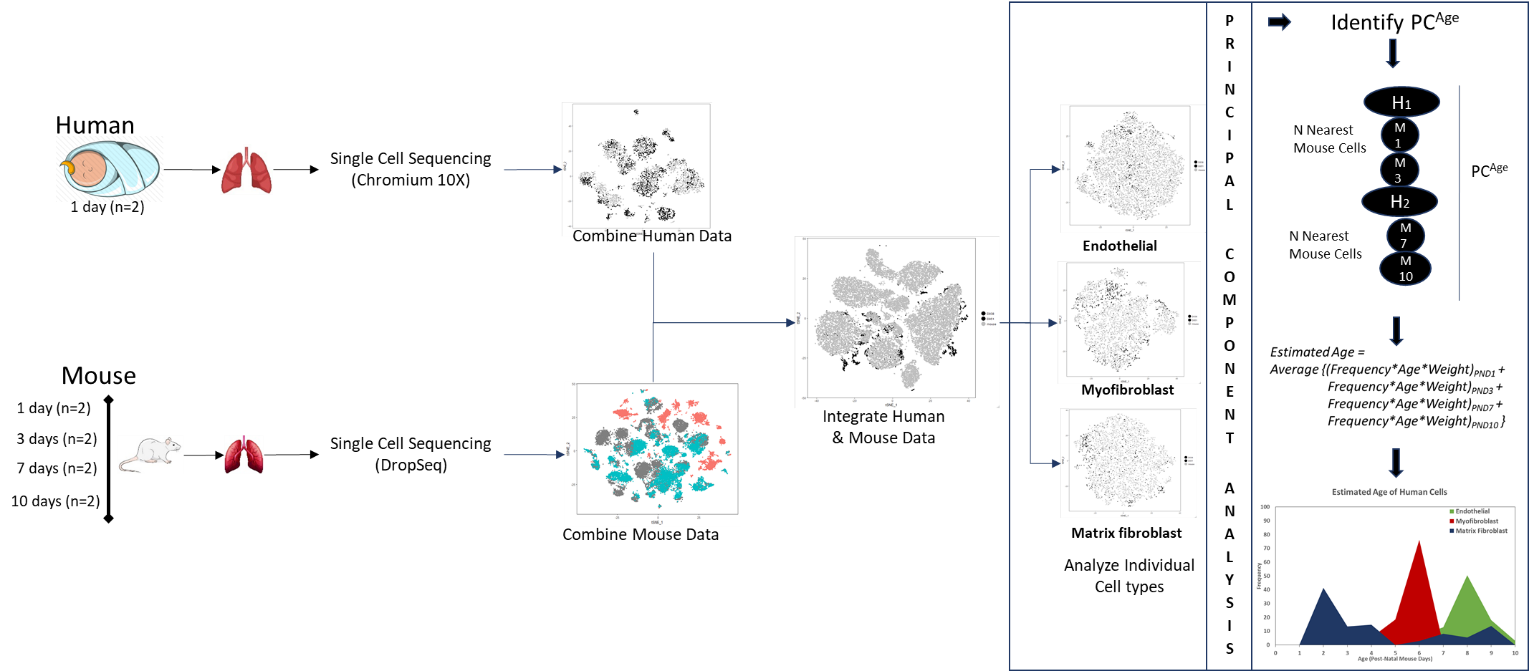

Figure S2: Quality Control metrics. We visualized gene and molecule counts, and mitochondrial genes present to filter cells for quality, and potentially exclude doublets.

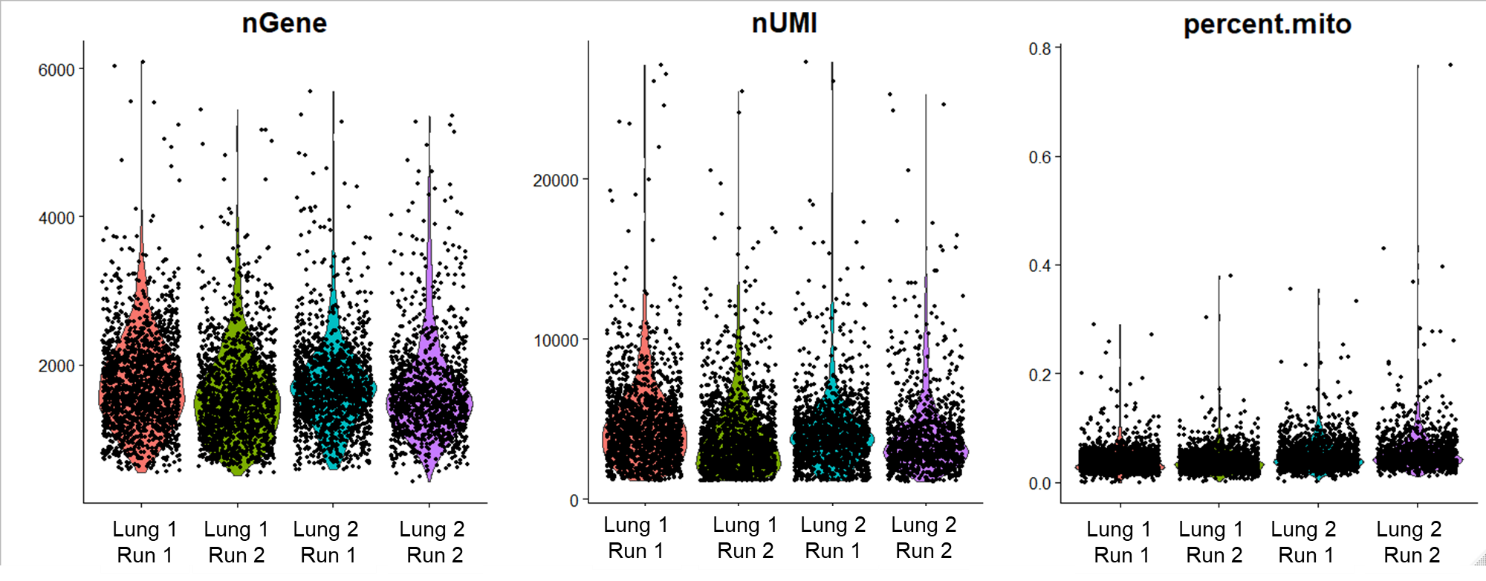

Figure S3: Identification of lung cell types using single cell RNA sequencing of newborn human lung. t-distributed Stochastic Neighbor Embedding (tSNE) analysis of cells leads to assignment of cell types to distinct tSNE clusters. Visualization of distinct cell clusters in tSNE plot from donor 1 (Full Term, born at 38 weeks of gestation), run 1 (A), run 2 (B), donor 2 (Pre-Term, born at 31 weeks of gestation), run 1 (C), and run 2 (D).

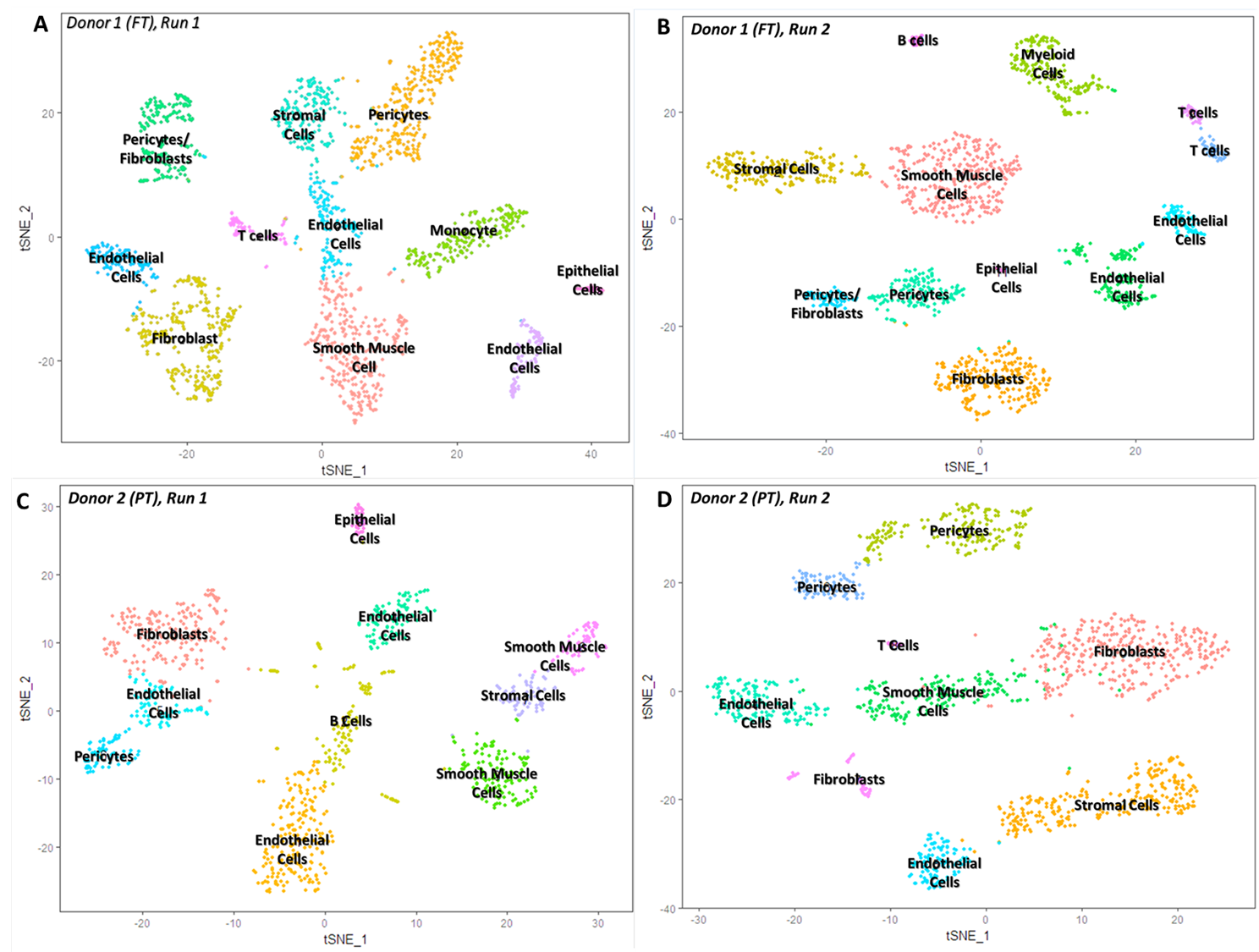

Figure S4 Identification of mouse lung cell types from Dropseq analysis of mouse lung. (A) Expression of known cell type markers in tSNE plot of cells. (B) The assignment of cell clusters to four major cell types, including endothelial cells, mesenchymal cells, immune cells, and epithelial cells. (C) Genes expression patterns of select markers of individual mouse cell clusters.

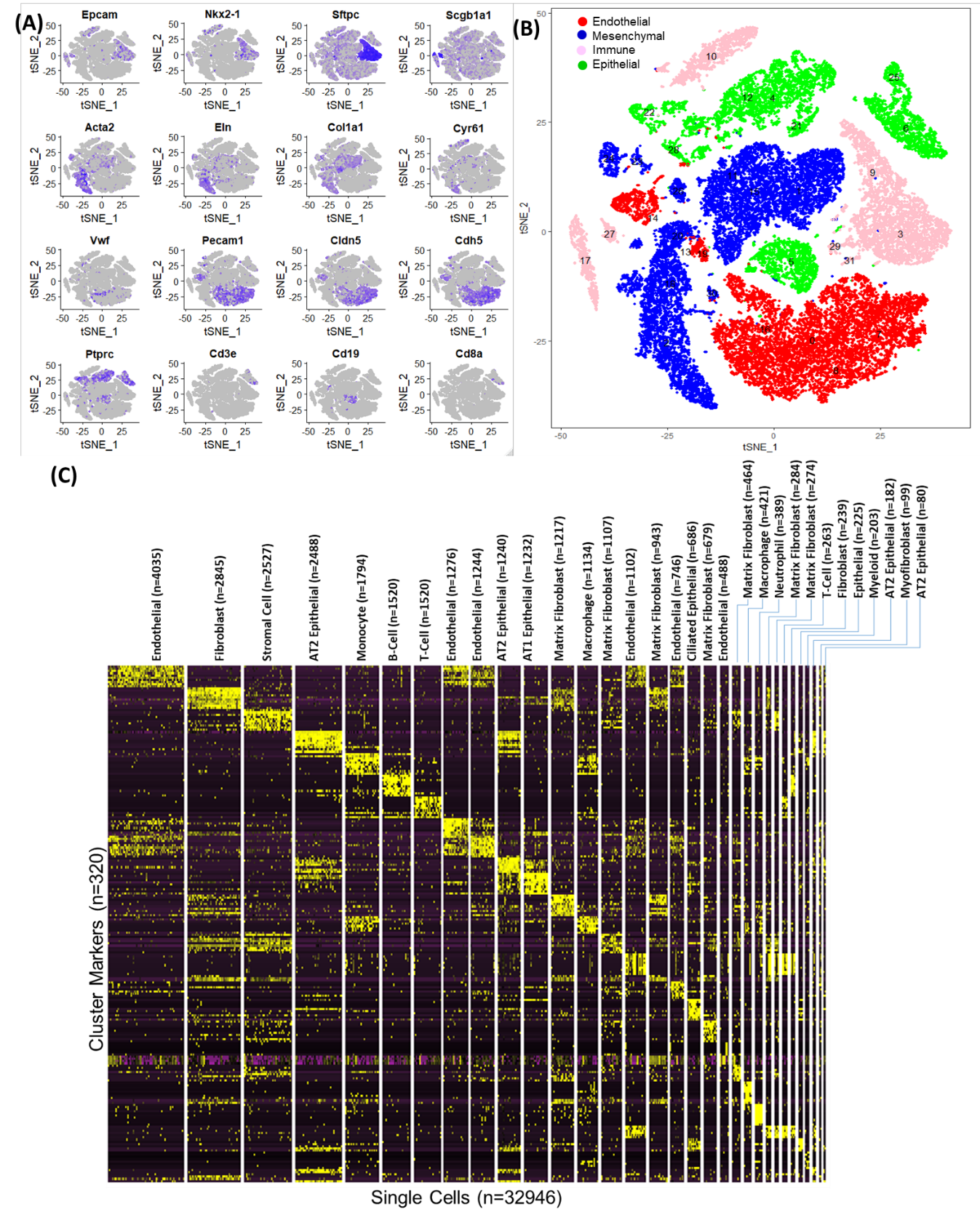

Figure S5. Relationship among the stages of lung development in human and mouse, focusing on the timeframe during which the lungs originated that were used to derive the single cell sequence data. The time points sampled in the transcriptomic analyses are indicated by arrows in the middle.

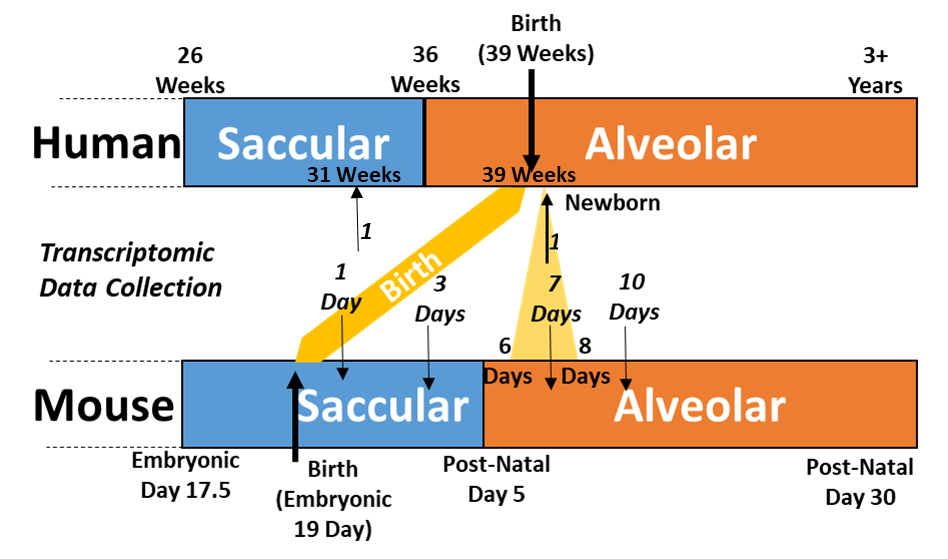

Figure S6: Presence of immune cells in newborn human lungs. (a) Expression of known immune cell markers PTPRC (CD45) and HLA-DRA in tSNE plot of cells. (b) Proportion of immune cell populations among all lung cells observed by sorting in four lungs obtained from one-day old donors, and (c) distributions of different immune cell populations in the same lung samples, as assessed by flow cytometry.

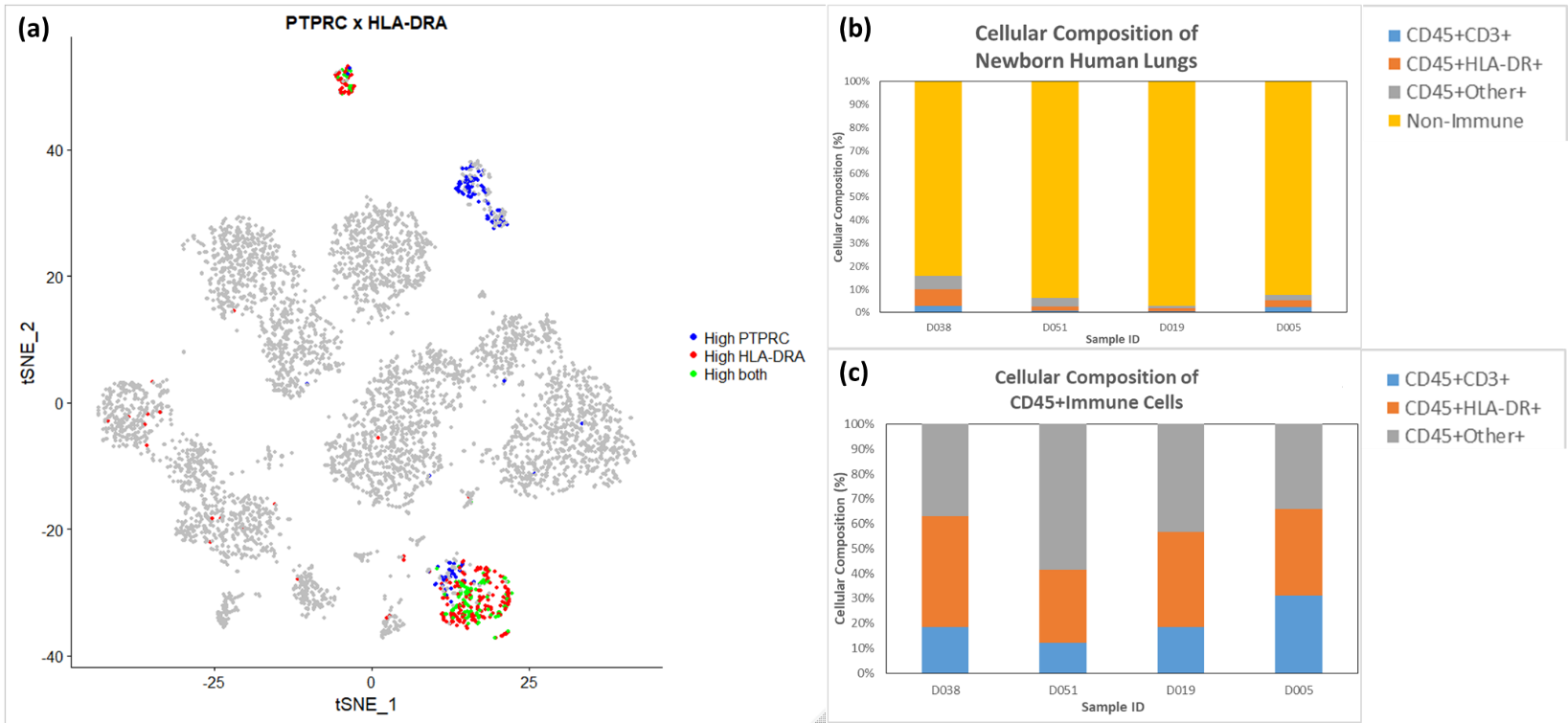

Figure S7: Differential expression analysis of developmentally early and later matrix fibroblasts identified 210 genes as differentially expressed. (a) Heatmap of 210 differentially expressed genes, where the rows indicate the genes, and columns indicate the individual cells. High expression is shown by yellow, and low expression is shown by purple. (b) Pathway analysis of 187 genes upregulated in immature fibroblasts identified 15 pathways significantly associated with those cells. Y-axis indicates the pathway names, and x-axis indicates -log(p-values). -log(p-values) greater than 1.3 indicates significance.

(a)

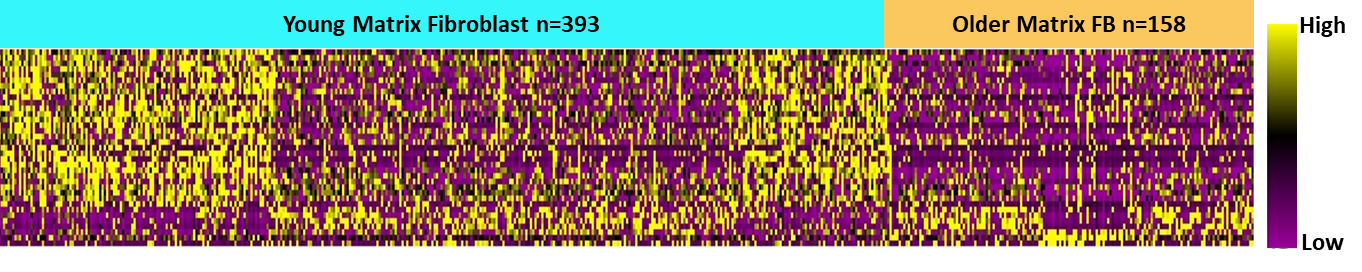

(b)

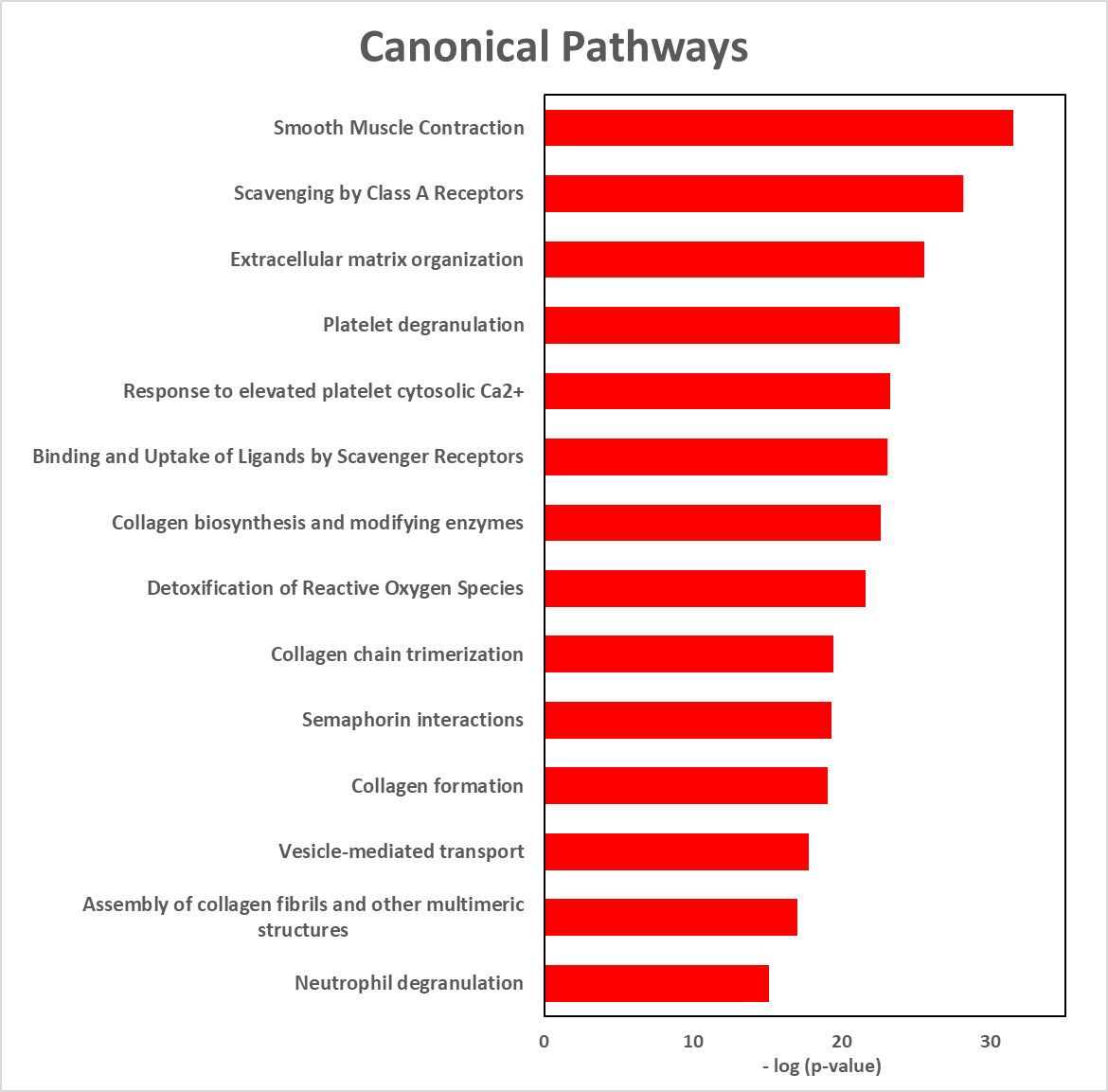

Figure S8: Location of cells expressing HES1. (A) Distribution of the estimated ages of the human cells among the different cell types, derived from post -natal age (PND) of 100 nearest mouse cells to each of the human cells. Shown here are proportion of cells of individual human cell types at each stage of maturity (represented by each circular ring) defined in terms of estimated post-natal day age of mouse. (B) Expression of younger matrix fibroblast marker HES1 in tSNE plot of cells. (C-F) tSNE plots showing coexpression of HES1 with known matrix fibroblast markers (COL6A1 and TCF1) serving as positive controls, and epithelial or endothelial cell markers (CDH1 and PECAM1) serving as negative controls. (G) Fluorescent in situ hybridization (FISH) combined with immunofluorescence of Immature Matrix Fibroblast marker HES1 (red), Non-Mesenchymal Cell Markers, PECAM1 or CDH1 cyan), and Mesenchymal Cell Markers COL6A3 or TCF21 (green) on newborn human lung sections from a donor lung of 1 day of age. Pink arrows indicated the presence of immature matrix fibroblasts shown by co-localization of HES1 (red) and Mesenchymal Cell Markers COL6A3 or TCF21 (green) while yellow arrows indicate non HES1 expressing cells. The scale bar is 50um.

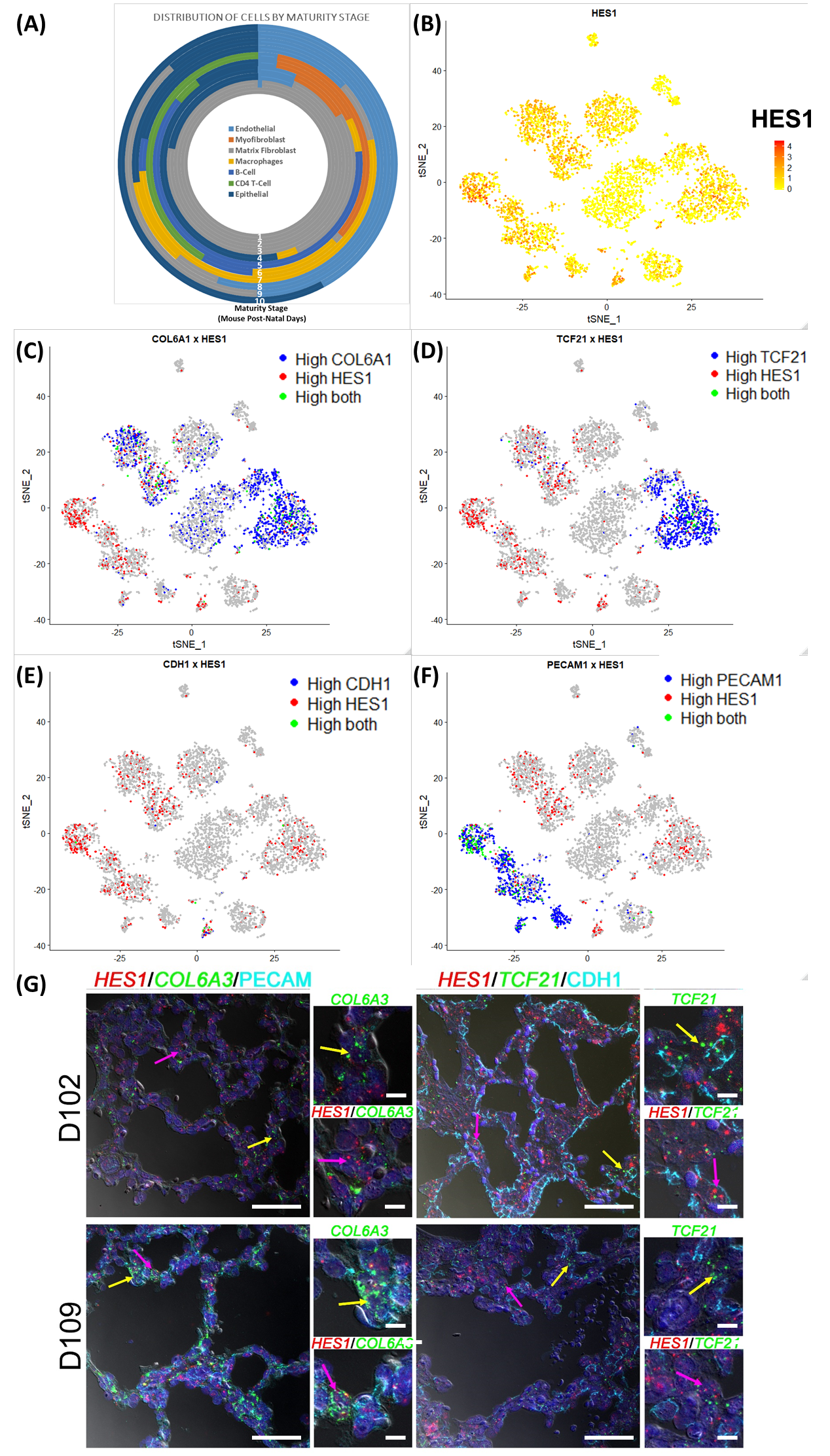

Figure S9: Location of cells expressing HES1 in Human and Mouse cells. (A) Expression of younger matrix fibroblast marker HES1 in tSNE plot of human and mouse cells. (B-E) tSNE plots showing coexpression of HES1 with known matrix fibroblast markers (COL6A1 and TCF1) serving as positive controls, and epithelial or endothelial cell markers (CDH1 and PECAM1) serving as negative controls.

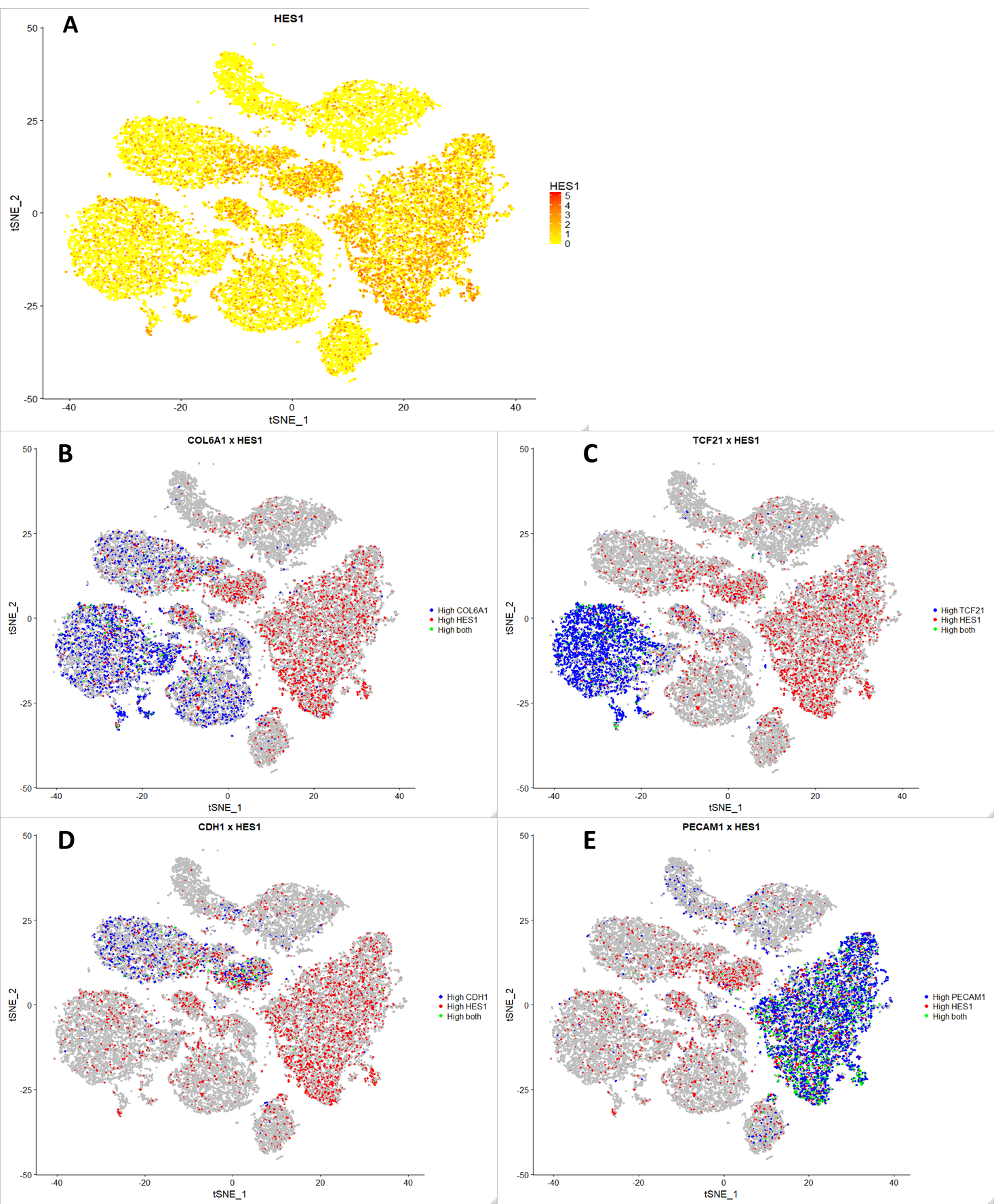
